# BioFabric Visualization of Network Alignments

**DOI:** 10.1101/2019.12.18.881664

**Authors:** Rishi M. Desai, William J.R. Longabaugh, Wayne B. Hayes

## Abstract

**Background:** Dozens of global network alignment algorithms have been developed over the past fifteen years. Effective network visualization tools are lacking and would enhance our ability to gain an intuitive understanding of the strengths and weaknesses of these algorithms.

**Results:** We have created a plugin to the existing network visualization tool BioFabric, called *VISNAB: Visualization of Network Alignments using BioFabric*. We leverage BioFabric’s unique approach to layout (nodes are horizontal lines connected by vertical lines representing edges) to improve understanding of network alignment performance. Our visualization tool allows the user to clearly spot deficiencies in alignments that cannot be detected through simply evaluating and comparing standard numerical topological measures such as the Edge Coverage (*EC*) or Symmetric Substructure Score (*S*^3^). Furthermore, we provide new automatic layouts that allow researchers to identify problem areas in an alignment. Finally, our new definitions of *node groups* and *link groups* that arise from our visualization technique allows us to also introduce novel numeric measures for assessing alignment quality.

**Conclusions:** Our new approach to visualize network alignments will allow researchers to gain a new, and better, understanding of the strengths and shortcomings of the many available network alignment algorithms.

## Background

Network alignment is the process of finding a mapping between the nodes of two networks. Alignments have been used in social networks to identify social structure and de-anonymize supposedly anonymous networks [1], brain connectomes to aid the creation of a standardized brain atlas [2, 3], and biomolecular networks to identify functional relationships and to transfer information between species [4]. The increasing availability of protein-protein interaction (PPI) networks over the last two decades has spurred the development of dozens of new network alignment algorithms. Particularly, the alignments of PPI networks reveal key insights in protein function and similarity [4], which in turn offers better understanding of mechanisms of human disease [5] and the process of aging in humans [6]; this may enable the transfer of biological information across species.

The BioFabric network visualization tool was created to help visualize and analyze large and complex networks [7]. While the traditional way of drawing nodes as dots and edges as lines (a method we refer to as *node link diagrams*), BioFabric depicts nodes as *horizontal lines*, while edges are vertical lines; small squares at the top and bottom of an edge denote the nodes (horizontal lines) the edge connects. There is one node line per row, and one edge line per column, arranged on a strictly regular grid. Thus, edges *never* overlap, completely eliminating the inevitable “hairball” that results from depicting dense networks with a traditional node-link diagram. Also, since links can originate and terminate anywhere along the appropriate node line segments, there is complete freedom to decide where a link is drawn. This flexibility for edge placement can be used to create many types of network visualizations that present meaningful semantic groupings of edges. The approach BioFabric uses has some similarities with “visibility representations” [8, 9], and a small example of using “nodes as lines” appeared in [10]. But BioFabric does not constrain nodes to be represented as discrete blocks and does not attempt to minimize link crossings at all. Indeed, in BioFabric it is possible to create useful network visualizations even when there are millions of intersections between node and edge lines.

There have been previous tools that have tackled the problem of visualizing network alignments [11, 12]. These tools still rely on variations of node-link diagrams and consequently suffer from the fundamental problems that plague this technique: alignments become harder to visualize as they become larger and more complex. The advantages provided by our new BioFabric plugin permit an entirely new approach to visualizing network alignments.

### Our Contributions

BioFabric’s ability to group and order *both* nodes and edges in meaningful ways provides a new way to visualize and understand network alignments. Here we will introduce a new system of node and edge groupings, with a new associated layout, that provides unique insights into alignments. We also provide a new layout technique that leverages the linear ordering generated by node misalignments that can highlight problems in the alignment. Finally, we introduce some new metrics, inspired by our visualization approach, that can be used to assess the quality of an alignment.

## Methodology

### Overview and Nomenclature

Figure 1 illustrates how BioFabric depicts network alignments and introduces the color-based nomenclature we use in this paper. Figure 1A depicts the alignment using a traditional node-link representation. Nodes from the smaller *blue* network (top left) are aligned one-to-one onto nodes from the larger *red* network (top center), creating a combined network (top right) that is the union of both under one of many possible alignments. When network elements (nodes or edges) are matched in this procedure, we refer to them as *purple*. We refer to unmatched elements from the smaller network as *blue*, and unmatched elements from the larger network are called *red*.

**Figure 1.**
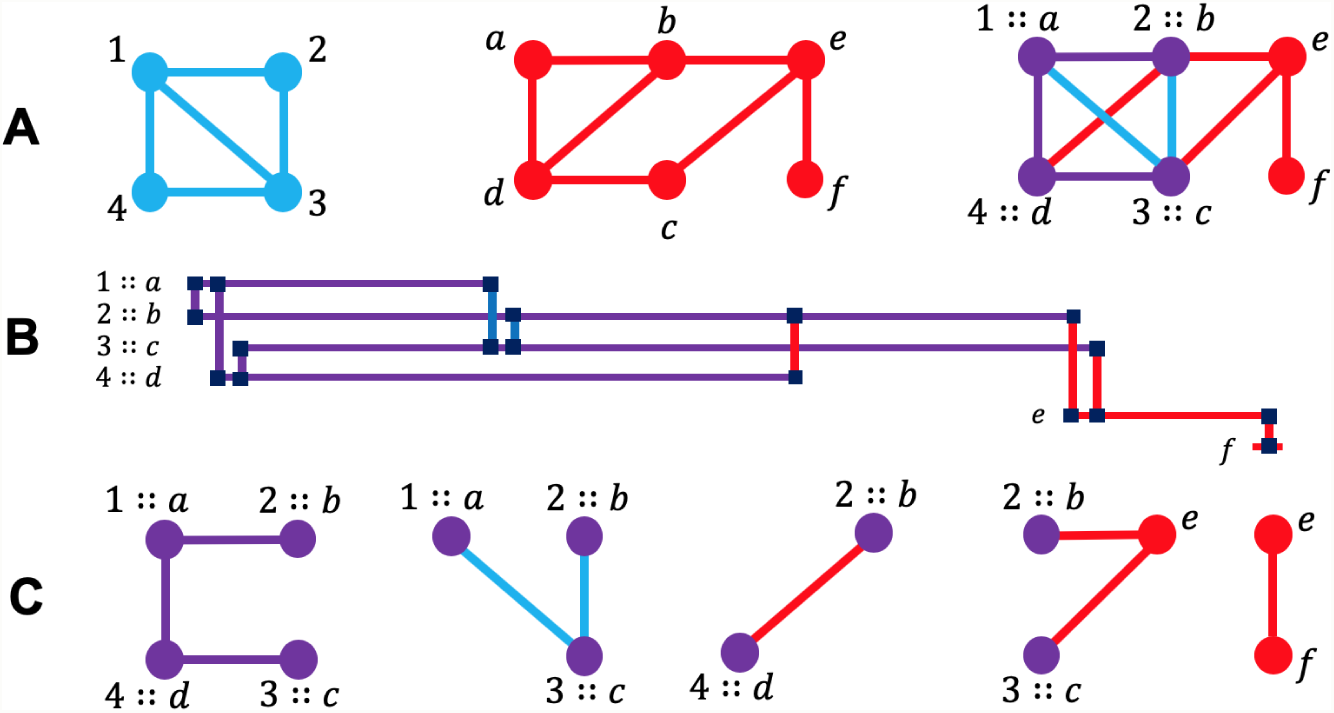
Network alignments with BioFabric. (A) A traditional node-link diagram to show a simple alignment, where a smaller blue network (upper left) is aligned onto a larger red network (upper center). The resulting aligned network (upper right) represents aligned nodes and edges as purple; node 1 aligned onto *a* is labeled as 1::*a*. Elements not aligned remain blue or red. (B) BioFabric visualizes the same upper-right network, where nodes are drawn as *horizontal lines* and edges as *vertical lines*. Purple horizontal lines are aligned nodes; red horizontal lines are unaligned nodes. There are no blue horizontal lines because we assume every node from *G*_1_ is aligned to some node in *G*_2_. (C) Each of the five different link groups described in the text, with the edges drawn directly under their corresponding BioFabric depiction. Not shown in this diagram are the node groups, which are not ordered in this depiction, for each node: 1::*a*: **(P:pBp)**; 2::*b*: **(P:P/pBp/pRp/pRr)**; 3::*c*: **(P:P/pBp/pRr)**; 4::*d*: **(P:P/pRp)**; *e*: **(R:pRr/rRr)**, *f* : **(R:rRr)**

Of course, in this example we are making the simplifying (and very common) assumption that *all* of the nodes in the smaller network are aligned to nodes in the larger network. In fact, in the most general alignment problem, we can also end up with *blue* nodes in the final alignment. In fact, VISNAB is fully capable of dealing with blue nodes in the final alignment. However, in our discussion of the technique, and in all the case studies we are discussing in this paper, we will deal with alignments with no blue nodes in the interest of simplicity. See the Appendix for additional information on how VISNAB handles blue nodes.

Figure 1B shows how BioFabric represents the combined network: the nodes are depicted as horizontal lines, and the edges are depicted as vertical lines. For this example, to make the comparison more concrete, this figure departs from the actual BioFabric presentation in two ways. First, the nodes here are drawn purple and red to match their counterparts in the node-link diagram; second, the different classes of edges are distinctly grouped. In BioFabric, in order to allow the network alignment to scale, node colors are cycled in a strict pattern, and edges are organized on a uniform regular grid.

### Node and Link Groupings

Consider the *rows* in Figure 1B: aligned purple nodes (1::*a* through 4::*d*) are grouped together exclusively in the top rows, while the unaligned red nodes (*e* and *f*) are grouped together exclusively in the bottom rows. Similarly, edges can be grouped in columns. As Figure 1C shows, there are *five* distinct classes of edges in the aligned network. Moving from left to right, we have purple *matched* edges, blue *orphan* edges, and three distinct classes of red *untouched* edges defined based upon the color of the two nodes incident on the edge: both purple, both red, or one of each. We refer to these five distinct classes as the *link groups* for a network alignment; they are enumerated in Table 1, along with the symbol we use to describe each one (e.g. **pRr**). For an alignment without blue nodes, these five link groups are a partition *P* of the set of edges in the union of the blue and red networks (where empty sets are allowed in *P*). The full enumeration of the seven link groups for the blue node case is provided in the Appendix.

**Table 1.** The enumeration of VISNAB’s link groups for the case of an alignment with no blue nodes. The link group numbering corresponds to the numbering used for the full blue node case, given in the Appendix.

| Link Group | Edge Color | Endpoint 1 | Endpoint 2 | Symbol |
| --- | --- | --- | --- | --- |
| 1 | Purple | Purple | Purple | P |
| 2 | Blue | Purple | Purple | pBp |
| 5 | Red | Purple | Purple | pRp |
| 6 | Red | Purple | Red | pRr |
| 7 | Red | Red | Red | rRr |

BioFabric’s link group separation allows visual quantification of the “common topology” discovered by an alignment. For example, one could argue that a better alignment would result by rotating the blue square 90° in either direction, combining the red and blue edges in the square’s interior to a single purple edge; the BioFabric layout of the resulting alignment would immediately and visually depict the elimination of the red and blue interior edges.

The two major classes of purple and red nodes can be further subdivided into *node groups* based upon the types of edges incident upon the node; these twenty groups and their symbols are enumerated in Table 2 for the case with no blue nodes. For example, red nodes that are not singletons (groups 37-39) can be classified as having node neighbors that are: (i) only purple (all incident edges are **pRr**), (ii) purple and red (incident edges are **pRr** and **rRr**), or (iii) only red (all incident edges are **rRr**). As with the link groups, for an alignment without blue nodes, these 20 groups are a partition *P* of the set of nodes in the union of the blue and red networks (where empty sets are allowed in *P*). The full enumeration of the 40 node groups for the blue node case are also provided in the Appendix.

**Table 2.** The enumeration of VISNAB’s node groups for the case of an alignment with no blue nodes. The node group numbering corresponds to the numbering used for the full blue node case, given in the Appendix.

| Node Group | Node Color | Incident Edges | Symbol |
| --- | --- | --- | --- |
| 1 | Purple | None | (P:0) |
| 2 | Purple | Purple | (P:P) |
| 3 | Purple | pBp | (P:pBp) |
| 6 | Purple | pRp | (P:pRp) |
| 7 | Purple | Purple, pBp | (P:P/pBp) |
| 10 | Purple | Purple, pRp | (P:P/pRp) |
| 11 | Purple | pBp, pRp | (P:pBp/pRp) |
| 14 | Purple | Purple, pBp, pRp | (P:P/pBp/pRp) |
| 17 | Purple | pRr | (P:pRr) |
| 18 | Purple | Purple, pRr | (P:P/pRr) |
| 19 | Purple | pBp, pRr | (P:pBp/pRr) |
| 22 | Purple | pRp, pRr | (P:pRp/pRr) |
| 23 | Purple | Purple, pBp, pRr | (P:P/pBp/pRr) |
| 26 | Purple | Purple, pRp, pRr | (P:P/pRp/pRr) |
| 27 | Purple | pBp, pRp, pRr | (P:pBp/pRp/pRr) |
| 30 | Purple | Purple, pBp, pRp, pRr | (P:P/pBp/pRp/pRr) |
| 37 | Red | pRr | (R:pRr) |
| 38 | Red | rRr | (R:rRr) |
| 39 | Red | pRr, rRr | (R:pRr/rRr) |
| 40 | Red | None | (R:0) |

### Network Merge

Most attempts to visualize an alignment provide side-by-side node-link diagrams of the input networks, with edges drawn between them to depict the alignment. The result rarely provides intuition about common topology discovered by the alignment. Instead, we *merge* the input networks (cf. Figure 1A, far right), characterizing the result with our node and link groups. Let *G*_1_ = (*V*_1_, *E*_1_) and *G*_2_ = (*V*_2_, *E*_2_) be two networks with |*V*_1_| ≤ |*V*_2_|. We define a pairwise global alignment from *G*_1_ to *G*_2_ is an injective function *a* : *V*_*a*_ → *V*_2_; every node in *V*_*a*_ ⊆ *V*_1_ is mapped to a distinct node in *V*_2_, and *V*_*a*_ is allowed to be a strict subset of *V*_1_, indicating some nodes in *V*_1_ (the blue nodes) may not be mapped. The merged network *G*_12_ = (*V*_12_, *E*_12_) consists of the nodes and edges in the two networks. All aligned nodes *u* ∈ *V*_*a*_, *v* ∈ *V*_2_ with *a*(*u*) = *v* are combined into one node *n* and added to *V*_12_. A combined node *n* is labeled in the format *u*::*v*. The unaligned nodes in *V*_1_ and *V*_2_ are also added to *V*_12_. *E*_12_ consists of all the edges in *E*_1_ and *E*_2_. An *aligned* edge is an edge (*u*_1_, *u*_2_) ∈ *E*_1_ : (*a*(*u*_1_), *a*(*u*_2_)) ∈ *E*_2_. Aligned edges (link group **P**) are only represented *once* in *E*_12_. Hence, the total number of edges |*E*_12_| varies with different alignments.

### New Metrics

Most topology-only network alignment approaches currently in use perform rather poorly across a wide range of biological test sets, particularly when the alignment is evaluated using **Node Correctness** (*NC*). Traditionally *NC* is the fraction of nodes *u* ∈ *V*_1_ aligned correctly when the correct alignment is known; we extended *NC* for alignments with unaligned nodes *u* ∈ *V*_1_. Let *a*′(*u*) be *a*(*u*) if *a*(*u*) is defined and *u* otherwise. Given an alignment *a* : *V*_*a*_ → *V*_2_ and the known correct alignment *a*_*c*_ : *V*_*c*_ → *V*_2_ with *V*_*a*_, *V*_*c*_ ⊆ *V*_1_, our *NC* measure is defined as the fraction of nodes 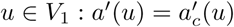.

In fact, slight differences in *NC* values, even if statistically significant, are arguably not very informative to assess pure topologically-driven alignments. We demonstrate here that other ways of assessing alignment performance can provide richer insights into the behavior of different techniques.

For example, if two nodes *u* and *v* in any graph *G* have an identical neighbor set (excluding the edge (*u, v*) if it exists), then it is impossible to distinguish between them using topology alone; thus, an alignment should not be penalized for swapping *u* and *v* if one of the two potential swaps is the “correct” one. Similarly, if *k* nodes all share the same neighbor set (excluding all edges between the *k* nodes themselves), any permutation of these *k* nodes should be scored equally if one of the permutations yields the “correct” alignment. This observation leads us to adapt a new **Jaccard Similarity** (*JS*) measure in the context of network alignment, as follows:

For some network *G* = (*V, E*), define the neighborhood of node *z*_1_ in *G* to be *N*_*G*_(*z*_1_) = {*z*_2_ ∈ *V* : (*z*_1_, *z*_2_) ∈ *E*}. For nodes *x, y* ∈ *V*, let *N*_*G*_(*x, y*) be the neighborhood of *x* disregarding *y*, and let *i*_*xy*_ be a corrective term accounting for a possible edge between the two, as follows: if *y* ∈ *N*_*G*_(*x*) then *N*_*G*_(*x, y*) = *N*_*G*_(*x*) − *y* and *i*_*xy*_ = 1, else *N*_*G*_(*x, y*) = *N*_*G*_(*x*) and *i*_*xy*_ = 0. Our extended *JS* definition *σ*_*G*_ : *V* × *V* → [0, 1] between two nodes is defined as:

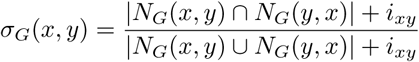

Intuitively, two aligned nodes score a 1.0 if they share an identical set of neighbors (ignoring a possible self-link between them) and are therefore impossible to distinguish using topology alone, and so a topology-driven alignment score shouldn’t penalize the misalignment of such pairs. (Note that when *x* and *y* are both singletons, we define *σ*_*G*_(*x, y*) ≡ 1.0 to avoid dividing by zero.)

Given node sets *V*_*a*_, *V*_*c*_ ⊆ *V*_1_, an alignment *a* : *V*_*a*_ → *V*_2_, and the correct alignment *a*_*c*_ : *V*_*c*_ → *V*_2_, our *JS* measure for the given alignment *a*, with respect to the correct alignment *a*_*c*_, in the case where |*V*_*a*_| = |*V*_*c*_| = |*V*_1_| is defined as:

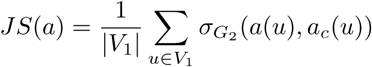

The *JS* measure for the “blue node” case, where |*V*_*a*_|, |*V*_*c*_| ≤ |*V*_1_|, is detailed in the Appendix. Significantly, Jaccard Similarity does *not* penalize a misalignment if the two nodes are topologically indistinguishable. Note also how *JS* gives “partial credit” for alignments where the nodes have nearly the same sets of neighbors.

The importance of node and link groups to VISNAB visualizations inspired us to design two new metrics that can be used to assess the performance of an alignment in the presence of a known correct alignment (*NC* = 1) between the two networks. For this, we use the node group and link group equivalence relations over the groupings of nodes and edges in an aligned network that we have developed. We can do this by comparing how the distribution of nodes and edges in the given alignment across the groupings compares to the distribution found in the correct alignment. To this end, we have created the **Node Group Similarity** (*NGS*) and **Link Group Similarity** (*LGS*) metrics. Both *NGS* and *LGS* are calculated using a vector representing the proportion of nodes and links distributed among the twenty node groups and five link groups, respectively. For example, the *NGS* vector **r** for *k* node groups ng_1_, ng_2_, …, ng*k* where |*V*_12_| = |*V*_1_| + |*V*_2_| − |*V*_*a*_| is

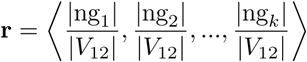

The angular similarity between the vector of the given alignment and the vector of the known correct alignment is calculated. For vectors with only positive elements, angular similarity = 1 − 2*θ/π*, where *θ* is the angle between the vector of the alignment and the vector of the correct alignment. Angular similarity was chosen (instead of e.g. cosine similarity) because it exhibits more rapid decay from the optimum value in the case of small angles.

All of these metrics, as well as other common measures, are displayed by VISNAB in a dialog when the user provides the correct alignment file as an optional part of the setup steps.

### Node and Link Group Layout Algorithm

Once we have created a merged network, we must provide an informative layout of the network that organizes the nodes and edges into a grouping framework that aids understanding the alignment. All BioFabric network layouts are fully defined by creating a linear ordering of the nodes, and a linear ordering of the edges. The default technique for laying out nodes is simple: within each node group it creates a linear ordering of the nodes using a breadth-first search from the node of highest degree, where neighboring nodes are visited in the order of highest to lowest degree. This basic technique, which uses a single queue for the breadth-first search, can be modified to create a layout that organizes the nodes using the alignment-based groupings we have defined. Specifically, we adapted it to use a *separate* queue for each of the twenty node groups. The resulting layout retains much of the same overall structure as the default algorithm, while ensuring that nodes are well-organized in each node group band. As nodes are visited during the search, they are added to the queue corresponding to their node group, and these node group queues are processed in the order listed in Table 2. A more complete description of the algorithm is provided in the Appendix.

Finally, if the correct alignment is provided, the user can choose to have correctly and incorrectly aligned nodes laid out separately in different node groups. The user can choose the criterion for the correct alignment to be based either on the traditional *NC* measure, or on our *JS* measure. If *JS* is chosen, the user can set the threshold value *β* ∈ [0, 1], so, in the case where |*V*_*a*_| = |*V*_*c*_| = |*V*_1_|, an aligned node *u*::*v* is denoted correct if *σ*_*G*_2 (*v, a*_*c*_(*u*)) ≥ *β*. If *JS* is chosen, the condition for correctly aligned nodes for the “blue node” case, where |*V*_*a*_|, |*V*_*c*_| ≤ |*V*_1_|, is detailed in the Appendix.

The default edge layout algorithm, using a slightly modified version of the existing link group feature described in [7], can be used to organize the edges into the five link groups. The new modification allows edges that are tagged with a particular relation to be grouped contiguously on a *per-network* basis; previously, this could only be done on a *per-node* basis. Using this new modification, one of five tags signifying the edge’s link group (Table 1) is assigned to each edge in the network. The edges are then partitioned based on these tags into five link groups, like those in Figure 1. Consequently, the upper-left corner of the layout contains the purple nodes that are either singletons or have only purple edges (**P:P**).

### Alignment Cycle Layout for Self-Alignments

A common approach for assessing the performance of alignment algorithms is to align a subset of a network to itself, where the larger network contains the same number of nodes but a larger number of edges [4, 13]. To allow researchers to better understand these analyses, we developed a special network alignment layout method that highlights alignment problems. Our technique leverages the fact that if a network is aligned to itself, then it can be viewed as a set of *cycles* each covering the set of *nodes*. Correctly aligned nodes will have a cycle length of one, *u* → *u*, but a misalignment of *u* → *v*, and *v* → *u*, will create a cycle of length two. More severe misalignments create longer cycles. Figure 2 shows a small example of this approach.

**Figure 2.**
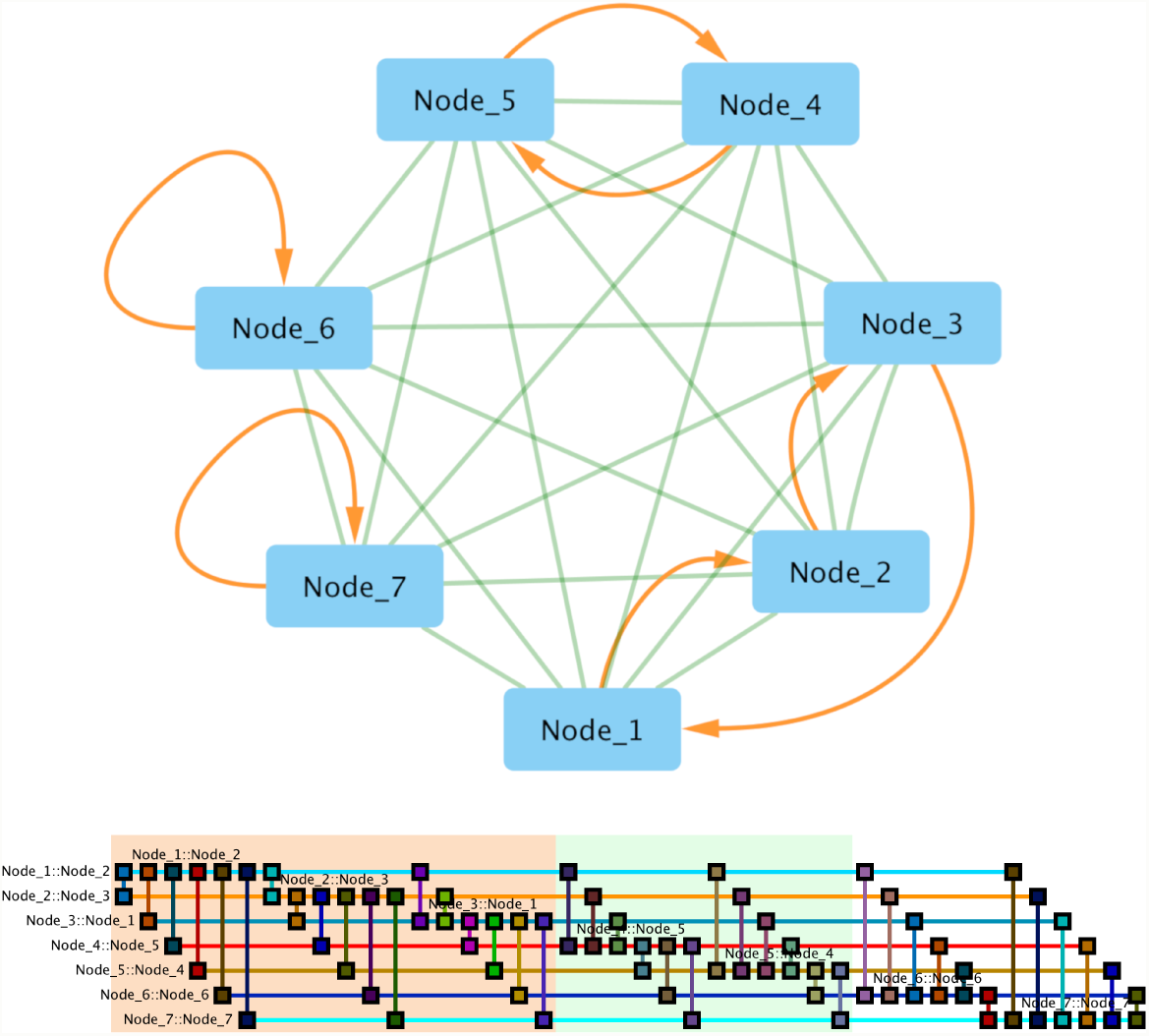
Understanding the Alignment Cycle layout. A network consisting of seven nodes has been aligned to itself (top); the orange directed edges indicate the results of the alignment, e.g. *Node 1* was incorrectly aligned to *Node 2*, while *Node 7* was correctly aligned. As shown, this results in a set of cycles (four in this example) which is then used to create the linear ordering needed for the BioFabric layout (bottom) that places aligned nodes in adjacent rows. The Alignment Cycle layout highlights incorrect alignment cycles with alternating colored blocks, as shown (block labels omitted).

Since a node layout in BioFabric is just a linear ordering of nodes, these alignment cycles provide a natural basis for specifying this ordering; the resulting layout vividly portrays the nature of the alignment. We start by placing a node in the first row, then use the alignment cycle containing that node to specify the row ordering of all remaining nodes in the cycle. The decision of how to order these cycles is again derived from a modification of BioFabric’s breadth-first search default layout algorithm [7]. For the alignment cycle layout, *all* the nodes in the cycle for a neighbor are simply placed first before the next neighbor is processed.

Though this technique is easiest to picture when a network is aligned to itself, the method has been generalized to handle network alignments from one network to another with *more* nodes. In this case, the alignment creates a set of *paths* instead of cycles, since not all nodes in the large network will map to a node in the small one. With paths, the algorithm places all nodes belonging to a path when any node in the path is first encountered in the search. Finally, the user can provide a mapping if the nodes in the two networks are in different namespaces (e.g. one network uses Saccharomyces Genome Database (SGD) IDs for node names, while the other network uses ENTREZ IDs).

### Interface with BioFabric

BioFabric Version 2 (now in beta release) provides a new plug-in architecture that allows developers to add new features to the program. A plug-in is written in Java, and when the .jar file containing the compiled code is placed in a directory specified by the user, the new functionality is available in BioFabric’s **Tools** menu. Our VISNAB plug-in uses this new architecture to provide the new functions that allow the user to load in two networks, an alignment file, and an optional correct reference alignment, and then process and lay out the desired view of the alignment. The plugin also uses the new node and link annotation feature that is now being introduced in BioFabric Version 2. This feature allows the user to specify spans of nodes or links that are then highlighted by drawing colored rectangles in the background.

## Results

### Case Study I: Simple Network Comparison

VISNAB can be used to directly compare two networks when aligning a network to itself. An example of this is where the nodes are proteins from the same organism and the networks are from two different studies. Our new node and link group definitions can provide visual insights into how the two networks compare. Notably, this technique can also be used to compare *any* two networks, as long as a 1:1 mapping of nodes between the two networks is provided.

Figure 3 shows a “correct” alignment between two datasets describing the *S*. *cerevisiae* (baker’s yeast) PPI network. The smaller *Yeast 2K* is derived from a network generated from data in [13] and used in [4], and contains 2,390 nodes and 16,127 edges, while the larger *SC* (BioGRID (v3.2.101, June 2013) [14]) contains 5,831 nodes and 77,149 edges. The networks are “aligned” based upon simply matching node names. Note these networks used different protein naming conventions; the Appendix provides details on how we created this correct alignment.

**Figure 3.**
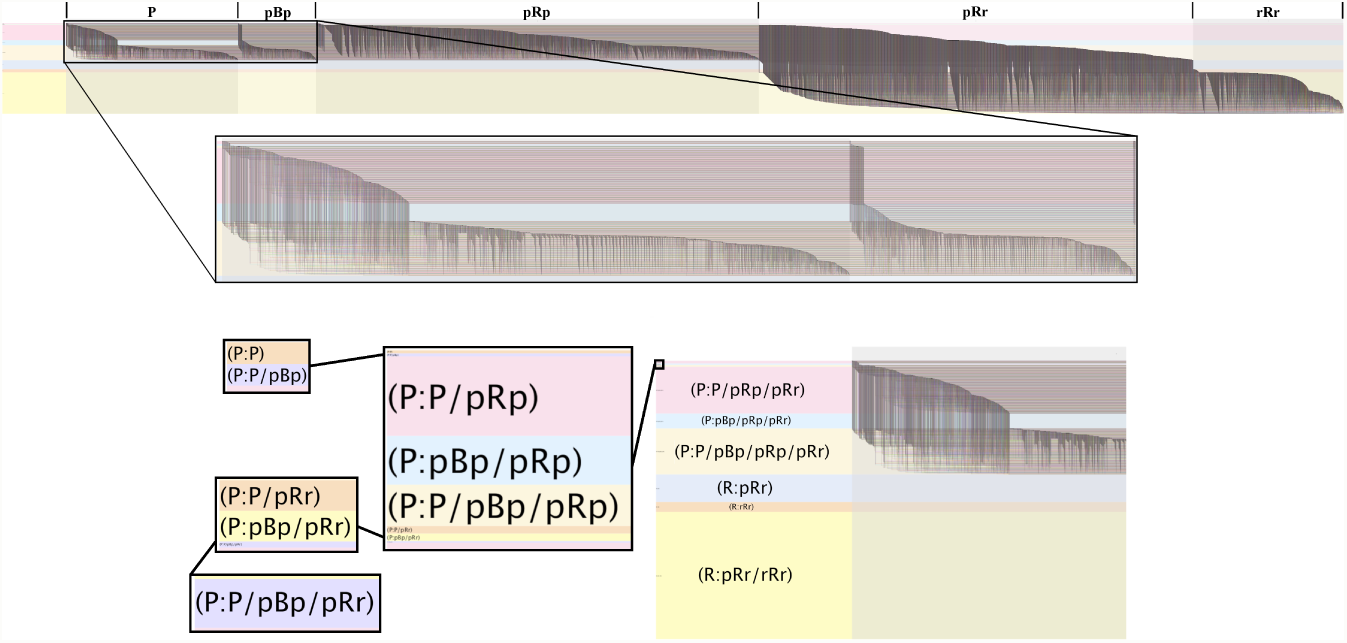
A direct comparison of network *Yeast 2K* [13] to *SC* [14]. Top: full network view. Middle: Detail of the subnetwork with purple (**P**) and blue (**pBp**) edges. Bottom: Detail of the node group assignments. The five link groups are denoted by alternating bands of light and dark shading; the node groups are each assigned fixed colors from BioFabric’s standard palette. Outsize tags for each link group have been added here for clarity along the top of the network; outsize tags for the largest node groups have been added here for clarity as well. As described in the text, common topological measures for alignment quality such as *S*^3^, *EC*, and *ICS* can be visually estimated using the *widths* of each link group. For example, *S*^3^ is the width of the leftmost link group **P** over the sum of the leftmost three link groups **P**, **pBp**, and **pRp**, i.e. about 1/4 (actual value: .248). Node groups can be used to gain insights into how the networks compare; see text for details.

The top of Figure 3 shows the entire merged network, while the detail below it displays the subnetwork with purple and blue edges. The lowest detail shows the distribution of nodes into our node groups, which are denoted by the horizontal colored bands. This display uses a recently added BioFabric Version 2 feature: node and link annotation displays.

Note how the five link groups and twenty node groups quickly reveal how these two networks compare. We immediately see that the number of nodes in the larger network are more than double the number in the original, as the purple nodes are in the colored bands down through the **(P:P/pBp/pRp/pRr)** (peach colored) node group; the node below that are red nodes only in the larger network. The edges in the two leftmost shaded link group bands (**P** and **pBp**) represent *exactly* the links in the *Yeast 2K* network. If the larger network were to be treated as a later, more complete, and more accurate survey, then the edges in group **pBp** represent the *false positives* from the earlier survey. The red edges in middle link group **pRp** are new edges from the more recent survey between old nodes from the original survey, while the edges in the right 40%, in the **pRr** and **rRr** link groups, represent new edges incident on at least one new red node.

Studying the relative sizes of the node groups also provides insights into how the networks differ. One thing to note right away is that the six node groups we do not expect to appear are in fact absent: **(P:0)**, **(P:pBp)**, **(P:pRp)**, **(P:pRr)**, **(P:pRp/pRr)**, and **(R:0)**. We expect this because these protein-protein interaction networks do not have any singleton nodes. Thus, for example, a node in the **(P:pBp)** class would require that the node be a singleton in the larger network. Conversely, a purple node with only red edges would require that the node be a singleton in the smaller network.

Another significant fact that is apparent by the node group distribution is that **over 90%** of the purple nodes are in the three node groups that have red edges going to both purple and red nodes: **(P:P/pRp/pRr)** (pink), **(P:pBp/pRp/pRr)** (powder blue), and **(P:P/pBp/pRp/pRr)** (peach). In contrast, there are only a tiny number of aligned nodes that have red edges going to *only* aligned or *only* unaligned nodes. Again, this seems consistent with a wider survey that would be expected to mostly find new interactions between both previously surveyed proteins as well as between those proteins and new proteins. The red nodes also show the same pattern of mostly (∼80%) having nodes with interactions to both new and old proteins. Finally, note how group **(P:pBp/pRp/pRr)** is the smallest of the major purple node groups. This tells us that there were fewer proteins in the original network where all the old edges were considered false positives, again a result that makes sense.

### Case Study II: Visualizing Common Topological Measures

There are several measures to assess the topological quality of an alignment. Depending on the context, these measures quantify how much topological similarity between the two PPI networks an alignment has exposed. Let *E*_*a*_ = {(*u*_1_, *u*_2_) ∈ *E*_1_ : (*a*(*u*_1_), *a*(*u*_2_)) ∈ *E*_2_} denote the edges in *G*_1_ aligned to edges in *G*_2_, i.e., purple edges. The ***EC*** measure (variously called Edge Coverage, Conservation, Correctness, or Correspondence by different authors) is the fraction of edges in the smaller network that are aligned to edges in the larger network: *EC*(*a*) = |*E*_*a*_|*/*|*E*_1_|. Let 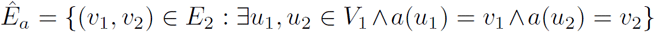 denote the set of edges of *G*_2_ induced on its aligned nodes. **Induced Conserved Structure** (*ICS*) is the ratio of aligned edges to induced edges: *ICS*(*a*) = |*E*_*a*_|*/*|*Êa*|. However, *EC* and *ICS* have the shortcoming that they can be high if the alignment maps sparse regions of one network to dense regions of the other. In the extreme, if *G*_2_ is a clique, any alignment has *EC* = 1. To overcome this drawback, [15] devised the **Symmetric Substructure Score** (*S*^3^), the ratio of aligned edges to all edges with aligned endpoint nodes:

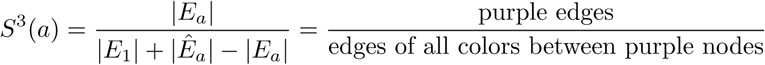

Since BioFabric lays out node rows and link columns on an absolute regular grid, the *proportions* of the widths of various node and link groups allow the user to rapidly visualize these topological metrics at a glance. For example, *S*^3^ can be estimated by comparing the *width* of the first link group (purple edges) over the *width* of the first three link groups (all links between purple nodes):

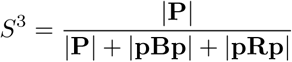

Looking at the top view of Figure 3, we visually estimate that this alignment has an *S*^3^ score of about 1/4 (actual value: .248). In the same fashion, *EC* is the width of the purple link group over the width of the purple and blue link groups:

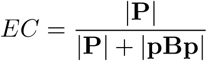

We can visually estimate *EC* to be about 2/3 (actual value: .689). Finally, *ICS* is the width of the purple link group over width of the purple group and red group with purple endpoint nodes:

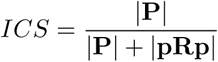

We can visually estimate *ICS* to be around 3/10 (actual value: .280). Thus, BioFabric’s node and link groups provide intuition on abstract topological measures.

### Case Study III: Visualizing Performance of Objective Functions

Choosing the best objective function to apply to a particular network alignment is currently a black art, and providing the researcher with a visual tool to assist in understanding the properties and performance of various objective functions helps to gain insights into this difficult problem.

Indeed, BioFabric’s presentation of network alignment allows the researcher to do just this. Figure 4 depicts four alignments between the *Yeast 2K* and *SC* networks previously introduced. The first row of the figure is the same gold-standard, correct alignment shown in Figure 3. For the other alignments, we used SANA [16], which allows us to select the objective functions we wish to use to create an alignment. These other three alignments were generated by running SANA for ten hours each (long enough for the random search algorithm to converge to a near-optimal score of the chosen objective function), optimizing the following objective functions: the second-row alignment was generated with an objective function utilizing only **Importance** (*I*) [17]; the third with .03 weight given to *S*^3^ and .97 weight given to *I*; and the fourth utilizing only *S*^3^.

**Figure 4.**
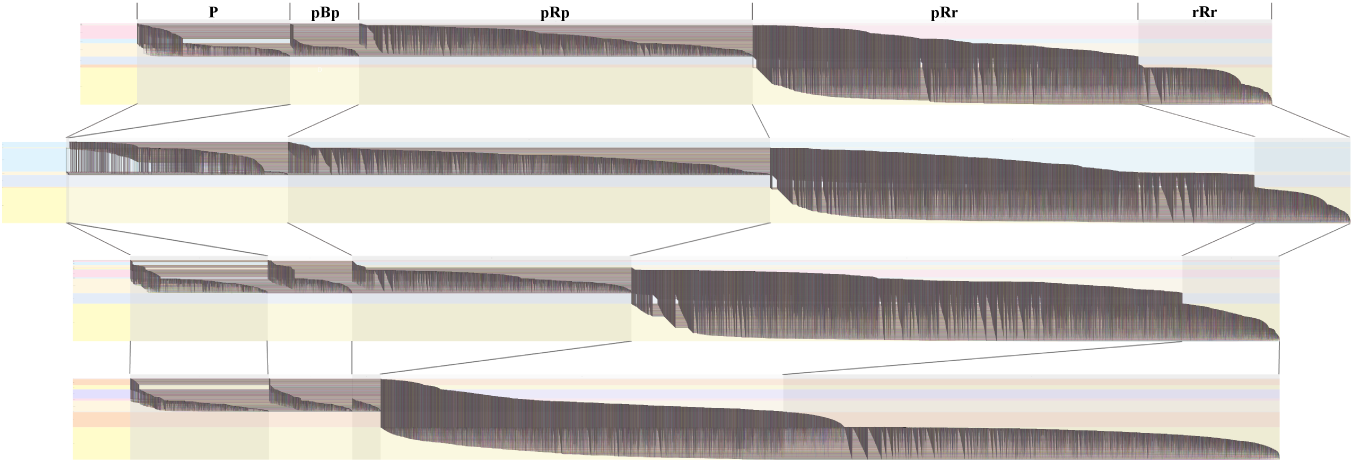
Visually assessing the performance of different objective functions with four alignments between *Yeast 2K* and *SC*. First row: gold-standard; second row: 1.0 *I* (Importance-only); third row: mixed with (.03 *S*^3^) + (.97 *I*); fourth row: 1.0 *S*^3^. We have drawn lines that follow each link group from the first row’s alignment to the respective link group in each successive row’s alignment. The alignment in the second row is noticeably wider than the others because aligned **P** edges represent *two* edges from the original network pair, and this alignment has very few **P** edges.

In addition to viewing the BioFabric plots of the four alignments, we can also look at our new *NGS, LGS*, and *JS* metrics for them, as well as the traditional *NC, S*^3^, and Resnik semantic similarity scores [18, 19, 20] calculated by FastSemSim [21]. These scores are shown in Table 3. The highest *S*^3^ value and the highest *NC* value are often the measures used to identify the “best” alignment. As the table shows, the pure *S*^3^ version provides the highest of both of these scores. Yet this “best” *NC* score is only 2.1%, which is not a very inspiring result. Even our new *JS* score of 6.9%, which is tolerant of mismatches that are topologically similar, is still very low.

**Table 3.** Scores for the four alignments between *Yeast 2K* and *SC*.

| Alignment | NGS | LGS | NC | JS | $S^3$ | Resnik |
| --- | --- | --- | --- | --- | --- | --- |
| Correct | 1.00 | 1.00 | 1.00 | 1.00 | 0.25 | 9.63 |
| $1.0 * I$ | 0.61 | 0.79 | 0.00042 | 0.021 | 0.0043 | 3.16 |
| $.03 * S^3 + .97 * I$ | 0.87 | 0.80 | 0.018 | 0.057 | 0.27 | 3.62 |
| $1.0 * S^3$ | 0.64 | 0.46 | 0.021 | 0.069 | 0.55 | 3.61 |

Simply *looking* at the four BioFabric plots in Figure 4 makes us question calling the pure *S*^3^ version the “best” alignment of these two networks. Recalling that *S*^3^ is the width of the first link group over the width of the first three link groups, we can instantly see that (i) the pure *S*^3^ version has a much higher value than any other alignment, even the correct alignment, and (ii) it achieves this by creating a alignment that forces *far* more edges into the two rightmost link groups **pRr** and **rRr**. Particularly worrisome is the very large increase in the size of **rRr** compared to the correct alignment; this set represents the edges and nodes that have been completely omitted from the alignment.

This fact that high-degree nodes were relegated to the red untouched nodes provides a clue to a way to improve the situation. Perhaps using Importance (in which the highest degree nodes tend to align to each other) as an objective function can remedy the problem? This approach gives rise to the “Pure Importance” version in row two of Figure 4. Indeed, this version has link groups **pRr** and **rRr** on the right end that appear to be *much* closer in size to the correct alignment, and the center link group **pRp** is closer to the correct alignment as well, albeit larger. But the leftmost group of purple **P** edges is so thin that it is almost non-existent. Hence, pure Importance yields an *S*^3^ value of merely .0043.

So, perhaps a simple linear mixture of the two objective functions can provide a compromise between these two extremes? To investigate this, we ran a series of alignment runs that used a linear mixture of the two objective functions, and nine combinations were scored. The scores for all nine are included in the Appendix. The mixture (.03 * *S*^3^) + (.97 * *I*) is shown in the third row, and provides a reasonable *visual match* of the gold-standard alignment in terms of the distribution of edges between the five link groups. This visual similarity is also borne from the metrics shown in Table 3.

Scanning across a variety of mixture values, this particular mix produced an alignment with a high *NGS*, a reasonably high *LGS*, a non-zero *NC*, and a respectable *JS* value. It also has the closest *S*^3^ value to the gold-standard, plus the highest functional similarity, i.e. the most biological relevance per the Resnik score, of all the alignments, though only marginally.

Another way to compare performance is by studying the node group distribution under the different objective functions, as shown in Figure 5. Notably, the “best” mixed alignment’s four largest node groups are the same as the four largest node groups of the correct alignment. The two largest purple node groups of each, **(P:P/pRp/pRr)** (pink) and **(P:P/pBp/pRp/pRr)** (peach), show a rich mixture of incident purple, red, and (in the latter group) blue edges. A significant difference between the two is the thicker bands above the pink **(P:P/pRp/pRr)** region for the mixed alignment compared to the correct alignment. The majority (76%) of nodes in those bands have no **pRr** incident edges, which reflect the fact that although there are *more* **pRr** edges in the mixed alignment (see Figure 3), those edges are concentrated across a *smaller* fraction of the purple nodes. Specifically, 23% of purple nodes in the mixed alignment have no **pRr** incident edges, compared to just 5% in the correct case. Table 10 in the Appendix provides the exact sizes of the different node groups for the four alignments.

**Figure 5.**
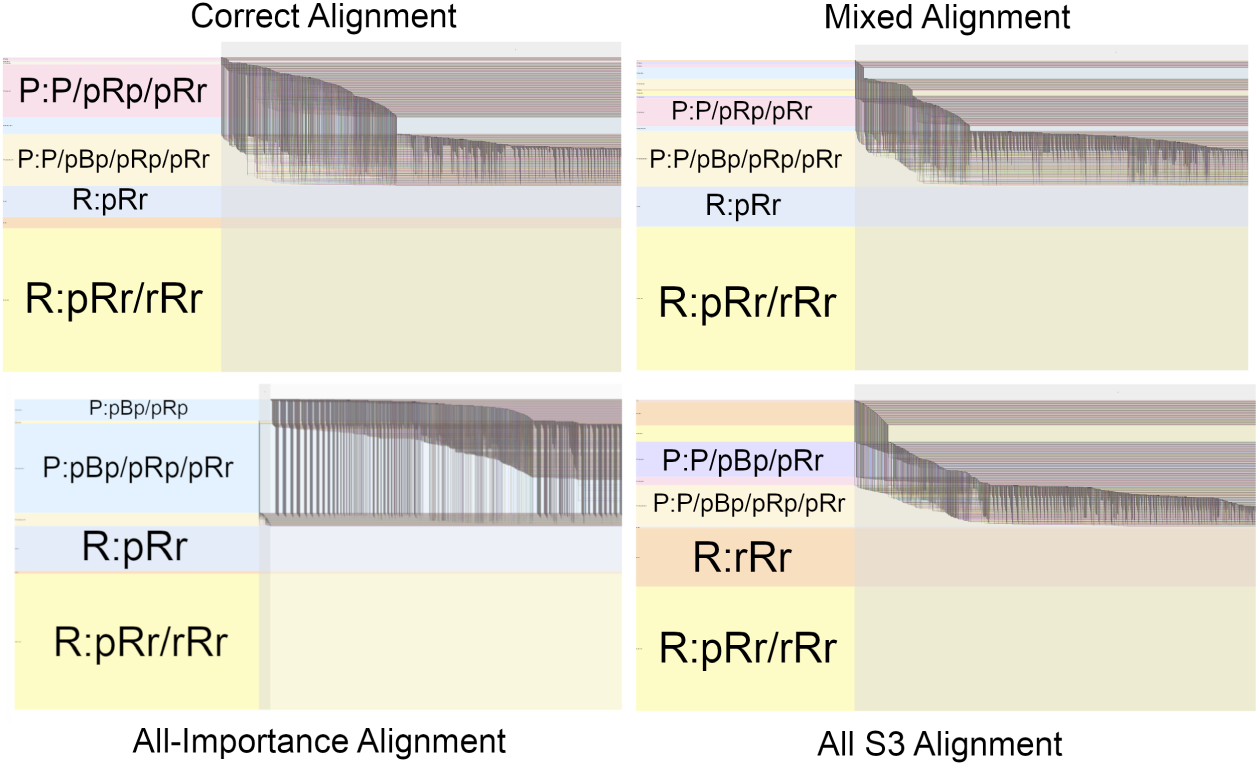
Visualization of node groups can also provide insights into understanding the performance of objective functions. Here we show the far left side of the four alignments in Figure 3, with the four largest node groups for each shown with added outsize tags for clarity. Note that four largest node groups of the “best” mixed alignment (upper right) are the same as those of the correct alignment (upper left), though there are size differences.

### Case Study IV: Finding Protein Cluster Misalignments

While looking at metric values for a particular alignment can provide some general idea of how an alignment performs, there is value in being able to quickly spot major problems in an alignment where a gold-standard alignment is available, and then be able to understand what the problem is.

Figure 6 shows a yeast network, *Yeast0*, of 1,004 nodes and 8,323 edges aligned to a network with the same set of nodes, but with 20% more (9,987) edges, *Yeast20*. This network is part of the *noisy yeast* variations on the network in [13]. The edges in the smaller network are a strict subset of the larger set. This dataset has been used in several previous studies [15, 16]. The particular alignment we are using here was part of an ensemble generated by SANA [16], but was unusually poor compared to other alignments in the ensemble; this lead us to ask what was so different about this poor alignment? This visualization is created using the same network merge technique described in **Network Merge**, above. The nodes in this network are labeled to represent the alignment result, e.g. *a*::*a* for a correct alignment, and *a*::*b* for an incorrect alignment. Edges are labeled **P** (for purple edges), **pBp** (blue), or **pRp** (red).

**Figure 6.**
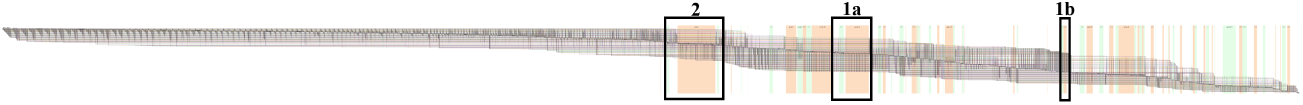
Full view of the problematic *Yeast0* to *Yeast20* alignment, arranged using the Alignment Cycle layout. BioFabric *shadow links* [7] are turned on. Alignment cycles of incorrectly aligned nodes are indicated with alternating orange and green blocks. The numbered boxes highlight the (noncontiguous) sections containing the one- and two-cycle cluster artifacts shown in Figure 7 and discussed in the text.

To be able to spot alignment problems, we view the network using our new Alignment Cycle layout method, described above. This layout does not partition nodes into node groups, but still generates link groups. Since the set of nodes is the same for both networks, there are no red nodes, so only three of the five link groups are present. For this type of presentation, we employ BioFabric’s *per-node* link group layout to separate links into the three distinct groups within each node’s dedicated *node zone*. This is in contrast to the *per-network* approach we used previously, which creates single global regions for each link group.

The large uncolored stretches of Figure 6 indicate that most of the nodes—particularly high-degree ones—are correctly aligned. However, cycles of misalignment, even those of length two, are shown with alternating orange and green blocks, generated using the link annotation feature. In order to understand these misalignments better, we zoom into each region to study them more closely.

One interesting region includes a misalignment cycle containing 15 nodes, as shown in Figure 7A. A node-link diagram version created using Cytoscape [22] is shown, where the protein-protein interactions are blue, and directed red edges represent the alignments. The BioFabric representation is shown at the top, with a inset detail shown as well. In addition to the large 15-node misalignment cycle on the left half of the node-link diagram, there are also correct alignments and small cycles of length two and three.

**Figure 7.**
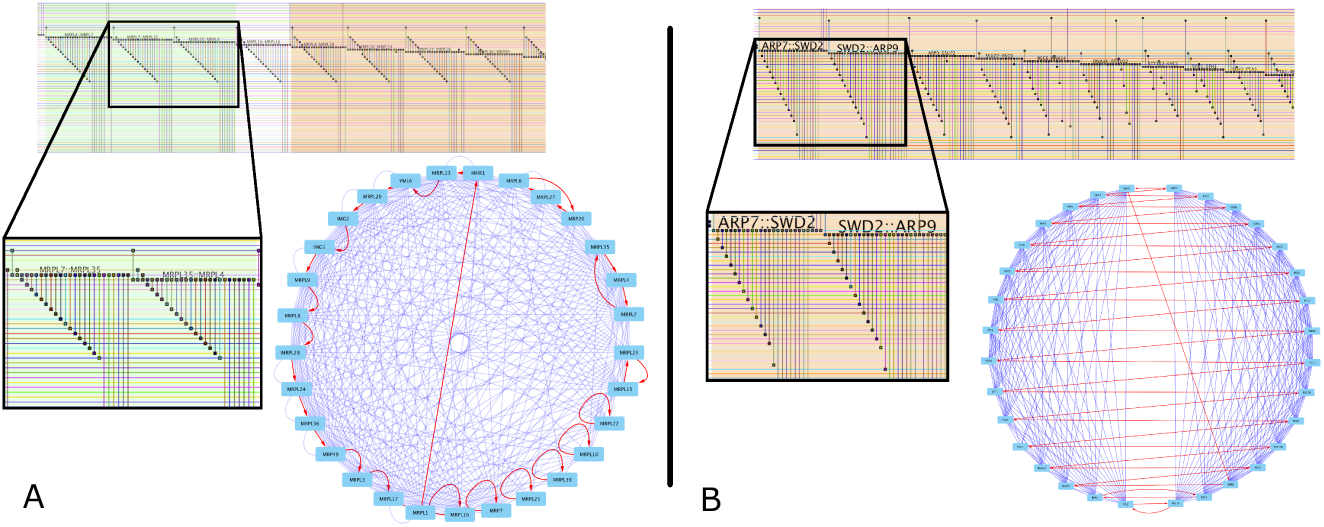
BioFabric allows the user to spot major alignment problems; in this case, we can see how two entire protein complexes were misaligned. In section **A**, left, we show proteins of the mitochondrial ribosomal large subunit; the traditional node-link diagram for this complex is shown with protein-protein interaction edges (colored blue) and alignments shown as directed edges (colored red). The misalignments (i.e. alignment edges that are not self loops) are to be expected, given the high topological similarity of the nodes, but nodes in this cluster are aligned with other nodes in the same cluster. Note in the BioFabric view (top left, with detail lower left), the repeating “edge wedge” pattern with the 45° angle is the canonical representation for a clique. The ordering of the nodes using alignment cycles has no visible effect when node misalignments are intra-cluster. In section **B**, compare the shape of the “edge wedges” in this BioFabric view to the one in **A**; note the wedges have a 60° (not 45°) angle and the edges for each node are incident on *alternating* and *disjoint* sets of node rows in a context where misaligned nodes are in adjacent rows. This is an unmistakable visual cue that there has been a misalignment between two completely distinct protein complexes. Note how in the node-link diagram (lower right) all the red node alignment links cross back and forth between the two sides. The left side has proteins of the CPF cleavage and polyadenylation factor, while the right side has components of the RSC chromatin remodeling complex.

This is clearly a single protein cluster; in fact, it is some of the protein components of the mitochondrial ribosomal protein of the large subunit. (Note that all characterizations of proteins in this paper were obtained from the SGD [23].) While not fully connected, it does approximate a clique, so the misalignments found here are unsurprising, given the objective function uses topology alone.

Crucially, we can instantly interpret the context of the misalignment with the BioFabric version of the network: the repeating pattern of “edge wedges” is the telltale sign in BioFabric of a clique when the nodes in the clique are laid out contiguously. Remembering that the order of the node rows has been determined using the order of the alignment cycles, we see that in this case the node order still maintains the typical clique pattern; thus the misalignments are actually *contained within the clique*. Since nodes within a clique are topologically indistinguishable, their misalignment is unavoidable (unless more information such as gene sequence data is used in the alignment objective function), and should not be penalized for an alignment which is driven by topology alone.

A more profound alignment problem is shown in Figure 7B. This misalignment involves 32 nodes in three alignment cycles. As the red edges in the node-link diagram shows, the alignment has incorrectly aligned one *entire* protein complex with a completely different complex! On the left are proteins of the CPF cleavage and polyadenylation factor, while on the right are the components of the RSC chromatin remodeling complex.

This problem stands out in the BioFabric version, where the user typically studies and compares the “edge wedge” shapes for nodes to better understand the network structure. While a typical clique pattern (as shown in Figure 7A) has edge wedges with a 45° lower margin, these wedges have a steeper, 60° lower margin; the edges for each node are actually incident on *alternating* nodes in this run. The explanation of this pattern is that the nodes belonging to the two separate protein clusters are intercalated in this run of nodes that have been laid out using the alignment cycles. Thus, our Alignment Cycle layout technique makes this problem stand out just by looking at it.

Even if one would argue that the traditional node-link diagram shows these effects well, the views shown had to be meticulously hand-crafted to illustrate the structure once we knew what we were looking at, while the BioFabric representations were automatically laid out. As we show in the Appendix, even a complex misalignment involving a cycle of *four* protein clusters can be quickly spotted.

## Discussion

Given the difficulty of viewing the quality and characteristics of network alignments using traditional node-link diagrams, it is unsurprising that researchers have concentrated on evaluating alignments by comparing and optimizing certain numerical metrics. But to be able to gain some broad intuition about network alignments, having effective and well-organized visualization techniques can provide huge benefits in understanding the problems.

While still providing the deep intuition of the node-link approach (albeit with the nodes depicted as lines), BioFabric additionally give the user the ability to group and order *both* nodes and edges in meaningful ways. This gives researchers a new way to tackle the problem of understanding network alignments. The organized grouping of *edges* that is made possible by nodes-as-lines is particularly unique. As we have shown, simple visual estimates of proportions of classes of link groups provides intuition into abstract measures like *S*^3^. Furthermore, as our Alignment Cycle layout demonstrates, it is possible to create new, specialized layout algorithms that exploit the linear ordering of nodes in ways that make it possible to quickly spot alignment pathologies.

Because of these features, VISNAB is a viable tool for studying objective functions and their performance. As discussed in [16], we strongly contend the network alignment community should focus not on developing new network alignment algorithms, but rather on devising more effective objective functions. Just as we experimented with combining *S*^3^ and Importance, researchers should design custom objective functions for their specific needs to yield optimum results. To accomplish this task, we propose VISNAB as a tool researchers should utilize to help design objective functions.

For example, as we have shown in Case Study III, researchers can work towards creating objective functions that distribute edges and links into appropriate proportions of our identified groupings. As was shown in Case Study I, the sizes of these groupings can reflect the character of the alignment problem at hand, e.g. a pair of experiments where the larger network represented a deeper, more thorough investigation of the system, where new elements were added, and false positives were identified. So an objective function that worked to take this topology into account would be an improvement. Note that this approach can be useful *even in the absence of a gold standard alignment* for a given problem, as the overall proportions of the groupings could arguably apply to other alignments in the same class of problems. In other words, target vectors for the *NGS* and *LGS* scores could be estimated a priori for a class of problems.

### Limitations

BioFabric’s presentation of “nodes as lines” is unfamiliar for network researchers who have grown accustomed to visualizing nodes as points and can take some time getting used to. For networks small enough, the traditional node-link diagram approach is adequate, so it would be beneficial to allow the user to select a subset of the BioFabric network and view it using the traditional presentation. Going the other way, taking a subset of a traditional node-link presentation and viewing it using BioFabric, would be valuable as well. But this capability is not currently available.

Another shortcoming of the current VISNAB implementation is that the ordering of the node groups is fixed, per the order in Table 2. A different order may be more helpful in visualizing a particular alignment, so being able to dynamically reorder the node groups would be a useful feature.

Finally, just like the existing *NC* score, our new *NGS, LGS*, and *JS* scores depend upon the availability of a gold-standard alignment. Thus, they can really only be used to evaluate the performance of alignment algorithms and objective functions when the correct answer is already known. To be incorporated into a useful objective function, it would be necessary that for certain classes of alignment problems, target vectors for the *NGS* and *LGS* scores could be estimated a priori. It is also an open question about the difficulties of developing objective functions based upon incremental evaluation for these measures.

### Future Work

An emerging area of investigation is the multiple network alignment problem, e.g. [24]. In these types of alignments, the fraction of the full set of networks that share a particular edge or node in the alignment is a crucial piece of information. This suggests that creating node and link groups for grouping elements with similar overlap percentages would be helpful in visualizing the quality and characteristics of a multiple network alignment.

Our new tool provides a platform to implement other new visualizations for exploring network alignments. Constructing and visualizing subgraphs of the full merged network (from **Network Merge**, above) can provide valuable insights as well. For example, one additional view that we support is the Orphan Edge layout, which shows the subgraph of *G*_1_ that consists all the nodes connected by the orphan blue **pBp** edges, plus the first degree neighbors of those nodes and corresponding connecting edges. In aligned networks where *E*_1_ ⊆ *E*_2_ (e.g. the *noisy yeast* network used in Case Study IV), blue edges will not exist at all in a correctly aligned network. Thus, viewing the context of the orphan blue **pBp** edges in the network can provide insights into alignment problems.

Similarly, we can visualize the common subgraph *CS*_*a*_ = (*V*_1_, *E*_*a*_) of the alignment by displaying only the aligned purple **P** edges and the purple nodes they are incident upon. If this subgraph’s nodes and edges are laid out using BioFabric’s default layout [7], the various connected components of *CS*_*a*_ can be assessed. One way some researchers evaluate alignments is to determine if the alignment has a common subgraph with large connected regions [4, 15]. To this end, the user can visually estimate the components of the measure **Largest Common Connected Subgraph** (LCCS) using the heights of the nodes and widths of the edges of the connected components.

## Conclusions

VISNAB provides a novel, powerful new way to visualize network alignments, and will allow researchers to gain a new and better understanding of the strengths and shortcomings of the many available network alignment algorithms.

## Supporting information

Case Study Files

VISNAB Source Code

BioFabric Source Code

## Appendix

### Link and Node Groups With Blue Nodes

In the main text, for simplicity, we limited the discussion to the common case of aligning of network *G*_1_ onto network *G*_2_ when every node in *G*_1_ is aligned onto a node in *G*_2_. In the nomenclature we have introduced, this is an alignment “without any blue nodes”. Table 1 and Table 2 enumerated the possible link and node groups for these alignments. However, VISNAB is capable of handling alignments where unaligned blue nodes are permitted. In that case, the five link groups expand to seven, adding in groups three and four, which account for the case where blue edges are incident on blue nodes. Table 4 enumerates all possible link groups in the presence of blue nodes in the alignment.

**Table 4.** Expansion of Link Groups When Unaligned (Blue) Nodes are Present.

| Link Group | Edge Color | Endpoint 1 | Endpoint 2 | Symbol |
| --- | --- | --- | --- | --- |
| 1 | Purple | Purple | Purple | P |
| 2 | Blue | Purple | Purple | pBp |
| 3 | Blue | Purple | Blue | pBb |
| 4 | Blue | Blue | Blue | bBb |
| 5 | Red | Purple | Purple | pRp |
| 6 | Red | Purple | Red | pRr |
| 7 | Red | Red | Red | rRr |

**Table 5.** Expansion of Node Groups When Unaligned (Blue) Nodes are Present.

| Node Group | Node Color | Incident Edges | Symbol |
| --- | --- | --- | --- |
| 1 | Purple | None | (P:0) |
| 2 | Purple | Purple | (P:P) |
| 3 | Purple | pBp | (P:pBp) |
| 4 | Purple | pBb | (P:pBb) |
| 5 | Purple | pBp, pBb | (P:pBp/pBb) |
| 6 | Purple | pRp | (P:pRp) |
| 7 | Purple | Purple, pBp | (P:P/pBp) |
| 8 | Purple | Purple, pBb | (P:P/pBb) |
| 9 | Purple | Purple, pBp, pBb | (P:P/pBp/pBb) |
| 10 | Purple | Purple, pRp | (P:P/pRp) |
| 11 | Purple | pBp, pRp | (P:pBp/pRp) |
| 12 | Purple | pBb, pRp | (P:pBb/pRp) |
| 13 | Purple | pBp, pBb, pRp | (P:pBp/pBb/pRp) |
| 14 | Purple | Purple, pBp, pRp | (P:P/pBp/pRp) |
| 15 | Purple | Purple, pBb, pRp | (P:P/pBb/pRp) |
| 16 | Purple | Purple, pBp, pBb, pRp | (P:P/pBp/pBb/pRp) |
| 17 | Purple | pRr | (P:pRr) |
| 18 | Purple | Purple, pRr | (P:P/pRr) |
| 19 | Purple | pBp, pRr | (P:pBp/pRr) |
| 20 | Purple | pBb, pRr | (P:pBb/pRr) |
| 21 | Purple | pBp, pBb, pRr | (P:pBp/pBb/pRr) |
| 22 | Purple | pRp, pRr | (P:pRp/pRr) |
| 23 | Purple | Purple, pBp, pRr | (P:P/pBp/pRr) |
| 24 | Purple | Purple, pBb, pRr | (P:P/pBb/pRr) |
| 25 | Purple | Purple, pBp, pBb, pRr | (P:P/pBp/pBb/pRr) |
| 26 | Purple | Purple, pRp, pRr | (P:P/pRp/pRr) |
| 27 | Purple | pBp, pRp, pRr | (P:pBp/pRp/pRr) |
| 28 | Purple | pBb, pRp, pRr | (P:pBb/pRp/pRr) |
| 29 | Purple | Blue, pRp, pRr | (P:pBp/pBb/pRp/pRr) |
| 30 | Purple | Purple, pBp, pRp, pRr | (P:P/pBp/pRp/pRr) |
| 31 | Purple | Purple, pBb, pRp, pRr | (P:P/pBb/pRp/pRr) |
| 32 | Purple | Purple, pBp, pBb, pRp, pRr | (P:P/pBp/pBb/pRp/pRr) |
| 33 | Blue | pBb | (B:pBb) |
| 34 | Blue | bBb | (B:bBb) |
| 35 | Blue | pBb, bBb | (B:pBb/bBb) |
| 36 | Blue | None | (B:0) |
| 37 | Red | pRr | (R:pRr) |
| 38 | Red | rRr | (R:rRr) |
| 39 | Red | pRr, rRr | (R:pRr/rRr) |
| 40 | Red | None | (R:0) |

In a similar fashion, the twenty node groups that can be present in an alignment without blue nodes expands to forty possible groups when blue nodes are allowed. Note how, for example, Node Group 3 (see Table 2) splits into three distinct groups (3, 4, and 5) with blue nodes, since blue edges can be incident on blue nodes as well as purple nodes. In a similar fashion, groups 7, 11, 14, 19, 23, 27, and 30 all split as well into three distinct groups. Groups 33 to 36 are introduced as well to account for blue nodes being present.

### Jaccard Similarity With Blue Nodes

Again, for simplicity, the main text only discussed the definition of our Jaccard Similarity (*JS*) score when there are no blue nodes present in the alignment. When blue nodes are not allowed, then for every node in *G*_1_, we can find (using the correct alignment) where that node is supposed to go in *G*_2_, and (using the given alignment) where it actually ends up in *G*_2_. These two nodes in *G*_2_ can then be compared to create the *JS* score for the node.

However, when blue nodes are allowed, there are four possible cases that can arise instead of one:

1. The node is supposed to be aligned, and it is (the case described above) (“purple node stays purple”)
2. The node is supposed to be aligned, and it is not (“purple node turns to blue”)
3. The node is not supposed to be aligned, and it is (“blue node turn to purple”)
4. The node is not supposed to be aligned, and it is not (“blue node stays blue”)

VISNAB handles case 4 by simply assigning a score of 1.0 to the node, since it is correctly left unaligned. To deal with cases 2 and 3, VISNAB instead compares two nodes in network *G*_1_. Specifically, for case 2, if a node a in *G*_1_ is supposed to (using the correct alignment) be aligned to node n in *G*_2_, but is instead unaligned, we look to see which node b in *G*_1_ is aligned (using the given alignment) to node n in *G*_2_. We then create the *JS* score for node a by comparing the neighborhoods of a and b in *G*_1_. If there is no node b (when nothing is aligned to node n in *G*_2_, i.e. it is “red”), then the *JS* score for node a is 0.0. Case 3 is handled analogously, again comparing two nodes a and b in *G*_1_ to obtain a *JS* score, with a 0.0 assigned if there is no matching node in *G*_1_.

For some network *G* = (*V, E*), let *N*_*G*_(*z*_1_) = {*z*_2_ ∈ *V* : (*z*_1_, *z*_2_) ∈ *E*} be the neighborhood of node *z*_1_ in *G*. For nodes *x, y* ∈ *V*, let *N*_*G*_(*x, y*) be the neighborhood of *x* disregarding *y*, and let *i*_*xy*_ be a corrective term accounting for a possible edge between the two. Accordingly, if *y* ∈ *N*_*G*_(*x*) then *N*_*G*_(*x, y*) = *N*_*G*_(*x*) − *y* and *i*_*xy*_ = 1, else *N*_*G*_(*x, y*) = *N*_*G*_(*x*) and *i*_*xy*_ = 0. Let *N*_*G*_(*y, x*) be defined analogously. Our extended *JS* definition *σ*_*G*_ : *V* × *V* → [0, 1] between two nodes is defined as:

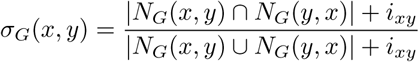

Note that when *x* and *y* are both singletons, we define *σ*(*x, y*) = 1.0 to avoid dividing by zero.

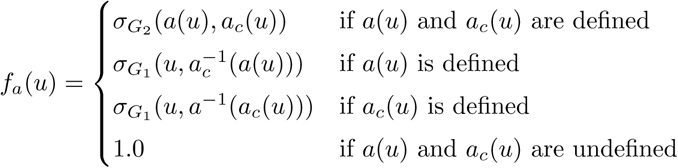

Given node sets *V*_*a*_, *V*_*c*_ ∈ *V*_1_, an alignment *a* : *V*_*a*_ → *V*_2_, and the correct alignment *a*_*c*_ : *V*_*c*_ → *V*_2_, our *JS* measure for the given alignment *a*, with respect to the correct alignment *a*_*c*_, is defined as:

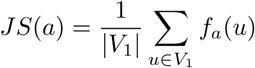

If the correct alignment is provided, the user can choose to have correctly and incorrectly aligned nodes laid out separately in different node groups. The user can choose the criterion for the correct alignment to be based either on the traditional *NC* measure, or on our *JS* measure. If *JS* is chosen, the user can set the threshold value *β* ∈ [0, 1], so, a node in the form of *u*::*v* or *u*:: is denoted correct if *f*_*a*_(*u*) ≥ *β*.

### Creation of the Correct Network Alignment

To create the “correct” alignment used in the case studies, we wanted to create two networks where all nodes in the smaller network had one and only one known matching node in the larger network. One consequence of this approach is that our correct alignment did not have any blue nodes. These case studies use two different protein-protein interaction datasets. The larger “SC” network, from *S*. *cerevisiae*, contains 5,831 nodes and 77,149 edges, and was originally obtained from BioGRID (v3.2.101, June 2013) [14]. The smaller “Yeast2” network, also from *S*. *cerevisiae*, has 2,390 nodes and 16,127 edges. It was originally generated from data in [13] and used in [4]. Both networks were previously used in [16].

Nodes in Yeast2 are tagged with a variety of gene symbols (e.g. *PSY4*), secondary identifiers, and synonyms, while nodes in SC were tagged with ENTREZ IDs (e.g. 852234). In order to generate the “correct” alignment file, it was necessary to find the mapping from the former to the latter. To do this, we first used the YeastMine API [25, 26] at https://yeastmine.yeastgenome.org/, provided by the Saccharomyces Genome Database (SGD) [23], in order to generate a mapping from the node names to the SGD IDs that we could then feed to the DAVID web tool [27, 28]. With a Java program employing libraries provided by `org.intermine`, we downloaded (11 Feb. 2018) tuples for `Gene.primaryIdentifier`, `Gene.secondaryIdentifier`, `Gene.symbol`, and `Gene.synonyms.value` for `Gene.organism.shortName=‘‘S. cerevisiae’’`, for all entries in the lists `Verified ORFs`, `Dubious ORFs` and `ALL Verified Uncharacterized Dubious ORFs`. Three remaining genes *YAR010C, YBR012W-B*, and *YHL009W-B* were not in any of these lists and were explicitly queried.

For each gene in Yeast2, we then matched the node name to a `Gene.synonyms.value`, and from this obtained a list of one or more `Gene.primaryIdentifiers`. In the cases where there was more than one, we chose the `Gene.primary-Identifier` that mapped to a Gene.symbol that matched the `Gene.synonyms.value`. For example, synonym *MSL1* mapped to SGD IDs S000004374 and S000001448. However, while the former SGD ID mapped to gene symbol *NAM2*, the latter mapped to *MSL1*, and thus was selected. With one exception, this approach resulted in an unambiguous mapping of all Yeast2 node names to SGD IDs. The exception was for gene names *EFG1* and *YGR272C*; the latter was merged into the former, giving both names the same SGD ID (S000007608). Thus, node *YGR272C* was dropped (see Table 6)

**Table 6.** Node that was merged, and dropped

| Gene Name | SGD ID |
| --- | --- |
| YGR272C | S000007608 |

These SGD IDs were then uploaded as a gene list to DAVID at:

https://david.ncifcrf.gov/conversion.jsp

(DAVID 6.8, accessed 18 Feb. 2018). Since DAVID has restrictions on large-scale queries through their web API, this was done manually. Upon uploading the list, DAVID’s Gene List Manager was not able to identify five IDs, so these nodes were dropped as well (see Table 7).

**Table 7.**
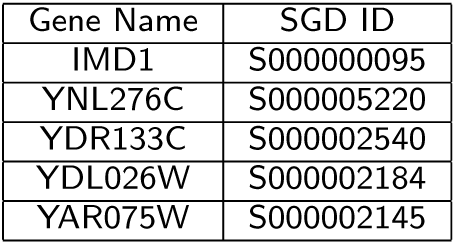
Nodes not identified by DAVID, and dropped

We instructed the tool to convert SGD IDs to ENTREZ GENE IDs, and downloaded the result. Thus, we had a mapping of 2,384 of the nodes in Yeast2 to ENTREZ IDs. However, not all of these ENTREZ IDs are present as nodes in the larger SC network. In order to create a correct alignment with no blue nodes, we then pruned the Yeast2 network to remove the small number of nodes that could not be mapped onto the SC network. This resulted in an additional five nodes that needed to be dropped; see Table 8.

**Table 8.** Nodes that could not be mapped to SC network, and dropped

| Gene Name | SGD ID |
| --- | --- |
| ATM1 | 855347 |
| PHM8 | 856759 |
| PUT1 | 850833 |
| CTM1 | 856509 |
| SBE2 | 851953 |

### Detailed Description of the Node Assignment Algorithm for the Node and Link Group Layout

The node assignment algorithm for the Node and Link Group Layout is a multi-queue breadth first search graph traversal. While a typical breadth first search utilizes a single queue, our multi-queue approach uses one queue for each node group, and the queues are processed in the order listed in Table 2. The traversal starts on the node of highest degree in the first queue; its neighbor nodes are then visited in order of decreasing degree. If a newly visited node is in the current node group, it will be placed onto the current queue; if it is not, it will be placed onto the queue of its node group. The traversal is finished with a queue when every node in that node group has been visited. If the queue is empty but there still are unvisited nodes in that group, the highest degree node from the set of unvisited nodes of that group is added to the queue; after the queue is traversed, if there still are unvisited nodes in the group, this step is repeated until all nodes in the group are visited. Once finished, the traversal moves to the next queue. If a queue is empty when first evaluated, the node of highest degree in that queue’s node group is added.

Thus, we created a “Yeast2-reduced” network consisting of 2,379 nodes and 16,063 edges, which was used in the case studies.

### Full Table of All Alignment Scores for Mixtures of Importance and Symmetric Substructure Score

Table 9 lists the scores for the ten-hour SANA [16] runs between Yeast2K-Reduced and SC, in which we used combinations of Importance (*I*) [17] and Symmetric Substructure Score (*S*^3^) [15] in the objective function. Note that all these scores, with the exception of Resnik, are available using the Alignment Measures tool in VISNAB. The Resnik scores [18, 19, 20] shown here are the means of the non-zero, non-”None” values computed separately using FastSemSim [21], incorporating Gene Ontology (GO) terms [29, 30] downloaded in February 2019.

**Table 9.** All Alignment Scores for Case Study III

| Alignment | NGS | LGS | NC | JS | $S^3$ | Resnik |
| --- | --- | --- | --- | --- | --- | --- |
| Correct | 1.00 | 1.00 | 1.00 | 1.00 | 0.25 | 9.63 |
| $1.0 * I$ | 0.61 | 0.79 | 0.00042 | 0.021 | 0.0043 | 3.16 |
| $.001 * S^3 + .999 * I$ | 0.88 | 0.86 | 0.00042 | 0.024 | 0.17 | 3.48 |
| $.003 * S^3 + .997 * I$ | 0.88 | 0.86 | 0.00042 | 0.021 | 0.18 | 3.39 |
| $.005 * S^3 + .995 * I$ | 0.72 | 0.85 | 0.00 | 0.025 | 0.10 | 3.27 |
| $.01 * S^3 + .99 * I$ | 0.88 | 0.86 | 0.00 | 0.024 | 0.19 | 3.32 |
| $.03 * S^3 + .97 * I$ | 0.87 | 0.80 | 0.018 | 0.057 | 0.27 | 3.62 |
| $.05 * S^3 + .95 * I$ | 0.73 | 0.53 | 0.022 | 0.063 | 0.49 | 3.44 |
| $.1 * S^3 + .9 * I$ | 0.67 | 0.48 | 0.017 | 0.067 | 0.54 | 3.50 |
| $1.0 * S^3$ | 0.64 | 0.46 | 0.021 | 0.069 | 0.55 | 3.61 |

### Table of Node Group Sizes for Case III

Table 10 provides the number of nodes in each node group for the four alignments discussed in Case III. Asterisks show the four largest node groups per alignment, which are labeled prominently in Figure 5. As called out in the text, of the 716 nodes in the top 13 rows above group **(P:P/pRp/pRr)** for the mixed alignment, 544 (76%) have no incident **pRr** edges.

### Percentage of Purple Nodes Without and With Incident **pRr** Edges between Correct and Mixed Alignments

The discussion of Figure 5 notes that while there are *more* **pRr** edges in the mixed alignment compared to the correct alignment, those edges are concentrated across a *smaller fraction* of the purple nodes in that mixed alignment. Table 11 compares the percentages of all purple nodes without [**(P:*)**] and with [**(P:*/pRr)**] **pRr** incident edges, between the correct and mixed alignments.

**Table 10.** Sizes of all Node Groups in Case III

| Symbol | Correct | All Importance | Mixed | All $S^3$ |
| --- | --- | --- | --- | --- |
| (P:0) | 0 | 0 | 0 | 0 |
| (P:P) | 2 | 0 | 23 | 16 |
| (P:pBp) | 0 | 0 | 0 | 0 |
| (P:pRp) | 0 | 0 | 0 | 0 |
| (P:P/pBp) | 2 | 0 | 59 | 23 |
| (P:P/pRp) | 52 | 0 | 62 | 3 |
| (P:pBp/pRp) | 32 | 403* | 207 | 0 |
| (P:P/pBp/pRp) | 27 | 9 | 193 | 1 |
| (P:pRr) | 0 | 0 | 0 | 0 |
| (P:P/pRr) | 5 | 0 | 31 | 439 |
| (P:pBp/pRr) | 5 | 35 | 98 | 303 |
| (P:pRp/pRr) | 0 | 0 | 0 | 0 |
| (P:P/pBp/pRr) | 1 | 9 | 43 | 662* |
| (P:P/pRp/pRr) | 981* | 0 | 530* | 157 |
| (P:pBp/pRp/pRr) | 310 | 1677* | 82 | 0 |
| (P:P/pBp/pRp/pRr) | 962* | 246 | 1051* | 775* |
| (R:pRr) | 578* | 843* | 752 * | 31 |
| (R:rRr) | 209 | 58 | 0 | 1087* |
| (R:pRr/rRr) | 2665* | 2551* | 2700* | 2334* |
| (R:0) | 0 | 0 | 0 | 0 |

**Table 11.** Comparison of Purple Node Fractions Without and With **pRr** Edges

| Alignment | (P:*) | (P:*/pRr) |
| --- | --- | --- |
| Correct Alignment | 4.83% | 95.17% |
| Mixed Alignment | 22.87% | 77.13% |

### Alignment Cycle Layout With Blue Nodes

When unaligned blue nodes are not allowed, there are four cases that must be handled by the Alignment Cycle layout, and these are indicated by a checkmark in the rightmost column of Table 12. When unaligned blue nodes are present, there are nine path types that must be handled. In the table, a network with nodes {*A, B, C*, … *L*} has been aligned onto a network with nodes {1, 2, 3, … 12}. Note that purple node runs can extend for any length of nodes, as shown by the …, but the matches and alignments given in this table are for the cases where there are none of these extra nodes. The Alignment Cycle layout will order the nodes in the path and cycle cases so that misaligned nodes are laid out next to their correct partners; see case 9 in particular to see this pattern.

**Table 12.**
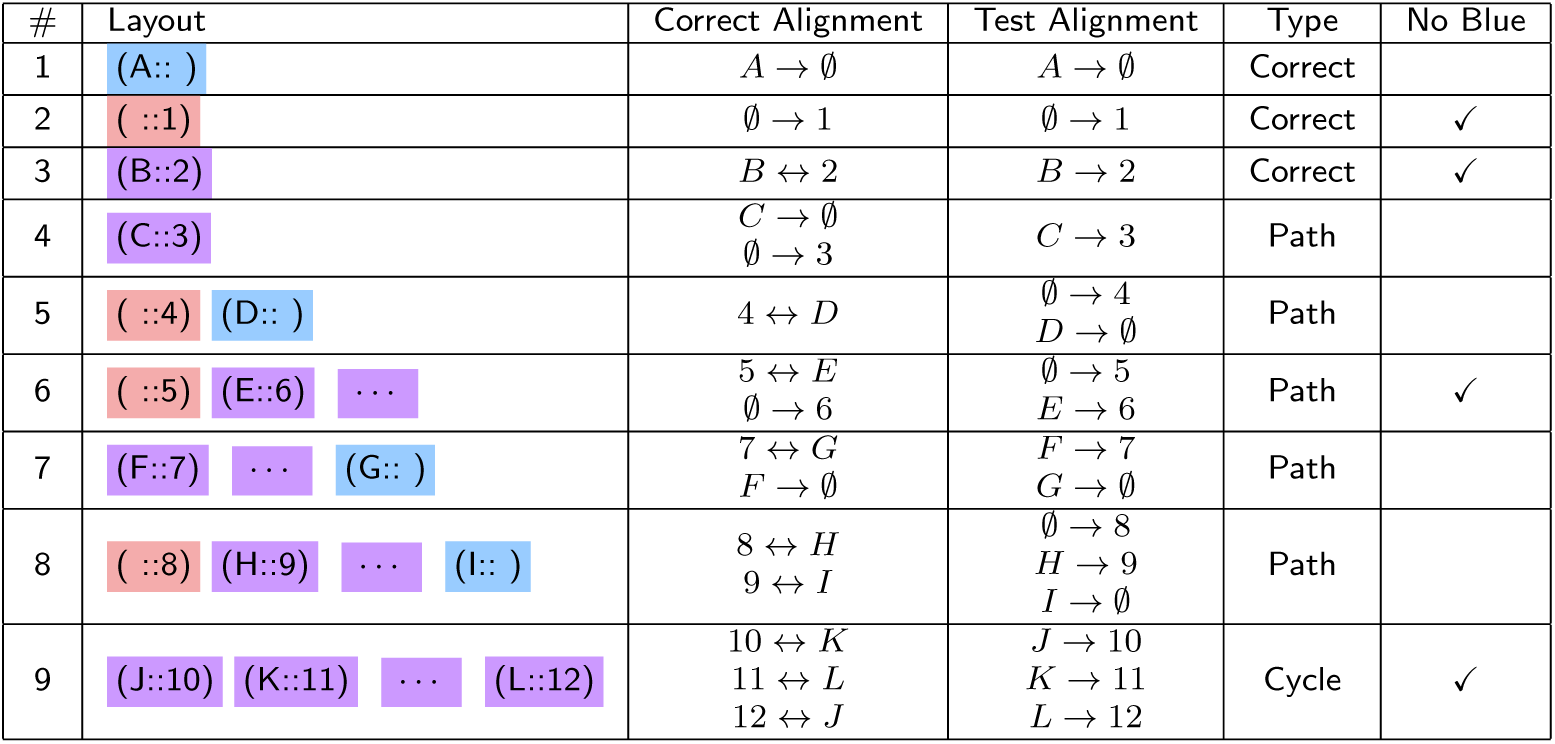
Alignment Cycle Layout Cases

### The Four-Cluster Misalignment

In Case Study IV, we showed how the Alignment Cycle layout could be used to spot alignment problems such as two entire protein clusters being swapped. Figure 8 shows a severe degeneracy for the same alignment run, where *four* separate protein clusters were misaligned in a cycle. The BioFabric depiction of this problem follows the same pattern shown in Figure 7, but the successive edge wedges are even steeper here, and show a clear pattern of cycling between four distinct sets of node rows.

**Figure 8.**
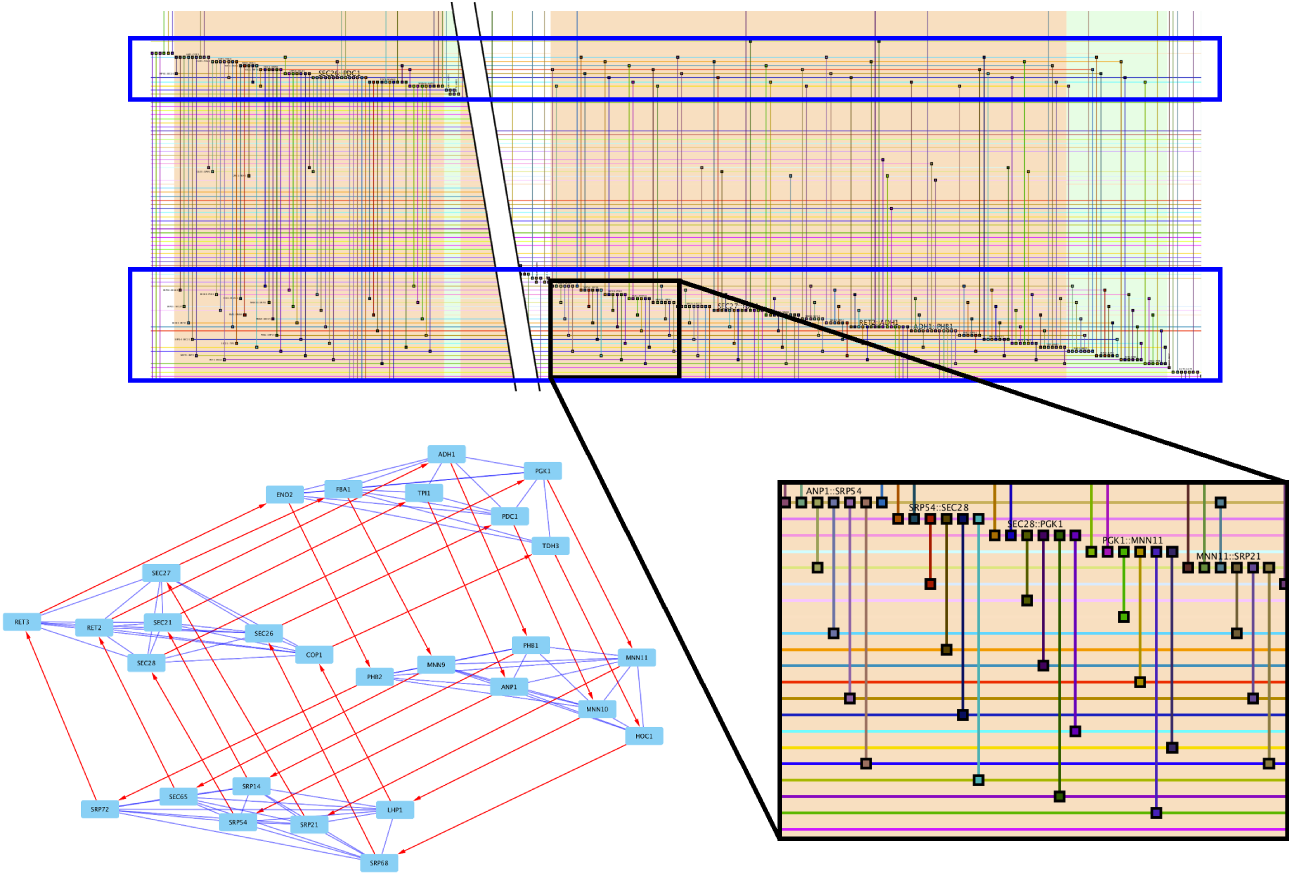
An even more striking misalignment, where four different protein complexes have been swapped in a round-robin fashion. The traditional node-link diagram is shown at the lower left with edges (colored blue) for the protein-protein interactions and directed edges (colored red) for the alignments. The four protein complexes clockwise from top (per SGD): 1) glycolysis and gluconeogenesis related genes, 2) mannosyltransferase complex and prohibitin complex, 3) signal recognition particle, and 4) the coatomer complex (COPI). The BioFabric layout on the top, shown in detail at the lower right, shows the distinct pattern displayed by this artifact, with adjacent edge wedges having edges cycling every fourth node. Note that a slice was removed from the upper view because the three separate cycles constituting this structure are not contiguous in the layout.

### Availability and requirements

**Project Name:** VISNAB: Visualization of Network Alignments using BioFabric

**Project Home Page** http://www.BioFabric.org/VISNAB/index.html. The VISNAB code repository is at https://github.com/wjrl/AlignmentPlugin, while the BioFabric code repository is at https://github.com/wjrl/BioFabric.

**Operating Systems** Cross-platform.

**Programming Language** Java

**Other Requirements** Plugin requires BioFabric Version 2 Beta Release 2 or above, available on GitHub and the BioFabric project Home Page. An OpenJDK Java runtime is bundled with BioFabric and does not need to be installed separately.

**License** LGPL V 2.1. Per the LGPL license, the VISNAB source code is provided in additional file 2.

## List of Abbreviations

EC: Edge Coverage, Conservation, Correctness, or Correspondence
ICS: Induced Conserved Structure
JS: Jaccard Similarity
LCCS: Largest Common Connected Subgraph
LGS: Link Group Similarity
NC: Node Correctness
NGS: Node Group Similarity
PPI: protein-protein interaction
(*S*^3^): Symmetric Substructure Score
SGD: Saccharomyces Genome Database
VISNAB: Visualization of Network Alignments using BioFabric

## Additional Files

Archived executables are available at: https://www.ics.uci.edu/~wayne/papers/BioFabricVISNAB/

Additional file 1: DesaiEtAl-2019-CaseStudyFiles.zip

Archive of .sif, .gw, .align, .resnik, and .bif (BioFabric) files for the four case studies. The latter can be loaded into BioFabric with VISNAB to view the case study results in detail.

Additional file 2: VISNAB-1.1.0.0-src.tar.gz

Contains the source code for the VISNAB V1.1 plugin. The most recent source code is available at the GitHub repository listed above.

Additional file 3: BioFabric-2.0.B.2-src-tar.gz

Contains the source code for the beta version of BioFabric 2.0 needed to run the plugin. The most recent recent source code is available at the GitHub repository listed above.

