## Supplementary material for "BioFabric Visualization of Network Alignments": Case Study Files: README-Manifest.txt

File Manifest for DesaiEtAl-2019-Case-Study-Results.zip

Accompanies R.M. Desai, W.J.R. Longabaugh, and W.B. Hayes. BioFabric Visualization of Network Alignments, 2019.

CaseStudies-I-II-III-Perfect.align : Perfect alignment file used in Case Studies I, II, and III.
CaseStudies-I-II-III-SC.sif : SIF file for S. cerevisiae “SC” network, for Case Studies I, II, and III.
CaseStudies-I-II-III-Yeast2KReduced.sif : SIF file for “Yeast2-reduced” network, for Case Studies I, II, and III.
CaseStudy-III: Directory containing alignment files, BioFabric BIF files, and Resnik results for all the objective function combinations in Case Study III. 
CaseStudy-IV-Alignment.align : The alignment file used in Case Study IV.
CaseStudy-IV-LargeYeast.gw : The larger yeast network used in Case Study IV.
CaseStudy-IV-SmallYeast.gw : The smaller yeast network used in Case Study IV.
CaseStudy-IV.bif : The BioFabric BIF file for in Case Study IV.
README-Manifest : This file.
