## Supplementary material for "BioFabric Visualization of Network Alignments": VISNAB Source Code: AlignCycleEdgeLayout.java

```
/*
**    Copyright (C) 2003-2018 Institute for Systems Biology 
**                            Seattle, Washington, USA. 
**
**    This library is free software; you can redistribute it and/or
**    modify it under the terms of the GNU Lesser General Public
**    License as published by the Free Software Foundation; either
**    version 2.1 of the License, or (at your option) any later version.
**
**    This library is distributed in the hope that it will be useful,
**    but WITHOUT ANY WARRANTY; without even the implied warranty of
**    MERCHANTABILITY or FITNESS FOR A PARTICULAR PURPOSE.  See the GNU
**    Lesser General Public License for more details.
**
**    You should have received a copy of the GNU Lesser General Public
**    License along with this library; if not, write to the Free Software
**    Foundation, Inc., 59 Temple Place, Suite 330, Boston, MA  02111-1307  USA
*/

package org.systemsbiology.biofabric.plugin.core.align;

import java.util.ArrayList;
import java.util.HashMap;
import java.util.HashSet;
import java.util.List;
import java.util.Map;
import java.util.TreeMap;

import org.systemsbiology.biofabric.api.io.BuildData;
import org.systemsbiology.biofabric.api.layout.DefaultEdgeLayout;
import org.systemsbiology.biofabric.api.model.AnnotationSet;
import org.systemsbiology.biofabric.api.model.NetLink;
import org.systemsbiology.biofabric.api.model.NetNode;
import org.systemsbiology.biofabric.api.model.Network;
import org.systemsbiology.biofabric.api.worker.AsynchExitRequestException;
import org.systemsbiology.biofabric.api.worker.BTProgressMonitor;
import org.systemsbiology.biofabric.api.worker.LoopReporter;
import org.systemsbiology.biofabric.plugin.PluginSupportFactory;


/****************************************************************************
**
** Like the default, but also calculates cycle bounds
*/

public class AlignCycleEdgeLayout extends DefaultEdgeLayout {
  
  ////////////////////////////////////////////////////////////////////////////
  //
  // PRIVATE INSTANCE MEMBERS
  //
  ////////////////////////////////////////////////////////////////////////////

  ////////////////////////////////////////////////////////////////////////////
  //
  // PUBLIC CONSTRUCTORS
  //
  ////////////////////////////////////////////////////////////////////////////

  /***************************************************************************
  **
  ** Constructor
  */

  public AlignCycleEdgeLayout() {
  }

  ////////////////////////////////////////////////////////////////////////////
  //
  // PUBLIC METHODS
  //
  ////////////////////////////////////////////////////////////////////////////
  
  /***************************************************************************
  **
  ** Do necessary pre-processing steps (e.g. automatic assignment to link groups)
  */
  
  @Override
  public void preProcessEdges(BuildData rbd, 
                              BTProgressMonitor monitor) throws AsynchExitRequestException {
    
    List<String> groupOrder = new ArrayList<String>();
    for (NetworkAlignment.EdgeType type : NetworkAlignment.LINK_GROUPS) {
      groupOrder.add(type.tag);
    }
    rbd.setGroupOrderAndMode(groupOrder, Network.LayoutMode.PER_NODE_MODE, true);
    return;
  } 
  
  /***************************************
  **
  ** Calc link annotations
  */ 
  
  @Override
  protected AnnotationSet calcGroupLinkAnnots(BuildData rbd, 
                                              List<NetLink> links, BTProgressMonitor monitor, 
                                              boolean shadow, List<String> linkGroups) throws AsynchExitRequestException { 
    
  	
  	NetworkAlignmentBuildData narbd = (NetworkAlignmentBuildData)rbd.getPluginBuildData();
  	if (narbd.useNodeGroups) {
  		return (PluginSupportFactory.buildAnnotationSet());
  	}
  	
  	//
  	// If shadows links are not shown, we are not going to do link groups for this, since we need every node
  	// to appear on the diagonal
  	//
  	
  	if (!shadow) {
  		return (PluginSupportFactory.buildAnnotationSet());
  	}

    TreeMap<Integer, NetNode> invert = new TreeMap<Integer, NetNode>();
    for (NetNode node : rbd.getNodeOrder().keySet()) {
      invert.put(rbd.getNodeOrder().get(node), node);
    }
    ArrayList<NetNode> order = new ArrayList<NetNode>(invert.values());
    return (calcGroupLinkAnnotsCycle(links, order, monitor, 
                                     shadow, narbd.cycleBounds, linkGroups));
  }

  /***************************************
  **
  ** Write out link annotations
  */
    
  private AnnotationSet calcGroupLinkAnnotsCycle(List<NetLink> links, List<NetNode> nodes,
                                                 BTProgressMonitor monitor, 
                                                 boolean shadow, 
                                                 List<AlignCycleLayout.CycleBounds> bounds, 
                                                 List<String> linkGroups) throws AsynchExitRequestException {
  
    String which = (shadow) ? "progress.linkAnnotationShad" : "progress.linkAnnotationNoShad";
    LoopReporter lr = new LoopReporter(links.size(), 20, monitor, 0, 1.0, which); 
      
    HashMap<NetNode, Integer> nodeOrder = new HashMap<NetNode, Integer>();
    for (int i = 0; i < nodes.size(); i++) {
      nodeOrder.put(nodes.get(i), Integer.valueOf(i));      
    }
    
    NetNode currZoner = null;
    AlignCycleLayout.CycleBounds currLooper = new AlignCycleLayout.CycleBounds(null, null, false, false);
    HashSet<NetNode> seen = new HashSet<NetNode>();
    int cycle = 0;

    AnnotationSet retval = PluginSupportFactory.buildAnnotationSet();
    int startPos = 0;
    int endPos = 0;
    int numLink = links.size();
    int count = 0;
    boolean first = true;
    for (int i = 0; i < numLink; i++) {
      NetLink link = links.get(i);
      lr.report();
      if (link.isShadow() && !shadow) {
        continue;
      }
      NetNode zoner = getZoneNode(link, nodeOrder, link.isShadow());
      if (first) {
        first = false;
      }
      if ((currZoner == null) || !currZoner.equals(zoner)) { // New Zone
        if (currZoner != null) { // i.e. currZoner != zoner
          if (currZoner.equals(currLooper.boundEnd)) {
            if (!currLooper.boundStart.equals(currLooper.boundEnd) || !currLooper.isCorrect) {
              String color = (cycle % 2 == 0) ? "Orange" : "Green";
              String type = currLooper.isCycle ? "cycle " : "path ";
              retval.addAnnot(PluginSupportFactory.buildAnnotation(type + cycle++, startPos, endPos, 0, color));
            }
          }
        }
        currZoner = zoner;
        for (AlignCycleLayout.CycleBounds bound : bounds) {
          if (!seen.contains(bound.boundStart) && bound.boundStart.equals(currZoner)) {
            startPos = count;
            seen.add(bound.boundStart);
            currLooper = bound;
          }
        }
      }
      endPos = count++;
    }
    //
    // Close out the last pending annotation
    //
    if (!currLooper.boundStart.equals(currLooper.boundEnd) || !currLooper.isCorrect) {
      String color = (cycle % 2 == 0) ? "Orange" : "Green";
      String type = currLooper.isCycle ? "cycle " : "path ";
      retval.addAnnot(PluginSupportFactory.buildAnnotation(type + cycle++, startPos, endPos, 0, color));
    }
    return (retval);
  }
  
  /***************************************
  **
  ** A zone node is the bottom node of a shadow link or the top node of
  ** a regular link
  */
    
  private NetNode getZoneNode(NetLink link, Map<NetNode, Integer> nodes, boolean isShadow) {
    int zeroIndex = nodes.get(link.getSrcNode()).intValue();
    int oneIndex = nodes.get(link.getTrgNode()).intValue();
    if (isShadow) {
      NetNode botnode = (zeroIndex < oneIndex) ? link.getTrgNode() : link.getSrcNode();
      return (botnode);
    } else {
      NetNode topnode = (zeroIndex < oneIndex) ? link.getSrcNode() : link.getTrgNode();
      return (topnode); 
    }
  }

  /***************************************
  **
  ** Get the color
  */
  
  @Override
  protected String getColor(String type, Map<String, String> colorMap) {
    String trimmed = type.trim();
    if (trimmed.equals("G12")) {
      return ("Purple");
    } else if (trimmed.equals("G1A")) {
      return ("PowderBlue");
    } else if (trimmed.equals("G2A")) {
      return ("Pink");
    }
    throw new IllegalArgumentException();
  } 
}
```
