## Supplementary material for "BioFabric Visualization of Network Alignments": VISNAB Source Code: AlignCycleLayout.java

```
/*
**    Copyright (C) 2003-2019 Institute for Systems Biology 
**                            Seattle, Washington, USA. 
**
**    This library is free software; you can redistribute it and/or
**    modify it under the terms of the GNU Lesser General Public
**    License as published by the Free Software Foundation; either
**    version 2.1 of the License, or (at your option) any later version.
**
**    This library is distributed in the hope that it will be useful,
**    but WITHOUT ANY WARRANTY; without even the implied warranty of
**    MERCHANTABILITY or FITNESS FOR A PARTICULAR PURPOSE.  See the GNU
**    Lesser General Public License for more details.
**
**    You should have received a copy of the GNU Lesser General Public
**    License along with this library; if not, write to the Free Software
**    Foundation, Inc., 59 Temple Place, Suite 330, Boston, MA  02111-1307  USA
*/

package org.systemsbiology.biofabric.plugin.core.align;

import java.util.ArrayList;
import java.util.Collections;
import java.util.HashMap;
import java.util.HashSet;
import java.util.Iterator;
import java.util.List;
import java.util.Map;
import java.util.Set;
import java.util.SortedSet;
import java.util.TreeMap;
import java.util.TreeSet;

import org.systemsbiology.biofabric.api.io.BuildData;
import org.systemsbiology.biofabric.api.layout.LayoutCriterionFailureException;
import org.systemsbiology.biofabric.api.layout.NodeLayout;
import org.systemsbiology.biofabric.api.model.AnnotationSet;
import org.systemsbiology.biofabric.api.model.NetLink;
import org.systemsbiology.biofabric.api.model.NetNode;
import org.systemsbiology.biofabric.api.worker.AsynchExitRequestException;
import org.systemsbiology.biofabric.api.worker.BTProgressMonitor;
import org.systemsbiology.biofabric.api.worker.LoopReporter;
import org.systemsbiology.biofabric.plugin.PluginSupportFactory;
import org.systemsbiology.biofabric.util.UiUtil;

/****************************************************************************
**
** This is a layout algorithm that highlights incorrect alignments by placing
** mis-aligned nodes next to each other in the layout.
** 
**
** With the consideration of blue nodes, what used to be four cases has expanded to nine.
**
** Case 1: Correctly unaligned single blue node: (A->null)
** Case 2: Correctly unaligned single red node: (null->1) (also in no-blue cases)
** Case 3: Correctly aligned single node purple cycle: (B->2) (also in no-blue cases)
** Case 4: Incorrect purple node singleton path: (C->3), but truth is (C->null, null->3)
** Case 5: Incorrect two node (red,blue) path: (null->4, D->null) but truth is (4->D)
** Case 6: Incorrect 1+n node (red, n*purple) path: purple run starting with a red (also in no-blue cases)
** Case 7: Incorrect 1+n node (n*purple, blue) path: purple run ending with a blue
** Case 8: Incorrect 2+n node (red, n*purple, blue) path: purple run starting with a red, ending with a blue
** Case 9: Incorrect 1+n node (n*purple) cycle: purple cycle (also in no-blue cases)
*/

public class AlignCycleLayout extends NodeLayout {
  
  ////////////////////////////////////////////////////////////////////////////
  //
  // PRIVATE INSTANCE MEMBERS
  //
  ////////////////////////////////////////////////////////////////////////////
  
  private NodeMaps maps_;
    
  ////////////////////////////////////////////////////////////////////////////
  //
  // PUBLIC CONSTRUCTORS
  //
  ////////////////////////////////////////////////////////////////////////////

  /***************************************************************************
  **
  ** Constructor
  */

  public AlignCycleLayout() {
    maps_ = null;
  }

  ////////////////////////////////////////////////////////////////////////////
  //
  // PUBLIC METHODS
  //
  ////////////////////////////////////////////////////////////////////////////

  /***************************************************************************
  **
  ** Find out if the necessary conditions for this layout are met. For this layout, we either
  ** need to have the source and target nodes of the alignment in the same namespace, or a
  ** perfect alignment file to map the two namespaces.
  ** If you reuse the same object, this layout will cache the calculation it uses to answer
  ** the question for the actual layout!
  */
  
  @Override
  public boolean criteriaMet(BuildData rbd,
                             BTProgressMonitor monitor) throws AsynchExitRequestException, 
                                                               LayoutCriterionFailureException {
    
    NetworkAlignmentBuildData narbd = (NetworkAlignmentBuildData)rbd.getPluginBuildData();
    maps_ = normalizeAlignMap(narbd.mapG1toG2, narbd.perfectG1toG2, narbd.allLargerNodes, 
                              narbd.allSmallerNodes, monitor);
    if (maps_ == null) {
      throw new LayoutCriterionFailureException();
    }
    return (true);
  }
  
  /***************************************************************************
  **
  ** If you reuse the same object, this layout will cache the calculation it uses to answer
  ** the question for the actual layout. If you need to clear the cache, use this:
  */
  
  public void clearCache() {
    maps_ = null;
    return;
  }
  
   
  /***************************************************************************
  **
  ** Relayout the network!
  */
  
  public List<NetNode> doNodeLayout(BuildData rbd, 
  		                              Params params,
  		                              BTProgressMonitor monitor) throws AsynchExitRequestException {
      
    List<NetNode> targetIDs = doNodeOrder(rbd, params, monitor);

    //
    // Now have the ordered list of targets we are going to display.
    // Build target->row maps and the inverse:
    //
    
    installNodeOrder(targetIDs, rbd, monitor);
    
    NetworkAlignmentBuildData narbd = (NetworkAlignmentBuildData)rbd.getPluginBuildData();

    rbd.setTurnOnShadows(narbd.turnShadowsOn);
		if (rbd.getSingletonNodes().size() > 0) {
			narbd.useNodeGroups = true;
		}
		
    if (narbd.useNodeGroups) {
	    TreeMap<Integer, NetNode> invert = new TreeMap<Integer, NetNode>();
	    for (NetNode node : rbd.getNodeOrder().keySet()) {
	      invert.put(rbd.getNodeOrder().get(node), node);
	    }
	    ArrayList<NetNode> order = new ArrayList<NetNode>(invert.values());
	    AnnotationSet nAnnots = generateNodeAnnotations(order, monitor, narbd.cycleBounds);
	    rbd.setNodeAnnotations(nAnnots);
    }
    
    return (targetIDs);
  }
  
  
  /***************************************
  **
  ** Generate Node annotations
  */
    
  private AnnotationSet generateNodeAnnotations(List<NetNode> nodes,
                                                BTProgressMonitor monitor, 
                                                List<AlignCycleLayout.CycleBounds> bounds) throws AsynchExitRequestException {
  
    LoopReporter lr = new LoopReporter(nodes.size(), 20, monitor, 0, 1.0, "progress.nodeAnnotation"); 
      
    HashMap<NetNode, Integer> nodeOrder = new HashMap<NetNode, Integer>();
    for (int i = 0; i < nodes.size(); i++) {
      nodeOrder.put(nodes.get(i), Integer.valueOf(i));      
    }
    
    int cycle = 0;

    AnnotationSet retval = PluginSupportFactory.buildAnnotationSet();
    for (CycleBounds bound : bounds) {
    	lr.report();
    	if (bound.isCorrect) {
    		continue;
    	}
    	String type = bound.isCycle ? "cycle " : "path ";
    	int startPos = nodeOrder.get(bound.boundStart).intValue();
    	int endPos = nodeOrder.get(bound.boundEnd).intValue();
      String color = (cycle % 2 == 0) ? "Orange" : "Green";
      retval.addAnnot(PluginSupportFactory.buildAnnotation(type + cycle++, startPos, endPos, 0, color));
    }
    lr.finish();
  
    return (retval);
  }
   
  /***************************************************************************
  **
  ** Relayout the network!
  */
  
  public List<NetNode> doNodeOrder(BuildData rbd, 
                                   Params params,
                                   BTProgressMonitor monitor) throws AsynchExitRequestException {
      
    
    //
    // The actual start node might be different since we unroll paths to find the first node.
    // Thus, skip allowing the user to mess with this for now.
    //
    
    List<NetNode> startNodeIDs = null;
    
    NetworkAlignmentBuildData narbd = (NetworkAlignmentBuildData)rbd.getPluginBuildData();
    
    if (maps_ == null) {
      maps_ = normalizeAlignMap(narbd.mapG1toG2, narbd.perfectG1toG2, narbd.allLargerNodes, 
                                narbd.allSmallerNodes, monitor);
    }
    
    Set<NetNode> allNodes = genAllNodes(rbd);
    Map<NetNode, PathElem> nodesToPathElem = genNodeToPathElem(allNodes);
    Map<PathElem, NetNode> pathElemToNode = genPathElemToNode(allNodes);
    Map<String, PathElem> smallToElem = new HashMap<String, PathElem>();
    Map<String, PathElem> largeToElem = new HashMap<String, PathElem>();
    Map<PathElem, PathElem> elemToNext = new HashMap<PathElem, PathElem>();
    genNamesToPathElem(maps_, nodesToPathElem, smallToElem, largeToElem, elemToNext); 
 
    Map<PathElem, AlignPath> alignPaths = calcAlignPathsV2(maps_, nodesToPathElem, elemToNext, 
    		                                                  smallToElem, largeToElem);
     
    List<CycleBounds> cycleBounds = new ArrayList<CycleBounds>();
        
    List<NetNode> targetIDs = alignPathNodeOrder(rbd.getLinks(), rbd.getSingletonNodes(), 
                                                 startNodeIDs, alignPaths, 
                                                 nodesToPathElem,
                                                 pathElemToNode,
                                                 cycleBounds,
                                                 monitor);
    narbd.cycleBounds = cycleBounds;
    return (targetIDs);
  }
  
  /***************************************************************************
  ** 
  ** Calculate alignPath node order.
  */

  private List<NetNode> alignPathNodeOrder(Set<NetLink> allLinks,
  		                                     Set<NetNode> loneNodes, 
  		                                     List<NetNode> startNodes,
  		                                     Map<PathElem, AlignPath> alignPaths,
  		                                     Map<NetNode, PathElem> nodesToPathElem,
  		                                     Map<PathElem, NetNode> pathElemToNode,
  		                                     List<CycleBounds> cycleBounds,
  		                                     BTProgressMonitor monitor) throws AsynchExitRequestException { 
    //
    // Note the allLinks Set has pruned out duplicates and synonymous non-directional links
    //
    // NOTE! If we are handed a perfect alignment, we can build cycles even if the network is not aligned
    // to itself!
    //
    // We are handed a data structure that points from each aligned node to the alignment cycle it belongs to.
    // Working with a start node (highest degree or from list), order neighbors by decreasing degree, but instead of
    // adding unseen nodes in that order to the queue, we add ALL the nodes in the cycle to the list, in cycle order.
    // If it is a true cycle (i.e. does not terminate in an unaligned node) we can start the cycle at the neighbor. 
    // If it is not a cycle but a path, we need to start at the beginning.
    //
  
    
    HashMap<NetNode, Integer> linkCounts = new HashMap<NetNode, Integer>();
    HashMap<NetNode, Set<NetNode>> targsPerSource = new HashMap<NetNode, Set<NetNode>>();
    ArrayList<NetNode> targets = new ArrayList<NetNode>();
         
    HashSet<NetNode> targsToGo = new HashSet<NetNode>();
    
    int numLink = allLinks.size();
    LoopReporter lr = new LoopReporter(numLink, 20, monitor, 0.0, 0.25, "progress.calculateNodeDegree");
    
    Iterator<NetLink> alit = allLinks.iterator();
    while (alit.hasNext()) {
      NetLink nextLink = alit.next();
      lr.report();
      NetNode sidwn = nextLink.getSrcNode();
      NetNode tidwn = nextLink.getTrgNode();
      Set<NetNode> targs = targsPerSource.get(sidwn);
      if (targs == null) {
        targs = new HashSet<NetNode>();
        targsPerSource.put(sidwn, targs);
      }
      targs.add(tidwn);
      targs = targsPerSource.get(tidwn);
      if (targs == null) {
        targs = new HashSet<NetNode>();
        targsPerSource.put(tidwn, targs);
      }
      targs.add(sidwn);
      targsToGo.add(sidwn);
      targsToGo.add(tidwn);        
      Integer srcCount = linkCounts.get(sidwn);
      linkCounts.put(sidwn, (srcCount == null) ? Integer.valueOf(1) : Integer.valueOf(srcCount.intValue() + 1));
      Integer trgCount = linkCounts.get(tidwn);
      linkCounts.put(tidwn, (trgCount == null) ? Integer.valueOf(1) : Integer.valueOf(trgCount.intValue() + 1));
    }
    lr.finish();
    
    //
    // Rank the nodes by link count:
    //
    
    lr = new LoopReporter(linkCounts.size(), 20, monitor, 0.25, 0.50, "progress.rankByDegree");
    
    TreeMap<Integer, SortedSet<NetNode>> countRank = new TreeMap<Integer, SortedSet<NetNode>>(Collections.reverseOrder());
    Iterator<NetNode> lcit = linkCounts.keySet().iterator();
    while (lcit.hasNext()) {
      NetNode src = lcit.next();
      lr.report();
      Integer count = linkCounts.get(src);
      SortedSet<NetNode> perCount = countRank.get(count);
      if (perCount == null) {
        perCount = new TreeSet<NetNode>();
        countRank.put(count, perCount);
      }
      perCount.add(src);
    }
    lr.finish();
    
    //
    // Get all kids added in.  Now doing this without recursion; seeing blown
    // stacks for huge networks!
    //
     
    while (!targsToGo.isEmpty()) {
      Iterator<Integer> crit = countRank.keySet().iterator();
      while (crit.hasNext()) {
        Integer key = crit.next();
        SortedSet<NetNode> perCount = countRank.get(key);
        Iterator<NetNode> pcit = perCount.iterator();
        while (pcit.hasNext()) {
          NetNode node = pcit.next();
          if (targsToGo.contains(node)) {
            PathElem nodeKey = nodesToPathElem.get(node);
            AlignPath ac = alignPaths.get(nodeKey);
            ArrayList<NetNode> queue = new ArrayList<NetNode>();
            List<PathElem> unlooped = ac.getReorderedKidsStartingAtKidOrStart(nodeKey);
            for (PathElem ulnode : unlooped) { 
              NetNode daNode = pathElemToNode.get(ulnode);
              targsToGo.remove(daNode);
              targets.add(daNode);
              queue.add(daNode);
            }
            NetNode boundsStart = pathElemToNode.get(unlooped.get(0));
            NetNode boundsEnd = pathElemToNode.get(unlooped.get(unlooped.size() - 1));
            cycleBounds.add(new CycleBounds(boundsStart, boundsEnd, ac.correct, ac.isCycle));
            flushQueue(targets, targsPerSource, linkCounts, targsToGo, queue, alignPaths, nodesToPathElem,
                       pathElemToNode, cycleBounds, monitor,  0.75, 1.0);
          }
        }
      }
    }
    
    //
    //
    // Tag on lone nodes.  If a node is by itself, but also shows up in the links,
    // we drop it.
    //
    // Used to do a set removeAll() operation, but discovered the operation was
    // taking FOREVER, e.g. remains 190804 targets 281832. So do this in a loop
    // that can be monitored for progress:
    //
    
    LoopReporter lr2 = new LoopReporter(loneNodes.size(), 20, monitor, 0.0, 0.25, "progress.addSingletonsToTargets");
    HashSet<NetNode> targSet = new HashSet<NetNode>(targets);
    TreeSet<NetNode> orderedTargSet = new TreeSet<NetNode>(loneNodes);
    //
    // Even adding on singleton nodes, we need to retain the align cycle ordering:
    //
    for (NetNode lnod : orderedTargSet) {
    	if (!targSet.contains(lnod)) {
    		lr2.report();
    		PathElem nodeKey = nodesToPathElem.get(lnod);
    		AlignPath ac = alignPaths.get(nodeKey);
        List<PathElem> unlooped = ac.getReorderedKidsStartingAtKidOrStart(nodeKey);
        NetNode firstNode = null;
        NetNode lastNode = null;
        for (PathElem ulnode : unlooped) { 
          NetNode daNode = pathElemToNode.get(ulnode);
          if (firstNode == null) {
          	firstNode = daNode;
          }
          lastNode = daNode;
          targSet.add(daNode);
          targets.add(daNode);
        }
        cycleBounds.add(new CycleBounds(firstNode, lastNode, ac.correct, ac.isCycle));
    	}    	
    }
    lr2.finish();
     
    return (targets);
  }
        
  /***************************************************************************
  **
  ** Node ordering
  */
  
  private List<NetNode> orderMyKids(Map<NetNode, Set<NetNode>> targsPerSource, 
  		                                   Map<NetNode, Integer> linkCounts, 
                                         Set<NetNode> targsToGo, NetNode node) {
    Set<NetNode> targs = targsPerSource.get(node);
    if (targs == null) {
    	return (new ArrayList<NetNode>());
    }
    TreeMap<Integer, SortedSet<NetNode>> kidMap = new TreeMap<Integer, SortedSet<NetNode>>(Collections.reverseOrder());
    Iterator<NetNode> tait = targs.iterator();
    while (tait.hasNext()) {  
      NetNode nextTarg = tait.next(); 
      Integer count = linkCounts.get(nextTarg);
      SortedSet<NetNode> perCount = kidMap.get(count);
      if (perCount == null) {
        perCount = new TreeSet<NetNode>();
        kidMap.put(count, perCount);
      }
      perCount.add(nextTarg);
    }
    
    ArrayList<NetNode> myKidsToProc = new ArrayList<NetNode>();
    Iterator<SortedSet<NetNode>> kmit = kidMap.values().iterator();
    while (kmit.hasNext()) {  
      SortedSet<NetNode> perCount = kmit.next(); 
      Iterator<NetNode> pcit = perCount.iterator();
      while (pcit.hasNext()) {  
        NetNode kid = pcit.next();
        if (targsToGo.contains(kid)) { 
          myKidsToProc.add(kid);
        }
      }
    }
    return (myKidsToProc);
  }    
  
  /***************************************************************************
  **
  ** Node ordering, non-recursive:
  */
  
  private void flushQueue(List<NetNode> targets, 
  		                    Map<NetNode, Set<NetNode>> targsPerSource, 
                          Map<NetNode, Integer> linkCounts, 
                          Set<NetNode> targsToGo, List<NetNode> queue,
                          Map<PathElem, AlignPath> alignPaths,
                          Map<NetNode, PathElem> nodesToPathElem,
                          Map<PathElem, NetNode> pathElemToNode,
                          List<CycleBounds> cycleBounds,
                          BTProgressMonitor monitor, double startFrac, double endFrac) 
                            throws AsynchExitRequestException {
  	
  	LoopReporter lr = new LoopReporter(targsToGo.size(), 20, monitor, startFrac, endFrac, "progress.nodeOrdering");
  	int lastSize = targsToGo.size();	
    while (!queue.isEmpty()) {
      NetNode node = queue.remove(0);
      int ttgSize = targsToGo.size();
      lr.report(lastSize - ttgSize);
      lastSize = ttgSize;
      List<NetNode> myKids = orderMyKids(targsPerSource, linkCounts, targsToGo, node);
      Iterator<NetNode> ktpit = myKids.iterator(); 
      while (ktpit.hasNext()) {  
        NetNode kid = ktpit.next();
        if (targsToGo.contains(kid)) {
          PathElem kidKey = nodesToPathElem.get(kid);
          AlignPath ac = alignPaths.get(kidKey);
          targsToGo.removeAll(ac.pathNodes);
          List<PathElem> unlooped = ac.getReorderedKidsStartingAtKidOrStart(kidKey);
          for (PathElem ulnode : unlooped) { 
            NetNode daNode = pathElemToNode.get(ulnode);
            targsToGo.remove(daNode);
            targets.add(daNode);
            queue.add(daNode);
          }
          NetNode boundsStart = pathElemToNode.get(unlooped.get(0));
          NetNode boundsEnd = pathElemToNode.get(unlooped.get(unlooped.size() - 1));
          cycleBounds.add(new CycleBounds(boundsStart, boundsEnd, ac.correct, ac.isCycle));
        }
      }
    }
    lr.finish();
    return;
  }

  /***************************************************************************
  **
  ** In the alignment map we are provided, nodes in G1 and G2 that have the same
  ** name have different OIDs. Eliminate this difference. Also, if the source and
  ** target are not in the same namespace, we need to use the perfect alignment 
  ** (if available) to create the cycle/path map. 
  */
  
  private NodeMaps normalizeAlignMap(Map<NetNode, NetNode> align, 
                                     Map<NetNode, NetNode> perfectAlign,
                                     Set<NetNode> allLargerNodes,
                                     Set<NetNode> allSmallerNodes,                                     
                                     BTProgressMonitor monitor)  throws AsynchExitRequestException {
    
    //
    // Build sets of names. If the *names* are not unique, this layout cannot proceed. (Remembering that
  	// we can have distinct nodes with the same name but different node IDs.)
    //
    
    LoopReporter lr = new LoopReporter(align.size(), 20, monitor, 0.0, 1.00, "progress.normalizeAlignMapA");
    
    HashSet<String> keyNames = new HashSet<String>();
    for (NetNode key : align.keySet()) {
      if (keyNames.contains(key.getName())) {
      	lr.finish();
      	System.err.println("Duplicated key " + key.getName());
        return (null); 
      }
      keyNames.add(key.getName());
      lr.report();
    }
    lr.finish();
    
    LoopReporter lr2 = new LoopReporter(align.size(), 20, monitor, 0.0, 1.00, "progress.normalizeAlignMapB");
    
    HashSet<String> valNames = new HashSet<String>();
    for (NetNode value : align.values()) {
      if (valNames.contains(value.getName())) {
      	lr2.finish();
      	System.err.println("Duplicated value " + value.getName());
        return (null); 
      }
      valNames.add(value.getName());
      lr2.report();
    }
    lr2.finish();
    
    //
    // Reminder: a map takes elements in domain and spits out values in the range.
    // Alignment map A takes some of the nodes in smallNodes set S, and specifically ALL of the
    // purple nodes in S, into a subset of the nodes in largeNodes set L. To do the cycle layout, we 
    // need to have some inverse map F from L to S such the domain of F is a subset of L, and the range 
    // of F completely covers the subset of S that are purple nodes. If the nodes are in the same namespace, 
    // the identity map on the elements of L is sufficient. If not, the inverse of the perfect alignment map 
    // does the trick. Check first if the identity map can work.
    //
    
    LoopReporter lr3 = new LoopReporter(allLargerNodes.size(), 20, monitor, 0.0, 1.00, "progress.identityMapCheckA");
    
    HashSet<String> largeNames = new HashSet<String>();
    for (NetNode large : allLargerNodes) {
      if (largeNames.contains(large.getName())) {
      	lr3.finish();
        return (null); 
      }
      largeNames.add(large.getName());
      lr3.report();
    }
    lr3.finish();
    
    LoopReporter lr4 = new LoopReporter(allSmallerNodes.size(), 20, monitor, 0.0, 1.00, "progress.identityMapCheckB");
    
    HashSet<String> smallNames = new HashSet<String>();
    for (NetNode small : allSmallerNodes) {
      if (smallNames.contains(small.getName())) {
      	lr4.finish();
        return (null); 
      }
      smallNames.add(small.getName());
      lr4.report();
    }
    lr4.finish();
    
    //
    // Before blue nodes came along, it was enough to check that "largeNames.containsAll(smallNames)". But
    // it is sufficient for largeNames to cover the purple nodes, i.e. the keySet:
    //
    
    boolean identityOK = largeNames.containsAll(keyNames);
 
    // 
    // If identity map does not work, we need to build maps from the perfect alignment:
    //
    
    Map<String, String> backMap = null;

    if (!identityOK) {
      if ((perfectAlign == null) || perfectAlign.isEmpty()) {
        return (null);
      } 
      backMap = new HashMap<String, String>();
      
      LoopReporter lr5 = new LoopReporter(perfectAlign.size(), 20, monitor, 0.0, 1.00, "progress.namespaceMapBuilding");
      
      //
      // Again, with allowing blue nodes, we are OK with just covering the purple nodes in the smaller network, not
      // all the nodes in the smaller network. Inverting the perfect alignment does this.
      //
      
      for (NetNode key : perfectAlign.keySet()) {
        NetNode val = perfectAlign.get(key);
        backMap.put(val.getName(), key.getName());
        lr5.report();
      }
      lr5.finish();
    }

    return (new NodeMaps(backMap));
  } 
  
  /***************************************************************************
  **
  ** Get all nodes
  */
  
  private Set<NetNode> genAllNodes(BuildData narbd) { 

    Set<NetNode> allNodes = new HashSet<NetNode>();
     
     for (NetLink link : narbd.getLinks()) {
       allNodes.add(link.getSrcNode());
       allNodes.add(link.getTrgNode());
     }
     allNodes.addAll(narbd.getSingletonNodes());
     return (allNodes);
   } 
   
  /***************************************************************************
  **
  ** Get map from network NetNode (of form G1::G2) to path elem (G1)
  ** With handling of blue nodes, nodes names of the form A:: and ::B are
  ** possible.
  */
  
  private Map<NetNode, PathElem> genNodeToPathElem(Set<NetNode> allNodes) { 

     Map<NetNode, PathElem> n2pe = new HashMap<NetNode, PathElem>();
     for (NetNode key : allNodes) {	 
       PathElem elem = new PathElem(key);
       n2pe.put(key, elem);
     }
     return (n2pe);
   } 
  
  /***************************************************************************
  **
  ** Fill in maps taking small or large net node names to PathElems
  */
  
  private void genNamesToPathElem(NodeMaps maps,
  		                            Map<NetNode, PathElem> elemMap,   
  		                            Map<String, PathElem> smallToElem,
                                  Map<String, PathElem> largeToElem,
                                  Map<PathElem, PathElem> elemToNext) { 

     for (NetNode key : elemMap.keySet()) {	 
       PathElem elem = elemMap.get(key);
       if (elem.color != PathElem.NodeColor.RED) {
         smallToElem.put(elem.smallNodeName, elem);
       }
       if (elem.color != PathElem.NodeColor.BLUE) {
         largeToElem.put(elem.largeNodeName, elem);
       }
     }
     
     for (NetNode key : elemMap.keySet()) {	 
       PathElem elem = elemMap.get(key);
       if (elem.color != PathElem.NodeColor.BLUE) {
         String nextSmallKey = (maps.backMap != null) ? maps.backMap.get(elem.largeNodeName) : elem.largeNodeName;
         if (nextSmallKey != null) {
           elemToNext.put(elem, smallToElem.get(nextSmallKey));
         }
       }  
     } 
     return;
   } 

  /***************************************************************************
  **
  ** Inverse of above
  */
  
  private Map<PathElem, NetNode> genPathElemToNode(Set<NetNode> allNodes) { 

    Map<PathElem, NetNode> pe2n = new HashMap<PathElem, NetNode>();
    
    for (NetNode key : allNodes) {
    	PathElem elem = new PathElem(key);
      pe2n.put(elem, key);
    }
    return (pe2n);
  } 
  
  
  /***************************************************************************
  **
  ** Extract the paths in the alignment network.
  ** We have two orthogonal issues: 1) are nodes in same namespace, and 2) are there more
  ** nodes in the larger network? With different namespaces, we need a reverse map. If the
  ** larger network has more nodes, there might not be a reverse mapping, so not every path
  ** will be a cycle. Note that while A->A is an obvious correct match, A->1234 is not so
  ** clear. A->B B->A is a swap that needs to be annotated as a cycle, but A->B should obviously
  ** also be marked as a path if B is in the larger net and not aligned. Similarly, A->1234 should
  ** be unmarked if it is correct, but marked if is incorrect.
  */
  
  private Map<PathElem, AlignPath> calcAlignPathsV2(NodeMaps align, Map<NetNode, PathElem> nodesToPathElem,
  																									Map<PathElem, PathElem> elemToNext,
  																									Map<String, PathElem> smallToElem,
  																									Map<String, PathElem> largeToElem) { 

     Map<PathElem, AlignPath> pathsPerStart = new HashMap<PathElem, AlignPath>();
     
     HashSet<PathElem> working = new HashSet<PathElem>(nodesToPathElem.values());
     while (!working.isEmpty()) {
       PathElem startElem = working.iterator().next();
       working.remove(startElem);
       AlignPath path = new AlignPath();
       pathsPerStart.put(startElem, path);
       path.pathNodes.add(startElem);
       PathElem nextElem = elemToNext.get(startElem);

       // Not every cycle closes itself, so nextElem can == null:
       while (nextElem != null) {
         if (nextElem.equals(startElem)) {
           path.isCycle = true;
           path.correct = (path.pathNodes.size() == 1);
           break;
         }
         AlignPath existing = pathsPerStart.get(nextElem);
         // If there is an existing path for the next key, we just glue that
         // existing path onto the end of this new path head. Note we do not
         // bother with then getting the "nextKey", as we have already 
         // traversed that path tail.
         if (existing != null) {
           path.pathNodes.addAll(existing.pathNodes);
           pathsPerStart.remove(nextElem);
           if (working.contains(nextElem)) {
             throw new IllegalStateException();
           }
           if (existing.isCycle) {
             throw new IllegalStateException();
           }
           break;
         } else {
        	 path.pathNodes.add(nextElem);
        	 working.remove(nextElem);
        	 nextElem = elemToNext.get(nextElem);    
         }
       }
     }
     
     Map<PathElem, AlignPath> pathsPerEveryNode = new HashMap<PathElem, AlignPath>();
     for (PathElem keyName : pathsPerStart.keySet()) {
       AlignPath path = pathsPerStart.get(keyName);
       //
       // We want to get all length-one paths that are Blue or Red marked correct. (Cases 1 & 2)
       if (path.pathNodes.size() == 1) {
      	 PathElem oneElem = path.pathNodes.get(0);
      	 if (oneElem.color == PathElem.NodeColor.PURPLE) {
      		 if (path.isCycle) {
      			 if (!path.correct) { // Incorrect case 3
      				 throw new IllegalStateException();
      			 }
      		 } else {
      			 if (path.correct) { // Incorrect case 4
      				 throw new IllegalStateException();
      			 }
      		 }
      	 } else {
      		 path.correct = true;
      	 }
       }
       for (PathElem nextName : path.pathNodes) {
         pathsPerEveryNode.put(nextName, path); 
       }
     }

     return (pathsPerEveryNode);
   }
  
  /***************************************************************************
  **
  ** For passing around layout path
  */  
  
  private static class AlignPath  {
        
    List<PathElem> pathNodes;
    boolean isCycle;
    boolean correct;

    AlignPath() {
      pathNodes = new ArrayList<PathElem>();
      isCycle = false;
      correct = false;
    } 
    
    List<PathElem> getReorderedKidsStartingAtKidOrStart(PathElem start) {
      // If we have a loop, unroll it starting at the provided start:
      if (isCycle) {
        int startIndex = pathNodes.indexOf(start);
        int len = pathNodes.size();
        List<PathElem> retval = new ArrayList<PathElem>();
        for (int i = 0; i < len; i++) {
          int index = (startIndex + i) % len;
          retval.add(pathNodes.get(index));
        }
        return (retval);
      } else {
        return (pathNodes);
      }
    }
  }
  
  /***************************************************************************
  **
  ** Path elements. Previously used strings, but blue node complexities drive this
  ** to be more involved
  */  
  
  private static class PathElem  {
  	
  	enum NodeColor {PURPLE, RED, BLUE};
        
    String smallNodeName;
    String largeNodeName;
    NodeColor color;
   
    PathElem(NetNode node) {
    	// The -1 argument means we get an empty string if the :: is a suffix (blue node)
      String[] toks = node.getName().split("::", -1);
      if (toks.length != 2) {
      	throw new IllegalArgumentException();
      }
      boolean firstEmpty = toks[0].equals("");
      boolean secondEmpty = toks[1].equals("");
      //
      // If first element is empty, we have a red node. If last element is empty, 
      // we have a blue node. If neither, the node is purple:
      //
      if (firstEmpty && !secondEmpty) {
        color = NodeColor.RED;
        largeNodeName = toks[1];
      } else if (!firstEmpty && secondEmpty) {
        color = NodeColor.BLUE;
        smallNodeName = toks[0];
      } else if (!firstEmpty && !secondEmpty) {
      	color = NodeColor.PURPLE;
      	smallNodeName = toks[0];
        largeNodeName = toks[1];
      } else {
      	throw new IllegalArgumentException();
      }
    }
    
    @Override
    public String toString() {
    	switch (this.color) {
	      case RED:
	      	return (this.color + " node : ::" + largeNodeName);
	      case BLUE:
	      	return (this.color + " node : " + smallNodeName + "::");
	      case PURPLE:
	      	return (this.color + " node : " + smallNodeName + "::" + largeNodeName);
        default:
        	break;   	
	    }
    	throw new IllegalStateException();
    }
 
    @Override
    public int hashCode() {
    	int smallCode = (smallNodeName == null) ? 0 : smallNodeName.hashCode();
    	int largeCode = (largeNodeName == null) ? 0 : largeNodeName.hashCode();
      return (smallCode + largeCode + color.hashCode());
    }

    @Override
    public boolean equals(Object other) {
	    if (other == null) {
	      return (false);
	    }
	    if (other == this) {
	      return (true);
	    }
	    if (!(other instanceof PathElem)) {
	      return (false);
	    }
	    
	    PathElem otherElem = (PathElem)other;
	    if (otherElem.color != this.color) {
	    	return (false);
	    }
	    switch (this.color) {
	      case RED:
	      	return (this.largeNodeName.equals(otherElem.largeNodeName));
	      case BLUE:
	      	return (this.smallNodeName.equals(otherElem.smallNodeName));
	      case PURPLE:
	      	return (this.largeNodeName.equals(otherElem.largeNodeName) &&
	      	        this.smallNodeName.equals(otherElem.smallNodeName));
        default:
        	break;   	
	    }
	    throw new IllegalStateException();
	  }
  }

  /***************************************************************************
  **
  ** For passing around layout params
  */  
  
  public static class DefaultParams implements Params {
        
    public List<NetNode> startNodes;

    public DefaultParams(List<NetNode> startNodes) {
      this.startNodes = startNodes;
    } 
  }
  
  /***************************************************************************
  **
  ** For passing around node maps
  ** Reminder: a map takes elements in domain and spits out values in the range.
  ** Alignment map A takes some of the nodes in smallNodes set S, and specifically ALL of the
  ** purple nodes in S, into a subset of the nodes in largeNodes set L. To do the cycle layout, we 
  ** need to have some inverse map F from L to S such the domain of F is a subset of L, and the range 
  ** of F completely covers the subset of S that are purple nodes. If the nodes are in the same namespace, 
  ** the identity map on the elements of L is sufficient. If not, the inverse of the perfect alignment map 
  ** does the trick. Check first if the identity map can work.
  */  
  
  private static class NodeMaps  {
        
    Map<String, String> backMap;  // The correct back alignment of nodes in L that go to purple nodes of S 

    NodeMaps(Map<String, String> backMap) {
      this.backMap = backMap;
    } 
  }
  
  /***************************************************************************
  **
  ** For passing around cycles. A cycle that starts and ends on the same node
  ** might be incorrect if a smaller network is overlaid on a larger one.
  */  
  
  public static class CycleBounds  {
        
    public NetNode boundStart;
    public NetNode boundEnd;
    public boolean isCorrect;
    public boolean isCycle;

    CycleBounds(NetNode boundStart, NetNode boundEnd, boolean isCorrect, boolean isCycle) {
      this.boundStart = boundStart;
      this.boundEnd = boundEnd;
      this.isCorrect = isCorrect;
      this.isCycle = isCycle;
    } 
  }
 
}
```
