## Supplementary material for "BioFabric Visualization of Network Alignments": VISNAB Source Code: AlignmentLoader.java

```
/*
**    Copyright (C) 2018 Rishi Desai
**
**    Copyright (C) 2003-2014 Institute for Systems Biology
**                            Seattle, Washington, USA.
**
**    This library is free software; you can redistribute it and/or
**    modify it under the terms of the GNU Lesser General Public
**    License as published by the Free Software Foundation; either
**    version 2.1 of the License, or (at your option) any later version.
**
**    This library is distributed in the hope that it will be useful,
**    but WITHOUT ANY WARRANTY; without even the implied warranty of
**    MERCHANTABILITY or FITNESS FOR A PARTICULAR PURPOSE.  See the GNU
**    Lesser General Public License for more details.
**
**    You should have received a copy of the GNU Lesser General Public
**    License along with this library; if not, write to the Free Software
**    Foundation, Inc., 59 Temple Place, Suite 330, Boston, MA  02111-1307  USA
*/

package org.systemsbiology.biofabric.plugin.core.align;

import java.text.MessageFormat;
import java.util.ArrayList;
import java.util.HashMap;
import java.util.HashSet;
import java.util.Map;
import java.util.Set;
import java.util.StringTokenizer;

import org.systemsbiology.biofabric.api.io.BuildExtractor;
import org.systemsbiology.biofabric.api.model.NetLink;
import org.systemsbiology.biofabric.api.model.NetNode;
import org.systemsbiology.biofabric.api.util.PluginResourceManager;
import org.systemsbiology.biofabric.api.worker.AsynchExitRequestException;
import org.systemsbiology.biofabric.plugin.PluginSupportFactory;

import java.io.BufferedReader;
import java.io.File;
import java.io.FileInputStream;
import java.io.IOException;
import java.io.InputStreamReader;

/****************************************************************************
 **
 ** This loads alignment (.align) files
 */

public class AlignmentLoader {
  
  ////////////////////////////////////////////////////////////////////////////
  //
  // PRIVATE INSTANCE MEMBERS
  //
  ////////////////////////////////////////////////////////////////////////////
  
	private PluginResourceManager rMan_;
	
  ////////////////////////////////////////////////////////////////////////////
  //
  // PUBLIC CONSTRUCTORS
  //
  ////////////////////////////////////////////////////////////////////////////
  
  /***************************************************************************
   **
   ** Constructor
   */
  
  public AlignmentLoader(String pluginClassName, PluginResourceManager rMan) {
  	rMan_ = rMan;
  }
  
  ////////////////////////////////////////////////////////////////////////////
  //
  // PUBLIC METHODS
  //
  ////////////////////////////////////////////////////////////////////////////
  
  /***************************************************************************
   **
   ** Process an alignment (.align) file
   */
  
  public String readAlignment(File infile, Map<NetNode, NetNode> mapG1ToG2, NetAlignFileStats stats,
                              ArrayList<NetLink> linksGraph1, HashSet<NetNode> loneNodesGraph1,
                              ArrayList<NetLink> linksGraph2, HashSet<NetNode> loneNodesGraph2)
          throws IOException {
      
    Map<String, NetNode> G1nameToNID, G2nameToNID;
    try {
    	BuildExtractor bex = PluginSupportFactory.getBuildExtractor();
      G1nameToNID = makeStringMap(bex.extractNodes(linksGraph1, loneNodesGraph1, null));
      G2nameToNID = makeStringMap(bex.extractNodes(linksGraph2, loneNodesGraph2, null));
    } catch (AsynchExitRequestException aere) {
      throw new IllegalStateException();
    }
    BufferedReader in = null;
    
    try {
    	in = new BufferedReader(new InputStreamReader(new FileInputStream(infile), "UTF-8"));
    
	    String line = null;
	    while ((line = in.readLine()) != null) {
	  
	      StringTokenizer st = new StringTokenizer(line);
	      if (st.countTokens() != 2) {
	        stats.badLines.add(line);
	        continue;
	      }
	  
	      //
	      // Checking if nodes are in graphs
	      //
	  
	      String strNameG1 = st.nextToken(), strNameG2 = st.nextToken();
	     
	      boolean existsInG1 = G1nameToNID.containsKey(strNameG1),
	              existsInG2 = G2nameToNID.containsKey(strNameG2);
      
	      if (!existsInG1) {
	        String msg = MessageFormat.format(rMan_.getPluginString("networkAlignment.nodeNotFoundG1"), strNameG1);
	        throw (new IOException(msg));
	      }
	      if (!existsInG2) {
	        String msg = MessageFormat.format(rMan_.getPluginString("networkAlignment.nodeNotFoundG2"), strNameG2);
	        throw (new IOException(msg));
	      }
	      
	      NetNode nodeG1 = G1nameToNID.get(strNameG1), nodeG2 = G2nameToNID.get(strNameG2);
	      
	      if (mapG1ToG2.containsKey(nodeG1)) {
	        
	        if (! mapG1ToG2.get(nodeG1).equals(nodeG2)) {
	          String msg = MessageFormat.format(rMan_.getPluginString("networkAlignment.mapError"), strNameG1);
	          throw (new IOException(msg));
	        } else {
	          stats.dupLines.add(line);
	        }
	      } else {
	        mapG1ToG2.put(nodeG1, nodeG2);
	      }
	    }
	  
//	    if (mapG1ToG2.size() != G1nameToNID.size()) {	// commented out for now - Rishi 11/20/2018
//	      String msg = MessageFormat.format(rMan_.getPluginString("networkAlignment.mapSizeError"), mapG1ToG2.size(), G1nameToNID.size());
//	      throw (new IOException(msg));
//	    }
    } finally {
    	if (in != null) in.close();
    }
    return (null);
  }
  
  /***************************************************************************
   **
   ** Process an alignment (.align) file for determining graph1 and graph2
   ** Nearly all error catching is done in other readAlignment method
   */
  
  public Map<String, String> readAlignment(File infile, NetAlignFileStats stats) throws IOException {
  	
    Map<String, String> mapG1ToG2Str = new HashMap<String, String>();
    BufferedReader in = null;
    
    try {
    	in = new BufferedReader(new InputStreamReader(new FileInputStream(infile), "UTF-8"));
  
	   
	    String line = null;
	    while ((line = in.readLine()) != null) {
	    
	      StringTokenizer st = new StringTokenizer(line);
	      if (st.countTokens() != 2) {
	        stats.badLines.add(line);
	        continue;
	      }
	    
	      String strNameG1 = st.nextToken(), strNameG2 = st.nextToken();
	      
	      if (mapG1ToG2Str.containsKey(strNameG1)) {
	        if (! mapG1ToG2Str.get(strNameG1).equals(strNameG2)) {
	          String msg = MessageFormat.format(rMan_.getPluginString("networkAlignment.mapError"), strNameG1);
	          throw (new IOException(msg));
	        } else {
	          stats.dupLines.add(line);
	        }
	      } else {
	        mapG1ToG2Str.put(strNameG1, strNameG2);
	      }
	    }
    } finally {
    	if (in != null) in.close();
    }

    return (mapG1ToG2Str);
  }
  
  /***************************************************************************
   **
   ** Map node name to NetNode using node set
   */
  
  private static Map<String,NetNode> makeStringMap(Set<NetNode> nodes) {
  
    Map<String,NetNode> retval = new HashMap<String, NetNode>();
    
    for (NetNode node : nodes) {
      retval.put(node.getName(), node);
    }
    return retval;
  }
  
  public static class NetAlignFileStats {
    public ArrayList<String> dupLines;
    public ArrayList<String> badLines;
    
    public NetAlignFileStats() {
      badLines = new ArrayList<String>();
      dupLines = new ArrayList<String>();
    }
  }
  
}
```
