## Supplementary material for "BioFabric Visualization of Network Alignments": VISNAB Source Code: JaccardSimilarity.java

```
/*
 **
 **    Copyright (C) 2016-2019 Rishi Desai
 **
 **    Copyright (C) 2003-2018 Institute for Systems Biology
 **                            Seattle, Washington, USA.
 **
 **    This library is free software; you can redistribute it and/or
 **    modify it under the terms of the GNU Lesser General Public
 **    License as published by the Free Software Foundation; either
 **    version 2.1 of the License, or (at your option) any later version.
 **
 **    This library is distributed in the hope that it will be useful,
 **    but WITHOUT ANY WARRANTY; without even the implied warranty of
 **    MERCHANTABILITY or FITNESS FOR A PARTICULAR PURPOSE.  See the GNU
 **    Lesser General Public License for more details.
 **
 **    You should have received a copy of the GNU Lesser General Public
 **    License along with this library; if not, write to the Free Software
 **    Foundation, Inc., 59 Temple Place, Suite 330, Boston, MA  02111-1307  USA
 */

package org.systemsbiology.biofabric.plugin.core.align;

import java.util.ArrayList;
import java.util.HashMap;
import java.util.HashSet;
import java.util.Map;
import java.util.Set;
import org.systemsbiology.biofabric.api.io.BuildExtractor;
import org.systemsbiology.biofabric.api.model.NetLink;
import org.systemsbiology.biofabric.api.model.NetNode;
import org.systemsbiology.biofabric.api.worker.AsynchExitRequestException;
import org.systemsbiology.biofabric.api.worker.BTProgressMonitor;
import org.systemsbiology.biofabric.plugin.PluginSupportFactory;

/***************************************************************************
 **
 ** Functions for Jaccard Similarity
 */

public class JaccardSimilarity {
  
  ////////////////////////////////////////////////////////////////////////////
  //
  // PRIVATE INSTANCE MEMBERS
  //
  ////////////////////////////////////////////////////////////////////////////
  
  //
  // These are from original untouched graphs and alignments
  //
  
  private ArrayList<NetLink> linksSmall_, linksLarge_;
  private HashSet<NetNode> lonersSmall_, lonersLarge_;
  private Map<NetNode, NetNode> mapG1toG2_, perfectG1toG2_;
  private Map<NetNode, NetNode> invMainG2toG1_, invPerfectG2toG1_;
  
  private Map<NetNode, Set<NetNode>> nodeToNeighSmall_, nodeToNeighLarge_;
  private Map<String, NetNode> nameToSmall_;
  
  //
  // These are from the processed, merged alignment - None are used
  //
  
  private Set<NetLink> allLinksMain_, allLinksPerfect_;
  private Set<NetNode> loneNodeIDsMain_, loneNodeIDsPerfect_;
  private NetworkAlignment.NodeColorMap colorMapMain_, colorMapPerfect_;
  private Map<NetNode, Set<NetNode>> nodeToNeighborsMain_, nodeToNeighborsPerfect_;
  private Map<NetNode, Set<NetLink>> nodeToLinksMain_, nodeToLinksPerfect_;
  
  private BTProgressMonitor monitor_;
  private final Double jaccSimThreshold_;
  
  ////////////////////////////////////////////////////////////////////////////
  //
  // CONSTRUCTORS
  //
  ////////////////////////////////////////////////////////////////////////////
  
  JaccardSimilarity(Set<NetLink> allLinksMain, Set<NetNode> loneNodeIDsMain,
                    NetworkAlignment.NodeColorMap colorMapMain,
                    Set<NetLink> allLinksPerfect, Set<NetNode> loneNodeIDsPerfect,
                    NetworkAlignment.NodeColorMap colorMapPerfect,
                    Map<NetNode, Set<NetNode>> nodeToNeighborsMain,
                    Map<NetNode, Set<NetLink>> nodeToLinksMain,
                    Map<NetNode, Set<NetNode>> nodeToNeighborsPerfect,
                    Map<NetNode, Set<NetLink>> nodeToLinksPerfect,
                    ArrayList<NetLink> linksSmall, HashSet<NetNode> lonersSmall,
                    ArrayList<NetLink> linksLarge, HashSet<NetNode> lonersLarge,
                    Map<NetNode, NetNode> mapG1toG2, Map<NetNode, NetNode> perfectG1toG2,
                    final Double jaccSimThreshold,
                    BTProgressMonitor monitor) throws AsynchExitRequestException {
    
    this.allLinksMain_ = allLinksMain;
    this.loneNodeIDsMain_ = loneNodeIDsMain;
    this.colorMapMain_ = colorMapMain;
    this.nodeToNeighborsMain_ = nodeToNeighborsMain;
    this.nodeToLinksMain_ = nodeToLinksMain;
    this.allLinksPerfect_ = allLinksPerfect;
    this.loneNodeIDsPerfect_ = loneNodeIDsPerfect;
    this.colorMapPerfect_ = colorMapPerfect;
    this.nodeToNeighborsPerfect_ = nodeToNeighborsPerfect;
    this.nodeToLinksPerfect_ = nodeToLinksPerfect;
    this.monitor_ = monitor;
    this.jaccSimThreshold_ = jaccSimThreshold;
  
    this.linksSmall_ = linksSmall;
    this.lonersSmall_ = lonersSmall;
    this.linksLarge_ = linksLarge;
    this.lonersLarge_ = lonersLarge;
    this.mapG1toG2_ = mapG1toG2;
    this.perfectG1toG2_ = perfectG1toG2;
    
    generateStructures();
    return;
  }
  
  private void generateStructures() throws AsynchExitRequestException {
    nodeToNeighSmall_ = new HashMap<NetNode, Set<NetNode>>();
    nodeToNeighLarge_ = new HashMap<NetNode, Set<NetNode>>();
    BuildExtractor bex = PluginSupportFactory.getBuildExtractor();
    bex.createNeighborLinkMap(linksSmall_, lonersSmall_, nodeToNeighSmall_, new HashMap<NetNode, Set<NetLink>>(), monitor_);
    bex.createNeighborLinkMap(linksLarge_, lonersLarge_, nodeToNeighLarge_, new HashMap<NetNode, Set<NetLink>>(), monitor_);
    
    invMainG2toG1_ = new HashMap<NetNode, NetNode>();
    invPerfectG2toG1_ = new HashMap<NetNode, NetNode>();
    makeInverseMap(mapG1toG2_, invMainG2toG1_);
    makeInverseMap(perfectG1toG2_, invPerfectG2toG1_);
    
    nameToSmall_ = new HashMap<String, NetNode>();
    Set<NetNode> smallNodes = PluginSupportFactory.getBuildExtractor().extractNodes(linksSmall_, lonersSmall_, monitor_);
    for (NetNode smallNode : smallNodes) {
      nameToSmall_.put(smallNode.getName(), smallNode);
    }
    return;
  }
  
  /***************************************************************************
   **
   ** @param nodeV12 must be an merged node, either A:: or A::B
   ** Checks if the JS value of the node is above set threshold
   */
  
  boolean isCorrectJS(NetNode nodeV12) {
    String smallName = StringUtilities.separateNodeOne(nodeV12.getName());
    
    NetNode smallNode = nameToSmall_.get(smallName);
    if (smallNode == null) {
      throw new IllegalStateException("Small node for " + nodeV12.getName() + " not found for Jaccard Similarity");
    }
    double jsVal = jaccSimDecision(smallNode);
    if (jaccSimThreshold_ == null) {
      throw new IllegalStateException("JS Threshold is null"); // should never happen
    }
    boolean isCorrect = Double.compare(jsVal, jaccSimThreshold_) >= 0;
    return (isCorrect);
  }
  
  /****************************************************************************
   **
   ** Jaccard Similarity Measure - Historically Adapted from NodeEQC.java
   ** Adapted for Blue Nodes Case - Now uses case work to see where nodes
   ** Should be compared.
   */
  
  double calcScore() throws AsynchExitRequestException {
    double totJ = 0.0;
    Set<NetNode> smallNodes = PluginSupportFactory.getBuildExtractor().extractNodes(linksSmall_, lonersSmall_, monitor_);
    for (NetNode node : smallNodes) {
      totJ += jaccSimDecision(node);
    }
    double measure = totJ / smallNodes.size();
    return (measure);
  }
  
  /***************************************************************************
   **
   ** For each node N in G1:
   **
   ** Perfect- Blue   &  Main- Blue    => JS = 1.0
   ** Perfect- Blue   &  Main- Purple  => JS over G1
   ** Perfect- Purple &  Main- Blue    => JS over G1
   ** Perfect- Purple &  Main- Purple  => JS over G2
   */
  
  private double jaccSimDecision(NetNode nodeG1) {
    boolean isBlueTest = (mapG1toG2_.get(nodeG1) == null), isBluePerf = (perfectG1toG2_.get(nodeG1) == null);
    double jsVal;
    if (isBlueTest && isBluePerf) { // blue and blue => 1.0
      jsVal = 1.0;
    } else if (isBlueTest) { // purple and blue  or  blue and purple => JS over smaller network
      NetNode match = findMatch(nodeG1, CaseType.PERFECT_PURPLE_MAIN_BLUE);
      jsVal = jaccSimValue(nodeG1, match, NetworkAlignment.GraphType.SMALL);
    } else if (isBluePerf) {
      NetNode match = findMatch(nodeG1, CaseType.PERFECT_BLUE_MAIN_PURPLE);
      jsVal = jaccSimValue(nodeG1, match, NetworkAlignment.GraphType.SMALL);
    } else { // purple and purple => JS over large network
      NetNode alignTest = mapG1toG2_.get(nodeG1), alignPerf = perfectG1toG2_.get(nodeG1);
      jsVal = jaccSimValue(alignTest, alignPerf, NetworkAlignment.GraphType.LARGE);
    }
    return (jsVal);
  }
  
  /***************************************************************************
   **
   ** Nodes must (obviously) be from the graph specified
   */
  
  private double jaccSimValue(NetNode nodeA, NetNode nodeB, NetworkAlignment.GraphType type) {
    int lenAdjust = 0;
  
    HashSet<NetNode> scratchA, scratchB;
    if (type == NetworkAlignment.GraphType.SMALL) {
      scratchA = new HashSet<NetNode>(nodeToNeighSmall_.get(nodeA));
      scratchB = new HashSet<NetNode>(nodeToNeighSmall_.get(nodeB));
    } else if (type == NetworkAlignment.GraphType.LARGE) {
      scratchA = new HashSet<NetNode>(nodeToNeighLarge_.get(nodeA));
      scratchB = new HashSet<NetNode>(nodeToNeighLarge_.get(nodeB));
    } else {
      throw (new IllegalArgumentException("Graph type not allowed"));
    }
    
    if (scratchA.contains(nodeB)) {
      scratchA.remove(nodeB);
      scratchB.remove(nodeA);
      lenAdjust = 1;
    }
    HashSet<NetNode> union = new HashSet<NetNode>(), intersect = new HashSet<NetNode>();
    union(scratchA, scratchB, union);
    intersection(scratchA, scratchB, intersect);
    
    int iSize = intersect.size() + lenAdjust;
    int uSize = union.size() + lenAdjust;
    Double jaccard = (double) (iSize) / (double) uSize;
    if (jaccard.isNaN()) {  // case of 0/0 for two singletons
      jaccard = 1.0;
    }
    return (jaccard);
  }
  
  /***************************************************************************
   **
   ** Finds the JS-match of a node in the Purple - Blue and Blue - Purple case
   */
  
  private NetNode findMatch(NetNode nodeG1, CaseType type) {
    NetNode nodeG1Match;
    if (type == CaseType.PERFECT_BLUE_MAIN_PURPLE) {
      NetNode nodeG2Main = mapG1toG2_.get(nodeG1);
      nodeG1Match = invPerfectG2toG1_.get(nodeG2Main);
    } else if (type == CaseType.PERFECT_PURPLE_MAIN_BLUE) {
      NetNode nodeG2Perf = perfectG1toG2_.get(nodeG1);
      nodeG1Match = invMainG2toG1_.get(nodeG2Perf);
    } else {
      throw (new IllegalArgumentException("Incorrect case type"));
    }
    return (nodeG1Match);
  }
  
  /***************************************************************************
   **
   ** Inverse of an alignment map
   */
  
  private void makeInverseMap(Map<NetNode, NetNode> in, Map<NetNode, NetNode> out) {
    for (Map.Entry<NetNode, NetNode> entry : in.entrySet()) {
      out.put(entry.getValue(), entry.getKey());
    }
    return;
  }
  
  /***************************************************************************
   **
   ** Set intersection helper
   */
  
  private <T> void intersection(Set<T> one, Set<T> two, Set<T> result) {
    result.clear();
    result.addAll(one);
    result.retainAll(two);
    return;
  }
  
  /***************************************************************************
   **
   ** Set union helper
   */
  
  private <T> void union(Set<T> one, Set<T> two, Set<T> result) {
    result.clear();
    result.addAll(one);
    result.addAll(two);
    return;
  }
  
  /***************************************************************************
   **
   ** The Purple-Blue and Blue-Purple cases
   */
  
  private enum CaseType {
    PERFECT_BLUE_MAIN_PURPLE,
    PERFECT_PURPLE_MAIN_BLUE
  }
  
  /****************************************************************************
   **
   ** String Utilities
   */

  private static class StringUtilities {

    /****************************************************************************
     **
     ** For "A::" and "A::B" this returns "A"
     */

    static String separateNodeOne(String in) {
      if (! in.contains("::")) {
        throw (new IllegalArgumentException("Node name needs ::"));
      }
      String ret = (in.split("::"))[0];
      return (ret);
    }

    static String separateNodeTwo(String in) {
      if (! in.contains("::")) {
        throw (new IllegalArgumentException("Node name needs ::"));
      }
      String ret = (in.split("::"))[1];
      return (ret);
    }


  }
  
}
```
