## Supplementary material for "BioFabric Visualization of Network Alignments": VISNAB Source Code: LICENSE

		  GNU LESSER GENERAL PUBLIC LICENSE
		       Version 2.1, February 1999

 Copyright (C) 1991, 1999 Free Software Foundation, Inc.
     59 Temple Place, Suite 330, Boston, MA  02111-1307  USA
 Everyone is permitted to copy and distribute verbatim copies
 of this license document, but changing it is not allowed.

[This is the first released version of the Lesser GPL.  It also counts
 as the successor of the GNU Library Public License, version 2, hence
 the version number 2.1.]

			    Preamble

  The licenses for most software are designed to take away your
freedom to share and change it.  By contrast, the GNU General Public
Licenses are intended to guarantee your freedom to share and change
free software--to make sure the software is free for all its users.

  This license, the Lesser General Public License, applies to some
specially designated software packages--typically libraries--of the
Free Software Foundation and other authors who decide to use it.  You
can use it too, but we suggest you first think carefully about whether
this license or the ordinary General Public License is the better
strategy to use in any particular case, based on the explanations below.

  When we speak of free software, we are referring to freedom of use,
not price.  Our General Public Licenses are designed to make sure that
you have the freedom to distribute copies of free software (and charge
for this service if you wish); that you receive source code or can get
it if you want it; that you can change the software and use pieces of
it in new free programs; and that you are informed that you can do
these things.

  To protect your rights, we need to make restrictions that forbid
distributors to deny you these rights or to ask you to surrender these
rights.  These restrictions translate to certain responsibilities for
you if you distribute copies of the library or if you modify it.

  For example, if you distribute copies of the library, whether gratis
or for a fee, you must give the recipients all the rights that we gave
you.  You must make sure that they, too, receive or can get the source
code.  If you link other code with the library, you must provide
complete object files to the recipients, so that they can relink them
with the library after making changes to the library and recompiling
it.  And you must show them these terms so they know their rights.

  We protect your rights with a two-step method: (1) we copyright the
library, and (2) we offer you this license, which gives you legal
permission to copy, distribute and/or modify the library.

  To protect each distributor, we want to make it very clear that
there is no warranty for the free library.  Also, if the library is
modified by someone else and passed on, the recipients should know
that what they have is not the original version, so that the original
author's reputation will not be affected by problems that might be
introduced by others.

  Finally, software patents pose a constant threat to the existence of
any free program.  We wish to make sure that a company cannot
effectively restrict the users of a free program by obtaining a
restrictive license from a patent holder.  Therefore, we insist that
any patent license obtained for a version of the library must be
consistent with the full freedom of use specified in this license.

  Most GNU software, including some libraries, is covered by the
ordinary GNU General Public License.  This license, the GNU Lesser
General Public License, applies to certain designated libraries, and
is quite different from the ordinary General Public License.  We use
this license for certain libraries in order to permit linking those
libraries into non-free programs.

  When a program is linked with a library, whether statically or using
a shared library, the combination of the two is legally speaking a
combined work, a derivative of the original library.  The ordinary
General Public License therefore permits such linking only if the
entire combination fits its criteria of freedom.  The Lesser General
Public License permits more lax criteria for linking other code with
the library.

  We call this license the "Lesser" General Public License because it
does Less to protect the user's freedom than the ordinary General
Public License.  It also provides other free software developers Less
of an advantage over competing non-free programs.  These disadvantages
are the reason we use the ordinary General Public License for many
libraries.  However, the Lesser license provides advantages in certain
special circumstances.

  For example, on rare occasions, there may be a special need to
encourage the widest possible use of a certain library, so that it becomes
a de-facto standard.  To achieve this, non-free programs must be
allowed to use the library.  A more frequent case is that a free
library does the same job as widely used non-free libraries.  In this
case, there is little to gain by limiting the free library to free
software only, so we use the Lesser General Public License.

  In other cases, permission to use a particular library in non-free
programs enables a greater number of people to use a large body of
free software.  For example, permission to use the GNU C Library in
non-free programs enables many more people to use the whole GNU
operating system, as well as its variant, the GNU/Linux operating
system.

  Although the Lesser General Public License is Less protective of the
users' freedom, it does ensure that the user of a program that is
linked with the Library has the freedom and the wherewithal to run
that program using a modified version of the Library.

  The precise terms and conditions for copying, distribution and
modification follow.  Pay close attention to the difference between a
"work based on the library" and a "work that uses the library".  The
former contains code derived from the library, whereas the latter must
be combined with the library in order to run.

		  GNU LESSER GENERAL PUBLIC LICENSE
   TERMS AND CONDITIONS FOR COPYING, DISTRIBUTION AND MODIFICATION

  0. This License Agreement applies to any software library or other
program which contains a notice placed by the copyright holder or
other authorized party saying it may be distributed under the terms of
this Lesser General Public License (also called "this License").
Each licensee is addressed as "you".

  A "library" means a collection of software functions and/or data
prepared so as to be conveniently linked with application programs
(which use some of those functions and data) to form executables.

  The "Library", below, refers to any such software library or work
which has been distributed under these terms.  A "work based on the
Library" means either the Library or any derivative work under
copyright law: that is to say, a work containing the Library or a
portion of it, either verbatim or with modifications and/or translated
straightforwardly into another language.  (Hereinafter, translation is
included without limitation in the term "modification".)

  "Source code" for a work means the preferred form of the work for
making modifications to it.  For a library, complete source code means
all the source code for all modules it contains, plus any associated
interface definition files, plus the scripts used to control compilation
and installation of the library.

  Activities other than copying, distribution and modification are not
covered by this License; they are outside its scope.  The act of
running a program using the Library is not restricted, and output from
such a program is covered only if its contents constitute a work based
on the Library (independent of the use of the Library in a tool for
writing it).  Whether that is true depends on what the Library does
and what the program that uses the Library does.
  
  1. You may copy and distribute verbatim copies of the Library's
complete source code as you receive it, in any medium, provided that
you conspicuously and appropriately publish on each copy an
appropriate copyright notice and disclaimer of warranty; keep intact
all the notices that refer to this License and to the absence of any
warranty; and distribute a copy of this License along with the
Library.

  You may charge a fee for the physical act of transferring a copy,
and you may at your option offer warranty protection in exchange for a
fee.

  2. You may modify your copy or copies of the Library or any portion
of it, thus forming a work based on the Library, and copy and
distribute such modifications or work under the terms of Section 1
above, provided that you also meet all of these conditions:

    a) The modified work must itself be a software library.

    b) You must cause the files modified to carry prominent notices
    stating that you changed the files and the date of any change.

    c) You must cause the whole of the work to be licensed at no
    charge to all third parties under the terms of this License.

    d) If a facility in the modified Library refers to a function or a
    table of data to be supplied by an application program that uses
    the facility, other than as an argument passed when the facility
    is invoked, then you must make a good faith effort to ensure that,
    in the event an application does not supply such function or
    table, the facility still operates, and performs whatever part of
    its purpose remains meaningful.

    (For example, a function in a library to compute square roots has
    a purpose that is entirely well-defined independent of the
    application.  Therefore, Subsection 2d requires that any
    application-supplied function or table used by this function must
    be optional: if the application does not supply it, the square
    root function must still compute square roots.)

These requirements apply to the modified work as a whole.  If
identifiable sections of that work are not derived from the Library,
and can be reasonably considered independent and separate works in
themselves, then this License, and its terms, do not apply to those
sections when you distribute them as separate works.  But when you
distribute the same sections as part of a whole which is a work based
on the Library, the distribution of the whole must be on the terms of
this License, whose permissions for other licensees extend to the
entire whole, and thus to each and every part regardless of who wrote
it.

Thus, it is not the intent of this section to claim rights or contest
your rights to work written entirely by you; rather, the intent is to
exercise the right to control the distribution of derivative or
collective works based on the Library.

In addition, mere aggregation of another work not based on the Library
with the Library (or with a work based on the Library) on a volume of
a storage or distribution medium does not bring the other work under
the scope of this License.

  3. You may opt to apply the terms of the ordinary GNU General Public
License instead of this License to a given copy of the Library.  To do
this, you must alter all the notices that refer to this License, so
that they refer to the ordinary GNU General Public License, version 2,
instead of to this License.  (If a newer version than version 2 of the
ordinary GNU General Public License has appeared, then you can specify
that version instead if you wish.)  Do not make any other change in
these notices.

  Once this change is made in a given copy, it is irreversible for
that copy, so the ordinary GNU General Public License applies to all
subsequent copies and derivative works made from that copy.

  This option is useful when you wish to copy part of the code of
the Library into a program that is not a library.

  4. You may copy and distribute the Library (or a portion or
derivative of it, under Section 2) in object code or executable form
under the terms of Sections 1 and 2 above provided that you accompany
it with the complete corresponding machine-readable source code, which
must be distributed under the terms of Sections 1 and 2 above on a
medium customarily used for software interchange.

  If distribution of object code is made by offering access to copy
from a designated place, then offering equivalent access to copy the
source code from the same place satisfies the requirement to
distribute the source code, even though third parties are not
compelled to copy the source along with the object code.

  5. A program that contains no derivative of any portion of the
Library, but is designed to work with the Library by being compiled or
linked with it, is called a "work that uses the Library".  Such a
work, in isolation, is not a derivative work of the Library, and
therefore falls outside the scope of this License.

  However, linking a "work that uses the Library" with the Library
creates an executable that is a derivative of the Library (because it
contains portions of the Library), rather than a "work that uses the
library".  The executable is therefore covered by this License.
Section 6 states terms for distribution of such executables.

  When a "work that uses the Library" uses material from a header file
that is part of the Library, the object code for the work may be a
derivative work of the Library even though the source code is not.
Whether this is true is especially significant if the work can be
linked without the Library, or if the work is itself a library.  The
threshold for this to be true is not precisely defined by law.

  If such an object file uses only numerical parameters, data
structure layouts and accessors, and small macros and small inline
functions (ten lines or less in length), then the use of the object
file is unrestricted, regardless of whether it is legally a derivative
work.  (Executables containing this object code plus portions of the
Library will still fall under Section 6.)

  Otherwise, if the work is a derivative of the Library, you may
distribute the object code for the work under the terms of Section 6.
Any executables containing that work also fall under Section 6,
whether or not they are linked directly with the Library itself.

  6. As an exception to the Sections above, you may also combine or
link a "work that uses the Library" with the Library to produce a
work containing portions of the Library, and distribute that work
under terms of your choice, provided that the terms permit
modification of the work for the customer's own use and reverse
engineering for debugging such modifications.

  You must give prominent notice with each copy of the work that the
Library is used in it and that the Library and its use are covered by
this License.  You must supply a copy of this License.  If the work
during execution displays copyright notices, you must include the
copyright notice for the Library among them, as well as a reference
directing the user to the copy of this License.  Also, you must do one
of these things:

    a) Accompany the work with the complete corresponding
    machine-readable source code for the Library including whatever
    changes were used in the work (which must be distributed under
    Sections 1 and 2 above); and, if the work is an executable linked
    with the Library, with the complete machine-readable "work that
    uses the Library", as object code and/or source code, so that the
    user can modify the Library and then relink to produce a modified
    executable containing the modified Library.  (It is understood
    that the user who changes the contents of definitions files in the
    Library will not necessarily be able to recompile the application
    to use the modified definitions.)

    b) Use a suitable shared library mechanism for linking with the
    Library.  A suitable mechanism is one that (1) uses at run time a
    copy of the library already present on the user's computer system,
    rather than copying library functions into the executable, and (2)
    will operate properly with a modified version of the library, if
    the user installs one, as long as the modified version is
    interface-compatible with the version that the work was made with.

    c) Accompany the work with a written offer, valid for at
    least three years, to give the same user the materials
    specified in Subsection 6a, above, for a charge no more
    than the cost of performing this distribution.

    d) If distribution of the work is made by offering access to copy
    from a designated place, offer equivalent access to copy the above
    specified materials from the same place.

    e) Verify that the user has already received a copy of these
    materials or that you have already sent this user a copy.

  For an executable, the required form of the "work that uses the
Library" must include any data and utility programs needed for
reproducing the executable from it.  However, as a special exception,
the materials to be distributed need not include anything that is
normally distributed (in either source or binary form) with the major
components (compiler, kernel, and so on) of the operating system on
which the executable runs, unless that component itself accompanies
the executable.

  It may happen that this requirement contradicts the license
restrictions of other proprietary libraries that do not normally
accompany the operating system.  Such a contradiction means you cannot
use both them and the Library together in an executable that you
distribute.

  7. You may place library facilities that are a work based on the
Library side-by-side in a single library together with other library
facilities not covered by this License, and distribute such a combined
library, provided that the separate distribution of the work based on
the Library and of the other library facilities is otherwise
permitted, and provided that you do these two things:

    a) Accompany the combined library with a copy of the same work
    based on the Library, uncombined with any other library
    facilities.  This must be distributed under the terms of the
    Sections above.

    b) Give prominent notice with the combined library of the fact
    that part of it is a work based on the Library, and explaining
    where to find the accompanying uncombined form of the same work.

  8. You may not copy, modify, sublicense, link with, or distribute
the Library except as expressly provided under this License.  Any
attempt otherwise to copy, modify, sublicense, link with, or
distribute the Library is void, and will automatically terminate your
rights under this License.  However, parties who have received copies,
or rights, from you under this License will not have their licenses
terminated so long as such parties remain in full compliance.

  9. You are not required to accept this License, since you have not
signed it.  However, nothing else grants you permission to modify or
distribute the Library or its derivative works.  These actions are
prohibited by law if you do not accept this License.  Therefore, by
modifying or distributing the Library (or any work based on the
Library), you indicate your acceptance of this License to do so, and
all its terms and conditions for copying, distributing or modifying
the Library or works based on it.

  10. Each time you redistribute the Library (or any work based on the
Library), the recipient automatically receives a license from the
original licensor to copy, distribute, link with or modify the Library
subject to these terms and conditions.  You may not impose any further
restrictions on the recipients' exercise of the rights granted herein.
You are not responsible for enforcing compliance by third parties with
this License.

  11. If, as a consequence of a court judgment or allegation of patent
infringement or for any other reason (not limited to patent issues),
conditions are imposed on you (whether by court order, agreement or
otherwise) that contradict the conditions of this License, they do not
excuse you from the conditions of this License.  If you cannot
distribute so as to satisfy simultaneously your obligations under this
License and any other pertinent obligations, then as a consequence you
may not distribute the Library at all.  For example, if a patent
license would not permit royalty-free redistribution of the Library by
all those who receive copies directly or indirectly through you, then
the only way you could satisfy both it and this License would be to
refrain entirely from distribution of the Library.

If any portion of this section is held invalid or unenforceable under any
particular circumstance, the balance of the section is intended to apply,
and the section as a whole is intended to apply in other circumstances.

It is not the purpose of this section to induce you to infringe any
patents or other property right claims or to contest validity of any
such claims; this section has the sole purpose of protecting the
integrity of the free software distribution system which is
implemented by public license practices.  Many people have made
generous contributions to the wide range of software distributed
through that system in reliance on consistent application of that
system; it is up to the author/donor to decide if he or she is willing
to distribute software through any other system and a licensee cannot
impose that choice.

This section is intended to make thoroughly clear what is believed to
be a consequence of the rest of this License.

  12. If the distribution and/or use of the Library is restricted in
certain countries either by patents or by copyrighted interfaces, the
original copyright holder who places the Library under this License may add
an explicit geographical distribution limitation excluding those countries,
so that distribution is permitted only in or among countries not thus
excluded.  In such case, this License incorporates the limitation as if
written in the body of this License.

  13. The Free Software Foundation may publish revised and/or new
versions of the Lesser General Public License from time to time.
Such new versions will be similar in spirit to the present version,
but may differ in detail to address new problems or concerns.

Each version is given a distinguishing version number.  If the Library
specifies a version number of this License which applies to it and
"any later version", you have the option of following the terms and
conditions either of that version or of any later version published by
the Free Software Foundation.  If the Library does not specify a
license version number, you may choose any version ever published by
the Free Software Foundation.

  14. If you wish to incorporate parts of the Library into other free
programs whose distribution conditions are incompatible with these,
write to the author to ask for permission.  For software which is
copyrighted by the Free Software Foundation, write to the Free
Software Foundation; we sometimes make exceptions for this.  Our
decision will be guided by the two goals of preserving the free status
of all derivatives of our free software and of promoting the sharing
and reuse of software generally.

			    NO WARRANTY

  15. BECAUSE THE LIBRARY IS LICENSED FREE OF CHARGE, THERE IS NO
WARRANTY FOR THE LIBRARY, TO THE EXTENT PERMITTED BY APPLICABLE LAW.
EXCEPT WHEN OTHERWISE STATED IN WRITING THE COPYRIGHT HOLDERS AND/OR
OTHER PARTIES PROVIDE THE LIBRARY "AS IS" WITHOUT WARRANTY OF ANY
KIND, EITHER EXPRESSED OR IMPLIED, INCLUDING, BUT NOT LIMITED TO, THE
IMPLIED WARRANTIES OF MERCHANTABILITY AND FITNESS FOR A PARTICULAR
PURPOSE.  THE ENTIRE RISK AS TO THE QUALITY AND PERFORMANCE OF THE
LIBRARY IS WITH YOU.  SHOULD THE LIBRARY PROVE DEFECTIVE, YOU ASSUME
THE COST OF ALL NECESSARY SERVICING, REPAIR OR CORRECTION.

  16. IN NO EVENT UNLESS REQUIRED BY APPLICABLE LAW OR AGREED TO IN
WRITING WILL ANY COPYRIGHT HOLDER, OR ANY OTHER PARTY WHO MAY MODIFY
AND/OR REDISTRIBUTE THE LIBRARY AS PERMITTED ABOVE, BE LIABLE TO YOU
FOR DAMAGES, INCLUDING ANY GENERAL, SPECIAL, INCIDENTAL OR
CONSEQUENTIAL DAMAGES ARISING OUT OF THE USE OR INABILITY TO USE THE
LIBRARY (INCLUDING BUT NOT LIMITED TO LOSS OF DATA OR DATA BEING
RENDERED INACCURATE OR LOSSES SUSTAINED BY YOU OR THIRD PARTIES OR A
FAILURE OF THE LIBRARY TO OPERATE WITH ANY OTHER SOFTWARE), EVEN IF
SUCH HOLDER OR OTHER PARTY HAS BEEN ADVISED OF THE POSSIBILITY OF SUCH
DAMAGES.

		     END OF TERMS AND CONDITIONS

           How to Apply These Terms to Your New Libraries

  If you develop a new library, and you want it to be of the greatest
possible use to the public, we recommend making it free software that
everyone can redistribute and change.  You can do so by permitting
redistribution under these terms (or, alternatively, under the terms of the
ordinary General Public License).

  To apply these terms, attach the following notices to the library.  It is
safest to attach them to the start of each source file to most effectively
convey the exclusion of warranty; and each file should have at least the
"copyright" line and a pointer to where the full notice is found.

    <one line to give the library's name and a brief idea of what it does.>
    Copyright (C) <year>  <name of author>

    This library is free software; you can redistribute it and/or
    modify it under the terms of the GNU Lesser General Public
    License as published by the Free Software Foundation; either
    version 2.1 of the License, or (at your option) any later version.

    This library is distributed in the hope that it will be useful,
    but WITHOUT ANY WARRANTY; without even the implied warranty of
    MERCHANTABILITY or FITNESS FOR A PARTICULAR PURPOSE.  See the GNU
    Lesser General Public License for more details.

    You should have received a copy of the GNU Lesser General Public
    License along with this library; if not, write to the Free Software
    Foundation, Inc., 59 Temple Place, Suite 330, Boston, MA  02111-1307  USA

Also add information on how to contact you by electronic and paper mail.

You should also get your employer (if you work as a programmer) or your
school, if any, to sign a "copyright disclaimer" for the library, if
necessary.  Here is a sample; alter the names:

  Yoyodyne, Inc., hereby disclaims all copyright interest in the
  library `Frob' (a library for tweaking knobs) written by James Random Hacker.

  <signature of Ty Coon>, 1 April 1990
  Ty Coon, President of Vice

That's all there is to it!
