## Supplementary material for "BioFabric Visualization of Network Alignments": VISNAB Source Code: NetAlignMeasureDialog.java

```
/*
**    Copyright (C) 2018 Rishi Desai
**
**    Copyright (C) 2003-2018 Institute for Systems Biology
**                            Seattle, Washington, USA.
**
**    This library is free software; you can redistribute it and/or
**    modify it under the terms of the GNU Lesser General Public
**    License as published by the Free Software Foundation; either
**    version 2.1 of the License, or (at your option) any later version.
**
**    This library is distributed in the hope that it will be useful,
**    but WITHOUT ANY WARRANTY; without even the implied warranty of
**    MERCHANTABILITY or FITNESS FOR A PARTICULAR PURPOSE.  See the GNU
**    Lesser General Public License for more details.
**
**    You should have received a copy of the GNU Lesser General Public
**    License along with this library; if not, write to the Free Software
**    Foundation, Inc., 59 Temple Place, Suite 330, Boston, MA  02111-1307  USA
*/

package org.systemsbiology.biofabric.plugin.core.align;

import java.awt.Dimension;
import java.awt.GridBagLayout;
import javax.swing.JFrame;
import javax.swing.JLabel;
import javax.swing.JPanel;
import javax.swing.SwingConstants;
import javax.swing.border.EmptyBorder;

import org.systemsbiology.biofabric.api.dialog.BTStashResultsDialog;
import org.systemsbiology.biofabric.api.dialog.DialogObj;
import org.systemsbiology.biofabric.api.util.PluginResourceManager;

public class NetAlignMeasureDialog extends BTStashResultsDialog {
  
  ////////////////////////////////////////////////////////////////////////////
  //
  // PRIVATE FIELDS
  //
  ////////////////////////////////////////////////////////////////////////////
  
	private static final long serialVersionUID = 1L;
	
  private JFrame parent_;
  private NetworkAlignmentPlugIn.NetAlignStats netAlignStats_;
  
  ////////////////////////////////////////////////////////////////////////////
  //
  // CONSTRUCTOR
  //
  ////////////////////////////////////////////////////////////////////////////
  
  public NetAlignMeasureDialog(JFrame parent, NetworkAlignmentPlugIn.NetAlignStats stats, PluginResourceManager rMan) {
    super(parent, rMan.getPluginString("networkAlignment.measures"), new Dimension(700, 400), 2);
    
    this.parent_ = parent;
    this.netAlignStats_ = stats;
  
    JPanel cp = (JPanel) getContentPane();
    cp.setBorder(new EmptyBorder(20, 20, 20, 20));
    cp.setLayout(new GridBagLayout());
 
    String msg = rMan.getPluginString("networkAlignment.measureMessage");
    addWidgetFullRow(new JLabel(msg), false);
    
    for (NetworkAlignmentPlugIn.NetAlignMeasure measure : netAlignStats_.getMeasures()) {
      String label = String.format("%s=\t%4.4f", measure.name, measure.val);
      addWidgetFullRow(new JLabel(label, SwingConstants.LEFT), false);
    }
    
    if (!netAlignStats_.hasStats()) {
      // should not happen because all topological measures are always calculable
      String noM = rMan.getPluginString("networkAlignment.noMeasuresAvailable");
      addWidgetFullRow(new JLabel(noM), false);
    }
    
    DialogObj.Buttons buttons = finishConstruction();
    buttons.cancelButton.setVisible(false);
  
    setLocationRelativeTo(parent);
  }
  
  @Override
  protected boolean stashForOK() {
    return (true);
  }
}
```
