## Supplementary material for "BioFabric Visualization of Network Alignments": VISNAB Source Code: NetworkAlignment.java

```
/*
**
**    Copyright (C) 2016-2019 Rishi Desai
**
**    Copyright (C) 2003-2018 Institute for Systems Biology
**                            Seattle, Washington, USA.
**
**    This library is free software; you can redistribute it and/or
**    modify it under the terms of the GNU Lesser General Public
**    License as published by the Free Software Foundation; either
**    version 2.1 of the License, or (at your option) any later version.
**
**    This library is distributed in the hope that it will be useful,
**    but WITHOUT ANY WARRANTY; without even the implied warranty of
**    MERCHANTABILITY or FITNESS FOR A PARTICULAR PURPOSE.  See the GNU
**    Lesser General Public License for more details.
**
**    You should have received a copy of the GNU Lesser General Public
**    License along with this library; if not, write to the Free Software
**    Foundation, Inc., 59 Temple Place, Suite 330, Boston, MA  02111-1307  USA
*/

package org.systemsbiology.biofabric.plugin.core.align;

import java.util.ArrayList;
import java.util.Arrays;
import java.util.Collections;
import java.util.Comparator;
import java.util.HashMap;
import java.util.HashSet;
import java.util.List;
import java.util.Map;
import java.util.Set;
import java.util.TreeSet;
import org.systemsbiology.biofabric.api.model.NetLink;
import org.systemsbiology.biofabric.api.model.NetNode;
import org.systemsbiology.biofabric.api.util.NID;
import org.systemsbiology.biofabric.api.util.UniqueLabeller;
import org.systemsbiology.biofabric.api.worker.AsynchExitRequestException;
import org.systemsbiology.biofabric.api.worker.BTProgressMonitor;
import org.systemsbiology.biofabric.api.worker.LoopReporter;
import org.systemsbiology.biofabric.io.BuildExtractorImpl;
import org.systemsbiology.biofabric.plugin.PluginSupportFactory;

/***************************************************************************
 **
 ** This class merges two individual graphs G1 and G2 with an alignment to form the
 ** network-alignment network.
 **
 ** LG = LINK GROUP
 ** NG = NODE GROUP
 **
 ** FIRST LG   = COVERED EDGE             (PURPLE EDGES)
 ** SECOND LG  = INDUCED_GRAPH1           (BLUE EDGES)
 ** THIRD LG   = HALF_ORPHAN_GRAPH1       (BLUE EDGES  - sometimes referred to in speech as CYAN edges)
 ** FOURTH LG  = FULL_ORPHAN_GRAPH1       (BLUE EDGES  - sometimes referred to in speech as GREEN edges)
 ** FIFTH LG   = INDUCED_GRAPH2           (RED EDGES)
 ** SIXTH LG   = HALF_UNALIGNED_GRAPH2    (RED EDGES)  - sometimes referred to in speech as ORANGE edges)
 ** SEVENTH LG = FULL_UNALIGNED_GRAPH2    (RED EDGES)  - sometimes referred to in speech as YELLOW edges)
 **
 ** PURPLE NODE =  ALIGNED G1-G2 NODE - often referred to as MERGED or COMBINED nodes
 ** BLUE NODE   =  UNALIGNED G1 NODE  - often referred to as ORPHAN nodes
 ** RED NODE    =  UNALIGNED G2 NODE
 **
 */

public class NetworkAlignment {
  
  ////////////////////////////////////////////////////////////////////////////
  //
  // PUBLIC STATIC MEMBERS
  //
  ////////////////////////////////////////////////////////////////////////////
  
  public enum EdgeType {
    COVERED("P", 0),
    INDUCED_GRAPH1("pBp", 1), HALF_ORPHAN_GRAPH1("pBb", 2), FULL_ORPHAN_GRAPH1("bBb", 3),
    INDUCED_GRAPH2("pRp", 4), HALF_UNALIGNED_GRAPH2("pRr", 5), FULL_UNALIGNED_GRAPH2("rRr", 6);
    
    public final String tag;
    public final int index;
    
    EdgeType(String tag, int index) {
      this.tag = tag;
      this.index = index;
    }
    
  }
  
  public static final EdgeType[] LINK_GROUPS = {
          EdgeType.COVERED,
          EdgeType.INDUCED_GRAPH1, EdgeType.HALF_ORPHAN_GRAPH1, EdgeType.FULL_ORPHAN_GRAPH1,
          EdgeType.INDUCED_GRAPH2, EdgeType.HALF_UNALIGNED_GRAPH2, EdgeType.FULL_UNALIGNED_GRAPH2
  };
  
  public enum NodeColor {
    PURPLE("P"), BLUE("B"), RED("R");
    
    public final String tag;
    
    NodeColor(String tag) {
      this.tag = tag;
    }
    
  }
  
  public enum GraphType {SMALL, LARGE}
  
  ////////////////////////////////////////////////////////////////////////////
  //
  // PRIVATE INSTANCE MEMBERS
  //
  ////////////////////////////////////////////////////////////////////////////
  
  //
  // G1 is the small (#nodes) network, G2 is the large network
  //
  
  private Map<NetNode, NetNode> mapG1toG2_, perfectG1toG2_;
  private ArrayList<NetLink> linksG1_;
  private HashSet<NetNode> lonersG1_;
  private ArrayList<NetLink> linksG2_;
  private HashSet<NetNode> lonersG2_;
  private NetworkAlignmentBuildData.ViewType outType_;
  private UniqueLabeller idGen_;
  private BTProgressMonitor monitor_;
  private final String TEMPORARY = "TEMP";
  
  //
  // largeToMergedID only contains aligned nodes
  //
  
  private Map<NetNode, NetNode> smallToMergedID_, largeToMergedID_;
  private Map<NetNode, NetNode> mergedIDToSmall_;
  private Map<NetNode, NetNode> smallToUnmergedID_, largeToUnmergedID_;
  
  //
  // mergedToCorrect only has aligned nodes
  //
  
  private ArrayList<NetLink> mergedLinks_;
  private Set<NetNode> mergedLoners_;
  private Map<NetNode, Boolean> mergedToCorrectNC_;
  private NodeColorMap nodeColorMap_;
  
  ////////////////////////////////////////////////////////////////////////////
  //
  // PUBLIC CONSTRUCTORS
  //
  ////////////////////////////////////////////////////////////////////////////
  
  public NetworkAlignment(ArrayList<NetLink> mergedLinks, Set<NetNode> mergedLoneNodeIDs,
                          Map<NetNode, NetNode> mapG1toG2, Map<NetNode, NetNode> perfectG1toG2_,
                          ArrayList<NetLink> linksG1, HashSet<NetNode> lonersG1,
                          ArrayList<NetLink> linksG2, HashSet<NetNode> lonersG2,
                          Map<NetNode, Boolean> mergedToCorrectNC, NodeColorMap nodeColorMap,
                          NetworkAlignmentBuildData.ViewType outType, UniqueLabeller idGen, BTProgressMonitor monitor) {
    
    this.mapG1toG2_ = mapG1toG2;
    this.perfectG1toG2_ = perfectG1toG2_;
    this.linksG1_ = linksG1;
    this.lonersG1_ = lonersG1;
    this.linksG2_ = linksG2;
    this.lonersG2_ = lonersG2;
    this.outType_ = outType;
    this.idGen_ = idGen;
    this.monitor_ = monitor;
    
    this.mergedLinks_ = mergedLinks;
    this.mergedLoners_ = mergedLoneNodeIDs;
    this.mergedToCorrectNC_ = mergedToCorrectNC;
    this.nodeColorMap_ = nodeColorMap;
  
    this.smallToUnmergedID_ = new HashMap<NetNode, NetNode>();
    this.largeToUnmergedID_ = new HashMap<NetNode, NetNode>();
  }
  
  /****************************************************************************
   **
   ** Merge the Network!
   */
  
  public void mergeNetworks() throws AsynchExitRequestException {
    
    //
    // Create merged and unmerged nodes and Correctness
    //
    
    createMergedNodes();
    createUnmergedNodes(GraphType.SMALL);
    createUnmergedNodes(GraphType.LARGE);
    
    //
    // Create individual link sets; "old" refers to pre-merged networks, "new" is merged network
    //
    
    List<NetLink> newLinksG1 = new ArrayList<NetLink>();
    Set<NetNode> newLonersG1 = new HashSet<NetNode>();
    
    createNewLinkList(newLinksG1, newLonersG1, GraphType.SMALL);
    
    List<NetLink> newLinksG2 = new ArrayList<NetLink>();
    Set<NetNode> newLonersG2 = new HashSet<NetNode>();
    
    createNewLinkList(newLinksG2, newLonersG2, GraphType.LARGE);
    
    //
    // Give each link its respective link relation
    //
    
    createMergedLinkList(newLinksG1, newLinksG2);
    
    finalizeLoneNodeIDs(newLonersG1, newLonersG2);
    
    //
    // POST processing
    //
    
    createNodeColorMap(newLinksG1, newLonersG1, newLinksG2, newLonersG2);
    
    //
    // Orphan Edges: All unaligned edges; plus all of their endpoint nodes' edges
    //
    
    if (outType_ == NetworkAlignmentBuildData.ViewType.ORPHAN) {
      (new OrphanEdgeLayout()).process(mergedLinks_, mergedLoners_, monitor_);
    }
    
    return;
  }
  
  ////////////////////////////////////////////////////////////////////////////
  //
  // PRIVATE METHODS
  //
  ////////////////////////////////////////////////////////////////////////////
  
  /****************************************************************************
   **
   ** Create merged nodes (Purple), install into maps; Correctness for Purple nodes
   */
  
  private void createMergedNodes() {
    
    smallToMergedID_ = new HashMap<NetNode, NetNode>();
    largeToMergedID_ = new HashMap<NetNode, NetNode>();
    mergedIDToSmall_ = new HashMap<NetNode, NetNode>();
    
    boolean doingPerfectGroup = (outType_ == NetworkAlignmentBuildData.ViewType.GROUP) &&
                                (perfectG1toG2_ != null);
     
    for (Map.Entry<NetNode, NetNode> entry : mapG1toG2_.entrySet()) {
      
      NetNode smallNode = entry.getKey(), largeNode = entry.getValue();
      String smallName = smallNode.getName(), largeName = largeNode.getName();
      
      //
      // Aligned nodes merge name in the form small::large
      //
      
      String mergedName = String.format("%s::%s", smallName, largeName);
      
      NID nid = idGen_.getNextOID();
      NetNode merged_node = PluginSupportFactory.buildNode(nid, mergedName);
      
      smallToMergedID_.put(smallNode, merged_node);
      largeToMergedID_.put(largeNode, merged_node);
      mergedIDToSmall_.put(merged_node, smallNode);
      
      //
      // Nodes are correctly aligned map
      //
      
      if (doingPerfectGroup) { // perfect alignment must be provided
        NetNode perfectLarge = perfectG1toG2_.get(smallNode);
        boolean alignedCorrect = (perfectLarge != null) && perfectLarge.equals(largeNode);
        mergedToCorrectNC_.put(merged_node, alignedCorrect);
      }
    }
    return;
  }
  
  /****************************************************************************
   **
   ** Create unmerged nodes (Blue or Red), install into maps; Correctness for Blue nodes
   */
  
  private void createUnmergedNodes(GraphType type) throws AsynchExitRequestException {
  
    boolean doingPerfectGroup = (outType_ == NetworkAlignmentBuildData.ViewType.GROUP) &&
            (perfectG1toG2_ != null);
    Map<NetNode, NetNode> oldToUnmerged, oldToMerged;
    Set<NetNode> nodes;
    switch (type) {
      case SMALL:
        nodes = PluginSupportFactory.getBuildExtractor().extractNodes(linksG1_, lonersG1_, monitor_);
        oldToMerged = smallToMergedID_;
        oldToUnmerged = smallToUnmergedID_;
        break;
      case LARGE:
        nodes = PluginSupportFactory.getBuildExtractor().extractNodes(linksG2_, lonersG2_, monitor_);
        oldToMerged = largeToMergedID_;
        oldToUnmerged = largeToUnmergedID_;
        break;
      default:
        throw (new IllegalArgumentException("Incorrect graph type"));
    }
  
    for (NetNode node : nodes) {
      if (oldToMerged.get(node) != null) {
        continue;
      }
      NetNode unalignedNewNode = modifyName(node, type);
      oldToUnmerged.put(node, unalignedNewNode);
      
      // We are dealing with Blue nodes, so if perfect alignment is not aligning
      // the node either, it is correct
  
      if (type == GraphType.SMALL && doingPerfectGroup) { // perfect alignment must be provided
        NetNode perfectLarge = perfectG1toG2_.get(node);
        boolean unalignedCorrectly = (perfectLarge == null);
        mergedToCorrectNC_.put(unalignedNewNode, unalignedCorrectly);
      }
    }
    return;
  }
  
  /****************************************************************************
   **
   ** Add the accompanying '::' after Blue nodes and before Red nodes
   */
  
  private NetNode modifyName(NetNode node, GraphType type) {
    
    NID newID = idGen_.getNextOID();
    String newName;
    if (type == NetworkAlignment.GraphType.SMALL) {
      newName = String.format("%s::", node.getName());  // A:: for blue nodes
    } else if (type == NetworkAlignment.GraphType.LARGE) {
      newName = String.format("::%s", node.getName());  // ::B for red nodes
    } else {
      throw (new IllegalArgumentException("Incorrect graph type"));
    }
    return (PluginSupportFactory.buildNode(newID, newName));
  }
  
  /****************************************************************************
   **
   ** Create new link lists based on merged nodes for both networks
   */
  
  private void createNewLinkList(List<NetLink> newLinks, Set<NetNode> newLoners, GraphType type)
          throws AsynchExitRequestException {
    
    List<NetLink> oldLinks;
    Set<NetNode> oldLoners;
    Map<NetNode, NetNode> oldToNewMergedID, oldToNewUnmergedID;
    String msg;
    
    switch (type) {
      case SMALL:
        oldLinks = linksG1_;
        oldLoners = lonersG1_;
        oldToNewMergedID = smallToMergedID_;
        oldToNewUnmergedID = smallToUnmergedID_;
        msg = "progress.mergingSmallLinks";
        break;
      case LARGE:
        oldLinks = linksG2_;
        oldLoners = lonersG2_;
        oldToNewMergedID = largeToMergedID_;
        oldToNewUnmergedID = largeToUnmergedID_;
        msg = "progress.mergingLargeLinks";
        break;
      default:
        throw new IllegalArgumentException();
    }
    
    LoopReporter lr = new LoopReporter(oldLinks.size(), 20, monitor_, 0.0, 1.0, msg);
    Set<NetLink> newLinkSet = new HashSet<NetLink>();
    
    for (NetLink oldLink : oldLinks) {
      
      NetNode oldA = oldLink.getSrcNode();
      NetNode oldB = oldLink.getTrgNode();
      
      //
      // Not all nodes are mapped in the small or large graph
      //
      
      NetNode newA = (oldToNewMergedID.containsKey(oldA)) ? oldToNewMergedID.get(oldA) : oldToNewUnmergedID.get(oldA);
      NetNode newB = (oldToNewMergedID.containsKey(oldB)) ? oldToNewMergedID.get(oldB) : oldToNewUnmergedID.get(oldB);
      
      NetLink newLink = PluginSupportFactory.buildLink(newA, newB, TEMPORARY, false, Boolean.valueOf(false));
      // 'directed' must be false
      newLinkSet.add(newLink);
      lr.report();
    }
    newLinks.addAll(newLinkSet);
    lr.finish();
    
    for (NetNode oldLoner : oldLoners) {
      
      NetNode newLoner = (oldToNewMergedID.containsKey(oldLoner)) ? oldToNewMergedID.get(oldLoner) : oldToNewUnmergedID.get(oldLoner);
      
      newLoners.add(newLoner);
    }
    return;
  }
  
  /****************************************************************************
   **
   ** Combine the two link lists into one, with G2,CC,G1 tags accordingly
   */
  
  private void createMergedLinkList(List<NetLink> newLinksG1, List<NetLink> newLinksG2)
          throws AsynchExitRequestException {
    
    LoopReporter lr = new LoopReporter(newLinksG2.size(), 20, monitor_, 0.0, 1.0, "progress.separatingLinksA");

    NetAlignFabricLinkLocator comp = new NetAlignFabricLinkLocator();
    sortLinks(newLinksG1);
  
    Set<NetNode> alignedNodesG1 = new HashSet<NetNode>(smallToMergedID_.values());
    Set<NetNode> alignedNodesG2 = new HashSet<NetNode>(largeToMergedID_.values());
    // contains all aligned nodes; contains() works in O(1)
  
    for (NetLink linkG2 : newLinksG2) {
      
      int index = Collections.binarySearch(newLinksG1, linkG2, comp);
      
      NetNode src = linkG2.getSrcNode(), trg = linkG2.getTrgNode();
      
      if (index >= 0) {
        addMergedLink(src, trg, EdgeType.COVERED.tag);
      } else {
        boolean containsSRC = alignedNodesG2.contains(src), containsTRG = alignedNodesG2.contains(trg);
        if (containsSRC && containsTRG) {
          addMergedLink(src, trg, EdgeType.INDUCED_GRAPH2.tag);
        } else if (containsSRC || containsTRG) {
          addMergedLink(src, trg, EdgeType.HALF_UNALIGNED_GRAPH2.tag);
        } else {
          addMergedLink(src, trg, EdgeType.FULL_UNALIGNED_GRAPH2.tag);
        }
      }
      lr.report();
    }
    lr = new LoopReporter(newLinksG1.size(), 20, monitor_, 0.0, 1.0, "progress.separatingLinksB");
    sortLinks(newLinksG2);
    
    for (NetLink linkG1 : newLinksG1) {
      
      int index = Collections.binarySearch(newLinksG2, linkG1, comp);
  
      NetNode src = linkG1.getSrcNode(), trg = linkG1.getTrgNode();
  
      if (index < 0) {
        boolean containsSRC = alignedNodesG1.contains(src), containsTRG = alignedNodesG1.contains(trg);
        if (containsSRC && containsTRG) {
          addMergedLink(src, trg, EdgeType.INDUCED_GRAPH1.tag);
        } else if (containsSRC || containsTRG) {
          addMergedLink(src, trg, EdgeType.HALF_ORPHAN_GRAPH1.tag);
        } else {
          addMergedLink(src, trg, EdgeType.FULL_ORPHAN_GRAPH1.tag);
        }
      }
      lr.report();
    }
    return; // This method is not ideal. . . but shall stay (6/19/18)
  }
  
  /****************************************************************************
   **
   ** Add both non-shadow and shadow links to merged link-list
   */
  
  private void addMergedLink(NetNode src, NetNode trg, String tag) {
    NetLink newMergedLink = PluginSupportFactory.buildLink(src, trg, tag, false);
    mergedLinks_.add(newMergedLink);
    
    // We never create shadow feedback links!
    if (!src.equals(trg)) {
      NetLink newMergedLinkShadow = PluginSupportFactory.buildLink(src, trg, tag, true);
      mergedLinks_.add(newMergedLinkShadow);
    }
    return;
  }
  
  /****************************************************************************
   **
   ** Combine loneNodeIDs lists into one
   */
  
  private void finalizeLoneNodeIDs(Set<NetNode> newLonersG1, Set<NetNode> newLonersG2) {
    mergedLoners_.addAll(newLonersG1);
    mergedLoners_.addAll(newLonersG2);
    return;
  }
  
  /****************************************************************************
   **
   ** POST processing: Create NodeColorMap map
   */
  
  private void createNodeColorMap(List<NetLink> newLinksG1, Set<NetNode> newLonersG1,
                                  List<NetLink> newLinksG2, Set<NetNode> newLonersG2) throws AsynchExitRequestException {
    
    Set<NetNode> nodesG1 = (new BuildExtractorImpl()).extractNodes(newLinksG1, newLonersG1, monitor_);
    Set<NetNode> nodesG2 = (new BuildExtractorImpl()).extractNodes(newLinksG2, newLonersG2, monitor_);
  
    Set<NetNode> alignedNodes = mergedIDToSmall_.keySet();
    Map<NetNode, NodeColor> map = new HashMap<NetNode, NodeColor>();
    
    for (NetNode node : nodesG1) {
      if (alignedNodes.contains(node)) {
        map.put(node, NodeColor.PURPLE);
      } else {
        map.put(node, NodeColor.BLUE);
      }
    }
    for (NetNode node : nodesG2) {
      if (alignedNodes.contains(node)) {
        // essentially re-assigns purple nodes for no reason
        map.put(node, NodeColor.PURPLE);
      } else {
        map.put(node, NodeColor.RED);
      }
    }
    nodeColorMap_.setMap(map);
    return;
  }
  
  /****************************************************************************
   **
   ** Sort list of FabricLinks
   */
  
  private void sortLinks(List<NetLink> newLinks) throws AsynchExitRequestException {
    NetAlignFabricLinkLocator comp = new NetAlignFabricLinkLocator();
    Set<NetLink> sorted = new TreeSet<NetLink>(comp);
    LoopReporter lr = new LoopReporter(newLinks.size(), 20, monitor_, 0.0, 1.0, "progress.sortingLinks");
  
    for (NetLink link : newLinks) {
      sorted.add(link);
      lr.report();
    }
    newLinks.clear();
    newLinks.addAll(sorted);  // assume to have no loop reporter here
    return;
  }
  
  ////////////////////////////////////////////////////////////////////////////
  //
  // INNER CLASSES
  //
  ////////////////////////////////////////////////////////////////////////////
  
  /****************************************************************************
   **
   ** This maps every node in V12 to its color: Blue, Purple, or Red
   */
  
  public static class NodeColorMap {
    
    private Map<NetNode, NodeColor> map;
    
    public NodeColorMap() {}
  
    public void setMap(Map<NetNode, NodeColor> map) {
      this.map = map;
    }
  
    public NodeColor getColor(NetNode node) {
      return (map.get(node));
    }
    
  }
  
  /****************************************************************************
   **
   ** All unaligned edges plus all of their endpoint nodes' edges
   */
  
  private static class OrphanEdgeLayout {
    
    public OrphanEdgeLayout() {
    }
    
    private void process(List<NetLink> mergedLinks, Set<NetNode> mergedLoneNodeIDs,
                         BTProgressMonitor monitor)
            throws AsynchExitRequestException {
      
      LoopReporter reporter = new LoopReporter(mergedLinks.size(), 20, monitor, 0.0, 1.0,
              "progress.findingOrphanEdges");
      
      Set<NetNode> blueNodesG1 = new TreeSet<NetNode>();
      for (NetLink link : mergedLinks) { // find the nodes of interest
        if (link.getRelation().equals(EdgeType.INDUCED_GRAPH1.tag)) {
          blueNodesG1.add(link.getSrcNode()); // it's a set - so with shadows no duplicates
          blueNodesG1.add(link.getTrgNode());
        }
        reporter.report();
      }
      
      reporter = new LoopReporter(mergedLinks.size(), 20, monitor, 0.0, 1.0,
              "progress.orphanEdgesContext");
      
      List<NetLink> blueEdgesPlusContext = new ArrayList<NetLink>();
      for (NetLink link : mergedLinks) { // add the edges connecting to the nodes of interest (one hop away)
        
        NetNode src = link.getSrcNode(), trg = link.getTrgNode();
        
        if (blueNodesG1.contains(src) || blueNodesG1.contains(trg)) {
          blueEdgesPlusContext.add(link);
        }
        reporter.report();
      }
  
      mergedLinks.clear();
      mergedLoneNodeIDs.clear();
      mergedLinks.addAll(blueEdgesPlusContext);
      return;
    }
    
  }
  
  /***************************************************************************
   **
   ** Used ONLY to order links for creating the merged link set in Network Alignments
   */
  
  private static class NetAlignFabricLinkLocator implements Comparator<NetLink> {
    
    /***************************************************************************
     **
     ** For any different links in the two separate network link sets, this
     ** says which comes first
     */
    
    public int compare(NetLink link1, NetLink link2) {
      
      if (link1.synonymous(link2)) {
        return (0);
      }
      
      //
      // Must sort the node names because A-B must be equivalent to B-A
      //
      
      String[] arr1 = {link1.getSrcNode().getName(), link1.getTrgNode().getName()};
      Arrays.sort(arr1);
      
      String[] arr2 = {link2.getSrcNode().getName(), link2.getTrgNode().getName()};
      Arrays.sort(arr2);
      
      String concat1 = String.format("%s___%s", arr1[0], arr1[1]);
      String concat2 = String.format("%s___%s", arr2[0], arr2[1]);
      
      //
      // This cuts the merge-lists algorithm to O(eloge) because binary search
      //
      
      return concat1.compareTo(concat2);
    }
  }
  
}
```
