## Supplementary material for "BioFabric Visualization of Network Alignments": VISNAB Source Code: NetworkAlignmentBuildData.java

```
/*
**
**    Copyright (C) 2018 Rishi Desai
**
**    Copyright (C) 2003-2018 Institute for Systems Biology
**                            Seattle, Washington, USA.
**
**    This library is free software; you can redistribute it and/or
**    modify it under the terms of the GNU Lesser General Public
**    License as published by the Free Software Foundation; either
**    version 2.1 of the License, or (at your option) any later version.
**
**    This library is distributed in the hope that it will be useful,
**    but WITHOUT ANY WARRANTY; without even the implied warranty of
**    MERCHANTABILITY or FITNESS FOR A PARTICULAR PURPOSE.  See the GNU
**    Lesser General Public License for more details.
**
**    You should have received a copy of the GNU Lesser General Public
**    License along with this library; if not, write to the Free Software
**    Foundation, Inc., 59 Temple Place, Suite 330, Boston, MA  02111-1307  USA
*/

package org.systemsbiology.biofabric.plugin.core.align;

import java.util.ArrayList;
import java.util.HashSet;
import java.util.List;
import java.util.Map;
import java.util.Set;

import org.systemsbiology.biofabric.api.layout.DefaultEdgeLayout;
import org.systemsbiology.biofabric.api.layout.DefaultLayout;
import org.systemsbiology.biofabric.api.layout.EdgeLayout;
import org.systemsbiology.biofabric.api.layout.NodeLayout;
import org.systemsbiology.biofabric.api.model.NetLink;
import org.systemsbiology.biofabric.api.model.NetNode;
import org.systemsbiology.biofabric.plugin.PluginBuildData;

/***************************************************************************
 **
 ** For passing around Network Alignment data
 */

public class NetworkAlignmentBuildData implements PluginBuildData {
  
  public enum ViewType {GROUP, ORPHAN, CYCLE}
  
  //
  // Note the absence of the alignment's link set and lone node set
  // They are both in the BuildData-of-Plugin Object
  //
  
  public NetworkAlignment.NodeColorMap colorMapMain;
  
  public Set<NetLink> allLinksPerfect;
  public Set<NetNode> loneNodeIDsPerfect;
  public NetworkAlignment.NodeColorMap colorMapPerfect;
  
  public Map<NetNode, Boolean> mergedToCorrectNC;
  public NodeGroupMap.PerfectNGMode mode;
  public final Double jaccSimThreshold;
  
  public ArrayList<NetLink> linksSmall, linksLarge;
  public HashSet<NetNode> lonersSmall, lonersLarge;
  public Set<NetNode> allLargerNodes, allSmallerNodes;
  public Map<NetNode, NetNode> mapG1toG2, perfectG1toG2;
  
  public ViewType view;
  public NetworkAlignmentPlugIn.NetAlignStats netAlignStats;
  public List<AlignCycleLayout.CycleBounds> cycleBounds;
  public boolean useNodeGroups; 
  public boolean turnShadowsOn;

  public NetworkAlignmentBuildData(NetworkAlignment.NodeColorMap colorMapMain,
                                   Set<NetLink> allLinksPerfect, Set<NetNode> loneNodeIDsPerfect,
                                   NetworkAlignment.NodeColorMap colorMapPerfect,
                                   Map<NetNode, Boolean> mergedToCorrectNC,
                                   Set<NetNode> allSmallerNodes, ArrayList<NetLink> linksSmall, HashSet<NetNode> loneNodeIDsSmall,
                                   Set<NetNode> allLargerNodes, ArrayList<NetLink> linksLarge, HashSet<NetNode> loneNodeIDsLarge,
                                   Map<NetNode, NetNode> mapG1toG2,
                                   Map<NetNode, NetNode> perfectG1toG2,
                                   NetworkAlignmentPlugIn.NetAlignStats netAlignStats, ViewType view,
                                   NodeGroupMap.PerfectNGMode mode, final Double jaccSimThreshold,
                                   boolean useNodeGroups, boolean turnShadowsOn) {
    this.colorMapMain = colorMapMain;
    this.allLinksPerfect = allLinksPerfect;
    this.loneNodeIDsPerfect = loneNodeIDsPerfect;
    this.colorMapPerfect = colorMapPerfect;
    this.mergedToCorrectNC = mergedToCorrectNC;
    this.allLargerNodes = allLargerNodes;
    this.linksLarge = linksLarge;
    this.lonersLarge = loneNodeIDsLarge;
    this.allSmallerNodes = allSmallerNodes;
    this.linksSmall = linksSmall;
    this.lonersSmall = loneNodeIDsSmall;
    this.mapG1toG2 = mapG1toG2;
    this.perfectG1toG2 = perfectG1toG2;
    
    this.netAlignStats = netAlignStats;
    this.view = view;
    this.mode = mode;
    this.jaccSimThreshold = jaccSimThreshold;
    this.useNodeGroups = useNodeGroups;
    this.turnShadowsOn = turnShadowsOn;
  }

  public NodeLayout getNodeLayout() {
    switch (view) {
      case GROUP:
        return (new NetworkAlignmentLayout());
      case ORPHAN:
        return (new DefaultLayout());
      case CYCLE:
        return (new AlignCycleLayout());
      default:
        throw new IllegalStateException();
    }
  }

  public EdgeLayout getEdgeLayout() {
    switch (view) {
      case GROUP:
        return (new NetworkAlignmentEdgeLayout());
      case ORPHAN:
        return (new DefaultEdgeLayout());
      case CYCLE:
        return (new AlignCycleEdgeLayout());
      default:
        throw new IllegalStateException();
    } 
  }
}
```
