## Supplementary material for "BioFabric Visualization of Network Alignments": VISNAB Source Code: NetworkAlignmentDialog.java

```
/*
**    Copyright (C) 2018 Rishi Desai
**
**    Copyright (C) 2003-2019 Institute for Systems Biology
**                            Seattle, Washington, USA.
**
**    This library is free software; you can redistribute it and/or
**    modify it under the terms of the GNU Lesser General Public
**    License as published by the Free Software Foundation; either
**    version 2.1 of the License, or (at your option) any later version.
**
**    This library is distributed in the hope that it will be useful,
**    but WITHOUT ANY WARRANTY; without even the implied warranty of
**    MERCHANTABILITY or FITNESS FOR A PARTICULAR PURPOSE.  See the GNU
**    Lesser General Public License for more details.
**
**    You should have received a copy of the GNU Lesser General Public
**    License along with this library; if not, write to the Free Software
**    Foundation, Inc., 59 Temple Place, Suite 330, Boston, MA  02111-1307  USA
*/

package org.systemsbiology.biofabric.plugin.core.align;

import java.awt.Dimension;
import java.awt.FlowLayout;
import java.awt.GridBagLayout;
import java.awt.event.ItemEvent;
import java.awt.event.ItemListener;
import javax.swing.JButton;
import javax.swing.JCheckBox;
import javax.swing.JComboBox;
import javax.swing.JFrame;
import javax.swing.JLabel;
import javax.swing.JOptionPane;
import javax.swing.JPanel;
import javax.swing.JTextField;
import javax.swing.SwingConstants;
import javax.swing.event.DocumentEvent;
import javax.swing.event.DocumentListener;

import org.systemsbiology.biofabric.api.dialog.BTStashResultsDialog;
import org.systemsbiology.biofabric.api.dialog.DialogObj;
import org.systemsbiology.biofabric.api.io.FileLoadFlows;
import org.systemsbiology.biofabric.api.util.ExceptionHandler;
import org.systemsbiology.biofabric.api.util.FixedJButton;
import org.systemsbiology.biofabric.api.util.MatchingJLabel;
import org.systemsbiology.biofabric.api.util.PluginResourceManager;
import org.systemsbiology.biofabric.plugin.PluginSupportFactory;

import javax.swing.border.EmptyBorder;
import java.awt.event.ActionEvent;
import java.awt.event.ActionListener;
import java.io.File;

public class NetworkAlignmentDialog extends BTStashResultsDialog {
  
  ////////////////////////////////////////////////////////////////////////////
  //
  // PRIVATE INSTANCE MEMBERS
  //
  ////////////////////////////////////////////////////////////////////////////
  
  private enum FileIndex {
    GRAPH_ONE_FILE, GRAPH_TWO_FILE, ALIGNMENT_FILE, PERFECT_FILE
  }
  
  private static final long serialVersionUID = 1L;
  private final JFrame parent_;
  private final NetworkAlignmentBuildData.ViewType analysisType_;
  private JButton perfectBrowse;
  private MatchingJLabel perfectFileMatch_;
  private JTextField graph1Field_, graph2Field_, alignField_, perfectField_;
  private File graph1File_, graph2File_, alignmentFile_, perfectAlignFile_; // perfect Alignment is optional
  private FixedJButton buttonOK_;
  private JCheckBox undirectedConfirm_;
  private JComboBox perfectNGsCombo_;
  private JLabel jaccSimLabel_;
  private JTextField jaccSimField_;
  private FileLoadFlows flf_;
  private PluginResourceManager rMan_;
  
  private final int NO_PERFECT_IDX = 0, WITH_PERFECT_IDX = 1, NC_IDX = 2, JS_IDX = 3; // indices on combo box
  private final double JACCARD_SIMILARITY_DEFAULT = .75;  // default jaccard similarity threshold
  
  public NetworkAlignmentDialog(JFrame parent, NetworkAlignmentBuildData.ViewType analysisType, 
  		                          String pluginClassName, FileLoadFlows flf, PluginResourceManager rMan) {
    super(parent, rMan.getPluginString("networkAlignment.title"), new Dimension(700, 500), 3);
        
    this.parent_ = parent;
    this.analysisType_ = analysisType;
    this.flf_ = flf;
    this.rMan_ = rMan;
    
    JPanel cp = (JPanel) getContentPane();
    cp.setBorder(new EmptyBorder(20, 20, 20, 20));
    cp.setLayout(new GridBagLayout());
    
    //
    // File Buttons and File Labels
    //
    
    JButton graph1Browse = new JButton(rMan_.getPluginString("networkAlignment.browse"));
    JButton graph2Browse = new JButton(rMan_.getPluginString("networkAlignment.browse"));
    JButton alignmentBrowse = new JButton(rMan_.getPluginString("networkAlignment.browse"));
    perfectBrowse = new JButton(rMan_.getPluginString("networkAlignment.browse"));
    
    initBrowseButtons(graph1Browse, graph2Browse, alignmentBrowse, perfectBrowse);
  
    graph1Field_ = new JTextField(30);
    graph2Field_= new JTextField(30);
    alignField_ = new JTextField(30);
    perfectField_ = new JTextField(30);
    undirectedConfirm_ = new JCheckBox(rMan_.getPluginString("networkAlignment.confirmUndirected"));
    
    initTextFields();

    MatchingJLabel graph1FileMatch, graph2FileMatch, alignFileMatch;
    JLabel jaccSimLabelMatch = new JLabel(rMan_.getPluginString("networkAlignment.jaccardSimilarityLabel")); // only to use as a reference, not in dialog
    perfectFileMatch_ = new MatchingJLabel(rMan_.getPluginString("networkAlignment.perfect"), jaccSimLabelMatch);
    graph1FileMatch = new MatchingJLabel(rMan_.getPluginString("networkAlignment.graph1"), jaccSimLabelMatch);
    graph2FileMatch = new MatchingJLabel(rMan_.getPluginString("networkAlignment.graph2"), jaccSimLabelMatch);
    alignFileMatch = new MatchingJLabel(rMan_.getPluginString("networkAlignment.alignment"), jaccSimLabelMatch);
    
    graph1FileMatch.setHorizontalAlignment(SwingConstants.CENTER);
    graph2FileMatch.setHorizontalAlignment(SwingConstants.CENTER);
    alignFileMatch.setHorizontalAlignment(SwingConstants.CENTER);
    perfectFileMatch_.setHorizontalAlignment(SwingConstants.CENTER);
    
    undirectedConfirm_.addActionListener(new ActionListener() {
      public void actionPerformed(ActionEvent e) {
        try {
          manageOKButton();
        } catch (Exception ex) {
          ExceptionHandler.getHandler().displayException(ex);
        }
      }
    });
    
    JPanel panGraphInfo = new JPanel(new FlowLayout(FlowLayout.LEFT));
    JPanel panGraphInfoTwo = null;
    switch (analysisType_) {
      case ORPHAN:
        panGraphInfo.add(new JLabel(rMan_.getPluginString("networkAlignment.message")));
        break;
      case CYCLE:
        panGraphInfo.add(new JLabel(rMan_.getPluginString("networkAlignment.message")));
        panGraphInfoTwo = new JPanel(new FlowLayout(FlowLayout.LEFT));
        panGraphInfoTwo.add(new JLabel(rMan_.getPluginString("networkAlignment.messageCycleTwo")));
        break;
      case GROUP:
        panGraphInfo.add(new JLabel(rMan_.getPluginString("networkAlignment.message")));
        break;
      default:
        throw new IllegalStateException();
    }
   
    JPanel panGraphConfirm = new JPanel(new FlowLayout(FlowLayout.LEFT));
    panGraphConfirm.add(undirectedConfirm_);
     
    addWidgetFullRow(panGraphInfo, true);
    if (panGraphInfoTwo != null) {
      addWidgetFullRow(panGraphInfoTwo, true);
    }
    addWidgetFullRow(panGraphConfirm, true);
    addLabeledFileBrowse(graph1FileMatch, graph1Field_, graph1Browse);
    addWidgetFullRow(new JLabel(rMan_.getPluginString("networkAlignment.hasBeen"), SwingConstants.CENTER), true);
    addLabeledFileBrowse(graph2FileMatch, graph2Field_, graph2Browse);
    addLabeledFileBrowse(alignFileMatch, alignField_, alignmentBrowse);
  
    JLabel perfectNGLabel = new MatchingJLabel(rMan_.getPluginString("networkAlignment.perfectNodeGroups"), jaccSimLabelMatch);
    perfectNGLabel.setHorizontalAlignment(SwingConstants.CENTER);
    
    int numChoices = (analysisType_ == NetworkAlignmentBuildData.ViewType.CYCLE) ? 2 : 4;
   
    String[] choices = new String[numChoices];
    choices[NO_PERFECT_IDX] = rMan_.getPluginString("networkAlignment.nonePerfect");
    choices[WITH_PERFECT_IDX] = rMan_.getPluginString("networkAlignment.noneWithPerfect");
    if (analysisType_ != NetworkAlignmentBuildData.ViewType.CYCLE) {
    	choices[NC_IDX] = rMan_.getPluginString("networkAlignment.nodeCorrectnessGroupOption");
    	choices[JS_IDX] = rMan_.getPluginString("networkAlignment.jaccardSimilarityGroupOption");
    }
    
    perfectNGsCombo_ = new JComboBox(choices); // have to use unchecked for v1.6
    
    //
    // Perfect File and Combo box functionality
    //
    
    perfectNGsCombo_.setEnabled(true);
    perfectNGsCombo_.setSelectedIndex(NO_PERFECT_IDX);
    perfectNGsCombo_.addItemListener(new ItemListener() {
      public void itemStateChanged(ItemEvent e) {
        try {
          managePerfectButtons();
          manageOKButton();
        } catch (Exception ex) {
          ExceptionHandler.getHandler().displayException(ex);
        }
      }
    });
  
    //
    // Jaccard Similarity Label and Field
    //
  
    jaccSimLabel_ = new MatchingJLabel(rMan_.getPluginString("networkAlignment.jaccardSimilarityLabel"), jaccSimLabelMatch);
    jaccSimLabel_.setHorizontalAlignment(SwingConstants.CENTER);
    jaccSimField_ = new JTextField(Double.toString(JACCARD_SIMILARITY_DEFAULT));
    
    //
    // No Perfect Alignment for Orphan Layout
    // 'Correct' node groups enabling
    //
    
    if (analysisType_ != NetworkAlignmentBuildData.ViewType.ORPHAN) { // add perfect alignment button
      addLabeledWidget(perfectNGLabel, perfectNGsCombo_, true, true);
      addLabeledFileBrowse(perfectFileMatch_, perfectField_, perfectBrowse);
    }
    
    if (analysisType_ == NetworkAlignmentBuildData.ViewType.GROUP) {
      addLabeledWidget(jaccSimLabel_, jaccSimField_, true, true);
    }
    
    managePerfectButtons();
  
    //
    // OK button
    //
    
    DialogObj.Buttons buttons = finishConstruction();
    
    buttonOK_ = buttons.okButton;
    buttonOK_.setEnabled(false);
    
    setLocationRelativeTo(parent);
  }
  
  /**
   *  Add Action Listeners to Browse Buttons
   */
  
  private void initBrowseButtons(JButton graph1Browse, JButton graph2Browse, JButton alignmentBrowse, JButton perfectBrowse) {
    graph1Browse.addActionListener(new ActionListener() {
      public void actionPerformed(ActionEvent e) {
        try {
          loadFromFile(FileIndex.GRAPH_ONE_FILE);
        } catch (Exception ex) {
          ExceptionHandler.getHandler().displayException(ex);
        }
      }
    });
    graph2Browse.addActionListener(new ActionListener() {
      public void actionPerformed(ActionEvent e) {
        try {
          loadFromFile(FileIndex.GRAPH_TWO_FILE);
        } catch (Exception ex) {
          ExceptionHandler.getHandler().displayException(ex);
        }
      }
    });
    alignmentBrowse.addActionListener(new ActionListener() {
      public void actionPerformed(ActionEvent e) {
        try {
          loadFromFile(FileIndex.ALIGNMENT_FILE);
        } catch (Exception ex) {
          ExceptionHandler.getHandler().displayException(ex);
        }
      }
    });
    perfectBrowse.addActionListener(new ActionListener() {
      public void actionPerformed(ActionEvent e) {
        try {
          loadFromFile(FileIndex.PERFECT_FILE);
        } catch (Exception ex) {
          ExceptionHandler.getHandler().displayException(ex);
        }
      }
    });
    return;
  }
  
  /**
   * Add DocumentListeners to TextFields
   */
  
  private void initTextFields() {
    graph1Field_.getDocument().addDocumentListener(new DocumentListener() {
      public void insertUpdate(DocumentEvent e) {
        try {
          manageOKButton();
        } catch (Exception ex) {
          ExceptionHandler.getHandler().displayException(ex);
        }
      }
  
      public void removeUpdate(DocumentEvent e) {
        try {
          manageOKButton();
        } catch (Exception ex) {
          ExceptionHandler.getHandler().displayException(ex);
        }
      }
  
      public void changedUpdate(DocumentEvent e) {}
    });
    graph2Field_.getDocument().addDocumentListener(new DocumentListener() {
      public void insertUpdate(DocumentEvent e) {
        try {
          manageOKButton();
        } catch (Exception ex) {
          ExceptionHandler.getHandler().displayException(ex);
        }
      }
    
      public void removeUpdate(DocumentEvent e) {
        try {
          manageOKButton();
        } catch (Exception ex) {
          ExceptionHandler.getHandler().displayException(ex);
        }
      }
    
      public void changedUpdate(DocumentEvent e) {}
    });
    alignField_.getDocument().addDocumentListener(new DocumentListener() {
      public void insertUpdate(DocumentEvent e) {
        try {
          manageOKButton();
        } catch (Exception ex) {
          ExceptionHandler.getHandler().displayException(ex);
        }
      }
    
      public void removeUpdate(DocumentEvent e) {
        try {
          manageOKButton();
        } catch (Exception ex) {
          ExceptionHandler.getHandler().displayException(ex);
        }
      }
    
      public void changedUpdate(DocumentEvent e) {}
    });
    perfectField_.getDocument().addDocumentListener(new DocumentListener() {
      public void insertUpdate(DocumentEvent e) {
        try {
          manageOKButton();
        } catch (Exception ex) {
          ExceptionHandler.getHandler().displayException(ex);
        }
      }
    
      public void removeUpdate(DocumentEvent e) {
        try {
          manageOKButton();
        } catch (Exception ex) {
          ExceptionHandler.getHandler().displayException(ex);
        }
      }
    
      public void changedUpdate(DocumentEvent e) {}
    });
    return;
  }
  
  /**
   * Check whether OK button should be activated or deactivated
   */
  
  private void manageOKButton () {
    if (hasMinRequirements()) {
      buttonOK_.setEnabled(true);
    } else {
      buttonOK_.setEnabled(false);
    }
    return;
  }
  
  /**
   * Check whether Perfect text-field and browse should be activated or deactivated
   */
  
  private void managePerfectButtons() {
    if (perfectNGsCombo_.getSelectedIndex() == NO_PERFECT_IDX) {
      perfectField_.setEnabled(false);
      perfectBrowse.setEnabled(false);
      perfectFileMatch_.setEnabled(false);  // gray out label
    } else {
      perfectField_.setEnabled(true);
      perfectBrowse.setEnabled(true);
      perfectFileMatch_.setEnabled(true);
    }
    if (perfectNGsCombo_.getSelectedIndex() == JS_IDX) {
      jaccSimLabel_.setEnabled(true);
      jaccSimField_.setEnabled(true);
    } else {
      jaccSimLabel_.setEnabled(false);
      jaccSimField_.setEnabled(false);
    }
    return;
  }
  
  /**
   * Check whether any text entered in field should be made into a File
   */
  
  private void manageFieldToFile() {
    String g1Field = graph1Field_.getText().trim(), g2Field = graph2Field_.getText().trim(),
            alignField = alignField_.getText().trim(), perfectField = perfectField_.getText().trim();
    
    // if user used only text-field OR entered file through browse but then typed something else in text-field
    if (graph1File_ == null || !g1Field.equals(graph1File_.getAbsolutePath())) {
      graph1File_ = new File(g1Field);
    }
    if (graph2File_ == null || !g2Field.equals(graph2File_.getAbsolutePath())) {
      graph2File_ = new File(g2Field);
    }
    if (alignmentFile_ == null || !alignField.equals(alignmentFile_.getAbsolutePath())) {
      alignmentFile_ = new File(alignField);
    }
    // if user used only text-field after activating drop-down OR
    // entered file through browse but then typed something else in text-field
    if ((perfectNGsCombo_.getSelectedIndex() != NO_PERFECT_IDX && perfectAlignFile_ == null) ||
            perfectAlignFile_ != null && !perfectField.equals(perfectAlignFile_.getAbsolutePath())) {
      perfectAlignFile_ = new File(perfectField);
    }
    return;
  }
  
  /**
   * If user entered in minimum required files in terms of text-fields and non-directed confirmation
   */
  
  private boolean hasMinRequirements() {
    boolean ret = undirectedConfirm_.isSelected();
    // text-field should have text whether or not user typed in text-field or used browse
    ret = ret && (!graph1Field_.getText().isEmpty()) && (!graph2Field_.getText().isEmpty()) && (!alignField_.getText().isEmpty());
  
    // if a perfect NG mode is activated but no perfect alignment is entered yet (through text-field or browse)
    if (perfectNGsCombo_.getSelectedIndex() != NO_PERFECT_IDX && perfectField_.getText().isEmpty()) {
      ret = false;
    }
    return (ret);
  }
  
  /**
   * * Loads the file and update UI
   */
  
  private void loadFromFile(FileIndex mode) {
    
    File file;
    
    switch (mode) {
      case GRAPH_ONE_FILE:
      case GRAPH_TWO_FILE:
        file = flf_.getTheFile(".gw", ".sif", "LoadDirectory", "filterName.graph", this);
        break;
      case ALIGNMENT_FILE:
      case PERFECT_FILE:
        file = flf_.getTheFile(".align", null, "LoadDirectory", "filterName.align", this);
        break;
      default:
        throw new IllegalArgumentException();
    }
    
    if (file == null) {
      return;
    }
    
    PluginSupportFactory.getPreferenceStorage().setPreference("LoadDirectory", file.getAbsoluteFile().getParent());
    
    switch (mode) {
      case GRAPH_ONE_FILE:
        graph1Field_.setText(file.getAbsolutePath());
        graph1File_ = file;
        break;
      case GRAPH_TWO_FILE:
        graph2Field_.setText(file.getAbsolutePath());
        graph2File_ = file;
        break;
      case ALIGNMENT_FILE:
        alignField_.setText(file.getAbsolutePath());
        alignmentFile_ = file;
        break;
      case PERFECT_FILE:
        perfectField_.setText(file.getAbsolutePath());
        perfectAlignFile_ = file;
        break;
      default:
        throw new IllegalArgumentException();
    }
    manageOKButton();
    return;
  }
  
  /**
   ** Checks value in JS text-field
   */
  
  private boolean jaccSimThresholdOK() {
    try {
      String text = jaccSimField_.getText();
      double val = Double.parseDouble(text);
      boolean ok = Double.compare(val, 1.0) <= 0 && Double.compare(val, 0.0) >= 0; // must be in [0,1]
      return (ok);
    } catch (NumberFormatException nfe) {
      return (false);
    }
  }
  
  /**
   ** Returns value in JS text-field after check
   */
  
  private Double getJaccSimThreshold() {
    return (jaccSimThresholdOK()) ? (Double.parseDouble(jaccSimField_.getText())) : null;
  }
  
  ////////////////////////////////////////////////////////////////////////////
  //
  // PUBLIC METHODS AND CLASSES
  //
  ////////////////////////////////////////////////////////////////////////////
  
  @Override
  public void okAction() {
    if (perfectNGsCombo_.getSelectedIndex() == JS_IDX) {
      if (! jaccSimThresholdOK()) {
        JOptionPane.showMessageDialog(parent_, rMan_.getPluginString("networkAlignment.jaccardSimilarityMsg"),
                rMan_.getPluginString("networkAlignment.jaccardSimilarityMsgTitle"),
                JOptionPane.ERROR_MESSAGE);
        return;
      }
    }
    try {
      manageFieldToFile();
      super.okAction();
    } catch (Exception ex) {
      // should never happen because OK button is disabled without correct files.
      ExceptionHandler.getHandler().displayException(ex);
    }
    return;
  }
  
  @Override
  public void closeAction() {
    try {
      super.closeAction();
    } catch (Exception ex) {
      ExceptionHandler.getHandler().displayException(ex);
    }
  }
  
  @Override
  protected boolean stashForOK() {
    return (hasMinRequirements());
  }
  
  public NetworkAlignmentDialogInfo getNAInfo() {
    if (! hasMinRequirements()) {
      // should never happen
      throw new IllegalStateException("Graph file(s) or alignment file missing.");
    }
    
    Double jaccSimThreshold = null;
    NodeGroupMap.PerfectNGMode mode;
    switch (perfectNGsCombo_.getSelectedIndex()) {
      case NO_PERFECT_IDX:
        mode = NodeGroupMap.PerfectNGMode.NONE;
        perfectAlignFile_ = null; // RishiDesai issue #36 fix; user adds perfect file but changes combo box to no file
        break;
      case WITH_PERFECT_IDX:
        mode = NodeGroupMap.PerfectNGMode.NONE;
        break;
      case NC_IDX:
        mode = NodeGroupMap.PerfectNGMode.NODE_CORRECTNESS;
        break;
      case JS_IDX:
        mode = NodeGroupMap.PerfectNGMode.JACCARD_SIMILARITY;
        jaccSimThreshold = getJaccSimThreshold();
        break;
      default:
        // should never happen
        throw (new IllegalStateException("Illegal perfect NG mode"));
    }
    return (new NetworkAlignmentDialogInfo(graph1File_, graph2File_, alignmentFile_, perfectAlignFile_, analysisType_, mode, jaccSimThreshold));
  }
  
  /**
   * The unread files for G1, G2, main alignment, and (possibly) perfect alignment
   */
  
  public static class NetworkAlignmentDialogInfo {
    
    public final File graph1, graph2, align, perfect; // graphs must be entered in correct order (G1 aligned to G2)

    public final NetworkAlignmentBuildData.ViewType analysisType;
    public final NodeGroupMap.PerfectNGMode mode;
    public final Double jaccSimThreshold;
    // Extra settings used for AlignCycleLayout:
    public boolean turnOnShadows;
    public boolean useNodeGroups;    
    
    public NetworkAlignmentDialogInfo(File graph1, File graph2, File align, File perfect,
                                      NetworkAlignmentBuildData.ViewType analysisType, NodeGroupMap.PerfectNGMode mode,
                                      Double jaccSimThreshold) {
      this.graph1 = graph1;
      this.graph2 = graph2;
      this.align = align;
      this.perfect = perfect;
      this.analysisType = analysisType;
      this.mode = mode;
      this.jaccSimThreshold = jaccSimThreshold;
    }
    
  }
  
}
```
