## Supplementary material for "BioFabric Visualization of Network Alignments": VISNAB Source Code: NetworkAlignmentEdgeLayout.java

```
/*
**    Copyright (C) 2018 Rishi Desai
**
**    Copyright (C) 2003-2017 Institute for Systems Biology
**                            Seattle, Washington, USA.
**
**    This library is free software; you can redistribute it and/or
**    modify it under the terms of the GNU Lesser General Public
**    License as published by the Free Software Foundation; either
**    version 2.1 of the License, or (at your option) any later version.
**
**    This library is distributed in the hope that it will be useful,
**    but WITHOUT ANY WARRANTY; without even the implied warranty of
**    MERCHANTABILITY or FITNESS FOR A PARTICULAR PURPOSE.  See the GNU
**    Lesser General Public License for more details.
**
**    You should have received a copy of the GNU Lesser General Public
**    License along with this library; if not, write to the Free Software
**    Foundation, Inc., 59 Temple Place, Suite 330, Boston, MA  02111-1307  USA
*/

package org.systemsbiology.biofabric.plugin.core.align;

import java.util.ArrayList;
import java.util.Collections;
import java.util.Comparator;
import java.util.HashSet;
import java.util.List;
import java.util.Map;
import java.util.Set;

import org.systemsbiology.biofabric.api.io.BuildData;
import org.systemsbiology.biofabric.api.layout.DefaultEdgeLayout;
import org.systemsbiology.biofabric.api.model.NetLink;
import org.systemsbiology.biofabric.api.model.Network;
import org.systemsbiology.biofabric.api.worker.AsynchExitRequestException;
import org.systemsbiology.biofabric.api.worker.BTProgressMonitor;
import org.systemsbiology.biofabric.api.worker.LoopReporter;

public class NetworkAlignmentEdgeLayout extends DefaultEdgeLayout {
  
  ////////////////////////////////////////////////////////////////////////////
  //
  // PRIVATE INSTANCE MEMBERS
  //
  ////////////////////////////////////////////////////////////////////////////
  
  ////////////////////////////////////////////////////////////////////////////
  //
  // PUBLIC CONSTRUCTORS
  //
  ////////////////////////////////////////////////////////////////////////////
  
  /***************************************************************************
   **
   ** Constructor
   */
  
  public NetworkAlignmentEdgeLayout() {
  }
  
  ////////////////////////////////////////////////////////////////////////////
  //
  // PUBLIC METHODS
  //
  ////////////////////////////////////////////////////////////////////////////
  
  /***************************************************************************
   **
   ** Do necessary pre-processing steps (e.g. automatic assignment to link groups)
   */
  
  @Override
  public void preProcessEdges(BuildData rbd,
                              BTProgressMonitor monitor) throws AsynchExitRequestException {
    installLinkGroups(rbd, monitor);
    return;
  }
  
  /***************************************
   **
   ** Get the color
   */
  
  @Override
  protected String getColor(String type, Map<String, String> colorMap) {
    return (null); // because there are node annots already, only grays link annots allowed
  }
  
  /***************************************************************************
   **
   ** Install Link groups for network alignments in this order:
   ** 1) All Covered Edges  2) uncovered G1  3) induced G2  4) half aligned half unaligned G2  5) full unaligned G2
   ** Note: some link groups may not be present.
   */
  
  private void installLinkGroups(BuildData rbd, BTProgressMonitor monitor) throws AsynchExitRequestException {
    
    LoopReporter lr = new LoopReporter(rbd.getLinks().size(), 20, monitor, 0.0, 1.0, "progress.orderingLinkGroups");
    Set<String> relations = new HashSet<String>();
    for (NetLink link : rbd.getLinks()) {
      relations.add(link.getRelation());
      lr.report();
    }
    lr.finish();
    
    List<String> groupOrder = new ArrayList<String>(relations);
    
    // trivial operation (group order is at most length 5)
    Collections.sort(groupOrder, new NetAlignLinkGroupLocator());
    
    rbd.setGroupOrderAndMode(groupOrder, Network.LayoutMode.PER_NETWORK_MODE, true);
    return;
  }
  
  /***************************************************************************
   **
   ** Link groups for network alignments in this order:
   ** 1) All Covered Edges  2) uncovered G1  3) induced G2  4) half aligned half unaligned G2  5) full unaligned G2
   */
  
  private static class NetAlignLinkGroupLocator implements Comparator<String> {
    
    /***************************************************************************
     **
     ** Use fabricated indexes to enforce default link group order
     */
    
    public int compare(String rel1, String rel2) {
      return (getIndex(rel1) - getIndex(rel2));
    }
    
    /***************************************************************************
     **
     ** Get index for default Link Group order
     */
    
    private int getIndex(String rel) {
      
      for (NetworkAlignment.EdgeType type : NetworkAlignment.LINK_GROUPS) {
        if (rel.equals(type.tag)) {
          return (type.index);
        }
      }
      throw (new IllegalArgumentException());
    }
    
  }
  
}
```
