## Supplementary material for "BioFabric Visualization of Network Alignments": VISNAB Source Code: NetworkAlignmentLayout.java

```
/*
**    Copyright (C) 2018 Rishi Desai
**
**    Copyright (C) 2003-2018 Institute for Systems Biology
**                            Seattle, Washington, USA.
**
**    This library is free software; you can redistribute it and/or
**    modify it under the terms of the GNU Lesser General Public
**    License as published by the Free Software Foundation; either
**    version 2.1 of the License, or (at your option) any later version.
**
**    This library is distributed in the hope that it will be useful,
**    but WITHOUT ANY WARRANTY; without even the implied warranty of
**    MERCHANTABILITY or FITNESS FOR A PARTICULAR PURPOSE.  See the GNU
**    Lesser General Public License for more details.
**
**    You should have received a copy of the GNU Lesser General Public
**    License along with this library; if not, write to the Free Software
**    Foundation, Inc., 59 Temple Place, Suite 330, Boston, MA  02111-1307  USA
*/

package org.systemsbiology.biofabric.plugin.core.align;

import java.util.ArrayList;
import java.util.Collections;
import java.util.HashMap;
import java.util.HashSet;
import java.util.Iterator;
import java.util.List;
import java.util.Map;
import java.util.Set;
import java.util.SortedMap;
import java.util.SortedSet;
import java.util.TreeMap;
import java.util.TreeSet;

import org.systemsbiology.biofabric.api.io.BuildData;
import org.systemsbiology.biofabric.api.layout.DefaultLayout;
import org.systemsbiology.biofabric.api.layout.NodeLayout;
import org.systemsbiology.biofabric.api.model.Annot;
import org.systemsbiology.biofabric.api.model.AnnotationSet;
import org.systemsbiology.biofabric.api.model.NetLink;
import org.systemsbiology.biofabric.api.model.NetNode;
import org.systemsbiology.biofabric.api.worker.AsynchExitRequestException;
import org.systemsbiology.biofabric.api.worker.BTProgressMonitor;
import org.systemsbiology.biofabric.api.worker.LoopReporter;
import org.systemsbiology.biofabric.plugin.PluginSupportFactory;

/****************************************************************************
 **
 ** This is the default layout algorithm
 */

public class NetworkAlignmentLayout extends NodeLayout {
  
  ////////////////////////////////////////////////////////////////////////////
  //
  // PRIVATE INSTANCE MEMBERS
  //
  ////////////////////////////////////////////////////////////////////////////
  
  ////////////////////////////////////////////////////////////////////////////
  //
  // PUBLIC CONSTRUCTORS
  //
  ////////////////////////////////////////////////////////////////////////////
  
  /***************************************************************************
   **
   ** Constructor
   */
  
  public NetworkAlignmentLayout() {
  }
  
  ////////////////////////////////////////////////////////////////////////////
  //
  // PUBLIC METHODS
  //
  ////////////////////////////////////////////////////////////////////////////
  
  /***************************************************************************
   **
   ** Relayout the network!
   */
  
  public List<NetNode> doNodeLayout(BuildData rbd, Params params, BTProgressMonitor monitor)
          throws AsynchExitRequestException {
    
    NetworkAlignmentBuildData nabd = (NetworkAlignmentBuildData)rbd.getPluginBuildData();
    
    List<NetNode> targetIDs;
    
    switch (nabd.view) {
      case GROUP:
        targetIDs = bfsNodeGroupLayout(rbd, monitor);
        break;
      case ORPHAN:
        targetIDs = (new DefaultLayout()).defaultNodeOrder(rbd.getLinks(), rbd.getSingletonNodes(), null, monitor);
        break;
      case CYCLE:
        targetIDs = (new AlignCycleLayout()).doNodeOrder(rbd, params, monitor);
        break;
      default:
        throw new IllegalStateException();
    }
    
    installNodeOrder(targetIDs, rbd, monitor);
    return (new ArrayList<NetNode>(targetIDs));
  }
  
  /***************************************************************************
   **
   ** Breadth first search based on node groups
   */

  private List<NetNode> bfsNodeGroupLayout(BuildData bd,
                                           BTProgressMonitor monitor) throws AsynchExitRequestException {
    //
    // Note the allLinks Set has pruned out duplicates and synonymous non-directional links
    //
    //
    // Build a target list, top to bottom, that adds the node with the most
    // links first, and adds those link targets ASAP. If caller supplies a start node,
    // we go there first:
    //
    
    HashMap<NetNode, Integer> linkCounts = new HashMap<NetNode, Integer>();
    HashMap<NetNode, Set<NetNode>> targsPerSource = new HashMap<NetNode, Set<NetNode>>();
    
    HashSet<NetNode> targsToGo = new HashSet<NetNode>();
    
    int numLink = bd.getLinks().size();
    LoopReporter lr = new LoopReporter(numLink, 20, monitor, 0.0, 0.25, "progress.calculateNodeDegree");
    
    Iterator<NetLink> alit = bd.getLinks().iterator();
    while (alit.hasNext()) {
      NetLink nextLink = alit.next();
      lr.report();
      NetNode sidwn = nextLink.getSrcNode();
      NetNode tidwn = nextLink.getTrgNode();
      Set<NetNode> targs = targsPerSource.get(sidwn);
      if (targs == null) {
        targs = new HashSet<NetNode>();
        targsPerSource.put(sidwn, targs);
      }
      targs.add(tidwn);
      targs = targsPerSource.get(tidwn);
      if (targs == null) {
        targs = new HashSet<NetNode>();
        targsPerSource.put(tidwn, targs);
      }
      targs.add(sidwn);
      targsToGo.add(sidwn);
      targsToGo.add(tidwn);
      Integer srcCount = linkCounts.get(sidwn);
      linkCounts.put(sidwn, (srcCount == null) ? Integer.valueOf(1) : Integer.valueOf(srcCount.intValue() + 1));
      Integer trgCount = linkCounts.get(tidwn);
      linkCounts.put(tidwn, (trgCount == null) ? Integer.valueOf(1) : Integer.valueOf(trgCount.intValue() + 1));
    }
    lr.finish();
    
    //
    // Initialize data structures for layout
    //
    
    NodeGroupMap grouper;
    NetworkAlignmentBuildData nabd = (NetworkAlignmentBuildData)bd.getPluginBuildData();
    
    if (nabd.mode == NodeGroupMap.PerfectNGMode.NODE_CORRECTNESS ||
            nabd.mode == NodeGroupMap.PerfectNGMode.JACCARD_SIMILARITY) {
      grouper = new NodeGroupMap(bd, NodeGroupMap.nodeGroupOrderPerfectNG, NodeGroupMap.nodeGroupAnnotsPerfectNG, monitor);
    } else {
      grouper = new NodeGroupMap(bd, NodeGroupMap.nodeGroupOrder, NodeGroupMap.nodeGroupAnnots, monitor);
    }
    
    // master list of nodes in each group
    SortedMap<Integer, List<NetNode>> classToGroup = new TreeMap<Integer, List<NetNode>>();
    for (int i = 0; i < grouper.numGroups(); i++) {
      classToGroup.put(i, new ArrayList<NetNode>());
    }
    // fill the master list with nodes
    Set<NetNode> allNodes = PluginSupportFactory.getBuildExtractor().extractNodes(bd.getLinks(), bd.getSingletonNodes(), monitor);
    for (NetNode node : allNodes) {
      int nodeClass = grouper.getIndex(node);
      classToGroup.get(nodeClass).add(node);
    }
    // sort by decreasing degree
    for (List<NetNode> group : classToGroup.values()) {
      Collections.sort(group, grouper.sortDecrDegree());
    }
    
    SortedMap<Integer, List<NetNode>> targetsGroup = new TreeMap<Integer, List<NetNode>>(),
            queueGroup = new TreeMap<Integer, List<NetNode>>(),
            targsLeftToGoGroup = new TreeMap<Integer, List<NetNode>>();
    
    // each node group (singletons too) gets queue and targets list
    for (int i = 0; i < grouper.numGroups(); i++) {
      targetsGroup.put(i, new ArrayList<NetNode>());
      queueGroup.put(i, new ArrayList<NetNode>());
      targsLeftToGoGroup.put(i, new ArrayList<NetNode>());
      for (NetNode node : classToGroup.get(i)) {
        targsLeftToGoGroup.get(i).add(node);
      }
    }
    
    //
    // Start breadth-first-search on first node group
    //
    
    LoopReporter lr2 = new LoopReporter(targsToGo.size(), 20, monitor, 0.0, 1.0, "progress.nodeOrdering");

    int currGroup = 0;
    while (currGroup < grouper.numGroups()) {
      
      if (targsLeftToGoGroup.get(currGroup).isEmpty()) {
        currGroup++;
        continue; // continue only after each node in group has been visited
      }
      // if queue is empty, pull head node from list
      if (queueGroup.get(currGroup).isEmpty()) {
        NetNode head = targsLeftToGoGroup.get(currGroup).remove(0);
        queueGroup.get(currGroup).add(head);
      }
      
      flushQueue(targetsGroup, targsPerSource, linkCounts, targsToGo, targsLeftToGoGroup, queueGroup, currGroup, grouper, lr2);
    }
    
    lr2.finish();
    
    //
    // Add lone nodes and "flatten" out the targets into one list
    //

    List<NetNode> targets = new ArrayList<NetNode>();
    for (int i = 0; i < grouper.numGroups(); i++) {
      List<NetNode> group = targetsGroup.get(i);
      for (NetNode node : group) {
        targets.add(node);
      }
    }
    
    if (targets.size() != allNodes.size()) {
      throw new IllegalStateException("target numGroups not equal to all-nodes numGroups");
    }

    installAnnotations(bd, targetsGroup, grouper);
    
    return (targets);
  }
  
  /***************************************************************************
   **
   ** Node ordering, non-recursive:
   */
  
  private void flushQueue(SortedMap<Integer, List<NetNode>> targetsGroup,
                          Map<NetNode, Set<NetNode>> targsPerSource,
                          Map<NetNode, Integer> linkCounts,
                          Set<NetNode> targsToGo, SortedMap<Integer, List<NetNode>> targsLeftToGoGroup,
                          SortedMap<Integer, List<NetNode>> queuesGroup,
                          final int currGroup, NodeGroupMap grouper, LoopReporter lr)
          throws AsynchExitRequestException {
    
    List<NetNode> queue = queuesGroup.get(currGroup);
    List<NetNode> leftToGo = targsLeftToGoGroup.get(currGroup);
      
    while (! queue.isEmpty()) {
      NetNode node = queue.remove(0);
      
      if (targetsGroup.get(currGroup).contains(node)) {
        continue; // visited each node only once
      }
      targetsGroup.get(currGroup).add(node);
      lr.report();
      
      if (grouper.getIndex(node) != currGroup) {
        throw new IllegalStateException("Node of incorrect group in queue");
      }
      
      List<NetNode> myKids = orderMyKids(targsPerSource, linkCounts, targsToGo, node);
      for (NetNode kid : myKids) {
        
        if (! targsToGo.contains(kid)) {
          throw new IllegalStateException("kid not in targsToGo");
        }
        
        int kidGroup = grouper.getIndex(kid);
        
        if (kidGroup == currGroup) {
          if (leftToGo.contains(kid)) {
            queue.add(kid);
            leftToGo.remove(kid);
            targsToGo.remove(kid);
          }
        } else {
          if (! queuesGroup.get(kidGroup).contains(kid)) {
            queuesGroup.get(kidGroup).add(kid); // if node from another group, put it in its queue
          }
        }
      }
    }
    return;
  }
  
  /***************************************************************************
   **
   ** Node ordering
   */
  
  private List<NetNode> orderMyKids(final Map<NetNode, Set<NetNode>> targsPerSource,
                                         Map<NetNode, Integer> linkCounts,
                                         Set<NetNode> targsToGo, final NetNode node) {
    Set<NetNode> targs = targsPerSource.get(node);
    if (targs == null) {
      return (new ArrayList<NetNode>());
    }
    TreeMap<Integer, SortedSet<NetNode>> kidMap = new TreeMap<Integer, SortedSet<NetNode>>(Collections.reverseOrder());
    Iterator<NetNode> tait = targs.iterator();
    while (tait.hasNext()) {
      NetNode nextTarg = tait.next();
      Integer count = linkCounts.get(nextTarg);
      SortedSet<NetNode> perCount = kidMap.get(count);
      if (perCount == null) {
        perCount = new TreeSet<NetNode>();
        kidMap.put(count, perCount);
      }
      perCount.add(nextTarg);
    }
    
    ArrayList<NetNode> myKidsToProc = new ArrayList<NetNode>();
    Iterator<SortedSet<NetNode>> kmit = kidMap.values().iterator();
    while (kmit.hasNext()) {
      SortedSet<NetNode> perCount = kmit.next();
      Iterator<NetNode> pcit = perCount.iterator();
      while (pcit.hasNext()) {
        NetNode kid = pcit.next();
        if (targsToGo.contains(kid)) {
          myKidsToProc.add(kid);
        }
      }
    }
    return (myKidsToProc);
  }
  
  /***************************************************************************
   **
   ** Install Layer Zero Node Annotations
   */
  
  private void installAnnotations(BuildData bd,
                                  SortedMap<Integer, List<NetNode>> targetsGroup, NodeGroupMap grouper) {
    
    AnnotationSet layerZeroAnnots = PluginSupportFactory.buildAnnotationSet();
    int min = 0;
    
    for (int i = 0; i < grouper.numGroups(); i++) {
      List<NetNode> group = targetsGroup.get(i);
      if (group.isEmpty()) {
        continue;
      }
      int max = min + group.size() - 1;
  
      Annot annot = PluginSupportFactory.buildAnnotation(grouper.getKey(i), min, max, 0, grouper.getColor(i));
      layerZeroAnnots.addAnnot(annot);
  
      min += group.size(); // update current minimum
    }
    bd.setNodeAnnotations(layerZeroAnnots);
    return;
  }
  
}
```
