## Supplementary material for "BioFabric Visualization of Network Alignments": VISNAB Source Code: NetworkAlignmentPlugIn.java

```
/*
**    Copyright (C) 2003-2018 Institute for Systems Biology 
**                            Seattle, Washington, USA. 
**
**    This library is free software; you can redistribute it and/or
**    modify it under the terms of the GNU Lesser General Public
**    License as published by the Free Software Foundation; either
**    version 2.1 of the License, or (at your option) any later version.
**
**    This library is distributed in the hope that it will be useful,
**    but WITHOUT ANY WARRANTY; without even the implied warranty of
**    MERCHANTABILITY or FITNESS FOR A PARTICULAR PURPOSE.  See the GNU
**    Lesser General Public License for more details.
**
**    You should have received a copy of the GNU Lesser General Public
**    License along with this library; if not, write to the Free Software
**    Foundation, Inc., 59 Temple Place, Suite 330, Boston, MA  02111-1307  USA
*/

package org.systemsbiology.biofabric.plugin.core.align;

import java.io.File;
import java.io.IOException;
import java.io.PrintWriter;
import java.text.MessageFormat;
import java.util.ArrayList;
import java.util.HashMap;
import java.util.HashSet;
import java.util.List;
import java.util.Map;
import java.util.Set;
import java.util.SortedMap;
import java.util.TreeMap;

import javax.swing.JFrame;
import javax.swing.JOptionPane;

import org.systemsbiology.biofabric.api.io.AttributeExtractor;
import org.systemsbiology.biofabric.api.io.BuildData;
import org.systemsbiology.biofabric.api.io.BuildExtractor;
import org.systemsbiology.biofabric.api.io.CharacterEntityMapper;
import org.systemsbiology.biofabric.api.io.FileLoadFlows;
import org.systemsbiology.biofabric.api.io.Indenter;
import org.systemsbiology.biofabric.api.io.PluginWhiteboard;
import org.systemsbiology.biofabric.api.model.AugRelation;
import org.systemsbiology.biofabric.api.model.NetLink;
import org.systemsbiology.biofabric.api.model.NetNode;
import org.systemsbiology.biofabric.api.model.Network;
import org.systemsbiology.biofabric.api.parser.AbstractFactoryClient;
import org.systemsbiology.biofabric.api.parser.GlueStick;
import org.systemsbiology.biofabric.api.util.ExceptionHandler;
import org.systemsbiology.biofabric.api.util.PluginResourceManager;
import org.systemsbiology.biofabric.api.util.UniqueLabeller;
import org.systemsbiology.biofabric.api.worker.AsynchExitRequestException;
import org.systemsbiology.biofabric.api.worker.BFWorker;
import org.systemsbiology.biofabric.api.worker.BTProgressMonitor;
import org.systemsbiology.biofabric.api.worker.BackgroundCore;
import org.systemsbiology.biofabric.api.worker.BackgroundWorkerControlManager;
import org.systemsbiology.biofabric.api.worker.BackgroundWorkerOwner;
import org.systemsbiology.biofabric.plugin.BioFabricToolPlugIn;
import org.systemsbiology.biofabric.plugin.BioFabricToolPlugInCmd;
import org.systemsbiology.biofabric.plugin.BioFabricToolPlugInData;
import org.systemsbiology.biofabric.plugin.PlugInManager;
import org.systemsbiology.biofabric.plugin.PluginSupportFactory;
import org.systemsbiology.biofabric.plugin.PlugInNetworkModelAPI;
import org.systemsbiology.biotapestry.biofabric.FabricCommands;

import org.xml.sax.Attributes;

/****************************************************************************
**
** Class for doing network alignment
*/

public class NetworkAlignmentPlugIn implements BioFabricToolPlugIn {
  
  private ArrayList<BioFabricToolPlugInCmd> myCmds_;
  private String myTag_;
  private NetAlignStats publishedNetAlignStats_;
  private NetAlignStats pendingNetAlignStats_;
  private FileLoadFlows flf_;
  private JFrame topWindow_;
  private BackgroundWorkerControlManager bwcm_;
  private String className_;
  private PluginResourceManager rMan_;
  private PlugInNetworkModelAPI api_;
  
  
  ////////////////////////////////////////////////////////////////////////////
  //
  // PUBLIC METHODS
  //
  ////////////////////////////////////////////////////////////////////////////
  
  /***************************************************************************
  **
  ** Null constructor (required)
  */
  
  public NetworkAlignmentPlugIn() {
    myCmds_ = new ArrayList<BioFabricToolPlugInCmd>();
    myCmds_.add(new LoadNetAlignGroupsCmd());
    myCmds_.add(new LoadNetAlignOrphanCmd());
    myCmds_.add(new LoadNetAlignCaseIICmd());
    myCmds_.add(new NetAlignMeasuresCmd());
    publishedNetAlignStats_ = new NetAlignStats();
    pendingNetAlignStats_ = new NetAlignStats();
    
    className_ = getClass().getName();
   
  }
  
  ////////////////////////////////////////////////////////////////////////////
  //
  // PUBLIC METHODS
  //
  ////////////////////////////////////////////////////////////////////////////
  
  /***************************************************************************
  **
  ** Set the unique tag
  */
  
  public void setUniquePlugInTag(String tag) {
    myTag_ = tag;
    return;
  }
  
  /***************************************************************************
  **
  ** Get the unique tag
  */
  
  public String getUniquePlugInTag() {
    return (myTag_);
  }

  /***************************************************************************
  **
  ** Install a new network
  */
  
  public void newNetworkInstalled(Network bfn) {
    // If we are working on an alignment, we will retain the pending stats we are generating. Else, they get dropped.
    if (pendingNetAlignStats_.hasStats()) {
      publishedNetAlignStats_ = pendingNetAlignStats_;
      pendingNetAlignStats_ = new NetAlignStats();
  	} else {
      publishedNetAlignStats_ = new NetAlignStats();
    }
    for (BioFabricToolPlugInCmd cmd : myCmds_) {
      ((Enabler)cmd).setEnabled(true);
    }
    return;
  }
  
  /***************************************************************************
  **
  ** Install PluginManager
  */
  
  public void installManager(PlugInManager pMan) {
  	rMan_ = PluginSupportFactory.getResourceManager(className_, pMan);
    rMan_.setPluginBundle("org.systemsbiology.biofabric.plugin.core.align.NetworkAlignment");  	
  }

  /***************************************************************************
  **
  ** InstallAPI
  */
  
  public void installAPI(PlugInNetworkModelAPI api) {
    flf_ = api.getFileUtilities();
    topWindow_ = api.getTopWindow();
    bwcm_ = api.getBWCtrlMgr();
    api_ = api;

    for (BioFabricToolPlugInCmd cmd : myCmds_) {
      ((Enabler)cmd).setEnabled(true);
    }
    return;
  }
  
  /***************************************************************************
  **
  ** Get name of tool menu to display
  */
  
  public String getToolMenu() {
    return (rMan_.getPluginString("command.alignmentCommands"));
  }
  
  /***************************************************************************
  **
  ** Get count of commands
  */
  
  public int getCommandCount() {
    return (myCmds_.size());   
  }
  
  /***************************************************************************
  **
  ** Get the nth command
  */
  
  public BioFabricToolPlugInCmd getCommand(int index) {
    return (myCmds_.get(index));
  }
 
  /***************************************************************************
  **
  ** Write session data to given output
  */
  
  public void writeXML(PrintWriter out, Indenter ind) {
    if (!publishedNetAlignStats_.hasStats()) {
      return;
    }
    ind.up();
    ind.indent();
    String name = getClass().getName();
    
    out.print("<");
    out.print(name);
    out.println(">");

    ind.up();
    publishedNetAlignStats_.writeXML(out, ind);
    ind.down();
    ind.indent();
    out.print("</");
    out.print(name);
    out.println(">");
    ind.down();
    return;
  }  
  
  /***************************************************************************
  **
  ** Get XML Reader
  */
 
  public AbstractFactoryClient getXMLWorker(PluginWhiteboard board) {
    return (new PlugInWorker(board, this));
  }
  
  /***************************************************************************
  **
  ** Attach session data read from XML
  */
 
  public void attachXMLData(BioFabricToolPlugInData data) {
    pendingNetAlignStats_ = (NetAlignStats)data;
    return;   
  }
 
  /***************************************************************************
   **
   ** Create individual networks from two files (.gw or .sif) and one .align file
   */
  
  private boolean networkAlignmentFromSources(NetworkAlignmentDialog.NetworkAlignmentDialogInfo nadi,
                                              NetworkAlignmentBuildData.ViewType outType) {
    
    UniqueLabeller idGen = new UniqueLabeller();
    
    //
    // create the individual networks (links + lone nodes)
    //
  
    ArrayList<NetLink> linksSmall = new ArrayList<NetLink>();
    HashSet<NetNode> lonersSmall = new HashSet<NetNode>();
    
    FileLoadFlows.FileLoadType type1 = flf_.getFileLoadType(nadi.graph1);
    FileLoadFlows.FileLoadResult flr = flf_.loadFromASource(nadi.graph1, linksSmall, lonersSmall, null, idGen, true, type1, false);
    File cacheFile = flr.getCacheFile();
    
    
    ArrayList<NetLink> linksLarge = new ArrayList<NetLink>();
    HashSet<NetNode> lonersLarge = new HashSet<NetNode>();
    
    FileLoadFlows.FileLoadType type2 = flf_.getFileLoadType(nadi.graph2);
    flf_.loadFromASource(nadi.graph2, linksLarge, lonersLarge, null, idGen, true, type2, false);
  
    return (networkAlignmentStepTwo(nadi, linksSmall, lonersSmall, linksLarge, lonersLarge,
    		                            nadi.jaccSimThreshold, idGen, outType, cacheFile));
  }
  
  /**************************************************************************
   **
   ** Load the alignment file
   */
  
  private Map<NetNode, NetNode> loadTheAlignmentFile(File file,
                                                              ArrayList<NetLink> linksGraph1, HashSet<NetNode> loneNodesGraph1,
                                                              ArrayList<NetLink> linksGraph2, HashSet<NetNode> loneNodesGraph2) {
    
    Map<NetNode, NetNode> mapG1toG2 = new HashMap<NetNode, NetNode>();
    try {
  
      AlignmentLoader.NetAlignFileStats stats = new AlignmentLoader.NetAlignFileStats();
      AlignmentLoader alod = new AlignmentLoader(className_, rMan_);
      
      alod.readAlignment(file, mapG1toG2, stats, linksGraph1, loneNodesGraph1, linksGraph2, loneNodesGraph2);
  
      if (!stats.badLines.isEmpty()) {
        String badLineFormat = rMan_.getPluginString("netAlignRead.badLineFormat");
        String badLineMsg = MessageFormat.format(badLineFormat, new Object[] {Integer.valueOf(stats.badLines.size())});
        JOptionPane.showMessageDialog(topWindow_, badLineMsg,
                rMan_.getPluginString("netAlignRead.badLineTitle"),
                JOptionPane.WARNING_MESSAGE);
      }
      if (!stats.dupLines.isEmpty()) {
        String dupLineFormat = rMan_.getPluginString("netAlignRead.dupLineFormat");
        String dupLineMsg = MessageFormat.format(dupLineFormat, new Object[] {Integer.valueOf(stats.dupLines.size())});
        JOptionPane.showMessageDialog(topWindow_, dupLineMsg,
                rMan_.getPluginString("netAlignRead.dupLineTitle"),
                JOptionPane.WARNING_MESSAGE);
      }
    } catch (IOException ioe) {
      flf_.displayFileInputError(ioe);
      return (null);
    }
    FabricCommands.setPreference("AttribDirectory", file.getAbsoluteFile().getParent());
    return (mapG1toG2);
  }
  
  /***************************************************************************
   **
   ** Merges individual networks using alignment
   */
  
  private boolean networkAlignmentStepTwo(NetworkAlignmentDialog.NetworkAlignmentDialogInfo nadi,
                                          ArrayList<NetLink> linksSmall, HashSet<NetNode> lonersSmall,
                                          ArrayList<NetLink> linksLarge, HashSet<NetNode> lonersLarge,
                                          Double jaccSimThreshold, UniqueLabeller idGen,
                                          NetworkAlignmentBuildData.ViewType outType, File cacheFile) {
    // Alignment processing
    Map<NetNode, NetNode> mapG1toG2 =
            loadTheAlignmentFile(nadi.align, linksSmall, lonersSmall, linksLarge, lonersLarge);
    if (mapG1toG2 == null) {
      return (true);
    }
    
    Map<NetNode, NetNode> perfectG1toG2;
    if (nadi.perfect != null) {
      perfectG1toG2 = loadTheAlignmentFile(nadi.perfect, linksSmall, lonersSmall, linksLarge, lonersLarge);
      if (perfectG1toG2 == null) {
        return (true);
      }
    } else {
      perfectG1toG2 = null;
    }
    
    //
    // Make sure G1's nodes are subset of G2's if perfect align not provided in CaseII cycle
    //
    
    if (outType == NetworkAlignmentBuildData.ViewType.CYCLE && perfectG1toG2 == null) {
      boolean isSubset = isG1subsetG2(linksSmall, lonersSmall, linksLarge, lonersLarge);
      if (!isSubset) {
        JOptionPane.showMessageDialog(topWindow_, rMan_.getPluginString("networkAlignment.cycleSubsetErrorMessage"),
                rMan_.getPluginString("networkAlignment.cycleSubsetErrorMessageTitle"),
                JOptionPane.ERROR_MESSAGE);
        return (true);
      }
    }
    
    //
    // The CaseII cycle alignment can use the perfect alignment file, so we need to be more
    // specific about when the perfect/Group analysis is being done:
    //
    
    boolean doingPerfectGroup = (outType == NetworkAlignmentBuildData.ViewType.GROUP) && 
                                (perfectG1toG2 != null);
           
    //
    // First process the given (main) alignment
    //
    
    NetworkAlignmentBuilder nab = new NetworkAlignmentBuilder();
    
    ArrayList<NetLink> mergedLinks = new ArrayList<NetLink>();
    Set<NetNode> mergedLoneNodeIDs = new HashSet<NetNode>();
    SortedMap<AugRelation, Boolean> relMap = new TreeMap<AugRelation, Boolean>();
    Set<NetLink> reducedLinks = new HashSet<NetLink>();
    Map<NetNode, Boolean> mergedToCorrectNC = null;
    NetworkAlignment.NodeColorMap nodeColorMap = new NetworkAlignment.NodeColorMap();
    if (doingPerfectGroup) {
      mergedToCorrectNC = new HashMap<NetNode, Boolean>();
    }
    
    boolean finished = nab.processNetAlign(mergedLinks, mergedLoneNodeIDs, mapG1toG2, perfectG1toG2, mergedToCorrectNC,
                                           nodeColorMap, linksSmall, lonersSmall, linksLarge, lonersLarge,
                                           relMap, outType, idGen, cacheFile);
  
    //
    // Second process the perfect alignment (if given)
    //
    
    nab = null;
    ArrayList<NetLink> mergedLinksPerfect = null;
    Set<NetNode> mergedLoneNodeIDsPerfect = null;
    SortedMap<AugRelation, Boolean> relMapPerfect = null;
    Set<NetLink> reducedLinksPerfect = null;
    NetworkAlignment.NodeColorMap nodeColorMapPerfect = null;
    
    if (finished && doingPerfectGroup) {
      //
      // We now have to process the Perfect alignment so we can compare the links/nodes (topology, etc)
      // between the given alignment and the perfect alignment. The added -'Perfect' on the variable names
      // signifies it.
      //
      nab = new NetworkAlignmentBuilder();
      mergedLinksPerfect = new ArrayList<NetLink>();
      mergedLoneNodeIDsPerfect = new HashSet<NetNode>();
      relMapPerfect = new TreeMap<AugRelation, Boolean>();
      reducedLinksPerfect = new HashSet<NetLink>();
      nodeColorMapPerfect = new NetworkAlignment.NodeColorMap();
      
      finished = nab.processNetAlign(mergedLinksPerfect, mergedLoneNodeIDsPerfect, perfectG1toG2, null, null,
              nodeColorMapPerfect, linksSmall, lonersSmall, linksLarge, lonersLarge, relMapPerfect,
              NetworkAlignmentBuildData.ViewType.GROUP, idGen, cacheFile);
    }
  
    // Warn user of alignment with only singletons
    if (finished && mergedLinks.isEmpty()) {
      JOptionPane.showMessageDialog(topWindow_, rMan_.getPluginString("networkAlignment.emptyNetwork"),
              rMan_.getPluginString("networkAlignment.emptyNetworkTitle"),
              JOptionPane.WARNING_MESSAGE);
    }
    
    if (finished) { // for main alignment
    	// If we have just singletons, we can skip the shadow question (though with no links it will not
    	// be asked anyway. More importantly, if user forced it on previously, don't ask again.
    	boolean skip_shadows = (mergedLinks.isEmpty() || nadi.turnOnShadows);
      finished = flf_.handleDirectionsDupsAndShadows(mergedLinks, mergedLoneNodeIDs, false, relMap, reducedLinks, cacheFile, true, skip_shadows);
    }
    
    if (finished && doingPerfectGroup) { // for perfect alignment
    	// Doing this a second time, we want to shut up the question about turning on shadow links (just got asked)
      finished = flf_.handleDirectionsDupsAndShadows(mergedLinksPerfect, mergedLoneNodeIDsPerfect, false, relMapPerfect, 
      																							 reducedLinksPerfect, cacheFile, true, true);
    }
  
    if (finished) { // Score Report
      finished = networkAlignmentStepFour(reducedLinks, mergedLoneNodeIDs, nodeColorMap, mergedToCorrectNC,
              reducedLinksPerfect, mergedLoneNodeIDsPerfect, nodeColorMapPerfect, pendingNetAlignStats_,
              linksSmall, lonersSmall, linksLarge, lonersLarge, mapG1toG2, perfectG1toG2);
    }
   
    if (finished) { // Load the alignments
      
      //
      // If we are doing a CaseII Cycle layout, we want to have a full list of the nodes in both
      // networks:
      //
  
      BuildExtractor bex = PluginSupportFactory.getBuildExtractor();
      Set<NetNode> allLargerNodes, allSmallerNodes;
      try {
        allLargerNodes = bex.extractNodes(linksLarge, lonersLarge,null);
        allSmallerNodes = bex.extractNodes(linksSmall,lonersSmall, null);
      } catch (AsynchExitRequestException aere) {
        throw (new IllegalStateException("Error met while extracting nodes"));
        // shouldn't happen
      }
      NetworkAlignmentBuildData nabd = new NetworkAlignmentBuildData(nodeColorMap, reducedLinksPerfect, mergedLoneNodeIDsPerfect, nodeColorMapPerfect,
              mergedToCorrectNC, allSmallerNodes, linksSmall, lonersSmall, allLargerNodes, linksLarge, lonersLarge, mapG1toG2, perfectG1toG2,
              pendingNetAlignStats_, outType, nadi.mode, jaccSimThreshold, nadi.useNodeGroups, nadi.turnOnShadows);
  
      networkAlignmentStepFive(reducedLinks, mergedLoneNodeIDs, nabd, idGen, nadi.align, cacheFile);
    }
    pendingNetAlignStats_ = new NetAlignStats();
    return (true);
  }
  

  
  /***************************************************************************
   **
   ** Process NetAlign Score Reports
   */
  
  private boolean networkAlignmentStepFour(Set<NetLink> reducedLinks, Set<NetNode> loneNodeIDs, NetworkAlignment.NodeColorMap nodeColorMap,
                                           Map<NetNode, Boolean> mergedToCorrectNC, Set<NetLink> reducedLinksPerfect,
                                           Set<NetNode> loneNodeIDsPerfect, NetworkAlignment.NodeColorMap nodeColorMapPerfect,
                                           NetAlignStats report,
                                           ArrayList<NetLink> linksSmall, HashSet<NetNode> lonersSmall,
                                           ArrayList<NetLink> linksLarge, HashSet<NetNode> lonersLarge,
                                           Map<NetNode, NetNode> mapG1toG2, Map<NetNode, NetNode> perfectG1toG2) {
    File holdIt;
    try {
      holdIt = File.createTempFile("BioFabricHold", ".zip");
      holdIt.deleteOnExit();
    } catch (IOException ioex) {
      holdIt = null;
    }
    NetAlignMeasureBuilder namb = new NetAlignMeasureBuilder();
    
    boolean finished = namb.processNetAlignMeasures(reducedLinks, loneNodeIDs, nodeColorMap, mergedToCorrectNC,
            reducedLinksPerfect, loneNodeIDsPerfect, nodeColorMapPerfect, report, linksSmall, lonersSmall,
            linksLarge, lonersLarge, mapG1toG2, perfectG1toG2, holdIt);
    
    return (finished);
  }
  
  /***************************************************************************
   **
   ** Build the network alignment
   */
  
    private boolean networkAlignmentStepFive(Set<NetLink> reducedLinks, Set<NetNode> loneNodeIDs,
            NetworkAlignmentBuildData nabd, UniqueLabeller idGen, File align, File holdIt) {

    HashMap<NetNode, String> emptyClustMap = new HashMap<NetNode, String>();

    BuildData bd = PluginSupportFactory.getBuildDataForPlugin(idGen, reducedLinks, loneNodeIDs, emptyClustMap, null);
    bd.setLayoutMode(Network.LayoutMode.PER_NETWORK_MODE);
    bd.setPluginBuildData(nabd);
  
    try {
    	// Let us know that the network we are about to hear about is an alignment:
      flf_.buildNetworkForPlugIn(bd, holdIt, rMan_); 
    } catch (OutOfMemoryError oom) {
      ExceptionHandler.getHandler().displayOutOfMemory(oom);
      return (false);
    }
    flf_.setCurrentXMLFile(null);
    FabricCommands.setPreference("LoadDirectory", align.getAbsoluteFile().getParent());
    flf_.manageWindowTitle(align.getName());
    return true;
  }
  
  /***************************************************************************
   **
   ** Check if G1's nodes are subset of G2's
   */
  
  private boolean isG1subsetG2(ArrayList<NetLink> linksSmall, HashSet<NetNode> lonersSmall,
                               ArrayList<NetLink> linksLarge, HashSet<NetNode> lonersLarge) {
  
    Set<NetNode> nodesG1 = null, nodesG2 = null;
    try {
      BuildExtractor bex = PluginSupportFactory.getBuildExtractor();
      nodesG1 = bex.extractNodes(linksSmall, lonersSmall, null);
      nodesG2 = bex.extractNodes(linksLarge, lonersLarge, null);
    } catch (AsynchExitRequestException aere) {
      // should never happen
    }
    
    if (nodesG1 == null || nodesG2 == null) {
      throw (new IllegalStateException("Node Set null in subset test"));
    }
    
    Set<String> namesG1 = new HashSet<String>(), namesG2 = new HashSet<String>();
    for (NetNode node : nodesG1) {
      namesG1.add(node.getName());
    }
    for (NetNode node : nodesG2) {
      namesG2.add(node.getName());
    }
    
    return (namesG2.containsAll(namesG1));
  }
  
  ////////////////////////////////////////////////////////////////////////////
  //
  // INNER CLASSES
  //
  ////////////////////////////////////////////////////////////////////////////
  
  interface Enabler {
    public void setEnabled(boolean isEnabled);
  }
  
  
  /***************************************************************************
  **
  ** Command
  */
  
  private class NetAlignMeasuresCmd implements BioFabricToolPlugInCmd, Enabler {
   
  	private boolean enabled_;
  	
    public void setEnabled(boolean isEnabled) {
    	enabled_ = isEnabled;
      return;
    }
      
    public String getCommandName() {
      return (rMan_.getPluginString("command.netAlignMeasures"));
    }   
    
    public boolean performOperation(JFrame topFrame) {
      if (!publishedNetAlignStats_.hasStats() || !enabled_) {
        return (false);
      }    
      
      NetAlignMeasureDialog scoreDialog = new NetAlignMeasureDialog(topFrame, publishedNetAlignStats_, rMan_);
      scoreDialog.setVisible(true);
      return (true);
    }
  
    public boolean isEnabled() {
      return (enabled_ && publishedNetAlignStats_.hasStats());
    } 
  }
 
  /***************************************************************************
  **
  ** Command
  */
  
  private class LoadNetAlignGroupsCmd implements BioFabricToolPlugInCmd, Enabler {
  
    public void setEnabled(boolean isEnabled) {
      return; // Always enabled  
    }
    
    public String getCommandName() {
      return (rMan_.getPluginString("command.netAlignGroupLayout"));  
    }
    
    public boolean performOperation(JFrame topFrame) {
      
      NetworkAlignmentDialog nad = new NetworkAlignmentDialog(topFrame,  
                                                              NetworkAlignmentBuildData.ViewType.GROUP, 
                                                              className_, flf_, rMan_);
      nad.setVisible(true);
      
      if(!nad.haveResult()) {
        return (false);
      }
  
      NetworkAlignmentDialog.NetworkAlignmentDialogInfo nai = nad.getNAInfo();
      
      boolean filesNotOkay =
              !flf_.standardFileChecks(nai.graph1, FileLoadFlows.FILE_MUST_EXIST, FileLoadFlows.FILE_CAN_CREATE_DONT_CARE,
                                       FileLoadFlows.FILE_DONT_CHECK_OVERWRITE, FileLoadFlows.FILE_MUST_BE_FILE,
                                       FileLoadFlows.FILE_CAN_WRITE_DONT_CARE, FileLoadFlows.FILE_CAN_READ) ||
              !flf_.standardFileChecks(nai.graph2, FileLoadFlows.FILE_MUST_EXIST, FileLoadFlows.FILE_CAN_CREATE_DONT_CARE,
                                       FileLoadFlows.FILE_DONT_CHECK_OVERWRITE, FileLoadFlows.FILE_MUST_BE_FILE,
                                       FileLoadFlows.FILE_CAN_WRITE_DONT_CARE, FileLoadFlows.FILE_CAN_READ) ||
              !flf_.standardFileChecks(nai.align, FileLoadFlows.FILE_MUST_EXIST, FileLoadFlows.FILE_CAN_CREATE_DONT_CARE,
                  FileLoadFlows.FILE_DONT_CHECK_OVERWRITE, FileLoadFlows.FILE_MUST_BE_FILE,
                  FileLoadFlows.FILE_CAN_WRITE_DONT_CARE, FileLoadFlows.FILE_CAN_READ);
      if (nai.perfect != null) {
        filesNotOkay = !flf_.standardFileChecks(nai.perfect, FileLoadFlows.FILE_MUST_EXIST, FileLoadFlows.FILE_CAN_CREATE_DONT_CARE,
                                                FileLoadFlows.FILE_DONT_CHECK_OVERWRITE, FileLoadFlows.FILE_MUST_BE_FILE,
                                                FileLoadFlows.FILE_CAN_WRITE_DONT_CARE, FileLoadFlows.FILE_CAN_READ);
      }
      
      if (filesNotOkay) {
        return (false);
      }
      return (networkAlignmentFromSources(nai, NetworkAlignmentBuildData.ViewType.GROUP));
    }
    
    public boolean isEnabled() {
      return (true);  // Always enabled  
    }
  
  }
  
  /***************************************************************************
  **
  ** Command
  */
  
  private class LoadNetAlignCaseIICmd implements BioFabricToolPlugInCmd, Enabler {
  

    public void setEnabled(boolean isEnabled) {
      return; // Always enabled  
    }
    
    public String getCommandName() {
      return (rMan_.getPluginString("command.netAlignCaseIILayout"));  
    }
    
    public boolean performOperation(JFrame topFrame) {
    
      NetworkAlignmentDialog nad = new NetworkAlignmentDialog(topFrame, NetworkAlignmentBuildData.ViewType.CYCLE, 
      		                                                    className_, flf_, rMan_);
      nad.setVisible(true);
      
      if(!nad.haveResult()) {
        return (false);
      }
  
      NetworkAlignmentDialog.NetworkAlignmentDialogInfo nai = nad.getNAInfo();
      
      boolean filesNotOkay =
              !flf_.standardFileChecks(nai.graph1, FileLoadFlows.FILE_MUST_EXIST, FileLoadFlows.FILE_CAN_CREATE_DONT_CARE,
                                       FileLoadFlows.FILE_DONT_CHECK_OVERWRITE, FileLoadFlows.FILE_MUST_BE_FILE,
                                       FileLoadFlows.FILE_CAN_WRITE_DONT_CARE, FileLoadFlows.FILE_CAN_READ) ||
              !flf_.standardFileChecks(nai.graph2, FileLoadFlows.FILE_MUST_EXIST, FileLoadFlows.FILE_CAN_CREATE_DONT_CARE,
                                       FileLoadFlows.FILE_DONT_CHECK_OVERWRITE, FileLoadFlows.FILE_MUST_BE_FILE,
                                       FileLoadFlows.FILE_CAN_WRITE_DONT_CARE, FileLoadFlows.FILE_CAN_READ) ||
              !flf_.standardFileChecks(nai.align, FileLoadFlows.FILE_MUST_EXIST, FileLoadFlows.FILE_CAN_CREATE_DONT_CARE,
                  FileLoadFlows.FILE_DONT_CHECK_OVERWRITE, FileLoadFlows.FILE_MUST_BE_FILE,
                  FileLoadFlows.FILE_CAN_WRITE_DONT_CARE, FileLoadFlows.FILE_CAN_READ);

      if (filesNotOkay) {
        return (false);
      }
      
      //
      // With this layout, shadow links can really help, since we do not draw link group annotations if shadows
      // are not present. So provide the user with the opportunity to force shadow links to be on.
      //
      
      if (!api_.getDisplayShadows()) {
        ShadowsAndGroupsDialog sagd = new ShadowsAndGroupsDialog(topWindow_, rMan_);
        sagd.setVisible(true);
      
        if (!sagd.haveResult()) {
          return (false);
        }
        nai.turnOnShadows = sagd.turnShadowsOn();
        nai.useNodeGroups = !nai.turnOnShadows;
      } else {
      	nai.useNodeGroups = false;
      }

      return (networkAlignmentFromSources(nai, NetworkAlignmentBuildData.ViewType.CYCLE));
    }
    
    public boolean isEnabled() {
      return (true);  // Always enabled     
    }
  
  }

  
  /***************************************************************************
   **
   ** Command
   */
  
  private class LoadNetAlignOrphanCmd implements BioFabricToolPlugInCmd, Enabler {
    
    public void setEnabled(boolean isEnabled) {
      return; // Always enabled  
    }
    
    public String getCommandName() {
      return (rMan_.getPluginString("command.orphanLayout"));  
    }
    
    public boolean performOperation(JFrame topFrame) {
      
      NetworkAlignmentDialog nad = new NetworkAlignmentDialog(topFrame, NetworkAlignmentBuildData.ViewType.ORPHAN, 
      		                                                    className_, flf_, rMan_);
      nad.setVisible(true);
      
      if(!nad.haveResult()) {
        return (false);
      }
      
      NetworkAlignmentDialog.NetworkAlignmentDialogInfo nai = nad.getNAInfo();
      
      boolean filesNotOkay =
              !flf_.standardFileChecks(nai.graph1, FileLoadFlows.FILE_MUST_EXIST, FileLoadFlows.FILE_CAN_CREATE_DONT_CARE,
                                       FileLoadFlows.FILE_DONT_CHECK_OVERWRITE, FileLoadFlows.FILE_MUST_BE_FILE,
                                       FileLoadFlows.FILE_CAN_WRITE_DONT_CARE, FileLoadFlows.FILE_CAN_READ) ||
              !flf_.standardFileChecks(nai.graph2, FileLoadFlows.FILE_MUST_EXIST, FileLoadFlows.FILE_CAN_CREATE_DONT_CARE,
                                       FileLoadFlows.FILE_DONT_CHECK_OVERWRITE, FileLoadFlows.FILE_MUST_BE_FILE,
                                       FileLoadFlows.FILE_CAN_WRITE_DONT_CARE, FileLoadFlows.FILE_CAN_READ) ||
              !flf_.standardFileChecks(nai.align, FileLoadFlows.FILE_MUST_EXIST, FileLoadFlows.FILE_CAN_CREATE_DONT_CARE,
                                       FileLoadFlows.FILE_DONT_CHECK_OVERWRITE, FileLoadFlows.FILE_MUST_BE_FILE,
                                       FileLoadFlows.FILE_CAN_WRITE_DONT_CARE, FileLoadFlows.FILE_CAN_READ);
      
      if (filesNotOkay) {
        return (false);
      }
      return (networkAlignmentFromSources(nai, NetworkAlignmentBuildData.ViewType.ORPHAN));
    }
    
    public boolean isEnabled() {
      return (true); // Always enabled      
    }
  }
  
  /****************************************************************************
  **
  ** Contains common network alignment scores
  */
  
  public static class NetAlignStats implements BioFabricToolPlugInData {
    
    private List<NetAlignMeasure> measures_;
    
    public NetAlignStats() {
      measures_ = new ArrayList<NetAlignMeasure>();
    }
    
    public NetAlignStats(List<NetAlignMeasure> measures) {
      this.measures_ = measures;
    }
     
    public void addAMeasure(NetAlignMeasure measure) {
      measures_.add(measure);
      return;
    }
    
    public boolean hasStats() {
      return (!measures_.isEmpty());
    }
    
    public List<NetAlignMeasure> getMeasures() {
      return (measures_);
    }
     
    @Override
    public String toString() {
      StringBuilder ret = new StringBuilder("Measures");
      for (NetAlignMeasure msr : measures_) {
        ret.append('\n').append(msr.name).append(':').append(String.format("%4.4f", msr.val));
      }
      return (ret.toString());
    }
    
    public void replaceValuesTo(NetAlignStats other) {
      measures_ = new ArrayList<NetAlignMeasure>(other.measures_);
    }
    
    public void writeXML(PrintWriter out, Indenter ind) {
      ind.indent();  
      out.println("<NetAlignStats>");
      ind.up();
      for (NetAlignMeasure msr : measures_) {
        msr.writeXML(out, ind);
      }
      ind.down();
      ind.indent();
      out.println("</NetAlignStats>");
      return;
    }
  }
  
  public static class NetAlignMeasure {
    
    public final Double val;
    public final String name;
    
    public NetAlignMeasure(String name, Double val) {
      this.val = val;
      this.name = name;
    }
  
    public void writeXML(PrintWriter out, Indenter ind) {
      ind.indent(); 
      out.print("<NetAlignMeasure name=\"");
      out.print(CharacterEntityMapper.mapEntities(name, false));
      out.print("\" val=\"");
      out.print(val);
      out.println("\"/>");
      return;
    }
 
    @Override
    public String toString() {
      return ("NetAlignMeasure{" + "val=" + val + ", name='" + name + '\'' + '}');
    }
  }

  /***************************************************************************
   **
   ** Class for building network alignments
   */
  
  private class NetworkAlignmentBuilder implements BackgroundWorkerOwner {
    
    private File holdIt_;
    private boolean finished_;
    
    public boolean processNetAlign(ArrayList<NetLink> mergedLinks, Set<NetNode> mergedLoneNodeIDs,
                                   Map<NetNode, NetNode> mapG1toG2,
                                   Map<NetNode, NetNode> perfectG1toG2,
                                   Map<NetNode, Boolean> mergedToCorrect,
                                   NetworkAlignment.NodeColorMap nodeColorMap,
                                   ArrayList<NetLink> linksG1, HashSet<NetNode> lonersG1,
                                   ArrayList<NetLink> linksG2, HashSet<NetNode> lonersG2,
                                   SortedMap<AugRelation, Boolean> relMap,
                                   NetworkAlignmentBuildData.ViewType outType, UniqueLabeller idGen, File holdIt) {
      finished_= true;
      holdIt_ = holdIt;
      try {    	
      	BFWorker bfw = PluginSupportFactory.getBFWorker(this, topWindow_, bwcm_, "fileLoad.waitTitle", "fileLoad.wait", true, rMan_);
        NetworkAlignmentRunner runner = new NetworkAlignmentRunner(mergedLinks, mergedLoneNodeIDs, mapG1toG2, perfectG1toG2,
                                                                   mergedToCorrect, nodeColorMap, linksG1, lonersG1, linksG2,
                                                                   lonersG2, relMap, outType, idGen, bfw);
        bfw.setCore(runner);
        bfw.launchWorker();
      } catch (Exception ex) {
        ExceptionHandler.getHandler().displayException(ex);
      }
      return (finished_);
    }
    
    public boolean handleRemoteException(Exception remoteEx) {
      finished_ = false;
      return (false);
    }
    
    public boolean handleCancellation() {
      finished_ = false;
      return (flf_.cancelAndRestore(holdIt_));
    }
    
    public void cleanUpPreEnable(Object result) {
      return;
    }
    
    public void cleanUpPostRepaint(Object result, boolean skipImage) {
      return;
    }
  }  
  
  /***************************************************************************
   **
   ** Background network alignment processing
   */
  
  private class NetworkAlignmentRunner implements BackgroundCore {
    
    private ArrayList<NetLink> mergedLinks_;
    private Set<NetNode> mergedLoneNodeIDs_;
    private Map<NetNode, NetNode> mapG1toG2_, perfectG1toG2_;
    private Map<NetNode, Boolean> mergedToCorrect_;
    private NetworkAlignment.NodeColorMap nodeColorMap_;
    private ArrayList<NetLink> linksG1_, linksG2_;
    private HashSet<NetNode> lonersG1_, lonersG2_;
    private SortedMap<AugRelation, Boolean> relMap_;
    private NetworkAlignmentBuildData.ViewType outType_;
    private UniqueLabeller idGen_;
    private BFWorker bfwk_;
    
    public NetworkAlignmentRunner(ArrayList<NetLink> mergedLinks, Set<NetNode> mergedLoners,
                                  Map<NetNode, NetNode> mapG1toG2,
                                  Map<NetNode, NetNode> perfectG1toG2,
                                  Map<NetNode, Boolean> mergedToCorrect,
                                  NetworkAlignment.NodeColorMap nodeColorMap,
                                  ArrayList<NetLink> linksG1, HashSet<NetNode> lonersG1,
                                  ArrayList<NetLink> linksG2, HashSet<NetNode> lonersG2,
                                  SortedMap<AugRelation, Boolean> relMap,
                                  NetworkAlignmentBuildData.ViewType outType, 
                                  UniqueLabeller idGen, BFWorker bfwk) {
      
      this.bfwk_ = bfwk;
      this.mergedLinks_ = mergedLinks;
      this.mergedLoneNodeIDs_ = mergedLoners;
      this.mapG1toG2_ = mapG1toG2;
      this.perfectG1toG2_ = perfectG1toG2;
      this.mergedToCorrect_ = mergedToCorrect;
      this.nodeColorMap_ = nodeColorMap;
      this.linksG1_ = linksG1;
      this.lonersG1_ = lonersG1;
      this.linksG2_ = linksG2;
      this.lonersG2_ = lonersG2;
      this.relMap_ = relMap;
      this.outType_ = outType;
      this.idGen_ = idGen;
    }
    
    public Object getEarlyResult() {
      return (new Boolean(false));
    }

    public Object runCore() throws AsynchExitRequestException {
      
    	BTProgressMonitor monitor = bfwk_.getMonitor();
      NetworkAlignment netAlign = new NetworkAlignment(mergedLinks_, mergedLoneNodeIDs_, mapG1toG2_, perfectG1toG2_,
              linksG1_, lonersG1_, linksG2_, lonersG2_, mergedToCorrect_, nodeColorMap_, outType_, idGen_, monitor);
      
      netAlign.mergeNetworks();
      BuildExtractor bex = PluginSupportFactory.getBuildExtractor();
      bex.extractRelations(mergedLinks_, relMap_, monitor);
      return (new Boolean(true));
    }
    
    public Object postRunCore() {
      return null;
    }
  }
  
  /***************************************************************************
   **
   ** Class for calculating network alignment measures
   */
  
  private class NetAlignMeasureBuilder implements BackgroundWorkerOwner {
    
    private File holdIt_;
    private boolean finished_;
    
    public boolean processNetAlignMeasures(Set<NetLink> reducedLinks, Set<NetNode> loneNodeIDs, NetworkAlignment.NodeColorMap nodeColorMap,
                                           Map<NetNode, Boolean> mergedToCorrectNC, Set<NetLink> reducedLinksPerfect,
                                           Set<NetNode> loneNodeIDsPerfect, NetworkAlignment.NodeColorMap nodeColorMapPerfect,
                                           NetAlignStats report,
                                           ArrayList<NetLink> linksSmall, HashSet<NetNode> lonersSmall,
                                           ArrayList<NetLink> linksLarge, HashSet<NetNode> lonersLarge,
                                           Map<NetNode, NetNode> mapG1toG2, Map<NetNode, NetNode> perfectG1toG2, File holdIt) {
      finished_ = true;
      holdIt_ = holdIt;
      try {
      	
      	BFWorker bfw = PluginSupportFactory.getBFWorker(this, topWindow_, bwcm_, "fileLoad.waitTitle", "fileLoad.wait", true, rMan_);
        NetAlignMeasureRunner runner = new NetAlignMeasureRunner(reducedLinks, loneNodeIDs, nodeColorMap, mergedToCorrectNC,
        		                                                     reducedLinksPerfect, loneNodeIDsPerfect, nodeColorMapPerfect,
        		                                                     report, linksSmall, lonersSmall, linksLarge, 
        		                                                     lonersLarge, mapG1toG2, perfectG1toG2, bfw, rMan_);
        bfw.setCore(runner);
        bfw.launchWorker();
      } catch (Exception ex) {
        ExceptionHandler.getHandler().displayException(ex);
      }
      return (finished_);
    }
    
    public boolean handleRemoteException(Exception remoteEx) {
      finished_ = false;
      return (false);
    }
  
    public boolean handleCancellation() {
      finished_ = false;
      return (flf_.cancelAndRestore(holdIt_));
    }
  
    public void cleanUpPreEnable(Object result) {
      return;
    }
  
    public void cleanUpPostRepaint(Object result, boolean skipImage) {
      return;
    }
  }
  
  /***************************************************************************
   **
   ** Background network alignment measure processing
   */
  
  private class NetAlignMeasureRunner implements BackgroundCore {
  
    private Map<NetNode, NetNode> mapG1toG2_;
    private Set<NetLink> reducedLinks_;
    private Set<NetNode> loneNodeIDs_;
    private NetworkAlignment.NodeColorMap nodeColorMap_;
    private Map<NetNode, Boolean> mergedToCorrectNC_;
    private Map<NetNode, NetNode> perfectG1toG2_;
    private Set<NetLink> reducedLinksPerfect_;
    private Set<NetNode> loneNodeIDsPerfect_;
    private NetworkAlignment.NodeColorMap nodeColorMapPerfect_;
  
    private ArrayList<NetLink> linksSmall_, linksLarge_;
    private HashSet<NetNode> lonersSmall_, lonersLarge_;
    private NetAlignStats report_;
    private BFWorker bfwk_;
    private PluginResourceManager rMan_;
    
    
    public NetAlignMeasureRunner(Set<NetLink> reducedLinks, Set<NetNode> loneNodeIDs, NetworkAlignment.NodeColorMap nodeColorMap,
                                 Map<NetNode, Boolean> mergedToCorrectNC, Set<NetLink> reducedLinksPerfect,
                                 Set<NetNode> loneNodeIDsPerfect, NetworkAlignment.NodeColorMap nodeColorMapPerfect, NetAlignStats report,
                                 ArrayList<NetLink> linksSmall, HashSet<NetNode> lonersSmall,
                                 ArrayList<NetLink> linksLarge, HashSet<NetNode> lonersLarge,
                                 Map<NetNode, NetNode> mapG1toG2, 
                                 Map<NetNode, NetNode> perfectG1toG2, BFWorker bfwk, PluginResourceManager rMan) {
      
    	rMan_ = rMan;
    	this.bfwk_ = bfwk;
      this.reducedLinks_ = reducedLinks;
      this.loneNodeIDs_ = loneNodeIDs;
      this.nodeColorMap_ = nodeColorMap;
      this.nodeColorMapPerfect_ = nodeColorMapPerfect;
      this.mergedToCorrectNC_ = mergedToCorrectNC;
      this.reducedLinksPerfect_ = reducedLinksPerfect;
      this.loneNodeIDsPerfect_ = loneNodeIDsPerfect;
      this.report_ = report;
      this.linksSmall_ = linksSmall;
      this.lonersSmall_ = lonersSmall;
      this.linksLarge_ = linksLarge;
      this.lonersLarge_ = lonersLarge;
      this.mapG1toG2_ = mapG1toG2;
      this.perfectG1toG2_ = perfectG1toG2;
    }
    
    public Object getEarlyResult() {
      return (new Boolean(false));
    }
  
    public Object runCore() throws AsynchExitRequestException {
  
      NetworkAlignmentScorer scorer = new NetworkAlignmentScorer(reducedLinks_, loneNodeIDs_, mergedToCorrectNC_,
              nodeColorMap_, nodeColorMapPerfect_, reducedLinksPerfect_, loneNodeIDsPerfect_,
              linksSmall_, lonersSmall_, linksLarge_, lonersLarge_, 
              mapG1toG2_, perfectG1toG2_, bfwk_.getMonitor(), rMan_);
  
      this.report_.replaceValuesTo(scorer.getNetAlignStats());
      
      return (new Boolean(true));
    }
  
    public Object postRunCore() {
      return (null);
    }
  }
  
  /***************************************************************************
  **
  ** For XML I/O
  */  
      
  public static class PlugInWorker extends AbstractFactoryClient {
   
    private NetworkAlignmentPlugIn plugin_;
   
    public PlugInWorker(PluginWhiteboard board, NetworkAlignmentPlugIn plugin) {
      super(board);
      plugin_ = plugin;
      String name = plugin.getClass().getName();
      myKeys_.add(name);
      installWorker(new NetAlignStatsWorker(board), new NetAlignStatsGlue());
    }
    
    protected Object localProcessElement(String elemName, Attributes attrs) throws IOException {
      Object retval = null;
      PluginWhiteboard board = (PluginWhiteboard)this.sharedWhiteboard_;
      if (myKeys_.contains(elemName)) {
        board.setCurrentPlugIn(plugin_);
        retval = board.getCurrentPlugIn();
      }
      return (retval);     
    }  
  }
  
  /***************************************************************************
  **
  ** For XML I/O
  */
  
  public static class NetAlignStatsWorker extends AbstractFactoryClient {
        
    public NetAlignStatsWorker(PluginWhiteboard board) {
      super(board);
      myKeys_.add("NetAlignStats");
      installWorker(new NetAlignMeasureWorker(board), null);
    }
    
    protected Object localProcessElement(String elemName, Attributes attrs) throws IOException {
      Object retval = null;
      PluginWhiteboard board = (PluginWhiteboard) this.sharedWhiteboard_;
      board.setCurrentPlugInData(new NetAlignStats());
      retval = board.getCurrentPlugInData();
      return (retval);
    }
  }
  
  public static class NetAlignStatsGlue implements GlueStick {    
    public Object glueKidToParent(Object kidObj, AbstractFactoryClient parentWorker, Object optionalArgs) throws IOException {
      PluginWhiteboard board = (PluginWhiteboard) optionalArgs;
      board.getCurrentPlugIn().attachXMLData(board.getCurrentPlugInData());
      return null;
    }
  } 
  
  /***************************************************************************
  **
  ** For XML I/O
  */
  
  public static class NetAlignMeasureWorker extends AbstractFactoryClient {
        
    public NetAlignMeasureWorker(PluginWhiteboard board) {
      super(board);
      myKeys_.add("NetAlignMeasure");
    }
    
    protected Object localProcessElement(String elemName, Attributes attrs) throws IOException {
      Object retval = null;
      PluginWhiteboard board = (PluginWhiteboard) this.sharedWhiteboard_;    
      ((NetAlignStats)board.getCurrentPlugInData()).addAMeasure(buildFromXML(elemName, attrs));
      return (retval);
    }
    
    private NetAlignMeasure buildFromXML(String elemName, Attributes attrs) throws IOException {
      String name = AttributeExtractor.extractAttribute(elemName, attrs, "NetAlignMeasure", "name", true);
      name = CharacterEntityMapper.unmapEntities(name, false);
      String valStr = AttributeExtractor.extractAttribute(elemName, attrs, "NetAlignMeasure", "val", true);
      
      NetAlignMeasure retval;
      try {
        Double value = Double.valueOf(valStr);
        retval = new NetAlignMeasure(name, value);
      } catch (NumberFormatException nfex) {
        throw new IOException();
      }
      return (retval);
    }  
  }
  
}
```
