## Supplementary material for "BioFabric Visualization of Network Alignments": VISNAB Source Code: NetworkAlignmentScorer.java

```
/*
**
**    Copyright (C) 2018 Rishi Desai
**
**    Copyright (C) 2003-2018 Institute for Systems Biology
**                            Seattle, Washington, USA.
**
**    This library is free software; you can redistribute it and/or
**    modify it under the terms of the GNU Lesser General Public
**    License as published by the Free Software Foundation; either
**    version 2.1 of the License, or (at your option) any later version.
**
**    This library is distributed in the hope that it will be useful,
**    but WITHOUT ANY WARRANTY; without even the implied warranty of
**    MERCHANTABILITY or FITNESS FOR A PARTICULAR PURPOSE.  See the GNU
**    Lesser General Public License for more details.
**
**    You should have received a copy of the GNU Lesser General Public
**    License along with this library; if not, write to the Free Software
**    Foundation, Inc., 59 Temple Place, Suite 330, Boston, MA  02111-1307  USA
*/

package org.systemsbiology.biofabric.plugin.core.align;

import java.util.ArrayList;
import java.util.Arrays;
import java.util.HashMap;
import java.util.HashSet;
import java.util.List;
import java.util.Map;
import java.util.Set;

import org.systemsbiology.biofabric.api.model.NetLink;
import org.systemsbiology.biofabric.api.model.NetNode;
import org.systemsbiology.biofabric.api.util.PluginResourceManager;
import org.systemsbiology.biofabric.api.worker.AsynchExitRequestException;
import org.systemsbiology.biofabric.api.worker.BTProgressMonitor;
import org.systemsbiology.biofabric.api.worker.LoopReporter;
import org.systemsbiology.biofabric.plugin.PluginSupportFactory;

/****************************************************************************
 **
 ** Calculates topological scores of network alignments: Edge Coverage (EC),
 ** Symmetric Substructure score (S3), Induced Conserved Substructure (ICS);
 **
 ** Node Correctness (NC) and Jaccard Similarity (JS) are calculable
 ** only if we have the perfect alignment.
 **
 ** NGS and LGS are the angular similarity between the normalized ratio vectors
 ** of the respective node groups and link groups of the main
 ** alignment and the perfect alignment.
 */

public class NetworkAlignmentScorer {
  
  ////////////////////////////////////////////////////////////////////////////
  //
  // PRIVATE INSTANCE MEMBERS
  //
  ////////////////////////////////////////////////////////////////////////////
  
  //
  // Keep track of both the main alignment and perfect alignment's info
  //
  
  private Set<NetLink> linksMain_, linksPerfect_;
  private Set<NetNode> loneNodeIDsMain_, loneNodeIDsPerfect_;
  private NetworkAlignment.NodeColorMap nodeColorMapMain_, nodeColorMapPerfect_;
  private Map<NetNode, Boolean> mergedToCorrectNC_;
  
  private Map<NetNode, Set<NetLink>> nodeToLinksMain_, nodeToLinksPerfect_;
  private Map<NetNode, Set<NetNode>> nodeToNeighborsMain_, nodeToNeighborsPerfect_;
  private NodeGroupMap groupMapMain_, groupMapPerfect_;
  
  //
  // This are from original untouched graphs and alignments
  //
  
  private ArrayList<NetLink> linksSmall_, linksLarge_;
  private HashSet<NetNode> lonersSmall_, lonersLarge_;
  private Map<NetNode, NetNode> mapG1toG2_, perfectG1toG2_;
  
  private BTProgressMonitor monitor_;
  private PluginResourceManager rMan_;
  
  //
  // The scores
  //
  
  private Double EC, S3, ICS, NC, NGS, LGS, JaccSim;
  private NetworkAlignmentPlugIn.NetAlignStats netAlignStats_;
  
  ////////////////////////////////////////////////////////////////////////////
  //
  // CONSTRUCTOR
  //
  ////////////////////////////////////////////////////////////////////////////
  
  public NetworkAlignmentScorer(Set<NetLink> reducedLinks, Set<NetNode> loneNodeIDs,
                                Map<NetNode, Boolean> mergedToCorrectNC,
                                NetworkAlignment.NodeColorMap nodeColorMap, NetworkAlignment.NodeColorMap nodeColorMapPerfect,
                                Set<NetLink> linksPerfect, Set<NetNode> loneNodeIDsPerfect,
                                ArrayList<NetLink> linksSmall, HashSet<NetNode> lonersSmall,
                                ArrayList<NetLink> linksLarge, HashSet<NetNode> lonersLarge,
                                Map<NetNode, NetNode> mapG1toG2, Map<NetNode, NetNode> perfectG1toG2,
                                BTProgressMonitor monitor, PluginResourceManager rMan) throws AsynchExitRequestException {
  	this.rMan_ = rMan;
    this.linksMain_ = new HashSet<NetLink>(reducedLinks);
    this.loneNodeIDsMain_ = new HashSet<NetNode>(loneNodeIDs);
    this.nodeColorMapMain_ = nodeColorMap;
    this.nodeColorMapPerfect_ = nodeColorMapPerfect;
    this.mergedToCorrectNC_ = mergedToCorrectNC;
    this.linksPerfect_ = linksPerfect;
    this.loneNodeIDsPerfect_ = loneNodeIDsPerfect;
    this.monitor_ = monitor;
    this.nodeToLinksMain_ = new HashMap<NetNode, Set<NetLink>>();
    this.nodeToNeighborsMain_ = new HashMap<NetNode, Set<NetNode>>();
    this.nodeToLinksPerfect_ = new HashMap<NetNode, Set<NetLink>>();
    this.nodeToNeighborsPerfect_ = new HashMap<NetNode, Set<NetNode>>();
    
    this.linksSmall_ = linksSmall;
    this.lonersSmall_ = lonersSmall;
    this.linksLarge_ = linksLarge;
    this.lonersLarge_ = lonersLarge;
    this.mapG1toG2_ = mapG1toG2;
    this.perfectG1toG2_ = perfectG1toG2;
    
    // Create Node Group Map to use for NGS/LGS
    this.groupMapMain_ = new NodeGroupMap(linksMain_, loneNodeIDsMain_, nodeColorMapMain_,
            linksPerfect_, loneNodeIDsPerfect_, nodeColorMapPerfect_,
            mergedToCorrectNC_, NodeGroupMap.PerfectNGMode.NONE, null,
            linksSmall, lonersSmall, linksLarge, lonersLarge, mapG1toG2, perfectG1toG2,
            NodeGroupMap.nodeGroupOrder, NodeGroupMap.nodeGroupAnnots, monitor_);
    if (mergedToCorrectNC != null) { // perfect alignment processing does not need a 'perfect' alignment
      this.groupMapPerfect_ = new NodeGroupMap(linksPerfect_, loneNodeIDsPerfect_, nodeColorMapPerfect_, null, null, null, null, NodeGroupMap.PerfectNGMode.NONE, null,
              linksSmall, lonersSmall, linksLarge, lonersLarge, mapG1toG2, perfectG1toG2,
              NodeGroupMap.nodeGroupOrder, NodeGroupMap.nodeGroupAnnots, monitor_);
    }
    
    removeDuplicateAndShadow();
    // Generate Structures
    PluginSupportFactory.getBuildExtractor().createNeighborLinkMap(linksMain_, loneNodeIDsMain_,
            nodeToNeighborsMain_, nodeToLinksMain_, monitor_);
  
    if (mergedToCorrectNC != null) {
      PluginSupportFactory.getBuildExtractor().createNeighborLinkMap(linksPerfect_, loneNodeIDsPerfect_,
              nodeToNeighborsPerfect_, nodeToLinksPerfect_, monitor_);
    }
    calcScores();
    finalizeMeasures();
    return;
  }
  
  ////////////////////////////////////////////////////////////////////////////
  //
  // PRIVATE METHODS
  //
  ////////////////////////////////////////////////////////////////////////////
  
  private void removeDuplicateAndShadow() throws AsynchExitRequestException {
    LoopReporter lr = new LoopReporter(linksMain_.size(), 20, monitor_, 0.0, 1.0, "progress.filteringLinksA");
    Set<NetLink> nonShdwLinks = new HashSet<NetLink>();
    for (NetLink link : linksMain_) {
      lr.report();
      if (! link.isShadow()) { // remove shadow links
        nonShdwLinks.add(link);
      }
    }
    
    //
    // We have to remove synonymous links (a->b) same as (b->a), and keep one;
    // Sort the names and concat into string (the key), so they are the same key in the map.
    // This means (a->b) and (b->a) should make the same string key.
    // If the key already has a value, we got a duplicate link.
    //
  
    lr = new LoopReporter(nonShdwLinks.size(), 20, monitor_, 0.0, 1.0, "progress.filteringLinksB");
    Map<String, NetLink> map = new HashMap<String, NetLink>();
    for (NetLink link : nonShdwLinks) {
      lr.report();
      String[] arr1 = {link.getSrcNode().getName(), link.getTrgNode().getName()};
      Arrays.sort(arr1);
      String concat = String.format("%s___%s", arr1[0], arr1[1]);
      
      if (map.get(concat) == null) {
        map.put(concat, link);
      } // skip duplicates
    }
    
    linksMain_.clear();
    for (Map.Entry<String, NetLink> entry : map.entrySet()) {
      linksMain_.add(entry.getValue());
    }
    return;
  }
  
  /****************************************************************************
   **
   ** Calculate the scores!
   */
  
  private void calcScores() throws AsynchExitRequestException {
    calcTopologicalMeasures();
  
    if (mergedToCorrectNC_ != null) { // must have perfect alignment for these measures
      calcNodeCorrectness();
      calcGroupSimilarity();
      calcJaccardSimilarity();
    }
  }
  
  /****************************************************************************
   **
   ** Create the Measure list and filter out 'null' measures
   */
  
  private void finalizeMeasures() {

    String
            ECn = rMan_.getPluginString("networkAlignment.edgeCoverage"),
            S3n = rMan_.getPluginString("networkAlignment.symmetricSubstructureScore"),
            ICSn = rMan_.getPluginString("networkAlignment.inducedConservedStructure"),
            NCn = rMan_.getPluginString("networkAlignment.nodeCorrectness"),
            NGSn = rMan_.getPluginString("networkAlignment.nodeGroupSimilarity"),
            LGSn = rMan_.getPluginString("networkAlignment.linkGroupSimilarity"),
            JSn = rMan_.getPluginString("networkAlignment.jaccardSimilarity");
    
    NetworkAlignmentPlugIn.NetAlignMeasure[] possibleMeasures = {
            new NetworkAlignmentPlugIn.NetAlignMeasure(ECn, EC),
            new NetworkAlignmentPlugIn.NetAlignMeasure(S3n, S3),
            new NetworkAlignmentPlugIn.NetAlignMeasure(ICSn, ICS),
            new NetworkAlignmentPlugIn.NetAlignMeasure(NCn, NC),
            new NetworkAlignmentPlugIn.NetAlignMeasure(NGSn, NGS),
            new NetworkAlignmentPlugIn.NetAlignMeasure(LGSn, LGS),
            new NetworkAlignmentPlugIn.NetAlignMeasure(JSn, JaccSim),
    };
  
    List<NetworkAlignmentPlugIn.NetAlignMeasure> measures = new ArrayList<NetworkAlignmentPlugIn.NetAlignMeasure>();
    for (NetworkAlignmentPlugIn.NetAlignMeasure msr : possibleMeasures) {
      if (msr.val != null) { // no point having null measures
        measures.add(msr);
      }
    }
    this.netAlignStats_ = new NetworkAlignmentPlugIn.NetAlignStats(measures);
    return;
  }
  
  private void calcTopologicalMeasures() throws AsynchExitRequestException{
    LoopReporter lr = new LoopReporter(linksMain_.size(), 20, monitor_, 0.0, 1.0, "progress.topologicalMeasures");
    int numCoveredEdge = 0, numInducedGraph1 = 0, numInducedGraph2 = 0;
    
    for (NetLink link : linksMain_) {
      lr.report();
      if (link.getRelation().equals(NetworkAlignment.EdgeType.COVERED.tag)) {
        numCoveredEdge++;
      } else if (link.getRelation().equals(NetworkAlignment.EdgeType.INDUCED_GRAPH1.tag)) {
        numInducedGraph1++;
      } else if (link.getRelation().equals(NetworkAlignment.EdgeType.INDUCED_GRAPH2.tag)) {
        numInducedGraph2++;
      }
    }
    
    EC = ((double) numCoveredEdge) / (numCoveredEdge + numInducedGraph1);
    S3 = ((double) numCoveredEdge) / (numCoveredEdge + numInducedGraph1 + numInducedGraph2);
    ICS = ((double) numCoveredEdge) / (numCoveredEdge + numInducedGraph2);
    
    // These three conditionals account for singleton node networks alignments where you get 0/0 for all three
    if (EC.isNaN()) {
      EC = null;
    }
    if (S3.isNaN()) {
      S3 = null;
    }
    if (ICS.isNaN()) {
      ICS = null;
    }
    return;
  }
  
  private void calcNodeCorrectness() {
    if (mergedToCorrectNC_ == null) {
      NC = null;
      return;
    }
    
    int numCorrect = 0;
    for (Map.Entry<NetNode, Boolean> node : mergedToCorrectNC_.entrySet()) {
      if (node.getValue()) {
        numCorrect++;
      }
    }
    NC = ((double)numCorrect) / (mergedToCorrectNC_.size());
    return;
  }
  
  private void calcGroupSimilarity() {
    GroupSimilarityMeasure gd = new GroupSimilarityMeasure();
    NGS = gd.calcNGS(groupMapMain_, groupMapPerfect_);
    LGS = gd.calcLGS(groupMapMain_, groupMapPerfect_);
    return;
  }
  
  private void calcJaccardSimilarity() throws AsynchExitRequestException {
    this.JaccSim = (new JaccardSimilarity(linksMain_, loneNodeIDsMain_, nodeColorMapMain_,
            linksPerfect_, loneNodeIDsPerfect_, nodeColorMapPerfect_, nodeToNeighborsMain_, nodeToLinksMain_,
            nodeToNeighborsPerfect_, nodeToLinksPerfect_, linksSmall_, lonersSmall_, linksLarge_, lonersLarge_,
            mapG1toG2_, perfectG1toG2_, null, monitor_)).calcScore();
    return;
  }
  
  public NetworkAlignmentPlugIn.NetAlignStats getNetAlignStats() {
    return (netAlignStats_);
  }
  
  ////////////////////////////////////////////////////////////////////////////
  //
  // INNER CLASSES
  //
  ////////////////////////////////////////////////////////////////////////////
  
  /****************************************************************************
   **
   ** N-dimensional vector used for scores
   */

  private static class VectorND {
  
    private double[] values_;
  
    public VectorND(int size) {
      this.values_ = new double[size];
    }
  
    public double get(int index) {
      return (values_[index]);
    }
  
    public void set(int index, double val) {
      values_[index] = val;
      return;
    }
  
    /****************************************************************************
     **
     ** Euclidean distance between two vectors
     */
  
    public double distance(VectorND vector) {
      if (this.values_.length!= vector.values_.length) {
        throw new IllegalArgumentException("score vector length not equal");
      }
      double ret = 0;
      for (int i = 0; i < values_.length; i++) {
        ret += (this.values_[i] - vector.values_[i]) * (this.values_[i] - vector.values_[i]);
      }
      ret = Math.pow(ret, .5);
      return (ret);
    }
  
    /****************************************************************************
     **
     ** Magnitude
     */
  
    public double magnitude() {
      double ret = this.dot(this);
      ret = Math.sqrt(ret);
      return (ret);
    }
  
    /****************************************************************************
     **
     ** Normalize
     */
  
    public void normalize() {
      double mag = magnitude();
      if (mag == 0) {
        return;
      }
      for (int i = 0; i < values_.length; i++) {
        values_[i] /= mag;
      }
      return;
    }
  
    /****************************************************************************
     **
     ** Dot product
     */
  
    public double dot(VectorND vector) {
      if (this.values_.length != vector.values_.length) {
        throw new IllegalArgumentException("score vector length not equal");
      }
      double ret = 0;
      for (int i = 0; i < this.values_.length; i++) {
        ret += this.values_[i] * vector.values_[i];
      }
      return (ret);
    }
  
    /****************************************************************************
     **
     ** Cosine similarity = returns cos(angle)
     */
  
    public double cosSim(VectorND vector) {
      double cosTheta = dot(vector) / (this.magnitude() * vector.magnitude());
      return (cosTheta);
    }
  
    /****************************************************************************
     **
     ** Angular Similarity = 1 - (2 * arccos(similarity) / pi)
     **
     ** similarity = cos(angle)
     */
    
    public double angSim(VectorND vector) {
      Double cosT = cosSim(vector);
      
      if (Double.compare(cosT, 1.0) > 0) { // fix for RishiDesai issue #36 (NaN was appearing)
        cosT = 1.0;
      } else if (Double.compare(cosT, 0.0) < 0) {
        cosT = 0.0;
      }
      Double sim = 1 - (2 * Math.acos(cosT) / Math.PI);
      if (sim.isNaN()) {
        // should not happen
      }
      return (sim);
    }
    
    @Override
    public String toString() {
      return "VectorND{" +
              "values_=" + Arrays.toString(values_) +
              '}';
    }
  
    @Override
    public boolean equals(Object o) {
      if (this == o) return (true);
      if (! (o instanceof VectorND)) {
        return (false);
      }
      VectorND vectorND = (VectorND) o;
      if (! Arrays.equals(values_, vectorND.values_)) {
        return (false);
      }
      return (true);
    }
  
    @Override
    public int hashCode() {
      return (Arrays.hashCode(values_));
    }
    
  }
  
  /****************************************************************************
   **
   ** NGS and LGS - with Angular similarity
   */
  
  private static class GroupSimilarityMeasure {
  
    /***************************************************************************
     **
     ** Calculated the score
     */
  
    double calcNGS(NodeGroupMap groupMapMain, NodeGroupMap groupMapPerfect) {
      VectorND main = getNGVector(groupMapMain), perfect = getNGVector(groupMapPerfect);
      double score = main.angSim(perfect);
      return (score);
    }
  
    /***************************************************************************
     **
     ** Convert ratio to vector
     */
    
    private VectorND getNGVector(NodeGroupMap groupMap) {
      VectorND vector = new VectorND(groupMap.numGroups());
      Map<String, Double> ngRatios = groupMap.getNodeGroupRatios();
  
      for (Map.Entry<String, Double> entry : ngRatios.entrySet()) {
        int index = groupMap.getIndex(entry.getKey());
        vector.set(index, entry.getValue());
      }
      vector.normalize();
      return (vector);
    }
  
    /***************************************************************************
     **
     ** Calculated the score
     */
  
    double calcLGS(NodeGroupMap groupMapMain, NodeGroupMap groupMapPerfect) {
      VectorND main = getLGVector(groupMapMain), perfect = getLGVector(groupMapPerfect);
      double score = main.angSim(perfect);
      return (score);
    }
  
    /***************************************************************************
     **
     ** Convert ratio to vector
     */
    
    private VectorND getLGVector(NodeGroupMap groupMap) {

      Map<String, Integer> relToIndex = new HashMap<String, Integer>();
      for (NetworkAlignment.EdgeType type : NetworkAlignment.LINK_GROUPS) {
        relToIndex.put(type.tag, type.index);
      }

      VectorND vector = new VectorND(NetworkAlignment.LINK_GROUPS.length);
      Map<String, Double> lgRatios = groupMap.getLinkGroupRatios();
      
      for (Map.Entry<String, Double> entry : lgRatios.entrySet()) {
        int index = relToIndex.get(entry.getKey());
        vector.set(index, entry.getValue());
      }
      vector.normalize();
      return (vector);
    }
    
  }

}
```
