## Supplementary material for "BioFabric Visualization of Network Alignments": VISNAB Source Code: NodeGroupMap.java

```
/*
**
**    Copyright (C) 2018-2019 Rishi Desai
**
**    Copyright (C) 2003-2018 Institute for Systems Biology
**                            Seattle, Washington, USA.
**
**    This library is free software; you can redistribute it and/or
**    modify it under the terms of the GNU Lesser General Public
**    License as published by the Free Software Foundation; either
**    version 2.1 of the License, or (at your option) any later version.
**
**    This library is distributed in the hope that it will be useful,
**    but WITHOUT ANY WARRANTY; without even the implied warranty of
**    MERCHANTABILITY or FITNESS FOR A PARTICULAR PURPOSE.  See the GNU
**    Lesser General Public License for more details.
**
**    You should have received a copy of the GNU Lesser General Public
**    License along with this library; if not, write to the Free Software
**    Foundation, Inc., 59 Temple Place, Suite 330, Boston, MA  02111-1307  USA
*/

package org.systemsbiology.biofabric.plugin.core.align;

import java.util.ArrayList;
import java.util.Comparator;
import java.util.HashMap;
import java.util.HashSet;
import java.util.List;
import java.util.Map;
import java.util.Set;
import org.systemsbiology.biofabric.api.io.BuildData;
import org.systemsbiology.biofabric.api.model.NetLink;
import org.systemsbiology.biofabric.api.model.NetNode;
import org.systemsbiology.biofabric.api.worker.AsynchExitRequestException;
import org.systemsbiology.biofabric.api.worker.BTProgressMonitor;
import org.systemsbiology.biofabric.api.worker.LoopReporter;
import org.systemsbiology.biofabric.plugin.PluginSupportFactory;

/***************************************************************************
 **
 ** HashMap based data structure
 **
 ** There are 40 DISTINCT Node Groups for aligned and unaligned nodes
 ** There are 76 DISTINCT Node Groups for aligned and unaligned nodes with Perfect NGs
 **
 ** With Perfect NodeGroups activated, the correctness of a node is determined by either NC or JS
 */

public class NodeGroupMap {
  
  ////////////////////////////////////////////////////////////////////////////
  //
  // PUBLIC MEMBERS
  //
  ////////////////////////////////////////////////////////////////////////////
  
  public enum PerfectNGMode {
    NONE, NODE_CORRECTNESS, JACCARD_SIMILARITY
  }
  
  ////////////////////////////////////////////////////////////////////////////
  //
  // PRIVATE INSTANCE MEMBERS
  //
  ////////////////////////////////////////////////////////////////////////////
  
  private final PerfectNGMode mode_;
  private Set<NetLink> links_;
  private Set<NetNode> loners_;
  private Map<NetNode, Boolean> mergedToCorrectNC_;
  private NetworkAlignment.NodeColorMap nodeColorMap_;
  
  private Map<NetNode, Set<NetLink>> nodeToLinks_;
  private Map<NetNode, Set<NetNode>> nodeToNeighbors_;
  
  private Map<GroupID, Integer> groupIDtoIndex_;
  private Map<Integer, GroupID> indexToGroupID_;
  private Map<GroupID, String> groupIDtoColor_;
  private final int numGroups_;
  
  private Map<String, Double> nodeGroupRatios_, linkGroupRatios_;
  private JaccardSimilarity funcJS_;
  private BTProgressMonitor monitor_;
  
  ////////////////////////////////////////////////////////////////////////////
  //
  // CONSTRUCTORS
  //
  ////////////////////////////////////////////////////////////////////////////
  
  public NodeGroupMap(BuildData bd, String[] nodeGroupOrder, String[][] colorMap,
                      BTProgressMonitor monitor) throws AsynchExitRequestException {
    this(bd.getLinks(), 
    		 bd.getSingletonNodes(), 
    		 ((NetworkAlignmentBuildData)bd.getPluginBuildData()).colorMapMain,
    		 ((NetworkAlignmentBuildData)bd.getPluginBuildData()).allLinksPerfect,
    		 ((NetworkAlignmentBuildData)bd.getPluginBuildData()).loneNodeIDsPerfect,
    		 ((NetworkAlignmentBuildData)bd.getPluginBuildData()).colorMapPerfect,
             ((NetworkAlignmentBuildData)bd.getPluginBuildData()).mergedToCorrectNC,
             ((NetworkAlignmentBuildData)bd.getPluginBuildData()).mode,
             ((NetworkAlignmentBuildData)bd.getPluginBuildData()).jaccSimThreshold,
             ((NetworkAlignmentBuildData)bd.getPluginBuildData()).linksSmall,
             ((NetworkAlignmentBuildData)bd.getPluginBuildData()).lonersSmall,
             ((NetworkAlignmentBuildData)bd.getPluginBuildData()).linksLarge,
             ((NetworkAlignmentBuildData)bd.getPluginBuildData()).lonersLarge,
             ((NetworkAlignmentBuildData)bd.getPluginBuildData()).mapG1toG2,
             ((NetworkAlignmentBuildData)bd.getPluginBuildData()).perfectG1toG2,
         nodeGroupOrder,
         colorMap, 
         monitor);
  }
  
  public NodeGroupMap(Set<NetLink> allLinksMain, Set<NetNode> loneNodeIDsMain,
                        NetworkAlignment.NodeColorMap colorMapMain,
                        Set<NetLink> allLinksPerfect, Set<NetNode> loneNodeIDsPerfect,
                        NetworkAlignment.NodeColorMap colorMapPerfect,
                        Map<NetNode, Boolean> mergedToCorrectNC,
                        PerfectNGMode mode, final Double jaccSimThreshold,
                        ArrayList<NetLink> linksSmall, HashSet<NetNode> lonersSmall,
                        ArrayList<NetLink> linksLarge, HashSet<NetNode> lonersLarge,
                        Map<NetNode, NetNode> mapG1toG2, Map<NetNode, NetNode> perfectG1toG2,
                        String[] nodeGroupOrder, String[][] colorMap,
                        BTProgressMonitor monitor) throws AsynchExitRequestException {
    
    this.links_ = allLinksMain;
    this.loners_ = loneNodeIDsMain;
    this.mergedToCorrectNC_ = mergedToCorrectNC;
    this.nodeColorMap_ = colorMapMain;
    this.numGroups_ = nodeGroupOrder.length;
    this.mode_ = mode;
    this.monitor_ = monitor;
    this.nodeToNeighbors_ = new HashMap<NetNode, Set<NetNode>>();
    this.nodeToLinks_ = new HashMap<NetNode, Set<NetLink>>();
    PluginSupportFactory.getBuildExtractor().createNeighborLinkMap(links_, loners_, nodeToNeighbors_, nodeToLinks_, monitor_);
    
    if (mode == PerfectNGMode.JACCARD_SIMILARITY) { // create structures for JS involving perfect alignment
      Map<NetNode, Set<NetNode>> nodeToNeighborsPerfect = new HashMap<NetNode, Set<NetNode>>();
      Map<NetNode, Set<NetLink>> nodeToLinksPerfect = new HashMap<NetNode, Set<NetLink>>();
      PluginSupportFactory.getBuildExtractor().createNeighborLinkMap(allLinksPerfect, loneNodeIDsPerfect,
              nodeToNeighborsPerfect, nodeToLinksPerfect, monitor_);
      
      this.funcJS_ = new JaccardSimilarity(allLinksMain, loneNodeIDsMain, colorMapMain, allLinksPerfect, loneNodeIDsPerfect,
              colorMapPerfect, nodeToNeighbors_, nodeToLinks_, nodeToNeighborsPerfect, nodeToLinksPerfect,
              linksSmall, lonersSmall, linksLarge, lonersLarge, mapG1toG2, perfectG1toG2, jaccSimThreshold, monitor);
    }
    generateOrderMap(nodeGroupOrder);
    generateColorMap(colorMap);
    calcNGRatios();
    calcLGRatios();
    return;
  }
  
  //////////////////////////////////////////////////////////////////////////
  //
  //  PRIVATE METHODS
  //
  //////////////////////////////////////////////////////////////////////////
  
  private void generateOrderMap(String[] nodeGroupOrder) {
    groupIDtoIndex_ = new HashMap<GroupID, Integer>();
    indexToGroupID_ = new HashMap<Integer, GroupID>();
    for (int index = 0; index < nodeGroupOrder.length; index++) {
      GroupID gID = new GroupID(nodeGroupOrder[index]);
      groupIDtoIndex_.put(gID, index);
      indexToGroupID_.put(index, gID);
    }
    return;
  }
  
  private void generateColorMap(String[][] colorMap) {
    groupIDtoColor_ = new HashMap<GroupID, String>();
    
    for (String[] ngCol : colorMap) {
      GroupID groupID = new GroupID(ngCol[0]);
      String color = ngCol[1];
      groupIDtoColor_.put(groupID, color);
    }
    return;
  }
  
  /***************************************************************************
   **
   ** Hash function
   */
  
  private GroupID generateID(NetNode node) {
    //
    // See which types of link groups the node's links are in
    //
    
    boolean[] inLG = new boolean[NetworkAlignment.LINK_GROUPS.length];
    
    
    for (NetLink link : nodeToLinks_.get(node)) {
      for (int rel = 0; rel < inLG.length; rel++) {
        if (link.getRelation().equals(NetworkAlignment.LINK_GROUPS[rel].tag)) {
          inLG[rel] = true;
        }
      }
    }
  
    List<String> tags = new ArrayList<String>();
    
    for (NetworkAlignment.EdgeType type : NetworkAlignment.LINK_GROUPS) {
      if (inLG[type.index]) {
        tags.add(type.tag);
      }
    }
    
    if (tags.isEmpty()) { // singletons
      tags.add("0");
    }
    
    StringBuilder sb = new StringBuilder();
    sb.append("(");
    sb.append(nodeColorMap_.getColor(node).tag);  // node color (P,B,R)
    sb.append(":");
  
    for (int i = 0; i < tags.size(); i++) {
      sb.append(tags.get(i));    // link group tags
      if (i != tags.size() - 1) {
        sb.append("/");
      }
    }
    
    if (mode_ != PerfectNGMode.NONE) {   // perfect NG mode is activated
      sb.append("/");
      if (mergedToCorrectNC_.get(node) == null) {
        sb.append(0);
      } else {
        boolean isCorrect;
        if (mode_ == PerfectNGMode.NODE_CORRECTNESS) {
          isCorrect = mergedToCorrectNC_.get(node);
        } else if (mode_ == PerfectNGMode.JACCARD_SIMILARITY) {
          isCorrect = funcJS_.isCorrectJS(node);
        } else {
          throw new IllegalStateException("Incorrect mode for Perfect NGs Group Map");
        }
        sb.append((isCorrect) ? 1 : 0);
      }
    }
    sb.append(")");
    
    return (new GroupID(sb.toString()));
  }
  
  /***************************************************************************
   **
   ** Calculate node group size to total #nodes for each group
   */
  
  private void calcNGRatios() {
    Set<NetNode> nodes = nodeToLinks_.keySet();
    double size = nodes.size();
    Set<GroupID> tags = groupIDtoIndex_.keySet();
  
    Map<GroupID, Integer> counts = new HashMap<GroupID, Integer>(); // initial vals
    for (GroupID gID : tags) {
      counts.put(gID, 0);
    }
    
    for (NetNode node : nodes) {
      GroupID gID = generateID(node);
      counts.put(gID, counts.get(gID) + 1);
    }
  
    nodeGroupRatios_ = new HashMap<String, Double>();
    for (Map.Entry<GroupID, Integer> count : counts.entrySet()) {
      String tag = count.getKey().getKey();
      double ratio = count.getValue() / size;
      nodeGroupRatios_.put(tag, ratio);
    }
    return;
  }
  
  /***************************************************************************
   **
   ** Calculate link group size to total #links for each group
   */
  
  private void calcLGRatios() throws AsynchExitRequestException {
    
    double size = links_.size();
    
    Map<String, Integer> counts = new HashMap<String, Integer>(); // initial vals
    for (NetworkAlignment.EdgeType type : NetworkAlignment.LINK_GROUPS) {
      counts.put(type.tag, 0);
    }
    
    LoopReporter lr = new LoopReporter(links_.size(), 20, monitor_, 0.0, 1.0, "progress.calculatingLinkRatios");
    for (NetLink link : links_) {
      lr.report();
      String rel = link.getRelation();
      counts.put(rel, counts.get(rel) + 1);
    }
    
    linkGroupRatios_ = new HashMap<String, Double>();
    for (Map.Entry<String, Integer> count : counts.entrySet()) {
      String tag = count.getKey();
      double ratio = count.getValue() / size;
      linkGroupRatios_.put(tag, ratio);
    }
    return;
  }
  
  ////////////////////////////////////////////////////////////////////////////
  //
  // PUBLIC METHODS
  //
  ////////////////////////////////////////////////////////////////////////////
  
  /***************************************************************************
   **
   ** Return the index in given ordering given node
   */
  
  public int getIndex(NetNode node) {
    GroupID groupID = generateID(node);
    Integer index = groupIDtoIndex_.get(groupID);
    if (index == null) {
      throw new IllegalArgumentException("GroupID " + groupID + " not found in given order list; given node " + node.getName());
    }
    return (index);
  }
  
  /***************************************************************************
   **
   ** Return the index from the given ordering given tag
   */
  
  public int getIndex(String key) {
    Integer index = groupIDtoIndex_.get(new GroupID(key));
    if (index == null) {
      throw new IllegalArgumentException("GroupID " + key + " not found in given order list; given key " + key);
    }
    return (index);
  }

  /***************************************************************************
   **
   ** Return the GroupID from the given node group ordering
   */
  
  public String getKey(Integer index) {
    if (indexToGroupID_.get(index) == null) {
      throw new IllegalArgumentException("Index not found in given order list; given index " + index);
    }
    return (indexToGroupID_.get(index).getKey());
  }
 
  /***************************************************************************
   **
   ** Return the Annot Color for index of NG label
   */
  
  public String getColor(Integer index) {
    GroupID groupID = indexToGroupID_.get(index);
    return (groupIDtoColor_.get(groupID));
  }
  
  public int numGroups() {
    return (numGroups_);
  }
  
  public Map<String, Double> getNodeGroupRatios() {
    return (nodeGroupRatios_);
  }
  
  public Map<String, Double> getLinkGroupRatios() {
    return (linkGroupRatios_);
  }
  
  /***************************************************************************
   **
   ** Sorts in decreasing node degree - method is here for convenience
   */
  
  public Comparator<NetNode> sortDecrDegree() {
    return (new Comparator<NetNode>() {
      public int compare(NetNode node1, NetNode node2) {
        int diffSize = nodeToNeighbors_.get(node2).size() - nodeToNeighbors_.get(node1).size();
        return (diffSize != 0) ? diffSize : node1.getName().compareTo(node2.getName());
      }
    });
  }

  ////////////////////////////////////////////////////////////////////////////
  //
  // PRIVATE CLASSES
  //
  ////////////////////////////////////////////////////////////////////////////
  
  /***************************************************************************
   **
   ** Hash
   */
  
  private static class GroupID {
    
    private final String key_;
    
    public GroupID(String key) {
      this.key_ = key;
    }
    
    public String getKey() {
      return (key_);
    }
    
    @Override
    public String toString() {
      return (key_);
    }
    
    @Override
    public boolean equals(Object o) {
      if (this == o) return (true);
      if (! (o instanceof GroupID)) return (false);
      
      GroupID groupID = (GroupID) o;
      
      if (key_ != null ? (! key_.equals(groupID.key_)) : (groupID.key_ != null)) return (false);
      return (true);
    }
    
    @Override
    public int hashCode() {
      return (key_ != null ? key_.hashCode() : 0);
    }
    
  }
  
  ////////////////////////////////////////////////////////////////////////////
  //
  // PUBLIC CONSTANTS
  //
  ////////////////////////////////////////////////////////////////////////////
  
  public static final String[] nodeGroupOrder;
  public static final String[][] nodeGroupAnnots = {  // Dark colors used for Purple nodes with any blue neighbors
          {"(P:0)",         "GrayBlue"},              // Normal colors used for original purple and red nodes (no blue nodes)
          {"(P:P)",         "Orange"},
          {"(P:pBp)",       "Yellow"},
          {"(P:pBb)",       "DarkPeach"},
          {"(P:pBp/pBb)",   "DarkPowderBlue"},
          {"(P:pRp)",           "Green"},
          {"(P:P/pBp)",         "Purple"},
          {"(P:P/pBb)",         "DarkOrange"},
          {"(P:P/pBp/pBb)",     "DarkYellow"},
          {"(P:P/pRp)",         "Pink"},
          {"(P:pBp/pRp)",       "PowderBlue"},
          {"(P:pBb/pRp)",       "DarkGreen"},
          {"(P:pBp/pBb/pRp)",   "DarkPurple"},
          {"(P:P/pBp/pRp)",     "Peach"},
          {"(P:P/pBb/pRp)",     "DarkGrayBlue"},
          {"(P:P/pBp/pBb/pRp)", "DarkPink"},
          {"(P:pRr)",           "GrayBlue"},
          {"(P:P/pRr)",         "Orange"},
          {"(P:pBp/pRr)",       "Yellow"},
          {"(P:pBb/pRr)",       "DarkPowderBlue"},
          {"(P:pBp/pBb/pRr)",   "DarkPeach"},
          {"(P:pRp/pRr)",           "Green"},
          {"(P:P/pBp/pRr)",         "Purple"},
          {"(P:P/pBb/pRr)",         "DarkGrayBlue"},
          {"(P:P/pBp/pBb/pRr)",     "DarkOrange"},
          {"(P:P/pRp/pRr)",         "Pink"},
          {"(P:pBp/pRp/pRr)",       "PowderBlue"},
          {"(P:pBb/pRp/pRr)",       "DarkYellow"},
          {"(P:pBp/pBb/pRp/pRr)",   "DarkPurple"},
          {"(P:P/pBp/pRp/pRr)",     "Peach"},
          {"(P:P/pBb/pRp/pRr)",     "DarkPink"},
          {"(P:P/pBp/pBb/pRp/pRr)", "DarkGreen"},
          {"(B:pBb)",               "PowderBlue"},
          {"(B:bBb)",               "Purple"},
          {"(B:pBb/bBb)",           "Pink"},
          {"(B:0)",                 "Peach"},
          {"(R:pRr)",           "GrayBlue"},
          {"(R:rRr)",           "Orange"},
          {"(R:pRr/rRr)",       "Yellow"},
          {"(R:0)",             "Green"}
  };
  public static final String[] nodeGroupOrderPerfectNG;
  public static final String[][] nodeGroupAnnotsPerfectNG = {  // Complement color used for 'incorrect' node group
          {"(P:0/1)",         "GrayBlue"},
          {"(P:0/0)",         "DarkGrayBlue"},
          {"(P:P/1)",         "Orange"},
          {"(P:P/0)",         "DarkOrange"},
          {"(P:pBp/1)",       "Yellow"},
          {"(P:pBp/0)",       "DarkYellow"},
          {"(P:pBb/1)",       "DarkPeach"},
          {"(P:pBb/0)",       "Peach"},
          {"(P:pBp/pBb/1)",   "DarkPowderBlue"},
          {"(P:pBp/pBb/0)",   "PowderBlue"},
          {"(P:pRp/1)",           "Green"},
          {"(P:pRp/0)",           "DarkGreen"},
          {"(P:P/pBp/1)",         "Purple"},
          {"(P:P/pBp/0)",         "DarkPurple"},
          {"(P:P/pBb/1)",         "DarkOrange"},
          {"(P:P/pBb/0)",         "Orange"},
          {"(P:P/pBp/pBb/1)",     "DarkYellow"},
          {"(P:P/pBp/pBb/0)",     "Yellow"},
          {"(P:P/pRp/1)",         "Pink"},
          {"(P:P/pRp/0)",         "DarkPink"},
          {"(P:pBp/pRp/1)",       "PowderBlue"},
          {"(P:pBp/pRp/0)",       "DarkPowderBlue"},
          {"(P:pBb/pRp/1)",       "DarkGreen"},
          {"(P:pBb/pRp/0)",       "Green"},
          {"(P:pBp/pBb/pRp/1)",   "DarkPurple"},
          {"(P:pBp/pBb/pRp/0)",   "Purple"},
          {"(P:P/pBp/pRp/1)",     "Peach"},
          {"(P:P/pBp/pRp/0)",     "DarkPeach"},
          {"(P:P/pBb/pRp/1)",     "DarkGrayBlue"},
          {"(P:P/pBb/pRp/0)",     "GrayBlue"},
          {"(P:P/pBp/pBb/pRp/1)", "DarkPink"},
          {"(P:P/pBp/pBb/pRp/0)", "Pink"},
          {"(P:pRr/1)",           "GrayBlue"},
          {"(P:pRr/0)",           "DarkGrayBlue"},
          {"(P:P/pRr/1)",         "Orange"},
          {"(P:P/pRr/0)",         "DarkOrange"},
          {"(P:pBp/pRr/1)",       "Yellow"},
          {"(P:pBp/pRr/0)",       "DarkYellow"},
          {"(P:pBb/pRr/1)",       "DarkPowderBlue"},
          {"(P:pBb/pRr/0)",       "PowderBlue"},
          {"(P:pBp/pBb/pRr/1)",   "DarkPeach"},
          {"(P:pBp/pBb/pRr/0)",   "Peach"},
          {"(P:pRp/pRr/1)",           "Green"},
          {"(P:pRp/pRr/0)",           "DarkGreen"},
          {"(P:P/pBp/pRr/1)",         "Purple"},
          {"(P:P/pBp/pRr/0)",         "DarkPurple"},
          {"(P:P/pBb/pRr/1)",         "DarkGrayBlue"},
          {"(P:P/pBb/pRr/0)",         "GrayBlue"},
          {"(P:P/pBp/pBb/pRr/1)",     "DarkOrange"},
          {"(P:P/pBp/pBb/pRr/0)",     "Orange"},
          {"(P:P/pRp/pRr/1)",         "Pink"},
          {"(P:P/pRp/pRr/0)",         "DarkPink"},
          {"(P:pBp/pRp/pRr/1)",       "PowderBlue"},
          {"(P:pBp/pRp/pRr/0)",       "DarkPowderBlue"},
          {"(P:pBb/pRp/pRr/1)",       "DarkYellow"},
          {"(P:pBb/pRp/pRr/0)",       "Yellow"},
          {"(P:pBp/pBb/pRp/pRr/1)",   "DarkPurple"},
          {"(P:pBp/pBb/pRp/pRr/0)",   "Purple"},
          {"(P:P/pBp/pRp/pRr/1)",     "Peach"},
          {"(P:P/pBp/pRp/pRr/0)",     "DarkPeach"},
          {"(P:P/pBb/pRp/pRr/1)",     "DarkPink"},
          {"(P:P/pBb/pRp/pRr/0)",     "Pink"},
          {"(P:P/pBp/pBb/pRp/pRr/1)", "DarkGreen"},
          {"(P:P/pBp/pBb/pRp/pRr/0)", "Green"},
          {"(B:pBb/1)",               "PowderBlue"},
          {"(B:pBb/0)",               "DarkPowderBlue"},
          {"(B:bBb/1)",               "Purple"},
          {"(B:bBb/0)",               "DarkPurple"},
          {"(B:pBb/bBb/1)",           "Pink"},
          {"(B:pBb/bBb/0)",           "DarkPink"},
          {"(B:0/1)",                 "Peach"},
          {"(B:0/0)",                 "DarkPeach"},
          {"(R:pRr/0)",           "GrayBlue"},
          {"(R:rRr/0)",           "Orange"},
          {"(R:pRr/rRr/0)",       "Yellow"},
          {"(R:0/0)",             "Green"}
  };
  
  static {  // create the node group order from the annot color list
    nodeGroupOrder = new String[NodeGroupMap.nodeGroupAnnots.length];
    for (int i = 0; i < NodeGroupMap.nodeGroupOrder.length; i++) {
      NodeGroupMap.nodeGroupOrder[i] = NodeGroupMap.nodeGroupAnnots[i][0];
    }
    
    nodeGroupOrderPerfectNG = new String[NodeGroupMap.nodeGroupAnnotsPerfectNG.length];
    for (int i = 0; i < NodeGroupMap.nodeGroupOrderPerfectNG.length; i++) {
      NodeGroupMap.nodeGroupOrderPerfectNG[i] = NodeGroupMap.nodeGroupAnnotsPerfectNG[i][0];
    }
  }
  
}
```
