## Supplementary material for "BioFabric Visualization of Network Alignments": VISNAB Source Code: ShadowsAndGroupsDialog.java

```
/*
**    Copyright (C) 2003-2019 Institute for Systems Biology 
**                            Seattle, Washington, USA. 
**
**    This library is free software; you can redistribute it and/or
**    modify it under the terms of the GNU Lesser General Public
**    License as published by the Free Software Foundation; either
**    version 2.1 of the License, or (at your option) any later version.
**
**    This library is distributed in the hope that it will be useful,
**    but WITHOUT ANY WARRANTY; without even the implied warranty of
**    MERCHANTABILITY or FITNESS FOR A PARTICULAR PURPOSE.  See the GNU
**    Lesser General Public License for more details.
**
**    You should have received a copy of the GNU Lesser General Public
**    License along with this library; if not, write to the Free Software
**    Foundation, Inc., 59 Temple Place, Suite 330, Boston, MA  02111-1307  USA
*/

package org.systemsbiology.biofabric.plugin.core.align;

import java.awt.Dimension;
import java.awt.event.ActionEvent;
import java.awt.event.ActionListener;

import javax.swing.JCheckBox;
import javax.swing.JFrame;
import javax.swing.JLabel;

import org.systemsbiology.biofabric.api.dialog.BTStashResultsDialog;
import org.systemsbiology.biofabric.api.util.PluginResourceManager;
import org.systemsbiology.biofabric.api.util.ExceptionHandler;

/****************************************************************************
**
** Dialog box for specifying details of the AlignCycle layout
*/

public class ShadowsAndGroupsDialog extends BTStashResultsDialog {

  ////////////////////////////////////////////////////////////////////////////
  //
  // PRIVATE CONSTANTS
  //
  ////////////////////////////////////////////////////////////////////////////  
 
  private static final long serialVersionUID = 1L;
    
  ////////////////////////////////////////////////////////////////////////////
  //
  // PRIVATE INSTANCE MEMBERS
  //
  ////////////////////////////////////////////////////////////////////////////  
  
  private JLabel message_;
  private boolean userWantsShadowsOn_;
  private JCheckBox turnOnShadows_;
   
  ////////////////////////////////////////////////////////////////////////////
  //
  // PUBLIC CONSTRUCTORS
  //
  ////////////////////////////////////////////////////////////////////////////    

  /***************************************************************************
  **
  ** Constructor 
  */ 
  
  public ShadowsAndGroupsDialog(JFrame parent, PluginResourceManager rMan) {     
    super(parent, rMan.getPluginString("shadowAndGroups.title"), new Dimension(800, 200), 2);
   
   final String ngMsg = rMan.getPluginString("shadowAndGroups.nodeGroups");
   final String lgMsg = rMan.getPluginString("shadowAndGroups.linkGroups");
      
    turnOnShadows_ = new JCheckBox(rMan.getPluginString("shadowAndGroups.turnOnShadows"), true);
    turnOnShadows_.addActionListener(new ActionListener() {
      public void actionPerformed(ActionEvent ev) {
        try {
        	boolean shadSel = turnOnShadows_.isSelected();        	
          message_.setText((shadSel) ? lgMsg : ngMsg);
        	message_.invalidate();
        	ShadowsAndGroupsDialog.this.getContentPane().validate();
        } catch (Exception ex) {
          ExceptionHandler.getHandler().displayException(ex);
        }
      }
    });
    addWidgetFullRow(turnOnShadows_, false);
    
    message_ = new JLabel(lgMsg);
    addWidgetFullRow(message_, false);
    
    
    finishConstruction();
  }
    
  ////////////////////////////////////////////////////////////////////////////
  //
  // PUBLIC METHODS
  //
  ////////////////////////////////////////////////////////////////////////////  
  
  /***************************************************************************
  **
  ** Get results
  */
  
  public boolean turnShadowsOn() {
    return (userWantsShadowsOn_);
  }
  
  ////////////////////////////////////////////////////////////////////////////
  //
  // PROTECTED METHODS
  //
  ////////////////////////////////////////////////////////////////////////////

  /***************************************************************************
  **
  ** Stash our results for later interrogation.
  ** 
  */
  
  protected boolean stashForOK() { 
    userWantsShadowsOn_ = turnOnShadows_.isSelected();
    return (true);
  }
}
```
