## Supplementary material for "BioFabric Visualization of Network Alignments": BioFabric Source Code: CycleFinder.java

```
/*
**    Copyright (C) 2003-2018 Institute for Systems Biology 
**                            Seattle, Washington, USA. 
**
**    This library is free software; you can redistribute it and/or
**    modify it under the terms of the GNU Lesser General Public
**    License as published by the Free Software Foundation; either
**    version 2.1 of the License, or (at your option) any later version.
**
**    This library is distributed in the hope that it will be useful,
**    but WITHOUT ANY WARRANTY; without even the implied warranty of
**    MERCHANTABILITY or FITNESS FOR A PARTICULAR PURPOSE.  See the GNU
**    Lesser General Public License for more details.
**
**    You should have received a copy of the GNU Lesser General Public
**    License along with this library; if not, write to the Free Software
**    Foundation, Inc., 59 Temple Place, Suite 330, Boston, MA  02111-1307  USA
*/

package org.systemsbiology.biofabric.analysis;

import java.util.HashMap;
import java.util.Map;
import java.util.Set;
import java.util.HashSet;
import java.util.Iterator;

import org.systemsbiology.biofabric.api.model.NetLink;
import org.systemsbiology.biofabric.api.model.NetNode;
import org.systemsbiology.biofabric.api.worker.AsynchExitRequestException;
import org.systemsbiology.biofabric.api.worker.BTProgressMonitor;
import org.systemsbiology.biofabric.api.worker.LoopReporter;

/****************************************************************************
**
** A Class
*/

public class CycleFinder {
  
  ////////////////////////////////////////////////////////////////////////////
  //
  // PRIVATE CONSTANTS
  //
  //////////////////////////////////////////////////////////////////////////// 
   
  ////////////////////////////////////////////////////////////////////////////
  //
  // PUBLIC CONSTANTS
  //
  //////////////////////////////////////////////////////////////////////////// 
    
  ////////////////////////////////////////////////////////////////////////////
  //
  // PRIVATE VARIABLES
  //
  ////////////////////////////////////////////////////////////////////////////
  
  private Set<NetNode> nodes_;
  private Set<NetLink> links_;
  private HashMap<NetNode, Set<NetLink>> linksForNode_;  
  private Integer white_;
  private Integer grey_;
  private Integer black_;
  
  ////////////////////////////////////////////////////////////////////////////
  //
  // PUBLIC CONSTRUCTORS
  //
  ////////////////////////////////////////////////////////////////////////////

  /***************************************************************************
  **
  ** Constructor
  */

  public CycleFinder(Set<NetNode> nodes, Set<NetLink> links, 
  		               BTProgressMonitor monitor) throws AsynchExitRequestException {
    nodes_ = nodes;
    links_ = links;
    linksForNode_ = new HashMap<NetNode, Set<NetLink>>();
    
    LoopReporter lr = new LoopReporter(links.size(), 20, monitor, 0.0, 1.0, "progress.cycleFinderSetup");

    Iterator<NetLink> lit = links_.iterator();
    while (lit.hasNext()) {
      NetLink link = lit.next();
      lr.report();
      Set<NetLink> linksForSrc = linksForNode_.get(link.getSrcNode());
      if (linksForSrc == null) {
        linksForSrc = new HashSet<NetLink>();
        linksForNode_.put(link.getSrcNode(), linksForSrc);
      }
      linksForSrc.add(link);
    }
    white_ = Integer.valueOf(0);
    grey_ = Integer.valueOf(1);
    black_ = Integer.valueOf(2);
    lr.finish();
  }

  ////////////////////////////////////////////////////////////////////////////
  //
  // PUBLIC METHODS
  //
  ////////////////////////////////////////////////////////////////////////////
  
  /***************************************************************************
  ** 
  ** Answer if there is a cycle
  */

  public boolean hasACycle(BTProgressMonitor monitor) throws AsynchExitRequestException {
    
    //
    // Color vertices white:
    //
 
    LoopReporter lr0 = new LoopReporter(nodes_.size(), 20, monitor, 0.0, 1.0, "progress.cycleFinderVisitPass1");
    
    
    HashMap<NetNode, Integer> colors = new HashMap<NetNode, Integer>();
    Iterator<NetNode> vit = nodes_.iterator();
    while (vit.hasNext()) {
      NetNode node = vit.next();
      lr0.report();
      colors.put(node, white_);
    }
    lr0.finish();
    
    //
    // Visit each white vertex:
    //
    
    LoopReporter lr = new LoopReporter(nodes_.size(), 20, monitor, 0.0, 1.0, "progress.cycleFinderVisitPass2");
 
    vit = nodes_.iterator();
    while (vit.hasNext()) {
      NetNode node = vit.next();      
      Integer color = colors.get(node);
      if (color.equals(white_)) {
        if (visit(node, colors, lr)) {
          lr.finish();
          return (true);
        }
      }
    }
    
    lr.finish();
    return (false);
  }
 
  /***************************************************************************
  ** 
  ** Visit a node.  Return true if a cycle
  */

  private boolean visit(NetNode vertex, Map<NetNode, Integer> colors, LoopReporter lr) throws AsynchExitRequestException {
    colors.put(vertex, grey_);
    lr.report();
    Set<NetLink> linksForVertex = linksForNode_.get(vertex);
    if (linksForVertex != null) {
      Iterator<NetLink> lit = linksForVertex.iterator();
      while (lit.hasNext()) {
        NetLink link = lit.next();
        Integer targColor = colors.get(link.getTrgNode());
        if (targColor.equals(grey_)) {
          return (true);          
        } else if (targColor.equals(white_)) {
          if (visit(link.getTrgNode(), colors, lr)) {
            return (true);
          }
        }
      }
    }
    colors.put(vertex, black_);
    return (false);
  }
}
```
