## Supplementary material for "BioFabric Visualization of Network Alignments": BioFabric Source Code: GraphSearcher.java

```
/*
**    Copyright (C) 2003-2018 Institute for Systems Biology 
**                            Seattle, Washington, USA. 
**
**    This library is free software; you can redistribute it and/or
**    modify it under the terms of the GNU Lesser General Public
**    License as published by the Free Software Foundation; either
**    version 2.1 of the License, or (at your option) any later version.
**
**    This library is distributed in the hope that it will be useful,
**    but WITHOUT ANY WARRANTY; without even the implied warranty of
**    MERCHANTABILITY or FITNESS FOR A PARTICULAR PURPOSE.  See the GNU
**    Lesser General Public License for more details.
**
**    You should have received a copy of the GNU Lesser General Public
**    License along with this library; if not, write to the Free Software
**    Foundation, Inc., 59 Temple Place, Suite 330, Boston, MA  02111-1307  USA
*/

package org.systemsbiology.biofabric.analysis;

import java.util.HashMap;
import java.util.Map;
import java.util.HashSet;
import java.util.Set;
import java.math.BigInteger;
import java.util.ArrayList;
import java.util.Collections;
import java.util.List;
import java.util.Iterator;
import java.util.SortedSet;
import java.util.TreeMap;
import java.util.TreeSet;

import org.systemsbiology.biofabric.api.model.NetLink;
import org.systemsbiology.biofabric.api.model.NetNode;
import org.systemsbiology.biofabric.api.worker.AsynchExitRequestException;
import org.systemsbiology.biofabric.api.worker.BTProgressMonitor;
import org.systemsbiology.biofabric.model.FabricLink;
import org.systemsbiology.biofabric.util.UiUtil;

/****************************************************************************
**
** A Class
*/

public class GraphSearcher {
  
  ////////////////////////////////////////////////////////////////////////////
  //
  // PRIVATE CONSTANTS
  //
  //////////////////////////////////////////////////////////////////////////// 
   
  ////////////////////////////////////////////////////////////////////////////
  //
  // PUBLIC CONSTANTS
  //
  //////////////////////////////////////////////////////////////////////////// 
    
  ////////////////////////////////////////////////////////////////////////////
  //
  // PRIVATE VARIABLES
  //
  ////////////////////////////////////////////////////////////////////////////
  
  private HashSet<NetNode> allNodes_;
  private HashSet<NetLink> allEdges_;
  private ArrayList<NetNode> nodeOrder_;
  private ArrayList<NetLink> edgeOrder_;
  
  ////////////////////////////////////////////////////////////////////////////
  //
  // PUBLIC CONSTRUCTORS
  //
  ////////////////////////////////////////////////////////////////////////////

  /***************************************************************************
  **
  ** Constructor
  */

  public GraphSearcher(Set<NetNode> nodes, Set<NetLink> links) {
    allNodes_ = new HashSet<NetNode>(nodes);
    allEdges_ = new HashSet<NetLink>(links);
    edgeOrder_ = null;
    nodeOrder_ = null;
  }  
  
  /***************************************************************************
  **
  ** Constructor.  Used to create a depth-first order that 
  ** retains original sibling order.  If link appears multiple
  ** times, the order is based on first appearance.
  */

  public GraphSearcher(List<NetNode> nodes, List<NetLink> links) {

    allNodes_ = new HashSet<NetNode>(nodes);
    allEdges_ = new HashSet<NetLink>();
    edgeOrder_ = new ArrayList<NetLink>();
    nodeOrder_ = new ArrayList<NetNode>(nodes);

    for (NetLink link : links) {
      if (!allEdges_.contains(link)) {
        edgeOrder_.add(link);
      }
      allEdges_.add(link);
    }
  }  
  
  ////////////////////////////////////////////////////////////////////////////
  //
  // PUBLIC METHODS
  //
  ////////////////////////////////////////////////////////////////////////////

  /***************************************************************************
  ** 
  ** Map of node degree. If relCollapse, multigraph edges collapsed, and equals neighbor count.
  */

  public Map<NetNode, Integer> nodeDegree(boolean inOnly, boolean relCollapse, BTProgressMonitor monitor) throws AsynchExitRequestException {
    return ((relCollapse) ? nodeNeighborCount(inOnly, allEdges_, monitor) : nodeDegree(inOnly, allEdges_, monitor));
  }
 
  /***************************************************************************
  ** 
  ** Map of node degree. If inOnly == true, we only calculate in-degree. If false,
  ** the degree number for a node is a combined in/out degree. Both sources and
  ** targets are included.
  */

  public static Map<NetNode, Integer> nodeDegree(boolean inOnly, Set<NetLink> edges, 
                                                      BTProgressMonitor monitor) throws AsynchExitRequestException {
    
    HashMap<NodeAndRel, Integer> retval0 = new HashMap<NodeAndRel, Integer>();

    for (NetLink link : edges) {
      NetNode src = link.getSrcNode();
      NetNode trg = link.getTrgNode();
      String relation = link.getRelation();
      NodeAndRel sar = new NodeAndRel(src, relation);
      if (!inOnly) {
        Integer deg = retval0.get(sar);
        if (deg == null) {
          retval0.put(sar, Integer.valueOf(1));
        } else {
          retval0.put(sar, Integer.valueOf(deg.intValue() + 1));
        }
      }
      NodeAndRel tar = new NodeAndRel(trg, relation);
      Integer deg = retval0.get(trg);
      if (deg == null) {
        retval0.put(tar, Integer.valueOf(1));
      } else {
        retval0.put(tar, Integer.valueOf(deg.intValue() + 1));
      }
    }
    
    HashMap<NetNode, Integer> retval = new HashMap<NetNode, Integer>();
    for (NodeAndRel nar : retval0.keySet()) {
      Integer count = retval0.get(nar);
      Integer nco = retval.get(nar.getNode());
      nco = (nco == null) ? Integer.valueOf(count) : Integer.valueOf(count + nco);		
      retval.put(nar.getNode(), nco);
    }  		

    return (retval);
  }
  
  /***************************************************************************
  ** 
  ** Map of node neighbor count. If inOnly == true, we only calculate in-degree. If false,
  ** the degree number for a node is a combined in/out degree. Both sources and
  ** targets are included. Note this differs from degree with multigraphs, where there can be
  ** multiple links between two nodes, but each one adds to degree
  */

  public static Map<NetNode, Integer> nodeNeighborCount(boolean inOnly, Set<NetLink> edges, 
                                                             BTProgressMonitor monitor) throws AsynchExitRequestException {
    
    HashMap<NetNode, Integer> retval = new HashMap<NetNode, Integer>();

    for (NetLink link : edges) {
      NetNode src = link.getSrcNode();
      NetNode trg = link.getTrgNode();
      if (!inOnly) {
        Integer deg = retval.get(src);
        if (deg == null) {
          retval.put(src, Integer.valueOf(1));
        } else {
          retval.put(src, Integer.valueOf(deg.intValue() + 1));
        }
      }
      Integer deg = retval.get(trg);
      if (deg == null) {
        retval.put(trg, Integer.valueOf(1));
      } else {
        retval.put(trg, Integer.valueOf(deg.intValue() + 1));
      }
    }
    return (retval);
  }
  
  /***************************************************************************
  ** 
  ** Sorted set of node degree
  */

  private SortedSet<NodeDegree> nodeDegreeSet(boolean relCollapse, BTProgressMonitor monitor) throws AsynchExitRequestException {
    return (nodeDegreeSet(allEdges_, relCollapse, monitor));
  }
  
  /***************************************************************************
  ** 
  ** Nodes annotated by full degree as NodeDegree objects are ordered by degree.
  ** When equal degree, sorted by name:
  */

  private static SortedSet<NodeDegree> nodeDegreeSet(Set<NetLink> edges, boolean relCollapse, 
                                                     BTProgressMonitor monitor) throws AsynchExitRequestException {
    TreeSet<NodeDegree> retval = new TreeSet<NodeDegree>();   
    // Map of node degree. Since the first argument is false,
    // the provided degree number is the combined in/out degree.
    Map<NetNode, Integer> nds = (relCollapse) ? nodeNeighborCount(false, edges, monitor)
    	                                             : nodeDegree(false, edges, monitor);
    // Adding to sorted set orders by degree:
    for (NetNode nar : nds.keySet()) {
      NodeDegree ndeg = new NodeDegree(nar, nds.get(nar).intValue());
      retval.add(ndeg);
    }
    return (retval);
  }
  
  /***************************************************************************
  ** 
  ** Sorts the provided set of nodes by node degree, then name.
  */

  public static List<NetNode> nodesByDegree(Set<NetNode> nodes, SortedSet<NodeDegree> nds) {
    
    ArrayList<NetNode> retval = new ArrayList<NetNode>();
    
    Iterator<NodeDegree> li = nds.iterator();
    while (li.hasNext()) {
      NodeDegree nd = li.next();
      NetNode node = nd.getNodeID();
      if (nodes.contains(node)) {
        retval.add(node);
      }
    }
    return (retval);
  }
  
  /***************************************************************************
  ** 
  ** Sorted set of node degree
  */

  public SortedSet<SourcedNodeDegree> nodeDegreeSetWithSource(List<NetNode> sourceOrder) {
 
    HashMap<NetNode, Set<NetNode>> allSrcs = new HashMap<NetNode, Set<NetNode>>();

    for (NetLink nextLink : allEdges_) {
      NetNode trg = nextLink.getTrgNode();
      Set<NetNode> trgSources = allSrcs.get(trg);
      if (trgSources == null) {
        trgSources = new HashSet<NetNode>();
        allSrcs.put(trg, trgSources);
      }
      trgSources.add(nextLink.getSrcNode());
    } 
    
    TreeSet<SourcedNodeDegree> retval = new TreeSet<SourcedNodeDegree>();

    for (NetNode node : allSrcs.keySet()) {
      Set<NetNode> trgSources = allSrcs.get(node);
      SourcedNodeDegree ndeg = new SourcedNodeDegree(node, sourceOrder, trgSources);
      retval.add(ndeg);
    }
    return (retval);
  }
  
  /***************************************************************************
  ** 
  ** Sorted set of node degree, taking into account multigraph relations
  */

  public SortedSet<SourcedNodeAndRelDegree> nodeDegreeSetWithSourceMultigraph(List<NetNode> sourceOrder) {
 
    HashMap<NetNode, Set<NodeAndRel>> allSrcs = new HashMap<NetNode, Set<NodeAndRel>>();

    for (NetLink nextLink : allEdges_) {
      NetNode trg = nextLink.getTrgNode();
      String rel = nextLink.getAugRelation().relation;
      Set<NodeAndRel> trgSrcAndRels = allSrcs.get(trg);
      if (trgSrcAndRels == null) {
        trgSrcAndRels = new HashSet<NodeAndRel>();
        allSrcs.put(trg, trgSrcAndRels);
      }
      trgSrcAndRels.add(new NodeAndRel(nextLink.getSrcNode(), rel));
    } 
    
    TreeSet<SourcedNodeAndRelDegree> retval = new TreeSet<SourcedNodeAndRelDegree>();

    for (NetNode node : allSrcs.keySet()) {
      Set<NodeAndRel> trgSources = allSrcs.get(node);
      SourcedNodeAndRelDegree ndeg = new SourcedNodeAndRelDegree(node, sourceOrder, trgSources);
      retval.add(ndeg);
    }
    return (retval);
  }

  /***************************************************************************
  ** 
  ** Sorted set of nodes via gray code
  */

  public SortedSet<SourcedNodeGray> nodeGraySetWithSource(List<NetNode> sourceOrder) {
    HashMap<NetNode, Set<NetNode>> allSrcs = new HashMap<NetNode, Set<NetNode>>();
    
    for (NetLink nextLink : allEdges_) {
      NetNode trg = nextLink.getTrgNode();
      Set<NetNode> trgSources = allSrcs.get(trg);
      if (trgSources == null) {
        trgSources = new HashSet<NetNode>();
        allSrcs.put(trg, trgSources);
      }
      trgSources.add(nextLink.getSrcNode());
    } 
    
    TreeSet<SourcedNodeGray> retval = new TreeSet<SourcedNodeGray>();

    for (NetNode node : allSrcs.keySet()) {
      Set<NetNode> trgSources = allSrcs.get(node);
      SourcedNodeGray ndeg = new SourcedNodeGray(node, sourceOrder, trgSources);
      retval.add(ndeg);
    }
    return (retval);
  }
  
  /***************************************************************************
  ** 
  ** Sorted set of nodes via gray code
  */

  public static SortedSet<SourcedNodeGray> nodeGraySetWithSourceFromMap(List<NetNode> sourceOrder, 
                                                                        Map<NetNode, Set<NetNode>> srcsPerTarg) {
     
    TreeSet<SourcedNodeGray> retval = new TreeSet<SourcedNodeGray>();
    for (NetNode node : srcsPerTarg.keySet()) {
      SourcedNodeGray ndeg = new SourcedNodeGray(node, sourceOrder, srcsPerTarg.get(node));
      retval.add(ndeg);
    }
    return (retval);
  }
    
  /***************************************************************************
  ** 
  ** Sorted list of nodes, ordered by degree. 
  */

  public List<NetNode> nodeDegreeOrder(boolean relCollapse, BTProgressMonitor monitor) throws AsynchExitRequestException {
    ArrayList<NetNode> retval = new ArrayList<NetNode>();
    SortedSet<NodeDegree> nds = nodeDegreeSet(relCollapse, monitor);
    for (NodeDegree ndeg : nds) {
      retval.add(ndeg.getNodeID());
    }
    return (retval);
  }
   
  /***************************************************************************
  ** 
  ** Topo sort
  */

  public Map<NetNode, Integer> topoSort(boolean compress) {
    
    if (edgeOrder_ != null) {
      throw new IllegalStateException();
    }
    
    //
    // Assign all roots to level 0, add them to return list at that level.
    // Delete them and edges from them.  Recalulate root list and continue.
    //
    
    HashMap<NetNode, Integer> retval = new HashMap<NetNode, Integer>();
    Set<NetNode> currentNodes = new HashSet<NetNode>(allNodes_);
    //
    // Deep copy:
    //
    Set<NetLink> currentEdges = new HashSet<NetLink>();
    Iterator<NetLink> li = allEdges_.iterator();
    while (li.hasNext()) {
      NetLink link = li.next();
      currentEdges.add(((FabricLink)link).clone());
    }
      
    Map<NetNode, Set<NetLink>> outEdges = calcOutboundEdges(currentEdges);
    Set<NetNode> rootNodes = buildRootList(currentNodes, currentEdges);
  
    int level = 0;
    while (!rootNodes.isEmpty()) {
      Integer ilevel = new Integer(level++);
      Iterator<NetNode> rit = rootNodes.iterator();
      while (rit.hasNext()) {
        NetNode nodeID = rit.next();
        retval.put(nodeID, ilevel);
        outEdges.remove(nodeID);
        currentNodes.remove(nodeID);
      }
      currentEdges = invertOutboundEdges(outEdges);
      rootNodes = buildRootList(currentNodes, currentEdges);
    }
    
    if (compress) {
      contractTopoSort(retval);
    }
    return (retval);
  }
 
  /***************************************************************************
  ** 
  ** Depth-First Search
  */

  public List<QueueEntry> depthSearch() {
    //
    // Do until roots are exhausted
    //
    HashSet<NetNode> visited = new HashSet<NetNode>();
    
    Set<NetNode> rootNodes = buildRootList(allNodes_, allEdges_);
    Map<NetNode, Set<NetLink>> outEdges = calcOutboundEdges(allEdges_); 

    List<QueueEntry> retval = new ArrayList<QueueEntry>();
    if (edgeOrder_ != null) {
      HashSet<NetNode> seenRoots = new HashSet<NetNode>();
      for (NetNode currNode : nodeOrder_) {
        if (!rootNodes.contains(currNode)) {
          continue;
        }
        boolean gottaLink = false;
        for (NetLink link : edgeOrder_) {
          NetNode src = link.getSrcNode();
          if (!currNode.equals(src)) {
            continue;
          }
          if (seenRoots.contains(src)) {
            continue;
          }
          seenRoots.add(src);
          gottaLink = true;
          searchGutsDepth(src, visited, outEdges, 0, edgeOrder_, retval);
        }
        if (!gottaLink) {
          visited.add(currNode);
          retval.add(new QueueEntry(0, currNode));
        }
      }
    } else {
      for (NetNode currNode : rootNodes) {
        searchGutsDepth(currNode, visited, outEdges, 0, null, retval);
      }
    }
    return (retval);
  }
 
  /***************************************************************************
  ** 
  ** Breadth-First Search, ordered by degree
  */

  public List<QueueEntry> breadthSearch(List<NetNode> startNodes, boolean relCollapse, BTProgressMonitor monitor) throws AsynchExitRequestException {
    
    if (edgeOrder_ != null) {
      throw new IllegalStateException();
    }
    
    List<NetNode> byDeg = nodeDegreeOrder(relCollapse, monitor);
    Collections.reverse(byDeg);
    ArrayList<NetNode> toProcess = new ArrayList<NetNode>(byDeg);
    
    if ((startNodes != null) && !startNodes.isEmpty()) {
    	toProcess.removeAll(startNodes);
    	ArrayList<NetNode> useDeg = new ArrayList<NetNode>(startNodes);	
    	useDeg.addAll(toProcess);
    	toProcess = useDeg;
    }
 
    //
    // Do until everybody is visited
    //
    
    HashSet<NetNode> visited = new HashSet<NetNode>();
    ArrayList<QueueEntry> queue = new ArrayList<QueueEntry>();
    List<QueueEntry> retval = new ArrayList<QueueEntry>();
    Map<NetNode, Set<NetLink>> outEdges = calcOutboundEdges(allEdges_);    
       
    while (!toProcess.isEmpty()) {  
	    queue.add(new QueueEntry(0, toProcess.get(0)));
	    searchGutsBreadth(visited, queue, outEdges, retval, null, byDeg);
	    toProcess.removeAll(visited);
    }
    return (retval);
  }
  
  /***************************************************************************
  ** 
  ** Breadth-First Search
  */

  public List<QueueEntry> breadthSearch(boolean byDegree, boolean relCollapse, List<NetNode> useRoots, 
                                        BTProgressMonitor monitor) throws AsynchExitRequestException {
    
    if (edgeOrder_ != null) {
      throw new IllegalStateException();
    }
    
    List<NetNode> byDeg = null;
    if (byDegree) {
      byDeg = nodeDegreeOrder(relCollapse, monitor);
      Collections.reverse(byDeg);
    }   
    
    //
    // Do until roots are exhausted
    //
    HashSet<NetNode> visited = new HashSet<NetNode>();
    ArrayList<QueueEntry> queue = new ArrayList<QueueEntry>();
    List<QueueEntry> retval = new ArrayList<QueueEntry>();
    Map<NetNode, Set<NetLink>> outEdges = calcOutboundEdges(allEdges_);    
   
    
    List<NetNode> rootNodes;
    if (useRoots == null) {
      rootNodes = new ArrayList<NetNode>(buildRootList(allNodes_, allEdges_));
    } else {
      rootNodes = useRoots;
    }
    
    
    Iterator<NetNode> rit = rootNodes.iterator();
    while (rit.hasNext()) {
      queue.add(new QueueEntry(0, rit.next()));
    }
  
    searchGutsBreadth(visited, queue, outEdges, retval, null, byDeg);
    return (retval);
  }
  
  /***************************************************************************
  ** 
  ** Breadth-First Search
  */

  public List<QueueEntry> breadthSearchUntilStopped(Set<NetNode> startNodes, CriteriaJudge judge) {
    
    if (edgeOrder_ != null) {
      throw new IllegalStateException();
    }
    
    //
    // Do until roots are exhausted
    //
    
    HashSet<NetNode> visited = new HashSet<NetNode>();
    ArrayList<QueueEntry> queue = new ArrayList<QueueEntry>();
    List<QueueEntry> retval = new ArrayList<QueueEntry>();
    
    Map<NetNode, Set<NetLink>> outEdges = calcOutboundEdges(allEdges_);    
    
    Iterator<NetNode> rit = startNodes.iterator();
    while (rit.hasNext()) {
      queue.add(new QueueEntry(0, rit.next()));
    }
    searchGutsBreadth(visited, queue, outEdges, retval, judge, null);
    return (retval);
  }
   
  /***************************************************************************
  ** 
  ** A topo sort map gives column for each node.  Invert that to give a list
  ** of nodes for each column.
  */
  
  public int invertTopoSort(Map<NetNode, Integer> topoSort, Map<Integer, List<NetNode>> invert) {
        
    Iterator<NetNode> kit = topoSort.keySet().iterator();
    int maxLevel = -1;
    while (kit.hasNext()) {
      NetNode key = kit.next();
      Integer level = topoSort.get(key);
      int currLev = level.intValue();
      if (currLev > maxLevel) {
        maxLevel = currLev;
      }
      List<NetNode> nodeList = invert.get(level);
      if (nodeList == null) {
        nodeList = new ArrayList<NetNode>();
        invert.put(level, nodeList);
      }
      nodeList.add(key);
    }
    return (maxLevel);
  }
  
  /***************************************************************************
  ** 
  ** Take a sort to a simple listing
  */
  
  public List<NetNode> topoSortToPartialOrdering(Map<NetNode, Integer> topoSort) {
    
    ArrayList<NetNode> retval = new ArrayList<NetNode>();
    TreeMap<Integer, List<NetNode>> invert = new TreeMap<Integer, List<NetNode>>();
      
    invertTopoSort(topoSort, invert);    
    Iterator<List<NetNode>> kit = invert.values().iterator();
    while (kit.hasNext()) {
      List<NetNode> listForLevel = kit.next();
      // Want reproducibility between VfA and Vfg layouts.  This is required.
      Collections.sort(listForLevel);
      retval.addAll(listForLevel);
    }
    return (retval);
  }  
  
  /***************************************************************************
  ** 
  ** Take a sort to a simple listing
  */
  
  public List<NetNode> topoSortToPartialOrdering(Map<NetNode, Integer> topoSort, Set<NetLink> allLinks,
                                                      boolean relCollapse, BTProgressMonitor monitor) throws AsynchExitRequestException {
    
    ArrayList<NetNode> retval = new ArrayList<NetNode>();
    TreeMap<Integer, List<NetNode>> invert = new TreeMap<Integer, List<NetNode>>();
    
    SortedSet<NodeDegree> nds = nodeDegreeSet(allLinks, relCollapse, monitor);
    
    invertTopoSort(topoSort, invert);    
    Iterator<List<NetNode>> kit = invert.values().iterator();
    while (kit.hasNext()) {
      List<NetNode> listForLevel = kit.next();
      List<NetNode> l4lbyDeg = nodesByDegree(new HashSet<NetNode>(listForLevel), nds);
      Collections.reverse(l4lbyDeg);
      retval.addAll(l4lbyDeg);
    }
    return (retval);
  }  
  
  /***************************************************************************
  ** 
  ** Prune edge list to only those from given source nodes
  */

  public List<NetLink> onlyLinksFromSources(List<NetLink> linkList, Set<NetNode> nodes) {    
    
    ArrayList<NetLink> retval = new ArrayList<NetLink>();
    int numLinks = linkList.size();
    for (int j = 0; j < numLinks; j++) {
      NetLink link = linkList.get(j);
      if (nodes.contains(link.getSrcNode())) {
        retval.add(link);
      }
    }  
    return (retval);
  } 
  
  ////////////////////////////////////////////////////////////////////////////
  //
  // PUBLIC CLASS METHODS
  //
  //////////////////////////////////////////////////////////////////////////// 

  /***************************************************************************
  ** 
  ** With a given topo sort, force the given moveID to one end
  */

  public static Map<NetNode, Integer> topoSortReposition(Map<NetNode, Integer> origSort, 
                                                              NetNode moveID, boolean moveMin) {
    
    Integer origCol = origSort.get(moveID);
    if (origCol == null) {
      return (new HashMap<NetNode, Integer>(origSort));
    }
    
    int colVal = origCol.intValue();
    boolean colDup = false;
    int minVal = Integer.MAX_VALUE;
    int maxVal = Integer.MIN_VALUE;
    Iterator<NetNode> oskit = origSort.keySet().iterator();
    while (oskit.hasNext()) {
      NetNode key = oskit.next();
      if (key.equals(moveID)) {
        continue;
      }
      Integer checkCol = origSort.get(key);
      int chekVal = checkCol.intValue();
      if (chekVal < minVal) {
        minVal = chekVal;
      }
      if (chekVal > maxVal) {
        maxVal = chekVal;
      }
      if (chekVal == colVal) {
        colDup = true;
      }
    }
    
    int minMove;
    int maxMove;
    int moveCol;
    int inc;
    if (moveMin) {      
      if (colDup) { // to front, everybody up
        moveCol = minVal; 
        minMove = minVal;
        maxMove = maxVal;
        inc = 1;
      } else { // to front, guys below move up
        moveCol = (colVal < minVal) ? colVal : minVal; 
        minMove = minVal;
        maxMove = colVal - 1;
        inc = 1;
      }
    } else {
      if (colDup) { // to end, nobody moves
        moveCol = maxVal + 1;
        minMove = minVal - 1;
        maxMove = minVal - 1;
        inc = 0;       
      } else { // to end, guys above move down
        moveCol = (colVal > maxVal) ? colVal : maxVal;
        minMove = minVal - 1;
        maxMove = minVal - 1;
        inc = 0;      
      } 
    }
      
    HashMap<NetNode, Integer> retval = new HashMap<NetNode, Integer>();
  
    oskit = origSort.keySet().iterator();
    while (oskit.hasNext()) {
      NetNode key = oskit.next();
      if (key.equals(moveID)) {
        retval.put(moveID, Integer.valueOf(moveCol));
      } else {   
        Integer checkCol = origSort.get(key);
        int chekVal = checkCol.intValue();
        if ((chekVal >= minMove) && (chekVal <= maxMove)) {
          retval.put(key, Integer.valueOf(chekVal + inc));
        } else {
          retval.put(key, checkCol);
        }
      }
    }
    return (retval);
  }
  
  /***************************************************************************
  ** 
  ** Take a sort to a simple listing
  */
  
  public List<NetNode> topoSortToPartialOrderingByDegree(Map<NetNode, Integer> topoSort, Map<NetNode, Integer> degree) {
    
    ArrayList<NetNode> retval = new ArrayList<NetNode>();
    TreeMap<Integer, List<NetNode>> invert = new TreeMap<Integer, List<NetNode>>();
    
    invertTopoSort(topoSort, invert);    
    
    boolean first = true;
    Iterator<List<NetNode>> kit = invert.values().iterator();
    while (kit.hasNext()) {
      List<NetNode> listForLevel = kit.next();
      if (first) {
        first = false;
        SortedSet<NodeDegree> ssnd = new TreeSet<NodeDegree>(Collections.reverseOrder());
        for (int i = 0; i < listForLevel.size(); i++) {
          NetNode node = listForLevel.get(i);
          ssnd.add(new NodeDegree(node, degree.get(node).intValue()));
        } 
        for (NodeDegree nd : ssnd) {
          retval.add(nd.getNodeID());
        }
      } else {
        HashSet<NetNode> snSortSet = new HashSet<NetNode>(retval);
        List<NetLink> justFromSrc = onlyLinksFromSources(new ArrayList<NetLink>(allEdges_), new HashSet<NetNode>(retval));
        ArrayList<NetNode> working = new ArrayList<NetNode>(retval);
        working.addAll(listForLevel);
        GraphSearcher gs = new GraphSearcher(working, justFromSrc); 
        SortedSet<GraphSearcher.SourcedNodeDegree> snds = gs.nodeDegreeSetWithSource(retval);
        Iterator<GraphSearcher.SourcedNodeDegree> dfit = snds.iterator();
        while (dfit.hasNext()) {
          GraphSearcher.SourcedNodeDegree node = dfit.next();
          if (!snSortSet.contains(node.getNode())) {
            retval.add(node.getNode());
          }
        } 
      }
    }
    return (retval);
  } 
  
  /***************************************************************************
  ** 
  ** Take a sort to a simple listing
  */
  
  public List<NetNode> topoSortToPartialOrderingByGray(Map<NetNode, Integer> topoSort, Map<NetNode, Integer> degree) {
    
    ArrayList<NetNode> retval = new ArrayList<NetNode>();
    TreeMap<Integer, List<NetNode>> invert = new TreeMap<Integer, List<NetNode>>();
    
    invertTopoSort(topoSort, invert);    
    
    boolean first = true;
    Iterator<List<NetNode>> kit = invert.values().iterator();
    while (kit.hasNext()) {
      List<NetNode> listForLevel = kit.next();
      if (first) {
        first = false;
        SortedSet<NodeDegree> ssnd = new TreeSet<NodeDegree>(Collections.reverseOrder());
        for (int i = 0; i < listForLevel.size(); i++) {
          NetNode node = listForLevel.get(i);
          ssnd.add(new NodeDegree(node, degree.get(node).intValue()));
        } 
        for (NodeDegree nd : ssnd) {
          retval.add(nd.getNodeID());
        }
      } else {
        HashSet<NetNode> snSortSet = new HashSet<NetNode>(retval);
        List<NetLink> justFromSrc = onlyLinksFromSources(new ArrayList<NetLink>(allEdges_), new HashSet<NetNode>(retval));
        ArrayList<NetNode> working = new ArrayList<NetNode>(retval);
        working.addAll(listForLevel);
        GraphSearcher gs = new GraphSearcher(working, justFromSrc); 
        SortedSet<GraphSearcher.SourcedNodeGray> snds = gs.nodeGraySetWithSource(retval);
        Iterator<GraphSearcher.SourcedNodeGray> dfit = snds.iterator();
        while (dfit.hasNext()) {
          GraphSearcher.SourcedNodeGray node = dfit.next();
          if (!snSortSet.contains(node.getNodeID())) {
            retval.add(node.getNodeID());
          }
        } 
      }
    }
    return (retval);
  } 
  
  ////////////////////////////////////////////////////////////////////////////
  //
  // PUBLIC INNER CLASSES
  //
  ////////////////////////////////////////////////////////////////////////////

  public class QueueEntry {
    public int depth;
    public NetNode name;
 
    QueueEntry(int depth, NetNode name) {
      this.depth = depth;
      this.name = name;
    }
    
    @Override
    public String toString() {
      return (name + " depth = " + depth);
    }
    
  }
  
  /***************************************************************************
  **
  ** Tell us when to stop looking
  */  
      
  public static interface CriteriaJudge {
    public boolean stopHere(NetNode nodeID);  
  } 
 
  /****************************************************************************
  **
  ** Node annotated by degree. The comparator orders by degree. For equal
  ** degree, uses lexicographic node name order. 
  */
  
  public static class NodeDegree implements Comparable<NodeDegree> {
    
    private NetNode node_;
    private int degree_;
  
    public NodeDegree(NetNode node, int degree) {
      node_ = node;
      degree_ = degree;
    }
    
    public NetNode getNodeID() {
      return (node_);
    }
    
    @Override
    public int hashCode() {
      return (node_.hashCode() + degree_);
    }
  
    @Override
    public String toString() {
      return (" node = " + node_ + " degree = " + degree_);
    }
    
    @Override
    public boolean equals(Object other) {
      if (!(other instanceof NodeDegree)) {
        return (false);
      }
      NodeDegree otherDeg = (NodeDegree)other;    
      if (this.degree_ != otherDeg.degree_) {
        return (false);
      }
      return ((this.node_ == null) ? (otherDeg.node_ == null) : this.node_.equals(otherDeg.node_));
    }
    
    public int compareTo(NodeDegree otherDeg) {
      if (this.degree_ != otherDeg.degree_) {
        return (this.degree_ - otherDeg.degree_);
      }
      if (this.node_ == null) {
        return ((otherDeg.node_ == null) ? 0 : -1);
      }
      return (this.node_.compareTo(otherDeg.node_));
    } 
  }
  
  /****************************************************************************
  **
  ** Node annotated by degree and by sources. Ordering such that higher degree
  ** nodes some first, and equal degree nodes are ordered by order of source
  ** nodes.
  */
  
  public static class SourcedNodeDegree implements Comparable<SourcedNodeDegree> {
    
    private NetNode node_;
    private int degree_;
    private boolean[] srcs_;
    
    public SourcedNodeDegree(NetNode node, List<NetNode> srcOrder, Set<NetNode> mySrcs) {
      node_ = node;
      degree_ = mySrcs.size();
      int numSrc = srcOrder.size();
      srcs_ = new boolean[numSrc];
      for (int i = 0; i < numSrc; i++) {
        if (mySrcs.contains(srcOrder.get(i))) {
          srcs_[i] = true;
        }
      }
    }
    
    public NetNode getNode() {
      return (node_);
    }
    
    @Override
    public int hashCode() {
      return (node_.hashCode() + degree_);
    }
  
    @Override
    public String toString() {
      return (" node = " + node_ + " degree = " + degree_);
    }
    
    @Override
    public boolean equals(Object other) {
      if (!(other instanceof SourcedNodeDegree)) {
        return (false);
      }
      SourcedNodeDegree otherDeg = (SourcedNodeDegree)other;    
      if (this.degree_ != otherDeg.degree_) {
        return (false);
      }
      return ((this.node_ == null) ? (otherDeg.node_ == null) : this.node_.equals(otherDeg.node_));
    }
    
    public int compareTo(SourcedNodeDegree otherDeg) {
      
      if (this.node_.equals(otherDeg.node_)) {
        return (0);
      }    
      
      if (this.degree_ != otherDeg.degree_) {
        // Higher degree is first:
        return (otherDeg.degree_ - this.degree_);
      }
      
      boolean iAmBigger = false;
      boolean heIsBigger = false;
      for (int i = 0; i < this.srcs_.length; i++) {

        if (this.srcs_[i]) {
          if (otherDeg.srcs_[i] == false) {
            iAmBigger = true;
            heIsBigger = false;
            break;
          }
        }
        if (otherDeg.srcs_[i]) {
          if (this.srcs_[i] == false) {
            iAmBigger = false;
            heIsBigger = true;
            break;
          } 
        }
      }
      if (iAmBigger) {
        return (-1);
      } else if (heIsBigger) {
        return (1);
      }
      if (this.node_ == null) {
        return ((otherDeg.node_ == null) ? 0 : -1);
      }
      return (this.node_.compareTo(otherDeg.node_));
    } 
  }
  
  /****************************************************************************
  **
  ** Node annotated by degree and by (source, relation) tuples. Ordering such that higher degree
  ** nodes some first, and equal degree nodes are ordered by order of source
  ** nodes. Note this version will reflect actual degree in a multigraph, with multiple edges
  ** between a source and a target.
  */
  
  public static class SourcedNodeAndRelDegree implements Comparable<SourcedNodeAndRelDegree> {
    
    private NetNode node_;
    private int degree_;
    private boolean[] srcs_;
    
    public SourcedNodeAndRelDegree(NetNode node, List<NetNode> srcOrder, Set<NodeAndRel> mySARs) {
      node_ = node;
      degree_ = mySARs.size();
      int numSrc = srcOrder.size();
      srcs_ = new boolean[numSrc];
      Set<NetNode> snSet = new HashSet<NetNode>();    
      for (NodeAndRel sar : mySARs) {
      	snSet.add(sar.getNode());
      }
      for (int i = 0; i < numSrc; i++) {
        if (snSet.contains(srcOrder.get(i))) {
          srcs_[i] = true;
        }
      }
    }
    
    public NetNode getNode() {
      return (node_);
    }
    
    @Override
    public int hashCode() {
      return (node_.hashCode() + degree_);
    }
  
    @Override
    public String toString() {
      return (" node = " + node_ + " degree = " + degree_);
    }
    
    @Override
    public boolean equals(Object other) {
      if (!(other instanceof SourcedNodeAndRelDegree)) {
        return (false);
      }
      SourcedNodeAndRelDegree otherDeg = (SourcedNodeAndRelDegree)other;    
      if (this.degree_ != otherDeg.degree_) {
        return (false);
      }
      return ((this.node_ == null) ? (otherDeg.node_ == null) : this.node_.equals(otherDeg.node_));
    }
    
    public int compareTo(SourcedNodeAndRelDegree otherDeg) {
      
      if (this.node_.equals(otherDeg.node_)) {
        return (0);
      }    
      
      if (this.degree_ != otherDeg.degree_) {
        // Higher degree is first:
        return (otherDeg.degree_ - this.degree_);
      }
      
      boolean iAmBigger = false;
      boolean heIsBigger = false;
      for (int i = 0; i < this.srcs_.length; i++) {

        if (this.srcs_[i]) {
          if (otherDeg.srcs_[i] == false) {
            iAmBigger = true;
            heIsBigger = false;
            break;
          }
        }
        if (otherDeg.srcs_[i]) {
          if (this.srcs_[i] == false) {
            iAmBigger = false;
            heIsBigger = true;
            break;
          } 
        }
      }
      if (iAmBigger) {
        return (-1);
      } else if (heIsBigger) {
        return (1);
      }
      if (this.node_ == null) {
        return ((otherDeg.node_ == null) ? 0 : -1);
      }
      return (this.node_.compareTo(otherDeg.node_));
    } 
  }
   
  /****************************************************************************
  **
  ** A Class
  */
  
  public static class SourcedNodeGray implements Comparable<SourcedNodeGray> {
    
    private NetNode node_;
    private int degree_;
    private char[] srcs_;
    private String binNum_;
    private BigInteger bi_;
    private BigInteger gray_;
    private List<NetNode> srcOrder_;
    private Set<NetNode> mySrcs_;
    
    public SourcedNodeGray(NetNode node, List<NetNode> srcOrder, Set<NetNode> mySrcs) {
      node_ = node;
      degree_ = mySrcs.size();
      mySrcs_ = mySrcs;
      int numSrc = srcOrder.size();
      srcOrder_ = new ArrayList<NetNode>(srcOrder);
      srcs_ = new char[numSrc];
      for (int i = 0; i < numSrc; i++) {
        srcs_[i] = (mySrcs.contains(srcOrder.get(i))) ? '1' : '0';
      }
      binNum_ = String.copyValueOf(srcs_);
   
      gray_ = new BigInteger(binNum_, 2);
      bi_ = gray_;
      BigInteger working = gray_;
      while (true) {
        working = working.shiftRight(1); // sign extension, but our numbers are always positive.
        if (working.equals(BigInteger.ZERO)) {
          break;
        }
        bi_ = bi_.xor(working);
      }
    }
     
    public Set<NetNode> getSrcs() {
      return (mySrcs_);
    }
    
    public NetNode getNodeID() {
      return (node_);
    }

    public String getBinNum() {
      return (binNum_);
    }    

    @Override
    public int hashCode() {
      return (node_.hashCode() + degree_);
    }
  
    @Override
    public String toString() {
      return (" node = " + node_ + " degree = " + degree_ + " gray_ = " + gray_.toString(2) + " bi = " + bi_.toString(2));
    }
    
    public String toStringAsSets() {
      StringBuffer ret = new StringBuffer();
      ret.append(node_.getName());
      ret.append(": ");
          
      boolean first = true;
      for (int i = 0; i < srcs_.length; i++) {
        if (srcs_[i] == '1') {
          if (!first) {
            ret.append(" + ");
          } else {
            first = false;
          }
          ret.append(srcOrder_.get(i));
        }
      }   
      return (ret.toString());
    }
    
    @Override
    public boolean equals(Object other) {
      if (!(other instanceof SourcedNodeGray)) {
        return (false);
      }
      SourcedNodeGray otherDeg = (SourcedNodeGray)other;    
      if (this.degree_ != otherDeg.degree_) {
        return (false);
      }
      return ((this.node_ == null) ? (otherDeg.node_ == null) : this.node_.equals(otherDeg.node_));
    }
    
    public int compareTo(SourcedNodeGray otherDeg) {
      int nameDiff = (this.node_.compareTo(otherDeg.node_));
      if (nameDiff == 0) {
        return (0);
      }
    
      int grayComp = -this.bi_.compareTo(otherDeg.bi_);
      if (grayComp != 0) {
        return (grayComp);
      }
  
      return (nameDiff);
    } 
  }
  
  /****************************************************************************
  **
  ** A Class
  */
  
  public static class SourcedNode implements Comparable<SourcedNode> {
    
    private NetNode node_;
    private int myIn_;
    private ArrayList<Integer> mySourceOrderList_;
   
    public SourcedNode(NetNode node, Map<NetNode, Integer> inDegs, 
                       Map<NetNode, Integer> nameToRow, Map<NetNode, Set<NetNode>> l2s) {
      node_ = node;
      TreeSet<Integer> myOrder = new TreeSet<Integer>(); 
      Set<NetNode> mySet = l2s.get(node_);
      for (NetNode nextNode : mySet) {
        Integer nextSourceRow = nameToRow.get(nextNode);
        myOrder.add(nextSourceRow);       
      }
      mySourceOrderList_ = new ArrayList<Integer>(myOrder);
      myIn_ = inDegs.get(node_).intValue();
    }
    
    public NetNode getNode() {
      return (node_);
    }
    
    @Override
    public int hashCode() {
      return (node_.hashCode());
    }
  
    @Override
    public String toString() {
      return (" node = " + node_);
    }
    
    @Override
    public boolean equals(Object other) {
      if (!(other instanceof SourcedNode)) {
        return (false);
      }
      SourcedNode otherDeg = (SourcedNode)other;    
      return ((this.node_ == null) ? (otherDeg.node_ == null) : this.node_.equals(otherDeg.node_));
    }
    
    public int compareTo(SourcedNode otherDeg) {
      
      //
      // Same name, same node:
      //
      
      if (this.node_.equals(otherDeg.node_)) {
        return (0);
      }
          
      int mySize = this.mySourceOrderList_.size();
      int hisSize = otherDeg.mySourceOrderList_.size();
      int min = Math.min(mySize, hisSize);
      for (int i = 0; i < min; i++) {
        int myVal = this.mySourceOrderList_.get(i).intValue();
        int hisVal = otherDeg.mySourceOrderList_.get(i).intValue();
        int diff = hisVal - myVal;
        if (diff != 0) {
          return (diff);
        }
      }
      
      int diffSize = hisSize - mySize;
      if (diffSize != 0) {
        return (diffSize);
      }
      
      int diffIn = this.myIn_ - otherDeg.myIn_;
      if (diffIn != 0) {
        return (diffIn);
      }
      
      if (this.node_ == null) {
        return ((otherDeg.node_ == null) ? 0 : -1);
      }
      return (this.node_.compareTo(otherDeg.node_));
    } 
  }
  
  /****************************************************************************
  **
  ** A class that allows us to sort nodes based on input order
  */

  public static class MMDSourcedNode implements Comparable<MMDSourcedNode> {

    private NetNode node_;
    private int myIn_;
    private ArrayList<Integer> mySourceOrderList_;
   
    public MMDSourcedNode(NetNode node, Map<NetNode, Integer> inDegs, 
                          List<NetNode> placeList, Map<NetNode, Set<NetNode>> l2s) {
      node_ = node;
      Set<NetNode> mySet = l2s.get(node_);
      TreeSet<Integer> mySourceOrder = new TreeSet<Integer>();
      int numNode = placeList.size();
      for (int i = 0; i < numNode; i++) {
        NetNode nodex = placeList.get(i);
        if (mySet.contains(nodex)) {
          mySourceOrder.add(Integer.valueOf(i));
        }
      }
      mySourceOrderList_ = new ArrayList<Integer>();
      myIn_ = inDegs.get(this.node_).intValue();
    }

    public NetNode getNode() {
      return (node_);
    }

    @Override
    public int hashCode() {
      return (node_.hashCode());
    }

    @Override
    public String toString() {
      return (" node = " + node_);
    }

    @Override
    public boolean equals(Object other) {
      if (!(other instanceof MMDSourcedNode)) {
        return (false);
      }
      MMDSourcedNode otherDeg = (MMDSourcedNode)other;
      return ((this.node_ == null) ? (otherDeg.node_ == null) : this.node_.equals(otherDeg.node_));
    }

    public int compareTo(MMDSourcedNode otherDeg) {

      //
      // Same name, same node:
      //

      if (this.node_.equals(otherDeg.node_)) {
        return (0);
      }

      int mySize = this.mySourceOrderList_.size();
      int hisSize = otherDeg.mySourceOrderList_.size();
      int min = Math.min(mySize, hisSize);
      for (int i = 0; i < min; i++) {
        int myVal = this.mySourceOrderList_.get(i).intValue();
        int hisVal = otherDeg.mySourceOrderList_.get(i).intValue();
        int diff = hisVal - myVal;
        if (diff != 0) {
          return (diff);
        }
      }

      int diffSize = hisSize - mySize;
      if (diffSize != 0) {
        return (diffSize);
      }

      int myIn = this.myIn_;
      int hisIn = otherDeg.myIn_;
      int diffIn = myIn - hisIn;
      if (diffIn != 0) {
        return (diffIn);
      }

      if (this.node_ == null) {
        return ((otherDeg.node_ == null) ? 0 : -1);
      }
      return (this.node_.compareTo(otherDeg.node_));
    }
  }
  
  /****************************************************************************
  **
  ** When ordering target nodes by degree, we get tripped up in a multigraph case when
  ** only counting sources. We need to count source-relation tuples with a multigraph: 
  */

  public static class NodeAndRel implements Comparable<NodeAndRel> {

    private NetNode node_;
    private String relation_;
   
    public NodeAndRel(NetNode node, String relation) {
      node_ = node;
      relation_ = relation;
    }

    public NetNode getNode() {
      return (node_);
    }
    
    public String getRelation() {
      return (relation_);
    }

    @Override
    public int hashCode() {
      return (node_.hashCode() + relation_.hashCode());
    }

    @Override
    public String toString() {
      return (" node = " + node_ + " relation = " + relation_);
    }

    @Override
    public boolean equals(Object other) {
      if (!(other instanceof NodeAndRel)) {
        return (false);
      }
      NodeAndRel otherSR = (NodeAndRel)other;
      if (!this.node_.equals(otherSR.node_)) {
      	return (false);
      }
      return (this.relation_.equals(otherSR.relation_));
    }

    public int compareTo(NodeAndRel otherSR) {

      //
      // Same name, same node:
      //

    	int cn = this.node_.compareTo(otherSR.node_);
    	if (cn != 0) {
    		return (cn);
    	}
    	
      return this.relation_.compareTo(otherSR.relation_);
    }
  }
  
  
  
  
  
  ////////////////////////////////////////////////////////////////////////////
  //
  // PRIVATE INSTANCE METHODS
  //
  ////////////////////////////////////////////////////////////////////////////
  
  /***************************************************************************
  **
  ** Build map from node to outbound edges
  */

  private Map<NetNode, Set<NetLink>> calcOutboundEdges(Set<NetLink> edges) {
    
    HashMap<NetNode, Set<NetLink>> retval = new HashMap<NetNode, Set<NetLink>>();
    Iterator<NetLink> li = edges.iterator();

    while (li.hasNext()) {
      NetLink link = li.next();
      addaLink(link, link.getSrcNode(), retval);
      if (!link.isDirected()) {
      	addaLink(link, link.getTrgNode(), retval);
      }
    }
    return (retval);
  }

  /***************************************************************************
  **
  ** Add a link to a bin
  */

  private void addaLink(NetLink link, NetNode bin, Map<NetNode, Set<NetLink>> collect) {
    Set<NetLink> forBin = collect.get(bin);
    if (forBin == null) {
      forBin = new HashSet<NetLink>();
      collect.put(bin, forBin);
    }
    forBin.add(link);
    return;
  }
  
  /***************************************************************************
  **
  ** Build a root list
  */

  private Set<NetNode> buildRootList(Set<NetNode> nodes, Set<NetLink> edges) {
  
    HashSet<NetNode> retval = new HashSet<NetNode>();
    retval.addAll(nodes);
    
    Iterator<NetLink> ei = edges.iterator();
    while (ei.hasNext()) {
      NetLink link = ei.next();
      NetNode trg = link.getTrgNode();
      retval.remove(trg);
    }
    return (retval);
  }
  
  /***************************************************************************
  **
  ** Invert
  */

  private Set<NetLink> invertOutboundEdges(Map<NetNode, Set<NetLink>> outEdges) {
    
    HashSet<NetLink> retval = new HashSet<NetLink>();
    Iterator<NetNode> ki = outEdges.keySet().iterator();

    while (ki.hasNext()) {
      NetNode src = ki.next();
      Set<NetLink> links = outEdges.get(src);
      Iterator<NetLink> sit = links.iterator();
      while (sit.hasNext()) {
        NetLink lnk = sit.next();
        if (lnk.isFeedback()) {
          retval.add(((FabricLink)lnk).clone());
        } else {
          retval.add(lnk.flipped());
        }
      }
    }
    return (retval);
  }  

  /***************************************************************************
  ** 
  ** Depth-First Search guts
  */

  private void searchGutsDepth(NetNode vertexID, HashSet<NetNode> visited, 
  		                         Map<NetNode, Set<NetLink>> edgesFromSrc,
                               int depth, List<NetLink> edgeOrder, List<QueueEntry> results) {

    if (visited.contains(vertexID)) {
      return;
    }
    visited.add(vertexID);
    results.add(new QueueEntry(depth, vertexID));
    Set<NetLink> outEdges = edgesFromSrc.get(vertexID);
    if (outEdges == null) {
      return;
    }
    
    if (edgeOrder != null) {
      Iterator<NetLink> eit = edgeOrder.iterator();
      while (eit.hasNext()) {
        NetLink link = eit.next();
        if (!vertexID.equals(link.getSrcNode())) {
          continue;
        }
        NetNode targ = link.getTrgNode();
        if (!visited.contains(targ)) {
          searchGutsDepth(targ, visited, edgesFromSrc, depth + 1, edgeOrder, results);
        }
      }
    } else {
      Iterator<NetLink> eit = outEdges.iterator();
      while (eit.hasNext()) {
        NetLink link = eit.next();
        NetNode targ = link.getTrgNode();
        if (!visited.contains(targ)) {
          searchGutsDepth(targ, visited, edgesFromSrc, depth + 1, edgeOrder, results);
        }
      }
    }
    return;
  }
  
  /***************************************************************************
  ** 
  ** Breadth-First Search guts
  */

  private void searchGutsBreadth(HashSet<NetNode> visited, ArrayList<QueueEntry> queue, Map<NetNode, Set<NetLink>> edgesFromSrc, 
                                 List<QueueEntry> results, CriteriaJudge judge, List<NetNode> byDegree) {

    while (queue.size() > 0) {
      QueueEntry curr = queue.remove(0);
      if (visited.contains(curr.name)) {
        continue;
      }
      visited.add(curr.name);
      results.add(curr);
      
      if (judge != null) {
        if (judge.stopHere(curr.name)) {
          continue;
        }     
      }
      
      Set<NetLink> outEdges = edgesFromSrc.get(curr.name);
      if (outEdges == null) {
        continue;
      }
      
      if (byDegree != null) {
      	HashSet<NetNode> fltrg = new HashSet<NetNode>();
      	for (NetLink fl : outEdges) {
      		if (fl.isDirected()) {
      		  fltrg.add(fl.getTrgNode());
      		} else {
      		  NetNode flSrc = fl.getSrcNode();
      		  fltrg.add((curr.name.equals(flSrc)) ? fl.getTrgNode() : flSrc);
      		}
      	}
        Iterator<NetNode> bdit = byDegree.iterator();
        while (bdit.hasNext()) { 
          NetNode trg = bdit.next();
          if (fltrg.contains(trg)) {
            queue.add(new QueueEntry(curr.depth + 1, trg));
          }
        }
        
      } else {
        Iterator<NetLink> oit = outEdges.iterator();
        while (oit.hasNext()) { 
          NetLink flink = oit.next();
          UiUtil.fixMePrintout("What should we be doing here???");
        }
      }
    }
    return;
  }
  
  /***************************************************************************
  ** 
  ** Contract the topo sort by moving nodes as far downstream as possible without
  ** breaking the partial ordering.
  */

  private void contractTopoSort(Map<NetNode, Integer> topoSort) {
    
    //
    // Make a list of nodes for each level.  Starting at the highest level,
    // get a node, and go through all the outbound links from that node.
    // Get the minimum level of all targets, and then move the node to
    // level min - 1.
    //
    // Iterate this process until no more changes can occur.
    //
    
    HashMap<Integer, List<NetNode>> nodesAtLevel = new HashMap<Integer, List<NetNode>>();
    int maxLevel = invertTopoSort(topoSort, nodesAtLevel);    
    
    if (maxLevel == -1) {  // nothing to do
      return;
    }
    
    Map<NetNode, Set<NetLink>> outEdges = calcOutboundEdges(allEdges_);    
    
    while (true) {
      boolean changed = false;
      for (int i = maxLevel; i >= 0; i--) {
        List<NetNode> nodeList = nodesAtLevel.get(Integer.valueOf(i));
        List<NetNode> listCopy = new ArrayList<NetNode>(nodeList);
        int numNodes = nodeList.size();
        for (int j = 0; j < numNodes; j++) {
          NetNode currNode = listCopy.get(j);
          Set<NetLink> linksForNode = outEdges.get(currNode);
          HashSet<NetNode> targsForNode = new HashSet<NetNode>();
          Iterator<NetLink> lfnit = linksForNode.iterator();
          while (lfnit.hasNext()) {
            NetLink link = lfnit.next();
            NetNode targ = link.getTrgNode();
            targsForNode.add(targ);
          }
          int min = getMinLevel(targsForNode, topoSort, i, maxLevel);
          if (min > i + 1) {
            List<NetNode> higherNodeList = nodesAtLevel.get(Integer.valueOf(min - 1));
            higherNodeList.add(currNode);
            nodeList.remove(currNode);
            topoSort.put(currNode, Integer.valueOf(min - 1));
            changed = true;
          }
        }
      }
      if (!changed) {
        return;
      }
    }
  }  
  
  /***************************************************************************
  ** 
  ** Get the minimum level of all the target nodes
  */

  private int getMinLevel(Set<NetNode> targs, Map<NetNode, Integer> topoSort, int currLevel, int maxLevel) {
    if (targs == null) {
      return (currLevel);
    }
    int min = maxLevel;    
    Iterator<NetNode> trgit = targs.iterator();
    while (trgit.hasNext()) {
      NetNode trg = trgit.next();
      Integer level = topoSort.get(trg);
      int currLev = level.intValue();
      if (min > currLev) {
        min = currLev;
      }
    }
    return (min);
  }  
  
  ////////////////////////////////////////////////////////////////////////////
  //
  // PRIVATE INNER CLASSES
  //
  //////////////////////////////////////////////////////////////////////////// 
}
```
