## Supplementary material for "BioFabric Visualization of Network Alignments": BioFabric Source Code: Link.java

```
/*
**    Copyright (C) 2003-2018 Institute for Systems Biology 
**                            Seattle, Washington, USA. 
**
**    This library is free software; you can redistribute it and/or
**    modify it under the terms of the GNU Lesser General Public
**    License as published by the Free Software Foundation; either
**    version 2.1 of the License, or (at your option) any later version.
**
**    This library is distributed in the hope that it will be useful,
**    but WITHOUT ANY WARRANTY; without even the implied warranty of
**    MERCHANTABILITY or FITNESS FOR A PARTICULAR PURPOSE.  See the GNU
**    Lesser General Public License for more details.
**
**    You should have received a copy of the GNU Lesser General Public
**    License along with this library; if not, write to the Free Software
**    Foundation, Inc., 59 Temple Place, Suite 330, Boston, MA  02111-1307  USA
*/

package org.systemsbiology.biofabric.analysis;

import java.util.Collection;
import java.util.Iterator;

/****************************************************************************
**
** A Class
*/

public class Link implements Cloneable, Comparable<Link> {
  protected String src_;
  protected String trg_;

  public Link(String src, String trg) {
    if ((src == null) || (trg == null)) {
      throw new IllegalArgumentException();
    }
    this.src_ = src;
    this.trg_ = trg;
  }

  public Link(Link other) {
    this.src_ = other.src_;
    this.trg_ = other.trg_;
  }
  
  @Override
  public Link clone() {
    try {
      return ((Link)super.clone());
    } catch (CloneNotSupportedException cnse) {
      throw new IllegalStateException();
    }
  }    
  
  public String getTrg() {
    return (trg_);
  }

  public String getSrc() {
    return (src_);
  }    

  @Override
  public int hashCode() {
    return (src_.hashCode() + trg_.hashCode());
  }

  @Override
  public String toString() {
    return ("src = " + src_ + " trg = " + trg_);
  }
  
  public String toSymbolicString() {
    return (src_ + signSymbol() + trg_);
  }
  
  public static String toSymbolicString(Collection<Link> links) {
    StringBuffer buf = new StringBuffer();
    Iterator<Link> lit = links.iterator();
    while (lit.hasNext()) {
      Link lnk = lit.next();
      buf.append(lnk.toSymbolicString());
      if (lit.hasNext()) {
        buf.append(", ");
      }
    }
    return (buf.toString());
  }
  
  public String signSymbol() {
    return ("--");
  }
  
  @Override
  public boolean equals(Object other) {
    if (other == null) {
      return (false);
    }
    if (other == this) {
      return (true);
    }
    if (!(other instanceof Link)) {
      return (false);
    }
    Link otherLink = (Link)other;
    return (this.src_.equals(otherLink.src_) && this.trg_.equals(otherLink.trg_));
  }
  
  public int compareTo(Link otherLink) {
    if (this == otherLink) {
      return (0);
    }

    if (!this.src_.equals(otherLink.src_)) {
      return (this.src_.compareToIgnoreCase(otherLink.src_));
    }
    
    return (this.trg_.compareToIgnoreCase(otherLink.trg_));
  } 
}
```
