## Supplementary material for "BioFabric Visualization of Network Alignments": BioFabric Source Code: NIDLink.java

```
/*
**    Copyright (C) 2003-2018 Institute for Systems Biology 
**                            Seattle, Washington, USA. 
**
**    This library is free software; you can redistribute it and/or
**    modify it under the terms of the GNU Lesser General Public
**    License as published by the Free Software Foundation; either
**    version 2.1 of the License, or (at your option) any later version.
**
**    This library is distributed in the hope that it will be useful,
**    but WITHOUT ANY WARRANTY; without even the implied warranty of
**    MERCHANTABILITY or FITNESS FOR A PARTICULAR PURPOSE.  See the GNU
**    Lesser General Public License for more details.
**
**    You should have received a copy of the GNU Lesser General Public
**    License along with this library; if not, write to the Free Software
**    Foundation, Inc., 59 Temple Place, Suite 330, Boston, MA  02111-1307  USA
*/

package org.systemsbiology.biofabric.analysis;

import java.util.Comparator;
import java.util.Map;

import org.systemsbiology.biofabric.api.util.NID;

/****************************************************************************
**
** A Class
*/

public class NIDLink implements Cloneable, Comparable<NIDLink> {
  protected NID.WithName src_;
  protected NID.WithName trg_;

  public NIDLink(NID.WithName src, NID.WithName trg) {
    if ((src == null) || (trg == null)) {
      throw new IllegalArgumentException();
    }
    this.src_ = src;
    this.trg_ = trg;
  }

  public NIDLink(NIDLink other) {
    this.src_ = other.src_;
    this.trg_ = other.trg_;
  }
  
  @Override
  public NIDLink clone() {
    try {
      return ((NIDLink)super.clone());
    } catch (CloneNotSupportedException cnse) {
      throw new IllegalStateException();
    }
  }    
  
  public NID.WithName getTrg() {
    return (trg_);
  }

  public NID.WithName getSrc() {
    return (src_);
  }    

  @Override
  public int hashCode() {
    return (src_.hashCode() + trg_.hashCode());
  }

  @Override
  public String toString() {
    return ("src = " + src_ + " trg = " + trg_);
  }
  
  @Override
  public boolean equals(Object other) {
    if (other == null) {
      return (false);
    }
    if (other == this) {
      return (true);
    }
    if (!(other instanceof NIDLink)) {
      return (false);
    }
    NIDLink otherLink = (NIDLink)other;
    return (this.src_.equals(otherLink.src_) && this.trg_.equals(otherLink.trg_));
  }
  
  public int compareTo(NIDLink otherLink) {
    if (this == otherLink) {
      return (0);
    }

    if (!this.src_.equals(otherLink.src_)) {
      return (this.src_.compareTo(otherLink.src_));
    }
    
    return (this.trg_.compareTo(otherLink.trg_));
  }
  
  public static class NIDLinkLocationOrder implements Comparator<NIDLink> {
  	
  	private Map<NID.WithName, Integer> nodeToRow_;
  	
  	public NIDLinkLocationOrder(Map<NID.WithName, Integer> nodeToRow) {
  		nodeToRow_ = nodeToRow;
  	}
  	
  	public int compare(NIDLink link1, NIDLink link2) {
  		NID.WithName l1s = link1.getSrc();
  		NID.WithName l1t = link1.getTrg();
  		NID.WithName l2s = link2.getSrc();
  		NID.WithName l2t = link2.getTrg();
  		Integer l1sR = nodeToRow_.get(l1s);
  	  Integer l1tR = nodeToRow_.get(l1t);  			
  		Integer l2sR = nodeToRow_.get(l2s);
  		Integer l2tR = nodeToRow_.get(l2t);

  		if (l1sR.equals(l2sR)) {
  			return (l1tR.intValue() - l2tR.intValue());  			
  		} else if (l1sR.equals(l2tR)) {
  			return (l1tR.intValue() - l2sR.intValue());			
  	  } else if (l1tR.equals(l2sR)) {
  			return (l1sR.intValue() - l2tR.intValue());
  	  } else if (l1tR.equals(l2tR)) {
  			return (l1sR.intValue() - l2sR.intValue());
  	  } else {
  	  	throw new IllegalArgumentException();
  	  }	
  	}	
  } 
}
```
