## Supplementary material for "BioFabric Visualization of Network Alignments": BioFabric Source Code: BTStashResultsDialog.java

```
/*
**    Copyright (C) 2003-2018 Institute for Systems Biology 
**                            Seattle, Washington, USA. 
**
**    This library is free software; you can redistribute it and/or
**    modify it under the terms of the GNU Lesser General Public
**    License as published by the Free Software Foundation; either
**    version 2.1 of the License, or (at your option) any later version.
**
**    This library is distributed in the hope that it will be useful,
**    but WITHOUT ANY WARRANTY; without even the implied warranty of
**    MERCHANTABILITY or FITNESS FOR A PARTICULAR PURPOSE.  See the GNU
**    Lesser General Public License for more details.
**
**    You should have received a copy of the GNU Lesser General Public
**    License along with this library; if not, write to the Free Software
**    Foundation, Inc., 59 Temple Place, Suite 330, Boston, MA  02111-1307  USA
*/

package org.systemsbiology.biofabric.api.dialog;

import java.awt.Dimension;
import java.awt.GridBagLayout;
import java.awt.GridBagConstraints;
import java.util.List;
import javax.swing.JButton;
import javax.swing.JComponent;
import javax.swing.JDialog;
import javax.swing.JFrame;
import javax.swing.JLabel;
import javax.swing.JPanel;
import javax.swing.border.EmptyBorder;

import org.systemsbiology.biofabric.ui.dialogs.utils.DialogSupport;
import org.systemsbiology.biofabric.util.ResourceManager;
import org.systemsbiology.biofabric.util.UiUtil;

/****************************************************************************
**
** Abstract base class for stash results dialogs
*/

public abstract class BTStashResultsDialog extends JDialog implements DialogObj.DialogSupportClient {

  ////////////////////////////////////////////////////////////////////////////
  //
  // PRIVATE INSTANCE MEMBERS
  //
  ////////////////////////////////////////////////////////////////////////////  

  private boolean haveResult_;
  protected JPanel cp_;
  protected ResourceManager rMan_;
  protected JFrame parent_;
  protected DialogSupport ds_;
  protected GridBagConstraints gbc_;
  protected int rowNum_;
  protected int columns_;
  private static final long serialVersionUID = 1L;
  
  ////////////////////////////////////////////////////////////////////////////
  //
  // PROTECTED CONSTRUCTORS
  //
  ////////////////////////////////////////////////////////////////////////////    

  /***************************************************************************
  **
  ** Constructor 
  */ 
  
  protected BTStashResultsDialog(JFrame parent, String titleResource, Dimension size, int columns) {     
    super(parent, titleResource, true);
    haveResult_ = false;
    parent_ = parent;
    rMan_ = ResourceManager.getManager();    
    setSize(size.width, size.height);
    cp_ = (JPanel)getContentPane();
    cp_.setBorder(new EmptyBorder(20, 20, 20, 20));
    cp_.setLayout(new GridBagLayout());
    gbc_ = new GridBagConstraints();
    rowNum_ = 0;
    ds_ = new DialogSupport(this, rMan_, gbc_);
    columns_ = columns;
  }

  ////////////////////////////////////////////////////////////////////////////
  //
  // PUBLIC METHODS
  //
  //////////////////////////////////////////////////////////////////////////// 

  /***************************************************************************
  **
  ** Standard apply
  ** 
  */
 
  public void applyAction() {
    throw new UnsupportedOperationException();
  }
  
  /***************************************************************************
  **
  ** Standard ok
  ** 
  */
 
  public void okAction() {
    if (stashResults(true)) {
      setVisible(false);
      dispose();
    }
    return;   
  }
   
  /***************************************************************************
  **
  ** Standard close
  ** 
  */
  
  public void closeAction() {
    if (stashResults(false)) {
      setVisible(false);
      dispose();
    }
    return;
  }  

  /***************************************************************************
  **
  ** Answer if we have a result
  ** 
  */
  
  public boolean haveResult() {
    return (haveResult_);
  }

  ////////////////////////////////////////////////////////////////////////////
  //
  // PROTECTED METHODS
  //
  ////////////////////////////////////////////////////////////////////////////   
  
  /***************************************************************************
  **
  ** Add a full row component
  */ 
  
  protected void addWidgetFullRow(JComponent comp, boolean fixHeight) {
    addWidgetFullRow(comp, fixHeight, false);
    return;
  } 
  
  /***************************************************************************
  **
  ** Add a full row component
  */ 
  
  protected void addWidgetFullRow(JComponent comp, boolean fixHeight, boolean flushLeft) {
    rowNum_ = ds_.addWidgetFullRow(cp_, comp, fixHeight, flushLeft, rowNum_, columns_);
    return;
  } 
  
  /***************************************************************************
  **
  ** Add a full row component
  */ 
  
  protected void addTallWidgetFullRow(JComponent comp, boolean fixHeight, boolean flushLeft, int height) {
    rowNum_ = ds_.addTallWidgetFullRow(cp_, comp, fixHeight, flushLeft, height, rowNum_, columns_); 
    return;
  } 

   /***************************************************************************
  **
  ** Add a full row component
  */ 
  
  protected void addWidgetFullRowWithInsets(JComponent comp, boolean fixHeight, int inst, int insl, int insb, int insr) {
    rowNum_ = ds_.addWidgetFullRowWithInsets(cp_, comp, fixHeight, inst, insl, insb, insr, rowNum_, columns_);
    return;
  } 

  /***************************************************************************
  **
  ** Add a full row component with a label
  */ 
  
  protected void addLabeledWidget(JLabel label, JComponent comp, boolean fixHeight, boolean flushLeft) {
    rowNum_ = ds_.addLabeledWidget(cp_, label, comp, fixHeight, flushLeft, rowNum_, columns_);
    return;
  }
  
  /***************************************************************************
  **
  ** Add a labeled component with a right button
  */ 
  
  public void addLabeledFileBrowse(JLabel label, JComponent comp, JButton button) {
  	
  	JPanel pan = new JPanel();
    pan.setLayout(new GridBagLayout());
    UiUtil.gbcSet(gbc_, 0, 0, 1, 1, UiUtil.HOR, 0, 0, 5, 5, 5, 5, UiUtil.E, 1.0, 0.0);
    pan.add(comp, gbc_);
    UiUtil.gbcSet(gbc_, 1, 0, 1, 1, UiUtil.NONE, 0, 0, 5, 5, 5, 5, UiUtil.CEN, 0.0, 0.0);    
    pan.add(button, gbc_);
  	
  	addLabeledWidget(label, pan, true, false);
  	return;
  }

  /***************************************************************************
  **
  ** Add a table
  */ 
  
  protected void addTable(JComponent tablePan, int rowHeight) { 
    rowNum_ = ds_.addTable(cp_, tablePan, rowHeight, rowNum_, columns_);
    return;
  }
  
  /***************************************************************************
  **
  ** Add a table
  */ 
  
  protected void addTableNoInset(JComponent tablePan, int rowHeight) {
    rowNum_ = ds_.addTableNoInset(cp_, tablePan, rowHeight, rowNum_, columns_);
    return;
  } 
    
  /***************************************************************************
  **
  ** Finish building
  */ 
  
  protected DialogObj.Buttons finishConstruction() {      
    DialogObj.Buttons but = ds_.buildAndInstallButtonBox(cp_, rowNum_, columns_, false, true);
    setLocationRelativeTo(parent_);
    return (but);
  } 
  
  /***************************************************************************
  **
  ** Finish building
  */ 
  
  protected void finishConstructionWithExtraLeftButton(JButton xtraButton) { 
    ds_.buildAndInstallButtonBoxWithExtra(cp_, rowNum_, columns_, false, xtraButton, true);    
    setLocationRelativeTo(parent_);
    return;
  }   
  
  /***************************************************************************
  **
  ** Finish building
  */ 
  
  protected void finishConstructionWithMultiExtraLeftButtons(List<JButton> xtraButtonList) { 
    ds_.buildAndInstallButtonBoxWithMultiExtra(cp_, rowNum_, columns_, false, xtraButtonList, true);    
    setLocationRelativeTo(parent_);
    return;
  }   

  /***************************************************************************
  **
  ** Stash our results for later interrogation. 
  ** 
  */
  
  protected boolean stashResults(boolean ok) {
    if (ok) {
      if (stashForOK()) {
        haveResult_ = true;
        return (true);
      } else {
        haveResult_ = false;
        return (false);
      }
    } else {
      haveResult_ = false;
      return (true);
    }
  }
  
  /***************************************************************************
  **
  ** Do the stashing 
  ** 
  */
  
  protected abstract boolean stashForOK();

}
```
