## Supplementary material for "BioFabric Visualization of Network Alignments": BioFabric Source Code: AttributeExtractor.java

```
/*
**    Copyright (C) 2003-2018 Institute for Systems Biology 
**                            Seattle, Washington, USA. 
**
**    This library is free software; you can redistribute it and/or
**    modify it under the terms of the GNU Lesser General Public
**    License as published by the Free Software Foundation; either
**    version 2.1 of the License, or (at your option) any later version.
**
**    This library is distributed in the hope that it will be useful,
**    but WITHOUT ANY WARRANTY; without even the implied warranty of
**    MERCHANTABILITY or FITNESS FOR A PARTICULAR PURPOSE.  See the GNU
**    Lesser General Public License for more details.
**
**    You should have received a copy of the GNU Lesser General Public
**    License along with this library; if not, write to the Free Software
**    Foundation, Inc., 59 Temple Place, Suite 330, Boston, MA  02111-1307  USA
*/

package org.systemsbiology.biofabric.api.io;

import java.io.IOException;
import org.xml.sax.Attributes;

/****************************************************************************
**
** Utility for extracting attributes
*/

public final class AttributeExtractor {

  ////////////////////////////////////////////////////////////////////////////
  //
  // PRIVATE INSTANCE FIELDS
  //
  ////////////////////////////////////////////////////////////////////////////
  
  ////////////////////////////////////////////////////////////////////////////
  //
  // PUBLIC CONSTRUCTORS
  //
  ////////////////////////////////////////////////////////////////////////////
  
  ////////////////////////////////////////////////////////////////////////////
  //
  // PUBLIC CLASS METHODS
  //
  ////////////////////////////////////////////////////////////////////////////

  /***************************************************************************
  **
  ** Handle the attributes for the keyword
  **
  */
  
  public static String extractAttribute(String elemName, 
                                        Attributes attrs,
                                        String elemTarget,
                                        String attribTarget,
                                        boolean required) throws IOException {
    if (!elemName.equals(elemTarget)) {
      return (null);
    }
    
    String name = null;    
    
    if (attrs != null) {
      int count = attrs.getLength();
      for (int i = 0; i < count; i++) {
        String key = attrs.getQName(i);
        if (key == null) {
          continue;
        }
        String val = attrs.getValue(i);
        if (key.equals(attribTarget)) {
          name = val;
        }
      }
    }
    
    if (required && (name == null)) {
      throw new IOException();
    }
    return (name);
  }
}
```
