## Supplementary material for "BioFabric Visualization of Network Alignments": BioFabric Source Code: BuildData.java

```
/*
**    Copyright (C) 2003-2018 Institute for Systems Biology 
**                            Seattle, Washington, USA. 
**
**    This library is free software; you can redistribute it and/or
**    modify it under the terms of the GNU Lesser General Public
**    License as published by the Free Software Foundation; either
**    version 2.1 of the License, or (at your option) any later version.
**
**    This library is distributed in the hope that it will be useful,
**    but WITHOUT ANY WARRANTY; without even the implied warranty of
**    MERCHANTABILITY or FITNESS FOR A PARTICULAR PURPOSE.  See the GNU
**    Lesser General Public License for more details.
**
**    You should have received a copy of the GNU Lesser General Public
**    License along with this library; if not, write to the Free Software
**    Foundation, Inc., 59 Temple Place, Suite 330, Boston, MA  02111-1307  USA
*/

package org.systemsbiology.biofabric.api.io;

import java.util.List;
import java.util.Map;
import java.util.Set;
import java.util.SortedMap;

import org.systemsbiology.biofabric.api.layout.EdgeLayout;
import org.systemsbiology.biofabric.api.layout.NodeLayout;
import org.systemsbiology.biofabric.api.model.AnnotationSet;
import org.systemsbiology.biofabric.api.model.NetLink;
import org.systemsbiology.biofabric.api.model.NetNode;
import org.systemsbiology.biofabric.api.model.Network;

import org.systemsbiology.biofabric.plugin.PluginBuildData;


/****************************************************************************
**
** This is data that is used to build and rebuild the network model
*/

public interface BuildData {
  
  ////////////////////////////////////////////////////////////////////////////
  //
  // PRIVATE CONSTANTS
  //
  //////////////////////////////////////////////////////////////////////////// 

  ////////////////////////////////////////////////////////////////////////////
  //
  // PUBLIC METHODS
  //
  //////////////////////////////////////////////////////////////////////////// 
                                           
  public PluginBuildData getPluginBuildData();
 
  public void setPluginBuildData(PluginBuildData plugInData);
  
  public Set<NetNode> getSingletonNodes();
  
  public Set<NetNode> getAllNodes();
 
  public Set<NetLink> getLinks();
  
  public Map<NetNode, Integer> getNodeOrder();

  public SortedMap<Integer, NetLink> getLinkOrder() ;

  public void setNodeOrder(Map<NetNode, Integer> nodeOrder);

  public void setLinkOrder(SortedMap<Integer, NetLink> linkOrder);
  
  public void setGroupOrderAndMode(List<String> groupOrder, Network.LayoutMode mode, 
                                    boolean showLinkGroupAnnotations);
  
  public void setLayoutMode(Network.LayoutMode mode);
   
  public List<String> getGroupOrder();
 
  public Network.LayoutMode getGroupOrderMode();
  
  public boolean getShowLinkGroupAnnotations();
   
  public void setNodeAnnotations(AnnotationSet annots); 
  
  public void setLinkAnnotations(Map<Boolean, AnnotationSet> annots);
  
  public AnnotationSet getNodeAnnotations();
  
  public Map<Boolean, AnnotationSet> getLinkAnnotations();

  public boolean needsLayoutForRelayout();

  public NodeLayout getNodeLayout();
  
  public EdgeLayout getEdgeLayout();
  
  public boolean getTurnOnShadows();
  
  public void setTurnOnShadows(boolean turnOn);
  

}
```
