## Supplementary material for "BioFabric Visualization of Network Alignments": BioFabric Source Code: BuildExtractor.java

```
/*
**    Copyright (C) 2003-2018 Institute for Systems Biology 
**                            Seattle, Washington, USA. 
**
**    This library is free software; you can redistribute it and/or
**    modify it under the terms of the GNU Lesser General Public
**    License as published by the Free Software Foundation; either
**    version 2.1 of the License, or (at your option) any later version.
**
**    This library is distributed in the hope that it will be useful,
**    but WITHOUT ANY WARRANTY; without even the implied warranty of
**    MERCHANTABILITY or FITNESS FOR A PARTICULAR PURPOSE.  See the GNU
**    Lesser General Public License for more details.
**
**    You should have received a copy of the GNU Lesser General Public
**    License along with this library; if not, write to the Free Software
**    Foundation, Inc., 59 Temple Place, Suite 330, Boston, MA  02111-1307  USA
*/

package org.systemsbiology.biofabric.api.io;

import java.util.Collection;
import java.util.List;
import java.util.Map;
import java.util.Set;
import java.util.SortedMap;

import org.systemsbiology.biofabric.api.model.AugRelation;
import org.systemsbiology.biofabric.api.model.NetLink;
import org.systemsbiology.biofabric.api.model.NetNode;
import org.systemsbiology.biofabric.api.worker.AsynchExitRequestException;
import org.systemsbiology.biofabric.api.worker.BTProgressMonitor;

/****************************************************************************
**
** Methods for extracting info while building networks
*/

public interface BuildExtractor {
  
  ////////////////////////////////////////////////////////////////////////////
  //
  // PUBLIC STATIC METHODS
  //
  ////////////////////////////////////////////////////////////////////////////
  
  /***************************************************************************
  **
  ** Extract nodes
  */
  
  public Set<NetNode> extractNodes(Collection<NetLink> allLinks, Set<NetNode> loneNodeIDs,
                                   BTProgressMonitor monitor) throws AsynchExitRequestException;
  
 
  /***************************************************************************
  ** 
  ** Extract relations
  */

  public void extractRelations(List<NetLink> allLinks, 
  		                         SortedMap<AugRelation, Boolean> relMap, 
  		                         BTProgressMonitor monitor) throws AsynchExitRequestException;
  
  /***************************************************************************
  ** 
  ** Helper to drop to map to single name: useful
  */

  public Map<String, NetNode> reduceNameSetToOne(Map<String, Set<NetNode>> mapsToSets);
  
  /***************************************************************************
  ** 
  ** Process a link set that has not had directionality established
  */

  public void assignDirections(List<NetLink> allLinks, 
  		                         Map<AugRelation, Boolean> relMap,
  		                         BTProgressMonitor monitor) throws AsynchExitRequestException; 
     
  /***************************************************************************
  ** 
  ** This culls a set of links to remove non-directional synonymous and
  ** duplicate links.  Note that shadow links have already been created
  ** and added to the allLinks list. 
  */

  public void preprocessLinks(List<NetLink> allLinks, Set<NetLink> retval, Set<NetLink> culled,
  		                        BTProgressMonitor monitor) throws AsynchExitRequestException;
  
  /***************************************************************************
   **
   ** Generates a Node->Neighbors and Node->Links Map
   */
  
  public void createNeighborLinkMap(Collection<NetLink> allLinks, Set<NetNode> loneNodeIDs, Map<NetNode,
          Set<NetNode>> nodeToNeighbors, Map<NetNode, Set<NetLink>> nodeToLinks, BTProgressMonitor monitor)
          throws AsynchExitRequestException;
  
}
```
