## Supplementary material for "BioFabric Visualization of Network Alignments": BioFabric Source Code: CharacterEntityMapper.java

```
/*
**    Copyright (C) 2003-2018 Institute for Systems Biology 
**                            Seattle, Washington, USA. 
**
**    This library is free software; you can redistribute it and/or
**    modify it under the terms of the GNU Lesser General Public
**    License as published by the Free Software Foundation; either
**    version 2.1 of the License, or (at your option) any later version.
**
**    This library is distributed in the hope that it will be useful,
**    but WITHOUT ANY WARRANTY; without even the implied warranty of
**    MERCHANTABILITY or FITNESS FOR A PARTICULAR PURPOSE.  See the GNU
**    Lesser General Public License for more details.
**
**    You should have received a copy of the GNU Lesser General Public
**    License along with this library; if not, write to the Free Software
**    Foundation, Inc., 59 Temple Place, Suite 330, Boston, MA  02111-1307  USA
*/

package org.systemsbiology.biofabric.api.io;

import java.util.regex.Pattern;
import java.util.regex.Matcher;

/****************************************************************************
**
** Utility for rewriting character entities
*/

public final class CharacterEntityMapper {

  ////////////////////////////////////////////////////////////////////////////
  //
  // PRIVATE INSTANCE FIELDS
  //
  ////////////////////////////////////////////////////////////////////////////
  
  ////////////////////////////////////////////////////////////////////////////
  //
  // PUBLIC CONSTRUCTORS
  //
  ////////////////////////////////////////////////////////////////////////////
  
  ////////////////////////////////////////////////////////////////////////////
  //
  // PUBLIC CLASS METHODS
  //
  ////////////////////////////////////////////////////////////////////////////
  
  /***************************************************************************
  **
  ** Maps characters to entities for XML output.
  */
  
  public static String mapEntities(String input, boolean doSpaces) {

    //
    // We are only doing common conversions.  FIX ME to be more general?
    //

    String retval = input;
    
    // Fix for BT-10-27-09:10.  Allow quotes in names.
    if (input.indexOf('"') != -1) {
      retval = retval.replaceAll("\"", "&quot;");
    }    
    if (input.indexOf('<') != -1) {
      retval = retval.replaceAll("<", "&lt;");
    }
    if (input.indexOf('>') != -1) {
      retval = retval.replaceAll(">", "&gt;");
    }    
    if (input.indexOf('&') != -1) {
      retval = retval.replaceAll("&", "&amp;");
    }
    if (doSpaces && (input.indexOf(' ') != -1)) {
      retval = retval.replaceAll(" ", "&nbsp;");
    }
    //
    // Get greek letters into escape sequences:
    //
    StringBuffer buf = new StringBuffer();
    int len = retval.length();
    for (int i = 0; i < len; i++) {
      char ch = retval.charAt(i);
      int intval = (int)ch;
      if (ch > 127) {
        buf.append("&#x");
        if (ch < 0x1000) {
          buf.append('0');
        }
        buf.append(Integer.toString(ch, 16));
        buf.append(";");
      } else {
        buf.append(ch);
      }
    }
    retval = buf.toString();
    return (retval);
  }

  /***************************************************************************
  **
  ** Maps entities to characters for XML input
  */
  
  public static String unmapEntities(String input, boolean doSpaces) {

  	if (input == null) {
  		return (input);
  	}
    //
    // We are only doing common conversions.  FIX ME to be more general?
    //
    
    String retval = input;
    // Fix for BT-10-27-09:10.  Allow quotes in names.  
    if (input.indexOf("&quot;") != -1) {
      retval = retval.replaceAll("&quot;", "\"");
    }        
    if (input.indexOf("&lt;") != -1) {
      retval = retval.replaceAll("&lt;", "<");
    }
    if (input.indexOf("&gt;") != -1) {
      retval = retval.replaceAll("&gt;", ">");
    }    
    if (input.indexOf("&amp;") != -1) {
      retval = retval.replaceAll("&amp;", "&");
    }
    if (doSpaces && (input.indexOf("&nbsp;") != -1)) {
      retval = retval.replaceAll("&nbsp;", " ");
    }
    
    //
    // This was intended to replace legacy Greek letter conversion code,
    // but it appears it is handled automatically by the XML parser code
    // anyway these days, and is not exercised:  (I think early XML parser 
    // code < 1.4.x did not do this automatically???)
    //
    
    StringBuffer buf = new StringBuffer();
    int len = retval.length();
    int i = 0;
    while (i < len) {
      String chopped = retval.substring(i);
      if (chopped.indexOf("&#x") == 0) {
        int semiIndex = chopped.indexOf(';');
        if (semiIndex > 7) {
          buf.append(chopped.charAt(0));
          i++;
        } else {
          String numStr = chopped.substring(3, semiIndex);
          try {
            int numVal = Integer.parseInt(numStr, 16);
            buf.append((char)numVal);
            i += 8;
          } catch (NumberFormatException nfe) {
            buf.append(chopped.charAt(0));
            i++;
          }
        }
      } else {
        buf.append(chopped.charAt(0));
        i++;
      }
    }
    retval = buf.toString();
    return (retval);
  }  

  /***************************************************************************
  **
  ** Maps characters to entities for XML output, keeping tags
  */
  
  public static String mapEntitiesNotTags(String input, boolean doSpaces) {
    //
    // Map them, then restore tags we want:
    //
    
    String retval = mapEntities(input, doSpaces);
    
    Pattern iopen = Pattern.compile("&lt;i&gt;");
    Pattern iclose = Pattern.compile("&lt;/i&gt;");
    Pattern supOpen = Pattern.compile("&lt;sup&gt;");
    Pattern supClose = Pattern.compile("&lt;/sup&gt;");
    Pattern subOpen = Pattern.compile("&lt;sub&gt;");
    Pattern subClose = Pattern.compile("&lt;/sub&gt;");   
    Matcher m = iopen.matcher(retval);
    retval = m.replaceAll("<i>");
    m = iclose.matcher(retval);
    retval = m.replaceAll("</i>");
    m = supOpen.matcher(retval);
    retval = m.replaceAll("<sup>");
    m = supClose.matcher(retval);
    retval = m.replaceAll("</sup>");
    m = subOpen.matcher(retval);
    retval = m.replaceAll("<sub>");
    m = subClose.matcher(retval);
    retval = m.replaceAll("</sub>");
    return (retval);
  }  

}
```
