## Supplementary material for "BioFabric Visualization of Network Alignments": BioFabric Source Code: FileLoadFlows.java

```
/*
**    Copyright (C) 2003-2018 Institute for Systems Biology 
**                            Seattle, Washington, USA. 
**
**    This library is free software; you can redistribute it and/or
**    modify it under the terms of the GNU Lesser General Public
**    License as published by the Free Software Foundation; either
**    version 2.1 of the License, or (at your option) any later version.
**
**    This library is distributed in the hope that it will be useful,
**    but WITHOUT ANY WARRANTY; without even the implied warranty of
**    MERCHANTABILITY or FITNESS FOR A PARTICULAR PURPOSE.  See the GNU
**    Lesser General Public License for more details.
**
**    You should have received a copy of the GNU Lesser General Public
**    License along with this library; if not, write to the Free Software
**    Foundation, Inc., 59 Temple Place, Suite 330, Boston, MA  02111-1307  USA
*/

package org.systemsbiology.biofabric.api.io;


import java.awt.Component;
import java.io.File;
import java.io.IOException;
import java.util.List;
import java.util.Map;
import java.util.Set;
import java.util.SortedMap;

import org.systemsbiology.biofabric.api.model.AugRelation;
import org.systemsbiology.biofabric.api.model.NetLink;
import org.systemsbiology.biofabric.api.model.NetNode;
import org.systemsbiology.biofabric.api.util.PluginResourceManager;
import org.systemsbiology.biofabric.api.util.UniqueLabeller;
import org.systemsbiology.biofabric.api.io.AttributeKey;

/****************************************************************************
**
** Handles File Loading Tasks
*/

public interface FileLoadFlows {


  ////////////////////////////////////////////////////////////////////////////
  //
  // PUBLIC CONSTANTS
  //
  //////////////////////////////////////////////////////////////////////////// 

  public static final int LINK_COUNT_FOR_BACKGROUND_WRITE = 5000;
  public static final int FILE_LENGTH_FOR_BACKGROUND_FILE_READ = 500000;
  public static final int SIZE_TO_ASK_ABOUT_SHADOWS = 100000;
  public static final int XML_SIZE_FOR_BACKGROUND_READ = 1000000;
  
  /***************************************************************************
  **
  ** For standard file checks
  */

  public static final boolean FILE_MUST_EXIST_DONT_CARE = false;
  public static final boolean FILE_MUST_EXIST           = true;
  
  public static final boolean FILE_CAN_CREATE_DONT_CARE = false;
  public static final boolean FILE_CAN_CREATE           = true;
  
  public static final boolean FILE_DONT_CHECK_OVERWRITE = false;
  public static final boolean FILE_CHECK_OVERWRITE      = true;
  
  public static final boolean FILE_MUST_BE_FILE         = false;
  public static final boolean FILE_MUST_BE_DIRECTORY    = true;  
          
  public static final boolean FILE_CAN_WRITE_DONT_CARE  = false;
  public static final boolean FILE_CAN_WRITE            = true;
  
  public static final boolean FILE_CAN_READ_DONT_CARE   = false;
  public static final boolean FILE_CAN_READ             = true;
  
  public static enum FileLoadType {GW, SIF};
      
  ////////////////////////////////////////////////////////////////////////////
  //
  // PUBLIC METHODS
  //
  ////////////////////////////////////////////////////////////////////////////  

  /***************************************************************************
  **
  ** Do network build for a plug-in that provides the needed custom BuildData
  */
  
  public void buildNetworkForPlugIn(BuildData pluginData, File holdIt, PluginResourceManager rMan);
 
  /***************************************************************************
  **
  ** Set the current file
  */ 
    
  public void setCurrentXMLFile(File file);
  
  /***************************************************************************
  **
  ** Second step for loading from SIF (and GW)
  */
    
  public boolean handleDirectionsDupsAndShadows(List<NetLink> links, Set<NetNode> loneNodeIDs, 
  		                                           boolean binMag, SortedMap<AugRelation, Boolean> relaMap,
  		                                           Set<NetLink> reducedLinks, File holdIt, 
  		                                           boolean doForceUndirected, boolean skipShadowQuestion);
   
  /***************************************************************************
   **
   ** Load from file and directly receive link set and loners set
   */
  
  public FileLoadResult loadFromASource(File file, List<NetLink> links,
                                        Set<NetNode> loneNodes, Integer magBins,
                                        UniqueLabeller idGen, boolean loadOnly, 
                                        FileLoadType type, boolean skipShadowQuestion);
  
  /***************************************************************************
   **
   ** Common load operation for gw or sif. Returns the file that caches the previous network, which is
   ** useful for restoring original network during a multi-file-loading workflow. If there was a problem,
   ** it returns null.
   */
  
  public FileLoadResult loadFromASource(File file, Map<AttributeKey, String> nameMap,
                                        Integer magBins, UniqueLabeller idGen, 
                                        FileLoadType type, boolean skipShadowQuestion);
  
  /***************************************************************************
  **
  ** Do standard file checks and warnings
  */
 
  public boolean standardFileChecks(File target, boolean mustExist, boolean canCreate,
                                    boolean checkOverwrite, boolean mustBeDirectory, 
                                    boolean canWrite, boolean canRead);
  
  /***************************************************************************
  **
  ** Get readable attribute file
  */
  
  public File getTheFile(String ext1, String ext2, String prefTag, String desc, Component currUI);
  
  
  /***************************************************************************
  **
  ** Get a directory
  */
  
  public File getTheDirectory(String prefTag);
  
  /***************************************************************************
  **
  ** Displays file reading error message
  */ 
       
  public void displayFileInputError(IOException ioex);
  
  /***************************************************************************
  **
  ** Do window title
  */ 

  public void manageWindowTitle(String fileName);
  
  /***************************************************************************
  **
  ** Routine for handling cancellation/restore operation. True if user has restored.
  */
  
  public boolean cancelAndRestore(File restoreFile);

  /***************************************************************************
  **
  ** Load the file. Map keys are strings or Links
  */
     
  public Map<AttributeKey, String> loadTheFile(File file, Map<String, Set<NetNode>> nameToIDs, boolean forNodes);
 
  /***************************************************************************
  **
  ** Do new model operations all on AWT thread!
  */ 

  public void newModelOperations(BuildData bfnbd, boolean forMain) throws IOException;  
  
  /***************************************************************************
  **
  ** Get the file load type.
  */ 

  public FileLoadType getFileLoadType(File toCheck);
  
  /***************************************************************************
  **
  ** Return value for file loads.
  */  
  	
	public interface FileLoadResult {
		public boolean getSuccess();
		public File getCacheFile();
	}
  
  
}
```
