## Supplementary material for "BioFabric Visualization of Network Alignments": BioFabric Source Code: Indenter.java

```
/*
**    Copyright (C) 2003-2004 Institute for Systems Biology 
**                            Seattle, Washington, USA. 
**
**    This library is free software; you can redistribute it and/or
**    modify it under the terms of the GNU Lesser General Public
**    License as published by the Free Software Foundation; either
**    version 2.1 of the License, or (at your option) any later version.
**
**    This library is distributed in the hope that it will be useful,
**    but WITHOUT ANY WARRANTY; without even the implied warranty of
**    MERCHANTABILITY or FITNESS FOR A PARTICULAR PURPOSE.  See the GNU
**    Lesser General Public License for more details.
**
**    You should have received a copy of the GNU Lesser General Public
**    License along with this library; if not, write to the Free Software
**    Foundation, Inc., 59 Temple Place, Suite 330, Boston, MA  02111-1307  USA
*/

package org.systemsbiology.biofabric.api.io;

import java.io.PrintWriter;

/****************************************************************************
**
** Handles details of indenting
*/

public final class Indenter {

  ////////////////////////////////////////////////////////////////////////////
  //
  // PUBLIC CONSTANTS
  //
  ////////////////////////////////////////////////////////////////////////////

  public static final int DEFAULT_INDENT = 2; 
  
  ////////////////////////////////////////////////////////////////////////////
  //
  // PRIVATE MEMBERS
  //
  ////////////////////////////////////////////////////////////////////////////

  private int indentSpaces_;
  private PrintWriter out_;
  private StringBuffer buf_;
  private int currLevel_;
  
  ////////////////////////////////////////////////////////////////////////////
  //
  // PUBLIC CONSTRUCTORS
  //
  ////////////////////////////////////////////////////////////////////////////
  
  /***************************************************************************
  **
  ** Constructor
  */

  public Indenter(PrintWriter out, int indentSpaces) {
    indentSpaces_ = indentSpaces;
    out_ = out;
    currLevel_ = 0;
  }
  
  /***************************************************************************
  **
  ** Constructor
  */

  public Indenter(StringBuffer buf, int indentSpaces) {
    indentSpaces_ = indentSpaces;
    buf_ = buf;
    currLevel_ = 0;
  }  
   
  ////////////////////////////////////////////////////////////////////////////
  //
  // PUBLIC METHODS
  //
  ////////////////////////////////////////////////////////////////////////////

  /***************************************************************************
  ** 
  ** Indent the given number of levels
  */

  public void indent() {
    int numSpaces = currLevel_ * indentSpaces_;
    for (int i = 0; i < numSpaces; i++) {
      if (out_ != null) {
        out_.print(' ');
      } else {
        buf_.append(' ');
      }
    }
    return;
  }

  /***************************************************************************
  ** 
  ** Increment one level
  */

  public Indenter up() {
    currLevel_++;
    return (this);
  }
  
  /***************************************************************************
  ** 
  ** Decrement one level
  */

  public Indenter down() {
    currLevel_--;
    return (this);
  }
  
  /***************************************************************************
  ** 
  ** Get the current level
  */

  public int getCurrLevel() {
    return (currLevel_);
  }
  
  /***************************************************************************
  ** 
  ** Set the current level
  */

  public void setCurrLevel(int currLevel) {
    currLevel_ = currLevel;
    return;
  }
  
  /***************************************************************************
  ** 
  ** Get the indent
  */

  public int getIndent() {
    return (indentSpaces_);
  }
    
  ////////////////////////////////////////////////////////////////////////////
  //
  // PRIVATE CONSTRUCTOR
  //
  ////////////////////////////////////////////////////////////////////////////

  /***************************************************************************
  ** 
  ** Do not use
  */

  private Indenter() {
  }
}
```
