## Supplementary material for "BioFabric Visualization of Network Alignments": BioFabric Source Code: AnnotColorSource.java

```
/*
**    Copyright (C) 2003-2018 Institute for Systems Biology 
**                            Seattle, Washington, USA. 
**
**    This library is free software; you can redistribute it and/or
**    modify it under the terms of the GNU Lesser General Public
**    License as published by the Free Software Foundation; either
**    version 2.1 of the License, or (at your option) any later version.
**
**    This library is distributed in the hope that it will be useful,
**    but WITHOUT ANY WARRANTY; without even the implied warranty of
**    MERCHANTABILITY or FITNESS FOR A PARTICULAR PURPOSE.  See the GNU
**    Lesser General Public License for more details.
**
**    You should have received a copy of the GNU Lesser General Public
**    License along with this library; if not, write to the Free Software
**    Foundation, Inc., 59 Temple Place, Suite 330, Boston, MA  02111-1307  USA
*/

package org.systemsbiology.biofabric.api.layout;

import java.awt.Color;
import java.util.HashMap;
import java.util.List;
import java.util.Map;

/****************************************************************************
**
** This provides annotation colors
*/

public class AnnotColorSource {
	
	public enum AnnotColor {

	  GRAY_BLUE("GrayBlue", 0, new Color(228, 236, 248, 255)),
	  ORANGE("Orange", 1, new Color(253, 222, 195, 255)),
	  YELLOW("Yellow", 2, new Color(255, 252, 203, 255)),
	  GREEN("Green", 3, new Color(227, 253, 230, 255)),
	  PURPLE("Purple", 4, new Color(227, 224, 253, 255)),
	  PINK("Pink", 5, new Color(253, 224, 235, 255)),
	  POWDER_BLUE("PowderBlue", 6, new Color(224, 243, 253, 255)),
	  PEACH("Peach", 7, new Color(254, 246, 225, 255)),
	  
	  // 90 %
	  DARK_GRAY_BLUE("DarkGrayBlue", 8, new Color(205, 212, 223, 255)),
	  DARK_ORANGE("DarkOrange", 9, new Color(228, 200, 176, 255)),
	  DARK_YELLOW("DarkYellow", 10, new Color(230, 227, 183, 255)),
	  DARK_GREEN("DarkGreen", 11, new Color(205, 228, 207, 255)),
	  DARK_PURPLE("DarkPurple", 12, new Color(205, 202, 228, 255)),
	  DARK_PINK("DarkPink", 13, new Color(228, 202, 212, 255)),
	  DARK_POWDER_BLUE("DarkPowderBlue", 14, new Color(202, 219, 228, 255)),
	  DARK_PEACH("DarkPeach", 15, new Color(229, 221, 203, 255));
	  
	  /* 80 %
	  DARK_GRAY_BLUE("DarkGrayBlue", 8, new Color(182, 189, 198, 255)),
	  DARK_ORANGE("DarkOrange", 9, new Color(202, 178, 156, 255)),
	  DARK_YELLOW("DarkYellow", 10, new Color(204, 202, 162, 255)),
	  DARK_GREEN("DarkGreen", 11, new Color(182, 202, 184, 255)),
	  DARK_PURPLE("DarkPurple", 12, new Color(182, 179, 202, 255)),
	  DARK_PINK("DarkPink", 13, new Color(202, 179, 188, 255)),
	  DARK_POWDER_BLUE("DarkPowderBlue", 14, new Color(179, 194, 202, 255)),
	  DARK_PEACH("DarkPeach", 15, new Color(203, 197, 180, 255));
	  */
	
	  private final String name_;
	  private final int cycle_;
	  private final Color col_;
	 
	  private AnnotColor(String name, int cycle, Color col) {
	    this.name_ = name; 
	    this.cycle_ = cycle;
	    this.col_ = col;
	  }
	  
	  public Color getColor() {
	    return (col_);
	  }
	  
	  public String getName() {
	    return (name_);
	  }
	  
	  public int getCycle() {
	    return (cycle_);
	  }
	      
	  public static AnnotColor getColor(String name) {
	    String collapseName = name.replaceAll(" ", "");
	    for (AnnotColor ac : AnnotColor.values()) {
	      if (ac.name_.equalsIgnoreCase(collapseName)) {
	        return (ac);
	      }
	    }
	    throw new IllegalArgumentException(); 
	  }
	}
	
	public static AnnotColor[] getColorCycle() {
    AnnotColor[] ancs = AnnotColor.values();
    AnnotColor[] annotColors = new AnnotColor[ancs.length];
    for (AnnotColor ac : ancs) {
      annotColors[ac.getCycle()] = ac;
    }
    return (ancs);
	}

	public static Color[] getGrayCycle() {
	  Color[] annotGrays = new Color[2];
    annotGrays[0] = new Color(220, 220, 220, 127);
    annotGrays[1] = new Color(245, 245, 245, 127);
    return (annotGrays);
	}

	public static Map<String, String> getColorMap(List<String> linkGroups) {
	  HashMap<String, String> colorMap = new HashMap<String, String>();
    int numLg = linkGroups.size();
    int numColor = AnnotColor.values().length;
    AnnotColor[] myCycle = getColorCycle();

    for (int i = 0; i < numLg; i++) {
      String linkGroup = linkGroups.get(i);
      AnnotColorSource.AnnotColor ac = myCycle[i % numColor];
      colorMap.put(linkGroup, ac.getName());
    }
    return (colorMap);
	}
	
  /***************************************************************************
  **
  ** Get number of annotation colors
  */
  
  public int getAnnotColorCount() {
    return (AnnotColor.values().length);
  }
  
  /***************************************************************************
  **
  ** Get number of link annotation colors
  */
  
  public int getLinkAnnotGrayCount() {
    return (2);
  }  
  
  /***************************************************************************
  **
  ** Get annotation color
 
  
  public AnnotColorSource.AnnotColor getAnnotColor(int i) {
    return (annotColors_[i]);
  }
  
  /***************************************************************************
  **
  ** Get link annotation color
  
  
  public Color getLinkAnnotGray(int i) {
    return (annotGrays_[i]);
  }	*/
}
```
