## Supplementary material for "BioFabric Visualization of Network Alignments": BioFabric Source Code: DefaultEdgeLayout.java

```
/*
**    Copyright (C) 2003-2018 Institute for Systems Biology 
**                            Seattle, Washington, USA. 
**
**    This library is free software; you can redistribute it and/or
**    modify it under the terms of the GNU Lesser General Public
**    License as published by the Free Software Foundation; either
**    version 2.1 of the License, or (at your option) any later version.
**
**    This library is distributed in the hope that it will be useful,
**    but WITHOUT ANY WARRANTY; without even the implied warranty of
**    MERCHANTABILITY or FITNESS FOR A PARTICULAR PURPOSE.  See the GNU
**    Lesser General Public License for more details.
**
**    You should have received a copy of the GNU Lesser General Public
**    License along with this library; if not, write to the Free Software
**    Foundation, Inc., 59 Temple Place, Suite 330, Boston, MA  02111-1307  USA
*/

package org.systemsbiology.biofabric.api.layout;

import java.util.ArrayList;
import java.util.Comparator;
import java.util.HashMap;
import java.util.Iterator;
import java.util.List;
import java.util.Map;
import java.util.Set;
import java.util.SortedMap;
import java.util.TreeMap;
import java.util.TreeSet;

import org.systemsbiology.biofabric.api.io.BuildData;
import org.systemsbiology.biofabric.api.model.AnnotationSet;
import org.systemsbiology.biofabric.api.model.AugRelation;
import org.systemsbiology.biofabric.api.model.NetLink;
import org.systemsbiology.biofabric.api.model.NetNode;
import org.systemsbiology.biofabric.api.model.Network;
import org.systemsbiology.biofabric.api.worker.AsynchExitRequestException;
import org.systemsbiology.biofabric.api.worker.BTProgressMonitor;
import org.systemsbiology.biofabric.api.worker.LoopReporter;
import org.systemsbiology.biofabric.plugin.PluginSupportFactory;

/****************************************************************************
**
** This is the default layout algorithm for edges. Actually usable in combination 
** with a wide variety of different node layout algorithms, not just the default
*/

public class DefaultEdgeLayout implements EdgeLayout {
  
  ////////////////////////////////////////////////////////////////////////////
  //
  // PRIVATE INSTANCE MEMBERS
  //
  ////////////////////////////////////////////////////////////////////////////

  ////////////////////////////////////////////////////////////////////////////
  //
  // PUBLIC CONSTRUCTORS
  //
  ////////////////////////////////////////////////////////////////////////////

  /***************************************************************************
  **
  ** Constructor
  */

  public DefaultEdgeLayout() {
  }

  ////////////////////////////////////////////////////////////////////////////
  //
  // PUBLIC METHODS
  //
  ////////////////////////////////////////////////////////////////////////////
  
  /***************************************************************************
  **
  ** Do necessary pre-processing steps (e.g. automatic assignment to link groups)
  */
  
  public void preProcessEdges(BuildData rbd, 
  		                        BTProgressMonitor monitor) throws AsynchExitRequestException {
  	return;
  }

  /***************************************************************************
  **
  ** Relayout the network, but can accept a subset of network links and nodes.
  */
  
  public SortedMap<Integer, NetLink> layoutEdges(Map<NetNode, Integer> nodeOrder,
  		                                           Set<NetLink> allLinks,
  		                                           List<String> linkGroups,
  		                                           Network.LayoutMode layoutMode,
  		                                           BTProgressMonitor monitor) throws AsynchExitRequestException {
    //
    // Build target->row map:
    //
    
    HashMap<NetNode, Integer> targToRow = new HashMap<NetNode, Integer>();
    Iterator<NetNode> trit = nodeOrder.keySet().iterator();
    while (trit.hasNext()) {
      NetNode target = trit.next();
      Integer rowObj = nodeOrder.get(target);
      targToRow.put(target, rowObj);
    }
    
    //
    // For link groups, we need to find which group suffix is the best match to each augmented relation.
    // Do this only once per relation, and store the results:
    //
    
    Map<String, String> augToRel = new HashMap<String, String>();  
    for (NetLink link : allLinks) {	
    	AugRelation augRel = link.getAugRelation();
    	String match = augToRel.get(augRel.relation);
    	if (match == null) {
    		for (String rel : linkGroups) {
    			if (bestSuffixMatch(augRel.relation, rel, linkGroups)) {
    				augToRel.put(augRel.relation, rel);   				
    			}
    		}	
    	}
    }
    
    //
    // This is where the action is! All the work is done by the DefaultFabricLinkLocater, which uses comparison tests
    // between any two link to set the order. Since this is a TreeSet, this operation is roughly O(e * log e).
    //
    
    DefaultFabricLinkLocater dfll = new DefaultFabricLinkLocater(targToRow, linkGroups, augToRel, layoutMode);
    TreeSet<NetLink> order = new TreeSet<NetLink>(dfll);
   
    //
    // Do this discretely to allow progress bar:
    //
    
    LoopReporter lr = new LoopReporter(allLinks.size(), 20, monitor, 0.0, 1.0, "progress.linkLayout");
    for (NetLink link : allLinks) {
    	lr.report();
      order.add(link);
    }
    
    SortedMap<Integer, NetLink> retval = new TreeMap<Integer, NetLink>();
    int count = 0;
    for (NetLink link : order) {
    	retval.put(Integer.valueOf(count++), link);
    	lr.report();
    }    
    lr.finish();

    return (retval);
  }

  /***************************************************************************
  **
  ** Relayout the whole network!
  */
  
  public void layoutEdges(BuildData rbd, 
  		                    BTProgressMonitor monitor) throws AsynchExitRequestException {
   
    SortedMap<Integer, NetLink> retval = layoutEdges(rbd.getNodeOrder(), rbd.getLinks(), 
    		                                             rbd.getGroupOrder(), rbd.getGroupOrderMode(), monitor);
    rbd.setLinkOrder(retval);
    if (rbd.getShowLinkGroupAnnotations()) {
      installLinkAnnotations(rbd, monitor);
    } else {
      rbd.setLinkAnnotations(null);
    }
    return;
  }
  
  /***************************************************************************
  **
  ** Relayout the whole network. Same as above, but the above is part of the API.
  */
  
  public SortedMap<Integer, NetLink> layoutAllEdges(BuildData rbd, BTProgressMonitor monitor) throws AsynchExitRequestException {
   
    SortedMap<Integer, NetLink> retval = layoutEdges(rbd.getNodeOrder(), rbd.getLinks(), 
    		                                             rbd.getGroupOrder(), rbd.getGroupOrderMode(), monitor);
    rbd.setLinkOrder(retval);
    if (rbd.getShowLinkGroupAnnotations()) {
      installLinkAnnotations(rbd, monitor);
    } else {
      rbd.setLinkAnnotations(null);
    }
    return (retval);
  }

  /***************************************************************************
  **
  ** Install link annotations for link groups
  */
  
  protected void installLinkAnnotations(BuildData rbd, BTProgressMonitor monitor)
    throws AsynchExitRequestException {
  
    LoopReporter lr = new LoopReporter(rbd.getLinkOrder().size(), 20, monitor, 0, 1.0, "progress.linkAnnotationPrep");  
    List<NetLink> linkList = new ArrayList<NetLink>();  
    for (NetLink link : rbd.getLinkOrder().values()) {   
      linkList.add(link);
      lr.report();
    }
    lr.finish();
    
    AnnotationSet nonShdwAnnots = calcGroupLinkAnnots(rbd, linkList, monitor, false, rbd.getGroupOrder());
    AnnotationSet withShdwAnnots = calcGroupLinkAnnots(rbd, linkList, monitor, true, rbd.getGroupOrder());
    
    Map<Boolean, AnnotationSet> linkAnnots = new HashMap<Boolean, AnnotationSet>();
    linkAnnots.put(true, withShdwAnnots);
    linkAnnots.put(false, nonShdwAnnots);
    
    rbd.setLinkAnnotations(linkAnnots);
    return;
  }
 
  /*********************************************************************************************
  **
  ** Calculate link group link annotations 
  */
    
  protected AnnotationSet calcGroupLinkAnnots(BuildData rbd, 
                                              List<NetLink> links, BTProgressMonitor monitor, 
                                              boolean shadow, List<String> linkGroups) throws AsynchExitRequestException {   
    
    String which = (shadow) ? "progress.linkAnnotationShad" : "progress.linkAnnotationNoShad";
    LoopReporter lr = new LoopReporter(links.size(), 20, monitor, 0, 1.0, which); 
    Map<String, String> colorMap = AnnotColorSource.getColorMap(linkGroups);
  
    AnnotationSet retval = PluginSupportFactory.buildAnnotationSet();
    String lastType = null;
    int startPos = 0;
    int endPos = 0;
    int numLink = links.size();
    int count = 0;
    for (int i = 0; i < numLink; i++) {
      NetLink link = links.get(i);
      lr.report();
      if (link.isShadow() && !shadow) {
        continue;
      }
      String thisType = link.getRelation();
      if (lastType == null) {
        lastType = thisType;      
        startPos = count;
      } else if (lastType.equals(thisType)) {
        // do nothing              
      } else {
        retval.addAnnot(PluginSupportFactory.buildAnnotation(lastType, startPos, endPos, 0, getColor(lastType, colorMap)));
        lastType = thisType;
        startPos = count;
      }
      endPos = count++;
    }
    if (lastType != null) {
      retval.addAnnot(PluginSupportFactory.buildAnnotation(lastType, startPos, endPos, 0, getColor(lastType, colorMap)));
    }
    lr.finish();
    return (retval);
  }

  /***************************************************************************
  **
  ** Answer if the given relation has the best suffix match with the given match,
  ** given all the options. Thus, "430" should match "30" instead of "0" if both
  ** are present.
  */

  private boolean bestSuffixMatch(String augR, String relToMatch, List<String> allRels) {
  	if (relToMatch == null) {
  		return (true);
  	}
  	int topLen = 0;
  	String topRel = null;
  	for (String aRel : allRels) {
  		int matchLen = aRel.length();
  		if (matchLen < topLen) {
  			continue;
  		}
  		int ioaRel = augR.indexOf(aRel);
  		if ((ioaRel >= 0) && ((ioaRel == (augR.length() - matchLen)))) {
        if (topLen == matchLen) {
        	throw new IllegalStateException();
  		  } else if (topLen < matchLen) {
  		    topLen = matchLen;
          topRel = aRel;	
  		  }
      }	 
  	}
  	if (topRel == null) {
  		throw new IllegalStateException();
  	}
    return (topRel.equals(relToMatch));
  }

  /***************************************************************************
  **
  ** Used to order links for default link layout
  */
   
  public static class DefaultFabricLinkLocater implements Comparator<NetLink> {
  	
  	private Map<NetNode, Integer> nodeToRow_;
  	private List<String> relOrder_;
  	private Map<String, String> augToRel_;
  	Network.LayoutMode layMode_;
  	
  	public DefaultFabricLinkLocater(Map<NetNode, Integer> nodeToRow, List<String> relOrder,
  			                            Map<String, String> augToRel, Network.LayoutMode layMode) {
  		nodeToRow_ = nodeToRow;
  		augToRel_ = augToRel;
  		relOrder_ = relOrder;
  		layMode_ = layMode;
  	}
  	
  	/***************************************************************************
	  ** 
	  ** For ANY two different links in the network, this says which comes first:
	  */
  	  	
  	public int compare(NetLink link1, NetLink link2) {
  	  
  	  if (link1.equals(link2)) {
  	    return (0);
  	  }
  	  
  		NetNode l1s = link1.getSrcNode();
  		NetNode l1t = link1.getTrgNode();
  		NetNode l2s = link2.getSrcNode();
  		NetNode l2t = link2.getTrgNode();
  		
  		Integer l1sR = nodeToRow_.get(l1s);
  	  Integer l1tR = nodeToRow_.get(l1t);  			
  		Integer l2sR = nodeToRow_.get(l2s);
  		Integer l2tR = nodeToRow_.get(l2t);
  		
  		int link1Top = Math.min(l1sR.intValue(), l1tR.intValue());
  		int link1Bot = Math.max(l1sR.intValue(), l1tR.intValue());
  		int link2Top = Math.min(l2sR.intValue(), l2tR.intValue());
  		int link2Bot = Math.max(l2sR.intValue(), l2tR.intValue());
  		
  		boolean link1Reg = !link1.isShadow();
  		boolean link2Reg = !link2.isShadow();
  	
  		//
  		// If layout mode is global, then the augmented relation order is what counts the most!
  		//
  		
  		if (layMode_ == Network.LayoutMode.PER_NETWORK_MODE) {
  			AugRelation link1Rel = link1.getAugRelation();
  			AugRelation link2Rel = link2.getAugRelation();
  			int ord1 = relOrder_.indexOf(augToRel_.get(link1Rel.relation));
  			int ord2 = relOrder_.indexOf(augToRel_.get(link2Rel.relation));
  			int ordDiff = ord1 - ord2;
  			if (ordDiff != 0) {
  				return (ordDiff);				
  			}	
  		}
  		
  		// If the two links are regular, the top node row controls the order.
  		if (link1Reg && link2Reg) { // Both regular
  			int topDiff = link1Top - link2Top;
  			if (topDiff != 0) {
  				return (topDiff);
  			}
  			// Same top node, link groups are next up, if they are defined:
  			Integer perGroup = orderUsingGroups(link1, link2);
  			if (perGroup != null) {
  				return (perGroup.intValue());
  			}
  			// Same top, no (or same) link group. So comparison depends on the bottom node row:
  			int botDiff = link1Bot - link2Bot;
  			if (botDiff != 0) {
  				return (botDiff);
  			}
  			// Same extent.
  			return (orderForNode(link1, link2, l1sR, l1tR, l2sR, l2tR));
  			
  		// If the two links are shadow, the bottom row determines the order:	
  		} else if (!link1Reg && !link2Reg) { // Both shadow
  		  int botDiff = link1Bot - link2Bot;
  			if (botDiff != 0) {
  				return (botDiff);
  			}
  		  // Same bottom node, link groups are next up, if they are defined:
  			Integer perGroup = orderUsingGroups(link1, link2);
  			if (perGroup != null) {
  				return (perGroup.intValue());
  			}
  			// Same bottom, no (or same) link group. So comparison depends on the top node row:
  			int topDiff = link1Top - link2Top;
  			if (topDiff != 0) {
  				return (topDiff);
  			}
  			// Same extent.
        return (orderForNode(link1, link2, l1sR, l1tR, l2sR, l2tR));	
  		// Link1 regular, Link2 shadow. Compare regular top to shadow bottom. If they are equal, they are associated
  	  // to the same node, where shadow ALWAYS comes first, unless we are using per-node groups
  		} else if (link1Reg && !link2Reg) { 
  			int rsDiff = link1Top - link2Bot;
  		  if (rsDiff != 0) {
  				return (rsDiff);
  			}
  		  // Same bottom node, link groups are next up, if they are defined:
  			Integer perGroup = orderUsingGroups(link1, link2);
  			if (perGroup != null) {
  				return (perGroup.intValue());
  			}  
  			return (1);
  		} else if (!link1Reg && link2Reg) { 
  			int rsDiff = link1Bot - link2Top;
  		  if (rsDiff != 0) {
  				return (rsDiff);
  			}
  		  // Same bottom node, link groups are next up, if they are defined:
  			Integer perGroup = orderUsingGroups(link1, link2);
  			if (perGroup != null) {
  				return (perGroup.intValue());
  			} 
  			return (-1);
  	  } else {
  	  	throw new IllegalArgumentException();
  	  }	
  	}
  	
  	/***************************************************************************
	  ** 
	  ** When per-node link groups are in effect, we us this to sort the links
	  ** for the node into their groups. Returns null if link groups not in effect,
	  ** or if the links are in the same group. I.e., keep on plugging!
	  */
  	
  	public Integer orderUsingGroups(NetLink link1, NetLink link2) { 	
   		AugRelation link1Rel = link1.getAugRelation();
  	  AugRelation link2Rel = link2.getAugRelation();	
  	  //
  	  // If we are in per node mode, the link group order has first precedence:
  	  //
  		if (layMode_ == Network.LayoutMode.PER_NODE_MODE) {
  			int ord1 = relOrder_.indexOf(augToRel_.get(link1Rel.relation));
  			int ord2 = relOrder_.indexOf(augToRel_.get(link2Rel.relation));
  			int ordDiff = ord1 - ord2;
  			if (ordDiff != 0) {
  				return (Integer.valueOf(ordDiff));				
  			}
  		}
  		return (null);
  	}
	
  	/***************************************************************************
	  ** 
	  ** When we get into a node where and are dealing with links of the same extent,
	  ** we resort to directionality, and finally to the lexicographic ordering of
	  ** the link relation tag.
	  */
  	
   	public int orderForNode(NetLink link1, NetLink link2, Integer l1sR, Integer l1tR, Integer l2sR, Integer l2tR) { 	
   		AugRelation link1Rel = link1.getAugRelation();
  	  AugRelation link2Rel = link2.getAugRelation();
  			
  		//
  		// Extents are the same. Undirected come first, then down links, then up links. Note this allows us
  	  // to treat directed feedbacks consistently, since undirected will always come before ambiguous direction.
  		//
  		
  		boolean link1IsDir = link1.isDirected();
  	  // If both are false, we have a feedback loop on one node:
  		Boolean link1PointsUp = (link1IsDir) ? Boolean.valueOf(l1sR.intValue() > l1tR.intValue()) : null;
  		Boolean link1PointsDown = (link1IsDir) ? Boolean.valueOf(l1sR.intValue() < l1tR.intValue()) : null;

  		boolean link2IsDir = link2.isDirected();
  	  // If both are false, we have a feedback loop on one node:
  		Boolean link2PointsUp = (link2IsDir) ? Boolean.valueOf(l2sR.intValue() > l2tR.intValue()) : null;
  		Boolean link2PointsDown = (link2IsDir) ? Boolean.valueOf(l2sR.intValue() < l2tR.intValue()) : null;

  		//
  		// Feedback. Undirected first. If 
      //
  		boolean undirFeed1 = !link1IsDir && (l1sR.intValue() == l1tR.intValue());
  		boolean undirFeed2 = !link2IsDir && (l2sR.intValue() == l2tR.intValue());
  	  boolean dirFeed1 = link1IsDir && !link1PointsUp && !link1PointsDown;
  		boolean dirFeed2 = link2IsDir && !link2PointsUp && !link2PointsDown;
  		
  		if (undirFeed1 && dirFeed2) {
  			return (-1);
  		} else if (dirFeed1 && undirFeed2) {
  			return (1);
  		} // Otherwise we leave for relation compare...
  		
  		// Note if neither is directed, we will skip to next test:
  		
  		if (link1IsDir && !link2IsDir) {
  			return (1);	
   	  } else if (!link1IsDir && link2IsDir) {
  			return (-1);	
  		} else if (link1IsDir && link2IsDir) { // both directed
  			if (link1PointsDown && link2PointsUp) {	
  			  return (-1);
  		  } else if (link1PointsUp && link2PointsDown) {
  		  	return (1);
  		  } // else both point the same way
  		} // else both undirected
  		
  		//
  		// Now the augmented relation tags drive the order:
  		//
  		
  		int relComp = link1Rel.compareTo(link2Rel);
  	  return (relComp);
   	}
  } 
 
  /***************************************
  **
  ** Get the color
  */
     
  protected String getColor(String type, Map<String, String> colorMap) {
    return (colorMap.get(type));
  }
}
```
