## Supplementary material for "BioFabric Visualization of Network Alignments": BioFabric Source Code: DefaultLayout.java

```
/*
**    Copyright (C) 2003-2018 Institute for Systems Biology 
**                            Seattle, Washington, USA. 
**
**    This library is free software; you can redistribute it and/or
**    modify it under the terms of the GNU Lesser General Public
**    License as published by the Free Software Foundation; either
**    version 2.1 of the License, or (at your option) any later version.
**
**    This library is distributed in the hope that it will be useful,
**    but WITHOUT ANY WARRANTY; without even the implied warranty of
**    MERCHANTABILITY or FITNESS FOR A PARTICULAR PURPOSE.  See the GNU
**    Lesser General Public License for more details.
**
**    You should have received a copy of the GNU Lesser General Public
**    License along with this library; if not, write to the Free Software
**    Foundation, Inc., 59 Temple Place, Suite 330, Boston, MA  02111-1307  USA
*/

package org.systemsbiology.biofabric.api.layout;

import java.util.ArrayList;
import java.util.Collections;
import java.util.HashMap;
import java.util.HashSet;
import java.util.Iterator;
import java.util.List;
import java.util.Map;
import java.util.Set;
import java.util.SortedSet;
import java.util.TreeMap;
import java.util.TreeSet;

import org.systemsbiology.biofabric.api.io.BuildData;
import org.systemsbiology.biofabric.api.model.NetLink;
import org.systemsbiology.biofabric.api.model.NetNode;
import org.systemsbiology.biofabric.api.worker.AsynchExitRequestException;
import org.systemsbiology.biofabric.api.worker.BTProgressMonitor;
import org.systemsbiology.biofabric.api.worker.LoopReporter;

/****************************************************************************
**
** This is the default layout algorithm
*/

public class DefaultLayout extends NodeLayout {
  
  ////////////////////////////////////////////////////////////////////////////
  //
  // PRIVATE INSTANCE MEMBERS
  //
  ////////////////////////////////////////////////////////////////////////////

  ////////////////////////////////////////////////////////////////////////////
  //
  // PUBLIC CONSTRUCTORS
  //
  ////////////////////////////////////////////////////////////////////////////

  /***************************************************************************
  **
  ** Constructor
  */

  public DefaultLayout() {

  }

  ////////////////////////////////////////////////////////////////////////////
  //
  // PUBLIC METHODS
  //
  ////////////////////////////////////////////////////////////////////////////

  /***************************************************************************
  **
  ** Relayout the network!
  */
  
  public List<NetNode> doNodeLayout(BuildData rbd, 
  		                              Params params,
  		                              BTProgressMonitor monitor) throws AsynchExitRequestException {
      
    List<NetNode> startNodeIDs = (params == null) ? null : ((DefaultParams)params).startNodes;
    List<NetNode> targetIDs = defaultNodeOrder(rbd.getLinks(), rbd.getSingletonNodes(), startNodeIDs, monitor);

    //
    // Now have the ordered list of targets we are going to display.
    // Build target->row maps and the inverse:
    //
    
    installNodeOrder(targetIDs, rbd, monitor);
    return (targetIDs);
  }

  /***************************************************************************
  ** 
  ** Calculate default node order. Used by several other layout classes
  */

  public List<NetNode> defaultNodeOrder(Set<NetLink> allLinks,
	                                      Set<NetNode> loneNodes, 
	                                      List<NetNode> startNodes, 
	                                      BTProgressMonitor monitor) throws AsynchExitRequestException { 
    //
    // Note the allLinks Set has pruned out duplicates and synonymous non-directional links
    //
    //
    // Build a target list, top to bottom, that adds the node with the most
    // links first, and adds those link targets ASAP. If caller supplies a start node,
    // we go there first:
    // 
    
    HashMap<NetNode, Integer> linkCounts = new HashMap<NetNode, Integer>();
    HashMap<NetNode, Set<NetNode>> targsPerSource = new HashMap<NetNode, Set<NetNode>>();
    ArrayList<NetNode> targets = new ArrayList<NetNode>();
         
    HashSet<NetNode> targsToGo = new HashSet<NetNode>();
    
    int numLink = allLinks.size();
    LoopReporter lr = new LoopReporter(numLink, 20, monitor, 0.0, 0.25, "progress.calculateNodeDegree");
    
    Iterator<NetLink> alit = allLinks.iterator();
    while (alit.hasNext()) {
      NetLink nextLink = alit.next();
      lr.report();
      NetNode sidwn = nextLink.getSrcNode();
      NetNode tidwn = nextLink.getTrgNode();
      Set<NetNode> targs = targsPerSource.get(sidwn);
      if (targs == null) {
        targs = new HashSet<NetNode>();
        targsPerSource.put(sidwn, targs);
      }
      targs.add(tidwn);
      targs = targsPerSource.get(tidwn);
      if (targs == null) {
        targs = new HashSet<NetNode>();
        targsPerSource.put(tidwn, targs);
      }
      targs.add(sidwn);
      targsToGo.add(sidwn);
      targsToGo.add(tidwn);        
      Integer srcCount = linkCounts.get(sidwn);
      linkCounts.put(sidwn, (srcCount == null) ? Integer.valueOf(1) : Integer.valueOf(srcCount.intValue() + 1));
      Integer trgCount = linkCounts.get(tidwn);
      linkCounts.put(tidwn, (trgCount == null) ? Integer.valueOf(1) : Integer.valueOf(trgCount.intValue() + 1));
    }
    lr.finish();
    
    //
    // Rank the nodes by link count:
    //
    
    lr = new LoopReporter(linkCounts.size(), 20, monitor, 0.25, 0.50, "progress.rankByDegree");
    
    TreeMap<Integer, SortedSet<NetNode>> countRank = new TreeMap<Integer, SortedSet<NetNode>>(Collections.reverseOrder());
    Iterator<NetNode> lcit = linkCounts.keySet().iterator();
    while (lcit.hasNext()) {
      NetNode src = lcit.next();
      lr.report();
      Integer count = linkCounts.get(src);
      SortedSet<NetNode> perCount = countRank.get(count);
      if (perCount == null) {
        perCount = new TreeSet<NetNode>();
        countRank.put(count, perCount);
      }
      perCount.add(src);
    }
    lr.finish();
    
    //
    // Handle the specified starting nodes case:
    //
    
    if ((startNodes != null) && !startNodes.isEmpty()) {
      ArrayList<NetNode> queue = new ArrayList<NetNode>();
      targsToGo.removeAll(startNodes);
      targets.addAll(startNodes);
      queue.addAll(startNodes);
      flushQueue(targets, targsPerSource, linkCounts, targsToGo, queue, monitor, 0.50, 0.75);
    }   
    
    //
    // Get all kids added in.  Now doing this without recursion; seeing blown
    // stacks for huge networks!
    //
    // While we still have nodes to place, find the highest degree *unplaced* node, add it to order list,
    // then handle all its children:
    //
     
    while (!targsToGo.isEmpty()) {
      Iterator<Integer> crit = countRank.keySet().iterator();
      while (crit.hasNext()) {
        Integer key = crit.next();
        SortedSet<NetNode> perCount = countRank.get(key);
        Iterator<NetNode> pcit = perCount.iterator();
        while (pcit.hasNext()) {
          NetNode node = pcit.next();
          if (targsToGo.contains(node)) {
            ArrayList<NetNode> queue = new ArrayList<NetNode>();
            targsToGo.remove(node);
            targets.add(node);
            addMyKidsNR(targets, targsPerSource, linkCounts, targsToGo, node, queue, monitor, 0.75, 1.0);
          }
        }
      }
    }
    
    //
    //
    // Tag on lone nodes.  If a node is by itself, but also shows up in the links,
    // we drop it.
    //
    // Used to do a set removeAll() operation, but discovered the operation was
    // taking FOREVER, e.g. remains 190804 targets 281832. So do this in a loop
    // that can be monitored for progress:
    //
    
    LoopReporter lr2 = new LoopReporter(loneNodes.size(), 20, monitor, 0.0, 0.25, "progress.addSingletonsToTargets");
    HashSet<NetNode> targSet = new HashSet<NetNode>(targets);
    TreeSet<NetNode> remains = new TreeSet<NetNode>();
    
    for (NetNode lnod : loneNodes) {
    	if (!targSet.contains(lnod)) {
    		lr2.report();
    		remains.add(lnod); 		
    	}    	
    }
    lr2.finish();
    targets.addAll(remains);
    return (targets);
  }
        
  /***************************************************************************
  **
  ** Ordering of neighbor nodes. 
  */
  
  private List<NetNode> orderMyKids(Map<NetNode, Set<NetNode>> targsPerSource, 
  		                                   Map<NetNode, Integer> linkCounts, 
                                         Set<NetNode> targsToGo, NetNode node) {
    Set<NetNode> targs = targsPerSource.get(node);
    if (targs == null) {
    	return (new ArrayList<NetNode>());
    }
    //
    // Get the kids ordered highest degree to lowest, with lex ordering if equal degree:
    //
    TreeMap<Integer, SortedSet<NetNode>> kidMap = new TreeMap<Integer, SortedSet<NetNode>>(Collections.reverseOrder());
    Iterator<NetNode> tait = targs.iterator();
    while (tait.hasNext()) {  
      NetNode nextTarg = tait.next(); 
      Integer count = linkCounts.get(nextTarg);
      SortedSet<NetNode> perCount = kidMap.get(count);
      if (perCount == null) {
        perCount = new TreeSet<NetNode>();
        kidMap.put(count, perCount);
      }
      perCount.add(nextTarg);
    }
    
    //
    // Go through that map and return an ordered list of neighbors *that have not yet been placed!!*
    //
    
    ArrayList<NetNode> myKidsToProc = new ArrayList<NetNode>();
    Iterator<SortedSet<NetNode>> kmit = kidMap.values().iterator();
    while (kmit.hasNext()) {  
      SortedSet<NetNode> perCount = kmit.next(); 
      Iterator<NetNode> pcit = perCount.iterator();
      while (pcit.hasNext()) {  
        NetNode kid = pcit.next();
        if (targsToGo.contains(kid)) { 
          myKidsToProc.add(kid);
        }
      }
    }
    return (myKidsToProc);
  }    
  
  /***************************************************************************
  **
  ** Handle all kids of the given node by adding it to the queue and flushing the queue:
  */
  
  private void addMyKidsNR(List<NetNode> targets, Map<NetNode, Set<NetNode>> targsPerSource, 
                           Map<NetNode, Integer> linkCounts, 
                           Set<NetNode> targsToGo, NetNode node, List<NetNode> queue,
                           BTProgressMonitor monitor, double startFrac, double endFrac) 
                          	 throws AsynchExitRequestException {
    queue.add(node);
    flushQueue(targets, targsPerSource, linkCounts, targsToGo, queue, monitor, startFrac, endFrac);
    return;
  }
  
  /***************************************************************************
  **
  ** Node ordering, non-recursive:
  */
  
  private void flushQueue(List<NetNode> targets, 
  		                    Map<NetNode, Set<NetNode>> targsPerSource, 
                          Map<NetNode, Integer> linkCounts, 
                          Set<NetNode> targsToGo, List<NetNode> queue, 
                          BTProgressMonitor monitor, double startFrac, double endFrac) 
                            throws AsynchExitRequestException {
  	
  	LoopReporter lr = new LoopReporter(targsToGo.size(), 20, monitor, startFrac, endFrac, "progress.nodeOrdering");
  	int lastSize = targsToGo.size();	
    while (!queue.isEmpty()) {
      NetNode node = queue.remove(0);
      int ttgSize = targsToGo.size();
      lr.report(lastSize - ttgSize);
      lastSize = ttgSize;
      List<NetNode> myKids = orderMyKids(targsPerSource, linkCounts, targsToGo, node);
      Iterator<NetNode> ktpit = myKids.iterator(); 
      while (ktpit.hasNext()) {  
        NetNode kid = ktpit.next();
        if (targsToGo.contains(kid)) {
          targsToGo.remove(kid);
          targets.add(kid);
          queue.add(kid);
        }
      }
    }
    lr.finish();
    return;
  }
  
  /***************************************************************************
  **
  ** For passing around layout params
  */  
  
  public static class DefaultParams implements Params {
        
    public List<NetNode> startNodes;

    public DefaultParams(List<NetNode> startNodes) {
      this.startNodes = startNodes;
    } 
  }

}
```
