## Supplementary material for "BioFabric Visualization of Network Alignments": BioFabric Source Code: NodeLayout.java

```
/*
**    Copyright (C) 2003-2018 Institute for Systems Biology 
**                            Seattle, Washington, USA. 
**
**    This library is free software; you can redistribute it and/or
**    modify it under the terms of the GNU Lesser General Public
**    License as published by the Free Software Foundation; either
**    version 2.1 of the License, or (at your option) any later version.
**
**    This library is distributed in the hope that it will be useful,
**    but WITHOUT ANY WARRANTY; without even the implied warranty of
**    MERCHANTABILITY or FITNESS FOR A PARTICULAR PURPOSE.  See the GNU
**    Lesser General Public License for more details.
**
**    You should have received a copy of the GNU Lesser General Public
**    License along with this library; if not, write to the Free Software
**    Foundation, Inc., 59 Temple Place, Suite 330, Boston, MA  02111-1307  USA
*/

package org.systemsbiology.biofabric.api.layout;

import java.util.ArrayList;
import java.util.HashMap;
import java.util.List;
import java.util.TreeMap;

import org.systemsbiology.biofabric.api.io.BuildData;
import org.systemsbiology.biofabric.api.model.NetNode;
import org.systemsbiology.biofabric.api.worker.AsynchExitRequestException;
import org.systemsbiology.biofabric.api.worker.BTProgressMonitor;
import org.systemsbiology.biofabric.api.worker.LoopReporter;
import org.systemsbiology.biofabric.io.BuildDataImpl;

/****************************************************************************
**
** This is the abstract base class for node layout algorithms
*/

public abstract class NodeLayout {

	////////////////////////////////////////////////////////////////////////////
  //
  // PUBLIC METHODS
  //
  ////////////////////////////////////////////////////////////////////////////
   
  /***************************************************************************
  **
  ** Find out if the necessary conditions for this layout are met. By default, a
  ** layout handles anything.
  */
   
  public boolean criteriaMet(BuildData rbd, BTProgressMonitor monitor) throws AsynchExitRequestException, 
                                                                              LayoutCriterionFailureException {
    return (true);
  }

  
  /***************************************************************************
  **
  ** Relayout the nodes!
  */
  
  public abstract List<NetNode> doNodeLayout(BuildData rbd, Params params,
												  		               BTProgressMonitor monitor) throws AsynchExitRequestException;
  
  
  /***************************************************************************
  **
  ** Install node orders
  */
  
  public void installNodeOrder(List<NetNode> targetIDs, BuildData rbd, 
  		                         BTProgressMonitor monitor) throws AsynchExitRequestException {
    int currRow = 0;
    LoopReporter lr = new LoopReporter(targetIDs.size(), 20, monitor, 0.0, 1.0, "progress.installOrdering");
    
    HashMap<NetNode, Integer> nodeOrder = new HashMap<NetNode, Integer>();
    for (NetNode target : targetIDs) {
      lr.report();
      Integer rowTag = Integer.valueOf(currRow++);
      nodeOrder.put(target, rowTag);
    }  
    rbd.setNodeOrder(nodeOrder);
    lr.finish();
    return;
  }
  
  /***************************************************************************
  ** 
  ** Utility conversion
  */

  protected List<NetNode> convertOrderToMap(BuildData rbd, 
  		                                      List<Integer> orderedStringRows) { 
    HashMap<NetNode, Integer> nOrd = new HashMap<NetNode, Integer>();
    TreeMap<Integer, NetNode> rev = new TreeMap<Integer, NetNode>();
    int numOsr = orderedStringRows.size();
    for (int i = 0; i < numOsr; i++) {
      Integer intval = orderedStringRows.get(i);
      NetNode nid = ((BuildDataImpl)rbd).getNodeIDForRowForNetwork(intval);
      nOrd.put(nid, Integer.valueOf(i));
      rev.put(Integer.valueOf(i), nid);      
    }
    rbd.setNodeOrder(nOrd);
    return (new ArrayList<NetNode>(rev.values()));
  }
  
  /***************************************************************************
  **
  ** For passing around layout params
  */  
  
  public interface Params {
  }
}
```
