## Supplementary material for "BioFabric Visualization of Network Alignments": BioFabric Source Code: AugRelation.java

```
/*
**    Copyright (C) 2003-2018 Institute for Systems Biology 
**                            Seattle, Washington, USA. 
**
**    This library is free software; you can redistribute it and/or
**    modify it under the terms of the GNU Lesser General Public
**    License as published by the Free Software Foundation; either
**    version 2.1 of the License, or (at your option) any later version.
**
**    This library is distributed in the hope that it will be useful,
**    but WITHOUT ANY WARRANTY; without even the implied warranty of
**    MERCHANTABILITY or FITNESS FOR A PARTICULAR PURPOSE.  See the GNU
**    Lesser General Public License for more details.
**
**    You should have received a copy of the GNU Lesser General Public
**    License along with this library; if not, write to the Free Software
**    Foundation, Inc., 59 Temple Place, Suite 330, Boston, MA  02111-1307  USA
*/

package org.systemsbiology.biofabric.api.model;


/***************************************************************************
**
** Augmented relation
*/  

public final class AugRelation implements Cloneable, Comparable<AugRelation> {
  public String relation;
  public boolean isShadow;
  
  public AugRelation(String relation, boolean isShadow) {
    this.relation = relation;
    this.isShadow = isShadow;
  }
  
  @Override
  public AugRelation clone() {
    try {
      return ((AugRelation)super.clone());
    } catch (CloneNotSupportedException cnse) {
      throw new IllegalStateException();
    }
  }   
 
  public int compareTo(AugRelation otherAug) {
    if (this.equals(otherAug)) {
      return (0);
    }

    if (this.isShadow != otherAug.isShadow) {
      return ((this.isShadow) ? -1 : 1);
    }    
    if (!this.relation.toUpperCase().equals(otherAug.relation.toUpperCase())) {
      return (this.relation.compareToIgnoreCase(otherAug.relation));
    }
    throw new IllegalStateException();
  }
    
  @Override  
  public boolean equals(Object other) {    
    if (other == null) {
      return (false);
    }
    if (other == this) {
      return (true);
    }
    if (!(other instanceof AugRelation)) {
      return (false);
    }
    AugRelation otherAug = (AugRelation)other;

    if (this.isShadow != otherAug.isShadow) {
      return (false);
    }
  
    return (this.relation.toUpperCase().equals(otherAug.relation.toUpperCase()));
  }
   
  @Override
  public int hashCode() {
    return (relation.toUpperCase().hashCode() + ((isShadow) ? 17 : 31));
  }

  @Override
  public String toString() {
    return ("rel = " + relation + " isShadow = " + isShadow);
  }    
}
```
