## Supplementary material for "BioFabric Visualization of Network Alignments": BioFabric Source Code: LinkComparator.java

```
/*
**    Copyright (C) 2003-2018 Institute for Systems Biology 
**                            Seattle, Washington, USA. 
**
**    This library is free software; you can redistribute it and/or
**    modify it under the terms of the GNU Lesser General Public
**    License as published by the Free Software Foundation; either
**    version 2.1 of the License, or (at your option) any later version.
**
**    This library is distributed in the hope that it will be useful,
**    but WITHOUT ANY WARRANTY; without even the implied warranty of
**    MERCHANTABILITY or FITNESS FOR A PARTICULAR PURPOSE.  See the GNU
**    Lesser General Public License for more details.
**
**    You should have received a copy of the GNU Lesser General Public
**    License along with this library; if not, write to the Free Software
**    Foundation, Inc., 59 Temple Place, Suite 330, Boston, MA  02111-1307  USA
*/

package org.systemsbiology.biofabric.api.model;

import java.util.Comparator;

import org.systemsbiology.biofabric.util.DataUtil;


/***************************************************************************
**
** Comparator for Fabric Links
** This is kinda bogus and old, and is just used to cleanse input. Replace
** with something newer?
*/  

public final class LinkComparator implements Comparator<NetLink> {
  
  public int compare(NetLink one, NetLink otherLink) {
    if (one.equals(otherLink)) {
      return (0);
    }
    
    String srcOne = DataUtil.normKey(one.getSrcNode().getName());
    String trgOne = DataUtil.normKey(one.getTrgNode().getName());
    String srcTwo = DataUtil.normKey(otherLink.getSrcNode().getName());
    String trgTwo = DataUtil.normKey(otherLink.getTrgNode().getName());
    
    if (!srcOne.equals(srcTwo)) {
      return (srcOne.compareTo(srcTwo));
    }    
    if (!trgOne.equals(trgTwo)) {
      return (trgOne.compareTo(trgTwo));
    } 
    if (one.isShadow() != otherLink.isShadow()) {
      return ((one.isShadow()) ? -1 : 1);
    }  
    if (!DataUtil.normKey(one.getRelation()).equals(DataUtil.normKey(otherLink.getRelation()))) {
      return (one.getRelation().compareToIgnoreCase(otherLink.getRelation()));
    }
   
    //
    // Gatta catch the far-out case where two links with same source and target node names but different
    // internal IDs get to here. Can't let it fall through
    //
    
    if (!one.getSrcNode().equals(otherLink.getSrcNode())) {
       return (one.getSrcNode().compareTo(otherLink.getSrcNode()));
    }
    if (!one.getTrgNode().equals(otherLink.getTrgNode())) {
      return (one.getTrgNode().compareTo(otherLink.getTrgNode()));
    }

    throw new IllegalStateException();
  }
  	
}
```
