## Supplementary material for "BioFabric Visualization of Network Alignments": BioFabric Source Code: NetLink.java

```
/*
**    Copyright (C) 2003-2018 Institute for Systems Biology 
**                            Seattle, Washington, USA. 
**
**    This library is free software; you can redistribute it and/or
**    modify it under the terms of the GNU Lesser General Public
**    License as published by the Free Software Foundation; either
**    version 2.1 of the License, or (at your option) any later version.
**
**    This library is distributed in the hope that it will be useful,
**    but WITHOUT ANY WARRANTY; without even the implied warranty of
**    MERCHANTABILITY or FITNESS FOR A PARTICULAR PURPOSE.  See the GNU
**    Lesser General Public License for more details.
**
**    You should have received a copy of the GNU Lesser General Public
**    License along with this library; if not, write to the Free Software
**    Foundation, Inc., 59 Temple Place, Suite 330, Boston, MA  02111-1307  USA
*/

package org.systemsbiology.biofabric.api.model;

/****************************************************************************
**
** This is the interface for network links that plugins can use
*/

public interface NetLink {

  public NetLink flipped();
  public AugRelation getAugRelation();
  public String getRelation();
  public boolean isFeedback();
  public void installDirection(Boolean isDirected);
  public boolean isDirected();
  public NetNode getTrgNode();
  public NetNode getSrcNode();
  public boolean isShadow();
  public boolean synonymous(NetLink other);
}
```
