## Supplementary material for "BioFabric Visualization of Network Alignments": BioFabric Source Code: Network.java

```
/*
**    Copyright (C) 2003-2018 Institute for Systems Biology 
**                            Seattle, Washington, USA. 
**
**    This library is free software; you can redistribute it and/or
**    modify it under the terms of the GNU Lesser General Public
**    License as published by the Free Software Foundation; either
**    version 2.1 of the License, or (at your option) any later version.
**
**    This library is distributed in the hope that it will be useful,
**    but WITHOUT ANY WARRANTY; without even the implied warranty of
**    MERCHANTABILITY or FITNESS FOR A PARTICULAR PURPOSE.  See the GNU
**    Lesser General Public License for more details.
**
**    You should have received a copy of the GNU Lesser General Public
**    License along with this library; if not, write to the Free Software
**    Foundation, Inc., 59 Temple Place, Suite 330, Boston, MA  02111-1307  USA
*/

package org.systemsbiology.biofabric.api.model;

import java.io.IOException;


/****************************************************************************
**
** This is the Network model.
*/

public interface Network {
  
  ////////////////////////////////////////////////////////////////////////////
  //
  // PRIVATE CONSTANTS
  //
  //////////////////////////////////////////////////////////////////////////// 

  ////////////////////////////////////////////////////////////////////////////
  //
  // PUBLIC CONSTANTS
  //
  //////////////////////////////////////////////////////////////////////////// 
  
  public enum LayoutMode {
    UNINITIALIZED_MODE("notSet"),
    PER_NODE_MODE("perNode"),
    PER_NETWORK_MODE("perNetwork");

    private String text;

    LayoutMode(String text) {
      this.text = text;
    }

    public String getText() {
      return (text);
    }

	  public static LayoutMode fromString(String text)  throws IOException {
	    if (text != null) {
	      for (LayoutMode lm : LayoutMode.values()) {
	        if (text.equalsIgnoreCase(lm.text)) {
	          return (lm);
	        }
	      }
	    }
	    throw new IOException();
	  }
  }
                    

  ////////////////////////////////////////////////////////////////////////////
  //
  // PUBLIC METHODS
  //
  ////////////////////////////////////////////////////////////////////////////

  public int getNodeCount();
  public int getLinkCount(boolean forShadow);
  
  
}
```
