## Supplementary material for "BioFabric Visualization of Network Alignments": BioFabric Source Code: AbstractFactoryClient.java

```
/*
**    Copyright (C) 2003-2014 Institute for Systems Biology 
**                            Seattle, Washington, USA. 
**
**    This library is free software; you can redistribute it and/or
**    modify it under the terms of the GNU Lesser General Public
**    License as published by the Free Software Foundation; either
**    version 2.1 of the License, or (at your option) any later version.
**
**    This library is distributed in the hope that it will be useful,
**    but WITHOUT ANY WARRANTY; without even the implied warranty of
**    MERCHANTABILITY or FITNESS FOR A PARTICULAR PURPOSE.  See the GNU
**    Lesser General Public License for more details.
**
**    You should have received a copy of the GNU Lesser General Public
**    License along with this library; if not, write to the Free Software
**    Foundation, Inc., 59 Temple Place, Suite 330, Boston, MA  02111-1307  USA
*/

package org.systemsbiology.biofabric.api.parser;

import java.util.Set;
import java.util.HashSet;
import java.io.IOException;
import java.util.HashMap;
import java.util.Iterator;

import org.xml.sax.Attributes;

/****************************************************************************
**
** This is used to build objects from XML
*/

public abstract class AbstractFactoryClient implements ParserClient {
  
  ////////////////////////////////////////////////////////////////////////////
  //
  // PRIVATE CONSTANTS
  //
  ////////////////////////////////////////////////////////////////////////////
 
  ////////////////////////////////////////////////////////////////////////////
  //
  // PRIVATE INSTANCE VARIABLES
  //
  ////////////////////////////////////////////////////////////////////////////
  
  protected Object container_;
 
  protected HashMap<String, AbstractFactoryClient> workerMap_;
  protected HashMap<String, GlueStick> glueStickMap_;  
  protected AbstractFactoryClient currWorker_;
  protected String currElem_;
  protected Object sharedWhiteboard_;
   
  protected HashSet<String> myKeys_;
  
  ////////////////////////////////////////////////////////////////////////////
  //
  // PUBLIC CONSTRUCTORS
  //
  ////////////////////////////////////////////////////////////////////////////

  /***************************************************************************
  **
  ** Constructor for a factory client
  */

  public AbstractFactoryClient(Object sharedWhiteboard) {
    myKeys_ = new HashSet<String>();
    workerMap_ = new HashMap<String, AbstractFactoryClient>();
    glueStickMap_ = new HashMap<String, GlueStick>();
    sharedWhiteboard_ = sharedWhiteboard;
  }

  ///////////////////////////////////////////////////////////////////////////
  //
  // PUBLIC METHODS
  //
  ////////////////////////////////////////////////////////////////////////////

  /***************************************************************************
  ** 
  ** Set the container
  */

  public void setContainer(Object container) {
    container_ = container;
    return;
  }
 
  /***************************************************************************
  **
  ** Callback for completion of the element
  **
  */
  
  public boolean finishElement(String elemName) throws IOException {
    //
    // If we have a current worker handling the element, let them know.
    // If they say they are finished, drop them as the current worker
    //
    if (currWorker_ != null) {
      boolean dropWorker = currWorker_.finishElement(elemName);
      if (dropWorker) {
        if (!elemName.equals(currElem_)) {
          throw new IllegalStateException();
        }
        currWorker_ = null;
        currElem_ = null;
      }
      // It turns out if we do not have a single top-level enclosing element, we may
      // need to release several times during our tenure:
      // return (false);
      return (myKeys_.contains(elemName));
    }
     
    localFinishElement(elemName);
    
    //
    // If the element was one of our top-level routing keys, we notify
    // our caller:
    //
    
    return (myKeys_.contains(elemName));
  }
  
  /***************************************************************************
  ** 
  ** Handle incoming characters
  */

  public void processCharacters(char[] chars, int start, int length) {
    if (currWorker_ != null) {
      currWorker_.processCharacters(chars, start, length);
      return;
    }
     
    localProcessCharacters(chars, start, length);
    return;
  }  
  
  /***************************************************************************
  **
  ** Return the element keywords that define the top-level objects we are
  ** handling.  Other keywords are not important to advertise.  We will
  ** route then when we get callled.
  */
  
  public Set<String> keywordsOfInterest() {
    return (myKeys_);
  }
  
  /***************************************************************************
  **
  ** Handle the attributes for the keyword
  **
  */
  
  public Object processElement(String elemName, Attributes attrs) throws IOException {
    //
    // If we have a current subworker, just pass it off to him.  If not, see if
    // we have a subworker that wants to process the tag, and make him the current
    // subworker.  If not, we process it ourselves.
    //
    
    if (currWorker_ != null) {
      return (currWorker_.processElement(elemName, attrs));
    }
     
    AbstractFactoryClient assignedWorker = workerMap_.get(elemName);
    if (assignedWorker != null) {
      currElem_ = elemName;
      currWorker_ = assignedWorker;
      Object createdElement = currWorker_.processElement(elemName, attrs);
      if (createdElement != null) {
        GlueStick glue = glueStickMap_.get(elemName);
        if (glue != null) {
          glue.glueKidToParent(createdElement, this, sharedWhiteboard_);
        }
      }
      return (createdElement);
    }
    
    return (localProcessElement(elemName, attrs));
  }
  
  /***************************************************************************
  ** 
  ** Install a worker
  */

  protected void installWorker(AbstractFactoryClient worker, GlueStick glue) {
    Set<String> uniqueKeys = worker.keywordsOfInterest();
 
    Iterator<String> ukit = uniqueKeys.iterator();
    while (ukit.hasNext()) {
      String key = ukit.next();
      workerMap_.put(key, worker);
      if (glue != null) {
        glueStickMap_.put(key, glue);
      }
    }
 
    return;
  }  
  
  /***************************************************************************
  **
  ** Handle processing my element
  **
  */
  
  protected abstract Object localProcessElement(String elemName, Attributes attrs) throws IOException;
  
  /***************************************************************************
  **
  ** Handle closing my elements
  **
  */
  
  protected void localFinishElement(String elemName) throws IOException {
    return;
  }

  /***************************************************************************
  **
  ** Handle my local characters
  **
  */
  
  protected void localProcessCharacters(char[] chars, int start, int length) {
    return;
  }  
 
}
```
