## Supplementary material for "BioFabric Visualization of Network Alignments": BioFabric Source Code: ParserClient.java

```
/*
**    Copyright (C) 2003-2018 Institute for Systems Biology 
**                            Seattle, Washington, USA. 
**
**    This library is free software; you can redistribute it and/or
**    modify it under the terms of the GNU Lesser General Public
**    License as published by the Free Software Foundation; either
**    version 2.1 of the License, or (at your option) any later version.
**
**    This library is distributed in the hope that it will be useful,
**    but WITHOUT ANY WARRANTY; without even the implied warranty of
**    MERCHANTABILITY or FITNESS FOR A PARTICULAR PURPOSE.  See the GNU
**    Lesser General Public License for more details.
**
**    You should have received a copy of the GNU Lesser General Public
**    License along with this library; if not, write to the Free Software
**    Foundation, Inc., 59 Temple Place, Suite 330, Boston, MA  02111-1307  USA
*/

package org.systemsbiology.biofabric.api.parser;

import java.util.Set;
import java.io.IOException;

import org.xml.sax.Attributes;

/****************************************************************************
**
** Interface implemented by classes that want to build themselves from
** XML input.
*/

public interface ParserClient {

  ////////////////////////////////////////////////////////////////////////////
  //
  // PUBLIC METHODS
  //
  ////////////////////////////////////////////////////////////////////////////

  /***************************************************************************
  ** 
  ** Return the element keywords (Strings) that we are interested in
  */

  public Set<String> keywordsOfInterest();

  /***************************************************************************
  ** 
  ** Set the container
  */

  public void setContainer(Object container);  
  
  /***************************************************************************
  ** 
  ** Handle the attributes for the keyword
  */

  public Object processElement(String elemName, Attributes attrs) throws IOException;
  
  /***************************************************************************
  ** 
  ** Handle incoming characters
  */

  public void processCharacters(char[] chars, int start, int length);  
  
  /***************************************************************************
  ** 
  ** Callback for completion of the element
  */

  public boolean finishElement(String elemName) throws IOException; 
}
```
