## Supplementary material for "BioFabric Visualization of Network Alignments": BioFabric Source Code: ExceptionHandler.java

```
/*
**    Copyright (C) 2003-2018 Institute for Systems Biology 
**                            Seattle, Washington, USA. 
**
**    This library is free software; you can redistribute it and/or
**    modify it under the terms of the GNU Lesser General Public
**    License as published by the Free Software Foundation; either
**    version 2.1 of the License, or (at your option) any later version.
**
**    This library is distributed in the hope that it will be useful,
**    but WITHOUT ANY WARRANTY; without even the implied warranty of
**    MERCHANTABILITY or FITNESS FOR A PARTICULAR PURPOSE.  See the GNU
**    Lesser General Public License for more details.
**
**    You should have received a copy of the GNU Lesser General Public
**    License along with this library; if not, write to the Free Software
**    Foundation, Inc., 59 Temple Place, Suite 330, Boston, MA  02111-1307  USA
*/

package org.systemsbiology.biofabric.api.util;

import javax.swing.JFrame;
import javax.swing.JOptionPane;

import org.systemsbiology.biofabric.ui.dialogs.ExceptionDialog;
import org.systemsbiology.biofabric.util.ResourceManager;

/****************************************************************************
**
** Exception Handler.  This is a Singleton.
*/

public final class ExceptionHandler {
  
  ////////////////////////////////////////////////////////////////////////////
  //
  // PRIVATE MEMBERS
  //
  ////////////////////////////////////////////////////////////////////////////

  private static ExceptionHandler singleton_;
  private JFrame topWindow_;
  private boolean paintException_;
  private boolean headless_;
  private boolean inProcess_;
  private String outOfMemoryMessage_;
  private String outOfMemoryTitle_;
  
  ////////////////////////////////////////////////////////////////////////////
  //
  // PUBLIC METHODS
  //
  ////////////////////////////////////////////////////////////////////////////

  /***************************************************************************
  ** 
  ** Report the exception
  */

  public void displayException(Exception ex) {
    //
    // Add version to allow for better error analysis: 
    //
    String jVer = System.getProperty("java.version");
    String jVend = System.getProperty("java.vendor");
    String os = System.getProperty("os.name");
    String version = "Java version = " + jVer + " vendor = " + jVend + " operating system = " + os;
     
    if (headless_) {
      if (inProcess_) {
        throw new HeadlessException(ex);
      } else {
        System.err.println(version);
        ex.printStackTrace();
        return;
      }
    }      
    if (topWindow_ == null) {
      System.err.println("Exception reporter not initialized");
      System.err.println(version);
      ex.printStackTrace();
      return;
    }
    System.err.println(version);
    ex.printStackTrace();
    ExceptionDialog ed = new ExceptionDialog(topWindow_, ex, version);
    ed.setVisible(true);
    return;
  }

  /***************************************************************************
  ** 
  ** Report out of memory error
  */

  public void displayOutOfMemory(OutOfMemoryError oom) {
    if (headless_) {
      oom.printStackTrace();
      throw oom;  
    }    
    if (topWindow_ == null) {
      System.err.println("Out of memory (Exception reporter not initialized)");
      oom.printStackTrace();
      throw oom;
    }
    oom.printStackTrace();
    JOptionPane.showMessageDialog(topWindow_, outOfMemoryMessage_,
                                  outOfMemoryTitle_, JOptionPane.ERROR_MESSAGE);
    throw oom;
  }  

  /***************************************************************************
  ** 
  ** Report the paint exception (only once per execution)
  */

  public void displayPaintException(Exception ex) {
    if (paintException_) {
      System.err.println("Extra paint exceptions continue");
      ex.printStackTrace();
      return;
    }
    paintException_ = true;
    displayException(ex);
    return;
  }  

  /***************************************************************************
  ** 
  ** Initialize
  */

  public void initialize(JFrame topWindow) {
    headless_ = false;
    topWindow_ = topWindow;
    return;
  }

  /***************************************************************************
  ** 
  ** Initialize for headless operation
  */

  public void initializeForHeadless(boolean inProcess) {
    headless_ = true;
    inProcess_ = inProcess;
    topWindow_ = null;
    return;
  }
  
  ////////////////////////////////////////////////////////////////////////////
  //
  // PUBLIC STATIC METHODS
  //
  ////////////////////////////////////////////////////////////////////////////

  /***************************************************************************
  ** 
  ** Get the singleton
  */

  public static synchronized ExceptionHandler getHandler() {
    if (singleton_ == null) {
      singleton_ = new ExceptionHandler();
    }
    return (singleton_);
  }

  ////////////////////////////////////////////////////////////////////////////
  //
  // PRIVATE CONSTRUCTORS
  //
  ////////////////////////////////////////////////////////////////////////////

  /***************************************************************************
  **
  ** Null Constructor
  */

  private ExceptionHandler() {
    paintException_ = false;
    ResourceManager rMan = ResourceManager.getManager();
    outOfMemoryMessage_ = rMan.getString("errorMsg.outOfMemory");
    outOfMemoryTitle_ = rMan.getString("errorMsg.outOfMemoryTitle");
  }
  
  ////////////////////////////////////////////////////////////////////////////
  //
  // PUBLIC INNER CLASSES
  //
  ////////////////////////////////////////////////////////////////////////////

  /***************************************************************************
  ** 
  ** When running headless as a standalone program, we issue error messages
  ** to System.err.  But when operating in-process, we want to propagate the
  ** exception back to the caller.  So we need to throw a RuntimeException
  ** to hide this from everybody who doesn't need to know.
  */

  public static class HeadlessException extends RuntimeException {
    
    private Exception wrapped_;
    
    HeadlessException(Exception wrapped) {
      wrapped_ = wrapped;
    }
    
    public Exception getWrappedException() {
      return (wrapped_);
    }
  }
}
```
