## Supplementary material for "BioFabric Visualization of Network Alignments": BioFabric Source Code: FixedJButton.java

```
/*
**    Copyright (C) 2003-2018 Institute for Systems Biology 
**                            Seattle, Washington, USA. 
**
**    This library is free software; you can redistribute it and/or
**    modify it under the terms of the GNU Lesser General Public
**    License as published by the Free Software Foundation; either
**    version 2.1 of the License, or (at your option) any later version.
**
**    This library is distributed in the hope that it will be useful,
**    but WITHOUT ANY WARRANTY; without even the implied warranty of
**    MERCHANTABILITY or FITNESS FOR A PARTICULAR PURPOSE.  See the GNU
**    Lesser General Public License for more details.
**
**    You should have received a copy of the GNU Lesser General Public
**    License along with this library; if not, write to the Free Software
**    Foundation, Inc., 59 Temple Place, Suite 330, Boston, MA  02111-1307  USA
*/

package org.systemsbiology.biofabric.api.util;

import java.awt.Dimension;
import java.awt.Insets;
import java.net.URL;
import javax.swing.BorderFactory;
import javax.swing.JButton;
import javax.swing.Icon;
import javax.swing.ImageIcon;
import javax.swing.border.Border;
import javax.swing.border.CompoundBorder;


/****************************************************************************
**
** Fixed Size Button
*/

public class FixedJButton extends JButton {
  
  ////////////////////////////////////////////////////////////////////////////
  //
  // PUBLIC CONSTANTS
  //
  ////////////////////////////////////////////////////////////////////////////
  
  public static final int BOTH_FIXED      = 0;
  public static final int VERT_ONLY_FIXED = 1;
  public static final int HORZ_ONLY_FIXED = 2;
  
  ////////////////////////////////////////////////////////////////////////////
  //
  // PRIVATE INSTANCE MEMBERS
  //
  ////////////////////////////////////////////////////////////////////////////

  private int which_;
  
  private static final long serialVersionUID = 1L;

  ////////////////////////////////////////////////////////////////////////////
  //
  // PUBLIC CONSTRUCTORS
  //
  ////////////////////////////////////////////////////////////////////////////
 
  /***************************************************************************
  **
  ** Constructor
  */

  public FixedJButton(String name) {
    super(name);
    which_ = BOTH_FIXED;
  }  
   
  /***************************************************************************
  **
  ** Constructor
  */

  public FixedJButton(int which, String name) {
    super(name);
    which_ = which;
  }  
  
  /***************************************************************************
  **
  ** Constructor for reducing label pad
  */

  public FixedJButton(String name, int reduce) {
    super(name);
    which_ = BOTH_FIXED;
    Border bord = getBorder();
    if (bord instanceof CompoundBorder) {
      CompoundBorder comp = (CompoundBorder)bord;
      Insets bIn = comp.getInsideBorder().getBorderInsets(this);
      Border outBord = comp.getOutsideBorder();
      Border eBord = BorderFactory.createEmptyBorder(bIn.top, bIn.left - reduce, bIn.bottom, bIn.right - reduce);
      CompoundBorder nComp = BorderFactory.createCompoundBorder(outBord, eBord);
      this.setBorder(nComp);
    } else {
      Insets bIn = bord.getBorderInsets(this);
      Border eBord = BorderFactory.createEmptyBorder(bIn.top, bIn.left - reduce, bIn.bottom, bIn.right - reduce);
      this.setBorder(eBord);
    }
  }  
   
  /***************************************************************************
  **
  ** Constructor
  */

  public FixedJButton(Icon icon) {
    super(icon);
    which_ = BOTH_FIXED;
  }
  
  /***************************************************************************
  **
  ** Constructor for reducing label pad
  */

  public FixedJButton(Icon icon, int reduce) {
    super(icon);
    which_ = BOTH_FIXED;
    Border bord = getBorder();
    if (bord instanceof CompoundBorder) {
      CompoundBorder comp = (CompoundBorder)bord;
      Insets bIn = comp.getInsideBorder().getBorderInsets(this);
      Border outBord = comp.getOutsideBorder();
      Border eBord = BorderFactory.createEmptyBorder(bIn.top, bIn.left - reduce, bIn.bottom, bIn.right - reduce);
      CompoundBorder nComp = BorderFactory.createCompoundBorder(outBord, eBord);
      this.setBorder(nComp);
    } else {
      Insets bIn = bord.getBorderInsets(this);
      Border eBord = BorderFactory.createEmptyBorder(bIn.top, bIn.left - reduce, bIn.bottom, bIn.right - reduce);
      this.setBorder(eBord);
    }
  } 
  
  ////////////////////////////////////////////////////////////////////////////
  //
  // PUBLIC METHODS
  //
  ////////////////////////////////////////////////////////////////////////////

  /***************************************************************************
  **
  ** Make pictures square
  */
  
  public Dimension getPreferredSize() {
    Dimension ps = super.getPreferredSize();
    if (getIcon() != null) {
      ps.width = ps.height;
    }
    return (ps);
  }  
 
  /***************************************************************************
  **
  ** Fixed minimum
  */
  
  public Dimension getMinimumSize() {
    if (which_ == BOTH_FIXED) {
      return (getPreferredSize());
    } else if (which_ == VERT_ONLY_FIXED) {
      int w = super.getMinimumSize().width;
      int h = getPreferredSize().height;
      return (new Dimension(w, h));
    } else if (which_ == HORZ_ONLY_FIXED) {
      int h = super.getMinimumSize().height;
      int w = getPreferredSize().width;
      return (new Dimension(w, h));
    }
    throw new IllegalArgumentException();
  }
  
  /***************************************************************************
  **
  ** Fixed Maximum
  */
  
  public Dimension getMaximumSize() {
    if (which_ == BOTH_FIXED) {
      return (getPreferredSize());
    } else if (which_ == VERT_ONLY_FIXED) {
      int w = super.getMaximumSize().width;
      int h = getPreferredSize().height;
      return (new Dimension(w, h));
    } else if (which_ == HORZ_ONLY_FIXED) {
      int h = super.getMaximumSize().height;
      int w = getPreferredSize().width;
      return (new Dimension(w, h));
    }
    throw new IllegalArgumentException();
  }

  ////////////////////////////////////////////////////////////////////////////
  //
  // PUBLIC METHODS
  //
  ////////////////////////////////////////////////////////////////////////////  
  
  /***************************************************************************
  **
  ** Mac Button sizing is a joke:
  */

  public static FixedJButton miniFactory(String name, int reduce) {    
    boolean isAMac = System.getProperty("os.name").toLowerCase().startsWith("mac os x");
    if (isAMac) {
      String letter = name.substring(0, 1);
      if (letter.equals("<")) {
        letter = "LT";
      } else if (letter.equals(">")) {
        letter = "GT";
      }
      URL img = FixedJButton.class.getResource("/org/systemsbiology/biofabric/images/Mac" + letter + ".gif");
      if (img == null) {
        img = FixedJButton.class.getResource("/org/systemsbiology/biofabric/images/MacBlank.gif");
      }
      ImageIcon pix = new ImageIcon(img);
      FixedJButton retval = new FixedJButton(pix);
      retval.setOpaque(false);
      return (retval);
    } else {
      return (new FixedJButton(name, reduce));
    }   
  }    
}
```
