## Supplementary material for "BioFabric Visualization of Network Alignments": BioFabric Source Code: MatchingJLabel.java

```
/*
**    Copyright (C) 2003-2018 Institute for Systems Biology 
**                            Seattle, Washington, USA. 
**
**    This library is free software; you can redistribute it and/or
**    modify it under the terms of the GNU Lesser General Public
**    License as published by the Free Software Foundation; either
**    version 2.1 of the License, or (at your option) any later version.
**
**    This library is distributed in the hope that it will be useful,
**    but WITHOUT ANY WARRANTY; without even the implied warranty of
**    MERCHANTABILITY or FITNESS FOR A PARTICULAR PURPOSE.  See the GNU
**    Lesser General Public License for more details.
**
**    You should have received a copy of the GNU Lesser General Public
**    License along with this library; if not, write to the Free Software
**    Foundation, Inc., 59 Temple Place, Suite 330, Boston, MA  02111-1307  USA
*/

package org.systemsbiology.biofabric.api.util;

import java.awt.Dimension;

import javax.swing.JLabel;


/****************************************************************************
**
** Fixed Size Button
*/

public class MatchingJLabel extends JLabel {
  
  ////////////////////////////////////////////////////////////////////////////
  //
  // PRIVATE INSTANCE MEMBERS
  //
  ////////////////////////////////////////////////////////////////////////////

  private JLabel reference_;

  
  private static final long serialVersionUID = 1L;
  
  ////////////////////////////////////////////////////////////////////////////
  //
  // PUBLIC CONSTRUCTORS
  //
  ////////////////////////////////////////////////////////////////////////////
 
  /***************************************************************************
  **
  ** Constructor
  */

  public MatchingJLabel(String name, JLabel reference) {
    super(name);
    reference_ = reference;
  }  
 
  ////////////////////////////////////////////////////////////////////////////
  //
  // PUBLIC METHODS
  //
  ////////////////////////////////////////////////////////////////////////////

  /***************************************************************************
  **
  ** Make pictures square
  */
  
  @Override
  public Dimension getPreferredSize() {
    Dimension ps = super.getPreferredSize();
    Dimension rps = reference_.getPreferredSize();
    if (rps.width > ps.width) {
      ps.width = rps.width;
    }
    return (ps);
  }  
 
  /***************************************************************************
  **
  ** Fixed minimum
  */
  
  @Override
  public Dimension getMinimumSize() {
    return (getPreferredSize());
  }
  
  /***************************************************************************
  **
  ** Fixed Maximum
  */
  
  @Override
  public Dimension getMaximumSize() {
    return (getPreferredSize());
  }
}
```
