## Supplementary material for "BioFabric Visualization of Network Alignments": BioFabric Source Code: NID.java

```
/*
**    Copyright (C) 2003-2018 Institute for Systems Biology 
**                            Seattle, Washington, USA. 
**
**    This library is free software; you can redistribute it and/or
**    modify it under the terms of the GNU Lesser General Public
**    License as published by the Free Software Foundation; either
**    version 2.1 of the License, or (at your option) any later version.
**
**    This library is distributed in the hope that it will be useful,
**    but WITHOUT ANY WARRANTY; without even the implied warranty of
**    MERCHANTABILITY or FITNESS FOR A PARTICULAR PURPOSE.  See the GNU
**    Lesser General Public License for more details.
**
**    You should have received a copy of the GNU Lesser General Public
**    License along with this library; if not, write to the Free Software
**    Foundation, Inc., 59 Temple Place, Suite 330, Boston, MA  02111-1307  USA
*/

package org.systemsbiology.biofabric.api.util;

import java.util.Collection;
import java.util.HashSet;
import java.util.Map;
import java.util.Set;

public final class NID implements Cloneable, Comparable<NID> {
  
  private final String id_;
  
  public NID(String id) {
    this.id_ = id;
  }  
 
  public NID(NID other) {
    this.id_ = other.id_;
  }
  
  public String getInternal() {
    return (this.id_);
  }
  
  @Override
  public NID clone() {
    try {
      NID newVal = (NID)super.clone();
      return (newVal);
    } catch (CloneNotSupportedException ex) {
      throw new IllegalStateException();     
    }    
  }
  
  @Override
  public int hashCode() {
    return (id_.hashCode());
  }
  
  @Override
  public boolean equals(Object other) {
    if (other == null) {
      return (false);
    }
    if (other == this) {
      return (true);
    }
    if (!(other instanceof NID)) {
      return (false);
    }
    NID otherMM = (NID)other;
    return (this.id_.equals(otherMM.id_));
  }  
 
  @Override
  public String toString() {
    return ("OID: " + id_);
  }
  
  public int compareTo(NID other) {
    return (this.id_.compareTo(other.id_));
  }
  
  //
  // A class that can be used to order nodes using their names, with the NID only as a tie-breaker if names match.
  //
  
  public final static class WithName implements Cloneable, Comparable<WithName> {
  
	  private final NID nid_;
	  private final String name_;
	  
	  public WithName(NID id, String name) {
	  	this.name_ = name;
	    this.nid_ = id;
	  }  
	 
	  public WithName(WithName other) {
	    this.nid_ = other.nid_;
	    this.name_ = other.name_;
	  }
	  
	  @Override
	  public NID.WithName clone() {
	    try {
	      NID.WithName newVal = (NID.WithName)super.clone();
	      return (newVal);
	    } catch (CloneNotSupportedException ex) {
	      throw new IllegalStateException();     
	    }    
	  }
	  
	  @Override
	  public int hashCode() {
	    return (nid_.hashCode());
	  }
	  
	  public NID getNID() {
	    return (nid_);
	  }
	  
	  public String getName() {
	    return (name_);
	  }
	  
	  @Override
	  public boolean equals(Object other) {
	    if (other == null) {
	      return (false);
	    }
	    if (other == this) {
	      return (true);
	    }
	    if (!(other instanceof WithName)) {
	      return (false);
	    }
	    WithName otherMM = (WithName)other;
	    boolean eq = this.nid_.equals(otherMM.nid_);
	    boolean neq = this.name_.equals(otherMM.name_);
	    if (eq && !neq) {
	    	throw new IllegalStateException();
	    }    
	    return (eq);
	  }  
	 
	  @Override
	  public String toString() {
	    return ("WithName: " + nid_ + " " + name_);
	  }
	  
	  //
	  // This is the meat of the class. Comparison depends on name. If names actually equal, we fall back on NID. Note we do
	  // NOT normalize the names: if there are case differences, those are better to use than arbitrary (but consistent) 
	  // internal ID.
	  public int compareTo(WithName other) {
	  	boolean eq = this.nid_.equals(other.nid_);
	    boolean neq = this.name_.equals(other.name_);
	    if (eq && !neq) {
	    	throw new IllegalStateException();
	    }	
	  	int nDiff = this.name_.compareTo(other.name_);
	  	if (nDiff != 0) {
	  		return (nDiff);
	  	}
	    return (this.nid_.compareTo(other.nid_));
	  }
  }
  
  public static Collection<WithName> addNames(Collection<NID> nids, Map<NID, String> idToName, Collection<WithName> target) {
  	for (NID nid : nids) {
  		target.add(new WithName(nid, idToName.get(nid)));
  	}
  	return (target);
  }
  
  
  public static Set<String> justNames(Collection<NID.WithName> nids) {
  	HashSet<String> retval = new HashSet<String>();
  	for (NID.WithName nid : nids) {
  		String name = nid.getName();
  		if (retval.contains(name)) {
  			throw new IllegalStateException();
  		}
  		retval.add(name);
  	}
  	return (retval);
  }

}
```
