## Supplementary material for "BioFabric Visualization of Network Alignments": BioFabric Source Code: UniqueLabeller.java

```
/*
**    Copyright (C) 2003-2010 Institute for Systems Biology 
**                            Seattle, Washington, USA. 
**
**    This library is free software; you can redistribute it and/or
**    modify it under the terms of the GNU Lesser General Public
**    License as published by the Free Software Foundation; either
**    version 2.1 of the License, or (at your option) any later version.
**
**    This library is distributed in the hope that it will be useful,
**    but WITHOUT ANY WARRANTY; without even the implied warranty of
**    MERCHANTABILITY or FITNESS FOR A PARTICULAR PURPOSE.  See the GNU
**    Lesser General Public License for more details.
**
**    You should have received a copy of the GNU Lesser General Public
**    License along with this library; if not, write to the Free Software
**    Foundation, Inc., 59 Temple Place, Suite 330, Boston, MA  02111-1307  USA
*/

package org.systemsbiology.biofabric.api.util;

import java.io.BufferedReader;
import java.io.FileReader;
import java.util.Comparator;
import java.util.HashSet;
import java.util.TreeSet;
import java.util.Iterator;

/****************************************************************************
**
** Utility for getting unique labelling
*/

public final class UniqueLabeller implements Cloneable {

  ////////////////////////////////////////////////////////////////////////////
  //
  // PRIVATE INSTANCE FIELDS
  //
  ////////////////////////////////////////////////////////////////////////////

  private TreeSet existing_;
  private HashSet legacy_;
  private boolean isNumeric_;
  private String fixedPrefix_;
  
  ////////////////////////////////////////////////////////////////////////////
  //
  // PUBLIC CONSTRUCTORS
  //
  ////////////////////////////////////////////////////////////////////////////

  /***************************************************************************
  **
  ** Null constructor
  */

  public UniqueLabeller() {
    existing_ = new TreeSet(new KeyComparator());
    legacy_ = new HashSet();
    isNumeric_ = true;
    fixedPrefix_ = null;
  }
  
  /***************************************************************************
  **
  ** Copy constructor
  */

  public UniqueLabeller(UniqueLabeller other) {
    this.existing_ = (TreeSet)other.existing_.clone(); // Immutable contents; shallow OK
    this.isNumeric_ = other.isNumeric_;
    this.fixedPrefix_ = other.fixedPrefix_;
    this.legacy_ = (HashSet)other.legacy_.clone(); // Immutable contents; shallow OK
  }  
  
  ////////////////////////////////////////////////////////////////////////////
  //
  // PUBLIC INSTANCE METHODS
  //
  ////////////////////////////////////////////////////////////////////////////

  /***************************************************************************
  **
  ** Force to use String keys
  */

  public void setToStringKeys() {
    if (!existing_.isEmpty() || !legacy_.isEmpty()) {
      throw new IllegalStateException();
    }
    isNumeric_ = false;
    return;
  }  
 
  /***************************************************************************
  **
  ** Set the fixed prefix
  */

  public void setFixedPrefix(String prefix) {
    if (!existing_.isEmpty() || !legacy_.isEmpty()) {
      throw new IllegalStateException();
    }
    isNumeric_ = false;
    fixedPrefix_ = prefix;
    return;
  }  
  
  /***************************************************************************
  **
  ** Clone
  */

  public Object clone() {
    try {
      UniqueLabeller retval = (UniqueLabeller)super.clone();
      retval.existing_ = (TreeSet)this.existing_.clone(); // Contents are immutable; shallow OK
      if (!(retval.existing_.comparator() instanceof KeyComparator)) {
        throw new IllegalStateException();
      }
      retval.legacy_ = (HashSet)this.legacy_.clone(); // Contents are immutable; shallow OK
      return (retval);
    } catch (CloneNotSupportedException cnse) {
      throw new IllegalStateException();
    }
  }
  
  /***************************************************************************
  **
  ** Create a copy that changes the prefixes of those keys containing the prefix
  ** 4/15/09 WJRL: With overhaul, we assume ALL keys must have the prefix, as per
  ** analysis of existing usages...
  */
  
  public UniqueLabeller mappedPrefixCopy(String oldPrefix, String newPrefix) {
   
    UniqueLabeller retval = (UniqueLabeller)this.clone();
    if (this.fixedPrefix_ == null) {
      throw new IllegalStateException();
    }
    if (!this.legacy_.isEmpty()) {
      throw new IllegalStateException();
    }    
    if (!this.fixedPrefix_.equals(oldPrefix)) {
      throw new IllegalArgumentException();
    }
    retval.fixedPrefix_ = newPrefix;
    return (retval);   
  }  
  
  /***************************************************************************
  **
  ** Add a preexisting label.  We are either a numeric label or an alphanumeric
  ** label type, based on what we see.  Returns false if the label already
  ** exists.
  */
  
  public boolean addExistingLabel(String label) {

    boolean retval;
    int intval = -1;
    String stringVal = null;
    
    if ((label == null) || label.trim().equals("")) {
      System.err.println("bad label " + label);
      throw new IllegalArgumentException();
    }
    
    if (label.toLowerCase().equals(label.toUpperCase())) {
      try {
        intval = Integer.parseInt(label);
      } catch (NumberFormatException nfe) {
        stringVal = label;
      }
    } else {
      stringVal = label;
    }
    
    //
    // If we have a fixed prefix, label better have the prefix, unless
    // it is a leacy label:
    //
    if (fixedPrefix_ != null) {
      if (label.indexOf(fixedPrefix_) != 0) {
        if (!legacy_.contains(label)) {
          System.err.println("label " + label + " fixedPrefix = " + fixedPrefix_);
          throw new IllegalArgumentException();
        } else {
          return (false);
        }
      }
      label = label.substring(fixedPrefix_.length());
    }

    if (legacy_.contains(label)) {
      return (false);
    }

    if (isNumeric_) {
      if (stringVal == null) {
        retval = existing_.add(new Integer(intval));
      } else {
        convertToAlpha();
        retval = existing_.add(stringVal);
      }
    } else {
      retval = existing_.add(label);
    }
    return (retval);   
  }
  
  /***************************************************************************
  **
  ** Due to the overhaul of the key issue algorithm, the longest key names
  ** form the basis of the next key, not at "zzz..." tag.  Add the concept
  ** of legacy labels to prexisting hold long names
  ** 
  */
  
  public boolean addExistingLegacyLabel(String label) {

    boolean retval;
    String stringVal = null;
    
    if ((label == null) || label.trim().equals("")) {
      System.err.println("bad label " + label);
      throw new IllegalArgumentException();
    }
    if (isNumeric_) {
      throw new IllegalStateException();
    }
    retval = legacy_.add(label);
    return (retval);   
  }    

  /***************************************************************************
  **
  ** Answer of ew are empty
  */
  
  public boolean isEmpty() {
    return (legacy_.isEmpty() && existing_.isEmpty());
  }
 
  /***************************************************************************
  **
  ** Get a unique node ID. 
  */
  
  public NID getNextOID() {
  	return (new NID(getNextLabel()));
  }
  
  /***************************************************************************
  **
  ** Get a unique label. 
  */
  
  public String getNextLabel() {    
    //
    // If we are numeric, bump up the last value.  If we are alpha,
    // increment the last character    //
    
    if (isNumeric_) {
      Integer newVal = null;
      if (existing_.size() == 0) {
        newVal = new Integer(0);
      } else {
        Integer highVal = (Integer)existing_.last();
        // Make the algorithm super-robust!
        if (highVal.intValue() == Integer.MAX_VALUE) {
          convertToAlpha();
          if (!legacy_.isEmpty() || (fixedPrefix_ != null)) {
            throw new IllegalStateException();
          }
          String highStrVal = (String)existing_.last();
          String newStrVal = genNextAlpha(highStrVal);
          boolean replaced = existing_.add(newStrVal);
          if (!replaced) {
            throw new IllegalStateException();
          }
          return (newStrVal); 
        }
        newVal = new Integer(highVal.intValue() + 1);
      }
      boolean replaced = existing_.add(newVal);
      if (!replaced) {
        throw new IllegalStateException();
      } 
      return (newVal.toString());
    } else {
      String highVal = (existing_.isEmpty()) ? null : (String)existing_.last();
      String newVal = null;
      // If we have legacy keys, keep bumping up if we have a collision:
      while (true) {
        newVal = genNextAlpha(highVal);
        if (legacy_.isEmpty()) {
          break;
        }
        String checkVal = (fixedPrefix_ != null) ? fixedPrefix_ + newVal : newVal;
        if (!legacy_.contains(checkVal)) {
          break;
        }
        highVal = newVal;
      }
      boolean replaced = existing_.add(newVal);
      if (!replaced) {
        System.err.println("Not replaced " + newVal);
        throw new IllegalStateException();
      }
      if (fixedPrefix_ != null) {
        newVal = fixedPrefix_ + newVal;
      }
      return (newVal);
    }   
  }  

  /***************************************************************************
  **
  ** Remove a label.  Nothing happens if it is not there 
  */
  
  public void removeLabel(String label) {
    if (!legacy_.isEmpty() && legacy_.contains(label)) {
      legacy_.remove(label);
      return;
    }
    
    if (existing_.size() == 0) {
      return;
    }     
    Object first = existing_.iterator().next();
    if (first instanceof Integer) {
      try {
        Integer intval = Integer.valueOf(label);
        if (!existing_.contains(intval)) {
          return;
        }        
      } catch (NumberFormatException nfe) {
        return;
      }
    } else {
      if (fixedPrefix_ != null) {
        if (label.indexOf(fixedPrefix_) != 0) {
          return;
        }
        label = label.substring(fixedPrefix_.length());
      }
      if (!existing_.contains(label)) {  // WAS: Class cast exception if set contains Integers, so do above junk
        return;
      }
    }
    if (isNumeric_) {
      try {
        Integer intval = Integer.valueOf(label);
        existing_.remove(intval);
      } catch (NumberFormatException nfe) {
        // FIX ME: SEEN an exception here removing rmic/delta link segment 11-18-03
        // BECAUSE attempting to remove from an empty list does that! 4-27-04 (See above fix)
        System.err.println(existing_ + " " + label);
        throw new IllegalArgumentException();
      }
    } else {
      existing_.remove(label);  // once alpha, always alpha!
    }
  }
  
  /***************************************************************************
  **
  ** Standard to String
  */
  
  public String toString() {
    return ("UniqueLabeller: " + existing_ + " legacy = " + legacy_ + " isNumeric = " + isNumeric_ + " fixedPrefix_ = " + fixedPrefix_);
  }
 
  ////////////////////////////////////////////////////////////////////////////
  //
  // PUBLIC CLASS METHODS
  //
  ////////////////////////////////////////////////////////////////////////////  
  
  /***************************************************************************
  **
  ** Change key prefixes 
  */
  
  public static String mapKeyPrefix(String key, String oldPrefix, String newPrefix, StringBuffer buf) {
    int startIndex = oldPrefix.length();
    if (buf == null) {
      buf = new StringBuffer();
    }
    int index = key.indexOf(oldPrefix);
    if (index == 0) {
      buf.setLength(0);
      buf.append(newPrefix);
      buf.append(key.substring(startIndex));
      key = buf.toString();
    }
    return (key);   
  }    
  
  ////////////////////////////////////////////////////////////////////////////
  //
  // PRIVATE INSTANCE METHODS
  //
  ////////////////////////////////////////////////////////////////////////////
  
  /***************************************************************************
  **
  ** Can't be just numeric anymore; convert to alpha
  */
  
  private void convertToAlpha() {
    if (!isNumeric_) {
      throw new IllegalStateException();
    }
    TreeSet newSet = new TreeSet(new KeyComparator());
    Iterator eit = existing_.iterator();
    while (eit.hasNext()) {
      Integer oldval = (Integer)eit.next();
      boolean replaced = newSet.add(oldval.toString());
      if (!replaced) {
        throw new IllegalStateException();
      }
    }
    isNumeric_ = false;
    existing_ = newSet;
    return;   
  }
  
  /***************************************************************************
  **
  ** Generate next alpha key.  This is the original 2003 bogus version that
  ** creates long "zzzzzzzzzz...." strings 
  
  
  private String genNextAlphaObsolete() {
    String newVal = null;
    if (existing_.size() == 0) {
      newVal = "a";
    } else {
      String highVal = (String)existing_.last();
      char lastChar = highVal.charAt(highVal.length() - 1);
      if ((lastChar == 'Z') || (lastChar == 'z') || !Character.isLetter(lastChar)) {
        newVal = highVal + "a";
      } else {
        Character newLast = new Character((char)((int)lastChar + 1)); 
        newVal = highVal.substring(0, highVal.length() - 1) + newLast;
      }
    }
    return (newVal);   
  }  
  
  /***************************************************************************
  **
  ** Generate next alpha key.  This is the new version that is smarter
  */
  
  public static String genNextAlpha(String highVal) {
    String newVal;
    if (highVal == null) {
      newVal = "A";
    } else {
      newVal = recursiveCharBump(highVal);
      if (newVal == null) {
        newVal = nextLonger(highVal);
      }
    }
    return (newVal);   
  } 

  /***************************************************************************
  **
  ** Get next highest string
  */
  
  public static String nextLonger(String currHigh) {
    int currLen = currHigh.length();
    StringBuffer buf = new StringBuffer();
    for (int j = 0; j <= currLen; j++) {
      buf.append("A");
    }
    return (buf.toString());
  }
   
  /***************************************************************************
  **
  ** Recursive string bump
  */
  
  public static String recursiveCharBump(String keyFrag) {
    StringBuffer buf = new StringBuffer();
    int fragLen = keyFrag.length();
    if (fragLen == 0) {
      return (null);
    }
    
    char lastChar = keyFrag.charAt(fragLen - 1); 
    char replaceChar = getNextChar(lastChar);
    String currRoot = keyFrag.substring(0, fragLen - 1);
    if (((int)replaceChar) == 0) {
      String newRoot = recursiveCharBump(currRoot);

      if (newRoot == null) {
        return (null);
      } else {
        buf.append(newRoot);
      }
      buf.append('A');
    } else {
      buf.append(currRoot);
      buf.append(replaceChar);
    }
    return (buf.toString());
  }
 
  /***************************************************************************
  **
  ** Generate next character.
  */
  
  public static char getNextChar(char currChar) {
    if (currChar == 'z') {
      return (0);
    } else if (currChar == 'Z') {
      return ('a');
    } else if (!Character.isLetter(currChar)) {
      return (0);
    } else {  
      return ((char)((int)currChar + 1));
    }
  }  

  ////////////////////////////////////////////////////////////////////////////
  //
  // TEST METHODS
  //
  ////////////////////////////////////////////////////////////////////////////  
  
  /***************************************************************************
  **
  ** Test frame
  */

  public static void main(String[] argv) {
    try {
      HashSet holdLegLab = new HashSet();
      HashSet holdLabNew = new HashSet();
      BufferedReader in = new BufferedReader(new FileReader("/users/wlongaba/tmp/justColKeys"));
      String line = null;
      while ((line = in.readLine()) != null) {
        holdLegLab.add(line.trim());
      }
      holdLegLab.add("zz_newColor_0z");
      holdLabNew.add("zz_newColor_0d");
      in.close();
      for (int j = 0; j < 80; j++) {
        System.out.println("Reload " + j);
        UniqueLabeller lab = new UniqueLabeller();
        lab.setFixedPrefix("zz_newColor_0");
        Iterator hlit = holdLegLab.iterator();
        while (hlit.hasNext()) {
          String nextEx = (String)hlit.next();
          lab.addExistingLegacyLabel(nextEx);
        }
        Iterator hlnit = holdLabNew.iterator();
        while (hlnit.hasNext()) {
          String nextEx = (String)hlnit.next();
          lab.addExistingLabel(nextEx);
        }
        for (int i = 0; i < 50; i++) {
          String nextLab = lab.getNextLabel();
          System.out.println("Next Label " + nextLab);
          holdLabNew.add(nextLab);
        }      
      }
      
    } catch (Exception ex) {
      System.err.println("Exception " + ex);
    }
  }
  
 /***************************************************************************
  **
  ** Sorts nodes for output
  **
  */
  
  static class KeyComparator implements Comparator {
    public int compare(Object o1, Object o2) {
      if ((o1 instanceof Integer) && (o2 instanceof Integer)) {
        Integer i1 = (Integer)o1;
        Integer i2 = (Integer)o2;
        return (i1.compareTo(i2));
      }
      String key1 = (String)o1;
      String key2 = (String)o2;
      int len1 = key1.length();
      int len2 = key2.length();
      if (len1 != len2) {
        return ((len1 < len2) ? -1 : 1);
      }
      return (key1.compareTo(key2));
    }
  }
}
```
