## Supplementary material for "BioFabric Visualization of Network Alignments": BioFabric Source Code: BackgroundCore.java

```
/*
**    Copyright (C) 2003-2018 Institute for Systems Biology 
**                            Seattle, Washington, USA. 
**
**    This library is free software; you can redistribute it and/or
**    modify it under the terms of the GNU Lesser General Public
**    License as published by the Free Software Foundation; either
**    version 2.1 of the License, or (at your option) any later version.
**
**    This library is distributed in the hope that it will be useful,
**    but WITHOUT ANY WARRANTY; without even the implied warranty of
**    MERCHANTABILITY or FITNESS FOR A PARTICULAR PURPOSE.  See the GNU
**    Lesser General Public License for more details.
**
**    You should have received a copy of the GNU Lesser General Public
**    License along with this library; if not, write to the Free Software
**    Foundation, Inc., 59 Temple Place, Suite 330, Boston, MA  02111-1307  USA
*/


package org.systemsbiology.biofabric.api.worker;

/****************************************************************************
**
** Class to support running background threads
*/

public interface BackgroundCore  {

	/****************************************************************************
  **
  ** Value to return with an early cancellation
  */   
   
  public Object getEarlyResult();
		
  /****************************************************************************
  **
  ** This routine is where all the work gets done.  It is overridden by
  ** child classes.  All work occurs on a background thread (unless this is explictly a foregound worker).
  */   
   
  public Object runCore() throws AsynchExitRequestException;

  /****************************************************************************
  **
  ** This work occurs on the background thread AFTER we have sent off 
  ** the final update progress report to the UI thread, but before
  ** we report on the UI thread that we are finished (unless this is explictly a foregound worker).
  */  
    
  public Object postRunCore(); 
  
} 
```
