## Supplementary material for "BioFabric Visualization of Network Alignments": BioFabric Source Code: BackgroundWorkerControlManager.java

```
/*
**    Copyright (C) 2003-2008 Institute for Systems Biology 
**                            Seattle, Washington, USA. 
**
**    This library is free software; you can redistribute it and/or
**    modify it under the terms of the GNU Lesser General Public
**    License as published by the Free Software Foundation; either
**    version 2.1 of the License, or (at your option) any later version.
**
**    This library is distributed in the hope that it will be useful,
**    but WITHOUT ANY WARRANTY; without even the implied warranty of
**    MERCHANTABILITY or FITNESS FOR A PARTICULAR PURPOSE.  See the GNU
**    Lesser General Public License for more details.
**
**    You should have received a copy of the GNU Lesser General Public
**    License along with this library; if not, write to the Free Software
**    Foundation, Inc., 59 Temple Place, Suite 330, Boston, MA  02111-1307  USA
*/

package org.systemsbiology.biofabric.api.worker;


/****************************************************************************
**
** Class implemented by classes that disable and enable UI controls
** during asynch operations
*/

public interface BackgroundWorkerControlManager { 

  /****************************************************************************
  **
  ** disable
  */  
  
  public void disableControls();

  /****************************************************************************
  **
  ** enable
  */  
  
  public void reenableControls();  
  
  /****************************************************************************
  **
  ** also redraw....
  */  
  
  public void redraw();  

} 
```
