## Supplementary material for "BioFabric Visualization of Network Alignments": BioFabric Source Code: BackgroundWorkerOwner.java

```
/*
**    Copyright (C) 2003-2007 Institute for Systems Biology 
**                            Seattle, Washington, USA. 
**
**    This library is free software; you can redistribute it and/or
**    modify it under the terms of the GNU Lesser General Public
**    License as published by the Free Software Foundation; either
**    version 2.1 of the License, or (at your option) any later version.
**
**    This library is distributed in the hope that it will be useful,
**    but WITHOUT ANY WARRANTY; without even the implied warranty of
**    MERCHANTABILITY or FITNESS FOR A PARTICULAR PURPOSE.  See the GNU
**    Lesser General Public License for more details.
**
**    You should have received a copy of the GNU Lesser General Public
**    License along with this library; if not, write to the Free Software
**    Foundation, Inc., 59 Temple Place, Suite 330, Boston, MA  02111-1307  USA
*/

package org.systemsbiology.biofabric.api.worker;


/****************************************************************************
**
** Class implemented by owners of Background workers
*/

public interface BackgroundWorkerOwner { 

  /****************************************************************************
  **
  ** We get a change to process a background thread exception before it
  ** gets popped up in a window.  If we return true, the exception will
  ** not get displayed.
  */  
  
  public boolean handleRemoteException(Exception remoteEx);
  
  /****************************************************************************
  **
  ** Things to do if we get cancelled. Returns true if user chooses to restore.
  */  
  
  public boolean handleCancellation();  
  
  /****************************************************************************
  **
  ** These are the routines to be executed, on the UI thread, after the 
  ** background is done, but before the UI controls are enabled and
  ** the new model is painted (e.g. zoom to worksheet center).
  */
  
  public void cleanUpPreEnable(Object result);
  
  /****************************************************************************
  **
  ** These are the routines to be executed, on the UI thread, after the 
  ** background is done, and after the UI has been repainted (e.g. warning
  ** dialogs).
  */
  
  public void cleanUpPostRepaint(Object result, boolean skipImage);   
} 
```
