## Supplementary material for "BioFabric Visualization of Network Alignments": BioFabric Source Code: BTProgressMonitor.java

```
/*
**    Copyright (C) 2003-2018 Institute for Systems Biology 
**                            Seattle, Washington, USA. 
**
**    This library is free software; you can redistribute it and/or
**    modify it under the terms of the GNU Lesser General Public
**    License as published by the Free Software Foundation; either
**    version 2.1 of the License, or (at your option) any later version.
**
**    This library is distributed in the hope that it will be useful,
**    but WITHOUT ANY WARRANTY; without even the implied warranty of
**    MERCHANTABILITY or FITNESS FOR A PARTICULAR PURPOSE.  See the GNU
**    Lesser General Public License for more details.
**
**    You should have received a copy of the GNU Lesser General Public
**    License along with this library; if not, write to the Free Software
**    Foundation, Inc., 59 Temple Place, Suite 330, Boston, MA  02111-1307  USA
*/

package org.systemsbiology.biofabric.api.worker;

import java.util.SortedMap;

/****************************************************************************
**
** An interface for monitoring progress
*/

public interface BTProgressMonitor {
 
  ////////////////////////////////////////////////////////////////////////////
  //
  // PUBLIC METHODS
  //
  ////////////////////////////////////////////////////////////////////////////

  /***************************************************************************
  **
  ** set total
  */
  
  public void setTotal(int total);  
  
  /***************************************************************************
  **
  ** Get total
  */
  
  public int getTotal();
  
  /***************************************************************************
  **
  ** Callback
  */
  
  public boolean updateUnknownProgress();
   
  /***************************************************************************
  **
  ** Callback
  */
  
  public boolean updateProgress(int done);
  
  /***************************************************************************
  **
  ** Callback
  */
  
  public boolean updateProgressAndPhase(int done, String message);
  
  /***************************************************************************
  **
  ** Callback
  */
  
  public boolean updateRankings(SortedMap<Integer, Double> chartVals);
    
  /***************************************************************************
  **
  ** Callback
  */
  
  public boolean keepGoing();  
  

  /***************************************************************************
  **
  ** Get progress
  */  
  
  public int getProgress();  

}
```
