## Supplementary material for "BioFabric Visualization of Network Alignments": BioFabric Source Code: LoopReporter.java

```
/*
**    Copyright (C) 2003-2018 Institute for Systems Biology 
**                            Seattle, Washington, USA. 
**
**    This library is free software; you can redistribute it and/or
**    modify it under the terms of the GNU Lesser General Public
**    License as published by the Free Software Foundation; either
**    version 2.1 of the License, or (at your option) any later version.
**
**    This library is distributed in the hope that it will be useful,
**    but WITHOUT ANY WARRANTY; without even the implied warranty of
**    MERCHANTABILITY or FITNESS FOR A PARTICULAR PURPOSE.  See the GNU
**    Lesser General Public License for more details.
**
**    You should have received a copy of the GNU Lesser General Public
**    License along with this library; if not, write to the Free Software
**    Foundation, Inc., 59 Temple Place, Suite 330, Boston, MA  02111-1307  USA
*/

package org.systemsbiology.biofabric.api.worker;

public final class LoopReporter {
  
  private long max_;
  private long skipLines_;
  private BTProgressMonitor monitor_;
  private double startFrac_;
  private double endFrac_;
  private String message_;
  private long count_;
  private long skipProg_;
  private boolean indeterminate_;
  
  public LoopReporter(long max, int bins, BTProgressMonitor monitor, double startFrac, double endFrac,
  		                String message) {
  	indeterminate_ = false;
    max_ = (max == 0) ? 1 : max;
    monitor_ = monitor;
    startFrac_ = startFrac;
    endFrac_ = endFrac;
    message_ = message;
    skipLines_ = (bins == 0) ? 0 : max_ / bins;
    count_ = 0L;
    skipProg_ = skipLines_;
  }  
  
  public LoopReporter(long max, int bins, BTProgressMonitor monitor) {
  	indeterminate_ = true;
    monitor_ = monitor;
    count_ = 0L;
    max_ = (max == 0) ? 1 : max;
    skipLines_ = (bins == 0) ? 0 : max_ / bins;
    count_ = 0L;
    skipProg_ = skipLines_;
  } 
  

  public void report(long progress) throws AsynchExitRequestException {
    skipProg_ -= progress;
    count_ += progress;
	  if (skipProg_ <= 0) {
	  	if (indeterminate_) {
	  	  if (monitor_ != null) {
	        if (!monitor_.updateUnknownProgress()) {
	          throw new AsynchExitRequestException();
	        }
	      }	  		
	  	} else {
		    double currFrac = startFrac_ + ((endFrac_ - startFrac_) * (count_ / (double)max_));
		    // If you want detailed progress output for debug, here you go:
		    // System.out.println("CP: " + message_ + ": " + currFrac + " [" + startFrac_ + " - " + endFrac_ + "]");
		    // System.out.println("CPMEM " + Runtime.getRuntime().freeMemory() + " " + System.currentTimeMillis());
	    	if (monitor_ != null) {
	        if (!monitor_.updateProgressAndPhase((int)(currFrac * 100.0), message_)) {
	          throw new AsynchExitRequestException();
	        }
	      }
	  	}
      skipProg_ = skipLines_;
	  }
	  return;
  }
  
  public void finish() throws AsynchExitRequestException {
    if (monitor_ != null) {
    	if (indeterminate_) {
    			if (!monitor_.updateUnknownProgress()) {
    					throw new AsynchExitRequestException();
    			}
    	} else {
    		if (!monitor_.updateProgressAndPhase((int)(100.0), message_)) {
    			throw new AsynchExitRequestException();
    	  }    		
    	}
    }
	  return;
  }
  
  
  
  public void report() throws AsynchExitRequestException {
  	report(1);
	  return;
  } 
}
```
