## Supplementary material for "BioFabric Visualization of Network Alignments": BioFabric Source Code: ArgParser.java

```
/*
**    Copyright (C) 2003-2018 Institute for Systems Biology 
**                            Seattle, Washington, USA. 
**
**    This library is free software; you can redistribute it and/or
**    modify it under the terms of the GNU Lesser General Public
**    License as published by the Free Software Foundation; either
**    version 2.1 of the License, or (at your option) any later version.
**
**    This library is distributed in the hope that it will be useful,
**    but WITHOUT ANY WARRANTY; without even the implied warranty of
**    MERCHANTABILITY or FITNESS FOR A PARTICULAR PURPOSE.  See the GNU
**    Lesser General Public License for more details.
**
**    You should have received a copy of the GNU Lesser General Public
**    License along with this library; if not, write to the Free Software
**    Foundation, Inc., 59 Temple Place, Suite 330, Boston, MA  02111-1307  USA
*/


package org.systemsbiology.biofabric.app;

import java.util.ArrayList;
import java.util.List;
import java.util.HashMap;
import java.util.Map;

import org.systemsbiology.biofabric.util.ResourceManager;

/****************************************************************************
**
** Argument parser
*/

public class ArgParser {

  ////////////////////////////////////////////////////////////////////////////
  //
  // PUBLIC STATIC CONSTANTS
  //
  ////////////////////////////////////////////////////////////////////////////  

  public enum AppType {VIEWER, PIPELINE};
   
  public static final String FILE         = "file";
  public static final String PLUG_IN_DIR  = "plugInDir";  
  
  public static final String SIF_BATCH_INPUT    = "sifImport";
  public static final String IMAGE_BATCH_OUTPUT = "pngExport";
  public static final String IMAGE_OUTPUT_WIDTH = "pngWidth";

  
  ////////////////////////////////////////////////////////////////////////////
  //
  // PUBLIC CONSTRUCTORS
  //
  ////////////////////////////////////////////////////////////////////////////    
        
  public ArgParser() {
  }  

  ////////////////////////////////////////////////////////////////////////////
  //
  // PUBLIC METHODS
  //
  ////////////////////////////////////////////////////////////////////////////  
 
  /***************************************************************************
  ** 
  ** Parse the args.  Returns null on failure.
  */

  public Map<String, Object> parse(AppType type, String argv[]) {
    // Parse arguments here...    
    ArrayList<String> list = new ArrayList<String>();
    for (int i = 0; i < argv.length; i++) {
      String arg = argv[i];
      list.add((arg.startsWith("-")) ? arg.toLowerCase() : arg);
    }
    
    HashMap<String, Object> argMap = new HashMap<String, Object>();
    String finalArg = (list.size() > 0) ? list.get(list.size() - 1) : null;
    List<ArgInfo> targs = loadForType(type);
    
    int tNum = targs.size();
    for (int i = 0; i < tNum; i++) {
      ArgInfo ai = targs.get(i);
      int index = list.indexOf("-" + ai.key.toLowerCase());
      if (!ai.followOn) {
        Boolean argVal;
        if (index != -1) {
          argVal = Boolean.valueOf(true);
          list.remove(index);
        } else {
          argVal = Boolean.valueOf(false);
        }      
        argMap.put(ai.key, argVal);
      } else {
        String argVal;
        if (index != -1) {
          if (index == (list.size() - 1)) {
            return (null);
          }
          argVal = list.get(index + 1);
          if (argVal.startsWith("-")) {
            return (null);
          }
          list.remove(index + 1);
          list.remove(index);
        } else {
          argVal = null;
        }      
        argMap.put(ai.key, argVal);
      }
    }
    
    if (list.size() == 0) {
      return (argMap);  
    }
    
    if (list.size() > 1) {
      return (null);
    }
    
    String remains = list.get(0);
    if (remains.equals(finalArg)) {
      argMap.put(FILE, remains);
    }
   
    return (argMap);
  }
  
  /***************************************************************************
  ** 
  ** Parse the args.  Returns null on failure.
  */

  public String getUsage(AppType type) {
    List<ArgInfo> targs = loadForType(type);         
    return (generateUsage(targs));    
  }
  
  ////////////////////////////////////////////////////////////////////////////
  //
  // PRIVATE INNER CLASSES
  //
  ////////////////////////////////////////////////////////////////////////////  

  /***************************************************************************
  **
  ** Holds argument info
  */  
  
  private class ArgInfo {
    String key;
    boolean followOn;
    boolean opKey;
    
    ArgInfo(String key, boolean followOn, boolean opKey) {
      this.key = key;
      this.followOn = followOn;
      this.opKey = opKey;
    }
  }
 
  ////////////////////////////////////////////////////////////////////////////
  //
  // PRIVATE METHODS
  //
  ////////////////////////////////////////////////////////////////////////////    

  /***************************************************************************
  ** 
  ** Prepare the target list
  */

  private List<ArgInfo> loadForType(AppType type) {
    ArrayList<ArgInfo> retval = new ArrayList<ArgInfo>();
    if (type == AppType.VIEWER) {
      retval.add(new ArgInfo(PLUG_IN_DIR, true, true));
      retval.add(new ArgInfo(FILE, true, true));
    } else if (type == AppType.PIPELINE) {      
      retval.add(new ArgInfo(SIF_BATCH_INPUT, true, false));   
      retval.add(new ArgInfo(IMAGE_BATCH_OUTPUT, true, false)); 
      retval.add(new ArgInfo(IMAGE_OUTPUT_WIDTH, true, false));   
    } else {
      throw new IllegalArgumentException();  
    }
    return (retval);
  }
  
  /***************************************************************************
  ** 
  ** Generate a usage string
  */

  private String generateUsage(List<ArgInfo> targs) {
    StringBuffer buf = new StringBuffer();
    ResourceManager rMan = ResourceManager.getManager();
    
    buf.append(rMan.getString("argParser.arguments"));
    buf.append("\n");    
    int tNum = targs.size();
    for (int i = 0; i < tNum; i++) {
      ArgInfo ai = targs.get(i);
      buf.append((ai.opKey) ? " [ -" : " -");
      buf.append(ai.key);
      buf.append((ai.opKey) ? " ] " : " ");
      if (ai.followOn) {
        buf.append(" argument");
        buf.append((ai.opKey) ? " ] " : "");
      }
      buf.append("\n");
    }
    return (buf.toString());
  }
}
```
