## Supplementary material for "BioFabric Visualization of Network Alignments": BioFabric Source Code: BioFabricApplication.java

```
/*
**    Copyright (C) 2003-2018 Institute for Systems Biology 
**                            Seattle, Washington, USA. 
**
**    This library is free software; you can redistribute it and/or
**    modify it under the terms of the GNU Lesser General Public
**    License as published by the Free Software Foundation; either
**    version 2.1 of the License, or (at your option) any later version.
**
**    This library is distributed in the hope that it will be useful,
**    but WITHOUT ANY WARRANTY; without even the implied warranty of
**    MERCHANTABILITY or FITNESS FOR A PARTICULAR PURPOSE.  See the GNU
**    Lesser General Public License for more details.
**
**    You should have received a copy of the GNU Lesser General Public
**    License along with this library; if not, write to the Free Software
**    Foundation, Inc., 59 Temple Place, Suite 330, Boston, MA  02111-1307  USA
*/

package org.systemsbiology.biofabric.app;

import java.awt.Dimension;
import java.awt.event.WindowAdapter;
import java.awt.event.WindowEvent;
import java.util.HashMap;
import java.util.Map;
import javax.swing.JFrame;
import javax.swing.SwingUtilities;
import javax.swing.WindowConstants;

import org.systemsbiology.biofabric.api.util.ExceptionHandler;
import org.systemsbiology.biofabric.cmd.CommandSet;
import org.systemsbiology.biofabric.plugin.PlugInManager;
import org.systemsbiology.biofabric.ui.dialogs.UpdateJavaDialog;
import org.systemsbiology.biofabric.util.ResourceManager;
import org.systemsbiology.biofabric.util.UiUtil;

/****************************************************************************
**
** The top-level BioFabric Application
*/

public class BioFabricApplication {

  ////////////////////////////////////////////////////////////////////////////
  //
  // PUBLIC STATIC CONSTANTS
  //
  //////////////////////////////////////////////////////////////////////////// 
  
  ////////////////////////////////////////////////////////////////////////////
  //
  // PRIVATE VARIABLES
  //
  ////////////////////////////////////////////////////////////////////////////  
 
  private boolean forCyto_;
  private BioFabricWindow bfw_;
  private BioFabricWindow selectionWindow_;
  private PlugInManager plum_;
  
  ////////////////////////////////////////////////////////////////////////////
  //
  // PUBLIC CONSTRUCTORS
  //
  ////////////////////////////////////////////////////////////////////////////    
 
  /***************************************************************************
  **
  ** Now supporting Cytoscape App usage
  */
  
  public BioFabricApplication(boolean forCyto, Map<String, Object> args) { 
    forCyto_ = forCyto;
    ResourceManager.initManager("org.systemsbiology.biofabric.props.BioFabric");
    plum_ = new PlugInManager();
    boolean ok = plum_.loadPlugIns(args);   
    if (!ok) {
      System.err.println("Problems loading plugins");
    }
  }
  
  ////////////////////////////////////////////////////////////////////////////
  //
  // PUBLIC METHODS
  //
  ////////////////////////////////////////////////////////////////////////////
    
  /***************************************************************************
  **
  ** Shutdown operations
  */
  
  public void shutdownFabric() {
    bfw_.stopBufferBuilding();
    bfw_.getWindow().dispose();
    if (selectionWindow_ != null) {
      selectionWindow_.getWindow().dispose();
    }
    if (!forCyto_) {
      System.exit(0);
    }
  }
   
  /***************************************************************************
  **
  ** Close selection window
  */
  
  public void closeSelection() {
    bfw_.getFabricPanel().setSelectionPanel(null);
    selectionWindow_.getWindow().setVisible(false);
    return;
  }
        
  /***************************************************************************
  **
  ** Launch selection window 
  */
     
  public BioFabricWindow launchSelection() {
    bfw_.getFabricPanel().setSelectionPanel(selectionWindow_.getFabricPanel());
    selectionWindow_.getWindow().setSize((int)(bfw_.getWindow().getWidth() * .80), 
                                         (int)(bfw_.getWindow().getHeight() * .80));
    selectionWindow_.getWindow().setVisible(true);
    raiseSelection();
   // selectionWindow_.show();
    return (selectionWindow_);
  }
  
 /***************************************************************************
  **
  ** Raise existing selection window 
  */
     
  public void raiseSelection() {
    selectionWindow_.getWindow().setExtendedState(JFrame.NORMAL);
    selectionWindow_.getWindow().toFront();
    return;
  }

  /***************************************************************************
  **
  ** Launch operations. Now public to support Cytoscape App usage
  */
   
  public BioFabricWindow launch(Map<String, Object> args) { 
    boolean isAMac = System.getProperty("os.name").toLowerCase().startsWith("mac os x");
    if (isAMac) {
      String verNum = System.getProperty("java.version").toLowerCase();
      if ((verNum.indexOf("1.4") == 0) || (verNum.indexOf("1.5") == 0)) {
        UpdateJavaDialog ujw = new UpdateJavaDialog();
        ujw.setVisible(true);
        if (!ujw.keepGoingAnyway()) {
          return (null);
        } 
      }
    }

    bfw_ = new BioFabricWindow(args, this, true, false);
    ExceptionHandler.getHandler().initialize(bfw_.getWindow());
    Dimension cbf = UiUtil.centerBigFrame(bfw_.getWindow(), 1600, 1200, 1.0, 0);
    bfw_.getWindow().setDefaultCloseOperation(WindowConstants.DO_NOTHING_ON_CLOSE);   
    bfw_.getWindow().addWindowListener(new WindowAdapter() {
      public void windowClosing(WindowEvent e) {
        shutdownFabric();
      }
    });
    CommandSet.initCmds("mainWindow", this, bfw_, true, plum_, null);
    bfw_.initWindow(cbf);
    bfw_.getWindow().setVisible(true);
    initSelection();  
    return (bfw_);
  }
    
  ////////////////////////////////////////////////////////////////////////////
  //
  // PUBLIC STATIC METHODS
  //
  ////////////////////////////////////////////////////////////////////////////
  
  /***************************************************************************
  ** 
  ** Main entry point
  */

  public static void main(String argv[]) {
    ArgParser ap = new ArgParser();
    final Map<String, Object> argMap = ap.parse(ArgParser.AppType.VIEWER, argv);
    if (argMap == null) {
      System.err.print(ap.getUsage(ArgParser.AppType.VIEWER));
      System.exit(0);
    }

    final BioFabricApplication su = new BioFabricApplication(false, argMap);    
    SwingUtilities.invokeLater(new Runnable() {
      public void run() {    
        su.launch(argMap);
      }
    });
  }
  
  ////////////////////////////////////////////////////////////////////////////
  //
  // PRIVATE METHODS
  //
  ////////////////////////////////////////////////////////////////////////////    
  
  /***************************************************************************
  **
  ** init selection window 
  */
     
  private void initSelection() {  
    selectionWindow_ = new BioFabricWindow(new HashMap<String, Object>(), this, false, false);
    Dimension swDim = new Dimension((int)(bfw_.getWindow().getWidth() * .80), (int)(bfw_.getWindow().getHeight() * .80));
    selectionWindow_.getWindow().setSize(swDim.width, swDim.height);
    selectionWindow_.getWindow().setLocationRelativeTo(bfw_.getWindow());
    selectionWindow_.getWindow().setDefaultCloseOperation(WindowConstants.DO_NOTHING_ON_CLOSE);   
    selectionWindow_.getWindow().addWindowListener(new WindowAdapter() {
      public void windowClosing(WindowEvent e) {
        closeSelection();
      }
    });
    CommandSet.initCmds("selectionWindow", this, selectionWindow_, false, plum_, null);
    selectionWindow_.initWindow(swDim);
    return;
  }
}
```
