## Supplementary material for "BioFabric Visualization of Network Alignments": BioFabric Source Code: BioFabricWindow.java

```
/*
**    Copyright (C) 2003-2018 Institute for Systems Biology 
**                            Seattle, Washington, USA. 
**
**    This library is free software; you can redistribute it and/or
**    modify it under the terms of the GNU Lesser General Public
**    License as published by the Free Software Foundation; either
**    version 2.1 of the License, or (at your option) any later version.
**
**    This library is distributed in the hope that it will be useful,
**    but WITHOUT ANY WARRANTY; without even the implied warranty of
**    MERCHANTABILITY or FITNESS FOR A PARTICULAR PURPOSE.  See the GNU
**    Lesser General Public License for more details.
**
**    You should have received a copy of the GNU Lesser General Public
**    License along with this library; if not, write to the Free Software
**    Foundation, Inc., 59 Temple Place, Suite 330, Boston, MA  02111-1307  USA
*/

package org.systemsbiology.biofabric.app;

import java.awt.BorderLayout;
import java.awt.CardLayout;
import java.awt.Color;
import java.awt.Dimension;
import java.awt.event.ActionEvent;
import java.net.URL;
import java.util.HashMap;
import java.util.List;
import java.util.Map;
import javax.swing.AbstractAction;
import javax.swing.Action;
import javax.swing.ImageIcon;
import javax.swing.JCheckBoxMenuItem;
import javax.swing.JComponent;
import javax.swing.JFrame;
import javax.swing.JMenu;
import javax.swing.JMenuBar;
import javax.swing.JPanel;
import javax.swing.JScrollPane;
import javax.swing.JSeparator;
import javax.swing.JSplitPane;
import javax.swing.JToolBar;
import javax.swing.KeyStroke;

import org.systemsbiology.biofabric.api.util.ExceptionHandler;
import org.systemsbiology.biofabric.api.worker.BackgroundWorkerControlManager;
import org.systemsbiology.biofabric.cmd.CommandSet;
import org.systemsbiology.biofabric.plugin.BioFabricToolPlugIn;
import org.systemsbiology.biofabric.ui.display.BioFabricNavAndControl;
import org.systemsbiology.biofabric.ui.display.BioFabricOverview;
import org.systemsbiology.biofabric.ui.display.BioFabricPanel;
import org.systemsbiology.biofabric.ui.display.FabricMagnifyingTool;
import org.systemsbiology.biofabric.ui.render.BucketRenderer;
import org.systemsbiology.biofabric.util.ResourceManager;

/****************************************************************************
**
** This is the BioFabric Window!
*/

public class BioFabricWindow implements BackgroundWorkerControlManager {
  
  ////////////////////////////////////////////////////////////////////////////
  //
  // PRIVATE INSTANCE MEMBERS
  //
  //////////////////////////////////////////////////////////////////////////// 
   
  private MyTopWindow myWindow_;
  private BioFabricPanel cp_;
  private BioFabricApplication bfa_;
  private FabricMagnifyingTool fmt_;
  private HashMap<Integer, Action> actionMap_;
  private BioFabricNavAndControl nac_;
  private boolean isMain_;
  private JPanel hidingPanel_;
  private CardLayout myCard_;
  private JSplitPane sp_;
  private double savedSplitFrac_;
  private boolean isHeadless_;
   
  ////////////////////////////////////////////////////////////////////////////
  //
  // PUBLIC CONSTRUCTORS
  //
  ////////////////////////////////////////////////////////////////////////////

  /***************************************************************************
  **
  ** Constructor
  */

  public BioFabricWindow(Map<String, Object> args, BioFabricApplication bfa, 
                         boolean isMain, boolean isHeadless) {
    isHeadless_ = isHeadless;
    if (!isHeadless) {
      myWindow_ = new MyTopWindow(isMain);
    }
    bfa_ = bfa;
    isMain_ = isMain;
    actionMap_ = new HashMap<Integer, Action>();
  }
    
  ////////////////////////////////////////////////////////////////////////////
  //
  // PUBLIC METHODS
  //
  ////////////////////////////////////////////////////////////////////////////

  /***************************************************************************
  ** 
  ** Get the window
  */

  public JFrame getWindow() { 
    return (myWindow_);
  }
 
  /****************************************************************************
  **
  ** disable
  */  
  
  public void disableControls() {
    disableControls(CommandSet.GENERAL_PUSH, true);
    return;
  }
  
  /***************************************************************************
  **
  ** Disable the main controls
  */ 
  
  public void disableControls(int pushFlags, boolean displayToo) {
    CommandSet fc = CommandSet.getCmds((isMain_) ? "mainWindow" : "selectionWindow");
    if (displayToo) {
      myCard_.show(hidingPanel_, "Hiding");
      fmt_.enableControls(false);
      nac_.getOverview().showView(false);
    }
    nac_.getNavTool().enableControls(false);  
    myWindow_.getContentPane().validate();
    fc.pushDisabled(pushFlags);
  }

  /****************************************************************************
  **
  ** enable
  */  
  
  public void reenableControls() {
    CommandSet fc = CommandSet.getCmds((isMain_) ? "mainWindow" : "selectionWindow");
    fc.popDisabled();
    myCard_.show(hidingPanel_, "SUPanel");
    fmt_.enableControls(true);
    nac_.getOverview().showView(true);
    nac_.getNavTool().enableControls(true);
    myWindow_.getContentPane().validate();    
    
    //
    // Following background thread operations, sometimes we need to
    // get keyboard focus back to the network panel:
    //
    // We make this conditional to keep it from being called in normal operation as 
    // the genome is changed, which causes the time slider to lose focus EVERY 
    // TIME IT IS MOVED!!!!!!!
    
  //  if (withFocus) {
   //   sup_.requestFocus();
  //  }
    
    return;
  } 
  
  /****************************************************************************
  **
  ** also redraw....
  */  
  
  public void redraw() {
    cp_.repaint();
    return;
  } 
    
  /***************************************************************************
  **
  ** Get it up and running
  */

  public void initWindow(Dimension dim) {
    CommandSet fc = CommandSet.getCmds((isMain_) ? "mainWindow" : "selectionWindow");
    
    JPanel cpane = null;
    JToolBar toolBar = null; 
    if (!isHeadless_) {
      cpane = (JPanel)myWindow_.getContentPane();
      ((JComponent)cpane).getInputMap(JComponent.WHEN_ANCESTOR_OF_FOCUSED_COMPONENT).put(KeyStroke.getKeyStroke("ESCAPE"), "BioTapCancel");
      ((JComponent)cpane).getActionMap().put("BioTapCancel", new AbstractAction() {
        private static final long serialVersionUID = 1L;
        public void actionPerformed(ActionEvent e) {
          try {
            AbstractAction aa = (AbstractAction)actionMap_.get(Integer.valueOf(CommandSet.CANCEL));
            aa.actionPerformed(null);
            return;
          } catch (Exception ex) {
            ExceptionHandler.getHandler().displayException(ex);
          }
        }
      });        
        
      menuInstall(fc, isMain_);
      toolBar = new JToolBar();
      stockActionMap(fc, isMain_);
      stockToolBar(toolBar, isMain_, fc);
    }
    nac_ = new BioFabricNavAndControl(isMain_, myWindow_, isHeadless_);
    fmt_ = nac_.getFMT();
    BucketRenderer bucketRend = new BucketRenderer(fc.getColorGenerator());
    cp_ = new BioFabricPanel(fc.getColorGenerator(), bfa_, fmt_, 
                             nac_.getOverview(), nac_.getNavTool(), isMain_, this, bucketRend, isHeadless_);  
    fc.setFabricPanel(cp_);
    nac_.setFabricPanel(cp_);
    cp_.setFabricLocation(nac_.getFabricLocation(), nac_.getMouseOverView());
    if (isHeadless_) {
      return;
    }
    
    cp_.getPanel().setBackground(Color.white);
    
    JScrollPane jsp = new JScrollPane(cp_.getPanel());
    
    //
    // "Zoom to full model" has problems if scrollbars come and go, since we check viewport
    // size to calculate the zoom:
    //
    jsp.setHorizontalScrollBarPolicy(JScrollPane.HORIZONTAL_SCROLLBAR_ALWAYS);
    jsp.setVerticalScrollBarPolicy(JScrollPane.VERTICAL_SCROLLBAR_ALWAYS);
    cp_.setScroll(jsp);
    // GOTTA USE THIS ON MY LINUX BOX, BUT NOWHERE ELSE!!!!
    //jsp.getViewport().setScrollMode(JViewport.BACKINGSTORE_SCROLL_MODE);
    cp_.getZoomController().registerScrollPaneAndZoomTarget(jsp, cp_);
     
    cpane.setLayout(new BorderLayout());
    
    if (toolBar != null) {
      cpane.add(toolBar, BorderLayout.NORTH);
    }
        
    hidingPanel_ = new JPanel();
    myCard_ = new CardLayout();
    hidingPanel_.setLayout(myCard_);
    hidingPanel_.add(jsp, "SUPanel");
    JPanel blankPanel = new JPanel();
    blankPanel.setBackground(Color.white);
    hidingPanel_.add(blankPanel, "Hiding");

    sp_ = new JSplitPane(JSplitPane.VERTICAL_SPLIT, hidingPanel_, nac_.getPanel());
    sp_.setDividerLocation((int)(dim.height * 0.50));
    sp_.setResizeWeight(1.0);

    cpane.add(sp_, BorderLayout.CENTER);    
   // cpane.add(nac_, BorderLayout.SOUTH);
           
    URL ugif = getClass().getResource("/org/systemsbiology/biofabric/images/BioFab16White.gif");  
    myWindow_.setIconImage(new ImageIcon(ugif).getImage());
    myWindow_.setResizable(true);
    fc.checkForChanges();
    // Zoom buttons need to be inactive with no model:
    fc.handleZoomButtons();
    return;
  } 

 /***************************************************************************
  **
  ** Drawing core
  */
  
  public void stopBufferBuilding() {
    cp_.shutdown();
    return;
  }
    
  /***************************************************************************
  **
  ** Get fabric panel
  */
  
  public BioFabricPanel getFabricPanel() {
    return (cp_);
  }
  
  /***************************************************************************
  **
  ** Get overvoew panel
  */
  
  public BioFabricOverview getOverview() {
    return (nac_.getOverview());
  }
  
  /***************************************************************************
  **
  ** Hide/show nav and controls
  */
  
  public void showNavAndControl(boolean show) {
    if (isHeadless_) {
      return;
    }
    if (show) {
      sp_.setEnabled(true);
      nac_.setToBlank(!show);
      sp_.setDividerLocation(savedSplitFrac_);
    } else {
      nac_.setToBlank(!show);
      int lastLoc = sp_.getDividerLocation();
      savedSplitFrac_ = (double)lastLoc / (double)sp_.getHeight();
      sp_.setDividerLocation(1.0);
      sp_.setEnabled(false);
    }
    return;
  }
  
  /***************************************************************************
  **
  ** Hide/show nav and controls
  */
  
  public void showTour(boolean show) {
    if (isHeadless_) {
      return;
    }
    if (nac_.showTour(show)) {
      sp_.resetToPreferredSizes();
    }
    return;
  }

  /***************************************************************************
  **
  ** Menu install
  */
  
  private void menuInstall(CommandSet fc, boolean isMain) {
    ResourceManager rMan = ResourceManager.getManager();
    JMenuBar menuBar = new JMenuBar();

    if (isMain) {
      JMenu fMenu = new JMenu(rMan.getString("command.File"));
      fMenu.setMnemonic(rMan.getChar("command.FileMnem"));
      menuBar.add(fMenu);
      fMenu.add(fc.getAction(CommandSet.LOAD_XML, false, null));
      fMenu.add(fc.getAction(CommandSet.SAVE, false, null));
      fMenu.add(fc.getAction(CommandSet.SAVE_AS, false, null));
      fMenu.add(new JSeparator());
      JMenu importMenu = new JMenu(rMan.getString("command.importMenu"));
      importMenu.setMnemonic(rMan.getChar("command.importMenuMnem"));
      fMenu.add(importMenu);    
      importMenu.add(fc.getAction(CommandSet.LOAD_FROM_SIF, false, null));
      importMenu.add(fc.getAction(CommandSet.LOAD_WITH_NODE_ATTRIBUTES, false, null));       
      //importMenu.add(fc.getAction(CommandSet.LOAD_WITH_EDGE_WEIGHTS, false, null)); Not supported in V2
      importMenu.add(fc.getAction(CommandSet.LOAD_FROM_GW, false, null));
  
     // JMenu netAlignMenu = new JMenu(rMan.getString("command.LoadNetAlign"));
     // netAlignMenu.add(fc.getAction(CommandSet.LOAD_NET_ALIGN_GROUPS, false, null));
      //netAlignMenu.add(fc.getAction(CommandSet.LOAD_NET_ALIGN_ORPHAN_EDGES, false, null));
     //importMenu.add(netAlignMenu);
      
      JMenu exportMenu = new JMenu(rMan.getString("command.exportMenu"));
      exportMenu.setMnemonic(rMan.getChar("command.exportMenuMnem"));
      fMenu.add(exportMenu);    
      exportMenu.add(fc.getAction(CommandSet.EXPORT_IMAGE, false, null)); 
      exportMenu.add(fc.getAction(CommandSet.EXPORT_IMAGE_PUBLISH, false, null));      
      exportMenu.add(new JSeparator());
      exportMenu.add(fc.getAction(CommandSet.EXPORT_NODE_ORDER, false, null)); 
      exportMenu.add(fc.getAction(CommandSet.EXPORT_LINK_ORDER, false, null)); 
      exportMenu.add(new JSeparator()); 
      exportMenu.add(fc.getAction(CommandSet.EXPORT_SELECTED_NODES, false, null));       
      fMenu.add(new JSeparator());
      fMenu.add(fc.getAction(CommandSet.EMPTY_NETWORK, false, null));
      fMenu.add(new JSeparator());
      fMenu.add(fc.getAction(CommandSet.PRINT, false, null));
      fMenu.add(new JSeparator());
      fMenu.add(fc.getAction(CommandSet.SET_PLUGIN_DIR, false, null));
      fMenu.add(new JSeparator());
      // Not supporting yet...
      //fMenu.add(fc.getAction(CommandSet.PRINT_PDF, false, null));
      //fMenu.add(new JSeparator());
      fMenu.add(fc.getAction(CommandSet.CLOSE, false, null));
    } else {
      JMenu fMenu = new JMenu(rMan.getString("command.File"));
      fMenu.setMnemonic(rMan.getChar("command.FileMnem"));
      menuBar.add(fMenu);
      JMenu exportMenu = new JMenu(rMan.getString("command.exportMenu"));
      exportMenu.setMnemonic(rMan.getChar("command.exportMenuMnem"));
      fMenu.add(exportMenu);    
      exportMenu.add(fc.getAction(CommandSet.EXPORT_IMAGE, false, null)); 
      exportMenu.add(fc.getAction(CommandSet.EXPORT_IMAGE_PUBLISH, false, null));
      exportMenu.add(new JSeparator());
      exportMenu.add(fc.getAction(CommandSet.EXPORT_SELECTED_NODES, false, null));       
    }
    
    JMenu eMenu = new JMenu(rMan.getString("command.Edit"));
    eMenu.setMnemonic(rMan.getChar("command.EditMnem"));
    menuBar.add(eMenu);
    eMenu.add(fc.getAction(CommandSet.CLEAR_SELECTIONS, false, null));    
    eMenu.add(fc.getAction(CommandSet.ADD_FIRST_NEIGHBORS, false, null));
    if (isMain) {
      eMenu.add(fc.getAction(CommandSet.PROPAGATE_DOWN, false, null));
    }
    Action bsa = fc.getAction(CommandSet.BUILD_SELECT, false, null);
    JCheckBoxMenuItem jcb = new JCheckBoxMenuItem(bsa);
    jcb.setSelected(true);
    eMenu.add(jcb);
    eMenu.add(new JSeparator());    
    eMenu.add(fc.getAction(CommandSet.SET_DISPLAY_OPTIONS, false, null));

    JMenu vMenu = new JMenu(rMan.getString("command.View"));
    vMenu.setMnemonic(rMan.getChar("command.ViewMnem"));
    menuBar.add(vMenu);
    vMenu.add(fc.getAction(CommandSet.ZOOM_OUT, false, null));    
    vMenu.add(fc.getAction(CommandSet.ZOOM_IN, false, null));
    vMenu.add(fc.getAction(CommandSet.ZOOM_TO_MODEL, false, null)); 
    vMenu.add(fc.getAction(CommandSet.ZOOM_TO_RECT, false, null));   
    vMenu.add(fc.getAction(CommandSet.ZOOM_TO_CURRENT_MOUSE, false, null));
    vMenu.add(fc.getAction(CommandSet.ZOOM_TO_CURRENT_MAGNIFY, false, null));
    vMenu.add(fc.getAction(CommandSet.ZOOM_TO_SELECTIONS, false, null));
    vMenu.add(new JSeparator());    
    vMenu.add(fc.getAction(CommandSet.CENTER_ON_PREVIOUS_SELECTION, false, null));
    vMenu.add(fc.getAction(CommandSet.ZOOM_TO_CURRENT_SELECTION, false, null));
    vMenu.add(fc.getAction(CommandSet.CENTER_ON_NEXT_SELECTION, false, null));
    
    //
    // Tools Menu
    //
    
    JMenu sMenu = new JMenu(rMan.getString("command.Tools"));
    sMenu.setMnemonic(rMan.getChar("command.ToolsMnem"));
    menuBar.add(sMenu);
    sMenu.add(fc.getAction(CommandSet.SEARCH, false, null));
    sMenu.add(fc.getAction(CommandSet.COMPARE_NODES, false, null));
    
    
     List<String> piks = fc.getPlugInKeys();
     for (String pik : piks) {
       BioFabricToolPlugIn tpi = fc.getPlugIn(pik);
       JMenu plugMenu = new JMenu(tpi.getToolMenu());
       sMenu.add(plugMenu);
       int numAction = tpi.getCommandCount();
       for (int i = 0; i < numAction; i++) {
         plugMenu.add(fc.getPluginAction(pik, i));
       }
     }
   
    //
    // Layout Menu
    //
    
    JMenu lMenu = new JMenu(rMan.getString("command.Layout"));
    lMenu.setMnemonic(rMan.getChar("command.LayoutMnem"));
    menuBar.add(lMenu);
    lMenu.add(fc.getAction(CommandSet.DEFAULT_LAYOUT, false, null));
    lMenu.add(new JSeparator());
    lMenu.add(fc.getAction(CommandSet.LAYOUT_TOP_CONTROL, false, null)); 
    lMenu.add(fc.getAction(CommandSet.HIER_DAG_LAYOUT, false, null));
    lMenu.add(fc.getAction(CommandSet.LAYOUT_VIA_NODE_CLUSTER_ASSIGN, false, null));   
    lMenu.add(fc.getAction(CommandSet.SET_LAYOUT, false, null)); 
    lMenu.add(fc.getAction(CommandSet.WORLD_BANK_LAYOUT, false, null)); 
    lMenu.add(new JSeparator());
    lMenu.add(fc.getAction(CommandSet.RELAYOUT_USING_CONNECTIVITY, false, null));
    lMenu.add(fc.getAction(CommandSet.RELAYOUT_USING_SHAPE_MATCH, false, null));
    lMenu.add(new JSeparator());
    lMenu.add(fc.getAction(CommandSet.LAYOUT_NODES_VIA_ATTRIBUTES, false, null));
    lMenu.add(fc.getAction(CommandSet.LAYOUT_LINKS_VIA_ATTRIBUTES, false, null));
    lMenu.add(new JSeparator());
    lMenu.add(fc.getAction(CommandSet.SET_LINK_GROUPS, false, null));
    lMenu.add(new JSeparator());
    lMenu.add(fc.getAction(CommandSet.ADD_NODE_ANNOTATIONS, false, null));
    lMenu.add(fc.getAction(CommandSet.ADD_LINK_ANNOTATIONS, false, null));
    
    //
    // Windows Menu
    //
    
    JMenu wMenu = new JMenu(rMan.getString("command.Windows"));
    wMenu.setMnemonic(rMan.getChar("command.ToolsMnem"));
    menuBar.add(wMenu);
    JCheckBoxMenuItem jcbS = new JCheckBoxMenuItem(fc.getAction(CommandSet.SHOW_NAV_PANEL, false, null));
    jcbS.setSelected(true);
    wMenu.add(jcbS);
    JCheckBoxMenuItem jcbT = new JCheckBoxMenuItem(fc.getAction(CommandSet.SHOW_TOUR, false, null));
    jcbT.setSelected(true);
    wMenu.add(jcbT);
    
    JMenu hMenu = new JMenu(rMan.getString("command.Help"));
    hMenu.setMnemonic(rMan.getChar("command.HelpMnem"));
    menuBar.add(hMenu);
    hMenu.add(fc.getAction(CommandSet.ABOUT, false, null));
    
    myWindow_.setJMenuBar(menuBar);
    return;
  }
    
  /***************************************************************************
  **
  ** Stock the action map
  */ 
  
  private void stockActionMap(CommandSet fc, boolean isMain) {  
    actionMap_.put(Integer.valueOf(CommandSet.SEARCH), fc.getAction(CommandSet.SEARCH, true, null));
    actionMap_.put(Integer.valueOf(CommandSet.ZOOM_OUT), fc.getAction(CommandSet.ZOOM_OUT, true, null));
    actionMap_.put(Integer.valueOf(CommandSet.ZOOM_IN), fc.getAction(CommandSet.ZOOM_IN, true, null));
    actionMap_.put(Integer.valueOf(CommandSet.ADD_FIRST_NEIGHBORS), fc.getAction(CommandSet.ADD_FIRST_NEIGHBORS, true, null));
    actionMap_.put(Integer.valueOf(CommandSet.CLEAR_SELECTIONS), fc.getAction(CommandSet.CLEAR_SELECTIONS, true, null));
    actionMap_.put(Integer.valueOf(CommandSet.ZOOM_TO_MODEL), fc.getAction(CommandSet.ZOOM_TO_MODEL, true, null));
    actionMap_.put(Integer.valueOf(CommandSet.ZOOM_TO_RECT), fc.getAction(CommandSet.ZOOM_TO_RECT, true, null));
    actionMap_.put(Integer.valueOf(CommandSet.TOGGLE_SHADOW_LINKS), fc.getAction(CommandSet.TOGGLE_SHADOW_LINKS, true, null));
    actionMap_.put(Integer.valueOf(CommandSet.CANCEL), fc.getAction(CommandSet.CANCEL, true, null));
    actionMap_.put(Integer.valueOf(CommandSet.ZOOM_TO_CURRENT_SELECTION), fc.getAction(CommandSet.ZOOM_TO_CURRENT_SELECTION, true, null));
    actionMap_.put(Integer.valueOf(CommandSet.CENTER_ON_NEXT_SELECTION), fc.getAction(CommandSet.CENTER_ON_NEXT_SELECTION, true, null));
    actionMap_.put(Integer.valueOf(CommandSet.CENTER_ON_PREVIOUS_SELECTION), fc.getAction(CommandSet.CENTER_ON_PREVIOUS_SELECTION, true, null));
    
    if (isMain) {
      actionMap_.put(Integer.valueOf(CommandSet.PROPAGATE_DOWN), fc.getAction(CommandSet.PROPAGATE_DOWN, true, null));
    }
    return;
  }

  /***************************************************************************
  **
  ** Stock the tool bar
  */ 
  
  private void stockToolBar(JToolBar toolBar, boolean isMain, CommandSet fc) {
    toolBar.removeAll();  
    toolBar.add(actionMap_.get(Integer.valueOf(CommandSet.ZOOM_OUT)));
    toolBar.add(actionMap_.get(Integer.valueOf(CommandSet.ZOOM_IN)));
    toolBar.add(actionMap_.get(Integer.valueOf(CommandSet.ZOOM_TO_MODEL)));
    toolBar.add(actionMap_.get(Integer.valueOf(CommandSet.ZOOM_TO_RECT)));
    toolBar.addSeparator();
    toolBar.add(actionMap_.get(Integer.valueOf(CommandSet.CENTER_ON_PREVIOUS_SELECTION)));
    toolBar.add(actionMap_.get(Integer.valueOf(CommandSet.ZOOM_TO_CURRENT_SELECTION)));
    toolBar.add(actionMap_.get(Integer.valueOf(CommandSet.CENTER_ON_NEXT_SELECTION)));
    toolBar.addSeparator();        
    toolBar.add(actionMap_.get(Integer.valueOf(CommandSet.ADD_FIRST_NEIGHBORS)));
    toolBar.add(actionMap_.get(Integer.valueOf(CommandSet.CLEAR_SELECTIONS)));
    toolBar.addSeparator();        
    toolBar.add(actionMap_.get(Integer.valueOf(CommandSet.CANCEL)));
    toolBar.addSeparator();    
    toolBar.add(actionMap_.get(Integer.valueOf(CommandSet.SEARCH)));
    toolBar.addSeparator();    
    toolBar.add(actionMap_.get(Integer.valueOf(CommandSet.TOGGLE_SHADOW_LINKS)));
    if (isMain) {
      toolBar.addSeparator();  
      toolBar.add(actionMap_.get(Integer.valueOf(CommandSet.PROPAGATE_DOWN)));
    }
    return;
  }
  
  /***************************************************************************
  **
  ** Now the actual frame to use
  */  
      
  public class MyTopWindow extends JFrame {
    
    private static final long serialVersionUID = 1L;
    
    /***************************************************************************
    **
    ** Constructor
    */

    public MyTopWindow(boolean isMain) {
      super((isMain) ? "BioFabric" : "BioFabric: Selected Submodel View");
    }
  }
}
```
