## Supplementary material for "BioFabric Visualization of Network Alignments": BioFabric Source Code: ImageGeneratorApplication.java

```
/*
**    Copyright (C) 2003-2018 Institute for Systems Biology 
**                            Seattle, Washington, USA. 
**
**    This library is free software; you can redistribute it and/or
**    modify it under the terms of the GNU Lesser General Public
**    License as published by the Free Software Foundation; either
**    version 2.1 of the License, or (at your option) any later version.
**
**    This library is distributed in the hope that it will be useful,
**    but WITHOUT ANY WARRANTY; without even the implied warranty of
**    MERCHANTABILITY or FITNESS FOR A PARTICULAR PURPOSE.  See the GNU
**    Lesser General Public License for more details.
**
**    You should have received a copy of the GNU Lesser General Public
**    License along with this library; if not, write to the Free Software
**    Foundation, Inc., 59 Temple Place, Suite 330, Boston, MA  02111-1307  USA
*/

package org.systemsbiology.biofabric.app;

import java.awt.Dimension;
import java.awt.Rectangle;
import java.io.File;
import java.util.List;
import java.util.Map;

import org.systemsbiology.biofabric.api.util.ExceptionHandler;
import org.systemsbiology.biofabric.cmd.CommandSet;
import org.systemsbiology.biofabric.cmd.HeadlessOracle;
import org.systemsbiology.biofabric.plugin.PlugInManager;
import org.systemsbiology.biofabric.ui.ImageExporter;
import org.systemsbiology.biofabric.ui.dialogs.ExportSettingsDialog;
import org.systemsbiology.biofabric.ui.display.BioFabricPanel;
import org.systemsbiology.biofabric.util.ResourceManager;

/****************************************************************************
**
** The top-level application or in-process entry point for headless image generation
*/

public class ImageGeneratorApplication {
 
  ////////////////////////////////////////////////////////////////////////////
  //
  // PUBLIC STATIC CONSTANTS
  //
  ////////////////////////////////////////////////////////////////////////////   
  
  /****************************************************************************
  **
  ** These are the allowed input and output options:
  */  
   
  public final static int SIF_INPUT  = 0;
  
  public final static int PNG_OUTPUT = 0;
  
  ////////////////////////////////////////////////////////////////////////////
  //
  // PRIVATE MEMBERS
  //
  ////////////////////////////////////////////////////////////////////////////   
  
  private Map<String, Object> args_;
  
  ////////////////////////////////////////////////////////////////////////////
  //
  // PUBLIC CONSTRUCTORS
  //
  ////////////////////////////////////////////////////////////////////////////    
  
  ////////////////////////////////////////////////////////////////////////////
  //
  // PUBLIC METHODS
  //
  ////////////////////////////////////////////////////////////////////////////
  
  ////////////////////////////////////////////////////////////////////////////
  //
  // PRIVATE VARIABLES
  //
  ////////////////////////////////////////////////////////////////////////////
    
  ////////////////////////////////////////////////////////////////////////////
  //
  // PUBLIC STATIC METHODS
  //
  ////////////////////////////////////////////////////////////////////////////
  
  /***************************************************************************
  ** 
  ** Main entry point for running as a command-line argument
  */

  public static void main(String argv[]) {
    ResourceManager.initManager("org.systemsbiology.biofabric.props.BioFabric");
    ArgParser ap = new ArgParser(); 
    Map<String, Object> argMap = ap.parse(ArgParser.AppType.PIPELINE, argv);
    if ((argMap == null) || (argMap.size() != 3)) {
      System.err.print(ap.getUsage(ArgParser.AppType.PIPELINE));
      System.exit(1);
    }
    ImageGeneratorApplication iga = new ImageGeneratorApplication(argMap);
    try {
      String errMsg = iga.generate();
      if (errMsg != null) {
        System.err.println(errMsg);
        System.exit(1);
      }
    } catch (GeneratorException gex) {
      System.err.println(gex.getMessage());  
    } catch (MissingArgException maex) {    
      System.err.print(ap.getUsage(ArgParser.AppType.PIPELINE));
      System.exit(1);
    }
    return;
  }

  ////////////////////////////////////////////////////////////////////////////
  //
  // PRIVATE CONSTRUCTOR
  //
  ////////////////////////////////////////////////////////////////////////////  

  /***************************************************************************
  ** 
  ** Used for command-line version
  */   
  
  private ImageGeneratorApplication(Map<String, Object> args) {
    args_ = args;
  }  
 
  ////////////////////////////////////////////////////////////////////////////
  //
  // PRIVATE METHODS
  //
  ////////////////////////////////////////////////////////////////////////////    
      
  /***************************************************************************
  ** 
  ** Handle command-line operation:
  */  
  
  private String generate() throws GeneratorException, MissingArgException {
    
    //
    // At this time, core code must run on one thread when in batch mode.
    //

    synchronized(CommandSet.class) {
      ResourceManager rMan = ResourceManager.getManager();    
      System.setProperty("java.awt.headless", "true");
      BioFabricWindow bfw = new BioFabricWindow(args_, null, true, true);
      ExceptionHandler.getHandler().initializeForHeadless(false);
      PlugInManager plum = new PlugInManager();
      boolean ok = plum.loadPlugIns(args_);   
      if (!ok) {
        System.err.println("Problems loading plugins");
      }    
      CommandSet.initCmds("mainWindow", null, bfw, true, plum, new HeadlessOracle());
      bfw.initWindow(null); 
      CommandSet cmd = CommandSet.getCmds("mainWindow");
           
      //
      // Currently Hardwired:
      //
      
      int inputType = SIF_INPUT;
      int outputType = PNG_OUTPUT;
  
      //
      // Input operations.  Gotta have one and only one
      //

      boolean haveInput = false;

      switch (inputType) {
        case SIF_INPUT:          
          String sifFileName = (String)args_.get(ArgParser.SIF_BATCH_INPUT);
          if (sifFileName == null) {
            throw new MissingArgException();
          }
          File sifFile = new File(sifFileName);
          CommandSet.HeadlessImportSIFAction sifLoader = cmd.new HeadlessImportSIFAction(sifFile);
          Object[] osArgs = null;
          boolean okl = sifLoader.performOperation(osArgs);
          if (!okl) {
            throw new GeneratorException(rMan.getString("headless.csvInputFailure"));
          }
          haveInput = true;
          break;
        default:
          throw new IllegalArgumentException();
      }   
      if (!haveInput) {
        throw new GeneratorException(rMan.getString("headless.noInputFailure"));
      }   

      boolean aSuccess = false;

      switch (outputType) {
        case PNG_OUTPUT:
          String imageFileName = (String)args_.get(ArgParser.IMAGE_BATCH_OUTPUT);
          if (imageFileName == null) {
            throw new MissingArgException();
          }
          CommandSet.HeadlessExportAction hexa = cmd.new HeadlessExportAction();
          Object[] osArgs = imageExportPrepForFile(args_, imageFileName, cmd);
          if (osArgs != null) {
            boolean hok = hexa.performOperation(osArgs);
            if (!hok) {
              System.err.println(rMan.getString("headless.imageExportFailure"));
            } else {
              aSuccess = true;
            }
          } else {
            System.err.println(rMan.getString("headless.imageExportFailure"));
          }
          break;
        default:
          throw new IllegalArgumentException();
      }
      
      if (!aSuccess) {
        System.err.println(rMan.getString("headless.totalExportFailure"));
        return (rMan.getString("headless.earlyExit"));
      } 
    }
    return (null);
  }
 
  /***************************************************************************
  ** 
  ** Image export argument prep for file output
  */
  
  
  private Object[] imageExportPrepForFile(Map args, String imageFileName, CommandSet cmd) throws MissingArgException{
    Object[] osArgs = new Object[3];
    osArgs[1] = new Boolean(true); // This is a file
    osArgs[2] = imageFileName;         
    ExportSettingsDialog.ExportSettings settings = imageExportPrepGuts(args, cmd);
    osArgs[0] = settings;    
    return (osArgs);
  }  
  
  /***************************************************************************
  ** 
  ** Image export argument prep
  */
   
  private ExportSettingsDialog.ExportSettings imageExportPrepGuts(Map<String, Object> args, 
                                                                  CommandSet cmd) throws MissingArgException {

    ResourceManager rMan = ResourceManager.getManager();
    String outputWidthStr = (String)args_.get(ArgParser.IMAGE_OUTPUT_WIDTH);
    if (outputWidthStr == null) {
      throw new MissingArgException();
    }
    int outputWidth = 0;
    try {
      outputWidth = Integer.parseInt(outputWidthStr);
    } catch (NumberFormatException nfex) {
      System.err.println(rMan.getString("headless.imageExportBadWidth"));
      return (null);           
    }

    List suppRes = ImageExporter.getSupportedResolutions(false);
    List suppForms = ImageExporter.getSupportedExports();    
    
    ExportSettingsDialog.ExportSettings settings = new ExportSettingsDialog.ExportSettings();

    // Make this an argument!
    if (suppForms.contains("PNG")) {
      settings.formatType = "PNG";
    } else {
      System.err.println(rMan.getString("headless.imageExportPrepFailure"));
      return (null);
    }
    
    if (ImageExporter.formatRequiresResolution(settings.formatType)) {
      List resList = ImageExporter.getSupportedResolutions(false);    
      if (resList.size() == 0) {
        throw new IllegalStateException();
      }
      Object[] resVal = (Object[])resList.get(0);
      ImageExporter.RationalNumber ratNum = (ImageExporter.RationalNumber)resVal[ImageExporter.CM];
      settings.res = new ImageExporter.ResolutionSettings();
      settings.res.dotsPerUnit = ratNum;
      settings.res.units = ImageExporter.CM;
    } else {
      settings.res = null;
    }
   
    BioFabricPanel bfp = cmd.getBFW().getFabricPanel();
    Rectangle wr = bfp.getRequiredSize();
    
    settings.zoomVal = (double)outputWidth / (double)wr.width; 

    double currentZoomHeight = Math.round(((double)wr.height) * settings.zoomVal);
    double currentZoomWidth = Math.round(((double)wr.width) * settings.zoomVal);    
    settings.size = new Dimension((int)currentZoomWidth, (int)currentZoomHeight);    
    return (settings);
  }

  ////////////////////////////////////////////////////////////////////////////
  //
  // PUBLIC INNER CLASSES
  //
  ////////////////////////////////////////////////////////////////////////////

  /***************************************************************************
  ** 
  ** Thrown exceptions include a message as well as an optional wrapped
  ** exception from deep down in the program...
  */

  public static class GeneratorException extends Exception {
    
    private Exception wrapped_;
    
    GeneratorException(String message, Exception wrapped) {
      super(message);
      wrapped_ = wrapped;
    }
    
    GeneratorException(String message) {
      super(message);
      wrapped_ = null;
    }
   
    public Exception getWrappedException() {
      return (wrapped_);
    }
  }  
  
  /***************************************************************************
  ** 
  ** Thrown exceptions include a message as well as an optional wrapped
  ** exception from deep down in the program...
  */

  public static class MissingArgException extends Exception {

  }
   
}
```
