## Supplementary material for "BioFabric Visualization of Network Alignments": BioFabric Source Code: CommandSet.java

```
/*
**    Copyright (C) 2003-2018 Institute for Systems Biology 
**                            Seattle, Washington, USA. 
**
**    This library is free software; you can redistribute it and/or
**    modify it under the terms of the GNU Lesser General Public
**    License as published by the Free Software Foundation; either
**    version 2.1 of the License, or (at your option) any later version.
**
**    This library is distributed in the hope that it will be useful,
**    but WITHOUT ANY WARRANTY; without even the implied warranty of
**    MERCHANTABILITY or FITNESS FOR A PARTICULAR PURPOSE.  See the GNU
**    Lesser General Public License for more details.
**
**    You should have received a copy of the GNU Lesser General Public
**    License along with this library; if not, write to the Free Software
**    Foundation, Inc., 59 Temple Place, Suite 330, Boston, MA  02111-1307  USA
*/

package org.systemsbiology.biofabric.cmd;

import java.awt.Color;
import java.awt.Event;
import java.awt.GridBagConstraints;
import java.awt.GridBagLayout;
import java.awt.Point;
import java.awt.Rectangle;
import java.awt.event.ActionEvent;
import java.awt.event.ActionListener;
import java.awt.event.WindowAdapter;
import java.awt.event.WindowEvent;
import java.awt.geom.Point2D;
import java.awt.print.PageFormat;
import java.awt.print.PrinterException;
import java.awt.print.PrinterJob;
import java.io.BufferedWriter;
import java.io.File;
import java.io.FileOutputStream;
import java.io.IOException;
import java.io.OutputStream;
import java.io.OutputStreamWriter;
import java.io.PrintWriter;
import java.net.URL;
import java.text.MessageFormat;
import java.util.ArrayList;
import java.util.HashMap;
import java.util.HashSet;
import java.util.Iterator;
import java.util.List;
import java.util.Map;
import java.util.Set;
import java.util.SortedMap;
import java.util.TreeMap;

import javax.swing.AbstractAction;
import javax.swing.Action;
import javax.swing.Box;
import javax.swing.ImageIcon;
import javax.swing.JEditorPane;
import javax.swing.JFileChooser;
import javax.swing.JFrame;
import javax.swing.JLabel;
import javax.swing.JOptionPane;
import javax.swing.JPanel;
import javax.swing.JScrollPane;
import javax.swing.KeyStroke;
import javax.swing.border.EmptyBorder;
import javax.swing.event.HyperlinkEvent;
import javax.swing.event.HyperlinkListener;
import javax.swing.filechooser.FileFilter;

import org.systemsbiology.biofabric.api.io.AttributeKey;
import org.systemsbiology.biofabric.api.io.FileLoadFlows;
import org.systemsbiology.biofabric.api.layout.DefaultLayout;
import org.systemsbiology.biofabric.api.model.AnnotationSet;
import org.systemsbiology.biofabric.api.model.AugRelation;
import org.systemsbiology.biofabric.api.model.NetLink;
import org.systemsbiology.biofabric.api.model.NetNode;
import org.systemsbiology.biofabric.api.util.ExceptionHandler;
import org.systemsbiology.biofabric.api.util.FixedJButton;
import org.systemsbiology.biofabric.api.util.UniqueLabeller;
import org.systemsbiology.biofabric.api.worker.AsynchExitRequestException;
import org.systemsbiology.biofabric.app.BioFabricApplication;
import org.systemsbiology.biofabric.app.BioFabricWindow;
import org.systemsbiology.biofabric.event.EventManager;
import org.systemsbiology.biofabric.event.SelectionChangeEvent;
import org.systemsbiology.biofabric.event.SelectionChangeListener;
import org.systemsbiology.biofabric.io.BuildDataImpl;
import org.systemsbiology.biofabric.io.FileLoadFlowsImpl;
import org.systemsbiology.biofabric.layouts.ControlTopLayout;
import org.systemsbiology.biofabric.layouts.NodeClusterLayout;
import org.systemsbiology.biofabric.layouts.NodeSimilarityLayout;

import org.systemsbiology.biofabric.model.BioFabricNetwork;
import org.systemsbiology.biofabric.plugin.BioFabricToolPlugIn;
import org.systemsbiology.biofabric.plugin.BioFabricToolPlugInCmd;
import org.systemsbiology.biofabric.plugin.PluginSupportFactory;
import org.systemsbiology.biofabric.plugin.PlugInManager;
import org.systemsbiology.biofabric.ui.FabricColorGenerator;
import org.systemsbiology.biofabric.ui.FabricDisplayOptions;
import org.systemsbiology.biofabric.ui.FabricDisplayOptionsManager;
import org.systemsbiology.biofabric.ui.ImageExporter;
import org.systemsbiology.biofabric.ui.dialogs.BreadthFirstLayoutDialog;
import org.systemsbiology.biofabric.ui.dialogs.ClusterLayoutSetupDialog;
import org.systemsbiology.biofabric.ui.dialogs.CompareNodesSetupDialog;
import org.systemsbiology.biofabric.ui.dialogs.ControlTopLayoutSetupDialog;
import org.systemsbiology.biofabric.ui.dialogs.DirectoryChooserDialog;
import org.systemsbiology.biofabric.ui.dialogs.ExportSettingsDialog;
import org.systemsbiology.biofabric.ui.dialogs.ExportSettingsPublishDialog;
import org.systemsbiology.biofabric.ui.dialogs.FabricDisplayOptionsDialog;
import org.systemsbiology.biofabric.ui.dialogs.FabricSearchDialog;
import org.systemsbiology.biofabric.ui.dialogs.LinkGroupingSetupDialog;
import org.systemsbiology.biofabric.ui.dialogs.NodeSimilarityLayoutSetupDialog;
import org.systemsbiology.biofabric.ui.dialogs.PointUpOrDownDialog;
import org.systemsbiology.biofabric.ui.dialogs.ReorderLayoutParamsDialog;
import org.systemsbiology.biofabric.ui.dialogs.SetRelationSemanticsDialog;
import org.systemsbiology.biofabric.ui.display.BioFabricPanel;
import org.systemsbiology.biofabric.ui.display.FabricMagnifyingTool;
import org.systemsbiology.biofabric.util.FileExtensionFilters;
import org.systemsbiology.biofabric.util.InvalidInputException;
import org.systemsbiology.biofabric.util.ResourceManager;
import org.systemsbiology.biofabric.util.UiUtil;
import org.systemsbiology.biotapestry.biofabric.FabricCommands;

/****************************************************************************
**
** Collection of primary commands for the application
*/

public class CommandSet implements ZoomChangeTracker, SelectionChangeListener, FabricDisplayOptionsManager.DisplayOptionTracker {
 
  ////////////////////////////////////////////////////////////////////////////
  //
  // PUBLIC CONSTANTS
  //
  //////////////////////////////////////////////////////////////////////////// 

  public static final int EMPTY_NETWORK     = 0; 
  public static final int CLOSE             = 1; 
  public static final int LOAD_FROM_SIF = 2;
  public static final int SEARCH            = 3; 
  public static final int ZOOM_OUT          = 4; 
  public static final int ZOOM_IN           = 5; 
  public static final int CLEAR_SELECTIONS  = 6; 
  public static final int SAVE_AS           = 7; 

  public static final int ZOOM_TO_MODEL     = 8; 
  public static final int ZOOM_TO_SELECTIONS = 9;
  public static final int PROPAGATE_DOWN    = 10; 
  public static final int ZOOM_TO_RECT      = 11;
  public static final int CANCEL            = 12;
  public static final int ZOOM_TO_CURRENT_SELECTION = 13;  
  public static final int ADD_FIRST_NEIGHBORS = 14;
  public static final int BUILD_SELECT        = 15;
  public static final int SET_DISPLAY_OPTIONS = 16;
  public static final int SET_LAYOUT          = 17;
  public static final int TOGGLE_SHADOW_LINKS = 18;
  
  // Former Gaggle Commands 17-24 dropped
  
  public static final int ABOUT                  = 25;
  public static final int CENTER_ON_NEXT_SELECTION = 26;  
  public static final int CENTER_ON_PREVIOUS_SELECTION = 27;  
  public static final int LAYOUT_NODES_VIA_ATTRIBUTES  = 28;
  public static final int LAYOUT_LINKS_VIA_ATTRIBUTES  = 29;
  public static final int LOAD_WITH_NODE_ATTRIBUTES = 30;
  public static final int LOAD_XML                  = 31;
  public static final int RELAYOUT_USING_CONNECTIVITY  = 32;
  public static final int RELAYOUT_USING_SHAPE_MATCH   = 33;
  public static final int SET_LINK_GROUPS              = 34;
  public static final int COMPARE_NODES                = 35;
  public static final int ZOOM_TO_CURRENT_MOUSE        = 36;  
  public static final int ZOOM_TO_CURRENT_MAGNIFY      = 37; 
  public static final int EXPORT_NODE_ORDER            = 38; 
  public static final int EXPORT_LINK_ORDER            = 39; 
  public static final int EXPORT_IMAGE                 = 40; 
  public static final int EXPORT_IMAGE_PUBLISH         = 41; 
  public static final int PRINT                        = 42;
  public static final int DEFAULT_LAYOUT               = 43;
  public static final int EXPORT_SELECTED_NODES        = 44;
  public static final int SAVE                         = 45;
  public static final int LAYOUT_VIA_NODE_CLUSTER_ASSIGN = 46;
  public static final int PRINT_PDF                    = 47;
  public static final int SHOW_TOUR                    = 48;
  public static final int SHOW_NAV_PANEL               = 49;
  public static final int LAYOUT_TOP_CONTROL           = 50;
  public static final int HIER_DAG_LAYOUT              = 51;
  public static final int WORLD_BANK_LAYOUT            = 52;
  public static final int LOAD_WITH_EDGE_WEIGHTS       = 53;
  public static final int LOAD_FROM_GW                 = 54;
  
  public static final int ADD_NODE_ANNOTATIONS         = 55;
  public static final int ADD_LINK_ANNOTATIONS         = 56;
  
  public static final int SET_PLUGIN_DIR               = 57;
 
  public static final int GENERAL_PUSH   = 0x01;
  public static final int ALLOW_NAV_PUSH = 0x02;
      
  ////////////////////////////////////////////////////////////////////////////
  //
  // MEMBERS
  //
  ////////////////////////////////////////////////////////////////////////////  

  private static HashMap<String, CommandSet> perClass_;
  private JFrame topWindow_;
  private BioFabricWindow bfw_;
  private BioFabricApplication bfa_;
  private BioFabricPanel bfp_;
  private PlugInManager pMan_;

  private boolean isAMac_;
  private boolean isForMain_;
  private boolean showNav_;
  
  private HashMap<Integer, ChecksForEnabled> withIcons_;
  private HashMap<Integer, ChecksForEnabled> noIcons_;
  private HashMap<String, ChecksForEnabled> plugInCmds_;
  private FabricColorGenerator colGen_;
  private FileLoadFlowsImpl flf_;

  ////////////////////////////////////////////////////////////////////////////
  //
  // PUBLIC METHODS
  //
  ////////////////////////////////////////////////////////////////////////////    
 
  /***************************************************************************
  **
  ** Needed for Cytoscape app support
  */ 
  
  public BioFabricWindow getBFW() {
    return (bfw_);
  }
   
  /***************************************************************************
  **
  ** Needed for refactoring
  */ 
  
  public FileLoadFlows getFileLoader() {
    return (flf_);
  }
   
  /***************************************************************************
  **
  ** Let us know we are on a mac
  */ 
  
  public boolean isAMac() {
    return (isAMac_);
  }
  
  /***************************************************************************
  **
  ** Notify listener of selection change
  */ 
  
  public void selectionHasChanged(SelectionChangeEvent scev) {
    checkForChanges();
    return;
  }
   
  /***************************************************************************
  **
  ** Trigger the enabled checks
  */ 
  
  public void checkForChanges() {
    Iterator<ChecksForEnabled> wiit = withIcons_.values().iterator();
    while (wiit.hasNext()) {
      ChecksForEnabled cfe = wiit.next();
      cfe.checkIfEnabled();
    }
    Iterator<ChecksForEnabled> niit = noIcons_.values().iterator();
    while (niit.hasNext()) {
      ChecksForEnabled cfe = niit.next();
      cfe.checkIfEnabled();
    }
    Iterator<ChecksForEnabled> piit = plugInCmds_.values().iterator();
    while (piit.hasNext()) {
      ChecksForEnabled cfe = piit.next();
      cfe.checkIfEnabled();
    }
    return;
  }
  
  /***************************************************************************
  **
  ** Push a disabled condition
  */ 
  
  public void pushDisabled(int pushCondition) {
    Iterator<ChecksForEnabled> wiit = withIcons_.values().iterator();
    while (wiit.hasNext()) {
      ChecksForEnabled cfe = wiit.next();
      cfe.pushDisabled(pushCondition);
    }
    Iterator<ChecksForEnabled> niit = noIcons_.values().iterator();
    while (niit.hasNext()) {
      ChecksForEnabled cfe = niit.next();
      cfe.pushDisabled(pushCondition);
    }
    Iterator<ChecksForEnabled> piit = plugInCmds_.values().iterator();
    while (piit.hasNext()) {
      ChecksForEnabled cfe = piit.next();
      cfe.pushDisabled(pushCondition);
    }

    return;
  }
  
  /***************************************************************************
  **
  ** Pop the disabled condition
  */ 
  
   public void popDisabled() {
    Iterator<ChecksForEnabled> wiit = withIcons_.values().iterator();
    while (wiit.hasNext()) {
      ChecksForEnabled cfe = wiit.next();
      cfe.popDisabled();
    }
    Iterator<ChecksForEnabled> niit = noIcons_.values().iterator();
    while (niit.hasNext()) {
      ChecksForEnabled cfe = niit.next();
      cfe.popDisabled();
    }
    Iterator<ChecksForEnabled> piit = plugInCmds_.values().iterator();
    while (piit.hasNext()) {
      ChecksForEnabled cfe = piit.next();
      cfe.popDisabled();
    }

    return;
  } 

  /***************************************************************************
  **
  ** Display options have changed!
  */   
  
  public void optionsHaveChanged(boolean needRebuild, boolean needRecolor) {
    if (!needRebuild && !needRecolor) {
      bfp_.repaint();
      return;
    }
    
    // No model? Nothing to do...
    // Fix for issue #89
    if (!bfp_.hasAModel()) {
    	return;
    }
    
    
    if (needRecolor && !needRebuild) {  
      flf_.doRecolor(isForMain_);
    } else if (needRebuild) {
      flf_.doDisplayOptionChange();  
    }
    return;   
  }
    
  /***************************************************************************
  **
  ** Get ColorGenerator
  */ 
  
  public FabricColorGenerator getColorGenerator() {
    return (colGen_);
  }
  
  /***************************************************************************
  **
  ** Get ordered keys for plugins that have commands to offer
  */ 
  
  public List<String> getPlugInKeys() {
    return (pMan_.getOrderedToolPlugInKeys());
  }
  
  /***************************************************************************
  **
  ** Get plugIn
  */ 
  
  public BioFabricToolPlugIn getPlugIn(String key) {
    return (pMan_.getToolPlugIn(key));
  }
  
  /***************************************************************************
  **
  ** Get an action
  */ 
  
  public Action getAction(int actionKey, boolean withIcon, Object[] optionArgs) {
    HashMap<Integer, ChecksForEnabled> useMap = (withIcon) ? withIcons_ : noIcons_;
    Integer actionKeyObject = Integer.valueOf(actionKey);
    ChecksForEnabled retval = useMap.get(actionKeyObject);
    if (retval != null) {
      return (retval);
    } else {
      switch (actionKey) { 
        case ABOUT:
          retval = new AboutAction(withIcon); 
          break;
        case EMPTY_NETWORK:
          retval = new EmptyNetworkAction(withIcon); 
          break;
        case CLOSE:
          retval = new CloseAction(withIcon); 
          break;
        case LOAD_XML:
          retval = new LoadXMLAction(withIcon); 
          break;
        case LOAD_FROM_SIF:
          retval = new ImportSIFAction(withIcon, false); 
          break;
        case LOAD_WITH_EDGE_WEIGHTS:
        	throw new IllegalArgumentException(); // Not supported in V2
          //retval = new ImportSIFAction(withIcon, true); 
          //break;          
        case LOAD_WITH_NODE_ATTRIBUTES:
          retval = new LoadWithNodeAttributesAction(withIcon); 
          break;
        case LOAD_FROM_GW:
          retval = new ImportGWAction(withIcon);
          break;
        case SAVE_AS:
          retval = new SaveAsAction(withIcon); 
          break;
        case SAVE:
          retval = new SaveAction(withIcon); 
          break;
        case EXPORT_NODE_ORDER:
          retval = new ExportNodeOrderAction(withIcon); 
          break;  
        case EXPORT_LINK_ORDER:
          retval = new ExportLinkOrderAction(withIcon); 
          break;  
        case EXPORT_SELECTED_NODES:
          retval = new ExportSelectedNodesAction(withIcon); 
          break;  
        case EXPORT_IMAGE:
          retval = new ExportSimpleAction(withIcon); 
          break;  
        case EXPORT_IMAGE_PUBLISH:
          retval = new ExportPublishAction(withIcon); 
          break;    
        case PRINT:
          retval = new PrintAction(withIcon); 
          break; 
        case PRINT_PDF: // Not yet supported
        	throw new IllegalArgumentException();
          //retval = new PrintPDFAction(withIcon); 
          //break;           
        case SEARCH:
          retval = new SearchAction(withIcon); 
          break;
        case ZOOM_IN:
          retval = new InOutZoomAction(withIcon, '+'); 
          break;
        case ZOOM_OUT:
          retval = new InOutZoomAction(withIcon, '-'); 
          break;
        case CLEAR_SELECTIONS:
          retval = new ClearSelectionsAction(withIcon); 
          break;
        case ZOOM_TO_MODEL:
          retval = new ZoomToModelAction(withIcon); 
          break;
        case ZOOM_TO_SELECTIONS:
          retval = new ZoomToSelected(withIcon); 
          break;  
        case PROPAGATE_DOWN:
          retval = new PropagateDownAction(withIcon); 
          break; 
        case ZOOM_TO_RECT:
          retval = new ZoomToRect(withIcon); 
          break;
        case CANCEL:
          retval = new CancelAction(withIcon); 
          break;
        case CENTER_ON_NEXT_SELECTION:
          retval = new CenterOnNextSelected(withIcon); 
          break; 
        case CENTER_ON_PREVIOUS_SELECTION:
          retval = new CenterOnPreviousSelected(withIcon); 
          break; 
        case ZOOM_TO_CURRENT_SELECTION:
          retval = new ZoomToCurrentSelected(withIcon); 
          break;
        case ZOOM_TO_CURRENT_MOUSE:
          retval = new ZoomToCurrentMouse(withIcon); 
          break;
        case ZOOM_TO_CURRENT_MAGNIFY:
          retval = new ZoomToCurrentMagnify(withIcon); 
          break;  
        case ADD_FIRST_NEIGHBORS:
          retval = new AddFirstNeighborsAction(withIcon); 
          break;          
        case BUILD_SELECT:
          retval = new BuildSelectAction(withIcon); 
          break;  
        case SET_DISPLAY_OPTIONS:
          retval = new SetDisplayOptionsAction(withIcon); 
          break;
        case LAYOUT_NODES_VIA_ATTRIBUTES:
          retval = new LayoutNodesViaAttributesAction(withIcon); 
          break;
        case LAYOUT_LINKS_VIA_ATTRIBUTES:
          retval = new LayoutLinksViaAttributesAction(withIcon); 
          break;
        case RELAYOUT_USING_CONNECTIVITY:
          retval = new LayoutViaConnectivityAction(withIcon); 
          break;
        case DEFAULT_LAYOUT:
          retval = new DefaultLayoutAction(withIcon); 
          break;
        case HIER_DAG_LAYOUT:
          retval = new HierDAGLayoutAction(withIcon); 
          break;
        case SET_LAYOUT:
          retval = new SetLayoutAction(withIcon); 
          break;
        case TOGGLE_SHADOW_LINKS:
          retval = new ToggleShadowLinks(withIcon); 
          break;                  
        case RELAYOUT_USING_SHAPE_MATCH:
          retval = new LayoutViaShapeMatchAction(withIcon); 
          break;  
        case SET_LINK_GROUPS:
          retval = new SetLinkGroupsAction(withIcon);
          break;
        case COMPARE_NODES:
          retval = new CompareNodesAction(withIcon);           
          break;
        case LAYOUT_VIA_NODE_CLUSTER_ASSIGN:
          retval = new LayoutViaNodeClusterAction(withIcon); 
          break;
        case LAYOUT_TOP_CONTROL:
          retval = new LayoutTopControlAction(withIcon); 
          break;          
        case SHOW_TOUR:
          retval = new ToggleShowTourAction(withIcon); 
          break;
        case SHOW_NAV_PANEL:
          retval = new ToggleShowNavPanelAction(withIcon); 
          break;
        case WORLD_BANK_LAYOUT:
          retval = new WorldBankLayoutAction(withIcon); 
          break;
        case ADD_NODE_ANNOTATIONS:
          retval = new AddNodeAnnotations(withIcon); 
          break;
        case ADD_LINK_ANNOTATIONS:
          retval = new AddLinkAnnotations(withIcon); 
          break;
        case SET_PLUGIN_DIR:
          retval = new SetPluginDirectory(withIcon); 
          break;          
          
        default:
          throw new IllegalArgumentException();
      }
      useMap.put(actionKeyObject, retval);
    }
    return (retval);
  }
  
  /***************************************************************************
  **
  ** Get an action
  */ 
    
  public Action getPluginAction(String plugin, int cmd) {
    String key = plugin + ":::" + cmd;
    ChecksForEnabled retval = plugInCmds_.get(key);
    if (retval != null) {
      return (retval);
    } else {
      BioFabricToolPlugIn pi = pMan_.getToolPlugIn(plugin);
      BioFabricToolPlugInCmd pic = pi.getCommand(cmd);
      retval = new PlugInAction(pic);
    }
    plugInCmds_.put(key, retval);
    return (retval);
  } 

  
  
  /***************************************************************************
  **
  ** Displays file reading error message for invalid input
  */ 
       
  public void displayInvalidInputError(InvalidInputException iiex) { 
    ResourceManager rMan = ResourceManager.getManager(); 
    String errKey = iiex.getErrorKey();
    boolean haveKey = (errKey != null) && (!errKey.equals(InvalidInputException.UNSPECIFIED_ERROR)); 
    int lineno = iiex.getErrorLineNumber();
    boolean haveLine = (lineno != InvalidInputException.UNSPECIFIED_LINE);
    String outMsg;
    if (haveKey && haveLine) { 
      String format = rMan.getString("fileRead.inputErrorMessageForLineWithDesc");
      String keyedErr = rMan.getString("invalidInput." + errKey);
      outMsg = MessageFormat.format(format, new Object[] {Integer.valueOf(lineno + 1), keyedErr});
    } else if (haveKey && !haveLine) {
      String format = rMan.getString("fileRead.inputErrorMessageWithDesc");
      String keyedErr = rMan.getString("invalidInput." + errKey);
      outMsg = MessageFormat.format(format, new Object[] {keyedErr});
    } else if (!haveKey && haveLine) {
      String format = rMan.getString("fileRead.inputErrorMessageForLine");
      outMsg = MessageFormat.format(format, new Object[] {Integer.valueOf(lineno + 1)});      
    } else {
      outMsg = rMan.getString("fileRead.inputErrorMessage");      
    } 
    JOptionPane.showMessageDialog(topWindow_, outMsg, 
                                  rMan.getString("fileRead.errorTitle"),
                                  JOptionPane.ERROR_MESSAGE);
  return;
  }
 
  /***************************************************************************
  **
  ** Set the fabric panel
  */ 
  
  public void setFabricPanel(BioFabricPanel bfp) {
    bfp_ = bfp;
    flf_.setFabricPanel(bfp);
    return;
  }  
   
  /***************************************************************************
  **
  ** Tell us the zoom state has changed
  */ 
  
  public void zoomStateChanged(boolean scrollOnly) {
   // if (!scrollOnly) { // Want nav panel resize to check zoom out ability
      handleZoomButtons();
   // }
    bfw_.getOverview().setViewInWorld(bfp_.getViewInWorld());
    return;
  }  
  
  /***************************************************************************
  **
  ** Handle zoom buttons
  */ 

  public void handleZoomButtons() {  
    //
    // Enable/disable zoom actions based on zoom limits:
    //

    InOutZoomAction zaOutWI = (InOutZoomAction)withIcons_.get(Integer.valueOf(ZOOM_OUT));
    InOutZoomAction zaOutNI = (InOutZoomAction)noIcons_.get(Integer.valueOf(ZOOM_OUT));
    InOutZoomAction zaInWI = (InOutZoomAction)withIcons_.get(Integer.valueOf(ZOOM_IN));
    InOutZoomAction zaInNI = (InOutZoomAction)noIcons_.get(Integer.valueOf(ZOOM_IN));
    if (!bfp_.hasAModel()) {
      zaOutWI.setConditionalEnabled(false);
      if (zaOutNI != null) zaOutNI.setConditionalEnabled(false);
      zaInWI.setConditionalEnabled(false);
      if (zaInNI != null) zaInNI.setConditionalEnabled(false);
      return;
    } 
    
    boolean downOn = bfp_.getZoomController().canZoomOut();
    zaOutWI.setConditionalEnabled(downOn);
    if (zaOutNI != null) zaOutNI.setConditionalEnabled(downOn);
 
    boolean upOn = !bfp_.getZoomController().zoomIsMax();
    zaInWI.setConditionalEnabled(upOn);
    if (zaInNI != null) zaInNI.setConditionalEnabled(upOn);
  
    return;
  }

  ////////////////////////////////////////////////////////////////////////////
  //
  // PUBLIC STATIC METHODS
  //
  ////////////////////////////////////////////////////////////////////////////
    
  /***************************************************************************
  ** 
  ** Get the commands for the given tag
  */

  public static synchronized CommandSet getCmds(String className) {
    if (perClass_ == null) {
      throw new IllegalStateException();
    }
    CommandSet fc = perClass_.get(className);
    if (fc == null) {
      throw new IllegalStateException();
    }
    return (fc);
  }  
 
  /***************************************************************************
  ** 
  ** Init the commands for the given tag
  */

  public static synchronized CommandSet initCmds(String className, BioFabricApplication bfa, 
                                                 BioFabricWindow topWindow, boolean isForMain, 
                                                 PlugInManager pMan, HeadlessOracle headlessOracle) {
    if (perClass_ == null) {
      perClass_ = new HashMap<String, CommandSet>();
    }
    CommandSet fc = perClass_.get(className);
    if (fc != null) {
      throw new IllegalStateException();
    }
    fc = new CommandSet(bfa, topWindow, isForMain, pMan, headlessOracle);
    perClass_.put(className, fc);
    return (fc);
  }
  
  ////////////////////////////////////////////////////////////////////////////
  //
  // PRIVATE CONSTRUCTORS
  //
  ////////////////////////////////////////////////////////////////////////////
  
  /***************************************************************************
  **
  ** Constructor 
  */ 
  
  private CommandSet(BioFabricApplication bfa, BioFabricWindow bfw, 
                     boolean isMain, PlugInManager pMan, HeadlessOracle headlessOracle) {
    bfa_ = bfa;
    bfw_ = bfw;
    topWindow_ = bfw.getWindow();
    withIcons_ = new HashMap<Integer, ChecksForEnabled>();
    noIcons_ = new HashMap<Integer, ChecksForEnabled>();
    plugInCmds_ = new HashMap<String, ChecksForEnabled>();
    colGen_ = new FabricColorGenerator();
    colGen_.newColorModel();
    isForMain_ = isMain;
    pMan_ = pMan;
    flf_ = new FileLoadFlowsImpl(bfw_, pMan_, colGen_, this, headlessOracle, isMain);
    isAMac_ = System.getProperty("os.name").toLowerCase().startsWith("mac os x");
    FabricDisplayOptionsManager.getMgr().addTracker(this);
    EventManager mgr = EventManager.getManager();
    mgr.addSelectionChangeListener(this);
  }    

  ////////////////////////////////////////////////////////////////////////////
  //
  // INNER CLASSES
  //
  ////////////////////////////////////////////////////////////////////////////

  /***************************************************************************
  **
  ** Command
  */ 
   
  public class HeadlessImportSIFAction {
    
    private File inputFile_;
    
    public HeadlessImportSIFAction(File inputFile) {
      inputFile_ = inputFile;
    }
    
    public boolean performOperation(Object[] args) {
    	FileLoadFlows.FileLoadResult flr = flf_.loadFromASource(inputFile_, null, null, new UniqueLabeller(), 
      		                                                    FileLoadFlows.FileLoadType.SIF, false);
      return (flr.getSuccess());
    }
  }
  
  /***************************************************************************
  **
  ** Command
  */ 
    
  public class HeadlessExportAction extends ExportImageAction {
 
    private static final long serialVersionUID = 1L;
    
    public HeadlessExportAction() {
      super(false, "command.HeadlessExport", "command.HeadlessExportMnem");
    }
   
    protected ExportSettingsDialog.ExportSettings getExportSettings() {
      return (null); 
    }
  }
  
  /***************************************************************************
  **
  ** Checks if it is enabled or not
  */
  
  public abstract class ChecksForEnabled extends AbstractAction {
    
    private static final long serialVersionUID = 1L;
    
    protected static final int IGNORE   = -1;
    protected static final int DISABLED =  0;
    protected static final int ENABLED  =  1;
    
    protected boolean enabled_ = true;
    protected boolean pushed_ = false;
    
    public void checkIfEnabled() {
      enabled_ = checkGuts();
      if (!pushed_) {
        this.setEnabled(enabled_);
      }
      return;
    }
    
    public void pushDisabled(int pushCondition) {
      pushed_ = canPush(pushCondition);
      boolean reversed = reversePush(pushCondition);
      if (pushed_) {
        this.setEnabled(reversed);
      }
    }
    
    public void setConditionalEnabled(boolean enabled) {
      //
      // If we are pushed, just stash the value.  If not
      // pushed, stash and apply.
      //
      enabled_ = enabled;
      if (!pushed_) {
        this.setEnabled(enabled_);
      }
    }    
    
    public boolean isPushed() {
      return (pushed_);
    }    
 
    public void popDisabled() {
      if (pushed_) {
        this.setEnabled(enabled_);
        pushed_ = false;
      }
      return;
    }
    
    // Default can always be enabled:
    protected boolean checkGuts() {
      return (true);
    }
 
    // Default can always be pushed:
    protected boolean canPush(int pushCondition) {
      return (true);
    }
    
    // Signals we are reverse pushed (enabled when others disabled)
    protected boolean reversePush(int pushCondition) {
      return (false);
    }    
 
  }

  /***************************************************************************
  **
  ** Command
  */ 
    
  private class PropagateDownAction extends ChecksForEnabled {
    
    private static final long serialVersionUID = 1L;
    
    PropagateDownAction(boolean doIcon) {
      ResourceManager rMan = ResourceManager.getManager(); 
      putValue(Action.NAME, rMan.getString("command.PropagateDown"));
      if (doIcon) {
        putValue(Action.SHORT_DESCRIPTION, rMan.getString("command.PropagateDown"));      
        URL ugif = getClass().getResource("/org/systemsbiology/biofabric/images/PropagateSelected24.gif");  
        putValue(Action.SMALL_ICON, new ImageIcon(ugif));
      } else {
        char mnem = rMan.getChar("command.PropagateDownMnem"); 
        putValue(Action.MNEMONIC_KEY, Integer.valueOf(mnem));
      }
    }

    public void actionPerformed(ActionEvent e) {
      try {
        bfp_.selectionsToSubmodel();
      } catch (Exception ex) {
        ExceptionHandler.getHandler().displayException(ex);
      }
      return;
    }
    
    protected boolean checkGuts() {
      return (bfp_.haveASelection());
    }
    
  }
    
  /***************************************************************************
  **
  ** Command
  */ 
    
  private class ZoomToSelected extends ChecksForEnabled {
    
    private static final long serialVersionUID = 1L;
    
    ZoomToSelected(boolean doIcon) {
      ResourceManager rMan = ResourceManager.getManager(); 
      putValue(Action.NAME, rMan.getString("command.ZoomToSelected"));
      if (doIcon) {
        putValue(Action.SHORT_DESCRIPTION, rMan.getString("command.ZoomToSelected"));
        URL ugif = getClass().getResource("/org/systemsbiology/biofabric/images/ZoomToAllFabricSelected24.gif");  
        putValue(Action.SMALL_ICON, new ImageIcon(ugif));
      } else {
        char mnem = rMan.getChar("command.ZoomToSelectedMnem"); 
        putValue(Action.MNEMONIC_KEY, Integer.valueOf(mnem));
      }
    }

    public void actionPerformed(ActionEvent e) {
      try {
        bfp_.getZoomController().zoomToSelected();
      } catch (Exception ex) {
        ExceptionHandler.getHandler().displayException(ex);
      }
      return;
    }
    
    protected boolean checkGuts() {
      return (bfp_.haveASelection());
    }    
  }
   
  /***************************************************************************
  **
  ** Command
  */ 
    
  private class ZoomToModelAction extends ChecksForEnabled {
    
    private static final long serialVersionUID = 1L;
    
    ZoomToModelAction(boolean doIcon) {
      ResourceManager rMan = ResourceManager.getManager(); 
      putValue(Action.NAME, rMan.getString("command.ZoomToModel"));
      if (doIcon) {
        putValue(Action.SHORT_DESCRIPTION, rMan.getString("command.ZoomToModel"));
        URL ugif = getClass().getResource("/org/systemsbiology/biofabric/images/ZoomToAllFabric24.gif");  
        putValue(Action.SMALL_ICON, new ImageIcon(ugif));
      } else {
        char mnem = rMan.getChar("command.ZoomToModelMnem"); 
        putValue(Action.MNEMONIC_KEY, Integer.valueOf(mnem));
      }
    }

    public void actionPerformed(ActionEvent e) {
      try {
        bfp_.getZoomController().zoomToModel(false);
      } catch (Exception ex) {
        ExceptionHandler.getHandler().displayException(ex);
      }
      return;
    }
    
    protected boolean checkGuts() {
      return (bfp_.hasAModel());
    }    
  }  

  /***************************************************************************
  **
  ** Command
  */ 
    
  private class ZoomToCurrentSelected extends ChecksForEnabled {
    
    private static final long serialVersionUID = 1L;
    
    ZoomToCurrentSelected(boolean doIcon) {
      ResourceManager rMan = ResourceManager.getManager(); 
      putValue(Action.NAME, rMan.getString("command.ZoomToCurrentSelected"));
      if (doIcon) {
        putValue(Action.SHORT_DESCRIPTION, rMan.getString("command.ZoomToCurrentSelected"));
        URL ugif = getClass().getResource("/org/systemsbiology/biofabric/images/ZoomToFabricSelected24.gif");  
        putValue(Action.SMALL_ICON, new ImageIcon(ugif));
      } else {
        char mnem = rMan.getChar("command.ZoomToCurrentSelectedMnem"); 
        putValue(Action.MNEMONIC_KEY, Integer.valueOf(mnem));
      }
    }

    public void actionPerformed(ActionEvent e) {
      try {
        bfp_.getZoomController().zoomToCurrentSelected();
      } catch (Exception ex) {
        ExceptionHandler.getHandler().displayException(ex);
      }
      return;
    }
    
    protected boolean checkGuts() {
      return (bfp_.haveASelection());
    }   
  }
  
  /***************************************************************************
  **
  ** Command
  */ 
    
  private class ZoomToCurrentMouse extends ChecksForEnabled {
    
    private static final long serialVersionUID = 1L;
    
    ZoomToCurrentMouse(boolean doIcon) {
      ResourceManager rMan = ResourceManager.getManager(); 
      putValue(Action.NAME, rMan.getString("command.ZoomToCurrentMouse"));
      if (doIcon) {
        putValue(Action.SHORT_DESCRIPTION, rMan.getString("command.ZoomToCurrentMouse"));
        URL ugif = getClass().getResource("/org/systemsbiology/biofabric/images/FIXME24.gif");  
        putValue(Action.SMALL_ICON, new ImageIcon(ugif));
      } else {
        char mnem = rMan.getChar("command.ZoomToCurrentMouseMnem"); 
        putValue(Action.MNEMONIC_KEY, Integer.valueOf(mnem));
        char accel = rMan.getChar("command.ZoomToCurrentMouseAccel");
        putValue(Action.ACCELERATOR_KEY, KeyStroke.getKeyStroke(accel, Event.CTRL_MASK, false));
      }
    }

    public void actionPerformed(ActionEvent e) {
      try {
        FabricMagnifyingTool fmt = bfp_.getMagnifier();
        Point2D pt = fmt.getMouseLoc();
        Point rcPoint = bfp_.worldToRowCol(pt);
        Rectangle rect = bfp_.buildFocusBox(rcPoint);
        bfp_.getZoomController().zoomToRect(rect);
      } catch (Exception ex) {
        ExceptionHandler.getHandler().displayException(ex);
      }
      return;
    }
    
    protected boolean checkGuts() {
      return (bfp_.hasAModel());
    }   
  }
  
  /***************************************************************************
  **
  ** Command
  */ 
    
  private class ZoomToCurrentMagnify extends ChecksForEnabled {
    
    private static final long serialVersionUID = 1L;
    
    ZoomToCurrentMagnify(boolean doIcon) {
      ResourceManager rMan = ResourceManager.getManager(); 
      putValue(Action.NAME, rMan.getString("command.ZoomToCurrentMagnify"));
      if (doIcon) {
        putValue(Action.SHORT_DESCRIPTION, rMan.getString("command.ZoomToCurrentMagnify"));
        URL ugif = getClass().getResource("/org/systemsbiology/biofabric/images/ZoomToFabricSelected24.gif");  
        putValue(Action.SMALL_ICON, new ImageIcon(ugif));
      } else {
        char mnem = rMan.getChar("command.ZoomToCurrentMagnifyMnem"); 
        putValue(Action.MNEMONIC_KEY, Integer.valueOf(mnem));
        char accel = rMan.getChar("command.ZoomToCurrentMagnifyAccel");
        putValue(Action.ACCELERATOR_KEY, KeyStroke.getKeyStroke(accel, Event.CTRL_MASK, false));
      }
    }

    public void actionPerformed(ActionEvent e) {
      try {        
        FabricMagnifyingTool fmt = bfp_.getMagnifier();
        Rectangle rect = fmt.getClipRect();
        bfp_.getZoomController().zoomToRect(rect);
      } catch (Exception ex) {
        ExceptionHandler.getHandler().displayException(ex);
      }
      return;
    }
    
    protected boolean checkGuts() {
      return (bfp_.hasAModel());
    }   
  }

  /***************************************************************************
  **
  ** Command
  */ 
    
  private class CenterOnNextSelected extends ChecksForEnabled {
    
    private static final long serialVersionUID = 1L;
    
    CenterOnNextSelected(boolean doIcon) {
      ResourceManager rMan = ResourceManager.getManager(); 
      putValue(Action.NAME, rMan.getString("command.CenterOnNextSelected"));
      if (doIcon) {
        putValue(Action.SHORT_DESCRIPTION, rMan.getString("command.CenterOnNextSelected"));
        URL ugif = getClass().getResource("/org/systemsbiology/biofabric/images/Forward24.gif");  
        putValue(Action.SMALL_ICON, new ImageIcon(ugif));
      } else {
        char mnem = rMan.getChar("command.CenterOnNextSelectedMnem"); 
        putValue(Action.MNEMONIC_KEY, Integer.valueOf(mnem));
      }
    }

    public void actionPerformed(ActionEvent e) {
      try {
        bfp_.getZoomController().centerToNextSelected();
      } catch (Exception ex) {
        ExceptionHandler.getHandler().displayException(ex);
      }
      return;
    }
    
    protected boolean checkGuts() {
      return (bfp_.haveASelection());
    }   
  }
  
  
  /***************************************************************************
  **
  ** Command
  */ 
    
  private class CenterOnPreviousSelected extends ChecksForEnabled {
    
    private static final long serialVersionUID = 1L;
    
    CenterOnPreviousSelected(boolean doIcon) {
      ResourceManager rMan = ResourceManager.getManager(); 
      putValue(Action.NAME, rMan.getString("command.CenterOnPreviousSelected"));
      if (doIcon) {
        putValue(Action.SHORT_DESCRIPTION, rMan.getString("command.CenterOnPreviousSelected"));
        URL ugif = getClass().getResource("/org/systemsbiology/biofabric/images/Back24.gif");  
        putValue(Action.SMALL_ICON, new ImageIcon(ugif));
      } else {
        char mnem = rMan.getChar("command.CenterOnPreviousSelectedMnem"); 
        putValue(Action.MNEMONIC_KEY, Integer.valueOf(mnem));
      }
    }

    public void actionPerformed(ActionEvent e) {
      try {
        bfp_.getZoomController().centerToPreviousSelected();
      } catch (Exception ex) {
        ExceptionHandler.getHandler().displayException(ex);
      }
      return;
    }
    
    protected boolean checkGuts() {
      return (bfp_.haveASelection());
    }   
  }
  
  /***************************************************************************
  **
  ** Command
  */ 
    
  private class AddFirstNeighborsAction extends ChecksForEnabled {
    
    private static final long serialVersionUID = 1L;
    
    AddFirstNeighborsAction(boolean doIcon) {
      ResourceManager rMan = ResourceManager.getManager(); 
      putValue(Action.NAME, rMan.getString("command.AddFirstNeighbors"));
      if (doIcon) {
        putValue(Action.SHORT_DESCRIPTION, rMan.getString("command.AddFirstNeighbors"));
        URL ugif = getClass().getResource("/org/systemsbiology/biofabric/images/PlusOneDeg24.gif");  
        putValue(Action.SMALL_ICON, new ImageIcon(ugif));
      } else {
        char mnem = rMan.getChar("command.AddFirstNeighborsMnem"); 
        putValue(Action.MNEMONIC_KEY, Integer.valueOf(mnem));
      }
    }

    public void actionPerformed(ActionEvent e) {
      try {
        bfp_.addFirstNeighbors();
      } catch (Exception ex) {
        ExceptionHandler.getHandler().displayException(ex);
      }
      return;
    }
    
    protected boolean checkGuts() {
      return (bfp_.haveASelection());
    }      
  }
  
  /***************************************************************************
  **
  ** Command
  */ 
    
  private class BuildSelectAction extends ChecksForEnabled {
    
    private static final long serialVersionUID = 1L;
    
    BuildSelectAction(boolean doIcon) {
      ResourceManager rMan = ResourceManager.getManager(); 
      putValue(Action.NAME, rMan.getString("command.BuildSelect"));
      if (doIcon) {
        putValue(Action.SHORT_DESCRIPTION, rMan.getString("command.BuildSelect"));
        URL ugif = getClass().getResource("/org/systemsbiology/biofabric/images/FIXME.gif");  
        putValue(Action.SMALL_ICON, new ImageIcon(ugif));
      } else {
        char mnem = rMan.getChar("command.BuildSelectMnem"); 
        putValue(Action.MNEMONIC_KEY, Integer.valueOf(mnem));
      }
    }

    public void actionPerformed(ActionEvent e) {
      try {
        bfp_.toggleBuildSelect();
      } catch (Exception ex) {
        ExceptionHandler.getHandler().displayException(ex);
      }
      return;
    }
  }
  
  /***************************************************************************
  **
  ** Command
  */ 
  
  private class InOutZoomAction extends ChecksForEnabled {
   
    private static final long serialVersionUID = 1L;
    private char sign_;
    
    InOutZoomAction(boolean doIcon, char sign) {
      sign_ = sign;
      ResourceManager rMan = ResourceManager.getManager();
      String iconName;
      String stringName;
      String mnemName;
      String accelName;      
      if (sign == '+') {
        iconName = "ZoomIn24.gif";
        stringName = "command.ZoomIn";
        mnemName = "command.ZoomInMnem";
        accelName = "command.ZoomInAccel";        
      } else {
        iconName = "ZoomOut24.gif";
        stringName = "command.ZoomOut";
        mnemName = "command.ZoomOutMnem";
        accelName = "command.ZoomOutAccel";
      }      
      putValue(Action.NAME, rMan.getString(stringName));
      if (doIcon) {
        putValue(Action.SHORT_DESCRIPTION, rMan.getString(stringName));      
        URL ugif = getClass().getResource("/org/systemsbiology/biofabric/images/" + iconName);  
        putValue(Action.SMALL_ICON, new ImageIcon(ugif));
      } else {
        char mnem = rMan.getChar(mnemName); 
        putValue(Action.MNEMONIC_KEY, Integer.valueOf(mnem));
      }
      char accel = rMan.getChar(accelName);
      putValue(Action.ACCELERATOR_KEY, KeyStroke.getKeyStroke(accel, Event.CTRL_MASK, false));
    }
    
    public void actionPerformed(ActionEvent e) {
      try {
        bfp_.getZoomController().bumpZoomWrapper(sign_);
      } catch (Exception ex) {
        ExceptionHandler.getHandler().displayException(ex);
      }
      return;
    }
    
    public void setConditionalEnabled(boolean enabled) {
      setEnabled(enabled);
      return;
    } 
    
    //
    // Override: We handle this internally.
    //
    public void checkIfEnabled() {
    }    
        
    protected boolean canPush(int pushCondition) {
      return ((pushCondition & CommandSet.ALLOW_NAV_PUSH) == 0x00);
    }     
    
  }
    
  /***************************************************************************
  **
  ** Command
  */ 
  
  private class ClearSelectionsAction extends ChecksForEnabled {
    
    private static final long serialVersionUID = 1L;
   
    ClearSelectionsAction(boolean doIcon) {
      ResourceManager rMan = ResourceManager.getManager(); 
      putValue(Action.NAME, rMan.getString("command.ClearSel"));
      if (doIcon) {
        putValue(Action.SHORT_DESCRIPTION, rMan.getString("command.ClearSel"));        
        URL ugif = getClass().getResource("/org/systemsbiology/biofabric/images/ClearFabricSelected24.gif");  
        putValue(Action.SMALL_ICON, new ImageIcon(ugif));
      } else {
        char mnem = rMan.getChar("command.ClearSelMnem"); 
        putValue(Action.MNEMONIC_KEY, Integer.valueOf(mnem)); 
      }
    }
    
    public void actionPerformed(ActionEvent e) {
      try {
        bfp_.clearSelections();        
      } catch (Exception ex) {
        ExceptionHandler.getHandler().displayException(ex);
      }
      return;
    }
    
    protected boolean checkGuts() {
      return (bfp_.haveASelection());
    }       
  } 
 
  /***************************************************************************
  **
  ** Command
  */ 
  
  private class ZoomToRect extends ChecksForEnabled {
    
    private static final long serialVersionUID = 1L;
   
    ZoomToRect(boolean doIcon) {
      ResourceManager rMan = ResourceManager.getManager(); 
      putValue(Action.NAME, rMan.getString("command.ZoomToRect"));
      if (doIcon) {
        putValue(Action.SHORT_DESCRIPTION, rMan.getString("command.ZoomToRect"));        
        URL ugif = getClass().getResource("/org/systemsbiology/biofabric/images/ZoomToFabricRect24.gif");  
        putValue(Action.SMALL_ICON, new ImageIcon(ugif));
      } else {
        char mnem = rMan.getChar("command.ZoomToRectMnem"); 
        putValue(Action.MNEMONIC_KEY, Integer.valueOf(mnem));
        char accel = rMan.getChar("command.ZoomToRectAccel");
        putValue(Action.ACCELERATOR_KEY, KeyStroke.getKeyStroke(accel, Event.CTRL_MASK, false));
      }
      
    }
    
    public void actionPerformed(ActionEvent e) {
      try {    
        bfp_.setToCollectZoomRect();
      } catch (Exception ex) {
        ExceptionHandler.getHandler().displayException(ex);
      }
      return;
    }
    
    protected boolean checkGuts() {
      return (bfp_.hasAModel());
    }
       
  }
  
  /***************************************************************************
  **
  ** Command
  */ 
    
  private class CancelAction extends ChecksForEnabled {
    
    private static final long serialVersionUID = 1L;
   
    CancelAction(boolean doIcon) {
      ResourceManager rMan = ResourceManager.getManager(); 
      putValue(Action.NAME, rMan.getString("command.CancelAddMode"));
      if (doIcon) {
        putValue(Action.SHORT_DESCRIPTION, rMan.getString("command.CancelAddMode"));        
        URL ugif = getClass().getResource("/org/systemsbiology/biofabric/images/Stop24.gif");  
        putValue(Action.SMALL_ICON, new ImageIcon(ugif));
      } else {
        char mnem = rMan.getChar("command.CancelAddModeMnem"); 
        putValue(Action.MNEMONIC_KEY, Integer.valueOf(mnem));
      }
    }

    public void actionPerformed(ActionEvent e) {
      try {
         bfp_.cancelModals();
      } catch (Exception ex) {
        ExceptionHandler.getHandler().displayException(ex);
      }
      return;
    }
    
    protected boolean checkGuts() {
      return (false);      
    }
 
    protected boolean reversePush(int pushCondition) {
      return ((pushCondition & CommandSet.ALLOW_NAV_PUSH) != 0x00);
    }    
      
  }

  /***************************************************************************
  **
  ** Command
  */ 
  
  private class SearchAction extends ChecksForEnabled {
    
    private static final long serialVersionUID = 1L;
    
    SearchAction(boolean doIcon) {
      ResourceManager rMan = ResourceManager.getManager(); 
      putValue(Action.NAME, rMan.getString("command.Search"));
      if (doIcon) {
        putValue(Action.SHORT_DESCRIPTION, rMan.getString("command.Search"));        
        URL ugif = getClass().getResource("/org/systemsbiology/biofabric/images/Find24.gif");  
        putValue(Action.SMALL_ICON, new ImageIcon(ugif));
      } else {
        char mnem = rMan.getChar("command.SearchMnem"); 
        putValue(Action.MNEMONIC_KEY, Integer.valueOf(mnem)); 
      }
    }
    
    public void actionPerformed(ActionEvent e) {
      try {
        boolean haveSelection = bfp_.haveASelection();
        boolean buildingSels = bfp_.amBuildingSelections();
        FabricSearchDialog fsd = new FabricSearchDialog(topWindow_, bfw_.getFabricPanel().getNetwork(), 
                                                        haveSelection, buildingSels);      
        fsd.setVisible(true);
        if (fsd.itemWasFound()) {
          Set<NetNode> matches = fsd.getMatches();
          boolean doDiscard = fsd.discardSelections();
          bfw_.getFabricPanel().installSearchResult(matches, doDiscard);
        }
        return;
      } catch (Exception ex) {
        ExceptionHandler.getHandler().displayException(ex);
      }
    }
    
    protected boolean checkGuts() {
      return (bfp_.hasAModel());
    }
    
  }
  
  /***************************************************************************
  **
  ** Command
  */ 
  
  private class CompareNodesAction extends ChecksForEnabled {
    
    private static final long serialVersionUID = 1L;
    
    CompareNodesAction(boolean doIcon) {
      ResourceManager rMan = ResourceManager.getManager(); 
      putValue(Action.NAME, rMan.getString("command.CompareNodes"));
      if (doIcon) {
        putValue(Action.SHORT_DESCRIPTION, rMan.getString("command.CompareNodes"));        
        URL ugif = getClass().getResource("/org/systemsbiology/biofabric/images/FIXME24.gif");  
        putValue(Action.SMALL_ICON, new ImageIcon(ugif));
      } else {
        char mnem = rMan.getChar("command.CompareNodesMnem"); 
        putValue(Action.MNEMONIC_KEY, Integer.valueOf(mnem)); 
      }
    }
    
    public void actionPerformed(ActionEvent e) {
      try {
        Set<NetNode> allNodeIDs = bfp_.getNetwork().getNodeSetIDs();
        Map<String, Set<NetNode>> normNameToIDs = bfp_.getNetwork().getNormNameToIDs();
        CompareNodesSetupDialog fsd = new CompareNodesSetupDialog(topWindow_, allNodeIDs, normNameToIDs);      
        fsd.setVisible(true);
        if (fsd.haveResult()) {
          Set<NetNode> result = fsd.getResults();
          bfp_.installSearchResult(result, true);          
          bfp_.addFirstNeighbors();
          bfp_.selectionsToSubmodel();
        }
        return;
      } catch (Exception ex) {
        ExceptionHandler.getHandler().displayException(ex);
      }
    }
    
    protected boolean checkGuts() {
      return (bfp_.hasAModel());
    }
    
  }
  
  /***************************************************************************
  **
  ** Command
  */ 
  
  private class CloseAction extends ChecksForEnabled {
    
    private static final long serialVersionUID = 1L;
    
    CloseAction(boolean doIcon) {
      ResourceManager rMan = ResourceManager.getManager(); 
      putValue(Action.NAME, rMan.getString("command.Close"));
      if (doIcon) {
        putValue(Action.SHORT_DESCRIPTION, rMan.getString("command.Close"));        
        URL ugif = getClass().getResource("/org/systemsbiology/biofabric/images/Stop24.gif");  
        putValue(Action.SMALL_ICON, new ImageIcon(ugif));
      } else {
        char mnem = rMan.getChar("command.CloseMnem"); 
        putValue(Action.MNEMONIC_KEY, Integer.valueOf(mnem)); 
      }
    }
    
    public void actionPerformed(ActionEvent e) {
      try {
        bfa_.shutdownFabric();
      } catch (Exception ex) {
        // Going down (usually) so don't show exception in UI
        ex.printStackTrace();
      }
    }
  }
  
  /***************************************************************************
  **
  ** Command
  */ 
   
  private abstract class LayoutViaAttributesAction extends ChecksForEnabled {
     
    private static final long serialVersionUID = 1L;
    
    LayoutViaAttributesAction(boolean doIcon, String name, String mnemStr) {
      ResourceManager rMan = ResourceManager.getManager(); 
      putValue(Action.NAME, rMan.getString(name));
      if (doIcon) {
        putValue(Action.SHORT_DESCRIPTION, rMan.getString(name));       
        URL ugif = getClass().getResource("/org/systemsbiology/biofabric/images/FIXME24.gif");  
        putValue(Action.SMALL_ICON, new ImageIcon(ugif));
      } else {
        char mnem = rMan.getChar(mnemStr); 
        putValue(Action.MNEMONIC_KEY, Integer.valueOf(mnem)); 
      }
    } 
    
    public void actionPerformed(ActionEvent e) {
      try {
        performOperation();
      } catch (Exception ex) {
        ExceptionHandler.getHandler().displayException(ex);
      }      
      return;
    }
   
    protected abstract boolean performOperation();
    
    protected boolean checkGuts() {
      return (bfp_.hasAModel());
    }
 
  }        
  
  /***************************************************************************
  **
  ** Command
  */ 
   
  private class LayoutNodesViaAttributesAction extends LayoutViaAttributesAction {
    
    private static final long serialVersionUID = 1L;
    
    LayoutNodesViaAttributesAction(boolean doIcon) {
      super(doIcon, "command.LayoutNodesViaAttributes", "command.LayoutNodesViaAttributesMnem");
    }
    
    protected boolean performOperation() {
      File file = flf_.getTheFile(".noa", ".na", "AttribDirectory", "filterName.noa", null);
      if (file == null) {
        return (true);
      }
      Map<AttributeKey, String> nodeAttributes = flf_.loadTheFile(file, null, true);
      if (nodeAttributes == null) {
        return (true);
      }
      if (!bfp_.getNetwork().checkNewNodeOrder(nodeAttributes)) { 
        ResourceManager rMan = ResourceManager.getManager();
        JOptionPane.showMessageDialog(topWindow_, rMan.getString("attribRead.badRowMessage"),
                                      rMan.getString("attribRead.badRowSemanticsTitle"),
                                      JOptionPane.WARNING_MESSAGE);
        return (true);
      } 
      flf_.doNetworkRelayout(nodeAttributes);    
      return (true);
    }
    
    @Override
    protected boolean checkGuts() {
      return (bfp_.hasAModel() && (bfp_.getNetwork().getLinkCount(true) != 0));
    }
  }      
  
  /***************************************************************************
  **
  ** Command
  */ 
   
  private class AddNodeAnnotations extends ChecksForEnabled  {
    
    private static final long serialVersionUID = 1L;
    
    AddNodeAnnotations(boolean doIcon) {
      
      ResourceManager rMan = ResourceManager.getManager(); 
      putValue(Action.NAME, rMan.getString("command.AddNodeAnnotations"));
      if (doIcon) {
        putValue(Action.SHORT_DESCRIPTION, rMan.getString("command.AddNodeAnnotations"));
        URL ugif = getClass().getResource("/org/systemsbiology/biofabric/images/FIXME24.gif");  
        putValue(Action.SMALL_ICON, new ImageIcon(ugif));
      } else {
        char mnem = rMan.getChar("command.AddNodeAnnotationsMnem"); 
        putValue(Action.MNEMONIC_KEY, Integer.valueOf(mnem));
      }
    } 
    
    public void actionPerformed(ActionEvent e) {
      try {
        performOperation();
      } catch (Exception ex) {
        ExceptionHandler.getHandler().displayException(ex);
      }      
      return;
    }

    protected boolean performOperation() {
      File file = flf_.getTheFile(".tsv", null, "AnnotDirectory", "filterName.tsv", null);
      if (file == null) {
        return (true);
      }
      Map<Boolean, AnnotationSet> aSet = flf_.loadAnnotations(file, true);
      if (aSet == null) {
        return (true);
      }
      bfp_.getNetwork().setNodeAnnotations(aSet.get(Boolean.TRUE));
      flf_.doRecolor(isForMain_);      
      return (true);
    }
    
    @Override
    protected boolean checkGuts() {
      return (bfp_.hasAModel() && (bfp_.getNetwork().getLinkCount(true) != 0));
    }
  }
 
  
  /***************************************************************************
  **
  ** Command
  */ 
   
  private class AddLinkAnnotations extends ChecksForEnabled  {
    
    private static final long serialVersionUID = 1L;
    
    AddLinkAnnotations(boolean doIcon) {
      
      ResourceManager rMan = ResourceManager.getManager(); 
      putValue(Action.NAME, rMan.getString("command.AddLinkAnnotations"));
      if (doIcon) {
        putValue(Action.SHORT_DESCRIPTION, rMan.getString("command.AddLinkAnnotations"));
        URL ugif = getClass().getResource("/org/systemsbiology/biofabric/images/FIXME24.gif");  
        putValue(Action.SMALL_ICON, new ImageIcon(ugif));
      } else {
        char mnem = rMan.getChar("command.AddLinkAnnotationsMnem"); 
        putValue(Action.MNEMONIC_KEY, Integer.valueOf(mnem));
      }
    } 
    
    public void actionPerformed(ActionEvent e) {
      try {
        performOperation();
      } catch (Exception ex) {
        ExceptionHandler.getHandler().displayException(ex);
      }      
      return;
    }

    protected boolean performOperation() {
      File file = flf_.getTheFile(".tsv", null, "AnnotDirectory", "filterName.tsv", null);
      if (file == null) {
        return (true);
      }
      Map<Boolean, AnnotationSet> aSet = flf_.loadAnnotations(file, false);
      if (aSet == null) {
        return (true);
      }
      bfp_.getNetwork().setLinkAnnotations(aSet.get(Boolean.TRUE), true);
      bfp_.getNetwork().setLinkAnnotations(aSet.get(Boolean.FALSE), false);
      flf_.doRecolor(isForMain_);
      
      return (true);
    }
    
    @Override
    protected boolean checkGuts() {
      return (bfp_.hasAModel() && (bfp_.getNetwork().getLinkCount(true) != 0));
    }
  }
  
  /***************************************************************************
  **
  ** Command
  */ 
   
  private class SetPluginDirectory extends ChecksForEnabled  {
    
    private static final long serialVersionUID = 1L;
    
    SetPluginDirectory(boolean doIcon) {
      
      ResourceManager rMan = ResourceManager.getManager(); 
      putValue(Action.NAME, rMan.getString("command.SetPluginDirectory"));
      if (doIcon) {
        putValue(Action.SHORT_DESCRIPTION, rMan.getString("command.SetPluginDirectory"));
        URL ugif = getClass().getResource("/org/systemsbiology/biofabric/images/FIXME24.gif");  
        putValue(Action.SMALL_ICON, new ImageIcon(ugif));
      } else {
        char mnem = rMan.getChar("command.SetPluginDirectoryMnem"); 
        putValue(Action.MNEMONIC_KEY, Integer.valueOf(mnem));
      }
    } 
    
    public void actionPerformed(ActionEvent e) {
      try {
        performOperation();
      } catch (Exception ex) {
        ExceptionHandler.getHandler().displayException(ex);
      }      
      return;
    }

    protected boolean performOperation() {
      // prompt the user to enter their name
    	String currentChoice = pMan_.getDirectory();
    	DirectoryChooserDialog dcd = new DirectoryChooserDialog(topWindow_, flf_, currentChoice);
    	dcd.setVisible(true);
    	if (dcd.haveResult()) {
    		ResourceManager rMan = ResourceManager.getManager();
        JOptionPane.showMessageDialog(topWindow_, rMan.getString("setPlugin.RestartMessage"),
                                      rMan.getString("setPlugin.RestartMessageTitle"),
                                      JOptionPane.INFORMATION_MESSAGE);	
        File directory = dcd.getDirectory();
        pMan_.setDirectory(directory);
      }
      return (true);
    }
  }
  
  /***************************************************************************
  **
  ** Command
  */ 
   
  private class LayoutLinksViaAttributesAction extends LayoutViaAttributesAction {
       
    private static final long serialVersionUID = 1L;
    
    LayoutLinksViaAttributesAction(boolean doIcon) {
      super(doIcon, "command.LayoutLinksViaAttributes", "command.LayoutLinksViaAttributesMnem");
    }
    
    protected boolean performOperation() {   
      File file = flf_.getTheFile(".eda", ".ed", "AttribDirectory", "filterName.eda", null);
      if (file == null) {
        return (true);
      }
      Map<String, Set<NetNode>> nameToIDs = bfp_.getNetwork().getNormNameToIDs();
      Map<AttributeKey, String> edgeAttributes = flf_.loadTheFile(file, nameToIDs, false);
      if (edgeAttributes == null) {
        return (true);
      }
      SortedMap<Integer, NetLink> modifiedAndChecked = bfp_.getNetwork().checkNewLinkOrder(edgeAttributes);      
      if (modifiedAndChecked == null) { 
        ResourceManager rMan = ResourceManager.getManager();
        JOptionPane.showMessageDialog(topWindow_, rMan.getString("attribRead.badColMessage"),
                                      rMan.getString("attribRead.badColSemanticsTitle"),
                                      JOptionPane.WARNING_MESSAGE);
        return (true);
      }     
      flf_.doNetworkLinkRelayout(modifiedAndChecked);        
      return (true);
    }
    
    @Override
    protected boolean checkGuts() {
      return (bfp_.hasAModel() && (bfp_.getNetwork().getLinkCount(true) != 0));
    } 
  }
  
  /***************************************************************************
  **
  ** Command
  */ 
   
  private class ToggleShowTourAction extends ChecksForEnabled {
    
    private static final long serialVersionUID = 1L;
    
    private boolean showTour_;
    
    ToggleShowTourAction(boolean doIcon) {
      ResourceManager rMan = ResourceManager.getManager(); 
      putValue(Action.NAME, rMan.getString("command.ShowTourAction"));
      if (doIcon) {
        putValue(Action.SHORT_DESCRIPTION, rMan.getString("command.ShowTourAction"));        
        URL ugif = getClass().getResource("/org/systemsbiology/biofabric/images/FIXME24.gif");  
        putValue(Action.SMALL_ICON, new ImageIcon(ugif));
      } else {
        char mnem = rMan.getChar("command.ShowTourActionMnem"); 
        putValue(Action.MNEMONIC_KEY, Integer.valueOf(mnem)); 
      }
      showTour_ = true;
    }
    
    public void actionPerformed(ActionEvent ev) {
      try {
        showTour_ = !showTour_;
        bfw_.showTour(showTour_);
      } catch (Exception ex) {
        ExceptionHandler.getHandler().displayException(ex);
      }
    }
    
    @Override
    protected boolean checkGuts() {
      return (showNav_);
    }   
  }  
  
  /***************************************************************************
  **
  ** Command
  */ 
   
  private class ToggleShowNavPanelAction extends ChecksForEnabled {
     
    private static final long serialVersionUID = 1L;
    
    ToggleShowNavPanelAction(boolean doIcon) {
      ResourceManager rMan = ResourceManager.getManager(); 
      putValue(Action.NAME, rMan.getString("command.ShowNavPanel"));
      if (doIcon) {
        putValue(Action.SHORT_DESCRIPTION, rMan.getString("command.ShowNavPanel"));        
        URL ugif = getClass().getResource("/org/systemsbiology/biofabric/images/FIXME24.gif");  
        putValue(Action.SMALL_ICON, new ImageIcon(ugif));
      } else {
        char mnem = rMan.getChar("command.ShowNavPanelMnem"); 
        putValue(Action.MNEMONIC_KEY, Integer.valueOf(mnem)); 
      }
      showNav_ = true;
    }
    
    public void actionPerformed(ActionEvent ev) {
      try {
        showNav_ = !showNav_;
        bfw_.showNavAndControl(showNav_);
        checkForChanges(); // To disable/enable tour hiding
      } catch (Exception ex) {
        ExceptionHandler.getHandler().displayException(ex);
      }
      return;
    }
  }  

  /***************************************************************************
  **
  ** Command
  */ 
   
  private class LayoutViaNodeClusterAction extends LayoutViaAttributesAction {
     
    private static final long serialVersionUID = 1L;
    
    LayoutViaNodeClusterAction(boolean doIcon) {
      super(doIcon, "command.LayoutViaNodeClusterAction", "command.LayoutViaNodeClusterAction");
    }
    
    protected boolean performOperation() {
    	Set<NetNode> sels = bfp_.getNodeSelections();
      NetNode selNode = (sels.size() == 1) ? sels.iterator().next() : null;
    	
    	ClusterLayoutSetupDialog clsd = new ClusterLayoutSetupDialog(topWindow_, bfp_.getNetwork(), selNode);
      clsd.setVisible(true);      
      if (clsd.haveResult()) {
        NodeClusterLayout.ClusterParams params = clsd.getParams();
        if (params.needsFile()) {
        	if (!ClusterLayoutSetupDialog.askForFileInfo(params, CommandSet.this, flf_, bfp_.getNetwork())) {
        		return (true);
        	}
        }
        flf_.doNetworkClusterRelayout(params);
      }
      return (true);
    }
    
    @Override
    protected boolean checkGuts() {
      return (bfp_.hasAModel() && (bfp_.getNetwork().getLinkCount(true) != 0));
    }
  }      
  
  /***************************************************************************
  **
  ** Command
  */ 
   
  private class LayoutTopControlAction extends BasicLayoutAction {
     
    private static final long serialVersionUID = 1L;
    
    LayoutTopControlAction(boolean doIcon) {
      super(doIcon, "command.TopControlLayout", "command.TopControlLayoutMnem", BuildDataImpl.BuildMode.CONTROL_TOP_LAYOUT);
    }
    
    @Override
    public void actionPerformed(ActionEvent e) {
      try { 
        ControlTopLayoutSetupDialog ctlsud = new ControlTopLayoutSetupDialog(topWindow_);
        ctlsud.setVisible(true);
        if (ctlsud.haveResult()) {
          List<String> fixedList = null;
          ControlTopLayout.CtrlMode cMode = ctlsud.getCMode();
          ControlTopLayout.TargMode tMode = ctlsud.getTMode();
          if (cMode == ControlTopLayout.CtrlMode.FIXED_LIST) {
            File fileEda = flf_.getTheFile(".txt", null, "AttribDirectory", "filterName.txt", null);
            if (fileEda == null) {
              return;
            }
            fixedList = UiUtil.simpleFileRead(fileEda);
            if (fixedList == null) {
              return;
            }
          }  
          flf_.doControlTopRelayout(cMode, tMode, fixedList);
        }
      } catch (Exception ex) {
        ExceptionHandler.getHandler().displayException(ex);
      }
      return;
    } 
  }  
  
  /***************************************************************************
  **
  ** Command
  */ 
   
  private class HierDAGLayoutAction extends BasicLayoutAction {
     
    private static final long serialVersionUID = 1L;
    
    HierDAGLayoutAction(boolean doIcon) {
      super(doIcon, "command.HierDAGLayout", "command.HierDAGLayoutMnem", BuildDataImpl.BuildMode.HIER_DAG_LAYOUT);
    }
    
    @Override
    public void actionPerformed(ActionEvent e) {
      try {
        PointUpOrDownDialog puodd = new PointUpOrDownDialog(topWindow_);
        puodd.setVisible(true);
        boolean pointUp = false;
        if (puodd.haveResult()) {
          pointUp = puodd.getPointUp();
          flf_.doHierDagRelayout(pointUp);
        }
      } catch (Exception ex) {
        ExceptionHandler.getHandler().displayException(ex);
      }
      return;
    }
  }
  
  /***************************************************************************
  **
  ** Command
  */ 
   
  private class SetLayoutAction extends BasicLayoutAction {
     
    private static final long serialVersionUID = 1L;
    
    SetLayoutAction(boolean doIcon) {
      super(doIcon, "command.SetLayout", "command.SetLayoutMnem", BuildDataImpl.BuildMode.SET_LAYOUT);
    }
    
    @Override
    public void actionPerformed(ActionEvent e) {
      try {
      	HashSet<String> rels = new HashSet<String>();
      	Set<NetLink> links = bfp_.getNetwork().getAllLinks(false);
      	for (NetLink link : links) {
      		rels.add(link.getRelation());
      	}
      	if (rels.size() != 1) { 
          ResourceManager rMan = ResourceManager.getManager();
          JOptionPane.showMessageDialog(topWindow_, rMan.getString("setLayout.oneRelationAllowed"),
                                        rMan.getString("setLayout.oneRelationAllowedTitle"),
                                        JOptionPane.WARNING_MESSAGE);
          return;
        }
        SetRelationSemanticsDialog puodd = new SetRelationSemanticsDialog(topWindow_, rels.iterator().next());
        puodd.setVisible(true);
        if (puodd.haveResult()) {
          flf_.doSetRelayout(puodd.getLinkMeaning());
        }
      } catch (Exception ex) {
        ExceptionHandler.getHandler().displayException(ex);
      }
      return;
    }
 
  } 
  
  /***************************************************************************
  **
  ** Command
  */ 
   
  private class ToggleShadowLinks extends ChecksForEnabled {
     
    private static final long serialVersionUID = 1L;
    
    ToggleShadowLinks(boolean doIcon) {
      ResourceManager rMan = ResourceManager.getManager(); 
      putValue(Action.NAME, rMan.getString("command.ToggleShadowLinks"));
      if (doIcon) {
        putValue(Action.SHORT_DESCRIPTION, rMan.getString("command.ToggleShadowLinks"));        
        URL ugif = getClass().getResource("/org/systemsbiology/biofabric/images/S24.gif");  
        putValue(Action.SMALL_ICON, new ImageIcon(ugif));
      } else {
        char mnem = rMan.getChar("command.ToggleShadowLinksMnem"); 
        putValue(Action.MNEMONIC_KEY, Integer.valueOf(mnem)); 
      }
      showNav_ = true;
    }
    
    public void actionPerformed(ActionEvent e) {
    	try {	
      	if (bfp_.getNetwork().getLinkCount(false) > FileLoadFlows.SIZE_TO_ASK_ABOUT_SHADOWS) {
  	    	ResourceManager rMan = ResourceManager.getManager(); 
  	    	int keepGoing =
  		      JOptionPane.showConfirmDialog(topWindow_, rMan.getString("toggleShadow.bigFileLongTime"),
  		                                    rMan.getString("toggleShadow.bigFileLongTimeTitle"),
  		                                    JOptionPane.YES_NO_OPTION);        
  		    if (keepGoing != JOptionPane.YES_OPTION) {
  		    	return;
  		    }
      	}
  
      	FabricDisplayOptionsManager dopmgr = FabricDisplayOptionsManager.getMgr();
      	FabricDisplayOptions dop = dopmgr.getDisplayOptions();
      	FabricDisplayOptions newDop = dop.clone();
      	newDop.setDisplayShadows(!dop.getDisplayShadows());
        dopmgr.setDisplayOptions(newDop, true, false);
      } catch (Exception ex) {
        ExceptionHandler.getHandler().displayException(ex);
      }
    	return;
    }
    
    @Override
    protected boolean checkGuts() {
      return (bfp_.hasAModel() && (bfp_.getNetwork().getLinkCount(true) != 0));
    }

  } 
  
  /***************************************************************************
  **
  ** Command
  */ 
   
  private class WorldBankLayoutAction extends BasicLayoutAction {
     
    private static final long serialVersionUID = 1L;
    
    WorldBankLayoutAction(boolean doIcon) {
      super(doIcon, "command.WorldBankLayout", "command.WorldBankLayoutMnem", BuildDataImpl.BuildMode.WORLD_BANK_LAYOUT);
    }
  }

  /***************************************************************************
  **
  ** Command
  */ 
   
  private class DefaultLayoutAction extends ChecksForEnabled {
    
    private static final long serialVersionUID = 1L;
    
    DefaultLayoutAction(boolean doIcon) {
      ResourceManager rMan = ResourceManager.getManager(); 
      putValue(Action.NAME, rMan.getString("command.DefaultLayout"));
      if (doIcon) {
        putValue(Action.SHORT_DESCRIPTION, rMan.getString("command.DefaultLayout"));        
        URL ugif = getClass().getResource("/org/systemsbiology/biofabric/images/FIXME24.gif");  
        putValue(Action.SMALL_ICON, new ImageIcon(ugif));
      } else {
        char mnem = rMan.getChar("command.DefaultLayoutMnem"); 
        putValue(Action.MNEMONIC_KEY, Integer.valueOf(mnem)); 
      }
    }
        
    public void actionPerformed(ActionEvent e) {
      try {
        Set<NetNode> sels = bfp_.getNodeSelections();
        NetNode selNode = (sels.size() == 1) ? sels.iterator().next() : null;
        BreadthFirstLayoutDialog bfl = new BreadthFirstLayoutDialog(topWindow_, selNode, bfw_.getFabricPanel().getNetwork());
        bfl.setVisible(true);          
        if (bfl.haveResult()) {
          DefaultLayout.Params params = bfl.getParams(); 
          flf_.doDefaultRelayout(params);
        }
      } catch (Exception ex) {
        ExceptionHandler.getHandler().displayException(ex);
      }      
      return;
    }
    
    @Override
    protected boolean checkGuts() {
      return (bfp_.hasAModel() && (bfp_.getNetwork().getLinkCount(true) != 0));
    }   
  }
  
  /***************************************************************************
  **
  ** Command
  */ 
   
  private abstract class BasicLayoutAction extends ChecksForEnabled {
    
    private static final long serialVersionUID = 1L;
    private BuildDataImpl.BuildMode bMode_;
    
    BasicLayoutAction(boolean doIcon, String nameTag, String mnemTag, BuildDataImpl.BuildMode bMode) {
      ResourceManager rMan = ResourceManager.getManager(); 
      putValue(Action.NAME, rMan.getString(nameTag));
      if (doIcon) {
        putValue(Action.SHORT_DESCRIPTION, rMan.getString(nameTag));        
        URL ugif = getClass().getResource("/org/systemsbiology/biofabric/images/FIXME24.gif");  
        putValue(Action.SMALL_ICON, new ImageIcon(ugif));
      } else {
        char mnem = rMan.getChar(mnemTag); 
        putValue(Action.MNEMONIC_KEY, Integer.valueOf(mnem)); 
      }
      bMode_ = bMode;
    }
        
    public void actionPerformed(ActionEvent e) {
      try {
        flf_.doBasicRelayout(bMode_); 
      } catch (Exception ex) {
        ExceptionHandler.getHandler().displayException(ex);
      }      
      return;
    }
    
    @Override
    protected boolean checkGuts() {
      return (bfp_.hasAModel() && (bfp_.getNetwork().getLinkCount(true) != 0));
    }   
  }
  
  
  /***************************************************************************
  **
  ** Command
  */ 
   
  private class LayoutViaConnectivityAction extends ChecksForEnabled {
     
    private static final long serialVersionUID = 1L;
    
    LayoutViaConnectivityAction(boolean doIcon) {
      ResourceManager rMan = ResourceManager.getManager(); 
      putValue(Action.NAME, rMan.getString("command.LayoutViaConnectivity"));
      if (doIcon) {
        putValue(Action.SHORT_DESCRIPTION, rMan.getString("command.LayoutViaConnectivity"));        
        URL ugif = getClass().getResource("/org/systemsbiology/biofabric/images/FIXME24.gif");  
        putValue(Action.SMALL_ICON, new ImageIcon(ugif));
      } else {
        char mnem = rMan.getChar("command.LayoutViaConnectivityMnem"); 
        putValue(Action.MNEMONIC_KEY, Integer.valueOf(mnem)); 
      }
    }
    
    public void actionPerformed(ActionEvent e) {
      try {
        performOperation(null);
      } catch (Exception ex) {
        ExceptionHandler.getHandler().displayException(ex);
      }      
      return;
    }
    
    private boolean performOperation(Object[] args) {
          
      NodeSimilarityLayoutSetupDialog clpd = 
        new NodeSimilarityLayoutSetupDialog(topWindow_, new NodeSimilarityLayout.ClusterParams());  
  
      clpd.setVisible(true);
      if (!clpd.haveResult()) {
        return (false);
      }
  
      flf_.doConnectivityRelayout(clpd.getParams()); 
      return (true);   
    }
    
    @Override
    protected boolean checkGuts() {
      return (bfp_.hasAModel() && (bfp_.getNetwork().getLinkCount(true) != 0));
    }   
  }
  
   /***************************************************************************
  **
  ** Command
  */ 
   
  private class LayoutViaShapeMatchAction extends ChecksForEnabled {
    
    private static final long serialVersionUID = 1L;

    LayoutViaShapeMatchAction(boolean doIcon) {
      ResourceManager rMan = ResourceManager.getManager(); 
      putValue(Action.NAME, rMan.getString("command.LayoutViaShapeMatch"));
      if (doIcon) {
        putValue(Action.SHORT_DESCRIPTION, rMan.getString("command.LayoutViaShapeMatch"));        
        URL ugif = getClass().getResource("/org/systemsbiology/biofabric/images/FIXME24.gif");  
        putValue(Action.SMALL_ICON, new ImageIcon(ugif));
      } else {
        char mnem = rMan.getChar("command.LayoutViaShapeMatchMnem"); 
        putValue(Action.MNEMONIC_KEY, Integer.valueOf(mnem)); 
      }
    }
    
    public void actionPerformed(ActionEvent e) {
      try {
        performOperation(null);
      } catch (Exception ex) {
        ExceptionHandler.getHandler().displayException(ex);
      }      
      return;
    }
    
    private boolean performOperation(Object[] args) {
      
      ReorderLayoutParamsDialog clpd = 
        new ReorderLayoutParamsDialog(topWindow_, new NodeSimilarityLayout.ResortParams());  
  
      clpd.setVisible(true);
      if (!clpd.haveResult()) {
        return (false);
      }
  
      flf_.doShapeMatchRelayout(clpd.getParams());
      return (true);   
    }
    
    @Override
    protected boolean checkGuts() {
      return (bfp_.hasAModel() && (bfp_.getNetwork().getLinkCount(true) != 0));
    }   
  }
  
  /***************************************************************************
  **
  ** Command
  */ 
   
  private class SetLinkGroupsAction extends ChecksForEnabled {
      
    private static final long serialVersionUID = 1L;
    
    SetLinkGroupsAction(boolean doIcon) {
      ResourceManager rMan = ResourceManager.getManager(); 
      putValue(Action.NAME, rMan.getString("command.SetLinkGroups"));
      if (doIcon) {
        putValue(Action.SHORT_DESCRIPTION, rMan.getString("command.SetLinkGroups"));        
        URL ugif = getClass().getResource("/org/systemsbiology/biofabric/images/FIXME24.gif");  
        putValue(Action.SMALL_ICON, new ImageIcon(ugif));
      } else {
        char mnem = rMan.getChar("command.SetLinkGroupsMnem"); 
        putValue(Action.MNEMONIC_KEY, Integer.valueOf(mnem)); 
      }
    }
    
    public void actionPerformed(ActionEvent e) {
      try {
        performOperation(null);
      } catch (Exception ex) {
        ExceptionHandler.getHandler().displayException(ex);
      }      
      return;
    }
    
    private boolean performOperation(Object[] args) {
          
      BioFabricNetwork bfn = bfp_.getNetwork();
      List<String> currentTags = bfn.getLinkGroups();
      boolean showGroupAnnot = bfn.getShowLinkGroupAnnotations();
      ArrayList<NetLink> links = new ArrayList<NetLink>(bfn.getAllLinks(true));
      TreeMap<AugRelation, Boolean> relMap = new TreeMap<AugRelation, Boolean>();
      try {
        PluginSupportFactory.getBuildExtractor().extractRelations(links, relMap, null);
      } catch (AsynchExitRequestException aerx) {
      	// Should not happen...
      }
      Set<AugRelation> allRelations = relMap.keySet(); 
      BioFabricNetwork.LayoutMode currMode = bfn.getLayoutMode();
      LinkGroupingSetupDialog lgsd = 
        new LinkGroupingSetupDialog(topWindow_, currentTags, showGroupAnnot, currMode, allRelations);
      lgsd.setVisible(true);
      if (!lgsd.haveResult()) {
        return (false);
      }

      BioFabricNetwork.LayoutMode mode = lgsd.getChosenMode();
      boolean showLinkAnnotations = lgsd.showLinkAnnotations();

      BuildDataImpl.BuildMode bmode; 
      if (mode == BioFabricNetwork.LayoutMode.PER_NODE_MODE) {
        bmode = BuildDataImpl.BuildMode.GROUP_PER_NODE_CHANGE;
      } else if (mode == BioFabricNetwork.LayoutMode.PER_NETWORK_MODE) {
      	bmode = BuildDataImpl.BuildMode.GROUP_PER_NETWORK_CHANGE;
      } else {
        throw new IllegalStateException();
      }

      flf_.doLinkGroupRelayout(bfn, lgsd.getGroups(), mode, showLinkAnnotations, bmode);
      
      return (true);
    }

    @Override
    protected boolean checkGuts() {
      return (bfp_.hasAModel());
    }  
  }
  

  
  /***************************************************************************
  **
  ** Command
  */ 
   
  private class ImportSIFAction extends ChecksForEnabled {
    
    private static final long serialVersionUID = 1L;
    private boolean doWeights_;
    
    ImportSIFAction(boolean doIcon, boolean doWeights) {
      ResourceManager rMan = ResourceManager.getManager(); 
      putValue(Action.NAME, rMan.getString((doWeights) ? "command.LoadSIFWithWeights" :"command.LoadSIF"));
      if (doIcon) {
        putValue(Action.SHORT_DESCRIPTION, rMan.getString((doWeights) ? "command.LoadSIFWithWeights" :"command.LoadSIF"));        
        URL ugif = getClass().getResource("/org/systemsbiology/biofabric/images/FIXME24.gif");  
        putValue(Action.SMALL_ICON, new ImageIcon(ugif));
      } else {
        char mnem = rMan.getChar((doWeights) ? "command.LoadSIFWithWeightsMnem" :"command.LoadSIFMnem"); 
        putValue(Action.MNEMONIC_KEY, Integer.valueOf(mnem)); 
      }
      doWeights_ = doWeights;
    }
    
    public void actionPerformed(ActionEvent e) {
      try {
        performOperation(null);
      } catch (Exception ex) {
        ExceptionHandler.getHandler().displayException(ex);
      }      
      return;
    }
    
    private boolean performOperation(Object[] args) {
 
      File file = null;
      String filename = FabricCommands.getPreference("LoadDirectory");
      while (file == null) {
        JFileChooser chooser = new JFileChooser();
        FileExtensionFilters.SimpleFilter sf = new FileExtensionFilters.SimpleFilter(".sif", "filterName.sif");
        chooser.addChoosableFileFilter(sf);
        chooser.setAcceptAllFileFilterUsed(true);
        chooser.setFileFilter(sf);

        if (filename != null) {
          File startDir = new File(filename);
          if (startDir.exists()) {
            chooser.setCurrentDirectory(startDir);
          }
        }

        int option = chooser.showOpenDialog(topWindow_);
        if (option != JFileChooser.APPROVE_OPTION) {
          return (true);
        }
        file = chooser.getSelectedFile();
        if (file == null) {
          return (true);
        }
        if (!flf_.standardFileChecks(file, FileLoadFlows.FILE_MUST_EXIST, FileLoadFlows.FILE_CAN_CREATE_DONT_CARE,
                                           FileLoadFlows.FILE_DONT_CHECK_OVERWRITE, FileLoadFlows.FILE_MUST_BE_FILE,
                                           FileLoadFlows.FILE_CAN_WRITE_DONT_CARE, FileLoadFlows.FILE_CAN_READ)) {
          file = null;
          continue;
        }
      }
      Integer magBins = (doWeights_) ? Integer.valueOf(4) : null;
      FileLoadFlows.FileLoadResult flr = flf_.loadFromASource(file, null, magBins, new UniqueLabeller(), 
      		                                                    FileLoadFlows.FileLoadType.SIF, false);
      return (flr.getSuccess());
    }
  }
  
  /***************************************************************************
   **
   ** Command
   */
  
  private class ImportGWAction extends ChecksForEnabled {
  
    private static final long serialVersionUID = 1L;
  
    ImportGWAction(boolean doIcon) {
      ResourceManager rMan = ResourceManager.getManager();
      putValue(Action.NAME, rMan.getString("command.LoadGW"));
      if (doIcon) {
        putValue(Action.SHORT_DESCRIPTION, rMan.getString("command.LoadGW"));
        URL ugif = getClass().getResource("/org/systemsbiology/biofabric/images/FIXME24.gif");
        putValue(Action.SMALL_ICON, new ImageIcon(ugif));
      } else {
        char mnem = rMan.getChar("command.LoadGWMnem");
        putValue(Action.MNEMONIC_KEY, Integer.valueOf(mnem));
      }
    }
  
    public void actionPerformed(ActionEvent e) {
      try {
        performOperation(null);
      } catch (Exception ex) {
        ExceptionHandler.getHandler().displayException(ex);
      }
      return;
    }
  
    private boolean performOperation(Object[] args) {
      File file = null;
      String filename = FabricCommands.getPreference("LoadDirectory");
      while (file == null) {
        JFileChooser chooser = new JFileChooser();
        FileExtensionFilters.SimpleFilter sf = new FileExtensionFilters.SimpleFilter(".gw", "filterName.gw");
        chooser.addChoosableFileFilter(sf);
        chooser.setAcceptAllFileFilterUsed(true);
        chooser.setFileFilter(sf);
    
        if (filename != null) {
          File startDir = new File(filename);
          if (startDir.exists()) {
            chooser.setCurrentDirectory(startDir);
          }
        }
    
        int option = chooser.showOpenDialog(topWindow_);
        if (option != JFileChooser.APPROVE_OPTION) {
          return (true);
        }
        file = chooser.getSelectedFile();
        if (file == null) {
          return (true);
        }
        if (!flf_.standardFileChecks(file, FileLoadFlows.FILE_MUST_EXIST, FileLoadFlows.FILE_CAN_CREATE_DONT_CARE,
                FileLoadFlows.FILE_DONT_CHECK_OVERWRITE, FileLoadFlows.FILE_MUST_BE_FILE,
                FileLoadFlows.FILE_CAN_WRITE_DONT_CARE, FileLoadFlows.FILE_CAN_READ)) {
          file = null;
          continue;
        }
      }
      FileLoadFlows.FileLoadResult flr = flf_.loadFromASource(file, null, null, new UniqueLabeller(), 
      														                            FileLoadFlows.FileLoadType.GW, false);
      return (flr.getSuccess());
    }
  
  }
  
  /***************************************************************************
  **
  ** Command
  */ 
   
  private class LoadXMLAction extends ChecksForEnabled {
    
    private static final long serialVersionUID = 1L;
     
    LoadXMLAction(boolean doIcon) {
      ResourceManager rMan = ResourceManager.getManager(); 
      putValue(Action.NAME, rMan.getString("command.LoadXML"));
      if (doIcon) {
        putValue(Action.SHORT_DESCRIPTION, rMan.getString("command.LoadXML"));        
        URL ugif = getClass().getResource("/org/systemsbiology/biofabric/images/FIXME24.gif");  
        putValue(Action.SMALL_ICON, new ImageIcon(ugif));
      } else {
        char mnem = rMan.getChar("command.LoadXMLMnem"); 
        putValue(Action.MNEMONIC_KEY, Integer.valueOf(mnem)); 
      }
      char accel = rMan.getChar("command.LoadXMLAccel");
      putValue(Action.ACCELERATOR_KEY, KeyStroke.getKeyStroke(accel, Event.CTRL_MASK, false));
    }
    
    public void actionPerformed(ActionEvent e) {
      try {
        performOperation(null);
      } catch (Exception ex) {
        ExceptionHandler.getHandler().displayException(ex);
      }      
      return;
    }
    
    private boolean performOperation(Object[] args) {
 
      File file = null;      
      String filename = FabricCommands.getPreference("LoadDirectory");
      while (file == null) {
        JFileChooser chooser = new JFileChooser(); 
        FileExtensionFilters.SimpleFilter sf = new FileExtensionFilters.SimpleFilter(".bif", "filterName.bif");
        chooser.addChoosableFileFilter(sf);
        chooser.setAcceptAllFileFilterUsed(true);
        chooser.setFileFilter(sf);

        if (filename != null) {
          File startDir = new File(filename);
          if (startDir.exists()) {
            chooser.setCurrentDirectory(startDir);  
          }
        }

        int option = chooser.showOpenDialog(topWindow_);
        if (option != JFileChooser.APPROVE_OPTION) {
          return (true);
        }
        file = chooser.getSelectedFile();
        if (file == null) {
          return (true);
        }
        if (!flf_.standardFileChecks(file, FileLoadFlows.FILE_MUST_EXIST, FileLoadFlows.FILE_CAN_CREATE_DONT_CARE, 
                                           FileLoadFlows.FILE_DONT_CHECK_OVERWRITE, FileLoadFlows.FILE_MUST_BE_FILE, 
                                           FileLoadFlows.FILE_CAN_WRITE_DONT_CARE, FileLoadFlows.FILE_CAN_READ)) {
          file = null;
          continue; 
        }
      }
      File holdIt;  
	    try {
	    	holdIt = File.createTempFile("BioFabricHold", ".zip");
	    	holdIt.deleteOnExit();
	    } catch (IOException ioex) {
	    	holdIt = null;
	    }
      return (flf_.loadXMLFromSource(file, holdIt));
    }
  }

  /***************************************************************************
  **
  ** Command
  */ 
   
  private class LoadWithNodeAttributesAction extends ChecksForEnabled {
     
    private static final long serialVersionUID = 1L;
     
    LoadWithNodeAttributesAction(boolean doIcon) {
      ResourceManager rMan = ResourceManager.getManager(); 
      putValue(Action.NAME, rMan.getString("command.LoadSIFWithNodeAttributes"));
      if (doIcon) {
        putValue(Action.SHORT_DESCRIPTION, rMan.getString("command.LoadSIFWithNodeAttributes"));        
        URL ugif = getClass().getResource("/org/systemsbiology/biofabric/images/FIXME24.gif");  
        putValue(Action.SMALL_ICON, new ImageIcon(ugif));
      } else {
        char mnem = rMan.getChar("command.LoadSIFWithNodeAttributesMnem"); 
        putValue(Action.MNEMONIC_KEY, Integer.valueOf(mnem)); 
      }
    }
    
    public void actionPerformed(ActionEvent e) {
      try {
        performOperation(null);
      } catch (Exception ex) {
        ExceptionHandler.getHandler().displayException(ex);
      }      
      return;
    }    
 
    private boolean performOperation(Object[] args) {
      File file = null;      
      String filename = FabricCommands.getPreference("LoadDirectory");
      while (file == null) {
        JFileChooser chooser = new JFileChooser(); 
        FileExtensionFilters.SimpleFilter sf = new FileExtensionFilters.SimpleFilter(".sif", "filterName.sif");
        chooser.addChoosableFileFilter(sf);
        chooser.setAcceptAllFileFilterUsed(true);
        chooser.setFileFilter(sf);
        if (filename != null) {
          File startDir = new File(filename);
          if (startDir.exists()) {
            chooser.setCurrentDirectory(startDir);  
          }
        }

        int option = chooser.showOpenDialog(topWindow_);
        if (option != JFileChooser.APPROVE_OPTION) {
          return (true);
        }
        file = chooser.getSelectedFile();
        if (file == null) {
          return (true);
        }
        if (!flf_.standardFileChecks(file, FileLoadFlows.FILE_MUST_EXIST, FileLoadFlows.FILE_CAN_CREATE_DONT_CARE, 
                                           FileLoadFlows.FILE_DONT_CHECK_OVERWRITE, FileLoadFlows.FILE_MUST_BE_FILE, 
                                           FileLoadFlows.FILE_CAN_WRITE_DONT_CARE, FileLoadFlows.FILE_CAN_READ)) {
          file = null;
          continue; 
        }
      }
      
      File attribFile = flf_.getTheFile(".noa", ".na", "AttribDirectory", "filterName.noa", null);
      if (attribFile == null) {
        return (true);
      }
  
      Map<AttributeKey, String> attribs = flf_.loadTheFile(attribFile, null, true);
      if (attribs == null) {
        return (true);
      }
 
      FileLoadFlows.FileLoadResult flr = flf_.loadFromASource(file, attribs, null, new UniqueLabeller(), 
      														                            FileLoadFlows.FileLoadType.SIF, false);
      return (flr.getSuccess());
    }
  }
  
  /***************************************************************************
  **
  ** Command
  */ 
    
  private class SaveAsAction extends ChecksForEnabled {

    private static final long serialVersionUID = 1L;
    
    SaveAsAction(boolean doIcon) {
      ResourceManager rMan = ResourceManager.getManager(); 
      putValue(Action.NAME, rMan.getString("command.SaveAs"));
      if (doIcon) {
        putValue(Action.SHORT_DESCRIPTION, rMan.getString("command.SaveAs"));        
        URL ugif = getClass().getResource("/org/systemsbiology/biofabric/images/SaveAs24.gif");  
        putValue(Action.SMALL_ICON, new ImageIcon(ugif));
      } else {     
        char mnem = rMan.getChar("command.SaveAsMnem"); 
        putValue(Action.MNEMONIC_KEY, Integer.valueOf(mnem)); 
      } 
    }
    
    public void actionPerformed(ActionEvent e) {
      try {
        performOperation(null);
      } catch (Exception ex) {
        ExceptionHandler.getHandler().displayException(ex);
      }      
      return;
    }
    
    public boolean performOperation(Object[] args) {
      if (args == null) {
        return (flf_.saveToFile(null));
      } else {
        if (((Boolean)args[0]).booleanValue()) {
          String fileName = (String)args[1];
          return (flf_.saveToFile(fileName));
        } else {
        	OutputStream stream = (OutputStream)args[1];
          BioFabricNetwork bfn = bfp_.getNetwork();     
			    if (bfn.getLinkCount(true) > FileLoadFlows.LINK_COUNT_FOR_BACKGROUND_WRITE) {
			      flf_.doBackgroundWrite(stream);
			      return (true);
			    } else {
			      try {
			        flf_.saveToOutputStream(stream, false, null);
			        return (true);
			      } catch (AsynchExitRequestException aeex) {
			      	// Not on background thread; will not happen
			      	return (false);
			      } catch (IOException ioe) {
			        flf_.displayFileOutputError();
			        return (false);
			      }
			    }
        }
      }
    }
    
    @Override
    protected boolean checkGuts() {
      return (bfp_.hasAModel());
    }      
  }  
 
  /***************************************************************************
  **
  ** Command
  */ 
    
  private class SaveAction extends ChecksForEnabled {

    private static final long serialVersionUID = 1L;
    
    SaveAction(boolean doIcon) {
      ResourceManager rMan = ResourceManager.getManager(); 
      putValue(Action.NAME, rMan.getString("command.Save"));
      if (doIcon) {
        putValue(Action.SHORT_DESCRIPTION, rMan.getString("command.Save"));        
        URL ugif = getClass().getResource("/org/systemsbiology/biofabric/images/Save24.gif");  
        putValue(Action.SMALL_ICON, new ImageIcon(ugif));
      } else {     
        char mnem = rMan.getChar("command.SaveMnem"); 
        putValue(Action.MNEMONIC_KEY, Integer.valueOf(mnem)); 
      }
      char accel = rMan.getChar("command.SaveAccel");
      putValue(Action.ACCELERATOR_KEY, KeyStroke.getKeyStroke(accel, Event.CTRL_MASK, false));     
    }
  
    public void actionPerformed(ActionEvent e) { 
      try { 
        flf_.saveToCurrentFile();
      } catch (Exception ex) {
        ExceptionHandler.getHandler().displayException(ex);
      }
      return;
    }
    
    @Override
    protected boolean checkGuts() {
      return (bfp_.hasAModel());
    }   
   
  }
  
  /***************************************************************************
  **
  ** Command
  */ 
    
  private abstract class ExportOrderAction extends ChecksForEnabled {

    private static final long serialVersionUID = 1L;
    
    ExportOrderAction(boolean doIcon, String name, String mnemStr) {
      ResourceManager rMan = ResourceManager.getManager(); 
      putValue(Action.NAME, rMan.getString(name));
      if (doIcon) {
        putValue(Action.SHORT_DESCRIPTION, rMan.getString(name));        
        URL ugif = getClass().getResource("/org/systemsbiology/biofabric/images/FIXME24.gif");  
        putValue(Action.SMALL_ICON, new ImageIcon(ugif));
      } else {     
        char mnem = rMan.getChar(mnemStr); 
        putValue(Action.MNEMONIC_KEY, Integer.valueOf(mnem)); 
      }
    }
    
    public void actionPerformed(ActionEvent e) {
      try {
        performOperation(null);
      } catch (Exception ex) {
        ExceptionHandler.getHandler().displayException(ex);
      }      
      return;
    }
    
    protected abstract FileFilter getFilter();
    protected abstract List<String> getSuffixList();
    protected abstract String getPrefSuffix();
    protected abstract void writeItOut(File file) throws IOException;
 
    public boolean performOperation(Object[] args) {      
      File file = null;
      String dirName = FabricCommands.getPreference("AttribDirectory");
      while (file == null) {
        JFileChooser chooser = new JFileChooser();
        FileFilter sf = getFilter();
        chooser.addChoosableFileFilter(sf);
        chooser.setAcceptAllFileFilterUsed(true);
        chooser.setFileFilter(sf);
        
        chooser.addChoosableFileFilter(getFilter());    
        if (dirName != null) {
          File startDir = new File(dirName);
          if (startDir.exists()) {
            chooser.setCurrentDirectory(startDir);  
          }
        }  
        int option = chooser.showSaveDialog(topWindow_);
        if (option != JFileChooser.APPROVE_OPTION) {
          return (true);
        }
        file = chooser.getSelectedFile();
        if (file != null) {
          if (!file.exists()) {
            List<String> cand = getSuffixList();
            if (!FileExtensionFilters.hasASuffix(file.getName(), "", cand)) { 
              file = new File(file.getAbsolutePath() + getPrefSuffix());
            }
          }
          if (!flf_.standardFileChecks(file, FileLoadFlows.FILE_MUST_EXIST_DONT_CARE, FileLoadFlows.FILE_CAN_CREATE, 
                                             FileLoadFlows.FILE_CHECK_OVERWRITE, FileLoadFlows.FILE_MUST_BE_FILE, 
                                             FileLoadFlows.FILE_CAN_WRITE, FileLoadFlows.FILE_CAN_READ_DONT_CARE)) {
            file = null;
            continue; 
          }
        }       
      }     
       
      try {
        writeItOut(file);
      } catch (IOException ioe) {
        flf_.displayFileOutputError();
        return (false);
      }
      FabricCommands.setPreference("AttribDirectory", file.getAbsoluteFile().getParent());     
      return (true);
    }
   
    @Override
    protected boolean checkGuts() {
      return (bfp_.hasAModel());
    }
     
  }
  
  /***************************************************************************
  **
  ** Command
  */ 
    
  private class ExportNodeOrderAction extends ExportOrderAction {

    private static final long serialVersionUID = 1L;
      
    ExportNodeOrderAction(boolean doIcon) {
      super(doIcon, "command.ExportNodeOrder", "command.ExportNodeOrderMnem");
    }
    
    protected FileFilter getFilter() {
      return (new FileExtensionFilters.DoubleExtensionFilter(".noa", ".na", "filterName.noa"));
    }
    
    protected List<String> getSuffixList() {
      ArrayList<String> cand = new ArrayList<String>();
      cand.add(".noa");
      cand.add(".na");
      return (cand);
    }
    
    protected String getPrefSuffix() {
      return (".noa");    
    }
    
    protected void writeItOut(File file) throws IOException {
      PrintWriter out = new PrintWriter(new BufferedWriter(new OutputStreamWriter(new FileOutputStream(file), "UTF-8")));        
      bfp_.getNetwork().writeNOA(out);
      out.close();
      return;
    }    
  }
  
  /***************************************************************************
  **
  ** Command
  */ 
    
  private class ExportLinkOrderAction extends ExportOrderAction {
    
    private static final long serialVersionUID = 1L;
    
    ExportLinkOrderAction(boolean doIcon) {
      super(doIcon, "command.ExportLinkOrder", "command.ExportLinkOrderMnem");
    }
    
    protected FileFilter getFilter() {
      return (new FileExtensionFilters.DoubleExtensionFilter(".eda", ".ea", "filterName.eda"));
    }
    
    protected List<String> getSuffixList() {
      ArrayList<String> cand = new ArrayList<String>();
      cand.add(".eda");
      cand.add(".ea");
      return (cand);
    }
    
    protected String getPrefSuffix() {
      return (".eda");    
    }
    
    protected void writeItOut(File file) throws IOException {
      PrintWriter out = new PrintWriter(new BufferedWriter(new OutputStreamWriter(new FileOutputStream(file), "UTF-8")));        
      bfp_.getNetwork().writeEDA(out);
      out.close();
      return;
    }   
  }
  
   /***************************************************************************
  **
  ** Command
  */ 
    
  private class ExportSelectedNodesAction extends ExportOrderAction {
    
    private static final long serialVersionUID = 1L;
    
    ExportSelectedNodesAction(boolean doIcon) {
      super(doIcon, "command.ExportSelectedNodes", "command.ExportSelectedNodesMnem");
    }
    
    protected FileFilter getFilter() {
      return (new FileExtensionFilters.SimpleFilter(".txt", "filterName.txt"));
    }
    
    protected List<String> getSuffixList() {
      ArrayList<String> cand = new ArrayList<String>();
      cand.add(".txt");
      return (cand);
    }
    
    protected String getPrefSuffix() {
      return (".txt");    
    }
    
    protected void writeItOut(File file) throws IOException {
      PrintWriter out = new PrintWriter(new BufferedWriter(new OutputStreamWriter(new FileOutputStream(file), "UTF-8")));        
      Set<NetNode> sels = bfp_.getNodeSelections();
      Iterator<NetNode> sit = sels.iterator();
      while (sit.hasNext()) {
        NetNode node = sit.next();
        out.println(node.getName());
      }
      out.close();
      return;
    } 
    
    @Override
    protected boolean checkGuts() {
      return (super.checkGuts() && !bfp_.getNodeSelections().isEmpty());
    }
    
  }
 
  /***************************************************************************
  **
  ** Command
  */ 
      
  private class PrintAction extends ChecksForEnabled {

    private static final long serialVersionUID = 1L;
    
    PrintAction(boolean doIcon) {
      ResourceManager rMan = ResourceManager.getManager(); 
      putValue(Action.NAME, rMan.getString("command.Print"));
      if (doIcon) {
        putValue(Action.SHORT_DESCRIPTION, rMan.getString("command.Print"));        
        URL ugif = getClass().getResource("/org/systemsbiology/biofabric/images/Print24.gif");  
        putValue(Action.SMALL_ICON, new ImageIcon(ugif));
      } else {
        char mnem = rMan.getChar("command.PrintMnem"); 
        putValue(Action.MNEMONIC_KEY, Integer.valueOf(mnem)); 
      }
      char accel = rMan.getChar("command.PrintAccel");
      putValue(Action.ACCELERATOR_KEY, KeyStroke.getKeyStroke(accel, Event.CTRL_MASK, false));     
    }

    public void actionPerformed(ActionEvent e) {
      try {
        PrinterJob pj = PrinterJob.getPrinterJob();
        PageFormat pf = pj.defaultPage();
        pf.setOrientation(PageFormat.LANDSCAPE);
        // FIX ME: Needed for Win32?  Linux won't default to landscape without this?
        //PageFormat pf2 = pj.pageDialog(pf);
        pj.setPrintable(bfp_, pf);
        if (pj.printDialog()) {
          try {
            pj.print();
          } catch (PrinterException pex) {
            System.err.println(pex);
          }
        }
      } catch (Exception ex) {
        ExceptionHandler.getHandler().displayException(ex);
      }
      return;
    }
    
    @Override
    protected boolean checkGuts() {
      return (bfp_.hasAModel());
    }  
  }
  
  /***************************************************************************
  **
  ** Command: HOLDING OUT OF V2 FOR NOW
  
      
  private class PrintPDFAction extends ChecksForEnabled {

    private static final long serialVersionUID = 1L;
    
    PrintPDFAction(boolean doIcon) {
      ResourceManager rMan = ResourceManager.getManager(); 
      putValue(Action.NAME, rMan.getString("command.PrintPDF"));
      if (doIcon) {
        putValue(Action.SHORT_DESCRIPTION, rMan.getString("command.PrintPDF"));        
        URL ugif = getClass().getResource("/org/systemsbiology/biofabric/images/Print24.gif");  
        putValue(Action.SMALL_ICON, new ImageIcon(ugif));
      } else {
        char mnem = rMan.getChar("command.PrintPDFMnem"); 
        putValue(Action.MNEMONIC_KEY, Integer.valueOf(mnem)); 
      }
      char accel = rMan.getChar("command.PrintPDFAccel");
      putValue(Action.ACCELERATOR_KEY, KeyStroke.getKeyStroke(accel, Event.CTRL_MASK, false));     
    }

    public void actionPerformed(ActionEvent e) {
      try {
        //PrinterJob pj = PrinterJob.getPrinterJob();
        //// PageFormat pf = pj.defaultPage();
        //  pf.setOrientation(PageFormat.LANDSCAPE);
        Properties p = new Properties();
        p.setProperty("PageSize","A5");
       // p.setProperty(PSGraphics2D.ORIENTATION, PageConstants.LANDSCAPE);
       //     ExportDialog export = new ExportDialog();
       //   export.showExportDialog(topWindow_, "Export view as ...", bfp_, "export");
    
        
        Rectangle2D viewInWorld = bfp_.getViewInWorld();
        Dimension viw = new Dimension((int)(viewInWorld.getWidth() / 10.0), (int)(viewInWorld.getHeight() / 10.0));
      //  VectorGraphics g = new PDFGraphics2D(new File("/Users/bill/OutputAOa.pdf"), viw); 
     //   g.setProperties(p); 
     //   g.startExport(); 
     //   bfp_.print(g); 
     //   g.endExport();
       
             
        // FIX ME: Needed for Win32?  Linux won't default to landscape without this?
        //PageFormat pf2 = pj.pageDialog(pf);
      //  pj.setPrintable(bfp_, pf);
      //  if (pj.printDialog()) {
      //    try {
        //    pj.print();
       //   } catch (PrinterException pex) {
        //    System.err.println(pex);
        //  }
      //  }
      } catch (Exception ex) {
        ExceptionHandler.getHandler().displayException(ex);
      }
      return;
    }
    
    @Override
    protected boolean checkGuts() {
      return (bfp_.hasAModel());
    }  
  }
  
  /***************************************************************************
  **
  ** Command
  */ 
    
  private abstract class ExportImageAction extends ChecksForEnabled {

    private static final long serialVersionUID = 1L;
    
    ExportImageAction(boolean doIcon, String res, String mnemStr) {
      ResourceManager rMan = ResourceManager.getManager(); 
      putValue(Action.NAME, rMan.getString(res));
      if (doIcon) {
        putValue(Action.SHORT_DESCRIPTION, rMan.getString(res));        
        URL ugif = getClass().getResource("/org/systemsbiology/biofabric/images/FIXME24.gif");  
        putValue(Action.SMALL_ICON, new ImageIcon(ugif));
      } else {     
        char mnem = rMan.getChar(mnemStr); 
        putValue(Action.MNEMONIC_KEY, Integer.valueOf(mnem)); 
      }
    }
    
    public void actionPerformed(ActionEvent e) {
      try {
        performOperation(null);
      } catch (Exception ex) {
        ExceptionHandler.getHandler().displayException(ex);
      }      
      return;
    }
    
    protected abstract ExportSettingsDialog.ExportSettings getExportSettings();
    
    public boolean performOperation(Object[] args) { 
   
      ExportSettingsDialog.ExportSettings set;
      if (args == null) {
        set = getExportSettings();
        if (set == null) {
          return (true);
        }
      } else {
        set = (ExportSettingsDialog.ExportSettings)args[0];
      }
      
      File file = null;
      OutputStream stream = null;
      
      if (args == null) {  // not headless...      
        List<String> supported = ImageExporter.getSupportedFileSuffixes();
        String filename = FabricCommands.getPreference("ExportDirectory");
        while (file == null) {
          JFileChooser chooser = new JFileChooser(); 
          chooser.addChoosableFileFilter(new FileExtensionFilters.MultiExtensionFilter(supported, "filterName.img"));
          if (filename != null) {
            File startDir = new File(filename);
            if (startDir.exists()) {
              chooser.setCurrentDirectory(startDir);  
            }
          }

          int option = chooser.showSaveDialog(topWindow_);
          if (option != JFileChooser.APPROVE_OPTION) {
            return (true);
          }
          file = chooser.getSelectedFile();
          if (file == null) {
            continue;
          }
          if (!file.exists()) {
            List<String> suffs = ImageExporter.getFileSuffixesForType(set.formatType);
            if (!FileExtensionFilters.hasASuffix(file.getName(), "." , suffs)) {
              file = new File(file.getAbsolutePath() + "." + 
                              ImageExporter.getPreferredSuffixForType(set.formatType));
            }
          }
          if (!flf_.standardFileChecks(file, FileLoadFlows.FILE_MUST_EXIST_DONT_CARE, FileLoadFlows.FILE_CAN_CREATE, 
                                             FileLoadFlows.FILE_CHECK_OVERWRITE, FileLoadFlows.FILE_MUST_BE_FILE, 
                                             FileLoadFlows.FILE_CAN_WRITE, FileLoadFlows.FILE_CAN_READ_DONT_CARE)) {
            file = null;
            continue; 
          }
        }
      } else {
        if (((Boolean)args[1]).booleanValue()) {
          file = new File((String)args[2]);
          if (!flf_.standardFileChecks(file, FileLoadFlows.FILE_MUST_EXIST_DONT_CARE, FileLoadFlows.FILE_CAN_CREATE, 
                                             FileLoadFlows.FILE_CHECK_OVERWRITE, FileLoadFlows.FILE_MUST_BE_FILE, 
                                             FileLoadFlows.FILE_CAN_WRITE, FileLoadFlows.FILE_CAN_READ_DONT_CARE)) {
            return (false);
          }
        } else {
          stream = (OutputStream)args[2];
        }
      }  
         
      try {         
        if (file != null) {
          bfp_.exportToFile(file, set.formatType, set.res, set.zoomVal, set.size);  
        } else {
          bfp_.exportToStream(stream, set.formatType, set.res, set.zoomVal, set.size);  
        }
        if (args == null) {
          FabricCommands.setPreference("ExportDirectory", file.getAbsoluteFile().getParent());
        }  
      } catch (IOException ioe) {
        flf_.displayFileOutputError();
        return (false);
      }
 
      return (true);
    }
    
    @Override
    protected boolean checkGuts() {
      return (bfp_.hasAModel());
    }  
  }
  
  /***************************************************************************
  **
  ** Command
  */ 
    
  private class ExportSimpleAction extends ExportImageAction {
 
    private static final long serialVersionUID = 1L;
    
    ExportSimpleAction(boolean doIcon) {
      super(doIcon, "command.Export", "command.ExportMnem");
    }
   
    protected ExportSettingsDialog.ExportSettings getExportSettings() {
      Rectangle wr = bfp_.getRequiredSize();
      ExportSettingsDialog esd = new ExportSettingsDialog(topWindow_, wr.width, wr.height);
      esd.setVisible(true);
      ExportSettingsDialog.ExportSettings set = esd.getResults();
      return (set); 
    }
  }
    
  
  /***************************************************************************
  **
  ** Command
  */ 
    
  private class ExportPublishAction extends ExportImageAction {
 
    private static final long serialVersionUID = 1L;
    
    ExportPublishAction(boolean doIcon) {
      super(doIcon, "command.ExportPublish", "command.ExportPublishMnem");
    }
   
    protected ExportSettingsDialog.ExportSettings getExportSettings() {
      Rectangle wr = bfp_.getRequiredSize();
      ExportSettingsPublishDialog esd = new ExportSettingsPublishDialog(topWindow_, wr.width, wr.height);
      esd.setVisible(true);
      ExportSettingsDialog.ExportSettings set = esd.getResults();
      return (set); 
    }
  }
  
  /***************************************************************************
  **
  ** Command
  */ 
   
  private class EmptyNetworkAction extends ChecksForEnabled {
     
    private static final long serialVersionUID = 1L;
    
    EmptyNetworkAction(boolean doIcon) {
      ResourceManager rMan = ResourceManager.getManager(); 
      putValue(Action.NAME, rMan.getString("command.EmptyNet"));
      if (doIcon) {
        putValue(Action.SHORT_DESCRIPTION, rMan.getString("command.EmptyNet"));        
        URL ugif = getClass().getResource("/org/systemsbiology/biofabric/images/FIXME24.gif");  
        putValue(Action.SMALL_ICON, new ImageIcon(ugif));
      } else {
        char mnem = rMan.getChar("command.EmptyNetMnem"); 
        putValue(Action.MNEMONIC_KEY, Integer.valueOf(mnem)); 
      }
    }
    
    public void actionPerformed(ActionEvent e) {
      try {
        flf_.manageWindowTitle(null);
        flf_.buildEmptyNetwork();
      } catch (Exception ex) {
        ExceptionHandler.getHandler().displayException(ex);
      }      
      return;
    }
  }
  
  /***************************************************************************
  **
  ** Command
  */ 
    
  private class SetDisplayOptionsAction extends ChecksForEnabled {
    
    private static final long serialVersionUID = 1L;
    
    SetDisplayOptionsAction(boolean doIcon) {
      ResourceManager rMan = ResourceManager.getManager(); 
      putValue(Action.NAME, rMan.getString("command.SetDisplayOpts"));
      if (doIcon) {
        putValue(Action.SHORT_DESCRIPTION, rMan.getString("command.SetDisplayOpts"));        
        URL ugif = getClass().getResource("/org/systemsbiology/biofabric/images/FIXME.gif");  
        putValue(Action.SMALL_ICON, new ImageIcon(ugif));
      } else {
        char mnem = rMan.getChar("command.SetDisplayOptsMnem");
        putValue(Action.MNEMONIC_KEY, Integer.valueOf(mnem));
      }   
    }

    public void actionPerformed(ActionEvent e) {
      try {
        FabricDisplayOptionsDialog dod = new FabricDisplayOptionsDialog(topWindow_);
        dod.setVisible(true);
        if (dod.haveResult()) {
          FabricDisplayOptionsManager dopmgr = FabricDisplayOptionsManager.getMgr();
          dopmgr.setDisplayOptions(dod.getNewOpts(), dod.needsRebuild(), dod.needsRecolor());
        }
      } catch (Exception ex) {
        ExceptionHandler.getHandler().displayException(ex);
      }
      return;
    }
  } 

  /***************************************************************************
  **
  ** Command
  */ 
    
  private class PlugInAction extends ChecksForEnabled {
    
    private static final long serialVersionUID = 1L;
    private BioFabricToolPlugInCmd cmd_;
   
    PlugInAction(BioFabricToolPlugInCmd cmd) {
      cmd_ = cmd;
      putValue(Action.NAME, cmd_.getCommandName());
    }

    public void actionPerformed(ActionEvent e) {
      try {
        cmd_.performOperation(topWindow_);
      } catch (Exception ex) {
        ExceptionHandler.getHandler().displayException(ex);
      }
      return;
    }
    
    @Override
    protected boolean checkGuts() {
      return (cmd_.isEnabled());      
    }
  }
  
  /***************************************************************************
  **
  ** Command
  */
  
  public class AboutAction extends ChecksForEnabled {
    
    private static final long serialVersionUID = 1L;
    private URL aboutURL_;
    private JEditorPane pane_;
    private JFrame frame_;
    private FixedJButton buttonB_;
    private URL gnuUrl_;
    private URL sunUrl_;
        
    AboutAction(boolean doIcon) {
      ResourceManager rMan = ResourceManager.getManager(); 
      putValue(Action.NAME, rMan.getString("command.About"));
      if (doIcon) {
        putValue(Action.SHORT_DESCRIPTION, rMan.getString("command.About"));
        URL ugif = getClass().getResource("/org/systemsbiology/biofabric/images/About24.gif");  
        putValue(Action.SMALL_ICON, new ImageIcon(ugif));
      } else {
        char mnem = rMan.getChar("command.AboutMnem"); 
        putValue(Action.MNEMONIC_KEY, Integer.valueOf(mnem));
      }
      aboutURL_ = getClass().getResource("/org/systemsbiology/biofabric/license/about.html");
    }
    
    //
    // Having an image link in the html turns out to be problematic
    // starting Fall 2008 with URL security holes being plugged.  So
    // change the window.  Note we use a back button now too!
    
    public void actionPerformed(ActionEvent e) {
      try {
        if (frame_ != null) {      
          frame_.setExtendedState(JFrame.NORMAL);
          frame_.toFront();
          return;
        }
        try {
          pane_ = new JEditorPane(aboutURL_);
        } catch (IOException ioex) {
          ExceptionHandler.getHandler().displayException(ioex);
          return;
        }
        // 8/09: COMPLETELY BOGUS, but URLs are breaking everywhere in the latest JVMs, an I don't
        // have time to fix this in a more elegant fashion!
        gnuUrl_ = getClass().getResource("/org/systemsbiology/biofabric/license/LICENSE");
        sunUrl_ = getClass().getResource("/org/systemsbiology/biofabric/license/LICENSE-SUN");
        ResourceManager rMan = ResourceManager.getManager();
        pane_.setEditable(false);
        frame_ = new JFrame(rMan.getString("window.aboutTitle"));
        pane_.addHyperlinkListener(new HyperlinkListener() {
          public void hyperlinkUpdate(HyperlinkEvent ev) {
            try {
              if (ev.getEventType() == HyperlinkEvent.EventType.ACTIVATED) {
                URL toUse = (ev.getDescription().indexOf("-SUN") != -1) ? sunUrl_ : gnuUrl_;
                pane_.setPage(toUse);
                buttonB_.setEnabled(true);
              }
            } catch (IOException ex) {
            }
          }
        });
        frame_.addWindowListener(new WindowAdapter() {
          @Override
          public void windowClosing(WindowEvent e) {
            frame_ = null;
            e.getWindow().dispose();
          }
        });
               
        JPanel cp = (JPanel)frame_.getContentPane();
        cp.setBackground(Color.white);   
        cp.setBorder(new EmptyBorder(20, 20, 20, 20));
        cp.setLayout(new GridBagLayout());
        GridBagConstraints gbc = new GridBagConstraints();    
        // URL sugif = getClass().getResource(
        //  "/org/systemsbiology/biotapestry/images/BioTapestrySplash.gif");
        JLabel label = new JLabel(); //new ImageIcon(sugif));        
        
        UiUtil.gbcSet(gbc, 0, 0, 1, 3, UiUtil.BO, 0, 0, 5, 5, 5, 5, UiUtil.CEN, 1.0, 0.0);
        cp.add(label, gbc);
        
        JScrollPane jsp = new JScrollPane(pane_);
        UiUtil.gbcSet(gbc, 0, 3, 1, 2, UiUtil.BO, 0, 0, 5, 5, 5, 5, UiUtil.CEN, 1.0, 1.0);
        cp.add(jsp, gbc);
        
        
        buttonB_ = new FixedJButton(rMan.getString("dialogs.back"));
        buttonB_.setEnabled(false);
        buttonB_.addActionListener(new ActionListener() {
          public void actionPerformed(ActionEvent ev) {
            try {
              pane_.setPage(aboutURL_);
              buttonB_.setEnabled(false);
            } catch (Exception ex) {
              ExceptionHandler.getHandler().displayException(ex);
            }
          }
        });     
        FixedJButton buttonC = new FixedJButton(rMan.getString("dialogs.close"));
        buttonC.addActionListener(new ActionListener() {
          public void actionPerformed(ActionEvent ev) {
            try {
              frame_.setVisible(false);
              frame_.dispose();
              frame_ = null;
            } catch (Exception ex) {
              ExceptionHandler.getHandler().displayException(ex);
            }
          }
        });
        Box buttonPanel = Box.createHorizontalBox();
        buttonPanel.add(Box.createHorizontalGlue()); 
        buttonPanel.add(buttonB_);
        buttonPanel.add(Box.createHorizontalStrut(10));    
        buttonPanel.add(buttonC);
        UiUtil.gbcSet(gbc, 0, 5, 1, 1, UiUtil.HOR, 0, 0, 5, 5, 5, 5, UiUtil.SE, 1.0, 0.0);
        cp.add(buttonPanel, gbc);        
        frame_.setSize(700, 700);
        frame_.setLocationRelativeTo(topWindow_);
        frame_.setVisible(true);
      } catch (Exception ex) {
        ExceptionHandler.getHandler().displayException(ex);
      }
      return;
    }
  }
   

}
```
