## Supplementary material for "BioFabric Visualization of Network Alignments": BioFabric Source Code: HeadlessOracle.java

```
/*
**    Copyright (C) 2003-2018 Institute for Systems Biology 
**                            Seattle, Washington, USA. 
**
**    This library is free software; you can redistribute it and/or
**    modify it under the terms of the GNU Lesser General Public
**    License as published by the Free Software Foundation; either
**    version 2.1 of the License, or (at your option) any later version.
**
**    This library is distributed in the hope that it will be useful,
**    but WITHOUT ANY WARRANTY; without even the implied warranty of
**    MERCHANTABILITY or FITNESS FOR A PARTICULAR PURPOSE.  See the GNU
**    Lesser General Public License for more details.
**
**    You should have received a copy of the GNU Lesser General Public
**    License along with this library; if not, write to the Free Software
**    Foundation, Inc., 59 Temple Place, Suite 330, Boston, MA  02111-1307  USA
*/

package org.systemsbiology.biofabric.cmd;

/****************************************************************************
**
** Answers questions and consumes error messages for headless operation 
*/

public class HeadlessOracle {
 
  ////////////////////////////////////////////////////////////////////////////
  //
  // PRIVATE MEMBERS
  //
  ////////////////////////////////////////////////////////////////////////////   
  
  ////////////////////////////////////////////////////////////////////////////
  //
  // PUBLIC CONSTRUCTORS
  //
  ////////////////////////////////////////////////////////////////////////////    
  
  /***************************************************************************
  ** 
  ** Constructor
  */ 
  
  public HeadlessOracle() {
    
  }  

  ////////////////////////////////////////////////////////////////////////////
  //
  // PUBLIC METHODS
  //
  ////////////////////////////////////////////////////////////////////////////
  
  /***************************************************************************
  ** 
  ** In-process processing entry point.  This method is synchronized internally
  ** on a global basis to insure one process call at a time.
  */

  public void displayErrorMessage(String message) {
    System.err.println(message);
    return;
  }    
}
```
