## Supplementary material for "BioFabric Visualization of Network Alignments": BioFabric Source Code: ZoomCommandSupport.java

```
/*
**    Copyright (C) 2003-2018 Institute for Systems Biology 
**                            Seattle, Washington, USA. 
**
**    This library is free software; you can redistribute it and/or
**    modify it under the terms of the GNU Lesser General Public
**    License as published by the Free Software Foundation; either
**    version 2.1 of the License, or (at your option) any later version.
**
**    This library is distributed in the hope that it will be useful,
**    but WITHOUT ANY WARRANTY; without even the implied warranty of
**    MERCHANTABILITY or FITNESS FOR A PARTICULAR PURPOSE.  See the GNU
**    Lesser General Public License for more details.
**
**    You should have received a copy of the GNU Lesser General Public
**    License along with this library; if not, write to the Free Software
**    Foundation, Inc., 59 Temple Place, Suite 330, Boston, MA  02111-1307  USA
*/

package org.systemsbiology.biofabric.cmd;

import java.awt.Point;
import java.awt.Dimension;
import javax.swing.JScrollPane;
import javax.swing.JScrollBar;
import javax.swing.JViewport;
import java.awt.Rectangle;
import java.awt.event.ComponentAdapter;
import java.awt.event.ComponentEvent;
import java.awt.event.HierarchyEvent;
import java.awt.event.HierarchyListener;
import java.awt.geom.Point2D;

import javax.swing.event.ChangeEvent;
import javax.swing.event.ChangeListener;

import org.systemsbiology.biofabric.api.util.ExceptionHandler;
import org.systemsbiology.biofabric.ui.display.BioFabricPanel;
import org.systemsbiology.biofabric.util.UiUtil;


/****************************************************************************
**
** Handles zoom operations
*/

public class ZoomCommandSupport {
  

  ////////////////////////////////////////////////////////////////////////////
  //
  // MEMBERS
  //
  ////////////////////////////////////////////////////////////////////////////  

  private ZoomTarget sup_;
  private JScrollPane jsp_;
  private double[] zoomVals_;
  private int[] zooms_;
  private int currZoomIndex_;
  private double customZoom_;
  private ZoomResult fullModelZoom_;
  
  private Dimension currViewSize_;
  private double currViewXFrac_;
  private double currViewYFrac_;
  private ZoomChangeTracker tracker_;

  ////////////////////////////////////////////////////////////////////////////
  //
  // CONSTRUCTOR
  //
  ////////////////////////////////////////////////////////////////////////////
  
  /***************************************************************************
  **
  ** Constructor 
  */ 
  
  public ZoomCommandSupport(ZoomChangeTracker tracker) {
    tracker_ = tracker;
  }  
 
  ////////////////////////////////////////////////////////////////////////////
  //
  // PUBLIC METHODS
  //
  ////////////////////////////////////////////////////////////////////////////    

  /***************************************************************************
  **
  ** Set zooms (used to drive quadtree depth)
  */

  public void setZoomLevels(int[] zooms) {
    zooms_ = zooms;
    return;
  }
  
  /***************************************************************************
  **
  ** Get zooms (used to drive quadtree depth on recolor operations)
  */

  public int[] getZoomLevels() {
    return (zooms_);
  }
  
  /***************************************************************************
  **
  ** Set zoom vals (do only at the start)
  */

  public void setZoomPoints(double[] zoomVals) {
    zoomVals_ = zoomVals;
    currZoomIndex_ = 0;
    customZoom_ = 0.0;
    fullModelZoom_ = null;
    return;
  }
  
  /***************************************************************************
  **
  ** Get zoom indices
  */

  public int[] getZoomIndices() {
  	int len = (zoomVals_ == null) ? 0 : zoomVals_.length;
  	int[] retval = new int[len];  	
  	for (int i = 0; i < len; i++) {
  		retval[i] = i;
  	}
    return (retval);
  }

  /***************************************************************************
  **
  ** Register the scroll pane
  */

  public void registerScrollPaneAndZoomTarget(JScrollPane jsp, ZoomTarget sup) {
    jsp_ = jsp;
    sup_ = sup;
    //
    // Need to make view adjustments when the scroll pane size changes:
    //
    jsp.addComponentListener(new ComponentAdapter() {
      @Override
      public void componentResized(ComponentEvent e) {
        try {
          handlePanelResize();
        } catch (Exception ex) {
          ExceptionHandler.getHandler().displayException(ex);
        }
      }
    });
    //
    // Always keep track of where the view center is.  When we resize, we
    // maintain that center:
    //    
    ChangeListener barChange = new ChangeListener() {
      public void stateChanged(ChangeEvent e) {
        try {
          trackScrollBars();
        } catch (Exception ex) {
          ExceptionHandler.getHandler().displayException(ex);
        }
      }
    };
    
    //
    // Having scroll bars come and go makes the actual view extent
    // change without any window size change: this happens with
    // zooming, causing bug #16. Track when it comes and goes:
    
    HierarchyListener hChange = new HierarchyListener() {
      public void hierarchyChanged(HierarchyEvent e) {       
        try {
          if ((e.getID() == HierarchyEvent.HIERARCHY_CHANGED) && 
              ((e.getChangeFlags() & HierarchyEvent.SHOWING_CHANGED) != 0x00)) {
            checkBarVisible();
          }
        } catch (Exception ex) {
          ExceptionHandler.getHandler().displayException(ex);
        }
      }
    };
        
    JScrollBar vsb = jsp.getVerticalScrollBar();
    vsb.getModel().addChangeListener(barChange);
    vsb.addHierarchyListener(hChange);
    JScrollBar hsb = jsp.getHorizontalScrollBar();     
    hsb.getModel().addChangeListener(barChange); 
    hsb.addHierarchyListener(hChange);
    return;
  }
  
  /***************************************************************************
  **
  ** Figure out what zoom value to use for a given model
  */ 

  private ZoomResult calcOptimalZoom(Rectangle chosen, Dimension allDim) {
    
    Dimension dim;
    if (chosen == null) {
      dim = sup_.getBasicSize();
    } else {
      dim = new Dimension(chosen.width, chosen.height);
    }
    
    int vIndex = -1;
    int hIndex = -1;
    if (zoomVals_ != null) { // Used before this is set for full model zoom calc
	    for (int i = 0; i < zoomVals_.length; i++) {
	      double zoom = zoomVals_[i];
	      double modelHeight = dim.height * zoom;
	      if (modelHeight <= allDim.height) {
	        vIndex++;
	      } else {
	        break;
	      }
	    }
	    for (int i = 0; i < zoomVals_.length; i++) {
	      double zoom = zoomVals_[i];
	      double modelWidth = dim.width * zoom;
	      if (modelWidth <= allDim.width) {
	        hIndex++;
	      } else {
	        break;
	      }
	    }
    }
    int retval = (vIndex < hIndex) ? vIndex : hIndex;
    if (retval == -1) {
      double vZoom = (double)allDim.height / (double)dim.height;
      double hZoom = (double)allDim.width / (double)dim.width;
      double useZoom = (vZoom > hZoom) ? hZoom : vZoom;
      return (new ZoomResult(useZoom));
    }
   
    retval = (retval > zoomVals_.length - 1) ? zoomVals_.length - 1 : retval;
    return (new ZoomResult(retval));
  }  

  
  /***************************************************************************
  **
  ** do the zoom operations
  */ 

  private void doZoom(double newZoomVal) { 
     
    JViewport view = jsp_.getViewport();
    Point viewPos = view.getViewPosition();
    Dimension viewDim = view.getExtentSize();

    int vCenter = viewPos.y + (viewDim.height / 2);
    int hCenter = viewPos.x + (viewDim.width / 2);    
    Point2D worldCenter = sup_.viewToWorld(new Point(hCenter, vCenter)); 
    sup_.setZoomFactor(newZoomVal, viewDim);    
    Point newCenter = sup_.pointToViewport(new Point((int)Math.round(worldCenter.getX()), 
                                                     (int)Math.round(worldCenter.getY()))); 
    viewportUpdate(newCenter, viewDim);    
    return;
  }

  /***************************************************************************
  **
  ** Bound viewport position
  */ 
    
  private Point doBounding(int newH, int newV, JViewport view) {
    //
    // Don't let the value go outide actual dimensions.  Note that with wide views, the
    // preferred size is smaller than the actual view:
    //    
    Dimension allDim = view.getViewSize();
    Dimension viewDim = view.getExtentSize();
    int delH = (allDim.width - viewDim.width);
    int delV = (allDim.height - viewDim.height);
    if (newH < 0) {
      newH = 0;
    } else if ((delH > 0) && (newH > delH)) {
      newH = delH;
    }    
    if (newV < 0) {
      newV = 0;
    } else if ((delV > 0) && (newV > delV)) {
      newV = delV;
    }  
    return (new Point(newH, newV));
  }  

  /***************************************************************************
  **
  ** Common viewport update ops
  */ 
    
  private void viewportUpdate(Point center, Dimension vDim) {
    JViewport view = jsp_.getViewport(); 
    view.setViewSize(sup_.getPreferredSize());
    
    int newV = center.y - (vDim.height / 2);
    int newH = center.x - (vDim.width / 2);
    view.setViewPosition(doBounding(newH, newV, view));
    view.invalidate();
    jsp_.validate();
    return;
  }  
   
  /***************************************************************************
  **
  ** Get current zoom value
  */ 
    
  private double getCurrentZoom() {
    return ((currZoomIndex_ == -1) ? customZoom_ : zoomVals_[currZoomIndex_]);
  }
 
  /***************************************************************************
  **
  ** Bump zoom
  */ 
    
  public void bumpZoomWrapper(char direction) {  
    bumpZoom(direction);
    double newZoomVal = getCurrentZoom();

    //
    // Enable/disable zoom actions based on zoom limits:
    //

    if (tracker_ != null) tracker_.zoomStateChanged(false);
    doZoom(newZoomVal);
    return;
  }
  
  /***************************************************************************
  **
  ** Set current zoom value
  */ 
    
  public void setZoomIndex(int index) {
    setCurrentZoom(new ZoomResult(index));
    double newZoomVal = getCurrentZoom();
    //
    // Enable/disable zoom actions based on zoom limits:
    //
    if (tracker_ != null) tracker_.zoomStateChanged(false);
    doZoom(newZoomVal);
    return;
  }
  
  /***************************************************************************
  **
  ** Bump zoom value
  */ 
    
  private void bumpZoom(char direction) {
    // Sanity check:
    if ((currZoomIndex_ < -1) || (zoomVals_ == null) || (currZoomIndex_ >= zoomVals_.length)) {
      return;
    }
    if (direction == '+') {
      if (currZoomIndex_ == -1) {
        setCurrentZoom(new ZoomResult(0));
      } else if (zoomIsMax()) {  // Safety net; should not happen
        return;
      } else {
        setCurrentZoom(new ZoomResult(currZoomIndex_ + 1));
      }
    } else {
      if (currZoomIndex_ == -1) {
      	if (customZoom_ > fullModelZoom_.customZoom) {
      		setCurrentZoom(fullModelZoom_);
      	}
      } else if (currZoomIndex_ == 0) {
  	    if (fullModelZoom_ == null) {
  	    	UiUtil.fixMePrintout("Seeing this as null after recolor while zoomed!!");
  	    	UiUtil.fixMePrintout("And zoom out ");
  	    	Rectangle bounds = sup_.getCurrentBasicBounds();    
          Dimension vDim = jsp_.getViewport().getExtentSize();
          fullModelZoom_ = calcOptimalZoom(bounds, vDim);
  	    }
        setCurrentZoom(fullModelZoom_);
      } else {
        setCurrentZoom(new ZoomResult(currZoomIndex_ - 1));
      }      
    }
  }    

  /***************************************************************************
  **
  ** May not want to allow a "wide" zoom:
  */ 
    
  public boolean zoomIsFirstDefined() {
    return (currZoomIndex_ == 0);
  }  
  
  /***************************************************************************
  **
  ** Answer if we are max
  */ 
    
  public boolean zoomIsMax() {
    return (currZoomIndex_ == (zoomVals_.length - 1));
  }    

  /***************************************************************************
  **
  ** Ask if we can still zoom out:
  */ 
    
  public boolean canZoomOut() { 
  	if (currZoomIndex_ > 0) {
  		return (true); // Easy case!
  	}
  	//
  	// We can still zoom out even if we are *at* zero index if the full model zoom is less than index zero:
  	//
  	if ((currZoomIndex_ == 0) && (zoomVals_ != null) && (zoomVals_.length > 0) && (fullModelZoom_.customZoom < zoomVals_[0])) {
  		return (true);
  	}
  	//
  	// We can still zoom out if we are on a custom zoom, if the fullModelZoom is less than the current custom zoom:
  	//
  	if ((currZoomIndex_ == -1) && (fullModelZoom_.customZoom < customZoom_)) {
  		return (true);
  	}
  	return (false);
  }  
 
  /***************************************************************************
  **
  ** Set current zoom value
  */ 
    
  private void setCurrentZoom(ZoomResult zres) {
    if (zres.doCustom) {
      customZoom_ = zres.customZoom;
      currZoomIndex_ = -1;
    } else {
      currZoomIndex_ = zres.index;
      customZoom_ = 0.0;  // Doesn't really matter...
    }
    return;
  }  
  
  /***************************************************************************
  **
  ** Zoom to a rectangle operations
  */ 
    
  public void zoomToRect(Rectangle bounds) {
    zoomToSelectedGutsGuts(bounds, true);
    return;
  }  

  /***************************************************************************
  **
  ** Zoom to selected operations
  */ 
    
  public void zoomToSelected() {
    Rectangle selected = sup_.getSelectedBounds();
    zoomToRect(selected);
    return;
  }  
  
  /***************************************************************************
  **
  ** Center on rectangle operations
  */ 
    
  public void centerOnRectangle(Rectangle centerRect) {
    centerOnSelectedGuts(centerRect);
    return;
  }  
 
  /***************************************************************************
  **
  ** Center on selected operations
  */ 
    
  public void centerOnSelected() {
    Rectangle selected = sup_.getSelectedBounds();
    centerOnSelectedGuts(selected);
    return;
  }  
 
  /***************************************************************************
  **
  ** Zoom to selected operations
  */ 
    
  private void zoomToSelectedGutsGuts(Rectangle bounds, boolean doZoom) {
    if (bounds == null) {
      zoomToModel(false);
      return;
    }
    
    //
    // Previous BioTapestry-driven approach supported having network elements
    // outside of the workspace. So a zoom partially outside of the workspace
    // would show selected elements. This is not going to happen in BioFabric,
    // and the feature meant that a zoom rect that was larger than some workspace
    // dimensions would not zoom (think fat rectangle around a piece of a long
    // thin network). Use the intersection rectangle instead to fix this problem,
    // instead of the previous solution that made the vertical scrollbar useless
    // on long thin networks!
    //
    
    Rectangle wsBounds = sup_.getWorldRect();
    bounds = wsBounds.intersection(bounds);
    if (bounds.isEmpty()) {
    	return;
    }
       
    Dimension vDim = jsp_.getViewport().getExtentSize();
    if (doZoom) {
      ZoomResult zres = calcOptimalZoom(bounds, vDim);       	
      setCurrentZoom(zres);
      sup_.setZoomFactor(getCurrentZoom(), vDim);
    }
 
    int x = bounds.x + (bounds.width / 2);
    int y = bounds.y + (bounds.height / 2);    
    Point pt = new Point(x, y);
    pt = sup_.pointToViewport(pt);
    viewportUpdate(pt, vDim);
    sup_.repaint();
    if (tracker_ != null) tracker_.zoomStateChanged(false);
    return;
  }  

  /***************************************************************************
  **
  ** Center on selected operations
  */ 
    
  private void centerOnSelectedGuts(Rectangle selected) {
    zoomToSelectedGutsGuts(selected, false);
  } 
  
  /***************************************************************************
  **
  ** Center to selected operations
  */ 
    
  public void centerToNextSelected() {
    sup_.incrementToNextSelection();
    Rectangle selected = sup_.getCurrentSelectedBounds();
    centerOnSelectedGuts(selected);
    return;
  }
  
  /***************************************************************************
  **
  ** Center to selected operations
  */ 
    
  public void centerToPreviousSelected() {
    sup_.decrementToPreviousSelection();
    Rectangle selected = sup_.getCurrentSelectedBounds();
    centerOnSelectedGuts(selected);
    return;
  } 
  
  /***************************************************************************
  **
  ** Zoom to selected operations
  */ 
    
  public void zoomToCurrentSelected() {
    Rectangle selected = sup_.getCurrentSelectedBounds();
    zoomToRect(selected);
    return;
  } 

  /***************************************************************************
  **
  ** If we need to display a union of model bounds and worksheet (model is
  ** off edges of worksheet), we need to increase the union to be centered
  ** on the worksheet.
  */ 
    
  private Rectangle centeredUnion(Rectangle union, Point2D center) {
    int cenX = (int)Math.round(center.getX());
    int cenY = (int)Math.round(center.getY());
    int delmx = cenX - union.x;
    int delmy = cenY - union.y;
    int delpx = (union.x + union.width) - cenX;
    int delpy = (union.y + union.height) - cenY;    
    int rX = (delpx > delmx) ? cenX - delpx : cenX - delmx;
    int rW = (delpx > delmx) ? (2 * delpx) : (2 * delmx);
    int rY = (delpy > delmy) ? cenY - delpy : cenY - delmy;
    int rH = (delpy > delmy) ? (2 * delpy) : (2 * delmy);    
   
    return (new Rectangle(rX, rY, rW, rH));
  }
  
  /***************************************************************************
  **
  ** Zoom to show current model.
  */ 

  public void zoomToModel(boolean force) {
  	Rectangle bounds = sup_.getCurrentBasicBounds();    
    Dimension vDim = jsp_.getViewport().getExtentSize();
  	if ((fullModelZoom_ == null) || force) {
      fullModelZoom_ = calcOptimalZoom(bounds, vDim);
  	}
    setCurrentZoom(fullModelZoom_);    
    sup_.setZoomFactor(getCurrentZoom(), vDim);
    ((BioFabricPanel)sup_).setFullModelViewPos(jsp_.getViewport().getViewPosition());
    ((BioFabricPanel)sup_).setFullModelExtent(vDim);
  
    int x = bounds.x + (bounds.width / 2);
    int y = bounds.y + (bounds.height / 2);    
    Point pt = new Point(x, y);
    pt = sup_.pointToViewport(pt);
    viewportUpdate(pt, vDim);
    sup_.repaint();
    if (tracker_ != null) tracker_.zoomStateChanged(false);
    return;
  }

  /***************************************************************************
  **
  ** Changes to panel size result in changes to world->view mappings, so we
  ** do this here:
  */ 
    
  public void handlePanelResize() { 
    JViewport view = jsp_.getViewport();     
    Dimension viewExtent = view.getExtentSize();
    Dimension viewSize = view.getViewSize(); 
    
    // Keep track of what the full model zoom is at all times:
    Rectangle bounds = sup_.getCurrentBasicBounds();    
    fullModelZoom_ = calcOptimalZoom(bounds, viewExtent);

    int currX = (int)Math.round((currViewXFrac_ * viewSize.width) - (viewExtent.width / 2.0));
    int currY = (int)Math.round((currViewYFrac_ * viewSize.height) - (viewExtent.height / 2.0));
    view.setViewPosition(doBounding(currX, currY, view));            

    //
    // When the viewport size exceeds the worksheet area, we need to change the 
    // world->viewport transform when viewport size changes:
    // 
    sup_.adjustWideZoomForSize(viewExtent);
    currViewSize_ = viewExtent;
    if (tracker_ != null) tracker_.zoomStateChanged(true);
    return;
  }

  /***************************************************************************
  **
  ** Always track the changes in the scroll bars.  We are NOT interested
  ** in any changes due to resizes, so we toss out results when the extent
  ** size is changing:
  */ 
    
  public void trackScrollBars() { 
    
    JViewport view = jsp_.getViewport();
    Dimension viewExtent = view.getExtentSize();
    if (currViewSize_ == null) {
      currViewSize_ = viewExtent;
    }
    
    if (viewExtent.equals(currViewSize_)) {
      Point viewPos = view.getViewPosition();
      int currX = viewPos.x + (viewExtent.width / 2);
      int currY = viewPos.y + (viewExtent.height / 2);
      Dimension viewSize = view.getViewSize(); 
      currViewXFrac_ = (double)currX / (double)viewSize.width; 
      currViewYFrac_ = (double)currY / (double)viewSize.height;
    }
    if (tracker_ != null) tracker_.zoomStateChanged(true);
    return;
  }
  
  /***************************************************************************
  **
  ** We need to know when scroll bar visibility changes, since we need to
  ** always have an accurate read on extent size (Fixes issue #16)
  */ 
    
  public void checkBarVisible() {    
    JViewport view = jsp_.getViewport();
    currViewSize_ = view.getExtentSize();
    return;
  }

  ////////////////////////////////////////////////////////////////////////////
  //
  // INNER CLASSES
  //
  ////////////////////////////////////////////////////////////////////////////
  
  /***************************************************************************
  **
  ** Zoom calculation result
  */
  
  private class ZoomResult {
    boolean doCustom;
    int index;
    double customZoom;
    
    ZoomResult(int index) {
      this.doCustom = false;
      this.index = index;
    }
    
    ZoomResult(double customZoom) {
      this.doCustom = true;
      this.customZoom = customZoom;
    }    
   
    public String toString() {
      return ((doCustom) ? ("ZoomResult customZoom = " + customZoom) : ("ZoomResult indexed zoom = " + index));  
    }
  
  }
}
```
