## Supplementary material for "BioFabric Visualization of Network Alignments": BioFabric Source Code: ZoomTarget.java

```
/*
**    Copyright (C) 2003-2016 Institute for Systems Biology 
**                            Seattle, Washington, USA. 
**
**    This library is free software; you can redistribute it and/or
**    modify it under the terms of the GNU Lesser General Public
**    License as published by the Free Software Foundation; either
**    version 2.1 of the License, or (at your option) any later version.
**
**    This library is distributed in the hope that it will be useful,
**    but WITHOUT ANY WARRANTY; without even the implied warranty of
**    MERCHANTABILITY or FITNESS FOR A PARTICULAR PURPOSE.  See the GNU
**    Lesser General Public License for more details.
**
**    You should have received a copy of the GNU Lesser General Public
**    License along with this library; if not, write to the Free Software
**    Foundation, Inc., 59 Temple Place, Suite 330, Boston, MA  02111-1307  USA
*/

package org.systemsbiology.biofabric.cmd;

import java.awt.Dimension;
import java.awt.Point;
import java.awt.Rectangle;
import java.awt.geom.Point2D;
import java.awt.geom.Rectangle2D;

/****************************************************************************
**
** Used to abstract away zoom operations of SUPanel
*/

public interface ZoomTarget {

  ////////////////////////////////////////////////////////////////////////////
  //
  // PUBLIC CONSTANTS
  //
  ////////////////////////////////////////////////////////////////////////////  
  
  ////////////////////////////////////////////////////////////////////////////
  //
  // PUBLIC INSTANCE METHODS
  //
  ////////////////////////////////////////////////////////////////////////////
  
  /***************************************************************************
  **
  ** Methods
  */ 

  public Point pointToViewport(Point world); 
  public Point2D getRawCenterPoint();  
  public Rectangle getSelectedBounds();
  public Rectangle getCurrentSelectedBounds();
  public void incrementToNextSelection();
  public void decrementToPreviousSelection();
  public Rectangle getWorldRect();
  public Dimension getPreferredSize();
  public void repaint();
  public Dimension getBasicSize();
  public Point2D viewToWorld(Point vPt);
  public void setZoomFactor(double newZoomVal, Dimension vDim);
  public Rectangle getCurrentBasicBounds();
  public void adjustWideZoomForSize(Dimension dims);
  public Point getCenterPoint();
  public void setCurrClipRect(Rectangle2D clipRect);
  public double getZoomFactor();
}
```
