## Supplementary material for "BioFabric Visualization of Network Alignments": BioFabric Source Code: ColorGenerator.java

```
/*
**    Copyright (C) 2003-2011 Institute for Systems Biology 
**                            Seattle, Washington, USA. 
**
**    This library is free software; you can redistribute it and/or
**    modify it under the terms of the GNU Lesser General Public
**    License as published by the Free Software Foundation; either
**    version 2.1 of the License, or (at your option) any later version.
**
**    This library is distributed in the hope that it will be useful,
**    but WITHOUT ANY WARRANTY; without even the implied warranty of
**    MERCHANTABILITY or FITNESS FOR A PARTICULAR PURPOSE.  See the GNU
**    Lesser General Public License for more details.
**
**    You should have received a copy of the GNU Lesser General Public
**    License along with this library; if not, write to the Free Software
**    Foundation, Inc., 59 Temple Place, Suite 330, Boston, MA  02111-1307  USA
*/

package org.systemsbiology.biofabric.db;

import java.util.HashMap;
import java.util.HashSet;
import java.util.Map;
import java.util.Set;
import java.util.Iterator;
import java.util.List;
import java.util.ArrayList;
import java.awt.Color;
import java.io.PrintWriter;
import java.util.Arrays;

import org.systemsbiology.biofabric.api.io.Indenter;
import org.systemsbiology.biofabric.api.util.UniqueLabeller;
import org.systemsbiology.biofabric.ui.NamedColor;

/****************************************************************************
**
** Color generator
*/

public class ColorGenerator {
  
  ////////////////////////////////////////////////////////////////////////////
  //
  // PRIVATE MEMBERS
  //
  ////////////////////////////////////////////////////////////////////////////

  private UniqueLabeller colorLabels_;
  private Map<String, NamedColor> colors_;

  private String[] geneCol_ = new String[] {  
    "EX-blue",
    "EX-orange",
    "EX-dark-cyan",
    "EX-red",
    "EX-dark-orange",
    "EX-dark-gray-purple",
    "EX-cyan",
    "EX-yellow-orange",
    "EX-pure-blue",
    "EX-dark-yellow-green",
    "EX-dark-magenta",
    "EX-dark-green",
    "EX-blue-magenta",
    "EX-yellow-green",
    "EX-magenta",
    "EX-green",
    "EX-yellow",
    "EX-purple",
    "EX-dark-purple",
    "EX-dark-red",
    "EX-pale-green",
    "EX-pale-blue",
    "EX-dark-tan",
    "EX-pale-blue-magenta",
    "EX-pale-yellow orange",
    "EX-medium-magenta",
    "EX-pale-red",
    "EX-pale-cyan",
    "EX-pale-yellow-green",
    "EX-pale-purple",
    "EX-pale-magenta",
    "EX-pale-red-orange"
  };
  
  //
  // This is an alternative ordering that maximizes RGB separation:
  //
  
  /*
  private String[] geneCol_ = new String[] {    
    "EX-blue",
    "EX-red",
    "EX-green",
    "EX-magenta",
    "EX-cyan",
    "EX-yellow",
    "EX-pure-blue",
    "EX-yellow-green",
    "EX-blue-magenta",
    "EX-orange",
    "EX-dark-gray-purple",
    "EX-yellow-orange",
    "EX-dark-green",
    "EX-pale-magenta",
    "EX-pale-yellow-green",
    "EX-purple",
    "EX-dark-cyan",
    "EX-dark-orange",
    "EX-pale-cyan",
    "EX-dark-magenta",
    "EX-pale-purple",
    "EX-dark-yellow-green",
    "EX-pale-red-orange",
    "EX-pale-blue",
    "EX-dark-red",
    "EX-pale-yellow-orange",
    "EX-dark-purple",
    "EX-pale-green",
    "EX-pale-blue-magenta",
    "EX-pale-red",
    "EX-medium-magenta",
    "EX-dark-tan"
    };
  */
  
  ////////////////////////////////////////////////////////////////////////////
  //
  // PUBLIC CONSTRUCTORS
  //
  ////////////////////////////////////////////////////////////////////////////

  /***************************************************************************
  **
  ** Null constructor
  */

  public ColorGenerator() {
    colorLabels_ = new UniqueLabeller();
    colorLabels_.addExistingLabel("zz_newColor_0");
    // Can't do this: SpEndomes has custom tags e.g. "forest"
    //colorLabels_.setFixedPrefix("zz_newColor_0")
    colors_ = new HashMap<String, NamedColor>();
  }

  ////////////////////////////////////////////////////////////////////////////
  //
  // PUBLIC METHODS
  //
  ////////////////////////////////////////////////////////////////////////////
  
  /***************************************************************************
  ** 
  ** Start a new model
  */

  public void newColorModel() {
    buildDefaultColors();
    return;
  }  

  /***************************************************************************
  ** 
  ** Drop everything
  */

  public void dropColors() {
    colorLabels_ = new UniqueLabeller();
    colorLabels_.addExistingLabel("zz_newColor_0");
    // Can't do this: SpEndomes has custom tags e.g. "forest"
    //colorLabels_.setFixedPrefix("zz_newColor_0");    
    colors_.clear();
    buildDefaultColors();    
    return;
  }

  /***************************************************************************
  ** 
  ** Get the color
  */

  public Color getColor(String colorKey) {
    return (colors_.get(colorKey).color);
  }

  /***************************************************************************
  ** 
  ** Get the named color
  */

  public NamedColor getNamedColor(String colorKey) {
    return (colors_.get(colorKey));
  }  

  /***************************************************************************
  **
  ** Update the color set
  */
  
  public GlobalChange updateColors(Map<String, NamedColor> namedColors) {
    GlobalChange retval = new GlobalChange();
    retval.origColors = deepCopyColorMap(colors_);
    colors_ = deepCopyColorMap(namedColors);
    retval.newColors = deepCopyColorMap(namedColors);   
    return (retval);
  }
    
  /***************************************************************************
  **
  ** Dump the database to the given file using XML
  */
  
  public void writeXML(PrintWriter out, Indenter ind) {    
    writeColors(out, ind);
    return;
  }

  /***************************************************************************
  **
  ** Get the next color label
  */
  
  public String getNextColorLabel() {
    return (colorLabels_.getNextLabel());    
  }  
  
  /***************************************************************************
  **
  ** Set the color for the given name
  */
  
  public void setColor(String itemId, NamedColor color) {
    colorLabels_.addExistingLabel(itemId);    
    colors_.put(itemId, color);
    return;
  }
  
  /***************************************************************************
  **
  ** Return an iterator over all the color keys
  */
  
  public Iterator<String> getColorKeys() {
    return (colors_.keySet().iterator());
  }

  /***************************************************************************
  **
  ** Return gene colors as list
  */
  
  public List<String> getGeneColorsAsList() {    
    return (Arrays.asList(geneCol_));
  }
    
  /***************************************************************************
  **
  ** Undo a change
  */
  
  public void changeUndo(GlobalChange undo) {   
    if ((undo.origColors != null) || (undo.newColors != null)) {
      colorChangeUndo(undo);
    }
    return;
  }  
  
  /***************************************************************************
  **
  ** Redo a change
  */
  
  public void changeRedo(GlobalChange undo) {
    if ((undo.origColors != null) || (undo.newColors != null)) {
      colorChangeRedo(undo);
    }    
    return;
  }

  /***************************************************************************
  **
  ** Return the colors you cannot delete
  */
  
  public Set<String> cannotDeleteColors() {
    HashSet<String> retval = new HashSet<String>();
    retval.add("white");
    //retval.add("red");  // This is handled now by EX-red
    retval.add("black");
    
    retval.add("yellowGreen");
    retval.add("inactiveYellowGreen");  
    retval.add("lightBlue");
    retval.add("inactiveLightBlue");
    retval.add("lightOrange");
    retval.add("inactiveLightOrange");
    retval.add("lightGreen");
    retval.add("inactiveLightGreen"); 
    retval.add("lightPurple");
    retval.add("inactiveLightPurple");
    retval.add("lightGray");
    retval.add("inactiveLightGray");
 
    int size = geneCol_.length;
    for (int i = 0; i < size; i++) {
      retval.add(geneCol_[i]);
    }
    return (retval);
  }  

  /***************************************************************************
  **
  ** Return from a cycle of active colors
  */
  
  public String activeColorCycle(int i) {
    i = i % 5;
    switch (i) {
      case 0:
        return ("lightGreen");
      case 1:
        return ("yellowGreen");
      case 2:
        return ("lightOrange");
      case 3:
        return ("lightPurple");
      case 4:
        return ("lightGray");
    }
    throw new IllegalStateException();
  }
  
  /***************************************************************************
  **
  ** A distinct active color
  */
  
  public String distinctActiveColor() { 
    return ("lightBlue");
  }
  
  /***************************************************************************
  **
  ** Return a distinct inactive color
  */
  
  public String distinctInactiveColor() {  
    return ("inactiveLightBlue");
  }
   
  /***************************************************************************
  **
  ** Return from a cycle of inactive colors
  */
  
  public String inactiveColorCycle(int i) {
    i = i % 5;
    switch (i) {
      case 0:
        return ("inactiveLightGreen");
      case 1:
        return ("inactiveYellowGreen");
      case 2:
        return ("inactiveLightOrange");
      case 3:
        return ("inactiveLightPurple");
      case 4:
        return ("inactiveLightGray");
    }
    throw new IllegalStateException();
  }
   
  /***************************************************************************
  **
  ** Get Ith available color
  */
  
  public String getGeneColor(int i) {
    return (geneCol_[i]);
  }   
  
  /***************************************************************************
  **
  ** Get number of colors
  */
  
  public int getNumColors() {
    return (geneCol_.length);
  }
  
  /***************************************************************************
  **
  ** Build the default color set.
  */
  
  private void buildDefaultColors() {
    colors_ = new HashMap<String, NamedColor>();
    colors_.put("inactiveLightBlue", new NamedColor("inactiveLightBlue", new Color(235, 235, 250), "Very Light Blue"));
    colorLabels_.addExistingLegacyLabel("inactiveLightBlue");   
    colors_.put("white", new NamedColor("white", new Color(255, 255, 255), "White"));    
    colorLabels_.addExistingLegacyLabel("white");   
    colors_.put("inactiveLightPurple", new NamedColor("inactiveLightPurple", new Color(245, 229, 240), "Very Light Purple"));
    colorLabels_.addExistingLegacyLabel("inactiveLightPurple");   
    colors_.put("lightBlue", new NamedColor("lightBlue", new Color(220, 220, 240), "Light Blue"));
    colorLabels_.addExistingLegacyLabel("lightBlue");   
    colors_.put("black", new NamedColor("black", new Color(0, 0, 0), "Black"));
    colorLabels_.addExistingLegacyLabel("black");   
    colors_.put("inactiveLightOrange", new NamedColor("inactiveLightOrange", new Color(255, 230, 200), "Very Light Orange"));
    colorLabels_.addExistingLegacyLabel("inactiveLightOrange");   
    colors_.put("lightGray", new NamedColor("lightGray", new Color(240, 240, 240), "Light Gray"));
    colorLabels_.addExistingLegacyLabel("lightGray");   
    colors_.put("darkGray", new NamedColor("darkGray", new Color(150, 150, 150), "Dark Gray"));   
    colorLabels_.addExistingLegacyLabel("darkGray");    
    colors_.put("inactiveYellowGreen", new NamedColor("inactiveYellowGreen", new Color(255, 255, 220), "Light Yellow Green"));
    colorLabels_.addExistingLegacyLabel("inactiveYellowGreen");   
    colors_.put("yellowGreen", new NamedColor("yellowGreen", new Color(246, 249, 170), "Yellow Green"));
    colorLabels_.addExistingLegacyLabel("yellowGreen");   
    colors_.put("inactiveLightGreen", new NamedColor("inactiveLightGreen", new Color(230, 255, 220), "Very Light Green"));
    colorLabels_.addExistingLegacyLabel("inactiveLightGreen");   
    colors_.put("lightGreen", new NamedColor("lightGreen", new Color(214, 239, 209), "Light Green"));
    colorLabels_.addExistingLegacyLabel("lightGreen");   
    colors_.put("lightOrange", new NamedColor("lightOrange", new Color(244, 211, 170), "Light Orange"));
    colorLabels_.addExistingLegacyLabel("lightOrange");   
    colors_.put("inactiveLightGray", new NamedColor("inactiveLightGray", new Color(245, 245, 245), "Very Light Gray"));
    colorLabels_.addExistingLegacyLabel("inactiveLightGray");   
    colors_.put("lightPurple", new NamedColor("lightPurple", new Color(235, 219, 229), "Light Purple"));
    colorLabels_.addExistingLegacyLabel("lightPurple");
    
    List<NamedColor> geneColors = buildGeneColors();
    int geneColSize = geneColors.size();
    for (int i = 0; i < geneColSize; i++) {
      NamedColor col = geneColors.get(i);
      colors_.put(col.key, col);
      colorLabels_.addExistingLegacyLabel(col.key);
    }
    return;
  }
  
  ////////////////////////////////////////////////////////////////////////////
  //
  // PRIVATE INSTANCE METHODS
  //
  ////////////////////////////////////////////////////////////////////////////
    
  /***************************************************************************
  **
  ** Write the colors to XML
  **
  */
  
  private void writeColors(PrintWriter out, Indenter ind) {
    ind.indent();
    out.println("<colors>");
    ind.up();
    Iterator<String> colors = colors_.keySet().iterator();
    while (colors.hasNext()) {
      String key = colors.next();
      NamedColor nc = colors_.get(key);
      Color c = nc.color;
      ind.indent();
      out.print("<color ");
      out.print("color=\"");
      out.print(key);
      out.print("\" name=\"");
      out.print(nc.name);      
      out.print("\" r=\"");
      out.print(c.getRed());
      out.print("\" g=\"");
      out.print(c.getGreen());
      out.print("\" b=\"");
      out.print(c.getBlue());      
      out.println("\" />");
    }
    ind.down().indent();
    out.println("</colors>");
    return;
  }   
  

  
  /***************************************************************************
  ** 
  ** Build gene colors
  */

  private List<NamedColor> buildGeneColors() {

    ArrayList<NamedColor> colors = new ArrayList<NamedColor>();
    colors.add(new NamedColor("EX-red", Color.getHSBColor(0.0F, 1.0F, 1.0F), "Bright Red"));
    colors.add(new NamedColor("EX-pale-red-orange", Color.getHSBColor(0.033F, 0.4F, 0.9F), "Dark Salmon"));
    colors.add(new NamedColor("EX-orange", Color.getHSBColor(0.067F, 1.0F, 1.0F), "Pumpkin Orange"));            
    colors.add(new NamedColor("EX-yellow-orange", Color.getHSBColor(0.1F, 1.0F, 1.0F), "Tangerine"));
    colors.add(new NamedColor("EX-pale-yellow orange", Color.getHSBColor(0.12F, 0.5F, 0.8F), "Dark Wheat"));    
    colors.add(new NamedColor("EX-yellow", Color.getHSBColor(0.133F, 1.0F, 1.0F), "Gold"));
    colors.add(new NamedColor("EX-pale-yellow-green", Color.getHSBColor(0.183F, 0.4F, 0.9F), "Pale Goldenrod"));
    colors.add(new NamedColor("EX-yellow-green", Color.getHSBColor(0.233F, 1.0F, 1.0F), "Lime"));
    colors.add(new NamedColor("EX-pale-green", Color.getHSBColor(0.283F, 0.5F, 0.8F), "Pale Green"));                     
    colors.add(new NamedColor("EX-green", Color.getHSBColor(0.333F, 1.0F, 1.0F), "Bright Green"));
    colors.add(new NamedColor("EX-pale-cyan", Color.getHSBColor(0.413F, 0.4F, 0.9F), "Aquamarine"));                     
    colors.add(new NamedColor("EX-cyan", Color.getHSBColor(0.5F, 1.0F, 1.0F), "Cyan"));
    colors.add(new NamedColor("EX-pale-blue", Color.getHSBColor(0.534F, 0.5F, 0.8F), "Powder Blue"));                    
    colors.add(new NamedColor("EX-blue", Color.getHSBColor(0.567F, 1.0F, 1.0F), "Sky Blue"));
    colors.add(new NamedColor("EX-pale-purple", Color.getHSBColor(0.634F, 0.35F, 0.9F), "Cornflower Blue")); 
    colors.add(new NamedColor("EX-pure-blue", Color.getHSBColor(0.667F, 1.0F, 1.0F), "Blue"));
    colors.add(new NamedColor("EX-purple", Color.getHSBColor(0.708F, 0.8F, 1.0F), "Indigo"));
    colors.add(new NamedColor("EX-pale-blue-magenta", Color.getHSBColor(0.738F, 0.5F, 0.8F), "Lilac"));                    
    colors.add(new NamedColor("EX-blue-magenta", Color.getHSBColor(0.767F, 1.0F, 1.0F), "Bright Purple"));
    colors.add(new NamedColor("EX-pale-magenta", Color.getHSBColor(0.80F, 0.4F, 0.9F), "Light Plum"));                   
    colors.add(new NamedColor("EX-magenta", Color.getHSBColor(0.833F, 1.0F, 1.0F), "Fuchsia"));
    colors.add(new NamedColor("EX-pale-red", Color.getHSBColor(0.917F, 0.5F, 0.8F), "Rose"));                    
    colors.add(new NamedColor("EX-dark-red", Color.getHSBColor(0.0F, 0.6F, 0.55F), "Deep Ochre"));
    colors.add(new NamedColor("EX-dark-tan", Color.getHSBColor(0.1F, 0.5F, 0.65F), "Dark Tan"));
    colors.add(new NamedColor("EX-dark-orange", Color.getHSBColor(0.12F, 1.0F, 0.5F), "Sienna"));                   
    colors.add(new NamedColor("EX-dark-yellow-green", Color.getHSBColor(0.183F, 1.0F, 0.5F), "Olive Green"));
    colors.add(new NamedColor("EX-dark-green", Color.getHSBColor(0.283F, 1.0F, 0.5F), "Dark Green"));                    
    colors.add(new NamedColor("EX-dark-cyan", Color.getHSBColor(0.534F, 1.0F, 0.5F), "Dark Steel Blue"));                      
    colors.add(new NamedColor("EX-dark-gray-purple", Color.getHSBColor(0.634F, 1.0F, 0.5F), "Dark Blue"));        
    colors.add(new NamedColor("EX-dark-purple", Color.getHSBColor(0.708F, 0.6F, 0.55F), "Slate Blue"));
    colors.add(new NamedColor("EX-dark-magenta", Color.getHSBColor(0.80F, 1.0F, 0.5F), "Violet"));                    
    colors.add(new NamedColor("EX-medium-magenta", Color.getHSBColor(0.833F, 0.5F, 0.65F), "Mauve"));
    return (colors);
  }       
  
  /***************************************************************************
  **
  ** Write the note properties to XML
  */
  
  private HashMap<String, NamedColor> deepCopyColorMap(Map<String, NamedColor> otherMap) {
    HashMap<String, NamedColor> retval = new HashMap<String, NamedColor>();
    Set<String> keys = otherMap.keySet();
    Iterator<String> kit = keys.iterator();
    while (kit.hasNext()) {
      String key = kit.next();
      NamedColor col = otherMap.get(key);
      retval.put(new String(key), new NamedColor(col));
    }
    return (retval);
  }
  
  /***************************************************************************
  **
  ** Undo a color change
  */
  
  private void colorChangeUndo(GlobalChange undo) {
    if ((undo.origColors != null) && (undo.newColors != null)) {
      colors_ = undo.origColors;
    } else {
      throw new IllegalArgumentException();
    }
    return;
  }
    
  /***************************************************************************
  **
  ** Redo a color change
  */
  
  private void colorChangeRedo(GlobalChange undo) {
    if ((undo.origColors != null) && (undo.newColors != null)) {
      colors_ = undo.newColors;
    } else {
      throw new IllegalArgumentException();
    }   
    return;
  }
}
```
