## Supplementary material for "BioFabric Visualization of Network Alignments": BioFabric Source Code: EventManager.java

```
/*
**    Copyright (C) 2003-2018 Institute for Systems Biology 
**                            Seattle, Washington, USA. 
**
**    This library is free software; you can redistribute it and/or
**    modify it under the terms of the GNU Lesser General Public
**    License as published by the Free Software Foundation; either
**    version 2.1 of the License, or (at your option) any later version.
**
**    This library is distributed in the hope that it will be useful,
**    but WITHOUT ANY WARRANTY; without even the implied warranty of
**    MERCHANTABILITY or FITNESS FOR A PARTICULAR PURPOSE.  See the GNU
**    Lesser General Public License for more details.
**
**    You should have received a copy of the GNU Lesser General Public
**    License along with this library; if not, write to the Free Software
**    Foundation, Inc., 59 Temple Place, Suite 330, Boston, MA  02111-1307  USA
*/

package org.systemsbiology.biofabric.event;

import java.util.HashSet;
import java.util.Iterator;

/****************************************************************************
**
** Event Manager.  This is a Singleton.  Not currently thread-safe.
*/

public class EventManager {
  
  ////////////////////////////////////////////////////////////////////////////
  //
  // PRIVATE MEMBERS
  //
  ////////////////////////////////////////////////////////////////////////////

  private static EventManager singleton_;
  private HashSet layoutListeners_;
  private HashSet selectListeners_;
  
  ////////////////////////////////////////////////////////////////////////////
  //
  // PUBLIC METHODS
  //
  ////////////////////////////////////////////////////////////////////////////

  /***************************************************************************
  ** 
  ** Add an event listener
  */

  public void addLayoutChangeListener(LayoutChangeListener lcl) {
    layoutListeners_.add(lcl);
    return;
  }
  
  /***************************************************************************
  ** 
  ** Remove an event listener
  */

  public void removeLayoutChangeListener(LayoutChangeListener lcl) {
    layoutListeners_.remove(lcl);
    return;
  }  
 
  /***************************************************************************
  ** 
  ** Ask for an event to be sent.
  */

  public void sendLayoutChangeEvent(LayoutChangeEvent lcev) {
    Iterator lclit = layoutListeners_.iterator();
    while (lclit.hasNext()) {
      LayoutChangeListener lcl = (LayoutChangeListener)lclit.next();
      lcl.layoutHasChanged(lcev);
    }
    return;
  }  

  /***************************************************************************
  ** 
  ** Add an event listener
  */

  public void addSelectionChangeListener(SelectionChangeListener scl) {
    selectListeners_.add(scl);
    return;
  }
  
  /***************************************************************************
  ** 
  ** Remove an event listener
  */

  public void removeSelectionChangeListener(SelectionChangeListener scl) {
    selectListeners_.remove(scl);
    return;
  }  
 
  /***************************************************************************
  ** 
  ** Ask for an event to be sent.
  */

  public void sendSelectionChangeEvent(SelectionChangeEvent scev) {
    Iterator sclit = selectListeners_.iterator();
    while (sclit.hasNext()) {
      SelectionChangeListener scl = (SelectionChangeListener)sclit.next();
      scl.selectionHasChanged(scev);
    }
    return;
  }
  
  ////////////////////////////////////////////////////////////////////////////
  //
  // PUBLIC STATIC METHODS
  //
  ////////////////////////////////////////////////////////////////////////////

  /***************************************************************************
  ** 
  ** Get the singleton
  */

  public static synchronized EventManager getManager() {
    if (singleton_ == null) {
      singleton_ = new EventManager();
    }
    return (singleton_);
  }

  ////////////////////////////////////////////////////////////////////////////
  //
  // PRIVATE CONSTRUCTORS
  //
  ////////////////////////////////////////////////////////////////////////////

  /***************************************************************************
  **
  ** Null constructor
  */

  private EventManager() {
    layoutListeners_ = new HashSet();
    selectListeners_ = new HashSet();
  }
  
  ////////////////////////////////////////////////////////////////////////////
  //
  // PRIVATE INSTANCE METHODS
  //
  ////////////////////////////////////////////////////////////////////////////
}
```
