## Supplementary material for "BioFabric Visualization of Network Alignments": BioFabric Source Code: LayoutChangeEvent.java

```
/*
**    Copyright (C) 2003-2018 Institute for Systems Biology 
**                            Seattle, Washington, USA. 
**
**    This library is free software; you can redistribute it and/or
**    modify it under the terms of the GNU Lesser General Public
**    License as published by the Free Software Foundation; either
**    version 2.1 of the License, or (at your option) any later version.
**
**    This library is distributed in the hope that it will be useful,
**    but WITHOUT ANY WARRANTY; without even the implied warranty of
**    MERCHANTABILITY or FITNESS FOR A PARTICULAR PURPOSE.  See the GNU
**    Lesser General Public License for more details.
**
**    You should have received a copy of the GNU Lesser General Public
**    License along with this library; if not, write to the Free Software
**    Foundation, Inc., 59 Temple Place, Suite 330, Boston, MA  02111-1307  USA
*/


package org.systemsbiology.biofabric.event;

/****************************************************************************
**
** Used to signal changes in the current layout
*/

public class LayoutChangeEvent implements ChangeEvent {
  
  ////////////////////////////////////////////////////////////////////////////
  //
  // PUBLIC CONSTANTS
  //
  ////////////////////////////////////////////////////////////////////////////
  
  public final static int NO_CHANGE          = 0;
  public final static int UNSPECIFIED_CHANGE = 1;
  
  ////////////////////////////////////////////////////////////////////////////
  //
  // PRIVATE VARIABLES
  //
  ////////////////////////////////////////////////////////////////////////////  
  
  private String layoutKey_;
  private int changeType_;

  ////////////////////////////////////////////////////////////////////////////
  //
  // PUBLIC CONSTRUCTORS
  //
  ////////////////////////////////////////////////////////////////////////////    

  /***************************************************************************
  **
  ** Build the event
  */ 
  
  public LayoutChangeEvent(String layoutKey, int changeType) {
    layoutKey_ = layoutKey;
    changeType_ = changeType;
  }

  ////////////////////////////////////////////////////////////////////////////
  //
  // PUBLIC INSTANCE METHODS
  //
  ////////////////////////////////////////////////////////////////////////////
  
  /***************************************************************************
  **
  ** Get the layout key
  */ 
  
  public String getLayoutKey() {
    return (layoutKey_);
  }
  
  /***************************************************************************
  **
  ** Get the change type
  */ 
  
  public int getChangeType() {
    return (changeType_);
  } 

  /***************************************************************************
  **
  ** Standard equals
  */     
  
  public boolean equals(Object other) {
    if (this == other) {
      return (true);
    }
    if (other == null) {
      return (false);
    }
    if (!(other instanceof LayoutChangeEvent)) {
      return (false);
    }
    LayoutChangeEvent otherLCE = (LayoutChangeEvent)other;
    // Actually ran into a null layout key 06/11/08:
    if ((this.layoutKey_ == null) && (otherLCE.layoutKey_ != null)) {
      return (false);
    }
    if (!this.layoutKey_.equals(otherLCE.layoutKey_)) {
      return (false);
    }
    
    return (this.changeType_ == otherLCE.changeType_);
  }   
}
```
