## Supplementary material for "BioFabric Visualization of Network Alignments": BioFabric Source Code: AnnotationLoader.java

```
/*
**    Copyright (C) 2003-2018 Institute for Systems Biology 
**                            Seattle, Washington, USA. 
**
**    This library is free software; you can redistribute it and/or
**    modify it under the terms of the GNU Lesser General Public
**    License as published by the Free Software Foundation; either
**    version 2.1 of the License, or (at your option) any later version.
**
**    This library is distributed in the hope that it will be useful,
**    but WITHOUT ANY WARRANTY; without even the implied warranty of
**    MERCHANTABILITY or FITNESS FOR A PARTICULAR PURPOSE.  See the GNU
**    Lesser General Public License for more details.
**
**    You should have received a copy of the GNU Lesser General Public
**    License along with this library; if not, write to the Free Software
**    Foundation, Inc., 59 Temple Place, Suite 330, Boston, MA  02111-1307  USA
*/

package org.systemsbiology.biofabric.io;

import java.io.BufferedReader;
import java.io.File;
import java.io.FileInputStream;
import java.io.IOException;
import java.io.InputStreamReader;
import java.util.ArrayList;
import java.util.HashMap;
import java.util.List;
import java.util.Map;
import java.util.Set;
import java.util.regex.Matcher;
import java.util.regex.Pattern;

import org.systemsbiology.biofabric.api.model.Annot;
import org.systemsbiology.biofabric.api.model.AnnotationSet;
import org.systemsbiology.biofabric.api.model.NetNode;
import org.systemsbiology.biofabric.api.worker.AsynchExitRequestException;
import org.systemsbiology.biofabric.api.worker.BTProgressMonitor;
import org.systemsbiology.biofabric.api.worker.LoopReporter;
import org.systemsbiology.biofabric.model.BioFabricNetwork;
import org.systemsbiology.biofabric.model.FabricLink;
import org.systemsbiology.biofabric.plugin.PluginSupportFactory;
import org.systemsbiology.biofabric.util.DataUtil;
import org.systemsbiology.biofabric.util.UiUtil;

/****************************************************************************
**
** This loads Annotation files
*/

public class AnnotationLoader {
  
  ////////////////////////////////////////////////////////////////////////////
  //
  // PRIVATE CONSTANTS
  //
  //////////////////////////////////////////////////////////////////////////// 
  
  ////////////////////////////////////////////////////////////////////////////
  //
  // PUBLIC CONSTANTS
  //
  //////////////////////////////////////////////////////////////////////////// 
  
  ////////////////////////////////////////////////////////////////////////////
  //
  // PRIVATE INSTANCE MEMBERS
  //
  ////////////////////////////////////////////////////////////////////////////
   
  ////////////////////////////////////////////////////////////////////////////
  //
  // PUBLIC CONSTRUCTORS
  //
  ////////////////////////////////////////////////////////////////////////////

  /***************************************************************************
  **
  ** Constructor
  */

  public AnnotationLoader() { 
  }

  ////////////////////////////////////////////////////////////////////////////
  //
  // PUBLIC METHODS
  //
  ////////////////////////////////////////////////////////////////////////////
   
  /***************************************************************************
  **
  ** Process an Annotation file
  */
  
  public Map<Boolean, AnnotationSet> readAnnotations(File infile, ReadStats stats, boolean forNodes,
  		                                               BioFabricNetwork bfn, BTProgressMonitor monitor) throws AsynchExitRequestException, IOException {
    
    long fileLen = infile.length();    
    Map<String, Set<NetNode>> nameToIDs = bfn.getNormNameToIDs();
    BufferedReader in = null;
    ArrayList<String[]> tokSets = new ArrayList<String[]>();
    LoopReporter lr = new LoopReporter(fileLen, 20, monitor, 0.0, 1.0, "progress.readingFile");
    try {
      in = new BufferedReader(new InputStreamReader(new FileInputStream(infile), "UTF-8"));
      String line = null;
      while ((line = in.readLine()) != null) {
        lr.report(line.length() + 1);
        if (line.trim().equals("")) {
          continue;
        }
        String[] tokens = lineToToks(line, stats, forNodes);
        if (tokens != null) {
          tokSets.add(tokens);
        }
      }
    } finally {
      if (in != null) {
        in.close();
      }
    }
    lr.finish();
    
    int numLines = tokSets.size();
    lr = new LoopReporter(numLines, 20, monitor, 0.0, 1.0, "progress.buildingAnnotations");
    
    Map<Boolean, AnnotationSet> retval = new HashMap<Boolean, AnnotationSet>();
     
    if (forNodes) {
      //
	    // Need to build a map from node name to row so we can build the Annotations. Invert
	    // what we have:
	    //
      AnnotationSet aSet = PluginSupportFactory.buildAnnotationSet();
	    HashMap<NetNode, Integer> idToRow = new HashMap<NetNode, Integer>();
	    int numNodes = bfn.getRowCount();
	    for (int i = 0; i < numNodes; i++) {
	      NetNode nid = bfn.getNodeIDForRow(Integer.valueOf(i));
	      idToRow.put(nid, Integer.valueOf(i));
	    }
	    for (int i = 0; i < numLines; i++) {
	      String[] tokens = tokSets.get(i);
	      lr.report();
	      consumeTokens(tokens, idToRow, nameToIDs, aSet, stats);
	    }
	    retval.put(Boolean.FALSE, aSet);
	    retval.put(Boolean.TRUE, aSet);   
    } else {  
    	
      //
      // Need to build a map from FabricLink to column so we can build the Annotations. Invert
      // what we have:
      //
      
    	Map<FabricLink, Integer> linkToColWithShadow = buildLinkToCol(bfn, true);
    	Map<FabricLink, Integer> linkToColNoShadow = buildLinkToCol(bfn, false);
    	Map<Boolean, Map<FabricLink, Integer>> linksToCols = new HashMap<Boolean, Map<FabricLink, Integer>>();
    	linksToCols.put(Boolean.TRUE, linkToColWithShadow);
    	linksToCols.put(Boolean.FALSE, linkToColNoShadow);
    	 
      AnnotationSet aSetShad = PluginSupportFactory.buildAnnotationSet();
      AnnotationSet aSet = PluginSupportFactory.buildAnnotationSet();
      retval.put(Boolean.TRUE, aSetShad);
      retval.put(Boolean.FALSE, aSet);
    	
      Pattern linkPat = Pattern.compile("(.*\\S) (.*)\\((.*)\\) (\\S.*)"); 
      Matcher mainMatch = linkPat.matcher("");
    	for (int i = 0; i < numLines; i++) {
        String[] tokens = tokSets.get(i);
        lr.report();
        consumeTokensForLink(tokens, linksToCols, nameToIDs, retval, stats, mainMatch);
      }  	
    }
    lr.finish();   
    return (retval);
  }

  /***************************************************************************
  ** 
  ** Build link to col map
  */
  
  private Map<FabricLink, Integer> buildLinkToCol(BioFabricNetwork bfn, boolean forShadows) {
    Map<FabricLink, Integer> linkToCol = new HashMap<FabricLink, Integer>();
    List<BioFabricNetwork.LinkInfo> lis = bfn.getLinkDefList(forShadows);
    for (BioFabricNetwork.LinkInfo li : lis) {
      linkToCol.put(li.getLink(), li.getUseColumn(forShadows));
    }
    return (linkToCol);
  }
   
  /***************************************************************************
  ** 
  ** Parse a line to tokens
  */

  protected String[] lineToToks(String line, ReadStats stats, boolean forNodes) throws IOException {
    int expectedMin = (forNodes) ? 4 : 5;
    int expectedMax = (forNodes) ? 5 : 6;
  	if (line.trim().equals("")) {
  		return (null);
  	}
    String[] tokens = line.split("\\t");
    if ((tokens.length == 1) && (line.indexOf("\\t") == -1)) {
      tokens = line.split(" ");
    }
    
    if (tokens.length == 0) {
      return (null);
    } else if ((tokens.length < expectedMin) || (tokens.length > expectedMax)) {
      stats.badLine = line;
      stats.errStr = "annotLoad.incorrectTokenCount";
      throw new IOException();
    } else {        
      return (tokens);
    }
  }
  
  /***************************************************************************
  ** 
  ** Consume tokens, make Annotations
  */

  protected void consumeTokensForLink(String[] tokens, Map<Boolean, Map<FabricLink, Integer>> linksToCols, 
                                      Map<String, Set<NetNode>> nameToID, 
                                      Map<Boolean, AnnotationSet> aSets,
                                      ReadStats stats, Matcher mainMatch) throws IOException {

    boolean isShadow = tokens[4].trim().equalsIgnoreCase("isShadow");
    Map<FabricLink, Integer> linkToCol = linksToCols.get(Boolean.valueOf(isShadow));
 
    int layer;
    try {
      layer = Integer.valueOf(tokens[3]).intValue();
    } catch (NumberFormatException nfex) {
      stats.badTok = tokens[3];
      stats.errStr = "annotLoad.badLayer";
      throw new IOException();
    }
    
    if (layer < 0) {
      stats.badTok = tokens[3];
      stats.errStr = "annotLoad.negLayer";
      throw new IOException();
    }
    
    String name = tokens[0].trim();
    
    String startLink = tokens[1].trim();
    startLink = stripQuotes(startLink);
    FabricLink minLink = parseLink(startLink, nameToID, stats, mainMatch);
    
    String endLink = tokens[2].trim();
    endLink = stripQuotes(endLink);
    FabricLink maxLink = parseLink(endLink, nameToID, stats, mainMatch);
    
    Integer minCol = linkToCol.get(minLink);
    if (minCol == null) {
      stats.badTok = startLink;
      stats.errStr = "annotLoad.linkNotFound";
      throw new IOException();   
    }
   
    Integer maxCol = linkToCol.get(maxLink);
    if (maxCol == null) {
      stats.badTok = endLink;
      stats.errStr = "annotLoad.linkNotFound";
      throw new IOException();   
    }
    
    int min = minCol.intValue();
    int max = maxCol.intValue();
    
    String colorName = (tokens.length == 5) ? null : tokens[5];
    
    try {
      Annot annot = PluginSupportFactory.buildAnnotation(name, min, max, layer, colorName);
      aSets.get(Boolean.valueOf(isShadow)).addAnnot(annot);
    } catch (IllegalArgumentException iaex) {
      stats.badTok = tokens[1] + " " + tokens[2] + " " + tokens[3];
      if (tokens.length == 6) {
        stats.badTok += " " + tokens[5];
      }
      stats.errStr = "annotLoad.badAnnotDefinition";
      throw new IOException();
    }
    return;
  }
  
  /***************************************************************************
  ** 
  ** Consume token for a link, make a link
  */

  private FabricLink parseLink(String token, Map<String, Set<NetNode>> nameToID,
                               ReadStats stats, Matcher mainMatch) throws IOException {
    mainMatch.reset(token);
    if (!mainMatch.matches()) {
      stats.badTok = token;
      stats.errStr = "annotLoad.badLink";
      throw new IOException();
    }
 
    String src = mainMatch.group(1).trim();
    String sha = mainMatch.group(2).trim();
    String rel = mainMatch.group(3).trim();
    String trg = mainMatch.group(4).trim();
    boolean isShadow = sha.equals("shdw");
    Set<NetNode> srcIDs = nameToID.get(DataUtil.normKey(src));
    if (srcIDs == null) {
      stats.badTok = src;
      stats.errStr = "annotLoad.nodeNotFound";
      throw new IOException();   
    } else if (srcIDs.size() != 1) {
      stats.badTok = src;
      stats.errStr = "annotLoad.multiNodesForName";
      throw new IOException();   
    }
    NetNode srcID = srcIDs.iterator().next();
    Set<NetNode> trgIDs = nameToID.get(DataUtil.normKey(trg));
    if (trgIDs == null) {
      stats.badTok = trg;
      stats.errStr = "annotLoad.nodeNotFound";
      throw new IOException();   
    } else if (trgIDs.size() != 1) {
      stats.badTok = trg;
      stats.errStr = "annotLoad.multiNodesForName";
      throw new IOException();   
    }
    NetNode trgID = trgIDs.iterator().next();
    UiUtil.fixMePrintout("NO! Cannot assume undirected");
    FabricLink nextLink = new FabricLink(srcID, trgID, rel, isShadow, false);
    return (nextLink);
  }

 
  /***************************************************************************
  **
  ** Strip quoted string
  */
  
  protected String stripQuotes(String inString) {
    String procString = inString.trim();
    if ((procString.indexOf("\"") == 0) && (procString.lastIndexOf("\"") == (procString.length() - 1))) {
      procString = procString.replaceAll("\"", "");
    }
    return (procString);
  }
  
  /***************************************************************************
  ** 
  ** Consume tokens, make Annotations
  */

  protected void consumeTokens(String[] tokens, Map<NetNode, Integer> idToRow, 
                               Map<String, Set<NetNode>> nameToID, AnnotationSet aSet,
                               ReadStats stats) throws IOException {
    
    
    int layer;
    try {
      layer = Integer.valueOf(tokens[3]).intValue();
    } catch (NumberFormatException nfex) {
      stats.badTok = tokens[3];
      stats.errStr = "annotLoad.badLayer";
      throw new IOException();
    }
    
    if (layer < 0) {
      stats.badTok = tokens[3];
      stats.errStr = "annotLoad.negLayer";
      throw new IOException();
    }
    
    String name = tokens[0].trim();
    
    String startNode = tokens[1].trim();
    startNode = stripQuotes(startNode);
    
    String endNode = tokens[2].trim();
    endNode = stripQuotes(endNode);
    
    int min = nameToRow(startNode, idToRow, nameToID, stats);  
    int max = nameToRow(endNode, idToRow, nameToID, stats);
   
    String colorName = (tokens.length == 4) ? null : tokens[4];
     
    try {
      Annot annot = PluginSupportFactory.buildAnnotation(name, min, max, layer, colorName);
      aSet.addAnnot(annot);
    } catch (IllegalArgumentException iaex) {
      stats.badTok = tokens[1] + " " + tokens[2] + " " + tokens[3];
      if (tokens.length == 5) {
        stats.badTok += " " + tokens[4];
      }
      stats.errStr = "annotLoad.badAnnotDefinition";
      throw new IOException();
    }
    return;
  }

  /***************************************************************************
  ** 
  ** Node ID to row
  */

  private int nameToRow(String nodeID, Map<NetNode, Integer> idToRow, 
                        Map<String, Set<NetNode>> nameToID, 
                        ReadStats stats) throws IOException {
  
    Set<NetNode> forID = nameToID.get(DataUtil.normKey(nodeID));
    if (forID == null) {
      stats.badTok = nodeID;
      stats.errStr = "annotLoad.nodeNotFound";
      throw new IOException();   
    } else if (forID.size() != 1) {
      stats.badTok = nodeID;
      stats.errStr = "annotLoad.multiNodesForName";
      throw new IOException();   
    }
    NetNode nidwn = forID.iterator().next();
    Integer rowNum = idToRow.get(nidwn);
    if (rowNum == null) {
      throw new IllegalStateException();
    }
    
    return (rowNum.intValue());
  } 
  
  
  ////////////////////////////////////////////////////////////////////////////
  //
  // PUBLIC INNER CLASSES
  //
  ////////////////////////////////////////////////////////////////////////////
   
  /***************************************************************************
  **
  ** Problems with imports
  */  
 
  public static class ReadStats {
    public String badLine;
    public String badTok;
    public String errStr;
    
    public ReadStats() {
    }
  }  
}
```
