## Supplementary material for "BioFabric Visualization of Network Alignments": BioFabric Source Code: AttributeLoader.java

```
/*
**    Copyright (C) 2003-2018 Institute for Systems Biology 
**                            Seattle, Washington, USA. 
**
**    This library is free software; you can redistribute it and/or
**    modify it under the terms of the GNU Lesser General Public
**    License as published by the Free Software Foundation; either
**    version 2.1 of the License, or (at your option) any later version.
**
**    This library is distributed in the hope that it will be useful,
**    but WITHOUT ANY WARRANTY; without even the implied warranty of
**    MERCHANTABILITY or FITNESS FOR A PARTICULAR PURPOSE.  See the GNU
**    Lesser General Public License for more details.
**
**    You should have received a copy of the GNU Lesser General Public
**    License along with this library; if not, write to the Free Software
**    Foundation, Inc., 59 Temple Place, Suite 330, Boston, MA  02111-1307  USA
*/

package org.systemsbiology.biofabric.io;

import java.io.FileInputStream;
import java.io.BufferedReader;
import java.io.File;
import java.io.InputStreamReader;
import java.util.ArrayList;
import java.io.IOException;
import java.util.Map;
import java.util.regex.Matcher;
import java.util.regex.Pattern;

import org.systemsbiology.biofabric.api.io.AttributeKey;
import org.systemsbiology.biofabric.api.model.NetNode;
import org.systemsbiology.biofabric.model.FabricLink;
import org.systemsbiology.biofabric.util.DataUtil;

/****************************************************************************
**
** This loads attribute files
*/

public class AttributeLoader {
  
  ////////////////////////////////////////////////////////////////////////////
  //
  // PRIVATE CONSTANTS
  //
  //////////////////////////////////////////////////////////////////////////// 
  
  ////////////////////////////////////////////////////////////////////////////
  //
  // PUBLIC CONSTANTS
  //
  //////////////////////////////////////////////////////////////////////////// 
  
  ////////////////////////////////////////////////////////////////////////////
  //
  // PRIVATE INSTANCE MEMBERS
  //
  ////////////////////////////////////////////////////////////////////////////
   
  ////////////////////////////////////////////////////////////////////////////
  //
  // PUBLIC CONSTRUCTORS
  //
  ////////////////////////////////////////////////////////////////////////////

  /***************************************************************************
  **
  ** Constructor
  */

  public AttributeLoader() {
 
  }

  ////////////////////////////////////////////////////////////////////////////
  //
  // PUBLIC METHODS
  //
  ////////////////////////////////////////////////////////////////////////////

    
  /***************************************************************************
  ** 
  ** Process an attribute input
  */

  public String readAttributes(File infile, boolean forNodes, Map<AttributeKey, String> results, 
  		                         Map<String, NetNode> nodes, ReadStats stats) throws IOException {
    
    Pattern nodePat = Pattern.compile("(.*)=(.*)");
    Pattern linkPat = Pattern.compile("(.*\\S) (.*)\\((.*)\\) (\\S.*)=(.*)"); 
    Matcher mainMatch = (forNodes) ? nodePat.matcher("") : linkPat.matcher("");

    String retval = null;
    BufferedReader in = new BufferedReader(new InputStreamReader(new FileInputStream(infile), "UTF-8"));
    String line = null;
    boolean isFirst = true;
    while ((line = in.readLine()) != null) {
      if (isFirst) {
        isFirst = false;
        retval = line.trim();
        continue;
      }
      if (line.trim().equals("")) {
      	continue;
      }
      mainMatch.reset(line);
      if (!mainMatch.matches()) {
        stats.badLines.add(line);
        continue;
      }
      if (forNodes) {
        String node = mainMatch.group(1).trim().toUpperCase();
        if ((node.indexOf("\"") == 0) && (node.lastIndexOf("\"") == (node.length() - 1))) {
          node = node.replaceAll("\"", "");
        }
        if (results.containsKey(node)) {          
          stats.dupLines.add(line);
          continue;
        }
        results.put(new StringKey(node), mainMatch.group(2).trim());
      } else {
        String src = mainMatch.group(1).trim();
        String sha = mainMatch.group(2).trim();
        String rel = mainMatch.group(3).trim();
        String trg = mainMatch.group(4).trim();
        boolean isShadow = sha.equals("shdw");
        if (isShadow) {
          stats.shadowsPresent = true;
        }
        NetNode srcID = nodes.get(DataUtil.normKey(src));
        NetNode trgID = nodes.get(DataUtil.normKey(trg));
        FabricLink nextLink = new FabricLink(srcID, trgID, rel, isShadow);
        if (results.containsKey(nextLink)) {
          stats.dupLines.add(line);
          continue;
        }
        results.put(nextLink, mainMatch.group(5).trim());
      }
    }
    in.close();
    return (retval);
  }
     
  ////////////////////////////////////////////////////////////////////////////
  //
  // PUBLIC INNER CLASES
  //
  ////////////////////////////////////////////////////////////////////////////

  public static class StringKey implements AttributeKey {
    public String key;
    
    public StringKey(String key) {
      this.key = key;
    } 
    
    @Override
    public String toString() {
      return (key);
    }
    
    @Override
    public int hashCode() {
      return (key.toUpperCase().hashCode());
    }

    @Override
    public boolean equals(Object other) {
      if (other == null) {
        return (false);
      }
      if (other == this) {
        return (true);
      }
      if (!(other instanceof StringKey)) {
        return (false);
      }
      StringKey otherT = (StringKey)other;
      return (DataUtil.normKey(this.key).equals(DataUtil.normKey(otherT.key)));
    }
  }
   
  public static class ReadStats {
    public ArrayList<String> dupLines;
    public ArrayList<String> badLines;
    public boolean shadowsPresent;
    
    public ReadStats() {
      badLines = new ArrayList<String>();
      dupLines = new ArrayList<String>();
    }
  }    
}
```
