## Supplementary material for "BioFabric Visualization of Network Alignments": BioFabric Source Code: BuildDataImpl.java

```
/*
**    Copyright (C) 2003-2018 Institute for Systems Biology 
**                            Seattle, Washington, USA. 
**
**    This library is free software; you can redistribute it and/or
**    modify it under the terms of the GNU Lesser General Public
**    License as published by the Free Software Foundation; either
**    version 2.1 of the License, or (at your option) any later version.
**
**    This library is distributed in the hope that it will be useful,
**    but WITHOUT ANY WARRANTY; without even the implied warranty of
**    MERCHANTABILITY or FITNESS FOR A PARTICULAR PURPOSE.  See the GNU
**    Lesser General Public License for more details.
**
**    You should have received a copy of the GNU Lesser General Public
**    License along with this library; if not, write to the Free Software
**    Foundation, Inc., 59 Temple Place, Suite 330, Boston, MA  02111-1307  USA
*/

package org.systemsbiology.biofabric.io;

import java.util.ArrayList;
import java.util.HashMap;
import java.util.HashSet;
import java.util.Iterator;
import java.util.List;
import java.util.Map;
import java.util.Set;
import java.util.SortedMap;

import org.systemsbiology.biofabric.analysis.Link;
import org.systemsbiology.biofabric.api.io.AttributeKey;
import org.systemsbiology.biofabric.api.io.BuildData;
import org.systemsbiology.biofabric.api.layout.DefaultEdgeLayout;
import org.systemsbiology.biofabric.api.layout.DefaultLayout;
import org.systemsbiology.biofabric.api.layout.EdgeLayout;
import org.systemsbiology.biofabric.api.layout.NodeLayout;
import org.systemsbiology.biofabric.api.model.AnnotationSet;
import org.systemsbiology.biofabric.api.model.NetLink;
import org.systemsbiology.biofabric.api.model.NetNode;
import org.systemsbiology.biofabric.api.model.Network;
import org.systemsbiology.biofabric.api.util.UniqueLabeller;
import org.systemsbiology.biofabric.api.worker.AsynchExitRequestException;
import org.systemsbiology.biofabric.api.worker.BTProgressMonitor;
import org.systemsbiology.biofabric.layouts.ControlTopLayout;
import org.systemsbiology.biofabric.layouts.HierDAGLayout;
import org.systemsbiology.biofabric.layouts.NodeClusterLayout;
import org.systemsbiology.biofabric.layouts.NodeSimilarityLayout;
import org.systemsbiology.biofabric.layouts.SetLayout;
import org.systemsbiology.biofabric.layouts.WorldBankLayout;

import org.systemsbiology.biofabric.model.BioFabricNetwork;
import org.systemsbiology.biofabric.plugin.PluginBuildData;
import org.systemsbiology.biofabric.ui.FabricColorGenerator;
import org.systemsbiology.biofabric.util.DataUtil;
import org.systemsbiology.biofabric.util.UiUtil;

/****************************************************************************
**
** This is data that is used to build and rebuild the network model
*/

public class BuildDataImpl implements BuildData {
  
  ////////////////////////////////////////////////////////////////////////////
  //
  // PRIVATE CONSTANTS
  //
  //////////////////////////////////////////////////////////////////////////// 

  ////////////////////////////////////////////////////////////////////////////
  //
  // PUBLIC CONSTANTS
  //
  //////////////////////////////////////////////////////////////////////////// 
  
  public enum BuildMode {DEFAULT_LAYOUT,
                         REORDER_LAYOUT ,
                         CLUSTERED_LAYOUT ,
                         SHADOW_LINK_CHANGE,
                         GROUP_PER_NODE_CHANGE,
                         BUILD_FOR_SUBMODEL,
                         BUILD_FROM_XML,
                         BUILD_FROM_SIF,
                         NODE_ATTRIB_LAYOUT,
                         LINK_ATTRIB_LAYOUT,
                         NODE_CLUSTER_LAYOUT,
                         CONTROL_TOP_LAYOUT,
                         HIER_DAG_LAYOUT,
                         WORLD_BANK_LAYOUT,
                         SET_LAYOUT,
                         GROUP_PER_NETWORK_CHANGE,
                         BUILD_FROM_PLUGIN
                        };
                                                
  ////////////////////////////////////////////////////////////////////////////
  //
  // PRIVATE INSTANCE VARIABLES
  //
  ////////////////////////////////////////////////////////////////////////////
                         
  private BuildMode mode_;
  
  //
  // For selection builds:
  //
  public BioFabricNetwork fullNet;
  public List<BioFabricNetwork.NodeInfo> subNodes;
  public List<BioFabricNetwork.LinkInfo> subLinks;
  
  //
  // For pre-built build data:
  //
  
  private BioFabricNetwork bfn_;
  private List<NetNode> existingIDOrder_;
  
  //
  // Nodes and edges:
  //
  
  private Set<NetLink> allLinks_;
  private Set<NetNode> loneNodeIDs_;
  private Set<NetNode> allNodeIDs_;
  
  //
  // Specifies layout:
  //
  
  private SortedMap<Integer, NetLink> linkOrder_;
  private Map<NetNode, Integer> nodeOrder_;

  //
  // Link Groups:
  //
    
  private Network.LayoutMode layoutMode_;
  private List<String> linkGroups_;
  private boolean showLinkGroupAnnotations_;
  
  //
  // If layout sets annotations, this is what gets set:
  
  private AnnotationSet nodeAnnotForLayout_;
  private Map<Boolean, AnnotationSet> linkAnnotsForLayout_;
   
  //
  // Keep passing this around and sharing:
  //
  
  private FabricColorGenerator colGen_;

  public Map<NetNode, String> clustAssign;

  public UniqueLabeller idGen;
  
  public ControlTopLayout.CtrlMode cMode; 
  public ControlTopLayout.TargMode tMode;
  public List<String> fixedOrder; 
  public Map<String, Set<NetNode>> normNameToIDs;
  public Boolean pointUp;
  public SetLayout.LinkMeans linkMeaning;
  private boolean turnOnShadows_;
  
  private PluginBuildData plugInData_;
  
  ////////////////////////////////////////////////////////////////////////////
  //
  // PUBLIC CONSTRUCTORS
  //
  ////////////////////////////////////////////////////////////////////////////   
        
  public BuildDataImpl(BuildMode mode) {
    this.mode_ = mode;
    this.turnOnShadows_ = false;
  }
  
  public BuildDataImpl(BioFabricNetwork fullNet, List<BioFabricNetwork.NodeInfo> subNodes, List<BioFabricNetwork.LinkInfo> subLinks) {
    this(BuildMode.BUILD_FOR_SUBMODEL);
    this.fullNet = fullNet;
    this.subNodes = subNodes;
    this.subLinks = subLinks;
  }

  public BuildDataImpl(BioFabricNetwork bfn, BuildMode mode) {
    this(mode);
    this.bfn_ = bfn;
  }
  
  public BuildDataImpl(BioFabricNetwork fullNet, BuildMode mode, BTProgressMonitor monitor) throws AsynchExitRequestException {
    this(mode);
    this.bfn_ = fullNet;
    this.allLinks_ = fullNet.getAllLinks(true);
    this.colGen_ = fullNet.getColorGenerator();
    this.nodeOrder_ = null;
    this.existingIDOrder_ = fullNet.existingIDOrder();
    this.linkOrder_ = null;
    this.linkGroups_ = fullNet.getLinkGrouping();
    this.loneNodeIDs_ = fullNet.getLoneNodes(monitor);
    this.allNodeIDs_ = fullNet.getAllNodeDefinitions().keySet();
    this.clustAssign = (fullNet.nodeClustersAssigned()) ? fullNet.nodeClusterAssigment() : null;
    this.layoutMode_ = fullNet.getLayoutMode();
    this.idGen = fullNet.getGenerator();
    this.nodeAnnotForLayout_ = null;
    this.linkAnnotsForLayout_ = null;
  }
    
  public BuildDataImpl(UniqueLabeller idGen,
  		                 Set<NetLink> allLinks, Set<NetNode> loneNodeIDs, 
  		                 Map<NetNode, String> clustAssign, 
  		                 FabricColorGenerator colGen, BuildMode mode) {
    this(mode);
    this.bfn_ = null;
    this.allLinks_ = allLinks;
    this.colGen_ = colGen;
    this.nodeOrder_ = null;
    this.existingIDOrder_ = null;
    this.linkOrder_ = null;
    this.linkGroups_ = new ArrayList<String>();
    this.clustAssign = clustAssign;
    this.loneNodeIDs_ = loneNodeIDs;
    this.allNodeIDs_ = null;
    this.layoutMode_ = BioFabricNetwork.LayoutMode.PER_NODE_MODE;
    this.idGen = idGen;
    this.nodeAnnotForLayout_ = null;
    this.linkAnnotsForLayout_ = null;
  } 

  public PluginBuildData getPluginBuildData() {
    return (plugInData_);
  }  
  
  public void setPluginBuildData(PluginBuildData plugInData) {
    plugInData_ = plugInData;
    return;
  }  
  
  public Set<NetNode> getSingletonNodes() {
    return (loneNodeIDs_);
  } 
  
  public Set<NetNode> getAllNodes() {
    return (allNodeIDs_);
  } 
  
  public Set<NetLink> getLinks() {
    return (allLinks_);
  } 
 
  // Bogus hack to keep cluster layout working for now
  public void initAllNodesBogus(Set<NetNode> startSet) {
    allNodeIDs_ =  startSet;
    return;
  }
  
  // Bogus hack to keep cluster layout working for now
  public void addToAllNodesBogus(NetNode node) {
    allNodeIDs_.add(node);
    return;
  }
  
  public BuildMode getMode() {
    return (mode_);
  }  
  
  public boolean canRestore() {
    return (true);
  } 
  
  //
  // Allow for late binding of color generator:
  //
  
  public void setColorGen(FabricColorGenerator colGen) {
    this.colGen_ = colGen;
    return;
  }
   
  public FabricColorGenerator getColorGen() {
    return (colGen_);
  }
 
  public Map<String, Set<NetNode>> genNormNameToID() {
  	HashMap<String, Set<NetNode>> retval = new HashMap<String, Set<NetNode>>();
  	Iterator<NetNode> nit = this.allNodeIDs_.iterator();
  	while (nit.hasNext()) {
  		NetNode nodeID = nit.next();
  		String name = nodeID.getName();
		  String nameNorm = DataUtil.normKey(name);
	   	Set<NetNode> forName = retval.get(nameNorm);
		  if (forName == null) {
			  forName = new HashSet<NetNode>();
			  retval.put(nameNorm, forName);
		  }
		  forName.add(nodeID);
	  }
    return (retval);	
  }
  
  public void setNodeOrderFromAttrib(Map<AttributeKey, String> nodeOrderIn) {
    this.nodeOrder_ = new HashMap<NetNode, Integer>();
    Map<String, Set<NetNode>> nameToID = genNormNameToID();
    for (AttributeKey key : nodeOrderIn.keySet()) {
      try {
        Integer newRow = Integer.valueOf(nodeOrderIn.get(key));
        String keyName = ((AttributeLoader.StringKey)key).key;
        String normName = DataUtil.normKey(keyName);        
        Set<NetNode> ids = nameToID.get(normName);
        if (ids.size() != 1) {
        	throw new IllegalStateException();
        }
        NetNode id = ids.iterator().next();
        this.nodeOrder_.put(id, newRow);
      } catch (NumberFormatException nfex) {
        throw new IllegalStateException();
      }
    }
    return;
  }
  
  public BioFabricNetwork getExistingNetwork() {
    return (bfn_);
  }  
  
  public NetNode getNodeIDForRowForNetwork(int intval) {
    return (bfn_.getNodeIDForRow(intval));
  }
  
  public int getRowCountForNetwork() {
    return (bfn_.getRowCount());
  } 
  
  public List<NetNode> getExistingIDOrder() {
    return (existingIDOrder_);
  }  

  public Map<NetNode, Integer> getNodeOrder() {
    return (this.nodeOrder_);
  }

  public SortedMap<Integer, NetLink> getLinkOrder() {
    return (linkOrder_);
  }

  public void setNodeOrder(Map<NetNode, Integer> nodeOrder) {
    nodeOrder_ = nodeOrder;
    return;
  }

  public void setLinkOrder(SortedMap<Integer, NetLink> linkOrder) {
    linkOrder_ = linkOrder;
    return;
  }
  
  public void setNodeAnnotations(AnnotationSet annots) {
    nodeAnnotForLayout_ = annots;
    return;
  }
  
  public void setLinkAnnotations(Map<Boolean, AnnotationSet> annots) {
    linkAnnotsForLayout_ = annots;
    return;
  }
  
  public AnnotationSet getNodeAnnotations() {
    return (nodeAnnotForLayout_);
  }
  
  public Map<Boolean, AnnotationSet> getLinkAnnotations() {
    return (linkAnnotsForLayout_);
  }
    
  public void setGroupOrderAndMode(List<String> groupOrder, Network.LayoutMode mode, 
                                   boolean showLinkGroupAnnotations) {
    linkGroups_ = groupOrder;
    layoutMode_ = mode;
    showLinkGroupAnnotations_ = showLinkGroupAnnotations;
    return;
  }
  
  public void setLayoutMode(Network.LayoutMode mode) {
  	layoutMode_ = mode;
  	return;
  }
   
  public List<String> getGroupOrder() {
    return (linkGroups_);
  }
 
  public Network.LayoutMode getGroupOrderMode() {
    return (layoutMode_);
  } 
  
  public boolean getShowLinkGroupAnnotations() {
  	UiUtil.fixMePrintout("Not propagating into layout operation correctly");
    return (showLinkGroupAnnotations_);
  }
  
  public void setCTL(ControlTopLayout.CtrlMode cMode, ControlTopLayout.TargMode tMode, List<String> fixedOrder, BioFabricNetwork bfn) {
    this.normNameToIDs = (fixedOrder == null) ? null : bfn.getNormNameToIDs();
    this.cMode = cMode;
    this.tMode = tMode;
    this.fixedOrder = fixedOrder;
    return;
  }
  
  public void setPointUp(Boolean pointUp) {
    this.pointUp = pointUp;
    return;
  }
  
  public void setLinkMeaning(SetLayout.LinkMeans linkMeaning) {
    this.linkMeaning = linkMeaning;
    return;
  }
  
  public boolean getTurnOnShadows() {
  	return (this.turnOnShadows_);
  }
  
  public void setTurnOnShadows(boolean turnOn) {
    this.turnOnShadows_ = turnOn;
    return;
  }
  
  public boolean needsLayoutForRelayout() {
    switch (mode_) {
      case DEFAULT_LAYOUT:
      case WORLD_BANK_LAYOUT:
      case CONTROL_TOP_LAYOUT:
      case HIER_DAG_LAYOUT:
      case SET_LAYOUT:      
      case REORDER_LAYOUT:
      case CLUSTERED_LAYOUT:      
      case NODE_CLUSTER_LAYOUT:
      	return (true);
      case NODE_ATTRIB_LAYOUT:
      case LINK_ATTRIB_LAYOUT:
      case GROUP_PER_NODE_CHANGE:
      case GROUP_PER_NETWORK_CHANGE:
      	// Already installed!
        return (false);
      case SHADOW_LINK_CHANGE:
      case BUILD_FOR_SUBMODEL:
      case BUILD_FROM_XML:
      case BUILD_FROM_SIF:
      case BUILD_FROM_PLUGIN:
      default:
      	// Not legal!
        throw new IllegalStateException();
    }
  }

  public NodeLayout getNodeLayout() {
  	
  	if (plugInData_ != null) {
      NodeLayout nlp = plugInData_.getNodeLayout();
  	  if (nlp != null) {
  	  	return (nlp);
  	  }
    }

  	switch (mode_) {
  	  case DEFAULT_LAYOUT:
  	  case BUILD_FROM_SIF:
  	  	return (new DefaultLayout());
  	  case WORLD_BANK_LAYOUT:
  	  	return (new WorldBankLayout());
  	  case REORDER_LAYOUT:
      case CLUSTERED_LAYOUT:
        return (new NodeSimilarityLayout()); 	
      case NODE_CLUSTER_LAYOUT:
        return (new NodeClusterLayout());
      case CONTROL_TOP_LAYOUT:
        return (new ControlTopLayout(cMode, tMode, fixedOrder, normNameToIDs));
      case HIER_DAG_LAYOUT:  
        return (new HierDAGLayout(pointUp.booleanValue())); 
      case SET_LAYOUT:         
        return (new SetLayout(linkMeaning)); 
  	  default:
  	  	throw new IllegalStateException();
  	} 	
  }

  public EdgeLayout getEdgeLayout() {
  	
    if (plugInData_ != null) {
      EdgeLayout elp = plugInData_.getEdgeLayout();
  	  if (elp != null) {
  	  	return (elp);
  	  }
    }

  	switch (mode_) {
  	  case REORDER_LAYOUT:
      case CLUSTERED_LAYOUT:  
      case NODE_CLUSTER_LAYOUT:
      case SET_LAYOUT:
         // The above layouts do edge layout as part of node layout:
        return (null);
      case HIER_DAG_LAYOUT: 
      	return (new HierDAGLayout.EdgeLayout());	
  	  default:
  	  	return (new DefaultEdgeLayout());	
  	}
  }

  /***************************************************************************
  **
  ** For passing around ranked nodes
  */  
  
  static class DoubleRanked  {
     double rank;
     String id;
     Link byLink;

    DoubleRanked(double rank, String id, Link byLink) {
      this.rank = rank;
      this.id = id;
      this.byLink = byLink;
    } 
  }
}
```
