## Supplementary material for "BioFabric Visualization of Network Alignments": BioFabric Source Code: BuildExtractorImpl.java

```
/*
**    Copyright (C) 2003-2018 Institute for Systems Biology 
**                            Seattle, Washington, USA. 
**
**    This library is free software; you can redistribute it and/or
**    modify it under the terms of the GNU Lesser General Public
**    License as published by the Free Software Foundation; either
**    version 2.1 of the License, or (at your option) any later version.
**
**    This library is distributed in the hope that it will be useful,
**    but WITHOUT ANY WARRANTY; without even the implied warranty of
**    MERCHANTABILITY or FITNESS FOR A PARTICULAR PURPOSE.  See the GNU
**    Lesser General Public License for more details.
**
**    You should have received a copy of the GNU Lesser General Public
**    License along with this library; if not, write to the Free Software
**    Foundation, Inc., 59 Temple Place, Suite 330, Boston, MA  02111-1307  USA
*/

package org.systemsbiology.biofabric.io;

import java.util.Collection;
import java.util.HashMap;
import java.util.HashSet;
import java.util.Iterator;
import java.util.List;
import java.util.Map;
import java.util.Set;
import java.util.SortedMap;

import org.systemsbiology.biofabric.api.io.BuildExtractor;
import org.systemsbiology.biofabric.api.model.AugRelation;
import org.systemsbiology.biofabric.api.model.LinkComparator;
import org.systemsbiology.biofabric.api.model.NetLink;
import org.systemsbiology.biofabric.api.model.NetNode;
import org.systemsbiology.biofabric.api.worker.AsynchExitRequestException;
import org.systemsbiology.biofabric.api.worker.BTProgressMonitor;
import org.systemsbiology.biofabric.api.worker.LoopReporter;

/****************************************************************************
**
** Methods for extracting info while building networks
*/

public class BuildExtractorImpl implements BuildExtractor {
  
  ////////////////////////////////////////////////////////////////////////////
  //
  // PUBLIC STATIC METHODS
  //
  ////////////////////////////////////////////////////////////////////////////
  
  /***************************************************************************
  **
  ** Extract nodes
  */
  
  public Set<NetNode> extractNodes(Collection<NetLink> allLinks, Set<NetNode> loneNodeIDs,
                                        BTProgressMonitor monitor) throws AsynchExitRequestException {
    
    Set<NetNode> retval = new HashSet<NetNode>(loneNodeIDs);
    LoopReporter lr = new LoopReporter(allLinks.size(), 20, monitor, 0.0, 1.0, "progress.analyzingNodes");
    
    for (NetLink link : allLinks) {
      retval.add(link.getSrcNode());
      retval.add(link.getTrgNode());
      lr.report();
    }
    return (retval);
  }
 
  /***************************************************************************
  ** 
  ** Extract relations
  */

  public void extractRelations(List<NetLink> allLinks, 
  		                         SortedMap<AugRelation, Boolean> relMap, 
  		                         BTProgressMonitor monitor) throws AsynchExitRequestException {
    HashSet<NetLink> flipSet = new HashSet<NetLink>();
    HashSet<AugRelation> flipRels = new HashSet<AugRelation>();
    HashSet<AugRelation> rels = new HashSet<AugRelation>();
    int size = allLinks.size();
    LoopReporter lr = new LoopReporter(size, 20, monitor, 0.0, 1.0, "progress.analyzingRelations");
    Iterator<NetLink> alit = allLinks.iterator();
    while (alit.hasNext()) {
      NetLink nextLink = alit.next();
      lr.report();
      AugRelation relation = nextLink.getAugRelation();
      if (!nextLink.isFeedback()) {  // Autofeedback not flippable
        NetLink flipLink = nextLink.flipped();
        if (flipSet.contains(flipLink)) {
          flipRels.add(relation);
        } else {
          flipSet.add(nextLink);
        }
      }
      rels.add(relation);
    } 
    
    //
    // We have a hint that something is signed if there are two
    // separate links running in opposite directions!
    //
        
    Boolean noDir = new Boolean(false);
    Boolean haveDir = new Boolean(true);
    Iterator<AugRelation> rit = rels.iterator();
    while (rit.hasNext()) {
      AugRelation rel = rit.next();
      relMap.put(rel, (flipRels.contains(rel)) ? haveDir : noDir);
    }    
    return;
  }
  
  /***************************************************************************
  ** 
  ** Helper to drop to map to single name: useful
  */

  public Map<String, NetNode> reduceNameSetToOne(Map<String, Set<NetNode>> mapsToSets) { 
  	HashMap<String, NetNode> retval = new HashMap<String, NetNode>();
    Iterator<String> alit = mapsToSets.keySet().iterator();
    while (alit.hasNext()) {
      String nextName = alit.next();
      Set<NetNode> forName = mapsToSets.get(nextName);
      if (forName.size() != 1) {
      	throw new IllegalStateException();
      }
      retval.put(nextName, forName.iterator().next());
    }
    return (retval);
  }
  
  
  /***************************************************************************
  ** 
  ** Process a link set that has not had directionality established
  */

  public void assignDirections(List<NetLink> allLinks, 
  		                         Map<AugRelation, Boolean> relMap,
  		                         BTProgressMonitor monitor) throws AsynchExitRequestException { 
     
	  int numLink = allLinks.size();
	  LoopReporter lr = new LoopReporter(numLink, 20, monitor, 0.0, 1.0, "progress.installDirections");
	 
    Iterator<NetLink> alit = allLinks.iterator();
    while (alit.hasNext()) {
      NetLink nextLink = alit.next();
      lr.report();
      AugRelation rel = nextLink.getAugRelation();
      Boolean isDir = relMap.get(rel);
      nextLink.installDirection(isDir);
    }
    lr.finish();
    return;
  }

  /***************************************************************************
  ** 
  ** This culls a set of links to remove non-directional synonymous and
  ** duplicate links.  Note that shadow links have already been created
  ** and added to the allLinks list. 
  */

  public void preprocessLinks(List<NetLink> allLinks, Set<NetLink> retval, Set<NetLink> culled,
  		                        BTProgressMonitor monitor) throws AsynchExitRequestException {
  	LinkComparator flc = new LinkComparator();
  	int numLink = allLinks.size();
	  LoopReporter lr = new LoopReporter(numLink, 20, monitor, 0.0, 1.0, "progress.cullingAndFlipping");
  	
    Iterator<NetLink> alit = allLinks.iterator();
    while (alit.hasNext()) {
      NetLink nextLink = alit.next();
      lr.report();
      if (retval.contains(nextLink)) {
        culled.add(nextLink);
      } else if (!nextLink.isDirected()) {
        if (!nextLink.isFeedback()) {
          NetLink flipLink = nextLink.flipped();
          if (retval.contains(flipLink)) {
            // Make the order consistent for a given src & pair!
            if (flc.compare(nextLink, flipLink) < 0) {
              retval.remove(flipLink);
              culled.add(flipLink);
              retval.add(nextLink);
            } else {
              culled.add(nextLink);              
            }  
          } else {
            retval.add(nextLink);
          }
        } else {
          retval.add(nextLink);
        }
      } else {
        retval.add(nextLink);
      }
    }
    return;
  }
  
  /***************************************************************************
   **
   ** Generates a Node->Neighbors and Node->Links Map
   */
  
  public void createNeighborLinkMap(Collection<NetLink> allLinks, Set<NetNode> loneNodeIDs,
                                    Map<NetNode, Set<NetNode>> nodeToNeighbors, Map<NetNode, Set<NetLink>> nodeToLinks,
                                    BTProgressMonitor monitor) throws AsynchExitRequestException {
  
    LoopReporter lr = new LoopReporter(allLinks.size(), 20, monitor, 0.0, 1.0, "progress.generatingStructures");
  
    for (NetLink link : allLinks) {
      lr.report();
      NetNode src = link.getSrcNode(), trg = link.getTrgNode();
    
      if (nodeToLinks.get(src) == null) {
        nodeToLinks.put(src, new HashSet<NetLink>());
      }
      if (nodeToLinks.get(trg) == null) {
        nodeToLinks.put(trg, new HashSet<NetLink>());
      }
      if (nodeToNeighbors.get(src) == null) {
        nodeToNeighbors.put(src, new HashSet<NetNode>());
      }
      if (nodeToNeighbors.get(trg) == null) {
        nodeToNeighbors.put(trg, new HashSet<NetNode>());
      }
    
      nodeToLinks.get(src).add(link);
      nodeToLinks.get(trg).add(link);
      nodeToNeighbors.get(src).add(trg);
      nodeToNeighbors.get(trg).add(src);
    }
  
    for (NetNode node : loneNodeIDs) {
      nodeToLinks.put(node, new HashSet<NetLink>());
      nodeToNeighbors.put(node, new HashSet<NetNode>());
    }
    return;
  }
  
}
```
