## Supplementary material for "BioFabric Visualization of Network Alignments": BioFabric Source Code: FabricFactory.java

```
/*
**    Copyright (C) 2003-2017 Institute for Systems Biology 
**                            Seattle, Washington, USA. 
**
**    This library is free software; you can redistribute it and/or
**    modify it under the terms of the GNU Lesser General Public
**    License as published by the Free Software Foundation; either
**    version 2.1 of the License, or (at your option) any later version.
**
**    This library is distributed in the hope that it will be useful,
**    but WITHOUT ANY WARRANTY; without even the implied warranty of
**    MERCHANTABILITY or FITNESS FOR A PARTICULAR PURPOSE.  See the GNU
**    Lesser General Public License for more details.
**
**    You should have received a copy of the GNU Lesser General Public
**    License along with this library; if not, write to the Free Software
**    Foundation, Inc., 59 Temple Place, Suite 330, Boston, MA  02111-1307  USA
*/

package org.systemsbiology.biofabric.io;

import java.util.Set;
import java.util.HashSet;
import java.io.IOException;
import java.util.ArrayList;
import java.util.HashMap;
import java.util.Iterator;
import java.util.Map;

import org.xml.sax.Attributes;
import org.systemsbiology.biofabric.api.io.PluginWhiteboard;
import org.systemsbiology.biofabric.api.model.Annot;
import org.systemsbiology.biofabric.api.util.NID;
import org.systemsbiology.biofabric.api.util.UniqueLabeller;
import org.systemsbiology.biofabric.api.model.AnnotationSet;
import org.systemsbiology.biofabric.api.parser.ParserClient;
import org.systemsbiology.biofabric.model.BioFabricNetwork;
import org.systemsbiology.biofabric.plugin.BioFabricToolPlugIn;
import org.systemsbiology.biofabric.plugin.BioFabricToolPlugInData;
import org.systemsbiology.biofabric.plugin.PlugInManager;
import org.systemsbiology.biofabric.ui.FabricColorGenerator;
import org.systemsbiology.biofabric.ui.FabricDisplayOptions;
import org.systemsbiology.biofabric.ui.NamedColor;

/****************************************************************************
**
** This handles fabric creation from XML files
*/

public class FabricFactory implements ParserClient {

  ////////////////////////////////////////////////////////////////////////////
  //
  // PRIVATE INSTANCE VARIABLES
  //
  ////////////////////////////////////////////////////////////////////////////

  private HashSet<String> allKeys_; 
  private ParserClient currClient_;
  private HashMap<String, ParserClient> clients_;
  private FactoryWhiteboard whiteBoard_;
  
  ////////////////////////////////////////////////////////////////////////////
  //
  // PUBLIC CONSTRUCTORS
  //
  ////////////////////////////////////////////////////////////////////////////

  /***************************************************************************
  **
  ** Constructor
  */

  public FabricFactory(PlugInManager pMan) {
      
    allKeys_ = new HashSet<String>();
    whiteBoard_ = new FactoryWhiteboard();
   
    BioFabricNetwork.NetworkDataWorker ndw = new BioFabricNetwork.NetworkDataWorker(whiteBoard_, pMan);    
    ArrayList<ParserClient> alist = new ArrayList<ParserClient>();
    alist.add(ndw);
    
    allKeys_.addAll(ndw.keywordsOfInterest()); 

    Iterator<ParserClient> cit = alist.iterator();
    clients_ = new HashMap<String, ParserClient>();
    while (cit.hasNext()) {
      ParserClient pc = cit.next();
      Set<String> keys = pc.keywordsOfInterest();
      Iterator<String> ki = keys.iterator();
      while (ki.hasNext()) {
        String key = ki.next();
        Object prev = clients_.put(key, pc);
        if (prev != null) {
          throw new IllegalArgumentException();
        }
      }
    }
    currClient_ = null;
    
  }

  ////////////////////////////////////////////////////////////////////////////
  //
  // PUBLIC METHODS
  //
  ////////////////////////////////////////////////////////////////////////////

  /***************************************************************************
  ** 
  ** Get the network
  */

  public BioFabricNetwork getFabricNetwork() {
    return (whiteBoard_.bfn);    
  }
  
  /***************************************************************************
  ** 
  ** Set the container
  */

  public void setContainer(Object container) {
    if (currClient_ != null) {
      currClient_.setContainer(container);
    }    
    return;    
  }
    
  /***************************************************************************
  **
  ** Callback for completion of the element
  **
  */
  
  public boolean finishElement(String elemName) throws IOException {
    if (currClient_ == null) {
      return (false);
    }
    if (currClient_.finishElement(elemName)) {
      currClient_ = null;
    }
    return (allKeys_.contains(elemName));
  }
  
  /***************************************************************************
  ** 
  ** Handle incoming characters
  */

  public void processCharacters(char[] chars, int start, int length) {
    if (currClient_ != null) {
      currClient_.processCharacters(chars, start, length);
    }    
    return;
  }
  
  /***************************************************************************
  **
  ** Return the element keywords that we are interested in
  **
  */
  
  public Set<String> keywordsOfInterest() {
    return (allKeys_);
  }
  
  /***************************************************************************
  **
  ** Handle the attributes for the keyword
  **
  */
  
  public Object processElement(String elemName, Attributes attrs) throws IOException {

    if (currClient_ != null) {
      return (currClient_.processElement(elemName, attrs));
    }
    
    ParserClient pc = clients_.get(elemName);
    if (pc != null) {
      currClient_ = pc; 
      return (currClient_.processElement(elemName, attrs));
    }
    return (null);
  }

  public static class FactoryWhiteboard implements PluginWhiteboard {
  	public UniqueLabeller ulb;
    public BioFabricNetwork bfn;
    public BioFabricNetwork.LinkInfo linkInfo;
    public BioFabricNetwork.NodeInfo nodeInfo;
    public BioFabricNetwork.DrainZone drainZone;
    public FabricColorGenerator fcg;
    public NamedColor currCol;
    public int colTarg;
    public String groupTag;
    public FabricDisplayOptions displayOpts;
    public Map<String, NID.WithName> legacyMap;
  	public Map<NID, NID.WithName> wnMap;
  	public Annot annot;
  	public AnnotationSet currAnnots;
  	
  	public BioFabricToolPlugIn currPlugIn_;
  	public BioFabricToolPlugInData currPlugInData_;
  	
	
  	public BioFabricToolPlugIn getCurrentPlugIn() {
  		return (currPlugIn_);
  	}
  	
  	public void setCurrentPlugIn(BioFabricToolPlugIn pi) {
  		currPlugIn_ = pi;
  		return;
  	}
  	
  	public BioFabricToolPlugInData getCurrentPlugInData() {
  	  return (currPlugInData_);
  	}
  	
  	public void setCurrentPlugInData(BioFabricToolPlugInData pid) {
  		currPlugInData_ = pid;
  		return;
  	}
  	
    public FactoryWhiteboard() {
      colTarg = FabricColorGenerator.UNCHANGED;
      ulb = new UniqueLabeller();
      legacyMap = new HashMap<String, NID.WithName>();
      wnMap = new HashMap<NID, NID.WithName>();
    } 
  }
}
```
