## Supplementary material for "BioFabric Visualization of Network Alignments": BioFabric Source Code: FabricImportLoader.java

```
/*
**    Copyright (C) 2003-2018 Institute for Systems Biology 
**                            Seattle, Washington, USA. 
**
**    This library is free software; you can redistribute it and/or
**    modify it under the terms of the GNU Lesser General Public
**    License as published by the Free Software Foundation; either
**    version 2.1 of the License, or (at your option) any later version.
**
**    This library is distributed in the hope that it will be useful,
**    but WITHOUT ANY WARRANTY; without even the implied warranty of
**    MERCHANTABILITY or FITNESS FOR A PARTICULAR PURPOSE.  See the GNU
**    Lesser General Public License for more details.
**
**    You should have received a copy of the GNU Lesser General Public
**    License along with this library; if not, write to the Free Software
**    Foundation, Inc., 59 Temple Place, Suite 330, Boston, MA  02111-1307  USA
*/

package org.systemsbiology.biofabric.io;

import java.io.FileInputStream;
import java.io.BufferedReader;
import java.io.File;
import java.io.InputStreamReader;
import java.util.ArrayList;
import java.util.HashMap;
import java.io.IOException;
import java.util.List;
import java.util.Map;
import java.util.Set;

import org.systemsbiology.biofabric.api.model.NetLink;
import org.systemsbiology.biofabric.api.model.NetNode;
import org.systemsbiology.biofabric.api.util.NID;
import org.systemsbiology.biofabric.api.util.UniqueLabeller;
import org.systemsbiology.biofabric.api.worker.AsynchExitRequestException;
import org.systemsbiology.biofabric.api.worker.BTProgressMonitor;
import org.systemsbiology.biofabric.api.worker.LoopReporter;
import org.systemsbiology.biofabric.model.FabricLink;
import org.systemsbiology.biofabric.model.FabricNode;
import org.systemsbiology.biofabric.util.DataUtil;

/****************************************************************************
 **
 ** This loads SIF files
 */

public abstract class FabricImportLoader {
  
  ////////////////////////////////////////////////////////////////////////////
  //
  // PRIVATE CONSTANTS
  //
  ////////////////////////////////////////////////////////////////////////////
  
  ////////////////////////////////////////////////////////////////////////////
  //
  // PUBLIC CONSTANTS
  //
  ////////////////////////////////////////////////////////////////////////////
  
  ////////////////////////////////////////////////////////////////////////////
  //
  // PRIVATE INSTANCE MEMBERS
  //
  ////////////////////////////////////////////////////////////////////////////
  
  ////////////////////////////////////////////////////////////////////////////
  //
  // PUBLIC CONSTRUCTORS
  //
  ////////////////////////////////////////////////////////////////////////////
  
  /***************************************************************************
   **
   ** Constructor
   */
  
  public FabricImportLoader() {
  
  }
  
  ////////////////////////////////////////////////////////////////////////////
  //
  // PUBLIC METHODS
  //
  ////////////////////////////////////////////////////////////////////////////
  
  /***************************************************************************
   **
   ** Parse a line to tokens
   */
  
  protected abstract String[] lineToToks(String line, FileImportStats stats) throws IOException;
  
  /***************************************************************************
   **
   ** Consume tokens, make links
   */
  
  protected abstract void consumeTokens(String[] tokens, UniqueLabeller idGen, List<NetLink> links,
                                        Set<NetNode> loneNodeIDs, Map<String, String> nameMap, Integer magBins,
                                        Map<String, NetNode> nameToID, FileImportStats stats) throws IOException;
  
  /***************************************************************************
   **
   ** Process a SIF input
   */
  
  public FileImportStats importFabric(File infile, UniqueLabeller idGen, List<NetLink> links,
                                      Set<NetNode> loneNodeIDs, Map<String, String> nameMap, Integer magBins,
                                      BTProgressMonitor monitor) throws AsynchExitRequestException, IOException {
    
    FileImportStats retval = new FileImportStats();
    long fileLen = infile.length();
    HashMap<String, NetNode> nameToID = new HashMap<String, NetNode>();
    BufferedReader in = null;
    ArrayList<String[]> tokSets = new ArrayList<String[]>();
    LoopReporter lr = new LoopReporter(fileLen, 20, monitor, 0.0, 1.0, "progress.readingFile");
    try {
      in = new BufferedReader(new InputStreamReader(new FileInputStream(infile), "UTF-8"));
      String line = null;
      while ((line = in.readLine()) != null) {
        lr.report(line.length() + 1);
        if (line.trim().equals("")) {
          continue;
        }
        String[] tokens = lineToToks(line, retval);
        if (tokens != null) {
          tokSets.add(tokens);
        }
      }
    } finally {
      if (in != null) {
        in.close();
      }
    }
    lr.finish();
    
    int numLines = tokSets.size();
    lr = new LoopReporter(numLines, 20, monitor, 0.0, 1.0, "progress.buildingEdgesAndNodes");
    
    for (int i = 0; i < numLines; i++) {
      String[] tokens = tokSets.get(i);
      lr.report();
      consumeTokens(tokens, idGen, links, loneNodeIDs, nameMap, magBins, nameToID, retval);
    }
    lr.finish();
    return (retval);
  }
  
  ////////////////////////////////////////////////////////////////////////////
  //
  // PROTECTED METHODS
  //
  ////////////////////////////////////////////////////////////////////////////
  
  /***************************************************************************
   **
   ** Strip quoted string
   */
  
  protected String stripQuotes(String inString) {
    String procString = inString.trim();
    if ((procString.indexOf("\"") == 0) && (procString.lastIndexOf("\"") == (procString.length() - 1))) {
      procString = procString.replaceAll("\"", "");
    }
    return (procString);
  }
  
  /***************************************************************************
   **
   ** Strip Brackets |{ }| from String
   */
  
  protected String stripBrackets(String inString) {
    String procString = inString.trim();
    if ((procString.indexOf("|{") == 0) && (procString.lastIndexOf("}|") == (procString.length() - 2))) {
      procString = procString.replaceAll("\\|\\{", "");
      procString = procString.replaceAll("\\}\\|", "");
    }
    return (procString);
  }
  
  /***************************************************************************
   **
   ** Map name
   */
  
  protected String mapName(String inString, Map<String, String> nameMap) {
    String retval = inString;
    if (nameMap == null) {
      return (retval);
    }
    String mappedString = nameMap.get(inString);
    if (mappedString != null) {
      retval = mappedString;
    }
    return (retval);
  }
  
  /***************************************************************************
   **
   ** Get an actual node ID
   */
  
  protected NetNode nameToNode(String inString, UniqueLabeller idGen, Map<String, NetNode> nameToID) {
    
    String normName = DataUtil.normKey(inString);
    
    NetNode nodeID = nameToID.get(normName);
    if (nodeID == null) {
      NID nid = idGen.getNextOID();
      nodeID = new FabricNode(nid, inString);
      nameToID.put(normName, nodeID);
    }
    return (nodeID);
  }
  
  /***************************************************************************
   **
   ** Get an actual node ID
   */
  
  protected void buildLinkAndShadow(NetNode srcID, NetNode trgID, String rel, List<NetLink> links) {
    
    FabricLink nextLink = new FabricLink(srcID, trgID, rel, false);
    links.add(nextLink);
    
    // We never create shadow feedback links!
    if (!srcID.equals(trgID)) {
      FabricLink nextShadowLink = new FabricLink(srcID, trgID, rel, true);
      links.add(nextShadowLink);
    }
    
    return;
  }
  
  ////////////////////////////////////////////////////////////////////////////
  //
  // PUBLIC INNER CLASSES
  //
  ////////////////////////////////////////////////////////////////////////////
  
  public static class FileImportStats {
    public ArrayList<String> badLines;
    
    public FileImportStats() {
      badLines = new ArrayList<String>();
    }
    
    public void copyInto(FileImportStats other) {
      this.badLines = new ArrayList<String>(other.badLines);
      return;
    }
  }
}
```
