## Supplementary material for "BioFabric Visualization of Network Alignments": BioFabric Source Code: FileLoadFlowsImpl.java

```
/*
**    Copyright (C) 2003-2018 Institute for Systems Biology 
**                            Seattle, Washington, USA. 
**
**    This library is free software; you can redistribute it and/or
**    modify it under the terms of the GNU Lesser General Public
**    License as published by the Free Software Foundation; either
**    version 2.1 of the License, or (at your option) any later version.
**
**    This library is distributed in the hope that it will be useful,
**    but WITHOUT ANY WARRANTY; without even the implied warranty of
**    MERCHANTABILITY or FITNESS FOR A PARTICULAR PURPOSE.  See the GNU
**    Lesser General Public License for more details.
**
**    You should have received a copy of the GNU Lesser General Public
**    License along with this library; if not, write to the Free Software
**    Foundation, Inc., 59 Temple Place, Suite 330, Boston, MA  02111-1307  USA
*/

package org.systemsbiology.biofabric.io;

import java.awt.Component;
import java.awt.Dimension;
import java.awt.Toolkit;
import java.awt.image.BufferedImage;
import java.io.BufferedInputStream;
import java.io.BufferedWriter;
import java.io.File;
import java.io.FileInputStream;
import java.io.FileOutputStream;
import java.io.IOException;
import java.io.InputStream;
import java.io.OutputStream;
import java.io.OutputStreamWriter;
import java.io.PrintWriter;
import java.text.MessageFormat;
import java.util.ArrayList;
import java.util.HashMap;
import java.util.HashSet;
import java.util.Iterator;
import java.util.List;
import java.util.Map;
import java.util.Set;
import java.util.SortedMap;
import java.util.TreeMap;
import java.util.zip.GZIPInputStream;
import java.util.zip.GZIPOutputStream;

import javax.swing.JFileChooser;
import javax.swing.JFrame;
import javax.swing.JOptionPane;
import javax.swing.filechooser.FileFilter;

import org.systemsbiology.biofabric.api.io.AttributeKey;
import org.systemsbiology.biofabric.api.io.BuildData;
import org.systemsbiology.biofabric.api.io.BuildExtractor;
import org.systemsbiology.biofabric.api.io.FileLoadFlows;
import org.systemsbiology.biofabric.api.io.Indenter;
import org.systemsbiology.biofabric.api.layout.DefaultLayout;
import org.systemsbiology.biofabric.api.layout.EdgeLayout;
import org.systemsbiology.biofabric.api.layout.LayoutCriterionFailureException;
import org.systemsbiology.biofabric.api.layout.NodeLayout;
import org.systemsbiology.biofabric.api.model.AnnotationSet;
import org.systemsbiology.biofabric.api.model.AugRelation;
import org.systemsbiology.biofabric.api.model.NetLink;
import org.systemsbiology.biofabric.api.model.NetNode;
import org.systemsbiology.biofabric.api.parser.ParserClient;
import org.systemsbiology.biofabric.api.util.ExceptionHandler;
import org.systemsbiology.biofabric.api.util.PluginResourceManager;
import org.systemsbiology.biofabric.api.util.UniqueLabeller;
import org.systemsbiology.biofabric.api.worker.AsynchExitRequestException;
import org.systemsbiology.biofabric.api.worker.BFWorker;
import org.systemsbiology.biofabric.api.worker.BTProgressMonitor;
import org.systemsbiology.biofabric.api.worker.BackgroundCore;
import org.systemsbiology.biofabric.api.worker.BackgroundWorkerControlManager;
import org.systemsbiology.biofabric.api.worker.BackgroundWorkerOwner;
import org.systemsbiology.biofabric.app.BioFabricWindow;
import org.systemsbiology.biofabric.cmd.CommandSet;
import org.systemsbiology.biofabric.cmd.HeadlessOracle;
import org.systemsbiology.biofabric.layouts.ControlTopLayout;
import org.systemsbiology.biofabric.layouts.NodeClusterLayout;
import org.systemsbiology.biofabric.layouts.NodeSimilarityLayout;
import org.systemsbiology.biofabric.layouts.SetLayout;
import org.systemsbiology.biofabric.model.BioFabricNetwork;
import org.systemsbiology.biofabric.parser.ProgressFilterInputStream;
import org.systemsbiology.biofabric.parser.SUParser;
import org.systemsbiology.biofabric.plugin.BioFabricToolPlugInData;
import org.systemsbiology.biofabric.plugin.PluginSupportFactory;
import org.systemsbiology.biofabric.plugin.PlugInManager;
import org.systemsbiology.biofabric.plugin.PlugInNetworkModelAPI;
import org.systemsbiology.biofabric.ui.FabricColorGenerator;
import org.systemsbiology.biofabric.ui.FabricDisplayOptions;
import org.systemsbiology.biofabric.ui.FabricDisplayOptionsManager;
import org.systemsbiology.biofabric.ui.dialogs.RelationDirectionDialog;
import org.systemsbiology.biofabric.ui.display.BioFabricPanel;
import org.systemsbiology.biofabric.ui.render.BufferBuilder;
import org.systemsbiology.biofabric.util.FileExtensionFilters;
import org.systemsbiology.biofabric.util.GarbageRequester;
import org.systemsbiology.biofabric.util.InvalidInputException;
import org.systemsbiology.biofabric.util.ResourceManager;
import org.systemsbiology.biofabric.util.UiUtil;
import org.systemsbiology.biotapestry.biofabric.FabricCommands;

/****************************************************************************
**
** Handles File Loading Tasks
*/

public class FileLoadFlowsImpl implements FileLoadFlows {

  ////////////////////////////////////////////////////////////////////////////
  //
  // PUBLIC CONSTANTS
  //
  //////////////////////////////////////////////////////////////////////////// 

  public static final int LINK_COUNT_FOR_BACKGROUND_WRITE = 5000;
  public static final int FILE_LENGTH_FOR_BACKGROUND_FILE_READ = 500000;
  public static final int SIZE_TO_ASK_ABOUT_SHADOWS = 100000;
  public static final int XML_SIZE_FOR_BACKGROUND_READ = 1000000;
  
  /***************************************************************************
  **
  ** For standard file checks
  */

  public static final boolean FILE_MUST_EXIST_DONT_CARE = false;
  public static final boolean FILE_MUST_EXIST           = true;
  
  public static final boolean FILE_CAN_CREATE_DONT_CARE = false;
  public static final boolean FILE_CAN_CREATE           = true;
  
  public static final boolean FILE_DONT_CHECK_OVERWRITE = false;
  public static final boolean FILE_CHECK_OVERWRITE      = true;
  
  public static final boolean FILE_MUST_BE_FILE         = false;
  public static final boolean FILE_MUST_BE_DIRECTORY    = true;  
          
  public static final boolean FILE_CAN_WRITE_DONT_CARE  = false;
  public static final boolean FILE_CAN_WRITE            = true;
  
  public static final boolean FILE_CAN_READ_DONT_CARE   = false;
  public static final boolean FILE_CAN_READ             = true;
  
  public static enum FileLoadType {GW, SIF};
      
  ////////////////////////////////////////////////////////////////////////////
  //
  // MEMBERS
  //
  ////////////////////////////////////////////////////////////////////////////  
  
  private HeadlessOracle headlessOracle_;
  private JFrame topWindow_;
  private BioFabricWindow bfw_;
  private BioFabricPanel bfp_;
  private File currentFile_;
  private PlugInManager pMan_;
  private FabricColorGenerator colGen_;
  private CommandSet cSet_;
  private boolean isForMain_;

  ////////////////////////////////////////////////////////////////////////////
  //
  // CONSTRUCTORS
  //
  ////////////////////////////////////////////////////////////////////////////
  
  /***************************************************************************
  **
  ** Constructor 
  */ 
  
  public FileLoadFlowsImpl(BioFabricWindow bfw, PlugInManager pMan,
                           FabricColorGenerator colGen, CommandSet cSet,
                           HeadlessOracle headlessOracle, boolean isForMain) {
    bfw_ = bfw;
    pMan_ = pMan;
    topWindow_ = bfw.getWindow();
    headlessOracle_ = headlessOracle;
    colGen_ = colGen;
    cSet_ = cSet;
    isForMain_ = isForMain;
  }    

  ////////////////////////////////////////////////////////////////////////////
  //
  // PUBLIC METHODS
  //
  ////////////////////////////////////////////////////////////////////////////  

  /***************************************************************************
  **
  ** Set the fabric panel
  */ 
  
  public void setFabricPanel(BioFabricPanel bfp) {
    bfp_ = bfp;
    if (isForMain_) {
      pMan_.installAPI(new PlugInInfo(this, bfp_.getNetwork(), bfw_));
    }
    return;
  } 
  
  /***************************************************************************
  **
  ** Do network build for a plug-in that provides the needed custom BuildData
  */ 
     
  public void buildNetworkForPlugIn(BuildData pluginData, File holdIt, PluginResourceManager rMan) { 
    NetworkBuilder nb = new FileLoadFlowsImpl.NetworkBuilder(true, holdIt, rMan);
    nb.setForPlugInBuild(pluginData);
    nb.doNetworkBuild();
    return;
  }
  
  /***************************************************************************
  **
  ** Do basic relayout
  */ 
     
  public void doBasicRelayout(BuildDataImpl.BuildMode bMode) { 
    (new NetworkRelayout()).doNetworkRelayout(bfp_.getNetwork(), bMode); 
    return;
  }
   
  /***************************************************************************
  **
  ** Do default relayout
  */ 
     
  public void doDefaultRelayout(DefaultLayout.Params params) { 
    NetworkRelayout nb = new NetworkRelayout();
    nb.setParams(params);
    nb.doNetworkRelayout(bfp_.getNetwork(), BuildDataImpl.BuildMode.DEFAULT_LAYOUT); 
    return;
  }

  /***************************************************************************
  **
  ** Do shape match relayout
  */ 
     
  public void doConnectivityRelayout(NodeSimilarityLayout.ClusterParams params) { 
    NetworkRelayout nb = new NetworkRelayout();
    nb.setParams(params);
    nb.doNetworkRelayout(bfp_.getNetwork(), BuildDataImpl.BuildMode.CLUSTERED_LAYOUT);    
    return;
  }
    
  /***************************************************************************
  **
  ** Do recolor
  */ 
    
  public void doDisplayOptionChange() {
    File holdIt;  
    try {
      holdIt = File.createTempFile("BioFabricHold", ".zip");
      holdIt.deleteOnExit();
    } catch (IOException ioex) {
      holdIt = null;
    }
   
    BioFabricNetwork bfn = bfp_.getNetwork();
    if (bfn != null) {
      NetworkBuilder nb = new NetworkBuilder(true, holdIt, null);
      nb.setForDisplayOptionChange(bfn, BuildDataImpl.BuildMode.SHADOW_LINK_CHANGE);
      nb.doNetworkBuild();
    }
    return;
  } 
  
  /***************************************************************************
  **
  ** Do recolor
  */ 
     
  public void doRecolor(boolean isForMain) {
    File holdIt;  
    try {
      holdIt = File.createTempFile("BioFabricHold", ".zip");
      holdIt.deleteOnExit();
    } catch (IOException ioex) {
      holdIt = null;
    }
    NetworkRecolor nb = new NetworkRecolor(); 
    nb.doNetworkRecolor(isForMain, holdIt);
    return;
  }

  /***************************************************************************
  **
  ** Do shape match relayout
  */ 
     
  public void doShapeMatchRelayout(NodeSimilarityLayout.ResortParams params) {
    NetworkRelayout nb = new NetworkRelayout();
    nb.setParams(params);
    nb.doNetworkRelayout(bfp_.getNetwork(), BuildDataImpl.BuildMode.REORDER_LAYOUT);   
    return;
  }

  /***************************************************************************
  **
  ** Do Link relayout
  */ 
     
  public void doLinkGroupRelayout(BioFabricNetwork bfn, List<String> groupOrder, 
                                  BioFabricNetwork.LayoutMode mode, 
                                  boolean showGroupLinkAnnotations, 
                                  BuildDataImpl.BuildMode bmode) {
    NetworkRelayout nb = new NetworkRelayout();
    nb.setGroupOrderAndMode(groupOrder, mode, showGroupLinkAnnotations);
    nb.doNetworkRelayout(bfn, bmode);
    return;
  }

  /***************************************************************************
   **
   ** Do set relayout
   */ 
      
   public void doSetRelayout(SetLayout.LinkMeans linkMeaning) {
     NetworkRelayout nb = new NetworkRelayout();
     nb.setLinkMeaning(linkMeaning);
     nb.doNetworkRelayout(bfp_.getNetwork(), BuildDataImpl.BuildMode.SET_LAYOUT);
     return;
   }
  
  /***************************************************************************
  **
  ** Do HierDag relayout
  */ 
     
  public void doHierDagRelayout(boolean pointUp) {
    NetworkRelayout nb = new NetworkRelayout();
    nb.setPointUp(pointUp);
    nb.doNetworkRelayout(bfp_.getNetwork(), BuildDataImpl.BuildMode.HIER_DAG_LAYOUT); 
    return;
  }
  
  /***************************************************************************
  **
  ** Do Control Top relayout
  */ 
     
  public void doControlTopRelayout(ControlTopLayout.CtrlMode cMode,
                                   ControlTopLayout.TargMode tMode, List<String> fixedList) {
    NetworkRelayout nb = new NetworkRelayout();
    nb.setControlTopModes(cMode, tMode, fixedList);
    nb.doNetworkRelayout(bfp_.getNetwork(), BuildDataImpl.BuildMode.CONTROL_TOP_LAYOUT); 
    return;
  }
  
  /***************************************************************************
  **
  ** Do Cluster relayout
  */ 
     
  public void doNetworkClusterRelayout(NodeClusterLayout.ClusterParams params) {
    NetworkRelayout nb = new NetworkRelayout();
    nb.setParams(params);
    nb.doNetworkRelayout(bfp_.getNetwork(), BuildDataImpl.BuildMode.NODE_CLUSTER_LAYOUT);
    return;
  }

  /***************************************************************************
  **
  ** Do network link relayout
  */ 
     
  public void doNetworkLinkRelayout(SortedMap<Integer, NetLink> modifiedAndChecked) {
    NetworkRelayout nb = new NetworkRelayout();
    nb.setLinkOrder(modifiedAndChecked);
    nb.doNetworkRelayout(bfp_.getNetwork(), BuildDataImpl.BuildMode.LINK_ATTRIB_LAYOUT);  
    return;
  }

  /***************************************************************************
  **
  ** Do network relayout
  */ 
     
  public void doNetworkRelayout(Map<AttributeKey, String> nodeAttributes) {
    NetworkRelayout nb = new NetworkRelayout();
    nb.setNodeOrderFromAttrib(nodeAttributes);
    nb.doNetworkRelayout(bfp_.getNetwork(), BuildDataImpl.BuildMode.NODE_ATTRIB_LAYOUT);   
    return;
  }

  /***************************************************************************
  **
  ** Do a background write
  */ 
     
  public void doBackgroundWrite(OutputStream stream) {
    BackgroundFileWriter bw = new BackgroundFileWriter(); 
    bw.doBackgroundWrite(stream);
    return;
  }

  /***************************************************************************
  **
  ** Set the current file
  */ 
    
  public void setCurrentXMLFile(File file) {
    currentFile_ = file;
    if (currentFile_ == null) {
      return;
    }
    FabricCommands.setPreference("LoadDirectory", file.getAbsoluteFile().getParent());
    return;
  }
  
  /***************************************************************************
  **
  ** Third step for loading from SIF
  */
    
  private boolean buildTheNetworkFomLinks(File file, UniqueLabeller idGen,
  		                                    Set<NetNode> loneNodeIDs, 
  		                                    Set<NetLink> reducedLinks, File holdIt) {
  	try {
      NetworkBuilder nb = new NetworkBuilder(true, holdIt, null);
      nb.setForSifBuild(idGen, reducedLinks, loneNodeIDs, BuildDataImpl.BuildMode.BUILD_FROM_SIF);  
      if (this.headlessOracle_ == null) {
        nb.doNetworkBuild();            
      } else {
        nb.doNetworkBuildForeground();
      }
    } catch (OutOfMemoryError oom) {
      ExceptionHandler.getHandler().displayOutOfMemory(oom);
      return (false);  
    }
    currentFile_ = null;
    FabricCommands.setPreference("LoadDirectory", file.getAbsoluteFile().getParent());
    manageWindowTitle(file.getName());
    return (true);
  } 
 
  /***************************************************************************
  **
  ** Announce bad lines
  */
    
  private void announceBadLines(FabricImportLoader.FileImportStats sss) {
    ResourceManager rMan = ResourceManager.getManager();
    if (!sss.badLines.isEmpty()) {        
      String badLineFormat = rMan.getString("fabricRead.badLineFormat");
      String badLineMsg = MessageFormat.format(badLineFormat, new Object[] {Integer.valueOf(sss.badLines.size())});
      if (headlessOracle_ == null) {
        JOptionPane.showMessageDialog(topWindow_, badLineMsg,
                                      rMan.getString("fabricRead.badLineTitle"),
                                      JOptionPane.WARNING_MESSAGE);
      } else {
        headlessOracle_.displayErrorMessage(badLineMsg);
      }
    }
    return;
  }

  /***************************************************************************
  **
  ** Second step for loading from SIF
  */
    
  public boolean handleDirectionsDupsAndShadows(List<NetLink> links, Set<NetNode> loneNodeIDs, 
  		                                           boolean binMag, SortedMap<AugRelation, Boolean> relaMap,
  		                                           Set<NetLink> reducedLinks, File holdIt, 
  		                                           boolean doForceUndirected, boolean skipShadowQuestion) {
    
    
    ResourceManager rMan = ResourceManager.getManager(); 
    try {     
      if (doForceUndirected) {
        relaMap = RelationDirectionDialog.forceAllUndirected(relaMap);
      } else {
        if (headlessOracle_ == null) {
          RelationDirectionDialog rdd = new RelationDirectionDialog(topWindow_, relaMap);
          rdd.setVisible(true);
          if (!rdd.haveResult()) {
            return (false);
          }
          if (rdd.getFromFile()) {
            File fileEda = getTheFile(".rda", ".txt", "AttribDirectory", "filterName.rda", null);
            if (fileEda == null) {
              return (true);
            }
            Map<AttributeKey, String> relAttributes = loadTheFile(fileEda, null, true);  // Use the simple a = b format of node attributes
            if (relAttributes == null) {
              return (true);
            }
            
            HashSet<AugRelation> needed = new HashSet<AugRelation>(relaMap.keySet());
          
            boolean tooMany = false;
            Iterator<AttributeKey> rit = relAttributes.keySet().iterator();
            while (rit.hasNext()) {
              AttributeLoader.StringKey sKey = (AttributeLoader.StringKey)rit.next();
              String key = sKey.key;
              String val = relAttributes.get(sKey);
              Boolean dirVal = Boolean.valueOf(val);
              AugRelation forNorm = new AugRelation(key, false);
              AugRelation forShad = new AugRelation(key, true);
              boolean matched = false;
              if (needed.contains(forNorm)) {
                matched = true;
                relaMap.put(forNorm, dirVal);
                needed.remove(forNorm);
              }
              if (needed.contains(forShad)) {
                matched = true;
                relaMap.put(forShad, dirVal);
                needed.remove(forShad);
              }
              if (!matched) {
                tooMany = true;
                break;
              }          
            }
            if (!needed.isEmpty() || tooMany) {
              JOptionPane.showMessageDialog(topWindow_, rMan.getString("fabricRead.directionMapLoadFailure"),
                                            rMan.getString("fabricRead.directionMapLoadFailureTitle"),
                                            JOptionPane.ERROR_MESSAGE);
              return (false);
            }
          } else {
            relaMap = rdd.getRelationMap();
          }
        }
      }
      
      HashSet<NetLink> culledLinks = new HashSet<NetLink>();
      
      if (headlessOracle_ == null) {
        boolean didFinish = backPreprocess(links, relaMap, reducedLinks, culledLinks, holdIt);
        if (!didFinish) {
          return (false);
        }
      } else {   
        try {
          preprocess(links, relaMap, reducedLinks, culledLinks, null);
        } catch ( AsynchExitRequestException axex) {
          // Not going to happen
        }
      }

      if (!culledLinks.isEmpty()) {
        String dupLinkFormat = rMan.getString("fabricRead.dupLinkFormat");
        // Ignore shadow link culls: / 2
        String dupLinkMsg = MessageFormat.format(dupLinkFormat, new Object[] {Integer.valueOf(culledLinks.size() / 2)});
        JOptionPane.showMessageDialog(topWindow_, dupLinkMsg,
                                      rMan.getString("fabricRead.dupLinkTitle"),
                                      JOptionPane.WARNING_MESSAGE);
      }
      
      //
      // For big files, user may want to specify layout options before the default layout with no
      // shadows. Let them set this here:
      //
      if (!skipShadowQuestion) {
	      if (reducedLinks.size() > SIZE_TO_ASK_ABOUT_SHADOWS) {
		      String shadowMessage = rMan.getString("fabricRead.askAboutShadows");
		      int doShadow =
		        JOptionPane.showConfirmDialog(topWindow_, shadowMessage,
		                                      rMan.getString("fabricRead.askAboutShadowsTitle"),
		                                      JOptionPane.YES_NO_CANCEL_OPTION);        
		      if (doShadow == JOptionPane.CANCEL_OPTION) {
		        return (false);
		      }
		      FabricDisplayOptions dops = FabricDisplayOptionsManager.getMgr().getDisplayOptions();
		      dops.setDisplayShadows((doShadow == JOptionPane.YES_OPTION));
		    }
      }
      
      //
      // Handle magnitude bins. Not supported in yet in V2.
      //
          
      if (binMag) {
      	/*
      	HashSet<NetLink> binnedLinks = new HashSet<NetLink>();
        Pattern p = Pattern.compile("[-+]?[0-9]*\\.?[0-9]+([eE][-+]?[0-9]+)?");
        
        Iterator<NetLink> alit = reducedLinks.iterator();
        while (alit.hasNext()) {
          NetLink nextLink = alit.next();
          AugRelation rel = nextLink.getAugRelation();
          Matcher m = p.matcher(rel.relation);
          int magCount = 0;
          if (m.find()) {      
            double d = Double.parseDouble(m.group(0));
            magCount = (int)Math.floor((Math.abs(d) * 10.0) - 5.0);
          }
         // for (int k = 0; k < magCount; k++) {
        	//  String suf = ":" + Integer.toString(k);
	      //    FabricLink nextLink = new FabricLink(source, target, rel + suf, false);
	      //  links.add(nextLink);
	        // We never create shadow feedback links!
	      //  if (!source.equals(target)) {
	       //   FabricLink nextShadowLink = new FabricLink(source, target, rel + suf, true);
	       //   links.add(nextShadowLink);
	      //  }       
        }*/
      }
    } catch (OutOfMemoryError oom) {
      ExceptionHandler.getHandler().displayOutOfMemory(oom);
      return (false);  
    }
    return (true);
  }
  
  /***************************************************************************
  **
  ** Do preprocessing
  */
  
  public boolean backPreprocess(List<NetLink> links, SortedMap<AugRelation, Boolean> relMap,
                            Set<NetLink> reducedLinks, Set<NetLink> culledLinks, File holdIt) {
    PreprocessNetwork pn = new PreprocessNetwork();
    boolean didFinish = pn.doNetworkPreprocess(links, relMap, reducedLinks, culledLinks, holdIt);
    return (didFinish);
  }
  
  /***************************************************************************
   **
   ** Common load operation for gw or sif
   */
  
  public FileLoadFlows.FileLoadResult loadFromASource(File file, Map<AttributeKey, String> nameMap,
                                                      Integer magBins, UniqueLabeller idGen, 
                                                      FileLoadFlows.FileLoadType type, boolean skipShadowQuestion) {
    
    HashMap<String, String> nodeNames = null;
    if (nameMap != null) {
      nodeNames = new HashMap<String, String>();
      for (AttributeKey key : nameMap.keySet()) {
        nodeNames.put(((AttributeLoader.StringKey)key).key, nameMap.get(key));
      }
    }
  
    File holdIt;
    try {
      holdIt = File.createTempFile("BioFabricHold", ".zip");
      holdIt.deleteOnExit();
    } catch (IOException ioex) {
      holdIt = null;
    }
  
    ArrayList<NetLink> links = new ArrayList<NetLink>();
    HashSet<NetNode> loneNodes = new HashSet<NetNode>();
    TreeMap<AugRelation, Boolean> relMap = new TreeMap<AugRelation, Boolean>();
    HashSet<NetLink> reducedLinks = new HashSet<NetLink>();
    FabricImportLoader.FileImportStats sss;
    
    if ((file.length() > FILE_LENGTH_FOR_BACKGROUND_FILE_READ) && (headlessOracle_ == null)) {
      sss = new FabricImportLoader.FileImportStats();
      BackgroundFileReader br = new BackgroundFileReader();
      //
      // This gets file in:
      //
      boolean finished;
      if (type == FileLoadFlows.FileLoadType.SIF) {
        finished = br.doBackgroundSIFRead(file, idGen, links, loneNodes, nodeNames, sss, magBins, relMap, holdIt);
      } else if (type == FileLoadFlows.FileLoadType.GW) {
        // need to process gw links if no relations provided
        finished = br.doBackgroundGWRead(file, idGen, links, loneNodes, nodeNames, sss, true, magBins, relMap, holdIt);
      } else {
        throw (new IllegalArgumentException("File type not identified"));
      }
      if (!finished) {
        return (new FileLoadResultImpl(true, holdIt));
      }
    } else {
      try {
        if (type == FileLoadFlows.FileLoadType.SIF) {
          sss = (new SIFImportLoader()).importFabric(file, idGen, links, loneNodes, nodeNames, magBins, null);
        } else if (type == FileLoadFlows.FileLoadType.GW) {
          sss = (new GWImportLoader()).importFabric(file, idGen, links, loneNodes, nodeNames, magBins, null);
          boolean finished = (new GWImportLoader.GWRelationManager()).process(links, topWindow_, null);
          if (!finished) {
            // should not happen
            return (new FileLoadResultImpl(false, holdIt));
          }
        } else {
          throw (new IllegalArgumentException("File type not identified"));
        }
        PluginSupportFactory.getBuildExtractor().extractRelations(links, relMap, null);
      } catch (AsynchExitRequestException axex) {
        // Should never happen
        return (new FileLoadResultImpl(false, holdIt));
      } catch (IOException ioe) {
        displayFileInputError(ioe);
        return (new FileLoadResultImpl(false, holdIt));
      } catch (OutOfMemoryError oom) {
        ExceptionHandler.getHandler().displayOutOfMemory(oom);
        return (new FileLoadResultImpl(false, holdIt));
      }
    }
  
    //
    // This looks for dups to toss and prep work:
    //
    
    announceBadLines(sss);
    boolean finished = handleDirectionsDupsAndShadows(links, loneNodes, (magBins != null), 
    		                                              relMap, reducedLinks, holdIt, false,
    		                                              skipShadowQuestion);
    if (finished) {
      buildTheNetworkFomLinks(file, idGen, loneNodes, reducedLinks, holdIt);
    }
    return (new FileLoadResultImpl(true, holdIt));
  }
  
  /***************************************************************************
   **
   ** Load from file and directly receive link set and loners set
   */
  
  public FileLoadFlows.FileLoadResult loadFromASource(File file, List<NetLink> links,
                                                      Set<NetNode> loneNodes, Integer magBins,
                                                      UniqueLabeller idGen, boolean loadOnly, 
                                                      FileLoadFlows.FileLoadType type, 
                                                      boolean skipShadowQuestion) {
    
    File holdIt;
    try {
      holdIt = File.createTempFile("BioFabricHold", ".zip");
      holdIt.deleteOnExit();
    } catch (IOException ioex) {
      holdIt = null;
    }
  
    HashSet<NetLink> reducedLinks = new HashSet<NetLink>();
    TreeMap<AugRelation, Boolean> relMap = new TreeMap<AugRelation, Boolean>();
    FabricImportLoader.FileImportStats sss = new FabricImportLoader.FileImportStats();
  
    // Always do background read- not worth checking file size- most files will likely be large
    BackgroundFileReader br = new BackgroundFileReader();
    // This gets file in:
    boolean finished;
    if (type == FileLoadFlows.FileLoadType.SIF) {
      finished = br.doBackgroundSIFRead(file, idGen, links, loneNodes, null, sss, magBins, relMap, holdIt);
    } else if (type == FileLoadFlows.FileLoadType.GW) {
      // no need for gw processing because links will be post-processed anyways (e.g. net align)
      finished = br.doBackgroundGWRead(file, idGen, links, loneNodes, null, sss, false, magBins, relMap, holdIt);
    } else {
      throw (new IllegalArgumentException("File type not identified"));
    }
  
    if (!finished) {
      return (new FileLoadResultImpl(true, holdIt));
    }
  
    //
    // Let user know if there were formatting problems (but keep going...)
    //
  
    announceBadLines(sss);
  
    //
    // If caller just wants to get the file in without followup, exit now:
    //
  
    if (loadOnly) {
      return (new FileLoadResultImpl(true, holdIt));
    }
  
    //
    // This looks for dups to toss and prep work:
    //
  
    finished = handleDirectionsDupsAndShadows(links, loneNodes, (magBins != null), relMap, 
    		                                      reducedLinks, holdIt, false, skipShadowQuestion);
  
    if (finished) {
      buildTheNetworkFomLinks(file, idGen, loneNodes, reducedLinks, holdIt);
    }
    return (new FileLoadResultImpl(true, holdIt));
  }
  
  /***************************************************************************
  **
  ** Preprocess ops that are either run in forground or background:
  */ 
    
  private void preprocess(List<NetLink> links, 
  		                    SortedMap<AugRelation, Boolean> relaMap,
  	                      Set<NetLink> reducedLinks, Set<NetLink> culledLinks,  
  	                      BTProgressMonitor monitor) throws AsynchExitRequestException {
  	BuildExtractor bex = PluginSupportFactory.getBuildExtractor();
    bex.assignDirections(links, relaMap, monitor);
    bex.preprocessLinks(links, reducedLinks, culledLinks, monitor);
    return;
  }  
    
  /***************************************************************************
  **
  ** Common load operations.
  */ 
    
  public boolean loadXMLFromSource(File file, File holdIt) {  
    ArrayList<ParserClient> alist = new ArrayList<ParserClient>();
    FabricFactory ff = new FabricFactory(pMan_);
    alist.add(ff);
    SUParser sup = new SUParser(alist);   
    if (file.length() > XML_SIZE_FOR_BACKGROUND_READ) {
      BackgroundFileReader br = new BackgroundFileReader(); 
      boolean finished = br.doBackgroundRead(ff, sup, file, false, holdIt);
      if (finished) {
        setCurrentXMLFile(file);
        postXMLLoad(ff, file.getName(), holdIt);
      }
      return (true);
    } else {
      try {
        sup.parse(file);  
      } catch (IOException ioe) {
        displayFileInputError(ioe);
        return (false);              
      } catch (OutOfMemoryError oom) {
        ExceptionHandler.getHandler().displayOutOfMemory(oom);
        return (false);  
      }
    }
    setCurrentXMLFile(file);
    postXMLLoad(ff, file.getName(), holdIt);
    return (true);
  }
  
  /***************************************************************************
  **
  ** Restore a network from backup file following a cancellation.
  */ 
    
  private boolean restoreFromBackup(File file) {  
    ArrayList<ParserClient> alist = new ArrayList<ParserClient>();
    FabricFactory ff = new FabricFactory(pMan_);
    alist.add(ff);
    SUParser sup = new SUParser(alist);   
    BackgroundFileReader br = new BackgroundFileReader(); 
    br.doBackgroundRead(ff, sup, file, true, null);
    file.delete();
    // Fix for issue # 79
    String cfn = (currentFile_ == null) ? null : currentFile_.getName();
    postXMLLoad(ff, cfn, null);
    return (true);
  }
  
  /***************************************************************************
  **
  ** Common load operations.
  */ 
    
  public boolean postXMLLoad(FabricFactory ff, String fileName, File holdIt) {  
    NetworkBuilder nb = new NetworkBuilder(true, holdIt, null); 
    nb.setBuildDataForXMLLoad(ff.getFabricNetwork(), BuildDataImpl.BuildMode.BUILD_FROM_XML);
    nb.doNetworkBuild();
    manageWindowTitle(fileName);
    return (true);
  }
    
  /***************************************************************************
  **
  ** Do standard file checks and warnings
  */
 
  public boolean standardFileChecks(File target, boolean mustExist, boolean canCreate,
                                    boolean checkOverwrite, boolean mustBeDirectory, 
                                    boolean canWrite, boolean canRead) {
    ResourceManager rMan = ResourceManager.getManager();
    boolean doesExist = target.exists();
  
    if (mustExist) {
      if (!doesExist) {
        String noFileFormat = rMan.getString("fileChecks.noFileFormat");
        String noFileMsg = MessageFormat.format(noFileFormat, new Object[] {target.getName()});
        JOptionPane.showMessageDialog(topWindow_, noFileMsg,
                                      rMan.getString("fileChecks.noFileTitle"),
                                      JOptionPane.ERROR_MESSAGE);
        return (false);
      }
    }
    if (mustBeDirectory) {
      if (doesExist && !target.isDirectory()) {
        String notADirectoryFormat = rMan.getString("fileChecks.notADirectoryFormat");
        String notADirectoryMsg = MessageFormat.format(notADirectoryFormat, new Object[] {target.getName()});
        JOptionPane.showMessageDialog(topWindow_, notADirectoryMsg,
                                      rMan.getString("fileChecks.notADirectoryTitle"),
                                      JOptionPane.ERROR_MESSAGE);
        return (false);
      }
    } else { // gotta be a file
      if (doesExist && !target.isFile()) {
        String notAFileFormat = rMan.getString("fileChecks.notAFileFormat");
        String notAFileMsg = MessageFormat.format(notAFileFormat, new Object[] {target.getName()});
        JOptionPane.showMessageDialog(topWindow_, notAFileMsg,
                                      rMan.getString("fileChecks.notAFileTitle"),
                                      JOptionPane.ERROR_MESSAGE);
        return (false);
      }
    }

    if (!doesExist && canCreate) {
      if (mustBeDirectory) {
        throw new IllegalArgumentException();
      }
      boolean couldNotCreate = false;
      try {
        if (!target.createNewFile()) {
          couldNotCreate = true;
        }
      } catch (IOException ioex) {
        couldNotCreate = true;   
      }
      if (couldNotCreate) {
        String noCreateFormat = rMan.getString("fileChecks.noCreateFormat");
        String noCreateMsg = MessageFormat.format(noCreateFormat, new Object[] {target.getName()});
        JOptionPane.showMessageDialog(topWindow_, noCreateMsg,
                                      rMan.getString("fileChecks.noCreateTitle"),
                                      JOptionPane.ERROR_MESSAGE);
        return (false);
      }
    }
    
    boolean didExist = doesExist;
    doesExist = target.exists();    
    
    if (canWrite) {
      if (doesExist && !target.canWrite()) {
        String noWriteFormat = rMan.getString("fileChecks.noWriteFormat");
        String noWriteMsg = MessageFormat.format(noWriteFormat, new Object[] {target.getName()});
        JOptionPane.showMessageDialog(topWindow_, noWriteMsg,
                                      rMan.getString("fileChecks.noWriteTitle"),
                                      JOptionPane.ERROR_MESSAGE);
        return (false);
      }
    }
    if (canRead) {
      if (doesExist && !target.canRead()) {
        String noReadFormat = rMan.getString("fileChecks.noReadFormat");
        String noReadMsg = MessageFormat.format(noReadFormat, new Object[] {target.getName()});     
        JOptionPane.showMessageDialog(topWindow_, noReadMsg,
                                      rMan.getString("fileChecks.noReadTitle"),
                                      JOptionPane.ERROR_MESSAGE);
        return (false);
      }
    }
    
    if (didExist && checkOverwrite) {  // note we care about DID exist (before creation)
      String overFormat = rMan.getString("fileChecks.doOverwriteFormat");
      String overMsg = MessageFormat.format(overFormat, new Object[] {target.getName()});
      if (headlessOracle_ == null) {
        int overwrite =
          JOptionPane.showConfirmDialog(topWindow_, overMsg,
                                        rMan.getString("fileChecks.doOverwriteTitle"),
                                        JOptionPane.YES_NO_OPTION);        
        if (overwrite != JOptionPane.YES_OPTION) {
          return (false);
        }
      }
    }
    return (true);
  }
  
  /***************************************************************************
  **
  ** Get readable attribute file
  */
  
  public File getTheFile(String ext1, String ext2, String prefTag, String desc, Component useUI) { 
    File file = null;      
    String filename = FabricCommands.getPreference(prefTag);
    while (file == null) {
      JFileChooser chooser = new JFileChooser();       
      FileFilter filter;
      if (ext2 == null) {
        filter = new FileExtensionFilters.SimpleFilter(ext1, desc);
      } else {
        filter = new FileExtensionFilters.DoubleExtensionFilter(ext1, ext2, desc);
      }     
      chooser.addChoosableFileFilter(filter);
      chooser.setAcceptAllFileFilterUsed(true);
      chooser.setFileFilter(filter);
      if (filename != null) {
        File startDir = new File(filename);
        if (startDir.exists()) {
          chooser.setCurrentDirectory(startDir);  
        }
      }

      int option = chooser.showOpenDialog(useUI == (null) ? topWindow_ : useUI);
      if (option != JFileChooser.APPROVE_OPTION) {
        return (null);
      }
      file = chooser.getSelectedFile();
      if (file == null) {
        return (null);
      }
      if (!standardFileChecks(file, FILE_MUST_EXIST, FILE_CAN_CREATE_DONT_CARE, 
                                    FILE_DONT_CHECK_OVERWRITE, FILE_MUST_BE_FILE, 
                                    FILE_CAN_WRITE_DONT_CARE, FILE_CAN_READ)) {
        file = null;
        continue; 
      }
    }
    return (file);
  }
  
  /***************************************************************************
  **
  ** Get a directory
  */
  
  public File getTheDirectory(String prefTag) { 
    File file = null;      
    String filename = FabricCommands.getPreference(prefTag);
    while (file == null) {
      JFileChooser chooser = new JFileChooser();           
      chooser.setAcceptAllFileFilterUsed(true);
      chooser.setFileSelectionMode(JFileChooser.DIRECTORIES_ONLY);
      if (filename != null) {
        File startDir = new File(filename);
        if (startDir.exists()) {
          chooser.setCurrentDirectory(startDir);  
        }
      }

      int option = chooser.showOpenDialog(topWindow_);
      if (option != JFileChooser.APPROVE_OPTION) {
        return (null);
      }
      file = chooser.getSelectedFile();
      if (file == null) {
        return (null);
      }     
      
      if (!standardFileChecks(file, FILE_MUST_EXIST, FILE_CAN_CREATE_DONT_CARE, 
                                    FILE_DONT_CHECK_OVERWRITE, FILE_MUST_BE_DIRECTORY, 
                                    FILE_CAN_WRITE_DONT_CARE, FILE_CAN_READ)) {
        file = null;
        continue; 
      }
    }
    return (file);
  }

  /***************************************************************************
  **
  ** Load an annotation file
  */
     
  public Map<Boolean, AnnotationSet> loadAnnotations(File file, boolean forNodes) {
    AnnotationLoader.ReadStats stats = new AnnotationLoader.ReadStats();
    try {         
      AnnotationLoader alod = new AnnotationLoader();
      Map<Boolean, AnnotationSet> aSet = alod.readAnnotations(file, stats, forNodes, bfp_.getNetwork(), null);
      FabricCommands.setPreference("AnnotDirectory", file.getAbsoluteFile().getParent());
      return (aSet);
    } catch (IOException ioe) {
      if (stats.errStr != null) {
        ResourceManager rMan = ResourceManager.getManager();
        JOptionPane.showMessageDialog(topWindow_, rMan.getString("attribRead.IOException"),
                                      rMan.getString("attribRead.IOExceptionTitle"),
                                      JOptionPane.ERROR_MESSAGE);
        return (null);
      } else {
        displayFileInputError(ioe);
        return (null);              
      }  
    } catch (AsynchExitRequestException aerex) {
      UiUtil.fixMePrintout("Do this read on background thread");
      return (null);
    }
  }
  
  /***************************************************************************
  **
  ** Load the file. Map keys are strings or Links
  */
     
  public Map<AttributeKey, String> loadTheFile(File file, Map<String, Set<NetNode>> nameToIDs, boolean forNodes) {
    HashMap<AttributeKey, String> attributes = new HashMap<AttributeKey, String>();
    try {    
      AttributeLoader.ReadStats stats = new AttributeLoader.ReadStats();
      AttributeLoader alod = new AttributeLoader();
      BuildExtractor bex = PluginSupportFactory.getBuildExtractor();
      Map<String, NetNode> nameToID = (!forNodes) ? bex.reduceNameSetToOne(nameToIDs) : null;
      alod.readAttributes(file, forNodes, attributes, nameToID, stats);
      if (!stats.badLines.isEmpty()) {
        ResourceManager rMan = ResourceManager.getManager();
        String badLineFormat = rMan.getString("attribRead.badLineFormat");
        String badLineMsg = MessageFormat.format(badLineFormat, new Object[] {Integer.valueOf(stats.badLines.size())});
        if (headlessOracle_ == null) {
          JOptionPane.showMessageDialog(topWindow_, badLineMsg,
                                        rMan.getString("attribRead.badLineTitle"),
                                        JOptionPane.WARNING_MESSAGE);
        } else {
          headlessOracle_.displayErrorMessage(badLineMsg);
        }
      }
      if (!stats.dupLines.isEmpty()) {
        ResourceManager rMan = ResourceManager.getManager();
        String dupLineFormat = rMan.getString("attribRead.dupLineFormat");
        String dupLineMsg = MessageFormat.format(dupLineFormat, new Object[] {Integer.valueOf(stats.dupLines.size())});
        if (headlessOracle_ == null) {
          JOptionPane.showMessageDialog(topWindow_, dupLineMsg,
                                        rMan.getString("attribRead.dupLineTitle"),
                                        JOptionPane.WARNING_MESSAGE);
        } else {
          headlessOracle_.displayErrorMessage(dupLineMsg);
        }
      }
      if (!forNodes && !stats.shadowsPresent) {
        ResourceManager rMan = ResourceManager.getManager();
        if (headlessOracle_ == null) {
          JOptionPane.showMessageDialog(topWindow_, rMan.getString("attribRead.noShadowError"),
                                        rMan.getString("attribRead.noShadowTitle"),
                                        JOptionPane.ERROR_MESSAGE);
        } else {
          headlessOracle_.displayErrorMessage(rMan.getString("attribRead.noShadowError"));
        }
        return (null);
      }
     
    } catch (IOException ioe) {
      displayFileInputError(ioe);
      return (null);              
    }
    FabricCommands.setPreference("AttribDirectory", file.getAbsoluteFile().getParent());
    return (attributes);
  }
 
  /***************************************************************************
   **
   ** Common save activities
   */   
   
   public boolean saveToCurrentFile() {
     if (currentFile_ != null) {
       return (saveToFile(currentFile_.getAbsolutePath()));
     } else {
       return (saveToFile(null));
     }
   }
  
  /***************************************************************************
  **
  ** Common save activities
  */   
  
  public boolean saveToFile(String fileName) {
       
    File file = null;
    if (fileName == null) {
      String dirName = FabricCommands.getPreference("LoadDirectory");
      while (file == null) {
        JFileChooser chooser = new JFileChooser();
        FileExtensionFilters.SimpleFilter sf = new FileExtensionFilters.SimpleFilter(".bif", "filterName.bif");
        chooser.addChoosableFileFilter(sf);
        chooser.setAcceptAllFileFilterUsed(true);
        chooser.setFileFilter(sf);
        if (dirName != null) {
          File startDir = new File(dirName);
          if (startDir.exists()) {
            chooser.setCurrentDirectory(startDir);  
          }
        }  
        int option = chooser.showSaveDialog(topWindow_);
        if (option != JFileChooser.APPROVE_OPTION) {
          return (true);
        }
        file = chooser.getSelectedFile();
        if (file != null) {
          if (!file.exists()) {
            if (!FileExtensionFilters.hasSuffix(file.getName(), ".bif")) {
              file = new File(file.getAbsolutePath() + ".bif");
            }
          }
          if (!standardFileChecks(file, FILE_MUST_EXIST_DONT_CARE, FILE_CAN_CREATE, 
                                        FILE_CHECK_OVERWRITE, FILE_MUST_BE_FILE, 
                                        FILE_CAN_WRITE, FILE_CAN_READ_DONT_CARE)) {
            file = null;
            continue; 
          }
        }       
      }     
    } else {
      // given a name, we do not check overwrite:
      file = new File(fileName);
      if (!standardFileChecks(file, FILE_MUST_EXIST_DONT_CARE, FILE_CAN_CREATE, 
                                    FILE_DONT_CHECK_OVERWRITE, FILE_MUST_BE_FILE, 
                                    FILE_CAN_WRITE, FILE_CAN_READ_DONT_CARE)) {
        return (false);
      }        
    }

    
    BioFabricNetwork bfn = bfp_.getNetwork();    
    if (bfn.getLinkCount(true) > LINK_COUNT_FOR_BACKGROUND_WRITE) {
      BackgroundFileWriter bw = new BackgroundFileWriter(); 
      bw.doBackgroundWrite(file);
      return (true);
    } else {
      try {
        saveToOutputStream(new FileOutputStream(file), false, null);
        setCurrentXMLFile(file);
        manageWindowTitle(file.getName());
        return (true);
      } catch (AsynchExitRequestException aeex) {
      	// Not on background thread; will not happen
      	return (false);
      } catch (IOException ioe) {
        displayFileOutputError();
        return (false);
      }
    }  
  }
  
  /***************************************************************************
  **
  ** Save to output stream
  */   
  
  public void saveToOutputStream(OutputStream stream, boolean compress, BTProgressMonitor monitor) 
  	throws AsynchExitRequestException, IOException {

  	PrintWriter out = null;
  	if (compress) {
  		out = new PrintWriter(new OutputStreamWriter(new GZIPOutputStream(stream, 8 * 1024), "UTF-8"));
  	} else {
  	  out = new PrintWriter(new BufferedWriter(new OutputStreamWriter(stream, "UTF-8")));
  	}  	
  	try {
	    Indenter ind = new Indenter(out, Indenter.DEFAULT_INDENT);
	    BioFabricNetwork bfn = bfp_.getNetwork();
	    if (bfn != null) {
	      bfn.writeXML(out, ind, monitor, compress);
	    }
  	} finally {
  		if (out != null) {
  			out.close();
  		}  		
  	}  
    return;
  }
  
  /***************************************************************************
  **
  ** Displays file reading error message
  */ 
       
  public void displayFileInputError(IOException ioex) { 
    ResourceManager rMan = ResourceManager.getManager();
    
    if ((ioex == null) || (ioex.getMessage() == null) || (ioex.getMessage().trim().equals(""))) {
      if (headlessOracle_ == null) {
        JOptionPane.showMessageDialog(topWindow_, 
                                      rMan.getString("fileRead.errorMessage"), 
                                      rMan.getString("fileRead.errorTitle"),
                                      JOptionPane.ERROR_MESSAGE);
      } else {
        headlessOracle_.displayErrorMessage(rMan.getString("fileRead.errorMessage"));
      }
      return;
    }
    String errMsg = ioex.getMessage().trim();
    String format = rMan.getString("fileRead.inputErrorMessageForIOEx");
    String outMsg = MessageFormat.format(format, new Object[] {errMsg}); 
    if (headlessOracle_ == null) {
      JOptionPane.showMessageDialog(topWindow_, outMsg, 
                                    rMan.getString("fileRead.errorTitle"),
                                    JOptionPane.ERROR_MESSAGE);
    } else {
      headlessOracle_.displayErrorMessage(outMsg);
    }
    return;
  }
  
  /***************************************************************************
  **
  ** Displays file writing error message
  */ 
       
  public void displayFileOutputError() { 
    ResourceManager rMan = ResourceManager.getManager(); 
    JOptionPane.showMessageDialog(topWindow_, 
                                  rMan.getString("fileWrite.errorMessage"), 
                                  rMan.getString("fileWrite.errorTitle"),
                                  JOptionPane.ERROR_MESSAGE);
    return;
  }
 
  /***************************************************************************
  **
  ** Displays file reading error message for invalid input
  */ 
       
  public void displayInvalidInputError(InvalidInputException iiex) { 
    ResourceManager rMan = ResourceManager.getManager(); 
    String errKey = iiex.getErrorKey();
    boolean haveKey = (errKey != null) && (!errKey.equals(InvalidInputException.UNSPECIFIED_ERROR)); 
    int lineno = iiex.getErrorLineNumber();
    boolean haveLine = (lineno != InvalidInputException.UNSPECIFIED_LINE);
    String outMsg;
    if (haveKey && haveLine) { 
      String format = rMan.getString("fileRead.inputErrorMessageForLineWithDesc");
      String keyedErr = rMan.getString("invalidInput." + errKey);
      outMsg = MessageFormat.format(format, new Object[] {Integer.valueOf(lineno + 1), keyedErr});
    } else if (haveKey && !haveLine) {
      String format = rMan.getString("fileRead.inputErrorMessageWithDesc");
      String keyedErr = rMan.getString("invalidInput." + errKey);
      outMsg = MessageFormat.format(format, new Object[] {keyedErr});
    } else if (!haveKey && haveLine) {
      String format = rMan.getString("fileRead.inputErrorMessageForLine");
      outMsg = MessageFormat.format(format, new Object[] {Integer.valueOf(lineno + 1)});      
    } else {
      outMsg = rMan.getString("fileRead.inputErrorMessage");      
    } 
    JOptionPane.showMessageDialog(topWindow_, outMsg, 
                                  rMan.getString("fileRead.errorTitle"),
                                  JOptionPane.ERROR_MESSAGE);
  return;
  }
 


  ////////////////////////////////////////////////////////////////////////////
  //
  // INNER CLASSES
  //
  ///////////////////////////////////////////////////////////////////////////
  
  /***************************************************************************
  **
  ** Class for loading huge files in
  **
  ** Sequence: 1) Fires off either new SIFReaderRunner or new ReaderRunner.
  **           2) When they finish, via finishedLoad(), this either calls:
  **               a) finishLoadFromSIFSource or 
  *                b) postXMLLoad
  *                
  *            In finishLoadFromSIFSource, we present a dialog, then:
  *            
  *              PreprocessNetwork pn = new PreprocessNetwork();
                 boolean didFinish = pn.doNetworkPreprocess(links, relaMap, reducedLinks, culledLinks);
                 
                 then another optional dialog, 
                 
  *              then we call
  *               NetworkBuilder nb = new NetworkBuilder(true);
                   nb.setForSifBuild(idGen, reducedLinks, loneNodeIDs, BioFabricNetwork.BuildMode.BUILD_FROM_SIF);
                   nb.doNetworkBuild(); 
                   
                   do network build uses NewNetworkRunner(isMain);
                   which runs expensiveModelOperations()
                  *** currently no file is cached.
                   
  ** 
  */ 
    
  public class BackgroundFileReader implements BackgroundWorkerOwner {
    
  	private File holdIt_;
    private Exception ex_;  
    private boolean finished_;
    private boolean forRecovery_;
     
    public boolean doBackgroundSIFRead(File file, UniqueLabeller idGen,
		    		                           List<NetLink> links, Set<NetNode> loneNodeIDs, 
		    		                           Map<String, String> nameMap, FabricImportLoader.FileImportStats sss,
		    		                           Integer magBins, SortedMap<AugRelation, Boolean> relMap,
		    		                           File holdIt) {

    	holdIt_ = holdIt;
      finished_ = true;
      forRecovery_ = false;
      try {
        BFWorker bfw = PluginSupportFactory.getBFWorker(this, topWindow_, bfw_, "fileLoad.waitTitle", "fileLoad.wait", true, null);
        SIFReaderRunner runner = new SIFReaderRunner(file, idGen, links, loneNodeIDs, nameMap, sss, magBins, relMap, holdIt_, bfw);                                                        
        bfw.setCore(runner);
        bfw.launchWorker();       
      } catch (Exception ex) {
        ExceptionHandler.getHandler().displayException(ex);
      }
      return (finished_);
    }
    
    public boolean doBackgroundGWRead(File file, UniqueLabeller idGen,
                                      List<NetLink> links, Set<NetNode> loneNodeIDs,
                                      Map<String, String> nameMap, FabricImportLoader.FileImportStats gws,
                                      boolean needsGWProcess, Integer magBins,
                                      SortedMap<AugRelation, Boolean> relMap, File holdIt) {
      holdIt_ = holdIt;
      finished_ = true;
      forRecovery_ = false;
      try {
        BFWorker bfw = PluginSupportFactory.getBFWorker(this, topWindow_, bfw_, "fileLoad.waitTitle", "fileLoad.wait", true, null);
        GWReaderRunner runner = new GWReaderRunner(file, idGen, links, loneNodeIDs, nameMap, gws, magBins, relMap, needsGWProcess, holdIt_, bfw);
        bfw.setCore(runner);
        bfw.launchWorker();       
      } catch (Exception ex) {
        ExceptionHandler.getHandler().displayException(ex);
      }
      return (finished_);
    }
  
    public boolean doBackgroundRead(FabricFactory ff, SUParser sup, File file, boolean compressed, File holdIt) {
    	holdIt_ = holdIt;
      finished_ = true;
      forRecovery_ = (holdIt == null);
      try {
      	BFWorker bfw = PluginSupportFactory.getBFWorker(this, topWindow_, bfw_, "fileLoad.waitTitle", "fileLoad.wait", true, null);
        ReaderRunner runner = new ReaderRunner(sup, file, compressed, holdIt_, bfw);                                                      
        bfw.setCore(runner);
        bfw.launchWorker(); 
      } catch (Exception ex) {
        ExceptionHandler.getHandler().displayException(ex);
      }
      return (finished_);
    }

    public boolean handleRemoteException(Exception remoteEx) {
    	finished_ = false;
      if (remoteEx instanceof IOException) {
        ex_ = remoteEx;
        return (true);
      }
      return (false);
    }    
        
    public void cleanUpPreEnable(Object result) {
      return;
    }
    
    public boolean handleCancellation() {
    	boolean restored = false;
    	if (!forRecovery_) {
    	  restored = cancelAndRestore(holdIt_);
    	}
    	finished_ = false;
    	return (restored);
    }     
    
    public void cleanUpPostRepaint(Object result, boolean skipImage) { 
      finishedLoad();
      return;
    }
     
    private void finishedLoad() {     
      if (ex_ != null) {
        displayFileInputError((IOException)ex_);               
      }
      return;
    }
  }
  
  /***************************************************************************
  **
  ** Background file load
  */ 
    
  private class ReaderRunner implements BackgroundCore {
   
    private File myFile_;
    private SUParser myParser_;
    private boolean compressed_;
    private File holdIt_;
    private BFWorker bfwk_;
    
    public ReaderRunner(SUParser sup, File file, boolean compressed, File holdIt, BFWorker bfwk) {

    	bfwk_ = bfwk;
      myFile_ = file;
      myParser_ = sup;
      compressed_ = compressed;
      holdIt_ = holdIt;
    }
    
     public Object getEarlyResult() {
    	 return (new Boolean(false));
     }
 
    public Object runCore() throws AsynchExitRequestException {
    	BTProgressMonitor monitor = bfwk_.getMonitor();
      if ((holdIt_ != null) && (holdIt_.length() == 0)) {
        buildRestoreCache(holdIt_, monitor);
      }
      ProgressFilterInputStream pfis = null;
      try {
        long fileLen = myFile_.length();
        FileInputStream fis = new FileInputStream(myFile_);
        InputStream bis;
        if (compressed_) {
          bis = new GZIPInputStream(fis, 8 * 1024);
        } else {
          bis = new BufferedInputStream(new FileInputStream(myFile_));
        } 
        pfis = new ProgressFilterInputStream(bis, fileLen);   
        myParser_.parse(pfis, monitor, compressed_);
        return (new Boolean(true));
      } catch (IOException ioe) {
        bfwk_.stashException(ioe);
        return (null);
      } finally {
        if (pfis != null) { try { pfis.close(); } catch (IOException ioe) {} }
      }
    } 
    
    public Object postRunCore() {
      return (null);
    } 
  }  
 
  /***************************************************************************
  **
  ** Once directionality of link relations is established, we need to assign directions
  ** and to remove non-directional synonymous and duplicate links. This is run in the
  ** background, but must be preceded by the user providing directed relation info, and
  ** followed by (maybe) telling the user what is dropped.
  */ 
    
  public class PreprocessNetwork implements BackgroundWorkerOwner {
    
    private boolean finished_;
    private File holdIt_;
    
    public boolean doNetworkPreprocess(List<NetLink> links, 
                                       SortedMap<AugRelation, Boolean> relaMap,
                                       Set<NetLink> reducedLinks, Set<NetLink> culledLinks, File holdIt) {
      holdIt_ = holdIt;
      finished_ = true;
      try {
        BFWorker bfw = PluginSupportFactory.getBFWorker(this, topWindow_, bfw_, "netPreprocess.waitTitle", "netPreprocess.wait", true, null);
        PreprocessRunner runner = new PreprocessRunner(links, relaMap, reducedLinks, culledLinks, holdIt_, bfw);                                                        
        bfw.setCore(runner);
        bfw.launchWorker();  
      } catch (Exception ex) {
        ExceptionHandler.getHandler().displayException(ex);
      }
      return (finished_);
    }

    public boolean handleRemoteException(Exception remoteEx) {
      finished_ = false;
      return (false);
    }    
        
    public void cleanUpPreEnable(Object result) {
      return;
    }
    
    public boolean handleCancellation() {
      finished_ = false;
      return (cancelAndRestore(holdIt_));
    }     
    
    public void cleanUpPostRepaint(Object result, boolean skipImage) {   
      return;
    }
  }  
  
  
  /***************************************************************************
  **
  ** Once directionality of link relations is established, we need to assign directions
  ** and to remove non-directional synonymous and duplicate links. This is run in the
  ** background, but must be preceded by the user providing directed relation info, and
  ** followed by (maybe) telling the user what is dropped.
  */ 
    
  private class PreprocessRunner implements BackgroundCore {
   
    private List<NetLink> links_; 
    private SortedMap<AugRelation, Boolean> relaMap_;
    private Set<NetLink> reducedLinks_; 
    private Set<NetLink> culledLinks_;
    private File holdIt_;
    private BFWorker bfwk_; 
    
    PreprocessRunner(List<NetLink> links, SortedMap<AugRelation, Boolean> relaMap,
                     Set<NetLink> reducedLinks, Set<NetLink> culledLinks, File holdIt, BFWorker bfwk) {
     
      
      bfwk_ = bfwk;
      links_ = links;
      relaMap_ = relaMap;
      reducedLinks_ = reducedLinks;
      culledLinks_ = culledLinks;
      holdIt_ = holdIt;
    }
    
    public Object getEarlyResult() {
    	return (new Boolean(false));
    }
    
    public Object runCore() throws AsynchExitRequestException {
    	BTProgressMonitor monitor = bfwk_.getMonitor();
      if (holdIt_.length() == 0) {
        buildRestoreCache(holdIt_, monitor);
      }
      preprocess(links_, relaMap_, reducedLinks_, culledLinks_, monitor);
      return (new Boolean(true));  
    }
    
    public Object postRunCore() {
      return (null);
    } 
  }
  
  /***************************************************************************
  **
  ** Class for writing huge files out on a background thread.
  */ 
    
  public class BackgroundFileWriter implements BackgroundWorkerOwner {
    
    private Exception ex_;   
    private File file_;

    public void doBackgroundWrite(File file) {
    	file_ = file;
      try { 
      	BFWorker bfw = PluginSupportFactory.getBFWorker(this, topWindow_, bfw_, "fileWrite.waitTitle", "fileWrite.wait", true, null);
    	  WriterRunner runner = new WriterRunner(file, bfw);
        bfw.setCore(runner);
        bfw.launchWorker();       
      } catch (Exception ex) {
        ExceptionHandler.getHandler().displayException(ex);
      }
    	return;
    }
    
    public void doBackgroundWrite(OutputStream stream) {
    	file_ = null;
    	BFWorker bfw = PluginSupportFactory.getBFWorker(this, topWindow_, bfw_, "fileWrite.waitTitle", "fileWrite.wait", true, null);
      WriterRunner runner = new WriterRunner(stream, bfw);
      try {                                                                
        bfw.setCore(runner);
        bfw.launchWorker();       
      } catch (Exception ex) {
        ExceptionHandler.getHandler().displayException(ex);
      }
      return;
    }

    public boolean handleRemoteException(Exception remoteEx) {
      if (remoteEx instanceof IOException) {
        ex_ = remoteEx;
        return (true);
      }
      return (false);
    }    
        
    public void cleanUpPreEnable(Object result) {
      return;
    }
    
    public boolean handleCancellation() {
    	// Maybe a future enhancement to allow user to choose if they wish the cancelled file wrote to be deleted?
    	if (file_ != null) {
    	  file_.delete();
    	}
    	return (false);
    }     
    
    public void cleanUpPostRepaint(Object result, boolean skipImage) { 
      finishedOut();
      return;
    }
     
    private void finishedOut() {     
      if (ex_ != null) {
        displayFileOutputError();
        return;                
      }     
      setCurrentXMLFile(file_);
      manageWindowTitle(file_.getName());
      return;
    }
  }
  
  /***************************************************************************
   **
   ** This reads in GW files on the background thread, but does not build a network. That
   ** occurs in subsequent steps.
   */
  
  private class GWReaderRunner implements BackgroundCore {
  
    private File myFile_;
    private List<NetLink> links_;
    private Set<NetNode> loneNodeIDs_;
    private UniqueLabeller idGen_;
    private Map<String, String> nameMap_;
    private FabricImportLoader.FileImportStats sss_;
    private Integer magBins_;
    private SortedMap<AugRelation, Boolean> relaMap_;
    private boolean needsGWProcessing_;
    private File restoreCacheFile_;
    private BFWorker bfwk_;
    
    public GWReaderRunner(File file, UniqueLabeller idGen, List<NetLink> links,
                          Set<NetNode> loneNodeIDs, Map<String, String> nameMap,
                          FabricImportLoader.FileImportStats gws,
                          Integer magBins, SortedMap<AugRelation, Boolean> relaMap,
                          boolean needsGWProcessing, File restoreCacheFile, BFWorker bfwk) {
     
      bfwk_ = bfwk;
      myFile_ = file;
      links_ = links;
      loneNodeIDs_ = loneNodeIDs;
      idGen_ = idGen;
      nameMap_ = nameMap;
      sss_ = gws;
      magBins_ = magBins;
      relaMap_ = relaMap;
      needsGWProcessing_ = needsGWProcessing;
      restoreCacheFile_ = restoreCacheFile;
    }
  
    public Object getEarlyResult() {
    	return (new Boolean(false));
    }
    
    
    public Object runCore() throws AsynchExitRequestException {
    	BTProgressMonitor monitor = bfwk_.getMonitor();
      try {
        if (restoreCacheFile_.length() == 0) {
          buildRestoreCache(restoreCacheFile_, monitor);
        }
        preLoadOperations();
        FabricImportLoader.FileImportStats sss = 
        		(new GWImportLoader()).importFabric(myFile_, idGen_, links_, loneNodeIDs_, nameMap_, magBins_, monitor);
        sss_.copyInto(sss);
        
        if (needsGWProcessing_) {
          boolean finshed = (new GWImportLoader.GWRelationManager()).process(links_, topWindow_, monitor);
          if (!finshed) {
            // should not happen
            return (null);
          }
        }
        PluginSupportFactory.getBuildExtractor().extractRelations(links_, relaMap_, monitor);
        return (new Boolean(true));
      } catch (IOException ioe) {
        bfwk_.stashException(ioe);
        return (null);
      }
    }
  
    public Object postRunCore() {
      return null;
    }
  }

  /***************************************************************************
  **
  ** This reads in SIF files on the background thread, but does not build a network. That
  ** occurs in subsequent steps.
  */ 
    
  private class SIFReaderRunner implements BackgroundCore {
   
    private File myFile_;
    private List<NetLink> links_;
    private Set<NetNode> loneNodeIDs_;
    private UniqueLabeller idGen_; 
    private Map<String, String> nameMap_;
    private FabricImportLoader.FileImportStats sss_;
    private Integer magBins_;
    private SortedMap<AugRelation, Boolean> relaMap_;
    private File restoreCacheFile_;
    private BFWorker bfwk_;
    
    public SIFReaderRunner(File file, UniqueLabeller idGen, List<NetLink> links, 
    		                   Set<NetNode> loneNodeIDs, Map<String, String> nameMap, 
    		                   FabricImportLoader.FileImportStats sss,
    		                   Integer magBins, SortedMap<AugRelation, Boolean> relaMap,
    		                   File restoreCacheFile, BFWorker bfwk) {
      
    	bfwk_ = bfwk;
      myFile_ = file;
      links_ = links;
      loneNodeIDs_ = loneNodeIDs;
      idGen_ = idGen;
      nameMap_ = nameMap;
      sss_ = sss;
      magBins_ = magBins;
      relaMap_ = relaMap;
      restoreCacheFile_ = restoreCacheFile;
    }
    
    public Object getEarlyResult() {
    	return (new Boolean(false));
    }
    
    public Object runCore() throws AsynchExitRequestException {
      try {
      	BTProgressMonitor monitor = bfwk_.getMonitor();
      	if (restoreCacheFile_.length() == 0) {
          buildRestoreCache(restoreCacheFile_, monitor);
      	}
        preLoadOperations();
        FabricImportLoader.FileImportStats sss = 
          (new SIFImportLoader()).importFabric(myFile_, idGen_, links_, loneNodeIDs_, nameMap_, magBins_, monitor);
        sss_.copyInto(sss);
        PluginSupportFactory.getBuildExtractor().extractRelations(links_, relaMap_, monitor);     
        return (new Boolean(true));
      } catch (IOException ioe) {
        bfwk_.stashException(ioe);
        return (null);
      }
    } 
    public Object postRunCore() {
      return (null);
    } 
  } 
    
  /***************************************************************************
  **
  ** Background file write
  */ 
    
  private class WriterRunner implements BackgroundCore {
   
    private File myFile_;
    private OutputStream myStream_;
    private BFWorker bfwk_;
    
    public WriterRunner(File file, BFWorker bfwk) {
      myFile_ = file;
      myStream_ = null;
      bfwk_ = bfwk;
    }
    public WriterRunner(OutputStream stream, BFWorker bfwk) {
      myStream_ = stream;
      myFile_ = null;
      bfwk_ = bfwk;
    }
     
    public Object getEarlyResult() {
    	return (new Boolean(false));
    }

    public Object runCore() throws AsynchExitRequestException {
      try {
        saveToOutputStream((myStream_ == null) ? new FileOutputStream(myFile_) : myStream_, false, bfwk_.getMonitor());
        return (new Boolean(true));
      } catch (IOException ioe) {
        bfwk_.stashException(ioe);
        return (null);
      }
    } 
    public Object postRunCore() {
      return (null);
    } 
  }
  
  /***************************************************************************
  **
  ** Routine for handling cancellation/restore operation
  */
  
  public boolean cancelAndRestore(File restoreFile) {
  	if ((restoreFile != null) && restoreFile.exists() && (restoreFile.length() > 20)) { // empty ZIP has 20 bytes
	    ResourceManager rMan = ResourceManager.getManager();
	    int restore =
	      JOptionPane.showConfirmDialog(topWindow_, rMan.getString("progress.cancelled"),
	                                    rMan.getString("progress.cancelledTitle"),
	                                    JOptionPane.YES_NO_OPTION);        
	    if (restore == JOptionPane.YES_OPTION) {
	    	restoreFromBackup(restoreFile);
	    	return (true);
	    } else {
	      restoreFile.delete();
	      manageWindowTitle(null);
        buildEmptyNetwork();
	    }
  	}
    return (false);
  }
 
  /***************************************************************************
  **
  ** Routine for handling cancellation/restore operation
  */
  
  public void buildRestoreCache(File restoreFile, BTProgressMonitor btpm) throws AsynchExitRequestException {
  	boolean throwOut = false;
    try {
  	  saveToOutputStream(new FileOutputStream(restoreFile), true, btpm);
  	} catch (IOException ioex) {
  		System.err.println("bad write");
  		throwOut = true;
  	} catch (AsynchExitRequestException aex) {  
	    throwOut = true;
	    throw aex;
  	} finally {
  		if (throwOut) {
  	    restoreFile.delete();		
  		}
  	}
  	return;
  }

  
  /***************************************************************************
  **
  ** Class for building networks
  */ 
    
  public class NetworkBuilder implements BackgroundWorkerOwner {
    
    private NewNetworkRunner runner_;
    private boolean finished_;
    private File holdIt_;  // For recovery
    private BFWorker bfwk_;
    
    NetworkBuilder(boolean isMain, File holdIt, PluginResourceManager rMan) {
    	bfwk_ = PluginSupportFactory.getBFWorker(this, topWindow_, bfw_, "netBuild.waitTitle", "netBuild.wait", true, rMan); 
      runner_ = new NewNetworkRunner(isMain, holdIt, bfwk_);
      holdIt_ = holdIt;
    }
    
    void setForSifBuild(UniqueLabeller idGen, Set<NetLink> links, 
                        Set<NetNode> loneNodeIDs, BuildDataImpl.BuildMode bMode) {
      if (bMode != BuildDataImpl.BuildMode.BUILD_FROM_SIF) {
        throw new IllegalArgumentException();
      }
      runner_.setBuildDataForSIF(idGen, links, loneNodeIDs, bMode);
      return;
    }
    
    void setForPlugInBuild(BuildData bData) {
      runner_.setBuildDataForPlugin(bData);
      return;
    }
    
    void setForDisplayOptionChange(BioFabricNetwork bfn, BuildDataImpl.BuildMode bMode) {
      if (bMode != BuildDataImpl.BuildMode.SHADOW_LINK_CHANGE) {
        throw new IllegalArgumentException();
      }
      runner_.setBuildDataForOptionChange(bfn, bMode);
      return;
    }

    void setBuildDataForXMLLoad(BioFabricNetwork bfn, BuildDataImpl.BuildMode bMode) {
      if (bMode != BuildDataImpl.BuildMode.BUILD_FROM_XML) {
        throw new IllegalArgumentException();
      }
      runner_.setBuildDataForXMLLoad(bfn, bMode); 
    }
 
    public void doNetworkBuildForeground() {
      try {
        runner_.runCore();
      } catch (AsynchExitRequestException axex) {
        // will not happen
      }
    }

    public boolean doNetworkBuild() {
      finished_ = true;
      try {                                                
        bfwk_.setCore(runner_);
        bfwk_.launchWorker();      
      } catch (Exception ex) {
        ExceptionHandler.getHandler().displayException(ex);
      }
      return (finished_);
    }

    public boolean handleRemoteException(Exception remoteEx) {
      finished_ = false;
      if (remoteEx instanceof IOException) {
        finishedImport(null, (IOException)remoteEx, false);
        return (true);
      } else if (remoteEx instanceof LayoutCriterionFailureException) {
        ResourceManager rMan = ResourceManager.getManager();
        JOptionPane.showMessageDialog(topWindow_, 
                                      rMan.getString("netLayout.unmetCriteriaMessage"), 
                                      rMan.getString("netLayout.unmetCriteriaTitle"),
                                      JOptionPane.ERROR_MESSAGE);
        
        
        cancelAndRestore(holdIt_);     
        return (true);
      }
      return (false);
    }    
        
    public void cleanUpPreEnable(Object result) {
      return;
    }
    
    public boolean handleCancellation() {
      finished_ = false;
      return (cancelAndRestore(holdIt_));
    }     
    
    public void cleanUpPostRepaint(Object result, boolean skipImage) {   
      finishedImport(result, null, skipImage);
      return;
    }

    private void finishedImport(Object result, IOException ioEx, boolean skipImage) {     
      if (ioEx != null) {
        displayFileInputError(ioEx);
        return;                
      }
      postLoadOperations((BufferedImage)result, skipImage);
      return;
    }
  }
  
  /***************************************************************************
  **
  ** Build New Network
  */ 
    
  private class NewNetworkRunner implements BackgroundCore {
 
    private boolean forMain_;
    private UniqueLabeller idGen_;
    private Set<NetLink> links_; 
    private Set<NetNode> loneNodeIDs_;
    private BuildDataImpl.BuildMode bMode_;
    private BioFabricNetwork bfn_;
    private File holdIt_;
    private long linkCount_;
    private BuildDataImpl plugInBuildData_;
    private BFWorker bfwk_;
    

    public NewNetworkRunner(boolean forMain, File holdIt, BFWorker bfwk) { 
      forMain_ = forMain;
      holdIt_ = holdIt;
      bfwk_ = bfwk;
    }
     
    public Object getEarlyResult(){
    	return (new BufferedImage(1, 1, BufferedImage.TYPE_INT_RGB));      
    }
    
    void setBuildDataForSIF(UniqueLabeller idGen, Set<NetLink> links, Set<NetNode> loneNodeIDs,
                            BuildDataImpl.BuildMode bMode) {    
      idGen_ = idGen;
      links_ = links; 
      loneNodeIDs_ = loneNodeIDs;
      bMode_ = bMode;
      linkCount_ = links.size();
      return;
    }
    
    void setBuildDataForPlugin(BuildData bd) {
      plugInBuildData_ = (BuildDataImpl)bd;
      bMode_ = plugInBuildData_.getMode();
      return;
    }
    
    void setBuildDataForOptionChange(BioFabricNetwork bfn, BuildDataImpl.BuildMode bMode) {
      bfn_ = bfn;
      linkCount_ = bfn.getLinkCount(true);
      bMode_ = bMode;
      return;
    }
    
    void setBuildDataForXMLLoad(BioFabricNetwork bfn, BuildDataImpl.BuildMode bMode) {
      bfn_ = bfn;
      linkCount_ = bfn.getLinkCount(true);
      bMode_ = bMode;
      return;
    }
    
    private BuildData generateBuildData() { 
      switch (bMode_) {
        case BUILD_FROM_SIF:
          HashMap<NetNode, String> emptyMap = new HashMap<NetNode, String>();
          return (new BuildDataImpl(idGen_, links_, loneNodeIDs_, emptyMap, colGen_, bMode_));
        case BUILD_FROM_PLUGIN:
          plugInBuildData_.setColorGen(colGen_);
          return (plugInBuildData_);
        case SHADOW_LINK_CHANGE:
        case BUILD_FROM_XML:
          return (new BuildDataImpl(bfn_, bMode_));   
        default:
          throw new IllegalStateException();    
      }
    }

    public Object runCore() throws AsynchExitRequestException {
      try {
      	BTProgressMonitor monitor = bfwk_.getMonitor();
        if ((holdIt_ != null) && (holdIt_.length() == 0)) {
          buildRestoreCache(holdIt_, monitor);
        }   
        BuildData bd = generateBuildData();
        preLoadOperations();
        // This can be run on foreground thread (for headless operation): shut up progress monitor
        // if that is the case:
        if (headlessOracle_ != null) {
        	monitor = null;
        }
        
        BufferedImage bi = expensiveModelOperations(bd, forMain_, monitor);
        if (linkCount_ > 10000) {
          (new GarbageRequester()).askForGC(monitor);
        }
        return (bi);
      } catch (IOException ex) {
        bfwk_.stashException(ex);
        return (null);
      } catch (LayoutCriterionFailureException ex) {
        bfwk_.stashException(ex);
        return (null);
      }
    }
    
    public Object postRunCore() {
      return (null);
    } 
  }  

  /***************************************************************************
  **
  ** Class for relayout of networks
  */ 
    
  public class NetworkRelayout implements BackgroundWorkerOwner {
     
    private File holdIt_;
    NetworkRelayoutRunner runner_;
    private BFWorker bfwk_;
      
    public NetworkRelayout() {
    	bfwk_ = PluginSupportFactory.getBFWorker(this, topWindow_, bfw_, "netRelayout.waitTitle", "netRelayout.wait", true, null);
      runner_ = new NetworkRelayoutRunner(bfwk_); 
      bfwk_.setCore(runner_);
    }
    
    public void setGroupOrderAndMode(List<String> groupOrder, BioFabricNetwork.LayoutMode mode, 
                                     boolean showGroupLinkAnnotations) {
      runner_.setGroupOrderAndMode(groupOrder, mode, showGroupLinkAnnotations);
      return;
    }

    public void setNodeOrderFromAttrib(Map<AttributeKey, String> nodeAttributes) {
      runner_.setNodeOrderFromAttrib(nodeAttributes);
      return;
    }  
    
    public void setParams(NodeLayout.Params params) {
      runner_.setParams(params);
      return;
    }  
    
    public void setPointUp(boolean pointUp) {
      runner_.setPointUp(pointUp);
      return;
    }
    
    public void setLinkMeaning(SetLayout.LinkMeans linkMeaning) {
      runner_.setLinkMeaning(linkMeaning);
      return;
    }
     
    public void setControlTopModes(ControlTopLayout.CtrlMode cMode,  ControlTopLayout.TargMode tMode, List<String> fixedList) {
      runner_.setControlTopModes(cMode, tMode, fixedList);
      return;      
    }

    public void setLinkOrder(SortedMap<Integer, NetLink> linkOrder) {
      runner_.setLinkOrder( linkOrder);
      return;
    }
    
    public void doNetworkRelayout(BioFabricNetwork bfn, BuildDataImpl.BuildMode bMode) {
      try {
        holdIt_ = File.createTempFile("BioFabricHold", ".zip");
        holdIt_.deleteOnExit();
        runner_.setNetworkAndMode(holdIt_, bfn, bMode); 
        if (bMode == BuildDataImpl.BuildMode.REORDER_LAYOUT) {
          bfwk_.makeSuperChart();
        }
        bfwk_.launchWorker();         
      } catch (Exception ex) {
        ExceptionHandler.getHandler().displayException(ex);
      }
      return;
    }

    public boolean handleRemoteException(Exception remoteEx) {
      if (remoteEx instanceof IOException) {
        displayFileInputError((IOException)remoteEx);
        return (true);
      }
      if (remoteEx instanceof LayoutCriterionFailureException) {
        ResourceManager rMan = ResourceManager.getManager();
        JOptionPane.showMessageDialog(topWindow_, 
                                      rMan.getString("netLayout.unmetCriteriaMessage"), 
                                      rMan.getString("netLayout.unmetCriteriaTitle"),
                                      JOptionPane.ERROR_MESSAGE);   
        return (true);
      }
      return (false);
    }    
        
    public void cleanUpPreEnable(Object result) {
      return;
    }
    
    //
    // Cancellation takes place on the UI Thread:
    //
    public boolean handleCancellation() {
      return (cancelAndRestore(holdIt_));
    }     
    
    public void cleanUpPostRepaint(Object result, boolean skipImage) {   
      finishedImport(result, null, skipImage);
      return;
    }

    private void finishedImport(Object result, IOException ioEx, boolean skipImage) {     
      postLoadOperations((BufferedImage)result, skipImage);
      return;
    }
  }
  
  /***************************************************************************
  **
  ** Background network layout
  */ 
    
  private class NetworkRelayoutRunner implements BackgroundCore {
 
    private BuildDataImpl rbd_;
    private BuildDataImpl.BuildMode mode_;
    private NodeLayout.Params params_;
    private BioFabricNetwork bfn_;
    private Map<AttributeKey, String> nodeAttrib_;
    private File holdIt_;
    private List<String> groupOrder_; 
    private BioFabricNetwork.LayoutMode layMode_;
    private Boolean pointUp_;
    private SetLayout.LinkMeans linkMeaning_;
    private SortedMap<Integer, NetLink> linkOrder_;
    private ControlTopLayout.CtrlMode cMode_; 
    private ControlTopLayout.TargMode tMode_; 
    private List<String> fixedList_;
    private boolean showLinkGroupAnnotations_;
    private BFWorker bfwk_;
  
    NetworkRelayoutRunner(BFWorker bfwk) {
      bfwk_ = bfwk;
    }
    
    public Object getEarlyResult() {
    	return (new BufferedImage(1, 1, BufferedImage.TYPE_INT_RGB)); 
    }

    void setNetworkAndMode(File holdIt, BioFabricNetwork bfn, BuildDataImpl.BuildMode bMode) {
      holdIt_ = holdIt;
      bfn_ = bfn;
      mode_ = bMode;
      return;
    }

    void setGroupOrderAndMode(List<String> groupOrder, 
                              BioFabricNetwork.LayoutMode mode, 
                              boolean showGroupLinkAnnotations) {
      groupOrder_ = groupOrder;
      layMode_ = mode;
      showLinkGroupAnnotations_ = showGroupLinkAnnotations;
      return;
    }

    void setNodeOrderFromAttrib(Map<AttributeKey, String> nodeAttributes) {
      nodeAttrib_ = nodeAttributes;
      return;     
    }  

    void setParams(NodeLayout.Params params) {
      params_ = params;
      return;
    }
    
    void setPointUp(boolean pointUp) {
      pointUp_ = Boolean.valueOf(pointUp);
      return;
    }
    
    void setLinkMeaning(SetLayout.LinkMeans linkMeaning) {
      linkMeaning_ = linkMeaning;
      return;
    }
        
    void setControlTopModes(ControlTopLayout.CtrlMode cMode,  ControlTopLayout.TargMode tMode, List<String> fixedList) {
      cMode_ = cMode;
      tMode_ = tMode;
      fixedList_ = fixedList;
      return;      
    }

    void setLinkOrder(SortedMap<Integer, NetLink> linkOrder) {
      linkOrder_ = linkOrder;
      return;
    }
 
    public Object runCore() throws AsynchExitRequestException {
    	BTProgressMonitor monitor = bfwk_.getMonitor(); 
      if ((holdIt_ != null) && (holdIt_.length() == 0)) {
        buildRestoreCache(holdIt_, monitor);
      }
      rbd_ = new BuildDataImpl(bfn_, mode_, monitor);
      // Either link group info is coming in from a dialog, or we extract it from the network:
      if ((groupOrder_ != null) && (layMode_ != null)) {
        rbd_.setGroupOrderAndMode(groupOrder_, layMode_, showLinkGroupAnnotations_);
      } else {
        rbd_.setGroupOrderAndMode(bfn_.getLinkGrouping(), bfn_.getLayoutMode(), bfn_.getShowLinkGroupAnnotations());
      }
      
      if (nodeAttrib_ != null) {
        rbd_.setNodeOrderFromAttrib(nodeAttrib_);   
      } else if (linkOrder_ != null) {
        rbd_.setLinkOrder(linkOrder_);
      }
      rbd_.setCTL(cMode_, tMode_, fixedList_, bfn_);
      
      rbd_.setPointUp(pointUp_);
      rbd_.setLinkMeaning(linkMeaning_);
      
      bfn_ = null; // Let go so we get GC!
      preLoadOperations();
      
      try {
       if (rbd_.needsLayoutForRelayout()) {
          NodeLayout nl = rbd_.getNodeLayout();
          boolean nlok = nl.criteriaMet(rbd_, monitor);
          if (!nlok) {
            throw new IllegalStateException(); // Should not happen, failure throws exception
          }
          nl.doNodeLayout(rbd_, params_, monitor);
          // Some "Node" layouts do the whole ball of wax, don't need this step:
          EdgeLayout el = rbd_.getEdgeLayout();
          if (el != null) {
            el.layoutEdges(rbd_, monitor);
          }
        }
        BufferedImage bi = expensiveModelOperations(rbd_, true, monitor);
        (new GarbageRequester()).askForGC(monitor);
        return (bi);
      } catch (IOException ex) {
        bfwk_.stashException(ex);
        return (null);
      } catch (LayoutCriterionFailureException ex) {
        bfwk_.stashException(ex);
        return (null);
      }
    }
    
    public Object postRunCore() {
      return (null);
    } 
  }  
  
  /***************************************************************************
  **
  ** Class for recoloring networks
  */ 
    
  public class NetworkRecolor implements BackgroundWorkerOwner {
    
    private File holdIt_;
    
    public void doNetworkRecolor(boolean isMain, File holdIt) {
      try {
        holdIt_ = holdIt;
        bfp_.shutdown();
        BFWorker bfw = PluginSupportFactory.getBFWorker(this, topWindow_, bfw_, "netRecolor.waitTitle", "netRecolor.wait", true, null);
        RecolorNetworkRunner runner = new RecolorNetworkRunner(isMain, holdIt_, bfw);                                                              
        bfw.setCore(runner);
        bfw.launchWorker();      
      } catch (Exception ex) {
        ExceptionHandler.getHandler().displayException(ex);
      }
      return;
    }

    public boolean handleRemoteException(Exception remoteEx) {
      if (remoteEx instanceof IOException) {
        finishedRecolor(null, (IOException)remoteEx);
        return (true);
      }
      return (false);
    }    
        
    public void cleanUpPreEnable(Object result) {
      return;
    }
    
    public boolean handleCancellation() {
      return (cancelAndRestore(holdIt_));
    }     
    
    public void cleanUpPostRepaint(Object result, boolean skipImage) {   
      finishedRecolor(result, null);
      return;
    }

    private void finishedRecolor(Object result, IOException ioEx) {     
      postRecolorOperations((BufferedImage)result);
      return;
    }
  }
  
  /***************************************************************************
  **
  ** Background network recolor
  */ 
    
  private class RecolorNetworkRunner implements BackgroundCore {
 
    private boolean forMain_;
    private File holdIt_;
    private BFWorker bfwk_;
    
    public RecolorNetworkRunner(boolean forMain, File holdIt, BFWorker bfwk) {
      bfwk_ = bfwk;
      holdIt_ = holdIt;
      forMain_ = forMain;
    }
    
    public Object getEarlyResult() {
    	return (new BufferedImage(1, 1, BufferedImage.TYPE_INT_RGB)); 
    }
    
    public Object runCore() throws AsynchExitRequestException {
      try { 
      	BTProgressMonitor monitor = bfwk_.getMonitor();
        buildRestoreCache(holdIt_, monitor); 
        BufferedImage bi = expensiveRecolorOperations(forMain_, monitor);
        (new GarbageRequester()).askForGC(monitor);
        return (bi);
      } catch (IOException ex) {
        bfwk_.stashException(ex);
        return (null);
      }
    }
    
    public Object postRunCore() {
      return (null);
    } 
  }  
 
  /***************************************************************************
  **
  ** Build an empty network
  */
  
  public void buildEmptyNetwork() {
    BuildData obd = new BuildDataImpl(new UniqueLabeller(),
                                      new HashSet<NetLink>(), 
                                      new HashSet<NetNode>(), 
                                      new HashMap<NetNode, String>(),
                                      colGen_, 
                                      BuildDataImpl.BuildMode.BUILD_FROM_SIF);
    try {
      newModelOperations(obd, true);
    } catch (IOException ioex) {
      //Silent fail     
    }
    return;
  }
  
  
  /***************************************************************************
  **
  ** Do new model operations
  */ 

  public void preLoadOperations() { 
    bfp_.reset();
    return;
  }
  
  /***************************************************************************
  **
  ** Do new model operations
  */ 

  public BufferedImage expensiveModelOperations(BuildData bfnbd, 
                                                boolean forMain, 
                                                BTProgressMonitor monitor) throws IOException, 
  																																								AsynchExitRequestException,
  	                                                                              LayoutCriterionFailureException {
    Dimension screenSize = (forMain && (headlessOracle_ == null)) ? Toolkit.getDefaultToolkit().getScreenSize() : new Dimension(600, 800);
    // Possibly expensive network analysis preparation:
    BioFabricNetwork bfn = new BioFabricNetwork(bfnbd, pMan_, monitor);
    // Possibly expensive display object creation:
    bfp_.installModel(bfn, monitor);
    // Very expensive display buffer creation:
    int[] preZooms = bfp_.calcZoomSettings(screenSize);
    BufferedImage topImage = null;
    if (headlessOracle_ == null) {
      if (forMain) {
        BufferBuilder bb = new BufferBuilder(null, 100, bfp_, bfp_.getBucketRend(), bfp_.getBufImgStack());
        topImage = bb.buildBufs(preZooms, bfp_, 25, monitor);
        bfp_.setBufBuilder(bb);      
      } else {
        BufferBuilder bb = new BufferBuilder(bfp_, bfp_.getBucketRend(), bfp_.getBufImgStack());
        topImage = bb.buildOneBuf();      
        bfp_.setBufBuilder(null);
      }
    }
    return (topImage);
  }

  /***************************************************************************
  **
  ** Do new model operations
  */ 

  public BufferedImage expensiveRecolorOperations(boolean forMain,
                                                  BTProgressMonitor monitor) throws IOException, AsynchExitRequestException {
    Dimension screenSize = (forMain) ? Toolkit.getDefaultToolkit().getScreenSize() : new Dimension(800, 400);
    screenSize.setSize((int)(screenSize.getWidth() * 0.8), (int)(screenSize.getHeight() * 0.4));
    colGen_.newColorModel();
    bfp_.changePaint(monitor);
    int[] zoomLevels = bfp_.getZoomController().getZoomLevels();
    BufferedImage topImage = null;
    if (forMain) {
      BufferBuilder bb = new BufferBuilder(null, 100, bfp_, bfp_.getBucketRend(), bfp_.getBufImgStack());
      topImage = bb.buildBufs(zoomLevels, bfp_, 24, monitor);
      bfp_.setBufBuilder(bb);      
    } else {
      BufferBuilder bb = new BufferBuilder(bfp_, bfp_.getBucketRend(), bfp_.getBufImgStack());
      topImage = bb.buildOneBuf();      
      bfp_.setBufBuilder(null);
    }
    return (topImage);
  }
  
  /***************************************************************************
  **
  ** Handles post-recolor operations
  */ 
       
  public void postRecolorOperations(BufferedImage topImage) {
    bfw_.getOverview().installImage(topImage, bfp_.getWorldScreen());
    return;
  }
   
  /***************************************************************************
  **
  ** Handles post-loading operations
  */ 
       
  public void postLoadOperations(BufferedImage topImage, boolean skipImageLoad) {
  	// Sometimes when we cancel we need to not load an empty result over a restored image
  	if (!skipImageLoad) {
      bfw_.getOverview().installImage(topImage, bfp_.getWorldScreen());
  	}
    bfp_.installModelPost();
    bfp_.initZoom();
    // Doing this before checkForChanges lets the plugins know how to answer: 
    pMan_.newNetworkInstalled(bfp_.getNetwork());
    cSet_.checkForChanges();
    cSet_.handleZoomButtons();
    bfp_.repaint();
    return;
  }
  
  /***************************************************************************
  **
  ** Information available to the plugin
  */ 
    
  public static class PlugInInfo implements PlugInNetworkModelAPI {
   
    private BioFabricNetwork bfn_;
    private FileLoadFlows flf_;
    private BioFabricWindow bfw_;
    
    PlugInInfo(FileLoadFlowsImpl flf, BioFabricNetwork bfn, BioFabricWindow bfw) {
      flf_ = flf;
      bfn_ = bfn;
      bfw_ = bfw;
    }
    
    public int getLinkCount(boolean forShadow)  {
      return (bfn_.getLinkCount(forShadow));
    }
    
    public int getNodeCount() {
      return (bfn_.getNodeCount());
    }
    
    public FileLoadFlows getFileUtilities() {
      return (flf_);
    }

    public JFrame getTopWindow() {
      return (bfw_.getWindow());
    }
   
    public BackgroundWorkerControlManager getBWCtrlMgr() {
      return (bfw_);
    }
    
    public void stashPluginData(String keyword, BioFabricToolPlugInData data) {
      bfn_.stashPluginData(keyword, data);
      return;
    }
 
    public BioFabricToolPlugInData providePluginData(String keyword) {
      return (bfn_.providePluginData(keyword));
    }
    
    public boolean getDisplayShadows() {
    	FabricDisplayOptionsManager fdom = FabricDisplayOptionsManager.getMgr();
      return (fdom.getDisplayOptions().getDisplayShadows());
    }

  }

  /***************************************************************************
  **
  ** Do new model operations all on AWT thread!
  */ 

  public void newModelOperations(BuildData bfnbd, boolean forMain) throws IOException { 
    preLoadOperations();
    try {
      BufferedImage topImage = expensiveModelOperations(bfnbd, forMain, null);
      postLoadOperations(topImage, false);
    } catch (AsynchExitRequestException aex) {
      // Not being used in background; will not happen
    } catch (LayoutCriterionFailureException ex) {
      // No links to lay out; will not happen
    }
    return;
  }
    
  /***************************************************************************
  **
  ** Do window title
  */ 

  public void manageWindowTitle(String fileName) {
    if (headlessOracle_ != null) {
      return;
    }
    ResourceManager rMan = ResourceManager.getManager();
    String title;
    if (fileName == null) {
      title = rMan.getString("window.title");  
    } else {
      String titleFormat = rMan.getString("window.titleWithName");
      title = MessageFormat.format(titleFormat, new Object[] {fileName});
    }
    topWindow_.setTitle(title);
    return;
  }
  
  /***************************************************************************
  **
  ** Get the file load type.
  */ 

  public FileLoadFlows.FileLoadType getFileLoadType(File toCheck) {
    return ((GWImportLoader.isGWFile(toCheck)) ? FileLoadFlows.FileLoadType.GW : FileLoadFlows.FileLoadType.SIF);
  }
  
  /***************************************************************************
  **
  ** Return value for file load flows.
  */ 
    
  public static class FileLoadResultImpl implements FileLoadFlows.FileLoadResult {
  	
  	private boolean success_;
  	private File cacheFile_;
  	
  	public FileLoadResultImpl(boolean success, File cacheFile) {
  		success_ = success;
  		cacheFile_ = cacheFile;			
  	}
  	  	
 		public boolean getSuccess() {
 			return (success_);		
 		}
 		
		public File getCacheFile() {
			return (cacheFile_);
		}
  }
}
```
