## Supplementary material for "BioFabric Visualization of Network Alignments": BioFabric Source Code: GWImportLoader.java

```
/*
**    File created by Rishi Desai
**
**    Copyright (C) 2003-2018 Institute for Systems Biology
**                            Seattle, Washington, USA.
**
**    This library is free software; you can redistribute it and/or
**    modify it under the terms of the GNU Lesser General Public
**    License as published by the Free Software Foundation; either
**    version 2.1 of the License, or (at your option) any later version.
**
**    This library is distributed in the hope that it will be useful,
**    but WITHOUT ANY WARRANTY; without even the implied warranty of
**    MERCHANTABILITY or FITNESS FOR A PARTICULAR PURPOSE.  See the GNU
**    Lesser General Public License for more details.
**
**    You should have received a copy of the GNU Lesser General Public
**    License along with this library; if not, write to the Free Software
**    Foundation, Inc., 59 Temple Place, Suite 330, Boston, MA  02111-1307  USA
*/

package org.systemsbiology.biofabric.io;

import java.io.BufferedReader;
import java.io.File;
import java.io.FileInputStream;
import java.io.IOException;
import java.io.InputStreamReader;
import java.util.ArrayList;
import java.util.HashMap;
import java.util.List;
import java.util.Map;
import java.util.Set;
import javax.swing.JFrame;

import org.systemsbiology.biofabric.api.model.NetLink;
import org.systemsbiology.biofabric.api.model.NetNode;
import org.systemsbiology.biofabric.api.util.UniqueLabeller;
import org.systemsbiology.biofabric.api.worker.AsynchExitRequestException;
import org.systemsbiology.biofabric.api.worker.BTProgressMonitor;
import org.systemsbiology.biofabric.api.worker.LoopReporter;
import org.systemsbiology.biofabric.model.FabricLink;
import org.systemsbiology.biofabric.ui.dialogs.LinkRelationDialog;

/****************************************************************************
 **
 ** This loads GW (LEDA file format) files
 */

public class GWImportLoader extends FabricImportLoader {
  
  ////////////////////////////////////////////////////////////////////////////
  //
  // PRIVATE CONSTANTS
  //
  ////////////////////////////////////////////////////////////////////////////
  
  private static final String DEFAULT_RELATION = "default";
  private static final int HEADER_LINES = 4; // # of header/parameter/version-info lines at the beginning of file
  
  ////////////////////////////////////////////////////////////////////////////
  //
  // PRIVATE INSTANCE MEMBERS
  //
  ////////////////////////////////////////////////////////////////////////////
  
  private int lineToTokIndex_, consTokIndex_;
  private Integer numNodes_, numEdges_;
  private Map<Integer, String> indexToName_;
  private Map<Integer, Boolean> indexUsed_;
  
  ////////////////////////////////////////////////////////////////////////////
  //
  // PUBLIC CONSTRUCTORS
  //
  ////////////////////////////////////////////////////////////////////////////
  
  /***************************************************************************
   **
   ** Constructor
   */
  
  public GWImportLoader() {
    this.lineToTokIndex_ = 0; // counter while reading tokens
    this.consTokIndex_ = 0;   // counter while consuming tokens
    this.indexToName_ = new HashMap<Integer, String>();
    this.indexUsed_ = new HashMap<Integer, Boolean>();
  }
  
  /***************************************************************************
   **
   ** Parse a line to tokens
   */
  
  protected String[] lineToToks(String line, FileImportStats stats) throws IOException {
    if (line.trim().equals("")) {
      return (null);
    }
    
    //
    // length == 1: Node Name or parameters(lines 0-3); Length == 4: Edge
    //
  
    String[] tokens = line.split("\\t");
    if ((tokens.length == 1) && (line.indexOf("\\t") == -1)) {
      tokens = line.split(" ");
    }
    
    if (tokens.length == 0 || tokens.length == 2 || tokens.length == 3 || tokens.length > 4) {
      stats.badLines.add(line);
      return (null);
    } else {
      
      if (lineToTokIndex_ == HEADER_LINES) {
        try {
          numNodes_ = Integer.parseInt(tokens[0].trim());
        } catch (NumberFormatException ex) {
          throw new IOException("We assume 4 header lines");
        }
      }
      if (numNodes_ != null && lineToTokIndex_ == HEADER_LINES + numNodes_ + 1) {
        try {
          numEdges_ = Integer.parseInt(tokens[0].trim());
        } catch (NumberFormatException ex) {
          throw new IOException("We assume 4 header lines");
        }
      }
      
      lineToTokIndex_++;
      
      return (tokens);
    }
  }
  
  /***************************************************************************
   **
   ** Consume tokens, process nodes, make links
   */
  
  protected void consumeTokens(String[] tokens, UniqueLabeller idGen, List<NetLink> links,
                               Set<NetNode> loneNodeIDs, Map<String, String> nameMap,
                               Integer magBins, Map<String, NetNode> nameToID,
                               FileImportStats stats) throws IOException {
    
    if (tokens.length == 1) {
  
      if (consTokIndex_ > HEADER_LINES && consTokIndex_ != HEADER_LINES + numNodes_ + 1) {
        
        int index = consTokIndex_ - HEADER_LINES; // nodes index starts with one
        
        String nodeName = tokens[0].trim();
        nodeName = stripBrackets(nodeName);
        
        indexToName_.put(index, nodeName);
        indexUsed_.put(index, false);
      }
      
    } else if (tokens.length == 4) {
      
      String sourceIndexStr = tokens[0].trim();
      String targetIndexStr = tokens[1].trim();
      String edgeRelation = tokens[3].trim();
      
      int sourceIndex, targetIndex;
      try {
        sourceIndex = Integer.parseInt(sourceIndexStr);
        targetIndex = Integer.parseInt(targetIndexStr);
      } catch (NumberFormatException ex) {
        throw new IOException("Could not parse integer");
      }
      
      indexUsed_.put(sourceIndex, true);
      indexUsed_.put(targetIndex, true);
      
      String sourceName = indexToName_.get(sourceIndex);
      String targetName = indexToName_.get(targetIndex);
      
      sourceName = stripBrackets(sourceName);
      targetName = stripBrackets(targetName);
      
      NetNode srcID = nameToNode(sourceName, idGen, nameToID);
      NetNode trgID = nameToNode(targetName, idGen, nameToID);
  
      edgeRelation = stripBrackets(edgeRelation);
      
      if (edgeRelation.equals("")) {
        edgeRelation = DEFAULT_RELATION;
      }
  
      //
      // Build the link, plus shadow if not auto feedback:
      //
  
      buildLinkAndShadow(srcID, trgID, edgeRelation, links);
  
    } else {
      throw new IllegalArgumentException();
    }
    
    consTokIndex_++;
    
    if (consTokIndex_ == HEADER_LINES + numNodes_ + 1 + numEdges_ + 1) { // reached end of file
      addLoneNodes(idGen, loneNodeIDs, nameToID);
    }
    // We don't check if there may be more lines or edges after the specified number of edges
    return;
  }
  
  private void addLoneNodes(UniqueLabeller idGen, Set<NetNode> loneNodeIDs,
                            Map<String, NetNode> nameToID) {
    
    for (Map.Entry<Integer, Boolean> entry : indexUsed_.entrySet()) {
      
      if (! entry.getValue()) {
        String loner = indexToName_.get(entry.getKey());
        NetNode lonerID = nameToNode(loner, idGen, nameToID);
        loneNodeIDs.add(lonerID);
      }
    }
    return;
  }
  
  ////////////////////////////////////////////////////////////////////////////
  //
  // STATIC METHODS AND CLASSES
  //
  ////////////////////////////////////////////////////////////////////////////
  
  /***************************************************************************
   **
   ** Manages links if GW file did not provide link relations
   */
  
  public static class GWRelationManager {
    
    public GWRelationManager() {}
    
    public boolean process(final List<NetLink> links, JFrame parent, BTProgressMonitor monitor)
            throws AsynchExitRequestException {
  
      LoopReporter lr;
      lr = new LoopReporter(links.size(), 20, monitor, 0.0, .3, "progress.processingLinkRelations");
      List<NetLink> linksToChange = new ArrayList<NetLink>(),
              returnLinks = new ArrayList<NetLink>();
      
      for (NetLink link : links) {  // only add links that weren't given relation (usually all of them)
        lr.report();
        if (link.getRelation().equals(DEFAULT_RELATION)) {
          linksToChange.add(link);
        } else {
          returnLinks.add(link);
        }
      }
      lr.finish();
      
      if (linksToChange.isEmpty()) { // leave if all links have a normal relation
        return (true);
      }
      
      LinkRelationDialog dialog = new LinkRelationDialog(parent, DEFAULT_RELATION);
      dialog.setVisible(true);
      if (!dialog.haveResult()) {
        // should not happen, should always have relation returned
        return (false);
      }
      
      final String newRel = dialog.getRelation();
      
      if (newRel.equals(DEFAULT_RELATION)) { // if user chooses default no point continuing
        return (true);
      }
      
      //
      // Create new links to replace old ones with default relation
      //
      
      lr = new LoopReporter(linksToChange.size(), 20, monitor, 0.3, .6, "progress.changingLinkRelations");
      
      for (NetLink link : linksToChange) {
        lr.report();
        NetLink newLink = new FabricLink(link.getSrcNode(), link.getTrgNode(), newRel, link.isShadow());
        returnLinks.add(newLink);
      }
      lr.finish();
      
      if (links.size() != returnLinks.size()) {
        throw (new IllegalStateException("Return links different size in GWRelationManager"));
      }
      
      //
      // Re-add return set of links to original parameter set
      //
      
      lr = new LoopReporter(returnLinks.size(), 20, monitor, .6, 1.00, "progress.addingNewLinks");
      links.clear();
      
      for (NetLink link : returnLinks) {
        lr.report();
        links.add(link);
      }
      lr.finish();
      return (true);
    }
    
  }
  
  /***************************************************************************
   **
   ** Test if file is .gw file
   */
  
  public static boolean isGWFile(File file) {
  	BufferedReader in = null;
    try {
      in = new BufferedReader(new InputStreamReader(new FileInputStream(file), "UTF-8"));
  
      String line = in.readLine();
      if (line.equals("LEDA.GRAPH")) {
        return (true);
      }
      for (int i = 1; i < HEADER_LINES; i++) { // skip next three lines; We assume four header lines
        in.readLine();
      }
      line = in.readLine();
      if (line == null) { // is sif file that has very few lines
        return (false);
      }
      String[] tok = line.split(" ");
      if (tok.length != 1) {  // should be #nodes here
        return (false);
      }
      
      int numNodes = Integer.parseInt(tok[0]); // to test if #nodes is in fact integer
      if (numNodes > 0) {
        String aNode = in.readLine();
        int len = aNode.length();
        if (len < 4) { // No guarantee there are four characters to substring:
        	return (false);
        }
        if (aNode.substring(0,2).equals("|{") && aNode.substring(len - 2, len).equals("}|")) {
          return (true);
        }
      } else {
        return (false);
      }
  
    } catch (IOException ioe) {
      return (false);
    } catch (NumberFormatException nfe) {
      return (false);
    } finally {
    	if (in != null) {
    		try {
    		  in.close();
    		} catch (IOException ioe) {		
    		}
    	}
    }
    return (false);
  }
  
}
```
