## Supplementary material for "BioFabric Visualization of Network Alignments": BioFabric Source Code: SIFImportLoader.java

```
/*
**    Copyright (C) 2003-2018 Institute for Systems Biology 
**                            Seattle, Washington, USA. 
**
**    This library is free software; you can redistribute it and/or
**    modify it under the terms of the GNU Lesser General Public
**    License as published by the Free Software Foundation; either
**    version 2.1 of the License, or (at your option) any later version.
**
**    This library is distributed in the hope that it will be useful,
**    but WITHOUT ANY WARRANTY; without even the implied warranty of
**    MERCHANTABILITY or FITNESS FOR A PARTICULAR PURPOSE.  See the GNU
**    Lesser General Public License for more details.
**
**    You should have received a copy of the GNU Lesser General Public
**    License along with this library; if not, write to the Free Software
**    Foundation, Inc., 59 Temple Place, Suite 330, Boston, MA  02111-1307  USA
*/

package org.systemsbiology.biofabric.io;

import java.io.IOException;
import java.util.List;
import java.util.Map;
import java.util.Set;

import org.systemsbiology.biofabric.api.model.NetLink;
import org.systemsbiology.biofabric.api.model.NetNode;
import org.systemsbiology.biofabric.api.util.UniqueLabeller;

/****************************************************************************
**
** This loads SIF files
*/

public class SIFImportLoader extends FabricImportLoader {
  
  ////////////////////////////////////////////////////////////////////////////
  //
  // PRIVATE CONSTANTS
  //
  //////////////////////////////////////////////////////////////////////////// 
  
  ////////////////////////////////////////////////////////////////////////////
  //
  // PUBLIC CONSTANTS
  //
  //////////////////////////////////////////////////////////////////////////// 
  
  ////////////////////////////////////////////////////////////////////////////
  //
  // PRIVATE INSTANCE MEMBERS
  //
  ////////////////////////////////////////////////////////////////////////////
   
  ////////////////////////////////////////////////////////////////////////////
  //
  // PUBLIC CONSTRUCTORS
  //
  ////////////////////////////////////////////////////////////////////////////

  /***************************************************************************
  **
  ** Constructor
  */

  public SIFImportLoader() { 
  }

  ////////////////////////////////////////////////////////////////////////////
  //
  // PUBLIC METHODS
  //
  ////////////////////////////////////////////////////////////////////////////
   
  /***************************************************************************
  ** 
  ** Parse a line to tokens
  */

  protected String[] lineToToks(String line, FileImportStats stats) throws IOException {
  	if (line.trim().equals("")) {
  		return (null);
  	}
    String[] tokens = line.split("\\t");
    if ((tokens.length == 1) && (line.indexOf("\\t") == -1)) {
      tokens = line.split(" ");
    }
    
    if (tokens.length == 0) {
      return (null);
    } else if ((tokens.length == 2) || (tokens.length > 3)) {
      stats.badLines.add(line);
      return (null);
    } else {        
      return (tokens);
    }
  }

 /***************************************************************************
  ** 
  ** Consume tokens, make links
  */

  protected void consumeTokens(String[] tokens, UniqueLabeller idGen, List<NetLink> links, 
  		                         Set<NetNode> loneNodeIDs, Map<String, String> nameMap, Integer magBins, 
  		                         Map<String, NetNode> nameToID, FileImportStats stats) throws IOException {
    if (tokens.length == 3) {
      String source = tokens[0].trim();
      source = stripQuotes(source);
    
      String target = tokens[2].trim();
      target = stripQuotes(target);
  
      //
      // This name map is for the load SIF with node attributes feature:
      //
      
      source = mapName(source, nameMap);
      target = mapName(target, nameMap);
      
      //
      // Map the name to an ID, if none yet, get a new ID and assign it
      //
      
      NetNode srcID = nameToNode(source, idGen, nameToID);
      NetNode trgID = nameToNode(target, idGen, nameToID);

      String rel = tokens[1].trim();
      rel = stripQuotes(rel);
    
      //
      // Build the link, plus shadow if not auto feedback:
      //
      
      buildLinkAndShadow(srcID, trgID, rel, links);
      
    } else {
      String loner = tokens[0].trim();
      loner = stripQuotes(loner);
      loner = mapName(loner, nameMap);
       
      NetNode lonerID = nameToNode(loner, idGen, nameToID);  
      loneNodeIDs.add(lonerID); 
    }
    return;
  }
}
```
