## Supplementary material for "BioFabric Visualization of Network Alignments": BioFabric Source Code: ControlTopLayout.java

```
/*
**    Copyright (C) 2003-2018 Institute for Systems Biology 
**                            Seattle, Washington, USA. 
**
**    This library is free software; you can redistribute it and/or
**    modify it under the terms of the GNU Lesser General Public
**    License as published by the Free Software Foundation; either
**    version 2.1 of the License, or (at your option) any later version.
**
**    This library is distributed in the hope that it will be useful,
**    but WITHOUT ANY WARRANTY; without even the implied warranty of
**    MERCHANTABILITY or FITNESS FOR A PARTICULAR PURPOSE.  See the GNU
**    Lesser General Public License for more details.
**
**    You should have received a copy of the GNU Lesser General Public
**    License along with this library; if not, write to the Free Software
**    Foundation, Inc., 59 Temple Place, Suite 330, Boston, MA  02111-1307  USA
*/

package org.systemsbiology.biofabric.layouts;

import java.util.ArrayList;
import java.util.Collections;
import java.util.HashMap;
import java.util.HashSet;
import java.util.List;
import java.util.Map;
import java.util.Set;
import java.util.SortedSet;
import java.util.TreeSet;
import java.util.Vector;

import org.systemsbiology.biofabric.analysis.CycleFinder;
import org.systemsbiology.biofabric.analysis.GraphSearcher;
import org.systemsbiology.biofabric.api.io.BuildData;
import org.systemsbiology.biofabric.api.layout.DefaultEdgeLayout;
import org.systemsbiology.biofabric.api.layout.LayoutCriterionFailureException;
import org.systemsbiology.biofabric.api.layout.NodeLayout;
import org.systemsbiology.biofabric.api.model.NetLink;
import org.systemsbiology.biofabric.api.model.NetNode;
import org.systemsbiology.biofabric.api.worker.AsynchExitRequestException;
import org.systemsbiology.biofabric.api.worker.BTProgressMonitor;
import org.systemsbiology.biofabric.api.worker.LoopReporter;
import org.systemsbiology.biofabric.model.BioFabricNetwork;
import org.systemsbiology.biofabric.util.DataUtil;
import org.systemsbiology.biofabric.util.ResourceManager;
import org.systemsbiology.biofabric.util.TrueObjChoiceContent;
import org.systemsbiology.biofabric.util.UiUtil;


/****************************************************************************
**
** Control nodes at top
*/

public class ControlTopLayout extends NodeLayout {
  
  ////////////////////////////////////////////////////////////////////////////
  //
  // PRIVATE CONSTANTS
  //
  //////////////////////////////////////////////////////////////////////////// 

  ////////////////////////////////////////////////////////////////////////////
  //
  // PUBLIC CONSTANTS
  //
  //////////////////////////////////////////////////////////////////////////// 
  
  public enum CtrlMode {
    
    CTRL_PARTIAL_ORDER("ctrlTop.ctrlPartialOrder"), 
    CTRL_INTRA_DEGREE_ONLY("ctrlTop.ctrlIntraDegree"), 
    CTRL_MEDIAN_TARGET_DEGREE("ctrlTop.ctrlMedianTarg"),
    CTRL_DEGREE_ONLY("ctrlTop.ctrlDegreeOnly"), 
    FIXED_LIST("ctrlTop.ctrlInputList"),
    ;    
 
    private String resource_;
     
    CtrlMode(String resource) {
      resource_ = resource;  
    }
     
    public static Vector<TrueObjChoiceContent<CtrlMode>> getControlChoices() {
      ResourceManager rMan = ResourceManager.getManager();
      Vector<TrueObjChoiceContent<CtrlMode>> retval = new Vector<TrueObjChoiceContent<CtrlMode>>();
      for (CtrlMode cm : CtrlMode.values()) {
        retval.add(new TrueObjChoiceContent<CtrlMode>(rMan.getString(cm.resource_), cm));
      }
      return (retval);
    }
  }
  
  public enum TargMode {
    TARGET_DEGREE("ctrlTop.trgTargDegree"), 
    NODE_DEGREE_ODOMETER_SOURCE("ctrlTop.degreeOdometer"),
    GRAY_CODE("ctrlTop.trgGrayCode"), 
    BREADTH_ORDER("ctrlTop.trgBreadth"), 
    ;
    
    private String resource_;
    
    TargMode(String resource) {
      resource_ = resource;  
    }
     
    public static Vector<TrueObjChoiceContent<TargMode>> getTargChoices() {
      ResourceManager rMan = ResourceManager.getManager();
      Vector<TrueObjChoiceContent<TargMode>> retval = new Vector<TrueObjChoiceContent<TargMode>>();
      for (TargMode tm : TargMode.values()) {
        retval.add(new TrueObjChoiceContent<TargMode>(rMan.getString(tm.resource_), tm));
      }
      return (retval);
    }
  }
  
  ////////////////////////////////////////////////////////////////////////////
  //
  // PRIVATE INSTANCE MEMBERS
  //
  ////////////////////////////////////////////////////////////////////////////
  
  private CtrlMode ctrlMode_;
  private TargMode targMode_;
  private List<String> fixedOrder_;
  private Map<String, Set<NetNode>> normNameToIDs_;
  private Set<NetNode> ctrlNodes_;
  
  ////////////////////////////////////////////////////////////////////////////
  //
  // PUBLIC CONSTRUCTORS
  //
  ////////////////////////////////////////////////////////////////////////////

  /***************************************************************************
  **
  ** Constructor
  */

  public ControlTopLayout(CtrlMode cMode, TargMode tMode, List<String> fixedOrder, Map<String, Set<NetNode>> normNameToIDs) {
    ctrlMode_ = cMode;
    targMode_ = tMode;
    fixedOrder_ = fixedOrder;
    normNameToIDs_ = normNameToIDs;
    ctrlNodes_ = null;
  }

  ////////////////////////////////////////////////////////////////////////////
  //
  // PUBLIC METHODS
  //
  ////////////////////////////////////////////////////////////////////////////
  
  /***************************************************************************
  **
  ** Find out if the necessary conditions for this layout are met. 
  */
  
  @Override
  public boolean criteriaMet(BuildData rbd,
                             BTProgressMonitor monitor) throws AsynchExitRequestException, 
                                                               LayoutCriterionFailureException {
    //
    // 1) The network must be fully directed:
  	//
  	
  	Set<NetLink> links = rbd.getLinks();
  	LoopReporter lr = new LoopReporter(links.size(), 20, monitor, 0.0, 1.0, "progress.ControlTopLayoutCriteriaCheck1");
  	
  	for (NetLink link : links) {
  		lr.report();
  	  if (!link.isDirected()) {
  	  	lr.finish();
  	  	throw new LayoutCriterionFailureException();
  	  }   	  	
  	}
  	lr.finish();
  	
  	//
    // 2) If there are no nodes with outbound links, what the heck are we doing? Get outta here!
    //
    
    ctrlNodes_ = controlNodes(rbd.getAllNodes(), rbd.getLinks(), monitor);
    if ((ctrlNodes_ == null) || ctrlNodes_.isEmpty()) {
      throw new LayoutCriterionFailureException();
    }
  	
    //
    // 3) If we are given a fixed list, we need to map to nodes, and make sure it is 1:1 and onto for all
    // the control nodes:
    //
  	
    if ((fixedOrder_ != null) && (normNameToIDs_ == null)) {
      throw new LayoutCriterionFailureException();
    }
     
    if (fixedOrder_ != null) {
    	HashSet<NetNode> checkSet = new HashSet<NetNode>(ctrlNodes_);
	    LoopReporter lr2 = new LoopReporter(fixedOrder_.size(), 20, monitor, 0.0, 1.0, "progress.ControlTopLayoutCriteriaCheck2");
	    for (String name : fixedOrder_) {
	    	lr2.report();
	    	Set<NetNode> matches = normNameToIDs_.get(DataUtil.normKey(name));
	    	if ((matches == null) || (matches.size() != 1)) {
	    		throw new LayoutCriterionFailureException();
	    	}
	    	NetNode match = matches.iterator().next();
	    	if (!checkSet.contains(match)) {
	    		throw new LayoutCriterionFailureException();
	    	}
	    	checkSet.remove(match);	    	
	    }
	    lr2.finish();
	    
	    if (!checkSet.isEmpty()) {
        throw new LayoutCriterionFailureException();
      }
    }
    
    return (true);  
  }
 
  /***************************************************************************
  **
  ** Order the nodes
  */
  
  public List<NetNode> doNodeLayout(BuildData rbd,
  																  Params params,
  		                              BTProgressMonitor monitor) throws AsynchExitRequestException {
									    
    
    List<NetNode> ctrlList;
    if (ctrlNodes_ == null) {
    	ctrlNodes_ = controlNodes(rbd.getAllNodes(), rbd.getLinks(), monitor);
    }
    SortedSet<NetNode> cnSet = new TreeSet<NetNode>(ctrlNodes_);
    List<NetNode> dfo = null;
    
    switch (ctrlMode_) {
      case CTRL_PARTIAL_ORDER:
        dfo = allNodeOrder(rbd.getAllNodes(), rbd.getLinks(), false, monitor);
        ctrlList = controlSortPartialOrder(rbd.getAllNodes(), rbd.getLinks(), cnSet, dfo, false, monitor);
        break;
      case CTRL_INTRA_DEGREE_ONLY:       
        ctrlList = controlSortIntraDegreeOnly(rbd.getAllNodes(), rbd.getLinks(), cnSet, false, monitor);
        break;
      case CTRL_DEGREE_ONLY:
        dfo = allNodeOrder(rbd.getAllNodes(), rbd.getLinks(), false, monitor);
        ctrlList = listToSublist(cnSet, dfo, monitor);
        break;   
      case CTRL_MEDIAN_TARGET_DEGREE:
        ctrlList = orderCtrlMedianTargetDegree(rbd.getAllNodes(), rbd.getLinks(), false, monitor);
        break;        
      case FIXED_LIST:
        ctrlList = null; // forcedTop;
        break;
      default:
        throw new IllegalStateException();
    }
     
    List<NetNode> nodeOrder;
    
    switch (targMode_) {
      case GRAY_CODE:
        nodeOrder = targetsBySourceGrayCode(ctrlList,cnSet, rbd.getAllNodes(), rbd.getLinks(), monitor);
        break;
      case NODE_DEGREE_ODOMETER_SOURCE:
        nodeOrder = targetsByNodeDegreeOdometerSourceMultigraph(ctrlList, cnSet, rbd.getAllNodes(), rbd.getLinks(), monitor);
        break;
      case TARGET_DEGREE:
        if (dfo == null) {
          dfo = allNodeOrder(rbd.getAllNodes(), rbd.getLinks(), false, monitor);
        }
        Set<NetNode> targs = new HashSet<NetNode>(rbd.getAllNodes());
        targs.removeAll(cnSet);
        nodeOrder = new ArrayList<NetNode>(ctrlList);
        nodeOrder.addAll(listToSublist(targs, dfo, monitor));
        break;
      case BREADTH_ORDER:
        nodeOrder = orderTargetsBreadth(ctrlList, cnSet, rbd.getAllNodes(), rbd.getLinks(), false, monitor);
        break;
      default:
        throw new IllegalStateException();
    }
    
    //
    // Singletons? Add them at the end
    //
    
    Set<NetNode> loneNodes = rbd.getSingletonNodes();
    if ((loneNodes != null) && !loneNodes.isEmpty()) {
    	TreeSet<NetNode> orderedLones = new TreeSet<NetNode>(loneNodes);
    	nodeOrder.addAll(orderedLones);
    }   
    
    //
    // Now have the ordered list of targets we are going to display.
    // Build target->row maps and the inverse:
    //
    installNodeOrder(nodeOrder, rbd, monitor);
    return (nodeOrder);
  }
  
  
  /***************************************************************************
  ** 
  ** Since we define control nodes as those with outputs, getting the source nodes
  ** from all the links gives us the set.
  */

  private Set<NetNode> controlNodes(Set<NetNode> nodes, Set<NetLink> links, 
                                         BTProgressMonitor monitor) throws AsynchExitRequestException {
    
    
    LoopReporter lr = new LoopReporter(links.size(), 20, monitor, 0.0, 1.0, "progress.findControlNodes"); 
    
    HashSet<NetNode> srcs = new HashSet<NetNode>();
    for (NetLink nextLink : links) {
      lr.report();
      srcs.add(nextLink.getSrcNode());
    }
    lr.finish();
    return (srcs);
  }
  
  /***************************************************************************
  ** 
  ** This method orders the control nodes using a partial ordering generated using
  ** a DAG subset of the links between the control nodes. Warning! Cycles are currently broken arbitrarily:
  */

  private List<NetNode> controlSortPartialOrder(Set<NetNode> nodes, Set<NetLink> links, 
                                                     SortedSet<NetNode> cnSet, List<NetNode> dfo, boolean relCollapse,
                                                     BTProgressMonitor monitor) throws AsynchExitRequestException {
    
    //
    // Figure out the links between control nodes:
    //
    
    HashSet<NetLink> ctrlLinks = new HashSet<NetLink>();
    for (NetLink nextLink : links) {
      if (cnSet.contains(nextLink.getTrgNode())) {
        ctrlLinks.add(nextLink);
      }
    } 

    //
    // Get an ordering of nodes, based on degree, that we use to break cycles. Get the control-set
    // links at the same time:
    //
    
    List<NetNode> ctrlNodes = listToSublist(cnSet, dfo, monitor);
    
    //
    // Use the order to create a reverse mapping:
    // 
  
    HashMap<NetNode, Integer> nodeToRow = new HashMap<NetNode, Integer>();
    
    int numNodes = ctrlNodes.size();
    for (int i = 0; i < numNodes; i++) {
      NetNode ctrlNode = ctrlNodes.get(i);
      nodeToRow.put(ctrlNode, Integer.valueOf(i));
    }

    //
    // Need to create a DAG, so we need to break cycles. Given our ordering, we hold out
    // links that point "up" until the end, and start adding them in, one at a time. If
    // we get a cycle (and we may not) we pull it back out. Not "optimum", but not 
    // arbitrary:
    //
    
    HashSet<NetLink> downLinks = new HashSet<NetLink>();
    HashSet<NetLink> upLinks = new HashSet<NetLink>();
    HashSet<NetLink> autoFeedLinks = new HashSet<NetLink>();
    
    for (NetLink ctrlLink : ctrlLinks) {
      if (ctrlLink.isShadow()) {
        continue;
      }
      NetNode srcID = ctrlLink.getSrcNode();
      NetNode trgID = ctrlLink.getTrgNode();
      int srcRow = nodeToRow.get(srcID).intValue();
      int trgRow = nodeToRow.get(trgID).intValue();
      if (srcRow < trgRow) {
        downLinks.add(ctrlLink);
      } else if (srcRow > trgRow) {
        upLinks.add(ctrlLink);
      } else {
        autoFeedLinks.add(ctrlLink);
      }
    }
    
    //
    // This fixes the order of the testing of up links so it is deterministic:
    //
    
    DefaultEdgeLayout.DefaultFabricLinkLocater dfll = 
      new DefaultEdgeLayout.DefaultFabricLinkLocater(nodeToRow, null, null, BioFabricNetwork.LayoutMode.UNINITIALIZED_MODE);
    TreeSet<NetLink> upLinkOrder = new TreeSet<NetLink>(dfll);
    upLinkOrder.addAll(upLinks);

    //
    // Create a subset of the control links that form a DAG:
    // Note the order is arbitrary:
    //
    
    HashSet<NetLink> dagLinks = new HashSet<NetLink>(downLinks);
    HashSet<NetLink> testLinks = new HashSet<NetLink>(dagLinks);
    HashSet<NetLink> heldOut = new HashSet<NetLink>(dagLinks);
     
    for (NetLink testLink : upLinkOrder) {
      testLinks.add(testLink);
      CycleFinder cf = new CycleFinder(nodes, testLinks, monitor); 
      if (!cf.hasACycle(monitor)) {
        dagLinks.add(testLink);
      } else {
        testLinks.remove(testLink);
        heldOut.add(testLink);
      }
    }
    
    //
    // Topo sort the nodes:
    //
 
    UiUtil.fixMePrintout("NO! Still arbitrary? (HashSet iteration??)");
    GraphSearcher gs = new GraphSearcher(new HashSet<NetNode>(ctrlNodes), dagLinks);
    Map<NetNode, Integer> ts = gs.topoSort(false);
    List<NetNode> retval = gs.topoSortToPartialOrdering(ts, links, relCollapse, monitor);
    
    //
    // Nodes that were dropped due to cycles still need to be added as 
    // source nodes! FIXME: Should add in degree order, not alpha order!
    //
    //
    
    TreeSet<NetNode> remaining = new TreeSet<NetNode>(ctrlNodes);
    remaining.removeAll(retval);
    retval.addAll(remaining);
 
    return (retval);
  }  

  /***************************************************************************
  ** 
  ** Calculate an ordering of ALL (source, target) nodes that puts the highest degree nodes first:
  */

  private List<NetNode> allNodeOrder(Set<NetNode> nodes, Set<NetLink> links, boolean relCollapse,  
                                          BTProgressMonitor monitor) throws AsynchExitRequestException {  
    GraphSearcher gs = new GraphSearcher(nodes, links); 
    List<NetNode> retval = gs.nodeDegreeOrder(relCollapse, monitor);
    Collections.reverse(retval);
    return (retval);
  }

  /***************************************************************************
  ** 
  ** Calculate a control node order based purely on degree of *intra-control*
  ** block links:
  */

  private List<NetNode> controlSortIntraDegreeOnly(Set<NetNode> nodes, Set<NetLink> links, 
                                                        SortedSet<NetNode> ctrlNodes, boolean relCollapse,
                                                        BTProgressMonitor monitor) throws AsynchExitRequestException {    
    //
    // Sort the nodes so we can apply a standard order to the nodes of zero degree at the start:
    //
    
    HashSet<NetLink> ctrlLinks = new HashSet<NetLink>();
    for (NetLink nextLink : links) {
      if (ctrlNodes.contains(nextLink.getTrgNode())) {
        ctrlLinks.add(nextLink);
      }
    }   
 
    GraphSearcher gs = new GraphSearcher(ctrlNodes, ctrlLinks); 
    List<NetNode> retval = gs.nodeDegreeOrder(relCollapse, monitor);
    
    //
    // Not all of the nodes get returned, if they have no inbound control links. So
    // this step adds them:
    //
    
    ctrlNodes.removeAll(retval);
    retval.addAll(ctrlNodes);
  
    Collections.reverse(retval);
    return (retval);
  }
 
  /***************************************************************************
  **
  ** Pure target nodes are ordered by breadth-first search from the 
  ** provided control nodes. 
  */

  private List<NetNode> orderTargetsBreadth(List<NetNode> ctrlList,
                                                 Set<NetNode> cnSet,
                                                 Set<NetNode> nodes, 
                                                 Set<NetLink> links,
                                                 boolean relCollapse,
                                                 BTProgressMonitor monitor) throws AsynchExitRequestException {
   
    GraphSearcher gs = new GraphSearcher(nodes, links);
    List<GraphSearcher.QueueEntry> queue = gs.breadthSearch(ctrlList, relCollapse, monitor);
    
    ArrayList<NetNode> outList = new ArrayList<NetNode>(ctrlList);
    for (GraphSearcher.QueueEntry qe : queue) {
      if (!cnSet.contains(qe.name)) {
        outList.add(qe.name);
      }
    }
    return (outList);
  }
  
  /***************************************************************************
  **
  ** Target nodes are ordered by degree. Within degree, targets are arranged
  ** to "odometer" thru the inputs
  */
  
  private List<NetNode> targetsBySourceGrayCode(List<NetNode> ctrlList,
                                                     Set<NetNode> cnSet,
                                                     Set<NetNode> nodes, 
                                                     Set<NetLink> links,
                                                     BTProgressMonitor monitor) throws AsynchExitRequestException {
   
    
    ArrayList<NetNode> outList = new ArrayList<NetNode>(ctrlList);
    GraphSearcher gs = new GraphSearcher(nodes, links);
    SortedSet<GraphSearcher.SourcedNodeGray> sngr = gs.nodeGraySetWithSource(ctrlList);
    for (GraphSearcher.SourcedNodeGray node : sngr) {
      if (!cnSet.contains(node.getNodeID())) {
        outList.add(node.getNodeID());
      }
    } 
    return (outList);
  }
  
  /***************************************************************************
  **
  ** Target nodes are ordered by degree. Within degree, targets are arranged
  ** to "odometer" thru the inputs
  */
  
  private List<NetNode> targetsByNodeDegreeOdometerSource(List<NetNode> ctrlList,
                                                               Set<NetNode> cnSet,
                                                               Set<NetNode> nodes, 
                                                               Set<NetLink> links,
                                                               BTProgressMonitor monitor) throws AsynchExitRequestException {
   
    ArrayList<NetNode> outList = new ArrayList<NetNode>(ctrlList);
    GraphSearcher gs = new GraphSearcher(nodes, links); 
    SortedSet<GraphSearcher.SourcedNodeDegree> snds = gs.nodeDegreeSetWithSource(ctrlList);    
    for (GraphSearcher.SourcedNodeDegree node : snds) {
      if (!cnSet.contains(node.getNode())) {
        outList.add(node.getNode());
      }
    }
    return (outList);
  }
  
  /***************************************************************************
  **
  ** Target nodes are ordered by degree. Within degree, targets are arranged
  ** to "odometer" thru the inputs
  */
  
  private List<NetNode> targetsByNodeDegreeOdometerSourceMultigraph(List<NetNode> ctrlList,
                                                               					 Set<NetNode> cnSet,
                                                               					 Set<NetNode> nodes, 
                                                               					 Set<NetLink> links,
                                                               					 BTProgressMonitor monitor) throws AsynchExitRequestException {
   
    ArrayList<NetNode> outList = new ArrayList<NetNode>(ctrlList);
    GraphSearcher gs = new GraphSearcher(nodes, links); 
    SortedSet<GraphSearcher.SourcedNodeAndRelDegree> snds = gs.nodeDegreeSetWithSourceMultigraph(ctrlList);    
    for (GraphSearcher.SourcedNodeAndRelDegree nodeAndRel : snds) {
      if (!cnSet.contains(nodeAndRel.getNode())) {
        outList.add(nodeAndRel.getNode());
      }
    }
    return (outList);
  }
  

  /***************************************************************************
  **
  ** Provide a sublist of original list, containing nodes in given set.
  */

  private List<NetNode> listToSublist(Set<NetNode> nodeSet, List<NetNode> dfo,
                                           BTProgressMonitor monitor) throws AsynchExitRequestException {
             
    ArrayList<NetNode> outList = new ArrayList<NetNode>();
    for (NetNode node : dfo) {
      if (nodeSet.contains(node)) {
        outList.add(node);
      }
    }
    return (outList);
  }
  
  /***************************************************************************
  **
  ** This creates an ordering of control nodes based on their median degree of their
  ** target nodes. 
  */
  
  private List<NetNode> orderCtrlMedianTargetDegree(Set<NetNode> nodes, Set<NetLink> links, boolean relCollapse,
                                                         BTProgressMonitor monitor) throws AsynchExitRequestException {
        
    ArrayList<NetNode> outList = new ArrayList<NetNode>();
    SortedSet<GraphSearcher.NodeDegree> ctrlMed = medianTargetDegree(nodes, links, relCollapse, monitor);
    for (GraphSearcher.NodeDegree nodeDeg : ctrlMed) {
      outList.add(nodeDeg.getNodeID());
    }
    Collections.reverse(outList);
    return (outList);
  }
  
  /***************************************************************************
  ** 
  ** Creates an ordered set of source nodes ordered by their median target degree
  */

  private SortedSet<GraphSearcher.NodeDegree> medianTargetDegree(Set<NetNode> nodes, Set<NetLink> links, boolean relCollapse,
                                                                 BTProgressMonitor monitor) throws AsynchExitRequestException {
    
    GraphSearcher gs = new GraphSearcher(nodes, links); 
    Map<NetNode, Integer> nDeg = gs.nodeDegree(true, relCollapse, monitor);
    
    HashMap<NetNode, List<Integer>> deg = new HashMap<NetNode, List<Integer>>();
    for (NetLink nextLink : links) {
      NetNode src = nextLink.getSrcNode();
      NetNode trg = nextLink.getTrgNode();
      Integer trgDeg = nDeg.get(trg);
      List<Integer> forSrc = deg.get(src);
      if (forSrc == null) {
        forSrc = new ArrayList<Integer>();
        deg.put(src, forSrc);
      }
      forSrc.add(trgDeg);
    }  
        
    TreeSet<GraphSearcher.NodeDegree> retval = new TreeSet<GraphSearcher.NodeDegree>();
    for (NetNode src : deg.keySet()) {
      List<Integer> forSrc = deg.get(src);
      Collections.sort(forSrc);
      int size = forSrc.size();    
      int medI = size / 2;
      Integer med = forSrc.get(medI);
      retval.add(new GraphSearcher.NodeDegree(src, med.intValue()));
    }
    return (retval);
  } 
}
```
