## Supplementary material for "BioFabric Visualization of Network Alignments": BioFabric Source Code: HierDAGLayout.java

```
/*
**    Copyright (C) 2003-2018 Institute for Systems Biology 
**                            Seattle, Washington, USA. 
**
**    This library is free software; you can redistribute it and/or
**    modify it under the terms of the GNU Lesser General Public
**    License as published by the Free Software Foundation; either
**    version 2.1 of the License, or (at your option) any later version.
**
**    This library is distributed in the hope that it will be useful,
**    but WITHOUT ANY WARRANTY; without even the implied warranty of
**    MERCHANTABILITY or FITNESS FOR A PARTICULAR PURPOSE.  See the GNU
**    Lesser General Public License for more details.
**
**    You should have received a copy of the GNU Lesser General Public
**    License along with this library; if not, write to the Free Software
**    Foundation, Inc., 59 Temple Place, Suite 330, Boston, MA  02111-1307  USA
*/

package org.systemsbiology.biofabric.layouts;

import java.util.ArrayList;
import java.util.Collections;
import java.util.HashMap;
import java.util.HashSet;
import java.util.Iterator;
import java.util.List;
import java.util.Map;
import java.util.Set;
import java.util.SortedMap;
import java.util.TreeSet;

import org.systemsbiology.biofabric.analysis.CycleFinder;
import org.systemsbiology.biofabric.analysis.GraphSearcher;
import org.systemsbiology.biofabric.api.io.BuildData;
import org.systemsbiology.biofabric.api.layout.LayoutCriterionFailureException;
import org.systemsbiology.biofabric.api.layout.NodeLayout;
import org.systemsbiology.biofabric.api.layout.DefaultEdgeLayout;
import org.systemsbiology.biofabric.api.model.AnnotationSet;
import org.systemsbiology.biofabric.api.model.Annot;
import org.systemsbiology.biofabric.api.model.NetLink;
import org.systemsbiology.biofabric.api.model.NetNode;
import org.systemsbiology.biofabric.api.util.MinMax;
import org.systemsbiology.biofabric.api.worker.AsynchExitRequestException;
import org.systemsbiology.biofabric.api.worker.BTProgressMonitor;
import org.systemsbiology.biofabric.api.worker.LoopReporter;

import org.systemsbiology.biofabric.plugin.PluginSupportFactory;

/****************************************************************************
**
** This is a hierarchical layout for a directed acyclic graph
*/

public class HierDAGLayout extends NodeLayout {
  
  ////////////////////////////////////////////////////////////////////////////
  //
  // PRIVATE INSTANCE MEMBERS
  //
  ////////////////////////////////////////////////////////////////////////////

   private Map<NetNode, Set<NetNode>> l2s_;
   private Map<NetNode, Integer> inDegs_;
   private Map<NetNode, Integer> outDegs_;
   private ArrayList<NetNode> placeList_;
   private HashMap<NetNode, Integer> nameToRow_;
   private boolean pointUp_;
  
  ////////////////////////////////////////////////////////////////////////////
  //
  // PUBLIC CONSTRUCTORS
  //
  ////////////////////////////////////////////////////////////////////////////

  /***************************************************************************
  **
  ** Constructor
  */

  public HierDAGLayout(boolean pointUp) {
     l2s_ = new HashMap<NetNode, Set<NetNode>>();
     inDegs_ = new HashMap<NetNode, Integer>();
     outDegs_ = new HashMap<NetNode, Integer>();
     placeList_ = new ArrayList<NetNode>();
     nameToRow_ = new HashMap<NetNode, Integer>();
     pointUp_ = pointUp;
  }

  ////////////////////////////////////////////////////////////////////////////
  //
  // PUBLIC METHODS
  //
  ////////////////////////////////////////////////////////////////////////////
 
  /***************************************************************************
  **
  ** Find out if the necessary conditions for this layout are met. 
  */
  
  @Override
  public boolean criteriaMet(BuildData rbd,
  		                       BTProgressMonitor monitor) throws AsynchExitRequestException, 
                                                             LayoutCriterionFailureException {
  	//
  	// 1) All the relations in the network are directed
  	// 2) There are no cycles in the network
  	//
	  
	  LoopReporter lr = new LoopReporter(rbd.getLinks().size(), 20, monitor, 0.0, 1.0, "progress.hDagLayoutCriteriaCheck");
	  
    for (NetLink aLink : rbd.getLinks()) {
      lr.report();
      if (!aLink.isDirected()) {
    	  throw new LayoutCriterionFailureException();
      }
    }
    lr.finish();
	  
  	CycleFinder cf = new CycleFinder(rbd.getAllNodes(), rbd.getLinks(), monitor);
    if (cf.hasACycle(monitor)) {
      throw new LayoutCriterionFailureException();
    }
    return (true); 	
  }

  /***************************************************************************
  **
  ** Generate the Node ordering
  */
  
  public List<NetNode> doNodeLayout(BuildData rbd,
  		                              Params params,
  		   													  BTProgressMonitor monitor) throws AsynchExitRequestException {
    
    List<NetNode> targets = orderByNodeDegree(rbd, monitor);       

    //
    // Now have the ordered list of targets we are going to display.
    // Build target->row maps and the inverse:
    //
    
    installNodeOrder(targets, rbd, monitor);
    
   // AnnotationSet nAnnots = generateNodeAnnotations(rbd, params);
  //  rbd.setNodeAnnotations(nAnnots);
      
  //  Map<Boolean, AnnotationSet> lAnnots = generateLinkAnnotations(rbd, params);
   // rbd.setLinkAnnotations(lAnnots);
    
     
    return (targets);
  }
  
  /***************************************************************************
  **
  ** Generate node annotations to tag each cluster
 
    
  private AnnotationSet generateNodeAnnotations(BuildData rbd, Params params) {
    
    AnnotationSet retval = new AnnotationSet();  
    
    TreeMap<Integer, NetNode> invert = new TreeMap<Integer, NetNode>();
    
    Map<NetNode, Integer> nod = rbd.getNodeOrder(); 
    
    for (NetNode node : nod.keySet()) {
      invert.put(nod.get(node), node);
    }
     
    String currClust = null;
    Integer startRow = null;
    Integer lastKey = invert.lastKey();
    for (Integer row : invert.keySet()) {
      NetNode node = invert.get(row);
      String clust = params.getClusterForNode(node);
      if (currClust == null) {
        currClust = clust;
        startRow = row;
        if (row.equals(lastKey)) {
          AnnotationSet.Annot annot = new AnnotationSet.Annot(currClust, startRow.intValue(), row.intValue(), 0, null);
          retval.addAnnot(annot);
          break;
        }
        continue;
      }
      if (currClust.equals(clust)) {
        if (row.equals(lastKey)) {
          AnnotationSet.Annot annot = new AnnotationSet.Annot(currClust, startRow.intValue(), row.intValue(), 0, null);
          retval.addAnnot(annot);
          break;
        }
        continue;
      } else { 
        // We have just entered a new cluster
        AnnotationSet.Annot annot = new AnnotationSet.Annot(currClust, startRow.intValue(), row.intValue() - 1, 0, null);

        retval.addAnnot(annot);
        startRow = row;
        currClust = clust;
        if (row.equals(lastKey)) {
          annot = new AnnotationSet.Annot(currClust, startRow.intValue(), row.intValue(), 0, null);

          retval.addAnnot(annot);
          break;
        }
      }
    }
    
    return (retval);
  }
  
  /***************************************************************************
  **
  ** Generate link annotations to tag each cluster and intercluster links
 
    
  private Map<Boolean, AnnotationSet> generateLinkAnnotations(BuildData rbd, Params params) { 
  	HashMap<Boolean, AnnotationSet> retval = new HashMap<Boolean, AnnotationSet>();
    
  	SortedMap<Integer,NetLink> lod = rbd.getLinkOrder();
  	
  	List<NetLink> noShadows = new ArrayList<NetLink>();
  	List<NetLink> withShadows = new ArrayList<NetLink>();
  	for (Integer col : lod.keySet()) {
  		NetLink fl = lod.get(col);
  		withShadows.add(fl);
  		if (!fl.isShadow()) {
  			noShadows.add(fl);
  		}
  	}

  	retval.put(Boolean.FALSE, generateLinkAnnotationsForSet(noShadows, params));
  	retval.put(Boolean.TRUE, generateLinkAnnotationsForSet(withShadows, params));
  	
    return (retval);
  }
   
  /***************************************************************************
  **
  ** Generate link annotations to tag each cluster and intercluster links
  
    
  private AnnotationSet generateLinkAnnotationsForSet(List<NetLink> linkList, Params params) { 
    
  	HashMap<String, MinMax> clustRanges = new HashMap<String, MinMax>();
  	 	
  	for (int i = 0; i < linkList.size() ; i++) {
  		NetLink fl = linkList.get(i);
  
  		String srcClust = params.getClusterForNode(fl.getSrcNode());
      String trgClust = params.getClusterForNode(fl.getTrgNode());
  		if (srcClust.equals(trgClust)) {
  			MinMax mmc = clustRanges.get(srcClust);
  			if (mmc == null) {
  				mmc = new MinMax(i);
  				clustRanges.put(srcClust, mmc);
  			}
  			mmc.update(i);
  		} else {
  			String combo = (srcClust.compareTo(trgClust) < 0) ? srcClust + "-" + trgClust : trgClust + "-" + srcClust;
   			MinMax mmc = clustRanges.get(combo);
  			if (mmc == null) {
  				mmc = new MinMax(i);
  				clustRanges.put(combo, mmc);
  			}
  			mmc.update(i);
  		}
  	}
  	
  	AnnotationSet afns = new AnnotationSet();
  	TreeMap<Integer, String> ord = new TreeMap<Integer, String>();
  	for (String aName : clustRanges.keySet()) {
  		MinMax mm = clustRanges.get(aName);
  		ord.put(Integer.valueOf(mm.min), aName);		
  	}
  	
    for (String aName : ord.values()) {
  		MinMax mm = clustRanges.get(aName);
  		AnnotationSet.Annot annot = new AnnotationSet.Annot(aName, mm.min, mm.max, 0, null);
  		afns.addAnnot(annot);
  	}
  
    return (afns);
  }
 
  /***************************************************************************
  **
  ** Get the ordering of nodes by node degree:
  */
  
  private List<NetNode> orderByNodeDegree(BuildData rbd, 
  		                                    BTProgressMonitor monitor) throws AsynchExitRequestException {
    
    HashSet<NetNode> nodesToGo = new HashSet<NetNode>(rbd.getAllNodes());
    
    // Build map of sources to targets, also record in and out degrees of each node:
    linksToSources(rbd.getAllNodes(), rbd.getLinks(), monitor);
    
    List<NetNode> placeList = extractRoots(monitor);
   
    AnnotationSet nAnnots = PluginSupportFactory.buildAnnotationSet();
    Annot annot = PluginSupportFactory.buildAnnotation("Level 0", 0, placeList.size() - 1, 0, null);
    nAnnots.addAnnot(annot);
    
    addToPlaceList(placeList);
    nodesToGo.removeAll(placeList);
    
    //
    // Find the guys whose cites have already been placed and place them:
    //
  
    LoopReporter lr = new LoopReporter(nodesToGo.size(), 20, monitor, 0.0, 1.0, "progress.findingCandidates");
    
    int count = 1;
    while (!nodesToGo.isEmpty()) {
      List<NetNode> nextBatch = findNextCandidates();
      lr.report(nextBatch.size());
      int start = placeList_.size();
      addToPlaceList(nextBatch);
      annot = PluginSupportFactory.buildAnnotation("Level " + count++, start, placeList_.size() - 1, 0, null);
      nAnnots.addAnnot(annot);
      // Used to do removeAll, but this took FOREVER with the nextBatch as a List, and stalled the
      // progress counter.
      for (NetNode nbNod : nextBatch) {
    	  if (nodesToGo.contains(nbNod)) {
    		  nodesToGo.remove(nbNod);
    	  }
      }
    }
    lr.finish();
    
    rbd.setNodeAnnotations(nAnnots);
    
    return (placeList_);
    
  }
  
  /***************************************************************************
  ** 
  ** Construct a map of the targets of each node. Note that instance members
  */

  private void linksToSources(Set<NetNode> nodeList, Set<NetLink> linkList,
  		                        BTProgressMonitor monitor) throws AsynchExitRequestException {
    
  	//
  	// For each node, we initialize a map of nodes it is pointing at, and initialize the inDegree and outDegree
  	// entries for it as well:
  	//
  	
  	LoopReporter lr = new LoopReporter(nodeList.size(), 20, monitor, 0.0, 1.0, "progress.hDagLayoutInit");
  	
    Iterator<NetNode> nit = nodeList.iterator();
    while (nit.hasNext()) {
      NetNode node = nit.next();
      lr.report();
      l2s_.put(node, new HashSet<NetNode>());
      inDegs_.put(node, Integer.valueOf(0));
      outDegs_.put(node, Integer.valueOf(0));
    } 
    
    //
    // Crank thru the links, accumulate degrees and targets
    //
    
    LoopReporter lr2 = new LoopReporter(linkList.size(), 20, monitor, 0.0, 1.0, "progress.hDagDegAndTargs");
    
    Iterator<NetLink> llit = linkList.iterator();
    while (llit.hasNext()) {
      NetLink link = llit.next();
      lr2.report();
      //
      // By default, layout designed to have links point up. Quick way to switch this
      // is to reverse semantics of source and target:
      //
      NetNode src = (pointUp_) ? link.getSrcNode() : link.getTrgNode();
      NetNode trg = (pointUp_) ? link.getTrgNode() : link.getSrcNode();
      Set<NetNode> toTarg = l2s_.get(src);
      toTarg.add(trg);
      Integer deg = outDegs_.get(src);
      outDegs_.put(src, Integer.valueOf(deg.intValue() + 1));
      deg = inDegs_.get(trg);
      inDegs_.put(trg, Integer.valueOf(deg.intValue() + 1)); 
    } 
    return;
  }
  
  /***************************************************************************
  ** 
  ** Add to list to place
  */

  private void addToPlaceList(List<NetNode> nextBatch) {
    int nextRow = placeList_.size();
    for (NetNode nextNode : nextBatch) {
      placeList_.add(nextNode);
      nameToRow_.put(nextNode, Integer.valueOf(nextRow++));
    }
    return;
  }
  
  /***************************************************************************
  ** 
  ** Extract the root nodes in order from highest degree to low
  */

  private List<NetNode> extractRoots(BTProgressMonitor monitor) throws AsynchExitRequestException {
 
    LoopReporter lr = new LoopReporter(l2s_.size(), 20, monitor, 0.0, 1.0, "progress.rootExtractPass1");
    
    Map<NetNode, Integer> roots = new HashMap<NetNode, Integer>();
      
    Iterator<NetNode> lit = l2s_.keySet().iterator();
    while (lit.hasNext()) {
      NetNode node = lit.next();
      lr.report();
      Set<NetNode> fn = l2s_.get(node);
      if (fn.isEmpty()) {
        roots.put(node, Integer.valueOf(0));
      }
    }
    lr.finish();
    
    
    LoopReporter lr2 = new LoopReporter(l2s_.size(), 20, monitor, 0.0, 1.0, "progress.rootExtractPass2");
    lit = l2s_.keySet().iterator();
    while (lit.hasNext()) {
      NetNode node = lit.next();
      lr2.report();
      Set<NetNode> fn = l2s_.get(node);
      Iterator<NetNode> sit = fn.iterator();
      while (sit.hasNext()) {
        NetNode trg = sit.next();
        Integer rs = roots.get(trg);
        if (rs != null) {
          roots.put(trg, Integer.valueOf(rs.intValue() + 1));          
        }
      }
    }
    lr2.finish();
    
    ArrayList<NetNode> buildList = new ArrayList<NetNode>();
    
    LoopReporter lr3 = new LoopReporter(roots.size(), 20, monitor, 0.0, 1.0, "progress.rootExtractPass3");
    int count = 1;
    TreeSet<NetNode> alpha = new TreeSet<NetNode>(Collections.reverseOrder());
    alpha.addAll(roots.keySet());
    while (buildList.size() < roots.size()) {
      for (NetNode node : alpha) {
        Integer val = roots.get(node);
        if (val.intValue() == count) {
          buildList.add(node);
          lr3.report();
        }
      }
      count++;
    }
    lr3.finish();
    Collections.reverse(buildList);
    return (buildList);
  }
  
  /***************************************************************************
  ** 
  ** Find the next guys to go:
  */

  private List<NetNode> findNextCandidates() {
 
    HashSet<NetNode> quickie = new HashSet<NetNode>(placeList_);
     
    ArrayList<GraphSearcher.SourcedNode> nextOutList = new ArrayList<GraphSearcher.SourcedNode>();
    Iterator<NetNode> lit = l2s_.keySet().iterator();
    while (lit.hasNext()) {
      NetNode node = lit.next();
      if (quickie.contains(node)) {
        continue;
      }
      Set<NetNode> fn = l2s_.get(node);
      boolean allThere = true;
      Iterator<NetNode> sit = fn.iterator();
      while (sit.hasNext()) {
        NetNode trg = sit.next();
        if (!quickie.contains(trg)) {
          allThere = false;
          break;
        }
      }
      if (allThere) {
        nextOutList.add(new GraphSearcher.SourcedNode(node, inDegs_, nameToRow_, l2s_));
      }
    }
    
    //
    // Order the nodes:
    //
    
    TreeSet<GraphSearcher.SourcedNode> nextOut = new TreeSet<GraphSearcher.SourcedNode>(Collections.reverseOrder());
    nextOut.addAll(nextOutList);
    
    //
    // Make them a list:
    //
   
    ArrayList<NetNode> retval = new ArrayList<NetNode>();
    for (GraphSearcher.SourcedNode sn : nextOut) {
      retval.add(sn.getNode());
    }
    return (retval);
  }
  
  public static class EdgeLayout extends DefaultEdgeLayout {
  
	  ////////////////////////////////////////////////////////////////////////////
	  //
	  // PRIVATE INSTANCE MEMBERS
	  //
	  ////////////////////////////////////////////////////////////////////////////
	
	  ////////////////////////////////////////////////////////////////////////////
	  //
	  // PUBLIC CONSTRUCTORS
	  //
	  ////////////////////////////////////////////////////////////////////////////
	
	  /***************************************************************************
	  **
	  ** Constructor
	  */
	
	  public EdgeLayout() {
	  }
	
	  /***************************************************************************
	  **
	  ** Relayout the whole network!
	  */
	  
	  public void layoutEdges(BuildData rbd, 
	  		                    BTProgressMonitor monitor) throws AsynchExitRequestException {
	   
	    SortedMap<Integer, NetLink> retval = layoutEdges(rbd.getNodeOrder(), rbd.getLinks(), 
	    		                                             rbd.getGroupOrder(), rbd.getGroupOrderMode(), monitor);
	    rbd.setLinkOrder(retval);
	    installLinkAnnotations(rbd, monitor);
	    return;
	  }
	
	  /***************************************************************************
	  **
	  ** Install link annotations for levels
	  */
	  
	  protected void installLinkAnnotations(BuildData rbd, BTProgressMonitor monitor)
	    throws AsynchExitRequestException {
	  
	  	
	  	SortedMap<Integer, NetLink> lo = rbd.getLinkOrder();
	    LoopReporter lr = new LoopReporter(lo.size(), 20, monitor, 0, 1.0, "progress.linkAnnotationPrep");  
	    List<NetLink> linkList = new ArrayList<NetLink>();  
	    for (NetLink link : lo.values()) {   
	      linkList.add(link);
	      lr.report();
	    }
	    lr.finish();
	    
	    Map<NetNode, Integer> no = rbd.getNodeOrder();
	    LoopReporter lr2 = new LoopReporter(no.size(), 20, monitor, 0, 1.0, "progress.nodeOrderInvert");  
	    Map<Integer, NetNode> invNo = new HashMap<Integer, NetNode>();  
	    for (NetNode node : no.keySet()) {
	    	invNo.put(no.get(node), node);
	    	lr2.report();
	    }
	    lr2.finish();
	    
	    AnnotationSet withShdwAnnots = calcLevelLinkAnnots(rbd, linkList, monitor, invNo, rbd.getNodeAnnotations());
	    
	    Map<Boolean, AnnotationSet> linkAnnots = new HashMap<Boolean, AnnotationSet>();
	    linkAnnots.put(true, withShdwAnnots);
	    linkAnnots.put(false, PluginSupportFactory.buildAnnotationSet()); // No link annots if no shadows...
	    
	    rbd.setLinkAnnotations(linkAnnots);
	    return;
	  }
	 
	  /*********************************************************************************************
	  **
	  ** Calculate Level annotations. First link on first node in level starts, last link on last
	  ** node in level ends each annotation.
	  */
	    
	  protected AnnotationSet calcLevelLinkAnnots(BuildData rbd, 
	                                              List<NetLink> links, BTProgressMonitor monitor, 
	                                              Map<Integer, NetNode> nodeOrder, AnnotationSet nodeAnnot) throws AsynchExitRequestException {   
	    
	    LoopReporter lr = new LoopReporter(links.size(), 20, monitor, 0, 1.0, "progress.linkAnnotationShad"); 
	    HashMap<String, String> colorMap = new HashMap<String, String>();
	
	    //
	    // Go through the links, look for first shadow link into start of node annotation group. That's the break
	    // for the link annotation.
	    //
	    
	    AnnotationSet retval = PluginSupportFactory.buildAnnotationSet();
	    boolean first = true;
	    int startPos = 0;
	    int numLink = links.size();
	    int annotCount = 0;
	    for (Annot currAnnot : nodeAnnot) {
	    	if (first) {
	    		first = false;
	    		continue;
	    	}
	    	MinMax mm = currAnnot.getRange();
	    	NetNode start = nodeOrder.get(mm.min);
	    	
	      for (int i = 0; i < numLink; i++) {
	        NetLink link = links.get(i);
	        NetNode src = link.getSrcNode();
	        if (link.getAugRelation().isShadow && src.equals(start)) {
	        	retval.addAnnot(PluginSupportFactory.buildAnnotation("Level " + annotCount++, startPos, i - 1, 0, null));
	        	startPos = i;
	        	break;
	        }
	      }
	    }
	    retval.addAnnot(PluginSupportFactory.buildAnnotation("Level " + annotCount, startPos, numLink - 1, 0, null));
	    return (retval);
	  }
  }
}
```
