## Supplementary material for "BioFabric Visualization of Network Alignments": BioFabric Source Code: NodeClusterLayout.java

```
/*
**    Copyright (C) 2003-2018 Institute for Systems Biology 
**                            Seattle, Washington, USA. 
**
**    This library is free software; you can redistribute it and/or
**    modify it under the terms of the GNU Lesser General Public
**    License as published by the Free Software Foundation; either
**    version 2.1 of the License, or (at your option) any later version.
**
**    This library is distributed in the hope that it will be useful,
**    but WITHOUT ANY WARRANTY; without even the implied warranty of
**    MERCHANTABILITY or FITNESS FOR A PARTICULAR PURPOSE.  See the GNU
**    Lesser General Public License for more details.
**
**    You should have received a copy of the GNU Lesser General Public
**    License along with this library; if not, write to the Free Software
**    Foundation, Inc., 59 Temple Place, Suite 330, Boston, MA  02111-1307  USA
*/

package org.systemsbiology.biofabric.layouts;

import java.util.ArrayList;
import java.util.Collection;
import java.util.Collections;
import java.util.HashMap;
import java.util.HashSet;
import java.util.Iterator;
import java.util.List;
import java.util.Map;
import java.util.Set;
import java.util.SortedMap;
import java.util.SortedSet;
import java.util.TreeMap;
import java.util.TreeSet;
import java.util.Vector;

import org.systemsbiology.biofabric.analysis.GraphSearcher;
import org.systemsbiology.biofabric.api.io.AttributeKey;
import org.systemsbiology.biofabric.api.io.BuildData;
import org.systemsbiology.biofabric.api.layout.DefaultEdgeLayout;
import org.systemsbiology.biofabric.api.layout.DefaultLayout;
import org.systemsbiology.biofabric.api.layout.NodeLayout;
import org.systemsbiology.biofabric.api.model.Annot;
import org.systemsbiology.biofabric.api.model.AnnotationSet;
import org.systemsbiology.biofabric.api.model.NetLink;
import org.systemsbiology.biofabric.api.model.NetNode;
import org.systemsbiology.biofabric.api.util.MinMax;
import org.systemsbiology.biofabric.api.util.NID;
import org.systemsbiology.biofabric.api.util.UniqueLabeller;
import org.systemsbiology.biofabric.api.worker.AsynchExitRequestException;
import org.systemsbiology.biofabric.api.worker.BTProgressMonitor;
import org.systemsbiology.biofabric.io.AttributeLoader;
import org.systemsbiology.biofabric.io.BuildDataImpl;

import org.systemsbiology.biofabric.model.FabricLink;
import org.systemsbiology.biofabric.model.FabricNode;
import org.systemsbiology.biofabric.plugin.PluginSupportFactory;
import org.systemsbiology.biofabric.util.DataUtil;
import org.systemsbiology.biofabric.util.ResourceManager;
import org.systemsbiology.biofabric.util.TrueObjChoiceContent;
import org.systemsbiology.biofabric.util.UiUtil;

/****************************************************************************
**
** This does node clustering layout
*/

public class NodeClusterLayout extends NodeLayout {
  
	////////////////////////////////////////////////////////////////////////////
  //
  // PRIVATE INSTANCE MEMBERS
  //
  ////////////////////////////////////////////////////////////////////////////
	
  ////////////////////////////////////////////////////////////////////////////
  //
  // PUBLIC CONSTRUCTORS
  //
  ////////////////////////////////////////////////////////////////////////////

  /***************************************************************************
  **
  ** Constructor
  */

  public NodeClusterLayout() { 

  }

  ////////////////////////////////////////////////////////////////////////////
  //
  // PUBLIC METHODS
  //
  ////////////////////////////////////////////////////////////////////////////
  
  /***************************************************************************
  **
  */
  
   public List<NetNode> doNodeLayout(BuildData bd, 
							                       Params crParams,
							                       BTProgressMonitor monitor) throws AsynchExitRequestException { 
     
    //
    // Go through all the links. If a link source and target are both in the same cluster, we add the link to the cluster
    //
  	
  	BuildDataImpl rbd = (BuildDataImpl)bd;
  	ClusterParams params = (ClusterParams)crParams;
    
    TreeMap<String, BuildDataImpl> perClust = new TreeMap<String, BuildDataImpl>();
    HashMap<Tuple, List<NetLink>> interClust = new HashMap<Tuple, List<NetLink>>();
    HashSet<NetNode> interNodesOnly = new HashSet<NetNode>();
    
    HashMap<NetNode, Integer> fullNodeDegree = null;
    if (params.cLay == ClusterParams.ClustLayout.BREADTH_CONN_FIRST) {
      fullNodeDegree = new HashMap<NetNode, Integer>();
    } 	
    
    // Future...
    // BuildData.BuildMode intraLay = (params.cLay == ClusterParams.ClustLayout.SIMILAR) ? BuildData.BuildMode.CLUSTERED_LAYOUT
    //        																																					 : BuildData.BuildMode.DEFAULT_LAYOUT;
    
    BuildDataImpl.BuildMode intraLay = BuildDataImpl.BuildMode.DEFAULT_LAYOUT;    
      
    Iterator<NetLink> flit = rbd.getLinks().iterator();
    while (flit.hasNext()) {
      NetLink fl = flit.next();
      NetNode source = fl.getSrcNode();
      NetNode target = fl.getTrgNode();
      if (fullNodeDegree != null) {
      	Integer count = fullNodeDegree.get(source);
        count = (count == null) ? Integer.valueOf(1) : Integer.valueOf(count.intValue() + 1);
        fullNodeDegree.put(source, count);   
        count = fullNodeDegree.get(target);
        count = (count == null) ? Integer.valueOf(1) : Integer.valueOf(count.intValue() + 1);
        fullNodeDegree.put(target, count);      	       
      }    
      
      String srcClust = params.getClusterForNode(source);
      String trgClust = params.getClusterForNode(target);
      if (srcClust.equals(trgClust)) {
        BuildDataImpl rbdpc = perClust.get(srcClust);
        if (rbdpc == null) {
          rbdpc = new BuildDataImpl(new UniqueLabeller(), new HashSet<NetLink>(), new HashSet<NetNode>(),
    		                            new HashMap<NetNode, String>(), rbd.getColorGen(), intraLay);
          rbdpc.initAllNodesBogus(new HashSet<NetNode>());
          perClust.put(srcClust, rbdpc);
        }
        UiUtil.fixMePrintout("ADDING TO THE RETURNED SET");
        rbdpc.getLinks().add(fl);
        addClusterNode(rbdpc, source); 
        addClusterNode(rbdpc, target);
      } else {
        Tuple intup = new Tuple(srcClust, trgClust); // Tuples reordered so val1 < val2!
        List<NetLink> icfl = interClust.get(intup);
        if (icfl == null) {
          icfl = new ArrayList<NetLink>();
          interClust.put(intup, icfl);
        }
        icfl.add(fl);
        interNodesOnly.add(source);
        interNodesOnly.add(target);
      }
    }
    
    //
    // Need to deal with "clusters" of nodes that have no internal links!
    //  
    
    for (NetNode node : rbd.getAllNodes()) {
    	String clust = params.getClusterForNode(node);
    	BuildDataImpl rbdpc = perClust.get(clust);
    	if (rbdpc == null) {
    		rbdpc = new BuildDataImpl(new UniqueLabeller(),
    		                          new HashSet<NetLink>(), new HashSet<NetNode>(),
    		                          new HashMap<NetNode, String>(), rbd.getColorGen(), intraLay);
        rbdpc.initAllNodesBogus(new HashSet<NetNode>());
        perClust.put(clust, rbdpc);
    	}  		
      addClusterNode(rbdpc, node); 
    }
    
    String startClust = (params.startNode != null) ? params.getClusterForNode(params.startNode) : null;
    
    List<String> bfc;
    switch (params.order) {
    	case BREADTH:
        bfc = breadthFirstClustering(params, interClust.keySet(), startClust, monitor);
        break;
    	case LINK_SIZE:
    		bfc = clusterSizeOrder(perClust, false, startClust);
    		break;  		
    	case NODE_SIZE:
    	  bfc = clusterSizeOrder(perClust, true, startClust);
    		break;
    	case NAME:
    	  bfc = new ArrayList<String>(perClust.keySet());
    	  if (startClust != null) {
    	    bfc.remove(startClust);
    	    bfc.add(0, startClust);
        }
    	  break;
    	default:
    		throw new IllegalStateException();
    }
    
    Map<String, List<NetNode>> hubs = null;
    if (params.cLay == ClusterParams.ClustLayout.BREADTH_CONN_FIRST) {
      hubs = rankInterClustHubs(interClust, params, fullNodeDegree);	
    }

    ArrayList<NetNode> allTargets = new ArrayList<NetNode>();

    Iterator<String> pcit = bfc.iterator(); 
    while (pcit.hasNext()) {
    	String clustName = pcit.next();
    	BuildData pcrbd = perClust.get(clustName);
      if (pcrbd == null) {
      	continue;
      }
      List<NetNode> targets;
      if (intraLay == BuildDataImpl.BuildMode.DEFAULT_LAYOUT) {
        DefaultLayout dl = new DefaultLayout();
        List<NetNode> starts = (hubs == null) ? null : hubs.get(clustName);
        targets = dl.defaultNodeOrder(pcrbd.getLinks(), pcrbd.getSingletonNodes(), starts, monitor); 
      // Future enhancement:
      //} else if (intraLay == BuildData.BuildMode.CLUSTERED_LAYOUT) {      	
        //NodeSimilarityLayout.ClusterParams crp = new NodeSimilarityLayout.ClusterParams();
    	  //NodeSimilarityLayout nslLayout = new NodeSimilarityLayout();
    	  //pcrbd.existingIDOrder = new ArrayList<NetNode>(pcrbd.allNodeIDs);
        // The idea is to use the NodeSimilarityLayout on each cluster....
        //targets = nslLayout.doClusteredLayoutOrder(pcrbd, crp, monitor, startFrac, endFrac);
    	  //targets = new ArrayList<NetNode>();
      } else {
      	throw new IllegalStateException();
      }
      allTargets.addAll(targets);
    }
    interNodesOnly.removeAll(allTargets);
    allTargets.addAll(interNodesOnly);
    
    //
    // We are done getting nodes ordered. Install this:
    //
    
    installNodeOrder(allTargets, rbd, monitor);
    
    
     UiUtil.fixMePrintout("Inter cluster shadows are messed up and on the right");
     // Can have ALL links from cluster 1 to all clusters right after cluster 1, and
     // all links to cluster 2 right before cluster 2.
     
    
    //
    // We lay out the edges using default layout. Then, if we want edges to be positioned
    // between clusters, we move them:
    //
   
    (new DefaultEdgeLayout()).layoutEdges(rbd, monitor);
   
    UiUtil.fixMePrintout("How does this interact with network-wide link groups??");
    if (params.iLink == ClusterParams.InterLink.BETWEEN) {
    	int origNum = rbd.getLinkOrder().size();
    	TreeMap<Integer, NetLink> newOrder = new TreeMap<Integer, NetLink>();
    	HashMap<String, List<NetLink>> holdEm = new HashMap<String, List<NetLink>>();
    	Iterator<NetLink> ksit = rbd.getLinkOrder().values().iterator();
    	int colCount = 0;
    	String currClust = null;
    	boolean drainTime = false;
    	boolean interClustLink = false;
    	while (ksit.hasNext()) {
    		NetLink daLink = ksit.next();
    		String srcClust = params.getClusterForNode(daLink.getSrcNode());
        String trgClust = params.getClusterForNode(daLink.getTrgNode());
        if (srcClust.equals(trgClust)) {
        	if ((currClust != null) && !currClust.equals(srcClust)) {
        		drainTime = true;
        	}
        	currClust = srcClust;
        	interClustLink = false;
        } else {
        	interClustLink = true;
        }
        
        if (drainTime) {
        	List<NetLink> toDrain = holdEm.get(currClust);
        	if (toDrain != null) {
	        	for (NetLink ihe : toDrain) {
	    	  		newOrder.put(Integer.valueOf(colCount++), ihe);   	  		
	    	  	}
        	}
        	holdEm.remove(currClust);
    	  	drainTime = false; 
    	  	newOrder.put(Integer.valueOf(colCount++), daLink);
        } else if (!interClustLink) {
        	currClust = srcClust;
    		  newOrder.put(Integer.valueOf(colCount++), daLink);
    	  } else {
    	  	String otherClust = srcClust.equals(currClust) ? trgClust : srcClust;
    	  	List<NetLink> toDefer = holdEm.get(otherClust);
    	  	if (toDefer == null) {
    	  		toDefer = new ArrayList<NetLink>();
    	  		holdEm.put(otherClust, toDefer);
    	  	}
    	  	toDefer.add(daLink);  	  	
    	  }
    	}
    	
    	 UiUtil.fixMePrintout("Cluster drain order is not correct");
    	for (String daClust : holdEm.keySet()) {
	      List<NetLink> toDrain = holdEm.get(daClust);
	    	if (toDrain != null) {
	      	for (NetLink ihe : toDrain) {
	  	  		newOrder.put(Integer.valueOf(colCount++), ihe);   	  		
	  	  	}
	    	}
    	}
    	if (newOrder.size() != origNum) {
    		throw new IllegalStateException();
    	}
    	rbd.setLinkOrder(newOrder);
    }
    
    //
    // Did we ask to save cluster assignment? If so, do it now. The NodeInfo has a field to store this
    // information
    //
    
    if (params.saveAssign) {
    	rbd.clustAssign = params.getClusterAssign(); 
    }
    
    AnnotationSet nAnnots = generateNodeAnnotations(rbd, params);
    rbd.setNodeAnnotations(nAnnots);
      
    Map<Boolean, AnnotationSet> lAnnots = generateLinkAnnotations(rbd, params);
    rbd.setLinkAnnotations(lAnnots);

    return (allTargets);
  }
   
   
  /***************************************************************************
  **
  ** Generate node annotations to tag each cluster
  */
    
  private AnnotationSet generateNodeAnnotations(BuildData rbd, ClusterParams params) {
    
    AnnotationSet retval = PluginSupportFactory.buildAnnotationSet();
    
    TreeMap<Integer, NetNode> invert = new TreeMap<Integer, NetNode>();
    
    Map<NetNode, Integer> nod = rbd.getNodeOrder(); 
    
    for (NetNode node : nod.keySet()) {
      invert.put(nod.get(node), node);
    }
     
    String currClust = null;
    Integer startRow = null;
    Integer lastKey = invert.lastKey();
    for (Integer row : invert.keySet()) {
      NetNode node = invert.get(row);
      String clust = params.getClusterForNode(node);
      if (currClust == null) {
        currClust = clust;
        startRow = row;
        if (row.equals(lastKey)) {
          Annot annot = PluginSupportFactory.buildAnnotation(currClust, startRow.intValue(), row.intValue(), 0, null);
          retval.addAnnot(annot);
          break;
        }
        continue;
      }
      if (currClust.equals(clust)) {
        if (row.equals(lastKey)) {
          Annot annot = PluginSupportFactory.buildAnnotation(currClust, startRow.intValue(), row.intValue(), 0, null);
          retval.addAnnot(annot);
          break;
        }
        continue;
      } else { 
        // We have just entered a new cluster
        Annot annot = PluginSupportFactory.buildAnnotation(currClust, startRow.intValue(), row.intValue() - 1, 0, null);

        retval.addAnnot(annot);
        startRow = row;
        currClust = clust;
        if (row.equals(lastKey)) {
          annot = PluginSupportFactory.buildAnnotation(currClust, startRow.intValue(), row.intValue(), 0, null);

          retval.addAnnot(annot);
          break;
        }
      }
    }
    
    return (retval);
  }
  
  /***************************************************************************
  **
  ** Generate link annotations to tag each cluster and intercluster links
  */
    
  private Map<Boolean, AnnotationSet> generateLinkAnnotations(BuildData rbd, ClusterParams params) { 
  	HashMap<Boolean, AnnotationSet> retval = new HashMap<Boolean, AnnotationSet>();
    
  	SortedMap<Integer,NetLink> lod = rbd.getLinkOrder();
  	
  	List<NetLink> noShadows = new ArrayList<NetLink>();
  	List<NetLink> withShadows = new ArrayList<NetLink>();
  	for (Integer col : lod.keySet()) {
  		NetLink fl = lod.get(col);
  		withShadows.add(fl);
  		if (!fl.isShadow()) {
  			noShadows.add(fl);
  		}
  	}

  	retval.put(Boolean.FALSE, generateLinkAnnotationsForSet(rbd, noShadows, params));
  	UiUtil.fixMePrintout("Messedup link annotations with shadows");
  	retval.put(Boolean.TRUE, generateLinkAnnotationsForSet(rbd, withShadows, params));
  	
    return (retval);
  }
   
  /***************************************************************************
  **
  ** Generate link annotations to tag each cluster and intercluster links
  */
    
  private AnnotationSet generateLinkAnnotationsForSet(BuildData rbd, List<NetLink> linkList, ClusterParams params) { 
    
  	HashMap<String, MinMax> clustRanges = new HashMap<String, MinMax>();
  	
  	//
  	// If the interlinks are inline, we have nothing to do except annotate the clusters themselves. Cluster bounds
  	// are set by the cluster of the top node of regular links. If shadows are shown, we include the bottom node 
  	// of shadow links.
  	//
  	
  	Map<NetNode, Integer> nod = rbd.getNodeOrder();
  	
  	
  	if (params.iLink == ClusterParams.InterLink.INLINE) {	
	  	for (int i = 0; i < linkList.size() ; i++) {
	  		NetLink fl = linkList.get(i);
	  		boolean isShad = fl.isShadow();
  		  NetNode srcNode = fl.getSrcNode();
  		  NetNode trgNode = fl.getTrgNode();
  		  String srcClust = params.getClusterForNode(srcNode);
        String trgClust = params.getClusterForNode(trgNode);
	      int srcRow = nod.get(srcNode).intValue();
	      int trgRow = nod.get(trgNode).intValue();
	      String useClust;
	      if (srcRow <= trgRow) {
	      	useClust = (isShad) ? trgClust : srcClust;
	      } else {
	      	useClust = (isShad) ? srcClust : trgClust;
	      }
  			MinMax mmc = clustRanges.get(useClust);
  			if (mmc == null) {
  				mmc = new MinMax(i);
  				clustRanges.put(useClust, mmc);
  			}
  			mmc.update(i);
	  	}
  	} else {
  		for (int i = 0; i < linkList.size() ; i++) {
	  		NetLink fl = linkList.get(i);
	  		boolean isShad = fl.isShadow();
  		  NetNode srcNode = fl.getSrcNode();
  		  NetNode trgNode = fl.getTrgNode();
  		  String srcClust = params.getClusterForNode(srcNode);
        String trgClust = params.getClusterForNode(trgNode);
        if (!srcClust.equals(trgClust)) {
        	continue;
        }
	      //int srcRow = nod.get(srcNode).intValue();
	      //int trgRow = nod.get(trgNode).intValue();
	      String useClust = srcClust;
	      //if (srcRow <= trgRow) {
	      //	useClust = (isShad) ? trgClust : srcClust;
	      //} else {
	      //	useClust = (isShad) ? srcClust : trgClust;
	      //}
  			MinMax mmc = clustRanges.get(useClust);
  			if (mmc == null) {
  				mmc = new MinMax(i);
  				clustRanges.put(useClust, mmc);
  			}
  			mmc.update(i);
	  	}
  	}
  	
  	
  	/*	} else {
  			String combo = (srcClust.compareTo(trgClust) < 0) ? srcClust + "-" + trgClust : trgClust + "-" + srcClust;
   			MinMax mmc = clustRanges.get(combo);
  			if (mmc == null) {
  				mmc = new MinMax(i);
  				clustRanges.put(combo, mmc);
  			}
  			mmc.update(i); */
  	//	}
  //	}
  	
  	AnnotationSet afns = PluginSupportFactory.buildAnnotationSet();
  	TreeMap<Integer, String> ord = new TreeMap<Integer, String>();
  	for (String aName : clustRanges.keySet()) {
  		MinMax mm = clustRanges.get(aName);
  		ord.put(Integer.valueOf(mm.min), aName);		
  	}
  	
    for (String aName : ord.values()) {
  		MinMax mm = clustRanges.get(aName);
  		Annot annot = PluginSupportFactory.buildAnnotation(aName, mm.min, mm.max, 0, null);
  		afns.addAnnot(annot);
  	}
  
    return (afns);
  }
  
  /***************************************************************************
  **
  ** Helper
  */
  
  private void addClusterNode(BuildDataImpl rbd, NetNode nid) { 
    if (!rbd.getAllNodes().contains(nid)) {
    	boolean ok = rbd.idGen.addExistingLabel(nid.getNID().getNID().getInternal());
    	if (!ok) {
    		throw new IllegalStateException();
    	}
      rbd.addToAllNodesBogus(nid);
    }
    return;
  }

  /***************************************************************************
  **
  ** Count map maintenance
  */
  
  private void addToCount(Map<String, Map<NetNode, Integer>> countMap, String clust, NetNode node) { 
 	
    Map<NetNode, Integer> perClust = countMap.get(clust);
    if (perClust == null) {
    	perClust = new HashMap<NetNode, Integer>();
    	countMap.put(clust, perClust);
    }
    Integer count = perClust.get(node);
    count = (count == null) ? Integer.valueOf(1) : Integer.valueOf(count.intValue() + 1);
    perClust.put(node, count);
    return;
  }
   
   
  /***************************************************************************
  **
  ** The nodes that are responsible for inter-cluster links may be almost no-shows
  ** in the intra-cluster link competition. Find those guys so we can put them first
  ** in the pile.
  */
  
  private Map<String, List<NetNode>> rankInterClustHubs(Map<Tuple, List<NetLink>> interLinks, 
  		                                                       ClusterParams params, 
  		                                                       Map<NetNode, Integer> fullNodeDegree) { 
 	
  	// Map(ClusterID -> Map(NodeID, InterClustDegree)):
  	Map<String, Map<NetNode, Integer>> preRetval = new HashMap<String, Map<NetNode, Integer>>();
  	for (Tuple tup : interLinks.keySet()) {
  	  List<NetLink> forTup = interLinks.get(tup);
  	  for (NetLink link : forTup) {
  	  	// Tup order is NOT src, trg, but lo, hi:
  	    addToCount(preRetval, params.getClusterForNode(link.getSrcNode()), link.getSrcNode());
  	    addToCount(preRetval, params.getClusterForNode(link.getTrgNode()), link.getTrgNode());
  	  }
  	}
  	
  	// Map(ClusterID -> List(NodeID)):
    Map<String, List<NetNode>> retval = new HashMap<String, List<NetNode>>();  
    for (String clust: preRetval.keySet()) {
    	Map<NetNode, Integer> degMap = preRetval.get(clust);
    	// Map(SortedInterClustDegree->Map(SortedFullNodeDegree, Set(SortedNodeName))):
    	TreeMap<Integer, SortedMap<Integer, SortedSet<NetNode>>> invert = 
      	new TreeMap<Integer, SortedMap<Integer, SortedSet<NetNode>>>(Collections.reverseOrder());
    	for (NetNode node : degMap.keySet()) {
    		Integer interClustDeg = degMap.get(node);
    		SortedMap<Integer, SortedSet<NetNode>> perInterDeg = invert.get(interClustDeg);
    		if (perInterDeg == null) {
    			perInterDeg = new TreeMap<Integer, SortedSet<NetNode>>(Collections.reverseOrder());
    			invert.put(interClustDeg, perInterDeg);
    		}
    		Integer fnDeg = fullNodeDegree.get(node);
    		SortedSet<NetNode> perFullDeg = perInterDeg.get(fnDeg);
    		if (perFullDeg == null) {
    			perFullDeg = new TreeSet<NetNode>();
    			perInterDeg.put(fnDeg, perFullDeg);
    		}
    		perFullDeg.add(node);
    	}
    	List<NetNode> flat = new ArrayList<NetNode>();
      for (SortedMap<Integer, SortedSet<NetNode>> perInterDeg : invert.values()) {
      	for (SortedSet<NetNode> perFull : perInterDeg.values()) {
      	  flat.addAll(perFull);
        }
      }
    	retval.put(clust, flat);
    }
  	return (retval);
  }
  
  /***************************************************************************
  **
  */
  
  private List<String> clusterSizeOrder(TreeMap<String, BuildDataImpl> perClust, 
  		                                  boolean nodeFirst, String startClust) { 
  	
  	TreeMap<Integer, SortedMap<Integer, SortedSet<String>>> preRet = 
  		new TreeMap<Integer, SortedMap<Integer, SortedSet<String>>>(Collections.reverseOrder());
  	for (String clustName : perClust.keySet()) {
  	  BuildData rbdpc = perClust.get(clustName);
  	  Integer size1 = Integer.valueOf((nodeFirst) ? rbdpc.getAllNodes().size() : rbdpc.getLinks().size());
  	  SortedMap<Integer, SortedSet<String>> forSize1 = preRet.get(size1);
  	  if (forSize1 == null) {
  	  	forSize1 = new TreeMap<Integer, SortedSet<String>>(Collections.reverseOrder());
  	  	preRet.put(size1, forSize1);
  	  }
  	  Integer size2 = Integer.valueOf((nodeFirst) ? rbdpc.getLinks().size() : rbdpc.getAllNodes().size());
  	  SortedSet<String> forSize2 = forSize1.get(size2);
  	  if (forSize2 == null) {
  	  	forSize2 = new TreeSet<String>();
  	  	forSize1.put(size2, forSize2);
  	  }
  	  forSize2.add(clustName); 	
  	}
  	
  	List<String> flat = new ArrayList<String>();
    for (SortedMap<Integer, SortedSet<String>> perSize1 : preRet.values()) {
    	for (SortedSet<String> perSize2 : perSize1.values()) {
    	  flat.addAll(perSize2);
      }
    }
    
    if (startClust != null) {
    	flat.remove(startClust);
    	flat.add(0, startClust);
    }
    
    return (flat);
  }

  /***************************************************************************
  **
  */
  
  private List<String> breadthFirstClustering(ClusterParams params, Set<Tuple> iclinks, String startClust,
                                              BTProgressMonitor monitor) throws AsynchExitRequestException {
    
  	UniqueLabeller uLab = new UniqueLabeller();
  	Map<String, NetNode> fakeNodes = new HashMap<String, NetNode>();
  	
  	for (String clu : params.getClusters()) {
  	  NID pnid = uLab.getNextOID();
  	  fakeNodes.put(clu, new FabricNode(pnid, clu));	
  	}
  	
  	Set<NetLink> links = new HashSet<NetLink>();
  	for (Tuple tup : iclinks) {
  		NetLink pseudo = new FabricLink(fakeNodes.get(tup.getVal1()), fakeNodes.get(tup.getVal2()), "ic", Boolean.valueOf(false), Boolean.valueOf(false));
  		links.add(pseudo);
  		pseudo = new FabricLink(fakeNodes.get(tup.getVal2()), fakeNodes.get(tup.getVal1()), "ic", Boolean.valueOf(false), Boolean.valueOf(false));
  		links.add(pseudo);
  	}
  	
  	List<NetNode> useRoots = null;
  	if (startClust != null) {
  		useRoots = new ArrayList<NetNode>();
  		useRoots.add(fakeNodes.get(startClust));
  	}
    GraphSearcher gs = new GraphSearcher(new HashSet<NetNode>(fakeNodes.values()), links);
    List<GraphSearcher.QueueEntry> qes = gs.breadthSearch(useRoots, true, monitor);
    ArrayList<String> retval = new ArrayList<String>(); 
    for (GraphSearcher.QueueEntry aqe : qes) {
      retval.add(aqe.name.getName());
  	}
    return (retval);  
  }
   
  /***************************************************************************
  **
  ** For passing around layout params
  */  
  
  public static class ClusterParams implements NodeLayout.Params {
        
    public enum Source {STORED, FILE}; // , PLUGIN}; Future enhancement
    
    public enum Order {NAME, NODE_SIZE, LINK_SIZE, BREADTH};
    
    public enum InterLink {INLINE, BETWEEN};
    
    public enum ClustLayout {BREADTH_CONN_FIRST, BREADTH} // , SIMILAR}; Future enhancement
    
          
    public Source source;
    public Order order;
    public InterLink iLink;
    public ClustLayout cLay;
    public NetNode startNode;
    public boolean saveAssign;
    private Map<NetNode, String> nodeClusters_;

    public ClusterParams(Source source, Order order, InterLink iLink, ClustLayout cLay, String startNode, 
    		                 Map<AttributeKey, String> nodeClusterAttributes, 
    		                 Map<String, NetNode> nodes, boolean saveAssign) {
    	
    	if (nodeClusterAttributes != null) {
    	  nodeClusters_ = new HashMap<NetNode, String>();
        for (AttributeKey key : nodeClusterAttributes.keySet()) {
          NetNode nodeID = nodes.get(DataUtil.normKey(((AttributeLoader.StringKey)key).key));
          nodeClusters_.put(nodeID, nodeClusterAttributes.get(key));
        }
      }
      this.source = source;
      this.order = order;
      this.saveAssign = saveAssign;
    }
 
    public ClusterParams(boolean stored) {
      this.source = (stored) ? Source.STORED : Source.FILE;
      this.order = Order.BREADTH;
      this.iLink = InterLink.BETWEEN;
      this.cLay = ClustLayout.BREADTH_CONN_FIRST;
      this.startNode = null;
      this.saveAssign = true;
    }
    
    public boolean needsFile() {
    	return (source.equals(Source.FILE));
    }

    public void install(Map<AttributeKey, String> nodeClusterAttributes, Map<String, NetNode> nodes) {   	
    	if (nodeClusterAttributes != null) {
    	  nodeClusters_ = new HashMap<NetNode, String>();
        for (AttributeKey key : nodeClusterAttributes.keySet()) {
        	NetNode nodeID = nodes.get(DataUtil.normKey(((AttributeLoader.StringKey)key).key));    	
          nodeClusters_.put(nodeID, nodeClusterAttributes.get(key));
        }
      }
    }
    
    public void assign(Map<NetNode, String> nodeClusterAssign) {   	
    	nodeClusters_ = nodeClusterAssign;
      return;
    }
 
    public String getClusterForNode(NetNode nodeID) {   	
    	return (nodeClusters_.get(nodeID));
    }  
    
    public Map<NetNode, String> getClusterAssign() {   	
    	return (nodeClusters_);
    }  
    
    
    public Collection<String> getClusters() {   	
    	return (nodeClusters_.values());
    }    
    
    public static Vector<TrueObjChoiceContent<Source>> getSourceChoices(boolean stored) {
      ResourceManager rMan = ResourceManager.getManager();
      Vector<TrueObjChoiceContent<Source>> retval = new Vector<TrueObjChoiceContent<Source>>();
      if (stored) {
      	retval.add(new TrueObjChoiceContent<Source>(rMan.getString("nodeClusterParams.stored"), Source.STORED));
      }
      retval.add(new TrueObjChoiceContent<Source>(rMan.getString("nodeClusterParams.file"), Source.FILE));
      // Future enhancement...
      // retval.add(new TrueObjChoiceContent<Source>(rMan.getString("nodeClusterParams.plugin"), Source.PLUGIN));   
      return (retval);
    }
    
    public static Vector<TrueObjChoiceContent<InterLink>> getILinkChoices() {
      ResourceManager rMan = ResourceManager.getManager();
      Vector<TrueObjChoiceContent<InterLink>> retval = new Vector<TrueObjChoiceContent<InterLink>>();
      retval.add(new TrueObjChoiceContent<InterLink>(rMan.getString("nodeClusterParams.between"), InterLink.BETWEEN));
      retval.add(new TrueObjChoiceContent<InterLink>(rMan.getString("nodeClusterParams.inline"), InterLink.INLINE));   
      return (retval);
    }
    
    public static Vector<TrueObjChoiceContent<ClustLayout>> getClustLayoutChoices() {
      ResourceManager rMan = ResourceManager.getManager();
      Vector<TrueObjChoiceContent<ClustLayout>> retval = new Vector<TrueObjChoiceContent<ClustLayout>>();
      retval.add(new TrueObjChoiceContent<ClustLayout>(rMan.getString("nodeClusterParams.breadthConnFirst"), ClustLayout.BREADTH_CONN_FIRST));
      retval.add(new TrueObjChoiceContent<ClustLayout>(rMan.getString("nodeClusterParams.breadth"), ClustLayout.BREADTH));
      // Future
      // retval.add(new TrueObjChoiceContent<ClustLayout>(rMan.getString("nodeClusterParams.similar"), ClustLayout.SIMILAR));   
      return (retval);
    }
        
    public static Vector<TrueObjChoiceContent<Order>> getOrderChoices() {
      ResourceManager rMan = ResourceManager.getManager();
      Vector<TrueObjChoiceContent<Order>> retval = new Vector<TrueObjChoiceContent<Order>>();
      retval.add(new TrueObjChoiceContent<Order>(rMan.getString("nodeClusterParams.breadthOrder"), Order.BREADTH));   
      retval.add(new TrueObjChoiceContent<Order>(rMan.getString("nodeClusterParams.name"), Order.NAME));
      retval.add(new TrueObjChoiceContent<Order>(rMan.getString("nodeClusterParams.nodeSize"), Order.NODE_SIZE));   
      retval.add(new TrueObjChoiceContent<Order>(rMan.getString("nodeClusterParams.linkSize"), Order.LINK_SIZE));   
      return (retval);
    }     
  }    

  /***************************************************************************
  **
  ** Class for representing a tuple
  */ 
   
  public static class Tuple implements Cloneable, Comparable<Tuple> {
  
    private String val1_;
    private String val2_;
  
    public Tuple(String val1, String val2) {
      if ((val1 == null) || (val2 == null)) {
        throw new IllegalArgumentException();
      }
      if (val1.compareTo(val2) < 0) {
        val1_ = val1;
        val2_ = val2;
      } else {
        val1_ = val2;
        val2_ = val1;
      }
    }  
   
    @Override
    public Tuple clone() {
      try {
        Tuple newTup = (Tuple)super.clone();
        return (newTup);
      } catch (CloneNotSupportedException ex) {
        throw new IllegalStateException();     
      }    
    }
    
    @Override
    public int hashCode() {
      return (val1_.hashCode() + val2_.hashCode());
    }
    
    public String getVal1() {
      return (val1_);
    }
    
    public String getVal2() {
      return (val2_);
    }  
    
    @Override
    public boolean equals(Object other) {
      if (other == null) {
        return (false);
      }
      if (other == this) {
        return (true);
      }
      if (!(other instanceof Tuple)) {
        return (false);
      }
      Tuple otherT = (Tuple)other;
      return ((this.val1_.equals(otherT.val1_)) && (this.val2_.equals(otherT.val2_)));
    }
 
    @Override
    public String toString() {
      return ("Tuple: (" + val1_ + ", " + val2_ + ")");
    }
    
    public int compareTo(Tuple otherTup) {
      int val1Diff = this.val1_.compareTo(otherTup.val1_);
      if (val1Diff != 0) {
        return (val1Diff);
      }      
      return (this.val2_.compareTo(otherTup.val2_));
    }   
  }
}
```
