## Supplementary material for "BioFabric Visualization of Network Alignments": BioFabric Source Code: NodeSimilarityLayout.java

```
/*
**    Copyright (C) 2003-2018 Institute for Systems Biology 
**                            Seattle, Washington, USA. 
**
**    This library is free software; you can redistribute it and/or
**    modify it under the terms of the GNU Lesser General Public
**    License as published by the Free Software Foundation; either
**    version 2.1 of the License, or (at your option) any later version.
**
**    This library is distributed in the hope that it will be useful,
**    but WITHOUT ANY WARRANTY; without even the implied warranty of
**    MERCHANTABILITY or FITNESS FOR A PARTICULAR PURPOSE.  See the GNU
**    Lesser General Public License for more details.
**
**    You should have received a copy of the GNU Lesser General Public
**    License along with this library; if not, write to the Free Software
**    Foundation, Inc., 59 Temple Place, Suite 330, Boston, MA  02111-1307  USA
*/

package org.systemsbiology.biofabric.layouts;

import java.util.ArrayList;
import java.util.Collections;
import java.util.HashMap;
import java.util.HashSet;
import java.util.Iterator;
import java.util.List;
import java.util.Map;
import java.util.Set;
import java.util.SortedMap;
import java.util.SortedSet;
import java.util.TreeMap;
import java.util.TreeSet;
import java.util.Vector;

import org.systemsbiology.biofabric.analysis.Link;
import org.systemsbiology.biofabric.api.io.BuildData;
import org.systemsbiology.biofabric.api.layout.NodeLayout;
import org.systemsbiology.biofabric.api.model.NetLink;
import org.systemsbiology.biofabric.api.model.NetNode;
import org.systemsbiology.biofabric.api.util.MinMax;
import org.systemsbiology.biofabric.api.worker.AsynchExitRequestException;
import org.systemsbiology.biofabric.api.worker.BTProgressMonitor;
import org.systemsbiology.biofabric.api.worker.LoopReporter;
import org.systemsbiology.biofabric.io.BuildDataImpl;
import org.systemsbiology.biofabric.model.BioFabricNetwork;
import org.systemsbiology.biofabric.util.ChoiceContent;
import org.systemsbiology.biofabric.util.DataUtil;
import org.systemsbiology.biofabric.util.DoubMinMax;
import org.systemsbiology.biofabric.util.ResourceManager;
import org.systemsbiology.biofabric.util.UiUtil;

/****************************************************************************
**
** This does the node similarity layout
*/

public class NodeSimilarityLayout extends NodeLayout {
  
  ////////////////////////////////////////////////////////////////////////////
  //
  // PRIVATE CONSTANTS
  //
  //////////////////////////////////////////////////////////////////////////// 

  ////////////////////////////////////////////////////////////////////////////
  //
  // PUBLIC CONSTANTS
  //
  //////////////////////////////////////////////////////////////////////////// 
  
  ////////////////////////////////////////////////////////////////////////////
  //
  // PRIVATE INSTANCE MEMBERS
  //
  ////////////////////////////////////////////////////////////////////////////
  
  ////////////////////////////////////////////////////////////////////////////
  //
  // PUBLIC CONSTRUCTORS
  //
  ////////////////////////////////////////////////////////////////////////////

  /***************************************************************************
  **
  ** Constructor
  */

  public NodeSimilarityLayout() {
  }

  ////////////////////////////////////////////////////////////////////////////
  //
  // PUBLIC METHODS
  //
  ////////////////////////////////////////////////////////////////////////////

  /***************************************************************************
  ** 
  ** Node layout
  */

  public List<NetNode> doNodeLayout(BuildData rbd, 
							                      NodeLayout.Params params,
							                      BTProgressMonitor monitor) throws AsynchExitRequestException { 
  	if (params instanceof NodeSimilarityLayout.ResortParams) {
  		return (doReorderLayout(rbd, (NodeSimilarityLayout.ResortParams)params, monitor));		
  	} else if (params instanceof NodeSimilarityLayout.ClusterParams) {
  		return (doClusteredLayout(rbd, (NodeSimilarityLayout.ClusterParams)params, monitor));	
  	} else {
  		throw new IllegalArgumentException();
  	}
  }

  /***************************************************************************
  ** 
  ** Move nodes to match shapes
  */

  private List<NetNode> doReorderLayout(BuildData rbd, 
                                        NodeSimilarityLayout.ResortParams rp,
                                        BTProgressMonitor monitor) throws AsynchExitRequestException { 

  	HashMap<NetNode, Integer> targToRow = new HashMap<NetNode, Integer>();
    SortedMap<Integer, SortedSet<Integer>> connVecs = getConnectionVectors(rbd, targToRow, monitor);
     
    List<Integer> ordered = new ArrayList<Integer>();
    int numRows = ((BuildDataImpl)rbd).getRowCountForNetwork();
    for (int i = 0; i < numRows; i++) {
      ordered.add(Integer.valueOf(i));
    }
    
    double currStart = 0.0;
    double inc = 1.0 / rp.passCount;
    double currEnd = currStart + inc;
    
    TreeMap<Integer, Double> rankings = new TreeMap<Integer, Double>();
    NodeSimilarityLayout.ClusterPrep cprep = setupForResort(connVecs, ordered, rankings);
    Double lastRank = rankings.get(rankings.lastKey());
    
    for (int i = 0; i < rp.passCount; i++) {
      monitor.updateRankings(rankings);
      List<Integer> nextOrdered = resort(cprep, monitor, currStart, currEnd);
      currStart = currEnd;
      currEnd = currStart + inc;
      cprep = setupForResort(connVecs, nextOrdered, rankings);
      Integer lastKey = rankings.lastKey();
      Double nowRank = rankings.get(lastKey);
      if (rp.terminateAtIncrease) {
        if (lastRank.doubleValue() < nowRank.doubleValue()) {
          rankings.remove(lastKey);
          break;
        }
      }
      ordered = nextOrdered;
      lastRank = nowRank;
    }
    
    monitor.updateRankings(rankings);
    List<NetNode> retval = convertOrderToMap(rbd, ordered);  
    return (retval);
  }
  
  /***************************************************************************
  **
  ** Clustered Layout guts
  */   

  private List<NetNode> doClusteredLayout(BuildData rbd, 
                                          NodeSimilarityLayout.ClusterParams params,
                                          BTProgressMonitor monitor) throws AsynchExitRequestException {   
 
    List<Integer> ordered = doClusteredLayoutOrder(rbd, params, monitor);
    List<NetNode> retval = convertOrderToMap(rbd, ordered);
    return (retval);
  }
  
  /***************************************************************************
  **
  ** Clustered Layout Ordering only
  */   

  private List<Integer> doClusteredLayoutOrder(BuildData rbd, 
							                                 NodeSimilarityLayout.ClusterParams cp,
							                                 BTProgressMonitor monitor) throws AsynchExitRequestException {   
 
    HashMap<NetNode, Integer> targToRow = new HashMap<NetNode, Integer>();
    SortedMap<Integer, SortedSet<Integer>> connVecs = getConnectionVectors(rbd, targToRow, monitor);

    TreeMap<Double, SortedSet<Link>> dists = new TreeMap<Double, SortedSet<Link>>(Collections.reverseOrder());
    HashMap<Integer, Integer> degMag = new HashMap<Integer, Integer>();
    
    Integer highestDegree = (cp.distanceMethod == NodeSimilarityLayout.ClusterParams.COSINES) ?
                             getCosines(connVecs, dists, degMag, rbd, targToRow, monitor) :
                             getJaccard(connVecs, dists, degMag, rbd, targToRow, monitor);
    
    ArrayList<Link> linkTrace = new ArrayList<Link>();
    ArrayList<Integer> jumpLog = new ArrayList<Integer>();
    
    
    LoopReporter lr = new LoopReporter(targToRow.size(), 20, monitor, 0.0, 1.0, "progress.preparingToChain"); 
    HashMap<Integer, NetNode> rowToTarg = new HashMap<Integer, NetNode>();
    for (NetNode targ : targToRow.keySet()) {
    	lr.report();
    	rowToTarg.put(targToRow.get(targ), targ);
    }
    lr.finish();
    
    List<Integer> ordered = orderByDistanceChained(rowToTarg, highestDegree, dists, 
                                                   degMag, linkTrace, cp.chainLength, 
                                                   cp.tolerance, jumpLog, monitor);
    return (ordered);
  }

  ////////////////////////////////////////////////////////////////////////////
  //
  // PRIVATE METHODS
  //
  ////////////////////////////////////////////////////////////////////////////
  
  /***************************************************************************
  ** 
  ** Get connection vectors
  */

  private SortedMap<Integer, SortedSet<Integer>> getConnectionVectors(BuildData rbd, 
  		                                                                Map<NetNode, Integer> targToRow,
  		                                                                BTProgressMonitor monitor) throws AsynchExitRequestException {
    
    Iterator<NetNode> rtit = ((BuildDataImpl)rbd).getExistingIDOrder().iterator();
    int count = 0;
    while (rtit.hasNext()) {
      NetNode node = rtit.next();
      targToRow.put(node, Integer.valueOf(count++));
    }
    
    LoopReporter lr = new LoopReporter(rbd.getLinks().size(), 20, monitor, 0.0, 1.0, "progress.getConnectionVectors"); 
    
    TreeMap<Integer, SortedSet<Integer>> retval = new TreeMap<Integer, SortedSet<Integer>>();
    Iterator<NetLink> ldit = rbd.getLinks().iterator();
    while (ldit.hasNext()) {
      NetLink fl = ldit.next();
      lr.report();
      if (fl.isShadow()) {
      	continue;
      }
      NetNode srcName = fl.getSrcNode();
      Integer srcRow = targToRow.get(srcName);
      NetNode trgName = fl.getTrgNode();
      Integer trgRow = targToRow.get(trgName);

      SortedSet<Integer> forRetval = retval.get(srcRow);
      if (forRetval == null) {
        forRetval = new TreeSet<Integer>();
        retval.put(srcRow, forRetval);
      }
      forRetval.add(trgRow);

      forRetval = retval.get(trgRow);
      if (forRetval == null) {
        forRetval = new TreeSet<Integer>();
        retval.put(trgRow, forRetval);
      }
      forRetval.add(srcRow);
    }
    lr.finish();
    return (retval);
  }
  
  /***************************************************************************
  ** 
  ** Get cosines.
  */

  private Integer getCosines(SortedMap<Integer, SortedSet<Integer>> connVec, 
                             SortedMap<Double, SortedSet<Link>> retval, 
                             Map<Integer, Integer> connMag, 
                             BuildData rbd, 
                             Map<NetNode, Integer> targToRow,
                             BTProgressMonitor monitor) throws AsynchExitRequestException {
    int rowCount = rbd.getLinks().size();
    Integer[] icache = new Integer[rowCount];
    String[] scache = new String[rowCount];
    for (int i = 0; i < rowCount; i++) {
      icache[i] = Integer.valueOf(i);
      scache[i] = icache[i].toString();
    }
    Integer highestDegree = null;
    int biggestMag = Integer.MIN_VALUE;
    HashSet<Integer> intersect = new HashSet<Integer>();
    
    LoopReporter lr = new LoopReporter(rbd.getLinks().size(), 20, monitor, 0.0, 1.0, "progress.getCosines");  
    
    Iterator<NetLink> ldit = rbd.getLinks().iterator();
    while (ldit.hasNext()) {
      NetLink fl = ldit.next();
      lr.report();
      if (fl.isShadow()) {
      	continue;
      }
      NetNode srcName = fl.getSrcNode();
      Integer srcRow = targToRow.get(srcName);
      NetNode trgName = fl.getTrgNode();
      Integer trgRow = targToRow.get(trgName);
      SortedSet<Integer> srcVec = connVec.get(srcRow);     
      int srcSize = srcVec.size();
      if (srcSize > biggestMag) {
        biggestMag = srcSize;
        highestDegree = srcRow;
      }
      SortedSet<Integer> trgVec = connVec.get(trgRow);     
      int trgSize = trgVec.size();
      if (trgSize > biggestMag) {
        biggestMag = trgSize;
        highestDegree = trgRow;
      }
      connMag.put(icache[srcRow.intValue()], icache[srcSize]);
      connMag.put(icache[trgRow.intValue()], icache[trgSize]);
     
      double sqrs = Math.sqrt(srcSize);
      double sqrt = Math.sqrt(trgSize);
      intersect.clear();
      intersect.addAll(srcVec);
      intersect.retainAll(trgVec);
      
      // Since normalized and equal, this is the sum of products of (1 / sqrs) * (1 / sqrt),
      // where we sum over intersect identical terms.  Note that the "vectors" do not include
      // an entry for the node itself, so two nodes only connected to each other end up with
      // zero intersection.
      
      double val = intersect.size() / (sqrs * sqrt);
      Double dot = new Double(val);
      SortedSet<Link> forDot = retval.get(dot);
      if (forDot == null) {
        forDot = new TreeSet<Link>();
        retval.put(dot, forDot);
      }
      forDot.add(new Link(scache[srcRow.intValue()], scache[trgRow.intValue()]));
    }
    lr.finish();
    return (highestDegree);
  }
 
  /***************************************************************************
  ** 
  ** Per Wikipedia:
   * 
   * Jaccard coefficient measures similarity between sample sets, and is 
   * defined as the size of the intersection divided by the size of the
   * union of the sample sets:
   * 
   * |A I B| / |A U B|
  **
  ** The Jaccard distance, which measures dissimilarity between sample sets, is 
  ** complementary to the Jaccard coefficient and is obtained by subtracting 
  ** the Jaccard coefficient from 1, or, equivalently, by dividing the difference 
  ** of the sizes of the union and the intersection of two sets by the size of the union:
  ** 
  ** (|A U B| - |A I B|) / |A U B|
   * 
   * This distance is a proper metric
  */
  
  private Integer getJaccard(SortedMap<Integer, SortedSet<Integer>> connVec, 
                            SortedMap<Double, SortedSet<Link>> retval, 
                            Map<Integer, Integer> connMag, 
                            BuildData rbd, 
                            Map<NetNode, Integer> targToRow,
  	                        BTProgressMonitor monitor) throws AsynchExitRequestException {

    int rowCount = rbd.getAllNodes().size();
    Integer[] icache = new Integer[rowCount];
    String[] scache = new String[rowCount];
    for (int i = 0; i < rowCount; i++) {
      icache[i] = Integer.valueOf(i);
      scache[i] = icache[i].toString();
    }

    Integer highestDegree = null;
    int biggestMag = Integer.MIN_VALUE;
    HashSet<Integer> union = new HashSet<Integer>();
    HashSet<Integer> intersect = new HashSet<Integer>();
    
    LoopReporter lr = new LoopReporter(rbd.getLinks().size(), 20, monitor, 0.0, 1.0, "progress.getJaccard");  
      
    Iterator<NetLink> ldit = rbd.getLinks().iterator();
    while (ldit.hasNext()) {
      NetLink fl = ldit.next();
      lr.report();
      if (fl.isShadow()) {
      	continue;
      }
      NetNode srcName = fl.getSrcNode();
      Integer srcRow = targToRow.get(srcName);
      NetNode trgName = fl.getTrgNode();
      Integer trgRow = targToRow.get(trgName);
      SortedSet<Integer> srcVec = connVec.get(srcRow);     
      int srcSize = srcVec.size();
      if (srcSize > biggestMag) {
        biggestMag = srcSize;
        highestDegree = srcRow;
      }
      SortedSet<Integer> trgVec = connVec.get(trgRow);     
      int trgSize = trgVec.size();
      if (trgSize > biggestMag) {
        biggestMag = trgSize;
        highestDegree = trgRow;
      }
      connMag.put(icache[srcRow.intValue()], icache[srcSize]);
      connMag.put(icache[trgRow.intValue()], icache[trgSize]);
     
      union.clear();
      DataUtil.union(srcVec, trgVec, union);
      intersect.clear();
      DataUtil.intersection(srcVec, trgVec, intersect);
      int uSize = union.size();
      int iSize = intersect.size();
      
      double jaccard = (double)(iSize) / (double)uSize;    
      Double jaccardObj = new Double(jaccard);
   
      SortedSet<Link> forDot = retval.get(jaccardObj);
      if (forDot == null) {
        forDot = new TreeSet<Link>();
        retval.put(jaccardObj, forDot);
      }
      forDot.add(new Link(scache[srcRow.intValue()], scache[trgRow.intValue()]));
    } 
    lr.finish();
    return (highestDegree);
  }
   
  /***************************************************************************
  ** 
  ** Like the above, but we prefer to keep growing out off of the last used
  ** set of nodes.
  */

  private List<Integer> orderByDistanceChained(Map<Integer, NetNode> rowToTarg, 
  		                                             Integer start, SortedMap<Double, SortedSet<Link>> cosines, 
			                                             Map<Integer, Integer> connMag, List<Link> linkTrace, 
			                                             int limit, double tol, List<Integer> jumpLog,
			                                             BTProgressMonitor monitor) 
			                                               throws AsynchExitRequestException { 
    int rowCount = rowToTarg.size();
    ArrayList<Integer> retval = new ArrayList<Integer>();
    HashSet<Integer> seen = new HashSet<Integer>();  
    retval.add(start);
    seen.add(start);
    
    //
    // Tried running multiple chains, with the idea of being able to
    // pick up on a previously successful chain instead of starting
    // from scratch.  But all the chains got down to the tolerance
    // below the best-match and sat there until they were reused.
    // Or worse, a single match from an old chain fired off before
    // we went to reclaim another chain to start over.  Seems to
    // be little benefit, and big speed hit.
  
    ArrayList<Integer> currentChain = new ArrayList<Integer>();
    currentChain.add(start);
    int switchCount = 0;
    int stayCount = 0;

    LoopReporter lr = new LoopReporter(rowCount, 20, monitor, 0.0, 1.0, "progress.orderByDistanceChained");

    //
    // Keep adding until all nodes are accounted for:
    //
      
    int lastSize = retval.size();
    while (retval.size() < rowCount) {
    	int rtvSize = retval.size();
      lr.report(rtvSize - lastSize);
      lastSize = rtvSize;
      // Find best unconstrained hop:
      DoubleRanked bestHop = findBestUnseenHop(rowToTarg, cosines, connMag, seen, null, retval);  
      // Find best hop off the current search net:   
      DoubleRanked currentHop = findBestUnseenHop(rowToTarg, cosines, connMag, seen, currentChain, retval);
 
      if (bestHop == null) { // Not found, need to find the highest non-seen guy.
        if (currentHop != null) {
          throw new IllegalStateException();
        }
        handleFallbacks(rowToTarg, connMag, linkTrace, seen, retval);
        continue;
      } 
      //
      // If the CURRENT chained hop stuff sucks, we toss the chain and start over:
      //
            
      if ((currentHop == null) || (currentHop.rank <= (bestHop.rank * tol))) {
        seen.add(bestHop.id);
        linkTrace.add(bestHop.byLink);
        jumpLog.add(Integer.valueOf(retval.size()));
        retval.add(bestHop.id);       
        currentChain.clear();
        maintainChain(currentChain, bestHop, limit);
        switchCount++;      
      } else {     
        // As long as the CURRENT chained hop distance is not too bad compared to the unchained,
        // we prefer to use it:
        seen.add(currentHop.id);
        linkTrace.add(currentHop.byLink);
        retval.add(currentHop.id);
        maintainChain(currentChain, currentHop, limit);
        stayCount++;
      }
    }
    lr.finish();
    return (retval);
  }
 

  /***************************************************************************
  ** 
  ** Maintain  the recent used chain
  */

  private void maintainChain(List<Integer> chain, DoubleRanked bestChainedHop, int limit) {
    //
    // The guy who established the link goes first.  The newly added guy goes second.
    // If we hit the limit, the last guys are tossed
    //
    
    Integer bySrc = Integer.valueOf(bestChainedHop.byLink.getSrc());
    Integer byTrg = Integer.valueOf(bestChainedHop.byLink.getTrg());
        
    Integer bridge = (bySrc.equals(bestChainedHop.id)) ? byTrg : bySrc;    
    chain.remove(bridge);
    chain.add(0, bestChainedHop.id);
    chain.add(0, bridge);
            
    while (chain.size() > limit) {
      chain.remove(chain.size() - 1);
    }
    
    return;
  }
  
  /***************************************************************************
  ** 
  ** Start with highest degree node.  Find the highest cosine for it and
  ** another node.  That node then becomes the next to search on.
  */

  private DoubleRanked findBestUnseenHop(Map<Integer, NetNode> nodeForRow, SortedMap<Double, SortedSet<Link>> cosines, 
                                         Map<Integer, Integer> degMag, HashSet<Integer> seen, 
                                         List<Integer> launchNodes, List<Integer> currentOrder) { 
    if ((launchNodes != null) && launchNodes.isEmpty()) {
      return (null);
    }
   
    Iterator<Double> dotIt = cosines.keySet().iterator();
    //
    // Look in order of cosine magnitude, high to low:
    //
    
    String maxDegNodeName = null;
    Integer maxDegNode = null;
    Integer maxDegDeg = null;
    Integer maxDegMinOther = null;
    Integer maxDegOtherNode = null;
    ArrayList<Link> candConnects = new ArrayList<Link>();
    
    while (dotIt.hasNext()) {       
      Double dot = dotIt.next(); 
      SortedSet<Link> forDot = cosines.get(dot);
      maxDegNode = null;
      maxDegDeg = null;
      // Each cosine magnitude has a list of links.  Find ones that span from the
      // set of placed nodes to the set of unplaced nodes:
      for (Link nextLink : forDot) {
        Integer src = Integer.valueOf(nextLink.getSrc());
        Integer trg = Integer.valueOf(nextLink.getTrg());
        Integer cand = null;
        Integer other = null;
        if (seen.contains(src) && !seen.contains(trg)) {
          if ((launchNodes == null) || launchNodes.contains(src)) {
            cand = trg;
            other = src;
          }
        } else if (seen.contains(trg) && !seen.contains(src)) {
          if ((launchNodes == null) || launchNodes.contains(trg)) {
            cand = src;
            other = trg;
          }
        }
        //
        // Having found one, record who has the highest degree:
        //
        if (cand != null) {
          Integer degVal = degMag.get(cand);
          UiUtil.fixMePrintout("current degmag counts src->trg and trg->src directed links as only degree 1");
          String n4r = nodeForRow.get(cand).getName();
          Integer r4o = Integer.valueOf(currentOrder.indexOf(other));
          boolean gtCon = (maxDegDeg == null) || (maxDegDeg.intValue() < degVal.intValue());
          boolean eqCon = (maxDegDeg != null) && (maxDegDeg.intValue() == degVal.intValue());
          boolean ltORow = (maxDegMinOther == null) || (maxDegMinOther.compareTo(r4o) > 0);
          boolean eqORow = (maxDegMinOther != null) && (maxDegMinOther.compareTo(r4o) == 0);
          boolean ltName = (maxDegNodeName == null) || (maxDegNodeName.compareToIgnoreCase(n4r) > 0);
          boolean eqName = (maxDegNodeName != null) && (maxDegNodeName.compareToIgnoreCase(n4r) == 0);
          if (gtCon || (eqCon && (ltORow || eqORow && (ltName || eqName)))) {
            maxDegNode = cand;
            maxDegNodeName = n4r;
            maxDegDeg = degVal;
            maxDegMinOther = r4o;
            maxDegOtherNode = other;
            if (!eqName) {
              candConnects.clear();
            }
            candConnects.add(nextLink);
          }
        }          
      }
      //
      // Having found the one with the highest cosine, tie-breaking with highest degree, tie breaking
      // with connection to higher node, tie-breaking by alphabetical order, add it to the list, 
      // find the "best link" we used to get there:
      //    
     
      if (maxDegNode != null) {
        Link viaLink = null;
        for (Link aConnect : candConnects) {
          String other = (aConnect.getSrc().equals(maxDegNode.toString())) ? aConnect.getTrg() : aConnect.getSrc();
          if (other.equals(maxDegOtherNode.toString())) {
            viaLink = aConnect;
            break;
          }
        }
        if (viaLink == null) {
          throw new IllegalStateException();
        }
      
        return (new DoubleRanked(dot.doubleValue(), maxDegNode, viaLink));
      }
    }
        
    return (null);
  }  
  
  /***************************************************************************
  ** 
  ** Handle the fallback case.
  */

  private void handleFallbacks(Map<Integer, NetNode> rowToNode, Map<Integer, Integer> degMag, List<Link> linkTrace, 
                                HashSet<Integer> seen, List<Integer> retval) { 
    Integer nextBest = getHighestDegreeRemaining(rowToNode, seen, degMag);
    if (nextBest != null) {        
      retval.add(nextBest);
      seen.add(nextBest);
      linkTrace.add(new Link(nextBest.toString(), nextBest.toString()));
    } else {
      // Nodes not connected need to be flushed
      TreeSet<Integer> remainingTargs = new TreeSet<Integer>();
      Iterator<Integer> rtkit = rowToNode.keySet().iterator();
      while (rtkit.hasNext()) {       
        Integer row = rtkit.next();
        remainingTargs.add(row);
      }       
      remainingTargs.removeAll(retval);
      Iterator<Integer> rtit = remainingTargs.iterator();
      while (rtit.hasNext()) {
        Integer rTrg = rtit.next();
        retval.add(rTrg);
        linkTrace.add(new Link(rTrg.toString(), rTrg.toString()));
      }
    }
    return;
  }
  
  /***************************************************************************
  ** 
  ** When we run out of connected nodes, go get the best one remaining
  */

  private Integer getHighestDegreeRemaining(Map<Integer, NetNode> rowToNode, Set<Integer> seen, Map<Integer, Integer> degMag) { 
    Integer maxDegNode = null;
    String maxDegNodeName = null;
    Integer maxDegDeg = null;
    Iterator<Integer> degIt = degMag.keySet().iterator();
    while (degIt.hasNext()) {       
      Integer cand = degIt.next();
      if (seen.contains(cand)) {
        continue;
      }
      Integer degVal = degMag.get(cand);
      String n4r = rowToNode.get(cand).getName();
      boolean gtCon = (maxDegDeg == null) || (maxDegDeg.intValue() < degVal.intValue());
      boolean eqCon = (maxDegDeg != null) && (maxDegDeg.intValue() == degVal.intValue());
      boolean ltName = (maxDegNodeName == null) || (maxDegNodeName.compareToIgnoreCase(n4r) > 0);
      boolean eqName = (maxDegNodeName != null) && (maxDegNodeName.compareToIgnoreCase(n4r) == 0);
      if (gtCon || (eqCon && (ltName || eqName))) {
        maxDegNode = cand;
        maxDegNodeName = n4r;
        maxDegDeg = degVal;
      }
    }              
    return (maxDegNode);
  }
 
  /***************************************************************************
  ** 
  ** Utility conversion
  */

  private void orderToMaps(List<Integer> orderedStringRows, Map<Integer, Integer> forward, Map<Integer, Integer> backward) { 
    int numOsr = orderedStringRows.size();
    for (int i = 0; i < numOsr; i++) {
      Integer sval = orderedStringRows.get(i);
      try {
        Integer newPos = sval;
        Integer oldPos = Integer.valueOf(i);
        forward.put(oldPos, newPos);
        backward.put(newPos, oldPos);
      } catch (NumberFormatException nfex) {
        throw new IllegalStateException();
      }
    }
    return;
  }
  
  /***************************************************************************
  ** 
  ** Debug output:
  */

  @SuppressWarnings("unused")
  private void curveDebug(SortedMap<Integer, Double> curve) {
    Iterator<Integer> cit = curve.keySet().iterator();
    int count = 0;
    while (cit.hasNext()) {
      Integer key = cit.next();
      Double val = curve.get(key);
      System.out.print("(" + key + "," + val + ")");
      if (count++ == 5) {
        break;
      }
    }
    System.out.println();    
  }
 
  /***************************************************************************
  ** 
  ** Build curves and caches
  */

  private void buildCurvesAndCaches(TreeMap<Integer, Integer> oldToNew, TreeMap<Integer, Integer> newToOld, 
                                    SortedMap<Integer, SortedSet<Integer>> connVec, Integer[] icache, Double[] dcache, 
                                    HashMap<Integer, SortedMap<Integer, Double>> curves, 
                                    HashMap<Integer, Map<Double, Set<Integer>>> connectMap, int numRows) {
    for (int i = 0; i < numRows; i++) {
      Integer newRow = Integer.valueOf(i);
      icache[i] = newRow;
      dcache[i] = new Double(i);
      Integer oldRow = newToOld.get(newRow);  
      SortedSet<Integer> baseNodeVec = connVec.get(oldRow);        
      SortedMap<Integer, Double> curve = calcShapeCurve(baseNodeVec, oldToNew);
      curves.put(newRow, curve);
      double connLog = Math.log(baseNodeVec.size()) / Math.log(2.0);
      Double connLogKey = new Double(connLog);
      int logBin = (connLog < 4.0) ? 0 : (int)Math.floor(connLog);
      Integer logBinKey = Integer.valueOf(logBin);
      Map<Double, Set<Integer>> fineGrain = connectMap.get(logBinKey);
      if (fineGrain == null) {
        fineGrain = new HashMap<Double, Set<Integer>>();
        connectMap.put(logBinKey, fineGrain);
      }
      Set<Integer> perFine = fineGrain.get(connLogKey);
      if (perFine == null) {
        perFine = new HashSet<Integer>();
        fineGrain.put(connLogKey, perFine);
      }
      perFine.add(newRow);
    }
    return;
  }

  /***************************************************************************
  ** 
  ** Build the set to check:
  */

  private void buildCheckSet(SortedMap<Integer, Double> baseCurve, TreeSet<Integer> connBuf, 
                             SortedSet<Integer> logKeys, 
                             HashMap<Integer, Map<Double, Set<Integer>>> connectMap) {
    //
    // Chose who to check against based on having about the same number of neighbors:
    //
    double baseConnLog = Math.log(baseCurve.size()) / Math.log(2.0);
    double baseConnLogLo = 0.98 * baseConnLog;
    double baseConnLogHi = 1.02 * baseConnLog;
    logKeys.clear();
    int logBinLo = (baseConnLog < 4.0) ? 0 : (int)Math.floor(baseConnLogLo);
    Integer logBinLoKey = Integer.valueOf(logBinLo);
    logKeys.add(logBinLoKey);
    int logBinHi = (baseConnLog < 4.0) ? 0 : (int)Math.floor(baseConnLogHi);
    Integer logBinHiKey = Integer.valueOf(logBinHi);
    logKeys.add(logBinHiKey);
    logKeys = DataUtil.fillOutHourly(logKeys);
    connBuf.clear();
    Iterator<Integer> lit = logKeys.iterator();
    while (lit.hasNext()) {
      Integer logBinKey = lit.next();
      int logBinVal = logBinKey.intValue();
      Map<Double, Set<Integer>> fineGrain = connectMap.get(logBinKey);
      if (fineGrain == null) {
        continue;
      }
      Iterator<Double> pfg = fineGrain.keySet().iterator();
      while (pfg.hasNext()) {
        Double fineKey = pfg.next();
        double fineKeyVal = fineKey.doubleValue();
        if ((logBinVal == 0) || ((baseConnLogLo <= fineKeyVal) && (baseConnLogHi >= fineKeyVal))) {
          Set<Integer> fineGrains = fineGrain.get(fineKey);
          connBuf.addAll(fineGrains);
        }
      }
    }
    return;
  }
  
  /***************************************************************************
  ** 
  ** Just report current goodness:
  */

  private ClusterPrep setupForResort(SortedMap<Integer, SortedSet<Integer>> connVec, 
                                     List<Integer> orderedStringRows, SortedMap<Integer, Double> rankings) {

    ClusterPrep retval = new ClusterPrep(orderedStringRows.size()); 
 
    orderToMaps(orderedStringRows, retval.oldToNew, retval.newToOld);
      
    //
    // Create a curve cache.
    //
    
    buildCurvesAndCaches(retval.oldToNew, retval.newToOld, connVec, retval.icache, retval.dcache, retval.curves, retval.connectMap, retval.numRows);
    
    //
    // Expensive method did a row rearrangement after every swap and shift.  Note that meant
    // we were running O(n^3), maybe O(n^2logn) at best.  Even then, the first decisions were
    // stale (based on orginal order) by the time the last decisions were made.  Instead, we
    // keep the orginal curves, and instead do multiple passes to iterate to (hopefully) a better
    // fit.
    // Moving left to right, compare each node fit to righthand neighbor (following shift) to all 
    // the remaining righthand neighbors.
    //
    
    TreeSet<Integer> unionBuf = new TreeSet<Integer>();
    TreeSet<Integer> keyBuf = new TreeSet<Integer>();
    double deltaSum = 0.0;
    Iterator<Integer> n2oit = (new TreeSet<Integer>(retval.newToOld.keySet())).iterator();
    while (n2oit.hasNext()) {
      Integer newRow = n2oit.next();
      int currRow = newRow.intValue();
      if (currRow == (retval.numRows - 1)) {
        break;
      }
      SortedMap<Integer, Double> baseCurve = retval.curves.get(newRow);
      SortedMap<Integer, Double> nextCurve = retval.curves.get(retval.icache[currRow + 1]);
      double delt = calcShapeDeltaUsingCurveMaps(baseCurve, nextCurve, unionBuf, keyBuf);
      deltaSum += delt;  
    }
    Integer useKey = (rankings.isEmpty()) ? Integer.valueOf(0) : Integer.valueOf(rankings.lastKey().intValue() + 1);
    rankings.put(useKey, new Double(deltaSum));  
    return (retval);
  }

  /***************************************************************************
  ** 
  ** Resort to group shapes:
  */

  public List<Integer> resort(ClusterPrep prep, BTProgressMonitor monitor, double startFrac, double endFrac) 
                               throws AsynchExitRequestException { 

    /*
    int numRows = orderedStringRows.size();
    TreeMap oldToNew = new TreeMap();
    TreeMap newToOld = new TreeMap();    
    orderToMaps(orderedStringRows, oldToNew, newToOld);
      
    //
    // Create a curve cache.
    //
    
    Integer[] icache = new Integer[numRows];
    Double[] dcache = new Double[numRows];
    HashMap curves = new HashMap();
    HashMap connectMap = new HashMap();
    buildCurvesAndCaches(oldToNew, newToOld, connVec, icache, dcache, curves, connectMap, numRows);
    
    //
    // Expensive method did a row rearrangement after every swap and shift.  Note that meant
    // we were running O(n^3), maybe O(n^2logn) at best.  Even then, the first decisions were
    // stale (based on orginal order) by the time the last decisions were made.  Instead, we
    // keep the orginal curves, and instead do multiple passes to iterate to (hopefully) a better
    // fit.
    // Moving left to right, compare each node fit to righthand neighbor (following shift) to all 
    // the remaining righthand neighbors.
    //
    

    double deltaSum = 0.0;
    Iterator n2oit = (new TreeSet(newToOld.keySet())).iterator();
    while (n2oit.hasNext()) {
      Integer newRow = (Integer)n2oit.next();
      int currRow = newRow.intValue();
      if (currRow == (numRows - 1)) {
        break;
      }
      SortedMap baseCurve = (SortedMap)curves.get(newRow);
      SortedMap nextCurve = (SortedMap)curves.get(icache[currRow + 1]);
      double delt = calcShapeDeltaUsingCurveMaps(baseCurve, nextCurve, unionBuf, keyBuf);
      deltaSum += delt;
    }
    Integer useKey = (rankings.isEmpty()) ? Integer.valueOf(0) : Integer.valueOf(((Integer)rankings.lastKey()).intValue() + 1);
    rankings.put(useKey, new Double(deltaSum));
    
    */
    
    
    
    
    TreeSet<Integer> unionBuf = new TreeSet<Integer>();
    TreeSet<Integer> keyBuf = new TreeSet<Integer>();
    

    TreeSet<Integer> stillAvail = new TreeSet<Integer>(prep.newToOld.keySet());   
    TreeMap<Integer, Integer> results = new TreeMap<Integer, Integer>();
 
    TreeSet<Integer> connBuf = new TreeSet<Integer>();
    SortedSet<Integer> logKeys = new TreeSet<Integer>();
    SortedMap<Integer, Double> baseCurve = null;
    int currRow = 0;
    int startCheck = 1;
    int fillSlot = 0;
    results.put(prep.icache[0], prep.icache[fillSlot++]);
    stillAvail.remove(prep.icache[0]);
    baseCurve = prep.curves.get(prep.icache[0]);

    while (!stillAvail.isEmpty()) {  
 
      Integer newRow = stillAvail.first();
      startCheck = newRow.intValue();

      buildCheckSet(baseCurve, connBuf, logKeys, prep.connectMap);
      connBuf.retainAll(stillAvail);
      // Always include the start in the search!
      connBuf.add(prep.icache[startCheck]);
          
      DoubMinMax dmm = new DoubMinMax();
      dmm.inverseInit();
      //double baseDelt = 0.0;
      double minMatch = Double.POSITIVE_INFINITY;
      int minI = currRow;
      int numCheck = 0;
      Iterator<Integer> cit = connBuf.iterator();
      while (cit.hasNext()) {
        Integer swapCheck = cit.next();
        int swapCheckVal = swapCheck.intValue();
        numCheck++;
        SortedMap<Integer, Double> shiftCandCurve = prep.curves.get(swapCheck);
        double delt = calcShapeDeltaUsingCurveMaps(baseCurve, shiftCandCurve, unionBuf, keyBuf);
        //if (swapCheckVal == startCheck) {
        //  baseDelt = delt;
        //}
        dmm.update(delt);
        if (delt < minMatch) {
         // Need to see if there is a global improvement!
        //  Integer preRemIndx = (Integer)newToOld.get(Integer.valueOf(i - 1));
         // TreeSet preRemoveVec = (TreeSet)connVec.get(preRemIndx);
         // Integer postRemIndx = (Integer)newToOld.get(Integer.valueOf(i + 1));
        //  TreeSet postRemoveVec = (TreeSet)connVec.get(oldForSwap);
        //  double deltRight = calcShapeDeltaViaCurve(preRemoveVec, postRemoveVec, oldToNew);
          minI = swapCheckVal;
          minMatch = delt;
        }
      }
      
      if (minI > startCheck) {
        baseCurve = prep.curves.get(prep.icache[minI]);
        stillAvail.remove(prep.icache[minI]);
        results.put(prep.icache[minI], prep.icache[fillSlot++]);      
      } else {
        baseCurve = prep.curves.get(prep.icache[startCheck]);
        stillAvail.remove(prep.icache[startCheck]);
        results.put(prep.icache[startCheck], prep.icache[fillSlot++]);              
      }
            
      if (monitor != null) {
        double currProg = startFrac + ((endFrac - startFrac) * (1.0 - ((double)stillAvail.size() / (double)prep.numRows)));
        if (!monitor.updateProgress((int)(currProg * 100.0))) {
          throw new AsynchExitRequestException();
        }
      }
    }

    //
    // Convert and leave:
    //
    
    ArrayList<Integer> retval = new ArrayList<Integer>();
    Iterator<Integer> o2nit = prep.oldToNew.values().iterator();
    while (o2nit.hasNext()) {
      Integer newRow = o2nit.next();
      Integer mappedRow = results.get(newRow);
      retval.add(mappedRow);
    }
    return (retval);
  } 
  
  /***************************************************************************
  ** 
  ** Calculate the shape per node:
  */

  private SortedMap<Integer, Double> calcShapeCurve(SortedSet<Integer> vec1, Map<Integer, Integer> newOrder) {
                 
    TreeSet<Integer> reordered1 = new TreeSet<Integer>();
    Iterator<Integer> ub1it = vec1.iterator();
    while (ub1it.hasNext()) {
      Integer linkEnd = ub1it.next();
      Integer mappedEnd = newOrder.get(linkEnd);
      reordered1.add(mappedEnd);
    }    
    ArrayList<Integer> vec1Order = new ArrayList<Integer>(reordered1);
    int numPts = vec1Order.size();
    TreeMap<Integer, Double> vec1Points = new TreeMap<Integer, Double>();
    for (int i = 0; i < numPts; i++) {
      Integer linkEnd = vec1Order.get(i);
      calcCurveMap(reordered1, vec1Order, linkEnd, vec1Points);
    }
  
    return (vec1Points);
  }

  /***************************************************************************
  ** 
  ** Debug output:
  */

  private double curveAverage(SortedMap<Integer, Double> curve) {
    double retval = 0.0;
    Double firstVal = null;
    double lastX = 0.0;
    Iterator<Integer> cit = curve.keySet().iterator();
    while (cit.hasNext()) {
      Integer key = cit.next();
      Double val = curve.get(key);
      double thisKey = key.doubleValue();
      double thisVal = val.doubleValue();
      if (firstVal == null) {
        firstVal = val;
      } else {
        retval += ((thisKey - lastX) * thisVal);        
      }
      lastX = thisKey;
    }
    if (firstVal == null) {
      return (0.0); 
    } else if (lastX == 0.0) {
      return (firstVal.doubleValue()); 
    }
    retval /= lastX;
    return (retval);
  }

  /***************************************************************************
  ** 
  ** Calculate the shape distance:
  */

  private double calcShapeDeltaUsingCurveMaps(SortedMap<Integer, Double> vec1Points, SortedMap<Integer, Double> vec2Points, 
                                              SortedSet<Integer> unionBuf, SortedSet<Integer> keyBuf) {
          
    //
    // We want to find out how close the "link shapes" of two nodes are, given the current
    // node ordering given in newOrder.  We compare the left-hand profiles:
    //
   
    double deltaSqSum = 0.0;
    if (vec1Points.isEmpty() || vec2Points.isEmpty()) {
      return (Double.POSITIVE_INFINITY);
    }
    
    double ca1 = curveAverage(vec1Points);
    double ca2 = curveAverage(vec2Points);
    
    
    unionBuf.clear();
    DataUtil.union(vec1Points.keySet(), vec2Points.keySet(), unionBuf);
    Iterator<Integer> ubit = unionBuf.iterator();
    while (ubit.hasNext()) {
      Integer point = ubit.next();
      double v1Val = interpCurve(vec1Points, point, keyBuf) - ca1;
      double v2Val = interpCurve(vec2Points, point, keyBuf) - ca2;
      double yDelt = v1Val - v2Val;
      deltaSqSum += (yDelt * yDelt);
    }
    double retval = Math.sqrt(deltaSqSum);
    if (Double.isNaN(retval)) {
      System.err.println(vec1Points);
      System.err.println(vec2Points);
      System.err.println(unionBuf);
      throw new IllegalStateException();
    }
    return (retval);
  }
  
  /***************************************************************************
  ** 
  ** Calculate a curve.  Now only used for exact hits; interpolation code is obsolete!
  */

  private void calcCurveMap(SortedSet<Integer> vec, List<Integer> vecOrder, Integer linkEnd, Map<Integer, Double> curvePoints) {
    int numVec = vecOrder.size();      
    int vec1Pos = vecOrder.indexOf(linkEnd);
    if (vec1Pos != -1) {
      curvePoints.put(linkEnd, new Double(numVec - vec1Pos));
    } else {
      throw new IllegalArgumentException();
    }
    return;
  }  
  
  /***************************************************************************
  ** 
  ** Calculate a curve point:
  */

  private double interpCurve(SortedMap<Integer, Double> curve, Integer xVal, SortedSet<Integer> xBuf) {

    Double exact = curve.get(xVal);    
    if (exact != null) {
      return (exact.doubleValue());
    } else {
      double[] weights = new double[2];
      xBuf.clear();
      xBuf.addAll(curve.keySet());
      MinMax bounds = DataUtil.boundingInts(xBuf, xVal.intValue());
      boolean areDiff = getWeights(bounds.min, bounds.max, xVal.doubleValue(), weights);
      if (areDiff) {        
        Integer mappedEndLo = Integer.valueOf(bounds.min);
        Integer mappedEndHi = Integer.valueOf(bounds.max);
       // double vec1XVal = Math.round(weights[0] * (double)bounds.min + weights[1] * (double)bounds.max);     
        double vec1YVal = weights[0] * curve.get(mappedEndLo).doubleValue() + weights[1] * curve.get(mappedEndHi).doubleValue();        
        return (vec1YVal);
      } else if (bounds.min > xBuf.first().intValue())  {
        return (curve.get(curve.firstKey()).doubleValue());
      } else {
        return (0.0);
      }        
    }
  }

  /***************************************************************************
  **
  ** For passing around ranked nodes
  */  
  
  static class DoubleRanked  {
     double rank;
     Integer id;
     Link byLink;

    DoubleRanked(double rank, Integer id, Link byLink) {
      this.rank = rank;
      this.id = id;
      this.byLink = byLink;
    } 
  }
  
  
  /***************************************************************************
  **
  ** For passing around prep data
  */  
  
  public static class ClusterPrep  { 
    int numRows;
    TreeMap<Integer, Integer> oldToNew;
    TreeMap<Integer, Integer> newToOld;    
    Integer[] icache;
    Double[] dcache;
    HashMap<Integer, SortedMap<Integer, Double>> curves;
    HashMap<Integer, Map<Double, Set<Integer>>> connectMap;  

    ClusterPrep(int numRows) { 
      this.numRows = numRows;
      oldToNew = new TreeMap<Integer, Integer>();
      newToOld = new TreeMap<Integer, Integer>();    
      icache = new Integer[numRows];
      dcache = new Double[numRows];
      curves = new HashMap<Integer, SortedMap<Integer, Double>>();
      connectMap = new HashMap<Integer, Map<Double, Set<Integer>>>();  
    }
  }

  
  ////////////////////////////////////////////////////////////////////////////
  //
  // PUBLIC INNER CLASSES
  //
  ////////////////////////////////////////////////////////////////////////////

  /***************************************************************************
  **
  ** For passing around layout params
  */  
  
  public static class ClusterParams implements Params {
    
    // NOTE:
    //
    // JACCARD gives intersection size over "how many connections do WE have"
    // COSINES gives intersection size over:
    //   sqrt("how many connections do I have)" * sqrt("how many connections do you have)"
    //
     
    public static final int JACCARD = 0;
    public static final int COSINES = 1;
        
    public double tolerance;
    public int chainLength; 
    public int distanceMethod;

    public ClusterParams(double tolerance, int chainLength, int distanceMethod) {
      this.tolerance = tolerance;
      this.chainLength = chainLength;
      this.distanceMethod = distanceMethod;
    }
    
    public ClusterParams() {
      tolerance = 0.80;
      chainLength = 15;
      distanceMethod = JACCARD;    
    }
    
    public static Vector<ChoiceContent> getDistanceChoices() {
      ResourceManager rMan = ResourceManager.getManager();
      Vector<ChoiceContent> retval = new Vector<ChoiceContent>();
      retval.add(new ChoiceContent(rMan.getString("clusterParams.jaccard"), JACCARD));
      retval.add(new ChoiceContent(rMan.getString("clusterParams.cosines"), COSINES));   
      return (retval);
    }
  } 
  
  /***************************************************************************
  **
  ** For passing around resort params
  */  
  
  public static class ResortParams implements Params {        
    public int passCount; 
    public boolean terminateAtIncrease; 

    public ResortParams(double tolerance, int passCount, boolean terminateAtIncrease) {
      this.passCount = passCount;
      this.terminateAtIncrease = terminateAtIncrease;
    }    
    public ResortParams() {
      passCount = 10;
      terminateAtIncrease = false;    
    }
  }
  
  /***************************************************************************
  **
  ** Affine coords in a single dimension
  */
  
  private boolean getWeights(double val1, double val2, double calcForVal, double[] toFill) {
    if (val1 == val2) {
      return (false);
    }
    toFill[0] = (calcForVal - val2) / (val1 - val2);
    toFill[1] = 1.0 - toFill[0];
    return (true);
  }
  
  
  
}
```
