## Supplementary material for "BioFabric Visualization of Network Alignments": BioFabric Source Code: SetLayout.java

```
/*
**    Copyright (C) 2003-2018 Institute for Systems Biology 
**                            Seattle, Washington, USA. 
**
**    This library is free software; you can redistribute it and/or
**    modify it under the terms of the GNU Lesser General Public
**    License as published by the Free Software Foundation; either
**    version 2.1 of the License, or (at your option) any later version.
**
**    This library is distributed in the hope that it will be useful,
**    but WITHOUT ANY WARRANTY; without even the implied warranty of
**    MERCHANTABILITY or FITNESS FOR A PARTICULAR PURPOSE.  See the GNU
**    Lesser General Public License for more details.
**
**    You should have received a copy of the GNU Lesser General Public
**    License along with this library; if not, write to the Free Software
**    Foundation, Inc., 59 Temple Place, Suite 330, Boston, MA  02111-1307  USA
*/

package org.systemsbiology.biofabric.layouts;

import java.util.ArrayList;
import java.util.Collections;
import java.util.HashMap;
import java.util.HashSet;
import java.util.List;
import java.util.Map;
import java.util.Set;
import java.util.SortedMap;
import java.util.SortedSet;
import java.util.TreeMap;
import java.util.TreeSet;

import org.systemsbiology.biofabric.analysis.GraphSearcher;
import org.systemsbiology.biofabric.api.io.BuildData;
import org.systemsbiology.biofabric.api.layout.DefaultEdgeLayout;
import org.systemsbiology.biofabric.api.layout.DefaultLayout;
import org.systemsbiology.biofabric.api.layout.LayoutCriterionFailureException;
import org.systemsbiology.biofabric.api.layout.NodeLayout;
import org.systemsbiology.biofabric.api.model.Annot;
import org.systemsbiology.biofabric.api.model.AnnotationSet;
import org.systemsbiology.biofabric.api.model.NetLink;
import org.systemsbiology.biofabric.api.model.NetNode;
import org.systemsbiology.biofabric.api.worker.AsynchExitRequestException;
import org.systemsbiology.biofabric.api.worker.BTProgressMonitor;
import org.systemsbiology.biofabric.api.worker.LoopReporter;
import org.systemsbiology.biofabric.plugin.PluginSupportFactory;

/****************************************************************************
**
** This handles creating a layout for the network presentation of set membership
*/

public class SetLayout extends NodeLayout {
  
  ////////////////////////////////////////////////////////////////////////////
  //
  // PRIVATE CONSTANTS
  //
  //////////////////////////////////////////////////////////////////////////// 
   
  ////////////////////////////////////////////////////////////////////////////
  //
  // PUBLIC CONSTANTS
  //
  //////////////////////////////////////////////////////////////////////////// 
 
  public enum LinkMeans {BELONGS_TO, CONTAINS}
  
  ////////////////////////////////////////////////////////////////////////////
  //
  // PRIVATE INSTANCE MEMBERS
  //
  ////////////////////////////////////////////////////////////////////////////
  
  private Map<NetNode, Set<NetNode>> elemsPerSet_;
  private Map<NetNode, Set<NetNode>> setsPerElem_;
  private LinkMeans direction_;

  ////////////////////////////////////////////////////////////////////////////
  //
  // PUBLIC CONSTRUCTORS
  //
  ////////////////////////////////////////////////////////////////////////////

  /***************************************************************************
  **
  ** Constructor
  */

  public SetLayout(LinkMeans direction) {
    direction_ = direction;
  }

  ////////////////////////////////////////////////////////////////////////////
  //
  // PUBLIC METHODS
  //
  ////////////////////////////////////////////////////////////////////////////

  /***************************************************************************
  **
  ** Find out if the necessary conditions for this layout are met. 
  */
  
  @Override
  public boolean criteriaMet(BuildData rbd,
                             BTProgressMonitor monitor) throws AsynchExitRequestException, 
                                                               LayoutCriterionFailureException {
    //
    // 1) All the relations in the network are directed
    // 2) The network is bipartite (so self links are not legit)
    // 3) singleton nodes represent empty sets
    // 4) Not a multigraph. We currently insure that by enforcing only one link relation.
  	//
    
    LoopReporter lr = new LoopReporter(rbd.getLinks().size(), 20, monitor, 0.0, 1.0, "progress.setLayoutCriteriaCheck"); 
    
    HashSet<String> rels = new HashSet<String>();
    for (NetLink aLink : rbd.getLinks()) {
      lr.report();
      if (!aLink.isDirected()) {
        throw new LayoutCriterionFailureException();
      }
      rels.add(aLink.getRelation());
      if (rels.size() > 1) {
        throw new LayoutCriterionFailureException();
      }    
    }
    lr.finish();
    
    elemsPerSet_ = new HashMap<NetNode, Set<NetNode>>();
    setsPerElem_ = new HashMap<NetNode, Set<NetNode>>();    
    
    extractSets(rbd.getLinks(), rbd.getSingletonNodes(), monitor);
 
    return (true);  
  }
  
  /***************************************************************************
  **
  ** Order the nodes
  */
  
  public List<NetNode> doNodeLayout(BuildData rbd,
  		                              Params params,
                                    BTProgressMonitor monitor) throws AsynchExitRequestException {
        
    //
    // We order the sets by cardinality, largest first. Ties broken by lexicographic order:
    //
    
    Map<Integer, SortedSet<NetNode>> byDegree = new TreeMap<Integer, SortedSet<NetNode>>(Collections.reverseOrder());
    
    for (NetNode set : elemsPerSet_.keySet()) {
      Set<NetNode> elems = elemsPerSet_.get(set);
      Integer card = Integer.valueOf(elems.size());
      SortedSet<NetNode> setsWithCard = byDegree.get(card);
      if (setsWithCard == null) {
        setsWithCard = new TreeSet<NetNode>();
        byDegree.put(card, setsWithCard);
      }
      setsWithCard.add(set);
    }

    //
    // Now we create an ordered list of the sets. We stick empty sets up top so they do not get hidden:
    //
    
    ArrayList<NetNode> setList = new ArrayList<NetNode>();
    
    SortedSet<NetNode> emptySets = byDegree.get(Integer.valueOf(0));
    if (emptySets != null) {
    	setList.addAll(emptySets);
    }
    
    for (Integer size : byDegree.keySet()) {
    	if (size.intValue() == 0) {
    		continue;
    	}
    	SortedSet<NetNode> forDeg = byDegree.get(size);
      setList.addAll(forDeg);
    }

    ArrayList<NetNode> nodeOrder = new ArrayList<NetNode>(setList);
    Map<NetNode, Set<NetNode>> setsHoldingElement = new HashMap<NetNode, Set<NetNode>>();
    HashMap<Set<NetNode>, String> tagMap = new HashMap<Set<NetNode>, String>();
    StringBuffer buf = new StringBuffer();

    SortedSet<GraphSearcher.SourcedNodeGray> snds = GraphSearcher.nodeGraySetWithSourceFromMap(setList, setsPerElem_);
    for (GraphSearcher.SourcedNodeGray node : snds) {   
      nodeOrder.add(node.getNodeID());
      Set<NetNode> setNodes = node.getSrcs();
      setsHoldingElement.put(node.getNodeID(), setNodes); 
      String tag = tagMap.get(setNodes);
      if (tag == null) {
        tag = buildTag(setNodes, setList, buf);
        tagMap.put(setNodes, tag);
      } 
    }

    //
    // Now have the ordered list of targets we are going to display.
    // Build target->row maps and the inverse:
    //
    
    (new DefaultLayout()).installNodeOrder(nodeOrder, rbd, monitor);
    
    //
    // We lay out the edges using default layout. The only reason we do this here is because we want to do link annotations
    // as part of the layout:
    //
   
    (new DefaultEdgeLayout()).layoutEdges(rbd, monitor); 
    
    
    AnnotationSet nAnnots = generateNodeAnnotations(rbd, setsHoldingElement, tagMap);
    rbd.setNodeAnnotations(nAnnots);
    
   Map<Boolean, AnnotationSet> lAnnots = generateLinkAnnotations(rbd, setsHoldingElement, tagMap);
   rbd.setLinkAnnotations(lAnnots);
    
    return (nodeOrder);
  }

  /***************************************************************************
  **
  ** Generate set intersection tag
  */
    
  private String buildTag(Set<NetNode> setNodes, ArrayList<NetNode> setList, StringBuffer buf) {
     
  	buf.setLength(0);
  	boolean first = true;
    for (NetNode node : setList) {
      if (setNodes.contains(node)) {
	      if (first) {
	    		first = false;
	    	} else {
	    		buf.append("&");
	    	}
      	buf.append(node.getName());
      }
    }
    return (buf.toString());
  }

  /***************************************************************************
  **
  ** Generate node annotations to tag each cluster
  */
    
  private AnnotationSet generateNodeAnnotations(BuildData rbd, Map<NetNode, Set<NetNode>> setsHoldingElement, Map<Set<NetNode>, String> tagMap) {
    
    AnnotationSet retval = PluginSupportFactory.buildAnnotationSet();
    
    TreeMap<Integer, NetNode> invert = new TreeMap<Integer, NetNode>();
    
    Map<NetNode, Integer> nod = rbd.getNodeOrder(); 
    
    for (NetNode node : nod.keySet()) {
      invert.put(nod.get(node), node);
    }
     
    Set<NetNode> currIntersect = null;
    Integer startRow = null;
    Integer lastKey = invert.lastKey();
    for (Integer row : invert.keySet()) {
      NetNode node = invert.get(row);
      Set<NetNode> sets = setsHoldingElement.get(node);
      if (currIntersect == null) {
        currIntersect = sets;
        startRow = row;
        if (row.equals(lastKey)) {
          Annot annot = PluginSupportFactory.buildAnnotation(tagMap.get(currIntersect), startRow.intValue(), row.intValue(), 0, null);
          retval.addAnnot(annot);
          break;
        }
        continue;
      }
      if (currIntersect.equals(sets)) {
        if (row.equals(lastKey)) {
          Annot annot = PluginSupportFactory.buildAnnotation(tagMap.get(currIntersect), startRow.intValue(), row.intValue(), 0, null);
          retval.addAnnot(annot);
          break;
        }
        continue;
      } else { 
        // We have just entered a new cluster
        Annot annot = PluginSupportFactory.buildAnnotation(tagMap.get(currIntersect), startRow.intValue(), row.intValue() - 1, 0, null);

        retval.addAnnot(annot);
        startRow = row;
        currIntersect = sets;
        if (row.equals(lastKey)) {
          annot = PluginSupportFactory.buildAnnotation(tagMap.get(currIntersect), startRow.intValue(), row.intValue(), 0, null);
          retval.addAnnot(annot);
          break;
        }
      }
    }
    
    return (retval);
  }
  
   /***************************************************************************
  **
  ** Generate link annotations to tag each set node and intersection band
  */
    
  private Map<Boolean, AnnotationSet> generateLinkAnnotations(BuildData rbd, Map<NetNode, Set<NetNode>> setsHoldingElement, 
  																														Map<Set<NetNode>, String> tagMap) { 
  	HashMap<Boolean, AnnotationSet> retval = new HashMap<Boolean, AnnotationSet>();
    
  	SortedMap<Integer, NetLink> lod = rbd.getLinkOrder();
  
  	List<NetLink> shadowsOnly = new ArrayList<NetLink>();
  	List<NetLink> noShadows = new ArrayList<NetLink>();

  	int firstShadow = -1;
  	for (Integer col : lod.keySet()) {
  		NetLink fl = lod.get(col);
  		if (fl.isShadow()) {
  			if (firstShadow < 0) {
  				firstShadow = col.intValue();
  			}
  			shadowsOnly.add(fl);
  		} else {
  			noShadows.add(fl);
  		}
  	}

  	//
  	// Link annotations for regular links delineate sets. For shadow links, annotations delineate nodes with identical set
    // memberships:
  	//
  	
  	AnnotationSet forSets = generateLinkAnnotationsForSets(noShadows);
  	AnnotationSet noShad = PluginSupportFactory.buildAnnotationSet();
  	AnnotationSet withShad = PluginSupportFactory.buildAnnotationSet();
  	for (Annot an : forSets) {
  		noShad.addAnnot(an);
  		withShad.addAnnot(an);
  	}  	
  	
  	retval.put(Boolean.FALSE, noShad);
  	retval.put(Boolean.TRUE, appendLinkAnnotationsForIntersections(withShad, shadowsOnly, setsHoldingElement, tagMap, firstShadow));
  	    
    return (retval);
  }

  /***************************************************************************
  **
  ** Generate link annotations to tag each set intersection band
  */
    
  private AnnotationSet generateLinkAnnotationsForSets(List<NetLink> linkList) { 
    
    AnnotationSet retval = PluginSupportFactory.buildAnnotationSet();
  	
  	NetNode currSetNode = null;
    int startCol = 0;
    int lastCol = linkList.size() - 1;
    
    for (int i = 0; i < linkList.size(); i++) {
  		NetLink fl = linkList.get(i);
  		if (fl.isShadow()) {
  			throw new IllegalArgumentException();
  		}
  		NetNode nextSet = (direction_ == LinkMeans.CONTAINS) ? fl.getSrcNode() : fl.getTrgNode();
      if (currSetNode == null) {
        currSetNode = nextSet;
        startCol = i;
        if (i == lastCol) {
          Annot annot = PluginSupportFactory.buildAnnotation(currSetNode.getName(), startCol, i, 0, null);
          retval.addAnnot(annot);
          break;
        }
        continue;
      }
      if (currSetNode.equals(nextSet)) {
        if (i == lastCol) {
          Annot annot = PluginSupportFactory.buildAnnotation(currSetNode.getName(), startCol, i, 0, null);
          retval.addAnnot(annot);
          break;
        }
        continue;
      } else { 
        // We have just entered a new cluster
        Annot annot = PluginSupportFactory.buildAnnotation(currSetNode.getName(), startCol, i - 1, 0, null);

        retval.addAnnot(annot);
        startCol = i;
        currSetNode = nextSet;
        if (i == lastCol) {
          annot = PluginSupportFactory.buildAnnotation(currSetNode.getName(), startCol, i, 0, null);
          retval.addAnnot(annot);
          break;
        }
      }
    }
    
    return (retval);
  }
  
  /***************************************************************************
  **
  ** Generate link annotations to tag each set intersection band
  */
    
  private AnnotationSet appendLinkAnnotationsForIntersections(AnnotationSet retval, List<NetLink> linkList, 
  		                                                        Map<NetNode, Set<NetNode>> setsHoldingElement, 
  																									          Map<Set<NetNode>, String> tagMap, int offset) { 
    
  	Set<NetNode> currIntersect = null;
    int startCol = offset;
    int lastCol = offset + linkList.size() - 1;
    
    for (int i = 0; i < linkList.size(); i++) {
  		NetLink fl = linkList.get(i);
  		if (!fl.isShadow()) {
  			throw new IllegalArgumentException();
  		}
  		NetNode elem = (direction_ == LinkMeans.CONTAINS) ? fl.getTrgNode() : fl.getSrcNode();
      Set<NetNode> sets = setsHoldingElement.get(elem);
      if (currIntersect == null) {
        currIntersect = sets;
        startCol = i + offset;
        if (i + offset == lastCol) {
          Annot annot = PluginSupportFactory.buildAnnotation(tagMap.get(currIntersect), startCol, i + offset, 0, null);
          retval.addAnnot(annot);
          break;
        }
        continue;
      }
      if (currIntersect.equals(sets)) {
        if (i + offset == lastCol) {
          Annot annot = PluginSupportFactory.buildAnnotation(tagMap.get(currIntersect), startCol, i + offset, 0, null);
          retval.addAnnot(annot);
          break;
        }
        continue;
      } else { 
        // We have just entered a new cluster
        Annot annot = PluginSupportFactory.buildAnnotation(tagMap.get(currIntersect), startCol, i + offset - 1, 0, null);

        retval.addAnnot(annot);
        startCol = i + offset;
        currIntersect = sets;
        if (i + offset == lastCol) {
          annot = PluginSupportFactory.buildAnnotation(tagMap.get(currIntersect), startCol, i + offset, 0, null);
          retval.addAnnot(annot);
          break;
        }
      }
    }
    
    return (retval);
  }
  
  /***************************************************************************
  ** 
  ** Extract the set information from the network
  */

  private void extractSets(Set<NetLink> links, Set<NetNode> singletons,
                           BTProgressMonitor monitor) throws AsynchExitRequestException, 
                                                             LayoutCriterionFailureException {
  
    //
    // This graph has to be bipartite, with all directional links going from node set A to node set B
    // ("BELONGS_TO") or the opposite ("CONTAINS"). Any singleton nodes must be considered to be empty sets.
    //
    
    HashSet<NetNode> setNodes = new HashSet<NetNode>();
    HashSet<NetNode> elementNodes = new HashSet<NetNode>();     
    
    // Singletons are empty sets:
    
    if (singletons != null) {
	    setNodes.addAll(singletons);
	    for (NetNode emptySet : singletons) {
	      elemsPerSet_.put(emptySet, new HashSet<NetNode>());
	    }
    } 
       
    LoopReporter lr = new LoopReporter(links.size(), 20, monitor, 0.0, 1.0, "progress.setLayoutSetExtraction"); 
    
    for (NetLink link : links) {
      lr.report();
      NetNode set = (direction_ == LinkMeans.CONTAINS) ? link.getSrcNode() : link.getTrgNode();
      NetNode elem = (direction_ == LinkMeans.CONTAINS) ? link.getTrgNode() : link.getSrcNode();
      setNodes.add(set);
      elementNodes.add(elem);
      
      Set<NetNode> forSet = elemsPerSet_.get(set);
      if (forSet == null) {
        forSet = new HashSet<NetNode>();
        elemsPerSet_.put(set, forSet);
      }
      forSet.add(elem);
      
      Set<NetNode> forElem = setsPerElem_.get(elem);
      if (forElem == null) {
        forElem = new HashSet<NetNode>();
        setsPerElem_.put(elem, forElem);
      }
      forElem.add(set);  
    }
    lr.finish();
    
    HashSet<NetNode> intersect = new HashSet<NetNode>(setNodes);
    intersect.retainAll(elementNodes);
    if (!intersect.isEmpty()) {
      throw new LayoutCriterionFailureException();
    }
  
    return;
  }
}
```
