## Supplementary material for "BioFabric Visualization of Network Alignments": BioFabric Source Code: WorldBankLayout.java

```
/*
**    Copyright (C) 2003-2018 Institute for Systems Biology 
**                            Seattle, Washington, USA. 
**
**    This library is free software; you can redistribute it and/or
**    modify it under the terms of the GNU Lesser General Public
**    License as published by the Free Software Foundation; either
**    version 2.1 of the License, or (at your option) any later version.
**
**    This library is distributed in the hope that it will be useful,
**    but WITHOUT ANY WARRANTY; without even the implied warranty of
**    MERCHANTABILITY or FITNESS FOR A PARTICULAR PURPOSE.  See the GNU
**    Lesser General Public License for more details.
**
**    You should have received a copy of the GNU Lesser General Public
**    License along with this library; if not, write to the Free Software
**    Foundation, Inc., 59 Temple Place, Suite 330, Boston, MA  02111-1307  USA
*/

package org.systemsbiology.biofabric.layouts;

import java.util.ArrayList;
import java.util.Collections;
import java.util.HashMap;
import java.util.HashSet;
import java.util.Iterator;
import java.util.List;
import java.util.Map;
import java.util.Set;
import java.util.SortedMap;
import java.util.SortedSet;
import java.util.TreeMap;
import java.util.TreeSet;

import org.systemsbiology.biofabric.api.io.BuildData;
import org.systemsbiology.biofabric.api.layout.LayoutCriterionFailureException;
import org.systemsbiology.biofabric.api.layout.NodeLayout;
import org.systemsbiology.biofabric.api.model.NetLink;
import org.systemsbiology.biofabric.api.model.NetNode;
import org.systemsbiology.biofabric.api.worker.AsynchExitRequestException;
import org.systemsbiology.biofabric.api.worker.BTProgressMonitor;
import org.systemsbiology.biofabric.api.worker.LoopReporter;
import org.systemsbiology.biofabric.util.UiUtil;

/****************************************************************************
**
** This does the "world bank" layout
*/

public class WorldBankLayout extends NodeLayout {
  
  ////////////////////////////////////////////////////////////////////////////
  //
  // PRIVATE CONSTANTS
  //
  //////////////////////////////////////////////////////////////////////////// 
  
  ////////////////////////////////////////////////////////////////////////////
  //
  // PUBLIC CONSTANTS
  //
  //////////////////////////////////////////////////////////////////////////// 
  
  ////////////////////////////////////////////////////////////////////////////
  //
  // PRIVATE INSTANCE MEMBERS
  //
  ////////////////////////////////////////////////////////////////////////////
  
  ////////////////////////////////////////////////////////////////////////////
  //
  // PUBLIC CONSTRUCTORS
  //
  ////////////////////////////////////////////////////////////////////////////

  /***************************************************************************
  **
  ** Constructor
  */

  public WorldBankLayout() {
  }

  ////////////////////////////////////////////////////////////////////////////
  //
  // PUBLIC METHODS
  //
  ////////////////////////////////////////////////////////////////////////////

  /***************************************************************************
  **
  ** Find out if the necessary conditions for this layout are met. 
  */
  
  @Override
  public boolean criteriaMet(BuildData rbd,
                             BTProgressMonitor monitor) throws AsynchExitRequestException, 
                                                               LayoutCriterionFailureException {
    //
    // 1) This is going to have problems with two connected nodes of degree 1.
    // 2) Other topology issues???
    //
    // NOTE: If we want to reproduce the true world bank network, we need to assign in-country,
    // between two-country, and global links to different link groups. 
    // 
    // NOTE 2: Note the "popular" hubs in the world bank network ARE NOT THEMSELVES INTERACTING. 
    
    
    LoopReporter lr = new LoopReporter(rbd.getLinks().size(), 20, monitor, 0.0, 1.0, "progress.hDagLayoutCriteriaCheck");
    
    for (NetLink aLink : rbd.getLinks()) {
      lr.report();
      //if (!aLink.isDirected()) {
       // throw new LayoutCriterionFailureException();
     // }
    }
    lr.finish();
    
   // CycleFinder cf = new CycleFinder(rbd.allNodeIDs, rbd.allLinks, monitor);
  //  if (cf.hasACycle(monitor)) {
   //   throw new LayoutCriterionFailureException();
   // }
    return (true);  
  }
  
  /***************************************************************************
  **
  ** Relayout the network!
  */
  
  public List<NetNode> doNodeLayout(BuildData rbd,
  																  Params params,
  		                              BTProgressMonitor monitor) throws AsynchExitRequestException {
    
    List<NetNode> targets = calcNodeOrder(rbd.getLinks(), rbd.getSingletonNodes(), monitor);       

    //
    // Now have the ordered list of targets we are going to display.
    // Build target->row maps and the inverse:
    //
    
    installNodeOrder(targets, rbd, monitor);
    return (targets);
  }
  
  /***************************************************************************
  ** 
  ** Bump count
  */

  private void bumpDaCount(Map<NetNode, Integer> countMap, NetNode dNode) {    
    Integer inc = countMap.get(dNode);
    if (inc == null) {
      countMap.put(dNode, Integer.valueOf(1));
    } else {
      countMap.put(dNode, Integer.valueOf(inc.intValue() + 1));
    }
    return;
  }
  
  /***************************************************************************
  ** 
  ** Track neighbors
  */

  private void addANeighbor(Map<NetNode, SortedSet<NetNode>> neighMap, NetNode daNode, NetNode daNeigh) {    
    SortedSet<NetNode> forNode = neighMap.get(daNode);
    if (forNode == null) {
      forNode = new TreeSet<NetNode>();
      neighMap.put(daNode, forNode);
    }
    forNode.add(daNeigh);
    return;
  }

  /***************************************************************************
  ** 
  ** Take the node->degree map and invert to create a decreasing-order 
  ** degree->sortedSet(node) map:
  */

  private SortedMap<Integer, SortedSet<NetNode>> invertCountMap(Map<NetNode, Integer> countMap, 
  		                                                               BTProgressMonitor monitor) throws AsynchExitRequestException { 

    LoopReporter lr = new LoopReporter(countMap.size(), 20, monitor, 0.0, 1.0, "progress.sortingByDegree");
  	    
  	TreeMap<Integer, SortedSet<NetNode>> retval = new TreeMap<Integer, SortedSet<NetNode>>(Collections.reverseOrder());
    for (NetNode daKey : countMap.keySet()) {
      lr.report();
      Integer daCount = countMap.get(daKey);
      SortedSet<NetNode> forCount = retval.get(daCount);
      if (forCount == null) {
        forCount = new TreeSet<NetNode>();
        retval.put(daCount, forCount);
      }
      forCount.add(daKey);
    }
    lr.finish();
    return (retval);
  }
  
  /***************************************************************************
  ** 
  ** Flatten a map
  */

  private List<NetNode> flattenDaCount(SortedMap<Integer, SortedSet<NetNode>> invCountMap) { 
    ArrayList<NetNode> retval = new ArrayList<NetNode>();
    Iterator<SortedSet<NetNode>> icmit = invCountMap.values().iterator();
    while (icmit.hasNext()) {
      retval.addAll(icmit.next());
    }
    return (retval);
  }

  /***************************************************************************
  ** 
  ** Calculate node order
  */

  private List<NetNode> calcNodeOrder(Set<NetLink> allLinks, Set<NetNode> loneNodes,
  		                                     BTProgressMonitor monitor) throws AsynchExitRequestException {
 
    HashMap<NetNode, Integer> node2Degree = new HashMap<NetNode, Integer>();
    HashMap<NetNode, SortedSet<NetNode>> node2Neighbor = new HashMap<NetNode, SortedSet<NetNode>>(); 
    HashSet<NetNode> allNodes = new HashSet<NetNode>();
    
    LoopReporter lr = new LoopReporter(allLinks.size(), 20, monitor, 0.0, 1.0, "progress.wblCalcNodeOrder");

    for (NetLink nextLink : allLinks) {
      lr.report();
      NetNode source = nextLink.getSrcNode();
      NetNode target = nextLink.getTrgNode();
 
      //
      // Track all link-related nodes:
      //
      
      allNodes.add(source);
      allNodes.add(target);
      
      //
      // Build the node2Degree map:
      //
      
      bumpDaCount(node2Degree, source);
      bumpDaCount(node2Degree, target);
      
      //
      // Record the neighbors for all the node:
      //
      addANeighbor(node2Neighbor, source, target);
      addANeighbor(node2Neighbor, target, source);
    }
    lr.finish();
    
    SortedMap<Integer, SortedSet<NetNode>> degree2Nodes = invertCountMap(node2Degree, monitor);
    
    //
    // For nodes that have *one* neighbor, collect those popular neighbors (i.e. hubs):
    //
    
    LoopReporter lr2 = new LoopReporter(node2Neighbor.size(), 20, monitor, 0.0, 1.0, "progress.wblPopNeighbors");
    
    HashMap<NetNode, Set<NetNode>> oneNeighbor = new HashMap<NetNode, Set<NetNode>>();
    for (NetNode node : node2Neighbor.keySet()) {
      lr2.report();
      SortedSet<NetNode> nextDoor = node2Neighbor.get(node);
      if (nextDoor.size() == 1) {        
        NetNode popular = nextDoor.first();
        Set<NetNode> popFriends = oneNeighbor.get(popular);
        if (popFriends == null) {
          popFriends = new HashSet<NetNode>();
          oneNeighbor.put(popular, popFriends);
        }
        popFriends.add(node);
      }
    }
    lr2.finish();
    
    LoopReporter lr3 = new LoopReporter(degree2Nodes.size(), 0, monitor, 0.0, 1.0, "progress.addingSatellites");
    
    
    ArrayList<NetNode> targets = new ArrayList<NetNode>();
    HashSet<NetNode> tSet = new HashSet<NetNode>(); // For rapid lookup
    
    //
    // Do this in decreasing degree order for the popular neighbors (hubs):
    //
    
    Set<NetNode> populars = oneNeighbor.keySet();
    
    for (Integer deg : degree2Nodes.keySet()) {
      lr3.report();
      //
      if (deg.intValue() == 1) {
         
        
        
        UiUtil.fixMePrintout("DO SOMETHING: Don't want 1-deg node added twice!");
      }
      //
      // Crank thru the populars at a particular degree:
      //
      SortedSet<NetNode> forDeg = degree2Nodes.get(deg);
      for (NetNode degNode : forDeg) {
        if (populars.contains(degNode)) {
          targets.add(degNode); // Add the popular node
          tSet.add(degNode);
          HashMap<NetNode, Integer> forDaPop = new HashMap<NetNode, Integer>();
          // Get the unpopular friends:
          Set<NetNode> unpopFriends = oneNeighbor.get(degNode);
          for (NetNode unPop : unpopFriends) {
            Integer upd = node2Degree.get(unPop);
            forDaPop.put(unPop, upd);            
          }
          //
          // Unpopular friends are ordered by decreasing degree. Note that this is not separating
          // out the friends into two separate sets (e.g. in-country versus between country) as
          // is done in the classic world-bank layout. That requires link groups to be defined:
          //
          SortedMap<Integer, SortedSet<NetNode>> invFor = invertCountMap(forDaPop, null);
          List<NetNode> fdc = flattenDaCount(invFor);
          targets.addAll(fdc); 
          tSet.addAll(fdc);
        }
      }
    }
    lr3.finish();
 
    //
    // Handle all remaining unplaced (linked) nodes! This is purely in order of decreasing degree.
    //
    
    HashSet<NetNode> stillToPlace = new HashSet<NetNode>(allNodes);
    stillToPlace.removeAll(targets);

    LoopReporter lr4 = new LoopReporter(degree2Nodes.size(), 0, monitor, 0.0, 1.0, "progress.addingGlobalNodes");
    
    for (SortedSet<NetNode> fdeg : degree2Nodes.values()) {
      lr4.report();
      for (NetNode chkNode : fdeg) {
        if (stillToPlace.contains(chkNode)) {
          targets.add(chkNode);
        }
      }
    }
    lr4.finish();

    //
    //
    // Tag on lone nodes. If a node is by itself, but also shows up in the links,
    // we drop it:
    //
    
    HashSet<NetNode> remains = new HashSet<NetNode>(loneNodes);
    remains.removeAll(targets);
    targets.addAll(new TreeSet<NetNode>(remains));
    return (targets);
  } 
}
```
