## Supplementary material for "BioFabric Visualization of Network Alignments": BioFabric Source Code: about.html

About BioFabric


BioFabric Version 2.0.B.2 (Sun Jun 9 21:33:44 PDT 2019)

BioFabric software is Copyright (C) 2003-2019, Institute for Systems Biology.
  
Portions Copyright (C) 2018-2019 Rishi Desai.
  
It is released under the GNU Lesser General Public License, a copy
of which may be found here.

Some of the toolbar icon images in this package are covered by the
following free distribution license:

- Sun Microsystems
