## Supplementary material for "BioFabric Visualization of Network Alignments": BioFabric Source Code: LICENSE-SUN

Copyright 2000 by Sun Microsystems, Inc. All Rights Reserved.

Sun grants you ("Licensee") a non-exclusive, royalty free, license to use, 
and redistribute this software graphics artwork, as individual graphics 
or as a collection, as part of software code or programs that you develop, 
provided that i) this copyright notice and license accompany the software 
graphics artwork; and ii) you do not utilize the software graphics artwork 
in a manner which is disparaging to Sun. Unless enforcement is prohibited 
by applicable law, you may not modify the graphics, and must use them true 
to color and unmodified in every way.

This software graphics artwork is provided "AS IS," without a warranty of 
any kind. ALL EXPRESS OR IMPLIED CONDITIONS, REPRESENTATIONS AND WARRANTIES, 
INCLUDING ANY IMPLIED WARRANTY OF MERCHANTABILITY, FITNESS FOR A PARTICULAR 
PURPOSE OR NON-INFRINGEMENT, ARE HEREBY EXCLUDED. SUN AND ITS LICENSORS 
SHALL NOT BE LIABLE FOR ANY DAMAGES SUFFERED BY LICENSEE AS A RESULT OF 
USING, MODIFYING OR DISTRIBUTING THE SOFTWARE GRAPHICS ARTWORK.

IN NO EVENT WILL SUN OR ITS LICENSORS BE LIABLE FOR ANY LOST REVENUE, 
PROFIT OR DATA, OR FOR DIRECT, INDIRECT, SPECIAL, CONSEQUENTIAL, 
INCIDENTAL OR PUNITIVE DAMAGES, HOWEVER CAUSED AND REGARDLESS OF THE 
THEORY OF LIABILITY, ARISING OUT OF THE USE OF OR INABILITY TO USE 
SOFTWARE GRAPHICS ARTWORK, EVEN IF SUN HAS BEEN ADVISED OF THE 
POSSIBILITY OF SUCH DAMAGES.

If any of the above provisions are held to be in violation of applicable 
law, void, or unenforceable in any jurisdiction, then such provisions 
are waived to the extent necessary for this Disclaimer to be otherwise 
enforceable in such jurisdiction.
