## Supplementary material for "BioFabric Visualization of Network Alignments": BioFabric Source Code: AnnotationSetImpl.java

```
/*
**    Copyright (C) 2003-2018 Institute for Systems Biology 
**                            Seattle, Washington, USA. 
**
**    This library is free software; you can redistribute it and/or
**    modify it under the terms of the GNU Lesser General Public
**    License as published by the Free Software Foundation; either
**    version 2.1 of the License, or (at your option) any later version.
**
**    This library is distributed in the hope that it will be useful,
**    but WITHOUT ANY WARRANTY; without even the implied warranty of
**    MERCHANTABILITY or FITNESS FOR A PARTICULAR PURPOSE.  See the GNU
**    Lesser General Public License for more details.
**
**    You should have received a copy of the GNU Lesser General Public
**    License along with this library; if not, write to the Free Software
**    Foundation, Inc., 59 Temple Place, Suite 330, Boston, MA  02111-1307  USA
*/

package org.systemsbiology.biofabric.model;

import java.io.IOException;
import java.io.PrintWriter;
import java.util.Collections;
import java.util.Iterator;
import java.util.List;
import java.util.SortedSet;
import java.util.TreeMap;
import java.util.TreeSet;


import org.systemsbiology.biofabric.api.model.AnnotationSet;
import org.systemsbiology.biofabric.api.model.AnnotsForPos;
import org.systemsbiology.biofabric.api.model.Annot;
import org.systemsbiology.biofabric.api.io.AttributeExtractor;
import org.systemsbiology.biofabric.api.io.CharacterEntityMapper;
import org.systemsbiology.biofabric.api.io.Indenter;
import org.systemsbiology.biofabric.api.layout.AnnotColorSource;
import org.systemsbiology.biofabric.api.parser.AbstractFactoryClient;
import org.systemsbiology.biofabric.api.parser.GlueStick;
import org.systemsbiology.biofabric.api.util.MinMax;
import org.systemsbiology.biofabric.io.FabricFactory;
import org.xml.sax.Attributes;

/****************************************************************************
**
** Used for annotating contiguous sets of nodes
*/

public class AnnotationSetImpl implements AnnotationSet, Cloneable, Iterable<Annot> {
  private TreeSet<Annot> annots_;
 
  public AnnotationSetImpl() {
    annots_ = new TreeSet<Annot>();
  }
  
  @Override
  public AnnotationSetImpl clone() {
    try {
      AnnotationSetImpl retval = (AnnotationSetImpl)super.clone();
      retval.annots_ = new TreeSet<Annot>();
      for (Annot annot : this.annots_) {
        retval.annots_.add(((AnnotImpl)annot).clone());
      }
      return (retval);
    } catch (CloneNotSupportedException cnse) {
      throw new IllegalStateException();
    }
  }    
 
  public Iterator<Annot> iterator() {
    return (annots_.iterator());
  }
  
  public int size() {
    return (annots_.size());
  }
  
  public void addAnnot(Annot an) {
    annots_.add(an);
    return;
  }
  
  public void fillAnnots(AnnotsForPos fillIt, Integer whereObj) {
    for (Annot an : annots_) {
      if (an.getRange().contained(whereObj)) {
        fillIt.addAnnot(an);
      }
    }
    return;
  }
 
  /***************************************************************************
  **
  ** Class to hold all the annotations relevant for a given position (row or column)
  */  
  
  public static class AnnotsForPosImpl implements AnnotsForPos {
    private TreeMap<Integer, SortedSet<Annot>> perLayers_;
 
    public AnnotsForPosImpl() {
      perLayers_ = new TreeMap<Integer, SortedSet<Annot>>(Collections.reverseOrder());
      perLayers_.put(Integer.valueOf(0), new TreeSet<Annot>());
    }
    
    public void addAnnot(Annot toAdd) {
      int layer = toAdd.getLayer();
      SortedSet<Annot> forLayer = perLayers_.get(Integer.valueOf(layer));
      if (forLayer == null) {
        forLayer = new TreeSet<Annot>();
        perLayers_.put(Integer.valueOf(layer), forLayer);
      }
      forLayer.add(toAdd);
      return;
    }

    public void clear() {
      for (Integer layer : perLayers_.keySet()) {
        perLayers_.get(layer).clear();
      }
      return;
    }
    
    public void displayStrings(List<String> fill) {
      for (Integer layer : perLayers_.keySet()) {
        SortedSet<Annot> forLay = perLayers_.get(layer);
        for (Annot ant : forLay) {
          fill.add(ant.getName());
        }
      }
      return;
    }
  }
  
  /***************************************************************************
  **
  ** Class to hold a single annotation
  */  
  
  public static class AnnotImpl implements Annot, Cloneable {
    private String tag_;
    private MinMax range_;
    private int layer_;
    private AnnotColorSource.AnnotColor color_;
   
    public AnnotImpl(String tag, int startPos, int endPos, int layer, String colorName) {
      if ((startPos > endPos) || (layer < 0)) {
        throw new IllegalArgumentException();
      }
      tag_ = tag;
      range_ = new MinMax(startPos, endPos);
      layer_ = layer;
      color_ = (colorName == null) ? null : AnnotColorSource.AnnotColor.getColor(colorName);
    }
    
    @Override
    public AnnotImpl clone() {
      try {
        AnnotImpl retval = (AnnotImpl)super.clone();
        retval.range_ = this.range_.clone();
        return (retval);
      } catch (CloneNotSupportedException cnse) {
        throw new IllegalStateException();
      }
    }    
   
    public String getName() {
      return (tag_);
    } 
    
    public MinMax getRange() {
      return (range_);
    }
    
    public int getLayer() {
      return (layer_);
    } 
    
    // May be null if user wants automatic cycling:
    public AnnotColorSource.AnnotColor getColor() {
      return (color_);
    } 

    @Override
    public int hashCode() {
      return (tag_.hashCode() + range_.hashCode() + layer_);
    }

    @Override
    public String toString() {
      String retval = "Annot name = " + tag_ + " range = " + range_ + " layer = " + layer_;
      return ((color_ == null) ? retval : retval + "color = " + color_);
    }
     
    @Override
    public boolean equals(Object other) {    
      if (other == null) {
        return (false);
      }
      if (other == this) {
        return (true);
      }
      if (!(other instanceof Annot)) {
        return (false);
      }
      Annot otherAnnot = (Annot)other;
    
      if (!this.tag_.equalsIgnoreCase(otherAnnot.getName())) {
        return (false);
      }
      if (!this.range_.equals(otherAnnot.getRange())) {
        return (false);
      }
      
      if (this.color_ == null) {
        if (otherAnnot.getColor() != null) {
          return (false);
        }
      } else {
        if (otherAnnot.getColor() == null) {
          return (false);
        } else {
          if (!this.color_.equals(otherAnnot.getColor())) {
            return (false);
          }
        }
      }
      return (this.layer_ == otherAnnot.getLayer());
    }  

    public int compareTo(Annot otherAnnot) {
      if (this.equals(otherAnnot)) {
        return (0);
      }
      
      int startDiff = this.range_.min - otherAnnot.getRange().min;
      if (startDiff != 0) {
        return (startDiff);
      }
      
      int endDiff = this.range_.max - otherAnnot.getRange().min;
      if (endDiff != 0) {
        return (endDiff);
      }
      
      int layerDiff = this.layer_ - otherAnnot.getLayer();
      if (layerDiff != 0) {
        return (layerDiff);
      }
      
      if (this.color_ == null) {
        if (otherAnnot.getColor() != null) {
          return (-1);
        }
      } else {
        if (otherAnnot.getColor() == null) {
          return (1);
        } else {
          int compval = this.color_.compareTo(otherAnnot.getColor());
          if (compval != 0) {
            return (compval);
          }
        }
      }
       
      return (this.tag_.compareToIgnoreCase(otherAnnot.getName()));
    }
    
    /***************************************************************************
    **
    ** Dump an annotation using XML
    */
     
    public void writeXML(PrintWriter out, Indenter ind) {    
      ind.indent();
      out.print("<annot tag=\"");
      out.print(CharacterEntityMapper.mapEntities(tag_, false));
      out.print("\" start=\"");
      out.print(range_.min);
      out.print("\" end=\"");
      out.print(range_.max); 
      out.print("\" layer=\"");
      out.print(layer_);
      if (color_ != null) {
        out.print("\" color=\"");
        out.print(color_.getName());
      }
      out.println("\" />");
      return;
    }
  }
  
  public static class AnnotsWorker extends AbstractFactoryClient {
    private String tag_;
  
    public AnnotsWorker(FabricFactory.FactoryWhiteboard whiteboard, String tag) {
      super(whiteboard);
      tag_ = tag;
      myKeys_.add(tag_);
      installWorker(new AnnotWorker(whiteboard), new MyAnnotGlue());
    }
    
    protected Object localProcessElement(String elemName, Attributes attrs) throws IOException {
      Object retval = null;
      if (elemName.equals(tag_)) {
        FabricFactory.FactoryWhiteboard board = (FabricFactory.FactoryWhiteboard)this.sharedWhiteboard_;
        board.currAnnots = new AnnotationSetImpl();
        retval = board.currAnnots;
      }
      return (retval);     
    }
  }

  public static class MyAnnotGlue implements GlueStick {
    public Object glueKidToParent(Object kidObj, AbstractFactoryClient parentWorker, 
                                  Object optionalArgs) throws IOException {
      FabricFactory.FactoryWhiteboard board = (FabricFactory.FactoryWhiteboard)optionalArgs;
      board.currAnnots.addAnnot(board.annot);
      return (null);
    }
  }  
  public static class AnnotWorker extends AbstractFactoryClient {
     
    public AnnotWorker(FabricFactory.FactoryWhiteboard board) {
      super(board); 
      myKeys_.add("annot");
    }
     
    protected Object localProcessElement(String elemName, Attributes attrs) throws IOException {
      Object retval = null;
      if (elemName.equals("annot")) {
        FabricFactory.FactoryWhiteboard board = (FabricFactory.FactoryWhiteboard)this.sharedWhiteboard_;
        board.annot = buildFromXML(elemName, attrs);
        retval = board.annot;
      }
      return (retval);     
    }
    
    private Annot buildFromXML(String elemName, Attributes attrs) throws IOException {  
      String name = AttributeExtractor.extractAttribute(elemName, attrs, "annot", "tag", true);
      String minStr = AttributeExtractor.extractAttribute(elemName, attrs, "annot", "start", true);
      String maxStr = AttributeExtractor.extractAttribute(elemName, attrs, "annot", "end", true);
      String layerStr = AttributeExtractor.extractAttribute(elemName, attrs, "annot", "layer", true);
      String colorStr = AttributeExtractor.extractAttribute(elemName, attrs, "annot", "color", false);
   
      name = CharacterEntityMapper.unmapEntities(name, false);
      try {
        int min = Integer.valueOf(minStr).intValue();
        int max = Integer.valueOf(maxStr).intValue();
        int layer = Integer.valueOf(layerStr).intValue();
        Annot retval = new AnnotImpl(name, min, max, layer, colorStr);
        return (retval);
      } catch (NumberFormatException nfex) {
        throw new IOException();
      }      
    }
  }
}
```
