## Supplementary material for "BioFabric Visualization of Network Alignments": BioFabric Source Code: BioFabricNetwork.java

```
/*
**    Copyright (C) 2003-2018 Institute for Systems Biology 
**                            Seattle, Washington, USA. 
**
**    This library is free software; you can redistribute it and/or
**    modify it under the terms of the GNU Lesser General Public
**    License as published by the Free Software Foundation; either
**    version 2.1 of the License, or (at your option) any later version.
**
**    This library is distributed in the hope that it will be useful,
**    but WITHOUT ANY WARRANTY; without even the implied warranty of
**    MERCHANTABILITY or FITNESS FOR A PARTICULAR PURPOSE.  See the GNU
**    Lesser General Public License for more details.
**
**    You should have received a copy of the GNU Lesser General Public
**    License along with this library; if not, write to the Free Software
**    Foundation, Inc., 59 Temple Place, Suite 330, Boston, MA  02111-1307  USA
*/

package org.systemsbiology.biofabric.model;

import java.io.IOException;
import java.io.PrintWriter;
import java.util.ArrayList;
import java.util.TreeSet;
import java.util.HashMap;
import java.util.HashSet;
import java.util.Iterator;
import java.util.List;
import java.util.Map;
import java.util.Set;
import java.util.SortedMap;
import java.util.TreeMap;

import org.xml.sax.Attributes;

import org.systemsbiology.biofabric.analysis.Link;
import org.systemsbiology.biofabric.api.io.AttributeExtractor;
import org.systemsbiology.biofabric.api.io.AttributeKey;
import org.systemsbiology.biofabric.api.io.BuildData;
import org.systemsbiology.biofabric.api.io.CharacterEntityMapper;
import org.systemsbiology.biofabric.api.io.Indenter;
import org.systemsbiology.biofabric.api.layout.AnnotColorSource;
import org.systemsbiology.biofabric.api.layout.DefaultEdgeLayout;
import org.systemsbiology.biofabric.api.layout.EdgeLayout;
import org.systemsbiology.biofabric.api.layout.LayoutCriterionFailureException;
import org.systemsbiology.biofabric.api.layout.NodeLayout;
import org.systemsbiology.biofabric.api.model.Annot;
import org.systemsbiology.biofabric.api.model.AnnotationSet;
import org.systemsbiology.biofabric.api.model.AugRelation;
import org.systemsbiology.biofabric.api.model.NetLink;
import org.systemsbiology.biofabric.api.model.NetNode;
import org.systemsbiology.biofabric.api.model.Network;
import org.systemsbiology.biofabric.api.parser.AbstractFactoryClient;
import org.systemsbiology.biofabric.api.parser.GlueStick;
import org.systemsbiology.biofabric.api.util.MinMax;
import org.systemsbiology.biofabric.api.util.NID;
import org.systemsbiology.biofabric.api.util.UniqueLabeller;
import org.systemsbiology.biofabric.api.worker.AsynchExitRequestException;
import org.systemsbiology.biofabric.api.worker.BTProgressMonitor;
import org.systemsbiology.biofabric.api.worker.LoopReporter;
import org.systemsbiology.biofabric.io.BuildDataImpl;
import org.systemsbiology.biofabric.io.FabricFactory;
import org.systemsbiology.biofabric.plugin.BioFabricToolPlugIn;
import org.systemsbiology.biofabric.plugin.BioFabricToolPlugInData;
import org.systemsbiology.biofabric.plugin.PlugInManager;
import org.systemsbiology.biofabric.ui.FabricColorGenerator;
import org.systemsbiology.biofabric.ui.FabricDisplayOptions;
import org.systemsbiology.biofabric.ui.FabricDisplayOptionsManager;
import org.systemsbiology.biofabric.util.DataUtil;
import org.systemsbiology.biofabric.util.UiUtil;

/****************************************************************************
**
** This is the Network model.
*/

public class BioFabricNetwork implements Network {
  
  ////////////////////////////////////////////////////////////////////////////
  //
  // PRIVATE CONSTANTS
  //
  //////////////////////////////////////////////////////////////////////////// 

  ////////////////////////////////////////////////////////////////////////////
  //
  // PUBLIC CONSTANTS
  //
  //////////////////////////////////////////////////////////////////////////// 
                 
  ////////////////////////////////////////////////////////////////////////////
  //
  // PRIVATE INSTANCE MEMBERS
  //
  ////////////////////////////////////////////////////////////////////////////
   
  //
  // For mapping of selections:
  //
  
  private HashMap<Integer, NetNode> rowToTargID_;
  private int rowCount_;
  
  //
  // Link and node definitions:
  //
  
  private TreeMap<Integer, LinkInfo> fullLinkDefs_;
  private TreeMap<Integer, Integer> nonShadowedLinkMap_;
  private HashMap<NetNode, NodeInfo> nodeDefs_;
  
  //
  // Grouping for links:
  //
  
  private List<String> linkGrouping_;
  private boolean showLinkGroupAnnotations_;
  
  //
  // Columns assignments, shadow and non-shadow states:
  //
 
  private ColumnAssign normalCols_;
  private ColumnAssign shadowCols_;
  
  private FabricColorGenerator colGen_;
  
  
  //
  // Current Link Layout Mode, either PER_NODE or PER_NETWORK
  // Default value is PER_NODE
  //

  private Network.LayoutMode layoutMode_;
  
  private UniqueLabeller nodeIDGenerator_;
  
  private AnnotationSet nodeAnnot_;
  private Map<Boolean, AnnotationSet> linkAnnots_;

  private PlugInManager pMan_;
  
  ////////////////////////////////////////////////////////////////////////////
  //
  // PUBLIC CONSTRUCTORS
  //
  ////////////////////////////////////////////////////////////////////////////

  /***************************************************************************
  **
  ** Constructor
  */

  public BioFabricNetwork(BuildData rbd, PlugInManager pMan, BTProgressMonitor monitor) 
  		throws AsynchExitRequestException, LayoutCriterionFailureException {
  	nodeIDGenerator_ = new UniqueLabeller();
  	BuildDataImpl bd = (BuildDataImpl)rbd;
  	pMan_ = pMan;
  	layoutMode_ = Network.LayoutMode.UNINITIALIZED_MODE;
  	BuildDataImpl.BuildMode mode = bd.getMode();
    nodeAnnot_ = new AnnotationSetImpl();
    Map<Boolean, AnnotationSet> linkAnnots_ = new HashMap<Boolean, AnnotationSet>();
    linkAnnots_.put(Boolean.TRUE, new AnnotationSetImpl());
    linkAnnots_.put(Boolean.FALSE, new AnnotationSetImpl());
    switch (mode) {
      case DEFAULT_LAYOUT:  
      case REORDER_LAYOUT:
      case CLUSTERED_LAYOUT:
      case GROUP_PER_NODE_CHANGE:
      case GROUP_PER_NETWORK_CHANGE:
      case NODE_ATTRIB_LAYOUT:
      case LINK_ATTRIB_LAYOUT:
      case NODE_CLUSTER_LAYOUT:
      case CONTROL_TOP_LAYOUT: 
      case HIER_DAG_LAYOUT:
      case SET_LAYOUT:
      case WORLD_BANK_LAYOUT:
        standardBuildDataInit(bd);
        transferRelayoutBuildData(bd);
        relayoutNetwork(bd, monitor);
        // Some layouts can ask to force shadows to be activated:
		    if (bd.getTurnOnShadows()) {
		      FabricDisplayOptions dops = FabricDisplayOptionsManager.getMgr().getDisplayOptions();
				  dops.setDisplayShadows(true);
		    }
        break;
      case BUILD_FROM_PLUGIN:      	
        standardBuildDataInit(bd);
        transferRelayoutBuildData(bd);
        processLinks(bd, monitor);
        // Some layouts can ask to force shadows to be activated:
		    if (bd.getTurnOnShadows()) {
		      FabricDisplayOptions dops = FabricDisplayOptionsManager.getMgr().getDisplayOptions();
				  dops.setDisplayShadows(true);
		    }
        break;
      case BUILD_FOR_SUBMODEL:
        colGen_ = bd.fullNet.colGen_;
        this.linkGrouping_ = new ArrayList<String>(bd.fullNet.linkGrouping_);
        this.showLinkGroupAnnotations_ = bd.fullNet.showLinkGroupAnnotations_;
        this.layoutMode_ = bd.fullNet.layoutMode_;
        fillSubModel(bd.fullNet, bd.subNodes, bd.subLinks);
        break;
      case BUILD_FROM_XML:
      case SHADOW_LINK_CHANGE:
        standardBuildDataTransfer(bd.getExistingNetwork());
        break;
      case BUILD_FROM_SIF:
        standardBuildDataInit(bd);
        linkGrouping_ = new ArrayList<String>();
        showLinkGroupAnnotations_ = false;
        layoutMode_ = LayoutMode.UNINITIALIZED_MODE;
        colGen_ = bd.getColorGen();
        processLinks(bd, monitor);
        break;
      default:
        throw new IllegalArgumentException();
    }
  }

  ////////////////////////////////////////////////////////////////////////////
  //
  // PUBLIC METHODS
  //
  ////////////////////////////////////////////////////////////////////////////

  /***************************************************************************
  ** 
  ** Build support
  */
  
  private void standardBuildDataInit(BuildData bd) {
    this.normalCols_ = new ColumnAssign();
    this.shadowCols_ = new ColumnAssign();
    this.rowToTargID_ = new HashMap<Integer, NetNode>();
    this.fullLinkDefs_ = new TreeMap<Integer, LinkInfo>();
    this.nonShadowedLinkMap_ = new TreeMap<Integer, Integer>();
    this.nodeDefs_ = new HashMap<NetNode, NodeInfo>();
    return;
  }
  
  /***************************************************************************
  ** 
  ** Build support
  */
  
  private void transferRelayoutBuildData(BuildDataImpl bd) { 
    this.linkGrouping_ = new ArrayList<String>(bd.getGroupOrder());
    this.showLinkGroupAnnotations_ = bd.getShowLinkGroupAnnotations();
    this.colGen_ = bd.getColorGen();
    this.layoutMode_ = bd.getGroupOrderMode();
    return;
  }
 
  /***************************************************************************
  ** 
  ** Build support
  */
  
  private void standardBuildDataTransfer(BioFabricNetwork built) {
    this.normalCols_ = built.normalCols_;
    this.shadowCols_ = built.shadowCols_;
    this.rowToTargID_ = built.rowToTargID_; 
    this.fullLinkDefs_ = built.fullLinkDefs_;
    this.nonShadowedLinkMap_ = built.nonShadowedLinkMap_;
    this.nodeDefs_ = built.nodeDefs_;
    this.colGen_ = built.colGen_;
    this.rowCount_ = built.rowCount_;
    this.linkGrouping_ = built.linkGrouping_;
    this.showLinkGroupAnnotations_ = built.showLinkGroupAnnotations_;
    this.layoutMode_ = built.layoutMode_;
    this.nodeAnnot_ = built.nodeAnnot_;
    this.linkAnnots_= built.linkAnnots_;
 
    return;
  }
  
  /***************************************************************************
  ** 
  ** Build support
  */
  
  public UniqueLabeller getGenerator() { 
    return (nodeIDGenerator_);
  }
  
  /***************************************************************************
  ** 
  ** Build support
  */
  
  public FabricColorGenerator getColorGenerator() { 
    return (colGen_);
  }
  
  /***************************************************************************
  ** 
  ** Build support
  */
  
  public List<String> getLinkGrouping() { 
    return (linkGrouping_);
  }

  /***************************************************************************
  ** 
  ** Set node annotations
  */

  public void setNodeAnnotations(AnnotationSet aSet) {
    nodeAnnot_ = aSet;
    return;
  }
  
  /***************************************************************************
  ** 
  ** Set link annotations
  */

  public void setLinkAnnotations(AnnotationSet aSet, boolean forShadow) {
    if (linkAnnots_ == null) {
      linkAnnots_ = new HashMap<Boolean, AnnotationSet>();
    }
    linkAnnots_.put(Boolean.valueOf(forShadow), aSet);
    return;
  }
  
  /***************************************************************************
  ** 
  ** Get node annotations
  */

  public AnnotationSet getNodeAnnotations() {
    return (nodeAnnot_);
  }
  
  /***************************************************************************
  ** 
  ** Get link annotations
  */

  public AnnotationSet getLinkAnnotations(boolean forShadow) {
    return ((linkAnnots_ == null) ? null : linkAnnots_.get(Boolean.valueOf(forShadow)));
  }
 
  /***************************************************************************
  ** 
  ** Get map from normalized name to IDs (Moving to Cytoscape SUIDs, there
  *  can be multiple nodes for one name)
  */

  public Map<String, Set<NetNode>> getNormNameToIDs() {
  	HashMap<String, Set<NetNode>> retval = new HashMap<String, Set<NetNode>>();
  	Iterator<NetNode> kit = nodeDefs_.keySet().iterator();
  	while (kit.hasNext()) {
  		NetNode key = kit.next();
  		NodeInfo forKey = nodeDefs_.get(key);
  		String nameNorm = DataUtil.normKey(forKey.getNodeName());
  		Set<NetNode> forName = retval.get(nameNorm);
  		if (forName == null) {
  			forName = new HashSet<NetNode>();
  			retval.put(nameNorm, forName);
  		}
  		forName.add(key);
  	}
    return (retval);
  }
  
  /***************************************************************************
  ** 
  ** Install link grouping
  */

  public void installLinkGroups(List<String> linkTagList) {    
    linkGrouping_ = new ArrayList<String>(linkTagList);
    return;
  }
  
  /***************************************************************************
  ** 
  ** Set if link groups get annotations
  */

  public void setShowLinkGroupAnnotations(boolean show) {    
    showLinkGroupAnnotations_ = show;
    return;
  }
  
  /***************************************************************************
  ** 
  ** Get if link groups get annotations
  */

  public boolean getShowLinkGroupAnnotations() {    
    return (showLinkGroupAnnotations_);
  }

  /***************************************************************************
  ** 
  ** Get link grouping
  */

  public List<String> getLinkGroups() {    
    return (linkGrouping_);
  }

  /***************************************************************************
  ** 
  ** Given an attribute list giving node order, confirm it is valid:
  */

  public boolean checkNewNodeOrder(Map<AttributeKey, String> nodeAttributes) { 
    
    //
    // All existing targets must have a row, and all existing
    // rows need a target assigned!
    //
    
    HashSet<String> normedNames = new HashSet<String>();
    Iterator<NetNode> rttvit = rowToTargID_.values().iterator();
    while (rttvit.hasNext()) {
    	NetNode key = rttvit.next();
    	NodeInfo ni = nodeDefs_.get(key);
      normedNames.add(DataUtil.normKey(ni.getNodeName()));
    }
     
    HashSet<String> normedKeys = new HashSet<String>();
    Iterator<AttributeKey> akit = nodeAttributes.keySet().iterator();
    while (akit.hasNext()) {
    	AttributeKey key = akit.next();
      normedKeys.add(DataUtil.normKey(key.toString()));
    }
  
    if (!normedNames.equals(normedKeys)) {
      return (false);
    }
    
    TreeSet<Integer> asInts = new TreeSet<Integer>();
    Iterator<String> nrvit = nodeAttributes.values().iterator();
    while (nrvit.hasNext()) {
      String asStr = nrvit.next();
      try {
        asInts.add(Integer.valueOf(asStr));
      } catch (NumberFormatException nfex) {
        return (false);
      }
    }
    
    if (!asInts.equals(new TreeSet<Integer>(rowToTargID_.keySet()))) {
      return (false);
    }
    
    return (true);
  }
  
  
  /***************************************************************************
  ** 
  ** Given an attribute list giving link order, confirm it is valid, get back
  ** map to integer values with directed tag filled in!
  **
  ** FIX ME: A few problems need to be addressed.  First, there is no
  ** decent error messaging about what is missing or duplicated.  Second,
  ** we require shadow links to ALWAYS be specified, even if the user does
  ** not care.... 
  */

  public SortedMap<Integer, NetLink> checkNewLinkOrder(Map<AttributeKey, String> linkRows) { 
    
    //
    // Recover the mapping that tells us what link relationships are
    // directed:
    //
    
    HashMap<AugRelation, Boolean> relDir = new HashMap<AugRelation, Boolean>();
    Set<NetLink> allLinks = getAllLinks(true);
    Iterator<NetLink> alit = allLinks.iterator();
    while (alit.hasNext()) {
      NetLink link = alit.next();
      AugRelation rel = link.getAugRelation();
      boolean isDir = link.isDirected();
      Boolean myVal = new Boolean(isDir);
      Boolean currVal = relDir.get(rel);
      if (currVal != null) {
        if (!currVal.equals(myVal)) {
          throw new IllegalStateException();
        }
      } else {
        relDir.put(rel, myVal);
      }
    }
    
    //
    // Create a map that takes column Integer to a correctly
    // directed copy of the Fabric link:
    //
    
    TreeMap<Integer, NetLink> dirMap = new TreeMap<Integer, NetLink>();
    Iterator<AttributeKey> lrit = linkRows.keySet().iterator();
    while (lrit.hasNext()) {
      FabricLink link = (FabricLink)lrit.next();
      String colNumStr = linkRows.get(link);
      FabricLink dirCopy = link.clone();
      Boolean isDirected = relDir.get(dirCopy.getAugRelation());
      dirCopy.installDirection(isDirected);
      try {
        dirMap.put(Integer.valueOf(colNumStr), dirCopy);
      } catch (NumberFormatException nfex) {
        return (null);
      }
    }
    
    // Ordered set of all our existing links:
    TreeSet<NetLink> alks = new TreeSet<NetLink>(allLinks);
    // Ordered set of guys we have been handed:
    TreeSet<NetLink> dmvs = new TreeSet<NetLink>(dirMap.values());
    
    // Has to be the case that the link definitions are 1:1 and
    // onto, or we have an error:
    if (!alks.equals(dmvs)) {
      return (null);
    }
  
    //
    // Has to be the case that all columns are also 1:1 and onto:
    //
    
    TreeSet<Integer> ldks = new TreeSet<Integer>(fullLinkDefs_.keySet());
    TreeSet<Integer> dmks = new TreeSet<Integer>(dirMap.keySet());
    
    if (!ldks.equals(dmks)) {
      return (null);
    }
    
    return (dirMap);
  }
  
  /***************************************************************************
  **
  ** Get all links
  */
  
  public Set<NetLink> getAllLinks(boolean withShadows) {  
    HashSet<NetLink> allLinks = new HashSet<NetLink>();
    Iterator<Integer> ldit = fullLinkDefs_.keySet().iterator();
    while (ldit.hasNext()) {
      Integer col = ldit.next();
      LinkInfo li = getLinkDefinition(col, true);  // just get everybody...
      FabricLink link = li.getLink();
      if (withShadows || !link.isShadow()) {
        allLinks.add(link);
      }
    }
    return (allLinks);
  }
 
  /***************************************************************************
  **
  ** Get ordered linkInfo iterator
  */
  
  public Iterator<Integer> getOrderedLinkInfo(boolean withShadows) {  
    return ((withShadows) ? fullLinkDefs_.keySet().iterator() : nonShadowedLinkMap_.keySet().iterator());
  }
  
  /***************************************************************************
  ** 
  ** Process a link set
  */

  private void processLinks(BuildData bd, BTProgressMonitor monitor) 
  		throws AsynchExitRequestException, LayoutCriterionFailureException {
    //
    // Note the allLinks Set has pruned out duplicates and synonymous non-directional links
    //

  	BuildDataImpl rbd = (BuildDataImpl)bd;
  	NodeLayout layout = rbd.getNodeLayout();
    boolean nlok = layout.criteriaMet(rbd, monitor); // No?? throws LayoutCriterionFailureException
    if (!nlok) {
      throw new IllegalStateException(); // Should not happen, failure throws exception
    }
  	 
    List<NetNode> targetIDs = layout.doNodeLayout(rbd, null, monitor);
    
    //
    // Now have the ordered list of targets we are going to display.
    //

    fillNodesFromOrder(targetIDs, rbd.getColorGen(), rbd.clustAssign, monitor);

    //
    // This now assigns the link to its column. The RelayoutBuildData gives us
    // the algorithm to use; typically we order them so that the shortest vertical 
    // link is drawn first.
    // Before we do layout, we give the algorithm a chance to e.g. install link groups
    // that will be used in the link layout.
    //
    
    EdgeLayout edgeLayout = rbd.getEdgeLayout();
    edgeLayout.preProcessEdges(rbd, monitor);
    
    edgeLayout.layoutEdges(rbd, monitor);
    specifiedLinkToColumn(rbd.getColorGen(), rbd.getLinkOrder(), false, monitor);
    
    //
    // If node annotations, link annotations, or group order has been set, install those now:
    //
    
    nodeAnnot_ = rbd.getNodeAnnotations();
    linkAnnots_ = rbd.getLinkAnnotations();
    linkGrouping_ = rbd.getGroupOrder();
    layoutMode_ = rbd.getGroupOrderMode();
    

    //
    // Determine the start & end of each target row needed to handle the incoming
    // and outgoing links:
    //

    trimTargetRows(monitor);
        
    //
    // For the lone nodes, they are assigned into the last column:
    //
    loneNodesToLastColumn(rbd.getSingletonNodes(), monitor);
    return;
  }
  
  /***************************************************************************
  **
  ** Relayout the network!
  */
  
  private void relayoutNetwork(BuildData bd, BTProgressMonitor monitor) throws AsynchExitRequestException {
  	BuildDataImpl rbd = (BuildDataImpl)bd;
    BuildDataImpl.BuildMode mode = rbd.getMode();
    installLinkGroups(rbd.getGroupOrder());
    setShowLinkGroupAnnotations(rbd.getShowLinkGroupAnnotations());
    setLayoutMode(rbd.getGroupOrderMode());
    boolean specifiedNodeOrder = (mode == BuildDataImpl.BuildMode.NODE_ATTRIB_LAYOUT) || 
                                 (mode == BuildDataImpl.BuildMode.DEFAULT_LAYOUT) ||
                                 (mode == BuildDataImpl.BuildMode.CONTROL_TOP_LAYOUT) ||
                                 (mode == BuildDataImpl.BuildMode.HIER_DAG_LAYOUT) ||
                                 (mode == BuildDataImpl.BuildMode.SET_LAYOUT) ||
                                 (mode == BuildDataImpl.BuildMode.WORLD_BANK_LAYOUT) ||
                                 (mode == BuildDataImpl.BuildMode.NODE_CLUSTER_LAYOUT) || 
                                 (mode == BuildDataImpl.BuildMode.CLUSTERED_LAYOUT) || 
                                 (mode == BuildDataImpl.BuildMode.REORDER_LAYOUT); 

    List<NetNode> targetIDs;
    if (specifiedNodeOrder) {
      targetIDs = specifiedIDOrder(rbd.getAllNodes(), rbd.getNodeOrder());
    } else {       
      targetIDs = rbd.getExistingIDOrder();
    }
   
    //
    // Now have the ordered list of targets we are going to display.
    // Build target->row maps and the inverse:
    //

    fillNodesFromOrder(targetIDs, rbd.getColorGen(), rbd.clustAssign, monitor);

    //
    // Ordering of links:
    //
  
    SortedMap<Integer, NetLink> lor = rbd.getLinkOrder();
    Map<NetNode, Integer> nor = rbd.getNodeOrder();
    
    if ((lor == null) || lor.isEmpty() || (mode == BuildDataImpl.BuildMode.GROUP_PER_NODE_CHANGE) || (mode == BuildDataImpl.BuildMode.GROUP_PER_NETWORK_CHANGE)) {
      if ((nor == null) || nor.isEmpty()) {
        Map<NetNode, Integer> norNew = new HashMap<NetNode, Integer>();
        int numT = targetIDs.size();
        for (int i = 0; i < numT; i++) {
          NetNode targID = targetIDs.get(i);
          norNew.put(targID, Integer.valueOf(i));
        }
        rbd.setNodeOrder(norNew);
      }
      lor = (new DefaultEdgeLayout()).layoutAllEdges(rbd, monitor);
    }

    //
    // This now assigns the link to its column, based on user specification
    //
  
		specifiedLinkToColumn(rbd.getColorGen(), lor, ((mode == BuildDataImpl.BuildMode.LINK_ATTRIB_LAYOUT) || 
		  		                                         (mode == BuildDataImpl.BuildMode.NODE_CLUSTER_LAYOUT) ||
		  		                                         (mode == BuildDataImpl.BuildMode.GROUP_PER_NODE_CHANGE) ||
		  		                                         (mode == BuildDataImpl.BuildMode.GROUP_PER_NETWORK_CHANGE)), monitor);
      
    //
    // Determine the start & end of each target row needed to handle the incoming
    // and outgoing links:
    //

    trimTargetRows(monitor);
        
    //
    // For the lone nodes, they are assigned into the last column:
    //

    loneNodesToLastColumn(rbd.getSingletonNodes(), monitor);
    
    //
    // If we have node/link annotations, install them
    //
    
    nodeAnnot_ = rbd.getNodeAnnotations();
    linkAnnots_ = rbd.getLinkAnnotations();

    return;
  }
  
  /***************************************************************************
  **
  ** Get specified node ID order list from attribute map
  */
  
  private List<NetNode> specifiedIDOrder(Set<NetNode> allNodeIDs, Map<NetNode, Integer> newOrder) { 
    TreeMap<Integer, NetNode> forRetval = new TreeMap<Integer, NetNode>();
    Iterator<NetNode> rttvit = allNodeIDs.iterator();
    while (rttvit.hasNext()) {
      NetNode key = rttvit.next();
      Integer val = newOrder.get(key);
      forRetval.put(val, key);
    }
    return (new ArrayList<NetNode>(forRetval.values()));
  }
 
  /***************************************************************************
  **
  ** Get existing order
  */
  
  public List<NetNode> existingIDOrder() {  
    ArrayList<NetNode> retval = new ArrayList<NetNode>();
    Iterator<Integer> rtit = new TreeSet<Integer>(rowToTargID_.keySet()).iterator();
    while (rtit.hasNext()) {
      Integer row = rtit.next();
      NetNode nodeID = rowToTargID_.get(row);
      retval.add(nodeID);
    }
    return (retval);
  }
  
  /***************************************************************************
  **
  ** Get existing link order
  */
  
  public SortedMap<Integer, FabricLink> getExistingLinkOrder() {  
    TreeMap<Integer, FabricLink> retval = new TreeMap<Integer, FabricLink>();
    Iterator<Integer> ldit = fullLinkDefs_.keySet().iterator();
    while (ldit.hasNext()) {
      Integer col = ldit.next();
      LinkInfo li = fullLinkDefs_.get(col);
      FabricLink link = li.getLink();
      retval.put(col, link);
    }
    return (retval);
  }

  /***************************************************************************
  ** 
  ** Process a link set
  */

  private void specifiedLinkToColumn(FabricColorGenerator colGen, 
  		                               SortedMap<Integer, NetLink> linkOrder, 
  		                               boolean userSpec, BTProgressMonitor monitor) throws AsynchExitRequestException {
     
    normalCols_.columnCount = 0;
    shadowCols_.columnCount = 0;
    int numColors = colGen.getNumColors();
    Iterator<Integer> frkit = linkOrder.keySet().iterator();
   
    
    LoopReporter lr = new LoopReporter(linkOrder.size(), 20, monitor, 0.0, 1.0, "progress.linkToColumn");
    
    while (frkit.hasNext()) {
      Integer nextCol = frkit.next();
      lr.report();
      NetLink nextLink = linkOrder.get(nextCol);
      Integer[] colCounts = addLinkDef(nextLink, numColors, normalCols_.columnCount, shadowCols_.columnCount, colGen);
      shadowCols_.columnCount = colCounts[0].intValue();
      if (colCounts[1] != null) {
        normalCols_.columnCount = colCounts[1].intValue();
      }
    }
    lr.finish();
   
    setDrainZonesWithMultipleLabels(true, monitor, 0.0, 0.5);
    setDrainZonesWithMultipleLabels(false, monitor, 0.5, 1.0);
 
    return;
  }
  
  /***************************************************************************
   ** Calculates each MinMax region for every node's zero or more drain zones
  ** 
   */
  
  private void setDrainZonesWithMultipleLabels(boolean forShadow,
  		                                         BTProgressMonitor monitor, 
					                                     double startFrac, 
					                                     double endFrac) throws AsynchExitRequestException {
    
    List<LinkInfo> links = getLinkDefList(forShadow);
    
    if (links.isEmpty()) {
    	return;
    }
    
    int size = links.size() + nodeDefs_.size();
    
    String progressTag = (forShadow) ? "progress.findingDrainZonesWithShadow" : "progress.findingDrainZones";
    LoopReporter lr = new LoopReporter(size, 20, monitor, startFrac, endFrac, progressTag);

    Map<NetNode, List<DrainZone>> nodeToZones = new TreeMap<NetNode, List<DrainZone>>();

    if (links.size() == 0) {
      return;
    }
    LinkInfo current = links.get(0);  // these keep track of start of zone and zone's node
    int startIdx = 0;
    
    for (int index = 0; index <= links.size(); index++) {
      lr.report();
      if (index == links.size()) {
        
        int endIdx = links.size() - 1;
  
        MinMax mm = new MinMax(startIdx, endIdx);
        NetNode name = findZoneNode(mm, links);
  
        if (nodeToZones.get(name) == null) {
          nodeToZones.put(name, new ArrayList<DrainZone>());
        }
        nodeToZones.get(name).add(new DrainZone(mm, forShadow));
        
      } else if (! isContiguous(links.get(index), current)) {
        
        int last = index - 1;  // backtrack one position
  
        MinMax mm = new MinMax(startIdx, last);
        NetNode name = findZoneNode(mm, links);
  
        if (nodeToZones.get(name) == null) {
          nodeToZones.put(name, new ArrayList<DrainZone>());
        }
        nodeToZones.get(name).add(new DrainZone(mm, forShadow));
        
        startIdx = index;           // update the start index
        current = links.get(index); // update the current node whose zone we're calculating
      }    
    }

    for (Map.Entry<NetNode, List<DrainZone>> entry : nodeToZones.entrySet()) {     
      NodeInfo ni = getNodeDefinition(entry.getKey());
      lr.report();
      ni.setDrainZones(entry.getValue(), forShadow);
    }
    
    lr.finish();
  }
  
  /***************************************************************************
   ** Returns true if two contiguous links are part of the same drain zone
   **
   */
  
  private boolean isContiguous(LinkInfo A, LinkInfo B) {
    
    NetNode mainA;
    if (A.isShadow()) {
      mainA = getNodeIDForRow(A.bottomRow());
    } else {
      mainA = getNodeIDForRow(A.topRow());
    }
    
    NetNode mainB;
    if (B.isShadow()) {
      mainB = getNodeIDForRow(B.bottomRow());
    } else {
      mainB = getNodeIDForRow(B.topRow());
    }
    
    return (mainA.equals(mainB));
  }
  
  /***************************************************************************
   * finds the node of the drain zone with bounds = MinMax([A,B])
   */
  
  private NetNode findZoneNode(MinMax mm, List<LinkInfo> links) {
    
    LinkInfo li = links.get(mm.min); // checking min is enough

    if (li.isShadow()) {
      return (getNodeIDForRow(li.bottomRow()));
    } else {
      return (getNodeIDForRow(li.topRow()));
    }
  }
   
  /***************************************************************************
  **
  ** Dump the network using XML
  */
  
  public void writeXML(PrintWriter out, Indenter ind, BTProgressMonitor monitor, boolean forCache) throws AsynchExitRequestException {    
    ind.indent();
    
    int numNodes = rowToTargID_.size();
    int numLinks = fullLinkDefs_.size();
    int numLm = nonShadowedLinkMap_.size();
    int numNA = (nodeAnnot_ == null) ? 0 : nodeAnnot_.size();
    int numLAs = (linkAnnots_ == null) ? 0 : linkAnnots_.get(Boolean.TRUE).size();
    int numLAns = (linkAnnots_ == null) ? 0 : linkAnnots_.get(Boolean.FALSE).size();
      
    out.println("<BioFabric>");
    ind.up();
    String label = (forCache) ? "progress.cachingCurrentNetwork" : "progress.writingFile";
    LoopReporter lr = new LoopReporter(numNodes + numLinks + numLm + numNA + numLAs + numLAns , 20, monitor, 0.0, 1.0, label);   
    colGen_.writeXML(out, ind);
    
    //
    // Display options:
    //
    
    FabricDisplayOptionsManager.getMgr().writeXML(out, ind);
       
    //
    // Dump the nodes, then the links:
    //  
    
    Iterator<Integer> r2tit = (new TreeSet<Integer>(rowToTargID_.keySet())).iterator();
    ind.indent();
    out.println("<nodes>");
    ind.up();
    while (r2tit.hasNext()) {
      Integer row = r2tit.next();
      lr.report();
      NetNode nodeID = rowToTargID_.get(row);
      NodeInfo ni = getNodeDefinition(nodeID);
      ni.writeXML(out, ind, row.intValue());
    }
    ind.down().indent();
    out.println("</nodes>");
  
    if (!linkGrouping_.isEmpty()) {
      Iterator<String> lgit = linkGrouping_.iterator();
      ind.indent();
      out.print("<linkGroups mode=\"");
      out.print(layoutMode_.getText());
      out.print("\" annots=\"");
      out.print(showLinkGroupAnnotations_);
      out.println("\">");
      ind.up();
      while (lgit.hasNext()) {
        String grpTag = lgit.next();
        ind.indent();
        out.print("<linkGroup tag=\"");
        out.print(grpTag);
        out.println("\" />");
      }
      ind.down().indent();
      out.println("</linkGroups>");
    }
  
    HashMap<Integer, Integer> inverse = new HashMap<Integer, Integer>();
    Iterator<Integer> nsit = nonShadowedLinkMap_.keySet().iterator();
    while (nsit.hasNext()) {
      Integer key = nsit.next();
      lr.report();
      inverse.put(nonShadowedLinkMap_.get(key), key);
    }    
    
    Iterator<Integer> ldit = fullLinkDefs_.keySet().iterator();
    ind.indent();
    out.println("<links>");
    ind.up();
    while (ldit.hasNext()) {
      Integer col = ldit.next();
      LinkInfo li = getLinkDefinition(col, true);
      FabricLink link = li.getLink();
      lr.report();
      ind.indent();
      out.print("<link srcID=\"");
      out.print(link.getSrcNode().getNID().getNID().getInternal());
      out.print("\" trgID=\"");
      out.print(link.getTrgNode().getNID().getNID().getInternal());
      out.print("\" rel=\"");
      AugRelation augr = link.getAugRelation();
      out.print(CharacterEntityMapper.mapEntities(augr.relation, false));
      out.print("\" directed=\"");
      out.print(link.isDirected());
      out.print("\" shadow=\"");
      out.print(augr.isShadow);
      if (!augr.isShadow) {
        Integer nsCol = inverse.get(col); 
        out.print("\" column=\"");
        out.print(nsCol);
      }
      out.print("\" shadowCol=\"");
      out.print(col);     
      out.print("\" srcRow=\"");
      out.print(li.getStartRow());
      out.print("\" trgRow=\"");
      out.print(li.getEndRow());
      out.print("\" color=\"");
      out.print(li.getColorKey());
      out.println("\" />");
    }
    ind.down().indent();
    out.println("</links>");
      
    ind.indent();
    out.println("<nodeAnnotations>");
    ind.up();
    if (nodeAnnot_ != null) {
      for (Annot an : nodeAnnot_) {
        lr.report();
        ((AnnotationSetImpl.AnnotImpl)an).writeXML(out, ind);
      }
    }
    ind.down().indent();
    out.println("</nodeAnnotations>");
    
    ind.indent();
    out.println("<linkAnnotations>");
    ind.up();
    if (linkAnnots_ != null) {
      for (Annot an : linkAnnots_.get(Boolean.FALSE)) {
        lr.report();
         ((AnnotationSetImpl.AnnotImpl)an).writeXML(out, ind);
      }
    }
    ind.down().indent();
    out.println("</linkAnnotations>");
    
    ind.indent();
    out.println("<shadowLinkAnnotations>");
    ind.up();
    if (linkAnnots_ != null) {
      for (Annot an : linkAnnots_.get(Boolean.TRUE)) {
        lr.report();
         ((AnnotationSetImpl.AnnotImpl)an).writeXML(out, ind);
      }
    }
    ind.down().indent();
    out.println("</shadowLinkAnnotations>");

    //
    // Let the plugins write to XML
    
    ind.indent();
    out.println("<plugInDataSets>");
    List<String> keyList = pMan_.getOrderedToolPlugInKeys();
    for (String key : keyList) {
      BioFabricToolPlugIn plugin = pMan_.getToolPlugIn(key);
      plugin.writeXML(out, ind);
    }
    ind.indent();
    out.println("</plugInDataSets>");
    
    lr.finish();
    ind.down().indent();
    out.println("</BioFabric>"); 
    return;
  }
  
  /***************************************************************************
  **
  ** Dump a node attribute file with row assignments:
  */
  
  public void writeNOA(PrintWriter out) {    
    out.println("Node Row");
    Iterator<Integer> r2tit = new TreeSet<Integer>(rowToTargID_.keySet()).iterator();
    while (r2tit.hasNext()) {
      Integer row = r2tit.next();
      NetNode nodeID = rowToTargID_.get(row);
      NodeInfo ni = getNodeDefinition(nodeID);
      out.print(ni.getNodeName());
      out.print(" = ");
      out.println(row);
    } 
    return;
  }
  
  /***************************************************************************
  **
  ** Dump an edge attribute file with column assignments:
  */
  
  public void writeEDA(PrintWriter out) {    
    out.println("Link Column");
    Iterator<Integer> ldit = fullLinkDefs_.keySet().iterator();
    while (ldit.hasNext()) {
      Integer col = ldit.next();
      LinkInfo li = getLinkDefinition(col, true);
      FabricLink link = li.getLink();
      out.print(link.toEOAString(nodeDefs_));
      out.print(" = ");
      out.println(col);
    }
    return;
  }
    
  /***************************************************************************
  ** 
  ** Get Node Definition
  */

  public NodeInfo getNodeDefinition(NetNode targID) {
    NodeInfo node = nodeDefs_.get(targID);
    return (node);
  }
  
  /***************************************************************************
  ** 
  ** Get All Node Definition
  */

  public Map<NetNode, NodeInfo> getAllNodeDefinitions() {
    return (nodeDefs_);
  }
  
  /***************************************************************************
  ** 
  ** Link definition for column.  If no link in column (may happen for selected networks),
  ** returns null
  */

  public LinkInfo getLinkDefinition(Integer colObj, boolean forShadow) {
    if (forShadow) {
      return (fullLinkDefs_.get(colObj));
    } else {
      Integer mapped = nonShadowedLinkMap_.get(colObj);
      if (mapped != null) {
        return (fullLinkDefs_.get(mapped));
      } else {
        return (null);
      }     
    }
  }
  
  /***************************************************************************
  ** 
  ** Get Target For Column
  */

  public NetNode getTargetIDForColumn(Integer colVal, boolean forShadow) {
    ColumnAssign useCA = (forShadow) ? shadowCols_ : normalCols_;
    NetNode target = useCA.columnToTarget.get(colVal);
    return (target);
  }
  
  /***************************************************************************
  ** 
  ** Get Drain zone For Column
  */

  public NetNode getDrainForColumn(Integer colVal, boolean forShadow) {
  
    int col = colVal.intValue();
    ColumnAssign useCA = (forShadow) ? shadowCols_ : normalCols_;
    NetNode targetID = useCA.columnToTarget.get(colVal);
    NetNode sourceID = useCA.columnToSource.get(colVal);
    if (targetID != null) {
      NodeInfo nit = nodeDefs_.get(targetID);
      List<DrainZone> tdzs = nit.getDrainZones(forShadow);
      if (tdzs != null) {
        for (DrainZone tdz : tdzs)
          if ((col >= tdz.getMinMax().min) && (col <= tdz.getMinMax().max)) {
            return (targetID);
          }
      }
    }
    if (sourceID != null) {
      NodeInfo nis = nodeDefs_.get(sourceID);
      List<DrainZone> sdzs = nis.getDrainZones(forShadow);
      if (sdzs != null) {
        for (DrainZone sdz : sdzs)
          if ((col >= sdz.getMinMax().min) && (col <= sdz.getMinMax().max)) {
            return (sourceID);
          }
      }
    }
  
    return (null);
  }
  
  /***************************************************************************
  ** 
  ** Get Source For Column
  */

  public NetNode getSourceIDForColumn(Integer colVal, boolean forShadow) {
    ColumnAssign useCA = (forShadow) ? shadowCols_ : normalCols_;
    NetNode source = useCA.columnToSource.get(colVal);
    return (source);
  }
  
  /***************************************************************************
  ** 
  ** Get node for row
  */

  public NetNode getNodeIDForRow(Integer rowObj) {
    NetNode node = rowToTargID_.get(rowObj);
    return (node);
  }
  
  /***************************************************************************
  ** 
  ** Get link count
  */

  public int getLinkCount(boolean forShadow) {
    return ((forShadow) ? fullLinkDefs_.size() : nonShadowedLinkMap_.size());
  } 
  
  /***************************************************************************
  ** 
  ** Get column count
  */

  public int getColumnCount(boolean forShadow) {
    ColumnAssign useCA = (forShadow) ? shadowCols_ : normalCols_;
    return (useCA.columnCount);
  } 
  
  /***************************************************************************
  ** 
  ** Get node count
  */

  public int getNodeCount() {
    return (rowCount_);
  } 
  
  /***************************************************************************
  ** 
  ** Get Row Count
  */

  public int getRowCount() {
    return (rowCount_);
  }
  
  /***************************************************************************
  **
  ** Stash plugin data for extraction
  */
  
  public void stashPluginData(String keyword, BioFabricToolPlugInData data) {
    
  }

  /***************************************************************************
  **
  ** Pull plugin data for extraction
  */
  
  public BioFabricToolPlugInData providePluginData(String keyword) {
     return (null);
  }

  /***************************************************************************
  ** 
  ** Get node defs
  */

  public List<NodeInfo> getNodeDefList() {
    return (new ArrayList<NodeInfo>(nodeDefs_.values()));
  } 
  
  /***************************************************************************
  ** 
  ** Get all node names
  */

  public Set<NetNode> getNodeSetIDs() {
  	HashSet<NetNode> retval = new HashSet<NetNode>();
  	Iterator<NodeInfo> nsit = nodeDefs_.values().iterator();
    while (nsit.hasNext()) {
    	NodeInfo ni = nsit.next();
      retval.add(new FabricNode(ni.getNodeID(), ni.getNodeName()));
    }
    return (retval);
  }  
  
  /***************************************************************************
  ** 
  ** Get link defs
  */

  public List<LinkInfo> getLinkDefList(boolean forShadow) {
    if (forShadow) {
      return (new ArrayList<LinkInfo>(fullLinkDefs_.values()));
    } else {
      ArrayList<LinkInfo> retval = new ArrayList<LinkInfo>();
      Iterator<Integer> nsit = nonShadowedLinkMap_.keySet().iterator();
      while (nsit.hasNext()) {
        Integer linkID = nsit.next();
        Integer mappedID = nonShadowedLinkMap_.get(linkID);
        retval.add(fullLinkDefs_.get(mappedID));
      }
      return (retval);
    }
  } 
  
  /***************************************************************************
   **
   ** Set Layout Mode (PER_NODE or PER_NETWORK)
   */

  public void setLayoutMode(LayoutMode mode) {
    layoutMode_ = mode;
  }

  /***************************************************************************
   **
   ** Get Layout Mode (PER_NODE or PER_NETWORK)
   */

  public LayoutMode getLayoutMode() {
    return layoutMode_;
  }
 
  /***************************************************************************
  ** 
  ** Get node matches
  */

  public Set<NetNode> nodeMatches(boolean fullMatch, String searchString) {
    HashSet<NetNode> retval = new HashSet<NetNode>();
    Iterator<NetNode> nkit = nodeDefs_.keySet().iterator();
    while (nkit.hasNext()) {
      NetNode nodeID = nkit.next();
      String nodeName = nodeDefs_.get(nodeID).getNodeName();
      if (matches(searchString, nodeName, fullMatch)) {
        retval.add(nodeID);
      }
    }    
    return (retval);
  }
  
  /***************************************************************************
  ** 
  ** Get first neighbors of node, along with info blocks
  */

  public void getFirstNeighbors(NetNode nodeID, Set<NetNode> nodeSet, List<NodeInfo> nodes, List<LinkInfo> links) {
    Iterator<LinkInfo> ldit = fullLinkDefs_.values().iterator();
    while (ldit.hasNext()) {
      LinkInfo linf = ldit.next();
      if (linf.getSource().equals(nodeID)) {
        nodeSet.add(linf.getTarget());
        links.add(linf);
      } else if (linf.getTarget().equals(nodeID)) {
        nodeSet.add(linf.getSource());
        links.add(linf);
      }
    }
    Iterator<NetNode> nsit = nodeSet.iterator();
    while (nsit.hasNext()) {
      NetNode nextID = nsit.next();
      nodes.add(nodeDefs_.get(nextID));
    }
    return;
  }
  
  /***************************************************************************
  ** 
  ** Get first neighbors of node
  */

  public Set<NetNode> getFirstNeighbors(NetNode nodeID) {
    HashSet<NetNode> nodeSet = new HashSet<NetNode>();
    Iterator<LinkInfo> ldit = fullLinkDefs_.values().iterator();
    while (ldit.hasNext()) {
      LinkInfo linf = ldit.next();
      if (linf.getSource().equals(nodeID)) {
        nodeSet.add(linf.getTarget());
      } else if (linf.getTarget().equals(nodeID)) {
        nodeSet.add(linf.getSource());
      }
    }
    return (nodeSet);
  }
   
  /***************************************************************************
  ** 
  ** Add first neighbors of node set
  */

  public void addFirstNeighbors(Set<NetNode> nodeSet, Set<Integer> columnSet, Set<FabricLink> linkSet, boolean forShadow) {
    HashSet<NetNode> newNodes = new HashSet<NetNode>();
    Iterator<LinkInfo> ldit = fullLinkDefs_.values().iterator();
    while (ldit.hasNext()) {
      LinkInfo linf = ldit.next();
      if (!forShadow && linf.isShadow()) {
        continue;
      }
      if (nodeSet.contains(linf.getSource())) {
        newNodes.add(linf.getTarget());
        columnSet.add(Integer.valueOf(linf.getUseColumn(forShadow)));
        linkSet.add(linf.getLink());
      }
      if (nodeSet.contains(linf.getTarget())) {
        newNodes.add(linf.getSource());
        columnSet.add(Integer.valueOf(linf.getUseColumn(forShadow)));
        linkSet.add(linf.getLink());
      }
    }
    nodeSet.addAll(newNodes);
    return;
  }

  /***************************************************************************
  **
  ** Do the match
  ** 
  */
 
  private boolean matches(String searchString, String nodeName, boolean isFull) {
    String canonicalNodeName = DataUtil.normKey(nodeName);
    if (isFull) {
      return (canonicalNodeName.equals(searchString));
    } else {
      return (canonicalNodeName.indexOf(searchString) != -1);
    }
  }

  /***************************************************************************
  **
  ** Count of links per targ:
  */

  private Map<NetNode, Integer> linkCountPerTarg(Set<NetNode> targets, List<LinkInfo> linkList) {
    HashMap<NetNode, Integer> retval = new HashMap<NetNode, Integer>();
    Iterator<LinkInfo> lcit = linkList.iterator();
    while (lcit.hasNext()) {
      LinkInfo linf = lcit.next();
      NetNode src = linf.getSource();
      NetNode trg = linf.getTarget();      
      if (targets.contains(src)) {
        Integer count = retval.get(src);
        if (count == null) {
          retval.put(src, Integer.valueOf(1));
        } else {
          retval.put(src, Integer.valueOf(count.intValue() + 1));
        }
      }
      if (!src.equals(trg) && targets.contains(trg)) {
        Integer count = retval.get(trg);
        if (count == null) {
          retval.put(trg, Integer.valueOf(1));
        } else {
          retval.put(trg, Integer.valueOf(count.intValue() + 1));
        }
      }
    }
    return (retval);
  }
   
  /***************************************************************************
  **
  ** When creating submodel and shadow links are being displayed, we can choose
  ** to only show one copy of a link: either the main link, or the shadow link, unless
  ** both endpoint drains are on display, then both will be shown.
  */

  private List<LinkInfo> pruneToMinSubModel(BioFabricNetwork bfn, List<NodeInfo> targetList, List<LinkInfo> linkList) {
          
    HashSet<NetNode> targSet = new HashSet<NetNode>();
    Iterator<NodeInfo> tlit = targetList.iterator();
    while (tlit.hasNext()) {
      NodeInfo ninf = tlit.next();
      targSet.add(new FabricNode(ninf.getNodeID(), ninf.getNodeName()));
    }   

    Map<NetNode, Integer> subCounts = linkCountPerTarg(targSet, linkList);
    Map<NetNode, Integer> fullCounts = linkCountPerTarg(bfn.getNodeSetIDs(), bfn.getLinkDefList(true));
    
    HashSet<Integer> skipThem = new HashSet<Integer>();
    HashSet<Integer> ditchThem = new HashSet<Integer>();
    Iterator<LinkInfo> lcit = linkList.iterator();
    while (lcit.hasNext()) {
      LinkInfo linf = lcit.next();
      NetNode topNode = bfn.getNodeIDForRow(Integer.valueOf(linf.topRow()));
      NetNode botNode = bfn.getNodeIDForRow(Integer.valueOf(linf.bottomRow()));
      boolean topStays = subCounts.get(topNode).equals(fullCounts.get(topNode));
      boolean botStays = subCounts.get(botNode).equals(fullCounts.get(botNode));
      if ((topStays && botStays) || (!topStays && !botStays)) {  // Nobody gets ditched!
        continue;
      }
      FabricLink link1 = linf.getLink();
      int col1 = linf.getUseColumn(true);
      skipThem.add(Integer.valueOf(col1));
      Iterator<LinkInfo> lc2it = linkList.iterator();
      while (lc2it.hasNext()) {
        LinkInfo linf2 = lc2it.next();
        int col2 = linf2.getUseColumn(true);
        if (skipThem.contains(Integer.valueOf(col2))) {
          continue;
        }
        if (linf2.getLink().shadowPair(link1)) { // got a shadow pair! 
          int maxLink = Math.max(col1, col2);
          int minLink = Math.min(col1, col2);
          ditchThem.add(Integer.valueOf((topStays) ? maxLink : minLink));
          break;
        }
      }
    } 
    
    ArrayList<LinkInfo> retval = new ArrayList<LinkInfo>();
    Iterator<LinkInfo> lcit3 = linkList.iterator();
    while (lcit3.hasNext()) {
      LinkInfo linf = lcit3.next();
      int col = linf.getUseColumn(true);
      if (!ditchThem.contains(Integer.valueOf(col))) {
        retval.add(linf);
      }
    }
    return (retval);
  }

  /***************************************************************************
  **
  ** Fill out model with submodel data
  */

  private void fillSubModel(BioFabricNetwork bfn, List<NodeInfo> targetList, List<LinkInfo> linkList) {
    
    boolean doPrune = FabricDisplayOptionsManager.getMgr().getDisplayOptions().getMinShadowSubmodelLinks();
    if (doPrune) {
      linkList = pruneToMinSubModel(bfn, targetList, linkList);
    }
     
    //
    // Figure out the new node bounds. This figures out the current rightmost column of all nodes
    // needed to show the links we want to include in the subnet:
    //
    
    HashMap<NetNode, Integer> lastColForNode = new HashMap<NetNode, Integer>();
    HashMap<NetNode, Integer> lastShadColForNode = new HashMap<NetNode, Integer>();
    
    for (LinkInfo linf : linkList) {
      if (!linf.isShadow()) {
        Integer lastCol = lastColForNode.get(linf.getTarget());
        if ((lastCol == null) || (lastCol.intValue() < linf.getUseColumn(false))) {
          lastColForNode.put(linf.getTarget(), Integer.valueOf(linf.getUseColumn(false)));
        }
        lastCol = lastColForNode.get(linf.getSource());
        if ((lastCol == null) || (lastCol.intValue() < linf.getUseColumn(false))) {
          lastColForNode.put(linf.getSource(), Integer.valueOf(linf.getUseColumn(false)));
        }
      }
      Integer lastColShad = lastShadColForNode.get(linf.getTarget());
      if ((lastColShad == null) || (lastColShad.intValue() < linf.getUseColumn(true))) {
        lastShadColForNode.put(linf.getTarget(), Integer.valueOf(linf.getUseColumn(true)));
      }
      lastColShad = lastShadColForNode.get(linf.getSource());
      if ((lastColShad == null) || (lastColShad.intValue() < linf.getUseColumn(true))) {
        lastShadColForNode.put(linf.getSource(), Integer.valueOf(linf.getUseColumn(true)));
      }
    }

    //
    // Need to compress the rows and columns, throwing away all empty slots
    // First record the "full scale" entries:
    //

    TreeSet<Integer> needRows = new TreeSet<Integer>();
    TreeSet<Integer> needColumns = new TreeSet<Integer>();   
    TreeSet<Integer> needColumnsShad = new TreeSet<Integer>();   

    for (NodeInfo targetInf : targetList) {
      needRows.add(Integer.valueOf(targetInf.nodeRow));
      needColumns.add(Integer.valueOf(targetInf.getColRange(false).min));
      needColumnsShad.add(Integer.valueOf(targetInf.getColRange(true).min));
    }

    Iterator<LinkInfo> cit = linkList.iterator();
    while (cit.hasNext()) {
      LinkInfo linf = cit.next();
      needRows.add(Integer.valueOf(linf.getStartRow()));
      needRows.add(Integer.valueOf(linf.getEndRow()));
      if (!linf.isShadow()) {
        needColumns.add(Integer.valueOf(linf.getUseColumn(false)));
      }
      needColumnsShad.add(Integer.valueOf(linf.getUseColumn(true)));
    }

    //
    // Create full-scale to mini-scale mappings:
    //

    TreeMap<Integer, Integer> rowMap = new TreeMap<Integer, Integer>();
    TreeMap<Integer, Integer> columnMap = new TreeMap<Integer, Integer>();
    TreeMap<Integer, Integer> shadColumnMap = new TreeMap<Integer, Integer>();

    int rowCount = 0;
    for (Integer fullRow : needRows) {
      rowMap.put(fullRow, Integer.valueOf(rowCount++));
    }

    int colCount = 0;
    for (Integer fullCol : needColumns) {
      columnMap.put(fullCol, Integer.valueOf(colCount++));
    }
    
    int shadColCount = 0;
    for (Integer fullCol : needColumnsShad) {
      shadColumnMap.put(fullCol, Integer.valueOf(shadColCount++));
    }
  
    //
    // Create modified copies of the node and link info with
    // compressed rows and columns:
    //

    ArrayList<NodeInfo> modTargetList = new ArrayList<NodeInfo>();
    ArrayList<LinkInfo> modLinkList = new ArrayList<LinkInfo>();

    int maxShadLinkCol = Integer.MIN_VALUE;
    int maxLinkCol = Integer.MIN_VALUE;
    cit = linkList.iterator();
    while (cit.hasNext()) {
      BioFabricNetwork.LinkInfo linf = cit.next();
      Integer startRowObj = rowMap.get(Integer.valueOf(linf.getStartRow()));
      Integer endRowObj = rowMap.get(Integer.valueOf(linf.getEndRow()));
      Integer miniColObj = (linf.isShadow()) ? Integer.valueOf(Integer.MIN_VALUE) : columnMap.get(Integer.valueOf(linf.getUseColumn(false)));
      Integer miniColShadObj = shadColumnMap.get(Integer.valueOf(linf.getUseColumn(true)));
      int miniColVal = miniColObj.intValue();
      if (miniColVal > maxLinkCol) {
        maxLinkCol = miniColVal;
      }
      int miniColShadVal = miniColShadObj.intValue();
      if (miniColShadVal > maxShadLinkCol) {
        maxShadLinkCol = miniColShadVal;
      }
      
      LinkInfo miniLinf = new LinkInfo(linf.getLink(), startRowObj.intValue(), endRowObj.intValue(), 
                                       miniColObj.intValue(), miniColShadObj.intValue(), linf.getColorKey());
      modLinkList.add(miniLinf);
    }
    if (maxLinkCol == Integer.MIN_VALUE) {
      maxLinkCol = 1;
    } else {
      maxLinkCol++;
    }
    if (maxShadLinkCol == Integer.MIN_VALUE) {
      maxShadLinkCol = 1;
    } else {
      maxShadLinkCol++;
    }
 
    int minTrgCol = Integer.MAX_VALUE;
    int minShadTrgCol = Integer.MAX_VALUE;
    for (NodeInfo infoFull : targetList) {
      Integer miniRowObj = rowMap.get(Integer.valueOf(infoFull.nodeRow));
      NodeInfo infoMini = new NodeInfo(infoFull.getNodeID(), infoFull.getNodeName(), miniRowObj.intValue(), infoFull.colorKey);

      Integer minCol = columnMap.get(Integer.valueOf(infoFull.getColRange(false).min));
      infoMini.updateMinMaxCol(minCol.intValue(), false);
      int miniColVal = minCol.intValue();
      if (miniColVal < minTrgCol) {
        minTrgCol = miniColVal;
      }
      
      Integer minShadCol = shadColumnMap.get(Integer.valueOf(infoFull.getColRange(true).min));
      infoMini.updateMinMaxCol(minShadCol.intValue(), true);
      int miniShadColVal = minShadCol.intValue();
      if (miniShadColVal < minShadTrgCol) {
        minShadTrgCol = miniShadColVal;
      }
   
      int maxCol;        
      Integer lastCol = lastColForNode.get(infoFull.getNodeID());
      if (lastCol == null) {
        maxCol = maxLinkCol;
      } else {
        Integer maxColObj = columnMap.get(lastCol);     
        maxCol = maxColObj.intValue();
      }
      infoMini.updateMinMaxCol(maxCol, false);
      
      int maxShadCol;        
      Integer lastShadCol = lastShadColForNode.get(infoFull.getNodeID());
      if (lastShadCol == null) {
        maxShadCol = maxShadLinkCol;
      } else {
        Integer maxShadColObj = shadColumnMap.get(lastShadCol);     
        maxShadCol = maxShadColObj.intValue();
      }
      infoMini.updateMinMaxCol(maxShadCol, true);
  
      modTargetList.add(infoMini);
    }
    if (minTrgCol == Integer.MAX_VALUE) {
      minTrgCol = 0;
    }
    
    rowToTargID_ = new HashMap<Integer, NetNode>();
    nodeDefs_ = new HashMap<NetNode, NodeInfo>();
    rowCount_ = modTargetList.size();
    for (int i = 0; i < rowCount_; i++) {
      NodeInfo infoMini = modTargetList.get(i);
      NetNode nwn = new FabricNode(infoMini.getNodeID(), infoMini.getNodeName());
      rowToTargID_.put(Integer.valueOf(infoMini.nodeRow), nwn);
      nodeDefs_.put(nwn, infoMini);
    }
    
    normalCols_ = new ColumnAssign();
    shadowCols_ = new ColumnAssign();
    fullLinkDefs_ = new TreeMap<Integer, LinkInfo>();
    nonShadowedLinkMap_ = new TreeMap<Integer, Integer>();
    
    int numMll = modLinkList.size();
    for (int i = 0; i < numMll; i++) {
      BioFabricNetwork.LinkInfo infoMini = modLinkList.get(i);
      
      Integer useShadCol = Integer.valueOf(infoMini.getUseColumn(true));
      fullLinkDefs_.put(useShadCol, infoMini);      
      shadowCols_.columnToSource.put(useShadCol, infoMini.getSource());   
      shadowCols_.columnToTarget.put(useShadCol, infoMini.getTarget());
           
      if (!infoMini.isShadow()) {
        Integer useCol = Integer.valueOf(infoMini.getUseColumn(false));
        nonShadowedLinkMap_.put(useCol, useShadCol);      
        normalCols_.columnToSource.put(useCol, infoMini.getSource());   
        normalCols_.columnToTarget.put(useCol, infoMini.getTarget());
      }
    }
    
    normalCols_.columnCount = maxLinkCol - minTrgCol;
    shadowCols_.columnCount = maxShadLinkCol - minShadTrgCol;
    
    //
    // Need to build drain zones.  For each node, start at max col and back up.
    // Drain starts at max IF the source is the top node.  Continues until
    // we stop having nodes in the next slot that are the top nodes.
    //
    // We only keep a drain zone when it is at the top of the display.  We can
    // check this by seeing if the last link for the node has the node at the top
    // (keep drain) or the bottom (drop drain) of the link.
    //
    // That logic only applies with non-shadow presentation.  For shadowed, everybody 
    // with a link has a drain zone
    //
    
     UiUtil.fixMePrintout("Are multiple drain zones handled for submodels? NO");
    
    Iterator<NetNode> ndkit = nodeDefs_.keySet().iterator();
    while (ndkit.hasNext()) {
      NetNode node = ndkit.next();
      NodeInfo srcNI = nodeDefs_.get(node);
      
      //
      // Non-shadow calcs:
      //
      MinMax srcDrain = null;
      MinMax range = srcNI.getColRange(false);
      int startCol = range.max;
      int endCol = range.min;
      for (int i = startCol; i >= endCol; i--) {
        LinkInfo linf = getLinkDefinition(Integer.valueOf(i), false);
        // done when no longer contiguous:
        if ((linf == null) || (!linf.getSource().equals(node) && !linf.getTarget().equals(node))) {
          break;
        }
        // Second case allows feedback:
        if ((linf.bottomRow() == srcNI.nodeRow) && (linf.topRow() != srcNI.nodeRow)) {
          break;
        }
        if (linf.topRow() == srcNI.nodeRow) {
          if (srcDrain == null) {
            srcDrain = new MinMax();
            srcDrain.init();
          }
          srcDrain.update(i); 
        }        
      }
      if (srcDrain != null) {
        srcNI.addDrainZone(new DrainZone(srcDrain, false));
      }
      
      //
      // Shadow calcs:
      //
      
      MinMax shadowSrcDrain = null;
      MinMax shadowRange = srcNI.getColRange(true);
      int startColShad = shadowRange.min;
      int endColShad = shadowRange.max;
      for (int i = startColShad; i <= endColShad; i++) {
        LinkInfo linf = getLinkDefinition(Integer.valueOf(i), true);
        if (linf == null) {
          continue;
        }
        boolean isShadow = ((linf != null) && linf.isShadow());
   //     if (!isShadow && (shadowSrcDrain == null)) {
    //      continue;
    //    }
        // done when no longer contiguous:
        if (shadowSrcDrain != null) {
          if ((linf == null) || (!linf.getSource().equals(node) && !linf.getTarget().equals(node))) {
            break;
          }
        }
        if (((linf.topRow() == srcNI.nodeRow) && !isShadow) || 
             (linf.bottomRow() == srcNI.nodeRow) && isShadow) {
          if (shadowSrcDrain == null) {
            shadowSrcDrain = new MinMax();
            shadowSrcDrain.init();
          }
          shadowSrcDrain.update(i); 
        }        
      }
      if (shadowSrcDrain != null) {
        srcNI.addDrainZone(new DrainZone(shadowSrcDrain, true));
      }
    }
   
    UiUtil.fixMePrintout("Nodes are stretching all the way to right border in LesMizCluster subset.");
    
    //
    // Handle the subsetting of the node and link annotations
    //
    
    this.nodeAnnot_ = annotationsForSubNet(rowMap, bfn.nodeAnnot_);
    if (bfn.linkAnnots_ != null) {
    	this.linkAnnots_ = new HashMap<Boolean, AnnotationSet>();
    	if (bfn.linkAnnots_.get(Boolean.FALSE) != null) {
        this.linkAnnots_.put(Boolean.FALSE, annotationsForSubNet(columnMap, bfn.linkAnnots_.get(Boolean.FALSE)));
    	}
    	if (bfn.linkAnnots_.get(Boolean.TRUE) != null) {
        this.linkAnnots_.put(Boolean.TRUE, annotationsForSubNet(shadColumnMap, bfn.linkAnnots_.get(Boolean.TRUE)));
    	}
    }
    
    return;
  }

  /***************************************************************************
  **
  ** Fill out reduced node or link annotations
  */

  private AnnotationSet annotationsForSubNet(TreeMap<Integer, Integer> reduceMap, AnnotationSet origAnnots) {
  	AnnotationSet retval = new AnnotationSetImpl();
  	if ((origAnnots == null) || (origAnnots.size() == 0)) {
  		return (retval);
  	}
  	
  	//
  	// We need to go through each annotation and see if any of the nodes/links in the subnet are in it. If so, it
  	// needs to be in the subset, with min/max values taken from the rows/cols present in the subnet.
  	//
  	
  	TreeMap<Annot, MinMax> useAnnots = new TreeMap<Annot, MinMax>();
  	for (Integer origRow : reduceMap.keySet()) {
  		int orVal = origRow.intValue();
  		for (Annot origAnnot : origAnnots) { 
  		  MinMax anRange = origAnnot.getRange();
  		  if ((orVal >= anRange.min) && (orVal <= anRange.max)) {
  		  	MinMax useRange = useAnnots.get(origAnnot);
  		  	Integer newRowOrCol = reduceMap.get(origRow);
  		  	int nrVal = newRowOrCol.intValue();
  		  	if (useRange == null) {
  		  		useRange = new MinMax(nrVal, nrVal);
  		  		useAnnots.put(origAnnot, useRange);
  		  	} else {
  		  		useRange.update(nrVal);
  		  	}
  		  	break;
  		  }
  	  }
    }
  	
  	for (Annot oldAnnot : useAnnots.keySet()) {
  		MinMax useRange = useAnnots.get(oldAnnot);
  		AnnotColorSource.AnnotColor color = oldAnnot.getColor();
  		String colorName = (color == null) ? null : color.getName();
  		AnnotationSetImpl.AnnotImpl ai = new AnnotationSetImpl.AnnotImpl(oldAnnot.getName(), useRange.min, useRange.max, 
  																																		 oldAnnot.getLayer(), colorName);
  		retval.addAnnot(ai);
  	}  	
  	return (retval);
  }

  /***************************************************************************
  ** 
  ** Fill out node info from order
  */

  private void fillNodesFromOrder(List<NetNode> targetIDs, 
  		                            FabricColorGenerator colGen, Map<NetNode, String> clustAssign,
  		                            BTProgressMonitor monitor) throws AsynchExitRequestException {
    //
    // Now have the ordered list of targets we are going to display.
    // Build target->row maps and the inverse:
    //
    
    int numColors = colGen.getNumColors();

    LoopReporter lr = new LoopReporter(targetIDs.size(), 20, monitor, 0.0, 1.0, "progress.nodeInfo");
  
    int currRow = 0;
    Iterator<NetNode> trit = targetIDs.iterator();
    while (trit.hasNext()) {
      NetNode targetID = trit.next();
      lr.report();
      Integer rowObj = Integer.valueOf(currRow);
      rowToTargID_.put(rowObj, targetID);
      String colorKey = colGen.getGeneColor(currRow % numColors);
      if (targetID == null) {
      	UiUtil.fixMePrintout("SAW THIS FOR ROW OBJ 5245");
        UiUtil.fixMePrintout("targetID is Null " + rowObj);
      }

      NodeInfo nextNI = new NodeInfo(targetID.getNID().getNID(), targetID.getName(), currRow++, colorKey);
      if (clustAssign != null) {
      	nextNI.setCluster(clustAssign.get(targetID));
      }
      nodeDefs_.put(targetID, nextNI);
    }
    rowCount_ = targetIDs.size();
    return;
  }
  
  /***************************************************************************
  **
  ** Add a node def
  */
  
  void addNodeInfoForIO(NodeInfo nif) {
  	NetNode nwn = new FabricNode(nif.getNodeID(), nif.getNodeName());
    nodeDefs_.put(nwn, nif);
    rowCount_ = nodeDefs_.size();   
    rowToTargID_.put(Integer.valueOf(nif.nodeRow), nwn);
    return;
  }
  
  /***************************************************************************
  ** 
  ** Determine the start & end of each target row needed to handle the incoming
  ** and outgoing links:
  */

  private void trimTargetRows(BTProgressMonitor monitor) throws AsynchExitRequestException {
  	  	
  	LoopReporter lr = new LoopReporter(fullLinkDefs_.size(), 20, monitor, 0.0, 0.5, "progress.trimTargetRows1");
  	Iterator<Integer> fldit = fullLinkDefs_.keySet().iterator();
    while (fldit.hasNext()) {
      Integer colNum = fldit.next();
      lr.report();
      LinkInfo li = fullLinkDefs_.get(colNum);
      shadowCols_.columnToSource.put(colNum, li.getSource());
      shadowCols_.columnToTarget.put(colNum, li.getTarget());
      NodeInfo srcNI = nodeDefs_.get(li.getSource());    
      NodeInfo trgNI = nodeDefs_.get(li.getTarget()); 
      srcNI.updateMinMaxCol(colNum.intValue(), true);
      trgNI.updateMinMaxCol(colNum.intValue(), true);
    }
    lr.finish();
    
    LoopReporter lr2 = new LoopReporter(nonShadowedLinkMap_.size(), 20, monitor, 0.5, 1.0, "progress.trimTargetRows2");
    Iterator<Integer> nslit = nonShadowedLinkMap_.keySet().iterator();
    while (nslit.hasNext()) {
      Integer colNum = nslit.next();
      lr2.report();
      Integer mappedCol = nonShadowedLinkMap_.get(colNum);
      LinkInfo li = fullLinkDefs_.get(mappedCol);
      normalCols_.columnToSource.put(colNum, li.getSource());
      normalCols_.columnToTarget.put(colNum, li.getTarget());
      NodeInfo srcNI = nodeDefs_.get(li.getSource());    
      NodeInfo trgNI = nodeDefs_.get(li.getTarget());    
      srcNI.updateMinMaxCol(colNum.intValue(), false);
      trgNI.updateMinMaxCol(colNum.intValue(), false);
    }
    lr.finish();
    return;
  }
  
  /***************************************************************************
  ** 
  ** For the lone nodes, they are assigned into the last column:
  */

  private void loneNodesToLastColumn(Set<NetNode> loneNodeIDs,
  		                               BTProgressMonitor monitor) throws AsynchExitRequestException {
  	
  	LoopReporter lr = new LoopReporter(loneNodeIDs.size(), 20, monitor, 0.0, 1.0, "progress.loneNodes");
  	
    Iterator<NetNode> lnit = loneNodeIDs.iterator();
    while (lnit.hasNext()) {
      NetNode loneID = lnit.next();
      lr.report();
      NodeInfo loneNI = nodeDefs_.get(loneID);     
      loneNI.updateMinMaxCol(normalCols_.columnCount - 1, false);
      loneNI.updateMinMaxCol(shadowCols_.columnCount - 1, true);
    } 
    lr.finish();
    return;
  }
  
  /***************************************************************************
  ** 
  ** Previous icky hack was broken, and depended on default layout algorithm being used to work.
  ** This is more expensive, but works:
  */

  public Set<NetNode> getLoneNodes(BTProgressMonitor monitor) throws AsynchExitRequestException { 
    HashSet<NetNode> retval = new HashSet<NetNode>(nodeDefs_.keySet());
    LoopReporter lr = new LoopReporter(fullLinkDefs_.size(), 20, monitor, 0.0, 1.0, "progress.findingLoneNodes");   
    for (LinkInfo lif : fullLinkDefs_.values()) {
      lr.report();
      NetLink link = lif.getLink();
      retval.remove(link.getSrcNode());
      retval.remove(link.getTrgNode());
    }
    lr.finish();
    return (retval);
  }
 
  /***************************************************************************
  **
  ** Add a link def
  */
  
  private Integer[] addLinkDef(NetLink nextLink, int numColors, int noShadowCol, int shadowCol, FabricColorGenerator colGen) {
    Integer[] retval = new Integer[2]; 
    String key = colGen.getGeneColor(shadowCol % numColors);
    int srcRow = nodeDefs_.get(nextLink.getSrcNode()).nodeRow;
    int trgRow = nodeDefs_.get(nextLink.getTrgNode()).nodeRow;
    LinkInfo linf = new LinkInfo((FabricLink)nextLink, srcRow, trgRow, noShadowCol, shadowCol, key);
    Integer shadowKey = Integer.valueOf(shadowCol);
    fullLinkDefs_.put(shadowKey, linf);
    retval[0] = Integer.valueOf(shadowCol + 1); 
    if (!linf.isShadow()) {
      nonShadowedLinkMap_.put(Integer.valueOf(noShadowCol), shadowKey);
      retval[1] = Integer.valueOf(noShadowCol + 1); 
    }
    return (retval);
  }
  
  /***************************************************************************
  **
  ** Add a link def
  */
  
  void addLinkInfoForIO(LinkInfo linf) {
    int useColVal = linf.getUseColumn(true);
    Integer useCol = Integer.valueOf(useColVal);
    fullLinkDefs_.put(useCol, linf);    
    if (useColVal > shadowCols_.columnCount) {
      shadowCols_.columnCount = useColVal;
    }
    shadowCols_.columnToSource.put(useCol, linf.getSource());   
    shadowCols_.columnToTarget.put(useCol, linf.getTarget());
    
    if (!linf.isShadow()) {
      int useNColVal = linf.getUseColumn(false);
      Integer useNCol = Integer.valueOf(useNColVal);
      nonShadowedLinkMap_.put(useNCol, useCol);    
      if (useNColVal > normalCols_.columnCount) {
        normalCols_.columnCount = useNColVal;
      }
      normalCols_.columnToSource.put(useNCol, linf.getSource());   
      normalCols_.columnToTarget.put(useNCol, linf.getTarget());
    }
    return;
  }
  
  /***************************************************************************
  **
  ** Add a link group tag
  */
  
  void addLinkGroupForIO(String tag) {
    this.linkGrouping_.add(tag);
    return;
  }
 
  /***************************************************************************
  **
  ** Set color generator (I/0)
  */
  
  void setColorGenerator(FabricColorGenerator fcg) {
    colGen_ = fcg;
    return;
  } 
  
  /***************************************************************************
  **
  ** Answer if we have node cluster assignments
  */
  
  public boolean nodeClustersAssigned() {
  	if (nodeDefs_.isEmpty()) {
  		return (false);
  	}
    NodeInfo ni = nodeDefs_.values().iterator().next();
    return (ni.getCluster() != null);
  } 
  
  /***************************************************************************
  **
  ** Return node cluster assignment
  */
  
  public Map<NetNode, String> nodeClusterAssigment() {
  	HashMap<NetNode, String> retval = new HashMap<NetNode, String>();
  	if (nodeDefs_.isEmpty()) {
  		return (retval);
  	}
    for (NodeInfo ni : nodeDefs_.values()) {
    	String cluster = ni.getCluster();
    	if (cluster == null) {
    		throw new IllegalStateException();
    	}
    	retval.put(new FabricNode(ni.getNodeID(), ni.getNodeName()), cluster);
    }
    return (retval);    
  } 
  
  /***************************************************************************
  **
  ** Set node cluster assignment
  */
  
  public void setNodeClusterAssigment(Map<NID, String> assign) {
  	if (nodeDefs_.isEmpty()) {
  		throw new IllegalStateException();
  	}
    for (NodeInfo ni : nodeDefs_.values()) {
    	String cluster = assign.get(ni.getNodeID());
    	if (cluster == null) {
    		throw new IllegalArgumentException();
    	}
    	ni.setCluster(cluster);
    }
    return;    
  } 
  
  /***************************************************************************
  **
  ** Constructor for I/O only
  */

  BioFabricNetwork() { 
    normalCols_ = new ColumnAssign();
    shadowCols_ = new ColumnAssign();
    rowToTargID_ = new HashMap<Integer, NetNode>(); 
    fullLinkDefs_ = new TreeMap<Integer, LinkInfo>();
    nonShadowedLinkMap_ = new TreeMap<Integer, Integer>();
    nodeDefs_ = new HashMap<NetNode, NodeInfo>();
    linkGrouping_ = new ArrayList<String>();
    showLinkGroupAnnotations_ = false;
    colGen_ = null;
  }
  
  /***************************************************************************
  **
  ** Info for a link
  */  
  
  public static class LinkInfo {
    private FabricLink myLink_;
    private int startRow_;
    private int endRow_;
    private int noShadowColumn_;
    private int shadowColumn_;
    private String colorKey_;
    
    public LinkInfo(FabricLink flink, int startRow, int endRow, int noShadowColumn, int shadowColumn, String colorKey) {
      myLink_ = flink.clone();
      startRow_ = startRow;
      endRow_ = endRow;
      noShadowColumn_ = noShadowColumn;
      shadowColumn_ = shadowColumn;
      colorKey_ = colorKey;
    }
    
    public int getStartRow() {
      return (startRow_);
    }  
    
    public int getEndRow() {
      return (endRow_);
    }  
    
    public int getUseColumn(boolean shadowEnabled) {
      if (!shadowEnabled && myLink_.isShadow()) {
        // FIX ME: Seen this case with submodel creation?
        throw new IllegalStateException();
      }
      return ((shadowEnabled) ? shadowColumn_ : noShadowColumn_);
    }  
    
    public String getColorKey() {
      return (colorKey_);
    }  
       
    public FabricLink getLink() {
      return (myLink_);
    }    
    
    public NetNode getSource() {
      return (myLink_.getSrcNode());
    }
    
    public NetNode getTarget() {
      return (myLink_.getTrgNode());
    }
    
    public AugRelation getAugRelation() {
      return (myLink_.getAugRelation());
    }
    
    public boolean isShadow() {
      return (myLink_.isShadow());
    }
    
    public boolean isDirected() {
      return (myLink_.isDirected());
    }
        
    public int bottomRow() {
      return (Math.max(startRow_, endRow_));
    } 
    
    public int topRow() {
      return (Math.min(startRow_, endRow_));
    } 
    
    public boolean inLinkRowRange(int row) {
      return ((row >= topRow()) && (row <= bottomRow()));
    } 
  }  
  
  /***************************************************************************
  **
  ** Node info
  */  
  
  public static class NodeInfo {
  	private NID nodeID_;
    private String nodeName_;
    public int nodeRow;
    public String colorKey;
    private String cluster_;
        
    private MinMax colRangeSha_;
    private MinMax colRangePln_;
    
    private List<DrainZone> shadowDrainZones_;
    private List<DrainZone> plainDrainZones_;
    
    NodeInfo(NID nodeID, String nodeName, int nodeRow, String colorKey) {
    	nodeID_ = nodeID;
      nodeName_ = nodeName;
      this.nodeRow = nodeRow;
      this.colorKey = colorKey;
      colRangeSha_ = new MinMax();
      colRangeSha_.init();
      colRangePln_ = new MinMax();
      colRangePln_.init();
      shadowDrainZones_ = new ArrayList<DrainZone>();
      cluster_ = null;
      plainDrainZones_ = new ArrayList<DrainZone>();
    }
      
    public String getNodeName() { 
      return (nodeName_);
    }
    
    public NID getNodeID() { 
      return (nodeID_);
    } 
    
    public NetNode getNodeIDWithName() { 
      return (new FabricNode(nodeID_, nodeName_));
    } 
 
    public List<DrainZone> getDrainZones(boolean forShadow) {
      return (forShadow) ? new ArrayList<DrainZone>(shadowDrainZones_) : new ArrayList<DrainZone>(plainDrainZones_);
    }
    
    public void addDrainZone(DrainZone dz) {
      if (dz.isShadow()) {
        shadowDrainZones_.add(dz);
      } else {
        plainDrainZones_.add(dz);
      }
      return;
    }
  
    public void setDrainZones(List<DrainZone> zones, boolean forShadow) {
      if (forShadow) {
        shadowDrainZones_ = new ArrayList<DrainZone>(zones);
      } else {
        plainDrainZones_ = new ArrayList<DrainZone>(zones);
      }
    }
    
    public MinMax getColRange(boolean forShadow) { 
      return (forShadow) ? colRangeSha_ : colRangePln_;
    }
      
    void updateMinMaxCol(int i, boolean forShadow) {
      MinMax useMM = (forShadow) ? colRangeSha_ : colRangePln_;
      useMM.update(i);
      return;
    }
    
    public void setCluster(String cluster) {
      cluster_ = cluster;
      return;
    }
    
    public String getCluster() {
      return (cluster_);
    }
    
    /***************************************************************************
    **
     ** Dump the node using XML
    */
  
    public void writeXML(PrintWriter out, Indenter ind, int row) {
      ind.indent();
      out.print("<node name=\"");
      out.print(CharacterEntityMapper.mapEntities(nodeName_, false));
      out.print("\" nid=\"");
      out.print(nodeID_.getInternal());
      out.print("\" row=\"");
      out.print(row);
      MinMax nsCols = getColRange(false);
      out.print("\" minCol=\"");
      out.print(nsCols.min);
      out.print("\" maxCol=\"");
      out.print(nsCols.max);
      MinMax sCols = getColRange(true);
      out.print("\" minColSha=\"");
      out.print(sCols.min);
      out.print("\" maxColSha=\"");
      out.print(sCols.max);
      out.print("\" color=\"");
      out.print(colorKey);
      String clust = getCluster();
      if (clust != null) {
        out.print("\" cluster=\"");
        out.print(CharacterEntityMapper.mapEntities(clust, false));
      }

      out.println("\">");
      
      //
      // DRAIN ZONES XML
      //
      
      ind.up();
      ind.indent();
      if (this.plainDrainZones_.size() > 0) {
        out.println("<drainZones>");
        ind.up();
        for (DrainZone dz : this.plainDrainZones_) {
          dz.writeXML(out, ind);
        }
        ind.down();
        ind.indent();
        out.println("</drainZones>");
      } else {
        out.println("<drainZones/>");
      }
      
      ind.indent();
      if (this.shadowDrainZones_.size() > 0) {
        out.println("<drainZonesShadow>");
        ind.up();
        for (DrainZone dzSha : this.shadowDrainZones_) {
          dzSha.writeXML(out, ind);
        }
        ind.down();
        ind.indent();
        out.println("</drainZonesShadow>");
      } else {
        out.println("<drainZonesShadow/>");
      }
      
      ind.down();
      ind.indent();
      out.println("</node>");
    }
  }
  
  /***************************************************************************
   **
   ** Drain Zone
   */
  
  public static class DrainZone {
    
    private MinMax dzmm;
    private boolean isShadow;
    
    public DrainZone(MinMax dzmm, boolean isShadow){
      this.dzmm = dzmm.clone();
      this.isShadow = isShadow;
    }
    
    public void writeXML(PrintWriter out, Indenter ind) {
      if (isShadow) {
        ind.indent();
        out.print("<drainZoneShadow minCol=\"");
        out.print(dzmm.min);
        out.print("\" maxCol=\"");
        out.print(dzmm.max);
        out.println("\" />");
      } else {
        ind.indent();
        out.print("<drainZone minCol=\"");
        out.print(dzmm.min);
        out.print("\" maxCol=\"");
        out.print(dzmm.max);
        out.println("\" />");
      }
    }
    
    public boolean isShadow() {
      return isShadow;
    }
    
    public MinMax getMinMax() {
      return dzmm;
    }
  }
  
  /***************************************************************************
  **
  ** For passing around ranked nodes
  */  
  
  static class DoubleRanked  {
     double rank;
     String id;
     Link byLink;

    DoubleRanked(double rank, String id, Link byLink) {
      this.rank = rank;
      this.id = id;
      this.byLink = byLink;
    } 
  }
  
  /***************************************************************************
  **
  ** For storing column assignments
  */  
  
  public static class ColumnAssign  {
    public HashMap<Integer, NetNode> columnToSource;
    public HashMap<Integer, NetNode> columnToTarget;
    public int columnCount;

    ColumnAssign() {
      this.columnToSource = new HashMap<Integer, NetNode>();
      this.columnToTarget = new HashMap<Integer, NetNode>();
      this.columnCount = 0;
    } 
  }
 
  ////////////////////////////////////////////////////////////////////////////
  //
  // PUBLIC STATIC METHODS
  //
  ////////////////////////////////////////////////////////////////////////////
  
  /***************************************************************************
  **
  ** For XML I/O
  */  
      
  public static class NetworkDataWorker extends AbstractFactoryClient {
    
    public NetworkDataWorker(FabricFactory.FactoryWhiteboard whiteboard, PlugInManager pMan) {
      super(whiteboard);
      myKeys_.add("BioFabric");
      installWorker(new FabricColorGenerator.ColorSetWorker(whiteboard), new MyColorSetGlue());
      installWorker(new FabricDisplayOptions.FabricDisplayOptionsWorker(whiteboard), null);
      installWorker(new NodeInfoWorker(whiteboard), new MyNodeGlue());
      installWorker(new LinkInfoWorker(whiteboard), new MyLinkGlue());
      installWorker(new LinkGroupWorker(whiteboard), null);
      installWorker(new AnnotationSetImpl.AnnotsWorker(whiteboard, "nodeAnnotations"), new MyAnnotsGlue(true, false));
      installWorker(new AnnotationSetImpl.AnnotsWorker(whiteboard, "linkAnnotations"), new MyAnnotsGlue(false, false));
      installWorker(new AnnotationSetImpl.AnnotsWorker(whiteboard, "shadowLinkAnnotations"), new MyAnnotsGlue(false, true));     
      installWorker(new PlugInManager.PlugInWorker(whiteboard, pMan), null);
    }
    
    protected Object localProcessElement(String elemName, Attributes attrs) throws IOException {
      Object retval = null;
      if (elemName.equals("BioFabric")) {
        FabricFactory.FactoryWhiteboard board = (FabricFactory.FactoryWhiteboard)this.sharedWhiteboard_;
        board.bfn = new BioFabricNetwork();
        retval = board.bfn;
      }
      return (retval);     
    }
  }
  
  public static class MyColorSetGlue implements GlueStick {
    public Object glueKidToParent(Object kidObj, AbstractFactoryClient parentWorker, 
                                  Object optionalArgs) throws IOException {
      FabricFactory.FactoryWhiteboard board = (FabricFactory.FactoryWhiteboard)optionalArgs;
      board.bfn.setColorGenerator(board.fcg);
      return (null);
    }
  }  
  
  public static class MyAnnotsGlue implements GlueStick {
    
    private boolean forNodes_;
    private boolean forShadow_;
    
    public MyAnnotsGlue(boolean forNodes, boolean forShadow) {
      forNodes_ = forNodes;
      forShadow_ = forShadow;
    }
   
    public Object glueKidToParent(Object kidObj, AbstractFactoryClient parentWorker, 
                                  Object optionalArgs) throws IOException {
      FabricFactory.FactoryWhiteboard board = (FabricFactory.FactoryWhiteboard)optionalArgs;
      if (forNodes_) {
        board.bfn.setNodeAnnotations(board.currAnnots);
      } else {
        board.bfn.setLinkAnnotations(board.currAnnots, forShadow_);
      }
      return (null);
    }
  }  

  public static class MyNodeGlue implements GlueStick {
    public Object glueKidToParent(Object kidObj, AbstractFactoryClient parentWorker, 
                                  Object optionalArgs) throws IOException {
      FabricFactory.FactoryWhiteboard board = (FabricFactory.FactoryWhiteboard)optionalArgs;
      board.bfn.addNodeInfoForIO(board.nodeInfo);
      return (null);
    }
  }  
  
  public static class MyLinkGlue implements GlueStick {
    public Object glueKidToParent(Object kidObj, AbstractFactoryClient parentWorker, 
                                  Object optionalArgs) throws IOException {
      FabricFactory.FactoryWhiteboard board = (FabricFactory.FactoryWhiteboard)optionalArgs;
      board.bfn.addLinkInfoForIO(board.linkInfo);
      return (null);
    }
  } 
  
  public static class MyGroupGlue implements GlueStick {
    public Object glueKidToParent(Object kidObj, AbstractFactoryClient parentWorker, 
                                  Object optionalArgs) throws IOException {
      FabricFactory.FactoryWhiteboard board = (FabricFactory.FactoryWhiteboard)optionalArgs;
      board.bfn.addLinkGroupForIO(board.groupTag);
      return (null);
    }
  }
  
  public static class DrainZoneGlue implements GlueStick {
    
    public Object glueKidToParent(Object kidObj, AbstractFactoryClient parentWorker, Object optionalArgs) throws IOException {
      FabricFactory.FactoryWhiteboard board = (FabricFactory.FactoryWhiteboard) optionalArgs;
      board.nodeInfo.addDrainZone(board.drainZone);
      return null;
    }
  }
  
  /***************************************************************************
  **
  ** For XML I/O
  */
  
  public static class DrainZoneWorker extends AbstractFactoryClient {
    
    private boolean isShadow;
    
    public DrainZoneWorker(FabricFactory.FactoryWhiteboard board, boolean isShadow) {
      super(board);
      this.isShadow = isShadow;
      
      if (this.isShadow) {
        myKeys_.add("drainZoneShadow");
      } else {
        myKeys_.add("drainZone");
      }
    }
    
    protected Object localProcessElement(String elemName, Attributes attrs) throws IOException {
      Object retval = null;
      FabricFactory.FactoryWhiteboard board = (FabricFactory.FactoryWhiteboard) this.sharedWhiteboard_;
      board.drainZone = buildFromXML(elemName, attrs);
      retval = board.drainZone;
      return (retval);
    }
    
    private DrainZone buildFromXML(String elemName, Attributes attrs) throws IOException {
      
      String minCol, maxCol;
      
      if (this.isShadow) {
        minCol = AttributeExtractor.extractAttribute(elemName, attrs, "drainZoneShadow", "minCol", true);
        maxCol = AttributeExtractor.extractAttribute(elemName, attrs, "drainZoneShadow", "maxCol", true);
      } else {
        minCol = AttributeExtractor.extractAttribute(elemName, attrs, "drainZone", "minCol", true);
        maxCol = AttributeExtractor.extractAttribute(elemName, attrs, "drainZone", "maxCol", true);
      }
  
      int min = Integer.valueOf(minCol).intValue();
      int max = Integer.valueOf(maxCol).intValue();
      MinMax dzmm = new MinMax(min, max);
      
      return (new DrainZone(dzmm, isShadow));
    }
    
  }
  
  public static class LinkGroupWorker extends AbstractFactoryClient {
    
    public LinkGroupWorker(FabricFactory.FactoryWhiteboard board) {
      super(board);
      FabricFactory.FactoryWhiteboard whiteboard = (FabricFactory.FactoryWhiteboard)sharedWhiteboard_;   
      myKeys_.add("linkGroups");
      installWorker(new LinkGroupTagWorker(whiteboard), new MyGroupGlue());
    }
    
    protected Object localProcessElement(String elemName, Attributes attrs) throws IOException {
      LayoutMode retval = null;
      if (elemName.equals("linkGroups")) {
        String target = AttributeExtractor.extractAttribute(elemName, attrs, "linkGroups", "mode", false);
        String annots = AttributeExtractor.extractAttribute(elemName, attrs, "linkGroups", "view", false);
        if (target == null) {
        	target = LayoutMode.PER_NODE_MODE.getText();   	
        }
        boolean showAnnots = (annots != null) ? Boolean.valueOf(annots) : false; 
  
        FabricFactory.FactoryWhiteboard board = (FabricFactory.FactoryWhiteboard)this.sharedWhiteboard_;
        retval = LayoutMode.fromString(target);
        board.bfn.setLayoutMode(retval);
        board.bfn.setShowLinkGroupAnnotations(showAnnots);

      }
      return (retval);     
    }  
  }

  /***************************************************************************
  **
  ** For XML I/O
  */  
      
  public static class LinkInfoWorker extends AbstractFactoryClient {
    
    public LinkInfoWorker(FabricFactory.FactoryWhiteboard board) {
      super(board);
      myKeys_.add("link");
    }
    
    protected Object localProcessElement(String elemName, Attributes attrs) throws IOException {
      Object retval = null;
      FabricFactory.FactoryWhiteboard board = (FabricFactory.FactoryWhiteboard)this.sharedWhiteboard_;    
      board.linkInfo = buildFromXML(elemName, attrs, board);
      retval = board.linkInfo;
      return (retval);     
    }  
    
    private LinkInfo buildFromXML(String elemName, Attributes attrs, FabricFactory.FactoryWhiteboard board) throws IOException { 
      String src = AttributeExtractor.extractAttribute(elemName, attrs, "link", "src", false);
      src = CharacterEntityMapper.unmapEntities(src, false);
      String trg = AttributeExtractor.extractAttribute(elemName, attrs, "link", "trg", false);
      trg = CharacterEntityMapper.unmapEntities(trg, false);
        
      String srcID = AttributeExtractor.extractAttribute(elemName, attrs, "link", "srcID", false);
      
      String trgID = AttributeExtractor.extractAttribute(elemName, attrs, "link", "trgID", false);
      
      NID.WithName srcNID;
      if (src != null) {
      	// Previously used DataUtil.normKey(src) as argument, but as issue #41 shows,
      	// that is wrong. Need the original string:
      	srcNID = board.legacyMap.get(src);	
      } else if (srcID != null) {
      	srcNID = board.wnMap.get(new NID(srcID));
      } else {
      	throw new IOException();
      }
      
      NID.WithName trgNID;
      if (trg != null) {
      	// Previously used DataUtil.normKey(trg) as argument, but as issue #41 shows,
      	// that is wrong. Need the original string:
      	trgNID = board.legacyMap.get(trg);
      } else if (trgID != null) {
      	trgNID = board.wnMap.get(new NID(trgID));
      } else {
      	throw new IOException();
      }
      
      String rel = AttributeExtractor.extractAttribute(elemName, attrs, "link", "rel", true);
      rel = CharacterEntityMapper.unmapEntities(rel, false);
      String directed = AttributeExtractor.extractAttribute(elemName, attrs, "link", "directed", true);
      Boolean dirObj = Boolean.valueOf(directed);
      String shadow = AttributeExtractor.extractAttribute(elemName, attrs, "link", "shadow", true);
      Boolean shadObj = Boolean.valueOf(shadow);
      FabricLink flink = new FabricLink(new FabricNode(srcNID), new FabricNode(trgNID), rel, shadObj.booleanValue(), dirObj);
      String col = AttributeExtractor.extractAttribute(elemName, attrs, "link", "column", false);
      String shadowCol = AttributeExtractor.extractAttribute(elemName, attrs, "link", "shadowCol", true);
      String srcRow = AttributeExtractor.extractAttribute(elemName, attrs, "link", "srcRow", true);
      String trgRow = AttributeExtractor.extractAttribute(elemName, attrs, "link", "trgRow", true);
      String color = AttributeExtractor.extractAttribute(elemName, attrs, "link", "color", true);
      
      if (!shadObj.booleanValue() && (col == null)) {
        throw new IOException();
      }
         
      LinkInfo retval;
      try {
        int useColumn = (col == null) ? Integer.MIN_VALUE : Integer.valueOf(col).intValue();
        int shadowColumn = Integer.valueOf(shadowCol).intValue();
        int srcRowVal = Integer.valueOf(srcRow).intValue();
        int trgRowVal = Integer.valueOf(trgRow).intValue();
        retval = new LinkInfo(flink, srcRowVal, trgRowVal, useColumn, shadowColumn, color);
      } catch (NumberFormatException nfex) {
        throw new IOException();
      }
      return (retval);
    }
  }
  
   /***************************************************************************
  **
  ** For XML I/O
  */  
      
  public static class LinkGroupTagWorker extends AbstractFactoryClient {
    
    public LinkGroupTagWorker(FabricFactory.FactoryWhiteboard board) {
      super(board);
      myKeys_.add("linkGroup");
    }
    
    protected Object localProcessElement(String elemName, Attributes attrs) throws IOException {
      Object retval = null;
      FabricFactory.FactoryWhiteboard board = (FabricFactory.FactoryWhiteboard)this.sharedWhiteboard_;
      board.groupTag = buildFromXML(elemName, attrs);
      retval = board.groupTag;
      return (retval);     
    }  
    
    private String buildFromXML(String elemName, Attributes attrs) throws IOException { 
      String tag = AttributeExtractor.extractAttribute(elemName, attrs, "linkGroup", "tag", true);
      return (tag);
    }
  }
  
  /***************************************************************************
  **
  ** For XML I/O
  */  
      
  public static class NodeInfoWorker extends AbstractFactoryClient {
    
  	
    public NodeInfoWorker(FabricFactory.FactoryWhiteboard board) {
      super(board);
      myKeys_.add("node");
      installWorker(new DrainZoneWorker(board, false), new DrainZoneGlue());
      installWorker(new DrainZoneWorker(board, true), new DrainZoneGlue());
    }
    
    protected Object localProcessElement(String elemName, Attributes attrs) throws IOException {
      Object retval = null;
      FabricFactory.FactoryWhiteboard board = (FabricFactory.FactoryWhiteboard)this.sharedWhiteboard_;
      if (myKeys_.contains(elemName)) {
        board.nodeInfo = buildFromXML(elemName, attrs, board);
        retval = board.nodeInfo;
      }
      return (retval);     
    }  
    
    private NodeInfo buildFromXML(String elemName, Attributes attrs, FabricFactory.FactoryWhiteboard board) throws IOException {
      String name = AttributeExtractor.extractAttribute(elemName, attrs, "node", "name", true);
      name = CharacterEntityMapper.unmapEntities(name, false);
      String nidStr = AttributeExtractor.extractAttribute(elemName, attrs, "node", "nid", false);

      NID nid;
      NID.WithName nwn;
      if (nidStr != null) {
      	boolean ok = board.ulb.addExistingLabel(nidStr);
      	if (!ok) {
      		throw new IOException();
      	}
      	nid = new NID(nidStr);
      	nwn = new NID.WithName(nid, name);
      } else {
      	nid = board.ulb.getNextOID();
      	nwn = new NID.WithName(nid, name);

      	// Addresses Issue 41. Used DataUtil.normKey(name), but if a node made it
      	// as a separate entity in the past, we should keep them unique. Note that with
      	// NIDs now, we can support "identically" named nodes anyway:
      	board.legacyMap.put(name, nwn);
      }
      board.wnMap.put(nid, nwn);
 
      String row = AttributeExtractor.extractAttribute(elemName, attrs, "node", "row", true);
      String minCol = AttributeExtractor.extractAttribute(elemName, attrs, "node", "minCol", true);
      String maxCol = AttributeExtractor.extractAttribute(elemName, attrs, "node", "maxCol", true);
      String minColSha = AttributeExtractor.extractAttribute(elemName, attrs, "node", "minColSha", true);
      String maxColSha = AttributeExtractor.extractAttribute(elemName, attrs, "node", "maxColSha", true);
      String minDrain = AttributeExtractor.extractAttribute(elemName, attrs, "node", "drainMin", false);
      String maxDrain = AttributeExtractor.extractAttribute(elemName, attrs, "node", "drainMax", false);
      String minDrainSha = AttributeExtractor.extractAttribute(elemName, attrs, "node", "drainMinSha", false);
      String maxDrainSha = AttributeExtractor.extractAttribute(elemName, attrs, "node", "drainMaxSha", false);
      String color = AttributeExtractor.extractAttribute(elemName, attrs, "node", "color", true);
      String cluster = AttributeExtractor.extractAttribute(elemName, attrs, "node", "cluster", false);
      cluster = CharacterEntityMapper.unmapEntities(cluster, false);
      
      NodeInfo retval;
      try {
        int nodeRow = Integer.valueOf(row).intValue();
        retval = new NodeInfo(nid, name, nodeRow, color);
        if (cluster != null) {
        	UiUtil.fixMePrintout("Make cluster assign a list");
        	retval.setCluster(cluster);
        }
        
        int min = Integer.valueOf(minCol).intValue();
        int max = Integer.valueOf(maxCol).intValue();
        retval.updateMinMaxCol(min, false);
        retval.updateMinMaxCol(max, false);
        
        int minSha = Integer.valueOf(minColSha).intValue();
        int maxSha = Integer.valueOf(maxColSha).intValue();
        retval.updateMinMaxCol(minSha, true);
        retval.updateMinMaxCol(maxSha, true);
  
        if (minDrain != null) {
          int minDrVal = Integer.valueOf(minDrain).intValue();
          int maxDrVal = Integer.valueOf(maxDrain).intValue();
          DrainZone dz = new DrainZone(new MinMax(minDrVal, maxDrVal), false);
          retval.addDrainZone(dz);
        }
        if (minDrainSha != null) {
          int minDrValSha = Integer.valueOf(minDrainSha).intValue();
          int maxDrValSha = Integer.valueOf(maxDrainSha).intValue();
          DrainZone dz = new DrainZone(new MinMax(minDrValSha, maxDrValSha), true);
          retval.addDrainZone(dz);
        }
      } catch (NumberFormatException nfex) {
        throw new IOException();
      }
      return (retval);
    }
  }
  
  /***************************************************************************
  ** 
  ** Build extents, used for node shading, node annotations, link annotations
  */

  public static class Extents {
  	public HashMap<Boolean, HashMap<Integer, MinMax>> allLinkExtents;
  	public HashMap<Boolean, HashMap<Integer, MinMax>> allNodeExtents;
  	public HashMap<Boolean, MinMax> allLinkFullRange;
  	public HashMap<Boolean, MinMax> allNodeFullRange;
  	public HashMap<Boolean, Integer> singletonNodeStart;
  		
  	public Extents() { 
  		allLinkExtents = new HashMap<Boolean, HashMap<Integer, MinMax>>();
  	  allNodeExtents = new HashMap<Boolean, HashMap<Integer, MinMax>>();
  	  allLinkFullRange = new HashMap<Boolean, MinMax>();
  	  allNodeFullRange = new HashMap<Boolean, MinMax>();
  	  singletonNodeStart = new HashMap<Boolean, Integer>();
  	  boolean[] builds = new boolean[] {true, false};
  	  for (boolean build : builds) {
    	  MinMax linkFullRange = new MinMax();
    	  HashMap<Integer, MinMax> linkExtents = new HashMap<Integer, MinMax>();
    	  allLinkExtents.put(Boolean.valueOf(build), linkExtents);
    	  allLinkFullRange.put(Boolean.valueOf(build), linkFullRange);
    	  MinMax nodeFullRange = new MinMax();
    	  HashMap<Integer, MinMax> nodeExtents = new HashMap<Integer, MinMax>();
    	  allNodeExtents.put(Boolean.valueOf(build), nodeExtents);
    	  allNodeFullRange.put(Boolean.valueOf(build), nodeFullRange); 
  	  }
  	}
  	
  	public Extents(BioFabricNetwork bfn, BTProgressMonitor monitor) throws AsynchExitRequestException { 
  		this();
    	boolean[] builds = new boolean[] {true, false};
    	
    	HashMap<NID, Integer> singletons = new HashMap<NID, Integer>();
 
      for (boolean build : builds) {
      	Boolean boolKey = Boolean.valueOf(build);
      	MinMax nodeFullRange = allNodeFullRange.get(boolKey).init();
    	  HashMap<Integer, MinMax> nodeExtents = allNodeExtents.get(boolKey);
  
	    	List<BioFabricNetwork.NodeInfo> targets = bfn.getNodeDefList();
	      int numNodes = targets.size();
	      String tag = (build) ? "WithShadows" : "NoShadows";
	          
	      LoopReporter lr = new LoopReporter(targets.size(), 20, monitor, 0.0, 1.0, "progress.buildNodeExtents" + tag);
		    for (int i = 0; i < numNodes; i++) {
		      BioFabricNetwork.NodeInfo node = targets.get(i);
		      int num = node.nodeRow;
		      Integer numObj = Integer.valueOf(num);
		      lr.report();
		      MinMax cols = node.getColRange(build);
		      nodeExtents.put(numObj, cols);
		      nodeFullRange.update(num);
		      if (build) { // only need to do this once....
		        singletons.put(node.getNodeID(), numObj);
		      }
		    }
		    lr.finish();
    	}
      
      for (boolean build : builds) {
    	  MinMax linkFullRange = allLinkFullRange.get(Boolean.valueOf(build)).init();
    	  HashMap<Integer, MinMax> linkExtents = allLinkExtents.get(Boolean.valueOf(build));
    	
		    List<BioFabricNetwork.LinkInfo> links = bfn.getLinkDefList(build);
		    int numLinks = links.size();
		    String tag = (build) ? "WithShadows" : "NoShadows";
	    	LoopReporter lr0 = new LoopReporter(links.size(), 20, monitor, 0.0, 1.0, "progress.buildLinkExtents" + tag);
    
		    for (int i = 0; i < numLinks; i++) {
		      BioFabricNetwork.LinkInfo link = links.get(i);
		      lr0.report();    
		      int num = link.getUseColumn(build);
		      int sRow = link.topRow();
		      int eRow = link.bottomRow();
		      linkExtents.put(Integer.valueOf(num), new MinMax(sRow, eRow));
		      linkFullRange.update(num);
		      if (build) {
		        singletons.remove(link.getSource().getNID().getNID());
		        singletons.remove(link.getTarget().getNID().getNID());
		      }
		    }
		    lr0.finish();
    	}
    
      for (boolean build : builds) {
      	Boolean boolKey = Boolean.valueOf(build);  	
      	String tag = (build) ? "WithShadows" : "NoShadows";
        LoopReporter lr2 = new LoopReporter(singletons.size(), 20, monitor, 0.0, 1.0, "progress.buildSingletonExtents" + tag);
        // Singleton node! We want to know the top of all the singleton nodes so we can
		    // not draw link groups down into that region
        for (NID singNode : singletons.keySet()) {
		      Integer existing = singletonNodeStart.get(boolKey);
		      lr2.report();
		      Integer singMin = null;
		      Integer singNum = singletons.get(singNode);
		      if (existing == null) {
		      	singMin = singNum;
		      } else if (existing.intValue() > singNum.intValue()) {
		      	singMin = singNum;
		      }
		      if (singMin != null) {
		      	singletonNodeStart.put(boolKey, singMin);
		      }	
		    }
		    lr2.finish();
      }
	  }
  }
}
```
