## Supplementary material for "BioFabric Visualization of Network Alignments": BioFabric Source Code: FabricLink.java

```
/*
**    Copyright (C) 2003-2018 Institute for Systems Biology 
**                            Seattle, Washington, USA. 
**
**    This library is free software; you can redistribute it and/or
**    modify it under the terms of the GNU Lesser General Public
**    License as published by the Free Software Foundation; either
**    version 2.1 of the License, or (at your option) any later version.
**
**    This library is distributed in the hope that it will be useful,
**    but WITHOUT ANY WARRANTY; without even the implied warranty of
**    MERCHANTABILITY or FITNESS FOR A PARTICULAR PURPOSE.  See the GNU
**    Lesser General Public License for more details.
**
**    You should have received a copy of the GNU Lesser General Public
**    License along with this library; if not, write to the Free Software
**    Foundation, Inc., 59 Temple Place, Suite 330, Boston, MA  02111-1307  USA
*/

package org.systemsbiology.biofabric.model;

import java.util.Map;

import org.systemsbiology.biofabric.api.io.AttributeKey;
import org.systemsbiology.biofabric.api.model.AugRelation;
import org.systemsbiology.biofabric.api.model.NetLink;
import org.systemsbiology.biofabric.api.model.NetNode;
import org.systemsbiology.biofabric.util.DataUtil;

/****************************************************************************
**
** A Class
*/

public class FabricLink implements NetLink, Cloneable, AttributeKey {
  private NetNode srcID_;
  private NetNode trgID_;
  private String relation_;
  private Boolean directed_;
  private boolean isShadow_;

  public FabricLink(NetNode srcID, NetNode trgID, String relation, boolean isShadow, Boolean directed) {
    if ((srcID == null) || (trgID == null) || (relation == null)) {
      throw new IllegalArgumentException();
    }
    srcID_ = srcID;
    trgID_ = trgID;
    relation_ = relation;
    isShadow_ = isShadow;
    directed_ = directed;
  }
  
  public FabricLink(NetNode srcID, NetNode trgID, String relation, boolean isShadow) {
    this(srcID, trgID, relation, isShadow, null);
  }
    
  public FabricLink flipped() {
    if (isFeedback()) {
      throw new IllegalStateException();
    }
    return (new FabricLink(trgID_, srcID_, relation_, isShadow_, directed_)); 
  }
  
  public boolean directionFrozen() {
    return (directed_ != null); 
  }
  
  public void installDirection(Boolean isDirected) {
    if (directed_ != null) {
      throw new IllegalStateException();
    }
    directed_ = isDirected;
    return;
  }
  
  @Override
  public FabricLink clone() {
    try {
      return ((FabricLink)super.clone());
    } catch (CloneNotSupportedException cnse) {
      throw new IllegalStateException();
    }
  }    
   
  public boolean isShadow() {
    return (isShadow_);
  }
  
  public void dropShadowStatus() {
    isShadow_ = false;
    return;
  }
  
  public NetNode getTrgNode() {
    return (trgID_);
  }

  public NetNode getSrcNode() {
    return (srcID_);
  } 
  
  public String getRelation() {
    return (relation_);
  }
  
  public AugRelation getAugRelation() {
    return (new AugRelation(relation_, isShadow_));
  } 
  
  public boolean isDirected() {
    if (directed_ == null) {
      throw new IllegalStateException(); 
    }  
    return (directed_.booleanValue());
  } 
    
  public boolean isFeedback() {
    return (srcID_.equals(trgID_));
  } 
  
  @Override
  public int hashCode() {
    return (srcID_.hashCode() + trgID_.hashCode() + DataUtil.normKey(relation_).hashCode() + ((isShadow_) ? 17 : 31) + 
            ((directed_ == null) ? 0 : directed_.hashCode()));
  }

  @Override
  public String toString() {
    return ("srcID = " + srcID_ + " trgID = " + trgID_ + "rel = " + relation_ + " directed_ = " + directed_ + " isShadow_ = " + isShadow_);
  }
 
  public String toDisplayString() {
    StringBuffer buf = new StringBuffer();
    buf.append(srcID_.getName());
    buf.append((isDirected()) ? '-' : '\u2190');
    if (isShadow_) {
      buf.append("shdw");
    }
    buf.append("(");
    buf.append(relation_);
    buf.append(")");  
    buf.append('\u2192');  // For bidirectional '\u2194'
    buf.append(trgID_.getName());
    return (buf.toString());
  }
  
  public String toEOAString(Map<NetNode, BioFabricNetwork.NodeInfo> nodeInfo) {
    StringBuffer buf = new StringBuffer();
    buf.append(nodeInfo.get(srcID_).getNodeName());
    if (isShadow_) {
      buf.append(" shdw");
    } else {
      buf.append(" ");
    }
    buf.append("(");
    buf.append(relation_);
    buf.append(") ");
    buf.append(nodeInfo.get(trgID_).getNodeName());
    return (buf.toString());
  }
   
  @Override
  public boolean equals(Object other) {    
    if (other == null) {
      return (false);
    }
    if (other == this) {
      return (true);
    }
    if (!(other instanceof FabricLink)) {
      return (false);
    }
    FabricLink otherLink = (FabricLink)other;
  
    if (!this.srcID_.equals(otherLink.srcID_)) {
      return (false);
    }
    if (!this.trgID_.equals(otherLink.trgID_)) {
      return (false);
    }
    
    if (this.isShadow_ != otherLink.isShadow_) {
      return (false);
    }
    
    if (!DataUtil.normKey(this.relation_).equals(DataUtil.normKey(otherLink.relation_))) {
      return (false);
    }
    
    if (this.directed_ == null) {
      return (otherLink.directed_ == null);
    }   
       
    return (this.directed_.equals(otherLink.directed_));
  }  
 
  public boolean synonymous(NetLink other) {
    if (this.equals(other)) {
      return (true);
    }
    if (this.isDirected() || other.isDirected()) {
      return (false);
    }
    if (!DataUtil.normKey(this.relation_).equals(DataUtil.normKey(other.getRelation()))) {
      return (false);
    }
    if (this.isShadow_ != other.isShadow()) {
      return (false);
    }    
    if (!this.srcID_.equals(other.getTrgNode())) {
      return (false);
    }
    return (this.trgID_.equals(other.getSrcNode()));
  }
   
   public boolean shadowPair(FabricLink other) {
    if (this.equals(other)) {
      return (false);
    }
    if (!this.srcID_.equals(other.srcID_)) {
      return (false);
    }
    if (!this.trgID_.equals(other.trgID_)) {
      return (false);
    }   
    if (!DataUtil.normKey(this.relation_).equals(DataUtil.normKey(other.relation_))) {
      return (false);
    }
    
    if (this.directed_ == null) {
      if (other.directed_ != null) {
        return (false); 
      }
    } else if (!this.directed_.equals(other.directed_)) {
      return (false);
    }
    if (this.isShadow_ == other.isShadow_) {
      return (false);
    }
    return (true);   
  }
}
```
