## Supplementary material for "BioFabric Visualization of Network Alignments": BioFabric Source Code: FabricNode.java

```
/*
**    Copyright (C) 2003-2018 Institute for Systems Biology 
**                            Seattle, Washington, USA. 
**
**    This library is free software; you can redistribute it and/or
**    modify it under the terms of the GNU Lesser General Public
**    License as published by the Free Software Foundation; either
**    version 2.1 of the License, or (at your option) any later version.
**
**    This library is distributed in the hope that it will be useful,
**    but WITHOUT ANY WARRANTY; without even the implied warranty of
**    MERCHANTABILITY or FITNESS FOR A PARTICULAR PURPOSE.  See the GNU
**    Lesser General Public License for more details.
**
**    You should have received a copy of the GNU Lesser General Public
**    License along with this library; if not, write to the Free Software
**    Foundation, Inc., 59 Temple Place, Suite 330, Boston, MA  02111-1307  USA
*/

package org.systemsbiology.biofabric.model;

import java.util.Map;

import org.systemsbiology.biofabric.api.model.AugRelation;
import org.systemsbiology.biofabric.api.model.NetLink;
import org.systemsbiology.biofabric.api.model.NetNode;
import org.systemsbiology.biofabric.api.util.NID;
import org.systemsbiology.biofabric.api.util.NID.WithName;
import org.systemsbiology.biofabric.io.AttributeLoader;
import org.systemsbiology.biofabric.util.DataUtil;

/****************************************************************************
**
** A Class
*/

public class FabricNode implements NetNode {
  private NID.WithName id_;

  public FabricNode(NID id, String name) {
    id_ = new NID.WithName(id, name);
  } 
  
  public FabricNode(NID.WithName id) {
    id_ = id;
  }
   
  public String getName() {
    return (id_.getName());
  }

  public NID.WithName getNID() {
    return (id_);
  } 
  
	@Override
  public FabricNode clone() {
    try {
      FabricNode newVal = (FabricNode)super.clone();
      newVal.id_ = this.id_.clone();
      return (newVal);
    } catch (CloneNotSupportedException ex) {
      throw new IllegalStateException();     
    }    
  }

  public int compareTo(NetNode other) {
	  return (this.id_.compareTo(other.getNID()));
	}
	 
  @Override
  public int hashCode() {
    return (id_.hashCode());
  }

  @Override
  public String toString() {
    return ("id = " + id_);
  }
   
  @Override
  public boolean equals(Object other) {    
    if (other == null) {
      return (false);
    }
    if (other == this) {
      return (true);
    }
    if (!(other instanceof FabricNode)) {
      return (false);
    }
    FabricNode otherLink = (FabricNode)other;
  
    return (this.id_.equals(otherLink.id_));
  }
}
```
