## Supplementary material for "BioFabric Visualization of Network Alignments": BioFabric Source Code: ProgressFilterInputStream.java

```
/*
**    Copyright (C) 2003-2017 Institute for Systems Biology 
**                            Seattle, Washington, USA. 
**
**    This library is free software; you can redistribute it and/or
**    modify it under the terms of the GNU Lesser General Public
**    License as published by the Free Software Foundation; either
**    version 2.1 of the License, or (at your option) any later version.
**
**    This library is distributed in the hope that it will be useful,
**    but WITHOUT ANY WARRANTY; without even the implied warranty of
**    MERCHANTABILITY or FITNESS FOR A PARTICULAR PURPOSE.  See the GNU
**    Lesser General Public License for more details.
**
**    You should have received a copy of the GNU Lesser General Public
**    License along with this library; if not, write to the Free Software
**    Foundation, Inc., 59 Temple Place, Suite 330, Boston, MA  02111-1307  USA
*/

package org.systemsbiology.biofabric.parser;

import java.io.FilterInputStream;
import java.io.IOException;
import java.io.InputStream;

/****************************************************************************
**
** We need to track the processed byte count of the XML parser.
**
*/

public class ProgressFilterInputStream extends FilterInputStream {
  
  ////////////////////////////////////////////////////////////////////////////
  //
  // PRIVATE INSTANCES
  //
  ////////////////////////////////////////////////////////////////////////////
  
  private long count_;
  private long total_;
  private double progress_;
  
  ////////////////////////////////////////////////////////////////////////////
  //
  // PUBLIC CONSTRUCTORS
  //
  ////////////////////////////////////////////////////////////////////////////

  /***************************************************************************
  **
  ** Standard constructor:
  */

  public ProgressFilterInputStream(InputStream in, long total) {
    super(in);
    total_ = total;
  }

  ////////////////////////////////////////////////////////////////////////////
  //
  // PUBLIC METHODS
  //
  ////////////////////////////////////////////////////////////////////////////

  /***************************************************************************
  ** 
  ** Return current progress:
  */

  public double getProgress(){
    return (progress_);
  }
 
  /***************************************************************************
  ** 
  ** Override the core function to track the bytes passing by:
  */

  @Override
  public int read(byte[] b, int off, int len) throws IOException {
    int numRead = super.read(b, off, len);
    if (numRead == -1) {
    	progress_ = 1.0;
    } else {
      count_ += numRead;
      progress_ = (double)count_ / (double)total_;
      progress_ = Math.min(1.0, progress_);
    }
    return (numRead);
  }
}
```
