## Supplementary material for "BioFabric Visualization of Network Alignments": BioFabric Source Code: SUParser.java

```
/*
**    Copyright (C) 2003-2017 Institute for Systems Biology 
**                            Seattle, Washington, USA. 
**
**    This library is free software; you can redistribute it and/or
**    modify it under the terms of the GNU Lesser General Public
**    License as published by the Free Software Foundation; either
**    version 2.1 of the License, or (at your option) any later version.
**
**    This library is distributed in the hope that it will be useful,
**    but WITHOUT ANY WARRANTY; without even the implied warranty of
**    MERCHANTABILITY or FITNESS FOR A PARTICULAR PURPOSE.  See the GNU
**    Lesser General Public License for more details.
**
**    You should have received a copy of the GNU Lesser General Public
**    License along with this library; if not, write to the Free Software
**    Foundation, Inc., 59 Temple Place, Suite 330, Boston, MA  02111-1307  USA
*/

package org.systemsbiology.biofabric.parser;

import java.io.IOException;
import java.util.Iterator;
import java.util.HashMap;
import java.util.HashSet;
import java.util.Set;
import java.util.List;
import java.io.File;
import java.net.URL;
import java.text.MessageFormat;

import org.xml.sax.EntityResolver;
import org.xml.sax.Attributes;
import org.xml.sax.SAXException;
import org.xml.sax.SAXParseException;
import org.xml.sax.XMLReader;
import org.xml.sax.InputSource;
import org.xml.sax.helpers.XMLReaderFactory;
import org.xml.sax.helpers.DefaultHandler;
import org.systemsbiology.biofabric.api.parser.ParserClient;
import org.systemsbiology.biofabric.api.worker.AsynchExitRequestException;
import org.systemsbiology.biofabric.api.worker.BTProgressMonitor;
import org.systemsbiology.biofabric.api.worker.LoopReporter;
import org.systemsbiology.biofabric.util.ResourceManager;

/****************************************************************************
**
** Parse a BioTapestry file
**
**
*/

public class SUParser extends DefaultHandler {

	private final static String ASYNC_FLAG_ = "__AEXR__";
	
  ////////////////////////////////////////////////////////////////////////////
  //
  // PRIVATE INSTANCES
  //
  ////////////////////////////////////////////////////////////////////////////
  
  private XMLReader parser_;
  private HashSet<Double> seen_;
  private HashMap<String, ParserClient> clients_;
  private ParserClient currClient_;
  private String lastElement_;
  private ProgressFilterInputStream pfis_;  
  private BTProgressMonitor monitor_; 
  private double startFrac_;
  private double endFrac_;
  private LoopReporter lr_;
  
  ////////////////////////////////////////////////////////////////////////////
  //
  // PUBLIC CONSTRUCTORS
  //
  ////////////////////////////////////////////////////////////////////////////

  /***************************************************************************
  **
  ** Parse the given file using the given factories:
  */

  public SUParser(List<ParserClient> citList) {
    this();
    //
    // Crank thru the clients and stash them in a HashMap
    //
    
    Iterator<ParserClient> cit = citList.iterator();
    clients_ = new HashMap<String, ParserClient>();
    while (cit.hasNext()) {
      ParserClient pc = cit.next();
      Set<String> keys = pc.keywordsOfInterest();
      Iterator<String> ki = keys.iterator();
      while (ki.hasNext()) {
        String key = ki.next();
        Object prev = clients_.put(key, pc);
        if (prev != null) {
          throw new IllegalArgumentException();
        }
      }
    }
    currClient_ = null;
    lastElement_ = null;
  }

  ////////////////////////////////////////////////////////////////////////////
  //
  // PUBLIC METHODS
  //
  ////////////////////////////////////////////////////////////////////////////

  /***************************************************************************
  ** 
  ** Parse the given file
  */

  public void parse(File toParse) throws IOException {
    parse(toParse.getPath());
    return;
  }
  
  /***************************************************************************
  ** 
  ** Parse the given file
  */

  public void parse(String fileName) throws IOException {    
    if (fileName.indexOf("file:///") != 0) {
      fileName = "file:///" + fileName;
    }
    try {
      parser_.parse(fileName);
    } catch (SAXException e) {
      String msg = formatSAXExceptionMessage(e);
      System.err.println("Got a SAX exception: " + msg);
      throw new IOException(msg);
    } catch (IOException e) {
      System.err.println("Got an IO exception: " + e);
      throw rebundleIOException(e);
    }
  }

  /***************************************************************************
  ** 
  ** Parse the given URL
  */

  public void parse(URL source) throws IOException {
    try {
      parser_.parse(new InputSource(source.openStream()));
    } catch (SAXException e) {
      String msg = formatSAXExceptionMessage(e);
      System.err.println("Got a SAX exception: " + msg);
      throw new IOException(msg);
    } catch (IOException e) {
      System.err.println("Got an IO exception: " + e);
      throw rebundleIOException(e);
    }
  } 
  
  /***************************************************************************
  ** 
  ** Parse the given input stream
  */

  public void parse(ProgressFilterInputStream pfis, BTProgressMonitor monitor, boolean forCache) 
    throws AsynchExitRequestException, IOException {
  	
  	pfis_ = pfis;
  	seen_ = new HashSet<Double>();
  	monitor_ = monitor;
  	lr_ = new LoopReporter(20, 0, monitor_, 0.0, 1.0, (forCache) ? "progress.fromCache" : "progress.readXML");
  	
    try {
      parser_.parse(new InputSource(pfis_));
    } catch (SAXException e) {
    	if (e.getMessage().equals(ASYNC_FLAG_)) {
    		throw new AsynchExitRequestException();
    	}
      String msg = formatSAXExceptionMessage(e);
      System.err.println("Got a SAX exception: " + msg);
      throw new IOException(msg);
    } catch (IOException e) {
      System.err.println("Got an IO exception: " + e);
      throw rebundleIOException(e);
    }
    return;
  }  

  /***************************************************************************
  ** 
  ** Called at start of the document
  */

  @Override
  public void startDocument() throws SAXException {
    return;
  }

  /***************************************************************************
  ** 
  ** Called at start of an element
  */

  @Override
  public void startElement(String uri, String local, String raw,
                           Attributes attrs) throws SAXException {

  	if (pfis_ != null) {
	  	double prog = pfis_.getProgress();
	  	Double step = Double.valueOf(Math.floor(prog * 20.0));
	  	if (!seen_.contains(step)) {
	  		seen_.add(step);
	  		try {
	  		  lr_.report();
	  		} catch (AsynchExitRequestException aex) {
          throw (rebundleAsyncExceptionToSAX(aex));
        }
	  	}
  	}
  		
    //
    // If we have a current client, we send him the element.  If
    // not, we find the client that is interested in the element
    // and send it to him:
    //
  	
    if (currClient_ != null) {
      try {
        lastElement_ = local;
        Object target = currClient_.processElement(local, attrs);
        if (target != null) {
          setTargets(target);
        }
      } catch (IOException ioe) {
        throw (rebundleIOExceptionToSAX(ioe));
      }
      return;
    }
    
    //
    // Find the client that wants it:
    //
    
    ParserClient pc = clients_.get(local);
    if (pc != null) {
      currClient_ = pc; 
      try {
        lastElement_ = local;
        Object target = currClient_.processElement(local, attrs);
        if (target != null) {
          setTargets(target);
        }
      } catch (IOException ioe) {
        throw (rebundleIOExceptionToSAX(ioe));
      }
    }
    return;
  }

  /***************************************************************************
  ** 
  ** Called at the end of an element
  */  
  
  @Override
  public void endElement(String uri, String local, String raw) throws SAXException {
    if (currClient_ == null) {
      return;
    }
    try {
      if (currClient_.finishElement(local)) {
        currClient_ = null;
      }
    } catch (IOException ioe) {
      throw (rebundleIOExceptionToSAX(ioe));
    }
    return;
  } 
    
  /***************************************************************************
  ** 
  ** Called for characters
  */

  @Override
  public void characters(char ch[], int start, int length) throws SAXException {
    if (currClient_ != null) {
      currClient_.processCharacters(ch, start, length);
    }
    return; 
  }

  /***************************************************************************
  ** 
  ** Called for ignorable whitespace
  */

  @Override
  public void ignorableWhitespace(char ch[], int start, int length) throws SAXException {
    return;
  }

  /***************************************************************************
  ** 
  ** Error callbacks
  */

  @Override
  public void warning(SAXParseException ex) throws SAXException {
    printError("Warning", ex);
  }

  @Override
  public void error(SAXParseException ex) throws SAXException {
    printError("Error", ex);
  }

  @Override
  public void fatalError(SAXParseException ex) throws SAXException {
    printError("Fatal Error", ex);
  }

  ////////////////////////////////////////////////////////////////////////////
  //
  // PUBLIC CLASS METHODS
  //
  ////////////////////////////////////////////////////////////////////////////

  /***************************************************************************
  ** 
  ** Main test harness
  */

  public static void main(String argv[]) {
    try {
      SUParser sup = new SUParser();
      sup.parse("FIXME");
    } catch (IOException e) {
      System.err.println("Got an IO exception: " + e);
    }
  }
  
  ////////////////////////////////////////////////////////////////////////////
  //
  // PRIVATE CONSTRUCTORS
  //
  ////////////////////////////////////////////////////////////////////////////
    
  /***************************************************************************
  **
  ** Null Constructor.
  */

  private SUParser() {
    try {
      //SAXParserFactory saxParserFactory = SAXParserFactory.newInstance();
      //saxParserFactory.setNamespaceAware(true);
      //saxParserFactory.setValidating(true);
      //SAXParser saxParser = saxParserFactory.newSAXParser();
      //parser_ = saxParser.getXMLReader();           
      //String pName = "org.apache.xerces.parsers.SAXParser";
      
      //
      // Starting with 1.5, we need to use a different parser (BT-05-18-05:1):
      //
      
      String jVer = System.getProperty("java.version");
      String pName;
      if (jVer.startsWith("1.4")) {
        pName = "org.apache.crimson.parser.XMLReaderImpl";
      } else {
        pName = "com.sun.org.apache.xerces.internal.parsers.SAXParser";       
      }
      
      parser_ = XMLReaderFactory.createXMLReader(pName);
      //parser_ = XMLReaderFactory.createXMLReader();

      parser_.setContentHandler(this);
      parser_.setErrorHandler(this);      
      //
      // Uncomment to get validation(as well as adding DOCTYPE elements to XML file)
      //
      // String valFeature = "http://xml.org/sax/features/validation";
      // parser_.setFeature(valFeature, true);
      // parser_.setEntityResolver(
      //  new MyEntityResolver("/org/systemsbiology/biotapestry/util/qpcr.dtd"));
 
    } catch (SAXException e) {
      System.err.println("Got a SAX exception: " + e);
   // } catch (ParserConfigurationException pcex) {
    //  System.err.println("Got a ParserConfigurationException: " + pcex);
    }
  }
  
  ////////////////////////////////////////////////////////////////////////////
  //
  // PRIVATE METHODS
  //
  ////////////////////////////////////////////////////////////////////////////

  /***************************************************************************
  ** 
  ** Handles error printouts
  */

  private void printError(String type, SAXParseException ex) {
    System.err.print("Error: ");
    System.err.println(type);
    if (ex == null) {
      return;
    }
    String sid = ex.getSystemId();
    if (sid != null) {
      System.err.println(sid);
    }
    System.err.print("Line number: ");
    System.err.println(ex.getLineNumber());
    System.err.print("Column number:");
    System.err.println(ex.getColumnNumber());
    System.err.print("Message: ");
    System.err.println(ex.getMessage());
    return;
  }
  

  /***************************************************************************
  ** 
  ** Handles setting targets
  */
  
  private void setTargets(Object target) {
    Iterator<ParserClient> ci = clients_.values().iterator();
    while (ci.hasNext()) {
      ParserClient cli = ci.next();
      cli.setContainer(target);
    }
    return;
  }
  
  /***************************************************************************
  ** 
  ** Handles message formatting
  */
  
  private String formatSAXExceptionMessage(SAXException e) {
    ResourceManager rMan = ResourceManager.getManager();
    String exMsg = e.getMessage();
    Integer lineNo = null;
    Integer colNo = null;
    if (e instanceof SAXParseException) {
      SAXParseException spe = (SAXParseException)e;
      lineNo = new Integer(spe.getLineNumber());
      colNo = new Integer(spe.getColumnNumber());
    }
    String retval;
    if (lineNo == null) {
      String format = rMan.getString("fileRead.SAXErrorFormat");
      retval = MessageFormat.format(format, new Object[] {exMsg});  
    } else {
      String format = rMan.getString("fileRead.SAXParseErrorFormat");
      retval = MessageFormat.format(format, new Object[] {exMsg, lineNo, colNo});  
    }
    return (retval);
  }
  
  /***************************************************************************
  ** 
  ** Rebundle IO exception
  */
  
  private IOException rebundleIOException(IOException e) {
    return (new IOException(buildIOExceptionMsg(e)));       
  }
  
  /***************************************************************************
  ** 
  ** Rebundle IO exception to SAX
  */
  
  private SAXException rebundleIOExceptionToSAX(IOException e) {
    return (new SAXException(buildIOExceptionMsg(e)));       
  }  
   
  /***************************************************************************
  ** 
  ** Rebundle Async exception to SAX
  */
  
  private SAXException rebundleAsyncExceptionToSAX(AsynchExitRequestException amex) {
    return (new SAXException(ASYNC_FLAG_));       
  }  
  
  /***************************************************************************
  ** 
  ** Rebundle IO exception
  */
  
  private String buildIOExceptionMsg(IOException e) {
    ResourceManager rMan = ResourceManager.getManager();
    String msg = (e == null) ? null : e.getMessage();
    String retmsg;
    if ((msg == null) || msg.trim().equals("")) {
      if (lastElement_ == null) {
        retmsg = rMan.getString("fileRead.IOErrorNoDetails");
      } else {
        String format = rMan.getString("fileRead.IOErrorNoMsgWithElemFormat");
        retmsg = MessageFormat.format(format, new Object[] {lastElement_});
      }
    } else {
      if (lastElement_ == null) {
        String format = rMan.getString("fileRead.IOErrorMsgNoElemFormat");
        retmsg = MessageFormat.format(format, new Object[] {msg});
      } else {
        String format = rMan.getString("fileRead.IOExtraMsgWithElemFormat");
        retmsg = MessageFormat.format(format, new Object[] {msg, lastElement_});                    
      }
    }
    return (retmsg);       
  }  
  
  ////////////////////////////////////////////////////////////////////////////
  //
  // PRIVATE INNER CLASSES
  //
  ////////////////////////////////////////////////////////////////////////////

  /*****************************************************************************
  **
  ** Allows use of local character streams for DTDs.
  */
  
  class MyEntityResolver implements EntityResolver {

    private String resource_;

    public MyEntityResolver(String resource) {
      resource_ = resource;
    }

    public InputSource resolveEntity(String publicID, String systemId)
      throws SAXException, IOException {
      return (new InputSource(getClass().getResourceAsStream(resource_)));
    }
  }
}
```
