## Supplementary material for "BioFabric Visualization of Network Alignments": BioFabric Source Code: AbstractPlugInDirective.java

```
/*
**    Copyright (C) 2003-2018 Institute for Systems Biology 
**                            Seattle, Washington, USA. 
**
**    This library is free software; you can redistribute it and/or
**    modify it under the terms of the GNU Lesser General Public
**    License as published by the Free Software Foundation; either
**    version 2.1 of the License, or (at your option) any later version.
**
**    This library is distributed in the hope that it will be useful,
**    but WITHOUT ANY WARRANTY; without even the implied warranty of
**    MERCHANTABILITY or FITNESS FOR A PARTICULAR PURPOSE.  See the GNU
**    Lesser General Public License for more details.
**
**    You should have received a copy of the GNU Lesser General Public
**    License along with this library; if not, write to the Free Software
**    Foundation, Inc., 59 Temple Place, Suite 330, Boston, MA  02111-1307  USA
*/

package org.systemsbiology.biofabric.plugin;

import java.io.IOException;
import java.io.File;

import java.net.URL;
import java.net.URLClassLoader;
import java.net.MalformedURLException;

import org.systemsbiology.biofabric.api.io.AttributeExtractor;
import org.xml.sax.Attributes;

/***************************************************************************
**
** Used to specify plugins
*/
  
public abstract class AbstractPlugInDirective implements Comparable<AbstractPlugInDirective> {
 
  ////////////////////////////////////////////////////////////////////////////
  //
  // PUBLIC CONSTANTS
  //
  ////////////////////////////////////////////////////////////////////////////  
 
  ////////////////////////////////////////////////////////////////////////////
  //
  // PROTECTED VARIABLES
  //
  ////////////////////////////////////////////////////////////////////////////
  
  protected String className_;
  protected int order_;
  protected File jar_;  

  ////////////////////////////////////////////////////////////////////////////
  //
  // CONSTRUCTORS
  //
  ////////////////////////////////////////////////////////////////////////////    

  /***************************************************************************
  **
  ** Constructor for internal use
  **
  */
  
  public AbstractPlugInDirective(String className, String order, File jarFile) {
    stockCore(className, order);
    jar_ = jarFile;
  }  

  /***************************************************************************
  **
  ** Constructor for internal use
  **
  */
  
  protected AbstractPlugInDirective() {
    jar_ = null;
  }
 
  ////////////////////////////////////////////////////////////////////////////
  //
  // PUBLIC METHODS
  //
  ////////////////////////////////////////////////////////////////////////////  
  
  /***************************************************************************
  **
  ** Get the plugin order
  */
  
  public int getOrder() {
    return (order_);
  }
 
  /***************************************************************************
  **
  ** Get the plugin class name
  */
  
  public String getClassName() {
    return (className_);
  }
  
  /***************************************************************************
  **
  ** Get the plugin jar file
  */
  
  public File getJar() {
    return (jar_);
  }  
  
  /***************************************************************************
  **
  ** Build a plug-in from the directive:
  **
  */
  
  public BioFabricToolPlugIn buildPlugIn(PlugInManager pMan) {
    if (jar_ == null) {
      return (manufacture(pMan));
    } else {
      return (load(pMan));
    }
  }
  
  /***************************************************************************
  **
  ** Implement comparable interface:
  **
  */
    
  public int compareTo(AbstractPlugInDirective other) {
    
    if (this.order_ != other.order_) {
      return ((this.order_ > other.order_) ? 1 : -1);
    }
    //
    // Throw up my hands:
    //
    return (this.className_.compareTo(other.className_));
  }
  
  ////////////////////////////////////////////////////////////////////////////
  //
  // PROTECTED METHODS
  //
  ////////////////////////////////////////////////////////////////////////////
  
  /***************************************************************************
  **
  ** Handle the attributes for the keyword
  **
  */
  
  protected void stockCoreFromXML(String elemName, 
                                  Attributes attrs) throws IOException {
                                                
    String className = AttributeExtractor.extractAttribute(elemName, attrs, elemName, "class", true);
    String orderStr = AttributeExtractor.extractAttribute(elemName, attrs, elemName, "order", true);
    try {
      stockCore(className, orderStr);        
    } catch (IllegalArgumentException iex) {
      throw new IOException();
    }
    return; 
  }
  
  /***************************************************************************
  **
  ** Stock core fields
  **
  */
  
  private void stockCore(String className, String order) throws IllegalArgumentException {

    this.className_ = className;
    
    try {
      order_ = Integer.parseInt(order);
    } catch (NumberFormatException nfe) {
      throw new IllegalArgumentException();
    }
    
    return;
  }
  
  /***************************************************************************
  **
  ** Create a plug-in from the directive:
  **
  */
      
  private BioFabricToolPlugIn load(PlugInManager pMan) {
    try {
      URL purl = jar_.toURI().toURL(); // As recommended in docs, to create escape chars
      URLClassLoader loader = new URLClassLoader(new URL[] {purl},BioFabricToolPlugIn.class.getClassLoader());  
      
      Class<?> pluggedIn = Class.forName(className_, true, loader);

      BioFabricToolPlugIn instance = (BioFabricToolPlugIn)pluggedIn.newInstance();
      instance.setUniquePlugInTag(className_);
      instance.installManager(pMan);
      return (instance);
    } catch (MalformedURLException muex) {
      System.err.println("PlugIn " + className_ + " not loaded: " + muex);      
    } catch (ClassNotFoundException cnfex) {
      System.err.println("PlugIn " + className_ + " not loaded: " + cnfex);
    } catch (InstantiationException iex) {
      System.err.println("PlugIn " + className_ + " not loaded: " + iex);
    } catch (IllegalAccessException iex) {
      System.err.println("PlugIn " + className_ + " not loaded: " + iex);
    }  
    return (null);
  }
  
  /***************************************************************************
  **
  ** Create a plug-in from the directive:
  */

  private BioFabricToolPlugIn manufacture(PlugInManager pMan) {
    try {
      Class<?> plugClass = Class.forName(className_);
      BioFabricToolPlugIn instance = (BioFabricToolPlugIn)plugClass.newInstance();
      instance.setUniquePlugInTag(className_);
      instance.installManager(pMan);
      return (instance);      
    } catch (ClassNotFoundException cnfex) {
      System.err.println("PlugIn " + className_ + " not loaded: " + cnfex);
    } catch (InstantiationException iex) {
      System.err.println("PlugIn " + className_ + " not loaded: " + iex);
    } catch (IllegalAccessException iex) {
      System.err.println("PlugIn " + className_ + " not loaded: " + iex);
    }  
    return (null);
  }
  
}
```
