## Supplementary material for "BioFabric Visualization of Network Alignments": BioFabric Source Code: BioFabricToolPlugIn.java

```
/*
**    Copyright (C) 2003-2018 Institute for Systems Biology 
**                            Seattle, Washington, USA. 
**
**    This library is free software; you can redistribute it and/or
**    modify it under the terms of the GNU Lesser General Public
**    License as published by the Free Software Foundation; either
**    version 2.1 of the License, or (at your option) any later version.
**
**    This library is distributed in the hope that it will be useful,
**    but WITHOUT ANY WARRANTY; without even the implied warranty of
**    MERCHANTABILITY or FITNESS FOR A PARTICULAR PURPOSE.  See the GNU
**    Lesser General Public License for more details.
**
**    You should have received a copy of the GNU Lesser General Public
**    License along with this library; if not, write to the Free Software
**    Foundation, Inc., 59 Temple Place, Suite 330, Boston, MA  02111-1307  USA
*/

package org.systemsbiology.biofabric.plugin;

import java.io.PrintWriter;

import org.systemsbiology.biofabric.api.io.Indenter;
import org.systemsbiology.biofabric.api.io.PluginWhiteboard;
import org.systemsbiology.biofabric.api.model.Network;
import org.systemsbiology.biofabric.api.parser.AbstractFactoryClient;

/****************************************************************************
**
** Interface for plugins that can implement BioFabric workflows
*/

public interface BioFabricToolPlugIn {
  
  ////////////////////////////////////////////////////////////////////////////
  //
  // PUBLIC METHODS
  //
  ////////////////////////////////////////////////////////////////////////////
 
  /***************************************************************************
  **
  ** Set the unique plugin tag:
  */
  
  public void setUniquePlugInTag(String tag);
  
  /***************************************************************************
  **
  ** Get the unique tag
  */
  
  public String getUniquePlugInTag();
   
  /***************************************************************************
  **
  ** Get name of tool menu to display
  */
  
  public String getToolMenu();
  
  /***************************************************************************
  **
  ** Get count of commands
  */
  
  public int getCommandCount();
  
  /***************************************************************************
  **
  ** Get the nth command
  */
  
  public BioFabricToolPlugInCmd getCommand(int index);
   
  /***************************************************************************
  **
  ** Dump out using XML
  */
   
  public void writeXML(PrintWriter out, Indenter ind);   

  /***************************************************************************
  **
  ** Get XML Reader
  */
 
  public AbstractFactoryClient getXMLWorker(PluginWhiteboard board);
  
  /***************************************************************************
  **
  ** Attach session data read from XML
  */
 
  public void attachXMLData(BioFabricToolPlugInData data);
  
  /***************************************************************************
  **
  ** Install a new network
  */
  
  public void newNetworkInstalled(Network bfn);
  
  /***************************************************************************
  **
  ** Install API
  */
  
  public void installAPI(PlugInNetworkModelAPI bfn);
  
  /***************************************************************************
  **
  ** Install PluginManager
  */
  
  public void installManager(PlugInManager pMan);  
  
    
}
```
