## Supplementary material for "BioFabric Visualization of Network Alignments": BioFabric Source Code: NetStatsPlugIn.java

```
/*
**    Copyright (C) 2003-2018 Institute for Systems Biology 
**                            Seattle, Washington, USA. 
**
**    This library is free software; you can redistribute it and/or
**    modify it under the terms of the GNU Lesser General Public
**    License as published by the Free Software Foundation; either
**    version 2.1 of the License, or (at your option) any later version.
**
**    This library is distributed in the hope that it will be useful,
**    but WITHOUT ANY WARRANTY; without even the implied warranty of
**    MERCHANTABILITY or FITNESS FOR A PARTICULAR PURPOSE.  See the GNU
**    Lesser General Public License for more details.
**
**    You should have received a copy of the GNU Lesser General Public
**    License along with this library; if not, write to the Free Software
**    Foundation, Inc., 59 Temple Place, Suite 330, Boston, MA  02111-1307  USA
*/

package org.systemsbiology.biofabric.plugin.core;

import java.io.IOException;
import java.io.PrintWriter;
import java.text.MessageFormat;
import java.util.ArrayList;

import javax.swing.JFrame;
import javax.swing.JOptionPane;

import org.systemsbiology.biofabric.api.io.AttributeExtractor;
import org.systemsbiology.biofabric.api.io.Indenter;
import org.systemsbiology.biofabric.api.io.PluginWhiteboard;
import org.systemsbiology.biofabric.api.model.Network;
import org.systemsbiology.biofabric.api.parser.AbstractFactoryClient;
import org.systemsbiology.biofabric.api.parser.GlueStick;
import org.systemsbiology.biofabric.api.util.PluginResourceManager;
import org.systemsbiology.biofabric.plugin.BioFabricToolPlugIn;
import org.systemsbiology.biofabric.plugin.BioFabricToolPlugInCmd;
import org.systemsbiology.biofabric.plugin.BioFabricToolPlugInData;
import org.systemsbiology.biofabric.plugin.PlugInManager;
import org.systemsbiology.biofabric.plugin.PlugInNetworkModelAPI;
import org.systemsbiology.biofabric.plugin.PluginSupportFactory;
import org.systemsbiology.biofabric.util.UiUtil;

import org.xml.sax.Attributes;

/****************************************************************************
**
** Class for the simple network statistics tool 
*/

public class NetStatsPlugIn implements BioFabricToolPlugIn {
  
  private ArrayList<BioFabricToolPlugInCmd> myCmds_;
  private String myTag_;
  private StatData myData_;
  private String className_;
  private PluginResourceManager rMan_;
  
  ////////////////////////////////////////////////////////////////////////////
  //
  // PUBLIC METHODS
  //
  ////////////////////////////////////////////////////////////////////////////
  
  /***************************************************************************
  **
  ** Null constructor (required)
  */
  
  public NetStatsPlugIn() {
    myCmds_ = new ArrayList<BioFabricToolPlugInCmd>();
    myCmds_.add(new NodeAndLinkCounterCmd());
    myData_ = new StatData(0, 0, 0);
    className_ = getClass().getName();
 
  }
  
  ////////////////////////////////////////////////////////////////////////////
  //
  // PUBLIC METHODS
  //
  ////////////////////////////////////////////////////////////////////////////
  
  /***************************************************************************
  **
  ** Set the unique tag
  */
  
  public void setUniquePlugInTag(String tag) {
    myTag_ = tag;
    return;
  }
  
  /***************************************************************************
  **
  ** Get the unique tag
  */
  
  public String getUniquePlugInTag() {
    return (myTag_);
  }

  /***************************************************************************
  **
  ** Install PluginManager
  */
  
  public void installManager(PlugInManager pMan) {
  	rMan_ = PluginSupportFactory.getResourceManager(className_, pMan);
    rMan_.setPluginBundle("org.systemsbiology.biofabric.plugin.core.NetStats"); 
    return;
  }
  
  
  /***************************************************************************
  **
  ** InstallAPI
  */
  
  public void installAPI(PlugInNetworkModelAPI bfn) {
    // Nothing to do...
    return;
  }
  
  /***************************************************************************
  **
  ** Install a new network
  */
  
  public void newNetworkInstalled(Network bfn) {
    for (BioFabricToolPlugInCmd cmd : myCmds_) {
      NodeAndLinkCounterCmd nalc = (NodeAndLinkCounterCmd)cmd;
      nalc.setNewNetwork(bfn);
    }
    return;
  }
  
  /***************************************************************************
  **
  ** Get name of tool menu to display
  */
  
  public String getToolMenu() {
    return (rMan_.getPluginString("command.pluginName"));
  }
  
  /***************************************************************************
  **
  ** Get count of commands
  */
  
  public int getCommandCount() {
    return (myCmds_.size());   
  }
  
  /***************************************************************************
  **
  ** Get the nth command
  */
  
  public BioFabricToolPlugInCmd getCommand(int index) {
    return (myCmds_.get(index));
  }
   
  /***************************************************************************
  **
  ** Write session data to given output
  */
  
  public void writeXML(PrintWriter out, Indenter ind) {  
    ind.up();
    ind.indent();
    
    String name = getClass().getName();
    
    out.print("<");
    out.print(name);
    out.println(">");
    ind.up();
    ind.indent();
    out.print("<netStats nodes=\"");
    out.print(myData_.nodeCount);
    out.print("\" links=\"");
    out.print(myData_.linkCount);
    out.print("\" fullShadowLinks=\"");
    out.print(myData_.fullShadowLinkCount);
    out.println("\" />");
    ind.down();
    ind.indent();
    out.print("</");
    out.print(name);
    out.println(">");
    ind.down();
    return;
  }  
  
  /***************************************************************************
  **
  ** Get XML Reader
  */
 
  public AbstractFactoryClient getXMLWorker(PluginWhiteboard board) {
    return (new PlugInWorker(board, this));
  }
  
  /***************************************************************************
  **
  ** Attach session data read from XML
  */
 
  public void attachXMLData(BioFabricToolPlugInData data) {
    myData_ = (StatData)data;
    return;   
  }
  
  ////////////////////////////////////////////////////////////////////////////
  //
  // INNER CLASSES
  //
  ////////////////////////////////////////////////////////////////////////////

  public class NodeAndLinkCounterCmd implements BioFabricToolPlugInCmd {    

    private boolean enabled_;
    
    /***************************************************************************
    **
    ** Set new network. In this case, process the network too
    */
    
    public void setNewNetwork(Network bfn) {
      if (bfn != null) {
        myData_.nodeCount = bfn.getNodeCount();
        myData_.linkCount = bfn.getLinkCount(false);
        myData_.fullShadowLinkCount = bfn.getLinkCount(true);
        enabled_ = (myData_.nodeCount > 0);
      } else {
        myData_.nodeCount = 0;
        myData_.linkCount = 0;
        myData_.fullShadowLinkCount = 0;
        enabled_ = false;      
      }
      return;
    }

    /***************************************************************************
    **
    ** Get the name
    */
    
    public String getCommandName() {
      return (rMan_.getPluginString("command.statsCommands"));
    }

    /***************************************************************************
    **
    ** Perform the operation
    */
    
    public boolean performOperation(JFrame topFrame) {
      if (!enabled_) {
        return (false);
      }
      
      String desc = MessageFormat.format(rMan_.getPluginString("modelCounts.message"), 
                                         new Object[] {new Integer(myData_.nodeCount), 
                                                       new Integer(myData_.linkCount), 
                                                       new Integer(myData_.fullShadowLinkCount)});  
      desc = UiUtil.convertMessageToHtml(desc);
      JOptionPane.showMessageDialog(topFrame, desc,
                                    rMan_.getPluginString("modelCounts.modelCountTitle"),
                                    JOptionPane.INFORMATION_MESSAGE);        
      return (true);
    }
  
    /***************************************************************************
    **
    ** Answer if command is enabled
    */
    
    public boolean isEnabled() {
      return (enabled_);    
    }
  }
  
  /***************************************************************************
  **
  ** For XML I/O
  */  
      
  public static class PlugInWorker extends AbstractFactoryClient {
   
    private NetStatsPlugIn plugin_;
   
    public PlugInWorker(PluginWhiteboard board, NetStatsPlugIn plugin) {
      super(board);
      plugin_ = plugin;
      String name = plugin.getClass().getName();
      myKeys_.add(name);
      installWorker(new NetStatsWorker(board), new StatsGlue());
    }
    
    protected Object localProcessElement(String elemName, Attributes attrs) throws IOException {
      Object retval = null;
      PluginWhiteboard board = (PluginWhiteboard)this.sharedWhiteboard_;
      if (myKeys_.contains(elemName)) {
        board.setCurrentPlugIn(plugin_);
        retval = board.getCurrentPlugIn();
      }
      return (retval);     
    }  
  }
  
  /***************************************************************************
  **
  ** For XML I/O
  */  
      
  public static class StatData implements BioFabricToolPlugInData {
    int nodeCount;
    int linkCount;
    int fullShadowLinkCount;
    
    public StatData(int nodeCount, int linkCount, int fullShadowLinkCount) {
      this.nodeCount = nodeCount;
      this.linkCount = linkCount;
      this.fullShadowLinkCount = fullShadowLinkCount;
    } 
  }
  
  /***************************************************************************
  **
  ** For XML I/O
  */
  
  public static class NetStatsWorker extends AbstractFactoryClient {
        
    public NetStatsWorker(PluginWhiteboard board) {
      super(board);
      myKeys_.add("netStats");
    }
    
    protected Object localProcessElement(String elemName, Attributes attrs) throws IOException {
      Object retval = null;
      PluginWhiteboard board = (PluginWhiteboard)this.sharedWhiteboard_;
      board.setCurrentPlugInData(buildFromXML(elemName, attrs));
      retval = board.getCurrentPlugInData();
      return (retval);
    }
    
    private StatData buildFromXML(String elemName, Attributes attrs) throws IOException {
      String nodeStr = AttributeExtractor.extractAttribute(elemName, attrs, "netStats", "nodes", true);
      String linkStr = AttributeExtractor.extractAttribute(elemName, attrs, "netStats", "links", true);
      String shadStr = AttributeExtractor.extractAttribute(elemName, attrs, "netStats", "fullShadowLinks", true);
  
      StatData retval;
      try {
        int nodeCount = Integer.valueOf(nodeStr).intValue();
        int linkCount = Integer.valueOf(linkStr).intValue();
        int fullCount = Integer.valueOf(shadStr).intValue();
        retval = new StatData(nodeCount, linkCount, fullCount);
      } catch (NumberFormatException nfex) {
        throw new IOException();
      }
      return (retval);
    }  
  }
  
  public static class StatsGlue implements GlueStick {    
    public Object glueKidToParent(Object kidObj, AbstractFactoryClient parentWorker, Object optionalArgs) throws IOException {
      PluginWhiteboard board = (PluginWhiteboard)optionalArgs;
      board.getCurrentPlugIn().attachXMLData(board.getCurrentPlugInData());
      return null;
    }
  }  
  
}
```
