## Supplementary material for "BioFabric Visualization of Network Alignments": BioFabric Source Code: PlugInManager.java

```
/*
**    Copyright (C) 2003-2018 Institute for Systems Biology 
**                            Seattle, Washington, USA. 
**
**    This library is free software; you can redistribute it and/or
**    modify it under the terms of the GNU Lesser General Public
**    License as published by the Free Software Foundation; either
**    version 2.1 of the License, or (at your option) any later version.
**
**    This library is distributed in the hope that it will be useful,
**    but WITHOUT ANY WARRANTY; without even the implied warranty of
**    MERCHANTABILITY or FITNESS FOR A PARTICULAR PURPOSE.  See the GNU
**    Lesser General Public License for more details.
**
**    You should have received a copy of the GNU Lesser General Public
**    License along with this library; if not, write to the Free Software
**    Foundation, Inc., 59 Temple Place, Suite 330, Boston, MA  02111-1307  USA
*/

package org.systemsbiology.biofabric.plugin;

import java.net.URL;
import java.util.List;
import java.util.ArrayList;
import java.util.TreeSet;
import java.util.Iterator;
import java.util.Enumeration;
import java.util.HashMap;
import java.util.Map;
import java.util.PropertyResourceBundle;
import java.util.ResourceBundle;
import java.io.File;
import java.io.FileFilter;
import java.io.IOException;
import java.util.jar.JarFile;
import java.util.jar.JarEntry;
import java.io.BufferedReader;
import java.io.InputStreamReader;
import java.io.PrintWriter;

import org.systemsbiology.biofabric.parser.SUParser;
import org.systemsbiology.biotapestry.biofabric.FabricCommands;
import org.systemsbiology.biofabric.api.io.Indenter;
import org.systemsbiology.biofabric.api.parser.AbstractFactoryClient;
import org.systemsbiology.biofabric.api.parser.GlueStick;
import org.systemsbiology.biofabric.api.parser.ParserClient;
import org.systemsbiology.biofabric.api.worker.AsynchExitRequestException;
import org.systemsbiology.biofabric.api.worker.BTProgressMonitor;
import org.systemsbiology.biofabric.app.ArgParser;
import org.systemsbiology.biofabric.io.FabricFactory;
import org.systemsbiology.biofabric.model.BioFabricNetwork;

import org.xml.sax.Attributes;

/****************************************************************************
**
** Plugin Manager.
*/

public class PlugInManager {
  
  ////////////////////////////////////////////////////////////////////////////
  //
  // PRIVATE MEMBERS
  //
  ////////////////////////////////////////////////////////////////////////////

  private ArrayList<BioFabricToolPlugIn> toolPlugIns_;
  private int maxCount_;
  private TreeSet<AbstractPlugInDirective> directives_;
  private HashMap<String, ResourceBundle> rBunds_;

  ////////////////////////////////////////////////////////////////////////////
  //
  // PUBLIC CONSTRUCTORS
  //
  ////////////////////////////////////////////////////////////////////////////

  /***************************************************************************
  **
  ** Basic Constructor
  */

  public PlugInManager() {
    toolPlugIns_ = new ArrayList<BioFabricToolPlugIn>();
    directives_ = new TreeSet<AbstractPlugInDirective>();
    maxCount_ = Integer.MIN_VALUE;
    rBunds_ = new HashMap<String, ResourceBundle>();
  }

  ////////////////////////////////////////////////////////////////////////////
  //
  // PUBLIC METHODS
  //
  ////////////////////////////////////////////////////////////////////////////

  /***************************************************************************
  **
  ** Get a resource bundle for a plugin
  */
  
  public ResourceBundle getResourceBundle(String name) {
  	//Loaded from a jar file:
    ResourceBundle rBund = rBunds_.get(name);
    if (rBund == null) {
    	// Part of classpath:
    	rBund = ResourceBundle.getBundle(name);
    }
    return (rBund);
  }

  /***************************************************************************
  **
  ** Install a new user-specified plugin directory
  */
  
  public void setDirectory(File directory) {
    FabricCommands.setPreference("PlugInDirectory", directory.getAbsolutePath());
    return;
  }
  
  /***************************************************************************
  **
  ** Install a new user-specified plugin directory
  */
  
  public String getDirectory() {
    return (FabricCommands.getPreference("PlugInDirectory"));
  }
  
  /***************************************************************************
  **
  ** Install a new network
  */
  
  public void installAPI(PlugInNetworkModelAPI api) {
    for (BioFabricToolPlugIn pi : toolPlugIns_) {
      pi.installAPI(api);
    }
    return;
  }
  
  /***************************************************************************
  **
  ** Install a new network
  */
  
  public void newNetworkInstalled(BioFabricNetwork bfn) {
    for (BioFabricToolPlugIn pi : toolPlugIns_) {
      pi.newNetworkInstalled(bfn);
    }
    return;
  }

  /***************************************************************************
  ** 
  ** Add a directive for nodes (can be either legacy or modern)
  */

  public void addDirective(AbstractPlugInDirective dir) {
    directives_.add(dir);
    int order = dir.getOrder();
    if (maxCount_ < order) {
      maxCount_ = order;
    }
    return;
  }   
  
  /***************************************************************************
  ** 
  ** Get an ordered list of plugin keys 
  */

  public List<String> getOrderedToolPlugInKeys() {
    ArrayList<String> retval = new ArrayList<String>();
    for (BioFabricToolPlugIn pi : toolPlugIns_) {
      retval.add(pi.getUniquePlugInTag());
    }
    return (retval);
  }  
  
  /***************************************************************************
  ** 
  ** Get BioFabricToolPlugIn for tag:
  */

  public BioFabricToolPlugIn getToolPlugIn(String key) {
    for (BioFabricToolPlugIn pi : toolPlugIns_) {
      if (pi.getUniquePlugInTag().equals(key)) {
        return (pi);
      }
    }
    throw new IllegalArgumentException();
  }  

  /***************************************************************************
  ** 
  ** Load plugins
  */

  public boolean loadPlugIns(Map<String, Object> args) {
    
    //
    // Load in the plugins specified in the resource file first:
    //

    if (!readPlugInListing()) {
      return (false);
    }
    
    //
    // Now load from jar file, if specified in command line argument:
    //
        
    String plugDirStr = (String)args.get(ArgParser.PLUG_IN_DIR);
    if (plugDirStr != null) {
      File plugDirectory = new File(plugDirStr);
      if (!plugDirectory.exists() || !plugDirectory.isDirectory() || !plugDirectory.canRead()) {
        return (false);
      }
      if (!readJarFiles(plugDirectory, maxCount_ + 1)) {
         return (false);
      }
    }
    
    //
    // Now load from jar files, if located in directory specified in preferences. We silently fail if nothing
    // is found:
    //
    
    String plugDirPref = FabricCommands.getPreference("PlugInDirectory");
    if (plugDirPref != null) {
      File plugDirectory = new File(plugDirPref);
      if (plugDirectory.exists() && plugDirectory.isDirectory() && plugDirectory.canRead()) {
      	readJarFiles(plugDirectory, maxCount_ + 1);
      }
    }

    Iterator<AbstractPlugInDirective> drit = directives_.iterator();
    while (drit.hasNext()) {
      AbstractPlugInDirective pid = drit.next();
      BioFabricToolPlugIn pi = pid.buildPlugIn(this);
      if (pi != null) {
        toolPlugIns_.add(pi);
      }
    }
     
    return (true);
  }

  /***************************************************************************
  **
  ** Dump plugin data using XML
  */
  
  public void writeXML(PrintWriter out, Indenter ind, BTProgressMonitor monitor, boolean forCache) throws AsynchExitRequestException {    
    //
    // Let the plugins write to XML
    //
    ind.indent();
    out.println("<plugInDataSets>");
    List<String> keyList = getOrderedToolPlugInKeys();
    for (String key : keyList) {
      BioFabricToolPlugIn plugin = getToolPlugIn(key);
      plugin.writeXML(out, ind);
    }
    ind.indent();
    out.println("</plugInDataSets>");
    return;
  }
  
  ////////////////////////////////////////////////////////////////////////////
  //
  // PRIVATE METHODS
  //
  ////////////////////////////////////////////////////////////////////////////

  /***************************************************************************
  **
  ** Read jars
  */

  private boolean readJarFiles(File plugInDir, int currMax) {
    try {
    	int propLen = ".properties".length();
      ExtensionFilter filter = new ExtensionFilter(".jar");
      if (plugInDir.isDirectory()) {
        File[] files = plugInDir.listFiles(filter);
        for (int i = 0; i < files.length; i++) {
          JarFile jar = new JarFile(files[i]);
        
          //
          // ToolPlugin:
          //        
          
          List<String> sl = getServiceList(jar, "org.systemsbiology.biofabric.plugin.BioFabricToolPlugIn");
          int numSvc = sl.size();
          for (int j = 0; j < numSvc; j++) {
            String plugin = sl.get(j);
            ToolPlugInDirective pid = new ToolPlugInDirective(plugin, Integer.toString(currMax++), files[i]);
            addDirective(pid);
          }
          
          //
          // Need to suck in any properties files for the PluginResourceManager to manage:
          //
          
          Enumeration<JarEntry> entries = jar.entries();
          while (entries.hasMoreElements()) {
          	JarEntry jarEntry = entries.nextElement();
          	String jarName = jarEntry.getName();
          	int jnlen = jarName.length();
          	if (jnlen <= propLen) {
          		continue;
          	}
          	String suffix = jarEntry.getName().substring(jnlen - propLen, jnlen);   	
          	if (suffix.equals(".properties")) {
          		String prefix = jarEntry.getName().substring(0, jnlen - propLen).replaceAll("/", ".");
          	  BufferedReader in = new BufferedReader(new InputStreamReader(jar.getInputStream(jarEntry)));
              rBunds_.put(prefix, new PropertyResourceBundle(in));
          	}
          }
        }
      }
    } catch (IOException ioex) {
      return (false);
    }
    return (true);
  }
  
  /***************************************************************************
  **
  ** Get the list of services
  */

  private List<String> getServiceList(JarFile jar, String svcInterface) throws IOException {
    svcInterface = "META-INF/services/" + svcInterface;
    Enumeration<JarEntry> entries = jar.entries();
    while (entries.hasMoreElements()) {
      JarEntry entry = entries.nextElement();
      String entryName = entry.getName();
      if (entryName.equals(svcInterface)) {
        return (readClassList(jar, entry));
      }
    }
    return (new ArrayList<String>());
  }

  /***************************************************************************
  **
  ** Get the list of classes from an entry
  */

  private List<String> readClassList(JarFile jar, JarEntry entry) throws IOException {
    ArrayList<String> retval = new ArrayList<String>();
    BufferedReader in = new BufferedReader(new InputStreamReader(jar.getInputStream(entry)));
    String newLine;
    while ((newLine = in.readLine()) != null) {
      newLine = newLine.trim();
      if (newLine.equals("") || newLine.startsWith("#")) {
        continue;
      }
      retval.add(newLine);
    }
    in.close();
    return (retval);
  }
  
  /***************************************************************************
  **
  ** Read the plugin listing
  */

  private boolean readPlugInListing() {
    URL url = getClass().getResource("/org/systemsbiology/biofabric/plugin/plugInListing.xml");
    if (url == null) {
      System.err.println("No plugIn directives file found");
      return (false);
    }
    ArrayList<ParserClient> alist = new ArrayList<ParserClient>();
    alist.add(new PlugInDirectiveFactory(this));
    SUParser sup = new SUParser(alist);
    try {
      sup.parse(url);
    } catch (IOException ioe) {
      System.err.println("Could not read plugIn directives file");
      return (false);              
    }
    return (true);
    
  }
  
  /***************************************************************************
  **
  ** For XML I/O
  */  
      
  public static class DirectiveWorker extends AbstractFactoryClient {
    
    public DirectiveWorker(PlugInDirectiveFactory.FactoryWhiteboard whiteboard) {
      super(whiteboard);
      myKeys_.add("BioFabricPlugIns");
      installWorker(new ToolPlugInDirective.DirectiveWorker(whiteboard), new MyDirectiveGlue());
    }
    
    protected Object localProcessElement(String elemName, Attributes attrs) throws IOException {
      // Nothing to do at this level
      return (null);     
    }
  }
  
  public static class MyDirectiveGlue implements GlueStick {
    public Object glueKidToParent(Object kidObj, AbstractFactoryClient parentWorker, 
                                  Object optionalArgs) throws IOException {
      PlugInDirectiveFactory.FactoryWhiteboard board = (PlugInDirectiveFactory.FactoryWhiteboard)optionalArgs;
      board.mgr.addDirective(board.dir);
      return (null);
    }
  }  
 
  /***************************************************************************
  **
  ** For XML I/O
  */  
      
  public static class PlugInWorker extends AbstractFactoryClient {
      
    public PlugInWorker(FabricFactory.FactoryWhiteboard board, PlugInManager pmg) {
      super(board);
      myKeys_.add("plugInDataSets");
      List<String> keyList = pmg.getOrderedToolPlugInKeys();
      for (String key : keyList) {
        BioFabricToolPlugIn plugin = pmg.getToolPlugIn(key);
        AbstractFactoryClient afc = plugin.getXMLWorker(board);
        installWorker(afc, null);
      }
    }
    
    protected Object localProcessElement(String elemName, Attributes attrs) throws IOException {
      Object retval = null;
      if (myKeys_.contains(elemName)) {
        // Nothing to do here
      }
      return (retval);     
    }  
  }

  ////////////////////////////////////////////////////////////////////////////
  //
  // INNER CLASSES
  //
  ////////////////////////////////////////////////////////////////////////////
  
  /***************************************************************************
  **
  ** File filter
  */ 
    
  class ExtensionFilter implements FileFilter {
    
    private String suffix_;
    
    ExtensionFilter(String suffix) {
      suffix_ = suffix;
    }
    
    private boolean hasSuffix(String fileName, String suffix) {
      int fnl = fileName.length();
      int sufl = suffix.length();   
      return ((fnl > sufl) && 
              (fileName.toLowerCase().lastIndexOf(suffix.toLowerCase()) == fnl - sufl));
    }  
  
    public boolean accept(File f) {
      String fileName = f.getName();
      return (hasSuffix(fileName, suffix_));
    }
  }
  
}
```
