## Supplementary material for "BioFabric Visualization of Network Alignments": BioFabric Source Code: PlugInNetworkModelAPI.java

```
/*
**    Copyright (C) 2003-2019 Institute for Systems Biology 
**                            Seattle, Washington, USA. 
**
**    This library is free software; you can redistribute it and/or
**    modify it under the terms of the GNU Lesser General Public
**    License as published by the Free Software Foundation; either
**    version 2.1 of the License, or (at your option) any later version.
**
**    This library is distributed in the hope that it will be useful,
**    but WITHOUT ANY WARRANTY; without even the implied warranty of
**    MERCHANTABILITY or FITNESS FOR A PARTICULAR PURPOSE.  See the GNU
**    Lesser General Public License for more details.
**
**    You should have received a copy of the GNU Lesser General Public
**    License along with this library; if not, write to the Free Software
**    Foundation, Inc., 59 Temple Place, Suite 330, Boston, MA  02111-1307  USA
*/

package org.systemsbiology.biofabric.plugin;

import javax.swing.JFrame;

import org.systemsbiology.biofabric.api.io.FileLoadFlows;
import org.systemsbiology.biofabric.api.worker.BackgroundWorkerControlManager;

/****************************************************************************
**
** Interface for information plugins can get from BioFabric
*/

public interface PlugInNetworkModelAPI {
  
  ////////////////////////////////////////////////////////////////////////////
  //
  // PUBLIC METHODS
  //
  ////////////////////////////////////////////////////////////////////////////
  
  /***************************************************************************
  **
  ** Get the link count, with and without shadows
  */
  
  public int getLinkCount(boolean forShadow);
  
  /***************************************************************************
  ** 
  ** Get node count
  */

  public int getNodeCount();
  
  /***************************************************************************
  ** 
  ** Get file loading utilities
  */

  public FileLoadFlows getFileUtilities();
 
  /***************************************************************************
  ** 
  ** Get top window
  */

  public JFrame getTopWindow();
  
  /***************************************************************************
  ** 
  ** Get the BackgroundWorkerControlManager
  */

  public BackgroundWorkerControlManager getBWCtrlMgr();

  /***************************************************************************
  **
  ** Stash plugin data for extraction
  */
  
  public void stashPluginData(String keyword, BioFabricToolPlugInData data);

  /***************************************************************************
  **
  ** Pull plugin data for extraction
  */
  
  public BioFabricToolPlugInData providePluginData(String keyword);
 
  /***************************************************************************
  **
  ** Get if we are currently set to display shadow links
  */

  public boolean getDisplayShadows();
  
}
```
