## Supplementary material for "BioFabric Visualization of Network Alignments": BioFabric Source Code: PluginSupportFactory.java

```
/*
**    Copyright (C) 2003-2018 Institute for Systems Biology 
**                            Seattle, Washington, USA. 
**
**    This library is free software; you can redistribute it and/or
**    modify it under the terms of the GNU Lesser General Public
**    License as published by the Free Software Foundation; either
**    version 2.1 of the License, or (at your option) any later version.
**
**    This library is distributed in the hope that it will be useful,
**    but WITHOUT ANY WARRANTY; without even the implied warranty of
**    MERCHANTABILITY or FITNESS FOR A PARTICULAR PURPOSE.  See the GNU
**    Lesser General Public License for more details.
**
**    You should have received a copy of the GNU Lesser General Public
**    License along with this library; if not, write to the Free Software
**    Foundation, Inc., 59 Temple Place, Suite 330, Boston, MA  02111-1307  USA
*/

package org.systemsbiology.biofabric.plugin;

import java.util.Map;
import java.util.Set;

import javax.swing.JFrame;

import org.systemsbiology.biofabric.api.io.BuildData;
import org.systemsbiology.biofabric.api.io.BuildExtractor;
import org.systemsbiology.biofabric.api.model.Annot;
import org.systemsbiology.biofabric.api.model.AnnotationSet;
import org.systemsbiology.biofabric.api.model.NetLink;
import org.systemsbiology.biofabric.api.model.NetNode;
import org.systemsbiology.biofabric.api.util.NID;
import org.systemsbiology.biofabric.api.util.PluginResourceManager;
import org.systemsbiology.biofabric.api.util.PreferenceStorage;
import org.systemsbiology.biofabric.api.util.UniqueLabeller;
import org.systemsbiology.biofabric.api.worker.BFWorker;
import org.systemsbiology.biofabric.api.worker.BackgroundWorkerControlManager;
import org.systemsbiology.biofabric.api.worker.BackgroundWorkerOwner;
import org.systemsbiology.biofabric.model.AnnotationSetImpl;
import org.systemsbiology.biofabric.model.FabricLink;
import org.systemsbiology.biofabric.model.FabricNode;
import org.systemsbiology.biofabric.io.BuildDataImpl;
import org.systemsbiology.biofabric.io.BuildExtractorImpl;
import org.systemsbiology.biofabric.ui.FabricColorGenerator;
import org.systemsbiology.biofabric.util.ResourceManager;
import org.systemsbiology.biofabric.worker.BackgroundWorker;
import org.systemsbiology.biofabric.worker.BackgroundWorkerClient;
import org.systemsbiology.biofabric.worker.WorkerClientBundle;

/****************************************************************************
**
** Factory for returning API Implementations
*/

public class PluginSupportFactory {
  
  ////////////////////////////////////////////////////////////////////////////
  //
  // PUBLIC METHODS
  //
  ////////////////////////////////////////////////////////////////////////////

  /***************************************************************************
  **
  ** Get a Network Node
  */
  
  public static NetNode buildNode(NID id, String name) {
  	return (new FabricNode(id, name));
  }
	
	/***************************************************************************
  **
  ** Get a Network Link
  */
  
  public static NetLink buildLink(NetNode srcID, NetNode trgID, String relation, boolean isShadow, Boolean directed) {
  	return (new FabricLink(srcID, trgID, relation, isShadow, directed));
  }
  
  /***************************************************************************
  **
  ** Get a Network Link
  */
  
  public static NetLink buildLink(NetNode srcID, NetNode trgID, String relation, boolean isShadow) {
  	return (new FabricLink(srcID, trgID, relation, isShadow));
  }
  
	/***************************************************************************
  **
  ** Build an empty Annotation Set
  */
  
  public static AnnotationSet buildAnnotationSet() {
  	return (new AnnotationSetImpl());
  }

	/***************************************************************************
  **
  ** Get an Annotation
  */
  
  public static Annot buildAnnotation(String tag, int startPos, int endPos, int layer, String colorName) {
  	return (new AnnotationSetImpl.AnnotImpl(tag, startPos, endPos, layer, colorName));
  }
	
  /***************************************************************************
  **
  ** Get a Build Extractor
  */
  
  public static PreferenceStorage getPreferenceStorage() {
  	return (new PreferenceStorage());
  }	

  /***************************************************************************
  **
  ** Get a Build Extractor
  */
  
  public static BuildExtractor getBuildExtractor() {
  	return (new BuildExtractorImpl());
  }
  
  /***************************************************************************
  **
  ** Get a BFWorker
  */
  
  public static BFWorker getBFWorker(BackgroundWorkerOwner owner,
                                     JFrame topWindow, BackgroundWorkerControlManager suw, String waitTitle, 
                                     String waitMsg, boolean allowCancels, PluginResourceManager rMan) {
  	
  	BackgroundWorker bw = new BackgroundWorker();
    BackgroundWorkerClient bwc = 
    		new BackgroundWorkerClient(owner, bw, topWindow, suw, waitTitle, waitMsg, allowCancels, rMan);
  	return (new WorkerClientBundle(bw, bwc));
  }
  
  /***************************************************************************
  **
  ** Get a Plugin Resource Manager
  */
  
  public static PluginResourceManager getResourceManager(String pluginName, PlugInManager pMan) {
  	return (new ResourceManager.ForPlugins(pluginName, pMan));
  } 
  
  /***************************************************************************
  **
  ** Get a BuildData
  */
  
  public static BuildData getBuildDataForPlugin(UniqueLabeller idGen,
										  		                      Set<NetLink> allLinks, Set<NetNode> loneNodeIDs, 
										  		                      Map<NetNode, String> clustAssign, 
										  		                      FabricColorGenerator colGen) {
  	 
    return (new BuildDataImpl(idGen, allLinks, loneNodeIDs, clustAssign, colGen, BuildDataImpl.BuildMode.BUILD_FROM_PLUGIN));
  }
}
```
