## Supplementary material for "BioFabric Visualization of Network Alignments": BioFabric Source Code: ToolPlugInDirective.java

```
/*
**    Copyright (C) 2003-2018 Institute for Systems Biology 
**                            Seattle, Washington, USA. 
**
**    This library is free software; you can redistribute it and/or
**    modify it under the terms of the GNU Lesser General Public
**    License as published by the Free Software Foundation; either
**    version 2.1 of the License, or (at your option) any later version.
**
**    This library is distributed in the hope that it will be useful,
**    but WITHOUT ANY WARRANTY; without even the implied warranty of
**    MERCHANTABILITY or FITNESS FOR A PARTICULAR PURPOSE.  See the GNU
**    Lesser General Public License for more details.
**
**    You should have received a copy of the GNU Lesser General Public
**    License along with this library; if not, write to the Free Software
**    Foundation, Inc., 59 Temple Place, Suite 330, Boston, MA  02111-1307  USA
*/

package org.systemsbiology.biofabric.plugin;

import java.io.IOException;
import java.io.File;
import java.util.Set;
import java.util.HashSet;

import org.systemsbiology.biofabric.api.parser.AbstractFactoryClient;
import org.xml.sax.Attributes;

/***************************************************************************
**
** Used to specify tool plugins
*/
  
public class ToolPlugInDirective extends AbstractPlugInDirective {
 
  ////////////////////////////////////////////////////////////////////////////
  //
  // PUBLIC CONSTANTS
  //
  ////////////////////////////////////////////////////////////////////////////  

  ////////////////////////////////////////////////////////////////////////////
  //
  // PRIVATE CONSTANTS
  //
  ////////////////////////////////////////////////////////////////////////////
  
  private static final String PLUGIN_TAG_ = "toolPlugIn";
  
  ////////////////////////////////////////////////////////////////////////////
  //
  // CONSTRUCTORS
  //
  ////////////////////////////////////////////////////////////////////////////    

  /***************************************************************************
  **
  ** Constructor for internal use
  **
  */
  
  public ToolPlugInDirective(String className, String order, File jarFile) {
    super(className, order, jarFile);
  }  

  /***************************************************************************
  **
  ** Constructor for XML input
  **
  */
  
  private ToolPlugInDirective() {
    super();
  }
 
  ////////////////////////////////////////////////////////////////////////////
  //
  // PUBLIC METHODS
  //
  ////////////////////////////////////////////////////////////////////////////  
  
  /***************************************************************************
  **
  ** Standard toString:
  **
  */
  
  @Override
  public String toString() { 
    return ("ToolPlugInDirective: " + className_ + " " + order_);
  }
  
  ////////////////////////////////////////////////////////////////////////////
  //
  // PUBLIC CLASS METHODS
  //
  ////////////////////////////////////////////////////////////////////////////
  
  /***************************************************************************
  **
  ** Return the element keywords that we are interested in
  **
  */
  
  public static Set<String> keywordsOfInterest() {
    HashSet<String> retval = new HashSet<String>();
    retval.add(PLUGIN_TAG_);
    return (retval);
  }
  
  /***************************************************************************
  **
  ** For XML I/O
  */  
      
  public static class DirectiveWorker extends AbstractFactoryClient {
    
    public DirectiveWorker(PlugInDirectiveFactory.FactoryWhiteboard whiteboard) {
      super(whiteboard);
      myKeys_.add(PLUGIN_TAG_);
    }
    
    protected Object localProcessElement(String elemName, Attributes attrs) throws IOException {
      Object retval = null;
      if (elemName.equals(PLUGIN_TAG_)) {
        PlugInDirectiveFactory.FactoryWhiteboard board = (PlugInDirectiveFactory.FactoryWhiteboard)this.sharedWhiteboard_;
        board.dir = new ToolPlugInDirective();
        board.dir.stockCoreFromXML(elemName, attrs);
        retval = board.dir;
      }
      return (retval);     
    }
  }
}
```
