## Supplementary material for "BioFabric Visualization of Network Alignments": BioFabric Source Code: BasicZoomTargetSupport.java

```
/*
**    Copyright (C) 2003-2017 Institute for Systems Biology 
**                            Seattle, Washington, USA. 
**
**    This library is free software; you can redistribute it and/or
**    modify it under the terms of the GNU Lesser General Public
**    License as published by the Free Software Foundation; either
**    version 2.1 of the License, or (at your option) any later version.
**
**    This library is distributed in the hope that it will be useful,
**    but WITHOUT ANY WARRANTY; without even the implied warranty of
**    MERCHANTABILITY or FITNESS FOR A PARTICULAR PURPOSE.  See the GNU
**    Lesser General Public License for more details.
**
**    You should have received a copy of the GNU Lesser General Public
**    License along with this library; if not, write to the Free Software
**    Foundation, Inc., 59 Temple Place, Suite 330, Boston, MA  02111-1307  USA
*/

package org.systemsbiology.biofabric.ui;

import java.awt.Point;
import java.awt.Dimension;
import java.awt.Rectangle;
import java.awt.geom.Point2D;
import java.awt.geom.AffineTransform;
import java.awt.geom.NoninvertibleTransformException;
import java.awt.geom.Rectangle2D;
import javax.swing.JPanel;

import org.systemsbiology.biofabric.cmd.ZoomTarget;
import org.systemsbiology.biofabric.util.UiUtil;

/***************************************************************************
** 
** Supports operations used in zooming
*/

public class BasicZoomTargetSupport implements ZoomTarget {
  
  ////////////////////////////////////////////////////////////////////////////
  //
  // PRIVATE MEMBERS
  //
  ////////////////////////////////////////////////////////////////////////////

  protected ZoomPresentation myGenomePre_;
  protected double zoom_;
  protected AffineTransform transform_;
  protected JPanel paintTarget_;
  protected Rectangle worldRect_;
  protected Rectangle2D clipRect_;
  
  ////////////////////////////////////////////////////////////////////////////
  //
  // PRIVATE CONSTANTS
  //
  //////////////////////////////////////////////////////////////////////////// 
  
  ////////////////////////////////////////////////////////////////////////////
  //
  // PUBLIC CONSTRUCTOR
  //
  ////////////////////////////////////////////////////////////////////////////  

  /***************************************************************************
  **
  ** Constructor
  */
  
  public BasicZoomTargetSupport(ZoomPresentation genomePre, JPanel paintTarget) { 
  	zoom_ = 1.0;
    transform_ = new AffineTransform();
    myGenomePre_ = genomePre;
    paintTarget_ = paintTarget;
    worldRect_ = new Rectangle(0, 0, 100, 100);
    clipRect_ = new Rectangle2D.Double(0.0, 0.0, 1.0, 1.0);
  }

  ////////////////////////////////////////////////////////////////////////////
  //
  // PUBLIC METHODS
  //
  ////////////////////////////////////////////////////////////////////////////

  /***************************************************************************
  ** 
  ** Set the world rect (formerly the "workspace")
  */

  public void setWorldRect(Rectangle worldRect) { 
    worldRect_ = (Rectangle)worldRect.clone();
    return;
  }

  /***************************************************************************
  ** 
  ** Let us know what the current clip rect is
  */

  public void setCurrClipRect(Rectangle2D clipRect) { 
    clipRect_.setRect(clipRect);
    return;
  }

  /***************************************************************************
  ** 
  ** Bump to next selection
  */

  public void incrementToNextSelection() { 
    myGenomePre_.bumpNextSelection();
    return;
  }
  
  /***************************************************************************
  ** 
  ** Bump to previous selection
  */

  public void decrementToPreviousSelection() { 
    myGenomePre_.bumpPreviousSelection();
    return;
  } 

  /***************************************************************************
  ** 
  ** repaint
  */
  
  public void repaint() {
    paintTarget_.repaint();
    return;
  }
 
  /***************************************************************************
  **
  ** Get the transform
  */
  
  public AffineTransform getTransform() {
    return (transform_);
  }  

  /***************************************************************************
  **
  ** Get the preferred size X
  */
  
  public Dimension getPreferredSize() {
    //
    // We always display a fixed-size working space in world coords.  This 
    // gives us the necessary component size we need to accommodate that world
    // at the given zoom level.
    //
    int width = (int)(worldRect_.width * zoom_); 
    int height = (int)(worldRect_.height * zoom_);
    return (new Dimension(width, height));
  }  
 
  /***************************************************************************
  **
  ** Get workspace bounds X
  */
  
  public Rectangle getWorldRect() {
    return ((Rectangle)worldRect_.clone());
  }  
  
  /***************************************************************************
  **
  ** Gets center point in viewport coordinates X
  */
  
  public Point getCenterPoint() {
    Point2D center = UiUtil.getRectCenter(worldRect_);
    Point2D centerPoint = (Point2D)center.clone();
    transform_.transform(centerPoint, centerPoint);
    int x = (int)Math.round(centerPoint.getX());
    int y = (int)Math.round(centerPoint.getY());
    Point pt = new Point(x, y);
    return (pt);
  }

  /***************************************************************************
  **
  ** Gets center point in world coordinates X
  */
  
  public Point2D getRawCenterPoint() {
    Point2D center = UiUtil.getRectCenter(worldRect_);
    return ((Point2D)center.clone());
  } 
  
  /***************************************************************************
  **
  ** Get the basic (unzoomed) image size: X Override for complex behavior
  */
  
  public Dimension getBasicSize() {
    Rectangle origRect = myGenomePre_.getRequiredSize();
    return (new Dimension(origRect.width, origRect.height));
  }
  
  /***************************************************************************
  **
  ** Get the basic model bounds X.  Override for complex behavior
  */
  
  public Rectangle getCurrentBasicBounds() {
    return (myGenomePre_.getRequiredSize());  
  }  
 
  /***************************************************************************
  **
  ** Get the bounds of the selected parts.  May be null. X
  */
  
  public Rectangle getSelectedBounds() {  
    return (myGenomePre_.getSelectionSize());  
  }
 
  /***************************************************************************
  **
  ** Get the bounds of the selected parts.  May be null. X
  */
  
  public Rectangle getCurrentSelectedBounds() {  
    return (myGenomePre_.getCurrentSelectionSize());  
  }
  
  /***************************************************************************
  **
  ** Gets given point in viewport coordinates X
  */
  
  public Point pointToViewport(Point worldPoint) {
  	Point2D newPoint = new Point2D.Double();
  	Point pt = new Point();
    return (pointToViewport(worldPoint, transform_, newPoint, pt));
  } 
  
  /***************************************************************************
  **
  ** Gets given point in viewport coordinates X
  */
  
  public static Point pointToViewport(Point worldPoint, AffineTransform trans, Point2D temp, Point res) {
    temp.setLocation(worldPoint.getX(), worldPoint.getY());
    trans.transform(temp, temp);
    int x = (int)Math.round(temp.getX());
    int y = (int)Math.round(temp.getY());
    res.setLocation(x, y);
    return (res);
  } 
  
  /***************************************************************************
  **
  ** Gets given viewpoint in world coordinates: Computes inverse; use sparingly! X
  */
  
  public Point2D viewToWorld(Point viewPoint) {
    Point2D ptSrc = new Point2D.Double(viewPoint.getX(), viewPoint.getY());
    try {
      Point2D ptDest = new Point2D.Double(0.0, 0.0);
      transform_.inverseTransform(ptSrc, ptDest);
      return (ptDest);
    } catch (NoninvertibleTransformException ex) {
      System.err.println("cannot invert: " + transform_ + " " + zoom_);
      throw new IllegalStateException();
    }
  }  
  
  /***************************************************************************
  **
  ** Set the zoom X
  */
 
  public void setZoomFactor(double zoom, Dimension viewportDim) {   
    //
    // If the viewport is larger than the preferred size, we center using that:
    //
    zoom_ = zoom;
    transform_ = getTransformForWideZoomFactor(zoom, viewportDim);
    myGenomePre_.setPresentationZoomFactor(zoom); 
    return;
  }
  
  /***************************************************************************
  **
  ** Set the zoom X
  */
 
  private AffineTransform getTransformForWideZoomFactor(double zoom, Dimension viewportDim) {   
    //
    // If the viewport is larger than the preferred size, we center using that:
    //
  	
  	UiUtil.fixMePrintout("READ ME");
  	//
  	// The viewport when the zoom button is clicked is checked for size. But if scrollbar is present, this
  	// messes stuff up, as the wide view will have no scrollbars.
  	//
  	
    AffineTransform transform = new AffineTransform();
   
    int rectWidth = (int)(worldRect_.getWidth() * zoom); 
    int rectHeight = (int)(worldRect_.getHeight() * zoom);
    
    int useWidth = (rectWidth < viewportDim.width) ? viewportDim.width : rectWidth;
    int useHeight = (rectHeight < viewportDim.height) ? viewportDim.height : rectHeight;
    Point2D center = UiUtil.getRectCenter(worldRect_); 
    transform = new AffineTransform();    
    transform.translate((useWidth / 2.0), (useHeight / 2.0));
    transform.scale(zoom, zoom);
    transform.translate(-center.getX(), -center.getY());    
    return (transform);
  }
  
  /***************************************************************************
  **
  ** Set the zoom X
  */
 
  public void adjustWideZoomForSize(Dimension viewportDim) {   
    //
    // If the viewport is larger than the preferred size, we center using that:
    //   
   
    int rectWidth = (int)(worldRect_.getWidth() * zoom_); 
    int rectHeight = (int)(worldRect_.getHeight() * zoom_);
    
    int useWidth = (rectWidth < viewportDim.width) ? viewportDim.width : rectWidth;
    int useHeight = (rectHeight < viewportDim.height) ? viewportDim.height : rectHeight;
    Point2D center = UiUtil.getRectCenter(worldRect_);
    
    transform_ = new AffineTransform();
    transform_.translate((useWidth / 2.0), (useHeight / 2.0));
    transform_.scale(zoom_, zoom_);
    transform_.translate(-center.getX(), -center.getY());    
    return;
  }  
  
  /***************************************************************************
  **
  ** Get the zoom
  */
  
  public double getZoomFactor() {
    return (zoom_);
  } 
  
  /***************************************************************************
  **
  ** Transforms screen point to model coordinates
  */  
  
  public void transformPoint(Point pt) { 
    try {
      transform_.inverseTransform(pt, pt);
    } catch (NoninvertibleTransformException nitex) {
      throw new IllegalStateException();
    }
    return;
  }  
 
  /***************************************************************************
  **
  ** Transforms screen point to model coordinates
  */  
  
  public void transformClick(int x, int y, Point pt) {
    Point2D clickPoint = new Point2D.Double(x, y);      
    try {
      transform_.inverseTransform(clickPoint, clickPoint);
    } catch (NoninvertibleTransformException nitex) {
      throw new IllegalStateException();
    }
    pt.x = (int)Math.round(clickPoint.getX());
    pt.y = (int)Math.round(clickPoint.getY());     
  }
  
  /***************************************************************************
  **
  ** Return the current pixel diameter
  */  
  
  public double currentPixelDiameter() {
    Point2D clickPoint0 = new Point2D.Double(0.0, 0.0); 
    Point2D clickPoint1 = new Point2D.Double(1.0, 0.0);
    try {
      transform_.inverseTransform(clickPoint0, clickPoint0);
      transform_.inverseTransform(clickPoint1, clickPoint1);
    } catch (NoninvertibleTransformException nitex) {
      throw new IllegalStateException();
    }
    return (Math.abs(clickPoint1.getX() - clickPoint0.getX()));
  }
  
  /***************************************************************************
  **
  ** Return the current clip rect (in world coordinates)
  */  
  
  public Rectangle2D getCurrentWorldClipRect() {
    Point2D clipUL = new Point2D.Double(clipRect_.getMinX(),  clipRect_.getMinY()); 
    Point2D clipLR = new Point2D.Double(clipRect_.getMaxX(),  clipRect_.getMaxY());
    try {
      transform_.inverseTransform(clipUL, clipUL);
      transform_.inverseTransform(clipLR, clipLR);
    } catch (NoninvertibleTransformException nitex) {
      throw new IllegalStateException();
    }
    return (new Rectangle2D.Double(clipUL.getX(), clipUL.getY(), clipLR.getX() - clipUL.getX(), clipLR.getY() - clipUL.getY()));
  }
  
  /***************************************************************************
  **
  ** Transforms screen point to model coordinates
  */  
  
  public void transformClick(double x, double y, Point pt) {
    Point2D clickPoint = new Point2D.Double(x, y);      
    try {
      transform_.inverseTransform(clickPoint, clickPoint);
    } catch (NoninvertibleTransformException nitex) {
      throw new IllegalStateException();
    }
    pt.x = (int)Math.round(clickPoint.getX());
    pt.y = (int)Math.round(clickPoint.getY());     
  }  
}
```
