## Supplementary material for "BioFabric Visualization of Network Alignments": BioFabric Source Code: BrowserLauncher.java

```
/*
**    Copyright (C) 2003-2018 Institute for Systems Biology 
**                            Seattle, Washington, USA. 
**
**    This library is free software; you can redistribute it and/or
**    modify it under the terms of the GNU Lesser General Public
**    License as published by the Free Software Foundation; either
**    version 2.1 of the License, or (at your option) any later version.
**
**    This library is distributed in the hope that it will be useful,
**    but WITHOUT ANY WARRANTY; without even the implied warranty of
**    MERCHANTABILITY or FITNESS FOR A PARTICULAR PURPOSE.  See the GNU
**    Lesser General Public License for more details.
**
**    You should have received a copy of the GNU Lesser General Public
**    License along with this library; if not, write to the Free Software
**    Foundation, Inc., 59 Temple Place, Suite 330, Boston, MA  02111-1307  USA
**
** NOTE:  This class contains Public Domain Software from Dem Pilafian: see
** comments below! 
*/

package org.systemsbiology.biofabric.ui;

import java.lang.reflect.Method;
import java.net.URI;

/****************************************************************************
**
** Class to launch a browser
*/

public class BrowserLauncher {
  
  ////////////////////////////////////////////////////////////////////////////
  //
  // PRIVATE MEMBERS
  //
  ////////////////////////////////////////////////////////////////////////////

  ////////////////////////////////////////////////////////////////////////////
  //
  // PUBLIC CONSTRUCTORS
  //
  ////////////////////////////////////////////////////////////////////////////
  
  public BrowserLauncher() {
  }
    
  ////////////////////////////////////////////////////////////////////////////
  //
  // PUBLIC METHODS
  //
  ////////////////////////////////////////////////////////////////////////////
 
  /***************************************************************************
  **
  ** Adapted from:
  **
  ** From http://www.centerkey.com/java/browser/
  ** Bare Bones Browser Launch 
  ** Version 3.1 (June 6, 2010) 
  ** By Dem Pilafian 
  ** Supports: 
  **   Mac OS X, GNU/Linux, Unix, Windows XP/Vista/7 
  ** Public Domain Software -- Free to Use as You Like
  */ 

  public void openBBURL(String url) { 
    try { 
      //attempt to use Desktop library from JDK 1.6+ 
      //code mimicks: java.awt.Desktop.getDesktop().browse() 
      URI myUri = new URI(url);
      Object[] uriArr = new Object[] {myUri};
           
      Class<?> d = Class.forName("java.awt.Desktop");      
      Method gdm = d.getDeclaredMethod("getDesktop", (Class[])null);      
      Object deskObj = gdm.invoke(null, (Object[])null);
      
      Class<?>[] uric = new Class[] {java.net.URI.class};
      Method meth = d.getDeclaredMethod("browse", uric);
      meth.invoke(deskObj, uriArr);
            
    } catch (Exception ex) { //library not available or failed 
      String osName = System.getProperty("os.name"); 
      try { 
        if (osName.startsWith("Mac OS")) { 
          Class<?> fm = Class.forName("com.apple.eio.FileManager");
          Class<?>[] sar = new Class[] {String.class};
          Method orl = fm.getDeclaredMethod("openURL", sar);
          orl.invoke(null, new Object[] {url}); 
        } else if (osName.startsWith("Windows")) {
          Runtime.getRuntime().exec("rundll32 url.dll,FileProtocolHandler " + url); 
        } else { //assume Unix or Linux 
          String used = null;
          String[] browsers = new String[] {"google-chrome", "firefox", "opera", "epiphany", "konqueror", "conkeror", "midori", "kazehakase", "mozilla", "netscape"}; 
          for (int i = 0; i < browsers.length ; i++) {
            String[] execArgs = new String[] {"which", browsers[i]};
            boolean found = (Runtime.getRuntime().exec(execArgs).getInputStream().read() != -1);
            if (found) {
              used = browsers[i];
              String[] execArgs2 = new String[] {used, url};
              Runtime.getRuntime().exec(execArgs2); 
              break;
            } 
          }
          if (used == null) {
            System.err.println("Browser launch failed");
          }
        }
      } catch (Exception e) { 
        System.err.println("Browser launch failed");
      }
    }
    return;
  } 
}
```
