## Supplementary material for "BioFabric Visualization of Network Alignments": BioFabric Source Code: CursorManager.java

```
/*
**    Copyright (C) 2003-2011 Institute for Systems Biology 
**                            Seattle, Washington, USA. 
**
**    This library is free software; you can redistribute it and/or
**    modify it under the terms of the GNU Lesser General Public
**    License as published by the Free Software Foundation; either
**    version 2.1 of the License, or (at your option) any later version.
**
**    This library is distributed in the hope that it will be useful,
**    but WITHOUT ANY WARRANTY; without even the implied warranty of
**    MERCHANTABILITY or FITNESS FOR A PARTICULAR PURPOSE.  See the GNU
**    Lesser General Public License for more details.
**
**    You should have received a copy of the GNU Lesser General Public
**    License along with this library; if not, write to the Free Software
**    Foundation, Inc., 59 Temple Place, Suite 330, Boston, MA  02111-1307  USA
*/

package org.systemsbiology.biofabric.ui;

import java.awt.event.ActionListener;
import java.awt.event.ActionEvent;
import java.awt.Cursor;
import java.awt.Image;
import java.awt.Point;
import java.awt.Toolkit;
import javax.swing.JPanel;
import javax.swing.Timer;
import java.net.URL;

/***************************************************************************
** 
** This class manages cursors
*/

public class CursorManager  {
  
  private Cursor currCursor_;
  private int errCount_;
  private JPanel pan_;
  private Cursor xCur_;
  private Cursor errCur_;  

  public CursorManager(JPanel pan, boolean isHeadless) {
    pan_ = pan;
    pan_.setCursor(Cursor.getPredefinedCursor(Cursor.DEFAULT_CURSOR));
    errCount_ = 0;
    if (!isHeadless) {
      buildCursors();
    }
  }

  public synchronized void showDefaultCursor() {
    currCursor_ = Cursor.getPredefinedCursor(Cursor.DEFAULT_CURSOR);
    if (errCount_ == 0) {
      pan_.setCursor(currCursor_);
    }
  }

  public synchronized void showModeCursor() {
    currCursor_ = xCur_;
    if (errCount_ == 0) {
      pan_.setCursor(currCursor_);
    }
  }    

  public synchronized void signalError() {
    if (errCount_++ == 0) { 
      pan_.setCursor(errCur_);
    }
    final Timer timer = new Timer(1000, null);
    timer.addActionListener(new ActionListener() {
      public void actionPerformed(ActionEvent ev) {
        clearError();
        timer.stop();
      }
    });
    timer.setRepeats(false);
    timer.start();
    return;
  }      

  public synchronized void clearError() {
    if (--errCount_ == 0) { 
      pan_.setCursor(currCursor_);
    }      
    return;
  } 

  private void buildCursors() {
    if (xCur_ == null) {
      Toolkit tkit = Toolkit.getDefaultToolkit();
      URL xCurGif = getClass().getResource("/org/systemsbiology/biofabric/images/CrossCursor32.gif");
      Point hot = new Point(15, 15);
      Image img = tkit.createImage(xCurGif);
      xCur_ = tkit.createCustomCursor(img, hot, "btCustomCross");
    }
    if (errCur_ == null) {
      Toolkit tkit = Toolkit.getDefaultToolkit();
      URL errCurGif = getClass().getResource("/org/systemsbiology/biofabric/images/ErrorCursor32.gif");
      Point hot = new Point(15, 15);
      Image img = tkit.createImage(errCurGif);
      errCur_ = tkit.createCustomCursor(img, hot, "btCustomError");
    }
    return;
  }       
}  
```
