## Supplementary material for "BioFabric Visualization of Network Alignments": BioFabric Source Code: BreadthFirstLayoutDialog.java

```
/*
**    Copyright (C) 2003-2014 Institute for Systems Biology 
**                            Seattle, Washington, USA. 
**
**    This library is free software; you can redistribute it and/or
**    modify it under the terms of the GNU Lesser General Public
**    License as published by the Free Software Foundation; either
**    version 2.1 of the License, or (at your option) any later version.
**
**    This library is distributed in the hope that it will be useful,
**    but WITHOUT ANY WARRANTY; without even the implied warranty of
**    MERCHANTABILITY or FITNESS FOR A PARTICULAR PURPOSE.  See the GNU
**    Lesser General Public License for more details.
**
**    You should have received a copy of the GNU Lesser General Public
**    License along with this library; if not, write to the Free Software
**    Foundation, Inc., 59 Temple Place, Suite 330, Boston, MA  02111-1307  USA
*/

package org.systemsbiology.biofabric.ui.dialogs;

import java.awt.Dimension;
import java.awt.event.ActionEvent;
import java.awt.event.ActionListener;
import java.util.ArrayList;
import java.util.Set;

import javax.swing.ButtonGroup;
import javax.swing.JFrame;
import javax.swing.JLabel;
import javax.swing.JOptionPane;
import javax.swing.JRadioButton;
import javax.swing.JTextField;

import org.systemsbiology.biofabric.api.dialog.BTStashResultsDialog;
import org.systemsbiology.biofabric.api.layout.DefaultLayout;
import org.systemsbiology.biofabric.api.model.NetNode;
import org.systemsbiology.biofabric.api.util.ExceptionHandler;
import org.systemsbiology.biofabric.model.BioFabricNetwork;
import org.systemsbiology.biofabric.util.DataUtil;
import org.systemsbiology.biofabric.util.ResourceManager;

/****************************************************************************
**
** Dialog box for setting breadth-first layout params
*/

public class BreadthFirstLayoutDialog extends BTStashResultsDialog {

  ////////////////////////////////////////////////////////////////////////////
  //
  // PRIVATE CONSTANTS
  //
  ////////////////////////////////////////////////////////////////////////////  
 
  private static final long serialVersionUID = 1L;
    
  ////////////////////////////////////////////////////////////////////////////
  //
  // PRIVATE INSTANCE MEMBERS
  //
  ////////////////////////////////////////////////////////////////////////////  
  
  
  private JRadioButton useDefault_;
  private JRadioButton userSpec_;
  private JTextField userName_;
  private DefaultLayout.Params params_;
  private JLabel nameLabel_;
  private BioFabricNetwork bfn_;
  
  ////////////////////////////////////////////////////////////////////////////
  //
  // PUBLIC CONSTRUCTORS
  //
  ////////////////////////////////////////////////////////////////////////////    

  /***************************************************************************
  **
  ** Constructor 
  */ 
  
  public BreadthFirstLayoutDialog(JFrame parent, NetNode currSel, BioFabricNetwork bfn) {     
    super(parent, ResourceManager.getManager().getString("breadthFirstLayout.title"), new Dimension(600, 350), 2);
    params_ = null;
    bfn_ = bfn;
    
    String highest = rMan_.getString("bFirst.useHighest");
    useDefault_ = new JRadioButton(highest, (currSel == null));
    useDefault_.addActionListener(new ActionListener() {
      public void actionPerformed(ActionEvent ev) {
        try {
          userName_.setEnabled(userSpec_.isSelected());
          nameLabel_.setEnabled(userSpec_.isSelected());
        } catch (Exception ex) {
          ExceptionHandler.getHandler().displayException(ex);
        }
      }
    });
    addWidgetFullRow(useDefault_, false);
        
    String spec = rMan_.getString("bFirst.userSelect");
    userSpec_ = new JRadioButton(spec, (currSel != null));
    userSpec_.addActionListener(new ActionListener() {
      public void actionPerformed(ActionEvent ev) {
        try {
          userName_.setEnabled(userSpec_.isSelected());
          nameLabel_.setEnabled(userSpec_.isSelected());
        } catch (Exception ex) {
          ExceptionHandler.getHandler().displayException(ex);
        }
      }
    });
    addWidgetFullRow(userSpec_, false);
    
    ButtonGroup group = new ButtonGroup();
    group.add(useDefault_);
    group.add(userSpec_);
    
    userName_ = new JTextField((currSel == null) ? "" : currSel.getName().trim());
    nameLabel_ = new JLabel(rMan_.getString("bFirst.selectName"));
    userName_.setEnabled(userSpec_.isSelected());
    nameLabel_.setEnabled(userSpec_.isSelected());   
    addLabeledWidget(nameLabel_, userName_, false, false); 
        
    finishConstruction();
  }
    
  ////////////////////////////////////////////////////////////////////////////
  //
  // PUBLIC METHODS
  //
  ////////////////////////////////////////////////////////////////////////////  

  /***************************************************************************
  **
  ** Get results
  */
  
  public DefaultLayout.Params getParams() {
    return (params_);
  }
  
  ////////////////////////////////////////////////////////////////////////////
  //
  // PRIVATE METHODS
  //
  ////////////////////////////////////////////////////////////////////////////

  /***************************************************************************
  **
  ** Stash our results for later interrogation.
  ** 
  */
  
  protected boolean stashForOK() {
 
    boolean useSelected = userSpec_.isSelected();
    
    if (useSelected) {
      String search = DataUtil.normKey(userName_.getText().trim());
      Set<NetNode> result = bfn_.nodeMatches(true, search);
      if (result.size() != 1) {
        ResourceManager rMan = ResourceManager.getManager();
        JOptionPane.showMessageDialog(parent_, 
                                      rMan.getString("bFirst.badNode"), 
                                      rMan.getString("bFirst.badNodeTitle"),
                                      JOptionPane.ERROR_MESSAGE);
        return (false);
      }
      ArrayList<NetNode> starts = new ArrayList<NetNode>();
      starts.add(result.iterator().next());
      params_ = new DefaultLayout.DefaultParams(starts);
    } else {
      params_ = new DefaultLayout.DefaultParams(null);
    }
   
    return (true);
  }
}
```
