## Supplementary material for "BioFabric Visualization of Network Alignments": BioFabric Source Code: ClusterLayoutSetupDialog.java

```
/*
**    Copyright (C) 2003-2018 Institute for Systems Biology 
**                            Seattle, Washington, USA. 
**
**    This library is free software; you can redistribute it and/or
**    modify it under the terms of the GNU Lesser General Public
**    License as published by the Free Software Foundation; either
**    version 2.1 of the License, or (at your option) any later version.
**
**    This library is distributed in the hope that it will be useful,
**    but WITHOUT ANY WARRANTY; without even the implied warranty of
**    MERCHANTABILITY or FITNESS FOR A PARTICULAR PURPOSE.  See the GNU
**    Lesser General Public License for more details.
**
**    You should have received a copy of the GNU Lesser General Public
**    License along with this library; if not, write to the Free Software
**    Foundation, Inc., 59 Temple Place, Suite 330, Boston, MA  02111-1307  USA
*/

package org.systemsbiology.biofabric.ui.dialogs;

import java.awt.Dimension;
import java.awt.event.ActionEvent;
import java.awt.event.ActionListener;
import java.io.File;
import java.util.HashSet;
import java.util.Iterator;
import java.util.Map;
import java.util.Set;

import javax.swing.JCheckBox;
import javax.swing.JComboBox;
import javax.swing.JFrame;
import javax.swing.JLabel;
import javax.swing.JOptionPane;
import javax.swing.JTextField;

import org.systemsbiology.biofabric.api.dialog.BTStashResultsDialog;
import org.systemsbiology.biofabric.api.io.AttributeKey;
import org.systemsbiology.biofabric.api.io.FileLoadFlows;
import org.systemsbiology.biofabric.api.model.NetNode;
import org.systemsbiology.biofabric.api.util.ExceptionHandler;
import org.systemsbiology.biofabric.api.util.FixedJButton;
import org.systemsbiology.biofabric.cmd.CommandSet;
import org.systemsbiology.biofabric.io.AttributeLoader;
import org.systemsbiology.biofabric.layouts.NodeClusterLayout;
import org.systemsbiology.biofabric.model.BioFabricNetwork;
import org.systemsbiology.biofabric.plugin.PluginSupportFactory;
import org.systemsbiology.biofabric.util.DataUtil;
import org.systemsbiology.biofabric.util.ResourceManager;
import org.systemsbiology.biofabric.util.TrueObjChoiceContent;
import org.systemsbiology.biofabric.util.UiUtil;

/****************************************************************************
**
** Dialog box for cluster layout setup
*/

public class ClusterLayoutSetupDialog extends BTStashResultsDialog {

  ////////////////////////////////////////////////////////////////////////////
  //
  // PRIVATE CONSTANTS
  //
  ////////////////////////////////////////////////////////////////////////////  
 
  private static final long serialVersionUID = 1L;
    
  ////////////////////////////////////////////////////////////////////////////
  //
  // PRIVATE INSTANCE MEMBERS
  //
  ////////////////////////////////////////////////////////////////////////////  

  private JComboBox sourceCombo_;
  private JComboBox orderCombo_;
  private JComboBox interCombo_;
  private JComboBox cLayoutCombo_;
  private JCheckBox saveAssignBox_;
  private NetNode currSel_;
  private JLabel nameLabel_;
  private JTextField userName_;
  private BioFabricNetwork bfn_;
  private NodeClusterLayout.ClusterParams results_;
  private boolean haveClusts_;
  
  ////////////////////////////////////////////////////////////////////////////
  //
  // PUBLIC CONSTRUCTORS
  //
  ////////////////////////////////////////////////////////////////////////////    

  /***************************************************************************
  **
  ** Constructor 
  */ 
  
  public ClusterLayoutSetupDialog(JFrame parent, BioFabricNetwork bfn, NetNode selNode) {     
    super(parent, ResourceManager.getManager().getString("nodeClusterLayout.title"), new Dimension(600, 350), 2);
    results_ = null;
    bfn_ = bfn;
    currSel_ = selNode;
    haveClusts_ = bfn.nodeClustersAssigned();
     
    sourceCombo_ = new JComboBox(NodeClusterLayout.ClusterParams.getSourceChoices(haveClusts_));
    orderCombo_ = new JComboBox(NodeClusterLayout.ClusterParams.getOrderChoices());
    interCombo_ = new JComboBox(NodeClusterLayout.ClusterParams.getILinkChoices());
    cLayoutCombo_ = new JComboBox(NodeClusterLayout.ClusterParams.getClustLayoutChoices());
    saveAssignBox_ = new JCheckBox(rMan_.getString("nodeClusterLayout.saveAssign"));
    
    NodeClusterLayout.ClusterParams params = new NodeClusterLayout.ClusterParams(haveClusts_);
    setToVals(params);
  
    JLabel label = new JLabel(rMan_.getString("nodeClusterLayout.source"));
    addLabeledWidget(label, sourceCombo_, false, false); 
    
    addWidgetFullRow(saveAssignBox_, false);
    
    label = new JLabel(rMan_.getString("nodeClusterLayout.order"));
    addLabeledWidget(label, orderCombo_, false, false); 
    
    label = new JLabel(rMan_.getString("nodeClusterLayout.interLink"));
    addLabeledWidget(label, interCombo_, false, false); 
    
    label = new JLabel(rMan_.getString("nodeClusterLayout.clusterLayout"));
    addLabeledWidget(label, cLayoutCombo_, false, false); 
       
    //
    // Build extra button:
    //

    FixedJButton buttonR = new FixedJButton(rMan_.getString("dialogs.resetDefaults"));
    buttonR.addActionListener(new ActionListener() {
      public void actionPerformed(ActionEvent ev) {
        try {
          NodeClusterLayout.ClusterParams params = new NodeClusterLayout.ClusterParams(haveClusts_);
          setToVals(params);
        } catch (Exception ex) {
          ExceptionHandler.getHandler().displayException(ex);
        }
      }
    });
    
    userName_ = new JTextField((currSel_ == null) ? "" : currSel_.getName().trim());
    nameLabel_ = new JLabel(rMan_.getString("bFirst.selectName"));
 //   userName_.setEnabled(userSpec_.isSelected());
   // nameLabel_.setEnabled(userSpec_.isSelected());   
    addLabeledWidget(nameLabel_, userName_, false, false); 

    finishConstructionWithExtraLeftButton(buttonR);

  }
    
  ////////////////////////////////////////////////////////////////////////////
  //
  // PUBLIC METHODS
  //
  ////////////////////////////////////////////////////////////////////////////  
  
  /***************************************************************************
  **
  ** Get results
  */
  
  public NodeClusterLayout.ClusterParams getParams() {
    return (results_);
  }
  
  ////////////////////////////////////////////////////////////////////////////
  //
  // PRIVATE METHODS
  //
  ////////////////////////////////////////////////////////////////////////////

  /***************************************************************************
  **
  ** Stash our results for later interrogation.
  ** 
  */
  
  protected boolean stashForOK() {
 
    results_ = new NodeClusterLayout.ClusterParams(haveClusts_);
    
    TrueObjChoiceContent<NodeClusterLayout.ClusterParams.Source> scc = 
    	(TrueObjChoiceContent<NodeClusterLayout.ClusterParams.Source>)sourceCombo_.getSelectedItem();
    results_.source = scc.val;
    
    TrueObjChoiceContent<NodeClusterLayout.ClusterParams.Order> occ = 
      	(TrueObjChoiceContent<NodeClusterLayout.ClusterParams.Order>)orderCombo_.getSelectedItem();    
    results_.order = occ.val;
    
    TrueObjChoiceContent<NodeClusterLayout.ClusterParams.InterLink> icc = 
      	(TrueObjChoiceContent<NodeClusterLayout.ClusterParams.InterLink>)interCombo_.getSelectedItem();    
    results_.iLink = icc.val;
    
    TrueObjChoiceContent<NodeClusterLayout.ClusterParams.ClustLayout> ccc = 
      	(TrueObjChoiceContent<NodeClusterLayout.ClusterParams.ClustLayout>)cLayoutCombo_.getSelectedItem();    
    results_.cLay = ccc.val;
    
    results_.saveAssign = saveAssignBox_.isSelected();
    
    if (results_.source.equals(NodeClusterLayout.ClusterParams.Source.STORED)) {
    	results_.assign(bfn_.nodeClusterAssigment());
    }

    String selName = userName_.getText();
    if ((selName != null) && !selName.trim().equals("")) {
    	String cand = selName.trim();
    	Map<String, Set<NetNode>> nn2ids = bfn_.getNormNameToIDs();
      Map<String, NetNode> nn2id = PluginSupportFactory.getBuildExtractor().reduceNameSetToOne(nn2ids);
      NetNode nidCand = nn2id.get(DataUtil.normKey(cand));
    	if (bfn_.getNodeDefinition(nidCand) == null) {
        ResourceManager rMan = ResourceManager.getManager();
        JOptionPane.showMessageDialog(parent_, 
                                      rMan.getString("nodeClusterLayout.nodeDoesNotExist"),
                                      rMan.getString("nodeClusterLayout.nodeDoesNotExistTitle"),
                                      JOptionPane.ERROR_MESSAGE);
        return (false);
    	}
    	results_.startNode = nidCand;
    } else {
    	results_.startNode = null;
    }
    return (true);
  }
  
  /***************************************************************************
  **
  ** Reset to default values
  ** 
  */
  
  private void setToVals(NodeClusterLayout.ClusterParams params) {
    int numSrc = sourceCombo_.getItemCount();
    for (int i = 0; i < numSrc; i++) {
      TrueObjChoiceContent<NodeClusterLayout.ClusterParams.Source> cc = 
      	(TrueObjChoiceContent<NodeClusterLayout.ClusterParams.Source>)sourceCombo_.getItemAt(i);
      if (cc.val == params.source) {
        sourceCombo_.setSelectedIndex(i);
        break;
      }
    } 
   
    int numOrd = orderCombo_.getItemCount();
    for (int i = 0; i < numOrd; i++) {
      TrueObjChoiceContent<NodeClusterLayout.ClusterParams.Order> cc = 
      	(TrueObjChoiceContent<NodeClusterLayout.ClusterParams.Order>)orderCombo_.getItemAt(i);
      if (cc.val == params.order) {
        orderCombo_.setSelectedIndex(i);
        break;
      }
    }   
    
    int numInt = interCombo_.getItemCount();
    for (int i = 0; i < numInt; i++) {
      TrueObjChoiceContent<NodeClusterLayout.ClusterParams.InterLink> cc = 
      	(TrueObjChoiceContent<NodeClusterLayout.ClusterParams.InterLink>)interCombo_.getItemAt(i);
      if (cc.val == params.iLink) {
        orderCombo_.setSelectedIndex(i);
        break;
      }
    }   
    
    int numClay = cLayoutCombo_.getItemCount();
    for (int i = 0; i < numClay; i++) {
      TrueObjChoiceContent<NodeClusterLayout.ClusterParams.ClustLayout> cc = 
      	(TrueObjChoiceContent<NodeClusterLayout.ClusterParams.ClustLayout>)cLayoutCombo_.getItemAt(i);
      if (cc.val == params.cLay) {
        orderCombo_.setSelectedIndex(i);
        break;
      }
    }
    
    saveAssignBox_.setSelected(params.saveAssign);
    return;
  }
    
  ////////////////////////////////////////////////////////////////////////////
  //
  // PUBLIC CONSTRUCTORS
  //
  ////////////////////////////////////////////////////////////////////////////    
    
    
  public static boolean askForFileInfo(NodeClusterLayout.ClusterParams params, CommandSet cset, 
                                       FileLoadFlows flf, BioFabricNetwork bfn)  {   
      
    File file = flf.getTheFile(".noa", ".na", "AttribDirectory", "filterName.noa", null);
    if (file == null) {
      return (false);
    }
    Map<AttributeKey, String> nodeAttributes = flf.loadTheFile(file, null, true);
    if (nodeAttributes == null) {
      return (false);
    }
    //
    // All existing targets must have a row, and all existing
    // rows need a target assigned!
    //
    
    HashSet<AttributeKey> asUpper = new HashSet<AttributeKey>();
    Iterator<NetNode> rttvit = bfn.getNodeSetIDs().iterator();
    while (rttvit.hasNext()) {
      asUpper.add(new AttributeLoader.StringKey(rttvit.next().getName()));
    }
    if (!asUpper.equals(new HashSet<AttributeKey>(nodeAttributes.keySet()))) {
      ResourceManager rMan = ResourceManager.getManager();
      JOptionPane.showMessageDialog(cset.getBFW().getWindow(), rMan.getString("attribRead.badRowMessage"),
                                    rMan.getString("attribRead.badRowSemanticsTitle"),
                                    JOptionPane.WARNING_MESSAGE);
      return (false);
    }
    UiUtil.fixMePrintout("Gotta make sure that no node name maps to two nodes (with different internal IDS)");
    Map<String, Set<NetNode>> nn2ids = bfn.getNormNameToIDs();
    
    
    Map<String, NetNode> nn2id = PluginSupportFactory.getBuildExtractor().reduceNameSetToOne(nn2ids);
    params.install(nodeAttributes, nn2id);
    return (true);
  }
}
```
