## Supplementary material for "BioFabric Visualization of Network Alignments": BioFabric Source Code: CompareNodesSetupDialog.java

```
/*
**    Copyright (C) 2003-2016 Institute for Systems Biology 
**                            Seattle, Washington, USA. 
**
**    This library is free software; you can redistribute it and/or
**    modify it under the terms of the GNU Lesser General Public
**    License as published by the Free Software Foundation; either
**    version 2.1 of the License, or (at your option) any later version.
**
**    This library is distributed in the hope that it will be useful,
**    but WITHOUT ANY WARRANTY; without even the implied warranty of
**    MERCHANTABILITY or FITNESS FOR A PARTICULAR PURPOSE.  See the GNU
**    Lesser General Public License for more details.
**
**    You should have received a copy of the GNU Lesser General Public
**    License along with this library; if not, write to the Free Software
**    Foundation, Inc., 59 Temple Place, Suite 330, Boston, MA  02111-1307  USA
*/

package org.systemsbiology.biofabric.ui.dialogs;

import java.awt.Dimension;
import java.awt.event.ActionEvent;
import java.awt.event.ActionListener;
import java.io.File;
import java.util.ArrayList;
import java.util.HashSet;
import java.util.Iterator;
import java.util.List;
import java.util.Map;
import java.util.Set;
import javax.swing.JFrame;
import javax.swing.JPanel;
import javax.swing.JOptionPane;

import org.systemsbiology.biofabric.api.dialog.BTStashResultsDialog;
import org.systemsbiology.biofabric.api.model.NetNode;
import org.systemsbiology.biofabric.api.util.ExceptionHandler;
import org.systemsbiology.biofabric.api.util.FixedJButton;
import org.systemsbiology.biofabric.api.util.NID;
import org.systemsbiology.biofabric.cmd.CommandSet;
import org.systemsbiology.biofabric.model.FabricNode;
import org.systemsbiology.biofabric.ui.dialogs.utils.EditableTable;
import org.systemsbiology.biofabric.util.DataUtil;
import org.systemsbiology.biofabric.util.ResourceManager;
import org.systemsbiology.biofabric.util.UiUtil;
import org.systemsbiology.biotapestry.biofabric.FabricCommands;

/****************************************************************************
**
** Dialog box for setting up a node comparison
*/

public class CompareNodesSetupDialog extends BTStashResultsDialog {
  
  ////////////////////////////////////////////////////////////////////////////
  //
  // PRIVATE INSTANCE MEMBERS
  //
  ////////////////////////////////////////////////////////////////////////////  

  private static final long serialVersionUID = 1L;
  private Set<NetNode> result_;
  private EditableTable<NodeListTableModel.TableRow> est_;
  private Set<NetNode> allNodeIDs_;
  private Map<String, Set<NetNode>> normNameToID_;
 
  ////////////////////////////////////////////////////////////////////////////
  //
  // PUBLIC CONSTRUCTORS
  //
  ////////////////////////////////////////////////////////////////////////////    

  /***************************************************************************
  **
  ** Constructor 
  */ 

  public CompareNodesSetupDialog(JFrame parent, Set<NetNode> allNodeIDs, Map<String, Set<NetNode>> normNameToID) { 
    super(parent, ResourceManager.getManager().getString("compareNodesSetup.title"), new Dimension(600, 500), 2);
    result_ = null;
    allNodeIDs_ = allNodeIDs;
    normNameToID_ = normNameToID;
    
    //
    // Build extra button:
    //

    FixedJButton buttonR = new FixedJButton(rMan_.getString("compareNodesSetup.loadFromFile"));
    buttonR.addActionListener(new ActionListener() {
      public void actionPerformed(ActionEvent ev) {
        try {
          loadFromFile();
        } catch (Exception ex) {
          ExceptionHandler.getHandler().displayException(ex);
        }
      }
    });   

    est_ = new EditableTable<NodeListTableModel.TableRow>(new NodeListTableModel(), parent_);
    EditableTable.TableParams etp = new EditableTable.TableParams();
    etp.addAlwaysAtEnd = false;
    etp.buttons = EditableTable.ALL_BUT_EDIT_BUTTONS;
    etp.singleSelectOnly = true;
    JPanel tablePan = est_.buildEditableTable(etp);
    
    addTable(tablePan, 5);
    finishConstructionWithExtraLeftButton(buttonR);

  }
  
  /***************************************************************************
  **
  ** Return results
  ** 
  */
  
  public Set<NetNode> getResults() {
    return (result_);
  }  

  ////////////////////////////////////////////////////////////////////////////
  //
  // PRIVATE METHODS
  //
  ////////////////////////////////////////////////////////////////////////////    
   
  ////////////////////////////////////////////////////////////////////////////
  //
  // INNER CLASSES
  //
  ////////////////////////////////////////////////////////////////////////////
  
  /***************************************************************************
  **
  ** The table
  */

  class NodeListTableModel extends EditableTable.TableModel<NodeListTableModel.TableRow> {

    private static final long serialVersionUID = 1L;
    private final static int NODE_NAME_  = 0;
    private final static int NUM_COL_    = 1;  
    
    private final static int NID_ = 0;
    private final static int NUM_HIDDEN_ = 1;
    
    NodeListTableModel() {
      super(NUM_COL_);
      colNames_ = new String[] {"compareNodesSetup.nodeName"};
      colClasses_ = new Class[] {String.class};
      addHiddenColumns(NUM_HIDDEN_);
    }    
   
    public class TableRow implements EditableTable.ATableRow {
      public String nodeName;
      public String nid;
      
      public TableRow() {
      }
      
      TableRow(int i) {
        nodeName = (String)columns_.get(NODE_NAME_).get(i);
        nid = (String)hiddenColumns_.get(NID_).get(i);
      }
      
      public void toCols() {
        columns_.get(NODE_NAME_).add(nodeName);  
        hiddenColumns_.get(NID_).add(nid);  
        return;
      }
    }
  
    protected TableRow constructARow(int i) {
      return (new TableRow(i));     
    }
    
    public TableRow constructARow() {
      return (new TableRow());     
    }
       
    List<NetNode> applyValues() {
      List<TableRow> vals = getValuesFromTable();
      
      ResourceManager rMan = ResourceManager.getManager();
      ArrayList<String> seenStrings = new ArrayList<String>();
      ArrayList<NetNode> seenTags = new ArrayList<NetNode>();
      int size = vals.size();
      if (size == 0) {
        return (seenTags);
      }
      
      for (int i = 0; i < size; i++) {
        TableRow row = vals.get(i);
        String tag = row.nodeName;
        if ((tag == null) || (tag.trim().equals(""))) {
          JOptionPane.showMessageDialog(parent_, rMan.getString("compareNodesSetup.badName"),
                                        rMan.getString("compareNodesSetup.badNameTitle"),
                                        JOptionPane.ERROR_MESSAGE);
          return (null);
        }
        
        tag = tag.trim();
        
        if (DataUtil.containsKey(seenStrings, tag)) {
          JOptionPane.showMessageDialog(parent_, rMan.getString("compareNodesSetup.dupName"),
                                        rMan.getString("compareNodesSetup.dupNameTitle"),
                                        JOptionPane.ERROR_MESSAGE);           
            
          return (null);
        }
                
        String normTag = DataUtil.normKey(tag);
        Set<NetNode> haveIDs = normNameToID_.get(normTag);
        boolean gotIt = (haveIDs != null);
        NetNode matchNode = null;
        if (gotIt) {
        	for (NetNode haveID : haveIDs) {
        		if (allNodeIDs_.contains(haveID)) {
        			if (matchNode != null) {
        				JOptionPane.showMessageDialog(parent_, rMan.getString("compareNodesSetup.ambiguousNode"),
                                              rMan.getString("compareNodesSetup.ambiguousTitle"),
                                              JOptionPane.ERROR_MESSAGE);           
                return (null);
        			} else {
        			  matchNode = haveID;
        			}
        		}
        	}
        }

        if (matchNode == null) {
          JOptionPane.showMessageDialog(parent_, rMan.getString("compareNodesSetup.notANode"),
                                        rMan.getString("compareNodesSetup.notANodeTitle"),
                                        JOptionPane.ERROR_MESSAGE);               
          return (null);
        }
        seenTags.add(matchNode);
        seenStrings.add(tag);
      }
      
      return (seenTags);
    }
  }  
  
  ////////////////////////////////////////////////////////////////////////////
  //
  // PROTECTED METHODS
  //
  ////////////////////////////////////////////////////////////////////////////

  /***************************************************************************
  **
  ** Stash our results for later interrogation.
  ** 
  */
  
  protected boolean stashForOK() {
    List<NetNode> av = ((NodeListTableModel)est_.getModel()).applyValues();    
    if (av == null) {
      result_ = null;
      return (false);
    }
    
    result_ = new HashSet<NetNode>(av);
    return (true);
  } 
  
  /***************************************************************************
  **
  ** Load names from a file
  ** 
  */
  
  void loadFromFile() {
    CommandSet cmd = CommandSet.getCmds("mainWindow");
    File fileEda = cmd.getFileLoader().getTheFile(".txt", null, "AttribDirectory", "filterName.txt", this);
    if (fileEda == null) {
      return;
    }
    List<String> nodes = UiUtil.simpleFileRead(fileEda);
    if (nodes == null) {
      return;
    }
    List<NodeListTableModel.TableRow> initRows = initTableRows(nodes);
    est_.updateTable(true, initRows);
    FabricCommands.setPreference("AttribDirectory", fileEda.getAbsoluteFile().getParent());
    return;   
  }
  
  ////////////////////////////////////////////////////////////////////////////
  //
  // PRIVATE METHODS
  //
  ////////////////////////////////////////////////////////////////////////////  
  
  /***************************************************************************
  **
  ** Get the list of table rows
  ** 
  */

  private List<NodeListTableModel.TableRow> initTableRows(List<String> inTags) {
    ArrayList<NodeListTableModel.TableRow> retval = new ArrayList<NodeListTableModel.TableRow>();
    EditableTable.TableModel<NodeListTableModel.TableRow> ecdtm = est_.getModel(); 
    Iterator<String> ceit = inTags.iterator();
    while (ceit.hasNext()) {
      String tag = ceit.next();
      NodeListTableModel.TableRow tr = ecdtm.constructARow();
      tr.nodeName = tag;
      retval.add(tr);
    }
    return (retval);
  } 
}
```
