## Supplementary material for "BioFabric Visualization of Network Alignments": BioFabric Source Code: ControlTopLayoutSetupDialog.java

```
/*
**    Copyright (C) 2003-2017 Institute for Systems Biology 
**                            Seattle, Washington, USA. 
**
**    This library is free software; you can redistribute it and/or
**    modify it under the terms of the GNU Lesser General Public
**    License as published by the Free Software Foundation; either
**    version 2.1 of the License, or (at your option) any later version.
**
**    This library is distributed in the hope that it will be useful,
**    but WITHOUT ANY WARRANTY; without even the implied warranty of
**    MERCHANTABILITY or FITNESS FOR A PARTICULAR PURPOSE.  See the GNU
**    Lesser General Public License for more details.
**
**    You should have received a copy of the GNU Lesser General Public
**    License along with this library; if not, write to the Free Software
**    Foundation, Inc., 59 Temple Place, Suite 330, Boston, MA  02111-1307  USA
*/

package org.systemsbiology.biofabric.ui.dialogs;

import java.awt.Dimension;

import javax.swing.JComboBox;
import javax.swing.JFrame;
import javax.swing.JLabel;

import org.systemsbiology.biofabric.api.dialog.BTStashResultsDialog;
import org.systemsbiology.biofabric.layouts.ControlTopLayout;
import org.systemsbiology.biofabric.util.ResourceManager;
import org.systemsbiology.biofabric.util.TrueObjChoiceContent;

/****************************************************************************
**
** Dialog box for setting layout params
*/

public class ControlTopLayoutSetupDialog extends BTStashResultsDialog {

  ////////////////////////////////////////////////////////////////////////////
  //
  // PRIVATE CONSTANTS
  //
  ////////////////////////////////////////////////////////////////////////////  
 
  private static final long serialVersionUID = 1L;
    
  ////////////////////////////////////////////////////////////////////////////
  //
  // PRIVATE INSTANCE MEMBERS
  //
  ////////////////////////////////////////////////////////////////////////////  
  
  private JComboBox controlTopChoice_;
  private JComboBox targetChoice_;
  private ControlTopLayout.CtrlMode ctMode_;
  private ControlTopLayout.TargMode trgMode_;
    
  
  ////////////////////////////////////////////////////////////////////////////
  //
  // PUBLIC CONSTRUCTORS
  //
  ////////////////////////////////////////////////////////////////////////////    

  /***************************************************************************
  **
  ** Constructor 
  */ 
  
  public ControlTopLayoutSetupDialog(JFrame parent) {     
    super(parent, ResourceManager.getManager().getString("controlTopLayout.title"), new Dimension(800, 200), 2);
       
    controlTopChoice_ = new JComboBox(ControlTopLayout.CtrlMode.getControlChoices());
    targetChoice_ = new JComboBox(ControlTopLayout.TargMode.getTargChoices());

    JLabel label = new JLabel(rMan_.getString("controlTopLayout.controlMode"));
    addLabeledWidget(label, controlTopChoice_, false, true); 
   
    label = new JLabel(rMan_.getString("controlTopLayout.targetMode"));
    addLabeledWidget(label, targetChoice_, false, false); 

    finishConstruction();
  }
    
  ////////////////////////////////////////////////////////////////////////////
  //
  // PUBLIC METHODS
  //
  ////////////////////////////////////////////////////////////////////////////  
  
  /***************************************************************************
  **
  ** Get results for control top mode
  */
  
  public ControlTopLayout.CtrlMode getCMode() {
    return (ctMode_);
  }
  
  /***************************************************************************
  **
  ** Get results for target mode
  */

  public ControlTopLayout.TargMode getTMode() {
    return (trgMode_); 
  }
  
  ////////////////////////////////////////////////////////////////////////////
  //
  // PRIVATE METHODS
  //
  ////////////////////////////////////////////////////////////////////////////

  /***************************************************************************
  **
  ** Stash our results for later interrogation.
  ** 
  */
  
  protected boolean stashForOK() {

    TrueObjChoiceContent<ControlTopLayout.CtrlMode> tocc =
      (TrueObjChoiceContent<ControlTopLayout.CtrlMode>)controlTopChoice_.getSelectedItem();
    ctMode_ = tocc.val;
    
    TrueObjChoiceContent<ControlTopLayout.TargMode> tocc2 =
        (TrueObjChoiceContent<ControlTopLayout.TargMode>)targetChoice_.getSelectedItem();
    trgMode_ = tocc2.val;
    
    return (true);
  }
}
```
