## Supplementary material for "BioFabric Visualization of Network Alignments": BioFabric Source Code: DirectoryChooserDialog.java

```
/*
**    Copyright (C) 2003-2018 Institute for Systems Biology
**                            Seattle, Washington, USA.
**
**    This library is free software; you can redistribute it and/or
**    modify it under the terms of the GNU Lesser General Public
**    License as published by the Free Software Foundation; either
**    version 2.1 of the License, or (at your option) any later version.
**
**    This library is distributed in the hope that it will be useful,
**    but WITHOUT ANY WARRANTY; without even the implied warranty of
**    MERCHANTABILITY or FITNESS FOR A PARTICULAR PURPOSE.  See the GNU
**    Lesser General Public License for more details.
**
**    You should have received a copy of the GNU Lesser General Public
**    License along with this library; if not, write to the Free Software
**    Foundation, Inc., 59 Temple Place, Suite 330, Boston, MA  02111-1307  USA
*/

package org.systemsbiology.biofabric.ui.dialogs;

import java.awt.Dimension;
import java.awt.GridBagLayout;
import java.awt.event.ActionEvent;
import java.awt.event.ActionListener;
import java.io.File;

import javax.swing.JFrame;
import javax.swing.JLabel;
import javax.swing.JPanel;
import javax.swing.JTextField;
import javax.swing.border.EmptyBorder;

import org.systemsbiology.biofabric.api.dialog.BTStashResultsDialog;
import org.systemsbiology.biofabric.api.io.FileLoadFlows;
import org.systemsbiology.biofabric.api.util.ExceptionHandler;
import org.systemsbiology.biofabric.api.util.FixedJButton;
import org.systemsbiology.biofabric.util.ResourceManager;

public class DirectoryChooserDialog extends BTStashResultsDialog {
  
  ////////////////////////////////////////////////////////////////////////////
  //
  // PRIVATE INSTANCE MEMBERS
  //
  ////////////////////////////////////////////////////////////////////////////
  
  private JFrame parent_;
  private JTextField nameField_;
  private File directory_;
  private static final long serialVersionUID = 1L;
  private FileLoadFlows flf_;
  

  public DirectoryChooserDialog(JFrame parent, FileLoadFlows flf, String currChoice) {
    super(parent, ResourceManager.getManager().getString("directoryChooser.title"), new Dimension(800, 150), 3);
    
    ResourceManager rMan = ResourceManager.getManager();
     
    parent_ = parent;
    flf_ = flf;
    
    JPanel cp = (JPanel) getContentPane();
    cp.setBorder(new EmptyBorder(20, 20, 20, 20));
    cp.setLayout(new GridBagLayout());
    
    //
    // File Buttons and File Labels
    //
    
    FixedJButton directoryBrowse = new FixedJButton(rMan.getString("directoryChooser.browse"));
    directoryBrowse.addActionListener(new ActionListener() {
      public void actionPerformed(ActionEvent e) {
        try {
        	File directory = flf_.getTheDirectory("PlugInDirectory");
        	if (directory != null) {
        	  nameField_.setText(directory.getAbsolutePath());
        	}
        } catch (Exception ex) {
          ExceptionHandler.getHandler().displayException(ex);
        }
      }
    });
    
  
    nameField_ = new JTextField(30);
    nameField_.setText((currChoice == null) ? "" : currChoice);
    JLabel directoryLabel = new JLabel(rMan.getString("directoryChooser.directory"));
    addLabeledFileBrowse(directoryLabel, nameField_, directoryBrowse);
    
    //
    // OK button
    //
    
    finishConstruction();
    setLocationRelativeTo(parent);
  }
 
  ////////////////////////////////////////////////////////////////////////////
  //
  // PUBLIC METHODS AND CLASSES
  //
  ////////////////////////////////////////////////////////////////////////////
  
  
  protected boolean stashForOK() {
  	directory_ = new File(nameField_.getText());
  	if (!flf_.standardFileChecks(directory_, FileLoadFlows.FILE_MUST_EXIST, FileLoadFlows.FILE_CAN_CREATE_DONT_CARE, 
                                 FileLoadFlows.FILE_DONT_CHECK_OVERWRITE, FileLoadFlows.FILE_MUST_BE_DIRECTORY, 
                                 FileLoadFlows.FILE_CAN_WRITE_DONT_CARE, FileLoadFlows.FILE_CAN_READ)) {
  		return (false);
  	}
    return (true);
  }
  
  public File getDirectory() {
    return (directory_);
  }  
}
```
