## Supplementary material for "BioFabric Visualization of Network Alignments": BioFabric Source Code: ExceptionDialog.java

```
/*
**    Copyright (C) 2003-2014 Institute for Systems Biology 
**                            Seattle, Washington, USA. 
**
**    This library is free software; you can redistribute it and/or
**    modify it under the terms of the GNU Lesser General Public
**    License as published by the Free Software Foundation; either
**    version 2.1 of the License, or (at your option) any later version.
**
**    This library is distributed in the hope that it will be useful,
**    but WITHOUT ANY WARRANTY; without even the implied warranty of
**    MERCHANTABILITY or FITNESS FOR A PARTICULAR PURPOSE.  See the GNU
**    Lesser General Public License for more details.
**
**    You should have received a copy of the GNU Lesser General Public
**    License along with this library; if not, write to the Free Software
**    Foundation, Inc., 59 Temple Place, Suite 330, Boston, MA  02111-1307  USA
*/

package org.systemsbiology.biofabric.ui.dialogs;

import java.awt.GridBagLayout;

import java.awt.Dimension;
import java.awt.GridBagConstraints;
import java.awt.event.ActionListener;
import java.awt.event.ActionEvent;
import javax.swing.JDialog;
import javax.swing.JTextArea;
import javax.swing.JFrame;
import javax.swing.JPanel;
import javax.swing.JLabel;
import javax.swing.JScrollPane;
import javax.swing.Box;
import javax.swing.border.EmptyBorder;
import java.io.StringWriter;
import java.io.PrintWriter;

import org.systemsbiology.biofabric.api.util.ExceptionHandler;
import org.systemsbiology.biofabric.api.util.FixedJButton;
import org.systemsbiology.biofabric.util.ResourceManager;
import org.systemsbiology.biofabric.util.UiUtil;
 
/****************************************************************************
**
** Dialog box for reporting an exception
*/

public class ExceptionDialog extends JDialog {
  
  ////////////////////////////////////////////////////////////////////////////
  //
  // PRIVATE CONSTANTS
  //
  ////////////////////////////////////////////////////////////////////////////       
  
  ////////////////////////////////////////////////////////////////////////////
  //
  // PRIVATE INSTANCE MEMBERS
  //
  //////////////////////////////////////////////////////////////////////////// 
  
  private static final long serialVersionUID = 1L;
  
  ////////////////////////////////////////////////////////////////////////////
  //
  // PUBLIC CONSTRUCTORS
  //
  ////////////////////////////////////////////////////////////////////////////    

  /***************************************************************************
  **
  ** Constructor 
  */ 
  
  public ExceptionDialog(JFrame parent, Exception ex, String version) {     
    super(parent, ResourceManager.getManager().getString("exception.title"), true);
       
    ResourceManager rMan = ResourceManager.getManager();    
    setSize(800, 300);
    JPanel cp = (JPanel)getContentPane();
    cp.setBorder(new EmptyBorder(20, 20, 20, 20));
    cp.setLayout(new GridBagLayout());
    cp.setMinimumSize(new Dimension(200, 200));
    cp.setPreferredSize(new Dimension(200, 200));    
    GridBagConstraints gbc = new GridBagConstraints();    

    JLabel label = new JLabel(rMan.getString("exception.report"));    
    UiUtil.gbcSet(gbc, 0, 0, 5, 1, UiUtil.BO, 0, 0, 0, 0, 0, 0, UiUtil.CEN, 1.0, 0.0);
    cp.add(label, gbc);
    
    label = new JLabel(rMan.getString("exception.pleaseExit"));    
    UiUtil.gbcSet(gbc, 0, 1, 5, 1, UiUtil.BO, 0, 0, 0, 0, 0, 0, UiUtil.CEN, 1.0, 0.0);
    cp.add(label, gbc);
          
    //
    // Stack trace:
    //
    
    StringWriter sw = new StringWriter();
    PrintWriter out = new PrintWriter(sw);
    
    out.println(version);
    ex.printStackTrace(out);
       
    JTextArea text = new JTextArea(sw.toString());
    text.setEditable(false);
    JScrollPane jsp = new JScrollPane(text); 

    UiUtil.gbcSet(gbc, 0, 2, 5, 8, UiUtil.BO, 0, 0, 5, 5, 5, 5, UiUtil.CEN, 1.0, 1.0);
    cp.add(jsp, gbc);    
    
    //
    // Build the button panel:
    //

    FixedJButton buttonC = new FixedJButton(rMan.getString("dialogs.close"));
    buttonC.addActionListener(new ActionListener() {
      public void actionPerformed(ActionEvent ev) {
        try {
        ExceptionDialog.this.setVisible(false);
        ExceptionDialog.this.dispose();
                  } catch (Exception ex) {
        ExceptionHandler.getHandler().displayException(ex);
      }
      }
    });
    
    Box buttonPanel = Box.createHorizontalBox();
    buttonPanel.add(Box.createHorizontalGlue());    
    buttonPanel.add(buttonC);
    buttonPanel.add(Box.createHorizontalStrut(10));     

    //
    // Build the dialog:
    //
  UiUtil.gbcSet(gbc, 0, 10, 5, 1, UiUtil.HOR, 0, 0, 5, 5, 5, 5, UiUtil.SE, 1.0, 0.0);
    cp.add(buttonPanel, gbc);
    setLocationRelativeTo(parent);
  }
  
  ////////////////////////////////////////////////////////////////////////////
  //
  // PUBLIC METHODS
  //
  ////////////////////////////////////////////////////////////////////////////
 
  ////////////////////////////////////////////////////////////////////////////
  //
  // INNER CLASSES
  //
  ////////////////////////////////////////////////////////////////////////////

  ////////////////////////////////////////////////////////////////////////////
  //
  // PRIVATE METHODS
  //
  ////////////////////////////////////////////////////////////////////////////  
  
}
```
