## Supplementary material for "BioFabric Visualization of Network Alignments": BioFabric Source Code: ExportSettingsDialog.java

```
/*
**    Copyright (C) 2003-2014 Institute for Systems Biology 
**                            Seattle, Washington, USA. 
**
**    This library is free software; you can redistribute it and/or
**    modify it under the terms of the GNU Lesser General Public
**    License as published by the Free Software Foundation; either
**    version 2.1 of the License, or (at your option) any later version.
**
**    This library is distributed in the hope that it will be useful,
**    but WITHOUT ANY WARRANTY; without even the implied warranty of
**    MERCHANTABILITY or FITNESS FOR A PARTICULAR PURPOSE.  See the GNU
**    Lesser General Public License for more details.
**
**    You should have received a copy of the GNU Lesser General Public
**    License along with this library; if not, write to the Free Software
**    Foundation, Inc., 59 Temple Place, Suite 330, Boston, MA  02111-1307  USA
*/

package org.systemsbiology.biofabric.ui.dialogs;

import java.awt.GridBagLayout;
import java.awt.GridBagConstraints;
import java.awt.event.ActionListener;
import java.awt.event.ActionEvent;
import javax.swing.event.CaretListener;
import javax.swing.event.CaretEvent;
import java.awt.event.FocusListener;
import java.awt.event.FocusEvent;
import java.awt.Toolkit;
import java.awt.Dimension;
import javax.swing.JDialog;
import javax.swing.JFrame;
import javax.swing.JLabel;
import javax.swing.JTextField;
import javax.swing.JPanel;
import javax.swing.JComboBox;
import javax.swing.JOptionPane;
import javax.swing.Box;
import javax.swing.border.EmptyBorder;
import java.util.prefs.Preferences;
import java.util.List;
import java.text.MessageFormat;

import org.systemsbiology.biofabric.api.util.ExceptionHandler;
import org.systemsbiology.biofabric.api.util.FixedJButton;
import org.systemsbiology.biofabric.ui.ImageExporter;
import org.systemsbiology.biofabric.util.ResourceManager;
import org.systemsbiology.biofabric.util.UiUtil;

/****************************************************************************
**
** Dialog box for setting up export
*/

public class ExportSettingsDialog extends JDialog {

  ////////////////////////////////////////////////////////////////////////////
  //
  // PUBLIC CONSTANTS
  //
  //////////////////////////////////////////////////////////////////////////// 
  
  public static final long HUGE_PIC = 3500 * 3500;  

  ////////////////////////////////////////////////////////////////////////////
  //
  // PRIVATE CONSTANTS
  //
  //////////////////////////////////////////////////////////////////////////// 

  private static final int NONE_   = 0;  
  private static final int ZOOM_   = 1;
  private static final int HEIGHT_ = 2;
  private static final int WIDTH_  = 3;

  ////////////////////////////////////////////////////////////////////////////
  //
  // PRIVATE INSTANCE MEMBERS
  //
  ////////////////////////////////////////////////////////////////////////////  

  private JTextField zoomField_;
  private double currentZoomVal_;
  private JTextField heightField_;
  private long currentZoomHeight_;
  private JTextField widthField_;
  private long currentZoomWidth_;
  private JComboBox combo_;
  private boolean processing_;
  private String currentFormat_;
     
  private int baseWidth_;
  private int baseHeight_;
  private ExportSettings finishedSettings_;  
  private Preferences prefs_;
  private static final long serialVersionUID = 1L;
    
  ////////////////////////////////////////////////////////////////////////////
  //
  // PUBLIC CONSTRUCTORS
  //
  ////////////////////////////////////////////////////////////////////////////    

  /***************************************************************************
  **
  ** Constructor 
  */ 
  
  public ExportSettingsDialog(JFrame parent, int baseWidth, int baseHeight) {     
    super(parent, ResourceManager.getManager().getString("exportDialog.title"), true);
    
    ResourceManager rMan = ResourceManager.getManager();
    setSize(400, 350);
    JPanel cp = (JPanel)getContentPane();
    cp.setBorder(new EmptyBorder(20, 20, 20, 20));
    cp.setLayout(new GridBagLayout());
    GridBagConstraints gbc = new GridBagConstraints();
    
    finishedSettings_ = null;
    prefs_ = Preferences.userNodeForPackage(this.getClass());
    currentZoomVal_ = prefs_.getDouble("ExportZoom", 1.0);
    List<String> types = ImageExporter.getSupportedExports();
    String maybeFormat = prefs_.get("ExportFormat", "PNG");
    if (types.contains(maybeFormat)) {
      currentFormat_ = maybeFormat;
    } else {
      currentFormat_ = types.get(0);
    }
    
    JPanel zoomPanel = buildZoomPanel(baseWidth, baseHeight, types);
    
    UiUtil.gbcSet(gbc, 0, 0, 10, 8, UiUtil.BO, 0, 0, 5, 5, 5, 5, UiUtil.CEN, 1.0, 1.0);    
    cp.add(zoomPanel, gbc);    

    //
    // Build the button panel:
    //

    FixedJButton buttonO = new FixedJButton(rMan.getString("exportDialog.ok"));
    buttonO.addActionListener(new ActionListener() {
      public void actionPerformed(ActionEvent ev) {
        try {
          if (extractProperties()) {
            ExportSettingsDialog.this.setVisible(false);
            ExportSettingsDialog.this.dispose();
          }
        } catch (Exception ex) {
          ExceptionHandler.getHandler().displayException(ex);
        }
      }
    });     
    FixedJButton buttonC = new FixedJButton(rMan.getString("exportDialog.cancel"));
    buttonC.addActionListener(new ActionListener() {
      public void actionPerformed(ActionEvent ev) {
        try {
          ExportSettingsDialog.this.setVisible(false);
          ExportSettingsDialog.this.dispose();
        } catch (Exception ex) {
          ExceptionHandler.getHandler().displayException(ex);
        }
      }
    });
    Box buttonPanel = Box.createHorizontalBox();
    buttonPanel.add(Box.createHorizontalGlue());    
    buttonPanel.add(buttonO);
    buttonPanel.add(Box.createHorizontalStrut(10));    
    buttonPanel.add(buttonC);    

    //
    // Build the dialog:
    //
    
    UiUtil.gbcSet(gbc, 0, 8, 10, 1, UiUtil.HOR, 0, 0, 5, 5, 5, 5, UiUtil.SE, 1.0, 0.0);
    cp.add(buttonPanel, gbc);    
    setLocationRelativeTo(parent);
  }
  
  /***************************************************************************
  **
  ** Get the dialog results.  Will be null on cancel.
  */
  
  public ExportSettings getResults() {
    return (finishedSettings_);
  }
    
  ////////////////////////////////////////////////////////////////////////////
  //
  // INNER CLASSES
  //
  ////////////////////////////////////////////////////////////////////////////  
 
  /***************************************************************************
  **
  ** These are the settings we return
  ** 
  */
  
  public static class ExportSettings {
    public double zoomVal;
    public String formatType;
    public ImageExporter.ResolutionSettings res;
    public Dimension size;
  }   

  ////////////////////////////////////////////////////////////////////////////
  //
  // PRIVATE METHODS
  //
  ////////////////////////////////////////////////////////////////////////////

  /***************************************************************************
  **
  ** Process a new zoom value
  */
  
  private void processZoomVal(boolean force, int whichVal) {

    //
    // While typing, blank fields are OK.  If that happens, we don't update anything else.
    // Bad or negative results give a beep, and we don't update anything either.  During
    // a force, bad results mean we replace the bad field with the current valid value,
    // as well as a beep.
    //
    
    if (processing_) {
      return;
    }
    processing_ = true;
    
    boolean fixVal = false;
    
    double newZoom = currentZoomVal_;
    if (whichVal == ZOOM_) {
      String zoomText = zoomField_.getText();
      boolean haveZoom = false;
      if (!zoomText.trim().equals("")) {
        try {
          double parsedZoom = Double.parseDouble(zoomText);
          if (parsedZoom > 0.0) {
            newZoom = parsedZoom;
            haveZoom = true;
          } else if (parsedZoom == 0) {
            if (force) {
              Toolkit.getDefaultToolkit().beep();
            }
          }
          if (haveZoom && tooSmall(newZoom, 0L, ZOOM_)) {
            newZoom = currentZoomVal_;
            haveZoom = false;
            Toolkit.getDefaultToolkit().beep();
          }
        } catch (NumberFormatException nfe) {
          Toolkit.getDefaultToolkit().beep();
        }
      } else if (force) {
        Toolkit.getDefaultToolkit().beep();  
      }

      if (force && !haveZoom) {
        fixVal = true;
      }
    }
    
    long newHeight = currentZoomHeight_;
    if (whichVal == HEIGHT_) {
      String heightText = heightField_.getText();
      boolean haveHeight = false;
      if (!heightText.trim().equals("")) {
        try {
          long parsedHeight = Long.parseLong(heightText);
          if (parsedHeight > 0) {
            newHeight = parsedHeight;
            haveHeight = true;
          } else {
            Toolkit.getDefaultToolkit().beep();
          }
          if (haveHeight && tooSmall(0.0, newHeight, HEIGHT_)) {
            newHeight = currentZoomHeight_;
            haveHeight = false;
            Toolkit.getDefaultToolkit().beep();
          }          
        } catch (NumberFormatException nfe) {
          Toolkit.getDefaultToolkit().beep();
        }
      } else if (force) {
        Toolkit.getDefaultToolkit().beep();  
      }

      if (force && !haveHeight) {
        fixVal = true;
      }
    }    

    long newWidth = currentZoomWidth_;
    if (whichVal == WIDTH_) {
      String widthText = widthField_.getText();
      boolean haveWidth = false;
      if (!widthText.trim().equals("")) {
        try {
          long parsedWidth = Long.parseLong(widthText);
          if (parsedWidth > 0) {
            newWidth = parsedWidth;
            haveWidth = true;
          } else {
            Toolkit.getDefaultToolkit().beep();
          }
          if (haveWidth && tooSmall(0.0, newWidth, WIDTH_)) {
            newWidth = currentZoomWidth_;
            haveWidth = false;
            Toolkit.getDefaultToolkit().beep();
          }  
        } catch (NumberFormatException nfe) {
          Toolkit.getDefaultToolkit().beep();
        }
      } else if (force) {
        Toolkit.getDefaultToolkit().beep();  
      }

      if (force && !haveWidth) {
        fixVal = true;
      }
    }    
    
    switch (whichVal) {
      case NONE_:
        break;  
      case ZOOM_:
        if (currentZoomVal_ != newZoom) {
          currentZoomVal_ = newZoom;
          currentZoomHeight_ = Math.round(baseHeight_ * currentZoomVal_);
          currentZoomWidth_ = Math.round(baseWidth_ * currentZoomVal_);
          widthField_.setText(Long.toString(currentZoomWidth_));
          heightField_.setText(Long.toString(currentZoomHeight_));
        }
        if (fixVal) {
          zoomField_.setText(Double.toString(currentZoomVal_));
        }
        break;
      case HEIGHT_:
        if (currentZoomHeight_ != newHeight) {
          currentZoomHeight_ = newHeight;
          currentZoomVal_ = currentZoomHeight_ /(double)baseHeight_;
          currentZoomWidth_ = Math.round(baseWidth_ * currentZoomVal_);
          zoomField_.setText(Double.toString(currentZoomVal_));
          widthField_.setText(Long.toString(currentZoomWidth_));
        }
        if (fixVal) {
          heightField_.setText(Long.toString(currentZoomHeight_));
        }
        break;        
      case WIDTH_:
        if (currentZoomWidth_ != newWidth) {
          currentZoomWidth_ = newWidth;
          currentZoomVal_ = currentZoomWidth_ /(double)baseWidth_;
          currentZoomHeight_ = Math.round(baseHeight_ * currentZoomVal_);
          zoomField_.setText(Double.toString(currentZoomVal_));
          heightField_.setText(Long.toString(currentZoomHeight_));
        }
        if (fixVal) {
          widthField_.setText(Long.toString(currentZoomWidth_));
        }
        break;        
      default:
        processing_ = false;
        throw new IllegalStateException();
    }
    processing_ = false;
    return;
  } 

  /***************************************************************************
  **
  ** fix zoom values
  */
  
  private void fixZoomVals() {    
    if (processing_) {
      return;
    }
    processing_ = true;
    widthField_.setText(Long.toString(currentZoomWidth_));
    heightField_.setText(Long.toString(currentZoomHeight_));
    zoomField_.setText(Double.toString(currentZoomVal_));
    processing_ = false;
    return;
  }  
 
  /***************************************************************************
  **
  ** Check if too small
  */
  
  private boolean tooSmall(double newZoom, long newDim, int whichVal) {
    switch (whichVal) {
      case ZOOM_:
        long newZoomHeight = Math.round(baseHeight_ * newZoom);
        long newZoomWidth = Math.round(baseWidth_ * newZoom);
        if ((newZoomHeight == 0) || (newZoomWidth == 0)) {
          return (true);
        }
        break;
      case HEIGHT_:
        double newZoomVal = newDim /(double)baseHeight_;
        newZoomWidth = Math.round(baseWidth_ * newZoomVal);
        if ((newZoomVal == 0) || (newZoomWidth == 0)) {
          return (true);
        }
        break;        
      case WIDTH_:
        newZoomVal = newDim /(double)baseWidth_;
        newZoomHeight = Math.round(baseHeight_ * newZoomVal);
        if ((newZoomVal == 0) || (newZoomHeight == 0)) {
          return (true);
        }
        break;        
      default:
        throw new IllegalStateException();
    }
    return (false);
  }    

  /***************************************************************************
  **
  ** Build zoom-based panel
  ** 
  */
  
  private JPanel buildZoomPanel(int baseWidth, int baseHeight, List types) {    
    //
    // Build the zoom setting input:
    //
    
    JPanel retval = new JPanel();
    retval.setLayout(new GridBagLayout());
    GridBagConstraints gbc = new GridBagConstraints();     
    ResourceManager rMan = ResourceManager.getManager();

    JLabel fixedLab = new JLabel(rMan.getString("exportDialog.fixedAspect"));    
    UiUtil.gbcSet(gbc, 0, 0, 2, 1, UiUtil.BO, 0, 0, 5, 5, 5, 5, UiUtil.E, 1.0, 1.0);    
    retval.add(fixedLab, gbc);    
   
    JLabel label = new JLabel(rMan.getString("exportDialog.zoom"));
    zoomField_ = new JTextField(Double.toString(currentZoomVal_));
    ZoomListener zl = new ZoomListener(ZOOM_);
    zoomField_.addActionListener(zl);
    zoomField_.addCaretListener(zl);
    zoomField_.addFocusListener(zl);
    
    UiUtil.gbcSet(gbc, 0, 1, 1, 1, UiUtil.BO, 0, 0, 5, 5, 5, 5, UiUtil.E, 1.0, 1.0);    
    retval.add(label, gbc);
    UiUtil.gbcSet(gbc, 1, 1, 1, 1, UiUtil.HOR, 0, 0, 5, 5, 5, 5, UiUtil.CEN, 0.5, 1.0);    
    retval.add(zoomField_, gbc);
        
    baseWidth_ = baseWidth;
    JLabel wlab = new JLabel(rMan.getString("exportDialog.width"));
    widthField_ = new JTextField();

    baseHeight_ = baseHeight;
    JLabel hlab = new JLabel(rMan.getString("exportDialog.height"));
    heightField_ = new JTextField();
    
    initZoomDims();

    ZoomListener zlw = new ZoomListener(WIDTH_);    
    widthField_.addActionListener(zlw);
    widthField_.addCaretListener(zlw);
    widthField_.addFocusListener(zlw);
    
    ZoomListener zlh = new ZoomListener(HEIGHT_);    
    heightField_.addActionListener(zlh);
    heightField_.addCaretListener(zlh);
    heightField_.addFocusListener(zlh);    
 
    UiUtil.gbcSet(gbc, 0, 2, 1, 1, UiUtil.BO, 0, 0, 5, 5, 5, 5, UiUtil.E, 1.0, 1.0);           
    retval.add(wlab, gbc);
    UiUtil.gbcSet(gbc, 1, 2, 1, 1, UiUtil.HOR, 0, 0, 5, 5, 5, 5, UiUtil.CEN, 0.5, 1.0);           
    retval.add(widthField_, gbc);
    
    UiUtil.gbcSet(gbc, 0, 3, 1, 1, UiUtil.BO, 0, 0, 5, 5, 5, 5, UiUtil.E, 1.0, 1.0);           
    retval.add(hlab, gbc);
    UiUtil.gbcSet(gbc, 1, 3, 1, 1, UiUtil.HOR, 0, 0, 5, 5, 5, 5, UiUtil.CEN, 0.5, 1.0);    
    retval.add(heightField_, gbc);    
      
    //
    // Build the export format selection:
    //
    
    JLabel flabel = new JLabel(rMan.getString("exportDialog.format"));
    combo_ = new JComboBox(types.toArray(new String[types.size()]));
    combo_.setSelectedItem(currentFormat_);
    Object currCombo = combo_.getSelectedItem();
    if ((currCombo == null) || (!currCombo.equals(currentFormat_))) {
      throw new IllegalArgumentException();
    }
    UiUtil.gbcSet(gbc, 0, 4, 1, 1, UiUtil.BO, 0, 0, 5, 5, 5, 5, UiUtil.E, 1.0, 1.0);           
    retval.add(flabel, gbc);
    UiUtil.gbcSet(gbc, 1, 4, 1, 1, UiUtil.HOR, 0, 0, 5, 5, 5, 5, UiUtil.CEN, 0.0, 1.0);    
    retval.add(combo_, gbc);
    return (retval);
  }

 /***************************************************************************
  **
  ** Init displayed zoom dimensions
  ** 
  */
  
  private void initZoomDims() {
    currentZoomHeight_ = Math.round(baseHeight_ * currentZoomVal_);
    currentZoomWidth_ = Math.round(baseWidth_ * currentZoomVal_);    
    widthField_.setText(Long.toString(currentZoomWidth_));
    heightField_.setText(Long.toString(currentZoomHeight_));
    return;
  }

  /***************************************************************************
  **
  ** Get ready for export
  ** 
  */
  
  private boolean extractProperties() {
    
    //
    // Get the dimensions:
    //
    
    long numPix = currentZoomWidth_ * currentZoomHeight_;
    if (numPix > HUGE_PIC) {
      ResourceManager rMan = ResourceManager.getManager();
      String desc = MessageFormat.format(rMan.getString("export.confirmBigExport"), 
                                         new Object[] {new Long(numPix)});
      desc = UiUtil.convertMessageToHtml(desc);                                         
      int result = JOptionPane.showConfirmDialog(this, desc,
                                                 rMan.getString("export.confirmBigExportTitle"),
                                                 JOptionPane.YES_NO_OPTION);
      if (result != 0) {
        return (false);
      }
    }
    
    finishedSettings_ = new ExportSettings();
    finishedSettings_.zoomVal = currentZoomVal_;
    finishedSettings_.formatType = (String)combo_.getSelectedItem();
 
    //
    // Get the resolution IF REQUIRED
    //
    
    if (ImageExporter.formatRequiresResolution(finishedSettings_.formatType)) {
      List resList = ImageExporter.getSupportedResolutions(false);    
      if (resList.size() == 0) {
        throw new IllegalStateException();
      }
      Object[] resVal = (Object[])resList.get(0);
      ImageExporter.RationalNumber ratNum = (ImageExporter.RationalNumber)resVal[ImageExporter.CM];

      finishedSettings_.res = new ImageExporter.ResolutionSettings();
      finishedSettings_.res.dotsPerUnit = ratNum;
      finishedSettings_.res.units = ImageExporter.CM;
    } else {
      finishedSettings_.res = null;
    }
   
    //
    // Get the dimensions:
    //
    
    finishedSettings_.size = new Dimension((int)currentZoomWidth_, (int)currentZoomHeight_);
    
    //
    // Save the preferences:
    //
   
    prefs_.putDouble("ExportZoom", currentZoomVal_);  
    prefs_.put("ExportFormat", finishedSettings_.formatType);    
    return (true);
  }
  
  ////////////////////////////////////////////////////////////////////////////
  //
  // PRIVATE INNER CLASSES
  //
  ////////////////////////////////////////////////////////////////////////////  
 
  /***************************************************************************
  **
  ** For handling zoom textField changes
  ** 
  */
  
  private class ZoomListener implements ActionListener, CaretListener, FocusListener {
    
    private int whichVal_;
    
    ZoomListener(int whichVal) {
      whichVal_ = whichVal;
    }
    
    public void actionPerformed(ActionEvent evt) {
      try {
        processZoomVal(true, whichVal_);
      } catch (Exception ex) {
        ExceptionHandler.getHandler().displayException(ex);
      }
    }
    public void caretUpdate(CaretEvent evt) {
      try {
        processZoomVal(false, whichVal_);
      } catch (Exception ex) {
        ExceptionHandler.getHandler().displayException(ex);
      }
    }
    public void focusGained(FocusEvent evt) {
    }    
    public void focusLost(FocusEvent evt) {
      try {
        fixZoomVals();
      } catch (Exception ex) {
        ExceptionHandler.getHandler().displayException(ex);
      }
    }        
  }   
}
```
