## Supplementary material for "BioFabric Visualization of Network Alignments": BioFabric Source Code: ExportSettingsPublishDialog.java

```
/*
**    Copyright (C) 2003-2012 Institute for Systems Biology 
**                            Seattle, Washington, USA. 
**
**    This library is free software; you can redistribute it and/or
**    modify it under the terms of the GNU Lesser General Public
**    License as published by the Free Software Foundation; either
**    version 2.1 of the License, or (at your option) any later version.
**
**    This library is distributed in the hope that it will be useful,
**    but WITHOUT ANY WARRANTY; without even the implied warranty of
**    MERCHANTABILITY or FITNESS FOR A PARTICULAR PURPOSE.  See the GNU
**    Lesser General Public License for more details.
**
**    You should have received a copy of the GNU Lesser General Public
**    License along with this library; if not, write to the Free Software
**    Foundation, Inc., 59 Temple Place, Suite 330, Boston, MA  02111-1307  USA
*/


package org.systemsbiology.biofabric.ui.dialogs;

import java.awt.GridBagLayout;
import java.awt.GridBagConstraints;
import java.awt.event.ActionListener;
import java.awt.event.ActionEvent;
import javax.swing.event.CaretListener;
import javax.swing.event.CaretEvent;
import java.awt.event.FocusListener;
import java.awt.event.FocusEvent;
import java.awt.Toolkit;
import java.awt.Dimension;
import javax.swing.JDialog;
import javax.swing.JFrame;
import javax.swing.JLabel;
import javax.swing.JTextField;
import javax.swing.JPanel;
import javax.swing.JComboBox;
import javax.swing.JCheckBox;
import javax.swing.Box;
import javax.swing.border.EmptyBorder;
import javax.swing.JRadioButton;
import javax.swing.ButtonGroup;
import javax.swing.JOptionPane;
import java.util.prefs.Preferences;
import java.util.List;
import java.text.MessageFormat;

import org.systemsbiology.biofabric.api.util.ExceptionHandler;
import org.systemsbiology.biofabric.api.util.FixedJButton;
import org.systemsbiology.biofabric.ui.ImageExporter;
import org.systemsbiology.biofabric.util.ResourceManager;
import org.systemsbiology.biofabric.util.UiUtil;

/****************************************************************************
**
** Dialog box for setting up export
*/

public class ExportSettingsPublishDialog extends JDialog {
  
  ////////////////////////////////////////////////////////////////////////////
  //
  // PRIVATE CONSTANTS
  //
  //////////////////////////////////////////////////////////////////////////// 

  private static final int NONE_   = 0;
  private static final int ZOOM_   = 1;  
  private static final int HEIGHT_ = 2;
  private static final int WIDTH_  = 3;
  private static final int ASPECT_ = 4;
  private static final int RES_    = 5;  
  
  private static final double CM_PER_INCH_ = 2.540;
  private static final double MIN_SIZE_INCH_ = 0.1;  
  private static final double MIN_SIZE_CM_ = CM_PER_INCH_ * MIN_SIZE_INCH_;
  private static final double MAX_SIZE_INCH_ = 72.0;  
  private static final double MAX_SIZE_CM_ = CM_PER_INCH_ * MAX_SIZE_INCH_;  
  
  private static final double DIM_MULT_  = 100.0;
  private static final double ZOOM_MULT_ = 100000.0;
 
  ////////////////////////////////////////////////////////////////////////////
  //
  // PRIVATE INSTANCE MEMBERS
  //
  ////////////////////////////////////////////////////////////////////////////  
 
  private JRadioButton useInches_;
  private JRadioButton useCm_;
  private JCheckBox keepAspect_;
  private JTextField pubWidth_;
  private JTextField pubHeight_;
  private JTextField pubZoom_;  
  private JComboBox pubResolution_;
  private JLabel pubWidthUnits_;
  private JLabel pubHeightUnits_;
  private JLabel pubResUnits_;
  private int currentUnits_;
  private JComboBox pubCombo_;
  private boolean processing_;
  private Settings currSettings_;
  
  private Settings origSettings_;
  private int origUnits_;
  private String origFormat_;
  private int origResIndex_;
  
  private List resList_;
  private String currentFormat_;
  private ExportSettingsDialog.ExportSettings finishedSettings_;  
  private Preferences prefs_; 
    
  ////////////////////////////////////////////////////////////////////////////
  //
  // PUBLIC CONSTRUCTORS
  //
  ////////////////////////////////////////////////////////////////////////////    

  /***************************************************************************
  **
  ** Constructor 
  */ 
  
  public ExportSettingsPublishDialog(JFrame parent, int baseWidth, int baseHeight) {     
    super(parent, ResourceManager.getManager().getString("exportDialog.title"), true);
    
    ResourceManager rMan = ResourceManager.getManager();
    setSize(400, 350);
    JPanel cp = (JPanel)getContentPane();
    cp.setBorder(new EmptyBorder(20, 20, 20, 20));
    cp.setLayout(new GridBagLayout());
    GridBagConstraints gbc = new GridBagConstraints();
    
    finishedSettings_ = null;
    currSettings_ = new Settings(baseHeight, baseWidth);    
    prefs_ = Preferences.userNodeForPackage(this.getClass());
    currSettings_.zoom = prefs_.getDouble("ExportPublishZoom", 1.0);
    
    List types = ImageExporter.getSupportedExports();
    String maybeFormat = prefs_.get("ExportPublishFormat", "TIFF");
    if (types.contains(maybeFormat)) {
      currentFormat_ = maybeFormat;
    } else if (types.contains("PNG")) {
      currentFormat_ = "PNG";
    } else {
      currentFormat_ = (String)types.get(0);
    }
    origFormat_ = currentFormat_;
    resList_ = ImageExporter.getSupportedResolutions(true);

    JPanel pubPanel = buildPublishPanel(baseWidth, baseHeight, types);
    origSettings_ = new Settings(currSettings_);
    origUnits_ = currentUnits_;
    origResIndex_ = pubResolution_.getSelectedIndex();
    
    UiUtil.gbcSet(gbc, 0, 0, 10, 8, UiUtil.BO, 0, 0, 5, 5, 5, 5, UiUtil.CEN, 1.0, 1.0);    
    cp.add(pubPanel, gbc);    

    //
    // Build the button panel:
    //
    
    FixedJButton buttonR = new FixedJButton(rMan.getString("dialogs.reset"));
    buttonR.addActionListener(new ActionListener() {
      public void actionPerformed(ActionEvent ev) {
        try {
          reset();
        } catch (Exception ex) {
          ExceptionHandler.getHandler().displayException(ex);
        }
      }
    });    
    
    FixedJButton buttonO = new FixedJButton(rMan.getString("exportDialog.ok"));
    buttonO.addActionListener(new ActionListener() {
      public void actionPerformed(ActionEvent ev) {
        try {
          if (extractProperties()) {
            ExportSettingsPublishDialog.this.setVisible(false);
            ExportSettingsPublishDialog.this.dispose();
          }
        } catch (Exception ex) {
          ExceptionHandler.getHandler().displayException(ex);
        }
      }
    });     
    FixedJButton buttonC = new FixedJButton(rMan.getString("exportDialog.cancel"));
    buttonC.addActionListener(new ActionListener() {
      public void actionPerformed(ActionEvent ev) {
        try {
          ExportSettingsPublishDialog.this.setVisible(false);
          ExportSettingsPublishDialog.this.dispose();
        } catch (Exception ex) {
          ExceptionHandler.getHandler().displayException(ex);
        }
      }
    });
    Box buttonPanel = Box.createHorizontalBox();
    buttonPanel.add(Box.createHorizontalStrut(10));    
    buttonPanel.add(buttonR);    
    buttonPanel.add(Box.createHorizontalGlue()); 
    buttonPanel.add(buttonO);
    buttonPanel.add(Box.createHorizontalStrut(10));    
    buttonPanel.add(buttonC);    

    //
    // Build the dialog:
    //
    
    UiUtil.gbcSet(gbc, 0, 8, 10, 1, UiUtil.HOR, 0, 0, 5, 5, 5, 5, UiUtil.SE, 1.0, 0.0);
    cp.add(buttonPanel, gbc);    
    setLocationRelativeTo(parent);
  }
  
  /***************************************************************************
  **
  ** Get the dialog results.  Will be null on cancel.
  */
  
  public ExportSettingsDialog.ExportSettings getResults() {
    return (finishedSettings_);
  }
    
  ////////////////////////////////////////////////////////////////////////////
  //
  // INNER CLASSES
  //
  ////////////////////////////////////////////////////////////////////////////  
 
  ////////////////////////////////////////////////////////////////////////////
  //
  // PRIVATE METHODS
  //
  ////////////////////////////////////////////////////////////////////////////

  /***************************************************************************
  **
  ** Process a new publish value
  */
  
  private void reset() {
    
    if (processing_) {
      return;
    }
    processing_ = true; 
    try {
      currentUnits_ = origUnits_;
      useInches_.setSelected(currentUnits_ == ImageExporter.INCHES);
      useCm_.setSelected(currentUnits_ == ImageExporter.CM);      
      updateUnits();
      pubCombo_.setSelectedItem(origFormat_);
      currSettings_.merge(origSettings_);
      pubWidth_.setText(Double.toString(currSettings_.width));
      pubHeight_.setText(Double.toString(currSettings_.height));
      pubZoom_.setText(Double.toString(currSettings_.zoom));
      keepAspect_.setSelected(currSettings_.aspectFixed);
      pubResolution_.setSelectedIndex(origResIndex_);
    } finally {
      processing_ = false;
    }
    return;
  }
  
  /***************************************************************************
  **
  ** Process a new publish value
  */
  
  private void processPublishVal(boolean force, int whichVal) {

    //
    // While typing, blank fields are OK.  If that happens, we don't update anything else.
    // Bad or negative results give a beep, and we don't update anything either.  During
    // a force, bad results mean we replace the bad field with the current valid value,
    // as well as a beep.
    //
    
    if (processing_) {
      return;
    }
    processing_ = true;    
    try {
      double min = (currentUnits_ == ImageExporter.INCHES) ? MIN_SIZE_INCH_ : MIN_SIZE_CM_;
      double max = (currentUnits_ == ImageExporter.INCHES) ? MAX_SIZE_INCH_ : MAX_SIZE_CM_;      

      boolean fixVal = false;

      double newZoom = currSettings_.zoom;
      if (whichVal == ZOOM_) {
        String zoomText = pubZoom_.getText().trim();
        boolean haveZoom = false;
        if (!zoomText.equals("") && !zoomText.equals(".")) {
          try {
            double parsedZoom = Double.parseDouble(zoomText);
            if (parsedZoom > 0.0) {
              newZoom = parsedZoom;
              haveZoom = true;
            } else if (parsedZoom == 0) {
              if (force) {
                Toolkit.getDefaultToolkit().beep();
              }
            } else {
              Toolkit.getDefaultToolkit().beep();
            }
            if (haveZoom && tooBigOrSmall(newZoom, ZOOM_)) {
              newZoom = currSettings_.zoom;
              haveZoom = false;
              Toolkit.getDefaultToolkit().beep();
            }
            if (haveZoom && (parsedZoom != rounded(parsedZoom, ZOOM_MULT_))) {
                            // Note!  Cannot do a fixVal during a cursor notification!
              newZoom = currSettings_.zoom;
              haveZoom = false;
              Toolkit.getDefaultToolkit().beep();
            }             
            
          } catch (NumberFormatException nfe) {
            Toolkit.getDefaultToolkit().beep();
          }
        } else if (force) {
          Toolkit.getDefaultToolkit().beep();  
        }

        if (force && !haveZoom) {
          fixVal = true;
        }
      }    

      double newHeight = currSettings_.height;
      if (whichVal == HEIGHT_) {
        String heightText = pubHeight_.getText().trim();
        boolean haveHeight = false;
        if (!heightText.equals("") && !heightText.equals(".")) {
          try {
            double parsedHeight = Double.parseDouble(heightText);
            if ((parsedHeight >= min) && (parsedHeight <= max)) {
              newHeight = parsedHeight;
              haveHeight = true;
            } else if (parsedHeight == 0) {
              if (force) {
                Toolkit.getDefaultToolkit().beep();
              }
            } else {
              Toolkit.getDefaultToolkit().beep();
            }
            if (haveHeight && tooBigOrSmall(newHeight, HEIGHT_)) {
              newHeight = currSettings_.height;
              haveHeight = false;
              Toolkit.getDefaultToolkit().beep();
            }
            if (haveHeight && (parsedHeight != rounded(parsedHeight, DIM_MULT_))) {
              // Note!  Cannot do a fixVal during a cursor notification!
              newHeight = currSettings_.height;
              haveHeight = false;
              Toolkit.getDefaultToolkit().beep();
            }          
          } catch (NumberFormatException nfe) {
            Toolkit.getDefaultToolkit().beep();
          }
        } else if (force) {
          Toolkit.getDefaultToolkit().beep();  
        }
        if (force && !haveHeight) {
          fixVal = true;
        }
      }    

      double newWidth = currSettings_.width;
      if (whichVal == WIDTH_) {
        String widthText = pubWidth_.getText().trim();
        boolean haveWidth = false;
        if (!widthText.equals("") && !widthText.equals(".")) {
          try {
            double parsedWidth = Double.parseDouble(widthText);
            if ((parsedWidth >= min) && (parsedWidth <= max)) {
              newWidth = parsedWidth;
              haveWidth = true;
            } else if (parsedWidth == 0) {
              if (force) {
                Toolkit.getDefaultToolkit().beep();
              }
            } else {
              Toolkit.getDefaultToolkit().beep();
            }
            if (haveWidth && tooBigOrSmall(newWidth, WIDTH_)) {
              newWidth = currSettings_.width;
              haveWidth = false;
              Toolkit.getDefaultToolkit().beep();
            }
            if (haveWidth && (parsedWidth != rounded(parsedWidth, DIM_MULT_))) {
              // Note!  Cannot do a fixVal during a cursor notification!
              newWidth = currSettings_.width;
              haveWidth = false;
              Toolkit.getDefaultToolkit().beep();
            }          
          } catch (NumberFormatException nfe) {
            Toolkit.getDefaultToolkit().beep();
          }
        } else if (force) {
          Toolkit.getDefaultToolkit().beep();  
        }
        if (force && !haveWidth) {
          fixVal = true;
        }
      }

      boolean newAspect = currSettings_.aspectFixed;
      if (whichVal == ASPECT_) {
        newAspect = keepAspect_.isSelected();
      }

      long newRes = currSettings_.resolution;
      if (whichVal == RES_) {
        newRes = resolutionForIndex(pubResolution_.getSelectedIndex());
      }

      //
      // Now go and set fields:
      //
      
      setFields(whichVal, newZoom, newHeight, newWidth, newAspect, newRes, fixVal); 
    } finally {
      processing_ = false;
    }
    return;
  } 
  
  /***************************************************************************
  **
  ** Set fields
  */
  
  private void setFields(int whichVal, double newZoom, double newHeight, double newWidth,
                         boolean newAspect, long newRes, boolean fixVal) {        

    //
    // Now go and set fields:
    //
                           
    Settings settings = new Settings(currSettings_);
    
    switch (whichVal) { 
      case ZOOM_:
        settings.changeZoom(newZoom);
        if (settings.zoomChanged(currSettings_)) {
          currSettings_.mergeZoom(settings);
        }
        if (settings.widthChanged(currSettings_)) {
          currSettings_.mergeWidth(settings);
          pubWidth_.setText(Double.toString(currSettings_.width)); 
        }
        if (settings.heightChanged(currSettings_)) {
          currSettings_.mergeHeight(settings);
          pubHeight_.setText(Double.toString(currSettings_.height)); 
        }        
        if (fixVal) {
          pubZoom_.setText(Double.toString(currSettings_.zoom));
        }
        break;      
      case HEIGHT_:
        settings.changeHeight(newHeight);
        if (settings.heightChanged(currSettings_)) {
          currSettings_.mergeHeight(settings);
        }
        if (settings.widthChanged(currSettings_)) {
          currSettings_.mergeWidth(settings);
          pubWidth_.setText(Double.toString(currSettings_.width)); 
        }
        if (settings.zoomChanged(currSettings_)) {
          currSettings_.mergeZoom(settings);
          pubZoom_.setText(Double.toString(currSettings_.zoom)); 
        }       
        if (fixVal) {
          pubHeight_.setText(Double.toString(currSettings_.height));
        }
        break;        
      case WIDTH_:
        settings.changeWidth(newWidth);
        if (settings.widthChanged(currSettings_)) {
          currSettings_.mergeWidth(settings);
        }
        currSettings_.mergeWidth(settings);
        if (settings.heightChanged(currSettings_)) {
          currSettings_.mergeHeight(settings);
          pubHeight_.setText(Double.toString(currSettings_.height)); 
        }
        if (settings.zoomChanged(currSettings_)) {
          currSettings_.mergeZoom(settings);
          pubZoom_.setText(Double.toString(currSettings_.zoom)); 
        }       
        if (fixVal) {
          pubWidth_.setText(Double.toString(currSettings_.width));
        }
        break;     
      case ASPECT_:
        settings.changeAspect(newAspect);
        if (settings.aspectChanged(currSettings_)) {
          currSettings_.mergeAspect(settings);
        }
        if (settings.heightChanged(currSettings_)) {
          currSettings_.mergeHeight(settings);
          pubHeight_.setText(Double.toString(currSettings_.height)); 
        }
        if (settings.widthChanged(currSettings_)) {
          currSettings_.mergeWidth(settings);
          pubWidth_.setText(Double.toString(currSettings_.width)); 
        }        
        break;
      case RES_:
        settings.changeResolution(newRes);
        if (settings.resolutionChanged(currSettings_)) {
          currSettings_.mergeResolution(settings);
        }
        if (settings.zoomChanged(currSettings_)) {
          currSettings_.mergeZoom(settings);
          pubZoom_.setText(Double.toString(currSettings_.zoom)); 
        } 
        break;        
      default:
        throw new IllegalStateException();
    }
    return;
  } 
  
  /***************************************************************************
  **
  ** fix publication values
  */
  
  private void fixPubVals() {    
    if (processing_) {
      return;
    }
    processing_ = true;
    try {
      currSettings_.roundVals();
      pubWidth_.setText(Double.toString(currSettings_.width));
      pubHeight_.setText(Double.toString(currSettings_.height));
      pubZoom_.setText(Double.toString(currSettings_.zoom));
    } finally {
      processing_ = false;
    }
    return;
  }  
 
  /***************************************************************************
  **
  ** Report if a value is too small
  */
  
  private boolean tooBigOrSmall(double newVal, int whichVal) {
    //
    // If we have a fixed aspect ratio and a dimension setting forces
    // the other to zero, it is too small:
    //
       
    double min = (currentUnits_ == ImageExporter.INCHES) ? MIN_SIZE_INCH_ : MIN_SIZE_CM_;
    double max = (currentUnits_ == ImageExporter.INCHES) ? MAX_SIZE_INCH_ : MAX_SIZE_CM_;    
    Settings settings = new Settings(currSettings_);
    
    switch (whichVal) {
      case ZOOM_:
        settings.changeZoom(newVal);
        return ((settings.width < min) || (settings.height < min) || 
                (settings.width > max) || (settings.height > max));
      case HEIGHT_:
        settings.changeHeight(newVal);
        return ((settings.width < min) || (settings.width > max) || (settings.zoom == 0.0));
      case WIDTH_:
        settings.changeWidth(newVal);
        return ((settings.height < min) || (settings.height > max ) || (settings.zoom == 0.0)); 
      default:
        throw new IllegalStateException();
    }
  }    
  
  /***************************************************************************
  **
  ** Build publish panel
  ** 
  */
  
  private JPanel buildPublishPanel(int baseWidth, int baseHeight, List types) {
    JPanel retval = new JPanel();
    retval.setLayout(new GridBagLayout());
    GridBagConstraints gbc = new GridBagConstraints();     
    ResourceManager rMan = ResourceManager.getManager();
    currentUnits_ = prefs_.getInt("ExportPublishUnits", ImageExporter.INCHES);
    currSettings_.aspectFixed = prefs_.getBoolean("ExportPublishAspect", true);

    String currUnits = rMan.getString((currentUnits_ == ImageExporter.INCHES) ? "exportDialog.inches" : "exportDialog.cm");
    
    int rowNum = addUnitChoices(retval, gbc, rMan);

        
    JLabel label = new JLabel(rMan.getString("exportDialog.pubSize"));
    JLabel wLab = new JLabel(rMan.getString("exportDialog.width"));
    pubWidth_ = new JTextField();
    pubWidthUnits_ = new JLabel(rMan.getString(currUnits));
    JLabel hLab = new JLabel(rMan.getString("exportDialog.height"));
    pubHeight_ = new JTextField();
    pubHeightUnits_ = new JLabel(rMan.getString(currUnits));
    JLabel zLab = new JLabel(rMan.getString("exportDialog.zoom"));
    pubZoom_ = new JTextField();
    
    keepAspect_ = new JCheckBox(rMan.getString("exportDialog.keepAspect"), currSettings_.aspectFixed);
    
    //
    // Resolution choices
    //
    
    JLabel reslabel = new JLabel(rMan.getString("exportDialog.resolution"));
    pubResolution_ = new JComboBox(buildResolutions());
    String savedResolution = prefs_.get("ExportPublishRes", "0");
    if (!savedResolution.equals("0")) {
      pubResolution_.setSelectedItem(savedResolution);
    }
    
    String prStr = rMan.getString((currentUnits_ == ImageExporter.INCHES) ? "exportDialog.dpi" : "exportDialog.dpc");
    pubResUnits_ = new JLabel(rMan.getString(prStr)); 
    currSettings_.resolution = resolutionForIndex(pubResolution_.getSelectedIndex());
    
    //
    // The relevant saved value is the zoom factor, and the height and width are
    // derived:
    //
    
    currSettings_.initDims();
    pubZoom_.setText(Double.toString(currSettings_.zoom));    
    pubWidth_.setText(Double.toString(currSettings_.width));
    pubHeight_.setText(Double.toString(currSettings_.height));
    
    //
    // Now add listeners:
    //

    PublishListener pl = new PublishListener(ZOOM_);
    pubZoom_.addActionListener(pl);
    pubZoom_.addCaretListener(pl);
    pubZoom_.addFocusListener(pl);    
    
    PublishListener pl1 = new PublishListener(WIDTH_);
    pubWidth_.addActionListener(pl1);
    pubWidth_.addCaretListener(pl1);
    pubWidth_.addFocusListener(pl1);
    
    PublishListener pl2 = new PublishListener(HEIGHT_);
    pubHeight_.addActionListener(pl2);
    pubHeight_.addCaretListener(pl2);
    pubHeight_.addFocusListener(pl2);
    
    keepAspect_.addActionListener(new ActionListener() {
      public void actionPerformed(ActionEvent ev) {
        try {
          processPublishVal(false, ASPECT_);
        } catch (Exception ex) {
          ExceptionHandler.getHandler().displayException(ex);
        }
      }
    });
    
    pubResolution_.addActionListener(new ActionListener() {
      public void actionPerformed(ActionEvent ev) {
        try {
          processPublishVal(false, RES_);
        } catch (Exception ex) {
          ExceptionHandler.getHandler().displayException(ex);
        }
      }
    });
    
    UiUtil.gbcSet(gbc, 0, rowNum, 3, 1, UiUtil.BO, 0, 0, 5, 5, 5, 5, UiUtil.E, 1.0, 1.0);           
    retval.add(reslabel, gbc);
    UiUtil.gbcSet(gbc, 3, rowNum, 2, 1, UiUtil.HOR, 0, 0, 5, 5, 5, 5, UiUtil.CEN, 0.0, 1.0);    
    retval.add(pubResolution_, gbc);    
    UiUtil.gbcSet(gbc, 5, rowNum++, 1, 1, UiUtil.BO, 0, 0, 5, 5, 5, 5, UiUtil.W, 0.0, 1.0);    
    retval.add(pubResUnits_, gbc);
    UiUtil.gbcSet(gbc, 3, rowNum++, 3, 1, UiUtil.BO, 0, 0, 5, 5, 5, 5, UiUtil.W, 0.0, 1.0);    
    retval.add(keepAspect_, gbc);     
    UiUtil.gbcSet(gbc, 0, rowNum, 3, 1, UiUtil.BO, 0, 0, 5, 5, 5, 5, UiUtil.E, 1.0, 1.0);           
    retval.add(label, gbc);
    UiUtil.gbcSet(gbc, 3, rowNum, 1, 1, UiUtil.BO, 0, 0, 5, 5, 5, 5, UiUtil.E, 0.0, 1.0);    
    retval.add(wLab, gbc); 
    UiUtil.gbcSet(gbc, 4, rowNum, 1, 1, UiUtil.HOR, 0, 0, 5, 5, 5, 5, UiUtil.CEN, 0.5, 1.0);    
    retval.add(pubWidth_, gbc);      
    UiUtil.gbcSet(gbc, 5, rowNum++, 1, 1, UiUtil.BO, 0, 0, 5, 5, 5, 5, UiUtil.W, 0.0, 1.0);    
    retval.add(pubWidthUnits_, gbc);     
    UiUtil.gbcSet(gbc, 3, rowNum, 1, 1, UiUtil.BO, 0, 0, 5, 5, 5, 5, UiUtil.E, 0.0, 1.0);    
    retval.add(hLab, gbc); 
    UiUtil.gbcSet(gbc, 4, rowNum, 1, 1, UiUtil.HOR, 0, 0, 5, 5, 5, 5, UiUtil.CEN, 0.5, 1.0);    
    retval.add(pubHeight_, gbc);      
    UiUtil.gbcSet(gbc, 5, rowNum++, 1, 1, UiUtil.BO, 0, 0, 5, 5, 5, 5, UiUtil.W, 0.0, 1.0);    
    retval.add(pubHeightUnits_, gbc);
    UiUtil.gbcSet(gbc, 3, rowNum, 1, 1, UiUtil.BO, 0, 0, 5, 5, 5, 5, UiUtil.E, 0.0, 1.0);    
    retval.add(zLab, gbc); 
    UiUtil.gbcSet(gbc, 4, rowNum++, 2, 1, UiUtil.HOR, 0, 0, 5, 5, 5, 5, UiUtil.CEN, 0.5, 1.0);    
    retval.add(pubZoom_, gbc);          
    
     
    //
    // Format choices
    //
    
    JLabel flabel = new JLabel(rMan.getString("exportDialog.format"));
    pubCombo_ = new JComboBox(types.toArray(new String[types.size()]));
    pubCombo_.setSelectedItem(currentFormat_);
    Object currCombo = pubCombo_.getSelectedItem();
    if ((currCombo == null) || (!currCombo.equals(currentFormat_))) {
      throw new IllegalArgumentException();
    }
    UiUtil.gbcSet(gbc, 0, rowNum, 3, 1, UiUtil.BO, 0, 0, 5, 5, 5, 5, UiUtil.E, 1.0, 1.0);           
    retval.add(flabel, gbc);
    UiUtil.gbcSet(gbc, 3, rowNum, 2, 1, UiUtil.HOR, 0, 0, 5, 5, 5, 5, UiUtil.CEN, 0.0, 1.0);    
    retval.add(pubCombo_, gbc);
    
    return (retval);
  }     

  /***************************************************************************
  **
  ** Add the unit choices to the panel
  */
  
  private int addUnitChoices(JPanel retval, GridBagConstraints gbc, ResourceManager rMan) {
     
    String inches = rMan.getString("exportDialog.inches");
    useInches_ = new JRadioButton(inches, (currentUnits_ == ImageExporter.INCHES));
    useInches_.addActionListener(new ActionListener() {
      public void actionPerformed(ActionEvent ev) {
        try {
          if (currentUnits_ != ImageExporter.INCHES) {
            currentUnits_ = ImageExporter.INCHES;
            updateUnits();
            validate();            
          }
        } catch (Exception ex) {
          ExceptionHandler.getHandler().displayException(ex);
        }
      }
    });         
        
    String cm = rMan.getString("exportDialog.cm");
    useCm_ = new JRadioButton(cm, (currentUnits_ == ImageExporter.CM));
    useCm_.addActionListener(new ActionListener() {
      public void actionPerformed(ActionEvent ev) {
        try {
          if (currentUnits_ != ImageExporter.CM) {
            currentUnits_ = ImageExporter.CM;
            updateUnits();
            validate();
          }
        } catch (Exception ex) {
          ExceptionHandler.getHandler().displayException(ex);
        }
      }
    });
    
    ButtonGroup group = new ButtonGroup();
    group.add(useInches_);
    group.add(useCm_);
    
    JLabel unitLabel = new JLabel(rMan.getString("exportDialog.useUnits"));
    UiUtil.gbcSet(gbc, 0, 0, 2, 1, UiUtil.BO, 0, 0, 5, 5, 5, 5, UiUtil.E, 1.0, 1.0);    
    retval.add(unitLabel, gbc);    
    UiUtil.gbcSet(gbc, 2, 0, 2, 1, UiUtil.BO, 0, 0, 5, 5, 5, 5, UiUtil.CEN, 0.0, 0.0);    
    retval.add(useInches_, gbc);
    UiUtil.gbcSet(gbc, 4, 0, 2, 1, UiUtil.BO, 0, 0, 5, 5, 5, 5, UiUtil.CEN, 0.0, 0.0);        
    retval.add(useCm_, gbc);
    return (1);
  }
  
  /***************************************************************************
  **
  ** Change the displayed units
  ** 
  */
  
  private void updateUnits() {
    ResourceManager rMan = ResourceManager.getManager();    
    String currUnits = rMan.getString((currentUnits_ == ImageExporter.INCHES) ? "exportDialog.inches" : "exportDialog.cm");
    pubWidthUnits_.setText(currUnits);
    pubHeightUnits_.setText(currUnits);
    pubResUnits_.setText(currUnits);    
    String prStr = rMan.getString((currentUnits_ == ImageExporter.INCHES) ? "exportDialog.dpi" : "exportDialog.dpc");
    pubResUnits_.setText(rMan.getString(prStr));    

    processing_ = true;
    try {
      int currRes = pubResolution_.getSelectedIndex();
      pubResolution_.removeAllItems();
      String[] res = buildResolutions();
      for (int i = 0; i < res.length; i++) {
        pubResolution_.addItem(res[i]);
      }
      pubResolution_.setSelectedIndex(currRes);
      currSettings_.changeUnits(resolutionForIndex(currRes), currentUnits_);

      pubWidth_.setText(Double.toString(currSettings_.width));
      pubHeight_.setText(Double.toString(currSettings_.height));
    } finally {
      processing_ = false;
    }
    return;
  }
  
  /***************************************************************************
  **
  ** Get a current resolution list
  */
  
  private String[] buildResolutions() {
    int numRes = resList_.size();
    String[] retval = new String[numRes];
    for (int i = 0; i < numRes; i++) {
      Object[] resVal = (Object[])resList_.get(i);
      retval[i] = Double.toString(((ImageExporter.RationalNumber)resVal[currentUnits_]).value);
      double res = ((ImageExporter.RationalNumber)resVal[currentUnits_]).value;
      retval[i] = Long.toString(Math.round(res));
    }
    return (retval);
  } 
  
  /***************************************************************************
  **
  ** Get the resolution for an index
  */
  
  private long resolutionForIndex(int i) {
    Object[] resVal = (Object[])resList_.get(i);
    return (Math.round(((ImageExporter.RationalNumber)resVal[currentUnits_]).value));
  } 
     
  /***************************************************************************
  **
  ** Get ready for export
  ** 
  */
  
  private boolean extractProperties() {
    
    long pixHeight = currSettings_.getPixHeight();
    long pixWidth = currSettings_.getPixWidth();
    
    //
    // Get the dimensions:
    //
    
    long numPix = pixHeight * pixWidth;
    if (numPix > ExportSettingsDialog.HUGE_PIC) {
      ResourceManager rMan = ResourceManager.getManager();
      String desc = MessageFormat.format(rMan.getString("export.confirmBigExport"), 
                                         new Object[] {new Long(numPix)});
      desc = UiUtil.convertMessageToHtml(desc);                                         
      int result = JOptionPane.showConfirmDialog(this, desc,
                                                 rMan.getString("export.confirmBigExportTitle"),
                                                 JOptionPane.YES_NO_OPTION);
      if (result != 0) {
        return (false);
      }
    }    
    
    finishedSettings_ = new ExportSettingsDialog.ExportSettings();

    finishedSettings_.formatType = (String)pubCombo_.getSelectedItem();
 
    //
    // Get the resolution:
    //

    int resIndex = pubResolution_.getSelectedIndex();
    Object[] resVal = (Object[])resList_.get(resIndex);
    ImageExporter.RationalNumber ratNum = (ImageExporter.RationalNumber)resVal[currentUnits_];
    
    finishedSettings_.res = new ImageExporter.ResolutionSettings();
    finishedSettings_.res.dotsPerUnit = ratNum;
    finishedSettings_.res.units = currentUnits_;    
    
    //
    // Zoom 
    //
    
    finishedSettings_.zoomVal = currSettings_.zoom;
    
    //
    // Get the dimensions:
    //
    
    finishedSettings_.size = new Dimension((int)pixWidth, (int)pixHeight);

    //
    // Save the preferences:
    //
    
    prefs_.putInt("ExportPublishUnits", currentUnits_);
    prefs_.putBoolean("ExportPublishAspect", currSettings_.aspectFixed);
    prefs_.putDouble("ExportPublishZoom", currSettings_.zoom);
    prefs_.put("ExportPublishRes", (String)pubResolution_.getSelectedItem());
    prefs_.put("ExportPublishFormat", finishedSettings_.formatType);   
    return (true);
  }
  
  ////////////////////////////////////////////////////////////////////////////
  //
  // STATIC METHODS
  //
  ////////////////////////////////////////////////////////////////////////////    

  /***************************************************************************
  **
  ** Round to 2 decimal places
  */
  
  static double rounded(double val, double mult) {
    double retval = (double)Math.round(val * mult);
    // Note!  Division by 100 has more numerical accuracy than multiplying by .01
    // (the latter cannot be represented precisely in binary...)
    return (retval / mult);
  }
    
  ////////////////////////////////////////////////////////////////////////////
  //
  // PRIVATE INNER CLASSES
  //
  ////////////////////////////////////////////////////////////////////////////  
 
  /***************************************************************************
  **
  ** For handling zoom textField changes
  ** 
  */
  
  private class PublishListener implements ActionListener, CaretListener, FocusListener {
    
    private int whichVal_;
    
    PublishListener(int whichVal) {
      whichVal_ = whichVal;
    }
    
    public void actionPerformed(ActionEvent evt) {
      try {
        processPublishVal(true, whichVal_);
      } catch (Exception ex) {
        ExceptionHandler.getHandler().displayException(ex);
      }
    }
    public void caretUpdate(CaretEvent evt) {
      try {
        processPublishVal(false, whichVal_);
      } catch (Exception ex) {
        ExceptionHandler.getHandler().displayException(ex);
      }
    }
    public void focusGained(FocusEvent evt) {
    }    
    public void focusLost(FocusEvent evt) {
      try {
        fixPubVals();
      } catch (Exception ex) {
        ExceptionHandler.getHandler().displayException(ex);
      }
    }        
  }
  
  /***************************************************************************
  **
  ** Degrees of Freedom
  ** 
  */
  
  private static class Settings {
    public long resolution;
    public double height;
    public double width; 
    public boolean aspectFixed;
    public double zoom;
    
    private double startAspect_;
    private int startHeight_;
    private int startWidth_;

    Settings(int baseHeight, int baseWidth) {    
      startHeight_ = baseHeight;
      startWidth_ = baseWidth;
      startAspect_ = (double)baseWidth / (double)baseHeight;
    }
    
    Settings(long resolution, double height, double width, 
             boolean aspectFixed, double zoom, 
             int baseHeight, int baseWidth, double baseAspect) {
    
      this.resolution = resolution;
      this.height = height;
      this.width = width;
      this.aspectFixed = aspectFixed;
      this.zoom = zoom;
    
      startHeight_ = baseHeight;
      startWidth_ = baseWidth;
      startAspect_ = baseAspect;
    }
    
    Settings() {
    }
    
    Settings(Settings other) {
      merge(other);
    }
    
    void merge(Settings other) {    
      this.resolution = other.resolution;
      this.height = other.height;
      this.width = other.width;
      this.aspectFixed = other.aspectFixed;
      this.zoom = other.zoom;
    
      startHeight_ = other.startHeight_;
      startWidth_ = other.startWidth_;
      startAspect_ = other.startAspect_;
      return;
    }    
    
    void initDims() {
      height = rounded((zoom * (double)startHeight_) / (double)resolution, 100.0);
      width = rounded((zoom * (double)startWidth_) / (double)resolution, DIM_MULT_);    
      return;
    }
    
    void roundVals() {
      height = rounded(height, DIM_MULT_);
      width = rounded(width, DIM_MULT_);
      zoom = rounded(zoom, ZOOM_MULT_);
      return;
    }
    
    void changeUnits(long res, int units) {
      resolution = res;
      double convert = (units == ImageExporter.INCHES) ? 1.0 / CM_PER_INCH_ : CM_PER_INCH_;
    
      height *= convert;
      width *= convert;
      height = rounded(height, DIM_MULT_);
      width = rounded(width, DIM_MULT_);    
      return;
    }
    
    long getPixHeight() {
      return (Math.round(height * resolution));
    }
    
    long getPixWidth() {
      return (Math.round(width * resolution));    
    }
        
    boolean resolutionChanged(Settings old) {
      return (resolution != old.resolution);
    }

    boolean heightChanged(Settings old) {
      return (height != old.height);
    }
    
    boolean widthChanged(Settings old) {
      return (width != old.width);
    }
    
    boolean aspectChanged(Settings old) {
      return (aspectFixed != old.aspectFixed);
    }
    
    boolean zoomChanged(Settings old) {
      return (zoom != old.zoom);
    }
    
    void mergeResolution(Settings src) {
      resolution = src.resolution;
      return;
    }

    void mergeHeight(Settings src) {
      height = src.height;
      return;      
    }
    
    void mergeWidth(Settings src) {
      width = src.width;
      return;      
    }
    
    void mergeAspect(Settings src) {
      aspectFixed = src.aspectFixed;
      return;      
    }
    
    void mergeZoom(Settings src) {
      zoom = src.zoom;
      return;      
    }     
    
    double calcZoomFromFixedDims() {
      long pixHeight = Math.round(height * resolution);   
      double zoomValForH = rounded((double)pixHeight / (double)startHeight_, ZOOM_MULT_);
    
      // Take the smaller of the two zooms.  If aspect ratio is not fixed, we will end
      // up padding the other dimension
    
      if (aspectFixed) {
        return (zoomValForH);
      } else {
        long pixWidth = Math.round(width * resolution);      
        double zoomValForW = rounded((double)pixWidth / (double)startWidth_, ZOOM_MULT_);      
        return ((zoomValForH > zoomValForW) ? zoomValForW : zoomValForH);
      }
    }
    
    void changeZoom(double newZoom) {
      if (zoom != newZoom) {
        zoom = newZoom;
        double oldAspect = (double)width / (double)height;
        if (aspectFixed || (oldAspect == startAspect_)) {
          height = rounded(((double)startHeight_ * zoom) / (double)resolution, DIM_MULT_);
          width = rounded(((double)startWidth_ * zoom) / (double)resolution, DIM_MULT_);
        } else if (oldAspect > startAspect_) {  // Width is currently padded
          height = rounded(((double)startHeight_ * zoom) / (double)resolution, DIM_MULT_);
          width = rounded(height * oldAspect, DIM_MULT_);
        } else { // Height is currently padded
          width = rounded(((double)startWidth_ * zoom) / (double)resolution, DIM_MULT_);
          height = rounded(width / oldAspect, DIM_MULT_);             
        }
      }
      return;
    }
    
    void changeHeight(double newHeight) {    
      if (height != newHeight) {
        double oldAspect = (double)width / (double)height;
        height = newHeight;
        if (aspectFixed) {  // Zoom, width both change
          long pixHeight = Math.round(height * resolution);
          zoom = rounded((double)pixHeight / (double)startHeight_, ZOOM_MULT_);
          width = rounded(((double)startWidth_ * zoom) / (double)resolution, DIM_MULT_);
        } else {  // width will stay fixed, zoom MAY change
          zoom = calcZoomFromFixedDims();
        }
      }
      return;
    }
    
    void changeWidth(double newWidth) {        
      if (width != newWidth) {
        double oldAspect = (double)width / (double)height;
        width = newWidth;
        if (aspectFixed) {  // Zoom, height both change
          long pixWidth = Math.round(width * resolution);
          zoom = rounded((double)pixWidth / (double)startWidth_, ZOOM_MULT_);
          height = rounded(((double)startHeight_ * zoom) / (double)resolution, DIM_MULT_);
        } else {  // height will stay fixed, zoom MAY change
          zoom = calcZoomFromFixedDims();
        }
      }
      return;
    }
    
    void changeAspect(boolean newAspect) {
      if (aspectFixed != newAspect) {
        aspectFixed = newAspect;
        if (aspectFixed) {
          // Change the one that will become bigger:
          double possibleWidth = rounded(height * startAspect_, DIM_MULT_);
          if (possibleWidth > width) {
            width = possibleWidth;
          } else {
            height = rounded(width / startAspect_, DIM_MULT_);
          }
        }
      }
      return;
    }
    
    void changeResolution(long newRes) {
      if (resolution != newRes) {
        resolution = newRes;
        // Changing resolution changes zoom
        zoom = calcZoomFromFixedDims();
      }
      return;
    }
  }   
}
```
