## Supplementary material for "BioFabric Visualization of Network Alignments": BioFabric Source Code: FabricDisplayOptionsDialog.java

```
/*
**    Copyright (C) 2003-2018 Institute for Systems Biology 
**                            Seattle, Washington, USA. 
**
**    This library is free software; you can redistribute it and/or
**    modify it under the terms of the GNU Lesser General Public
**    License as published by the Free Software Foundation; either
**    version 2.1 of the License, or (at your option) any later version.
**
**    This library is distributed in the hope that it will be useful,
**    but WITHOUT ANY WARRANTY; without even the implied warranty of
**    MERCHANTABILITY or FITNESS FOR A PARTICULAR PURPOSE.  See the GNU
**    Lesser General Public License for more details.
**
**    You should have received a copy of the GNU Lesser General Public
**    License along with this library; if not, write to the Free Software
**    Foundation, Inc., 59 Temple Place, Suite 330, Boston, MA  02111-1307  USA
*/


package org.systemsbiology.biofabric.ui.dialogs;

import java.awt.Dimension;
import java.awt.event.ActionListener;
import java.awt.event.ActionEvent;
import java.net.MalformedURLException;
import java.net.URL;

import javax.swing.JCheckBox;
import javax.swing.JComboBox;
import javax.swing.JFrame;
import javax.swing.JLabel;
import javax.swing.JOptionPane;
import javax.swing.JTextField;

import org.systemsbiology.biofabric.api.dialog.BTStashResultsDialog;
import org.systemsbiology.biofabric.api.util.ExceptionHandler;
import org.systemsbiology.biofabric.api.util.FixedJButton;
import org.systemsbiology.biofabric.ui.FabricDisplayOptions;
import org.systemsbiology.biofabric.ui.FabricDisplayOptionsManager;
import org.systemsbiology.biofabric.util.ResourceManager;

/****************************************************************************
**
** Dialog box for editing display properties
*/

public class FabricDisplayOptionsDialog extends BTStashResultsDialog {

  ////////////////////////////////////////////////////////////////////////////
  //
  // PRIVATE CONSTANTS
  //
  ////////////////////////////////////////////////////////////////////////////  
 
	private static final long serialVersionUID = 1L;
	
	private final Integer[] minDrainZoneChoices = {1, 2, 3, 4, 5, 10, 20};
	  
  ////////////////////////////////////////////////////////////////////////////
  //
  // PRIVATE INSTANCE MEMBERS
  //
  ////////////////////////////////////////////////////////////////////////////  
  
  private JTextField selectionOpaqueLevelField_;
  private JTextField nodeLighterLevelField_;
  private JTextField linkDarkerLevelField_;
  private JCheckBox displayShadowsBox_;
  private JCheckBox minShadSublinksBox_;
  private JCheckBox shadeNodesBox_;
  private JComboBox minDrainZoneBox_;
  private JCheckBox offerNodeBrowser_;
  private JTextField browserURLField_;
  private JCheckBox offerLinkBrowser_;
  private JTextField browserLinkURLField_;
  //private JCheckBox offerMouseOverView_;
  //private JTextField mouseOverTemplateField_;
  private FabricDisplayOptions newOpts_;
  
  private boolean needRebuild_;
  private boolean needRecolor_;
  
  ////////////////////////////////////////////////////////////////////////////
  //
  // PUBLIC CONSTRUCTORS
  //
  ////////////////////////////////////////////////////////////////////////////    

  /***************************************************************************
  **
  ** Constructor 
  */ 
  
  public FabricDisplayOptionsDialog(JFrame parent) {     
    super(parent, ResourceManager.getManager().getString("displayOptions.title"), new Dimension(1000, 600), 2);
    ResourceManager rMan = ResourceManager.getManager();
    parent_ = parent; 
    
    FabricDisplayOptionsManager dopmgr = FabricDisplayOptionsManager.getMgr();
    FabricDisplayOptions options = dopmgr.getDisplayOptions();
     
    selectionOpaqueLevelField_ = new JTextField(Double.toString(options.getSelectionOpaqueLevel()));
    JLabel label = new JLabel(rMan.getString("displayOptions.selectionOpaqueLevel"));
    addLabeledWidget(label, selectionOpaqueLevelField_, false, false); 
        
    nodeLighterLevelField_ = new JTextField(Double.toString(options.getNodeLighterLevel()));
    label = new JLabel(rMan.getString("displayOptions.nodeLighterLevel"));
    addLabeledWidget(label, nodeLighterLevelField_, false, false); 
    
    linkDarkerLevelField_ = new JTextField(Double.toString(options.getLinkDarkerLevel()));
    label = new JLabel(rMan.getString("displayOptions.linkDarkerLevel"));
    addLabeledWidget(label, linkDarkerLevelField_, false, false); 
    
    displayShadowsBox_ = new JCheckBox(rMan.getString("displayOptions.displayShadowLinks"));
    displayShadowsBox_.setSelected(options.getDisplayShadows());
    displayShadowsBox_.addActionListener(new ActionListener() {
      public void actionPerformed(ActionEvent ev) {
        try {
          minShadSublinksBox_.setEnabled(displayShadowsBox_.isSelected());
        } catch (Exception ex) {
          ExceptionHandler.getHandler().displayException(ex);
        }
        return;
      }
    });
   
    addWidgetFullRow(displayShadowsBox_, false); 
    
    minShadSublinksBox_ = new JCheckBox(rMan.getString("displayOptions.minShadowSubLinks"));
    minShadSublinksBox_.setSelected(options.getMinShadowSubmodelLinks());
    minShadSublinksBox_.setEnabled(options.getDisplayShadows());
    addWidgetFullRow(minShadSublinksBox_, false); 
 
    shadeNodesBox_ = new JCheckBox(rMan.getString("displayOptions.shadeNodes"));
    shadeNodesBox_.setSelected(options.getShadeNodes());
    addWidgetFullRow(shadeNodesBox_, false);
  
    minDrainZoneBox_ = new JComboBox(minDrainZoneChoices);  // have to use unchecked for v1.6
    minDrainZoneBox_.setSelectedIndex(getMinDrainZoneIndex(options.getMinDrainZone()));
    label = new JLabel(rMan.getString("displayOptions.minDrainZone"));
    addLabeledWidget(label, minDrainZoneBox_, false, false);
    
    offerNodeBrowser_ = new JCheckBox(rMan.getString("displayOptions.offerNodeBrowser"));
    offerNodeBrowser_.setSelected(options.getOfferNodeBrowser());
    addWidgetFullRow(offerNodeBrowser_, false); 
    
    browserURLField_ = new JTextField(options.getBrowserURL());
    label = new JLabel(rMan.getString("displayOptions.browserURL"));
    addLabeledWidget(label, browserURLField_, false, false); 
    
    offerLinkBrowser_ = new JCheckBox(rMan.getString("displayOptions.offerLinkBrowser"));
    offerLinkBrowser_.setSelected(options.getOfferLinkBrowser());
    addWidgetFullRow(offerLinkBrowser_, false); 
    
    browserLinkURLField_ = new JTextField(options.getBrowserLinkURL());
    label = new JLabel(rMan.getString("displayOptions.browserLinkURL"));
    addLabeledWidget(label, browserLinkURLField_, false, false); 
    
    //offerMouseOverView_ = new JCheckBox(rMan.getString("displayOptions.offerMouseOverView"));
    //offerMouseOverView_.setSelected(options.getOfferMouseOverView());
    //addWidgetFullRow(offerMouseOverView_, false); 
    
    //mouseOverTemplateField_ = new JTextField(options.getMouseOverURL());
    //label = new JLabel(rMan.getString("displayOptions.MouseOverURL"));
    //addLabeledWidget(label, mouseOverTemplateField_, false, false);    
    
    //
    // Build extra button:
    //

    FixedJButton buttonR = new FixedJButton(rMan.getString("dialogs.resetDefaults"));
    buttonR.addActionListener(new ActionListener() {
      public void actionPerformed(ActionEvent ev) {
        try {
          resetDefaults();
        } catch (Exception ex) {
          ExceptionHandler.getHandler().displayException(ex);
        }
      }
    });    
   
    finishConstructionWithExtraLeftButton(buttonR);

  }
    
  ////////////////////////////////////////////////////////////////////////////
  //
  // PUBLIC METHODS
  //
  ////////////////////////////////////////////////////////////////////////////  
  
  /***************************************************************************
  **
  ** Get results
  */
  
  public FabricDisplayOptions getNewOpts() {
    return (newOpts_);
  }
  
  /***************************************************************************
  **
  ** Get results
  */
  
  public boolean needsRecolor() {
    return (needRecolor_);
  }
  
  /***************************************************************************
  **
  ** Get results
  */
  
  public boolean needsRebuild() {
    return (needRebuild_);
  }
  
  ////////////////////////////////////////////////////////////////////////////
  //
  // PRIVATE METHODS
  //
  ////////////////////////////////////////////////////////////////////////////

  /***************************************************************************
  **
  ** Stash our results for later interrogation.
  ** 
  */
  
  protected boolean stashForOK() {
 
    FabricDisplayOptionsManager dopmgr = FabricDisplayOptionsManager.getMgr();
    FabricDisplayOptions oldOpts = dopmgr.getDisplayOptions();
    newOpts_ = new FabricDisplayOptions();
   
    needRecolor_ = false;
    needRebuild_ = false;
    
    String selOpqStr = selectionOpaqueLevelField_.getText();
    Double selOpq = parseDouble(selOpqStr, "displayOptions.badOpqLevel");
    if (selOpq == null) {
      return (false); 	
    } else if ((selOpq.doubleValue() < 0.0) || (selOpq.doubleValue() > 1.0))	{
        ResourceManager rMan = ResourceManager.getManager();
        JOptionPane.showMessageDialog(parent_, 
                                      rMan.getString("displayOptions.opacityOOB"), 
                                      rMan.getString("displayOptions.badLevelTitle"),
                                      JOptionPane.ERROR_MESSAGE);
        return (false);
    } else {
      newOpts_.setSelectionOpaqueLevel(selOpq.doubleValue());
    }
    
    String nodeLightStr = nodeLighterLevelField_.getText();
    Double nodeLight = parseDouble(nodeLightStr, "displayOptions.badNodeLightLevel");
    if (nodeLight == null) {
      return (false);
    } else if ((nodeLight.doubleValue() < 0.0) || (nodeLight.doubleValue() > 1.0))	{
        ResourceManager rMan = ResourceManager.getManager();
        JOptionPane.showMessageDialog(parent_, 
                                      rMan.getString("displayOptions.nodeLightOOB"), 
                                      rMan.getString("displayOptions.badLevelTitle"),
                                      JOptionPane.ERROR_MESSAGE);
        return (false);
    } else {
      newOpts_.setNodeLighterLevel(nodeLight.doubleValue());
      needRecolor_ = needRecolor_ || (oldOpts.getNodeLighterLevel() != newOpts_.getNodeLighterLevel());
    }
    
    String linkDrkStr = linkDarkerLevelField_.getText();
    Double linkDrk = parseDouble(linkDrkStr, "displayOptions.badLinkDarkLevel");
    if (linkDrk == null) {
      return (false);
    } else if ((linkDrk.doubleValue() < 0.0) || (linkDrk.doubleValue() > 1.0))	{
        ResourceManager rMan = ResourceManager.getManager();
        JOptionPane.showMessageDialog(parent_, 
                                      rMan.getString("displayOptions.linkDarkOOB"), 
                                      rMan.getString("displayOptions.badLevelTitle"),
                                      JOptionPane.ERROR_MESSAGE);
        return (false);
    } else {
      newOpts_.setLinkDarkerLevel(linkDrk.doubleValue());
      needRecolor_ = needRecolor_ || (oldOpts.getLinkDarkerLevel() != newOpts_.getLinkDarkerLevel());      
    }
    
    boolean currDisp = displayShadowsBox_.isSelected();
    newOpts_.setDisplayShadows(currDisp);
    needRebuild_ = needRebuild_ || (oldOpts.getDisplayShadows() != newOpts_.getDisplayShadows());
      
    boolean minShadSublinks = minShadSublinksBox_.isSelected();
    newOpts_.setMinShadowSubmodelLinks(currDisp && minShadSublinks);
 
    boolean currShade = shadeNodesBox_.isSelected();
    newOpts_.setShadeNodes(currShade);
    needRecolor_ = needRecolor_ || (oldOpts.getShadeNodes() != newOpts_.getShadeNodes());
    
    int newMinDrainZone = minDrainZoneChoices[minDrainZoneBox_.getSelectedIndex()];
    newOpts_.setMinDrainZone(newMinDrainZone);
    needRecolor_ = needRecolor_ || (oldOpts.getMinDrainZone() != newOpts_.getMinDrainZone());
    
    boolean offerNode = offerNodeBrowser_.isSelected();
    newOpts_.setOfferNodeBrowser(offerNode);
    
    boolean offerLink = offerLinkBrowser_.isSelected();
    newOpts_.setOfferLinkBrowser(offerLink);
    
    //boolean offerMO = offerMouseOverView_.isSelected();
    //newOpts_.setOfferMouseOverView(offerMO);    
    
    String browserURL = browserURLField_.getText().trim();
    if (!testURL(browserURL, true)) {
      return (false);    
    }
    newOpts_.setBrowserURL(browserURL);
    
    String browserLinkURL = browserLinkURLField_.getText().trim();
    if (!testURL(browserLinkURL, false)) {
      return (false);    
    }
    newOpts_.setBrowserLinkURL(browserLinkURL);
    
   // String mouseOverURL = this.mouseOverTemplateField_.getText().trim();
    //UiUtil.fixMePrintout("Test this URL for correctness");
    //newOpts_.setMouseOverURL(mouseOverURL);

    return (true);
  }
  
  /***************************************************************************
  **
  ** test URL
  ** 
  */
  
  private boolean testURL(String browserURL, boolean forNode) {
  
    if (!browserURL.equals("")) {
      boolean badURL = false;
      String testURL = null;
      
      if (forNode) {
        if (browserURL.indexOf(FabricDisplayOptions.NODE_NAME_PLACEHOLDER) == -1) {
          badURL = true;
        } else {
          testURL = browserURL.replaceFirst(FabricDisplayOptions.NODE_NAME_PLACEHOLDER, "WJRL");
        }
      } else {
        if ((browserURL.indexOf(FabricDisplayOptions.LINK_SRC_PLACEHOLDER) == -1) ||
            (browserURL.indexOf(FabricDisplayOptions.LINK_TRG_PLACEHOLDER) == -1) ||
            (browserURL.indexOf(FabricDisplayOptions.LINK_REL_PLACEHOLDER) == -1)) {
          badURL = true;
        } else {
          testURL = browserURL.replaceFirst(FabricDisplayOptions.LINK_SRC_PLACEHOLDER, "WJRL");
          testURL = testURL.replaceFirst(FabricDisplayOptions.LINK_TRG_PLACEHOLDER, "WJRL");
          testURL = testURL.replaceFirst(FabricDisplayOptions.LINK_REL_PLACEHOLDER, "WJRL");
       }        
      }
      
      if (testURL != null) {
        try {
          URL textURL = new URL(browserURL);
        } catch (MalformedURLException muex) {
          badURL = true;
        }
      }
      if (badURL) {
        ResourceManager rMan = ResourceManager.getManager();
        JOptionPane.showMessageDialog(parent_, 
                                      rMan.getString("displayOptions.badURL"), 
                                      rMan.getString("displayOptions.badURLTitle"),
                                      JOptionPane.ERROR_MESSAGE);
        return (false);
      }
    }
    return (true); 
  }
 
  /***************************************************************************
  **
  ** Parse the double
  ** 
  */
  
  private Double parseDouble(String doubleVal, String badMsg) {
    boolean badVal = false;
    double levelVal = 1.0;
    if ((doubleVal == null) || doubleVal.trim().equals("")) {
      badVal = true;
    } else {
      try {
        levelVal = Double.parseDouble(doubleVal);
        if (levelVal < 0.0) {
          badVal = true;
        }
      } catch (NumberFormatException ex) {
        badVal = true;
      }

    }
    if (badVal) {
      ResourceManager rMan = ResourceManager.getManager();
      JOptionPane.showMessageDialog(parent_, 
                                    rMan.getString(badMsg), 
                                    rMan.getString("displayOptions.badLevelTitle"),
                                    JOptionPane.ERROR_MESSAGE);
      return (null);
    }
    return (new Double(levelVal));
  }
  
  /***************************************************************************
   **
   **
   */
  
  private int getMinDrainZoneIndex(int choice) {
    for (int i = 0; i < minDrainZoneChoices.length; i++) {
      if (choice == minDrainZoneChoices[i]) {
        return (i);
      }
    }
    throw new IllegalArgumentException("Wrong argument: " + choice + " for min drain zone index request");
  }
 
  /***************************************************************************
  **
  ** Reset to default values
  ** 
  */
  
  private void resetDefaults() {
    FabricDisplayOptions defOptions = new FabricDisplayOptions();
    selectionOpaqueLevelField_.setText(Double.toString(defOptions.getSelectionOpaqueLevel()));
    nodeLighterLevelField_.setText(Double.toString(defOptions.getNodeLighterLevel()));
    linkDarkerLevelField_.setText(Double.toString(defOptions.getLinkDarkerLevel()));  
    displayShadowsBox_.setSelected(defOptions.getDisplayShadows());  
    minShadSublinksBox_.setSelected(defOptions.getMinShadowSubmodelLinks());  
    shadeNodesBox_.setSelected(defOptions.getShadeNodes());   
    browserURLField_.setText(defOptions.getBrowserURL());
    browserLinkURLField_.setText(defOptions.getBrowserLinkURL());
    //mouseOverTemplateField_.setText(defOptions.getMouseOverURL());
    return;
  }   
}
```
