## Supplementary material for "BioFabric Visualization of Network Alignments": BioFabric Source Code: FabricSearchDialog.java

```
/*
**    Copyright (C) 2003-2017 Institute for Systems Biology 
**                            Seattle, Washington, USA. 
**
**    This library is free software; you can redistribute it and/or
**    modify it under the terms of the GNU Lesser General Public
**    License as published by the Free Software Foundation; either
**    version 2.1 of the License, or (at your option) any later version.
**
**    This library is distributed in the hope that it will be useful,
**    but WITHOUT ANY WARRANTY; without even the implied warranty of
**    MERCHANTABILITY or FITNESS FOR A PARTICULAR PURPOSE.  See the GNU
**    Lesser General Public License for more details.
**
**    You should have received a copy of the GNU Lesser General Public
**    License along with this library; if not, write to the Free Software
**    Foundation, Inc., 59 Temple Place, Suite 330, Boston, MA  02111-1307  USA
*/


package org.systemsbiology.biofabric.ui.dialogs;

import java.awt.GridBagLayout;
import java.awt.GridBagConstraints;
import java.awt.event.ActionListener;
import java.awt.event.ActionEvent;
import java.util.Set;
import javax.swing.JDialog;
import javax.swing.JFrame;
import javax.swing.JLabel;
import javax.swing.JTextField;
import javax.swing.JPanel;
import javax.swing.Box;
import javax.swing.border.EmptyBorder;
import java.util.Vector;
import javax.swing.JCheckBox;
import javax.swing.JComboBox;

import javax.swing.JOptionPane;

import org.systemsbiology.biofabric.api.model.NetNode;
import org.systemsbiology.biofabric.api.util.ExceptionHandler;
import org.systemsbiology.biofabric.api.util.FixedJButton;
import org.systemsbiology.biofabric.model.BioFabricNetwork;
import org.systemsbiology.biofabric.util.ChoiceContent;
import org.systemsbiology.biofabric.util.DataUtil;
import org.systemsbiology.biofabric.util.ResourceManager;
import org.systemsbiology.biofabric.util.UiUtil;

/****************************************************************************
**
** Dialog box for searching for fabric nodes
*/

public class FabricSearchDialog extends JDialog {

  
  ////////////////////////////////////////////////////////////////////////////
  //
  // PRIVATE CONSTANTS
  //
  ////////////////////////////////////////////////////////////////////////////  
  
  private static final int FULL_MATCH_    = 0;
  private static final int PARTIAL_MATCH_ = 1;  
    
  ////////////////////////////////////////////////////////////////////////////
  //
  // PRIVATE INSTANCE MEMBERS
  //
  ////////////////////////////////////////////////////////////////////////////  
 
  private static final long serialVersionUID = 1L;
  private JTextField stringField_;
  private JFrame parent_;
  private boolean itemFound_;
  private JComboBox matchTypeCombo_;
  private JCheckBox discardSelections_;
  private BioFabricNetwork bfn_;
  private Set<NetNode> result_;
  private boolean doDiscard_;

  ////////////////////////////////////////////////////////////////////////////
  //
  // PUBLIC CONSTRUCTORS
  //
  ////////////////////////////////////////////////////////////////////////////    

  /***************************************************************************
  **
  ** Constructor 
  */ 
  
  public FabricSearchDialog(JFrame parent, BioFabricNetwork bfn, boolean haveSelection, boolean buildingSels) {     
    super(parent, ResourceManager.getManager().getString("nsearch.title"), true);

    parent_ = parent;
    itemFound_ = false;
    bfn_ = bfn;
    doDiscard_ = false;
  
    ResourceManager rMan = ResourceManager.getManager();    
    setSize(600, 300);
    JPanel cp = (JPanel)getContentPane();
    cp.setBorder(new EmptyBorder(20, 20, 20, 20));
    cp.setLayout(new GridBagLayout());
    GridBagConstraints gbc = new GridBagConstraints();    
     
    JPanel search = buildNodeSearchTab(haveSelection, buildingSels);
    int rowNum = 0;
    UiUtil.gbcSet(gbc, 0, rowNum, 10, 8, UiUtil.BO, 0, 0, 5, 5, 5, 5, UiUtil.CEN, 1.0, 1.0);    
    cp.add(search, gbc);    
    
    rowNum += 8;
    
    //
    // Build the button panel:
    //
  
    FixedJButton buttonO = new FixedJButton(rMan.getString("nsearch.search"));
    buttonO.addActionListener(new ActionListener() {
      public void actionPerformed(ActionEvent ev) {
        try {
          if (!doSearch()) {
            return;
          }
          FabricSearchDialog.this.setVisible(false);
          FabricSearchDialog.this.dispose();
        } catch (Exception ex) {
          ExceptionHandler.getHandler().displayException(ex);
        }
      }
    });     
    FixedJButton buttonC = new FixedJButton(rMan.getString("dialogs.cancel"));
    buttonC.addActionListener(new ActionListener() {
      public void actionPerformed(ActionEvent ev) {
        try {
          FabricSearchDialog.this.setVisible(false);
          FabricSearchDialog.this.dispose();
        } catch (Exception ex) {
          ExceptionHandler.getHandler().displayException(ex);
        }
      }
    });
    Box buttonPanel = Box.createHorizontalBox();
    buttonPanel.add(Box.createHorizontalGlue()); 
    buttonPanel.add(buttonO);
    buttonPanel.add(Box.createHorizontalStrut(10));    
    buttonPanel.add(buttonC);

    //
    // Build the dialog:
    //
    UiUtil.gbcSet(gbc, 0, rowNum, 4, 1, UiUtil.HOR, 0, 0, 5, 5, 5, 5, UiUtil.SE, 1.0, 0.0);
    cp.add(buttonPanel, gbc);
    setLocationRelativeTo(parent);
  }

  /***************************************************************************
  **
  ** Answer if something matched the search
  ** 
  */
  
  public boolean itemWasFound() {
    return (itemFound_);
  }
  
  /***************************************************************************
  **
  ** Get search matches
  ** 
  */
  
  public Set<NetNode> getMatches() {
    return (result_);
  }
  
  /***************************************************************************
  **
  ** Answer if we are to discard previous selections
  ** 
  */
  
  public boolean discardSelections() {
    return (doDiscard_);
  }  
 
  ////////////////////////////////////////////////////////////////////////////
  //
  // PRIVATE INNER CLASSES
  //
  ////////////////////////////////////////////////////////////////////////////  
 
  ////////////////////////////////////////////////////////////////////////////
  //
  // PRIVATE METHODS
  //
  ////////////////////////////////////////////////////////////////////////////

  /***************************************************************************
  **
  ** Node search tab
  */ 
  
  private JPanel buildNodeSearchTab(boolean haveSelection, boolean buildingSelections) {
    
    ResourceManager rMan = ResourceManager.getManager();    
    JPanel cp = new JPanel();
    cp.setLayout(new GridBagLayout());
    GridBagConstraints gbc = new GridBagConstraints();    
    //
    // Build the search string panel:
    //

    JLabel label = new JLabel(rMan.getString("nsearch.searchString"));   
    stringField_ = new JTextField();

    int rowNum = 0;
    UiUtil.gbcSet(gbc, 0, rowNum, 1, 1, UiUtil.NONE, 0, 0, 5, 5, 5, 5, UiUtil.E, 0.0, 1.0);
    cp.add(label, gbc);
    UiUtil.gbcSet(gbc, 1, rowNum, 1, 1, UiUtil.HOR, 0, 0, 5, 5, 5, 5, UiUtil.CEN, 1.0, 1.0);
    cp.add(stringField_, gbc); 
    
    matchTypeCombo_ = new JComboBox(getMatchChoices());
    
    UiUtil.gbcSet(gbc, 2, rowNum++, 1, 1, UiUtil.HOR, 0, 0, 5, 5, 5, 5, UiUtil.CEN, 0.0, 1.0);
    cp.add(matchTypeCombo_, gbc);    
      
    if (haveSelection) {
      discardSelections_ = new JCheckBox(rMan.getString("nsearch.discardCurrentSelections"));
      discardSelections_.setSelected(!buildingSelections);
      UiUtil.gbcSet(gbc, 2, rowNum++, 1, 1, UiUtil.HOR, 0, 0, 5, 5, 5, 5, UiUtil.CEN, 0.0, 1.0);
      cp.add(discardSelections_, gbc);
    }
       
    return (cp);
  } 
  
  /***************************************************************************
  **
  ** Get combo box guts
  */  
  
  private Vector<ChoiceContent> getMatchChoices() {
    ResourceManager rMan = ResourceManager.getManager();
    Vector<ChoiceContent> retval = new Vector<ChoiceContent>();
    retval.add(new ChoiceContent(rMan.getString("nsearch.fullMatch"), FULL_MATCH_));
    retval.add(new ChoiceContent(rMan.getString("nsearch.partialMatch"), PARTIAL_MATCH_));   
    return (retval);
  }

  /***************************************************************************
  **
  ** Do the search
  ** 
  */
  
  private boolean doSearch() {    
    int matchType = ((ChoiceContent)matchTypeCombo_.getSelectedItem()).val;
    String search = normalizeSearchString(stringField_.getText().trim());
    result_ = bfn_.nodeMatches(matchType == FULL_MATCH_, search);
    if (result_.isEmpty()) { 
      ResourceManager rMan = ResourceManager.getManager();
      JOptionPane.showMessageDialog(this, rMan.getString("nsearch.noMatchMessage"),
                                      rMan.getString("nsearch.noMatchTitle"),
                                      JOptionPane.WARNING_MESSAGE);
      return (false);
    }
    doDiscard_ = (discardSelections_ != null) ? discardSelections_.isSelected() : false;
    itemFound_ = true;
    return (true);
  }  
 
  /***************************************************************************
  **
  ** Normalize the search string
  ** 
  */
  
  private String normalizeSearchString(String searchString) {
    return (DataUtil.normKey(searchString));
  }    
}
```
