## Supplementary material for "BioFabric Visualization of Network Alignments": BioFabric Source Code: LinkGroupingSetupDialog.java

```
/*
**    Copyright (C) 2003-2018 Institute for Systems Biology 
**                            Seattle, Washington, USA. 
**
**    This library is free software; you can redistribute it and/or
**    modify it under the terms of the GNU Lesser General Public
**    License as published by the Free Software Foundation; either
**    version 2.1 of the License, or (at your option) any later version.
**
**    This library is distributed in the hope that it will be useful,
**    but WITHOUT ANY WARRANTY; without even the implied warranty of
**    MERCHANTABILITY or FITNESS FOR A PARTICULAR PURPOSE.  See the GNU
**    Lesser General Public License for more details.
**
**    You should have received a copy of the GNU Lesser General Public
**    License along with this library; if not, write to the Free Software
**    Foundation, Inc., 59 Temple Place, Suite 330, Boston, MA  02111-1307  USA
*/

package org.systemsbiology.biofabric.ui.dialogs;

import javax.swing.JCheckBox;
import javax.swing.JComboBox;
import javax.swing.JFrame;
import javax.swing.JLabel;
import javax.swing.JOptionPane;
import javax.swing.JPanel;

import org.systemsbiology.biofabric.api.dialog.BTStashResultsDialog;
import org.systemsbiology.biofabric.api.model.AugRelation;
import org.systemsbiology.biofabric.api.util.ExceptionHandler;
import org.systemsbiology.biofabric.api.util.FixedJButton;
import org.systemsbiology.biofabric.cmd.CommandSet;
import org.systemsbiology.biofabric.model.BioFabricNetwork;
import org.systemsbiology.biofabric.ui.dialogs.utils.EditableTable;
import org.systemsbiology.biofabric.util.DataUtil;
import org.systemsbiology.biofabric.util.ResourceManager;
import org.systemsbiology.biofabric.util.UiUtil;

import java.awt.Dimension;
import java.awt.event.ActionEvent;
import java.awt.event.ActionListener;
import java.io.File;
import java.util.ArrayList;
import java.util.Iterator;
import java.util.List;
import java.util.Set;

/****************************************************************************
 * *
 * * Dialog comboBox for specifying link groupings
 */

public class LinkGroupingSetupDialog extends BTStashResultsDialog {

  ////////////////////////////////////////////////////////////////////////////
  //
  // PRIVATE INSTANCE MEMBERS
  //
  ////////////////////////////////////////////////////////////////////////////
  
  private final int CHOICE_PER_NODE    = 0;
  private final int CHOICE_PER_NETWORK = 1;

  private List<String> linkGroupResult_;
  private EditableTable<LinkGroupingTableModel.TableRow> est_;
  private JFrame parent_;
  private Set<AugRelation> allRelations_;
  private JComboBox comboBox;
  private JCheckBox showAnnotationsBox_;
  private boolean showAnnotations_;
  private BioFabricNetwork.LayoutMode chosenMode;
  
  private static final long serialVersionUID = 1L;

  ////////////////////////////////////////////////////////////////////////////
  //
  // PUBLIC CONSTRUCTORS
  //
  ////////////////////////////////////////////////////////////////////////////    

  /***************************************************************************
   * *
   * * Constructor
   */

  public LinkGroupingSetupDialog(JFrame parent, List<String> currentTags, boolean showGroupAnnot, 
                                 BioFabricNetwork.LayoutMode mode,
                                 Set<AugRelation> allRelations) {
    super(parent, ResourceManager.getManager().getString("linkGroupEdit.title"), new Dimension(650, 450), 2);
    parent_ = parent;
    allRelations_ = allRelations;

    // bfn is only for pre-selecting the JComboBox to current LayoutMode
    installJComboBox(mode);
       
    showAnnotationsBox_ = new JCheckBox(rMan_.getString("linkGroupEdit.showLinkAnnotations"));
    addWidgetFullRow(showAnnotationsBox_, true, true);
    showAnnotationsBox_.setSelected(showGroupAnnot);

    FixedJButton fileButton =
            new FixedJButton(rMan_.getString("linkGroupEdit.loadFromFile"));

    fileButton.addActionListener(new ActionListener() {
      public void actionPerformed(ActionEvent e) {
        try {
          loadFromFile();
        } catch (Exception ex) {
          ExceptionHandler.getHandler().displayException(ex);
        }
      }
    });

    est_ = new EditableTable<LinkGroupingTableModel.TableRow>(new LinkGroupingTableModel(), parent_);
    EditableTable.TableParams etp = new EditableTable.TableParams();
    etp.addAlwaysAtEnd = false;
    etp.buttons = EditableTable.ALL_BUT_EDIT_BUTTONS;
    etp.singleSelectOnly = true;
    JPanel tablePan = est_.buildEditableTable(etp);

    addTable(tablePan, 4);

    List<LinkGroupingTableModel.TableRow> initRows = initTableRows((currentTags == null) ? new ArrayList<String>() : currentTags);
    
    est_.getModel().extractValues(initRows);
    setLocationRelativeTo(parent);

    finishConstructionWithExtraLeftButton(fileButton);
  }

  /***************************************************************************
   * * Install JComboBox with existing mode pre-selected from BioFabric Network's
   * * current layoutMode
   */

  private void installJComboBox(BioFabricNetwork.LayoutMode mode) {
    JLabel boxLabel = new JLabel(rMan_.getString("linkGroupEdit.mode"));

    String[] choices = new String[2];
    choices[CHOICE_PER_NODE] = rMan_.getString("linkGroupEdit.orderPerNode");
    choices[CHOICE_PER_NETWORK] = rMan_.getString("linkGroupEdit.orderNetwork");

    comboBox = new JComboBox(choices);

    if (mode == BioFabricNetwork.LayoutMode.UNINITIALIZED_MODE ||
            mode == BioFabricNetwork.LayoutMode.PER_NODE_MODE) {
      comboBox.setSelectedIndex(CHOICE_PER_NODE);

    } else if (mode == BioFabricNetwork.LayoutMode.PER_NETWORK_MODE) {
      comboBox.setSelectedIndex(CHOICE_PER_NETWORK);
    }

    addLabeledWidget(boxLabel, comboBox, true, true);
  }

  /***************************************************************************
   * *
   * * Return results
   * *
   */
  
  public List<String> getGroups() {
    return (linkGroupResult_);
  }

  /***************************************************************************
   * *
   * * Return chosen mode of comboBox
   * *
   */

  public BioFabricNetwork.LayoutMode getChosenMode() {
    return chosenMode;
  }
  
  /***************************************************************************
  ** 
  ** Return the choice to show link annotations or not 
  */  
  
  public boolean showLinkAnnotations() {
    return showAnnotations_;
  } 
  
  /***************************************************************************
   * *
   * * Stash our results for later interrogation.
   */
  
  @Override
  protected boolean stashForOK() {
    linkGroupResult_ = ((LinkGroupingTableModel) est_.getModel()).applyValues();
    if (linkGroupResult_ == null) {
      return (false);
    }
    
    int mode = comboBox.getSelectedIndex();
    
    if (mode == CHOICE_PER_NODE) {
      chosenMode =  BioFabricNetwork.LayoutMode.PER_NODE_MODE;
    } else if (mode == CHOICE_PER_NETWORK){
      chosenMode = BioFabricNetwork.LayoutMode.PER_NETWORK_MODE;
    } else {
      ExceptionHandler.getHandler()
              .displayException(new IllegalArgumentException("Illegal Selected Index"));
    }
    
    showAnnotations_ = showAnnotationsBox_.isSelected();
    
    return (true);
  }

  ////////////////////////////////////////////////////////////////////////////
  //
  // PRIVATE METHODS
  //
  ////////////////////////////////////////////////////////////////////////////
  
  /**
   * loads relation order from file and updates table with new order.
   * error checking done when user presses "ok"
   */
  
  private void loadFromFile() {
  	CommandSet cmd = CommandSet.getCmds("mainWindow");
    File fileEda = cmd.getFileLoader().getTheFile(".txt", null, "AttribDirectory", "filterName.txt", this);
    if (fileEda == null) {
      return;
    }
    List<String> groups = UiUtil.simpleFileRead(fileEda);
    if (groups == null) {
      return;
    }
    
    List<LinkGroupingTableModel.TableRow> initRows = initTableRows(groups);
    est_.updateTable(true, initRows);  // update table
    return;
  }
  
  /***************************************************************************
  **
  ** Get the list of table rows
  ** 
  */

  private List<LinkGroupingTableModel.TableRow> initTableRows(List<String> inTags) {
    ArrayList<LinkGroupingTableModel.TableRow> retval = new ArrayList<LinkGroupingTableModel.TableRow>();
    EditableTable.TableModel<LinkGroupingTableModel.TableRow> ecdtm = est_.getModel(); 
    Iterator<String> ceit = inTags.iterator();
    while (ceit.hasNext()) {
      String tag = ceit.next();
      LinkGroupingTableModel.TableRow tr = ecdtm.constructARow();
      tr.groupTag = tag;
      retval.add(tr);
    }
    return (retval);
  }  
  
  ////////////////////////////////////////////////////////////////////////////
  //
  // INNER CLASSES
  //
  ////////////////////////////////////////////////////////////////////////////
  
  /***************************************************************************
   * *
   * * The table
   */

  class LinkGroupingTableModel extends EditableTable.TableModel<LinkGroupingTableModel.TableRow> {

    private final static int GROUP_TAG_ = 0;
    private final static int NUM_COL_ = 1;
    
    private static final long serialVersionUID = 1L;
    
    LinkGroupingTableModel() {
      super(NUM_COL_);
      colNames_ = new String[]{"linkGroupEdit.suffix"};
      colClasses_ = new Class[]{String.class};
    }
    
    public class TableRow implements EditableTable.ATableRow {
      public String groupTag;
      
      public TableRow() {
      }
      
      public TableRow(String val) {
      	groupTag = val;
      	return;
      }
      
      TableRow(int i) {
        groupTag = (String)columns_.get(GROUP_TAG_).get(i);
      }
      
      public void toCols() {
        columns_.get(GROUP_TAG_).add(groupTag);  
        return;
      }
    }
    
    protected TableRow constructARow(int i) {
      return (new TableRow(i));     
    }
    
    public TableRow constructARow() {
      return (new TableRow());     
    }
     
    List<String> applyValues() {
      List<LinkGroupingTableModel.TableRow> vals = getValuesFromTable();
      
      //
      // Make sure the groups are OK. Names must be unique, non-blank, present as suffixes in the
      // provided set of link relations, and they must cover the set.
      //
      
      ResourceManager rMan = ResourceManager.getManager();
      ArrayList<String> seenTags = new ArrayList<String>();
      int size = vals.size();
      if (size == 0) {
        return (seenTags);
      }
      
      for (int i = 0; i < size; i++) {
        String tag = vals.get(i).groupTag;
        if ((tag == null) || (tag.trim().equals(""))) {
          JOptionPane.showMessageDialog(parent_, rMan.getString("rsedit.badRegion"),
                  rMan.getString("rsedit.badRegionTitle"),
                  JOptionPane.ERROR_MESSAGE);
          return (null);
        }
        
        tag = tag.trim();
        
        if (DataUtil.containsKey(seenTags, tag)) {
          JOptionPane.showMessageDialog(parent_, rMan.getString("rsedit.dupRegion"),
                  rMan.getString("rsedit.badRegionTitle"),
                  JOptionPane.ERROR_MESSAGE);

          return (null);
        }
        
        seenTags.add(tag);
      }
      
      
      boolean fail = false;
      int numST = seenTags.size();
      Iterator<AugRelation> arit = allRelations_.iterator();
      while (arit.hasNext()) {
        AugRelation relation =arit.next();
        boolean gotIt = false;
        for (int i = 0; i < numST; i++) {
          if (relation.relation.indexOf(seenTags.get(i)) != - 1) {
            gotIt = true;
            break;
          }
        }
        if (! gotIt) {
          fail = true;
          break;
        }
      }
      
      if (fail) {
        JOptionPane.showMessageDialog(parent_, rMan.getString("rsedit.notCovering"),
                rMan.getString("rsedit.badCoverageTitle"),
                JOptionPane.ERROR_MESSAGE);

        return (null);
      }
      return (seenTags);
    }
  }
  
}
```
