## Supplementary material for "BioFabric Visualization of Network Alignments": BioFabric Source Code: LinkRelationDialog.java

```
/*
 **    Copyright (C) 2018 Rishi Desai
 **
 **    Copyright (C) 2003-2018 Institute for Systems Biology
 **                            Seattle, Washington, USA.
 **
 **    This library is free software; you can redistribute it and/or
 **    modify it under the terms of the GNU Lesser General Public
 **    License as published by the Free Software Foundation; either
 **    version 2.1 of the License, or (at your option) any later version.
 **
 **    This library is distributed in the hope that it will be useful,
 **    but WITHOUT ANY WARRANTY; without even the implied warranty of
 **    MERCHANTABILITY or FITNESS FOR A PARTICULAR PURPOSE.  See the GNU
 **    Lesser General Public License for more details.
 **
 **    You should have received a copy of the GNU Lesser General Public
 **    License along with this library; if not, write to the Free Software
 **    Foundation, Inc., 59 Temple Place, Suite 330, Boston, MA  02111-1307  USA
 */

package org.systemsbiology.biofabric.ui.dialogs;

import java.awt.Dimension;
import java.awt.event.WindowEvent;
import java.awt.event.WindowListener;
import java.text.MessageFormat;
import javax.swing.JFrame;
import javax.swing.JLabel;
import javax.swing.JOptionPane;
import javax.swing.JTextField;
import javax.swing.event.DocumentEvent;
import javax.swing.event.DocumentListener;

import org.systemsbiology.biofabric.api.dialog.BTStashResultsDialog;
import org.systemsbiology.biofabric.api.dialog.DialogObj;
import org.systemsbiology.biofabric.api.util.ExceptionHandler;
import org.systemsbiology.biofabric.api.util.FixedJButton;
import org.systemsbiology.biofabric.util.ResourceManager;

public class LinkRelationDialog extends BTStashResultsDialog {
  
  ////////////////////////////////////////////////////////////////////////////
  //
  // PRIVATE FIELDS
  //
  ////////////////////////////////////////////////////////////////////////////
  
  private JFrame parent_;
  private FixedJButton buttonOK_;
  private JTextField textField_;
  private final String DEFAULT_RELATION_;
  
  private static final long serialVersionUID = 1L;
  
  ////////////////////////////////////////////////////////////////////////////
  //
  // CONSTRUCTOR
  //
  ////////////////////////////////////////////////////////////////////////////
  
  public LinkRelationDialog(JFrame parent, final String default_relation) {
    super(parent, ResourceManager.getManager().getString("dialog.relationTitle"), new Dimension(600, 200), 2);
    this.parent_ = parent;
    this.DEFAULT_RELATION_ = default_relation;
    
    final ResourceManager rMan = ResourceManager.getManager();
    
    String msg = MessageFormat.format(rMan.getString("dialog.relationMessage"), "GW");
    JLabel msgLabel = new JLabel(msg);
    addWidgetFullRow(msgLabel, false, false);
    
    textField_ = new JTextField(DEFAULT_RELATION_);
    textField_.getDocument().addDocumentListener(new DocumentListener() {
      public void insertUpdate(DocumentEvent e) {
        try {
          manageOKButton();
        } catch (Exception ex) {
          ExceptionHandler.getHandler().displayException(ex);
        }
      }
      
      public void removeUpdate(DocumentEvent e) {
        try {
          manageOKButton();
        } catch (Exception ex) {
          ExceptionHandler.getHandler().displayException(ex);
        }
      }
      
      public void changedUpdate(DocumentEvent e) {
        try {
          manageOKButton();
        } catch (Exception ex) {
          ExceptionHandler.getHandler().displayException(ex);
        }
      }
    });
    
    JLabel label = new JLabel(rMan.getString("dialog.linkRelation"));
    addLabeledWidget(label, textField_, false, false);
    
    //
    // OK and Cancel button
    //
    
    DialogObj.Buttons buttons = finishConstruction();
    
    buttonOK_ = buttons.okButton;
    manageOKButton();
    
    FixedJButton buttonCancel = buttons.cancelButton;
    buttonCancel.setEnabled(false);   // cancel button must always be disabled
    
    //
    // if user closes dialog, go with what is in text-field.
    // if text-field is empty or mis-formatted, use default relation
    //
    
    this.addWindowListener(new WindowListener() {
      public void windowClosing(WindowEvent e) {
        String msg = MessageFormat.format(rMan.getString("dialog.relationWarning"), getRelation());
        JOptionPane.showMessageDialog(parent_, msg,
                rMan.getString("dialog.relationWarningTitle"),
                JOptionPane.WARNING_MESSAGE);
        stashResults(true);
      }
      
      public void windowClosed(WindowEvent e) {}
      public void windowOpened(WindowEvent e) {}
      public void windowIconified(WindowEvent e) {}
      public void windowDeiconified(WindowEvent e) {}
      public void windowActivated(WindowEvent e) {}
      public void windowDeactivated(WindowEvent e) {}
    });
    
    setLocationRelativeTo(parent);
  }
  
  /**
   * * Return the relation (or default value)
   */
  
  public String getRelation() {
    String ret = textField_.getText().trim();
    return (hasMinRequirements() ? ret : DEFAULT_RELATION_);
  }
  
  /**
   * Check whether OK button should be activated or deactivated
   */
  
  private void manageOKButton() {
    if (hasMinRequirements()) {
      buttonOK_.setEnabled(true);
    } else {
      buttonOK_.setEnabled(false);
    }
    return;
  }
  
  /**
   * Check whether textfield has correct text
   */
  
  private boolean hasMinRequirements() {
    String text = textField_.getText().trim();
    return (! text.isEmpty() && text.split(" ").length == 1);
  }
  
  @Override
  public void okAction() {
    try {
      super.okAction();
    } catch (Exception ex) {
      ExceptionHandler.getHandler().displayException(ex);
    }
    return;
  }
  
  @Override
  protected boolean stashForOK() {
    return (hasMinRequirements());
  }
  
}
```
