## Supplementary material for "BioFabric Visualization of Network Alignments": BioFabric Source Code: NodeSimilarityLayoutSetupDialog.java

```
/*
**    Copyright (C) 2003-2014 Institute for Systems Biology 
**                            Seattle, Washington, USA. 
**
**    This library is free software; you can redistribute it and/or
**    modify it under the terms of the GNU Lesser General Public
**    License as published by the Free Software Foundation; either
**    version 2.1 of the License, or (at your option) any later version.
**
**    This library is distributed in the hope that it will be useful,
**    but WITHOUT ANY WARRANTY; without even the implied warranty of
**    MERCHANTABILITY or FITNESS FOR A PARTICULAR PURPOSE.  See the GNU
**    Lesser General Public License for more details.
**
**    You should have received a copy of the GNU Lesser General Public
**    License along with this library; if not, write to the Free Software
**    Foundation, Inc., 59 Temple Place, Suite 330, Boston, MA  02111-1307  USA
*/


package org.systemsbiology.biofabric.ui.dialogs;

import java.awt.Dimension;
import java.awt.event.ActionListener;
import java.awt.event.ActionEvent;
import javax.swing.JFrame;
import javax.swing.JTextField;
import javax.swing.JLabel;
import javax.swing.JComboBox;
import javax.swing.JOptionPane;

import org.systemsbiology.biofabric.api.dialog.BTStashResultsDialog;
import org.systemsbiology.biofabric.api.util.ExceptionHandler;
import org.systemsbiology.biofabric.api.util.FixedJButton;
import org.systemsbiology.biofabric.layouts.NodeSimilarityLayout;
import org.systemsbiology.biofabric.util.ChoiceContent;
import org.systemsbiology.biofabric.util.ResourceManager;

/****************************************************************************
**
** Dialog box for setting layout params
*/

public class NodeSimilarityLayoutSetupDialog extends BTStashResultsDialog {

  ////////////////////////////////////////////////////////////////////////////
  //
  // PRIVATE CONSTANTS
  //
  ////////////////////////////////////////////////////////////////////////////  
 
  private static final long serialVersionUID = 1L;
    
  ////////////////////////////////////////////////////////////////////////////
  //
  // PRIVATE INSTANCE MEMBERS
  //
  ////////////////////////////////////////////////////////////////////////////  
  
  private JTextField chainSizeField_;
  private JTextField jumpToleranceField_;
  private JComboBox distanceTypeCombo_;
  private NodeSimilarityLayout.ClusterParams results_;
  
  ////////////////////////////////////////////////////////////////////////////
  //
  // PUBLIC CONSTRUCTORS
  //
  ////////////////////////////////////////////////////////////////////////////    

  /***************************************************************************
  **
  ** Constructor 
  */ 
  
  public NodeSimilarityLayoutSetupDialog(JFrame parent, NodeSimilarityLayout.ClusterParams params) {     
    super(parent, ResourceManager.getManager().getString("clusteredLayout.title"), new Dimension(600, 350), 2);
    results_ = null;
       
    distanceTypeCombo_ = new JComboBox(NodeSimilarityLayout.ClusterParams.getDistanceChoices());
    int numSrc = distanceTypeCombo_.getItemCount();
    for (int i = 0; i < numSrc; i++) {
      ChoiceContent cc = (ChoiceContent)distanceTypeCombo_.getItemAt(i);
      if (cc.val == params.distanceMethod) {
        distanceTypeCombo_.setSelectedIndex(i);
        break;
      }
    }   

    JLabel label = new JLabel(rMan_.getString("clusteredLayout.distanceType"));
    addLabeledWidget(label, distanceTypeCombo_, false, false); 
   
    chainSizeField_ = new JTextField(Integer.toString(params.chainLength));
    label = new JLabel(rMan_.getString("clusteredLayout.chainSize"));
    addLabeledWidget(label, chainSizeField_, false, false); 
        
    jumpToleranceField_ = new JTextField(Double.toString(params.tolerance));
    label = new JLabel(rMan_.getString("clusteredLayout.jumpTolerance"));
    addLabeledWidget(label, jumpToleranceField_, false, false); 
    
    //
    // Build extra button:
    //

    FixedJButton buttonR = new FixedJButton(rMan_.getString("dialogs.resetDefaults"));
    buttonR.addActionListener(new ActionListener() {
      public void actionPerformed(ActionEvent ev) {
        try {
          resetDefaults();
        } catch (Exception ex) {
          ExceptionHandler.getHandler().displayException(ex);
        }
      }
    });    
   
    finishConstructionWithExtraLeftButton(buttonR);

  }
    
  ////////////////////////////////////////////////////////////////////////////
  //
  // PUBLIC METHODS
  //
  ////////////////////////////////////////////////////////////////////////////  
  
  /***************************************************************************
  **
  ** Get results
  */
  
  public NodeSimilarityLayout.ClusterParams getParams() {
    return (results_);
  }
  
  ////////////////////////////////////////////////////////////////////////////
  //
  // PRIVATE METHODS
  //
  ////////////////////////////////////////////////////////////////////////////

  /***************************************************************************
  **
  ** Stash our results for later interrogation.
  ** 
  */
  
  protected boolean stashForOK() {
 
    results_ = new NodeSimilarityLayout.ClusterParams();
    
    ChoiceContent cc = (ChoiceContent)distanceTypeCombo_.getSelectedItem();
    results_.distanceMethod = cc.val;
          
    String chainStr = chainSizeField_.getText();
    Integer chainLen = parseInteger(chainStr, "clusteredLayout.badChain");
    if (chainLen == null) {
      return (false);
    } else {
      results_.chainLength = chainLen.intValue();
    }
    
    String jumpStr = jumpToleranceField_.getText();
    Double jumpTol = parseDouble(jumpStr, "clusteredLayout.badTolerance");
    if (jumpTol == null) {
      return (false);
    } else {
      results_.tolerance = jumpTol.doubleValue();
    }
    
    return (true);
  }
   
  /***************************************************************************
  **
  ** Parse the double
  ** 
  */
  
  private Double parseDouble(String doubleVal, String badMsg) {
    boolean badVal = false;
    double levelVal = 1.0;
    if ((doubleVal == null) || doubleVal.trim().equals("")) {
      badVal = true;
    } else {
      try {
        levelVal = Double.parseDouble(doubleVal);
        if (levelVal < 0.0) {
          badVal = true;
        }
        if (levelVal > 1.0) {
          badVal = true;
        }
      } catch (NumberFormatException ex) {
        badVal = true;
      }

    }
    if (badVal) {
      ResourceManager rMan = ResourceManager.getManager();
      JOptionPane.showMessageDialog(parent_, 
                                    rMan.getString(badMsg), 
                                    rMan.getString("clusteredLayout.badValueTitle"),
                                    JOptionPane.ERROR_MESSAGE);
      return (null);
    }
    return (new Double(levelVal));
  } 
  
  /***************************************************************************
  **
  ** Parse the integer
  ** 
  */
  
  private Integer parseInteger(String intVal, String badMsg) {
    boolean badVal = false;
    int retVal = 1;
    if ((intVal == null) || intVal.trim().equals("")) {
      badVal = true;
    } else {
      try {
        retVal = Integer.parseInt(intVal);
        if (retVal < 1) {
          badVal = true;
        }
      } catch (NumberFormatException ex) {
        badVal = true;
      }

    }
    if (badVal) {
      ResourceManager rMan = ResourceManager.getManager();
      JOptionPane.showMessageDialog(parent_, 
                                    rMan.getString(badMsg), 
                                    rMan.getString("clusteredLayout.badValueTitle"),
                                    JOptionPane.ERROR_MESSAGE);
      return (null);
    }
    return (Integer.valueOf(retVal));
  } 
 
  /***************************************************************************
  **
  ** Reset to default values
  ** 
  */
  
  private void resetDefaults() {
    NodeSimilarityLayout.ClusterParams defaults = new NodeSimilarityLayout.ClusterParams();
   
    int numSrc = distanceTypeCombo_.getItemCount();
    for (int i = 0; i < numSrc; i++) {
      ChoiceContent cc = (ChoiceContent)distanceTypeCombo_.getItemAt(i);
      if (cc.val == defaults.distanceMethod) {
        distanceTypeCombo_.setSelectedIndex(i);
        break;
      }
    }  
    chainSizeField_.setText(Integer.toString(defaults.chainLength));
    jumpToleranceField_.setText(Double.toString(defaults.tolerance));
    return;
  }   
}
```
