## Supplementary material for "BioFabric Visualization of Network Alignments": BioFabric Source Code: PointUpOrDownDialog.java

```
/*
**    Copyright (C) 2003-2017 Institute for Systems Biology 
**                            Seattle, Washington, USA. 
**
**    This library is free software; you can redistribute it and/or
**    modify it under the terms of the GNU Lesser General Public
**    License as published by the Free Software Foundation; either
**    version 2.1 of the License, or (at your option) any later version.
**
**    This library is distributed in the hope that it will be useful,
**    but WITHOUT ANY WARRANTY; without even the implied warranty of
**    MERCHANTABILITY or FITNESS FOR A PARTICULAR PURPOSE.  See the GNU
**    Lesser General Public License for more details.
**
**    You should have received a copy of the GNU Lesser General Public
**    License along with this library; if not, write to the Free Software
**    Foundation, Inc., 59 Temple Place, Suite 330, Boston, MA  02111-1307  USA
*/

package org.systemsbiology.biofabric.ui.dialogs;

import java.awt.Dimension;

import javax.swing.ButtonGroup;
import javax.swing.JFrame;
import javax.swing.JRadioButton;

import org.systemsbiology.biofabric.api.dialog.BTStashResultsDialog;
import org.systemsbiology.biofabric.util.ResourceManager;

/****************************************************************************
**
** Dialog box for specifying if hierarchical DAG directed links point up or down
*/

public class PointUpOrDownDialog extends BTStashResultsDialog {

  ////////////////////////////////////////////////////////////////////////////
  //
  // PRIVATE CONSTANTS
  //
  ////////////////////////////////////////////////////////////////////////////  
 
  private static final long serialVersionUID = 1L;
    
  ////////////////////////////////////////////////////////////////////////////
  //
  // PRIVATE INSTANCE MEMBERS
  //
  ////////////////////////////////////////////////////////////////////////////  
  
  private JRadioButton pointUp_;
  private JRadioButton pointDown_;
  private boolean userWantsUp_;
  
  ////////////////////////////////////////////////////////////////////////////
  //
  // PUBLIC CONSTRUCTORS
  //
  ////////////////////////////////////////////////////////////////////////////    

  /***************************************************************************
  **
  ** Constructor 
  */ 
  
  public PointUpOrDownDialog(JFrame parent) {     
    super(parent, ResourceManager.getManager().getString("hierDagUpOrDown.title"), new Dimension(400, 200), 2);
   
    pointUp_ = new JRadioButton(rMan_.getString("hierDagUpOrDown.pointUp"), true);
    addWidgetFullRow(pointUp_, false);
    pointDown_ = new JRadioButton(rMan_.getString("hierDagUpOrDown.pointDown"), false);
    addWidgetFullRow(pointDown_, false);
          
    ButtonGroup group = new ButtonGroup();
    group.add(pointUp_);
    group.add(pointDown_);
    
    finishConstruction();
  }
    
  ////////////////////////////////////////////////////////////////////////////
  //
  // PUBLIC METHODS
  //
  ////////////////////////////////////////////////////////////////////////////  

  /***************************************************************************
  **
  ** Get results
  */
  
  public boolean getPointUp() {
    return (userWantsUp_);
  }
  
  ////////////////////////////////////////////////////////////////////////////
  //
  // PRIVATE METHODS
  //
  ////////////////////////////////////////////////////////////////////////////

  /***************************************************************************
  **
  ** Stash our results for later interrogation.
  ** 
  */
  
  protected boolean stashForOK() { 
    userWantsUp_ = pointUp_.isSelected();
    return (true);
  }
}
```
