## Supplementary material for "BioFabric Visualization of Network Alignments": BioFabric Source Code: RelationDirectionDialog.java

```
/*
**    Copyright (C) 2003-2018 Institute for Systems Biology 
**                            Seattle, Washington, USA. 
**
**    This library is free software; you can redistribute it and/or
**    modify it under the terms of the GNU Lesser General Public
**    License as published by the Free Software Foundation; either
**    version 2.1 of the License, or (at your option) any later version.
**
**    This library is distributed in the hope that it will be useful,
**    but WITHOUT ANY WARRANTY; without even the implied warranty of
**    MERCHANTABILITY or FITNESS FOR A PARTICULAR PURPOSE.  See the GNU
**    Lesser General Public License for more details.
**
**    You should have received a copy of the GNU Lesser General Public
**    License along with this library; if not, write to the Free Software
**    Foundation, Inc., 59 Temple Place, Suite 330, Boston, MA  02111-1307  USA
*/

package org.systemsbiology.biofabric.ui.dialogs;

import java.awt.Dimension;
import java.awt.event.ActionEvent;
import java.awt.event.ActionListener;

import javax.swing.JButton;
import javax.swing.JFrame;
import javax.swing.JPanel;

import java.util.Iterator;
import java.util.ArrayList;
import java.util.HashSet;
import java.util.List;
import java.util.SortedMap;
import java.util.TreeMap;

import org.systemsbiology.biofabric.api.dialog.BTStashResultsDialog;
import org.systemsbiology.biofabric.api.model.AugRelation;
import org.systemsbiology.biofabric.api.util.ExceptionHandler;
import org.systemsbiology.biofabric.api.util.FixedJButton;
import org.systemsbiology.biofabric.ui.dialogs.utils.EditableTable;
import org.systemsbiology.biofabric.util.ResourceManager;

/****************************************************************************
**
** Dialog box for editing relation directionality
*/

public class RelationDirectionDialog extends BTStashResultsDialog {

  ////////////////////////////////////////////////////////////////////////////
  //
  // PRIVATE INSTANCE MEMBERS
  //
  //////////////////////////////////////////////////////////////////////////// 
  
  private EditableTable<RelationDirTableModel.TableRow> est_;
  private SortedMap<AugRelation, Boolean> returnMap_;
  private boolean getFromFile_;  
  private HashSet<AugRelation> returnKeys_;  
  private SortedMap<AugRelation, Boolean> reducedMap_;
  private static final long serialVersionUID = 1L;
 
  ////////////////////////////////////////////////////////////////////////////
  //
  // PUBLIC CONSTRUCTORS
  //
  ////////////////////////////////////////////////////////////////////////////    

  /***************************************************************************
  **
  ** Constructor 
  */ 
  
  public RelationDirectionDialog(JFrame parent, SortedMap<AugRelation, Boolean> relationMap) {
    super(parent, ResourceManager.getManager().getString("relDir.title"), new Dimension(800, 400), 1);
    
    getFromFile_ = false;
    returnMap_ = null;
    returnKeys_ = new HashSet<AugRelation>(relationMap.keySet());
    reducedMap_ = new TreeMap<AugRelation, Boolean>();
    HashSet<String> seenRels = new HashSet<String>();
    Iterator<AugRelation> ceit = relationMap.keySet().iterator();
    while (ceit.hasNext()) {
      AugRelation aug = ceit.next();
      if (seenRels.contains(aug.relation)) {
        continue;
      }
      seenRels.add(aug.relation);
      Boolean isDir = relationMap.get(aug);
      reducedMap_.put(new AugRelation(aug.relation, false), isDir); 
    }   
 
    est_ = new EditableTable<RelationDirTableModel.TableRow>(new RelationDirTableModel(), parent_);
    EditableTable.TableParams etp = new EditableTable.TableParams();
    etp.addAlwaysAtEnd = true;
    etp.singleSelectOnly = true;
    etp.cancelEditOnDisable = true;
    etp.tableIsUnselectable = true;
    etp.perColumnEnums = null;
    etp.colWidths = new ArrayList<EditableTable.ColumnWidths>();  
    etp.colWidths.add(new EditableTable.ColumnWidths(RelationDirTableModel.RELATION, 50, 300, 5000));   
    etp.colWidths.add(new EditableTable.ColumnWidths(RelationDirTableModel.DIRECTION, 50, 100, 200));
   
    JPanel tablePan = est_.buildEditableTable(etp);
    addTable(tablePan, 8);  
    returnKeys_ = new HashSet<AugRelation>(relationMap.keySet());
 
    ArrayList<JButton> xtraButtonList = new ArrayList<JButton>();
    FixedJButton buttonSA = new FixedJButton(rMan_.getString("dialogs.selectAll"));
    buttonSA.addActionListener(new ActionListener() {
      public void actionPerformed(ActionEvent ev) {
        try {
          doMassUpdate(true);
        } catch (Exception ex) {
          ExceptionHandler.getHandler().displayException(ex);
        }
      }
    });
    xtraButtonList.add(buttonSA);
    
    FixedJButton buttonSN = new FixedJButton(rMan_.getString("dialogs.selectNone"));
    buttonSN.addActionListener(new ActionListener() {
      public void actionPerformed(ActionEvent ev) {
        try {
          doMassUpdate(false);
        } catch (Exception ex) {
          ExceptionHandler.getHandler().displayException(ex);
        }
      }
    });
    xtraButtonList.add(buttonSN);
    
    FixedJButton buttonFile = new FixedJButton(rMan_.getString("relDir.useFile"));
    buttonFile.addActionListener(new ActionListener() {
      public void actionPerformed(ActionEvent ev) {
        try {
          getFromFile_ = true;
          okAction();
        } catch (Exception ex) {
          ExceptionHandler.getHandler().displayException(ex);
        }
      }
    });
    xtraButtonList.add(buttonFile);

    finishConstructionWithMultiExtraLeftButtons(xtraButtonList);
    displayProperties();
  }
  
  /***************************************************************************
  **
  ** Mass check/uncheck
  ** 
  */
  
  void doMassUpdate(boolean select) {
    Boolean isDir = new Boolean(select);
    EditableTable.TableModel<RelationDirTableModel.TableRow> ecdtm = est_.getModel(); 
    List<RelationDirTableModel.TableRow> vals = ecdtm.getValuesFromTable();
    int numVals = vals.size();
    ArrayList<RelationDirTableModel.TableRow> upVals = new ArrayList<RelationDirTableModel.TableRow>();
    for (int i = 0; i < numVals; i++) {
      RelationDirTableModel.TableRow tr = vals.get(i);
      RelationDirTableModel.TableRow trc = ecdtm.constructARow();
      trc.relation = tr.relation;
      trc.isDir = isDir;
      upVals.add(trc);
    }
    est_.updateTable(true, upVals);
    return;
  }
  
  /***************************************************************************
  **
  ** Get the result
  ** 
  */
  
  public boolean getFromFile() {
    return (getFromFile_);
  }   
  
  /***************************************************************************
  **
  ** Get the result
  ** 
  */
  
  public SortedMap<AugRelation, Boolean> getRelationMap() {
    if (getFromFile_) {
      return (null);
    }
    returnMap_ = new TreeMap<AugRelation, Boolean>();
    Iterator<AugRelation> ceit = returnKeys_.iterator();
    while (ceit.hasNext()) {
      AugRelation aug = ceit.next();
      Boolean isDir = reducedMap_.get(new AugRelation(aug.relation, false));
      returnMap_.put(aug, isDir);
    }   
    return (returnMap_);
  }   
 
  /***************************************************************************
  **
  ** Provide a function to force all relations to undirected
  */
  
  public static SortedMap<AugRelation, Boolean> forceAllUndirected(SortedMap<AugRelation, Boolean> relationMap) {
    TreeMap<AugRelation, Boolean> returnMap = new TreeMap<AugRelation, Boolean>();
    for (AugRelation aug : relationMap.keySet()) {
      returnMap.put(aug, Boolean.FALSE); 
    } 
    return (returnMap); 
  }
  
  ////////////////////////////////////////////////////////////////////////////
  //
  // INNER CLASSES
  //
  ////////////////////////////////////////////////////////////////////////////  

  /***************************************************************************
  **
  ** Used for the data table
  */
  
  class RelationDirTableModel extends EditableTable.TableModel<RelationDirTableModel.TableRow> {
    
    private static final long serialVersionUID = 1L;
    
    final static int RELATION         = 0;
    final static int DIRECTION        = 1;
    private final static int NUM_COL_ = 2;
    
    class TableRow implements EditableTable.ATableRow {
      String relation;
      Boolean isDir;
   
      TableRow() {        
      }
      
      TableRow(int i) {
        relation = (String)columns_.get(RELATION).get(i);
        isDir = (Boolean)columns_.get(DIRECTION).get(i);
      }
      
      public void toCols() {
        columns_.get(RELATION).add(relation);  
        columns_.get(DIRECTION).add(isDir);
        return;
      }
      
      void replaceCols(int row) {
        columns_.get(RELATION).set(row, relation);  
        columns_.get(DIRECTION).set(row, isDir);
        return;
      }  
    }
 
    RelationDirTableModel() {
      super(NUM_COL_);
      colNames_ = new String[] {"relDir.relation",
                                "relDir.direction"};
      colClasses_ = new Class[] {String.class,
                                 Boolean.class};
      canEdit_ = new boolean[] {false,
                                true};
    }
 
    protected TableRow constructARow(int i) {
      return (new TableRow(i));     
    }
    public TableRow constructARow() {
      return (new TableRow());     
    }   
  }
    
  ////////////////////////////////////////////////////////////////////////////
  //
  // PRIVATE METHODS
  //
  ////////////////////////////////////////////////////////////////////////////

  /***************************************************************************
  **
  ** Apply the data values to our property table UI component
  ** 
  */
  
  private void displayProperties() {
    List<RelationDirTableModel.TableRow> tableRows = initTableRows();
    est_.getModel().extractValues(tableRows);
    return;
  }
  
  /***************************************************************************
  **
  ** Get the list of table rows
  ** 
  */

  private List<RelationDirTableModel.TableRow> initTableRows() {
    ArrayList<RelationDirTableModel.TableRow> retval = new ArrayList<RelationDirTableModel.TableRow>();
    EditableTable.TableModel<RelationDirTableModel.TableRow> ecdtm = est_.getModel(); 
    Iterator<AugRelation> ceit = reducedMap_.keySet().iterator();
    while (ceit.hasNext()) {
      AugRelation aug = ceit.next();
      RelationDirTableModel.TableRow tr = ecdtm.constructARow();
      tr.relation = aug.relation;
      tr.isDir = reducedMap_.get(aug);
      retval.add(tr);
    }
    return (retval);
  }
 
  /***************************************************************************
  **
  ** Stash our results for later interrogation.
  ** 
  */
  
  protected boolean stashForOK() {
    if (getFromFile_) {
      reducedMap_.clear();
      return (true);
    }
    EditableTable.TableModel<RelationDirTableModel.TableRow> ecdtm = est_.getModel();
    List<RelationDirTableModel.TableRow> vals = ecdtm.getValuesFromTable();
    int numVals = vals.size();
    for (int i = 0; i < numVals; i++) {
      RelationDirTableModel.TableRow tr = vals.get(i);
      reducedMap_.put(new AugRelation(tr.relation, false), tr.isDir);
    }
    return (true);
  }  
}
```
