## Supplementary material for "BioFabric Visualization of Network Alignments": BioFabric Source Code: ReorderLayoutParamsDialog.java

```
/*
**    Copyright (C) 2003-2012 Institute for Systems Biology 
**                            Seattle, Washington, USA. 
**
**    This library is free software; you can redistribute it and/or
**    modify it under the terms of the GNU Lesser General Public
**    License as published by the Free Software Foundation; either
**    version 2.1 of the License, or (at your option) any later version.
**
**    This library is distributed in the hope that it will be useful,
**    but WITHOUT ANY WARRANTY; without even the implied warranty of
**    MERCHANTABILITY or FITNESS FOR A PARTICULAR PURPOSE.  See the GNU
**    Lesser General Public License for more details.
**
**    You should have received a copy of the GNU Lesser General Public
**    License along with this library; if not, write to the Free Software
**    Foundation, Inc., 59 Temple Place, Suite 330, Boston, MA  02111-1307  USA
*/


package org.systemsbiology.biofabric.ui.dialogs;

import java.awt.Dimension;
import java.awt.event.ActionListener;
import java.awt.event.ActionEvent;
import javax.swing.JCheckBox;
import javax.swing.JFrame;
import javax.swing.JTextField;
import javax.swing.JLabel;
import javax.swing.JOptionPane;

import org.systemsbiology.biofabric.api.dialog.BTStashResultsDialog;
import org.systemsbiology.biofabric.api.util.ExceptionHandler;
import org.systemsbiology.biofabric.api.util.FixedJButton;
import org.systemsbiology.biofabric.layouts.NodeSimilarityLayout;
import org.systemsbiology.biofabric.util.ResourceManager;

/****************************************************************************
**
** Dialog box for setting layout params
*/

public class ReorderLayoutParamsDialog extends BTStashResultsDialog {

  ////////////////////////////////////////////////////////////////////////////
  //
  // PRIVATE CONSTANTS
  //
  ////////////////////////////////////////////////////////////////////////////  
 
  ////////////////////////////////////////////////////////////////////////////
  //
  // PRIVATE INSTANCE MEMBERS
  //
  ////////////////////////////////////////////////////////////////////////////  
  
  private JTextField numPassesField_;
  private JCheckBox termAtIncreaseBox_;
  private NodeSimilarityLayout.ResortParams results_;
  
  ////////////////////////////////////////////////////////////////////////////
  //
  // PUBLIC CONSTRUCTORS
  //
  ////////////////////////////////////////////////////////////////////////////    

  /***************************************************************************
  **
  ** Constructor 
  */ 
  
  public ReorderLayoutParamsDialog(JFrame parent, NodeSimilarityLayout.ResortParams params) {     
    super(parent, ResourceManager.getManager().getString("clusteredLayout.reorderTitle"), new Dimension(600, 300), 2);
    ResourceManager rMan = ResourceManager.getManager();
    parent_ = parent; 
    results_ = null;
     
    numPassesField_ = new JTextField(Integer.toString(params.passCount));
    JLabel label = new JLabel(rMan.getString("clusteredLayout.passCount"));
    addLabeledWidget(label, numPassesField_, false, false); 
        
    termAtIncreaseBox_ = new JCheckBox(rMan.getString("clusteredLayout.termAtIncrease"));
    termAtIncreaseBox_.setSelected(params.terminateAtIncrease);
    addWidgetFullRow(termAtIncreaseBox_, false); 
    
    //
    // Build extra button:
    //

    FixedJButton buttonR = new FixedJButton(rMan.getString("dialogs.resetDefaults"));
    buttonR.addActionListener(new ActionListener() {
      public void actionPerformed(ActionEvent ev) {
        try {
          resetDefaults();
        } catch (Exception ex) {
          ExceptionHandler.getHandler().displayException(ex);
        }
      }
    });    
   
    finishConstructionWithExtraLeftButton(buttonR);

  }
    
  ////////////////////////////////////////////////////////////////////////////
  //
  // PUBLIC METHODS
  //
  ////////////////////////////////////////////////////////////////////////////  
  
  /***************************************************************************
  **
  ** Get results
  */
  
  public NodeSimilarityLayout.ResortParams getParams() {
    return (results_);
  }
  
  ////////////////////////////////////////////////////////////////////////////
  //
  // PRIVATE METHODS
  //
  ////////////////////////////////////////////////////////////////////////////

  /***************************************************************************
  **
  ** Stash our results for later interrogation.
  ** 
  */
  
  protected boolean stashForOK() {
 
    results_ = new NodeSimilarityLayout.ResortParams();
       
    String passStr = numPassesField_.getText();
    Integer passes = parseInteger(passStr, "clusteredLayout.badPass");
    if (passes == null) {
      return (false);
    } else {
      results_.passCount = passes.intValue();
    }
    
    results_.terminateAtIncrease = termAtIncreaseBox_.isSelected();
    return (true);
  }
  
  /***************************************************************************
  **
  ** Parse the integer
  ** 
  */
  
  private Integer parseInteger(String intVal, String badMsg) {
    boolean badVal = false;
    int retVal = 1;
    if ((intVal == null) || intVal.trim().equals("")) {
      badVal = true;
    } else {
      try {
        retVal = Integer.parseInt(intVal);
        if (retVal < 1) {
          badVal = true;
        }
      } catch (NumberFormatException ex) {
        badVal = true;
      }

    }
    if (badVal) {
      ResourceManager rMan = ResourceManager.getManager();
      JOptionPane.showMessageDialog(parent_, 
                                    rMan.getString(badMsg), 
                                    rMan.getString("clusteredLayout.badValueTitle"),
                                    JOptionPane.ERROR_MESSAGE);
      return (null);
    }
    return (new Integer(retVal));
  } 
 
  /***************************************************************************
  **
  ** Reset to default values
  ** 
  */
  
  private void resetDefaults() {
    NodeSimilarityLayout.ResortParams defaults = new NodeSimilarityLayout.ResortParams();
    numPassesField_.setText(Integer.toString(defaults.passCount));
    termAtIncreaseBox_.setSelected(defaults.terminateAtIncrease);
    return;
  }   
}
```
