## Supplementary material for "BioFabric Visualization of Network Alignments": BioFabric Source Code: UpdateJavaDialog.java

```
/*
**    Copyright (C) 2003-2018 Institute for Systems Biology 
**                            Seattle, Washington, USA. 
**
**    This library is free software; you can redistribute it and/or
**    modify it under the terms of the GNU Lesser General Public
**    License as published by the Free Software Foundation; either
**    version 2.1 of the License, or (at your option) any later version.
**
**    This library is distributed in the hope that it will be useful,
**    but WITHOUT ANY WARRANTY; without even the implied warranty of
**    MERCHANTABILITY or FITNESS FOR A PARTICULAR PURPOSE.  See the GNU
**    Lesser General Public License for more details.
**
**    You should have received a copy of the GNU Lesser General Public
**    License along with this library; if not, write to the Free Software
**    Foundation, Inc., 59 Temple Place, Suite 330, Boston, MA  02111-1307  USA
*/


package org.systemsbiology.biofabric.ui.dialogs;

import java.awt.Dimension;
import java.awt.Frame;
import java.awt.GridBagConstraints;
import java.awt.GridBagLayout;
import java.awt.Toolkit;
import java.awt.event.ActionEvent;
import java.awt.event.ActionListener;
import javax.swing.Box;
import javax.swing.JButton;
import javax.swing.JDialog;
import javax.swing.JLabel;
import javax.swing.JPanel;
import javax.swing.border.EmptyBorder;

import org.systemsbiology.biofabric.util.UiUtil;

/****************************************************************************
**
** The frame for recommending Java update
*/

public class UpdateJavaDialog extends JDialog {
                                                          
                                                             
  ////////////////////////////////////////////////////////////////////////////
  //
  // PRIVATE MEMBERS
  //
  ////////////////////////////////////////////////////////////////////////////  

  private boolean keepGoing_;
  
  ////////////////////////////////////////////////////////////////////////////
  //
  // PUBLIC CONSTRUCTORS
  //
  ////////////////////////////////////////////////////////////////////////////    

  /***************************************************************************
  **
  ** Constructor 
  */ 
  
  public UpdateJavaDialog()  {
    super((Frame)null, true);
    keepGoing_ = false;
    Dimension screenSize = Toolkit.getDefaultToolkit().getScreenSize();
    setSize(640, 200);    

    Dimension frameSize = getSize();
    int x = (screenSize.width - frameSize.width) / 2;
    int y = (screenSize.height - frameSize.height) / 2;
    setLocation(x, y);
    JPanel cp = (JPanel)getContentPane();
    cp.setBorder(new EmptyBorder(20, 20, 20, 20));
    cp.setLayout(new GridBagLayout());
    GridBagConstraints gbc = new GridBagConstraints();    

    //
    // Build the message panel:
    //
    
    String message = "<html><center>BioFabric works best on Mac OS X with Java 6 or above.<br><br>(Java 5 " +
                      "renders BioFabric networks too lightly.) <br><br> It is recommended that you upgrade to at " +
                      "least Java version 6.</center></html>";
       

    JLabel label = new JLabel(message, JLabel.CENTER);
    UiUtil.gbcSet(gbc, 0, 0, 1, 1, UiUtil.BO, 0, 0, 5, 5, 5, 5, UiUtil.CEN, 1.0, 1.0);
    cp.add(label, gbc);
    
    //
    // Build the button panel:
    //

    JButton buttonO = new JButton("Continue Anyway");
    buttonO.addActionListener(new ActionListener() {
      public void actionPerformed(ActionEvent ev) {
        try {
          keepGoing_ = true;
          UpdateJavaDialog.this.setVisible(false);
          UpdateJavaDialog.this.dispose();
        } catch (Exception ex) {
          System.err.println("Caught exception");
        }
      }
    });     
    
    JButton buttonC = new JButton("Exit");
    buttonC.addActionListener(new ActionListener() {
      public void actionPerformed(ActionEvent ev) {
        try {
          UpdateJavaDialog.this.setVisible(false);
          UpdateJavaDialog.this.dispose();
        } catch (Exception ex) {
          System.err.println("Caught exception");
        }
      }
    });     
       
    Box buttonPanel = Box.createHorizontalBox();
    buttonPanel.add(Box.createHorizontalGlue());
    buttonPanel.add(buttonO);
    buttonPanel.add(Box.createHorizontalStrut(10)); 
    buttonPanel.add(buttonC);
    buttonPanel.add(Box.createHorizontalStrut(10)); 
  
    UiUtil.gbcSet(gbc, 0, 1, 1, 1, UiUtil.HOR, 0, 0, 5, 5, 5, 5, UiUtil.CEN, 1.0, 0.0);
    cp.add(buttonPanel, gbc); 
  }
  
  ////////////////////////////////////////////////////////////////////////////
  //
  // PUBLIC METHODS
  //
  ////////////////////////////////////////////////////////////////////////////    
  
  
  /***************************************************************************
  **
  ** Return results
  ** 
  */
  
  public boolean keepGoingAnyway() {
    return (keepGoing_);
  }  
}
```
