## Supplementary material for "BioFabric Visualization of Network Alignments": BioFabric Source Code: DialogSupport.java

```
/*
**    Copyright (C) 2003-2018 Institute for Systems Biology 
**                            Seattle, Washington, USA. 
**
**    This library is free software; you can redistribute it and/or
**    modify it under the terms of the GNU Lesser General Public
**    License as published by the Free Software Foundation; either
**    version 2.1 of the License, or (at your option) any later version.
**
**    This library is distributed in the hope that it will be useful,
**    but WITHOUT ANY WARRANTY; without even the implied warranty of
**    MERCHANTABILITY or FITNESS FOR A PARTICULAR PURPOSE.  See the GNU
**    Lesser General Public License for more details.
**
**    You should have received a copy of the GNU Lesser General Public
**    License along with this library; if not, write to the Free Software
**    Foundation, Inc., 59 Temple Place, Suite 330, Boston, MA  02111-1307  USA
*/

package org.systemsbiology.biofabric.ui.dialogs.utils;

import java.awt.GridBagConstraints;
import java.awt.event.ActionListener;
import java.awt.event.ActionEvent;
import java.util.ArrayList;
import java.util.List;
import javax.swing.JPanel;
import javax.swing.Box;
import javax.swing.JButton;
import javax.swing.JComponent;
import javax.swing.JDialog;
import javax.swing.JLabel;
import javax.swing.border.Border;

import org.systemsbiology.biofabric.api.dialog.DialogObj;
import org.systemsbiology.biofabric.api.util.ExceptionHandler;
import org.systemsbiology.biofabric.api.util.FixedJButton;
import org.systemsbiology.biofabric.util.ResourceManager;
import org.systemsbiology.biofabric.util.UiUtil;

/****************************************************************************
**
** Helper for common dialog tasks
*/

public class DialogSupport {
  
  ////////////////////////////////////////////////////////////////////////////
  //
  // PRIVATE INSTANCE MEMBERS
  //
  ////////////////////////////////////////////////////////////////////////////  

  private DialogObj.DialogSupportClient dsClient_;
  private ResourceManager rMan_; 
  private GridBagConstraints gbc_;
  private JDialog closeMe_;
 
  ////////////////////////////////////////////////////////////////////////////
  //
  // PUBLIC CLIENT INTERFACE
  //
  ////////////////////////////////////////////////////////////////////////////    

  ////////////////////////////////////////////////////////////////////////////
  //
  // PUBLIC CONSTRUCTORS
  //
  ////////////////////////////////////////////////////////////////////////////    
  
  /***************************************************************************
  **
  ** Constructor 
  */ 
  
  public DialogSupport(DialogObj.DialogSupportClient client, ResourceManager rMan, GridBagConstraints gbc) {
    dsClient_ = client;
    rMan_ = rMan;
    gbc_ = gbc;
  }
  
  /***************************************************************************
  **
  ** Constructor 
  */ 
  
  public DialogSupport(ResourceManager rMan, GridBagConstraints gbc, JDialog closeMe) {
    closeMe_ = closeMe;
    rMan_ = rMan;
    gbc_ = gbc;
  }  
  
  /***************************************************************************
  **
  ** Constructor 
  */ 
  
  public DialogSupport(ResourceManager rMan, GridBagConstraints gbc) {
    rMan_ = rMan;
    gbc_ = gbc;
  }  
  
  ////////////////////////////////////////////////////////////////////////////
  //
  // PUBLIC METHODS
  //
  ////////////////////////////////////////////////////////////////////////////    
  
  /***************************************************************************
  **
  ** Get stashed resource manager
  */ 
  
  public ResourceManager getRman() {
    return (rMan_); 
  }
  
  /***************************************************************************
  **
  ** Get stashed gb constraints
  */ 
  
  public GridBagConstraints getGbc() {
    return (gbc_); 
  }
 
  /***************************************************************************
  **
  ** Build a button panel
  */ 

  public DialogObj.ButtonsAndBox buildButtonBox(boolean doApply, boolean showAsCancel, boolean doCentering) {       
 
    DialogObj.Buttons myButtons = new DialogObj.Buttons();
    
    myButtons.applyButton = null;
    if (doApply) {
      myButtons.applyButton = new FixedJButton(rMan_.getString("dialogs.apply"));
      myButtons.applyButton.addActionListener(new ActionListener() {
        public void actionPerformed(ActionEvent ev) {
          try {
            dsClient_.applyAction();
          } catch (Exception ex) {
            ExceptionHandler.getHandler().displayException(ex);
          } catch (OutOfMemoryError oom) {
            ExceptionHandler.getHandler().displayOutOfMemory(oom);
          }
        }
      });
    }
    myButtons.okButton = new FixedJButton(rMan_.getString("dialogs.ok"));
    myButtons.okButton.addActionListener(new ActionListener() {
      public void actionPerformed(ActionEvent ev) {
        try {
          dsClient_.okAction();
        } catch (Exception ex) {
          ExceptionHandler.getHandler().displayException(ex);
        } catch (OutOfMemoryError oom) {
          ExceptionHandler.getHandler().displayOutOfMemory(oom);
        }
      }
    });     
    myButtons.cancelButton = new FixedJButton(rMan_.getString((showAsCancel) ? "dialogs.cancel" : "dialogs.close"));
    myButtons.cancelButton.addActionListener(new ActionListener() {
      public void actionPerformed(ActionEvent ev) {
        try {
          dsClient_.closeAction();
        } catch (Exception ex) {
          ExceptionHandler.getHandler().displayException(ex);
        } catch (OutOfMemoryError oom) {
          ExceptionHandler.getHandler().displayOutOfMemory(oom);
        }
      }
    });
    
    Box buttonPanel = Box.createHorizontalBox();
    buttonPanel.add(Box.createHorizontalGlue());
    if (doApply) {
      buttonPanel.add(myButtons.applyButton);
      buttonPanel.add(Box.createHorizontalStrut(10));
    }
    buttonPanel.add(myButtons.okButton);
    buttonPanel.add(Box.createHorizontalStrut(10));    
    buttonPanel.add(myButtons.cancelButton);
    if (doCentering) {
      buttonPanel.add(Box.createHorizontalGlue());
    }
    DialogObj.ButtonsAndBox retval = new DialogObj.ButtonsAndBox();
    retval.buttonBox = buttonPanel;
    retval.buttons = myButtons;
    return (retval);
  }
 
  /***************************************************************************
  **
  ** Build and install buttons
  */ 

  public DialogObj.Buttons buildAndInstallButtonBox(JPanel cp, int rowNum, int colWidth, boolean doApply, boolean showAsCancel, Border border) { 
    DialogObj.ButtonsAndBox bAndBox = buildButtonBox(doApply, showAsCancel, false);
    if (border != null) {
      bAndBox.buttonBox.setBorder(border);
    } 
    UiUtil.gbcSet(gbc_, 0, rowNum, colWidth, 1, UiUtil.HOR, 0, 0, 5, 5, 5, 5, UiUtil.SE, 1.0, 0.0);
    cp.add(bAndBox.buttonBox, gbc_);
    return (bAndBox.buttons);
  }
  
  /***************************************************************************
  **
  ** Build and install buttons
  */ 

  public DialogObj.Buttons buildAndInstallCenteredButtonBox(JPanel cp, int rowNum, int colWidth, boolean doApply, boolean showAsCancel) { 
    DialogObj.ButtonsAndBox bAndBox = buildButtonBox(doApply, showAsCancel, true);
    UiUtil.gbcSet(gbc_, 0, rowNum, colWidth, 1, UiUtil.HOR, 0, 0, 5, 5, 5, 5, UiUtil.SE, 1.0, 0.0);
    cp.add(bAndBox.buttonBox, gbc_);
    return (bAndBox.buttons);
  } 
  
  /***************************************************************************
  **
  ** Build and install buttons
  */ 

  public DialogObj.Buttons buildAndInstallButtonBox(JPanel cp, int rowNum, int colWidth, boolean doApply, boolean showAsCancel) { 
    DialogObj.Buttons retval = buildAndInstallButtonBox(cp, rowNum, colWidth, doApply, showAsCancel, null);
    return (retval);
  }
    
  /***************************************************************************
  **
  ** Build a button panel
  */ 

  public Box buildButtonBoxWithExtra(boolean doApply, JButton xtraButton, boolean showAsCancel) {
    ArrayList<JButton> xtraButtonList = new ArrayList<JButton>();
    xtraButtonList.add(xtraButton);
    return (buildButtonBoxWithMultiExtra(doApply, xtraButtonList, showAsCancel));
  }
  
  /***************************************************************************
  **
  ** Build a button panel with many left extras
  */ 

  public Box buildButtonBoxWithMultiExtra(boolean doApply, List<JButton> xtraButtonList, boolean showAsCancel) {       
 
    FixedJButton buttonA = null;
    if (doApply) {
      buttonA = new FixedJButton(rMan_.getString("dialogs.apply"));
      buttonA.addActionListener(new ActionListener() {
        public void actionPerformed(ActionEvent ev) {
          try {
            dsClient_.applyAction();
          } catch (Exception ex) {
            ExceptionHandler.getHandler().displayException(ex);
          } catch (OutOfMemoryError oom) {
            ExceptionHandler.getHandler().displayOutOfMemory(oom);
          }
        }
      });
    }
    FixedJButton buttonO = new FixedJButton(rMan_.getString("dialogs.ok"));
    buttonO.addActionListener(new ActionListener() {
      public void actionPerformed(ActionEvent ev) {
        try {
          dsClient_.okAction();
        } catch (Exception ex) {
          ExceptionHandler.getHandler().displayException(ex);
        } catch (OutOfMemoryError oom) {
          ExceptionHandler.getHandler().displayOutOfMemory(oom);
        }
      }
    });     
    FixedJButton buttonC = new FixedJButton(rMan_.getString((showAsCancel) ? "dialogs.cancel" : "dialogs.close"));
    buttonC.addActionListener(new ActionListener() {
      public void actionPerformed(ActionEvent ev) {
        try {
          dsClient_.closeAction();
        } catch (Exception ex) {
          ExceptionHandler.getHandler().displayException(ex);
        } catch (OutOfMemoryError oom) {
          ExceptionHandler.getHandler().displayOutOfMemory(oom);
        }
      }
    });
    
    Box buttonPanel = Box.createHorizontalBox();
    buttonPanel.add(Box.createHorizontalStrut(10));
    int numBut = xtraButtonList.size();
    for (int i = 0; i < numBut; i++) {
      buttonPanel.add(xtraButtonList.get(i));
      if (i != (numBut - 1)) {
        buttonPanel.add(Box.createHorizontalStrut(10));
      }
    }
    buttonPanel.add(Box.createHorizontalGlue());
    if (doApply) {
      buttonPanel.add(buttonA);
      buttonPanel.add(Box.createHorizontalStrut(10));
    }
    buttonPanel.add(buttonO);
    buttonPanel.add(Box.createHorizontalStrut(10));    
    buttonPanel.add(buttonC);
    return (buttonPanel);
  }
  
  /***************************************************************************
  **
  ** Build and install buttons
  */ 

  public void buildAndInstallButtonBoxWithExtra(JPanel cp, int rowNum, int colWidth, boolean doApply, 
                                                JButton xtraButton, boolean showAsCancel) { 
    Box bBox = buildButtonBoxWithExtra(doApply, xtraButton, showAsCancel);
    UiUtil.gbcSet(gbc_, 0, rowNum, colWidth, 1, UiUtil.HOR, 0, 0, 5, 5, 5, 5, UiUtil.SE, 1.0, 0.0);
    cp.add(bBox, gbc_);
    return;
  }
  
  /***************************************************************************
  **
  ** Build and install buttons
  */ 

  public void buildAndInstallButtonBoxWithMultiExtra(JPanel cp, int rowNum, int colWidth, boolean doApply, 
                                                     List<JButton> xtraButtonList, boolean showAsCancel) { 
    Box bBox = buildButtonBoxWithMultiExtra(doApply, xtraButtonList, showAsCancel);
    UiUtil.gbcSet(gbc_, 0, rowNum, colWidth, 1, UiUtil.HOR, 0, 0, 5, 5, 5, 5, UiUtil.SE, 1.0, 0.0);
    cp.add(bBox, gbc_);
    return;
  }
    
  /***************************************************************************
  **
  ** Build a close button panel
  */ 

  public Box buildCloseButtonBox() {       
    FixedJButton buttonC = new FixedJButton(rMan_.getString("dialogs.close"));
    buttonC.addActionListener(new ActionListener() {
      public void actionPerformed(ActionEvent ev) {
        try {
          closeMe_.setVisible(false);
          closeMe_.dispose();
        } catch (Exception ex) {
          ExceptionHandler.getHandler().displayException(ex);
        } catch (OutOfMemoryError oom) {
          ExceptionHandler.getHandler().displayOutOfMemory(oom);
        }
      }
    });
    
    Box buttonPanel = Box.createHorizontalBox();
    buttonPanel.add(Box.createHorizontalGlue());   
    buttonPanel.add(buttonC);
    buttonPanel.add(Box.createHorizontalStrut(10));       
    return (buttonPanel);
  }
  
  /***************************************************************************
  **
  ** Build and install buttons
  */ 

  public void buildAndInstallCloseButtonBox(JPanel cp, int rowNum, int colWidth, Border border) {  
    Box bBox = buildCloseButtonBox();
    if (border != null) {
      bBox.setBorder(border);
    }
    UiUtil.gbcSet(gbc_, 0, rowNum, colWidth, 1, UiUtil.HOR, 0, 0, 5, 5, 5, 5, UiUtil.SE, 1.0, 0.0);
    cp.add(bBox, gbc_);
    return;
  }  
   
  /***************************************************************************
  **
  ** Install a labeled component
  */ 

  public int installLabeledJComp(JComponent tf, JPanel cp, String labelRS, int rowNum, int colWidth) {  
    JLabel typeLab = new JLabel(rMan_.getString(labelRS));       
    UiUtil.gbcSet(gbc_, 0, rowNum, 1, 1, UiUtil.NONE, 0, 0, 5, 5, 5, 5, UiUtil.E, 0.0, 0.0);       
    cp.add(typeLab, gbc_);  
    UiUtil.gbcSet(gbc_, 1, rowNum++, colWidth - 1, 1, UiUtil.HOR, 0, 0, 5, 5, 5, 5, UiUtil.W, 1.0, 0.0);       
    cp.add(tf, gbc_);   
    return (rowNum);
  }
  
  /***************************************************************************
  **
  ** Install a labeled component
  */ 

  public int installPreLabeledJComp(JComponent tf, JPanel cp, JLabel typeLab, int rowNum, int colWidth) {      
    UiUtil.gbcSet(gbc_, 0, rowNum, 1, 1, UiUtil.NONE, 0, 0, 5, 5, 5, 5, UiUtil.E, 0.0, 0.0);       
    cp.add(typeLab, gbc_);  
    UiUtil.gbcSet(gbc_, 1, rowNum++, colWidth - 1, 1, UiUtil.HOR, 0, 0, 5, 5, 5, 5, UiUtil.W, 1.0, 0.0);       
    cp.add(tf, gbc_);   
    return (rowNum);
  }
    
  /***************************************************************************
  **
  ** Install a labeled component pair
  */ 

  public int installLabelJCompPair(JPanel cp, String labelRS1, JComponent tf1, 
                                   String labelRS2, JComponent tf2, int rowNum, int colWidth, boolean gottaGrow) { 
    int halfCol = colWidth / 2;
    
    double vert = (gottaGrow) ? 1.0 : 0.0;
    JLabel typeLab = new JLabel(rMan_.getString(labelRS1));       
    UiUtil.gbcSet(gbc_, 0, rowNum, 1, 1, UiUtil.NONE, 0, 0, 5, 5, 5, 5, UiUtil.E, 0.0, vert);       
    cp.add(typeLab, gbc_);  
    UiUtil.gbcSet(gbc_, 1, rowNum, halfCol - 1, 1, UiUtil.HOR, 0, 0, 5, 5, 5, 5, UiUtil.W, 0.5, vert);       
    cp.add(tf1, gbc_); 
    
    JLabel lab2 = new JLabel(rMan_.getString(labelRS2));       
    UiUtil.gbcSet(gbc_, halfCol, rowNum, 1, 1, UiUtil.NONE, 0, 0, 5, 5, 5, 5, UiUtil.E, 0.0, vert);       
    cp.add(lab2, gbc_);  
    UiUtil.gbcSet(gbc_, halfCol + 1, rowNum++, halfCol - 1, 1, UiUtil.HOR, 0, 0, 5, 5, 5, 5, UiUtil.W, 0.5, vert);       
    cp.add(tf2, gbc_);      
    return (rowNum);
  }

  /***************************************************************************
  **
  ** Install a labeled jCOmponent field triple
  */ 

  public int installLabelJCompTriple(JPanel cp, 
                                     Object labelRS1, JComponent tf1, 
                                     Object labelRS2, JComponent tf2, 
                                     Object labelRS3, JComponent tf3, 
                                     int rowNum, int colWidth) { 
    int thirdCol = colWidth / 3;
    int twoThirdCol = thirdCol * 2;
    
    int width;
    
    if (labelRS1 != null) {
      JLabel typeLab = (labelRS1 instanceof String) ? new JLabel(rMan_.getString((String)labelRS1)) : (JLabel)labelRS1;       
      UiUtil.gbcSet(gbc_, 0, rowNum, 1, 1, UiUtil.NONE, 0, 0, 5, 5, 5, 5, UiUtil.E, 0.0, 0.0);       
      cp.add(typeLab, gbc_);
      width = 1;
    } else {
      width = 0;
    }
    
    UiUtil.gbcSet(gbc_, width, rowNum, thirdCol - width, 1, UiUtil.HOR, 0, 0, 5, 5, 5, 5, UiUtil.W, 0.33, 0.0);       
    cp.add(tf1, gbc_); 
    
    if (labelRS2 != null) {
      JLabel lab2 = (labelRS2 instanceof String) ? new JLabel(rMan_.getString((String)labelRS2)) : (JLabel)labelRS2;
      UiUtil.gbcSet(gbc_, thirdCol, rowNum, 1, 1, UiUtil.NONE, 0, 0, 5, 5, 5, 5, UiUtil.E, 0.0, 0.0);       
      cp.add(lab2, gbc_);
      width = 1;
    } else {
      width = 0;
    }
    
    UiUtil.gbcSet(gbc_, thirdCol + width, rowNum, thirdCol - width, 1, UiUtil.HOR, 0, 0, 5, 5, 5, 5, UiUtil.W, 0.33, 0.0);       
    cp.add(tf2, gbc_);
  
     
    if (labelRS2 != null) {
      JLabel lab3 = (labelRS3 instanceof String) ? new JLabel(rMan_.getString((String)labelRS3)) : (JLabel)labelRS3;
      UiUtil.gbcSet(gbc_, twoThirdCol, rowNum, 1, 1, UiUtil.NONE, 0, 0, 5, 5, 5, 5, UiUtil.E, 0.0, 0.0);       
      cp.add(lab3, gbc_);  
      width = 1;
    } else {
      width = 0;
    }
    
    UiUtil.gbcSet(gbc_, twoThirdCol + width, rowNum++, thirdCol - width, 1, UiUtil.HOR, 0, 0, 5, 5, 5, 5, UiUtil.W, 0.34, 0.0);       
    cp.add(tf3, gbc_);
       
    return (rowNum);
  }
  
  /***************************************************************************
  **
  ** Add a full row component
  */ 
  
  public int addWidgetFullRow(JPanel cp, JComponent comp, boolean fixHeight, int rowNum, int columns) {
    return (addWidgetFullRow(cp, comp, fixHeight, false, rowNum, columns));
  } 
  
  /***************************************************************************
  **
  ** Add a full row component
  */ 
  
  public int addWidgetFullRow(JPanel cp, JComponent comp, boolean fixHeight, 
                                  boolean flushLeft, int rowNum, int columns) {
    int worc = (flushLeft) ? UiUtil.W : UiUtil.CEN;
    double vFac = (fixHeight) ? 0.0 : 1.0;
    UiUtil.gbcSet(gbc_, 0, rowNum++, columns, 1, UiUtil.HOR, 0, 0, 5, 5, 5, 5, worc, 1.0, vFac);
    cp.add(comp, gbc_);
    return (rowNum);
  } 
  
  /***************************************************************************
  **
  ** Add a full row component
  */ 
  
  public int addTallWidgetFullRow(JPanel cp, JComponent comp, boolean fixHeight, boolean flushLeft, 
                                     int height, int rowNum, int columns) {
    int worc = (flushLeft) ? UiUtil.W : UiUtil.CEN;
    double vFac = (fixHeight) ? 0.0 : 1.0;
    UiUtil.gbcSet(gbc_, 0, rowNum, columns, height, UiUtil.HOR, 0, 0, 5, 5, 5, 5, worc, 1.0, vFac);
    cp.add(comp, gbc_);
    rowNum += height;
    return (rowNum);
  } 

   /***************************************************************************
  **
  ** Add a full row component
  */ 
  
  public int addWidgetFullRowWithInsets(JPanel cp, JComponent comp, boolean fixHeight, 
                                           int inst, int insl, int insb, int insr, 
                                           int rowNum, int columns) {
    double vFac = (fixHeight) ? 0.0 : 1.0;
    UiUtil.gbcSet(gbc_, 0, rowNum++, columns, 1, UiUtil.HOR, 0, 0, inst, insl, insb, insr, UiUtil.CEN, 1.0, vFac);
    cp.add(comp, gbc_);
    return (rowNum);
  } 

  /***************************************************************************
  **
  ** Add a full row component with a label
  */ 
  
  public int addLabeledWidget(JPanel cp, JLabel label, JComponent comp, boolean fixHeight, boolean flushLeft,
                                 int rowNum, int columns) {
    double vFac = (fixHeight) ? 0.0 : 1.0;
    int eorw = (flushLeft) ? UiUtil.W : UiUtil.E;
    UiUtil.gbcSet(gbc_, 0, rowNum, 1, 1, UiUtil.NONE, 0, 0, 5, 5, 5, 5, eorw, 0.0, vFac);
    cp.add(label, gbc_);
    UiUtil.gbcSet(gbc_, 1, rowNum++, columns - 1, 1, UiUtil.HOR, 0, 0, 5, 5, 5, 5, UiUtil.CEN, 1.0, vFac);    
    cp.add(comp, gbc_);        
    return (rowNum);
  }
  
  /***************************************************************************
  **
  ** Add a table
  */ 
  
  public int addTable(JPanel cp, JComponent tablePan, int rowHeight, int rowNum, int columns) {  
    UiUtil.gbcSet(gbc_, 0, rowNum, columns, rowHeight, UiUtil.BO, 0, 0, 5, 5, 5, 5, UiUtil.CEN, 1.0, 1.0);
    rowNum += rowHeight;
    cp.add(tablePan, gbc_);
    return (rowNum);
  }
  
  /***************************************************************************
  **
  ** Add a table
  */ 
  
  public int addTableNoInset(JPanel cp, JComponent tablePan, int rowHeight, int rowNum, int columns) {  
    UiUtil.gbcSet(gbc_, 0, rowNum, columns, rowHeight, UiUtil.BO, 0, 0, 0, 0, 0, 0, UiUtil.CEN, 1.0, 1.0);
    rowNum += rowHeight;
    cp.add(tablePan, gbc_);
    return (rowNum);
  } 
}
```
