## Supplementary material for "BioFabric Visualization of Network Alignments": BioFabric Source Code: EditableTable.java

```
/*
**    Copyright (C) 2003-2014 Institute for Systems Biology 
**                            Seattle, Washington, USA. 
**
**    This library is free software; you can redistribute it and/or
**    modify it under the terms of the GNU Lesser General Public
**    License as published by the Free Software Foundation; either
**    version 2.1 of the License, or (at your option) any later version.
**
**    This library is distributed in the hope that it will be useful,
**    but WITHOUT ANY WARRANTY; without even the implied warranty of
**    MERCHANTABILITY or FITNESS FOR A PARTICULAR PURPOSE.  See the GNU
**    Lesser General Public License for more details.
**
**    You should have received a copy of the GNU Lesser General Public
**    License along with this library; if not, write to the Free Software
**    Foundation, Inc., 59 Temple Place, Suite 330, Boston, MA  02111-1307  USA
*/

package org.systemsbiology.biofabric.ui.dialogs.utils;

import java.awt.BorderLayout;
import java.awt.GridBagConstraints;
import java.awt.GridBagLayout;
import java.awt.Point;
import java.awt.Rectangle;
import java.awt.event.ActionListener;
import java.awt.event.ActionEvent;
import javax.swing.JPanel;
import javax.swing.Box;
import javax.swing.JTable;
import javax.swing.JScrollPane;
import java.util.ArrayList;
import java.util.HashMap;
import java.util.HashSet;
import java.util.Iterator;
import java.util.List;
import java.util.Map;
import java.util.Set;
import javax.swing.BorderFactory;
import javax.swing.JButton;
import javax.swing.JComponent;
import javax.swing.JFrame;
import javax.swing.JViewport;
import javax.swing.table.AbstractTableModel;
import javax.swing.ListSelectionModel;
import javax.swing.event.ListSelectionListener;
import javax.swing.event.ListSelectionEvent;
import javax.swing.table.DefaultTableCellRenderer;
import javax.swing.table.JTableHeader;
import javax.swing.table.TableCellEditor;
import javax.swing.table.TableCellRenderer;
import javax.swing.table.TableColumn;
import javax.swing.table.TableColumnModel;

import org.systemsbiology.biofabric.api.util.ExceptionHandler;
import org.systemsbiology.biofabric.api.util.FixedJButton;
import org.systemsbiology.biofabric.util.ComboBoxEditorTracker;
import org.systemsbiology.biofabric.util.ComboFinishedTracker;
import org.systemsbiology.biofabric.util.DoubleEditor;
import org.systemsbiology.biofabric.util.EditableComboBoxEditorTracker;
import org.systemsbiology.biofabric.util.EnumCell;
import org.systemsbiology.biofabric.util.IntegerEditor;
import org.systemsbiology.biofabric.util.NumberEditorTracker;
import org.systemsbiology.biofabric.util.ProtoDouble;
import org.systemsbiology.biofabric.util.ProtoInteger;
import org.systemsbiology.biofabric.util.ResourceManager;
import org.systemsbiology.biofabric.util.TextEditor;
import org.systemsbiology.biofabric.util.TextEditorTracker;
import org.systemsbiology.biofabric.util.TextFinishedTracker;
import org.systemsbiology.biofabric.util.TrackingUnit;
import org.systemsbiology.biofabric.util.UiUtil;

/****************************************************************************
**
** This is an editable table.  Rows are not sortable. The generic class is
** parameterized on the table row type.
*/

public class EditableTable<T extends EditableTable.ATableRow> {

  ////////////////////////////////////////////////////////////////////////////
  //
  // PUBLIC INTERFACE
  //
  ////////////////////////////////////////////////////////////////////////////   

  /***************************************************************************
  **
  ** Used for TableRows
  */

  public interface ATableRow {
    // Should also implement constructors for () and (int i)
    void toCols();
  }  
  
  ////////////////////////////////////////////////////////////////////////////
  //
  // PUBLIC CONSTANTS
  //
  ////////////////////////////////////////////////////////////////////////////   
  
  public final static int NO_BUTTONS    = 0x00;
  public final static int ADD_BUTTON    = 0x01;
  public final static int EDIT_BUTTON   = 0x02;
  public final static int DELETE_BUTTON = 0x04;
  public final static int RAISE_BUTTON  = 0x08; 
  public final static int LOWER_BUTTON  = 0x10;
  public final static int RAISE_AND_LOWER_BUTTONS = RAISE_BUTTON | LOWER_BUTTON;
  public final static int ALL_BUT_EDIT_BUTTONS = ADD_BUTTON | RAISE_AND_LOWER_BUTTONS | DELETE_BUTTON;
  public final static int ALL_BUTTONS = ALL_BUT_EDIT_BUTTONS | EDIT_BUTTON;
    
  ////////////////////////////////////////////////////////////////////////////
  //
  // PRIVATE INSTANCE MEMBERS
  //
  ////////////////////////////////////////////////////////////////////////////  
  
  private TableModel<T> estm_;
  private int[] selectedRows_;
  private boolean trueButtonDEnable_;
  private boolean trueButtonAEnable_;
  private boolean trueButtonREnable_;
  private boolean trueButtonLEnable_;
  private boolean trueButtonEEnable_;  
  private JButton tableButtonD_;
  private JButton tableButtonR_;
  private JButton tableButtonL_;  
  private JButton tableButtonA_;  
  private JButton tableButtonE_;    
  private TrackingJTable qt_;
  private SelectionTracker tracker_;
  private boolean addAlwaysAtEnd_;
  private TextEditor textEdit_;
  private Set<Class<?>> usedClasses_;
  private HashMap<Integer, TableCellEditor> editors_;
  private HashMap<Integer, TableCellRenderer> renderers_;
  private HashSet<Integer> editableEnums_;
  private JFrame parent_;
  private boolean cancelEditOnDisable_;
  private EditButtonHandler ebh_;
  
  ////////////////////////////////////////////////////////////////////////////
  //
  // PUBLIC CONSTRUCTORS
  //
  ////////////////////////////////////////////////////////////////////////////   
  
  
  public EditableTable(TableModel<T> atm, JFrame parent) {
    estm_ = atm;
    parent_ = parent;
  }
  
  ////////////////////////////////////////////////////////////////////////////
  //
  // PUBLIC METHODS
  //
  ////////////////////////////////////////////////////////////////////////////   
  

  public JTable getTable() {
    return (qt_);
  }

  public TableModel<T> getModel() {
    return (estm_);
  }
  
  /***************************************************************************
  **
  ** Get the selected rows
  */   

  public int[] getSelectedRows() {
    return (selectedRows_);
  } 

  /***************************************************************************
  **
  ** Install edit button handler
  */   

  public void setEditButtonHandler(EditButtonHandler ebh) {
    ebh_ = ebh;
    return;
  }
  
  /***************************************************************************
  **
  ** Enable or disable the table
  */   

  public void setEnabled(boolean enable) {
    qt_.setEnabled(enable);    
   
    //
    // Handle editing changes:
    //
    
    if (!enable) {
      stopTheEditing(cancelEditOnDisable_);
    }

    ((DefaultTableCellRenderer)qt_.getTableHeader().getDefaultRenderer()).setEnabled(enable);
    
    enableDefaultEditorsAndRenderers(enable);
    enableColumnEditorsAndRenderers(enable);
  
    //
    // Gathered up from TimeStage setup: use always?
    //
    
    qt_.getSelectionModel().clearSelection();
    
    //
    // Handle rendering changes:
    //
    
    if (!enable) {
      if (tableButtonA_ != null) tableButtonA_.setEnabled(false);
      if (tableButtonD_ != null) tableButtonD_.setEnabled(false);
      if (tableButtonR_ != null) tableButtonR_.setEnabled(false);
      if (tableButtonL_ != null) tableButtonL_.setEnabled(false);
      if (tableButtonE_ != null) tableButtonE_.setEnabled(false);
    } else {
      syncButtons();
    }
   
    qt_.getTableHeader().repaint();
    qt_.repaint();
    return;
  }
  
  /***************************************************************************
  **
  ** Sync buttons to true allowed state
  */   

  private void syncButtons() {
    if (tableButtonA_ != null) {
      tableButtonA_.setEnabled(trueButtonAEnable_);
      tableButtonA_.revalidate();
    }
    if (tableButtonD_ != null) {
      tableButtonD_.setEnabled(trueButtonDEnable_);
      tableButtonD_.revalidate();
    }
    if (tableButtonR_ != null) {
      tableButtonR_.setEnabled(trueButtonREnable_);
      tableButtonR_.revalidate();
    }
    if (tableButtonL_ != null) {
      tableButtonL_.setEnabled(trueButtonLEnable_);
      tableButtonL_.revalidate();  
    }
    if (tableButtonE_ != null) {
      tableButtonE_.setEnabled(trueButtonEEnable_);
      tableButtonE_.revalidate();  
    }
    return;
  }

  /***************************************************************************
  **
  ** Update the table
  */   

  public void updateTable(boolean fireChange, List<T> elements) {
    estm_.extractValues(elements);
   // estm_.modifyMap(estm_.buildFullClickList());
    if (fireChange) {
      estm_.fireTableDataChanged();
    }
    if (estm_.getRowCount() > 0) {
      qt_.getSelectionModel().setSelectionInterval(0, 0);
    } else {
      selectedRows_ = new int[0];
      trueButtonAEnable_ = estm_.canAddRow();
      trueButtonDEnable_ = false;
      trueButtonREnable_ = false;
      trueButtonLEnable_ = false;
      trueButtonEEnable_ = false;      
      syncButtons();      
    }
    return;
  }
  
  /***************************************************************************
  **
  ** Get stuff out of the table
  */   

  public List<T> getValuesFromTable() {
    return (estm_.getValuesFromTable());
  }

  /***************************************************************************
  **
  ** Clear current selections
  */   

  public void clearSelections() {    
    tracker_.setIgnore(true);
    qt_.getSelectionModel().clearSelection();
    selectedRows_ = null;
    tracker_.setIgnore(false);        
    return;
  }

  /***************************************************************************
  **
  ** Make selected row visible
  ** 
  */

  public void makeCurrentSelectionVisible() { 
    makeSelectionVisible(qt_.getSelectionModel().getMinSelectionIndex());
    return;
  }

  /***************************************************************************
  **
  ** Make specified row visible
  ** 
  */

  public void makeSelectionVisible(int selRow) { 
    JViewport viewport = (JViewport)qt_.getParent();
    Rectangle rect = qt_.getCellRect(selRow, 0, true);
    Point pt = viewport.getViewPosition();
    rect.setLocation(rect.x - pt.x, rect.y - pt.y);
    viewport.scrollRectToVisible(rect);
    viewport.validate();
    return;
  }

  /***************************************************************************
  **
  ** Used to build and install a JTable using the model
  */

  public JPanel buildEditableTable(TableParams etp) {
    
    usedClasses_ = estm_.usedClasses();   
    selectedRows_ = new int[0];
    addAlwaysAtEnd_ = etp.addAlwaysAtEnd;
    cancelEditOnDisable_ = etp.cancelEditOnDisable;
    qt_ = new TrackingJTable(estm_);
    estm_.setJTable(qt_);
    JTableHeader th = qt_.getTableHeader();
    // Not sorting these days:
    //th.addMouseListener(new ReadOnlyStringTableModel.SortTrigger(this));    
    th.setReorderingAllowed(false);

    tracker_ = new SelectionTracker();
    ListSelectionModel lsm = qt_.getSelectionModel();
    if (etp.singleSelectOnly) {
      lsm.setSelectionMode(ListSelectionModel.SINGLE_SELECTION);
    }
    lsm.addListSelectionListener(tracker_);
    if (etp.tableIsUnselectable) {
      qt_.setRowSelectionAllowed(false);
    }
 
    //
    // Set specialty widths:
    //

    if (etp.colWidths != null) {
      int numC = etp.colWidths.size();
      TableColumnModel tcm = qt_.getColumnModel();
      for (int i = 0; i < numC; i++) {
        ColumnWidths cw = etp.colWidths.get(i); 
        int viewColumn = qt_.convertColumnIndexToView(cw.colNum);
        TableColumn tfCol = tcm.getColumn(viewColumn);
        tfCol.setMinWidth(cw.min);
        tfCol.setPreferredWidth(cw.pref);
        tfCol.setMaxWidth(cw.max);
      }
    }

    UiUtil.installDefaultCellRendererForPlatform(qt_, String.class, true);
    UiUtil.installDefaultCellRendererForPlatform(qt_, ProtoInteger.class, true);
    UiUtil.installDefaultCellRendererForPlatform(qt_, Integer.class, true); // For non-edit number columns!
    
    editors_ = new HashMap<Integer, TableCellEditor>();
    renderers_ = new HashMap<Integer, TableCellRenderer>();
    editableEnums_ = new HashSet<Integer>();
    setDefaultEditors();   
    if (etp.perColumnEnums != null) {  
      setColumnEditorsAndRenderers(etp.perColumnEnums, renderers_, editors_, editableEnums_);
      estm_.setEnums(renderers_, editors_, editableEnums_);
    }

    UiUtil.platformTableRowHeight(qt_, true);
    
    return ((etp.buttonsOnSide) ? addButtonsOnSide(etp) : addButtonsOnBottom(etp));
  }

  
  /***************************************************************************
  **
  ** Refresh editors and renderers
  */       
 
  public void refreshEditorsAndRenderers(Map<Integer, EnumCellInfo> perColumnEnums) { 
    editors_.clear();
    renderers_.clear();
    editableEnums_.clear();
    setDefaultEditors();   
    if (perColumnEnums != null) {  
      setColumnEditorsAndRenderers(perColumnEnums, renderers_, editors_, editableEnums_);
      estm_.setEnums(renderers_, editors_, editableEnums_);
    }
    estm_.refreshColumnEditorsAndRenderers();
    return;
  }
    
  /***************************************************************************
  **
  ** Actually make the buttons
  */       
 
  private void createButtons(TableParams etp) { 
    
    ResourceManager rMan = ResourceManager.getManager(); 
    if (etp.buttons != NO_BUTTONS) {
      if ((etp.buttons & ADD_BUTTON) != 0x00) {
        String aTag = rMan.getString("dialogs.addEntry");
        tableButtonA_ = (etp.buttonsOnSide) ? new JButton(aTag) : new FixedJButton(aTag);
        tableButtonA_.addActionListener(new ActionListener() {
          public void actionPerformed(ActionEvent ev) {
            try {
              stopTheEditing(false);
              int[] addAt = (addAlwaysAtEnd_) ? new int[0] : selectedRows_;
              if (estm_.addRow(addAt)) {
                qt_.revalidate();
              }
            } catch (Exception ex) {
              ExceptionHandler.getHandler().displayException(ex);
            }
          }
        });
        //
        // FIX ME:  Should start off false, since table might not allow an add.  But too
        // many existing clients depend on this.  They just extract values, without calling
        // to update the table.  That needs to be fixed!
        trueButtonAEnable_ = true;
        tableButtonA_.setEnabled(true);
      }

      if ((etp.buttons & LOWER_BUTTON) != 0x00) {
        String lTag = rMan.getString("dialogs.lowerEntry");
        tableButtonL_ = (etp.buttonsOnSide) ? new JButton(lTag) : new FixedJButton(lTag);
        tableButtonL_.addActionListener(new ActionListener() {
          public void actionPerformed(ActionEvent ev) {
            try {
              stopTheEditing(false);
              estm_.bumpRowDown(selectedRows_);
              selectedRows_[0]++;
              qt_.getSelectionModel().setSelectionInterval(selectedRows_[0], selectedRows_[0]);
              qt_.revalidate();
            } catch (Exception ex) {
              ExceptionHandler.getHandler().displayException(ex);
            }
          }
        });
        tableButtonL_.setEnabled(false);
        trueButtonLEnable_ = false;
      }
      
      if ((etp.buttons & RAISE_BUTTON) != 0x00) {
        String rTag = rMan.getString("dialogs.raiseEntry");
        tableButtonR_ = (etp.buttonsOnSide) ? new JButton(rTag) : new FixedJButton(rTag);
        tableButtonR_.addActionListener(new ActionListener() {
          public void actionPerformed(ActionEvent ev) {
            try {
              stopTheEditing(false);
              estm_.bumpRowUp(selectedRows_);
              selectedRows_[0]--;
              qt_.getSelectionModel().setSelectionInterval(selectedRows_[0], selectedRows_[0]);
              qt_.revalidate();
            } catch (Exception ex) {
              ExceptionHandler.getHandler().displayException(ex);
            }
          }
        });
        tableButtonR_.setEnabled(false);
        trueButtonREnable_ = false;
      }

      if ((etp.buttons & DELETE_BUTTON) != 0x00) {
        String dTag = rMan.getString("dialogs.deleteEntry");
        tableButtonD_ = (etp.buttonsOnSide) ? new JButton(dTag) : new FixedJButton(dTag);
        tableButtonD_.addActionListener(new ActionListener() {
          public void actionPerformed(ActionEvent ev) {
            try {
              stopTheEditing(true);
              estm_.deleteRows(selectedRows_);
              qt_.getSelectionModel().clearSelection();        
              qt_.revalidate();
            } catch (Exception ex) {
              ExceptionHandler.getHandler().displayException(ex);
            }
          }
        });
        tableButtonD_.setEnabled(false);
        trueButtonDEnable_ = false;
      }
      
      if ((etp.buttons & EDIT_BUTTON) != 0x00) {
        String eTag = rMan.getString("dialogs.editEntry");
        tableButtonE_ = (etp.buttonsOnSide) ? new JButton(eTag) : new FixedJButton(eTag);
        tableButtonE_.addActionListener(new ActionListener() {
          public void actionPerformed(ActionEvent ev) {
            try {
              stopTheEditing(false);
              ebh_.pressed();
              qt_.revalidate();
            } catch (Exception ex) {
              ExceptionHandler.getHandler().displayException(ex);
            }
          }
        });
        tableButtonE_.setEnabled(false);
        trueButtonEEnable_ = false;
      }
    } 
    return;
  }  
  
 /***************************************************************************
  **
  ** Get buttons on the bottom
  */       
 
  private JPanel startPanel(TableParams etp) { 
    
    //
    // No scroll only implemented for NO_BUTTONS case!
    //
    
    if ((etp.buttons != NO_BUTTONS) && (etp.noScroll)) {
      throw new IllegalStateException();
    }
    
    JPanel retval = new JPanel();
    retval.setLayout(new GridBagLayout());
    GridBagConstraints gbc = new GridBagConstraints();   
    
    if (etp.noScroll) {
      JTableHeader jth = qt_.getTableHeader();
      retval.setBorder(BorderFactory.createEtchedBorder());
      retval.setLayout(new BorderLayout()); 
      retval.add(jth, BorderLayout.NORTH); 
      retval.add(qt_, BorderLayout.CENTER);        
    } else {
      JScrollPane jsp = new JScrollPane(qt_);
      UiUtil.gbcSet(gbc, 0, 0, 1, 10, UiUtil.BO, 0, 0, 5, 5, 5, 5, UiUtil.CEN, 1.0, 1.0);
      retval.add(jsp, gbc);
    }
    return (retval);
  }

  
  /***************************************************************************
  **
  ** Get buttons on the bottom
  */       
 
  private JPanel addButtonsOnBottom(TableParams etp) { 
    
    JPanel retval = startPanel(etp);
    
    if (etp.buttons != NO_BUTTONS) {
      GridBagConstraints gbc = new GridBagConstraints();
      createButtons(etp);
      int buttonCount = 0;
      
      Box tableButtonPanel = Box.createHorizontalBox();
      tableButtonPanel.add(Box.createHorizontalGlue()); 
      if ((etp.buttons & ADD_BUTTON) != 0x00) {
        tableButtonPanel.add(tableButtonA_);
        buttonCount++;
      }
      if ((etp.buttons & LOWER_BUTTON) != 0x00) {
        if (buttonCount != 0) {
          tableButtonPanel.add(Box.createHorizontalStrut(10));
        }
        tableButtonPanel.add(tableButtonL_);
        buttonCount++;
      }
       if ((etp.buttons & RAISE_BUTTON) != 0x00) {
        if (buttonCount != 0) {
          tableButtonPanel.add(Box.createHorizontalStrut(10));
        }
        tableButtonPanel.add(tableButtonR_);
        buttonCount++;
      }
      if ((etp.buttons & DELETE_BUTTON) != 0x00) {
        if (buttonCount != 0) {
          tableButtonPanel.add(Box.createHorizontalStrut(10));
        }
        tableButtonPanel.add(tableButtonD_);
        buttonCount++;
      }
      if ((etp.buttons & EDIT_BUTTON) != 0x00) {
        if (buttonCount != 0) {
          tableButtonPanel.add(Box.createHorizontalStrut(10));
        }
        tableButtonPanel.add(tableButtonE_);
        buttonCount++;
      }
      tableButtonPanel.add(Box.createHorizontalGlue());
 
      UiUtil.gbcSet(gbc, 0, 10, 1, 1, UiUtil.HOR, 0, 0, 5, 5, 5, 5, UiUtil.SE, 1.0, 0.0);
      retval.add(tableButtonPanel, gbc);
    }
    return (retval);
  }

  /***************************************************************************
  **
  ** Put the buttons on the side
  */       
 
  private JPanel addButtonsOnSide(TableParams etp) { 
    
    JPanel retval = startPanel(etp);
    
    if (etp.buttons != NO_BUTTONS) {
      GridBagConstraints gbc = new GridBagConstraints();
      createButtons(etp);
      int buttonCount = 0;  
      JPanel tableButtonPanel = new JPanel();
      tableButtonPanel.setLayout(new GridBagLayout());
      if ((etp.buttons & ADD_BUTTON) != 0x00) {
        UiUtil.gbcSet(gbc, 0, buttonCount++, 1, 1, UiUtil.HOR, 0, 0, 5, 5, 5, 5, UiUtil.N, 0.0, 0.0);
        tableButtonPanel.add(tableButtonA_, gbc);
      }
      if ((etp.buttons & LOWER_BUTTON) != 0x00) {
        UiUtil.gbcSet(gbc, 0, buttonCount++, 1, 1, UiUtil.HOR, 0, 0, 5, 5, 5, 5, UiUtil.N, 0.0, 0.0);
        tableButtonPanel.add(tableButtonL_, gbc);
      }
       if ((etp.buttons & RAISE_BUTTON) != 0x00) {
        UiUtil.gbcSet(gbc, 0, buttonCount++, 1, 1, UiUtil.HOR, 0, 0, 5, 5, 5, 5, UiUtil.N, 0.0, 0.0);
        tableButtonPanel.add(tableButtonR_, gbc);
      }
      if ((etp.buttons & DELETE_BUTTON) != 0x00) {
        UiUtil.gbcSet(gbc, 0, buttonCount++, 1, 1, UiUtil.HOR, 0, 0, 5, 5, 5, 5, UiUtil.N, 0.0, 0.0);
        tableButtonPanel.add(tableButtonD_, gbc);
      }
      if ((etp.buttons & EDIT_BUTTON) != 0x00) {
        UiUtil.gbcSet(gbc, 0, buttonCount++, 1, 1, UiUtil.HOR, 0, 0, 5, 5, 5, 5, UiUtil.N, 0.0, 0.0);
        tableButtonPanel.add(tableButtonE_, gbc);
      }

      UiUtil.gbcSet(gbc, 2, 0, 1, 1, UiUtil.VERT, 0, 0, 5, 5, 5, 5, UiUtil.CEN, 0.0, 1.0);
      retval.add(tableButtonPanel, gbc);
    }
    return (retval);
  }

  /***************************************************************************
  **
  ** Set the default editors
  ** 
  */

  private void setDefaultEditors() {
    Iterator<Class<?>> cit = usedClasses_.iterator();
    while (cit.hasNext()) {
      Class<?> used = cit.next();
      if (used.equals(ProtoDouble.class)) {
        DoubleEditor dEdit = new DoubleEditor(parent_);
        new NumberEditorTracker(dEdit, estm_);
        qt_.setDefaultEditor(ProtoDouble.class, dEdit);
      } else if (used.equals(String.class)) {
        textEdit_ = new TextEditor();
        new TextEditorTracker(textEdit_, estm_, estm_);
        qt_.setDefaultEditor(String.class, textEdit_);
      } else if (used.equals(ProtoInteger.class)) {
        IntegerEditor intEdit = new IntegerEditor(parent_);
        new NumberEditorTracker(intEdit, estm_);
        qt_.setDefaultEditor(ProtoInteger.class, intEdit);
      } else if (used.equals(Boolean.class)) {
        // Happy with the default!
       } else if (used.equals(Integer.class)) {
        // Happy with the default!  (actually gotta be non-editable!)
      } else if (!used.equals(EnumCell.class)) {
        throw new IllegalStateException();
      }
    }
    return;
  }  
  
  /***************************************************************************
  **
  ** Set the editor/renderer enabled state
  ** 
  */

  private void enableDefaultEditorsAndRenderers(boolean enable) {
    Iterator<Class<?>> cit = usedClasses_.iterator();
    while (cit.hasNext()) {
      Class<?> used = cit.next();
      if (used.equals(ProtoDouble.class)) {
        DoubleEditor dt = (DoubleEditor)qt_.getDefaultEditor(ProtoDouble.class);
        dt.setEnabled(enable);
        dt.setEditable(enable);
        // Sometimes a TrackingDoubleRenderer, not a default renderer:
        JComponent dtcr = (JComponent)qt_.getDefaultRenderer(ProtoDouble.class);
        dtcr.setEnabled(enable);
      } else if (used.equals(String.class)) {
        TextEditor te = (TextEditor)qt_.getDefaultEditor(String.class);
        te.setEnabled(enable);
        te.setEditable(enable);
        DefaultTableCellRenderer dtcr = (DefaultTableCellRenderer)qt_.getDefaultRenderer(String.class);
        dtcr.setEnabled(enable);
      } else if (used.equals(ProtoInteger.class)) {
        IntegerEditor ie = (IntegerEditor)qt_.getDefaultEditor(ProtoInteger.class);
        ie.setEnabled(enable);
        ie.setEditable(enable);
        DefaultTableCellRenderer dtcr = (DefaultTableCellRenderer)qt_.getDefaultRenderer(ProtoInteger.class);
        dtcr.setEnabled(enable);
      } else if (used.equals(Boolean.class)) {
        // leave alone!
      } else if (used.equals(Integer.class)) {
        // leave alone!
      } else if (!used.equals(EnumCell.class)) {
        throw new IllegalStateException();
      }
    }
    return;
  }  

  /***************************************************************************
  **
  ** Set the enabled/disabled state for the editors and renderers for specific columns
  ** 
  */

  private void enableColumnEditorsAndRenderers(boolean enable) {
    Iterator<Integer> rkit = renderers_.keySet().iterator();
    while (rkit.hasNext()) {
      Integer key = rkit.next();
      if (editableEnums_.contains(key)) {
        EnumCell.EditableComboBoxRenderer ecbr = (EnumCell.EditableComboBoxRenderer)renderers_.get(key);
        ecbr.setEnabled(enable);
      } else {
        EnumCell.ReadOnlyComboBoxRenderer rocbr = (EnumCell.ReadOnlyComboBoxRenderer)renderers_.get(key);
        rocbr.setEnabled(enable);
      }
    }
    Iterator<Integer> ekit = editors_.keySet().iterator();
    while (ekit.hasNext()) {
      Integer key = ekit.next();
      if (editableEnums_.contains(key)) {
        EnumCell.EditableEnumCellEditor eece = (EnumCell.EditableEnumCellEditor)editors_.get(key);
        eece.setEnabled(enable);
        eece.setEditable(enable);
      } else {
        EnumCell.ReadOnlyEnumCellEditor roene = (EnumCell.ReadOnlyEnumCellEditor)editors_.get(key);
        roene.setEnabled(enable);
      }
    }
    return;
  }
  
  /***************************************************************************
  **
  ** Set the editors and renderers for specific columns
  ** 
  */

  private void setColumnEditorsAndRenderers(Map<Integer, EnumCellInfo> contents, Map<Integer, TableCellRenderer> renderers, 
                                            Map<Integer, TableCellEditor> editors, Set<Integer> editableEnums) {  
    Iterator<Integer> cit = contents.keySet().iterator();
    while (cit.hasNext()) {
      Integer intObj = cit.next();
      EnumCellInfo eci = contents.get(intObj);
      TableColumn tc = qt_.getColumnModel().getColumn(intObj.intValue());   
      if (eci.editable) {
        EnumCell.EditableComboBoxRenderer ecbr = new EnumCell.EditableComboBoxRenderer(eci.values);
        tc.setCellRenderer(ecbr);
        renderers.put(intObj, ecbr);
        
        EnumCell.EditableEnumCellEditor re = new EnumCell.EditableEnumCellEditor(eci.values);
        new EditableComboBoxEditorTracker(re, estm_);
        tc.setCellEditor(re);
        editors.put(intObj, re);
        editableEnums.add(intObj);
      } else {
        EnumCell.ReadOnlyComboBoxRenderer rocbr;
        if (eci.trackingUnits == null) {
          rocbr = new EnumCell.ReadOnlyComboBoxRenderer(eci.values);
        } else {
          rocbr = new EnumCell.TrackingReadOnlyComboBoxRenderer(eci.values, eci.trackingUnits);
        }
        tc.setCellRenderer(rocbr);
        renderers.put(intObj, rocbr);
        
        EnumCell.ReadOnlyEnumCellEditor me = new EnumCell.ReadOnlyEnumCellEditor(eci.values);
        new ComboBoxEditorTracker(me, estm_, estm_);     
        tc.setCellEditor(me);
        editors.put(intObj, me);
      }
    }
    return;
  }
 
  /***************************************************************************
  **
  ** Stop all table editing.
  ** 
  */

  public void stopTheEditing(boolean doCancel) {
    Iterator<Class<?>> cit = usedClasses_.iterator();
    while (cit.hasNext()) {
      Class<?> used = cit.next();
      if (!used.equals(EnumCell.class)) {
        // Ignore default editors:
        if (!used.equals(Integer.class) && !used.equals(Boolean.class)) {
          TableCellEditor tce = qt_.getDefaultEditor(used);
          if (doCancel) {
            tce.cancelCellEditing();
          } else {
            tce.stopCellEditing();
          }
        }
      }
    }
    Iterator<Integer> ekit = editors_.keySet().iterator();
    while (ekit.hasNext()) {
      Integer key = ekit.next();
      TableCellEditor tce = editors_.get(key);
      if (doCancel) {
        tce.cancelCellEditing();
      } else {
        tce.stopCellEditing();
      }     
    }
    return;
  }  
  
  ////////////////////////////////////////////////////////////////////////////
  //
  // PUBLIC INNER CLASSES
  //
  ////////////////////////////////////////////////////////////////////////////

  /***************************************************************************
  **
  ** Used for tracking button presses
  */

  public static interface EditButtonHandler {
    public void pressed();
  }  
 
  /***************************************************************************
  **
  ** Used for setup
  */
  
  public static class TableParams {
    public boolean addAlwaysAtEnd;
    public boolean singleSelectOnly;
    public boolean cancelEditOnDisable;
    public boolean tableIsUnselectable;
    public int buttons;
    public Map<Integer, EnumCellInfo> perColumnEnums;
    public List<ColumnWidths> colWidths;
    public boolean buttonsOnSide;
    public boolean noScroll;
  }
  
  /***************************************************************************
  **
  ** Used for setup
  */
  
  public static class ColumnWidths {
    public int colNum;
    public int min;
    public int pref;
    public int max;
    
    public ColumnWidths() {   
    }
    
    public ColumnWidths(int colNum, int min, int pref, int max) {
      this.colNum = colNum;
      this.min = min;
      this.pref = pref;
      this.max = max;
    }     
  }
  
  /***************************************************************************
  **
  ** Used for enum cell definitions
  */
  
  public static class EnumCellInfo {
    public boolean editable;
    public List<EnumCell> values;
    public Map<Integer, TrackingUnit> trackingUnits;
    
    public EnumCellInfo(boolean editable, List<EnumCell> values) {
      this(editable, values, null);
    }
    
    public EnumCellInfo(boolean editable, List<EnumCell> values, Map<Integer, TrackingUnit> trackingUnits) {
      this.editable = editable;
      this.values = values;
      this.trackingUnits = trackingUnits;
      if ((trackingUnits != null) && editable) {
        throw new IllegalArgumentException();
      }
    }
  }
  
  /***************************************************************************
  **
  ** Used for the data table renderer
  */
  
  
  public static abstract class TableModel<T extends ATableRow> extends AbstractTableModel implements ComboFinishedTracker, TextFinishedTracker {

    protected JTable tmqt_;
    protected ArrayList<List<Object>> columns_;
    protected ArrayList<Integer> origIndex_;
    protected ArrayList<List<Object>> hiddenColumns_;
    protected int rowCount_;
    protected String[] colNames_;
    protected Class<?>[] colClasses_;
    protected boolean[] canEdit_;
    protected boolean[] colUseDirect_;
    protected Set<Integer> myEditableEnums_;
    protected HashMap<Integer, TableCellRenderer> myRenderers_;
    protected HashMap<Integer, TableCellEditor> myEditors_;
    protected int collapseCount_;
    protected boolean isCollapsed_;
    private static final long serialVersionUID = 1L;

    protected TableModel(int colNum) {
      columns_ = new ArrayList<List<Object>>();
      for (int i = 0; i < colNum; i++) {
        columns_.add(new ArrayList<Object>());
      }
      origIndex_ = new ArrayList<Integer>();
      hiddenColumns_ = new ArrayList<List<Object>>();
      canEdit_ = null;
      colUseDirect_ = null;
      collapseCount_ = -1;
      isCollapsed_ = false;
      myRenderers_ = new HashMap<Integer, TableCellRenderer>();
      myEditors_ = new HashMap<Integer, TableCellEditor>();
    }
    
    
    //
    // Warning! Only useful in special circumstances!
    //
    
    protected void resetColumnCount(int colNum) {
      columns_ = new ArrayList<List<Object>>();
      for (int i = 0; i < colNum; i++) {
        columns_.add(new ArrayList<Object>());
      }
    } 
    
    protected void addHiddenColumns(int colNum) {
      hiddenColumns_ = new ArrayList<List<Object>>();
      for (int i = 0; i < colNum; i++) {
        hiddenColumns_.add(new ArrayList<Object>());
      }
      return;
    }

    Set<Class<?>> usedClasses() {
      HashSet<Class<?>> retval = new HashSet<Class<?>>();
      for (int i = 0; i < colClasses_.length; i++) {
        retval.add(colClasses_[i]);
      }
      return (retval);
    }
    
    
    void setJTable(JTable tab) {
      tmqt_ = tab; 
    }
    
    void setEnums(Map<Integer, TableCellRenderer> renderers, Map<Integer, TableCellEditor> editors, Set<Integer> editableEnums) {
      myRenderers_.clear();
      myEditors_.clear();
      myRenderers_.putAll(renderers);
      myEditors_.putAll(editors);
      myEditableEnums_ = editableEnums;
      return;
    }

    public void deleteRows(int[] rows) {
      for (int i = 0; i < rows.length; i++) {
        int clen = columns_.size();
        for (int j = 0; j < clen; j++) {
          columns_.get(j).remove(rows[i]);
        }
        int hlen = hiddenColumns_.size();
        for (int j = 0; j < hlen; j++) {
          hiddenColumns_.get(j).remove(rows[i]);
        }
        origIndex_.remove(rows[i]);
        for (int j = i + 1; j < rows.length; j++) {
          if (rows[j] > rows[i]) {
            rows[j]--;
          }
        }
        rowCount_--;
      }
      return; 
    }

    public void bumpRowUp(int[] rows) {
      if ((rows.length != 1) || (rows[0] == 0)) {
        throw new IllegalStateException();
      }
      int clen = columns_.size();
      for (int j = 0; j < clen; j++) {
        Object remObj = columns_.get(j).remove(rows[0] - 1);
        columns_.get(j).add(rows[0], remObj);
      }
      int hlen = hiddenColumns_.size();
      for (int j = 0; j < hlen; j++) {
        Object remObj = hiddenColumns_.get(j).remove(rows[0] - 1);
        hiddenColumns_.get(j).add(rows[0], remObj);
      }
      Integer remObj = origIndex_.remove(rows[0] - 1);
      origIndex_.add(rows[0], remObj);
      return;
    }

    public void bumpRowDown(int[] rows) {
      if ((rows.length != 1) || (rows[0] == (origIndex_.size() - 1))) {
        throw new IllegalStateException();
      }
      int clen = columns_.size();
      for (int j = 0; j < clen; j++) {
        Object remObj = columns_.get(j).remove(rows[0]);
        columns_.get(j).add(rows[0] + 1, remObj);
      }
      int hlen = hiddenColumns_.size();
      for (int j = 0; j < hlen; j++) {
        Object remObj = hiddenColumns_.get(j).remove(rows[0]);
        hiddenColumns_.get(j).add(rows[0] + 1, remObj);
      }
      Integer remObj = origIndex_.remove(rows[0]);
      origIndex_.add(rows[0] + 1, remObj);
      return;
    }    

    public boolean addRow(int[] rows) {
      if (rows.length == 0) {
        origIndex_.add(null);
      } else if (rows.length == 1) {
        origIndex_.add(rows[0] + 1, null);
      } else {
        throw new IllegalStateException(); 
      }
      int clen = columns_.size();
      for (int j = 0; j < clen; j++) {
        if (rows.length == 0) {
          columns_.get(j).add(null);
        } else {
          columns_.get(j).add(rows[0] + 1, null);
        }
      }
      int hlen = hiddenColumns_.size();
      for (int j = 0; j < hlen; j++) {
        if (rows.length == 0) {
          hiddenColumns_.get(j).add(null);
        } else {
          hiddenColumns_.get(j).add(rows[0] + 1, null);
        }
      } 
      rowCount_++;
      return (true); 
    } 

    public int getRowCount() {
      return (rowCount_); 
    }  

    public int getColumnCount() {
      if (collapseCount_ == -1) {
        return (columns_.size());
      }
      return ((isCollapsed_) ? collapseCount_ : columns_.size());
    }

    protected List<Object> getListAt(int c) {
      try {
        if (c >= columns_.size()) {
          throw new IllegalArgumentException();
        }
        return (columns_.get(c));
      } catch (Exception ex) {
        ExceptionHandler.getHandler().displayException(ex);
      }
      return (null);
    }

    // NOT SORTABLE (YET!)

    protected int mapSelectionIndex(int r) {
      return (r);
    }

    @Override
    public String getColumnName(int c) {
      try {
        if (c >= colNames_.length) {
          throw new IllegalArgumentException();
        }
        if ((colUseDirect_ == null) || !colUseDirect_[c]) {
          ResourceManager rMan = ResourceManager.getManager();
          return (rMan.getString(colNames_[c]));
        } else {
          return (colNames_[c]);
        }
      } catch (Exception ex) {
        ExceptionHandler.getHandler().displayException(ex);
      }
      return (null);
    }

    @Override
    public Class<?> getColumnClass(int c) {
      try {
        if (c >= colClasses_.length) {
          throw new IllegalArgumentException();
        }
        return (colClasses_[c]);
      } catch (Exception ex) {
        ExceptionHandler.getHandler().displayException(ex);
      }
      return (null);
    }

    public Object getValueAt(int r, int c) {
      try {
        List<Object> list = getListAt(c);
        if (list.isEmpty()) {
          return (null);
        }
        return (list.get(mapSelectionIndex(r)));
      } catch (Exception ex) {
        ExceptionHandler.getHandler().displayException(ex);
      }
      return (null);
    }

    @Override
    public void setValueAt(Object value, int r, int c) {
      try { 
        List<Object> list = getListAt(c);
        if (list.isEmpty()) {
          return;
        }
        Object currVal = list.get(r);
        Class<?> colClass = getColumnClass(c);
        if (colClass.equals(ProtoInteger.class)) {
          ProtoInteger currNum = (ProtoInteger)currVal;
          if ((currNum != null) && currNum.textValue.equals(((ProtoInteger)value).textValue)) {
            return;
          }
        } else if (colClass.equals(String.class)) {
          String currStr = (String)currVal;
          if ((currStr != null) && currStr.equals(value)) {
            return;
          }    
        } else if (colClass.equals(ProtoDouble.class)) {
          ProtoDouble currNum = (ProtoDouble)currVal;
          if ((currNum != null) && currNum.textValue.equals(((ProtoDouble)value).textValue)) {
            return;
          }
        } else if (colClass.equals(Boolean.class)) {  
          Boolean currInclude = (Boolean)currVal;
          if ((currInclude != null) && (currInclude.equals(value))) {
            return;
          } 
        } else if (colClass.equals(EnumCell.class)) {
          EnumCell currEC = (EnumCell)currVal;
          if (myEditableEnums_.contains(new Integer(c))) {
            if ((currEC != null) && (currEC.internal != null) &&
                currEC.internal.equals(((EnumCell)value).internal)) {
              return;
            }
          } else {
            if ((currEC != null) && (currEC.value == ((EnumCell)value).value)) {
              return;
            }
          }
        } else {
          throw new IllegalStateException();
        }
        list.set(r, value);
      } catch (Exception ex) {
        ExceptionHandler.getHandler().displayException(ex);
      }
      return;
    } 

    @Override
    public boolean isCellEditable(int r, int c) {
      try { 
        if (canEdit_ == null) {
          return (true);
        } else {
          return (canEdit_[c]);
        }
      } catch (Exception ex) {
        ExceptionHandler.getHandler().displayException(ex);
      }
      return (false);
    }

    public void extractValues(List<T> rowElements) {
      origIndex_.clear();
      int clen = columns_.size();
      for (int i = 0; i < clen; i++) {
        columns_.get(i).clear();
      }
      int hlen = hiddenColumns_.size();
      for (int i = 0; i < hlen; i++) {
        hiddenColumns_.get(i).clear();
      }
      rowCount_ = 0;
      Iterator<T> iit = rowElements.iterator();
      while (iit.hasNext()) {
        iit.next();
        origIndex_.add(new Integer(rowCount_++));
      }
      Iterator<T> rit = rowElements.iterator();
      while (rit.hasNext()) { 
        T ent = rit.next();
        ent.toCols();
      }     
      return;  
    }
    
    public abstract T constructARow();
    protected abstract T constructARow(int i);
     
    public List<T> getValuesFromTable() {
      ArrayList<T> retval = new ArrayList<T>();
      for (int i = 0; i < rowCount_; i++) {
        T ent = constructARow(i);
        retval.add(ent);
      }
      return (retval); 
    }
    
    public void collapseView(boolean doCollapse) {
      if (collapseCount_ == -1) {
        throw new IllegalStateException();
      }
      isCollapsed_ = doCollapse;
      fireTableStructureChanged();
      refreshColumnEditorsAndRenderers();
      return;
    }
   
    //
    // In non-editable enums (e.g. OneEnumTableModel), an empty enum list means
    // you can't add rows to the table.  Provide this hook.
    //
    
    public boolean canAddRow() {
      return (true);
    }
        
    /***************************************************************************
    **
    ** If columns change, we gotta reinstall!
    ** 
    */

    private void refreshColumnEditorsAndRenderers() {  
      int currColumns = getColumnCount();
      TableColumnModel tcm = tmqt_.getColumnModel();
 
      Iterator<Integer> rkit = myRenderers_.keySet().iterator();
      while (rkit.hasNext()) {
        Integer key = rkit.next();
        if (key.intValue() >= currColumns) {
          continue;
        }
        TableColumn tc = tcm.getColumn(key.intValue());  
        if (myEditableEnums_.contains(key)) {
          EnumCell.EditableComboBoxRenderer ecbr = (EnumCell.EditableComboBoxRenderer)myRenderers_.get(key);
          tc.setCellRenderer(ecbr);
        } else {
          EnumCell.ReadOnlyComboBoxRenderer rocbr = (EnumCell.ReadOnlyComboBoxRenderer)myRenderers_.get(key);
          tc.setCellRenderer(rocbr);
        }
      }
      
      Iterator<Integer> ekit = myEditors_.keySet().iterator();
      while (ekit.hasNext()) {
        Integer key = ekit.next();
        if (key.intValue() >= currColumns) {
          continue;
        }
        TableColumn tc = tcm.getColumn(key.intValue());
        if (myEditableEnums_.contains(key)) {
          EnumCell.EditableEnumCellEditor eece = (EnumCell.EditableEnumCellEditor)myEditors_.get(key);
          tc.setCellEditor(eece);
        } else {
          EnumCell.ReadOnlyEnumCellEditor roene = (EnumCell.ReadOnlyEnumCellEditor)myEditors_.get(key);
          tc.setCellEditor(roene);
        }
      }
      return;
    }

    //
    // These methods allow the table to revise its contents when a field is
    // finished editing!
    //

    public boolean textEditingDone(int col, int row, Object val) {
      return (true);
    }
    
    public boolean comboEditingDone(int col, int row, Object val) {
      return (true);
    }
      
  }
  
  /***************************************************************************
  **
  ** Used for the data table renderer
  */
  
  class SelectionTracker implements ListSelectionListener {
    
    private boolean ignore_;
    
    SelectionTracker() {
      ignore_ = false;
    }
    
    public void valueChanged(ListSelectionEvent lse) {
      try {
        if (ignore_) {
          return;
        }
        if (lse.getValueIsAdjusting()) {
          return;
        }
        selectedRows_ = qt_.getSelectedRows();
        Object lastOrig = qt_.getLastOrig();
        int col = qt_.getLastColumn();
        if (lastOrig != null) {
          EnumCell.EditableEnumCellEditor fixit = (EnumCell.EditableEnumCellEditor)editors_.get(new Integer(col));
          fixit.getTableCellEditorComponent(qt_, lastOrig, 
                                            true, selectedRows_[0], qt_.getLastColumn());
          qt_.clearLastOrig();
        }
        boolean haveARow = (selectedRows_.length > 0);
        trueButtonDEnable_ = haveARow;
        trueButtonEEnable_ = haveARow;
        boolean haveZeroOrOne = (selectedRows_.length <= 1);
        trueButtonAEnable_ = (addAlwaysAtEnd_ || haveZeroOrOne) && estm_.canAddRow();
        boolean canLower = ((selectedRows_.length == 1) && 
                            (selectedRows_[0] < estm_.getRowCount() - 1));        
        trueButtonLEnable_ = canLower;
        boolean canRaise = ((selectedRows_.length == 1) && (selectedRows_[0] != 0));        
        trueButtonREnable_ = canRaise;  
        syncButtons();
      } catch (Exception ex) {
        ExceptionHandler.getHandler().displayException(ex);
      }
      return;             
    } 
    
    public void setIgnore(boolean ignore) {
      ignore_ = ignore;
      return;
    }
  } 
  
  /***************************************************************************
  **
  ** 4/19/10: NASTY Editable combo box bug in base Swing table implementation.
  ** If we edit a field, e.g. "AAA3" to "AAA34", then click to the _same_ column
  ** with an identical entry, "AAA3", it incorrectly pops to "AAA34".  Various
  ** attempts to circumvent this have failed.  So we track the editor selections
  ** in the table, and if the current request matches the previous editor column,
  ** and it is an editable combo box, we see if the original value of the previous
  ** editor matches our current value.  If so, we will replace it back to the original
  ** then the list selection event comes in; this appears to be consistently after the
  ** error has been introduced. 
  ** 
  ** Used to track editing bug
  */
  
  class TrackingJTable extends JTable {
    
    private static final long serialVersionUID = 1L;
    
    private int lastCol_ = -1;
    private Object lastOrig_;
           
    TrackingJTable(TableModel<T> estm) {
      super(estm);
    }
    
    Object getLastOrig() {
      return (lastOrig_);
    }
    
    void clearLastOrig() {
      lastOrig_ = null;
      return;
    }
    
    int getLastColumn() {
      return (lastCol_);
    }
    
    @Override
    public TableCellEditor getCellEditor(int row, int column) {
      lastOrig_ = null;
      if ((lastCol_ == column) && editableEnums_.contains(new Integer(column))) {
        EnumCell.EditableEnumCellEditor fixit = (EnumCell.EditableEnumCellEditor)editors_.get(new Integer(column));
        if (((EnumCell)estm_.getValueAt(row, column)).display.equals(((EnumCell)fixit.getOriginalValue()).display)) {
          lastOrig_ = estm_.getValueAt(row, column);
        }
      }
      lastCol_ = column;
      return (super.getCellEditor(row, column));
    }
  }
  
  /***************************************************************************
  **
  ** A Concrete instantiation of the TableModel that supports a single value.
  */
  
  public static class OneValueModel extends EditableTable.TableModel<OneValueModel.TableRow> {
    
    private final static int VALUE   = 0;
    private final static int NUM_COL_ = 1;
    
    private static final long serialVersionUID = 1L;
    
    public class TableRow implements ATableRow {
      public String value;
  
      public TableRow() {        
      }
      
      TableRow(int i) {
        value = (String)columns_.get(VALUE).get(i);
      }
      
      public void toCols() {
        columns_.get(VALUE).add(value);  
        return;
      }
    }
 
    public OneValueModel(String title, boolean edit) {
      super(NUM_COL_);
      colNames_ = new String[] {title};
      colClasses_ = new Class[] {String.class};
      canEdit_ = new boolean[] {edit};
    }
 
    protected TableRow constructARow(int i) {
      return (new TableRow(i));
    }
    
    public TableRow constructARow() {
      return (new TableRow());
    }
    
  }
  
  /***************************************************************************
  **
  ** Useful for single tagged values
  */

  public static class TaggedValueModel extends EditableTable.TableModel<TaggedValueModel.TableRow> {
    
    private final static int VALUE    = 0;
    private final static int NUM_COL_ = 1;
    
    private final static int HIDDEN_     = 0;
    private final static int NUM_HIDDEN_ = 1;
    
    private static final long serialVersionUID = 1L;
    
    public class TableRow implements ATableRow {
      public String hidden;
      public String value;
      
      public TableRow() {
      }
      
      TableRow(int i) {
        hidden = (String)hiddenColumns_.get(HIDDEN_).get(i);
        value = (String)columns_.get(VALUE).get(i);
      }
      
      public void toCols() {
        columns_.get(VALUE).add(value);
        hiddenColumns_.get(HIDDEN_).add(hidden);
        return;
      }
    }
 
    public TaggedValueModel(String title, boolean editable) {
      super(NUM_COL_);
      colNames_ = new String[] {title};
      colClasses_ = new Class[] {String.class};
      canEdit_ = new boolean[] {editable};
      addHiddenColumns(NUM_HIDDEN_);
    }
 
    protected TableRow constructARow(int i) {
      return (new TableRow(i));
    }
    
    public TableRow constructARow() {
      return (new TableRow());
    }
    
  }
  
  /***************************************************************************
  **
  ** Used for single enum tables
  */
  
  public static class OneEnumTableModel extends EditableTable.TableModel<OneEnumTableModel.TableRow> {
    
    public final static int ENUM_COL_ = 0;
    private final static int NUM_COL_ = 1;
    
    private final static int ORIG_ORDER_ = 0;
    private final static int NUM_HIDDEN_ = 1;
    
    private ArrayList<EnumCell> currEnums_;
    private static final long serialVersionUID = 1L;
    
    public class TableRow implements ATableRow {
      public Integer origOrder;
      public EnumCell enumChoice;
      
      public TableRow() {
      }
      
      TableRow(int i) {
        enumChoice = (EnumCell)columns_.get(ENUM_COL_).get(i);
        origOrder = (Integer)hiddenColumns_.get(ORIG_ORDER_).get(i);
      }
      
      public void toCols() {
        columns_.get(ENUM_COL_).add(enumChoice);  
        hiddenColumns_.get(ORIG_ORDER_).add(origOrder);
        return;
      }
    }
  
    public OneEnumTableModel(String title, List<EnumCell> currEnums) {
      super(NUM_COL_);
      colNames_ = new String[] {title};
      colClasses_ = new Class[] {EnumCell.class};
      canEdit_ = new boolean[] {true};
      addHiddenColumns(NUM_HIDDEN_);
      currEnums_ = new ArrayList<EnumCell>(currEnums);
    }
   
    protected TableRow constructARow(int i) {
      return (new TableRow(i));
    }
    
    public TableRow constructARow() {
      return (new TableRow());
    }
       
    //
    // This class does not allow editable enums, since we are
    // disabling the add button with an empty enum
    //
    
    @Override
    void setEnums(Map<Integer, TableCellRenderer> renderers, Map<Integer, TableCellEditor> editors, Set<Integer> editableEnums) {
      if (!editableEnums.isEmpty()) {
        throw new IllegalArgumentException();
      }
      super.setEnums(renderers, editors, editableEnums);
      return;
    }

    public void setCurrentEnums(List<EnumCell> prsList) {
      currEnums_.clear();
      currEnums_.addAll(prsList);
      return;
    }
    
    @Override
    public boolean canAddRow() {
      return (!currEnums_.isEmpty());
    }   
    
    @Override
    public boolean addRow(int[] rows) {
      super.addRow(rows);
      int lastIndex = rowCount_ - 1;
      int addIndex = (rows.length == 0) ? lastIndex : rows[0] + 1;
      if (currEnums_.isEmpty()) {
        throw new IllegalStateException();
      }
      EnumCell fillIn = new EnumCell(currEnums_.get(0));
      columns_.get(ENUM_COL_).set(addIndex, fillIn);
      hiddenColumns_.get(ORIG_ORDER_).set(addIndex, new Integer(lastIndex));      
      return (true);
    }    
  }
}
```
