## Supplementary material for "BioFabric Visualization of Network Alignments": BioFabric Source Code: BioFabricNavAndControl.java

```
/*
**    Copyright (C) 2003-2018 Institute for Systems Biology 
**                            Seattle, Washington, USA. 
**
**    This library is free software; you can redistribute it and/or
**    modify it under the terms of the GNU Lesser General Public
**    License as published by the Free Software Foundation; either
**    version 2.1 of the License, or (at your option) any later version.
**
**    This library is distributed in the hope that it will be useful,
**    but WITHOUT ANY WARRANTY; without even the implied warranty of
**    MERCHANTABILITY or FITNESS FOR A PARTICULAR PURPOSE.  See the GNU
**    Lesser General Public License for more details.
**
**    You should have received a copy of the GNU Lesser General Public
**    License along with this library; if not, write to the Free Software
**    Foundation, Inc., 59 Temple Place, Suite 330, Boston, MA  02111-1307  USA
*/

package org.systemsbiology.biofabric.ui.display;

import java.awt.BorderLayout;
import java.awt.CardLayout;
import java.awt.Color;
import java.awt.Dimension;
import java.awt.Font;

import javax.swing.JFrame;
import javax.swing.JLabel;
import javax.swing.JPanel;

import javax.swing.JSplitPane;
import javax.swing.border.EmptyBorder;
import javax.swing.border.LineBorder;

import org.systemsbiology.biofabric.cmd.CommandSet;
import org.systemsbiology.biofabric.util.ResourceManager;


/****************************************************************************
**
** This is the BioFabric Control dashboard wrapper
*/

public class BioFabricNavAndControl {
  
  ////////////////////////////////////////////////////////////////////////////
  //
  // PRIVATE INSTANCE MEMBERS
  //
  //////////////////////////////////////////////////////////////////////////// 
  
  private MyNavPanel myPanel_;
  private FabricMagnifyingTool fmt_;
  private BioFabricOverview bfo_;
  private MouseOverView mvo_;
  private FabricNavTool fnt_;
  private FabricNavTool.LabeledFabricNavTool lfnt_;
  private FabricLocation floc_;
  private CardLayout clay_;
  private boolean collapsed_;
  private JPanel withControls_;
  private JSplitPane spot_;
  private double savedSplitFrac_;
  
  ////////////////////////////////////////////////////////////////////////////
  //
  // PUBLIC CONSTRUCTORS
  //
  ////////////////////////////////////////////////////////////////////////////

  /***************************************************************************
  **
  ** Constructor
  */

  public BioFabricNavAndControl(boolean isMain, JFrame topWindow, boolean isHeadless) {

    if (isHeadless) {
      return;
    }
    myPanel_ = new MyNavPanel();
    
    floc_ = new FabricLocation();

    CommandSet fc = CommandSet.getCmds((isMain) ? "mainWindow" : "selectionWindow");
    fmt_ = new FabricMagnifyingTool(fc.getColorGenerator());
    fmt_.keyInstall((JPanel)topWindow.getContentPane());  
    JPanel fmpan = new JPanel();
    fmpan.setLayout(new BorderLayout());
    fmpan.setBorder(new LineBorder(Color.black, 2));
    Font labelFont = new Font("SansSerif", Font.BOLD, 20);
    JLabel magLab = new JLabel(ResourceManager.getManager().getString("biofabric.magnifier"));
    magLab.setBorder(new EmptyBorder(0, 5, 0, 0));
    magLab.setOpaque(true);
    magLab.setBackground(Color.white);
    magLab.setFont(labelFont);
    fmpan.add(magLab, BorderLayout.NORTH);
    fmpan.add(fmt_, BorderLayout.CENTER);

    bfo_ = new BioFabricOverview(isHeadless);
    fmt_.setFabricOverview(bfo_);

    // This is a new feature that has no place to live, and no UI to install the needed data.
    // It was used for showing brain region images corresponding to nodes in a demo. When the
    // user runs the mouse over a node, it would show an image corresponding to a node. Over a link,
    // the images corresponding to endpoints. One place to put this is to let the user choose whether
    // to show the tour panel or the overview. It will need the user to provide the location of where
    // to find the image files, plus the filename pieces before and after the node name.
    //
    mvo_ = new MouseOverView();
    mvo_.setIsAlive(false); 
    mvo_.setFileLocations("path", "filePrefix", "fileSuffix");
    JPanel fopan = new JPanel();
    fopan.setLayout(new BorderLayout());
    fopan.setBorder(new LineBorder(Color.black, 2));
    JLabel overLab = new JLabel(ResourceManager.getManager().getString("biofabric.overview"));
    overLab.setBorder(new EmptyBorder(0, 5, 0, 0));
    overLab.setOpaque(true);
    overLab.setBackground(Color.white);
    overLab.setFont(labelFont);
    fopan.add(overLab, BorderLayout.NORTH);
    fopan.add(bfo_.getPanel(), BorderLayout.CENTER);
    
    lfnt_ = new FabricNavTool.LabeledFabricNavTool(topWindow, labelFont);
    fnt_ = lfnt_.getFabricNavTool();
     
    spot_ = new JSplitPane(JSplitPane.HORIZONTAL_SPLIT, fopan, lfnt_);
    
    // See above comment about new mvo_ option. This is the code to directly replace the tour with the
    // mouse overview.
    // JPanel dmvo = new JPanel();
    // dmvo.setLayout(new GridLayout(1, 2));
    // dmvo.add(mvo_.getPanel(0));
    // dmvo.add(mvo_.getPanel(1));
    // spot_ = new JSplitPane(JSplitPane.HORIZONTAL_SPLIT, fopan, dmvo);
    
    spot_.setBorder(new EmptyBorder(0,0,0,0));
    spot_.setResizeWeight(1.0);

    JSplitPane sp = new JSplitPane(JSplitPane.HORIZONTAL_SPLIT, fmpan, spot_);
    withControls_ = new JPanel();
    withControls_.setLayout(new BorderLayout());
    withControls_.add(floc_, BorderLayout.NORTH);
    withControls_.add(sp, BorderLayout.CENTER);
    
    clay_ = new CardLayout();
    myPanel_.setLayout(clay_);
    myPanel_.add(withControls_, "cntrl");
    myPanel_.add(new JPanel(), "blank");
    clay_.show(myPanel_, "cntrl");
    collapsed_ = false;
    
    return;
  }
  
  ////////////////////////////////////////////////////////////////////////////
  //
  // PUBLIC METHODS
  //
  ////////////////////////////////////////////////////////////////////////////

  /***************************************************************************
  **
  ** Get actual panel
  */
  
  public MyNavPanel getPanel() {
    return (myPanel_);
  }
 
  /***************************************************************************
  **
  ** Hide/show nav and controls
  */
  
  public boolean showTour(boolean show) {
    if (show) {
      spot_.setEnabled(true);
      lfnt_.setToBlank(!show);
      double need = (double)(spot_.getWidth() - lfnt_.getMinimumSize().width) / (double)spot_.getWidth();
      spot_.setDividerLocation(Math.min(savedSplitFrac_, need));
      if (lfnt_.getMinimumSize().height > myPanel_.getHeight()) {
        return (true);
      }
    } else {
      lfnt_.setToBlank(!show);
      int lastLoc = spot_.getDividerLocation();
      savedSplitFrac_ = (double)lastLoc / (double)spot_.getWidth();
      spot_.setDividerLocation(1.0);
      spot_.setEnabled(false);
    }
    return (false);
  }
  
  /***************************************************************************
  **
  ** Get the FabricNavTool
  */

  public FabricNavTool getNavTool() {
    return (fnt_);
  }  
  
  /***************************************************************************
  **
  ** Set to blank or populated
  */

  public void setToBlank(boolean val) {
    clay_.show(myPanel_, (val) ? "blank" : "cntrl");
    collapsed_ = val;
    return;
  }  
    
  /***************************************************************************
  **
  ** Get the FabricLocation
  */

  public FabricLocation getFabricLocation() {
    return (floc_);
  }  
  
  /***************************************************************************
  **
  ** Get the Mouseover view:
  */

  public MouseOverView getMouseOverView() {
    return (mvo_);
  }  
   
  /***************************************************************************
  **
  ** Get the FMT
  */

  public FabricMagnifyingTool getFMT() {
    return (fmt_);
  }
  
  /***************************************************************************
  **
  ** Get the Overview
  */

  public BioFabricOverview getOverview() {
    return (bfo_);
  }
   
  /***************************************************************************
  **
  ** Set the fabric panel
  */

  public void setFabricPanel(BioFabricPanel cp) {
    if (fnt_ != null) {
      fnt_.setFabricPanel(cp);
    }
    return;
  }
  
  /***************************************************************************
  **
  ** Now the actual panel to use
  */  
      
  public class MyNavPanel extends JPanel {
    
    private static final long serialVersionUID = 1L;
    
    /***************************************************************************
    **
    ** Sizing
    */
     
    @Override
    public Dimension getPreferredSize() {
      if (collapsed_) {
        return (new Dimension(0, 0));    
      } else {
        return (withControls_.getPreferredSize());
      } 
    }

    @Override
    public Dimension getMinimumSize() {
      if (collapsed_) {
        return (new Dimension(0, 0));    
      } else {
        return (withControls_.getMinimumSize());
      } 
    }
     
    @Override
    public Dimension getMaximumSize() {
      if (collapsed_) {
        return (new Dimension(0, 0));    
      } else {
        return (withControls_.getMaximumSize());
      } 
    }

    /***************************************************************************
    **
    ** Set bounds
    */

    @Override
    public void setBounds(int x, int y, int width, int height) {
      super.setBounds(x, y, width, height);
      repaint();
      return;
    }
  }
}
```
