## Supplementary material for "BioFabric Visualization of Network Alignments": BioFabric Source Code: BioFabricOverview.java

```
/*
**    Copyright (C) 2003-2014 Institute for Systems Biology 
**                            Seattle, Washington, USA. 
**
**    This library is free software; you can redistribute it and/or
**    modify it under the terms of the GNU Lesser General Public
**    License as published by the Free Software Foundation; either
**    version 2.1 of the License, or (at your option) any later version.
**
**    This library is distributed in the hope that it will be useful,
**    but WITHOUT ANY WARRANTY; without even the implied warranty of
**    MERCHANTABILITY or FITNESS FOR A PARTICULAR PURPOSE.  See the GNU
**    Lesser General Public License for more details.
**
**    You should have received a copy of the GNU Lesser General Public
**    License along with this library; if not, write to the Free Software
**    Foundation, Inc., 59 Temple Place, Suite 330, Boston, MA  02111-1307  USA
*/

package org.systemsbiology.biofabric.ui.display;

import java.awt.BasicStroke;
import java.awt.CardLayout;
import java.awt.Color;
import java.awt.Dimension;
import java.awt.Graphics;
import java.awt.Graphics2D;
import java.awt.Point;
import java.awt.Rectangle;
import java.awt.RenderingHints;
import java.awt.geom.AffineTransform;
import java.awt.geom.Ellipse2D;
import java.awt.geom.Point2D;
import java.awt.geom.Rectangle2D;
import java.awt.image.BufferedImage;
import javax.swing.JPanel;

/****************************************************************************
**
** This is the display panel
*/

public class BioFabricOverview {
  
  ////////////////////////////////////////////////////////////////////////////
  //
  // PRIVATE CONSTANTS
  //
  //////////////////////////////////////////////////////////////////////////// 

  private final static float MOUSE_RADIUS_ = 10.0F;
  private final static float MOUSE_XHAIR_ = 15.0F;
  private final static float BOX_XHAIR_ = 15.0F;
    
  ////////////////////////////////////////////////////////////////////////////
  //
  // PUBLIC CONSTANTS
  //
  //////////////////////////////////////////////////////////////////////////// 
  
  ////////////////////////////////////////////////////////////////////////////
  //
  // PRIVATE INSTANCE MEMBERS
  //
  ////////////////////////////////////////////////////////////////////////////
  
  private JPanel myPanel_;
  private Dimension currSize_;
  private BufferedImage img_;
  private BufferedImage scaledImg_;
  @SuppressWarnings("unused")
  private Point navFocus_;
  private Point scaledImgOrigin_;
  private Rectangle worldRect_;
  private AffineTransform transform_;
  private Point2D mousePoint_;
  private double zoom_;
  private boolean mouseIn_;
  private Rectangle2D viewInWorld_;
  private Rectangle magInWorld_;
  private boolean hideMag_;
  private CardLayout myCard_;
  private PaintPanel pPan_;
  
  ////////////////////////////////////////////////////////////////////////////
  //
  // PUBLIC CONSTRUCTORS
  //
  ////////////////////////////////////////////////////////////////////////////

  /***************************************************************************
  **
  ** Constructor
  */

  public BioFabricOverview(boolean isHeadless) {
    if (isHeadless) {
      return;
    }
    myPanel_ = new MyOverPanel();
    myPanel_.setBackground(Color.white);
    scaledImg_ = null;
    img_ = null;
    navFocus_ = new Point(0, 0);
    scaledImgOrigin_ = new Point(0, 0);
    mouseIn_ = false;
    hideMag_ = true;
    
    worldRect_ = new Rectangle();
    mousePoint_ = new Point2D.Double();
    viewInWorld_= new Rectangle2D.Double();
    magInWorld_ = new Rectangle();
    
    myCard_ = new CardLayout();
    myPanel_.setLayout(myCard_);
    
    pPan_ = new PaintPanel();
    myPanel_.add(pPan_, "theView");
    JPanel blankPanel = new JPanel();
    blankPanel.setBackground(Color.white);
    myPanel_.add(blankPanel, "Hiding");   
  }

  ////////////////////////////////////////////////////////////////////////////
  //
  // PUBLIC METHODS
  //
  ////////////////////////////////////////////////////////////////////////////

  /***************************************************************************
  ** 
  ** Get the panel
  */

  public JPanel getPanel() { 
    return (myPanel_);
  }

  /***************************************************************************
  **
  ** Show or hide the view
  */

  public void showView(boolean enabled) {
    myCard_.show(myPanel_, (enabled) ? "theView" : "Hiding"); 
    return;
  }
  
  /***************************************************************************
  **
  ** Set current mouse location
  */

  public void setMouse(Point2D center, Point cprc) {
    navFocus_ = cprc;
    mousePoint_.setLocation(center);
    mouseIn_ = true;
    myPanel_.repaint();
    return;
  }
  
  /***************************************************************************
  **
  ** Set the viewport location
  */

  public void setViewInWorld(Rectangle2D viewInWorld) {
    viewInWorld_.setFrame(viewInWorld);
    myPanel_.repaint();
    return;
  }
  
  /***************************************************************************
  **
  ** Set magnifier location
  */

  public void setMagnifyView(Rectangle magInWorld) {
    magInWorld_.setBounds(magInWorld);
    myPanel_.repaint();
    return;
  }
    
  /***************************************************************************
  **
  ** Set if mouse in in view
  */

  public void setMouseIn(boolean isIn, boolean magIgnoring) {
    mouseIn_ = isIn;
    hideMag_ = (!isIn && !magIgnoring);
    myPanel_.repaint();
    return;
  }
  
  /***************************************************************************
  ** 
  ** Install a model
  */

  public void installImage(BufferedImage img, Rectangle worldRect) {
    img_ = img;
    scaledImg_ = null;
    worldRect_.setBounds(worldRect);
    resizeImage();
    myPanel_.repaint();
    return;
  }
  
  /***************************************************************************
  **
  ** resize image
  */

  private void resizeImage() {
    if (img_ == null) {
      return;
    }
    if (currSize_ == null) {
      currSize_ = myPanel_.getSize();
    }
    //
    // Don't do this step if the overview is not currently bring displayed
    //
    if ((currSize_.width <= 0) || (currSize_.height <= 0)) {
      return;
    }
    double imgAR = (double)img_.getWidth() / (double)img_.getHeight();
    double panelAR = currSize_.getWidth() / currSize_.getHeight();
    int imgHeight;
    int imgWidth;
    int startX;
    int startY;
    if (panelAR < imgAR) { // long image, tall panel
      imgWidth = currSize_.width;
      imgHeight = (int)(imgWidth / imgAR);
      if (imgHeight == 0) {
        imgHeight = 1;
      }
      startX = 0;
      startY = (currSize_.height - imgHeight) / 2;
    } else {
      imgHeight = currSize_.height;
      imgWidth = (int)(imgHeight * imgAR);
      if (imgWidth == 0) {
        imgWidth = 1;
      }
      startX = (currSize_.width - imgWidth) / 2;
      startY = 0;
    }
    scaledImgOrigin_.setLocation(startX, startY);
    scaledImg_ = new BufferedImage(imgWidth, imgHeight, BufferedImage.TYPE_INT_RGB);
    Graphics2D g2 = scaledImg_.createGraphics();
    g2.setRenderingHint(RenderingHints.KEY_INTERPOLATION, RenderingHints.VALUE_INTERPOLATION_BICUBIC);
    g2.drawImage(img_, 0, 0, imgWidth, imgHeight, null);
    g2.dispose();
  
    double zoomHW = imgWidth / worldRect_.getWidth();
    double zoomVW = imgHeight / worldRect_.getHeight();
    if (zoomHW < zoomVW) {
      zoom_ = zoomHW;
    } else {
      zoom_ = zoomVW;
    }
    
    transform_ = new AffineTransform();    
    transform_.translate((worldRect_.getWidth() / 2.0) * zoom_, (worldRect_.getHeight() / 2.0) * zoom_);
    transform_.scale(zoom_, zoom_);
    transform_.translate(-worldRect_.getCenterX(), -worldRect_.getCenterY()); 
    return;
  } 

  /***************************************************************************
  **
  ** Handles the drawing
  */  
  
  private class PaintPanel extends JPanel {
    private static final long serialVersionUID = 1L;
  
    /***************************************************************************
    **
    ** Constructor
    */
  
    public PaintPanel() {
      setBackground(Color.white);
    }
    
    /***************************************************************************
    **
    ** Drawing routine
    */
    
    public void paintComponent(Graphics g) {
      super.paintComponent(g);
      Graphics2D g2 = (Graphics2D)g;     
      if (img_ == null) {
        return;
      }
      if (scaledImg_ == null) {
        resizeImage();
      }
     
      g2.drawImage(scaledImg_, scaledImgOrigin_.x, scaledImgOrigin_.y, null);
      drawFloater(g);
      return;
    }
    
    /***************************************************************************
    **
    ** Gets given point in viewport coordinates
    */
    
    private Point pointToViewport(Point2D worldPoint) {
      Point2D newPoint = new Point2D.Double(worldPoint.getX(), worldPoint.getY());
      transform_.transform(newPoint, newPoint);
      int x = (int)(Math.round(newPoint.getX()));
      int y = (int)(Math.round(newPoint.getY()));
      Point pt = new Point(x, y);
      return (pt);
    } 
    
    /***************************************************************************
    **
    ** Drawing core
    */
    
    private void drawFloater(Graphics g) {
      Graphics2D g2 = (Graphics2D)g; 
      g2.setRenderingHint(RenderingHints.KEY_ANTIALIASING, RenderingHints.VALUE_ANTIALIAS_ON);
      g2.setRenderingHint(RenderingHints.KEY_FRACTIONALMETRICS, RenderingHints.VALUE_FRACTIONALMETRICS_ON);
      BasicStroke selectedStroke = new BasicStroke(1, BasicStroke.CAP_SQUARE, BasicStroke.JOIN_MITER);    
      g2.setStroke(selectedStroke);    
      g2.setPaint(Color.GREEN);
         
      if ((mousePoint_ != null) && mouseIn_) {  
        Point viewP = pointToViewport(mousePoint_); 
        Ellipse2D circ = new Ellipse2D.Double(viewP.x + scaledImgOrigin_.x - MOUSE_RADIUS_, 
                                              viewP.y + scaledImgOrigin_.y - MOUSE_RADIUS_, 
                                              2.0 * MOUSE_RADIUS_, 2.0 * MOUSE_RADIUS_);
        g2.draw(circ);
        g2.drawLine((int)(viewP.x + scaledImgOrigin_.x - MOUSE_XHAIR_), 
                    (viewP.y + scaledImgOrigin_.y), 
                    (int)(viewP.x + scaledImgOrigin_.x + MOUSE_XHAIR_),  
                    (viewP.y + scaledImgOrigin_.y));
        g2.drawLine((viewP.x + scaledImgOrigin_.x), 
                    (int)(viewP.y + scaledImgOrigin_.y - MOUSE_XHAIR_), 
                    (viewP.x + scaledImgOrigin_.x),  
                    (int)(viewP.y + scaledImgOrigin_.y + MOUSE_XHAIR_));
      }
      
      if (viewInWorld_ != null) {      
        Point2D viwUL = new Point2D.Double(viewInWorld_.getMinX(), viewInWorld_.getMinY());
        Point viewUL = pointToViewport(viwUL);
        Point2D viwLR = new Point2D.Double(viewInWorld_.getMaxX(), viewInWorld_.getMaxY());
        Point viewLR = pointToViewport(viwLR);
        int vw = viewLR.x - viewUL.x;
        int vh = viewLR.y - viewUL.y;
        int vcx = (viewUL.x + viewLR.x) / 2;
        int vcy = (viewUL.y + viewLR.y) / 2;
        g2.drawRect((viewUL.x + scaledImgOrigin_.x), 
                    (viewUL.y + scaledImgOrigin_.y), vw, vh);
        g2.drawLine((int)(vcx + scaledImgOrigin_.x - BOX_XHAIR_), 
                    (vcy + scaledImgOrigin_.y), 
                    (int)(vcx + scaledImgOrigin_.x + BOX_XHAIR_),  
                    (vcy + scaledImgOrigin_.y));
        g2.drawLine((vcx + scaledImgOrigin_.x), 
                    (int)(vcy + scaledImgOrigin_.y - BOX_XHAIR_), 
                    (vcx + scaledImgOrigin_.x),  
                    (int)(vcy + scaledImgOrigin_.y + BOX_XHAIR_));
        
        
        
      }
      
      if ((magInWorld_ != null) && !hideMag_) {
        g2.setPaint(Color.RED);
        Point2D viwUL = new Point2D.Double(magInWorld_.getMinX(), magInWorld_.getMinY());
        Point viewUL = pointToViewport(viwUL);
        Point2D viwLR = new Point2D.Double(magInWorld_.getMaxX(), magInWorld_.getMaxY());
        Point viewLR = pointToViewport(viwLR);
        int vw = viewLR.x - viewUL.x;
        int vh = viewLR.y - viewUL.y;
        int vcx = (viewUL.x + viewLR.x) / 2;
        int vcy = (viewUL.y + viewLR.y) / 2;
        g2.drawRect((viewUL.x + scaledImgOrigin_.x), 
                    (viewUL.y + scaledImgOrigin_.y), vw, vh);
        g2.drawLine((int)(vcx + scaledImgOrigin_.x - BOX_XHAIR_), 
                    (vcy + scaledImgOrigin_.y), 
                    (int)(vcx + scaledImgOrigin_.x + BOX_XHAIR_),  
                    (vcy + scaledImgOrigin_.y));
        g2.drawLine((vcx + scaledImgOrigin_.x), 
                    (int)(vcy + scaledImgOrigin_.y - BOX_XHAIR_), 
                    (vcx + scaledImgOrigin_.x),  
                    (int)(vcy + scaledImgOrigin_.y + BOX_XHAIR_));
      }
      return;
    }
  }
  
  /***************************************************************************
  **
  ** Now the actual panel to use
  */  
      
  public class MyOverPanel extends JPanel {
    
    private static final long serialVersionUID = 1L;
    
    /***************************************************************************
    **
    ** Sizing
    */

    @Override
    public Dimension getPreferredSize() {
      return (new Dimension(800, 100));    
    }
     
    @Override
    public Dimension getMinimumSize() {
      return (new Dimension(10, 10));    
    }
     
    @Override
    public Dimension getMaximumSize() {
      return (new Dimension(4000, 340));    
    }
       
    /***************************************************************************
    **
    ** Handle size change
    */

    @Override
    public void setBounds(int x, int y, int width, int height) {
      super.setBounds(x, y, width, height);
      currSize_ = new Dimension(width, height);
      resizeImage();
      repaint();
      return;
    } 
  } 
}
```
