## Supplementary material for "BioFabric Visualization of Network Alignments": BioFabric Source Code: BioFabricPanel.java

```
/*
**    Copyright (C) 2003-2018 Institute for Systems Biology 
**                            Seattle, Washington, USA. 
**
**    This library is free software; you can redistribute it and/or
**    modify it under the terms of the GNU Lesser General Public
**    License as published by the Free Software Foundation; either
**    version 2.1 of the License, or (at your option) any later version.
**
**    This library is distributed in the hope that it will be useful,
**    but WITHOUT ANY WARRANTY; without even the implied warranty of
**    MERCHANTABILITY or FITNESS FOR A PARTICULAR PURPOSE.  See the GNU
**    Lesser General Public License for more details.
**
**    You should have received a copy of the GNU Lesser General Public
**    License along with this library; if not, write to the Free Software
**    Foundation, Inc., 59 Temple Place, Suite 330, Boston, MA  02111-1307  USA
*/

package org.systemsbiology.biofabric.ui.display;

import java.awt.AlphaComposite;
import java.awt.BasicStroke;
import java.awt.Color;
import java.awt.Dimension;
import java.awt.Graphics;
import java.awt.Graphics2D;
import java.awt.Point;
import java.awt.Rectangle;
import java.awt.RenderingHints;
import java.awt.event.MouseAdapter;
import java.awt.event.MouseEvent;
import java.awt.event.MouseMotionAdapter;
import java.awt.event.MouseWheelEvent;
import java.awt.event.MouseWheelListener;
import java.awt.geom.AffineTransform;
import java.awt.geom.Point2D;
import java.awt.geom.Rectangle2D;
import java.awt.image.BufferedImage;
import java.awt.print.PageFormat;
import java.awt.print.Printable;
import java.io.File;
import java.util.ArrayList;
import java.io.IOException;
import java.io.OutputStream;
import java.util.HashMap;
import java.util.HashSet;
import java.util.Iterator;
import java.util.List;
import java.util.Map;
import java.util.Set;
import java.util.SortedMap;
import java.util.SortedSet;
import java.util.TreeMap;
import java.util.TreeSet;

import javax.swing.JPanel;
import javax.swing.JScrollBar;
import javax.swing.JScrollPane;
import javax.swing.JViewport;

import org.systemsbiology.biofabric.api.io.BuildData;
import org.systemsbiology.biofabric.api.model.AnnotationSet;
import org.systemsbiology.biofabric.api.model.AnnotsForPos;
import org.systemsbiology.biofabric.api.model.NetNode;
import org.systemsbiology.biofabric.api.util.ExceptionHandler;
import org.systemsbiology.biofabric.api.util.MinMax;
import org.systemsbiology.biofabric.api.util.NID;
import org.systemsbiology.biofabric.api.worker.AsynchExitRequestException;
import org.systemsbiology.biofabric.api.worker.BTProgressMonitor;
import org.systemsbiology.biofabric.api.worker.LoopReporter;
import org.systemsbiology.biofabric.app.BioFabricApplication;
import org.systemsbiology.biofabric.app.BioFabricWindow;
import org.systemsbiology.biofabric.cmd.CommandSet;
import org.systemsbiology.biofabric.cmd.ZoomCommandSupport;
import org.systemsbiology.biofabric.cmd.ZoomTarget;
import org.systemsbiology.biofabric.event.EventManager;
import org.systemsbiology.biofabric.event.SelectionChangeEvent;
import org.systemsbiology.biofabric.io.BuildDataImpl;
import org.systemsbiology.biofabric.model.AnnotationSetImpl;
import org.systemsbiology.biofabric.model.BioFabricNetwork;
import org.systemsbiology.biofabric.model.FabricLink;
import org.systemsbiology.biofabric.ui.BasicZoomTargetSupport;
import org.systemsbiology.biofabric.ui.CursorManager;
import org.systemsbiology.biofabric.ui.FabricColorGenerator;
import org.systemsbiology.biofabric.ui.FabricDisplayOptions;
import org.systemsbiology.biofabric.ui.FabricDisplayOptionsManager;
import org.systemsbiology.biofabric.ui.ImageExporter;
import org.systemsbiology.biofabric.ui.PopupMenuControl;
import org.systemsbiology.biofabric.ui.ZoomPresentation;
import org.systemsbiology.biofabric.ui.render.BucketRenderer;
import org.systemsbiology.biofabric.ui.render.BufBuildDrawer;
import org.systemsbiology.biofabric.ui.render.ImgAndBufPool;
import org.systemsbiology.biofabric.ui.render.BufferBuilder;
import org.systemsbiology.biofabric.ui.render.PaintCacheSmall;
import org.systemsbiology.biofabric.util.QuadTree;
import org.systemsbiology.biofabric.util.UiUtil;

/****************************************************************************
**
** This is the display panel
*/

public class BioFabricPanel implements ZoomTarget, ZoomPresentation, Printable,
                                       BufBuildDrawer, BufferBuilder.BufferBuilderClient {
  
  ////////////////////////////////////////////////////////////////////////////
  //
  // PRIVATE CONSTANTS
  //
  //////////////////////////////////////////////////////////////////////////// 

  private static final double PAD_MULT_ = 0.20; 
  
  ////////////////////////////////////////////////////////////////////////////
  //
  // PUBLIC CONSTANTS
  //
  //////////////////////////////////////////////////////////////////////////// 
  
  public static final int GRID_SIZE = 18;
  public static final int GRID_SIZE_SQ = GRID_SIZE * GRID_SIZE;
  
  ////////////////////////////////////////////////////////////////////////////
  //
  // PRIVATE INSTANCE MEMBERS
  //
  ////////////////////////////////////////////////////////////////////////////
  
  private MyPaintPanel myPanel_;
  
  private JScrollPane jsp_;

  //
  // Note on zoom implementation.  This one class is serving
  // as both the myGenomePre_ AND the sup_ classes!  Thus
  // you will have separate calls doing much the same thing!
  //
  
  private BasicZoomTargetSupport zoomer_;
  private ZoomCommandSupport zcs_; 
  
  private TreeMap<Double, Integer> zoomMap_;
  
  private Point lastPress_;  
  private Point lastView_;          
  private Point lastAbs_;
  private boolean collectingZoomMode_;
  private boolean collectingTourStart_;
  private boolean tourStartSelectionOnly_;
  private Point firstZoomPoint_;
  
  private boolean isAMac_;
  @SuppressWarnings("unused")
  private boolean lastShifted_;
  private boolean lastCtrl_;
  
  private BioFabricPanel myMini_;
  
  
  // Selection-related data:
  private HashSet<FabricLink> currLinkSelections_; 
  private HashSet<NetNode> currNodeSelections_; 
  private HashSet<Integer> currColSelections_;
  
  private ArrayList<BioFabricNetwork.NodeInfo> targetList_;
  private ArrayList<BioFabricNetwork.LinkInfo> linkList_;
  private ArrayList<Rectangle> rects_;
  private int currSel_;
  private PaintCacheSmall.Reduction selections_;
  // End Selection-related data
  
  private BufferBuilder bufferBuilder_;
  

  private FabricLocation myLocation_;
  private MouseOverView mov_;
  private Rectangle2D worldRectNetAR_;
  private Rectangle2D worldRectScreenAR_;
  private Dimension screenDim_;
  
  private PaintCacheSmall.FloaterSet floaterSet_;
  private Point selFocus_;
  private Point tourFocus_;
  
  private BioFabricNetwork bfn_;
  private PaintCacheSmall painter_;

  private BioFabricApplication bfa_;
  private FabricMagnifyingTool fmt_;
  private FabricNavTool fnt_;
  private BioFabricOverview bfo_;

  private BufferedImage bim_;
  private boolean doBuildSelect_;
  private CursorManager cursorMgr_;
  private BioFabricWindow bfw_;
  private Map<NetNode, Rectangle2D> nodeNameLocations_;
  private Map<NetNode, List<Rectangle2D>> drainNameLocations_;
  private QuadTree forSelections_;
   
  private PopupMenuControl popCtrl_;
  
  private BucketRenderer bucketRend_;

  
  private Rectangle clipRect_;
  private Rectangle clipRect2_;
  private ImgAndBufPool bis_;
  private List<BufferedImage> staleImages_;
    
  private Point	fullModelPos_;
  private Dimension	fullModelExtent_;
  
  ////////////////////////////////////////////////////////////////////////////
  //
  // PUBLIC CONSTRUCTORS
  //
  ////////////////////////////////////////////////////////////////////////////

  /***************************************************************************
  **
  ** Constructor
  */

  public BioFabricPanel(FabricColorGenerator colGen, BioFabricApplication bfa, 
                        FabricMagnifyingTool fmt, BioFabricOverview bfo, FabricNavTool fnt, 
                        boolean isForMain, BioFabricWindow bfw, BucketRenderer bRend, boolean headless) {
    bfa_ = bfa;
    fmt_ = fmt;
    fnt_ = fnt;
    bfo_ = bfo;
    bfw_ = bfw;
    myPanel_ = (headless) ? null : new MyPaintPanel();
    zoomer_ = new BasicZoomTargetSupport(this, myPanel_);
    CommandSet fc = CommandSet.getCmds((isForMain) ? "mainWindow" : "selectionWindow");
    zcs_ = new ZoomCommandSupport(fc);
    isAMac_ = fc.isAMac();
    painter_ = new PaintCacheSmall(colGen);
    if (fmt_ != null) {
      fmt_.setPainters(painter_, painter_);
    }
    zoomMap_ = new TreeMap<Double, Integer>();
    if (myPanel_ != null) {
      myPanel_.addMouseListener(new MouseHandler());
      myPanel_.addMouseMotionListener(new MouseMotionHandler());
    } 
    currLinkSelections_ = new HashSet<FabricLink>();
    currNodeSelections_ = new HashSet<NetNode>();
    currColSelections_ = new HashSet<Integer>();
    targetList_ = new ArrayList<BioFabricNetwork.NodeInfo>();
    linkList_ = new ArrayList<BioFabricNetwork.LinkInfo>();
    collectingZoomMode_ = false;
    collectingTourStart_ = false;
    tourStartSelectionOnly_ = false;
    firstZoomPoint_ = null;
    lastShifted_ = false;
    lastCtrl_ = false;
    clipRect_ = new Rectangle();
    clipRect2_ = new Rectangle();
   
    worldRectNetAR_ = new Rectangle2D.Double(0.0, 0.0, 100.0, 100.0); 
    zoomer_.setWorldRect(UiUtil.rectFromRect2D(worldRectNetAR_)); 
    bfn_ = null;
    
    floaterSet_ = new PaintCacheSmall.FloaterSet(null, null, null);
    if (fmt_ != null) {
      fmt_.setCurrentFloater(floaterSet_);  // Actually directly shared!
    }
    
    rects_ = new ArrayList<Rectangle>();
    currSel_ = -1;
    selFocus_ = null;
    tourFocus_ = null;   
 
    doBuildSelect_ = true;
    cursorMgr_ = (headless) ? null : new CursorManager(myPanel_, false);
    popCtrl_ = (headless) ? null : new PopupMenuControl(myPanel_);
    bucketRend_ = bRend;
    staleImages_ = new ArrayList<BufferedImage>();
  }

  ////////////////////////////////////////////////////////////////////////////
  //
  // PUBLIC METHODS
  //
  ////////////////////////////////////////////////////////////////////////////

  /***************************************************************************
  ** 
  ** Get the panel
  */

  public JPanel getPanel() { 
    return (myPanel_);
  }

  /***************************************************************************
  ** 
  ** Get the bucket renderer
  */

  public BucketRenderer getBucketRend() { 
    return (bucketRend_);
  }
  
  /***************************************************************************
  ** 
  ** Get the buffered image stack
  */

  public ImgAndBufPool getBufImgStack() { 
    return (bis_);
  }
  
  /***************************************************************************
  ** 
  ** Get the magnifying tool
  */

  public FabricMagnifyingTool getMagnifier() { 
    return (fmt_);
  }
 
  /***************************************************************************
  ** 
  ** Set the current clip rect
  */

  public void setCurrClipRect(Rectangle2D clipRect) { 
    zoomer_.setCurrClipRect(clipRect);
    return;
  }

  /***************************************************************************
  **
  ** Set if we accumulate selections
  */
  
  public void toggleBuildSelect() {
    doBuildSelect_ = !doBuildSelect_;
    return;
  } 

  /***************************************************************************
  **
  ** See if we accumulate selections
  */
  
  public boolean amBuildingSelections() {
    return (doBuildSelect_);
  } 
   
  /***************************************************************************
  **
  ** Answer if we have a model
  */
  
  public boolean hasAModel() {
    return ((bfn_ != null) && (bfn_.getRowCount() > 0));
  } 
  
  /***************************************************************************
  **
  ** Get the network
  */
  
  public BioFabricNetwork getNetwork() {
    return (bfn_);
  } 
   
  /***************************************************************************
  **
  ** Drawing routine
  */
  
  public void setSelectionPanel(BioFabricPanel mini) {
    myMini_ = mini;
    return;
  } 

  /***************************************************************************
  **
  ** Reset to clear model
  */
  
  public void reset() {
    if (bufferBuilder_ != null) {
      bufferBuilder_.release();
    }
    painter_.clear();
    bfn_ = null;
    if (fmt_ != null) {
      fmt_.setModel(null);
    }
    zoomMap_.clear();
    worldRectNetAR_ = new Rectangle2D.Double(0.0, 0.0, 100.0, 100.0); 
    zoomer_.setWorldRect(UiUtil.rectFromRect2D(worldRectNetAR_));
    clearSelections();
    if (fnt_ != null) {
      fnt_.haveAModel(false);
    }
    floaterSet_.clear();
    nodeNameLocations_ = null;
    drainNameLocations_ = null;
    return;
  } 
  
  /***************************************************************************
  **
  ** Shutdown
  */
  
  public void shutdown() {
    if (bufferBuilder_ != null) {
      bufferBuilder_.release();
    }
    return;
  }
  
  /***************************************************************************
  **
  ** Answer if ready
  */
  
  public void yourOrderIsReady(int size) {
    if (bfn_ == null) {
      return;
    }
    //System.out.println("OIR " + size + " " + System.currentTimeMillis() + " " + Runtime.getRuntime().freeMemory());
    Double zoomVal = new Double(zoomer_.getZoomFactor());
    Integer numObj = zoomMap_.get(zoomVal);
    if ((numObj != null) && (size == numObj.intValue())) {
      repaint();
    }   
    return;
  } 
 
  /***************************************************************************
  **
  ** Set the buffer builder
  */
  
  public void setBufBuilder(BufferBuilder bb) {
    bufferBuilder_ = bb;
    return;
  }
  
  /***************************************************************************
  **
  ** Get the buffer builder
  */
  
  public BufferBuilder getBufBuilder() {
    return (bufferBuilder_);
  }  
 
  /***************************************************************************
  **
  ** This is currently the entire contorted calculation of zooms
  */
  
  public int[] calcZoomSettings(Dimension screenSize) {
    
    //
    // Figure out the fundamental zoom to get the model to fill the screen. We will use this largest version
  	// as the baseline. Note that for the view with the nav panel showing, we will zoom out on this largest
  	// view.
    //

    double screenAR = screenSize.getWidth()  / screenSize.getHeight(); // > 1.0 for wide and flat screen
    double worldAR = worldRectNetAR_.getWidth()  / worldRectNetAR_.getHeight(); // > 1.0 for wide and flat network
    
    //
    // If the network has a higher aspect ratio than the screen, it will fill the available screen width and
    // need to be padded on the top and bottom with white space. If it has a lower aspect ratio, it will fill
    // the screen height, and need to be padded on the sides.
    //
    
    double worldWidthScreenAR;
    double worldHeightScreenAR;
    double zoom;    
    
    if (worldAR > screenAR) {
    	worldWidthScreenAR = worldRectNetAR_.getWidth();
    	worldHeightScreenAR = worldWidthScreenAR / screenAR;
    	zoom = screenSize.getWidth() / worldRectNetAR_.getWidth(); 	
    } else {
      worldHeightScreenAR = worldRectNetAR_.getHeight();
    	worldWidthScreenAR = worldHeightScreenAR * screenAR;
    	zoom = screenSize.getHeight() / worldRectNetAR_.getHeight();
    }
    
    screenDim_ = new Dimension(screenSize);
    bis_ = new ImgAndBufPool(50000);
    
    //
    // We create a world rectangle centered on the existing world, with the aspect ratio
    // of the screen:
    //
    
    double worldCenterX = worldRectNetAR_.getCenterX();
    double worldCenterY = worldRectNetAR_.getCenterY();
    
    double worldULX = worldCenterX - (worldWidthScreenAR / 2.0);
    double worldULY = worldCenterY - (worldHeightScreenAR / 2.0);
    worldRectScreenAR_ = new Rectangle2D.Double(worldULX, worldULY, worldWidthScreenAR, worldHeightScreenAR);
      
    //
    // OK, the start zoom has been calculated to show the whole network. We now calculate powers of
    // two until we basically fill the screen with a link glyph. Figure out that zoom level:
    //
    
    double maxZoom =  screenSize.getHeight() / (4.0 * GRID_SIZE);
    
    //
    // We terminate the image tiling and go to straight rendering when we hit 10E4 links across the screen.
    // In the case of tiny networks, we only generate a single image:
    //
    
    double transitionZoom =  screenSize.getWidth() / (1.0E4 * GRID_SIZE);

    //
    // Generate the zoom settings
    //
    
    double lastZoom = zoom;
    
    ArrayList<Double> zoomList = new ArrayList<Double>();
    ArrayList<Integer> forRet = new ArrayList<Integer>();
    int count = 0;
    while ((lastZoom < maxZoom) || (count < 1)) {
    	Double zObj = Double.valueOf(lastZoom);
    	zoomList.add(zObj);
    	// We generate at least two zoom levels, both of which are image-based:
    	if ((lastZoom < transitionZoom) || (count < 2)) {
    		Integer index = Integer.valueOf(count);
    	  zoomMap_.put(zObj, index); 
    	  forRet.add(index);
    	} 
    	count++;
      lastZoom *= 2.0;	
    }
      
    //
    // Companion renderer needs this info too:
    //
    
    bucketRend_.setModelDims(screenDim_, worldRectScreenAR_, bis_);
    
    int[] zooms = new int[forRet.size()];
    double[] zoomVals = new double[zoomList.size()];
    
    count = 0;
    for (Integer zo : forRet) {
    	zooms[count++] = zo.intValue();
    }
    count = 0;
    for (Double zo : zoomList) {
    	zoomVals[count++] = zo.doubleValue();
    }

    //
    // Install the calculated zoom levels:
    //
    
    zcs_.setZoomPoints(zoomVals); 
    zcs_.setZoomLevels(zooms); 
   
    return (zooms);
  }

  /***************************************************************************
  ** 
  ** Get the dimensions for the buffer
  */
  
   public void dimsForBuf(Dimension screenDim, Rectangle2D worldRect) {
     screenDim.setSize(screenDim_);
     worldRect.setRect(worldRectScreenAR_);    
     return;
   }

  /***************************************************************************
  ** 
  ** Clear selections
  */

  public void clearSelections() { 
    currLinkSelections_.clear();
    currNodeSelections_.clear();
    currColSelections_.clear();
    fmt_.setSelections(null);
    targetList_.clear();
    linkList_.clear();
    floaterSet_.currSelRect = null;
    rects_.clear();
    currSel_ = -1;
    selections_ = null;
    if (fnt_ != null) {
      fnt_.haveASelection(false);
    }
    handleFloaterChange();
    EventManager mgr = EventManager.getManager();
    SelectionChangeEvent ev = new SelectionChangeEvent(null, null, SelectionChangeEvent.SELECTED_ELEMENT);
    mgr.sendSelectionChangeEvent(ev);     
    return;
  }
  /***************************************************************************
  ** 
  ** Set location announcement
  */

  public void setFabricLocation(FabricLocation loc, MouseOverView mov) { 
    myLocation_ = loc;
    mov_ = mov;
    return;
  }  
  
  /***************************************************************************
  ** 
  ** Build a focus box
  */

  public Rectangle buildFocusBox(Point rcPoint) {
    Point ulp = (Point)rcPoint.clone();
    ulp.setLocation(ulp.x - 2, ulp.y - 2);
    Point2D wulp = rowColToWorld(ulp);    
    return (new Rectangle((int)wulp.getX(), (int)wulp.getY(), 4 * GRID_SIZE, 4 * GRID_SIZE));      
  }
  
  /***************************************************************************
  ** 
  ** Answer if we have a selection
  */

  public boolean haveASelection() {
    return (!(currLinkSelections_.isEmpty() &&
              currNodeSelections_.isEmpty() &&
              currColSelections_.isEmpty())); 
  }
  
  /***************************************************************************
  ** 
  ** Get node selections
  */

  public Set<NetNode> getNodeSelections() {
    return (currNodeSelections_);
  }
  
  /***************************************************************************
  ** 
  ** Get detail panel
  */

  public void installSearchResult(Set<NetNode> results, boolean doDiscard) {
    
    if (doDiscard) {
      currLinkSelections_.clear();
      currNodeSelections_.clear();
      currColSelections_.clear();
      fmt_.setSelections(null);
      currSel_ = -1;
      floaterSet_.currSelRect = null;
    }       
    if (results.isEmpty()) {
      buildSelectionGeometry(null, null);     
      return;
    }
    
    currNodeSelections_.addAll(results);
    buildSelectionGeometry(results.iterator().next(), null);
    zoomToSelected();    
    return;
  } 
  
  /***************************************************************************
  ** 
  ** Add first neighbors to selection
  */

  public void addFirstNeighbors() {
    boolean showShadows = FabricDisplayOptionsManager.getMgr().getDisplayOptions().getDisplayShadows();
    bfn_.addFirstNeighbors(currNodeSelections_, currColSelections_, currLinkSelections_, showShadows);
    buildSelectionGeometry(null, null);
    return;
  } 

  /***************************************************************************
  ** 
  ** Send selections to mini panel
  */

  private void transmitSelections(List<BioFabricNetwork.NodeInfo> targetList, List<BioFabricNetwork.LinkInfo> linkList) {      
    if (myMini_ == null) {
      bfa_.launchSelection();
      if (myMini_ == null) {
        throw new IllegalStateException();
      }
    } else {
      bfa_.raiseSelection();
    }
    CommandSet fc = CommandSet.getCmds("selectionWindow");
    try {
      BuildData bfnsbd = new BuildDataImpl(bfn_, targetList, linkList);
      fc.getFileLoader().newModelOperations(bfnsbd, false);
    } catch (IOException ioex) {
      throw new IllegalStateException();  // should not happen
    }
    return;
  }

   
  /***************************************************************************
  ** 
  ** Handle floater update
  */

  public void handleFloaterChange() {
    repaint();
    if (fmt_ != null) {
      fmt_.repaint();
    }
    return;
  } 
  
  /***************************************************************************
  ** 
  ** Repaint
  */

  public void repaint() {
    if (myPanel_ != null) {
      myPanel_.repaint();
    }
    if (fmt_ != null) {
      fmt_.repaint();
    }
    return;
  } 

  /***************************************************************************
  ** 
  ** Tour right
  */

  public TourStatus goRight(boolean selectedOnly) {
    return (goLeftRight(1, selectedOnly)); 
  } 

  /***************************************************************************
  ** 
  ** Tour far right
  */

  public TourStatus goFarRight(boolean selectedOnly) {
    return (goFarLeftRight(true, selectedOnly));
  }
  
  /***************************************************************************
  ** 
  ** Tour left
  */

  public TourStatus goLeft(boolean selectedOnly) {
    return (goLeftRight(-1, selectedOnly)); 
  } 
  
  /***************************************************************************
  ** 
  ** Tour far left
  */

  public TourStatus goFarLeft(boolean selectedOnly) {
    return (goFarLeftRight(false, selectedOnly));
  }

  /***************************************************************************
  ** 
  ** Tour up
  */

  public TourStatus goUp(boolean selectedOnly) {
    return (goUpDown(true, selectedOnly));
  }
  
  /***************************************************************************
  ** 
  ** Tour down
  */

  public TourStatus goDown(boolean selectedOnly) {
    return (goUpDown(false, selectedOnly));
  }  
  
  /***************************************************************************
  ** 
  ** Zoom to current tour stop
  */

  public void zoomToTourStop() {
    zoomToRectangle(floaterSet_.tourRect);
    return;
  }  
  
  /***************************************************************************
  ** 
  ** Tour to drain zone
  */

  public TourStatus tourToDrainZone(boolean selectedOnly) {
    return (goToDrainZone(selectedOnly));
  }  
  
  /***************************************************************************
  ** 
  ** Clear the tour
  */

  public void clearTour() {
    floaterSet_.tourRect = null;
    handleFloaterChange();
    return;
  }  

  /***************************************************************************
  ** 
  ** Find out updated tour directions
  */

  public TourStatus getTourDirections(boolean selectedOnly) {
    if (tourFocus_ == null) {
      return (null);
    }
    return (handleTourStop(tourFocus_.x, tourFocus_.y, Integer.valueOf(tourFocus_.x), 
                           (selectedOnly) ? bfn_.getNodeIDForRow(Integer.valueOf(tourFocus_.y)) : null));
  }   
    
  /***************************************************************************
  ** 
  ** Tour guts
  */

  private TourStatus goToDrainZone(boolean selectedOnly) {
    // FIX ME? Issues with this working with non-drain nodes?
     UiUtil.fixMePrintout("Handle cycling through multiple drain zones");
    NetNode nodeName = bfn_.getNodeIDForRow(Integer.valueOf(tourFocus_.y));
    if (nodeName == null) {
      return (null);
    }
    BioFabricNetwork.NodeInfo node = bfn_.getNodeDefinition(nodeName);
    boolean showShadows = FabricDisplayOptionsManager.getMgr().getDisplayOptions().getDisplayShadows();
    
    
    int colMin;
    int colMax;
    
    List<BioFabricNetwork.DrainZone> dropZone = node.getDrainZones(showShadows);
    MinMax nmm = null;
    if (dropZone == null) {
      nmm = node.getColRange(showShadows);
      colMin = nmm.max; // Yes, both are max
      colMax = nmm.max;
    } else {
      UiUtil.fixMePrintout("Crash here on go to drain zone in tour: array size 0");
      colMin = dropZone.get(0).getMinMax().min; // QUICK FIX WILL MAKE BETTER SOLUTION LATER
      colMax = dropZone.get(0).getMinMax().max;
    }
    
    int start = colMin;
    int end = colMax;
    for (int i = start; i <= end; i++) {
      Integer testCol = Integer.valueOf(i);
      if (selectedOnly) {
        if (!currColSelections_.contains(testCol)) {
          continue;
        }
      }
      NetNode target = bfn_.getTargetIDForColumn(testCol, showShadows);
      NetNode source = bfn_.getSourceIDForColumn(testCol, showShadows);
      if ((target != null) && (source != null)) {
        if (target.equals(nodeName) || source.equals(nodeName)) {
          return (handleTourStop(i, tourFocus_.y, testCol, (selectedOnly) ? nodeName : null));
        }
      }
    }

    return (null);
  }  
  
  /***************************************************************************
  ** 
  ** Tour guts
  */

  private TourStatus goLeftRight(int inc, boolean selectedOnly) {
    NetNode nodeName = bfn_.getNodeIDForRow(Integer.valueOf(tourFocus_.y));
    if (nodeName == null) {
      return (null);
    }
    BioFabricNetwork.NodeInfo node = bfn_.getNodeDefinition(nodeName);
    boolean showShadows = FabricDisplayOptionsManager.getMgr().getDisplayOptions().getDisplayShadows();
    MinMax nmm = node.getColRange(showShadows);
    int colMin = nmm.min;
    int colMax = nmm.max;
    int currCol = tourFocus_.x;
    if (currCol < colMin) {
      currCol = colMin - 1;  // So name starts begin at first link
    }
    if (currCol > colMax) {
      currCol = colMax;
    }

    int start = currCol + inc;
    int end = (inc == 1) ? colMax : colMin;
    for (int i = start; (inc == 1) ? (i <= end) : (i >= end); i += inc) {
      Integer testCol = Integer.valueOf(i);
      if (selectedOnly) {
        if (!currColSelections_.contains(testCol)) {
          continue;
        }
      }
      NetNode target = bfn_.getTargetIDForColumn(testCol, showShadows);
      NetNode source = bfn_.getSourceIDForColumn(testCol, showShadows);
      if ((target != null) && (source != null)) {
        if (target.equals(nodeName) || source.equals(nodeName)) {
          return (handleTourStop(i, tourFocus_.y, testCol, (selectedOnly) ? nodeName : null));
        }
      }
    }

    //
    // If we tried going left but had no success, we always have the target if we are
    // not-selection-bound or have a selected node:
    //
    boolean nodeAlive = (!selectedOnly) ? true : currNodeSelections_.contains(nodeName);
    if ((inc == -1) && nodeAlive) { 
      Rectangle2D targName = nodeNameLocations_.get(nodeName);
      Point targNameRC = worldToRowCol(new Point2D.Double(targName.getCenterX(), targName.getCenterY()));
      int useCol = targNameRC.x;
      return (handleTourStop(useCol, tourFocus_.y, Integer.valueOf(useCol), nodeName));
    }

    return (null);
  }

  /***************************************************************************
  ** 
  ** Tour guts
  */

  private TourStatus goFarLeftRight(boolean goRight, boolean selectedOnly) {
    NetNode nodeName = bfn_.getNodeIDForRow(Integer.valueOf(tourFocus_.y));
    if (nodeName == null) {
      return (null);
    }
    boolean showShadows = FabricDisplayOptionsManager.getMgr().getDisplayOptions().getDisplayShadows();
    BioFabricNetwork.NodeInfo node = bfn_.getNodeDefinition(nodeName);
    MinMax range = node.getColRange(showShadows);
    int useCol = (goRight) ? range.max : range.min;
    
    //
    // The far left is ALWAYS the target name if we are not-selection-bound or
    // have a selected node:
    
    boolean nodeAlive = (!selectedOnly) ? true : currNodeSelections_.contains(nodeName);
    if (!goRight && nodeAlive) { 
      Rectangle2D targName = nodeNameLocations_.get(nodeName);
      Point targNameRC = worldToRowCol(new Point2D.Double(targName.getCenterX(), targName.getCenterY()));
      useCol = targNameRC.x;
      return (handleTourStop(useCol, tourFocus_.y, Integer.valueOf(useCol), nodeName));
    }

    //
    // If we are working with selected only, we must find the left or rightmost
    // selected column for the target
    //
    
    if (selectedOnly) {
      int end = tourFocus_.y;
      int inc = (goRight) ? -1 : 1;
      for (int i = useCol; (inc == 1) ? (i <= end) : (i >= end); i += inc) {
        Integer testCol = Integer.valueOf(i);
        if (!currColSelections_.contains(testCol)) {
          continue;
        }
        NetNode target = bfn_.getTargetIDForColumn(testCol, showShadows);
        NetNode source = bfn_.getSourceIDForColumn(testCol, showShadows);
        if ((target != null) && (source != null)) {
          if (target.equals(node.getNodeIDWithName()) || source.equals(node.getNodeIDWithName())) {
            return (handleTourStop(useCol, tourFocus_.y, testCol, nodeName));
          }
        }
      }
    } else {
      Integer testCol = Integer.valueOf(useCol); 
      NetNode target = bfn_.getTargetIDForColumn(testCol, showShadows);
      NetNode source = bfn_.getSourceIDForColumn(testCol, showShadows);
      if ((target != null) && (source != null)) {
        return (handleTourStop(useCol, tourFocus_.y, testCol, null));
      }   
    }   
    return (null);
  }
  
  /***************************************************************************
  ** 
  ** For finding stops that are selected
  */

  private SortedSet<Integer> findSelectedLinkStops(NetNode nodeName) {
    TreeSet<Integer> retval = new TreeSet<Integer>();
    boolean showShadows = FabricDisplayOptionsManager.getMgr().getDisplayOptions().getDisplayShadows();
    BioFabricNetwork.NodeInfo node = bfn_.getNodeDefinition(nodeName);
    MinMax range = node.getColRange(showShadows);
    int start = range.min;
    int end = range.max;

    for (int i = start; i <= end; i++) {
      Integer testCol = Integer.valueOf(i);
      if (!currColSelections_.contains(testCol)) {
        continue;
      }
      NetNode target = bfn_.getTargetIDForColumn(testCol, showShadows);
      NetNode source = bfn_.getSourceIDForColumn(testCol, showShadows);
      if ((target != null) && (source != null)) {
        if (target.equals(node.getNodeIDWithName()) || source.equals(node.getNodeIDWithName())) { 
          retval.add(testCol);
        }
      }
    }
    return (retval);
  }  
  
  /***************************************************************************
  ** 
  ** Answer if the current tour stop is now unselected
  */

  public boolean tourStopNowUnselected() {
    if (tourFocus_ == null) {
      return (false);
    }
    return (tourStopIsUnselected(tourFocus_));
  }
  
  /***************************************************************************
  ** 
  ** Answer if the current tour stop is unselected
  */

  private boolean tourStopIsUnselected(Point focusPoint) {
    if (focusPoint == null) {
      return (false);
    }
    if (currColSelections_.isEmpty() && currNodeSelections_.isEmpty()) {
      return (false);
    }
    Integer rowObj = Integer.valueOf(focusPoint.y);
    Integer colObj = Integer.valueOf(focusPoint.x);
    
    //
    // Outside of row bounds:
    //
    NetNode nodeName = bfn_.getNodeIDForRow(rowObj);
    if (nodeName == null) {
      return (false);
    }
    
    //
    // On an name, depends on node selections:
    //
    Rectangle2D targName = nodeNameLocations_.get(nodeName);
    Point targNameRC = worldToRowCol(new Point2D.Double(targName.getCenterX(), targName.getCenterY()));
    boolean onTargName = focusPoint.equals(targNameRC);
    if (onTargName) {
      return (!currNodeSelections_.contains(nodeName));
    } 
    
    //
    // Else depends on links.  First, out of bounds:
    //

    boolean showShadows = FabricDisplayOptionsManager.getMgr().getDisplayOptions().getDisplayShadows();
    if (bfn_.getLinkDefinition(colObj, showShadows) == null) {
      return (false);
    }

    SortedSet<Integer> okStops = findSelectedLinkStops(nodeName);
    return (!okStops.contains(colObj));
  }  
   
 
  /***************************************************************************
  ** 
  ** Setup and bookkeeping after tour stop is changed
  */

  private TourStatus handleTourStop(int x, int y, Integer col, NetNode selectedOnlyNodeName) {      
    tourFocus_.setLocation(x, y);
    floaterSet_.tourRect = buildFocusBox(tourFocus_);
    handleFloaterChange();
    centerOnRectangle(floaterSet_.tourRect);
    fmt_.setCenter(rowColToWorld(tourFocus_), tourFocus_, true);
    MouseLocInfo vals = buildMouseLocation(tourFocus_);
    SortedSet<Integer> okStops = (selectedOnlyNodeName == null) ? null : findSelectedLinkStops(selectedOnlyNodeName);
    boolean nodeAlive = (selectedOnlyNodeName == null) ? true : currNodeSelections_.contains(selectedOnlyNodeName);
    boolean showShadows = FabricDisplayOptionsManager.getMgr().getDisplayOptions().getDisplayShadows();
    BioFabricNetwork.LinkInfo ld = bfn_.getLinkDefinition(col, showShadows);
    if ((ld != null) && !ld.inLinkRowRange(tourFocus_.y)) {
      ld = null;
    }
    return (new TourStatus(vals, bfn_, ld, tourFocus_, okStops, nodeAlive, tourStopIsUnselected(tourFocus_)));
  }  

  /***************************************************************************
  ** 
  ** Tour guts
  */

  private TourStatus goUpDown(boolean up, boolean selectedOnly) {
    if ((tourFocus_.y < 0) || (tourFocus_.y >= bfn_.getRowCount())) {
      return (null);
    }
    boolean showShadows = FabricDisplayOptionsManager.getMgr().getDisplayOptions().getDisplayShadows();
    Integer useCol = Integer.valueOf(tourFocus_.x);
    BioFabricNetwork.LinkInfo link = bfn_.getLinkDefinition(useCol, showShadows);
    if (link == null) {
      return (null);
    }
    int useX = tourFocus_.x;
    int useY;
    if (up) {
      if (tourFocus_.y > link.getEndRow()) {
        useY = link.getEndRow();
      } else if (tourFocus_.y > link.getStartRow()) {
        useY = link.getStartRow();
      } else {
        useY = link.getEndRow();
      }
    } else {
      if (tourFocus_.y < link.getStartRow()) {
        useY =  link.getStartRow();
      } else if (tourFocus_.y < link.getEndRow()) {
        useY = link.getEndRow();
      } else {
        useY = link.getStartRow();
      }       
    }
    return (handleTourStop(useX, useY, useCol, (selectedOnly) ? bfn_.getNodeIDForRow(Integer.valueOf(useY)) : null));
  }

  /***************************************************************************
  ** 
  ** Collect a zoom rectangle
  */

  public void setToCollectZoomRect() {
    bfw_.disableControls(CommandSet.ALLOW_NAV_PUSH, false);
    collectingZoomMode_ = true;
    firstZoomPoint_ = null;
    cursorMgr_.showModeCursor();
    return;
  }
  
  /***************************************************************************
  ** 
  ** Collect a tour start
  */
  
  public void setToCollectTourStart(boolean selectionOnly) {
    bfw_.disableControls(CommandSet.ALLOW_NAV_PUSH, false);
    collectingTourStart_ = true;
    tourStartSelectionOnly_ = selectionOnly;
    cursorMgr_.showModeCursor();
    return;
  }
    
  /***************************************************************************
  ** 
  ** Cancel current modal
  */

  public void cancelModals() { 
    collectingZoomMode_ = false;
    collectingTourStart_ = false;
    tourStartSelectionOnly_ = false;
    firstZoomPoint_ = null;
    floaterSet_.floater = null;
    handleFloaterChange();
    cursorMgr_.showDefaultCursor();
    bfw_.reenableControls();
    return;
  }

  /***************************************************************************
  ** 
  ** Bump to next selection
  */

  public void incrementToNextSelection() { 
    bumpNextSelection();
    return;
  }
  
  /***************************************************************************
  ** 
  ** Bump to next selection
  */

  public void decrementToPreviousSelection() { 
    bumpPreviousSelection();
    return;
  }  
  
 /***************************************************************************
  ** 
  ** reg pane
  */

  public void setScroll(JScrollPane jsp) { 
    jsp_ = jsp;
    // Now in V2, we use wheel scroll to zoom (thanks anonymous reviewer #1!) not
    // to do vertical scrolling, which is useless anyway.
    //
    jsp_.setWheelScrollingEnabled(false);
    jsp_.addMouseWheelListener(new MouseWheelListener() {
	    public void mouseWheelMoved(MouseWheelEvent e) {
	    	getZoomController().bumpZoomWrapper((e.getWheelRotation() > 0) ? '-' : '+');
	    }
    });
    return;
  }
  
  /***************************************************************************
  **
  ** Update zoom state
    
    
  void updateZoom(char sign) {
    try {
      getZoomController().bumpZoomWrapper(sign);
      repaint();
    } catch (Exception ex) {
      ExceptionHandler.getHandler().displayException(ex);
    }
    return;
  }     

  /***************************************************************************
  ** 
  ** Change the paint
  */

  public void changePaint(BTProgressMonitor monitor) throws AsynchExitRequestException {
    if (bufferBuilder_ != null) {
      bufferBuilder_.release();
    }    
    if (bfn_ == null) {
      return;
    }
    FabricDisplayOptions fdo = FabricDisplayOptionsManager.getMgr().getDisplayOptions();
    boolean shadeNodes = fdo.getShadeNodes();
    boolean showShadows = fdo.getDisplayShadows();
   
    BioFabricNetwork.Extents ext = new BioFabricNetwork.Extents(bfn_, monitor);
    painter_.buildObjCache(bfn_.getNodeDefList(), bfn_.getLinkDefList(showShadows), shadeNodes, 
									         showShadows, ext, new HashMap<NetNode, Rectangle2D>(), 
									         new HashMap<NetNode, List<Rectangle2D>>(), worldRectNetAR_, 
									         bfn_.getNodeAnnotations(), bfn_.getLinkAnnotations(Boolean.valueOf(showShadows)), monitor);
    
    handleFloaterChange();
    return;
  }
 
  /***************************************************************************
  ** 
  ** Install a model
  */

  public void installModel(BioFabricNetwork bfn, BTProgressMonitor monitor) throws AsynchExitRequestException {
    bfn_ = bfn;
    FabricDisplayOptions fdo = FabricDisplayOptionsManager.getMgr().getDisplayOptions();
    boolean shadeNodes = fdo.getShadeNodes();
    boolean showShadows = fdo.getDisplayShadows();
     
    int cols = bfn_.getColumnCount(showShadows);
    int rows = bfn_.getRowCount();
    
  
    double netWidth = cols * GRID_SIZE;
    double netHeight = rows * GRID_SIZE;
  
    Rectangle2D linksAndNodes = new Rectangle2D.Double(0.0, 0.0, netWidth, netHeight);
 
    nodeNameLocations_ = new HashMap<NetNode, Rectangle2D>();
    drainNameLocations_ = new HashMap<NetNode, List<Rectangle2D>>();
    BioFabricNetwork.Extents ext = new BioFabricNetwork.Extents(bfn_, monitor);
    Rectangle2D fullNetRect = painter_.buildObjCache(bfn_.getNodeDefList(), bfn_.getLinkDefList(showShadows), 
													    		                   shadeNodes, showShadows, ext, nodeNameLocations_, 
													    		                   drainNameLocations_, linksAndNodes, 
													    		                   bfn_.getNodeAnnotations(), 
													    		                   bfn_.getLinkAnnotations(Boolean.valueOf(showShadows)), monitor);
    
    double ulPtx = PAD_MULT_ * fullNetRect.getWidth();
    double ulPty = PAD_MULT_ * fullNetRect.getHeight();
    
    worldRectNetAR_ = new Rectangle2D.Double(fullNetRect.getX() - ulPtx, fullNetRect.getY() - ulPty,
    		                                     fullNetRect.getWidth() + (2.0 * ulPtx), 
    		                                     fullNetRect.getHeight()+ (2.0 * ulPty));
    UiUtil.force2DToGrid(worldRectNetAR_, GRID_SIZE);
    
    zoomer_.setWorldRect(UiUtil.rectFromRect2D(worldRectNetAR_));
 
    bucketRend_.buildBucketCache(bfn_.getNodeDefList(), bfn_.getLinkDefList(showShadows), 
    		                         bfn_.getNodeAnnotations(), bfn_.getLinkAnnotations(Boolean.valueOf(showShadows)),
    		                         ext, showShadows);
      
    LoopReporter lr = new LoopReporter(drainNameLocations_.size(), 20, monitor, 0.0, 1.0, "progress.drainsToQuad");

    forSelections_ = new QuadTree(fullNetRect, 5);
    Iterator<NetNode> kit = drainNameLocations_.keySet().iterator();
    int count = 0;
    while (kit.hasNext()) {
    	NetNode nid = kit.next();
    	lr.report();
    	List<Rectangle2D> rects = drainNameLocations_.get(nid);
    	int numR = rects.size();
    	for (int i = 0; i < numR; i++) {
    	  String key = Integer.toString(count++);
    	  QuadTree.Payload pay = new QuadTree.Payload(rects.get(i), key);
    	  forSelections_.insertPayload(pay);
    	}
    }
    lr.finish();
    if (fnt_ != null) {
      fnt_.haveAModel(true);
    }
    return;
  }
  
  /***************************************************************************
  ** 
  ** Install a model
  */

  public void installModelPost() { 
    fmt_.setModel(bfn_);
    if (myPanel_ != null) {
      myPanel_.requestFocus();
    }
    return;
  }
  
  /***************************************************************************
  ** 
  ** Needed? Yes.
  */

  public void initZoom() { 
    getZoomController().zoomToModel(true);
    // Prior to V2 Beta, we called this:
    // zoomer_.setWorldRect(UiUtil.rectFromRect2D(getFullScreenWorld()));   
    // Previously, "This is needed to make sure that the full-on view of the model
    // encapsulates the whole workspace.  Otherwise, drag rects on
    // e.g. long thin models go outside the workspace and we don't
    // actually zoom!"
    //
    // BUT! This is what "bulked up" the world workspace vertically so that the right scrollbar became
    // *completely useless* on long thin models! In that case e.g. 99% of the scroll travel
    // was for blank space above and below the model.
    // 
    // Instead, the correct approach was to ditch the BioTapestry-centric approach that
    // took the UNION of the world workspace and the selected rect, and if it was not equal,
    // it centered on the UNION. That was useless. That is fixed (see 
    // ZoomCommandSupport.zoomToSelectedGutsGuts(), so we abandon fullScreenWorld bit.
   
    return;
  }
  
  /***************************************************************************
  **
  ** Find the viewport in world coordinates
  */
  
  public Rectangle2D getViewInWorld(JViewport view) {
    Point viewPos = view.getViewPosition();
    Dimension viewDim = view.getExtentSize();
    Point2D vrul = viewToWorld(viewPos);
    Point2D vrlr = viewToWorld(new Point(viewPos.x + viewDim.width, viewPos.y + viewDim.height));
    Rectangle2D viewInWorld = new Rectangle2D.Double(vrul.getX(), vrul.getY(), vrlr.getX() - vrul.getX(), vrlr.getY() - vrul.getY());
    return (viewInWorld);
  } 
  
  /***************************************************************************
  **
  ** Find the viewport in world coordinates
  */
  
  public Rectangle2D getViewInWorld() {
    JViewport view = jsp_.getViewport();
    return (getViewInWorld(view));
  } 
  
  /***************************************************************************
  **
  ** Find the full extent of the view! Note we are not using this anymore
  
  public Rectangle getFullScreenWorld() {
    JViewport view = jsp_.getViewport();
    Point viewPos = new Point(0, 0);
    Dimension viewDim = view.getViewSize();
    Point2D vrul = viewToWorld(viewPos);
    Point2D vrlr = viewToWorld(new Point(viewPos.x + viewDim.width, viewPos.y + viewDim.height));
    Rectangle fullScreenWorld = new Rectangle((int)vrul.getX(), (int)vrul.getY(), 
                                              (int)(vrlr.getX() - vrul.getX()), 
                                              (int)(vrlr.getY() - vrul.getY()));
    return (fullScreenWorld);
  } 

  /***************************************************************************
  **
  ** Drawing routine
  */
  
  public void selectionsToSubmodel() { 
    transmitSelections(targetList_, linkList_);
    return;  
  }
 
  public void setFullModelViewPos(Point vp) {
  	fullModelPos_ = vp;
  	return;
  	
  }
  
  public void setFullModelExtent(Dimension dim) {
  	fullModelExtent_ = dim;
  	return;
  }

  /***************************************************************************
  **
  ** Drawing core
  */
  
  private void drawingGuts(Graphics g, Rectangle2D viewRect) {
    Graphics2D g2 = (Graphics2D)g;   
    g2.setRenderingHint(RenderingHints.KEY_ANTIALIASING, RenderingHints.VALUE_ANTIALIAS_ON);
    g2.setRenderingHint(RenderingHints.KEY_FRACTIONALMETRICS, RenderingHints.VALUE_FRACTIONALMETRICS_ON);
    AffineTransform saveTrans = g2.getTransform();   
    clipRect2_.setBounds((int)viewRect.getX(), (int)viewRect.getY(), (int)viewRect.getWidth(), (int)viewRect.getHeight());
    g2.transform(zoomer_.getTransform());
    BasicStroke selectedStroke = new BasicStroke(PaintCacheSmall.STROKE_SIZE, BasicStroke.CAP_SQUARE, BasicStroke.JOIN_MITER);    
    g2.setStroke(selectedStroke);
    
    // USE THIS INSTEAD???
    //g2.setBackground(Color.WHITE);
    //g2.clearRect(clip.x, clip.y, clip.width, clip.height);
    
    
    g2.setPaint(Color.WHITE);
    g2.drawRect(clipRect2_.x, clipRect2_.y, clipRect2_.width, clipRect2_.height); 
    painter_.paintIt(g2, clipRect2_, null);
    
    if (!targetList_.isEmpty() || !linkList_.isEmpty()) {
      g2.setTransform(saveTrans);
      drawSelections(g2, clipRect2_);
      g2.setTransform(saveTrans);
      g2.transform(zoomer_.getTransform());
    }
    drawFloater(g2, false);
    
    return;
  }
  
  /***************************************************************************
  **
  ** Drawing core
  */
  
  private void drawFloater(Graphics2D g2, boolean needInit) {
    if (floaterSet_.isEmpty()) {
      return;
    }        
    AffineTransform saveTrans = null;
    if (needInit) {
      g2.setRenderingHint(RenderingHints.KEY_ANTIALIASING, RenderingHints.VALUE_ANTIALIAS_ON);
      g2.setRenderingHint(RenderingHints.KEY_FRACTIONALMETRICS, RenderingHints.VALUE_FRACTIONALMETRICS_ON);
      saveTrans = g2.getTransform();   
      g2.transform(zoomer_.getTransform());  
    }
    painter_.drawFloater(g2, floaterSet_);    
    if (needInit) {
      g2.setTransform(saveTrans);
    } 
    return;
  }

  /***************************************************************************
  **
  ** Support printing
  */  
  
  public int print(Graphics g, PageFormat pf, int pageIndex) {
    if (pageIndex != 0) {
      return (NO_SUCH_PAGE);
    }
    double px = pf.getImageableX();
    double py = pf.getImageableY();
    double pw = pf.getImageableWidth();
    double ph = pf.getImageableHeight();
 
    Rectangle worldPiece = UiUtil.rectFromRect2D(worldRectNetAR_);
     
    double wFrac = worldPiece.width / pw;
    double hFrac = worldPiece.height / ph;
    double frac = 1.0 / ((hFrac > wFrac) ? hFrac : wFrac);
    
    AffineTransform trans = new AffineTransform();
    trans.translate(px  + (pw / 2.0), py + (ph / 2.0));
    trans.scale(frac, frac);
    trans.translate(-worldPiece.getCenterX(), -worldPiece.getCenterY());
    Graphics2D g2 = (Graphics2D)g;
    g2.setColor(Color.white);     
    g2.setRenderingHint(RenderingHints.KEY_ANTIALIASING, RenderingHints.VALUE_ANTIALIAS_ON);
    g2.setRenderingHint(RenderingHints.KEY_FRACTIONALMETRICS, RenderingHints.VALUE_FRACTIONALMETRICS_ON);
    BasicStroke selectedStroke = new BasicStroke(PaintCacheSmall.STROKE_SIZE, BasicStroke.CAP_SQUARE, BasicStroke.JOIN_MITER);    
    g2.setStroke(selectedStroke);
    g2.transform(trans); 
    g2.fillRect(worldPiece.x, worldPiece.y, worldPiece.width, worldPiece.height);
    painter_.paintIt(g2, worldPiece, null);
    return (PAGE_EXISTS);
  }
 
  /***************************************************************************
  **
  ** Drawing core
  */
  
  public boolean drawForBuffer(BufferedImage bi, Rectangle2D clip, Dimension screenDim, 
  		                         Rectangle2D worldRec, int heightPad, double linksPerPixel) { 
  	Graphics2D g2 = bi.createGraphics();
    g2.setColor(Color.WHITE);
    g2.fillRect(0, 0, screenDim.width, screenDim.height + heightPad);
    double zoomH = screenDim.getWidth() / worldRec.getWidth();
    double zoomV = screenDim.getHeight() / worldRec.getHeight();
    double zoom = Math.max(zoomH, zoomV); //Math.min(zoomH, zoomV);
    Point2D centerW = new Point2D.Double(worldRec.getX() + (worldRec.getWidth() / 2.0), worldRec.getY() + (worldRec.getHeight() / 2.0));
    AffineTransform transform = new AffineTransform();
    transform.translate(screenDim.getWidth() / 2.0, screenDim.getHeight() / 2.0);
    transform.scale(zoom, zoom);
    transform.translate(-centerW.getX(), -centerW.getY());

    g2.setRenderingHint(RenderingHints.KEY_ANTIALIASING, RenderingHints.VALUE_ANTIALIAS_ON);
    g2.setRenderingHint(RenderingHints.KEY_FRACTIONALMETRICS, RenderingHints.VALUE_FRACTIONALMETRICS_ON);
    BasicStroke selectedStroke = new BasicStroke(PaintCacheSmall.STROKE_SIZE, BasicStroke.CAP_SQUARE, BasicStroke.JOIN_MITER);    
    g2.setStroke(selectedStroke);
    g2.setTransform(transform); 
    boolean retval = painter_.paintIt(g2, UiUtil.rectFromRect2D(clip), null);
    
    // To debug sizing problems, this draws a green bounding rectangle:
    //AffineTransform transformx = new AffineTransform();
    //g2.setTransform(transformx);
    //BasicStroke selectedStrokex = new BasicStroke(1, BasicStroke.CAP_SQUARE, BasicStroke.JOIN_MITER);    
    //g2.setStroke(selectedStrokex);  
    //g2.setColor(Color.GREEN);
    //g2.drawRect(0, 0, screenDim.width - 1, screenDim.height - 1);

    g2.dispose();
    return (retval);
  }
  
  /***************************************************************************
  **
  ** Drawing core
  */
  
  public boolean drawForPrint(Graphics g, Rectangle clip, Dimension screenDim, Rectangle worldRec) {
    Graphics2D g2 = (Graphics2D)g;
    double zoomH = screenDim.getWidth() / worldRec.getWidth();
    double zoomV = screenDim.getHeight() / worldRec.getHeight();
    double zoom = Math.max(zoomH, zoomV); //Math.min(zoomH, zoomV);
    Point2D centerW = new Point2D.Double(worldRec.getX() + (worldRec.getWidth() / 2.0), worldRec.getY() + (worldRec.getHeight() / 2.0));
    AffineTransform transform = new AffineTransform();
    transform.translate(screenDim.getWidth() / 2.0, screenDim.getHeight() / 2.0);
    transform.scale(zoom, zoom);
    transform.translate(-centerW.getX(), -centerW.getY());

    g2.setRenderingHint(RenderingHints.KEY_ANTIALIASING, RenderingHints.VALUE_ANTIALIAS_ON);
    g2.setRenderingHint(RenderingHints.KEY_FRACTIONALMETRICS, RenderingHints.VALUE_FRACTIONALMETRICS_ON);
    BasicStroke selectedStroke = new BasicStroke(PaintCacheSmall.STROKE_SIZE, BasicStroke.CAP_SQUARE, BasicStroke.JOIN_MITER);    
    g2.setStroke(selectedStroke);
    g2.setTransform(transform);
    boolean retval = painter_.paintIt(g2, clip, null);
    // Debug sizing problems
    //g2.setColor(Color.red);
    //g2.drawRect(clip.x, clip.y, clip.width - 1, clip.height - 1);
    return (retval);
  }
  
  /***************************************************************************
  **
  ** Draw the selections as fully colored over a partially opaque overlay
  */
  
  public void drawSelections(Graphics2D g2, Rectangle clip) {
    
    FabricDisplayOptions fdo = FabricDisplayOptionsManager.getMgr().getDisplayOptions();
  
    Rectangle viewRect = jsp_.getViewport().getViewRect();
    
    if ((bim_ == null) || (bim_.getHeight() != viewRect.height) || (bim_.getWidth() != viewRect.width)) {
      bim_ = new BufferedImage(viewRect.width, viewRect.height, BufferedImage.TYPE_4BYTE_ABGR);
    }
    Graphics2D ig2 = bim_.createGraphics();
    ig2.setTransform(new AffineTransform());
    Color drawCol = new Color(1.0f, 1.0f, 1.0f, (float)fdo.getSelectionOpaqueLevel()); 
    ig2.setBackground(drawCol);
    ig2.clearRect(0, 0, viewRect.width, viewRect.height);
    ig2.setRenderingHint(RenderingHints.KEY_ANTIALIASING, RenderingHints.VALUE_ANTIALIAS_ON);
    ig2.setRenderingHint(RenderingHints.KEY_FRACTIONALMETRICS, RenderingHints.VALUE_FRACTIONALMETRICS_ON);
    BasicStroke selectedStroke = new BasicStroke(PaintCacheSmall.STROKE_SIZE, BasicStroke.CAP_SQUARE, BasicStroke.JOIN_MITER);    
    ig2.setStroke(selectedStroke);
          
    AffineTransform overTrans;
    AffineTransform otp = new AffineTransform();
    // The zoomer transform is a combination of the zoom, plus needed offset when we
    // have zoomed out and no longer have scrollbars:
    overTrans = new AffineTransform(zoomer_.getTransform());
     
    // Need to offset the origin to handle view rectangle offset:
    otp.translate(-viewRect.getX(), -viewRect.getY());
    // This needs to be pre-concatenated to have it "done first"
    overTrans.preConcatenate(otp);
    ig2.setTransform(overTrans);
  
    //
    // Render the image:
    //
    //ig2.setComposite(AlphaComposite.Clear);
    
    ig2.setComposite(AlphaComposite.Src);
    painter_.paintIt(ig2, clip, selections_);

    g2.drawImage(bim_, viewRect.x, viewRect.y, viewRect.width, viewRect.height, null);
    return;
  }
   
  /***************************************************************************
  **
  ** Zoom to selected
  */
  
  public void zoomToSelected() {
    zcs_.zoomToSelected();
    return;
  }  
  
  /***************************************************************************
  **
  ** Zoom to selected
  */
  
  public void zoomToRectangle(Rectangle rect) {
    zcs_.zoomToRect(rect);
    return;
  }  
 
  /***************************************************************************
  **
  ** Center on selected
  */
  
  public void centerOnSelected() {
    zcs_.centerOnSelected();
    return;
  }  
  
  /***************************************************************************
  **
  ** Center on rectangle
  */
  
  public void centerOnRectangle(Rectangle rect) {
    zcs_.centerOnRectangle(rect);
    return;
  }  
   
  /***************************************************************************
  **
  ** Get zoom support
  */
  
  public ZoomCommandSupport getZoomController() {
    return (zcs_);
  }  
  
  /***************************************************************************
  **
  ** Return the required size of the layout
  */
  
  public Rectangle getRequiredSize() {
    return (UiUtil.rectFromRect2D(worldRectNetAR_));
  }
  
  /***************************************************************************
  **
  ** Return the required size of the layout
  */
  
  public Rectangle getWorldScreen() {
    return (UiUtil.rectFromRect2D(this.worldRectScreenAR_));
  }

  /***************************************************************************
  **
  ** Return the required size of the selected items.  Null if nothing is
  ** selected.
  */
  
  public Rectangle getSelectionSize() {
    int numRect = rects_.size();
    if (numRect == 0) {
      return (null);
    }
    Rectangle retval = (Rectangle)(rects_.get(0).clone());
    for (int i = 1; i < numRect; i++) {
      Rectangle.union(retval, rects_.get(i), retval);
    }
    return (retval);
  }

  /***************************************************************************
  **
  ** Return the required size of the current zoom selection item.  Null if nothing is
  ** selected.
  */
  
  public Rectangle getCurrentSelectionSize() {
    if (currSel_ == -1) {
      return (null);
    }
    return (rects_.get(currSel_));    
  }  

  /***************************************************************************
  **
  ** Answer if we have a current selection
  */
  
  public boolean haveCurrentSelection() {
    return (currSel_ != -1);
  }
  
  /***************************************************************************
  **
  ** Answers if we multiple selections (i.e. can cycle through selections) 
  */
  
  public boolean haveMultipleSelections() {  
    return (rects_.size() > 1);
  }
 
  /***************************************************************************
  **
  ** Bump to the previous selection for zoom
  */
  
  public void bumpPreviousSelection() {
    currSel_--;
    if (currSel_ < 0) {
      currSel_ = rects_.size() - 1;
    }
    bumpGuts();
    return;     
  }
  
  /***************************************************************************
  **
  ** Bump to the next selection for zoom
  */
  
  public void bumpNextSelection() {
    currSel_ = (currSel_ + 1) % rects_.size();
    bumpGuts();
    return;
  }

  /***************************************************************************
  **
  ** Bump guts
  */
  
  private void bumpGuts() {
    if (rects_.isEmpty()) {
      fnt_.haveASelection(false);
    } else {
      // FIX ME:  Have seen cases where currSel_ is larger than rects_ on submodel build?
      if (currSel_ <= (rects_.size() - 1)) {
        Rectangle rect = rects_.get(currSel_);
        floaterSet_.currSelRect = (Rectangle)rect.clone();  // world coords
        selFocus_ = worldToRowCol(new Point2D.Double(floaterSet_.currSelRect.getCenterX(), floaterSet_.currSelRect.getCenterY()));
        fmt_.setCenter(rowColToWorld(selFocus_), selFocus_, true);
        fnt_.haveASelection(true);
      }
    }
    handleFloaterChange();  // else tour rect does not redraw
    return;
  }
    
  /***************************************************************************
  **
  ** We implement the ZoomPresentation interface!
  */
  
  public void setPresentationZoomFactor(double zoom) {
    // Don't need to do anything; this is a case where the target
    // interface, that we also implement, is doing the job already...
  }
 
  /***************************************************************************
  **
  ** Zoom support.  We implement the ZoomTarget interface!
  */  
  
  public Point pointToViewport(Point world) {
    return (zoomer_.pointToViewport(world));    
  }
  
  public Point2D getRawCenterPoint() {
    return (zoomer_.getRawCenterPoint());    
  }
  
  public Rectangle getSelectedBounds() {
    return (zoomer_.getSelectedBounds());
  }
  
  public Rectangle getCurrentSelectedBounds() {
    return (zoomer_.getCurrentSelectedBounds());    
  }  
  
  public Rectangle getWorldRect() {
    return (zoomer_.getWorldRect());    
  }
  
  public Dimension getPreferredSize() {
    return (this.myPanel_.getPreferredSize());    
  }
  
  public Dimension getBasicSize() {
    return (zoomer_.getBasicSize());    
  }
  
  public Point2D viewToWorld(Point vPt) {
    return (zoomer_.viewToWorld(vPt));    
  }
  
  public void setZoomFactor(double newZoomVal, Dimension vDim) {
    zoomer_.setZoomFactor(newZoomVal, vDim);
    return;
  }
  
  public Rectangle getCurrentBasicBounds() {
    return (zoomer_.getCurrentBasicBounds());    
  }

  public void adjustWideZoomForSize(Dimension dims) {
    zoomer_.adjustWideZoomForSize(dims);
    // Seeing lags where last resize not getting painted:
    repaint();
    return;
  }
   
  public Point getCenterPoint() {
    return (zoomer_.getCenterPoint());    
  }
   
  public double getZoomFactor() {
    return (zoomer_.getZoomFactor());    
  }

  /***************************************************************************
  **
  ** Useful for selection
  */  
  
  public Point transToRowCol(Point loc) {
    Point2D vrul = viewToWorld(loc);
    return (worldToRowCol(vrul));
  }
  
  /***************************************************************************
  **
  ** Useful for selection
  */  
  
  public Point worldToRowCol(Point2D wloc) {
    Point2D retval = (Point2D)wloc.clone();
    UiUtil.forceToGrid(wloc.getX(), wloc.getY(), retval, GRID_SIZE);
    return (new Point((int)retval.getX() / GRID_SIZE, (int)retval.getY() / GRID_SIZE));
  }
 
  /***************************************************************************
  **
  ** Useful for selection
  */  
  
  public Point2D rowColToWorld(Point loc) {
    int col = (loc.x * GRID_SIZE);
    int row = (loc.y * GRID_SIZE);
    return (new Point2D.Double(col, row));
  }
 
  /***************************************************************************
  **
  ** Useful for selection
  */  
  
  public Rectangle valsToRect(int sx, int sy, int ex, int ey, boolean convert) {           
    int x, y, width, height, endx, endy;
    if (convert) {      
      Point locs = new Point(sx, sy);
      Point spt = transToRowCol(locs);
      Point loce = new Point(ex, ey);
      Point ept = transToRowCol(loce);
      Point2D start = new Point2D.Double(spt.x, spt.y);
      Point2D end = new Point2D.Double(ept.x, ept.y);      
      x = (int)start.getX();
      y = (int)start.getY();
      width = (int)end.getX() - x;
      height = (int)end.getY() - y;
      endx = (int)end.getX();
      endy = (int)end.getY();
    } else {
      x = sx;
      y = sy;
      width = ex - sx;
      height = ey - sy;
      endx = ex;
      endy = ey;
    }

    int rx, ry, rw, rh;
    if ((width != 0) && (height != 0)) {
      if (width < 0) {
        rx = endx;
        rw = -width;
      } else {
        rx = x;
        rw = width;
      }
      if (height < 0) {
        ry = endy;
        rh = -height;
      } else {
        ry = y;
        rh = height;
      }
      return (new Rectangle(rx, ry, rw, rh));
    } else {
     return (null);
    }
  }  
 
  public static class MouseLocInfo {
  	public String nodeDesc;
  	public String linkDesc;
  	public String zoneDesc;
  	public String linkSrcDesc;
  	public String linkTrgDesc;
  	public ArrayList<String> nodeAnnotations;
  	public ArrayList<String> linkAnnotations;
  	
  	public MouseLocInfo(String nodeDesc, String linkDesc, String zoneDesc, 
  	                    String linkSrcDesc, String linkTrgDesc, 
  	                    String nodeAnnotation, String linkAnnotation) {
  		this.nodeDesc = nodeDesc;
  	  this.linkDesc = linkDesc;
  	  this.zoneDesc = zoneDesc;
  	  this.linkSrcDesc = linkSrcDesc;
  	  this.linkTrgDesc = linkTrgDesc;
  	  this.nodeAnnotations = new ArrayList<String>();
  	  this.nodeAnnotations.add(nodeAnnotation);
  	  this.linkAnnotations = new ArrayList<String>();
  	  this.linkAnnotations.add(linkAnnotation);
  	}
  	
  	public MouseLocInfo() {
  		this.nodeDesc = "<none>";
  	  this.linkDesc = "<none>";
  	  this.zoneDesc = "<none>";
  	  this.linkSrcDesc = "<none>";
  	  this.linkTrgDesc = "<none>";
  	  this.nodeAnnotations = new ArrayList<String>();
      this.nodeAnnotations.add("<none>");
      this.linkAnnotations = new ArrayList<String>();
      this.linkAnnotations.add("<none>");   
  	}  	
  }
  
  /***************************************************************************
  **
  ** build mouse location
  */ 
  
  MouseLocInfo buildMouseLocation(Point cprc) {
    MouseLocInfo retval = new MouseLocInfo();
    Integer colObj = Integer.valueOf(cprc.x);
    Integer rowObj = Integer.valueOf(cprc.y);
    NetNode target = bfn_.getNodeIDForRow(rowObj);
    boolean showShadows = FabricDisplayOptionsManager.getMgr().getDisplayOptions().getDisplayShadows();
    NetNode src = bfn_.getSourceIDForColumn(colObj, showShadows);
    NetNode trg = bfn_.getTargetIDForColumn(colObj, showShadows);
    NetNode drain = bfn_.getDrainForColumn(colObj, showShadows);

    retval.nodeAnnotations.clear();
    AnnotationSet ansn = bfn_.getNodeAnnotations();
    if (ansn != null) {
      AnnotsForPos a4pn = new AnnotationSetImpl.AnnotsForPosImpl(); // FIXME make an instance variable?
      ansn.fillAnnots(a4pn, rowObj);
      a4pn.displayStrings(retval.nodeAnnotations);
    }
    
    retval.linkAnnotations.clear();
    AnnotationSet ansl = bfn_.getLinkAnnotations(showShadows);
    if (ansl != null) {
      AnnotsForPos a4pl = new AnnotationSetImpl.AnnotsForPosImpl(); // FIXME make an instance variable?
      ansl.fillAnnots(a4pl, colObj);
      a4pl.displayStrings(retval.linkAnnotations);
    }
    
    int numRows = bfn_.getRowCount();
    if (target != null) {        
      BioFabricNetwork.NodeInfo ni = bfn_.getNodeDefinition(target);
      MinMax nimm = ni.getColRange(showShadows);
      if ((nimm.min <= cprc.x) && (nimm.max >= cprc.x)) {
        retval.nodeDesc = target.getName();           
      } else {
        Rectangle2D nnl = nodeNameLocations_.get(target);
        Point2D inWorld = rowColToWorld(cprc);
        UiUtil.fixMePrintout("zigg.sif plus zigg.noa see null ptr here *on mouse move during node-attribute relayout??*");
        if ((nnl == null) || (inWorld == null)) {
        	System.err.println("nnl " + nnl + " iw " + inWorld);
        	Thread.dumpStack();
        } else if (nnl.contains(inWorld)) {
          retval.nodeDesc = target.getName();
        }       
      }    
    }  
    if ((src != null) && (trg != null)) {
      BioFabricNetwork.LinkInfo li = bfn_.getLinkDefinition(colObj, showShadows);
      if (li != null) {
        int minRow = li.topRow();
        int maxRow = li.bottomRow();
        if ((minRow <= cprc.y) && (maxRow >= cprc.y)) {
        	FabricLink flink = li.getLink();
        	retval.linkDesc = flink.toDisplayString();
          retval.linkSrcDesc = flink.getSrcNode().getName();
          retval.linkTrgDesc = flink.getTrgNode().getName();
        }     
      }
    }
    if (drain != null) {        
      if ((0 <= cprc.x) && (numRows >= cprc.y)) {
        retval.zoneDesc = drain.getName();           
      }
    }  
    return (retval);
 }
  
  /***************************************************************************
  **
  ** Run the selection logic for point click. Point rcbp is for row-column, Point sloc
  ** is sub-row-column resolution.
  */  
  
  public void selectionLogicPoint(Point rcbp, Point sloc,
                                  boolean showShadows, Set<NetNode> nodes, Set<FabricLink> links, 
                                  Set<Integer> cols, boolean shiftPressed) { 


    if ((nodeNameLocations_ == null) || (drainNameLocations_ == null)) {
      return;
    }
  
    //
    // If shift pressed, we need ranges:
    //
    
    MinMax colRange = null;
    MinMax nodeRange = null;
    
    if (shiftPressed) {
      colRange = new MinMax();
      colRange.init();
      nodeRange = new MinMax();
      nodeRange.init(); 
      Iterator<Integer> cit = cols.iterator();
      while (cit.hasNext()) {
        Integer col = cit.next();
        colRange.update(col.intValue());
      }
      Iterator<NetNode> nit = nodes.iterator();
      while (nit.hasNext()) {
        NetNode node = nit.next();
        int row = bfn_.getNodeDefinition(node).nodeRow;
        nodeRange.update(row);
      }     
    }
 
    //
    // Clicking on one guy will make him the current selection!
    //

    int row = rcbp.y;
    int col = rcbp.x;

    //
    // One point clicks can select a drain name, but only if a link click
    // fails.  Note for this op, we need to use sub-row-col resolution to click
    // tiny lables, so we use sloc, not rcbp:
    //
    
    boolean nodeAdd = false;
    boolean linkAdd = false;

    //
    // Do we have a drain zone from a one point click?
    //
    
    NetNode gotDrain = null;
    Point2D worldPt = viewToWorld(sloc);
    Iterator<NetNode> dnlit = drainNameLocations_.keySet().iterator(); 
    while (dnlit.hasNext()) {
      NetNode target = dnlit.next();
      List<Rectangle2D> nameLocs = drainNameLocations_.get(target);
      for (Rectangle2D zone : nameLocs) {
        if (zone.contains(worldPt)) {
          gotDrain = target;
          break;
        }
      }
    }

    HashSet<String> foundKeys = new HashSet<String>();   
    forSelections_.getPayloadKeys(worldPt, foundKeys);

    UiUtil.fixMePrintout("READ ME FOR NEW POLICY");
    // Rectangle selection should pull in links with glyphs within the rectangle, along with nodes they
    // are incident upon. They should also always be additive.
    // Should be able to select a link (which includes endpoints) by clicking on it, if there is no
    // node that beats it (ambiguity favors the node). 

    boolean gotLink = false;
 
    //
    // Disaster! On Stanford net, a smallish box draw can capture 60,000 node rows and
    // 130,000 link rows. For those keeping track at home, that means this loop runs
    // 7.8 billion times.
    //
    
    Integer rowObj = Integer.valueOf(row);
    NetNode target = bfn_.getNodeIDForRow(rowObj);          
    Integer colObj = Integer.valueOf(col);
    
    //
    // We have hit a row with a node, but not a drain zone:
    //
    
    if ((target != null) && (gotDrain == null)) {
      //
      // See if we click on the node line directly:
      //
      BioFabricNetwork.NodeInfo tni = bfn_.getNodeDefinition(target);
      MinMax range = tni.getColRange(showShadows);
      if ((col >= range.min) && (col <= range.max)) { // Click within start and end of node line
        if (nodes.contains(target)) {
          nodes.remove(target); // If in, take out
        } else {
          if (nodeRange != null) {
            nodeRange.update(row); // Update node range, if we have it
          }
          nodes.add(target); // Add node to selection
          nodeAdd = true; // we have an addition
        } 
      } else {
        Point2D worldPt2 = rowColToWorld(new Point(col, row)); // Looking at the crossing of the node and the link line
        Rectangle2D nameLoc = nodeNameLocations_.get(target); 
        if (nameLoc.contains(worldPt2)) { // Does name location for target contain this world point??
          if (nodes.contains(target)) { // Same remove or add logic as above
            nodes.remove(target);
          } else {
            if (nodeRange != null) {
              nodeRange.update(row);
            }
            nodes.add(target);
            nodeAdd = true;
          }
        }
      }
    }
  
    BioFabricNetwork.LinkInfo linf = bfn_.getLinkDefinition(colObj, showShadows); // get link for the column
    if (linf != null) { // If we have a link in the column...
    	// if we are on a row where link starts or ends, we have an intersection. 
    	// WRONG! When both ends are in box, the link gets removed! 
      if ((rowObj.intValue() == linf.getStartRow()) || (rowObj.intValue() == linf.getEndRow())) {
      	// Usual add or remove ON COLUMNS
        boolean removeIt = false;
        if (cols.contains(colObj)) {
          cols.remove(colObj);
          removeIt = true;
        } else {
          if (colRange != null) {
            colRange.update(col);
          }
          cols.add(colObj);
        }
        // If told to remove, we remove the link. Otherwise, we add link, AND the node.
        NetNode src = bfn_.getSourceIDForColumn(colObj, showShadows); 
        NetNode trg = bfn_.getTargetIDForColumn(colObj, showShadows);
        if (removeIt) {
          links.remove(linf.getLink());
        } else {
          links.add(linf.getLink().clone());
          nodes.add(src);
          nodes.add(trg);
        } 
        gotLink = true;
      }
    }

    //
    // No link, but we do have a drain zone:
    //
    
    if (!gotLink && (gotDrain != null)) {
      if (nodes.contains(gotDrain)) {
        nodes.remove(gotDrain);
      } else {
        if (nodeRange != null) {
          int row2 = bfn_.getNodeDefinition(gotDrain).nodeRow;
          nodeRange.update(row2);
        }
        nodes.add(gotDrain);
        nodeAdd = true;
      }   
    }
 
    //
    // If shift is pressed, we are filling in the gaps:
    //
 
    if (shiftPressed) {
      if (linkAdd && (colRange.min != Integer.MAX_VALUE)) {
        for (int i = colRange.min; i < colRange.max; i++) {
          Integer colObj2 = Integer.valueOf(i);
          cols.add(colObj2);
          NetNode src = bfn_.getSourceIDForColumn(colObj2, showShadows); 
          NetNode trg = bfn_.getTargetIDForColumn(colObj2, showShadows);
          BioFabricNetwork.LinkInfo linf2 = bfn_.getLinkDefinition(colObj2, showShadows);
          links.add(linf2.getLink().clone());
          nodes.add(src);
          nodes.add(trg);
        }
      }
     if (nodeAdd && (nodeRange.min != Integer.MAX_VALUE)) {
        for (int i = nodeRange.min; i < nodeRange.max; i++) {
          Integer rowObj2 = Integer.valueOf(i);
          NetNode target2 = bfn_.getNodeIDForRow(rowObj2);
          nodes.add(target2);
        }
      }
    }
    return;
  }
  
  /***************************************************************************
  **
  ** Run the selection logic
  */  
  
  public void selectionLogicRect(Rectangle rect, boolean showShadows, Set<NetNode> nodes, Set<FabricLink> links, 
                                 Set<Integer> cols) { 


    if ((nodeNameLocations_ == null) || (drainNameLocations_ == null)) {
      return;
    }
 
    //
    // Figure out the bounds
    //

    int startRow = rect.y ;
    int endRow =  rect.y + rect.height;
    int startCol =  rect.x;
    int endCol = rect.x + rect.width;

    UiUtil.fixMePrintout("READ ME FOR NEW POLICY");
    // Rectangle selection should pull in links with glyphs within the rectangle, along with nodes they
    // are incident upon. They should also always be additive.
    // Should be able to select a link (which includes endpoints) by clicking on it, if there is no
    // node that beats it (ambiguity favors the node). 
    
    //
    // Given a rectangle, we iterate through the columns, get the links, and see if the start
    // or end of the links are within the box. If yes, we select the link, as well as the nodes
    // they are incident on:
    //
    
    //Integer rowObj = Integer.valueOf(row);
   // 	NetNode target = bfn_.getNodeIDForRow(rowObj);
        	
    for (int col = startCol; col <= endCol; col++) {               
      Integer colObj = Integer.valueOf(col);
      BioFabricNetwork.LinkInfo linf = bfn_.getLinkDefinition(colObj, showShadows); // get link for the column
      if (linf != null) { // If we have a link in the column...
      	int lstart = linf.getStartRow();
      	int lend = linf.getEndRow();
      	boolean startOK = (lstart >= startRow) && (lstart <= endRow);
        boolean endOK = (lend >= startRow) && (lend <= endRow);
        if (startOK || endOK) {
          cols.add(colObj);
          NetNode src = bfn_.getSourceIDForColumn(colObj, showShadows); 
          NetNode trg = bfn_.getTargetIDForColumn(colObj, showShadows);
     
          links.add(linf.getLink().clone());
          nodes.add(src);
          nodes.add(trg);
        }
      }
    }
    return;
  }

  /***************************************************************************
  **
  ** Build needed selection geometry
  */  
  
  public void buildSelectionGeometry(NetNode newStartName, Rectangle newStartRect) {     
    Point focus = new Point();
    boolean showShadows = FabricDisplayOptionsManager.getMgr().getDisplayOptions().getDisplayShadows(); 
    //
    // Create target focus boxes, sorted by row:
    //
    targetList_.clear();
    TreeMap<Integer, Rectangle> sortTargs = new TreeMap<Integer, Rectangle>();
    Iterator<NetNode> tgit = currNodeSelections_.iterator();
    while (tgit.hasNext()) {
      NetNode target = tgit.next();
      BioFabricNetwork.NodeInfo targetInf = bfn_.getNodeDefinition(target);
      targetList_.add(targetInf);
      Rectangle2D targName = nodeNameLocations_.get(target);
      Point targNameRC = worldToRowCol(new Point2D.Double(targName.getCenterX(), targName.getCenterY()));
      focus.setLocation(targNameRC.x, targetInf.nodeRow);
      sortTargs.put(Integer.valueOf(targetInf.nodeRow), buildFocusBox(focus));  
    }

    //
    // Create link focus boxes:
    //
    
    HashMap<Integer, SortedMap<Integer, Rectangle>> linksByRow = new HashMap<Integer, SortedMap<Integer, Rectangle>>();
    linkList_.clear();
    Iterator<Integer> cit = currColSelections_.iterator();
    while (cit.hasNext()) {
      Integer colObj = cit.next();
      BioFabricNetwork.LinkInfo linf = bfn_.getLinkDefinition(colObj, showShadows);
      linkList_.add(linf);
      Integer strtObj = Integer.valueOf(linf.getStartRow());
      SortedMap<Integer, Rectangle> lbr = linksByRow.get(strtObj);
      if (lbr == null) {
        lbr = new TreeMap<Integer, Rectangle>();
        linksByRow.put(strtObj, lbr);
      }
      focus.setLocation(linf.getUseColumn(showShadows), linf.getStartRow());
      lbr.put(colObj, buildFocusBox(focus));
      
      Integer endObj = Integer.valueOf(linf.getEndRow());
      lbr = linksByRow.get(endObj);
      if (lbr == null) {
        lbr = new TreeMap<Integer, Rectangle>();
        linksByRow.put(endObj, lbr);
      }
      focus.setLocation(linf.getUseColumn(showShadows), linf.getEndRow());
      lbr.put(colObj, buildFocusBox(focus));
    }
   
    // FIX ME: seeing currSel = 0 on empty rects!
    Rectangle currRect = ((currSel_ == -1) || (currSel_ >= rects_.size())) ? null : (Rectangle)rects_.get(currSel_);

    rects_.clear();
    Iterator<Integer> stit = sortTargs.keySet().iterator();
    while (stit.hasNext()) {
      Integer row = stit.next();
      Rectangle trgRect = sortTargs.get(row);
      if (trgRect.equals(currRect)) {
        currSel_ = rects_.size();
      } else if ((newStartName != null) && newStartName.equals(bfn_.getNodeIDForRow(row))) {
        currSel_ = rects_.size();
      }
      rects_.add(trgRect);
      SortedMap<Integer, Rectangle> ufr = linksByRow.get(row);
      if (ufr == null) {
        continue;
      }
      rects_.addAll(ufr.values());
    }
    
    //
    // New selections all built.  Set the current selection
    //
    
    if (newStartName == null) {
      if (newStartRect != null) {
        resetCurrentSelection(newStartRect);
      } else if (currRect != null) {
        resetCurrentSelection(currRect);
      }
    }
    if ((rects_.size() > 0) && (currSel_ == -1)) {
      currSel_ = 0;
    }
    bumpGuts();
    handleFloaterChange();
    UiUtil.fixMePrintout("This has gotta change");
    HashSet<Integer> targRows = new HashSet<Integer>();
    HashSet<Integer> targCols = new HashSet<Integer>(currColSelections_);
    HashSet<NID> targIDs = new HashSet<NID>();
    
    int numTarg = targetList_.size();
    for (int i = 0; i < numTarg; i++) {
    	BioFabricNetwork.NodeInfo targetInf = targetList_.get(i);
    	targRows.add(Integer.valueOf(targetInf.nodeRow));
    	targIDs.add(targetInf.getNodeID());
    }
    selections_ = new PaintCacheSmall.Reduction(targRows, targCols, targIDs); 
    fmt_.setSelections(selections_);
    EventManager mgr = EventManager.getManager();
    SelectionChangeEvent ev = new SelectionChangeEvent(null, null, SelectionChangeEvent.SELECTED_ELEMENT);
   
    mgr.sendSelectionChangeEvent(ev);  
    return;
  }
  
  /***************************************************************************
  **
  ** reset the current selection
  */  
  
  private void resetCurrentSelection(Rectangle newStart) {
    int numRect = rects_.size();
    for (int i = 0 ; i < numRect; i++) {
      Rectangle trgRect = rects_.get(i);
      if (trgRect.equals(newStart)) {
        currSel_ = i;
      }
    }
    return;
  }
  
  /***************************************************************************
  **
  ** Support image export
  */  
  
  public void exportToFile(File saveFile, String format, ImageExporter.ResolutionSettings res, double zoom, Dimension size) throws IOException {    
    exportGuts(saveFile, format, res, zoom, size);
    return;
  }  
  
  /***************************************************************************
  **
  ** Support image export
  */  
  
  public void exportToStream(OutputStream stream, String format, ImageExporter.ResolutionSettings res, double zoom, Dimension size) throws IOException {    
    exportGuts(stream, format, res, zoom, size);
    return;
  }   
  
  /***************************************************************************
  **
  ** Support image export.  Because of the way the image handler operates, it can take
  ** either an OutputStream or a File as the first argument:
  */  
  
  private void exportGuts(Object outObj, String format, ImageExporter.ResolutionSettings res, double zoom, Dimension size) throws IOException { 
    
    Rectangle worldPiece = UiUtil.rectFromRect2D(worldRectNetAR_);
    Dimension useSize = (size == null) ? new Dimension((int)(worldPiece.width * zoom), (int)(worldPiece.height * zoom)) : size;
           
    BufferedImage bi = new BufferedImage(useSize.width, useSize.height, BufferedImage.TYPE_INT_RGB);
    drawForBuffer(bi, worldPiece, useSize, worldPiece, 0, 0.0);  
    ImageExporter iex = new ImageExporter();
    iex.export(outObj, bi, format, res);    
    return;
  }
 
  /***************************************************************************
  **
  ** Get tour going from current selection
  */  
  
  public TourStatus startTourFromSelection(boolean selectionOnly) {
    if ((currSel_ == -1) || (floaterSet_.currSelRect == null)) {
      throw new IllegalStateException();
    }
    floaterSet_.tourRect = (Rectangle)floaterSet_.currSelRect.clone();  // world coords
    tourFocus_ = worldToRowCol(new Point2D.Double(floaterSet_.tourRect.getCenterX(), floaterSet_.tourRect.getCenterY()));
    centerOnRectangle(floaterSet_.tourRect);
    fmt_.setCenter(rowColToWorld(tourFocus_), tourFocus_, true);
    MouseLocInfo vals = buildMouseLocation(tourFocus_);
    handleFloaterChange();  // else tour rect does not redraw
    boolean showShadows = FabricDisplayOptionsManager.getMgr().getDisplayOptions().getDisplayShadows(); 
    Integer tfx = Integer.valueOf(tourFocus_.x);
    BioFabricNetwork.LinkInfo ld = bfn_.getLinkDefinition(tfx, showShadows);
    if ((ld != null) && !ld.inLinkRowRange(tourFocus_.y)) {
      ld = null;
    }
    NetNode nodeForRow = bfn_.getNodeIDForRow(Integer.valueOf(tourFocus_.y));
    SortedSet<Integer> okStops = (selectionOnly) ? null : findSelectedLinkStops(nodeForRow);
    boolean nodeAlive = (!selectionOnly) ? true : currNodeSelections_.contains(nodeForRow);
    return (new TourStatus(vals, bfn_, ld, tourFocus_, okStops, nodeAlive, false));    
  }
  
  /***************************************************************************
  ** 
  ** For finding nearest tour start within limits
  */

  private Point findClosestTourStart(Point prtc, int limit) {

    int start = prtc.x - limit;
    int end = prtc.x + limit;
    int minY = prtc.y - limit;
    int maxY = prtc.y + limit;
    
    boolean showShadows = FabricDisplayOptionsManager.getMgr().getDisplayOptions().getDisplayShadows(); 
    Point2D forVec = new Point2D.Double(prtc.x, prtc.y);
    double minDist = Double.POSITIVE_INFINITY;
    Point closestRC = new Point();
    for (int i = start; i <= end; i++) {
      Integer testCol = Integer.valueOf(i);
      BioFabricNetwork.LinkInfo linf = bfn_.getLinkDefinition(testCol, showShadows);
      if (linf != null) {     
        if ((linf.getStartRow() >= minY) && (linf.getStartRow() <= maxY)) {
          Point2D testPt = new Point2D.Double(i, linf.getStartRow());
          double dSq = forVec.distanceSq(testPt);
          if (dSq < minDist) {
            minDist = dSq;
            closestRC.setLocation(i, linf.getStartRow());
          }
        }
        if ((linf.getEndRow() >= minY) && (linf.getEndRow() <= maxY)) {
          Point2D testPt = new Point2D.Double(i, linf.getEndRow());
          double dSq = forVec.distanceSq(testPt);
          if (dSq < minDist) {
            minDist = dSq;
            closestRC.setLocation(i, linf.getEndRow());
          }
        }
      }
    }
  
    //
    // Now try names:
    //
    
    Point2D inWorldL = rowColToWorld(new Point(prtc.x - limit, prtc.y));
    Point2D inWorldR = rowColToWorld(new Point(prtc.x + limit, prtc.y));
    for (int i = minY; i <= maxY; i++) {
      Integer testRow = Integer.valueOf(i);
      NetNode nodeName = bfn_.getNodeIDForRow(testRow);
      if (nodeName != null) {
        Rectangle2D nnl = nodeNameLocations_.get(nodeName);
        Point2D nameCenter = new Point2D.Double(nnl.getCenterX(), nnl.getCenterY());
        if ((nameCenter.getX() >= inWorldL.getX()) && (nameCenter.getX() <= inWorldR.getX())) {
          Point2D ncRC = worldToRowCol(nameCenter);
          // Closest ROW is the one that really counts!
          double dSq = forVec.distanceSq(new Point2D.Double(forVec.getX(), ncRC.getY()));
          if (dSq < minDist) {
            minDist = dSq;
            closestRC.setLocation((int)ncRC.getX(), i);
          }
        }       
      }
    }

    return ((minDist < Double.POSITIVE_INFINITY) ? closestRC : null);
  }  
 
  /***************************************************************************
  **
  ** Draw Object
  */  
  
  private static class ImageToUse {
    BufferedImage image;
    int stX;
    int stY;
    
    ImageToUse(BufferedImage image, int stX, int stY) {
      this.image = image;
      this.stX = stX;
      this.stY = stY;
    }
  }  
  
  /***************************************************************************
  **
  ** Tour status
  */  
  
  public static class TourStatus implements Cloneable {    
     public String nodeName;
     public String linkName;
     boolean upEnabled;
     boolean downEnabled;
     boolean leftEnabled;
     boolean rightEnabled;
     boolean farLeftEnabled;
     boolean farRightEnabled;
     boolean currStopUnselected;
     
     TourStatus(MouseLocInfo vals, BioFabricNetwork bfn, BioFabricNetwork.LinkInfo link, 
                Point navFocus, SortedSet<Integer> selectedOnly, boolean nodeAlive, boolean stopUnselected) {
       nodeName = vals.nodeDesc;
       linkName = vals.linkDesc;
       boolean showShadows = FabricDisplayOptionsManager.getMgr().getDisplayOptions().getDisplayShadows(); 
       currStopUnselected = stopUnselected;
       // Null link means we are sitting on a node label
       if (link == null) {
         upEnabled = false;
         downEnabled = false;
         leftEnabled = false;
         farLeftEnabled = false;
         rightEnabled = true;
         farRightEnabled = true;         
       } else {
         BioFabricNetwork.NodeInfo src = bfn.getNodeDefinition(link.getSource());
         BioFabricNetwork.NodeInfo trg = bfn.getNodeDefinition(link.getTarget());
         BioFabricNetwork.NodeInfo useNode = (navFocus.y == src.nodeRow) ? src : trg;
         if ((selectedOnly != null) && selectedOnly.isEmpty()) {
           leftEnabled = false;
           farLeftEnabled = nodeAlive;
           rightEnabled = false;
           farRightEnabled = false;           
         } else {
         //  int minCol = (selectedOnly == null) ? useNode.minCol : ((Integer)selectedOnly.first()).intValue();
           MinMax range = useNode.getColRange(showShadows);
           int maxCol = (selectedOnly == null) ? range.max : selectedOnly.last().intValue();
           upEnabled = (navFocus.y != link.topRow());
           downEnabled = (navFocus.y != link.bottomRow());
           leftEnabled = (navFocus.x >= range.min) && nodeAlive;
           farLeftEnabled = (navFocus.x >= range.min) && nodeAlive;
           rightEnabled = (maxCol != navFocus.x);
           farRightEnabled = (maxCol != navFocus.x);
         }
       }
     }
     
     public TourStatus clone() {
      try {
        return ((TourStatus)super.clone());
      } catch (CloneNotSupportedException cnse) {
        throw new IllegalStateException();
      }
    }    
  }
    
  /***************************************************************************
  **
  ** Handles click events
  */  
  
  public class MouseHandler extends MouseAdapter { 
    
    private final static int CLICK_SLOP_  = 2;
    
    @Override
    public void mouseClicked(MouseEvent me) {
      if (me.isPopupTrigger()) {
        Point pscreenLoc = me.getComponent().getLocationOnScreen();
        Point abs = new Point(me.getX() + pscreenLoc.x, me.getY() + pscreenLoc.y);
        triggerPopup(me.getX(), me.getY(), abs);
      }
      return;
    }  

    private void handleClick(int lastX, int lastY, boolean shiftPressed) {    
      Point loc = new Point(lastX, lastY);
      Point rcp = transToRowCol(loc);
      handleSelection(rcp, null, loc, true, shiftPressed);
      return;
    }
     
    private void handleSelection(Point rcbp, Rectangle rect, Point sloc, boolean onePt, boolean shiftPressed) {
      if (bfn_ == null) {
        return;
      }
      boolean showShadows = FabricDisplayOptionsManager.getMgr().getDisplayOptions().getDisplayShadows(); 
      if (collectingTourStart_) {
        boolean tstatUS = false;
        boolean forSelOnly = tourStartSelectionOnly_;
        if (rcbp != null) {
          Point newFocus = findClosestTourStart(rcbp, 3);
          if (newFocus == null) {
            cursorMgr_.signalError();
            return;
          }
          if (tourStopIsUnselected(newFocus)) {
            tstatUS = true;
            forSelOnly = false;
            fnt_.resetSkipSelections();
          }          
          tourFocus_ = newFocus;
          floaterSet_.tourRect = buildFocusBox(tourFocus_);
          MouseLocInfo loc = buildMouseLocation(tourFocus_);
          BioFabricNetwork.LinkInfo linf = bfn_.getLinkDefinition(Integer.valueOf(tourFocus_.x), showShadows);
          if ((linf != null) && !linf.inLinkRowRange(tourFocus_.y)) {
            linf = null;
          }
          NetNode nodeForRow = bfn_.getNodeIDForRow(Integer.valueOf(tourFocus_.y));
          SortedSet<Integer> okStops = (forSelOnly) ? findSelectedLinkStops(nodeForRow) : null;
          boolean nodeAlive = (!forSelOnly) ? true : currNodeSelections_.contains(nodeForRow);
          fnt_.installNames(new TourStatus(loc, bfn_, linf, tourFocus_, okStops, nodeAlive, tstatUS));
        }
        collectingTourStart_ = false;
        tourStartSelectionOnly_ = false;
        cursorMgr_.showDefaultCursor();
        bfw_.reenableControls();
        return;
      }
      if (collectingZoomMode_) {
        if (firstZoomPoint_ == null) {
          firstZoomPoint_ = (Point)sloc.clone();
          return;
        } else {
          Rectangle rcRect = valsToRect(firstZoomPoint_.x, firstZoomPoint_.y, 
                                        sloc.x, sloc.y, false);
          if (rcRect == null) {
            return;
          }
          Point2D lpu = viewToWorld(new Point(rcRect.x, rcRect.y));
          Point2D rpl = viewToWorld(new Point((int)rcRect.getMaxX(), (int)rcRect.getMaxY()));
          Rectangle zoomTo = new Rectangle((int)lpu.getX(), (int)lpu.getY(), 
                                           (int)(rpl.getX() - lpu.getX()), (int)(rpl.getY() - lpu.getY()));
          if ((zoomTo.width > 0) && (zoomTo.height > 0)) {
            zoomToRectangle(zoomTo);
          }
          collectingZoomMode_ = false;
          firstZoomPoint_ = null;
          cursorMgr_.showDefaultCursor();
          bfw_.reenableControls();
          return;
        }
      }
          
      if (!doBuildSelect_) {
        currLinkSelections_.clear();
        currNodeSelections_.clear();
        currColSelections_.clear();
        fmt_.setSelections(null);
        currSel_ = -1;
        floaterSet_.currSelRect = null;
      }
      
      //
      // Clicking on one guy will make him the current selection!
      //
      Rectangle newStart;
      if (onePt) {
        newStart = buildFocusBox(rcbp);
        selectionLogicPoint(rcbp, sloc, showShadows, currNodeSelections_, currLinkSelections_, currColSelections_, shiftPressed); 
      } else {
      	newStart = null;
      	selectionLogicRect(rect, showShadows, currNodeSelections_, currLinkSelections_, currColSelections_); 
      }
      
      buildSelectionGeometry(null, newStart);  
      return;
    }  
    
    private void dragResult(int sx, int sy, int ex, int ey, boolean isCtrl) {
      if (!isCtrl) {  // Keep floater alive! (change to ! for V2)
        return;
      }
      if (collectingZoomMode_) {
        collectingZoomMode_ = false;
        cursorMgr_.showDefaultCursor();
        bfw_.reenableControls();
        return;
      } else if (collectingTourStart_) {
        collectingTourStart_ = false;
        tourStartSelectionOnly_ = false;
        cursorMgr_.showDefaultCursor();
        bfw_.reenableControls();
        return;
      } else {
        Rectangle rcRect = valsToRect(sx, sy, ex, ey, true);
        if (rcRect != null) {
          handleSelection(null, rcRect, null, false, false);
        }
      }
    }  
    
    @Override
    public void mousePressed(MouseEvent me) {
      try {
        if (me.isPopupTrigger()) {
          Point pscreenLoc = me.getComponent().getLocationOnScreen();
          Point abs = new Point(me.getX() + pscreenLoc.x, me.getY() + pscreenLoc.y);
          triggerPopup(me.getX(), me.getY(), abs);    
        } else {
          lastPress_ = new Point(me.getX(), me.getY());
          lastView_ = jsp_.getViewport().getViewPosition(); 
          Point screenLoc = me.getComponent().getLocationOnScreen();
          lastAbs_ = new Point(me.getX() + screenLoc.x, me.getY() + screenLoc.y);         
          lastShifted_ = me.isShiftDown();        
          lastCtrl_ = me.isControlDown() || (isAMac_ && me.isMetaDown());
        }
      } catch (Exception ex) {
        ExceptionHandler.getHandler().displayException(ex);
      }
      return;
    }   

    @Override
    public void mouseEntered(MouseEvent me) {
      try {
        bfo_.setMouseIn(true, fmt_.isIgnoring());
        fmt_.setMouseIn(true);
      } catch (Exception ex) {
        ExceptionHandler.getHandler().displayException(ex);
      }        
      return;
    }      
    
    @Override
    public void mouseExited(MouseEvent me) {
      try {
        bfo_.setMouseIn(false, fmt_.isIgnoring());  
        fmt_.setMouseIn(false);
        myLocation_.setNodeAndLink(new MouseLocInfo());
        mov_.showForNode(new MouseLocInfo());
      } catch (Exception ex) {
        ExceptionHandler.getHandler().displayException(ex);
      }        
      return;
    }      
    
    @Override
    public void mouseReleased(MouseEvent me) {
      try {
      	
      	//
      	// Now that we use a simple drag to scroll, we need to know if the user
      	// has done that (else it just looks like a click: no world coordinate change
      	// between press and release:
      	//
      	
      	Point screenLoc = me.getComponent().getLocationOnScreen();
        Point newAbs = new Point(me.getX() + screenLoc.x, me.getY() + screenLoc.y);
        boolean absChange = (lastAbs_ != null) && !newAbs.equals(lastAbs_);
  
        // Do this stuff NO MATTER WHAT!

        lastView_ = null;
        lastAbs_ = null;

        if (me.isPopupTrigger()) {
          Point pscreenLoc = me.getComponent().getLocationOnScreen();
          Point abs = new Point(me.getX() + pscreenLoc.x, me.getY() + pscreenLoc.y);
          triggerPopup(me.getX(), me.getY(), abs);
          return;
        }
        
        //
        // V2: Control is now used for boxing select, so this needs to go:
        //
        
        // if (me.isControlDown() || (isAMac_ && me.isMetaDown())) {
        //   lastPress_ = null;
        //  return;
        // }
        
        boolean shiftPressed = me.isShiftDown();
        int currX = me.getX();
        int currY = me.getY();

        if (lastPress_ == null) {
          return;
        }
        
        int lastX = lastPress_.x;
        int lastY = lastPress_.y;
        int diffX = Math.abs(currX - lastX);
        int diffY = Math.abs(currY - lastY); 
        
        // Note that a ctrl-drag leaves us with diff == 0, thus
        // it would select after the ctrl drag!
        
        if (!absChange && (diffX <= CLICK_SLOP_) && (diffY <= CLICK_SLOP_)) { 
          handleClick(lastX, lastY, shiftPressed);
        } else if (lastCtrl_ && ((diffX >= CLICK_SLOP_) || (diffY >= CLICK_SLOP_))) {
          dragResult(lastX, lastY, currX, currY, lastCtrl_);
        }

        // Pre-V2 code (ctrl pressed was drag)
        //if (!lastCtrl_ && (diffX <= CLICK_SLOP_) && (diffY <= CLICK_SLOP_)) { 
        //  handleClick(lastX, lastY, shiftPressed);
        //} else if ((diffX >= CLICK_SLOP_) || (diffY >= CLICK_SLOP_)) {
        //  dragResult(lastX, lastY, currX, currY, lastCtrl_);
        //}
        floaterSet_.floater = null;
        lastPress_ = null;  // DO THIS NO MATTER WHAT TOO       
        handleFloaterChange();
      } catch (Exception ex) {
        ExceptionHandler.getHandler().displayException(ex);
      }
         
      return;
    }
    
    private void triggerPopup(int x, int y, Point screenAbs) {
      try {
        Point loc = new Point(x, y);
        Point rcp = transToRowCol(loc);
        HashSet<NetNode> nodes = new HashSet<NetNode>();
        HashSet<FabricLink> links = new HashSet<FabricLink>();
        HashSet<Integer> cols = new HashSet<Integer>();
        boolean showShadows = FabricDisplayOptionsManager.getMgr().getDisplayOptions().getDisplayShadows(); 
        selectionLogicPoint(rcp, loc, showShadows, nodes, links, cols, false); 
        if (!links.isEmpty()) {
          FabricLink fabLink = links.iterator().next();
          popCtrl_.showLinkPopup(fabLink, loc); 
        } else if (!nodes.isEmpty()) {
          NetNode nodeName = nodes.iterator().next();
          popCtrl_.showNodePopup(nodeName, loc); 
        }  
      } catch (Exception ex) {
        ExceptionHandler.getHandler().displayException(ex);
      }
      return;
    }
  }
   
  /***************************************************************************
  **
  ** Handles mouse motion events
  */  
      
  public class MouseMotionHandler extends MouseMotionAdapter {
    
    @Override
    public void mouseDragged(MouseEvent me) {
      try {
        if (lastPress_ == null) {
          return;
        }
        Point currPt = me.getPoint();
        if (me.isControlDown() || (isAMac_ && me.isMetaDown())) {
        	Point2D lpw = viewToWorld(lastPress_);
          Point2D cupw = viewToWorld(currPt);
          floaterSet_.floater = valsToRect((int)lpw.getX(), (int)lpw.getY(), (int)cupw.getX(), (int)cupw.getY(), false);
          handleFloaterChange();
        } else if (collectingZoomMode_) {
          floaterSet_.floater = null;
          handleFloaterChange();        
        } else {
          Point compLoc = me.getComponent().getLocationOnScreen();
          Point currAbs = new Point(compLoc.x + currPt.x, compLoc.y + currPt.y); 

          JScrollBar hsb = jsp_.getHorizontalScrollBar();
          int hMax = hsb.getMaximum() - hsb.getVisibleAmount();
          int hMin = hsb.getMinimum();
          int newX = lastView_.x - (currAbs.x - lastAbs_.x);
          if (newX > hMax) newX = hMax;
          if (newX < hMin) newX = hMin;

          JScrollBar vsb = jsp_.getVerticalScrollBar();
          int vMax = vsb.getMaximum() - vsb.getVisibleAmount();
          int vMin = vsb.getMinimum();
          int newY = lastView_.y - (currAbs.y - lastAbs_.y);
          if (newY > vMax) newY = vMax;
          if (newY < vMin) newY = vMin;

          jsp_.getViewport().setViewPosition(new Point(newX, newY));
          jsp_.getViewport().invalidate(); 
          jsp_.revalidate();
        }
      } catch (Exception ex) {
        ExceptionHandler.getHandler().displayException(ex);
      }
    }
    
    @Override
    public void mouseMoved(MouseEvent me) {
      try {
        Point currPt = me.getPoint();
        Point2D cpw = viewToWorld(currPt);
        Point cprc = worldToRowCol(cpw);
        fmt_.setCenter(cpw, cprc, false);  
        bfo_.setMouse(cpw, cprc);
        if (bfn_ == null) {
          return;
        }
        MouseLocInfo vals = buildMouseLocation(cprc);
        myLocation_.setNodeAndLink(vals);
        mov_.showForNode(vals);
        if (collectingZoomMode_) {
          if (firstZoomPoint_ != null) {
            Point2D lpw = viewToWorld(firstZoomPoint_);
            Point2D cupw = viewToWorld(currPt);
            floaterSet_.floater = valsToRect((int)lpw.getX(), (int)lpw.getY(), (int)cupw.getX(), (int)cupw.getY(), false);
            handleFloaterChange();
          }
        }      
      } catch (Exception ex) {
        ExceptionHandler.getHandler().displayException(ex);
      }
    }         
  } 
  
  /***************************************************************************
  **
  ** Now the actual panel to use
  */  
      
  public class MyPaintPanel extends JPanel {
    
    private static final long serialVersionUID = 1L;
    
    /***************************************************************************
    **
    ** preferred size
    */
    
    @Override
    public Dimension getPreferredSize() {
      return (zoomer_.getPreferredSize());    
    }

    /***************************************************************************
    **
    ** Drawing routine for printing
    */

    @Override
    public void print(Graphics g) {
      if (bfn_ == null) {
        return;
      }  
      Rectangle2D viewInWorld = getViewInWorld();
      Graphics2D g2 = (Graphics2D)g;
      drawingGuts(g2, viewInWorld);
      return;
    }

    @Override
    public void paintComponent(Graphics g) {
      super.paintComponent(g); 
       
      //
      // Make sure to return any stale images we have lying around:
      //
       
      Iterator<BufferedImage> siit = staleImages_.iterator();
      while (siit.hasNext()) {
        BufferedImage bi = siit.next();
        bis_.returnImage(bi);
      }
      staleImages_.clear();
         
      if (bfn_ == null) {
        return;
      }
       
      Double zoomVal = Double.valueOf(zoomer_.getZoomFactor());
      Rectangle2D viewInWorld = getViewInWorld();
       
      //
      // When we zoom in far enough, we start to draw it instead. If our zoom is too wide, we need to use the
      // full-screen version and scale it.
      //
       
      Integer numObj = zoomMap_.get(zoomVal);
      boolean wideCase = false;
       
      // Submodel view does not have a buffer builder, thus second test here:
      if ((zoomVal.doubleValue() < zoomMap_.firstKey().doubleValue()) && (bufferBuilder_ != null)) {
        numObj = Integer.valueOf(0);
        wideCase = true;    
      } else {
        if ((numObj == null) || (bufferBuilder_ == null)) {
          Graphics2D g2 = (Graphics2D)g;
          drawingGuts(g2, viewInWorld);
          return;
        }
      }
      ArrayList<ImageToUse> imagesToUse = new ArrayList<ImageToUse>();
          
      //
      // Image boundary problems will be reduced if we just do the world to viewport transform
      // for the upper right and then just add the image tile size to that point?
      // 
      UiUtil.fixMePrintout("implement the above??");

      ArrayList<Rectangle2D> slicesToCover = new ArrayList<Rectangle2D>();
      bufferBuilder_.getSlicesToCover(numObj, viewInWorld, slicesToCover);
      int numSlice = slicesToCover.size();
      AffineTransform wideTrans = null;
      for (int i = 0; i < numSlice; i++) {
        Rectangle2D worldRect = slicesToCover.get(i);
        BufferedImage img = null;
        try {
          img = bufferBuilder_.getImageForPiece(numObj.intValue(), worldRect);
        } catch (IOException ioex) {
          System.err.println("Bad load");
        }
        if (img != null) {
          Point wtv = pointToViewport(new Point((int)Math.round(worldRect.getX()), (int)Math.round(worldRect.getY())));
          int stX = wtv.x;
          int stY = wtv.y;
           
          //
          // "Wide" Case:
          // We want to fill the current viewport with the model, using the full-screen baseline image. This typically means
          // shrinking the image. Note that when the nav panel is shown, a wide and thin network may not need any shrinking, 
          // but may need the top and bottom empty space chopped off. A tall and square network will need to be shrunk down
          // so the vertical dimension fits. 
          //
             
          if (wideCase) {
            wideTrans = new AffineTransform();
           
            // When last zoom selected was from fixed set, window resize DOES NOT CHANGE ZOOM. If last zoom
            // selected was whole model, the ops below are dynamically resizing the model to keep fitting into
            // viewport (because the fixed set zoom choice remains null) This is not consistent.
             
            //
            // This is the zoom level at which the baseline image is rendered:
            //
             
            double fkz = zoomMap_.firstKey().doubleValue();
             
            //
            // This is the zoom level that we need for the model to be displayed:
            //
             
            double zf = zoomer_.getZoomFactor();
            double izoom = zf / fkz;
             
    
            JViewport view = jsp_.getViewport();
            Dimension viewDim = view.getViewSize();       
            wideTrans.translate(viewDim.getWidth() / 2.0, viewDim.getHeight() / 2.0);
            wideTrans.scale(izoom, izoom);
            UiUtil.fixMePrintout("Scaling image " + izoom);
            wideTrans.translate(-img.getWidth() / 2.0, -img.getHeight() / 2.0);
            stX = 0;
            stY = 0;
          }

          ImageToUse itu = new ImageToUse(img, stX, stY);
          imagesToUse.add(itu);
        }
      }
      if (imagesToUse.isEmpty()) {
        return;
      }
     
      Graphics2D g2p = (Graphics2D)g;
      Iterator<ImageToUse> ituit = imagesToUse.iterator();
      while (ituit.hasNext()) {
        ImageToUse it = ituit.next();
        if (it.image != null) {
          int useX = it.stX;
          int useY = it.stY;
          AffineTransform stash = null;
          if (wideTrans != null) {
            stash = g2p.getTransform();
            g2p.transform(wideTrans);
          }
          g2p.drawImage(it.image, useX, useY, null);
          if (stash != null) {
            g2p.setTransform(stash); 
            staleImages_.add(it.image);
          }       
        }
      }
       
      clipRect_.setBounds((int)viewInWorld.getX(), (int)viewInWorld.getY(),
                          (int)viewInWorld.getWidth(), (int)viewInWorld.getHeight());
      if (!targetList_.isEmpty() || !linkList_.isEmpty()) {
        drawSelections(g2p, clipRect_);
      }
      drawFloater(g2p, true);
      g.dispose();
      return;
    }
  }
}
```
