## Supplementary material for "BioFabric Visualization of Network Alignments": BioFabric Source Code: FabricLocation.java

```
/*
**    Copyright (C) 2003-2018 Institute for Systems Biology 
**                            Seattle, Washington, USA. 
**
**    This library is free software; you can redistribute it and/or
**    modify it under the terms of the GNU Lesser General Public
**    License as published by the Free Software Foundation; either
**    version 2.1 of the License, or (at your option) any later version.
**
**    This library is distributed in the hope that it will be useful,
**    but WITHOUT ANY WARRANTY; without even the implied warranty of
**    MERCHANTABILITY or FITNESS FOR A PARTICULAR PURPOSE.  See the GNU
**    Lesser General Public License for more details.
**
**    You should have received a copy of the GNU Lesser General Public
**    License along with this library; if not, write to the Free Software
**    Foundation, Inc., 59 Temple Place, Suite 330, Boston, MA  02111-1307  USA
*/

package org.systemsbiology.biofabric.ui.display;

import java.awt.Color;
import java.awt.GridLayout;
import java.util.Iterator;
import java.util.List;

import javax.swing.JPanel;


/****************************************************************************
**
** This is location announcement
*/

public class FabricLocation extends JPanel {
    
  ////////////////////////////////////////////////////////////////////////////
  //
  // PRIVATE CONSTANTS
  //
  //////////////////////////////////////////////////////////////////////////// 
	
  private static final long serialVersionUID = 1L;
  
  ////////////////////////////////////////////////////////////////////////////
  //
  // PUBLIC CONSTANTS
  //
  //////////////////////////////////////////////////////////////////////////// 
  
  ////////////////////////////////////////////////////////////////////////////
  //
  // PRIVATE INSTANCE MEMBERS
  //
  ////////////////////////////////////////////////////////////////////////////

  private InfoPanel nodePanel_;
  private InfoPanel linkPanel_;
  private InfoPanel linkZonePanel_;
  private InfoPanel nodeAnnotPanel_;
  private InfoPanel linkAnnotPanel_;
  private StringBuffer scratch_;
  
  ////////////////////////////////////////////////////////////////////////////
  //
  // PUBLIC CONSTRUCTORS
  //
  ////////////////////////////////////////////////////////////////////////////

  /***************************************************************************
  **
  ** Constructor
  */

  public FabricLocation() {
    setBackground(Color.WHITE);
  	setLayout(new GridLayout(2, 1));
  	JPanel topPanel = new JPanel();
  	JPanel botPanel = new JPanel();
  	topPanel.setLayout(new GridLayout(1, 2));
  	topPanel.setBackground(Color.WHITE);
  	botPanel.setLayout(new GridLayout(1, 3));
  	botPanel.setBackground(Color.WHITE);
  	add(topPanel);
  	add(botPanel);
  	
    nodePanel_ = new InfoPanel(false, 15, 150, false, true);  
    linkPanel_ = new InfoPanel(false, 15, 150, false, true);
    linkZonePanel_ = new InfoPanel(false, 15, 150, false, true);
    
    nodeAnnotPanel_ = new InfoPanel(false, 15, 300, false, true);  
    linkAnnotPanel_ = new InfoPanel(false, 15, 300, false, true);
    
    nodePanel_.installName("Mouse Over Node Row: <none>");
    linkPanel_.installName("Mouse Over Link: <none>");
    linkZonePanel_.installName("Mouse Over Node Link Zone: <none>");
    nodeAnnotPanel_.installName("Mouse Over Node Annotations: <none>");
    linkAnnotPanel_.installName("Mouse Over Link Annotations: <none>");

    topPanel.add(nodeAnnotPanel_);
    topPanel.add(linkAnnotPanel_);
     
    botPanel.add(nodePanel_);
    botPanel.add(linkPanel_);
    botPanel.add(linkZonePanel_);
    scratch_ = new StringBuffer();
  }

  ////////////////////////////////////////////////////////////////////////////
  //
  // PUBLIC METHODS
  //
  ////////////////////////////////////////////////////////////////////////////
  
  /***************************************************************************
  **
  ** Set the values
  */

  public void setNodeAndLink(BioFabricPanel.MouseLocInfo mlo) {
     nodePanel_.installName("Mouse Over Node Row: " + mlo.nodeDesc);
	   linkPanel_.installName("Mouse Over Link: " + mlo.linkDesc);
	   linkZonePanel_.installName("Mouse Over Node Link Zone: " + mlo.zoneDesc);
	   nodeAnnotPanel_.installName("Mouse Over Node Annotations: " + displayString(scratch_, mlo.nodeAnnotations));
	   linkAnnotPanel_.installName("Mouse Over Link Annotations: " + displayString(scratch_, mlo.linkAnnotations));
	   
     repaint();   
    return;
  } 
  
  /***************************************************************************
  **
  ** Glue names together
  */
  
  private String displayString(StringBuffer buf, List<String> vals) {
    buf.setLength(0);
    Iterator<String> fli = vals.iterator();
    while (fli.hasNext()) {
      buf.append(fli.next());
      if (fli.hasNext()) {
        buf.append(", ");
      }
    }
    return (buf.toString());
  }

}
```
