## Supplementary material for "BioFabric Visualization of Network Alignments": BioFabric Source Code: FabricMagnifier.java

```
/*
**    Copyright (C) 2003-2018 Institute for Systems Biology 
**                            Seattle, Washington, USA. 
**
**    This library is free software; you can redistribute it and/or
**    modify it under the terms of the GNU Lesser General Public
**    License as published by the Free Software Foundation; either
**    version 2.1 of the License, or (at your option) any later version.
**
**    This library is distributed in the hope that it will be useful,
**    but WITHOUT ANY WARRANTY; without even the implied warranty of
**    MERCHANTABILITY or FITNESS FOR A PARTICULAR PURPOSE.  See the GNU
**    Lesser General Public License for more details.
**
**    You should have received a copy of the GNU Lesser General Public
**    License along with this library; if not, write to the Free Software
**    Foundation, Inc., 59 Temple Place, Suite 330, Boston, MA  02111-1307  USA
*/

package org.systemsbiology.biofabric.ui.display;

import java.awt.AlphaComposite;
import java.awt.BasicStroke;
import java.awt.Color;
import java.awt.Dimension;
import java.awt.Graphics;
import java.awt.Graphics2D;
import java.awt.Point;
import java.awt.Rectangle;
import java.awt.RenderingHints;
import java.awt.geom.AffineTransform;
import java.awt.geom.Point2D;
import java.awt.image.BufferedImage;
import javax.swing.JPanel;

import org.systemsbiology.biofabric.ui.FabricColorGenerator;
import org.systemsbiology.biofabric.ui.FabricDisplayOptions;
import org.systemsbiology.biofabric.ui.FabricDisplayOptionsManager;
import org.systemsbiology.biofabric.ui.render.PaintCacheSmall;

/****************************************************************************
**
** This is the magnifying glass!
*/

public class FabricMagnifier extends JPanel {
    
  ////////////////////////////////////////////////////////////////////////////
  //
  // PUBLIC CONSTANTS
  //
  //////////////////////////////////////////////////////////////////////////// 
  
  public static final int PREF_SIZE = 10;
  public static final int MIN_SIZE = 2;
  public static final int MAX_SIZE = 10;
  public static final int MAG_GRID = 20;
  
  ////////////////////////////////////////////////////////////////////////////
  //
  // PRIVATE INSTANCE MEMBERS
  //
  ////////////////////////////////////////////////////////////////////////////
   
  private int miniSize_;
  private boolean needInit_;
  private PaintCacheSmall painter_;
  private PaintCacheSmall selectionPainter_;
  private PaintCacheSmall.Reduction selections_;
  private AffineTransform miniTrans_;
  private Rectangle worldRec_;
  private Rectangle clipRec_;
  private Point2D center_;
  private Point centerRC_;
  private boolean ignore_;
  private boolean mouseIn_;
  private boolean byTour_;
  private BioFabricOverview bfo_;
  private BufferedImage bim_;
  private PaintCacheSmall.FloaterSet floaters_;
  private int currSize_;
  private static final long serialVersionUID = 1L;

  ////////////////////////////////////////////////////////////////////////////
  //
  // PUBLIC CONSTRUCTORS
  //
  ////////////////////////////////////////////////////////////////////////////

  /***************************************************************************
  **
  ** Constructor
  */

  public FabricMagnifier(FabricColorGenerator colGen) {
    currSize_ = PREF_SIZE;
    miniSize_ = PREF_SIZE;
    ignore_ = false;
    byTour_ = false;
    mouseIn_ = false;
    needInit_ = true;
    center_ = new Point2D.Double(0.0, 0.0);
    centerRC_ = new Point(0, 0);
    miniTrans_ = new AffineTransform();
    setBackground(Color.white);
    painter_ = new PaintCacheSmall(colGen);
    selectionPainter_ = new PaintCacheSmall(colGen);
    floaters_ = null;
    worldRec_ = new Rectangle();
    clipRec_= new Rectangle();
  }

  ////////////////////////////////////////////////////////////////////////////
  //
  // PUBLIC METHODS
  //
  ////////////////////////////////////////////////////////////////////////////

  /***************************************************************************
  **
  ** Set current selections
  */

  public void setSelections(PaintCacheSmall.Reduction selections) {
    selections_ = selections;
    return;
  } 
  
  /***************************************************************************
  **
  ** Set the current floater
  */

  public void setCurrentFloater(PaintCacheSmall.FloaterSet floaters) {
    floaters_ = floaters;
    return;
  } 

  /***************************************************************************
  **
  ** Let us know who the overview is:
  */

  public void setFabricOverview(BioFabricOverview bfo) {
    bfo_ = bfo;
    return;
  } 
  
  /***************************************************************************
  **
  ** Set the size. The value coming in the door is the number of rows & cols
  ** at maximum magnification.
  */

  public void setCurrSize(int size) {
    if ((size == miniSize_) && !needInit_) {
      return;
    }
    needInit_ = false;
    int oldSize = miniSize_;
    miniSize_ = (size < MIN_SIZE) ? MIN_SIZE : size;
    int newCurr = (currSize_ / oldSize) * miniSize_;
    setZoom(newCurr);
    return;
  } 
  
  /***************************************************************************
  **
  ** Sizing
  */
  
  @Override
  public Dimension getPreferredSize() {
    return (new Dimension(miniSize_ * MAG_GRID, miniSize_ * MAG_GRID));    
  }

  @Override
  public Dimension getMinimumSize() {
    return (new Dimension(miniSize_ * MAG_GRID, miniSize_ * MAG_GRID));    
  }
  
  @Override
  public Dimension getMaximumSize() {
    return (getPreferredSize());    
  }

  /***************************************************************************
  **
  ** Set bounds
  */

  @Override
  public void setBounds(int x, int y, int width, int height) {
    super.setBounds(x, y, width, height);
    setMiniZoom();
    repaint();
    return;
  }    

  /***************************************************************************
  **
  ** Paint
  */

  @Override
  public void paintComponent(Graphics g) {
    super.paintComponent(g);
    Graphics2D g2 = (Graphics2D)g;
    drawingGuts(g2);
    return;
  }

  /***************************************************************************
  **
  ** Bump grid size
  */

  public void bumpGridSize(boolean up) {
    int newSize = (up) ? currSize_ * 2 : currSize_ / 2;
    if (newSize < miniSize_) {
      newSize = miniSize_;
    }
    setZoom(newSize);
    return;
  }
  
  /***************************************************************************
  **
  ** Toggle freeze
  */

  public void toggleFreeze() {
    ignore_ = !ignore_;
    return;
  }
  
  /***************************************************************************
  **
  ** Go up
  */

  public void up() {
    int inc = getZoom() / currSize_;
    centerRC_.y -= inc;
    center_.setLocation(center_.getX(), center_.getY() - (BioFabricPanel.GRID_SIZE * inc));
    setMiniZoom();
    repaint();
    return;
  }
  
  /***************************************************************************
  **
  ** Go down
  */

  public void down() {
    int inc = getZoom() / currSize_;
    centerRC_.y += inc;
    center_.setLocation(center_.getX(), center_.getY() + (BioFabricPanel.GRID_SIZE * inc));
    setMiniZoom();
    repaint();
    return;
  }
  
  /***************************************************************************
  **
  ** Go left
  */

  public void left() {
    int inc = getZoom() / currSize_;
    centerRC_.x -= inc;
    center_.setLocation(center_.getX() - (BioFabricPanel.GRID_SIZE * inc), center_.getY());
    setMiniZoom();
    repaint();
    return;
  }
  
  /***************************************************************************
  **
  ** go right
  */

  public void right() {
    int inc = getZoom() / currSize_;
    centerRC_.x += inc;
    center_.setLocation(center_.getX() + (BioFabricPanel.GRID_SIZE * inc), center_.getY());
    setMiniZoom();
    repaint();
    return;
  }
      
  /***************************************************************************
  **
  ** Set zoom
  */

  public void setZoom(int gridsize) {
    currSize_ = gridsize;
    setMiniZoom();
    repaint();
    return;
  }
  
  /***************************************************************************
  **
  ** Get Zoom
  */

  public int getZoom() {
    return (currSize_);
  } 
   
  /***************************************************************************
  **
  ** Get transform
  */

  public AffineTransform getTransform() {
    return (miniTrans_);
  }

  /***************************************************************************
  **
  ** Get center
  */

  public Point getCenterRC() {
    return (centerRC_);
  }

  /***************************************************************************
  **
  ** Get the clip rect
  */

  public Rectangle getClipRect() {
    return (clipRec_);
  }
   
  /***************************************************************************
  **
  ** Set the center
  */

  public void setCenter(Point2D center, Point centerRC, boolean byTour) {
    if (ignore_) {
      return;
    }
    byTour_ = byTour;
    centerRC_.setLocation(centerRC);
    center_.setLocation(center);
    setMiniZoom();
    repaint();
    return;
  }
  
  /***************************************************************************
  **
  ** Tell us mouse is in
  */

  public void setMouseIn(boolean isIn) {
    if (ignore_) {
      return;
    }
    mouseIn_ = isIn;
    repaint();
    return;
  } 

  /***************************************************************************
  **
  ** Set the zoom
  */

  public void setMiniZoom() {
    Dimension screenDim = getSize();
    int worldWidth = (currSize_ * BioFabricPanel.GRID_SIZE);
    int worldHeight  = (currSize_ * BioFabricPanel.GRID_SIZE);
    worldRec_.setBounds((int)center_.getX() - (worldWidth / 2), 
                        (int)center_.getY() - (worldHeight / 2),
                              worldWidth, worldHeight);   
    clipRec_.setBounds((int)(worldRec_.getX() - BioFabricPanel.GRID_SIZE), (int)(worldRec_.getY() - BioFabricPanel.GRID_SIZE),
                       (int)(worldRec_.getWidth() + (BioFabricPanel.GRID_SIZE * 2)), (int)(worldRec_.getHeight() + (BioFabricPanel.GRID_SIZE * 2)));
    double zoomH = screenDim.getWidth() / worldRec_.getWidth();
    double zoomV = screenDim.getHeight() / worldRec_.getHeight();
    double zoom = Math.min(zoomH, zoomV);

    miniTrans_.setToIdentity();
    miniTrans_.translate(screenDim.getWidth() / 2, screenDim.getHeight() / 2);
    miniTrans_.scale(zoom, zoom);
    miniTrans_.translate(-center_.getX(), -center_.getY());
    bfo_.setMagnifyView(worldRec_);
    return;
  }

  /***************************************************************************
  **
  ** Set painters
  */

  public void setPainters(PaintCacheSmall painter, PaintCacheSmall selectionPainter) {
    painter_ = painter;
    selectionPainter_ = selectionPainter;
    repaint();
    return;
  }

  /***************************************************************************
  **
  ** Drawing core
  */

  private void drawingGuts(Graphics g) {
    if (painter_ == null) {
      return;
    }
    if (!mouseIn_ && !byTour_ && !ignore_) {
      return;
    }
    Graphics2D g2 = (Graphics2D)g;   
    g2.setRenderingHint(RenderingHints.KEY_ANTIALIASING, RenderingHints.VALUE_ANTIALIAS_ON);
    g2.setRenderingHint(RenderingHints.KEY_FRACTIONALMETRICS, RenderingHints.VALUE_FRACTIONALMETRICS_ON);
    AffineTransform saveTrans = g2.getTransform();    
    g2.transform(miniTrans_);
    BasicStroke selectedStroke = new BasicStroke(PaintCacheSmall.STROKE_SIZE, BasicStroke.CAP_SQUARE, BasicStroke.JOIN_MITER);    
    g2.setStroke(selectedStroke);
    painter_.paintIt(g2, clipRec_, null);
    g2.setTransform(saveTrans);
    
    if ((selections_ != null) && selections_.somethingToPaint()) {
      drawSelections(g2, clipRec_);
    }
    if (floaters_ != null) {
      g2.transform(miniTrans_);
      painter_.drawFloater(g2, floaters_); 
    }
    return;
  }
  
  /***************************************************************************
  **
  ** Draw the selections as cleared areas in a partially opaque overlay
  */
  
  public void drawSelections(Graphics2D g2, Rectangle clip) {
    
    FabricDisplayOptions fdo = FabricDisplayOptionsManager.getMgr().getDisplayOptions();
  
    Dimension viewDim = getSize();
    
    if ((bim_ == null) || (bim_.getHeight() != viewDim.height) || (bim_.getWidth() != viewDim.width)) {
      bim_ = new BufferedImage(viewDim.width, viewDim.height, BufferedImage.TYPE_INT_ARGB);
    }
    Graphics2D ig2 = bim_.createGraphics();
    ig2.setTransform(new AffineTransform());
    Color drawCol = new Color(1.0f, 1.0f, 1.0f, (float)fdo.getSelectionOpaqueLevel()); 
    ig2.setBackground(drawCol);
    ig2.clearRect(0, 0, viewDim.width, viewDim.height);
    ig2.setRenderingHint(RenderingHints.KEY_ANTIALIASING, RenderingHints.VALUE_ANTIALIAS_ON);
    ig2.setRenderingHint(RenderingHints.KEY_FRACTIONALMETRICS, RenderingHints.VALUE_FRACTIONALMETRICS_ON);
    BasicStroke selectedStroke = new BasicStroke(PaintCacheSmall.STROKE_SIZE, BasicStroke.CAP_SQUARE, BasicStroke.JOIN_MITER);    
    ig2.setStroke(selectedStroke);
    ig2.setTransform(miniTrans_);
    ig2.setComposite(AlphaComposite.Src);
    selectionPainter_.paintIt(ig2, clip, selections_);
    g2.drawImage(bim_, 0, 0, viewDim.width, viewDim.height, null);
    return;
  } 
}
```
