## Supplementary material for "BioFabric Visualization of Network Alignments": BioFabric Source Code: FabricMagnifyingTool.java

```
/*
**    Copyright (C) 2003-2018 Institute for Systems Biology 
**                            Seattle, Washington, USA. 
**
**    This library is free software; you can redistribute it and/or
**    modify it under the terms of the GNU Lesser General Public
**    License as published by the Free Software Foundation; either
**    version 2.1 of the License, or (at your option) any later version.
**
**    This library is distributed in the hope that it will be useful,
**    but WITHOUT ANY WARRANTY; without even the implied warranty of
**    MERCHANTABILITY or FITNESS FOR A PARTICULAR PURPOSE.  See the GNU
**    Lesser General Public License for more details.
**
**    You should have received a copy of the GNU Lesser General Public
**    License along with this library; if not, write to the Free Software
**    Foundation, Inc., 59 Temple Place, Suite 330, Boston, MA  02111-1307  USA
*/

package org.systemsbiology.biofabric.ui.display;

import java.awt.BasicStroke;
import java.awt.CardLayout;
import java.awt.Color;
import java.awt.Dimension;
import java.awt.Font;
import java.awt.Graphics;
import java.awt.Graphics2D;
import java.awt.GridBagConstraints;
import java.awt.GridBagLayout;
import java.awt.GridLayout;
import java.awt.Point;
import java.awt.Rectangle;
import java.awt.event.ActionEvent;
import java.awt.event.KeyEvent;
import java.awt.event.MouseAdapter;
import java.awt.event.MouseEvent;
import java.awt.font.FontRenderContext;
import java.awt.geom.AffineTransform;
import java.awt.geom.Line2D;
import java.awt.geom.NoninvertibleTransformException;
import java.awt.geom.Point2D;
import java.awt.geom.Rectangle2D;
import java.util.ArrayList;
import java.util.List;
import javax.swing.AbstractAction;
import javax.swing.ActionMap;
import javax.swing.InputMap;
import javax.swing.JComponent;
import javax.swing.JLabel;
import javax.swing.JList;
import javax.swing.JPanel;
import javax.swing.JScrollPane;
import javax.swing.KeyStroke;
import javax.swing.ListSelectionModel;

import org.systemsbiology.biofabric.api.model.NetNode;
import org.systemsbiology.biofabric.api.util.ExceptionHandler;
import org.systemsbiology.biofabric.api.util.MinMax;
import org.systemsbiology.biofabric.model.BioFabricNetwork;
import org.systemsbiology.biofabric.ui.FabricColorGenerator;
import org.systemsbiology.biofabric.ui.FabricDisplayOptionsManager;
import org.systemsbiology.biofabric.ui.render.PaintCacheSmall;
import org.systemsbiology.biofabric.util.ColorListRenderer;
import org.systemsbiology.biofabric.util.UiUtil;


/****************************************************************************
**
** This is the magnifying glass!
*/

public class FabricMagnifyingTool extends JPanel {
    
  ////////////////////////////////////////////////////////////////////////////
  //
  // PRIVATE CONSTANTS
  //
  //////////////////////////////////////////////////////////////////////////// 
  
  private static final float TOP_LINK_SPACING_ = 12.0F;
  
  ////////////////////////////////////////////////////////////////////////////
  //
  // PUBLIC CONSTANTS
  //
  //////////////////////////////////////////////////////////////////////////// 
  
  ////////////////////////////////////////////////////////////////////////////
  //
  // PRIVATE INSTANCE MEMBERS
  //
  ////////////////////////////////////////////////////////////////////////////

  private boolean ignore_;
  private FabricMagnifier myMag_;
  private JPanel lockPanel_;
  private JLabel lockStatus_;
  private JLabel toLock_;
  private NodeLabels nodeLabels_;
  private LinkLabels topLinkLabels_;
  private LinkDisplay linkDisplay_;
  private BioFabricNetwork model_;
  private Font tiny_;
  private FabricColorGenerator colGen_;
  private boolean mouseIn_;
  private boolean byTour_;
  private boolean accept_;
  private Point2D mouseCenter_;
  private int currSize_;
  private  boolean neverSet_;
  
  private CardLayout myCard_;
  private static final long serialVersionUID = 1L;

  ////////////////////////////////////////////////////////////////////////////
  //
  // PUBLIC CONSTRUCTORS
  //
  ////////////////////////////////////////////////////////////////////////////

  /***************************************************************************
  **
  ** Constructor
  */

  public FabricMagnifyingTool(FabricColorGenerator colGen) {
    mouseCenter_ = null;
    colGen_ = colGen;
    ignore_ = false;
    accept_ = true;
    neverSet_ = true;
    lockPanel_ = new JPanel();
    Font micro = new Font("SansSerif", Font.PLAIN, 10);
    lockStatus_ = new JLabel("Tracking", JLabel.CENTER);
    lockStatus_.setFont(micro);
    lockStatus_.setOpaque(true);
    lockStatus_.setBackground(Color.green);
    toLock_ = new JLabel("Lock: Ctrl-Space", JLabel.CENTER);
    toLock_.setFont(micro);
    JLabel toZoom = new JLabel("Zoom: Z,C,+,-", JLabel.CENTER);
    toZoom.setFont(micro);
    JLabel toPan = new JLabel("Pan: A,W,X,D", JLabel.CENTER);
    toPan.setFont(micro);
    JLabel toPan2 = new JLabel("Pan: \u2190,\u2191,\u2193,\u2192", JLabel.CENTER);
    toPan2.setFont(micro);
    lockPanel_.setLayout(new GridLayout(0,1));
    lockPanel_.add(lockStatus_);
    lockPanel_.add(toLock_);
    lockPanel_.add(toZoom);
    lockPanel_.add(toPan);   
    lockPanel_.add(toPan2);   
    myMag_ = new FabricMagnifier(colGen);
    myMag_.addMouseListener(new MouseHandler());
    nodeLabels_ = new NodeLabels();
    topLinkLabels_ = new LinkLabels();
    linkDisplay_ = new LinkDisplay();
    myMag_.setFocusable(true);
    mouseIn_ = false;
    byTour_ = false;
    currSize_ = FabricMagnifier.PREF_SIZE;
    
    JPanel realGuts = sizeGuts(currSize_);
    
    myCard_ = new CardLayout();
    setLayout(myCard_);
    add(realGuts, "SUPanel");
    JPanel blankPanel = new JPanel();
    blankPanel.setBackground(Color.white);
    add(blankPanel, "Hiding");
       
    tiny_ = new Font("SansSerif", Font.BOLD, 10);
    
  }

  ////////////////////////////////////////////////////////////////////////////
  //
  // PUBLIC METHODS
  //
  ////////////////////////////////////////////////////////////////////////////

  /***************************************************************************
  **
  ** Set selections
  */

  public void setSelections(PaintCacheSmall.Reduction selections) {
  	myMag_.setSelections(selections);
    return;
  }    

  /***************************************************************************
  **
  ** Are we ignoring the mouse?
  */

  public boolean isIgnoring() {
    return (ignore_);
  }    
 
  /***************************************************************************
  **
  ** Enable or disable the panel
  */

  public void enableControls(boolean enabled) {
    accept_ = enabled;    
    myCard_.show(this, (enabled) ? "SUPanel" : "Hiding"); 
    return;
  }    
  
  /***************************************************************************
  **
  ** Set the current floater
  */

  public void setCurrentFloater(PaintCacheSmall.FloaterSet floaters) {
    myMag_.setCurrentFloater(floaters);
    return;
  }  

  /***************************************************************************
  **
  ** Drawing core
  */
  
  public void keyInstall(JPanel cpane) {
    
    // Problem with coming up with a universal freeze key,
    // since EVERYBODY loves the space bar, and LOTS of left hand CTRL
    // keys are are already common
    
    InputMap iMapf = myMag_.getInputMap(JComponent.WHEN_FOCUSED);
    InputMap iMapw = myMag_.getInputMap(JComponent.WHEN_IN_FOCUSED_WINDOW);
    ActionMap aMap = myMag_.getActionMap();

    iMapw.put(KeyStroke.getKeyStroke(KeyEvent.VK_SPACE, KeyEvent.CTRL_DOWN_MASK), "MagnifyFreeze");

    aMap.put("MagnifyFreeze", new AbstractAction() {
      private static final long serialVersionUID = 1L;
      public void actionPerformed(ActionEvent e) {
        try {
          if (!accept_) {
            return;
          }
          myMag_.requestFocusInWindow();
          myMag_.toggleFreeze();
          ignore_ = !ignore_;
          lockStatus_.setText((ignore_) ? "Locked" : "Tracking");
          lockStatus_.setBackground((ignore_) ? Color.red : Color.green);      
          lockStatus_.invalidate();
          toLock_.setText((ignore_) ? "Unlock: Ctrl-Space" : "Lock: Ctrl-Space");
          toLock_.invalidate();
          lockPanel_.validate();
          repaint();
        } catch (Exception ex) {
          ExceptionHandler.getHandler().displayException(ex);
        }
      }
    });
    
    iMapw.put(KeyStroke.getKeyStroke(KeyEvent.VK_C, 0), "MagnifyMore");
    iMapw.put(KeyStroke.getKeyStroke(KeyEvent.VK_EQUALS, 0), "MagnifyMore");
    aMap.put("MagnifyMore", new AbstractAction() {
      private static final long serialVersionUID = 1L;
      public void actionPerformed(ActionEvent e) {
        try {
          if (!accept_) {
            return;
          }
          myMag_.bumpGridSize(false);
          linkSync();
          myMag_.requestFocusInWindow();
          repaint();
        } catch (Exception ex) {
          ExceptionHandler.getHandler().displayException(ex);
        }
      }
    });
    iMapw.put(KeyStroke.getKeyStroke(KeyEvent.VK_Z, 0), "MagnifyLess");
    iMapw.put(KeyStroke.getKeyStroke(KeyEvent.VK_MINUS, 0), "MagnifyLess");
    aMap.put("MagnifyLess", new AbstractAction() {
      private static final long serialVersionUID = 1L;
      public void actionPerformed(ActionEvent e) {
        try {
          if (!accept_) {
            return;
          }
          myMag_.bumpGridSize(true);
          linkSync();
          myMag_.requestFocusInWindow();
          repaint();
        } catch (Exception ex) {
          ExceptionHandler.getHandler().displayException(ex);
        }
      }
    });

    iMapf.put(KeyStroke.getKeyStroke(KeyEvent.VK_KP_UP, 0), "MagnifyUp");
    iMapf.put(KeyStroke.getKeyStroke(KeyEvent.VK_UP, 0), "MagnifyUp");
    iMapw.put(KeyStroke.getKeyStroke(KeyEvent.VK_W, 0), "MagnifyUp");
    aMap.put("MagnifyUp", new AbstractAction() {
      private static final long serialVersionUID = 1L;
      public void actionPerformed(ActionEvent e) {
        try { 
          if (!accept_) {
            return;
          }
          myMag_.up();
          linkSync();
          myMag_.requestFocusInWindow();
          repaint();
        } catch (Exception ex) {
          ExceptionHandler.getHandler().displayException(ex);
        }
      }
    });
    iMapf.put(KeyStroke.getKeyStroke(KeyEvent.VK_KP_DOWN, 0), "MagnifyDown");
    iMapf.put(KeyStroke.getKeyStroke(KeyEvent.VK_DOWN, 0), "MagnifyDown");
    iMapw.put(KeyStroke.getKeyStroke(KeyEvent.VK_X, 0), "MagnifyDown");
    aMap.put("MagnifyDown", new AbstractAction() {
      private static final long serialVersionUID = 1L;
      public void actionPerformed(ActionEvent e) {
        try {
          if (!accept_) {
            return;
          }
          myMag_.down();
          linkSync();
          myMag_.requestFocusInWindow();
          repaint();
        } catch (Exception ex) {
          ExceptionHandler.getHandler().displayException(ex);
        }
      }
    });
    iMapf.put(KeyStroke.getKeyStroke(KeyEvent.VK_KP_LEFT, 0), "MagnifyLeft");
    iMapf.put(KeyStroke.getKeyStroke(KeyEvent.VK_LEFT, 0), "MagnifyLeft");
    iMapw.put(KeyStroke.getKeyStroke(KeyEvent.VK_A, 0), "MagnifyLeft");
    aMap.put("MagnifyLeft", new AbstractAction() {
      private static final long serialVersionUID = 1L;
      public void actionPerformed(ActionEvent e) {
        try {
          if (!accept_) {
            return;
          }
          myMag_.left();
          linkSync();
          myMag_.requestFocusInWindow();
          repaint();
        } catch (Exception ex) {
          ExceptionHandler.getHandler().displayException(ex);
        }
      }
    });
    iMapf.put(KeyStroke.getKeyStroke(KeyEvent.VK_KP_RIGHT, 0), "MagnifyRight");
    iMapf.put(KeyStroke.getKeyStroke(KeyEvent.VK_RIGHT, 0), "MagnifyRight");
    iMapw.put(KeyStroke.getKeyStroke(KeyEvent.VK_D, 0), "MagnifyRight");
    aMap.put("MagnifyRight", new AbstractAction() {
      private static final long serialVersionUID = 1L;
      public void actionPerformed(ActionEvent e) {
        try {
          if (!accept_) {
            return;
          }
          myMag_.right();
          linkSync();
          myMag_.requestFocusInWindow();
          repaint();
        } catch (Exception ex) {
          ExceptionHandler.getHandler().displayException(ex);
        }
      }
    });
    return;
  }
 
  /***************************************************************************
  **
  ** Sizing
  */
  
  @Override
  public Dimension getPreferredSize() {
    return (getMinimumSize());    
  }

  @Override
  public Dimension getMinimumSize() {
    if (neverSet_) {
      return (new Dimension(550, (2 * (FabricMagnifier.PREF_SIZE * FabricMagnifier.MAG_GRID))));
    } else {
      return (new Dimension(550, (2 * (FabricMagnifier.MIN_SIZE * FabricMagnifier.MAG_GRID))));
    }
  }
  
  @Override
  public Dimension getMaximumSize() {
    return (new Dimension(4000, (2 * (FabricMagnifier.MAX_SIZE * FabricMagnifier.MAG_GRID)) + 40));    
  }

  /***************************************************************************
  **
  ** 
  */

  public void setFabricOverview(BioFabricOverview bfo) {
    myMag_.setFabricOverview(bfo);
    return;
  } 
  
  /***************************************************************************
  **
  ** 
  */

  public void setModel(BioFabricNetwork model) {
    model_ = model;
    return;
  } 
   
  /***************************************************************************
  **
  ** 
  */

  @Override
  public void setBounds(int x, int y, int width, int height) {
    neverSet_ = false;
    currSize_ = (int)Math.floor(height / (FabricMagnifier.MAG_GRID + TOP_LINK_SPACING_));
    myMag_.setCurrSize(currSize_);
    super.setBounds(x, y, width, height);
    linkSync();
    repaint();
    return;
  } 
  
  /***************************************************************************
  **
  ** 
  */

  public void setCenter(Point2D center, Point cprc, boolean byTour) {
    if (!byTour) {
      mouseCenter_ = center;
    }
    myMag_.setCenter(center, cprc, byTour);
    byTour_ = byTour;
    if (ignore_) {
      return;
    }
    linkSync();
    repaint();
    return;
  }
  
  /***************************************************************************
  **
  ** 
  */

  public void setMouseIn(boolean isIn) {
    myMag_.setMouseIn(isIn);
    mouseIn_ = isIn;
    if (ignore_) {
      return;
    }
    linkSync();
    repaint();
    return;
  }
 
  /***************************************************************************
  **
  ** 
  */

  public void linkSync() {
    List<LinkListElementDisplay> links;
    if (model_ != null) { 
      links = linkDisplay_.calcLinks();      
    } else {
      links = new ArrayList<LinkListElementDisplay>();
    }
    linkDisplay_.setLinks(links);
    return;
  }
    
  /***************************************************************************
  **
  **
  */

  public void setPainters(PaintCacheSmall painter, PaintCacheSmall selectionPainter) {
    myMag_.setPainters(painter, selectionPainter);
    return;
  }
 
  /***************************************************************************
  **
  ** 
  */

  public Rectangle getClipRect() {
    return (myMag_.getClipRect());
  }
  
  /***************************************************************************
  **
  ** 
  */

  public Point2D getMouseLoc() {
    return (mouseCenter_);
  }
  
  ////////////////////////////////////////////////////////////////////////////
  //
  // PRIVATE METHODS
  //
  ////////////////////////////////////////////////////////////////////////////  
  
  /***************************************************************************
  **
  ** Set sizing
  */
  
  private JPanel sizeGuts(int magSize) {
  
    JPanel realGuts = new JPanel();
    
    realGuts.setLayout(new GridBagLayout());
    GridBagConstraints gbc = new GridBagConstraints();    

    //
    // NOTE NO GROWTH ALLOWED IN VERTICAL DIRECTION.
    // Dividing real dims by 10 to make GridBag happy...
    //
    // 10C x 14R
    int col = 0;
    int row = 0;
    int wth = 10;
    int ht = (int)Math.ceil((((magSize + 1) * TOP_LINK_SPACING_) / 10) + 4);
    UiUtil.gbcSet(gbc, col, row, wth, ht, UiUtil.BO, 0, 0, 0, 0, 0, 0, UiUtil.N, 0.25, 0.0);
    realGuts.add(lockPanel_, gbc);
     
    // 45C x 14R
    col = wth;
    row = 0;
    wth = 45;
    ht = (int)Math.ceil((((magSize + 1) * TOP_LINK_SPACING_) / 10) + 4);
    UiUtil.gbcSet(gbc, col, row, wth, ht, UiUtil.BO, 0, 0, 0, 0, 0, 0, UiUtil.SW, 0.75, 0.0);
    realGuts.add(topLinkLabels_, gbc);
     
    // 10C x 20R
    col = 0;
    row = ht;
    wth = 10;
    ht = (magSize * FabricMagnifier.MAG_GRID) / 10;
    UiUtil.gbcSet(gbc, col, row, wth, ht, UiUtil.BO, 0, 0, 0, 0, 0, 0, UiUtil.CEN, 0.25, 0.0);
    realGuts.add(nodeLabels_, gbc);
    
    // 20C x 20R NO GROWTH AT ALL!
    col = wth;
    wth = (magSize * FabricMagnifier.MAG_GRID) / 10;
    ht = (magSize * FabricMagnifier.MAG_GRID) / 10;  
    UiUtil.gbcSet(gbc, col, row, wth, ht, UiUtil.NONE, 0, 0, 0, 0, 0, 0, UiUtil.CEN, 0.0, 0.0);
    realGuts.add(myMag_, gbc);
    
    // 25C x 20R
    col = col + wth;
    wth = 45 - wth;
    ht = (magSize * FabricMagnifier.MAG_GRID) / 10;     
    UiUtil.gbcSet(gbc, col, row, wth, ht, UiUtil.BO, 0, 0, 0, 0, 0, 0, UiUtil.CEN, 0.75, 0.0);
    realGuts.add(linkDisplay_, gbc);
    return (realGuts);
  }

  /***************************************************************************
  **
  ** Labels the nodes
  */  
  
  private class NodeLabels extends JPanel {
    private static final long serialVersionUID = 1L;
    
    NodeLabels() {
      setBackground(new Color(245, 245, 245));
    }
    @Override
    public Dimension getPreferredSize() {
      return (new Dimension(100, (currSize_ * FabricMagnifier.MAG_GRID)));    
    }
    @Override
    public Dimension getMinimumSize() {
      return (new Dimension(100, 10));    
    }
    @Override
    public Dimension getMaximumSize() {
      return (new Dimension(4000, (currSize_ * FabricMagnifier.MAG_GRID)));    
    }
    @Override
    public void paintComponent(Graphics g) {
      super.paintComponent(g);
      if (model_ == null) {
        return;
      }
      if (!mouseIn_ && !byTour_ && !ignore_) {
        return;
      }
      Graphics2D g2 = (Graphics2D)g;
      Point myCen = myMag_.getCenterRC();
      int myZoom = myMag_.getZoom();
      if (myZoom != currSize_) {
        return;
      }
      Point2D atPoint = new Point2D.Double(0.0, 0.0);
      Point2D drawPoint = new Point2D.Double(0.0, 0.0);
      boolean showShadows = FabricDisplayOptionsManager.getMgr().getDisplayOptions().getDisplayShadows();
      AffineTransform ac = myMag_.getTransform();
      g2.setFont(tiny_);
      g2.setPaint(Color.BLACK);
      FontRenderContext frc = g2.getFontRenderContext();
      int startCen = myCen.y - (currSize_ / 2) - 1;
      int endCen = myCen.y + (currSize_ / 2) + 1;   
      for (int i = startCen; i <= endCen; i++) {
        NetNode node = model_.getNodeIDForRow(Integer.valueOf(i));
        if (node == null) {
          continue;
        }
        BioFabricNetwork.NodeInfo ni = model_.getNodeDefinition(node);
        MinMax nimm = ni.getColRange(showShadows);
        if ((nimm.min <= myCen.x + (currSize_ / 2)) && (nimm.max >= myCen.x - (currSize_ / 2))) {
          Rectangle2D bounds = tiny_.getStringBounds(node.getName(), frc);
          atPoint.setLocation(0.0, i * BioFabricPanel.GRID_SIZE);
          ac.transform(atPoint, drawPoint);
          g2.drawString(node.getName(), getWidth() - (float)bounds.getWidth() - 5.0F, (float)drawPoint.getY() + ((float)bounds.getHeight() / 3.0F)); 
        } 
      }
      return;
    }    
  } 
  
  /***************************************************************************
  **
  ** Labels the links
  */  
  
  private class LinkLabels extends JPanel {
    private static final long serialVersionUID = 1L;
    
    LinkLabels() {
      setBackground(new Color(245, 245, 245));
    }
    
    @Override
    public Dimension getPreferredSize() {
      return (new Dimension(450, (int)((currSize_ + 1) * TOP_LINK_SPACING_)));   
    }
    
    @Override
    public Dimension getMinimumSize() {
      return (new Dimension(450, (int)((currSize_ + 1) * TOP_LINK_SPACING_)));
    }
    
    @Override
    public Dimension getMaximumSize() {
      return (new Dimension(4000, (int)((currSize_ + 1) * TOP_LINK_SPACING_)));    
    }
    
    @Override
    public void paintComponent(Graphics g) {
      super.paintComponent(g);
      if (model_ == null) {
        return;
      }
      if (!mouseIn_ && !byTour_ && !ignore_) {
        return;
      }
      Graphics2D g2 = (Graphics2D)g;
      Point myCen = myMag_.getCenterRC();
      int myZoom = myMag_.getZoom();
      if (myZoom != currSize_) {
        return;
      }
      boolean showShadows = FabricDisplayOptionsManager.getMgr().getDisplayOptions().getDisplayShadows();
      Point2D atPoint = new Point2D.Double(0.0, 0.0);
      Point2D drawPoint = new Point2D.Double(0.0, 0.0);
      AffineTransform ac = myMag_.getTransform();
      g2.setFont(tiny_);
      BasicStroke selectedStroke = new BasicStroke(1, BasicStroke.CAP_SQUARE, BasicStroke.JOIN_MITER);    
      g2.setStroke(selectedStroke);
     
      FontRenderContext frc = g2.getFontRenderContext();
      int numCol = model_.getColumnCount(showShadows);
      int startCen = myCen.x - (currSize_ / 2);
      if (startCen < 0) {
        startCen = 0;
      }
      if (startCen >= numCol) {
        return;
      }
    
      int endCen = myCen.x + (currSize_ / 2);
      if (endCen > numCol) {
        endCen = numCol;
      }
      int count = -1;
     
      float deltaY = TOP_LINK_SPACING_;
      float startY = TOP_LINK_SPACING_;
      for (int i = startCen; i <= endCen; i++) {
        count++;
        Integer colObj = Integer.valueOf(i);
        NetNode src = model_.getSourceIDForColumn(colObj, showShadows);
        NetNode trg = model_.getTargetIDForColumn(colObj, showShadows);
        if ((src == null) || (trg == null)) {
          continue;
        }
        BioFabricNetwork.LinkInfo li = model_.getLinkDefinition(colObj, showShadows);
        if (li == null) {
          continue;
        }
        int minRow = li.topRow();
        int maxRow = li.bottomRow();
        
        if ((minRow <= (myCen.y + (currSize_ / 2))) && (maxRow >= (myCen.y - (currSize_ / 2)))) {
          String linkDisp = li.getLink().toDisplayString();
          Rectangle2D bounds = tiny_.getStringBounds(linkDisp, frc);
          atPoint.setLocation((i * BioFabricPanel.GRID_SIZE), 0.0);
          ac.transform(atPoint, drawPoint);
          float baseptX = (float)drawPoint.getX();
          if ((baseptX < 0) || (baseptX > (currSize_ * FabricMagnifier.MAG_GRID))) {
            continue;
          }
          float baseptY = startY + (count * deltaY);
          g2.setPaint(Color.BLACK);
          g2.drawString(linkDisp, baseptX + 5.0F, baseptY);
          Color paintCol = colGen_.getModifiedColor(li.getColorKey(), FabricColorGenerator.DARKER);
          float boxBaseY = baseptY - ((float)bounds.getHeight() / 3.0F);
          Line2D line = new Line2D.Float(baseptX, boxBaseY, baseptX, 400.0F);
          g2.setColor(paintCol);
          g2.draw(line);
          Rectangle2D rect = new Rectangle2D.Float(baseptX - 2.0F, boxBaseY - 2.0F, 5.0F, 5.0F);
          g2.fill(rect);
        }
      }
      return;
    }    
  } 
  
  /***************************************************************************
  **
  ** Displays all the links in a JList
  */  
  
  private class LinkDisplay extends JPanel {
    
    private JList linkList_;
    private ColorListRenderer renderer_;
    private static final long serialVersionUID = 1L;
     
    LinkDisplay() {
      ArrayList myList = new ArrayList();
      linkList_ = new JList(myList.toArray());
      linkList_.setSelectionMode(ListSelectionModel.SINGLE_SELECTION);
      renderer_ = new ColorListRenderer(myList);
      linkList_.setCellRenderer(renderer_);
      JScrollPane jsp = new JScrollPane(linkList_);
      jsp.setHorizontalScrollBarPolicy(JScrollPane.HORIZONTAL_SCROLLBAR_AS_NEEDED);
      jsp.setVerticalScrollBarPolicy(JScrollPane.VERTICAL_SCROLLBAR_AS_NEEDED);    
      setLayout(new GridLayout(1, 1));
      add(jsp);
    }
    
    public Dimension getPreferredSize() {
      return (new Dimension(250, (currSize_ * FabricMagnifier.MAG_GRID)));    
    }

    public Dimension getMinimumSize() {
      return (new Dimension(200, (currSize_ * FabricMagnifier.MAG_GRID)));    
    }

    public Dimension getMaximumSize() {
      return (new Dimension(4000, (currSize_ * FabricMagnifier.MAG_GRID)));    
    }
    
    public void setLinks(List<LinkListElementDisplay> links) {
      linkList_.clearSelection();
      linkList_.setListData(links.toArray());
      renderer_.setValues(links);
      return;
    }
    
    public List<LinkListElementDisplay> calcLinks() {
      ArrayList<LinkListElementDisplay> list = new ArrayList<LinkListElementDisplay>();
      if (!mouseIn_ && !byTour_ && !ignore_) {
        return (list);
      }
    
      Point myCen = myMag_.getCenterRC();
      int zoom = myMag_.getZoom();
      int halfZoom = zoom / 2;
      int startCen = myCen.x - halfZoom;
      int endCen = myCen.x + halfZoom;
      boolean showShadows = FabricDisplayOptionsManager.getMgr().getDisplayOptions().getDisplayShadows();
   
      for (int i = startCen; i <= endCen; i++) {
        Integer colObj = Integer.valueOf(i);
        NetNode src = model_.getSourceIDForColumn(colObj, showShadows);
        if (src == null) {
          continue;
        }
        BioFabricNetwork.LinkInfo li = model_.getLinkDefinition(colObj, showShadows);
        if (li == null) {
          continue;
        }
        int minRow = li.topRow();
        int maxRow = li.bottomRow();
        
        if (((minRow >= (myCen.y - halfZoom)) && (minRow <= (myCen.y + halfZoom))) ||
            ((maxRow >= (myCen.y - halfZoom)) && (maxRow <= (myCen.y + halfZoom)))) {     
          Color paintCol = colGen_.getModifiedColor(li.getColorKey(), FabricColorGenerator.DARKER);
          list.add(new LinkListElementDisplay(paintCol, li.getLink().toDisplayString()));
        }
      }
      return (list);
    }
  }
  
  /***************************************************************************
  **
  ** Labels the links
  */  
  
  private class LinkListElementDisplay implements ColorListRenderer.ColorSource {
     
    private Color color_;
    private String desc_;
    
    LinkListElementDisplay(Color color, String desc) {
      color_ = color;
      desc_ = desc;
    }
    
    public Color getColor() {
      return (color_);
    }
    
    public String getDescription(){
      return (desc_);
    } 
  }
  
  /***************************************************************************
  **
  ** Handles click events WHY IS THIS EVEN HERE????
  */  
  
  public class MouseHandler extends MouseAdapter { 
    
    private Point lastPress_;
    
    private final static int CLICK_SLOP_  = 2;
      
    public void mouseClicked(MouseEvent me) {
      if (me.isPopupTrigger()) {
        // Do nothing....
      }
      return;
    }  
    private void handleClick(int lastX, int lastY) {    
      Point2D clickPoint = new Point2D.Double(lastX, lastY);
      Point2D worldPoint = new Point2D.Double(0.0, 0.0);
      AffineTransform ac = myMag_.getTransform();
      try {
        ac.inverseTransform(clickPoint, worldPoint);
      } catch (NoninvertibleTransformException nite) {
        return;
        // silent fail....
      }
      return;
    }
     
    public void mousePressed(MouseEvent me) {
      try {
        if (me.isPopupTrigger()) {
          // Do nothing....
          return;
        } else {
          lastPress_ = new Point(me.getX(), me.getY());
        }
      } catch (Exception ex) {
        ExceptionHandler.getHandler().displayException(ex);
      }
      return;
    }   

    public void mouseReleased(MouseEvent me) {
      try {
        if (me.isPopupTrigger()) {
          // Do nothing....
          return;
        }
        
        int currX = me.getX();
        int currY = me.getY();     
        if (lastPress_ == null) {
          return;
        }
        int lastX = lastPress_.x;
        int lastY = lastPress_.y;
        int diffX = Math.abs(currX - lastX);
        int diffY = Math.abs(currY - lastY); 
        
        // Note that a ctrl-drag leaves us with diff == 0, thus
        // it would select after the ctrl drag!
        
        if ((diffX <= CLICK_SLOP_) && (diffY <= CLICK_SLOP_)) { 
          handleClick(lastX, lastY);
        }
        lastPress_ = null;  // DO THIS NO MATTER WHAT TOO       
      } catch (Exception ex) {
        ExceptionHandler.getHandler().displayException(ex);
      }       
      return;
    }
  } 
}
```
