## Supplementary material for "BioFabric Visualization of Network Alignments": BioFabric Source Code: FabricNavTool.java

```
/*
**    Copyright (C) 2003-2017 Institute for Systems Biology 
**                            Seattle, Washington, USA. 
**
**    This library is free software; you can redistribute it and/or
**    modify it under the terms of the GNU Lesser General Public
**    License as published by the Free Software Foundation; either
**    version 2.1 of the License, or (at your option) any later version.
**
**    This library is distributed in the hope that it will be useful,
**    but WITHOUT ANY WARRANTY; without even the implied warranty of
**    MERCHANTABILITY or FITNESS FOR A PARTICULAR PURPOSE.  See the GNU
**    Lesser General Public License for more details.
**
**    You should have received a copy of the GNU Lesser General Public
**    License along with this library; if not, write to the Free Software
**    Foundation, Inc., 59 Temple Place, Suite 330, Boston, MA  02111-1307  USA
*/

package org.systemsbiology.biofabric.ui.display;

import java.awt.BorderLayout;
import java.awt.CardLayout;
import java.awt.Color;
import java.awt.Dimension;
import java.awt.Font;
import java.awt.GridLayout;
import java.awt.event.ActionEvent;
import java.awt.event.ActionListener;
import java.text.MessageFormat;

import javax.swing.Box;
import javax.swing.JCheckBox;
import javax.swing.JFrame;
import javax.swing.JLabel;
import javax.swing.JOptionPane;
import javax.swing.JPanel;
import javax.swing.border.EmptyBorder;
import javax.swing.border.LineBorder;

import org.systemsbiology.biofabric.api.util.ExceptionHandler;
import org.systemsbiology.biofabric.api.util.FixedJButton;
import org.systemsbiology.biofabric.util.ResourceManager;

/****************************************************************************
**
** This is the navigator
*/

public class FabricNavTool extends JPanel {
    
  ////////////////////////////////////////////////////////////////////////////
  //
  // PRIVATE INSTANCE MEMBERS
  //
  ////////////////////////////////////////////////////////////////////////////

  private static final int BIG_     = 0;
  private static final int SMALL_   = 1;
  private static final int NUM_PAN_ = 2;
  
  private BioFabricPanel bfp_;
  private FixedJButton[] startAtCurrent_;    
  private FixedJButton[] chooseAStart_;      
  private InfoPanel[] infoPanel_;  
  private FixedJButton[] buttonUp_;  
  private FixedJButton[] buttonRight_;    
  private FixedJButton[] buttonFarRight_;      
  private FixedJButton[] buttonLeft_;
  private FixedJButton[] buttonFarLeft_; 
  private FixedJButton[] buttonDown_; 
  private FixedJButton[] buttonZoom_;
  private FixedJButton[] buttonLinkZone_;
  private FixedJButton[] buttonClear_;  
  private JCheckBox[] selectedOnly_;  
  private BioFabricPanel.TourStatus currTstat_;
  private boolean haveSelection_;
  private boolean haveAModel_;
  private boolean controlsEnabled_;
  private JFrame topWindow_;
  private boolean doSOReset_;
  private int currPan_;
  private CardLayout clay_;
  private JPanel bigPan_;
  private JPanel smallPan_;
  private boolean neverSet_;
  private static final long serialVersionUID = 1L;
  
  ////////////////////////////////////////////////////////////////////////////
  //
  // PUBLIC CONSTRUCTORS
  //
  ////////////////////////////////////////////////////////////////////////////

  /***************************************************************************
  **
  ** Constructor
  */

  public FabricNavTool(JFrame topWindow) {

    setBackground(Color.WHITE);
    
    buildButtons();
    controlsEnabled_ = true;
    topWindow_ = topWindow;
    doSOReset_ = false;
    neverSet_ = true;
    
    clay_ = new CardLayout();
    setLayout(clay_);
    currPan_ = BIG_;
    haveAModel_ = false;
    haveSelection_ = false;
    smallPan_ = fillPanel(SMALL_);
    add(smallPan_, "small");
    bigPan_ = fillPanel(BIG_);
    add(bigPan_, "big");
    clay_.show(this, "big");
    clearTour();
  }

  ////////////////////////////////////////////////////////////////////////////
  //
  // PUBLIC METHODS
  //
  ////////////////////////////////////////////////////////////////////////////
  
  /***************************************************************************
  **
  ** Sizing
  */

  @Override
  public Dimension getPreferredSize() {
    return (bigPan_.getPreferredSize());
  }

  @Override
  public Dimension getMinimumSize() {
    if (neverSet_) {
      return (bigPan_.getMinimumSize());
    }
    return (smallPan_.getMinimumSize());
  }
  
  @Override
  public Dimension getMaximumSize() {
    return (bigPan_.getMaximumSize());
  }
    
  /***************************************************************************
  **
  ** 
  */

  @Override
  public void setBounds(int x, int y, int width, int height) {
    neverSet_ = false;
    Dimension minBig = bigPan_.getMinimumSize();
    if ((currPan_ == BIG_) && ((minBig.height > height) || (minBig.width > width))) {
      currPan_ = SMALL_;
      clay_.show(this, "small");
    } else if ((currPan_ == SMALL_) && (minBig.height < height) && (minBig.width < width)) {
      currPan_ = BIG_;
      clay_.show(this, "big");
    }
    super.setBounds(x, y, width, height);
    repaint();
    return;
  } 
  
  /***************************************************************************
  **
  ** Reset the skip selections
  */

  public void resetSkipSelections() {
    for (int i = 0; i < NUM_PAN_; i++) {
      selectedOnly_[i].setSelected(false);
    }
    return;
  }    
  
  /***************************************************************************
  **
  ** Enable or disable the panel
  */

  public void enableControls(boolean enabled) {
    controlsEnabled_ = enabled;
    if (!enabled) {
      disableControls();
    } else {
      syncToState();
    }
    return;
  }    
         
  /***************************************************************************
  **
  ** Let us know we have a model
  */

  public void haveAModel(boolean haveIt) {
    haveAModel_ = haveIt;
    for (int i = 0; i < NUM_PAN_; i++) {
      chooseAStart_[i].setEnabled(controlsEnabled_ && haveAModel_);
      infoPanel_[i].clear();
    }

    if (!haveAModel_) {
      currTstat_ = null;
      disableControls();
    }
    repaint();
    return;
  }
  /***************************************************************************
  **
  ** Let us know we have a selection
  */

  public void haveASelection(boolean haveIt) {
    for (int i = 0; i < NUM_PAN_; i++) {
      startAtCurrent_[i].setEnabled(haveIt);
      selectedOnly_[i].setEnabled(haveIt);
    }
    if (currTstat_ != null) {
      currTstat_.currStopUnselected = bfp_.tourStopNowUnselected();
    }
    haveSelection_ = haveIt;
    repaint();
    return;
  }
  
  /***************************************************************************
  **
  ** Enable/disable tour
  */

  private void clearTour() {
    for (int i = 0; i < NUM_PAN_; i++) {
      infoPanel_[i].clear();
    }
    revalidate();
    disableControls();
    return;
  } 
  
  /***************************************************************************
  **
  ** Give valid answer to use selected only
  */

  private boolean calcSelectedOnly() {
    return (selectedOnly_[currPan_].isEnabled() && selectedOnly_[currPan_].isSelected());
  } 

  /***************************************************************************
  **
  ** Drawing routine
  */

  public void setFabricPanel(BioFabricPanel bfp) {
    bfp_ = bfp;
    return;
  }
  
  /***************************************************************************
  **
  ** install Names
  */

  public void installNames(BioFabricPanel.TourStatus tstat) {
    if (tstat == null) {
      return;       
    } 
    currTstat_ = tstat.clone();
    syncToState();
    return;
  } 
  
  /***************************************************************************
  **
  ** Enable/disable tour
  */

  private void disableControls() {
    for (int i = 0; i < NUM_PAN_; i++) {
      buttonUp_[i].setEnabled(false); 
      buttonRight_[i].setEnabled(false); 
      buttonFarRight_[i].setEnabled(false); 
      buttonLeft_[i].setEnabled(false); 
      buttonFarLeft_[i].setEnabled(false); 
      buttonDown_[i].setEnabled(false);
      buttonClear_[i].setEnabled(false); 
      buttonZoom_[i].setEnabled(false);
      buttonLinkZone_[i].setEnabled(false);
      infoPanel_[i].setEnabled(false);
      selectedOnly_[i].setEnabled(false);
    }
    repaint();
    return;
  }   

  /***************************************************************************
  **
  ** Sync to state!
  */

  private void syncToState() {
    for (int i = 0; i < NUM_PAN_; i++) {
      chooseAStart_[i].setEnabled(haveAModel_);
    }
    if (currTstat_ == null) {
      disableControls();
      return;
    }
    ResourceManager rMan = ResourceManager.getManager();
    for (int i = 0; i < NUM_PAN_; i++) {
      buttonZoom_[i].setEnabled(true);
      buttonLinkZone_[i].setEnabled(true);
      buttonClear_[i].setEnabled(true);
      infoPanel_[i].setEnabled(true);
      String nodeName = MessageFormat.format(rMan.getString("navTool.currNode"), new Object[] {currTstat_.nodeName}); 
      String linkName = MessageFormat.format(rMan.getString("navTool.currLink"), new Object[] {currTstat_.linkName});
      infoPanel_[i].installNames(nodeName, linkName);
      buttonUp_[i].setEnabled(currTstat_.upEnabled);   
      buttonRight_[i].setEnabled(currTstat_.rightEnabled);    
      buttonFarRight_[i].setEnabled(currTstat_.farRightEnabled);   
      buttonLeft_[i].setEnabled(currTstat_.leftEnabled);   
      buttonFarLeft_[i].setEnabled(currTstat_.farLeftEnabled);   
      buttonDown_[i].setEnabled(currTstat_.downEnabled);
      selectedOnly_[i].setEnabled(haveSelection_);
      startAtCurrent_[i].setEnabled(haveSelection_);
    }
    revalidate();
    repaint();
    return;
  }
  
  
  /***************************************************************************
  **
  ** Create pairs of buttons
  */

  private void buildButtons() {
    infoPanel_ = new InfoPanel[NUM_PAN_];
    startAtCurrent_ = new FixedJButton[NUM_PAN_];    
    chooseAStart_ = new FixedJButton[NUM_PAN_] ;      
    
    buttonUp_ = new FixedJButton[NUM_PAN_];
    buttonRight_ = new FixedJButton[NUM_PAN_];    
    buttonFarRight_ = new FixedJButton[NUM_PAN_];      
    buttonLeft_ = new FixedJButton[NUM_PAN_];
    buttonFarLeft_ = new FixedJButton[NUM_PAN_]; 
    buttonDown_ = new FixedJButton[NUM_PAN_]; 
    buttonZoom_ = new FixedJButton[NUM_PAN_];
    buttonLinkZone_ = new FixedJButton[NUM_PAN_];
    buttonClear_ = new FixedJButton[NUM_PAN_];  
    selectedOnly_ = new JCheckBox[NUM_PAN_];
    return;
  }

  /***************************************************************************
  **
  ** Build correct buttons
  */

  private FixedJButton buildZeButton(int i, String label) {
    if (i == SMALL_) {
      return (FixedJButton.miniFactory(label, 12));
    } else {
      return (new FixedJButton(label));
    }  
  }   
  
  /***************************************************************************
  **
  ** Create a panel
  */

  private JPanel fillPanel(int i) {
  
    JPanel retval = new JPanel();
    retval.setBackground(Color.WHITE);
    ResourceManager rMan = ResourceManager.getManager();
    retval.setLayout(new GridLayout(1,1));
    
    String zeLabel = rMan.getString("navTool.startAtCurrent");
    startAtCurrent_[i] = buildZeButton(i, zeLabel);
    startAtCurrent_[i].setToolTipText(zeLabel);
    startAtCurrent_[i].addActionListener(new StartFromSel());
    startAtCurrent_[i].setEnabled(false);
     
    zeLabel = rMan.getString("navTool.chooseAStart");
    chooseAStart_[i] = buildZeButton(i, zeLabel);
    chooseAStart_[i].setToolTipText(zeLabel);
    chooseAStart_[i].addActionListener(new ChooseAStart());
    chooseAStart_[i].setEnabled(false);

    zeLabel = rMan.getString("navTool.up");
    buttonUp_[i] = buildZeButton(i, zeLabel);
    buttonUp_[i].setToolTipText(zeLabel);
    buttonUp_[i].addActionListener(new GoUp());
    
    zeLabel = rMan.getString("navTool.right");
    buttonRight_[i] = buildZeButton(i, zeLabel);
    buttonRight_[i].setToolTipText(zeLabel);
    buttonRight_[i].addActionListener(new GoRight());
    
    zeLabel = ">>";
    buttonFarRight_[i] = buildZeButton(i, ">>");
    buttonFarRight_[i].setToolTipText(zeLabel);
    buttonFarRight_[i].addActionListener(new GoFarRight());
    
    zeLabel = rMan.getString("navTool.left");
    buttonLeft_[i] = buildZeButton(i, zeLabel);
    buttonLeft_[i].setToolTipText(zeLabel);
    buttonLeft_[i].addActionListener(new GoLeft());
    
    zeLabel = "<<";
    buttonFarLeft_[i] = buildZeButton(i, "<<");
    buttonFarLeft_[i].setToolTipText(zeLabel);
    buttonFarLeft_[i].addActionListener(new GoFarLeft());
 
    zeLabel = rMan.getString("navTool.down");
    buttonDown_[i] = buildZeButton(i, zeLabel);
    buttonDown_[i].setToolTipText(zeLabel);
    buttonDown_[i].addActionListener(new GoDown());
  
    zeLabel = rMan.getString("navTool.clear");
    buttonClear_[i] = buildZeButton(i, zeLabel);
    buttonClear_[i].setToolTipText(zeLabel);
    buttonClear_[i].addActionListener(new ClearTour());
   
    zeLabel = rMan.getString("navTool.zoom");
    buttonZoom_[i] = buildZeButton(i, zeLabel);
    buttonZoom_[i].setToolTipText(zeLabel);
    buttonZoom_[i].addActionListener(new Zoom());
    
    zeLabel = rMan.getString("navTool.dropZone");
    buttonLinkZone_[i] = buildZeButton(i, zeLabel);
    buttonLinkZone_[i].setToolTipText(zeLabel);
    buttonLinkZone_[i].addActionListener(new ToDrain());
    
    selectedOnly_[i] = new JCheckBox(rMan.getString("navTool.skip"));
    selectedOnly_[i].setOpaque(true);
    selectedOnly_[i].setBackground(Color.white);
    selectedOnly_[i].addActionListener(new SelOnly());

    selectedOnly_[i].setEnabled(false);
          
    Box cbuttonPanel = Box.createHorizontalBox();
    cbuttonPanel.add(Box.createHorizontalGlue()); 
    cbuttonPanel.add(startAtCurrent_[i]);        
    cbuttonPanel.add(Box.createHorizontalStrut(5));
    cbuttonPanel.add(chooseAStart_[i]);        
    cbuttonPanel.add(Box.createHorizontalGlue());
       
    Box infoPanel = Box.createHorizontalBox();
    infoPanel.add(Box.createHorizontalStrut(5));
    infoPanel_[i] = new InfoPanel(true, 50, 100, true, false);
    infoPanel.add(infoPanel_[i]);
    infoPanel.add(Box.createHorizontalStrut(5));

    Box dbuttonPanel = Box.createHorizontalBox();
    dbuttonPanel.add(Box.createHorizontalGlue()); 
    dbuttonPanel.add(buttonUp_[i]);
    dbuttonPanel.add(Box.createHorizontalGlue());
    
    Box ebuttonPanel = Box.createHorizontalBox();
    ebuttonPanel.add(Box.createHorizontalGlue());
    ebuttonPanel.add(buttonFarLeft_[i]);
    ebuttonPanel.add(Box.createHorizontalStrut(5));
    ebuttonPanel.add(buttonLeft_[i]);
    ebuttonPanel.add(Box.createHorizontalStrut(5));
    ebuttonPanel.add(buttonRight_[i]);
    ebuttonPanel.add(Box.createHorizontalStrut(5));
    ebuttonPanel.add(buttonFarRight_[i]);
    ebuttonPanel.add(Box.createHorizontalGlue());

    Box fbuttonPanel = Box.createHorizontalBox();
    fbuttonPanel.add(Box.createHorizontalGlue()); 
    fbuttonPanel.add(buttonDown_[i]);
    fbuttonPanel.add(Box.createHorizontalGlue());
    
    Box hbuttonPanel = Box.createHorizontalBox();
    hbuttonPanel.add(Box.createHorizontalGlue()); 
    hbuttonPanel.add(buttonClear_[i]);        
    hbuttonPanel.add(Box.createHorizontalStrut(5));
    hbuttonPanel.add(buttonZoom_[i]);  
    hbuttonPanel.add(Box.createHorizontalStrut(5));
    hbuttonPanel.add(buttonLinkZone_[i]);  
    hbuttonPanel.add(Box.createHorizontalGlue());
 
    Box ibuttonPanel = Box.createHorizontalBox();
    ibuttonPanel.add(Box.createHorizontalGlue()); 
    ibuttonPanel.add(selectedOnly_[i]);
    ibuttonPanel.add(Box.createHorizontalGlue());    
    
    Box gbuttonPanel = Box.createVerticalBox();
    gbuttonPanel.add(Box.createVerticalGlue());
    gbuttonPanel.add(cbuttonPanel); 
   // gbuttonPanel.add(Box.createVerticalStrut(1));
    gbuttonPanel.add(Box.createVerticalGlue());
    gbuttonPanel.add(infoPanel);
  //  gbuttonPanel.add(Box.createVerticalStrut(1));
    gbuttonPanel.add(Box.createVerticalGlue());
    gbuttonPanel.add(dbuttonPanel);
 //   gbuttonPanel.add(Box.createVerticalStrut(1));
    gbuttonPanel.add(Box.createVerticalGlue());
    gbuttonPanel.add(dbuttonPanel);
  //  gbuttonPanel.add(Box.createVerticalStrut(1));
    gbuttonPanel.add(Box.createVerticalGlue());
    gbuttonPanel.add(ebuttonPanel);
 //   gbuttonPanel.add(Box.createVerticalStrut(1));
    gbuttonPanel.add(Box.createVerticalGlue());
    gbuttonPanel.add(fbuttonPanel);
  //  gbuttonPanel.add(Box.createVerticalStrut(1));
    gbuttonPanel.add(Box.createVerticalGlue());
    gbuttonPanel.add(hbuttonPanel);
 //   gbuttonPanel.add(Box.createVerticalStrut(1));
    gbuttonPanel.add(Box.createVerticalGlue());
    gbuttonPanel.add(ibuttonPanel);
 //   gbuttonPanel.add(Box.createVerticalStrut(1));
    gbuttonPanel.add(Box.createVerticalGlue());
      
    retval.add(gbuttonPanel);
    return (retval);
  }

  
  /***************************************************************************
  **
  ** Shared listener:
  */

  public class StartFromSel implements ActionListener {  
    public void actionPerformed(ActionEvent ev) {
      try {
        BioFabricPanel.TourStatus tstat = bfp_.startTourFromSelection(calcSelectedOnly());
        installNames(tstat);
      } catch (Exception ex) {
        ExceptionHandler.getHandler().displayException(ex);
      }
    }
  }
  
  /***************************************************************************
  **
  ** Shared listener:
  */

  public class ChooseAStart implements ActionListener {  
    public void actionPerformed(ActionEvent ev) {
      try {
        bfp_.setToCollectTourStart(calcSelectedOnly());
      } catch (Exception ex) {
        ExceptionHandler.getHandler().displayException(ex);
      }
    }
  }
  
  /***************************************************************************
  **
  ** Shared listener:
  */

  public class GoUp implements ActionListener {  
    public void actionPerformed(ActionEvent ev) {
      try {
        BioFabricPanel.TourStatus tstat = bfp_.goUp(calcSelectedOnly());
        installNames(tstat);
      } catch (Exception ex) {
        ExceptionHandler.getHandler().displayException(ex);
      }
    }
  }

  /***************************************************************************
  **
  ** Shared listener:
  */

  public class GoRight implements ActionListener { 
    public void actionPerformed(ActionEvent ev) {
      try {
        BioFabricPanel.TourStatus tstat = bfp_.goRight(calcSelectedOnly());
        installNames(tstat);
      } catch (Exception ex) {
        ExceptionHandler.getHandler().displayException(ex);
      }
    }
  }
  
  /***************************************************************************
  **
  ** Shared listener:
  */

  public class GoFarRight implements ActionListener {  
    public void actionPerformed(ActionEvent ev) {
      try {
        BioFabricPanel.TourStatus tstat = bfp_.goFarRight(calcSelectedOnly());
        installNames(tstat);
      } catch (Exception ex) {
        ExceptionHandler.getHandler().displayException(ex);
      }
    }
  }
      
  /***************************************************************************
  **
  ** Shared listener:
  */

  public class GoLeft implements ActionListener {  
    public void actionPerformed(ActionEvent ev) {
      try {
        BioFabricPanel.TourStatus tstat = bfp_.goLeft(calcSelectedOnly());
        installNames(tstat);
      } catch (Exception ex) {
        ExceptionHandler.getHandler().displayException(ex);
      }
    }
  }

  /***************************************************************************
  **
  ** Shared listener:
  */

  public class GoFarLeft implements ActionListener {  
    public void actionPerformed(ActionEvent ev) {
      try {
        BioFabricPanel.TourStatus tstat = bfp_.goFarLeft(calcSelectedOnly());
        installNames(tstat);
      } catch (Exception ex) {
        ExceptionHandler.getHandler().displayException(ex);
      }
    }
  }
           
  /***************************************************************************
  **
  ** Shared listener:
  */

  public class GoDown implements ActionListener {    
    public void actionPerformed(ActionEvent ev) {
      try {
        BioFabricPanel.TourStatus tstat = bfp_.goDown(calcSelectedOnly());
        installNames(tstat);
      } catch (Exception ex) {
        ExceptionHandler.getHandler().displayException(ex);
      }
    }
  } 
    
  /***************************************************************************
  **
  ** Shared listener:
  */

  public class ClearTour implements ActionListener {     
    public void actionPerformed(ActionEvent ev) {
      try {
        bfp_.clearTour();
        clearTour();
      } catch (Exception ex) {
        ExceptionHandler.getHandler().displayException(ex);
      }
    }
  }

  /***************************************************************************
  **
  ** Shared listener:
  */

  public class Zoom implements ActionListener {
    public void actionPerformed(ActionEvent ev) {
      try {
        bfp_.zoomToTourStop();
      } catch (Exception ex) {
        ExceptionHandler.getHandler().displayException(ex);
      }
    }
  }
    
  /***************************************************************************
  **
  ** Shared listener:
  */

  public class ToDrain implements ActionListener {
    public void actionPerformed(ActionEvent ev) {
      try {
        BioFabricPanel.TourStatus tstat = bfp_.tourToDrainZone(calcSelectedOnly());
        installNames(tstat);
      } catch (Exception ex) {
        ExceptionHandler.getHandler().displayException(ex);
      }
    }
  }
 
  /***************************************************************************
  **
  ** Shared listener:
  */

  public class SelOnly implements ActionListener { 
    public void actionPerformed(ActionEvent ev) {
      try {
        if (doSOReset_) {
          return;
        }
        if ((currTstat_ != null) && (currTstat_.currStopUnselected) && selectedOnly_[currPan_].isSelected()) {
          ResourceManager rMan = ResourceManager.getManager();    
          JOptionPane.showMessageDialog(topWindow_, rMan.getString("navTool.currLocUnselected"),
                                        rMan.getString("navTool.currLocUnselectedTitle"),
                                        JOptionPane.ERROR_MESSAGE);
          doSOReset_ = true;
          selectedOnly_[currPan_].setSelected(false);
          doSOReset_ = false;
          return;
        }
        boolean newVal = selectedOnly_[currPan_].isSelected();
        BioFabricPanel.TourStatus tstat = bfp_.getTourDirections(newVal);
        doSOReset_ = true;
        for (int i = 0; i < NUM_PAN_; i++) {
          if (i != currPan_) {
            selectedOnly_[i].setSelected(newVal);
          }
        }
        doSOReset_ = false;
        
        installNames(tstat);
      } catch (Exception ex) {
        ExceptionHandler.getHandler().displayException(ex);
      }
    }
  } 

  /***************************************************************************
  **
  ** Wrap a toggled panel around the basic tool:
  */

  public static class LabeledFabricNavTool extends JPanel {

    private FabricNavTool fnt_;
    private JPanel livePan_;
    private CardLayout clay_;
    private boolean collapsed_;
    
    private static final long serialVersionUID = 1L;

    public LabeledFabricNavTool(JFrame topWindow, Font labelFont) {
      fnt_ = new FabricNavTool(topWindow);
      livePan_ = new JPanel();
      livePan_.setLayout(new BorderLayout());

      livePan_.setBorder(new LineBorder(Color.black, 2));
      JLabel navLab = new JLabel(ResourceManager.getManager().getString("biofabric.tour"));
      navLab.setBorder(new EmptyBorder(0, 5, 0, 0));
      navLab.setOpaque(true);
      navLab.setBackground(Color.white);
      navLab.setFont(labelFont);
      livePan_.add(navLab, BorderLayout.NORTH);
      livePan_.add(fnt_, BorderLayout.CENTER);
    
      clay_ = new CardLayout();
      setLayout(clay_);
      add(livePan_, "live");
      add(new JPanel(), "blank");
      clay_.show(this, "live");
      collapsed_ = false;
    }
    
    /***************************************************************************
    **
    ** get actual tool
    */
  
    public FabricNavTool getFabricNavTool() {
      return (fnt_);
    }     
    
    /***************************************************************************
    **
    ** Set to blank or populated
    */
  
    public void setToBlank(boolean val) {
      clay_.show(this, (val) ? "blank" : "live");
      collapsed_ = val;
      return;
    } 
    
    /***************************************************************************
    **
    ** Sizing
    */
    
    @Override
    public Dimension getPreferredSize() {
      if (collapsed_) {
        return (new Dimension(0, 0));    
      } else {
        return (livePan_.getPreferredSize());
      } 
    }
  
    @Override
    public Dimension getMinimumSize() {
      if (collapsed_) {
        return (new Dimension(0, 0));    
      } else {
        return (livePan_.getMinimumSize());
      } 
    }
    
    @Override
    public Dimension getMaximumSize() {
      if (collapsed_) {
        return (new Dimension(0, 0));    
      } else {
        return (livePan_.getMaximumSize());
      } 
    }
  }
}
```
