## Supplementary material for "BioFabric Visualization of Network Alignments": BioFabric Source Code: InfoPanel.java

```
/*
**    Copyright (C) 2003-2017 Institute for Systems Biology 
**                            Seattle, Washington, USA. 
**
**    This library is free software; you can redistribute it and/or
**    modify it under the terms of the GNU Lesser General Public
**    License as published by the Free Software Foundation; either
**    version 2.1 of the License, or (at your option) any later version.
**
**    This library is distributed in the hope that it will be useful,
**    but WITHOUT ANY WARRANTY; without even the implied warranty of
**    MERCHANTABILITY or FITNESS FOR A PARTICULAR PURPOSE.  See the GNU
**    Lesser General Public License for more details.
**
**    You should have received a copy of the GNU Lesser General Public
**    License along with this library; if not, write to the Free Software
**    Foundation, Inc., 59 Temple Place, Suite 330, Boston, MA  02111-1307  USA
*/

package org.systemsbiology.biofabric.ui.display;

import java.awt.Color;
import java.awt.Dimension;
import java.awt.Font;
import java.awt.Graphics;
import java.awt.Graphics2D;
import java.awt.RenderingHints;
import java.awt.font.FontRenderContext;
import java.awt.geom.AffineTransform;
import java.awt.geom.Rectangle2D;

import javax.swing.JPanel;

/****************************************************************************
**
** This is a scalable text panel
*/

public class InfoPanel extends JPanel {
    
  ////////////////////////////////////////////////////////////////////////////
  //
  // PRIVATE INSTANCE MEMBERS
  //
  ////////////////////////////////////////////////////////////////////////////

  private static final long serialVersionUID = 1L;
  
  private Font myFont_;
  private boolean twoLines_;
  private String text1_;
  private String text2_;
  private int minHeight_;
  private int minWidth_;
  private boolean centered_;
  private boolean pad_;
	
  ////////////////////////////////////////////////////////////////////////////
  //
  // PUBLIC CONSTRUCTORS
  //
  ////////////////////////////////////////////////////////////////////////////

  /***************************************************************************
  **
  ** Constructor
  */
  
  public InfoPanel(boolean twoLines, int minHeight, int minWidth, boolean centered, boolean pad) {
    setBackground(new Color(255, 255, 255));
    myFont_ = new Font("SansSerif", Font.BOLD, 12);
    twoLines_ = twoLines;
    text1_ = "";
    if (twoLines_) {
      text2_ = "";
    }
    minHeight_ = minHeight;
    minWidth_ = minWidth;
    centered_ = centered;
    pad_ = pad;
  }
 
  ////////////////////////////////////////////////////////////////////////////
  //
  // METHODS
  //
  ////////////////////////////////////////////////////////////////////////////  
  
  /***************************************************************************
  **
  ** Clear text
  */
  
  public void clear() {
    text1_ = "";
    if (twoLines_) {
      text2_ = "";
    }
    return;
  }

  /***************************************************************************
  **
  ** Install text
  */
  
  public void installName(String text1) {
  	if (twoLines_) {
  		throw new IllegalStateException();
  	}
    text1_ = text1; 
    return;
  }
   
  /***************************************************************************
  **
  ** Install text
  */
  
  public void installNames(String text1, String text2) {
  	if (!twoLines_) {
  		throw new IllegalStateException();
  	}
    text1_ = text1; 
    text2_ = text2;
    return;
  }
    
  /***************************************************************************
  **
  ** The usual dimension methods
  */
  
  @Override
  public Dimension getPreferredSize() {
    return (new Dimension(200, minHeight_));    
  }

  @Override
  public Dimension getMinimumSize() {
    return (new Dimension(minWidth_, minHeight_));    
  }
  
  @Override
  public Dimension getMaximumSize() {
    return (new Dimension(4000, 200));    
  }
  
  /***************************************************************************
  **
  ** Paints
  */
  
  public void paintComponent(Graphics g) {
    super.paintComponent(g);
    Graphics2D g2 = (Graphics2D)g;
    g2.setRenderingHint(RenderingHints.KEY_ANTIALIASING, RenderingHints.VALUE_ANTIALIAS_ON);
    g2.setRenderingHint(RenderingHints.KEY_FRACTIONALMETRICS, RenderingHints.VALUE_FRACTIONALMETRICS_ON);
    AffineTransform saveTrans = g2.getTransform();
    g2.setFont(myFont_);
    g2.setPaint(Color.BLACK);
    FontRenderContext frc = g2.getFontRenderContext();
    
    double maxFactor = 0.0;
    Rectangle2D boundsL = null;
    
    Rectangle2D boundsN = myFont_.getStringBounds(text1_, frc);
    if (pad_) {
      boundsN.setRect(boundsN.getX(), boundsN.getY(), boundsN.getWidth() + 16, boundsN.getHeight()); 
    }
    double reduceN = boundsN.getWidth() / this.getWidth();
    
    if (twoLines_) {
	    boundsL = myFont_.getStringBounds(text2_, frc);
	    if (pad_) {
	      boundsL.setRect(boundsL.getX(), boundsL.getY(), boundsL.getWidth() + 16, boundsL.getHeight()); 
	    }
	    double reduceL = boundsL.getWidth() / this.getWidth();
	    maxFactor = Math.max(reduceN, reduceL);
    } else {
    	maxFactor = reduceN;
    }
  
    float useFactor = 1.0F;   
    if (maxFactor > 1.0) {
      double rmf = 1.0 / maxFactor;
      useFactor = (float)maxFactor;
      g2.translate(this.getWidth() / 2.0, this.getHeight() / 2.0);
      g2.scale(rmf, rmf);
      g2.translate(-this.getWidth() / 2.0, -this.getHeight() / 2.0);
    }

    float textX1;
    float textY1;
    float textX2 = 0.0F;
    float textY2 = 0.0F;
    
    if (centered_) {    
      textX1 = (getWidth() / 2.0F) - ((float)boundsN.getWidth() / 2.0F);
      textY1 = (twoLines_) ? (getHeight() / 2.0F) - (useFactor * getHeight() / 6.0F) + ((float)(boundsN.getHeight()) / 3.0F)
    		                        : getHeight() - (useFactor * getHeight() / 6.0F) + ((float)(boundsN.getHeight()) / 3.0F);
      if (twoLines_) {
        textX2 = (getWidth() / 2.0F) - ((float)boundsL.getWidth() / 2.0F); 
        textY2 = (getHeight() / 2.0F) + (useFactor * getHeight() / 6.0F) + ((float)(boundsL.getHeight()) / 3.0F);
      }
    } else {
    	textX1 = (getWidth() / 2.0F) - ((useFactor * getWidth()) / 2.0F);
      textY1 = (twoLines_) ? (getHeight() / 2.0F) - (useFactor * getHeight() / 6.0F) + ((float)(boundsN.getHeight()) / 3.0F)
    		                   : (getHeight() / 2.0F) + ((float)(boundsN.getHeight()) / 5.0F);

      if (twoLines_) {
        textX2 = (getWidth() / 2.0F) - ((useFactor * getWidth()) / 2.0F);
        textY2 = (getHeight() / 2.0F) + (useFactor * getHeight() / 6.0F) + ((float)(boundsL.getHeight()) / 3.0F);
      }  	
    }
   
    g2.drawString(text1_, textX1, textY1);
    if (twoLines_) {
      g2.drawString(text2_, textX2, textY2);
    }
    
    g2.setTransform(saveTrans);
    return;
  }    
}
```
