## Supplementary material for "BioFabric Visualization of Network Alignments": BioFabric Source Code: MouseOverView.java

```
/*
**    Copyright (C) 2003-2018 Institute for Systems Biology 
**                            Seattle, Washington, USA. 
**
**    This library is free software; you can redistribute it and/or
**    modify it under the terms of the GNU Lesser General Public
**    License as published by the Free Software Foundation; either
**    version 2.1 of the License, or (at your option) any later version.
**
**    This library is distributed in the hope that it will be useful,
**    but WITHOUT ANY WARRANTY; without even the implied warranty of
**    MERCHANTABILITY or FITNESS FOR A PARTICULAR PURPOSE.  See the GNU
**    Lesser General Public License for more details.
**
**    You should have received a copy of the GNU Lesser General Public
**    License along with this library; if not, write to the Free Software
**    Foundation, Inc., 59 Temple Place, Suite 330, Boston, MA  02111-1307  USA
*/

package org.systemsbiology.biofabric.ui.display;

import javax.swing.JPanel;


/****************************************************************************
**
** This panel gives a view for mouseovers
*/

public class MouseOverView {
  
  ////////////////////////////////////////////////////////////////////////////
  //
  // PRIVATE CONSTANTS
  //
  //////////////////////////////////////////////////////////////////////////// 
    
  ////////////////////////////////////////////////////////////////////////////
  //
  // PUBLIC CONSTANTS
  //
  //////////////////////////////////////////////////////////////////////////// 
  
  ////////////////////////////////////////////////////////////////////////////
  //
  // PRIVATE INSTANCE MEMBERS
  //
  ////////////////////////////////////////////////////////////////////////////
   
  private MouseOverViewPanel pan1_;
  private MouseOverViewPanel pan2_;
  private boolean isAlive_;

  ////////////////////////////////////////////////////////////////////////////
  //
  // PUBLIC CONSTRUCTORS
  //
  ////////////////////////////////////////////////////////////////////////////

  /***************************************************************************
  **
  ** Constructor
  */

  public MouseOverView() {
    pan1_ = new MouseOverViewPanel();
    pan2_ = new MouseOverViewPanel();
    isAlive_ = false;
  }

  ////////////////////////////////////////////////////////////////////////////
  //
  // PUBLIC METHODS
  //
  ////////////////////////////////////////////////////////////////////////////

  /***************************************************************************
  **
  ** Show or hide the view
  */

  public void setIsAlive(boolean alive) {
  	isAlive_ = alive;
    return;
  }
  
  /***************************************************************************
  **
  ** Set file locations to use for image files
  */

  public void setFileLocations(String path, String filePrefix, String fileSuffix)  {
  	pan1_.setFileLocations(path, filePrefix, fileSuffix);
    pan2_.setFileLocations(path, filePrefix, fileSuffix);
    return;
  } 
 
  /***************************************************************************
  **
  ** Show or hide the view
  */

  public void showView(boolean enabled) {
  	pan1_.showView(enabled);
    pan2_.showView(enabled);
    return;
  }
   
  /***************************************************************************
  **
  ** SGet the view
  */

  public JPanel getPanel(int num) {
  	return ((num == 0) ? pan1_ : pan2_);
  }

  /***************************************************************************
  ** 
  ** Install a model
  */

  public void showForNode(BioFabricPanel.MouseLocInfo mlo) {
  	if (!isAlive_) {
  		return;
  	}
  	pan1_.showForNode(mlo.zoneDesc.equals("<none>") ? null : mlo.zoneDesc);
  	if (mlo.linkDesc.equals("<none>")) {
  	  pan2_.showForNode(null);
  	} else {
  		String other = mlo.zoneDesc.equals(mlo.linkSrcDesc) ? mlo.linkTrgDesc : mlo.linkSrcDesc;
  		pan2_.showForNode(other);
  	}
  	return;
  }
}
```
