## Supplementary material for "BioFabric Visualization of Network Alignments": BioFabric Source Code: MouseOverViewPanel.java

```
/*
**    Copyright (C) 2003-2018 Institute for Systems Biology 
**                            Seattle, Washington, USA. 
**
**    This library is free software; you can redistribute it and/or
**    modify it under the terms of the GNU Lesser General Public
**    License as published by the Free Software Foundation; either
**    version 2.1 of the License, or (at your option) any later version.
**
**    This library is distributed in the hope that it will be useful,
**    but WITHOUT ANY WARRANTY; without even the implied warranty of
**    MERCHANTABILITY or FITNESS FOR A PARTICULAR PURPOSE.  See the GNU
**    Lesser General Public License for more details.
**
**    You should have received a copy of the GNU Lesser General Public
**    License along with this library; if not, write to the Free Software
**    Foundation, Inc., 59 Temple Place, Suite 330, Boston, MA  02111-1307  USA
*/

package org.systemsbiology.biofabric.ui.display;

import java.awt.CardLayout;
import java.awt.Color;
import java.awt.Dimension;
import java.awt.Graphics;
import java.awt.Graphics2D;
import java.awt.Point;
import java.awt.RenderingHints;
import java.awt.image.BufferedImage;
import java.io.File;
import java.io.FileInputStream;
import java.io.IOException;
import java.util.HashMap;
import java.util.Iterator;

import javax.imageio.ImageIO;
import javax.imageio.ImageReader;
import javax.imageio.stream.ImageInputStream;
import javax.swing.JPanel;

/****************************************************************************
**
** This panel gives a view for mouseovers
*/

public class MouseOverViewPanel extends JPanel {
  
  ////////////////////////////////////////////////////////////////////////////
  //
  // PRIVATE CONSTANTS
  //
  //////////////////////////////////////////////////////////////////////////// 
    
  ////////////////////////////////////////////////////////////////////////////
  //
  // PUBLIC CONSTANTS
  //
  //////////////////////////////////////////////////////////////////////////// 
  
  ////////////////////////////////////////////////////////////////////////////
  //
  // PRIVATE INSTANCE MEMBERS
  //
  ////////////////////////////////////////////////////////////////////////////
   
  private Dimension currSize_;
  private HashMap<String, BufferedImage> imgs_;
  private BufferedImage img_;
  private BufferedImage scaledImg_;
  private Point scaledImgOrigin_;
  private CardLayout myCard_;
  private ImagePanel pPan_;
  private String path_ ;
  private String filePrefix_;
  private String fileSuffix_ ;
  
  private static final long serialVersionUID = 1L;
  
  ////////////////////////////////////////////////////////////////////////////
  //
  // PUBLIC CONSTRUCTORS
  //
  ////////////////////////////////////////////////////////////////////////////

  /***************************************************************************
  **
  ** Constructor
  */

  public MouseOverViewPanel() {
    setBackground(Color.white);
    imgs_ = new HashMap<String, BufferedImage>();
    scaledImg_ = null;
    scaledImgOrigin_ = new Point(0, 0);
    img_ = null;
    myCard_ = new CardLayout();
    setLayout(myCard_);
    
    pPan_ = new ImagePanel();
    add(pPan_, "theView");
    JPanel blankPanel = new JPanel();
    blankPanel.setBackground(Color.white);
    add(blankPanel, "Hiding");   
  }

  ////////////////////////////////////////////////////////////////////////////
  //
  // PUBLIC METHODS
  //
  ////////////////////////////////////////////////////////////////////////////

  /***************************************************************************
  **
  ** Sizing
  */

  @Override
  public Dimension getPreferredSize() {
    return (new Dimension(800, 100));    
  }
  
  @Override
  public Dimension getMinimumSize() {
    return (new Dimension(10, 10));  
  }
  
  @Override
  public Dimension getMaximumSize() {
    return (new Dimension(4000, 340));    
  }
    
  /***************************************************************************
  **
  ** Handle size change
  */

  @Override
  public void setBounds(int x, int y, int width, int height) {
    super.setBounds(x, y, width, height);
    currSize_ = new Dimension(width, height);
    resizeImage();
    repaint();
    return;
  } 
 
  /***************************************************************************
  **
  ** Set the fragments of file to show: path + filePrefix + nodeName + fileSuffix.
  */

  public void setFileLocations(String path, String filePrefix, String fileSuffix) {
    path_ = path;
    filePrefix_ = filePrefix;
    fileSuffix_ = fileSuffix;
    return;
  }

  /***************************************************************************
  **
  ** Show or hide the view
  */

  public void showView(boolean enabled) {
    myCard_.show(this, (enabled) ? "theView" : "Hiding"); 
    return;
  }

  /***************************************************************************
  ** 
  ** Install a model
  */

  public void showForNode(String nodeName) {
  	if (nodeName == null) {
  		installImage(null);
  		return;
  	}
  	BufferedImage bi = imgs_.get(nodeName);
  	if (bi != null) {
  		installImage(bi);
  	} else {
	  	try {
	      loadAFile(nodeName);
	  	} catch (IOException ioex) {
	  		System.err.println("Handle file load failures: " + ioex.getLocalizedMessage());
	  	}
  	}
    return;
  }
 
  /***************************************************************************
  ** 
  ** Install a model
  */

  private void loadAFile(String nodeName) throws IOException {
  	File forFoo = new File(path_ + filePrefix_ + nodeName + fileSuffix_);
    BufferedImage mvi = readImageFromFile(forFoo);
    imgs_.put(nodeName, mvi);
    installImage(mvi);
    return;
  }
  
  /***************************************************************************
  ** 
  ** Install a model
  */

  private void installImage(BufferedImage img) {
    img_ = img;
    scaledImg_ = null;
    resizeImage();
    repaint();
    return;
  }
  
  /***************************************************************************
  **
  ** resize image
  */

  private void resizeImage() {
    if (img_ == null) {
      return;
    }
    if (currSize_ == null) {
      currSize_ = getSize();
    }
    //
    // Don't do this step if the overview is not currently bring displayed
    //
    if ((currSize_.width <= 0) || (currSize_.height <= 0)) {
      return;
    }
    double imgAR = (double)img_.getWidth() / (double)img_.getHeight();
    double panelAR = currSize_.getWidth() / currSize_.getHeight();
    int imgHeight;
    int imgWidth;
    int startX;
    int startY;
    if (panelAR < imgAR) { // long image, tall panel
      imgWidth = currSize_.width;
      imgHeight = (int)(imgWidth / imgAR);
      if (imgHeight == 0) {
        imgHeight = 1;
      }
      startX = 0;
      startY = (currSize_.height - imgHeight) / 2;
    } else {
      imgHeight = currSize_.height;
      imgWidth = (int)(imgHeight * imgAR);
      if (imgWidth == 0) {
        imgWidth = 1;
      }
      startX = (currSize_.width - imgWidth) / 2;
      startY = 0;
    }
    scaledImgOrigin_.setLocation(startX, startY);
    scaledImg_ = new BufferedImage(imgWidth, imgHeight, BufferedImage.TYPE_INT_RGB);
    Graphics2D g2 = scaledImg_.createGraphics();
    g2.setRenderingHint(RenderingHints.KEY_INTERPOLATION, RenderingHints.VALUE_INTERPOLATION_BICUBIC);
    g2.drawImage(img_, 0, 0, imgWidth, imgHeight, null);
    g2.dispose();
    return;
  } 
  
  /***************************************************************************
  **
  ** Read in an image.
  */
  
  public BufferedImage readImageFromFile(File readFile) throws IOException {
		FileInputStream fis = new FileInputStream(readFile);
		ImageInputStream iis = ImageIO.createImageInputStream(fis);
		Iterator<ImageReader> readers = ImageIO.getImageReaders(iis);
		if (!readers.hasNext()) {
		  throw new IOException();
		}
		ImageReader reader = readers.next();
		BufferedImage retval = ImageIO.read(iis);
		return (retval);
	}
  
  /***************************************************************************
  **
  ** Handles the drawing
  */  
  
  private class ImagePanel extends JPanel {
    private static final long serialVersionUID = 1L;
  
    /***************************************************************************
    **
    ** Constructor
    */
  
    public ImagePanel() {
      setBackground(Color.white);
    }
    
    /***************************************************************************
    **
    ** Drawing routine
    */
    
    public void paintComponent(Graphics g) {
      super.paintComponent(g);
      Graphics2D g2 = (Graphics2D)g;     
      if (img_ == null) {
        return;
      }
      if (scaledImg_ == null) {
        resizeImage();
      }
     
      g2.drawImage(scaledImg_, scaledImgOrigin_.x, scaledImgOrigin_.y, null);
      return;
    }
  }
}
```
