## Supplementary material for "BioFabric Visualization of Network Alignments": BioFabric Source Code: FabricColorGenerator.java

```
/*
**    Copyright (C) 2003-2017 Institute for Systems Biology 
**                            Seattle, Washington, USA. 
**
**    This library is free software; you can redistribute it and/or
**    modify it under the terms of the GNU Lesser General Public
**    License as published by the Free Software Foundation; either
**    version 2.1 of the License, or (at your option) any later version.
**
**    This library is distributed in the hope that it will be useful,
**    but WITHOUT ANY WARRANTY; without even the implied warranty of
**    MERCHANTABILITY or FITNESS FOR A PARTICULAR PURPOSE.  See the GNU
**    Lesser General Public License for more details.
**
**    You should have received a copy of the GNU Lesser General Public
**    License along with this library; if not, write to the Free Software
**    Foundation, Inc., 59 Temple Place, Suite 330, Boston, MA  02111-1307  USA
*/

package org.systemsbiology.biofabric.ui;

import java.awt.Color;
import java.io.IOException;
import java.io.PrintWriter;
import java.util.HashMap;
import java.util.Iterator;
import java.util.Map;

import org.xml.sax.Attributes;
import org.systemsbiology.biofabric.api.io.AttributeExtractor;
import org.systemsbiology.biofabric.api.io.Indenter;
import org.systemsbiology.biofabric.api.parser.AbstractFactoryClient;
import org.systemsbiology.biofabric.api.parser.GlueStick;
import org.systemsbiology.biofabric.db.ColorGenerator;
import org.systemsbiology.biofabric.io.FabricFactory;

/****************************************************************************
**
** Color generator
*/

public class FabricColorGenerator {
  
  ////////////////////////////////////////////////////////////////////////////
  //
  // PUBLIC CONSTANTS
  //
  ////////////////////////////////////////////////////////////////////////////

  public final static int UNCHANGED = 0;
  public final static int BRIGHTER  = 1;
  public final static int DARKER    = 2;
      
  ////////////////////////////////////////////////////////////////////////////
  //
  // PRIVATE MEMBERS
  //
  ////////////////////////////////////////////////////////////////////////////

  private ColorGenerator myColGen_;
  private HashMap<String, Color> brighter_;
  private HashMap<String, Color> darker_;

  ////////////////////////////////////////////////////////////////////////////
  //
  // PUBLIC CONSTRUCTORS
  //
  ////////////////////////////////////////////////////////////////////////////

  /***************************************************************************
  **
  ** Null constructor
  */

  public FabricColorGenerator() {
    myColGen_ = new ColorGenerator();
    brighter_ = new HashMap<String, Color>();
    darker_ = new HashMap<String, Color>();
  }

  ////////////////////////////////////////////////////////////////////////////
  //
  // PUBLIC METHODS
  //
  ////////////////////////////////////////////////////////////////////////////
  
  /***************************************************************************
  ** 
  ** Start a new model
  */

  public void newColorModel() {
    boolean useHSV = false;
    myColGen_.newColorModel();
    FabricDisplayOptions fdo = FabricDisplayOptionsManager.getMgr().getDisplayOptions();
    double light = fdo.getNodeLighterLevel() + 1.0;
    double dark = 1.0 / (fdo.getLinkDarkerLevel() + 1.0);
    float[] hsbvals = new float[3];
    int numCol = myColGen_.getNumColors();
    for (int i = 0; i < numCol; i++) {
      String gckey = myColGen_.getGeneColor(i);
      NamedColor nc = myColGen_.getNamedColor(gckey);
      int baseR = nc.color.getRed();    
      int baseG = nc.color.getGreen();  
      int baseB = nc.color.getBlue();
      if (useHSV) {
        Color.RGBtoHSB(baseR, baseG, baseB, hsbvals);
        float lb = Math.min(1.0F, hsbvals[2] * (float)light);
        brighter_.put(gckey, Color.getHSBColor(hsbvals[0], hsbvals[1], lb));
        float db = Math.min(1.0F, hsbvals[2] * (float)dark);
        darker_.put(gckey, Color.getHSBColor(hsbvals[0], hsbvals[1], db));
      } else {
      	// These three standard colors get TOO light if we apply the operation. So keep them as is:
      	if ((nc.name.indexOf("Gold") == -1) && (nc.name.indexOf("Dark Wheat") == -1) &&  (nc.name.indexOf("Pale Goldenrod") == -1)) {
	        int rb = Math.min(255, (int)Math.round(baseR * light));
	        int gb = Math.min(255, (int)Math.round(baseG * light));
	        int bb = Math.min(255, (int)Math.round(baseB * light));       
	        brighter_.put(gckey, new Color(rb, gb, bb));
      	} else {
      		brighter_.put(gckey, nc.color);
      	}
        int rd = Math.min(255, (int)Math.round(baseR * dark));
        int gd = Math.min(255, (int)Math.round(baseG * dark));
        int bd = Math.min(255, (int)Math.round(baseB * dark));
        darker_.put(gckey, new Color(rd, gd, bd));
      }
    }
  }   
  
  /***************************************************************************
  ** 
  ** Drop everything
  */

  public void dropColors() {
    myColGen_.dropColors();
    brighter_.clear();
    darker_.clear();
    return;
  }
  
  /***************************************************************************
  ** 
  ** Add color for IO
  */

  void addColorForIO(int target, NamedColor color) {
    switch (target) {
      case BRIGHTER:
        brighter_.put(color.name, color.color);
        break;
      case DARKER:
        darker_.put(color.name, color.color);
        break;
      case UNCHANGED:
      default:
        throw new IllegalArgumentException();
    }
    return;
  }
  
  /***************************************************************************
  ** 
  ** Get the modified color
  */

  public Color getModifiedColor(String colorKey, int which) {
    Color paintCol;
    switch (which) { 
      case UNCHANGED:
        NamedColor nc = myColGen_.getNamedColor(colorKey);
        paintCol = ((nc == null) || (nc.color == null)) ? null : nc.color;
        break;
      case DARKER:
        paintCol = darker_.get(colorKey);
        break;
      case BRIGHTER:
        paintCol = brighter_.get(colorKey);
        break;
      default:
        throw new IllegalArgumentException();
    }
    if (paintCol == null) {
      paintCol = Color.BLACK;
    }
    return (paintCol);
  }  

  /***************************************************************************
  **
  ** Get Ith available color
  */
  
  public String getGeneColor(int i) {
    return (myColGen_.getGeneColor(i));
  }   
  
  /***************************************************************************
  **
  ** Get number of colors
  */
  
  public int getNumColors() {
    return (myColGen_.getNumColors());
  }
 
  /***************************************************************************
  **
  ** Write the colors to XML
  **
  */
  
  public void writeXML(PrintWriter out, Indenter ind) {
    ind.indent();
    out.println("<colors>");
    ind.up();
    writeColorsForType(out, ind, "nodes", brighter_);
    writeColorsForType(out, ind, "links", darker_); 
    ind.down().indent();
    out.println("</colors>");
    return;
  }
   
  ////////////////////////////////////////////////////////////////////////////
  //
  // PRIVATE INSTANCE METHODS
  //
  ////////////////////////////////////////////////////////////////////////////
   
  /***************************************************************************
  **
  ** Write the colors to XML
  **
  */
  
  private void writeColorsForType(PrintWriter out, Indenter ind, String type, Map<String, Color> colMap) {
    ind.indent();
    out.print("<colorSet usage=\"");
    out.print(type);
    out.println("\">");
    ind.up();
    Iterator<String> colors = colMap.keySet().iterator();
    while (colors.hasNext()) {
      String key = colors.next();
      Color c = colMap.get(key);
      ind.indent();
      out.print("<color ");
      out.print("color=\"");
      out.print(key);
      out.print("\" r=\"");
      out.print(c.getRed());
      out.print("\" g=\"");
      out.print(c.getGreen());
      out.print("\" b=\"");
      out.print(c.getBlue());      
      out.println("\" />");
    }
    ind.down().indent();
    out.println("</colorSet>");
    return;
  }
  
  /***************************************************************************
  **
  ** For XML I/O
  */  
      
  public static class ColorSetWorker extends AbstractFactoryClient {
    
    public ColorSetWorker(FabricFactory.FactoryWhiteboard board) {
      super(board);
      FabricFactory.FactoryWhiteboard whiteboard = (FabricFactory.FactoryWhiteboard)sharedWhiteboard_;   
      myKeys_.add("colorSet");
      installWorker(new ColorWorker(whiteboard), new MyColorGlue());
    }
    
    protected Object localProcessElement(String elemName, Attributes attrs) throws IOException {
      Object retval = null;
      if (elemName.equals("colorSet")) {
        String target = AttributeExtractor.extractAttribute(elemName, attrs, "colorSet", "usage", true);
        int targVal;
        if (target.equals("nodes")) {
          targVal = BRIGHTER;
        } else if (target.equals("links")) {
          targVal = DARKER;
        } else {
          throw new IOException();
        }
        FabricFactory.FactoryWhiteboard board = (FabricFactory.FactoryWhiteboard)this.sharedWhiteboard_;
        if (board.colTarg == UNCHANGED) {
          board.fcg = new FabricColorGenerator();          
        }
        board.colTarg = targVal;
        retval = board.fcg;
      }
      return (retval);     
    }  
  }
  
  public static class MyColorGlue implements GlueStick {
    public Object glueKidToParent(Object kidObj, AbstractFactoryClient parentWorker, 
                                  Object optionalArgs) throws IOException {
      FabricFactory.FactoryWhiteboard board = (FabricFactory.FactoryWhiteboard)optionalArgs;
      board.fcg.addColorForIO(board.colTarg, board.currCol);
      return (null);
    }
  }
  
  /***************************************************************************
  **
  ** For XML I/O
  */  
      
  public static class ColorWorker extends AbstractFactoryClient {
    
    public ColorWorker(FabricFactory.FactoryWhiteboard board) {
      super(board);
      myKeys_.add("color");
    }
    
    protected Object localProcessElement(String elemName, Attributes attrs) throws IOException {
      Object retval = null;
      FabricFactory.FactoryWhiteboard board = (FabricFactory.FactoryWhiteboard)this.sharedWhiteboard_;
      board.currCol = buildFromXML(elemName, attrs);
      retval = board.currCol;
      return (retval);     
    }  
    
    private NamedColor buildFromXML(String elemName, Attributes attrs) throws IOException {  
      String name = AttributeExtractor.extractAttribute(elemName, attrs, "color", "color", true);
      String rStr = AttributeExtractor.extractAttribute(elemName, attrs, "color", "r", true);
      String gStr = AttributeExtractor.extractAttribute(elemName, attrs, "color", "g", true);
      String bStr = AttributeExtractor.extractAttribute(elemName, attrs, "color", "b", true);
      NamedColor retval;
      try {
        int r = Integer.valueOf(rStr).intValue();
        int g = Integer.valueOf(gStr).intValue();
        int b = Integer.valueOf(bStr).intValue();
        if (((r < 0) || (r > 255)) || ((g < 0) || (g > 255)) || ((b < 0) || (b > 255))) {
          throw new IOException();
        }
        retval = new NamedColor(name, new Color(r, g, b), name);
      } catch (NumberFormatException nfex) {
        throw new IOException();
      }
      return (retval);
    }
  } 
}
```
