## Supplementary material for "BioFabric Visualization of Network Alignments": BioFabric Source Code: FabricDisplayOptions.java

```
/*
**    Copyright (C) 2003-2012 Institute for Systems Biology 
**                            Seattle, Washington, USA. 
**
**    This library is free software; you can redistribute it and/or
**    modify it under the terms of the GNU Lesser General Public
**    License as published by the Free Software Foundation; either
**    version 2.1 of the License, or (at your option) any later version.
**
**    This library is distributed in the hope that it will be useful,
**    but WITHOUT ANY WARRANTY; without even the implied warranty of
**    MERCHANTABILITY or FITNESS FOR A PARTICULAR PURPOSE.  See the GNU
**    Lesser General Public License for more details.
**
**    You should have received a copy of the GNU Lesser General Public
**    License along with this library; if not, write to the Free Software
**    Foundation, Inc., 59 Temple Place, Suite 330, Boston, MA  02111-1307  USA
*/

package org.systemsbiology.biofabric.ui;

import java.io.IOException;
import java.io.PrintWriter;

import org.xml.sax.Attributes;
import org.systemsbiology.biofabric.api.io.AttributeExtractor;
import org.systemsbiology.biofabric.api.io.CharacterEntityMapper;
import org.systemsbiology.biofabric.api.io.Indenter;
import org.systemsbiology.biofabric.api.parser.AbstractFactoryClient;
import org.systemsbiology.biofabric.io.FabricFactory;


/****************************************************************************
**
** A class to specify display options
*/

public class FabricDisplayOptions implements Cloneable {
  
  ////////////////////////////////////////////////////////////////////////////    
  //
  // PUBLIC CONSTANTS
  //
  //////////////////////////////////////////////////////////////////////////// 
  
  public static final String NODE_NAME_PLACEHOLDER = "%_BIOFABRIC_NODE_NAME_%";
  
  public static final String LINK_SRC_PLACEHOLDER = "%_BIOFABRIC_LINK_SRC_%";
  public static final String LINK_TRG_PLACEHOLDER = "%_BIOFABRIC_LINK_TRG_%";
  public static final String LINK_REL_PLACEHOLDER = "%_BIOFABRIC_LINK_REL_%";
  
  ////////////////////////////////////////////////////////////////////////////    
  //
  // PRIVATE CONSTANTS
  //
  //////////////////////////////////////////////////////////////////////////// 

  private static final double DEFAULT_SELECTION_OPAQUE_LEVEL_ = 0.90; 
  private static final double DEFAULT_NODE_LIGHTER_LEVEL_ = 0.43; 
  private static final double DEFAULT_LINK_DARKER_LEVEL_ = 0.43;
  private static final boolean DEFAULT_DISPLAY_SHADOWS_ = false;
  private static final boolean DEFAULT_SHADE_NODES_ = false;
  private static final int DEFAULT_MIN_DRAIN_ZONE_ = 1;
  private static final boolean DEFAULT_OFFER_NODE_BROWSER_ = false;
  private static final boolean DEFAULT_OFFER_LINK_BROWSER_ = false;
  private static final boolean DEFAULT_OFFER_MOUSEOVER_VIEW_ = false;
  private static final boolean DEFAULT_MINIMIZE_SHADOW_SUBMODEL_LINKS_ = true;
  private static final String DEFAULT_BROWSER_URL_ = "http://www.genecards.org/cgi-bin/carddisp.pl?gene=" + NODE_NAME_PLACEHOLDER;
  private static final String DEFAULT_LINK_BROWSER_URL_ = "http://localhost:8080/setToYourPage.jsp?src=" + LINK_SRC_PLACEHOLDER  + "&rel=" + LINK_REL_PLACEHOLDER  + "&trg=" + LINK_TRG_PLACEHOLDER;
  private static final String DEFAULT_MOUSEOVER_URL_ = "/Users/bill/Desktop/ForBrains/" + NODE_NAME_PLACEHOLDER + ".jpg";

  ////////////////////////////////////////////////////////////////////////////
  //
  // PRIVATE VARIABLES
  //
  ////////////////////////////////////////////////////////////////////////////

  private double selectionOpaqueLevel_;
  private double nodeLighterLevel_;
  private double linkDarkerLevel_;
  private boolean displayShadows_;
  private boolean shadeNodes_;
  private boolean minShadSubLinks_;
  private boolean offerNodeBrowser_;
  private boolean offerLinkBrowser_;
  private boolean offerMouseOverView_;
  private String browserURL_;
  private String browserLinkURL_;
  private String mouseOverURL_;
  private int minDrainZone_;
 
  ////////////////////////////////////////////////////////////////////////////
  //
  // PUBLIC CONSTRUCTORS
  //
  ////////////////////////////////////////////////////////////////////////////

  /***************************************************************************
  **
  ** Constructor
  */

  public FabricDisplayOptions() {
    selectionOpaqueLevel_ = DEFAULT_SELECTION_OPAQUE_LEVEL_;
    nodeLighterLevel_ = DEFAULT_NODE_LIGHTER_LEVEL_;
    linkDarkerLevel_ = DEFAULT_LINK_DARKER_LEVEL_;
    displayShadows_ = DEFAULT_DISPLAY_SHADOWS_;
    shadeNodes_ = DEFAULT_SHADE_NODES_;
    minDrainZone_ = DEFAULT_MIN_DRAIN_ZONE_;
    browserURL_ = DEFAULT_BROWSER_URL_;
    browserLinkURL_ = DEFAULT_LINK_BROWSER_URL_;
    minShadSubLinks_ = DEFAULT_MINIMIZE_SHADOW_SUBMODEL_LINKS_;
    
    offerNodeBrowser_ = DEFAULT_OFFER_NODE_BROWSER_;
    offerLinkBrowser_ = DEFAULT_OFFER_LINK_BROWSER_;
    offerMouseOverView_ = DEFAULT_OFFER_MOUSEOVER_VIEW_;
    mouseOverURL_= DEFAULT_MOUSEOVER_URL_;
  }
  
  /***************************************************************************
  **
  ** Constructor for IO
  */

  FabricDisplayOptions(String selectionOpaqueLevelStr, String nodeLighterLevelStr, 
                       String linkDarkerLevelStr, String displayShadowsStr,
                       String shadeNodesStr, String minDrainZoneStr, String minShadSubLinksStr,
                       String offerNodeBrowser, String offerLinkBrowser, String offerMouseOverView,
                       String browserURL, String browserLinkURL, String mouseOverURL) throws IOException {
    
    if (selectionOpaqueLevelStr != null) {
      try {
        selectionOpaqueLevel_ = Double.parseDouble(selectionOpaqueLevelStr);
      } catch (NumberFormatException ex) {
        throw new IOException();
      }
    } else {
      selectionOpaqueLevel_ = DEFAULT_SELECTION_OPAQUE_LEVEL_;
    }
    
    if (nodeLighterLevelStr != null) {
      try {
        nodeLighterLevel_ = Double.parseDouble(nodeLighterLevelStr);
      } catch (NumberFormatException ex) {
        throw new IOException();
      }
    } else {
      nodeLighterLevel_ = DEFAULT_NODE_LIGHTER_LEVEL_;
    }
    
    if (linkDarkerLevelStr != null) {
      try {
        linkDarkerLevel_ = Double.parseDouble(linkDarkerLevelStr);
      } catch (NumberFormatException ex) {
        throw new IOException();
      }
    } else {
      linkDarkerLevel_ = DEFAULT_LINK_DARKER_LEVEL_;
    }
    
    if (displayShadowsStr != null) {
      displayShadows_ = Boolean.valueOf(displayShadowsStr).booleanValue();
    } else {
      displayShadows_ = DEFAULT_DISPLAY_SHADOWS_;
    }
    
    if (shadeNodesStr != null) {
      shadeNodes_ = Boolean.valueOf(shadeNodesStr).booleanValue();
    } else {
      shadeNodes_ = DEFAULT_SHADE_NODES_;
    }
    
    if (minDrainZoneStr != null) {
      minDrainZone_ = Integer.valueOf(minDrainZoneStr).intValue();
    } else {
      minDrainZone_ = DEFAULT_MIN_DRAIN_ZONE_;
    }
    
    if (minShadSubLinksStr != null) {
      minShadSubLinks_ = Boolean.valueOf(minShadSubLinksStr).booleanValue();
    } else {
      minShadSubLinks_ = DEFAULT_MINIMIZE_SHADOW_SUBMODEL_LINKS_;
    }
    
    if (offerNodeBrowser != null) {
      offerNodeBrowser_ = Boolean.valueOf(offerNodeBrowser).booleanValue();
    } else {
      offerNodeBrowser_ = DEFAULT_OFFER_NODE_BROWSER_;
    }
        
    if (offerLinkBrowser != null) {
      offerLinkBrowser_ = Boolean.valueOf(offerLinkBrowser).booleanValue();
    } else {
      offerLinkBrowser_ = DEFAULT_OFFER_LINK_BROWSER_;
    }
        
    if (offerMouseOverView != null) {
      offerMouseOverView_ = Boolean.valueOf(offerMouseOverView).booleanValue();
    } else {
      offerMouseOverView_ = DEFAULT_OFFER_MOUSEOVER_VIEW_;
    }
 
    if (browserURL != null) {
      browserURL_ = browserURL;
    } else {
      browserURL_ = DEFAULT_BROWSER_URL_;
    }
    
    if (browserLinkURL != null) {
      browserLinkURL_ = browserLinkURL;
    } else {
      browserLinkURL_ = DEFAULT_LINK_BROWSER_URL_;
    }
    
    if (mouseOverURL != null) {
      mouseOverURL_ = mouseOverURL;
    } else {
      mouseOverURL_ = DEFAULT_MOUSEOVER_URL_;
    }      
  }
  
  ////////////////////////////////////////////////////////////////////////////
  //
  // PUBLIC METHODS
  //
  //////////////////////////////////////////////////////////////////////////// 

  /***************************************************************************
  **
  ** Get browser URL
  */

  public String getBrowserURL() {
    return (browserURL_);
  }   
 
  /***************************************************************************
  **
  ** Set browser URL
  */

  public void setBrowserURL(String browserURL) {
    browserURL_ = browserURL;
    return;
  }   
  
  /***************************************************************************
  **
  ** Get browser Link URL
  */

  public String getBrowserLinkURL() {
    return (browserLinkURL_);
  }   
 
  /***************************************************************************
  **
  ** Set browser Link URL
  */

  public void setBrowserLinkURL(String browserLinkURL) {
    browserLinkURL_ = browserLinkURL;
    return;
  }   
 
  /***************************************************************************
  **
  ** Get mouseover URL
  */

  public String getMouseOverURL() {
    return (mouseOverURL_);
  }   
 
  /***************************************************************************
  **
  ** Set mouseover URL
  */

  public void setMouseOverURL(String mouseOverURL) {
    mouseOverURL_ = mouseOverURL;
    return;
  }     
   
  /***************************************************************************
  **
  ** Get Selection Opaque level
  */

  public double getSelectionOpaqueLevel() {
    return (selectionOpaqueLevel_);
  }   
 
  /***************************************************************************
  **
  ** Set Selection Opaque level
  */

  public void setSelectionOpaqueLevel(double opqLevel) {
    if ((opqLevel < 0.0) || (opqLevel > 1.0)) {
      throw new IllegalArgumentException();
    }
    selectionOpaqueLevel_ = opqLevel;
    return;
  }     

  /***************************************************************************
  **
  ** Get Node Lighter level
  */

  public double getNodeLighterLevel() {
    return (nodeLighterLevel_);
  }   
 
  /***************************************************************************
  **
  ** Set Node Lighter level
  */

  public void setNodeLighterLevel(double nodeLight) {
    if (nodeLight < 0.0) {
      throw new IllegalArgumentException();
    }
    nodeLighterLevel_ = nodeLight;
    return;
  }     
  
  /***************************************************************************
  **
  ** Get Link Darker level
  */

  public double getLinkDarkerLevel() {
    return (linkDarkerLevel_);
  }   
 
  /***************************************************************************
  **
  ** Set Link Darker level
  */

  public void setLinkDarkerLevel(double linkDark) {
    if (linkDark < 0.0) {
      throw new IllegalArgumentException();
    }
    linkDarkerLevel_ = linkDark;
    return;
  }        
  
  /***************************************************************************
  **
  ** Get if we are to minimize links in shadow submodels
  */

  public boolean getMinShadowSubmodelLinks() {
    return (minShadSubLinks_);
  }   
 
  /***************************************************************************
  **
  ** Set if we are to minimize links in shadow submodels
  */

  public void setMinShadowSubmodelLinks(boolean doMin) {
    minShadSubLinks_ = doMin;
    return;
  }     

  /***************************************************************************
  **
  ** Get if we are to display shadow links
  */

  public boolean getDisplayShadows() {
    return (displayShadows_);
  }   
 
  /***************************************************************************
  **
  ** Set if we are to display shadow links
  */

  public void setDisplayShadows(boolean display) {
    displayShadows_ = display;
    return;
  }     
  
  /***************************************************************************
  **
  ** Get if we are to shade nodes
  */

  public boolean getShadeNodes() {
    return (shadeNodes_);
  }   
 
  /***************************************************************************
  **
  ** Set if we are to shade nodes
  */

  public void setShadeNodes(boolean shade) {
    shadeNodes_ = shade;
    return;
  }
  
  /***************************************************************************
   **
   ** Get if we are changing min drain zone size
   */
  
  public int getMinDrainZone() {
    return (minDrainZone_);
  }
  
  /***************************************************************************
   **
   ** Set if we are changing min drain zone size
   */

  public void setMinDrainZone(int size) {
    minDrainZone_ = size;
    return;
  }
  
  /***************************************************************************
  **
  ** Get if we are to offer node browser option
  */

  public boolean getOfferNodeBrowser() {
    return (offerNodeBrowser_);
  }   
 
  /***************************************************************************
  **
  ** Set if we are to offer node browser option
  */

  public void setOfferNodeBrowser(boolean offerNodeBrowser) {
    offerNodeBrowser_ = offerNodeBrowser;
    return;
  }  
  
 /***************************************************************************
  **
  ** Get if we are to offer link browser option
  */

  public boolean getOfferLinkBrowser() {
    return (offerLinkBrowser_);
  }   
 
  /***************************************************************************
  **
  ** Set if we are to offer link browser option
  */

  public void setOfferLinkBrowser(boolean offerLinkBrowser) {
    offerLinkBrowser_ = offerLinkBrowser;
    return;
  }  
  
  /***************************************************************************
  **
  ** Get if we are to offer mouseover view
  */

  public boolean getOfferMouseOverView() {
    return (offerMouseOverView_);
  }   
 
  /***************************************************************************
  **
  ** Set if we are to offer mouseover view
  */

  public void setOfferMouseOverView(boolean offerMouseOverView) {
    offerMouseOverView_ = offerMouseOverView;
    return;
  }  
  
  /***************************************************************************
  **
  ** Clone
  */

  @Override
  public FabricDisplayOptions clone() {
    try {
      FabricDisplayOptions retval = (FabricDisplayOptions)super.clone();
      return (retval);
    } catch (CloneNotSupportedException cnse) {
      throw new IllegalStateException();
    }
  }
 
  /***************************************************************************
  **
  ** Write the item to XML
  **
  */
  
  public void writeXML(PrintWriter out, Indenter ind) {
    ind.indent();
    out.print("<displayOptions ");
    
    if (selectionOpaqueLevel_ != DEFAULT_SELECTION_OPAQUE_LEVEL_) {
      out.print("selOpq=\"");
      out.print(selectionOpaqueLevel_);    
      out.print("\" ");
    }
    
    if (nodeLighterLevel_ != DEFAULT_NODE_LIGHTER_LEVEL_) {
      out.print("nodeLight=\"");
      out.print(nodeLighterLevel_);    
      out.print("\" ");
    }
     
    if (linkDarkerLevel_ != DEFAULT_LINK_DARKER_LEVEL_) {
      out.print("linkDark=\"");
      out.print(linkDarkerLevel_);    
      out.print("\" ");
    }
  
    if (displayShadows_ != DEFAULT_DISPLAY_SHADOWS_) {
      out.print("shadows=\"");
      out.print(displayShadows_);    
      out.print("\" ");
    }
    
    if (shadeNodes_ != DEFAULT_SHADE_NODES_) {
      out.print("shading=\"");
      out.print(shadeNodes_);    
      out.print("\" ");
    }
  
    if (minDrainZone_ != DEFAULT_MIN_DRAIN_ZONE_) {
      out.print("minDrainZone=\"");
      out.print(minDrainZone_);
      out.print("\" ");
    }
    
    if (minShadSubLinks_ != DEFAULT_MINIMIZE_SHADOW_SUBMODEL_LINKS_) {
      out.print("minShadSubLinks=\"");
      out.print(minShadSubLinks_);    
      out.print("\" ");
    }
    
    if (offerNodeBrowser_ != DEFAULT_OFFER_NODE_BROWSER_) {
      out.print("offerNodeBrowser=\"");
      out.print(offerNodeBrowser_);    
      out.print("\" ");
    }
        
    if (offerLinkBrowser_ != DEFAULT_OFFER_LINK_BROWSER_) {
      out.print("offerLinkBrowser=\"");
      out.print(offerLinkBrowser_);    
      out.print("\" ");
    }  
    
    if (offerMouseOverView_ != DEFAULT_OFFER_MOUSEOVER_VIEW_) {
      out.print("offerMouseOverView=\"");
      out.print(offerMouseOverView_);    
      out.print("\" ");
    }
    
    if (!browserURL_.equals(DEFAULT_BROWSER_URL_)) {
      out.print("browserURL=\"");
      out.print(CharacterEntityMapper.mapEntities(browserURL_, false));    
      out.print("\" ");
    }
    
    if (!browserLinkURL_.equals(DEFAULT_LINK_BROWSER_URL_)) {
      out.print("browserLinkURL=\"");
      out.print(CharacterEntityMapper.mapEntities(browserLinkURL_, false));    
      out.print("\" ");
    }
    
    if (!mouseOverURL_.equals(DEFAULT_MOUSEOVER_URL_)) {
      out.print("mouseOverURL=\"");
      out.print(CharacterEntityMapper.mapEntities(mouseOverURL_, false));    
      out.print("\" ");
    }
         
    out.println("/>");  
    return;
  }  
  
 
  ////////////////////////////////////////////////////////////////////////////
  //
  // PUBLIC CLASS METHODS
  //
  ////////////////////////////////////////////////////////////////////////////
  
  /***************************************************************************
  **
  ** For XML I/O
  */  
      
  public static class FabricDisplayOptionsWorker extends AbstractFactoryClient {
 
    public FabricDisplayOptionsWorker(FabricFactory.FactoryWhiteboard whiteboard) {
      super(whiteboard);
      myKeys_.add("displayOptions");
    }
  
    protected Object localProcessElement(String elemName, Attributes attrs) throws IOException {
      Object retval = null;
      if (elemName.equals("displayOptions")) {
        FabricFactory.FactoryWhiteboard board = (FabricFactory.FactoryWhiteboard)this.sharedWhiteboard_;
        board.displayOpts = buildFromXML(elemName, attrs);
        FabricDisplayOptionsManager.getMgr().setDisplayOptionsForIO(board.displayOpts);
        retval = board.displayOpts;
      }
      return (retval);     
    }
    
    private FabricDisplayOptions buildFromXML(String elemName, Attributes attrs) throws IOException {  
      String selectionOpaqueLevelStr = AttributeExtractor.extractAttribute(elemName, attrs, "displayOptions", "selOpq", false);
      String nodeLighterLevelStr = AttributeExtractor.extractAttribute(elemName, attrs, "displayOptions", "nodeLight", false);
      String linkDarkerLevelStr = AttributeExtractor.extractAttribute(elemName, attrs, "displayOptions", "linkDark", false);
      String displayShadowsStr = AttributeExtractor.extractAttribute(elemName, attrs, "displayOptions", "shadows", false);
      String shadeNodesStr = AttributeExtractor.extractAttribute(elemName, attrs, "displayOptions", "shading", false);
      String minDrainZoneStr = AttributeExtractor.extractAttribute(elemName, attrs, "displayOptions", "minDrainZone",false);
      String minShadSubLinks = AttributeExtractor.extractAttribute(elemName, attrs, "displayOptions", "minShadSubLinks", false);
      String offerNodeBrowserStr = AttributeExtractor.extractAttribute(elemName, attrs, "displayOptions", "offerNodeBrowser", false);
      String offerLinkBrowserStr = AttributeExtractor.extractAttribute(elemName, attrs, "displayOptions", "offerLinkBrowser", false);
      String offerMouseOverViewStr = AttributeExtractor.extractAttribute(elemName, attrs, "displayOptions", "offerMouseOverView", false);
      String browserURL = AttributeExtractor.extractAttribute(elemName, attrs, "displayOptions", "browserURL", false);
      String browserLinkURL = AttributeExtractor.extractAttribute(elemName, attrs, "displayOptions", "browserLinkURL", false);
      String mouseOverURL = AttributeExtractor.extractAttribute(elemName, attrs, "displayOptions", "mouseOverURL", false);
   
      browserURL = (browserURL == null) ? null : CharacterEntityMapper.unmapEntities(browserURL, false);
      browserLinkURL =  (browserLinkURL == null) ? null : CharacterEntityMapper.unmapEntities(browserLinkURL, false);
      
      return (new FabricDisplayOptions(selectionOpaqueLevelStr, nodeLighterLevelStr, 
                                       linkDarkerLevelStr, displayShadowsStr,
                                       shadeNodesStr, minDrainZoneStr, minShadSubLinks,
                                       offerNodeBrowserStr, offerLinkBrowserStr, offerMouseOverViewStr,
                                       browserURL, browserLinkURL, mouseOverURL));        
    }
  }
 
}
```
