## Supplementary material for "BioFabric Visualization of Network Alignments": BioFabric Source Code: FabricDisplayOptionsManager.java

```
/*
**    Copyright (C) 2003-2014 Institute for Systems Biology 
**                            Seattle, Washington, USA. 
**
**    This library is free software; you can redistribute it and/or
**    modify it under the terms of the GNU Lesser General Public
**    License as published by the Free Software Foundation; either
**    version 2.1 of the License, or (at your option) any later version.
**
**    This library is distributed in the hope that it will be useful,
**    but WITHOUT ANY WARRANTY; without even the implied warranty of
**    MERCHANTABILITY or FITNESS FOR A PARTICULAR PURPOSE.  See the GNU
**    Lesser General Public License for more details.
**
**    You should have received a copy of the GNU Lesser General Public
**    License along with this library; if not, write to the Free Software
**    Foundation, Inc., 59 Temple Place, Suite 330, Boston, MA  02111-1307  USA
*/

package org.systemsbiology.biofabric.ui;

import java.io.PrintWriter;
import java.util.ArrayList;

import org.systemsbiology.biofabric.api.io.Indenter;

/****************************************************************************
**
** Display Options Manager.  This is a Singleton.
*/

public class FabricDisplayOptionsManager {
  
  ////////////////////////////////////////////////////////////////////////////
  //
  // PUBLIC CONSTANTS
  //
  //////////////////////////////////////////////////////////////////////////// 

  ////////////////////////////////////////////////////////////////////////////
  //
  // PRIVATE MEMBERS
  //
  ////////////////////////////////////////////////////////////////////////////

  private static FabricDisplayOptionsManager singleton_;
  private FabricDisplayOptions options_;
  private ArrayList<DisplayOptionTracker> trackers_;
  
  ////////////////////////////////////////////////////////////////////////////
  //
  // PUBLIC METHODS
  //
  ////////////////////////////////////////////////////////////////////////////

  
  /***************************************************************************
  ** 
  ** Get the display options
  */

  public FabricDisplayOptions getDisplayOptions() {
    return (options_);
  }
  
  /***************************************************************************
  ** 
  ** Set the display options
  */

  public void setDisplayOptions(FabricDisplayOptions opts, boolean needRebuild, boolean needRecolor) {
    options_ = opts.clone();
    int numTrack = trackers_.size();
    for (int i = 0; i < numTrack; i++) {
      DisplayOptionTracker dot = trackers_.get(i);
      dot.optionsHaveChanged(needRebuild, needRecolor);
    }
    return;
  }
  
  /***************************************************************************
  ** 
  ** Set the display options for IO.
  */

  public void setDisplayOptionsForIO(FabricDisplayOptions opts) {
    options_ = opts;
    return;
  }
 
  /***************************************************************************
  ** 
  ** Add a tracker
  */

  public void addTracker(DisplayOptionTracker dot) {
    trackers_.add(dot);
    return;
  }
  
  /***************************************************************************
  **
  ** Write the options to XML
  **
  */
  
  public void writeXML(PrintWriter out, Indenter ind) {
    options_.writeXML(out, ind);
    return;
  }
   
  ////////////////////////////////////////////////////////////////////////////
  //
  // PUBLIC STATIC METHODS
  //
  ////////////////////////////////////////////////////////////////////////////
  
  
  /***************************************************************************
  ** 
  ** Get the singleton
  */

  public static synchronized FabricDisplayOptionsManager getMgr() {
    if (singleton_ == null) {
      singleton_ = new FabricDisplayOptionsManager();
    }
    return (singleton_);
  }
  
  ////////////////////////////////////////////////////////////////////////////
  //
  // INNER CLASSES
  //
  ////////////////////////////////////////////////////////////////////////////

  public interface DisplayOptionTracker {  
    public void optionsHaveChanged(boolean needRebuild, boolean needRecolor);
  }

  ////////////////////////////////////////////////////////////////////////////
  //
  // PRIVATE CONSTRUCTORS
  //
  ////////////////////////////////////////////////////////////////////////////

  /***************************************************************************
  **
  ** Null constructor
  */
    
  private FabricDisplayOptionsManager() {
    options_ = new FabricDisplayOptions();
    trackers_ = new ArrayList<DisplayOptionTracker>();
  }
}
```
