## Supplementary material for "BioFabric Visualization of Network Alignments": BioFabric Source Code: ImageExporter.java

```
/*
**    Copyright (C) 2003-2018 Institute for Systems Biology 
**                            Seattle, Washington, USA. 
**
**    This library is free software; you can redistribute it and/or
**    modify it under the terms of the GNU Lesser General Public
**    License as published by the Free Software Foundation; either
**    version 2.1 of the License, or (at your option) any later version.
**
**    This library is distributed in the hope that it will be useful,
**    but WITHOUT ANY WARRANTY; without even the implied warranty of
**    MERCHANTABILITY or FITNESS FOR A PARTICULAR PURPOSE.  See the GNU
**    Lesser General Public License for more details.
**
**    You should have received a copy of the GNU Lesser General Public
**    License along with this library; if not, write to the Free Software
**    Foundation, Inc., 59 Temple Place, Suite 330, Boston, MA  02111-1307  USA
*/

package org.systemsbiology.biofabric.ui;

import java.util.Iterator;
import java.util.List;
import java.util.HashSet;
import java.util.Arrays;
import java.util.ArrayList;

import java.awt.image.BufferedImage;
import java.io.IOException;
import javax.imageio.ImageIO;
import javax.imageio.ImageWriter;
import javax.imageio.stream.ImageOutputStream;
import javax.imageio.metadata.IIOMetadata;
import javax.imageio.metadata.IIOMetadataNode;
import javax.imageio.ImageWriteParam;
import javax.imageio.ImageTypeSpecifier;
import javax.imageio.IIOImage;
import javax.imageio.metadata.IIOInvalidTreeException;

import org.w3c.dom.Node;
import org.w3c.dom.NodeList;

/****************************************************************************
**
** Do image IO testing
*/

public class ImageExporter {
  
  ////////////////////////////////////////////////////////////////////////////
  //
  // PRIVATE CONSTANTS
  //
  //////////////////////////////////////////////////////////////////////////// 
  
  ////////////////////////////////////////////////////////////////////////////
  //
  // PUBLIC CONSTANTS
  //
  //////////////////////////////////////////////////////////////////////////// 

  public static final int INCHES  = 0;
  public static final int CM      = 1;
  // Internal use only:
  private static final int METER_ = 2;  
  
  ////////////////////////////////////////////////////////////////////////////
  //
  // PRIVATE INSTANCE MEMBERS
  //
  ////////////////////////////////////////////////////////////////////////////

  ////////////////////////////////////////////////////////////////////////////
  //
  // PUBLIC CONSTRUCTORS
  //
  ////////////////////////////////////////////////////////////////////////////

  /***************************************************************************
  **
  ** Constructor
  */

  public ImageExporter() {
  }

  ////////////////////////////////////////////////////////////////////////////
  //
  // PUBLIC METHODS
  //
  ////////////////////////////////////////////////////////////////////////////

  /***************************************************************************
  **
  ** Support image export
  */  
  
  public void export(Object outObj, BufferedImage bi, String format, ResolutionSettings res) throws IOException {
  
    if (format.equalsIgnoreCase("TIFF")) {
      writeTIFFImage(bi, outObj, res);  
    } else if (format.equalsIgnoreCase("PNG")) {
      writePNGImage(bi, outObj, res);
    } else if (format.equalsIgnoreCase("JPEG")) {
      writeJPGImage(bi, outObj, res);
    } else {
      throw new IllegalArgumentException();
    }
    return;
  }

  ////////////////////////////////////////////////////////////////////////////
  //
  // PUBLIC INNER CLASSES
  //
  ////////////////////////////////////////////////////////////////////////////  
   
  /***************************************************************************
  **
  ** This is for a rational number
  ** 
  */
  
  public static class RationalNumber {
    public double value;
    public int numerator;
    public int denominator;
    
    public RationalNumber(double val, int num, int den) {
      value = val;
      numerator = num;
      denominator = den;
    }   
  }
  
  /***************************************************************************
  **
  ** These are the resolution settings we may need
  ** 
  */
  
  public static class ResolutionSettings {
    public RationalNumber dotsPerUnit;
    public int units;
  }
  
  ////////////////////////////////////////////////////////////////////////////
  //
  // PRIVATE METHODS
  //
  ////////////////////////////////////////////////////////////////////////////  

  /***************************************************************************
  ** 
  ** Set resolution metadata
  */

  private void setTIFFResolutionMetadata(IIOMetadata meta, ResolutionSettings res) {

    IIOMetadataNode topnode = (IIOMetadataNode)meta.getAsTree(meta.getNativeMetadataFormatName()); 
    NodeList nl = topnode.getElementsByTagName("TIFFField");
    String[] svals = new String[] {"ImageWidth","ImageLength","ResolutionUnit"};
    String[] rvals = new String[] {"XResolution","YResolution"};
    
    List smatches = Arrays.asList(svals);
    List rmatches = Arrays.asList(rvals);    
    int num = nl.getLength();
    for (int i = 0; i < num; i++) {
      IIOMetadataNode node = (IIOMetadataNode)nl.item(i);
      String nameVal = node.getAttribute("name");
      if (smatches.contains(nameVal)) {          
        NodeList snl = ((IIOMetadataNode)node).getElementsByTagName("TIFFShort");
        IIOMetadataNode shortNode = (IIOMetadataNode)snl.item(0);
        //Horizontal and vertical resolution unit
        //1 - none specified
        //2 - Inches (English units)
        //3 - Centimeters (SI units)  
        if (nameVal.equals("ResolutionUnit")) {
          String resUnits = (res.units == INCHES) ? "2" : "3";
          shortNode.setAttribute("value", resUnits);
        }
      } else if (rmatches.contains(nameVal)) {          
        NodeList rnl = ((IIOMetadataNode)node).getElementsByTagName("TIFFRational");
        IIOMetadataNode ratNode = (IIOMetadataNode)rnl.item(0);
        ratNode.setAttribute("value", res.dotsPerUnit.numerator + "/" + res.dotsPerUnit.denominator);
      }      
    }
    try {
      meta.setFromTree(meta.getNativeMetadataFormatName(), topnode);
    } catch (IIOInvalidTreeException itex) {
      throw new IllegalStateException();
    } 
    return;
  }
  
  /***************************************************************************
  ** 
  ** Set resolution metadata
  */

  private void setPNGResolutionMetadata(IIOMetadata meta, ResolutionSettings res) {
    if (res == null) {
      return;
    }
    IIOMetadataNode topnode = (IIOMetadataNode)meta.getAsTree(meta.getNativeMetadataFormatName());
    //<pHYs pixelsPerUnitXAxis="2835" pixelsPerUnitYAxis="2835" unitSpecifier="meter"/> 
    ResolutionSettings resMet = mapResolutionToMeters(res);
    long resVal = Math.round((double)resMet.dotsPerUnit.numerator / (double)resMet.dotsPerUnit.denominator);
    String resValStr = Long.toString(resVal);
    IIOMetadataNode pNode = new IIOMetadataNode("pHYs");
    pNode.setAttribute("pixelsPerUnitXAxis", resValStr);
    pNode.setAttribute("pixelsPerUnitYAxis", resValStr);
    pNode.setAttribute("unitSpecifier", "meter");
    topnode.appendChild(pNode);
    try {
      meta.setFromTree(meta.getNativeMetadataFormatName(), topnode);
    } catch (IIOInvalidTreeException itex) {
      throw new IllegalStateException();
    } 
    return;
  }  

  /***************************************************************************
  ** 
  ** Set resolution metadata
  */

  private void setJPGResolutionMetadata(IIOMetadata meta, ResolutionSettings res) {
    if (res == null) {
      return;
    }
    /*   
     FOR JPEG:   
      <!ELEMENT "app0JFIF" (JFXX?, app2ICC?)>
          <!ATTLIST "app0JFIF" "resUnits" ("0" | "1" | "2") "0">
            <!-- The resolution units for Xdensisty and Ydensity (0 = no 
                 units, just aspect ratio; 1 = dots/inch; 2 = dots/cm) --> 
          <!ATTLIST "app0JFIF" "Xdensity" #CDATA "1">
            <!-- The horizontal density or aspect ratio numerator --> 
            <!-- Data type: Integer -->
            <!-- Min value: 1 (inclusive) -->
            <!-- Max value: 65535 (inclusive) -->
          <!ATTLIST "app0JFIF" "Ydensity" #CDATA "1">
            <!-- The vertical density or aspect ratio denominator --> 
            <!-- Data type: Integer -->
            <!-- Min value: 1 (inclusive) -->
            <!-- Max value: 65535 (inclusive) --> 
    */    
    IIOMetadataNode topnode = (IIOMetadataNode)meta.getAsTree(meta.getNativeMetadataFormatName()); 
    NodeList nl = topnode.getElementsByTagName("app0JFIF");
    if (nl.getLength() < 1) {
      throw new IllegalStateException();
    }
    String resUnits = (res.units == INCHES) ? "1" : "2";
    long lResVal = Math.round((double)res.dotsPerUnit.numerator / (double)res.dotsPerUnit.denominator);
    String resValStr = Long.toString(lResVal);       
    IIOMetadataNode node = (IIOMetadataNode)nl.item(0);
    node.setAttribute("resUnits", resUnits);
    node.setAttribute("Xdensity", resValStr);
    node.setAttribute("Ydensity", resValStr);      
    try {
      meta.setFromTree(meta.getNativeMetadataFormatName(), topnode);
    } catch (IIOInvalidTreeException itex) {
      throw new IllegalStateException();
    } 
    return;
  }  

  /***************************************************************************
  ** 
  ** Write out an image
  */

  private void writeTIFFImage(BufferedImage bi, Object outObj, ResolutionSettings res) throws IOException {
    if (res == null) {
      throw new IllegalArgumentException();
    }
    Iterator writers = ImageIO.getImageWritersByFormatName("tiff");
    ImageWriter writer = (ImageWriter)writers.next();
    ImageOutputStream ios = ImageIO.createImageOutputStream(outObj);
    writer.setOutput(ios);
    ImageWriteParam param = writer.getDefaultWriteParam();
    param.setCompressionMode(ImageWriteParam.MODE_EXPLICIT);
    // This could be user-configurable instead...
    param.setCompressionType("LZW");
    ImageTypeSpecifier its = new ImageTypeSpecifier(bi);
    IIOMetadata meta = writer.getDefaultImageMetadata(its, param);
    setTIFFResolutionMetadata(meta, res);
    IIOImage img = new IIOImage(bi, null, meta);
    writer.write(writer.getDefaultStreamMetadata(param), img, param);
    ios.close();
    writer.dispose();
    return;
  }
  
  /***************************************************************************
  ** 
  ** Write out an PNG image
  */

  private void writePNGImage(BufferedImage bi, Object outObj, ResolutionSettings res) throws IOException {
    Iterator writers = ImageIO.getImageWritersByFormatName("png");
    ImageWriter writer = (ImageWriter)writers.next();
    ImageOutputStream ios = ImageIO.createImageOutputStream(outObj);
    writer.setOutput(ios);
    ImageWriteParam param = writer.getDefaultWriteParam();
    ImageTypeSpecifier its = new ImageTypeSpecifier(bi);
    IIOMetadata meta = writer.getDefaultImageMetadata(its, param);
    Node node = meta.getAsTree(meta.getNativeMetadataFormatName());
    setPNGResolutionMetadata(meta, res);    
    IIOImage img = new IIOImage(bi, null, meta);
    writer.write(writer.getDefaultStreamMetadata(param), img, param);
    ios.close();
    writer.dispose();
    return;
  }

  
  /***************************************************************************
  ** 
  ** Write out a JPG image
  */  
  
  private void writeJPGImage(BufferedImage bi, Object outObj, ResolutionSettings res) throws IOException {
    Iterator writers = ImageIO.getImageWritersByFormatName("jpg");
    ImageWriter writer = (ImageWriter)writers.next();
    ImageOutputStream ios = ImageIO.createImageOutputStream(outObj);
    writer.setOutput(ios);
    ImageWriteParam param = writer.getDefaultWriteParam();
    param.setCompressionMode(ImageWriteParam.MODE_EXPLICIT);
    param.setCompressionType("JPEG");
    // This could be user-configurable instead...
    param.setCompressionQuality(1.0F);
    ImageTypeSpecifier its = new ImageTypeSpecifier(bi);
    IIOMetadata meta = writer.getDefaultImageMetadata(its, param);
    Node node = meta.getAsTree(meta.getNativeMetadataFormatName());
    setJPGResolutionMetadata(meta, res);    
    IIOImage img = new IIOImage(bi, null, meta);
    writer.write(writer.getDefaultStreamMetadata(param), img, param);
    ios.close();
    writer.dispose();
    return;
  }


  
  ////////////////////////////////////////////////////////////////////////////
  //
  // PUBLIC STATIC METHODS
  //
  ////////////////////////////////////////////////////////////////////////////

  /***************************************************************************
  **
  ** Get supported export formats
  */
  
  public static List<String> getSupportedExports() {
     
    String[] names = ImageIO.getWriterFormatNames();
    // This is a little contorted, to insure presentation order:
    HashSet<String> present = new HashSet<String>();
    for (int i = 0; i < names.length; i++) {
      if (names[i].equalsIgnoreCase("TIF")) {
        present.add("TIFF");
      } else if (names[i].equalsIgnoreCase("PNG")) {
        present.add("PNG");
      } else if (names[i].equalsIgnoreCase("JPG")) {
        present.add("JPEG");
      }
    }
    ArrayList<String> retval = new ArrayList<String>();     
    if (present.contains("TIFF")) {
      retval.add("TIFF");
    }
    if (present.contains("PNG")) {
      retval.add("PNG");
    }
    if (present.contains("JPEG")) {
      retval.add("JPEG");
    }       
    return (retval);
  }
   
  /***************************************************************************
  **
  ** Get supported export file suffixes
  */
  
  public static List<String> getSupportedFileSuffixes() {
    List<String> exps = getSupportedExports();
    ArrayList<String> retval = new ArrayList<String>(); 
    if (exps.contains("TIFF")) {
      retval.add("tif");
      retval.add("tiff");
    }
    if (exps.contains("PNG")) {
      retval.add("png");
    }
    if (exps.contains("JPEG")) {
      retval.add("jpg");
      retval.add("jpeg");
    }
    return (retval);
  }
  
  /***************************************************************************
  **
  ** Map type to suffixes
  */
  
  public static List<String> getFileSuffixesForType(String type) {
    ArrayList<String> retval = new ArrayList<String>();
    if (type.equals("TIFF")) {
      retval.add("tif");
      retval.add("tiff");      
    } else if (type.equals("PNG")) {
      retval.add("png");
    } else if (type.equals("JPEG")) {
      retval.add("jpg");
      retval.add("jpeg");
    }
    return (retval);
  }  
  
  /***************************************************************************
  **
  ** Map type to preferred suffix
  */
  
  public static String getPreferredSuffixForType(String type) {
    if (type.equals("TIFF")) {
      return ("tif");      
    } else if (type.equals("PNG")) {
      return ("png");
    } else if (type.equals("JPEG")) {
      return ("jpg");
    }
    throw new IllegalArgumentException();
  }    
  
  /***************************************************************************
  **
  ** Get supported export resolutions
  */
  
  public static List getSupportedResolutions(boolean forPub) {
    ArrayList retval = new ArrayList();
    if (forPub) {
      Object[] res = new Object[2];
      res[INCHES] = new RationalNumber(150.0, 150, 1);
      res[CM] = new RationalNumber(59.0, 59, 1);
      retval.add(res);
      res = new Object[2];
      res[INCHES] = new RationalNumber(300.0, 300, 1);
      res[CM] = new RationalNumber(118.0, 118, 1);
      retval.add(res);
      res = new Object[2];
      res[INCHES] = new RationalNumber(600.0, 600, 1);
      res[CM] = new RationalNumber(236.0, 236, 1);
      retval.add(res);      
    } else {
      Object[] res = new Object[2];
      res[INCHES] = new RationalNumber(72.0, 72, 1);
      res[CM] = new RationalNumber(28.0, 28, 1);
      retval.add(res);            
    }
    return (retval);
  }
  
  /***************************************************************************
  **
  ** Map a resolution setting to meters
  */
  
  public static ResolutionSettings mapResolutionToMeters(ResolutionSettings res) {
    ResolutionSettings retval = new ResolutionSettings();
    retval.units = METER_;
    RationalNumber oldDpu = res.dotsPerUnit;
    int newNum;
    if (res.units == CM) {     
      newNum = oldDpu.numerator * 100;      
    } else {
      switch (oldDpu.numerator) {
        case 150:
          newNum = 5900;
          break;
        case 300:
          newNum = 11800;
          break;
        case 600:
          newNum = 23600;
          break;
        case 72:
          newNum = 2800; 
          break;
        default:
          throw new IllegalArgumentException();
      }
    }
    int newDen = oldDpu.denominator;
    double newRes = (double)newNum;
    retval.dotsPerUnit = new RationalNumber(newRes, newNum, newDen);
    return (retval);
  }
  
 /***************************************************************************
  **
  ** Answer if we must give a resolution
  */
  
  public static boolean formatRequiresResolution(String type) {
    return (type.equals("TIFF"));
  }
}
```
