## Supplementary material for "BioFabric Visualization of Network Alignments": BioFabric Source Code: NamedColor.java

```
/*
**    Copyright (C) 2003-2018 Institute for Systems Biology 
**                            Seattle, Washington, USA. 
**
**    This library is free software; you can redistribute it and/or
**    modify it under the terms of the GNU Lesser General Public
**    License as published by the Free Software Foundation; either
**    version 2.1 of the License, or (at your option) any later version.
**
**    This library is distributed in the hope that it will be useful,
**    but WITHOUT ANY WARRANTY; without even the implied warranty of
**    MERCHANTABILITY or FITNESS FOR A PARTICULAR PURPOSE.  See the GNU
**    Lesser General Public License for more details.
**
**    You should have received a copy of the GNU Lesser General Public
**    License along with this library; if not, write to the Free Software
**    Foundation, Inc., 59 Temple Place, Suite 330, Boston, MA  02111-1307  USA
*/

package org.systemsbiology.biofabric.ui;

import java.awt.Color;

import org.systemsbiology.biofabric.util.ColorListRenderer;

/****************************************************************************
**
** Represents a color with a name
*/

public class NamedColor implements Comparable<NamedColor>, ColorListRenderer.ColorSource {
 
  ////////////////////////////////////////////////////////////////////////////
  //
  // PUBLIC INSTANCE VARIABLES
  //
  ////////////////////////////////////////////////////////////////////////////  
   
  public Color color;
  public String name;
  public String key;
  
  ////////////////////////////////////////////////////////////////////////////
  //
  // PRIVATE CLASS VARIABLES
  //
  ////////////////////////////////////////////////////////////////////////////  
   
  private static final int H = 0;
  private static final int S = 1;
  private static final int B = 2;

  ////////////////////////////////////////////////////////////////////////////
  //
  // PUBLIC CONSTRUCTORS
  //
  ////////////////////////////////////////////////////////////////////////////
  
  /***************************************************************************
  **
  ** Constructor
  */

  public NamedColor(String key, Color color, String name) {
    this.key = key;
    this.color = color;
    this.name = name;
  }

  /***************************************************************************
  **
  ** Copy Constructor
  */

  public NamedColor(NamedColor other) {
    this.key = other.key;
    this.color = other.color;
    this.name = other.name;
  }  
  
  /***************************************************************************
  **
  ** Sorts named colors for output
  ** Compares this object with the specified object for order.  Returns a
   * negative integer, zero, or a positive integer as this object is less
   * than, equal to, or greater than the specified object.<p>
  */
  
  public int compareTo(NamedColor other) { 
    if (this == other) {
      return (0);
    }
    Color tCol = this.color;
    Color oCol = other.color;
    
    //
    // If the colors are equal, we sort by name only:
    //
    
    if (tCol.equals(oCol)) {
      return (this.name.compareToIgnoreCase(other.name));
    }    
    
    //
    // Sort by HSB.  If S = 0, it goes first, and we sort by
    // brightness only.  Otherwise, sort by (binned)hue, then saturation, then brighness.
    //
    
    float[] tHsb = Color.RGBtoHSB(tCol.getRed(), tCol.getGreen(), tCol.getBlue(), null);
    float[] oHsb = Color.RGBtoHSB(oCol.getRed(), oCol.getGreen(), oCol.getBlue(), null);    
    
    if ((oHsb[S] == 0.0) && (tHsb[S] == 0.0)) {
      float bDiff = tHsb[B] - oHsb[B];
      if (bDiff != 0.0F) {
        return (bDiff > 0) ? 1 : -1;
      } else {
        return (this.name.compareToIgnoreCase(other.name));
      }
    } else if (oHsb[S] == 0.0) {
      return (1);
    } else if (tHsb[S] == 0.0) {
      return (-1);
    }

    float hDiff = tHsb[H] - oHsb[H];
    if (Math.abs(hDiff) > 0.05F) {  // Bin hues by 20 bins
      return (hDiff > 0) ? 1 : -1;
    }
    
    float sDiff = tHsb[S] - oHsb[S];
    if (sDiff != 0.0F) {
      return (sDiff > 0) ? 1 : -1;
    }
    
    float bDiff = tHsb[B] - oHsb[B];
    if (bDiff != 0.0F) {
      return (bDiff > 0) ? 1 : -1;
    }    
    
    //
    // Still tied, drop back to exact hue:
    //
    
    if (hDiff != 0) {
      return (hDiff > 0) ? 1 : -1;
    }
    
    //
    // Same color, use name:
    //
    
    return (this.name.compareToIgnoreCase(other.name));    
  } 
  
  public Color getColor() {
    return (color);
  }
    
  public String getDescription(){
    return (name);
  } 
}
```
