## Supplementary material for "BioFabric Visualization of Network Alignments": BioFabric Source Code: PopupMenuControl.java

```
/*
**    Copyright (C) 2003-2012 Institute for Systems Biology 
**                            Seattle, Washington, USA. 
**
**    This library is free software; you can redistribute it and/or
**    modify it under the terms of the GNU Lesser General Public
**    License as published by the Free Software Foundation; either
**    version 2.1 of the License, or (at your option) any later version.
**
**    This library is distributed in the hope that it will be useful,
**    but WITHOUT ANY WARRANTY; without even the implied warranty of
**    MERCHANTABILITY or FITNESS FOR A PARTICULAR PURPOSE.  See the GNU
**    Lesser General Public License for more details.
**
**    You should have received a copy of the GNU Lesser General Public
**    License along with this library; if not, write to the Free Software
**    Foundation, Inc., 59 Temple Place, Suite 330, Boston, MA  02111-1307  USA
*/

package org.systemsbiology.biofabric.ui;

import java.awt.Point;
import java.awt.event.ActionEvent;
import javax.swing.JPopupMenu;
import javax.swing.event.PopupMenuListener;
import javax.swing.event.PopupMenuEvent;
import javax.swing.AbstractAction;
import javax.swing.Action;
import javax.swing.JPanel;

import org.systemsbiology.biofabric.api.model.NetNode;
import org.systemsbiology.biofabric.api.util.ExceptionHandler;
import org.systemsbiology.biofabric.model.FabricLink;
import org.systemsbiology.biofabric.util.ResourceManager;

/***************************************************************************
** 
** Handles building and controlling popup menus
*/

public class PopupMenuControl {
  
  ////////////////////////////////////////////////////////////////////////////
  //
  // PUBLIC CONSTANTS
  //
  //////////////////////////////////////////////////////////////////////////// 
  
  ////////////////////////////////////////////////////////////////////////////
  //
  // PRIVATE MEMBERS
  //
  ////////////////////////////////////////////////////////////////////////////
   
  private NodePopup popupGuts_;
  private LinkPopup popupGutsLink_;
  private NetNode currNode_;
  private FabricLink currLink_;
  private JPanel parent_;
  
  ////////////////////////////////////////////////////////////////////////////
  //
  // PRIVATE CONSTANTS
  //
  //////////////////////////////////////////////////////////////////////////// 
    
  ////////////////////////////////////////////////////////////////////////////
  //
  // PUBLIC CONSTRUCTOR
  //
  ////////////////////////////////////////////////////////////////////////////  

  /***************************************************************************
  **
  ** Constructor
  */
  
  public PopupMenuControl(JPanel parent) {
    parent_ = parent;
    popupGuts_ = new NodePopup();
    popupGutsLink_ = new LinkPopup();
  }    
  
  ////////////////////////////////////////////////////////////////////////////
  //
  // PUBLIC METHODS
  //
  ////////////////////////////////////////////////////////////////////////////

  /***************************************************************************
  **
  ** 
  */
  
  public boolean isVisible() {
    return (popupGuts_.isVisible() || popupGutsLink_.isVisible());   
  } 
 
  /***************************************************************************
  **
  ** Launch popup menu
  */
  
  public void showNodePopup(NetNode nodeName, Point pt) {
    currNode_ = nodeName;
    popupGuts_.showPopup(pt);       
    return;
  }
  
  /***************************************************************************
  **
  ** Launch popup menu
  */
  
  public void showLinkPopup(FabricLink link, Point pt) {
    currLink_ = link;
    popupGutsLink_.showPopup(pt);       
    return;
  }
  
  ////////////////////////////////////////////////////////////////////////////
  //
  // PRIVATE METHODS
  //
  ////////////////////////////////////////////////////////////////////////////
        
  ////////////////////////////////////////////////////////////////////////////
  //
  // PRIVATE INNER CLASSES
  //
  ////////////////////////////////////////////////////////////////////////////  
  
  /***************************************************************************
  **
  ** Used for node popup 
  */
  
  private class NodePopup {
    private JPopupMenu popMenu_;    
    private LaunchBrowserForNode lbn_;
         
    /***************************************************************************
    **
    ** Constructor
    */
  
    NodePopup() {
      ResourceManager rMan = ResourceManager.getManager();
      popMenu_ = new JPopupMenu();
      popMenu_.addPopupMenuListener(new PopupHandler());
      // Experimental Data
      lbn_ = new LaunchBrowserForNode(rMan.getString("nodePopup.launchBrowser"), rMan.getChar("nodePopup.launchBrowserMnem"));  
      popMenu_.add(lbn_);      
    }

    /***************************************************************************
    **
    ** Get the popup
    */
  
    JPopupMenu getPopup() {
      return (popMenu_);
    } 
 
    /***************************************************************************
    **
    ** Answer if visible
    */
  
    boolean isVisible() {
      return (popMenu_.isVisible());
    } 
 
    /***************************************************************************
    **
    ** Show popup menu
    */
  
    void showPopup(Point pt) {
      prepareMenu();
      popMenu_.show(parent_, pt.x, pt.y);
      popMenu_.requestFocusInWindow();
      return;
    }
         
    /***************************************************************************
    **
    ** Prepare the node menu for display
    */
  
    void prepareMenu() { 
    	FabricDisplayOptions fdo = FabricDisplayOptionsManager.getMgr().getDisplayOptions();
      String browserURL = fdo.getBrowserURL();
      boolean canShow = fdo.getOfferNodeBrowser();
      lbn_.setEnabled(canShow && !browserURL.equals(""));
      return;
    }
  }
  
  /***************************************************************************
  **
  ** Used for link popup 
  */
  
  private class LinkPopup {
    private JPopupMenu popMenu_;    
    private LaunchBrowserForLink lbl_;
         
    /***************************************************************************
    **
    ** Constructor
    */
  
    LinkPopup() {
      ResourceManager rMan = ResourceManager.getManager();
      popMenu_ = new JPopupMenu();
      popMenu_.addPopupMenuListener(new PopupHandler());
      // Experimental Data
      lbl_ = new LaunchBrowserForLink(rMan.getString("nodePopup.launchLinkBrowser"), rMan.getChar("nodePopup.launchLinkBrowserMnem"));  
      popMenu_.add(lbl_);      
    }

    /***************************************************************************
    **
    ** Get the popup
    */
  
    JPopupMenu getPopup() {
      return (popMenu_);
    } 
 
    /***************************************************************************
    **
    ** Answer if visible
    */
  
    boolean isVisible() {
      return (popMenu_.isVisible());
    } 
 
    /***************************************************************************
    **
    ** Show popup menu
    */
  
    void showPopup(Point pt) {
      prepareMenu();
      popMenu_.show(parent_, pt.x, pt.y);
      popMenu_.requestFocusInWindow();
      return;
    }
         
    /***************************************************************************
    **
    ** Prepare the link menu for display
    */
  
    void prepareMenu() { 
    	FabricDisplayOptions fdo = FabricDisplayOptionsManager.getMgr().getDisplayOptions();
    	boolean canShow = fdo.getOfferLinkBrowser();
      String browserURL = fdo.getBrowserLinkURL();
      lbl_.setEnabled(canShow && !browserURL.equals(""));
      return;
    }
  }

  /***************************************************************************
  **
  ** Listens for popup events
  */  
  
  private class PopupHandler implements PopupMenuListener {
    
    public void popupMenuCanceled(PopupMenuEvent e) {
    }
    
    public void popupMenuWillBecomeInvisible(PopupMenuEvent e) {
    }
    
    public void popupMenuWillBecomeVisible(PopupMenuEvent e) {
    }
  }
 
  /***************************************************************************
  **
  ** Command
  */ 
    
  public class LaunchBrowserForNode extends AbstractAction {
    
    private static final long serialVersionUID = 1L;
    
    public LaunchBrowserForNode(String text, char mnem) {
      super(text);
      putValue(Action.MNEMONIC_KEY, Integer.valueOf(mnem));
    }

    public void actionPerformed(ActionEvent e) {
      try {
      	FabricDisplayOptions fdo = FabricDisplayOptionsManager.getMgr().getDisplayOptions();
      	if (!fdo.getOfferNodeBrowser()) {
      		return;
      	}
        String browserURL = fdo.getBrowserURL();
        if (browserURL.indexOf(FabricDisplayOptions.NODE_NAME_PLACEHOLDER) == -1) {
          return;
        }
        browserURL = browserURL.replaceFirst(FabricDisplayOptions.NODE_NAME_PLACEHOLDER, currNode_.getName());
        (new BrowserLauncher()).openBBURL(browserURL);      
      } catch (Exception ex) {
        ExceptionHandler.getHandler().displayException(ex);
      }
      return;
    }
  }  
  
  /***************************************************************************
  **
  ** Command
  */ 
    
  public class LaunchBrowserForLink extends AbstractAction {
    
    private static final long serialVersionUID = 1L;
    
    public LaunchBrowserForLink(String text, char mnem) {
      super(text);
      putValue(Action.MNEMONIC_KEY, Integer.valueOf(mnem));
    }

    public void actionPerformed(ActionEvent e) {
      try {
      	FabricDisplayOptions fdo = FabricDisplayOptionsManager.getMgr().getDisplayOptions();
      	if (!fdo.getOfferLinkBrowser()) {
      		return;
      	}
        String browserURL = fdo.getBrowserLinkURL();
        if ((browserURL.indexOf(FabricDisplayOptions.LINK_SRC_PLACEHOLDER) == -1) ||
            (browserURL.indexOf(FabricDisplayOptions.LINK_TRG_PLACEHOLDER) == -1) ||
            (browserURL.indexOf(FabricDisplayOptions.LINK_REL_PLACEHOLDER) == -1)) {
          return;
        } else {          
          NetNode src = currLink_.getSrcNode();
          NetNode trg = currLink_.getTrgNode();
          String rel = currLink_.getAugRelation().relation;
          browserURL = browserURL.replaceFirst(FabricDisplayOptions.LINK_SRC_PLACEHOLDER, src.getName());
          browserURL = browserURL.replaceFirst(FabricDisplayOptions.LINK_TRG_PLACEHOLDER, trg.getName());
          browserURL = browserURL.replaceFirst(FabricDisplayOptions.LINK_REL_PLACEHOLDER, rel);
       }        
       (new BrowserLauncher()).openBBURL(browserURL);      
      } catch (Exception ex) {
        ExceptionHandler.getHandler().displayException(ex);
      }
      return;
    }
  }
}
```
