## Supplementary material for "BioFabric Visualization of Network Alignments": BioFabric Source Code: BoxPath.java

```
/*
**    Copyright (C) 2003-2017 Institute for Systems Biology 
**                            Seattle, Washington, USA. 
**
**    This library is free software; you can redistribute it and/or
**    modify it under the terms of the GNU Lesser General Public
**    License as published by the Free Software Foundation; either
**    version 2.1 of the License, or (at your option) any later version.
**
**    This library is distributed in the hope that it will be useful,
**    but WITHOUT ANY WARRANTY; without even the implied warranty of
**    MERCHANTABILITY or FITNESS FOR A PARTICULAR PURPOSE.  See the GNU
**    Lesser General Public License for more details.
**
**    You should have received a copy of the GNU Lesser General Public
**    License along with this library; if not, write to the Free Software
**    Foundation, Inc., 59 Temple Place, Suite 330, Boston, MA  02111-1307  USA
*/

package org.systemsbiology.biofabric.ui.render;

import java.awt.Color;
import java.awt.Graphics2D;
import java.awt.Rectangle;

/****************************************************************************
**
** This is a "Path" for drawing a box
*/

public class BoxPath {
 
  ////////////////////////////////////////////////////////////////////////////
  //
  // PUBLIC CONSTANTS
  //
  //////////////////////////////////////////////////////////////////////////// 
  
  private Color col;
  private Rectangle nodeShadeRect;
  private Rectangle scratchRect;
  
  ////////////////////////////////////////////////////////////////////////////
  //
  // PUBLIC CONSTRUCTORS
  //
  ////////////////////////////////////////////////////////////////////////////

  /***************************************************************************
  **
  ** Constructor
  */
  
  public BoxPath(Color col, Rectangle rect) {
    this.col = col;
    this.nodeShadeRect = rect;
    this.scratchRect = new Rectangle();
  }

  ////////////////////////////////////////////////////////////////////////////
  //
  // PUBLIC METHODS
  //
  ////////////////////////////////////////////////////////////////////////////

  /***************************************************************************
  **
  **  Draw it:
  */
 
  public int paint(Graphics2D g2, Rectangle bounds) {
    g2.setPaint(col);
    // No bounds? Rectangle fully in bounds? Just draw the actual rectangle!
    if ((bounds == null) || bounds.contains(nodeShadeRect)) {
      scratchRect.setRect(nodeShadeRect);
    } else {
      // Nothing to draw:
      if (!bounds.intersects(nodeShadeRect)) {
        return (0);
      } 
      // With very huge boxes (e.g. 100,000 units) we see fragmented draws on Mac. Make tiny:
      double xMinN = nodeShadeRect.getMinX();
      double xMaxN = nodeShadeRect.getMaxX();
      double yMinN = nodeShadeRect.getMinY();
      double yMaxN = nodeShadeRect.getMaxY();
      
      double xMinB = bounds.getMinX();
      double xMaxB = bounds.getMaxX();
      double yMinB = bounds.getMinY();
      double yMaxB = bounds.getMaxY();
      
      double xMinUse = (xMinN < xMinB) ? xMinB - 1000.0 : xMinN; 
      double xMaxUse = (xMaxN > xMaxB) ? xMaxB + 1000.0 : xMaxN; 
      double yMinUse = (yMinN < yMinB) ? yMinB - 1000.0 : yMinN; 
      double yMaxUse = (yMaxN > yMaxB) ? yMaxB + 1000.0 : yMaxN; 
      
      scratchRect.setRect(xMinUse, yMinUse, xMaxUse - xMinUse, yMaxUse - yMinUse);
    } 
    g2.fill(scratchRect);
    return (1);
  } 
} 
```
