## Supplementary material for "BioFabric Visualization of Network Alignments": BioFabric Source Code: BucketRenderer.java

```
/*
**    Copyright (C) 2003-2017 Institute for Systems Biology 
**                            Seattle, Washington, USA. 
**
**    This library is free software; you can redistribute it and/or
**    modify it under the terms of the GNU Lesser General Public
**    License as published by the Free Software Foundation; either
**    version 2.1 of the License, or (at your option) any later version.
**
**    This library is distributed in the hope that it will be useful,
**    but WITHOUT ANY WARRANTY; without even the implied warranty of
**    MERCHANTABILITY or FITNESS FOR A PARTICULAR PURPOSE.  See the GNU
**    Lesser General Public License for more details.
**
**    You should have received a copy of the GNU Lesser General Public
**    License along with this library; if not, write to the Free Software
**    Foundation, Inc., 59 Temple Place, Suite 330, Boston, MA  02111-1307  USA
*/

package org.systemsbiology.biofabric.ui.render;

import java.awt.Color;
import java.awt.Dimension;
import java.awt.Graphics2D;
import java.awt.Point;
import java.awt.geom.AffineTransform;
import java.awt.geom.Point2D;
import java.awt.geom.Rectangle2D;
import java.awt.image.BufferedImage;
import java.util.ArrayList;
import java.util.List;

import org.systemsbiology.biofabric.api.layout.AnnotColorSource;
import org.systemsbiology.biofabric.api.model.Annot;
import org.systemsbiology.biofabric.api.model.AnnotationSet;
import org.systemsbiology.biofabric.api.util.MinMax;
import org.systemsbiology.biofabric.model.AnnotationSetImpl;
import org.systemsbiology.biofabric.model.BioFabricNetwork;
import org.systemsbiology.biofabric.model.BioFabricNetwork.LinkInfo;
import org.systemsbiology.biofabric.model.BioFabricNetwork.NodeInfo;
import org.systemsbiology.biofabric.ui.BasicZoomTargetSupport;
import org.systemsbiology.biofabric.ui.FabricColorGenerator;
import org.systemsbiology.biofabric.ui.display.BioFabricPanel;
import org.systemsbiology.biofabric.util.UiUtil;

/****************************************************************************
**
** This is a fast network image builder for big networks.
*/

public class BucketRenderer implements BufBuildDrawer {
    
  ////////////////////////////////////////////////////////////////////////////
  //
  // PRIVATE INSTANCE MEMBERS
  //
  ////////////////////////////////////////////////////////////////////////////
  
  private List<BioFabricNetwork.NodeInfo> targetList_;
  private List<BioFabricNetwork.LinkInfo> linkList_;
  private BioFabricNetwork.Extents ext_;
  private AnnotationSet nodeAnnot_;
  private AnnotationSet linkAnnot_;
  
  private Dimension screenDim_;
  private boolean showShadows_;
  private Rectangle2D worldRect_;
  private ImgAndBufPool bis_;
  private AnnotColorSource.AnnotColor[] annotColors_;
  private Color[] linkAnnotGrays_;
  private final Color[] nodeCycle_;
  private final Color[] linkCycle_;
  
  private static final int LINK_MAX_ALPHA_ = 180;
  private static final int NODE_MAX_ALPHA_ = 180;


  ////////////////////////////////////////////////////////////////////////////
  //
  // PUBLIC CONSTRUCTORS
  //
  ////////////////////////////////////////////////////////////////////////////

  /***************************************************************************
  **
  ** Constructor
  */

  public BucketRenderer(FabricColorGenerator colGen) {

    targetList_ = new ArrayList<NodeInfo>();
    linkList_ = new ArrayList<LinkInfo>();
    ext_ = new BioFabricNetwork.Extents();
    worldRect_ = new Rectangle2D.Double(0.0, 0.0, 100.0, 100.0);
    nodeAnnot_ = new AnnotationSetImpl();
    linkAnnot_ = new AnnotationSetImpl();
    bis_ = null;
    
    //
    // Node color cycle sampled off of screen capture from regular renderer. This gives
    // regions of solid nodes a "cycling" pattern that has the same appearance of the
    // regular renderer. Same thing below for link cycle.
    //
    
    nodeCycle_ = new Color[] {
      new Color(145,141,126),
      new Color(194,201,213),
      new Color(160,141,129),
      new Color(177,191,189),
      new Color(162,129,134),
      new Color(189,166,148),
      new Color(163,162,157),
      new Color(148,163,165),
      new Color(182,164,165),
      new Color(144,155,140),
      new Color(202,160,178),
      new Color(133,144,129),
      new Color(208,183,185),
      new Color(153,142,127),
      new Color(215,193,204),
      new Color(145,141,125),
      new Color(194,201,213),
      new Color(160,141,129),
      new Color(186,168,166),
      new Color(164,147,137),
      new Color(166,150,153),
      new Color(175,168,152),
      new Color(161,170,148),
      new Color(184,166,159),
      new Color(144,156,141),
      new Color(202,160,178),
      new Color(133,144,129),
      new Color(209,190,194),
      new Color(165,125,143),
      new Color(219,200,203)
    };

    //
    // Link color cycle sampled off of screen capture from regular renderer:
    //
    
    linkCycle_ = new Color[] {
      new Color(123,104,111),
      new Color(92,92,90),
      new Color(129,124,117),
      new Color(99,95,93),
      new Color(137,122,128),
      new Color(109,106,84),
      new Color(133,119,125),
      new Color(103,95,101),
      new Color(130,106,109),
      new Color(106,100,96),
      new Color(113,95,100),
      new Color(109,108,102),
      new Color(114,110,101),
      new Color(115,103,108),
      new Color(99,90,102),
      new Color(124,114,116),
      new Color(96,83,103),
      new Color(130,126,123),
      new Color(111,85,105),
      new Color(139,125,129),
      new Color(109,106,84),
      new Color(117,114,121),
      new Color(106,90,95),
      new Color(126,104,98),
      new Color(105,107,104),
      new Color(102,105,112),
      new Color(111,105,107),
      new Color(102,98,112),
      new Color(119,106,109),
      new Color(99,90,102)
    };
    
    annotColors_ = AnnotColorSource.getColorCycle();
    linkAnnotGrays_ = AnnotColorSource.getGrayCycle();
  }

  ////////////////////////////////////////////////////////////////////////////
  //
  // PUBLIC METHODS
  //
  ////////////////////////////////////////////////////////////////////////////
  
  /***************************************************************************
  ** 
  ** Get the dimensions for the buffer
  */
  
   public void dimsForBuf(Dimension screenDim, Rectangle2D worldRect) {
  	 UiUtil.fixMePrintout("Seeing NPE here during recolor");
     screenDim.setSize(screenDim_);
     worldRect.setRect(worldRect_);    
     return;
   }

  /***************************************************************************
  ** 
  ** Install a model
  */
   
  public void buildBucketCache(List<BioFabricNetwork.NodeInfo> targets, 
  		                         List<BioFabricNetwork.LinkInfo> links,
  		                         AnnotationSet nodeAnnot, AnnotationSet linkAnnot,
  		                         BioFabricNetwork.Extents ext, boolean showShadows) { 
 
  	targetList_ = targets;
    linkList_ = links;
    ext_ = ext;
    nodeAnnot_ = nodeAnnot;
    linkAnnot_ = linkAnnot;
    showShadows_ = showShadows;
    return;
  }
  
  /***************************************************************************
  ** 
  ** Install a model size
  */
   
  public void setModelDims(Dimension screenDim, Rectangle2D worldRect, ImgAndBufPool bis) { 
    screenDim_ = screenDim;
    worldRect_.setRect(worldRect);
    bis_ = bis;
    return;
  }

  /***************************************************************************
  **
  ** Drawing core
  */
  
  public boolean drawForBuffer(BufferedImage bi, Rectangle2D clip, Dimension screenDim, 
  		                         Rectangle2D worldRec, int heightPad, double linksPerPixel) {
  	
  	int imgHeight = bi.getHeight();
    int imgWidth = bi.getWidth();
  	Graphics2D g2 = bi.createGraphics();
  	g2.setPaint(Color.WHITE);
    g2.fillRect(0, 0, imgWidth, imgHeight);
 
    if ((nodeAnnot_ != null) && nodeAnnot_.size() > 0){
	    BufferedImage bina = bis_.fetchImage(imgWidth, imgHeight, BufferedImage.TYPE_INT_ARGB);
	    drawNodeAnnotsForBuffer(bina, clip, screenDim, worldRec, heightPad, linksPerPixel);
	    g2.drawImage(bina, 0, 0, null);
	    bis_.returnImage(bina);
    }
    if ((linkAnnot_ != null) && linkAnnot_.size() > 0){
      BufferedImage binal = bis_.fetchImage(imgWidth, imgHeight, BufferedImage.TYPE_INT_ARGB);
      drawLinkAnnotsForBuffer(binal, clip, screenDim, worldRec, heightPad, linksPerPixel);
      g2.drawImage(binal, 0, 0, null);
    	bis_.returnImage(binal);
    }
    BufferedImage bin = bis_.fetchImage(imgWidth, imgHeight, BufferedImage.TYPE_INT_ARGB);
    drawNodesForBuffer(bin, clip, screenDim, worldRec, heightPad, linksPerPixel);
    g2.drawImage(bin, 0, 0, null);
  	BufferedImage bil = bis_.fetchImage(imgWidth, imgHeight, BufferedImage.TYPE_INT_ARGB);
  	drawLinksForBuffer(bil, clip, screenDim, worldRec, heightPad, linksPerPixel);
  	g2.drawImage(bil, 0, 0, null);
  	g2.dispose();
  	
  
  	bis_.returnImage(bin);
  	bis_.returnImage(bil);
  	
  	return (true);
  }

  /***************************************************************************
  **
  ** Drawing core
  */
  
  private boolean drawLinksForBuffer(BufferedImage bi, Rectangle2D clip, Dimension screenDim, Rectangle2D worldRec, int heightPad, double lpp) {
  	
    BufAndMeta bam = new BufAndMeta(bi, clip, screenDim, worldRec, heightPad, lpp, bis_);
    
	  int bufOffset = (bam.ulInV.x * bam.scrnHeight) + bam.ulInV.y;	  
	  
    for (BioFabricNetwork.LinkInfo lif : linkList_) {

	    bam.yStrt = lif.topRow() * BioFabricPanel.GRID_SIZE;
	    bam.yEnd = lif.bottomRow() * BioFabricPanel.GRID_SIZE;
	    bam.x = lif.getUseColumn(showShadows_) * BioFabricPanel.GRID_SIZE;
	   	if (!bam.clipForLinks(clip)) {
	    	continue;
	    }
	    int bufStart = (bam.startPtInV.x * bam.scrnHeight) + bam.startPtInV.y - bufOffset;
	    int bufEnd = (bam.endPtInV.x * bam.scrnHeight) + bam.endPtInV.y - bufOffset;
	    
	    bam.transToBuf(bufStart, bufEnd);
    }
 
    for (int i = 0; i < bam.bufLen; i++) {
    	int pix = bam.mybuf[i];
    	
    	int red = 0;
    	int green = 0;
    	int blue = 0;
    	int alpha = 0; 	
    	
    	int xval = i / bam.scrnHeight;
      
      if (pix != 0) {
        double val = pix / lpp; 
        int index = xval % linkCycle_.length;       
        red = linkCycle_[index].getRed();
        green = linkCycle_[index].getGreen();
        blue = linkCycle_[index].getBlue();
        alpha = Math.min(LINK_MAX_ALPHA_, (int)Math.round(val * LINK_MAX_ALPHA_));
      }
    	
      int rgb = alpha << 24 | red << 16 | green << 8 | blue;
      
    	if (xval >= bam.imgWidth) {
    		continue;
    	}
    	int yval = i % bam.scrnHeight;
    	if (yval >= bam.imgHeight) {
    		continue;
    	}
      bi.setRGB(xval, yval, rgb);
	  }
    
    bis_.returnBuf(bam.mybuf);
    return (true);
  }
 
  /***************************************************************************
  **
  ** Drawing core
  */
  
  private boolean drawNodesForBuffer(BufferedImage bi, Rectangle2D clip, Dimension screenDim, Rectangle2D worldRec, int heightPad, double lpp) {
  
  	BufAndMeta bam = new BufAndMeta(bi, clip, screenDim, worldRec, heightPad, lpp, bis_);

	  int bufOffset = (bam.ulInV.y * bam.scrnWidth) + bam.ulInV.x;	  
	  
    for (BioFabricNetwork.NodeInfo nif : targetList_) {

	    MinMax colRange = nif.getColRange(showShadows_);	    				
	    bam.xStrt = colRange.min * BioFabricPanel.GRID_SIZE;
	    bam.xEnd = colRange.max * BioFabricPanel.GRID_SIZE;
	    bam.y = nif.nodeRow * BioFabricPanel.GRID_SIZE;
	    if (!bam.clipForNodes(clip)) {
	    	continue;
	    }
	    int bufStart = (bam.startPtInV.y * bam.scrnWidth) + bam.startPtInV.x - bufOffset;
	    int bufEnd = (bam.endPtInV.y * bam.scrnWidth) + bam.endPtInV.x - bufOffset;
	       
	    bam.transToBuf(bufStart, bufEnd);
    }
    
    for (int i = 0; i < bam.bufLen; i++) {
    	int pix = bam.mybuf[i];
    	
    	int red = 0;
    	int green = 0;
    	int blue = 0;
    	int alpha = 0;
    	
    	int yval = i / bam.scrnWidth;
    	
    	if (pix != 0) {
    	  double val = pix / lpp; 
    	  int index = yval % nodeCycle_.length;    	  
    	  red = nodeCycle_[index].getRed();
    	  green = nodeCycle_[index].getGreen();
    	  blue = nodeCycle_[index].getBlue();
    	  alpha = Math.min(NODE_MAX_ALPHA_, (int)Math.round(val * NODE_MAX_ALPHA_));
    	}
    	
      int rgb = alpha << 24 | red << 16 | green << 8 | blue;
    //  mybuf[i] = rgb;
    	
    	
    	if (yval >= bam.imgHeight) {
    		continue;
    	}
    	int xval = i % bam.scrnWidth;
    	if (xval >= bam.imgWidth) {
    		continue;
    	}
      bi.setRGB(xval, yval, rgb);
	  }
    // Use this instead:
    // int[] a = ((DataBufferInt)bi.getRaster().getDataBuffer()).getData();
    // System.arraycopy(mybuf, 0, a, 0, Math.min(mybuf.length, a.length));
    bis_.returnBuf(bam.mybuf);
    return (true);
  }
  
  /***************************************************************************
  **
  ** Drawing core
  */
  
  private boolean drawNodeAnnotsForBuffer(BufferedImage bi, Rectangle2D clip, Dimension screenDim, Rectangle2D worldRec, int heightPad, double lpp) {
  
  	BufAndMeta bam = new BufAndMeta(bi, clip, screenDim, worldRec, heightPad, lpp, bis_);

	  int bufOffset = (bam.ulInV.y * bam.scrnWidth) + bam.ulInV.x;	  
	  
	  MinMax linkCols = ext_.allLinkFullRange.get(Boolean.valueOf(showShadows_));
	  int pad = PaintCacheSmall.calcAnnotationPad(linkCols);
	  
	  int colNum = 0;
    for (Annot annot : this.nodeAnnot_) {
	    MinMax rowRange = annot.getRange();
      for (int i = rowRange.min; i <= rowRange.max; i++) {
		    bam.xStrt = linkCols.min * BioFabricPanel.GRID_SIZE;
		    bam.xEnd = linkCols.max * BioFabricPanel.GRID_SIZE;
		    bam.y = i * BioFabricPanel.GRID_SIZE;
		    if (!bam.clipForNodes(clip)) {
		    	continue;
		    }
		    int bufStart = (bam.startPtInV.y * bam.scrnWidth) + bam.startPtInV.x - bufOffset;
		    int bufEnd = (bam.endPtInV.y * bam.scrnWidth) + bam.endPtInV.x - bufOffset;
		      
		    AnnotColorSource.AnnotColor acol = annot.getColor();
		    int useColorNum = (acol == null) ? colNum : acol.getCycle();
		    
		    bam.transColorToBuf(bufStart, bufEnd, useColorNum);
      }
      colNum = (colNum + 1) % annotColors_.length;
    }
    
    for (int i = 0; i < bam.bufLen; i++) {
    	int val = bam.mybuf[i];
    	if ((val == -1) || (val == 0)) {
    		continue;
    	}
    	Color col = annotColors_[val - 1].getColor();
    	int red = col.getRed();
    	int green = col.getGreen();
    	int blue = col.getBlue();
 
      int rgb = 255 << 24 | red << 16 | green << 8 | blue;
    //  mybuf[i] = rgb;
    	int yval = i / bam.scrnWidth;
    	
    	if (yval >= bam.imgHeight) {
    		continue;
    	}
    	int xval = i % bam.scrnWidth;
    	if (xval >= bam.imgWidth) {
    		continue;
    	}

      bi.setRGB(xval, yval, rgb);
	  }
    // Use this instead:
    // int[] a = ((DataBufferInt)bi.getRaster().getDataBuffer()).getData();
    // System.arraycopy(mybuf, 0, a, 0, Math.min(mybuf.length, a.length));
    bis_.returnBuf(bam.mybuf);
    return (true);
  }
  
  /***************************************************************************
  **
  ** Drawing core
  */
  
  private boolean drawLinkAnnotsForBuffer(BufferedImage bi, Rectangle2D clip, 
                                          Dimension screenDim, Rectangle2D worldRec, 
                                          int heightPad, double lpp) {
  
  	BufAndMeta bam = new BufAndMeta(bi, clip, screenDim, worldRec, heightPad, lpp, bis_);

	  int bufOffset = (bam.ulInV.y * bam.scrnWidth) + bam.ulInV.x;	  
	  
	  MinMax nodeRows = ext_.allNodeFullRange.get(Boolean.valueOf(showShadows_));
	  int pad = PaintCacheSmall.calcAnnotationPad(nodeRows);
	  
	  Color[] useColors;
	  if ((nodeAnnot_ != null) && (nodeAnnot_.size() > 0)) {
	  	useColors = linkAnnotGrays_;
	  	
	  } else {
	  	useColors = new Color[annotColors_.length];
	  	for (int i = 0; i < annotColors_.length; i++) {
	  		useColors[i] = annotColors_[i].getColor();
	  	}
	  }
	  
	  int colNum = 0;
    for (Annot annot : linkAnnot_) {
	    MinMax colRange = annot.getRange();
      for (int i = nodeRows.min; i <= nodeRows.max; i++) {
		    bam.xStrt = colRange.min * BioFabricPanel.GRID_SIZE;
		    bam.xEnd = colRange.max * BioFabricPanel.GRID_SIZE;
		    bam.y = i * BioFabricPanel.GRID_SIZE;
		    if (!bam.clipForNodes(clip)) {
		    	continue;
		    }
		    int bufStart = (bam.startPtInV.y * bam.scrnWidth) + bam.startPtInV.x - bufOffset;
		    int bufEnd = (bam.endPtInV.y * bam.scrnWidth) + bam.endPtInV.x - bufOffset;
		      
        AnnotColorSource.AnnotColor acol = annot.getColor();
        int useColorNum = (acol == null) ? colNum : acol.getCycle();

		    bam.transColorToBuf(bufStart, bufEnd, useColorNum);
      }
      colNum = (colNum + 1) % useColors.length;
    }
    
    for (int i = 0; i < bam.bufLen; i++) {
    	int val = bam.mybuf[i];
    	if ((val == -1) || (val == 0)) {
    		continue;
    	}
    	int red = useColors[val - 1].getRed();
    	int green = useColors[val - 1].getGreen();
    	int blue = useColors[val - 1].getBlue();
    	int alpha = useColors[val - 1].getAlpha();
 
      int rgb = alpha << 24 | red << 16 | green << 8 | blue;
    //  mybuf[i] = rgb;
    	int yval = i / bam.scrnWidth;
    	
    	if (yval >= bam.imgHeight) {
    		continue;
    	}
    	int xval = i % bam.scrnWidth;
    	if (xval >= bam.imgWidth) {
    		continue;
    	}

      bi.setRGB(xval, yval, rgb);
	  }
    // Use this instead:
    // int[] a = ((DataBufferInt)bi.getRaster().getDataBuffer()).getData();
    // System.arraycopy(mybuf, 0, a, 0, Math.min(mybuf.length, a.length));
    bis_.returnBuf(bam.mybuf);
    return (true);
  }
  
  ////////////////////////////////////////////////////////////////////////////
  //
  // PRIVATE INNER CLASSES
  //
  ////////////////////////////////////////////////////////////////////////////

  private static class BufAndMeta {
  	int imgHeight;
    int imgWidth;

    double zoomH;
    double zoomV;
    double zoom;
    AffineTransform transform;
     
    int scrnHeight;
    int scrnWidth;
     
    int bufLen;
    
    Point2D newPoint;
  	int[] mybuf;
	
  	Point pts;
    Point pte;    
    Point startPoint;
	  Point endPoint;
	  Point ulInV;
	  
	  Point startPtInV;
	  Point endPtInV;
	  
	  // For link drawing:
	  double yStrt;
	  double yEnd;
	  double x;
	  
	  // for node drawing:
	  double xStrt;
	  double xEnd;
	  double y;
 
	  BufAndMeta(BufferedImage bi, Rectangle2D clip, Dimension screenDim, Rectangle2D worldRec, 
	  		       int heightPad, double lpp, ImgAndBufPool bis) {
  	
	  	imgHeight = bi.getHeight();
	    imgWidth = bi.getWidth();
		
	    zoomH = screenDim.getWidth() / worldRec.getWidth();
	    zoomV = screenDim.getHeight() / worldRec.getHeight();
	    zoom = Math.max(zoomH, zoomV);
	    Point2D centerW = new Point2D.Double(worldRec.getX() + (worldRec.getWidth() / 2.0), worldRec.getY() + (worldRec.getHeight() / 2.0));
	    transform = new AffineTransform();
	    transform.translate(screenDim.getWidth() / 2.0, screenDim.getHeight() / 2.0);
	    transform.scale(zoom, zoom);
	    transform.translate(-centerW.getX(), -centerW.getY());
	      
	    scrnHeight = screenDim.height;
	    scrnWidth = screenDim.width;
	     
	    bufLen = scrnHeight * scrnWidth;
	    
	    newPoint = new Point2D.Double();
	   
	    mybuf = bis.fetchBuf(bufLen);
	    
	    pts = new Point();
	    pte = new Point();    
	    startPoint = new Point();
		  endPoint = new Point();
		  
		  startPoint.setLocation(worldRec.getX(), worldRec.getY());
		  ulInV = BasicZoomTargetSupport.pointToViewport(startPoint, transform, newPoint,pts); 
	  }
	  
	  boolean clipForLinks(Rectangle2D clip) {
	  	if ((x < clip.getX()) || (x > (clip.getX() + clip.getWidth()))) {
	    	return (false);
	    }
	    if (yStrt < clip.getY()) {
	    	yStrt = clip.getY();
	    }
	    if (yStrt > (clip.getY() + clip.getHeight())) {
	    	yStrt = clip.getY() + clip.getHeight();
	    }
	    if (yEnd < clip.getY()) {
	    	yEnd = clip.getY();
	    }
	    if (yEnd > (clip.getY() + clip.getHeight())) {
	    	yEnd = clip.getY() + clip.getHeight();
	    }
	
	    startPoint.setLocation(x, yStrt);
	    endPoint.setLocation(x, yEnd);
	        
	    startPtInV = BasicZoomTargetSupport.pointToViewport(startPoint, transform, newPoint, pts);
	    endPtInV = BasicZoomTargetSupport.pointToViewport(endPoint,transform, newPoint, pte);
	       
	    return (true);
	  }  
	  
	  boolean clipForNodes(Rectangle2D clip) {
	  	if ((y < clip.getY()) || (y > (clip.getY() + clip.getHeight()))) {
	    	return (false);
	    }
	    if (xStrt < clip.getX()) {
	    	xStrt = clip.getX();
	    }
	    if (xStrt > (clip.getX() + clip.getWidth())) {
	    	xStrt = clip.getX() + clip.getWidth();
	    }
	    if (xEnd < clip.getX()) {
	    	xEnd = clip.getX();
	    }
	    if (xEnd > (clip.getX() + clip.getWidth())) {
	    	xEnd = clip.getX() + clip.getWidth();
	    }   
	    startPoint.setLocation(xStrt, y);
	    endPoint.setLocation(xEnd, y);
	    
	    startPtInV = BasicZoomTargetSupport.pointToViewport(startPoint, transform, newPoint, pts);
	    endPtInV = BasicZoomTargetSupport.pointToViewport(endPoint,transform, newPoint, pte);
	       
	    return (true);
	  } 
	  
	  void transToBuf(int bufStart, int bufEnd) {
	  	for (int i = bufStart; i < bufEnd; i++) {
	    	int off = i;
	    	if (off < 0) {
	    		continue;
	    	}
	    	if (off >= mybuf.length) {
          continue;	
	    	}
	    	mybuf[off] += 1;    	
	    }
	  	return;
	  }
	  
	  void transColorToBuf(int bufStart, int bufEnd, int color) {
	  	for (int i = bufStart; i < bufEnd; i++) {
	    	int off = i;
	    	if (off < 0) {
	    		continue;
	    	}
	    	if (off >= mybuf.length) {
          continue;	
	    	}
	    	int currCol = mybuf[off];
	    	int slotCol = color + 1;
	    	if ((currCol == 0) || (currCol == slotCol)) {
	    	  mybuf[off] = slotCol;
	    	} else {
	    		mybuf[off] = -1;
	    	}
	    }
	  	return;
	  }  
  }  
}
```
