## Supplementary material for "BioFabric Visualization of Network Alignments": BioFabric Source Code: BufferBuilder.java

```
/*
**    Copyright (C) 2003-2018 Institute for Systems Biology 
**                            Seattle, Washington, USA. 
**
**    This library is free software; you can redistribute it and/or
**    modify it under the terms of the GNU Lesser General Public
**    License as published by the Free Software Foundation; either
**    version 2.1 of the License, or (at your option) any later version.
**
**    This library is distributed in the hope that it will be useful,
**    but WITHOUT ANY WARRANTY; without even the implied warranty of
**    MERCHANTABILITY or FITNESS FOR A PARTICULAR PURPOSE.  See the GNU
**    Lesser General Public License for more details.
**
**    You should have received a copy of the GNU Lesser General Public
**    License along with this library; if not, write to the Free Software
**    Foundation, Inc., 59 Temple Place, Suite 330, Boston, MA  02111-1307  USA
*/

package org.systemsbiology.biofabric.ui.render;

import java.awt.Dimension;
import java.awt.Rectangle;
import java.awt.geom.Rectangle2D;
import java.awt.image.BufferedImage;
import java.awt.image.DataBufferInt;
import java.awt.image.DirectColorModel;
import java.awt.image.WritableRaster;
import java.io.IOException;
import java.text.MessageFormat;
import java.util.ArrayList;
import java.util.HashMap;
import java.util.List;
import javax.swing.SwingUtilities;

import org.systemsbiology.biofabric.api.worker.AsynchExitRequestException;
import org.systemsbiology.biofabric.api.worker.BTProgressMonitor;
import org.systemsbiology.biofabric.api.worker.LoopReporter;
import org.systemsbiology.biofabric.ui.display.BioFabricPanel;
import org.systemsbiology.biofabric.util.QuadTree;
import org.systemsbiology.biofabric.util.UiUtil;

/****************************************************************************
**
** This builds and manages image chunks
*/

public class BufferBuilder {
  
  ////////////////////////////////////////////////////////////////////////////
  //
  // PRIVATE CONSTANTS
  //
  //////////////////////////////////////////////////////////////////////////// 

  //
  // Tweak the slice height up by one, else rounding/trunc errors introduce white
  // gridding on the image mosaic!
  //
  
  private static final int SLICE_HEIGHT_HACK_ = 1;
  
  //
  // At what level of links per pixel to we transition from bin renderer to drawing renderer?
  //
  
  private static final double TRANSITION_LPP_ = 20.0;
  
  ////////////////////////////////////////////////////////////////////////////
  //
  // PUBLIC CONSTANTS
  //
  //////////////////////////////////////////////////////////////////////////// 
  
  ////////////////////////////////////////////////////////////////////////////
  //
  // PRIVATE INSTANCE MEMBERS
  //
  ////////////////////////////////////////////////////////////////////////////
  
  private RasterCache cache_;
  private HashMap<Rectangle2D, WorldPieceOffering> allWorldsToImageName_;
  private QuadTree findWorldsQT_;
  private BufBuildDrawer drawRender_;
  private BufBuildDrawer binRender_;
  private int[] bbZooms_;
  private Dimension screenDim_;
  private Rectangle2D worldRect_;
  private BufferBuilderClient bbc_;
  private boolean timeToExit_;
  private BuildImageWorker biw_;
  private ImgAndBufPool bis_;

  
  ////////////////////////////////////////////////////////////////////////////
  //
  // PUBLIC CONSTRUCTORS
  //
  ////////////////////////////////////////////////////////////////////////////

  /***************************************************************************
  **
  ** Constructor
  */

  public BufferBuilder(String cachePref, int maxMeg, BufBuildDrawer drawRender, 
  		                 BufBuildDrawer binRender, ImgAndBufPool bis) {
  	BufferedImage forModel = new BufferedImage(1, 1, BufferedImage.TYPE_INT_RGB);
  	DirectColorModel dcm = (DirectColorModel)forModel.getColorModel();
    cache_ = new RasterCache(cachePref, maxMeg, dcm);
    allWorldsToImageName_ = new HashMap<Rectangle2D, WorldPieceOffering>();
    findWorldsQT_ = null;
    drawRender_ = drawRender;
    binRender_ = binRender;
    bbc_ = null;
    timeToExit_ = false;
    bis_ = bis;
  }
  
  /***************************************************************************
  **
  ** Constructor
  */

  public BufferBuilder(BufBuildDrawer drawRender, BufBuildDrawer binRender, ImgAndBufPool bis) {
    drawRender_ = drawRender;
    binRender_ = binRender;
    allWorldsToImageName_ = new HashMap<Rectangle2D, WorldPieceOffering>();
    findWorldsQT_ = null;
    bbc_ = null;
    timeToExit_ = false;
    bis_ = bis;
  }

  ////////////////////////////////////////////////////////////////////////////
  //
  // PUBLIC METHODS
  //
  ////////////////////////////////////////////////////////////////////////////
    
  /***************************************************************************
  **
  ** Kill it off
  */
  
  public void release() {
    synchronized (this) {
      timeToExit_ = true;
      bbc_ = null;
      this.notify();
    }
    if (findWorldsQT_ != null) {
    	findWorldsQT_.clear();
    }
    cache_.releaseResources();
    return;
  }
  
  /***************************************************************************
  **
  ** Simple overview one-shot
  */
  
  public BufferedImage buildOneBuf() {
    screenDim_ = new Dimension();
    worldRect_ = new Rectangle2D.Double();
    binRender_.dimsForBuf(screenDim_, worldRect_); // Both will give same answer... 
    Rectangle worldPiece = UiUtil.rectFromRect2D(worldRect_);
    BufferedImage bi = bis_.fetchImage(screenDim_.width, screenDim_.height, BufferedImage.TYPE_INT_RGB);
    double lpp = linksPerPix(screenDim_, worldPiece);
    BufBuildDrawer useDrawer = (lpp < TRANSITION_LPP_) ? drawRender_ : binRender_;
    useDrawer.drawForBuffer(bi, worldPiece, screenDim_, worldPiece, 0, lpp);  
    return (bi); 
  }
 
  /***************************************************************************
  **
  ** Get our slices in world to cover the given view in the world at the desired depth
  */
  
  public void getSlicesToCover(int depth, Rectangle2D viewInWorld, List<Rectangle2D>slicesToCover) {
  
  	ArrayList<QuadTree.QuadTreeNode> qtnList = new ArrayList<QuadTree.QuadTreeNode>();
  	findWorldsQT_.getNodes(viewInWorld, depth, qtnList);
  	int lsiz = qtnList.size();
  	for (int i = 0; i < lsiz; i++) {
  		QuadTree.QuadTreeNode qtn = qtnList.get(i);
      slicesToCover.add(qtn.getWorldExtent());
    }
  	return;
  }
  
  /***************************************************************************
  **
  **  Build all the bufs
  */
  
  public BufferedImage buildBufs(int[] zooms, BufferBuilderClient bbc, int maxSize, 
  		                           BTProgressMonitor monitor) throws IOException, AsynchExitRequestException {
    timeToExit_ = false;
    bbZooms_ = new int[zooms.length];
    System.arraycopy(zooms, 0, bbZooms_, 0, zooms.length);
    screenDim_ = new Dimension();
    worldRect_ = new Rectangle2D.Double();
    drawRender_.dimsForBuf(screenDim_, worldRect_); // These values are now ours
    Rectangle worldPiece = UiUtil.rectFromRect2D(worldRect_);   
    findWorldsQT_ = new QuadTree(worldPiece, zooms.length);
         
    //
    // Build the first two zoom levels before we even get started:
    //
    
    List<QueueRequest> requestQueuePre = buildQueue(0, 1, 10); 
    LoopReporter lr = new LoopReporter(requestQueuePre.size(), 20, monitor, 0.0, 1.0, "progress.stockingImageBufferTop");
    while (!requestQueuePre.isEmpty()) {
      QueueRequest qr = requestQueuePre.remove(0);
      lr.report();
      buildBuffer(new Dimension(qr.imageDim.width, qr.imageDim.height), qr);
    }
    
    //
    // Now build up the requests for the background thread:
    //   
    
    int useMax = 1; //maxSize; 
    List<QueueRequest> requestQueue = (zooms.length > 2) ? buildQueue(2, zooms.length - 1, useMax) : new ArrayList<QueueRequest>();
 
    bbc_ = bbc;
 
    if (!requestQueue.isEmpty()) {
      biw_ = new BuildImageWorker(screenDim_, requestQueue);
      Thread runThread = new Thread(biw_);
      runThread.setPriority(runThread.getPriority() - 2);
      runThread.start();
    }

    return (getTopImage()); 
  }

  /***************************************************************************
  **
  ** Get the top buffered image
  */
  
  public BufferedImage getTopImage() throws IOException {
  	ArrayList<QuadTree.QuadTreeNode> nodes = new ArrayList<QuadTree.QuadTreeNode>();
  	boolean found = findWorldsQT_.getAllNodesToDepth(0, 0, nodes);
  	if (!found || (nodes.size() != 1)) {
  		throw new IOException();
  	}
    WorldPieceOffering wpo = allWorldsToImageName_.get(nodes.get(0).getWorldExtent());
    if (wpo == null) { // After a new network is created....
    	return (null);
    }
    BufferedImage retval = null;
    
    //
    // It is true there are two locks floating around, one on this object, and one of the
    // bis_ BufImgStack. But since bis_ does not acquire any locks during its methods, we
    // do not need to worry about deadlock conditions.
    //
    synchronized (this) {
      retval = cache_.getAnImage(wpo.cacheHandle, bis_);
    }
    return (retval);
  }

  /***************************************************************************
  **
  ** Get an image at the given depth
  */
  
  public BufferedImage getImageForPiece(int depth, Rectangle2D worldRect) throws IOException {
  	WorldPieceOffering wpo = allWorldsToImageName_.get(worldRect);
  	if (wpo == null) {
  		wpo = new WorldPieceOffering(null, screenDim_, worldRect, false);
  		allWorldsToImageName_.put(worldRect, wpo);;
  	}
  	boolean needLoRes = false;
    synchronized (this) {  	
      needLoRes = (wpo.cacheHandle == null);
    }
    // Yeah, this could be stale. Not the end of the world though, and we are not going to
    // hold the lock until the lo-res slice is built...
    if (needLoRes) {
    	UiUtil.fixMePrintout("RACE CONDITION! Make sure low-res does not replace a high-res if it gets done second");
      buildLoResSlice(worldRect, wpo);
      if (biw_ != null) {
        biw_.bumpRequest(new QueueRequest(depth, screenDim_, worldRect));
      }
    }
    BufferedImage retval = null;
    synchronized (this) {
      if ((wpo.cacheHandle != null) && !wpo.cacheHandle.equals("")) {
        retval = cache_.getAnImage(wpo.cacheHandle, bis_);
      }
    }
    return (retval);
  }
 
  /***************************************************************************
  **
  ** Build the request queue
  */
  
  private List<QueueRequest> buildQueue(int startIndex, int endIndex, int maxCount)  {
    ArrayList<QueueRequest> retval = new ArrayList<QueueRequest>();
    ArrayList<QuadTree.QuadTreeNode> nodes = new ArrayList<QuadTree.QuadTreeNode>();
    findWorldsQT_.getAllNodesToDepth(startIndex, endIndex, nodes);
    Rectangle2D emptyExtent = new Rectangle2D.Double(0.0, 0.0, 0.0, 0.0);
    int numWorlds = nodes.size();
    for (int i = 0; i < numWorlds; i++) {
    	QuadTree.QuadTreeNode node = nodes.get(i);
    	Rectangle2D worldExtent = node.getWorldExtent();
    	if (worldExtent.equals(emptyExtent)) { // This is an empty model...
    		continue;
    	}
      WorldPieceOffering wpo = allWorldsToImageName_.get(worldExtent);
      if (wpo != null) {
      	UiUtil.fixMePrintout("Cancel of relayout puts us here");
      	System.err.println("Dup " + worldExtent);
      	throw new IllegalStateException();
      }
      wpo = new WorldPieceOffering(null, screenDim_, worldExtent, false);
      allWorldsToImageName_.put(worldExtent, wpo);        
      retval.add(new QueueRequest(node.getDepth(), screenDim_, wpo.worldRect));
      if (retval.size() >= maxCount) {
        return (retval);
      }
    }
    return (retval);
  }  

  /***************************************************************************
  **
  ** Build the buffers
  */
  
  private boolean buildBuffer(Dimension imageDim, QueueRequest qr) throws IOException {     
    //
    // To chunk the image, we parcel out pieces of world to pieces of screen:
    //
    synchronized (this) {
      if (timeToExit_) {
        return (false);
      }
    }
    buildHiResSlice(imageDim, qr.depth, qr.worldPiece);     
    return (true);
  }  
   
  /***************************************************************************
  **
  ** If we lack an image slice, we first create a lo-res version from existing images. NOTE
  ** THAT THIS IS RUNNING ON THE AWT THREAD.
  ** 
  */
  
  private String buildLoResSlice(Rectangle2D worldRect, WorldPieceOffering wpo) throws IOException {
  	// Useful to track memory usage:
    //long preRend = Runtime.getRuntime().freeMemory();  
    //System.out.println("preRend " + preRend);
    ArrayList<QuadTree.QuadTreeNode> path = new  ArrayList<QuadTree.QuadTreeNode>() ;  
    BufferedImage bi1 = null;

    if (!findWorldsQT_.getPath(worldRect, path)) {
    	throw new IllegalStateException();
    }
    
    //
    // Crank backwards up the chain to find an image to use:
    //
    
    WorldPieceOffering wpou = null;
    int pathLen = path.size();
    for (int i = pathLen - 1; i >= 0; i--) {
    	QuadTree.QuadTreeNode node = path.get(i);
      synchronized (this) {
        wpou = allWorldsToImageName_.get(node.getWorldExtent());  
        if ((wpou != null) && (wpou.cacheHandle != null) && !wpou.cacheHandle.equals("")) {
          bi1 = cache_.getAnImage(wpou.cacheHandle, bis_);
          break;
        }
      }
      
    }

    if (bi1 == null) {  // blank!
      return (null);
    }

    //
    // The piece of the image we use depends on how the target world rect fits inside the
    // world rect of the image we are using:
    //
    
    double subxFrac = (worldRect.getX() - wpou.worldRect.getX()) / wpou.worldRect.getWidth();
    int subxLoc = (int)Math.round(subxFrac * screenDim_.getWidth());
    double subyFrac = (worldRect.getY() - wpou.worldRect.getY()) / wpou.worldRect.getHeight();
    int subyLoc = (int)Math.round(subyFrac * screenDim_.getHeight());
    int subW = (int)Math.round((worldRect.getWidth() / wpou.worldRect.getWidth()) * bi1.getWidth());
    int subH = (int)Math.round((worldRect.getHeight() / wpou.worldRect.getHeight()) * bi1.getHeight());  
    
    //
    // Previous version created a BufferedImage from a chunk of the bigger image, which was not
    // so terrible, since it shared the data buffer. But we used a rendering context to draw the
    // piece into a full-size image, which generated large (multi-megabyte) piles of garbage.
    //
    // So now we do the data manipulation directly, pulling the bytes out of the small piece
    // and laying them into the larger byte array to scale them up.
    //

    // First, pull the smaller byte chunk out of the image we are going to enlarge:
    WritableRaster wr = bi1.getRaster();
    int pixNum = subW * subH;

    
    //
    // OK, should do scaling with int array, but stick with old code for the moment:
    //
    RasterCache.ShiftData sd = new RasterCache.ShiftData((DirectColorModel)bi1.getColorModel());
    UiUtil.fixMePrintout("OK, should do scaling with original int array, but stick with old code for the moment");
    int[] bbcI = bis_.fetchBuf(pixNum);
    wr.getDataElements(subxLoc, subyLoc, subW, subH, bbcI); 
    int smallLen = pixNum * 3;
    byte[] bbc = bis_.fetchByteBuf(smallLen);
    RasterCache.oneIntToThreeBytes(bbcI, bbc, sd);
    bis_.returnBuf(bbcI);
     
    //
    // Get the image we are going to produce:
    //
    
    BufferedImage scaled = bis_.fetchImage(screenDim_.width, screenDim_.height + SLICE_HEIGHT_HACK_, BufferedImage.TYPE_INT_RGB);
    int scaledIntBufSize = scaled.getRaster().getDataBuffer().getSize();
    byte[] bbs = bis_.fetchByteBuf(scaledIntBufSize * 3);    
    
    for (int i = 0; i < bbs.length; i++) {
    	bbs[i] = (byte)255;
    }
    
    //
    // Data in small-size bbc needs to be laid out in larger bbs buffer and duplicated:
    //
    // For a doubling, we copy each pixel twice, skipping rows in the target array. Then we duplicate the row
    // into the adjacent row. For 4X scaling, we do the same, but duplicate pixel four times, write it every four
    // rows, and copy the line three times. Same pattern with higher order scaling...
    //
    
    int factor = (int)Math.floor(screenDim_.width / subW);
    int facSq = factor * factor;
    
    for (int i = 0; i < subH; i++) { // do this for each row...
    	for (int j = 0; j < subW; j++) { // do this for each pixel in the row...
    		int currOffSmall = (i * (subW * 3)) + (j * 3); // This is always the same regardless of scaling...
    		int currOffLargeBase = ((i * ((subW * 3) * facSq)) + (j * 3 * factor));
    		int dupOffset = 0;
        for (int k = 0; k < factor; k++) { // Duplicate this pixel into the target...
          int currOffLargeDup = currOffLargeBase + dupOffset;
          dupOffset += 3;
	    		for (int m = 0; m < 3; m++) { // for the three channels of the pixel...
	    			byte b = bbc[currOffSmall + m];
	    	    bbs[currOffLargeDup + m] = b;
	    	  }
        }
    	}
    }
    //
    // Now duplicate each row:
    //
      
    for (int i = 0; i < screenDim_.height; i += factor) { // do this for each row...
    	int currOffBase = i * screenDim_.width * 3;
      for (int k = 1; k < factor; k++) { //duplicate the row...
      	int currOffDest = (i + k) * screenDim_.width * 3;
      	try {
  			  System.arraycopy(bbs, currOffBase, bbs, currOffDest, screenDim_.width * 3);
        } catch (ArrayIndexOutOfBoundsException aex) {
        	
        	UiUtil.fixMePrintout("Have seen array out of bounds here (collatz.sif->Hierarchical)!");
        	//ik 896 10
          //bbs 4,350,000
          //col 4,300,800
          //cold 4,348,800 (+ 1600 * 3) = 4,353,600
 
        	System.err.println("factor " + factor + " " + screenDim_.width + " " + subW);
    		  System.err.println("ik " + i + " " + k);
    		  System.err.println("bbs " + bbs.length);
    		  System.err.println("col " + currOffBase);
    		  System.err.println("cold " + currOffDest);
    		  i = 10000000;
    		  break;
    	  }
    	}
    }

    //
    // Write it into the destination:
    //
    
    WritableRaster bisRast = scaled.getRaster();
    
    //
    // We need an array of ints to set here:
    //
    int[] bbsI = bis_.fetchBuf(scaledIntBufSize);
    RasterCache.threeBytesToOneInt(bbs, bbsI, sd);
 
  	bisRast.setDataElements(0, 0, screenDim_.width, screenDim_.height + SLICE_HEIGHT_HACK_, bbsI);
 
    String handle = null;
    synchronized (this) {
      if (!wpo.isDrawn) {
        if (!isBlankImage(scaled)) {
        	// Caching recycles the image
          wpo.cacheHandle = cache_.cacheAnImage(scaled, bis_);
        } else {
          wpo.cacheHandle = "";
          // gotta manually recycle the image:
          bis_.returnImage(scaled);
        }
      } 
    }
    bis_.returnImage(bi1);
    bis_.returnByteBuf(bbc);
    bis_.returnByteBuf(bbs);
    bis_.returnBuf(bbsI);
    
    // Useful to track memory usage:
    //long po = Runtime.getRuntime().freeMemory();
    //long used = preRend - po;
    //System.out.println("Post getSub " + po);
    //System.out.println("rendering new scaled image used: " + used);
    return (handle);
  }
  
  /***************************************************************************
  **
  ** Works (not very well) for doubling scale
  */
  
  private void worksForDoubling(int subW, int subH, byte[] bbc, byte[] bbs) { 
  
    //System.out.println("c s " + bbc.length + " " + bbs.length + " -------------------------------------------");
    for (int i = 0; i < subH; i++) {
    	for (int j = 0; j < subW; j++) {
    		int currOffSmall = (i * (subW * 3)) + (j * 3); // This is always correct.....
    		// 
    		int currOffLarge = ((i * ((subW * 3) * 4)) + (j * 3 * 2));
    		int currOffLargeDup = currOffLarge + 3;
    		int currOffLargeNextRow = (i * (subW * 3 * 4)) + (subW * 3 * 2) + (j * 3 * 2);
    		int currOffLargeNextRowDup = currOffLargeNextRow + 3;
    		for (int k = 0; k < 3; k++) {
    			try {
    			  byte b = bbc[currOffSmall + k];
    	      bbs[currOffLarge + k] = b;
    	      bbs[currOffLargeDup + k] = b;
    	      bbs[currOffLargeNextRow + k] = b;
     	      bbs[currOffLargeNextRowDup + k] = b;
    			} catch (ArrayIndexOutOfBoundsException aex) {
    				System.err.println("ijk " + i + " " +  j + " " + k);
    			  System.err.println("cos " + currOffSmall);
    		    System.err.println("col " + currOffLarge);
    		    System.err.println("cold " + currOffLargeDup);
    		    System.err.println("coln " + currOffLargeNextRow);
    		    System.err.println("colnd " + currOffLargeNextRowDup);
    		    i = 100000;
    		    j = 100000;
    				break;
    			}
    	  }
    	}
    }
    return;
  }
 
  /***************************************************************************
  **
  ** How many links per pix for this current zoom?
  */
  
  private double linksPerPix(Dimension imageDim, Rectangle2D worldPiece) { 
    double zoom = Math.max(imageDim.getWidth() / worldPiece.getWidth(), imageDim.getHeight() / worldPiece.getHeight());
    double linksPerPix = 1.0 / (BioFabricPanel.GRID_SIZE * zoom);
    return (linksPerPix);
  }
  
  /***************************************************************************
  **
  ** While the lo-res slice is being displayed, we go out and render to a high-res
  ** slice that is cached:
  */
  
  private void buildHiResSlice(Dimension imageDim, int depth, Rectangle2D worldPiece) throws IOException {
    BufferedImage bi = bis_.fetchImage(imageDim.width, imageDim.height + SLICE_HEIGHT_HACK_, BufferedImage.TYPE_INT_RGB);

    double lpp = linksPerPix(imageDim, worldPiece);
    BufBuildDrawer useDrawer = (lpp < TRANSITION_LPP_) ? drawRender_ : binRender_;
   
    boolean didDraw = useDrawer.drawForBuffer(bi, worldPiece, imageDim, worldPiece, SLICE_HEIGHT_HACK_, lpp);
  
    BufferBuilderClient tellHim = null;
    synchronized (this) {
    	WorldPieceOffering wpo = allWorldsToImageName_.get(worldPiece);
      if (didDraw) {
        UiUtil.fixMePrintout("saw an NPE here. wpo must have been null!");
        if ((wpo.cacheHandle == null) || wpo.cacheHandle.equals("")) {
          wpo.cacheHandle = cache_.cacheAnImage(bi, bis_);
        } else {
          cache_.replaceAnImage(wpo.cacheHandle, bi, bis_);
        }
        wpo.isDrawn = true;
      } else {  // nothing drawn
        if (wpo.cacheHandle == null) {
          wpo.cacheHandle = "";
        } else if (!wpo.cacheHandle.equals("")) {
          cache_.dropAnImage(wpo.cacheHandle, bis_);
        }
      }
      if (bbc_ != null) {
        tellHim = bbc_;
      }
    }
    
    if (tellHim != null) {
      final int noteKey = depth;
      final BufferBuilderClient fth = tellHim;
      SwingUtilities.invokeLater(new Runnable() {
        public void run() {
          fth.yourOrderIsReady(noteKey);
        }
      });
    }
    return;
  }
 
  /***************************************************************************
  **
  ** Tell us if the image is all white.
  */

  private boolean isBlankImage(BufferedImage bi) {
  	
  	//
  	// First version of this created vast amount of garbage. Now, we just pull the
  	// ints out of the Image and make sure they are all full-on:
  	//
    int width = bi.getWidth();
    int height = bi.getHeight();
    int len = width * height;
    int[] bbs = ((DataBufferInt)bi.getRaster().getDataBuffer()).getData();
    DirectColorModel dcm = (DirectColorModel)bi.getColorModel();
    for (int i = 0; i < len; i++) {
    	if ((dcm.getRed(bbs[i]) != 255) || (dcm.getGreen(bbs[i]) != 255) || (dcm.getBlue(bbs[i]) != 255)) {
    		return (false);
    	}
    }
    return (true);
  }
  
  /***************************************************************************
  **
  ** Build images in the background:
  */

  public class BuildImageWorker implements Runnable {

    private Dimension screenDim_;
    @SuppressWarnings("unused")
    private String errString_;
    private ArrayList<QueueRequest> requests_;

    public BuildImageWorker(Dimension screenDim, List<QueueRequest> requests) {
      screenDim_ = (Dimension)screenDim.clone();
      requests_ = new ArrayList<QueueRequest>(requests);
    }

    public void run() {
      try {
        while (true) {
          QueueRequest qr = getNextRequest();
          if (qr == null) {
            break;
          }
          if (!buildBuffer(new Dimension(qr.imageDim.width, qr.imageDim.height), qr)) {
            return;
          }
        }
      } catch (IOException ex) {
        ex.printStackTrace();
        errString_ = "IOException";
      } catch (Throwable oom) {
        String format = "Other error : {1}";
        oom.printStackTrace();
        errString_ = MessageFormat.format(format, new Object[] {oom.getMessage()}); 
      }
   //   client_.retrievedResult(theUrl_, myResult_);
      return;
    }
    
    
    private QueueRequest getNextRequest() {
      synchronized (BufferBuilder.this) {
        while (requests_.isEmpty()) {
	        try {
		        BufferBuilder.this.wait();
	        } catch (InterruptedException e) {
	        }
          if (timeToExit_) {
            return (null);
          }
	      }
        return (requests_.remove(0));     
      }
    }
       
    //
    // When a lo-res result is all we can return, push the hi-res requests to the
    // front of the queue:
    //
    
    void bumpRequest(QueueRequest qr) {
      synchronized (BufferBuilder.this) {
        requests_.remove(qr);  // May not be there!!!!
        requests_.add(0, qr);
        BufferBuilder.this.notify(); 
      }
      return;
    }
  }
    
  /***************************************************************************
  **
  ** World Piece Offering
  */  
  
  private static class WorldPieceOffering implements Cloneable {
    String cacheHandle;
    boolean isDrawn;
    Dimension imageDim;
    Rectangle2D worldRect;
    
    WorldPieceOffering(String cacheHandle, Dimension imageDim, Rectangle2D worldRect, boolean isDrawn) {
      this.cacheHandle = cacheHandle;
      this.imageDim = imageDim;
      this.worldRect = worldRect;
      this.isDrawn = isDrawn;
    }
    
    @Override
    public WorldPieceOffering clone() {
      try {
        return ((WorldPieceOffering)super.clone()); // Don't copy rects; they are not changed
      } catch (CloneNotSupportedException cnse) {
        throw new IllegalStateException();
      }
    }   
  }
  
  /***************************************************************************
  **
  ** Request to draw an object
  */  
  
  private static class QueueRequest {
    int depth;
    Dimension imageDim;
    Rectangle2D worldPiece;
    
    QueueRequest(int depth, Dimension imageDim, Rectangle2D worldPiece) {
      this.depth = depth;
      this.worldPiece = worldPiece;
      this.imageDim = imageDim;
    }
    
    public int hashCode() {
      return (depth + worldPiece.hashCode() + imageDim.hashCode());
    }
      
    public boolean equals(Object other) {    
      if (other == null) {
        return (false);
      }
      if (other == this) {
        return (true);
      }
      if (!(other instanceof QueueRequest)) {
        return (false);
      }
      QueueRequest otherReq = (QueueRequest)other;

      if (this.depth != otherReq.depth) {
        return (false);
      }
      if (!this.imageDim.equals(otherReq.imageDim)) {
      	return (false);
      } 
      return (this.worldPiece.equals(otherReq.worldPiece));
    }  
    
  }
 
  /***************************************************************************
  **
  ** Interface for guys who wnat to listen for our results
  */  
  
  public interface BufferBuilderClient { 
    public void yourOrderIsReady(int sizeNum);
  } 
}
```
