## Supplementary material for "BioFabric Visualization of Network Alignments": BioFabric Source Code: GlyphPath.java

```
/*
**    Copyright (C) 2003-2017 Institute for Systems Biology 
**                            Seattle, Washington, USA. 
**
**    This library is free software; you can redistribute it and/or
**    modify it under the terms of the GNU Lesser General Public
**    License as published by the Free Software Foundation; either
**    version 2.1 of the License, or (at your option) any later version.
**
**    This library is distributed in the hope that it will be useful,
**    but WITHOUT ANY WARRANTY; without even the implied warranty of
**    MERCHANTABILITY or FITNESS FOR A PARTICULAR PURPOSE.  See the GNU
**    Lesser General Public License for more details.
**
**    You should have received a copy of the GNU Lesser General Public
**    License along with this library; if not, write to the Free Software
**    Foundation, Inc., 59 Temple Place, Suite 330, Boston, MA  02111-1307  USA
*/

package org.systemsbiology.biofabric.ui.render;

import java.awt.Color;
import java.awt.Graphics2D;
import java.awt.Rectangle;
import java.awt.geom.GeneralPath;
import java.awt.geom.Path2D;
import java.awt.geom.Rectangle2D;

/****************************************************************************
**
** These are paths used for link glyphs
*/

public class GlyphPath {
 
	////////////////////////////////////////////////////////////////////////////
  //
  // PRIVATE INSTANCE MEMBERS
  //
  ////////////////////////////////////////////////////////////////////////////
  
  private Color col_;
  private Path2D gp_;
  private Rectangle2D rect_;
  private Rectangle2D rect2_;
  private Rectangle2D gpBounds_;
  private boolean isDirected_;
  
  ////////////////////////////////////////////////////////////////////////////
  //
  // PACKAGE CONSTRUCTORS
  //
  ////////////////////////////////////////////////////////////////////////////

  /***************************************************************************
  **
  ** Used for reuse of the object
  */ 

  GlyphPath() {
  	col_ = null;
    rect_ = new Rectangle2D.Double();
    rect2_ = new Rectangle2D.Double();
    gp_ = new Path2D.Double();
    gpBounds_ = new Rectangle2D.Double();
    isDirected_ = false;
  }   
  
  /***************************************************************************
  **
  ** Used either directed or undirected links: two glyphs, one on each end. FIXME: Autolink drawn twice?
  */ 

  GlyphPath(Color col, int x, int yStrt, int yEnd, double halfWidth, boolean isDirected) {	
    col_ = col;
    isDirected_ = isDirected;
    if (isDirected) {
    	buildDirected(x, yStrt, yEnd, halfWidth);
    } else {
    	buildUndirected(x, yStrt, yEnd, halfWidth);
    }
  }   
 
  /***************************************************************************
  **
  ** Used for undirected links
  */ 

  private void buildUndirected(int x, int yStrt, int yEnd, double halfWidth) {
    rect_ = new Rectangle2D.Double();
    rect2_ = new Rectangle2D.Double(); 
    initUndir(x, yStrt, yEnd, halfWidth);
    return;
  }   
 
  /***************************************************************************
  **
  ** Used for drawing link ends and arrows for directed links.
  */ 

  private void buildDirected(int x, int yStrt, int yEnd, double halfWidth) {
  	rect_ = new Rectangle2D.Double();
    gp_ = new GeneralPath.Double();
    gpBounds_ = new Rectangle2D.Double();
    initDir(x, yStrt, yEnd, halfWidth);
    return;
  }   
  
  /***************************************************************************
  **
  ** Reuse the same object
  */ 

  GlyphPath reuse(Color col, int x, int yStrt, int yEnd, double halfWidth, boolean isDirected) {
    col_ = col;
    isDirected_ = isDirected;
    if (isDirected) {
    	initDir(x, yStrt, yEnd, halfWidth);
    } else {
    	initUndir(x, yStrt, yEnd, halfWidth);
    }
    return (this);
  }   
  	
  /***************************************************************************
  **
  ** Used for undirected links: two glyphs, one on each end. FIXME: Autolink drawn twice?
  */ 

  void initUndir(int x, int yStrt, int yEnd, double halfWidth) {
    rect_.setRect(x - halfWidth, yStrt - halfWidth, 2.0 * halfWidth, 2.0 * halfWidth);
    rect2_.setRect(x - halfWidth, yEnd - halfWidth, 2.0 * halfWidth, 2.0 * halfWidth);
    return;
  }   
 
  /***************************************************************************
  **
  ** Used for drawing link ends and arrows for directed links. FIXME: Autolink drawn twice?
  */ 

  void initDir(int x, int yStrt, int yEnd, double halfWidth) {
    rect_.setRect(x - halfWidth, yStrt - halfWidth, 2.0 * halfWidth, 2.0 * halfWidth);
    double yoff;
    double yMin;
    double yMax;
    if (yStrt < yEnd) {
    	yoff = -halfWidth;
    	yMin = yEnd - (2.0 * halfWidth);
      yMax = yEnd + halfWidth;
    } else {	
    	yoff = halfWidth;
      yMin = yEnd - halfWidth;
    	yMax = yEnd + (2.0 * halfWidth);
    }   
    gp_.reset();
    gp_.moveTo(x - halfWidth, yEnd + (2.0 * yoff));
    gp_.lineTo(x, (yEnd + yoff));
    gp_.lineTo((x - halfWidth), (yEnd + yoff));
    gp_.lineTo((x - halfWidth), (yEnd - yoff));
    gp_.lineTo((x + halfWidth), (yEnd - yoff));
    gp_.lineTo((x + halfWidth), (yEnd + yoff));
    gp_.lineTo(x, (yEnd + yoff));
    gp_.lineTo((x + halfWidth), (yEnd + (2.0 * yoff)));
    gp_.closePath();
    // Previously used gp_.getBounds2D(), but this creates a new Rectangle with each reuse.
    gpBounds_.setRect(x - halfWidth, yMin, 2.0 * halfWidth, yMax - yMin);
    return; 
  }   
  
  /***************************************************************************
  **
  ** Draw it:
  */ 

  int paint(Graphics2D g2, Rectangle bounds) {
    int didPaint = 0;
    g2.setPaint(col_);
 
    //
    // Second section: Link glyph drawing:
    //  
    if (rect_ != null) {
      // 2A: First glyph
      if ((bounds == null) ||
          ((rect_.getMaxX() > bounds.getMinX()) && 
           (rect_.getMinX() < bounds.getMaxX()) && 
           (rect_.getMaxY() > bounds.getMinY()) && 
           (rect_.getMinY() < bounds.getMaxY()))) {          
        g2.fill(rect_);     
        g2.setPaint(Color.BLACK);
        g2.draw(rect_);
        didPaint++;    
      }
      // 2B: Second glyph
      if (!isDirected_) {
        if ((bounds == null) ||
            ((rect2_.getMaxX() > bounds.getMinX()) && 
            (rect2_.getMinX() < bounds.getMaxX()) && 
            (rect2_.getMaxY() > bounds.getMinY()) && 
            (rect2_.getMinY() < bounds.getMaxY()))) {         
          g2.setPaint(col_);
          g2.fill(rect2_);     
          g2.setPaint(Color.BLACK);
          g2.draw(rect2_);
          didPaint++;
        }
      // 2C: Arrowhead glyph (uses path gp instead of rect2). Note we have saved the
      // bounding rect in rect:
      } else {
        if ((bounds == null) ||
            ((gpBounds_.getMaxX() > bounds.getMinX()) && 
             (gpBounds_.getMinX() < bounds.getMaxX()) && 
             (gpBounds_.getMaxY() > bounds.getMinY()) && 
             (gpBounds_.getMinY() < bounds.getMaxY()))) {         
          g2.setPaint(col_);
          g2.fill(gp_);     
          g2.setPaint(Color.BLACK);
          g2.draw(gp_);
          didPaint++;
        }
      }
    }
    return (didPaint);
  }
}
 
```
