## Supplementary material for "BioFabric Visualization of Network Alignments": BioFabric Source Code: ImgAndBufPool.java

```
/*
**    Copyright (C) 2003-2017 Institute for Systems Biology 
**                            Seattle, Washington, USA. 
**
**    This library is free software; you can redistribute it and/or
**    modify it under the terms of the GNU Lesser General Public
**    License as published by the Free Software Foundation; either
**    version 2.1 of the License, or (at your option) any later version.
**
**    This library is distributed in the hope that it will be useful,
**    but WITHOUT ANY WARRANTY; without even the implied warranty of
**    MERCHANTABILITY or FITNESS FOR A PARTICULAR PURPOSE.  See the GNU
**    Lesser General Public License for more details.
**
**    You should have received a copy of the GNU Lesser General Public
**    License along with this library; if not, write to the Free Software
**    Foundation, Inc., 59 Temple Place, Suite 330, Boston, MA  02111-1307  USA
*/

package org.systemsbiology.biofabric.ui.render;

import java.awt.Color;
import java.awt.Graphics2D;
import java.awt.image.BufferedImage;
import java.util.ArrayList;
import java.util.Arrays;
import java.util.HashMap;
import java.util.Iterator;
import java.util.List;


/****************************************************************************
**
** This is a cache for reusing buffered images
*/

public class ImgAndBufPool {
    
  ////////////////////////////////////////////////////////////////////////////
  //
  // PRIVATE INSTANCE MEMBERS
  //
  ////////////////////////////////////////////////////////////////////////////
  
  private HashMap<StackKey, List<BufferedImage>> biStack_;
  private HashMap<Integer, List<int[]>> bufStack_;
  private HashMap<Integer, List<byte[]>> byteStack_;
  private int binSize_;
  
  ////////////////////////////////////////////////////////////////////////////
  //
  // PUBLIC CONSTRUCTORS
  //
  ////////////////////////////////////////////////////////////////////////////

  /***************************************************************************
  **
  ** Constructor
  */

  public ImgAndBufPool(int binSize) {
    biStack_ = new HashMap<StackKey, List<BufferedImage>>();
    bufStack_ = new HashMap<Integer, List<int[]>>();
    byteStack_ = new HashMap<Integer, List<byte[]>>();
    binSize_ = binSize;
  }

  ////////////////////////////////////////////////////////////////////////////
  //
  // PUBLIC METHODS
  //
  ////////////////////////////////////////////////////////////////////////////
  
  /***************************************************************************
  ** 
  ** Get a buffered image
  */
  
  public synchronized BufferedImage fetchImage(int imgWidth, int imgHeight, int type) {
  	
  	List<BufferedImage> stack = null;
  	Iterator<StackKey> skit = biStack_.keySet().iterator();
  	while (skit.hasNext()) {
  		StackKey sk = skit.next();
  		if (sk.matches(imgWidth, imgHeight, type)) {
  			stack = biStack_.get(sk);
  			break;
  		}
  	}
  	
  	if (stack == null) {
  		stack = new ArrayList<BufferedImage>();
  		biStack_.put(new StackKey(imgWidth, imgHeight, type), stack);		
  	}
 
  	BufferedImage bi;
   	if (stack.isEmpty()) {
      //		System.out.println("Creating new BI");
  	  bi = new BufferedImage(imgWidth, imgHeight, type);	  
      //	  System.out.println("BI " + bi);
  	  /* 
  	  WritableRaster rast = bi.getRaster();
  	  SampleModel asm = rast.getSampleModel();
  	  if (asm instanceof PixelInterleavedSampleModel) {
  	    PixelInterleavedSampleModel sm = (PixelInterleavedSampleModel)asm;
	  	  int[] bo = sm.getBandOffsets();
	  	  System.out.println("BIrsmdt " + sm.getDataType());
	  	  System.out.println("BIrsmw " + sm.getWidth());
	  	  System.out.println("BIrsmh " + sm.getHeight());
	  	  System.out.println("BIrsmps " + sm.getPixelStride());
	  	  System.out.println("BIrsmss " +  sm.getScanlineStride());  
	  	  System.out.println("BIrsmbol " +  bo[0] + " " + bo[1] + " " + bo[2]); 
	  	  System.out.println("BIrw " + rast.getWidth());
	  	  System.out.println("BIrh " + rast.getHeight());
  	  }
  	  System.out.println("BIxt " +  bi.getNumXTiles());
  	  System.out.println("BIyt " +  bi.getNumYTiles());
  	 */
    } else {
    	// WJRL 11/25/17: Gotta blank out the image! Otherwise we get stuff that was drawn in last time.
      bi = stack.remove(stack.size() - 1);
      Graphics2D graphics = bi.createGraphics();
      graphics.setPaint(new Color(255, 255, 255));
      graphics.fillRect (0, 0, bi.getWidth(), bi.getHeight());
    }
    //  	System.out.println("fetching BI : size = " + stack.size() + " " +  + bi.hashCode());
   	
    return (bi);
  }

  
  /***************************************************************************
  ** 
  ** Get an integer array
  */
  
  public synchronized int[] fetchBuf(int size) {
  	
  	List<int[]> stack = bufStack_.get(Integer.valueOf(size));
  	if (stack == null) {
  		stack = new ArrayList<int[]>();
  		bufStack_.put(Integer.valueOf(size), stack);		
  	}
 
    int[] retval;
  	if (stack.isEmpty()) {
//  		System.out.println("Creating new buf");
  	  retval = new int[size];
  	} else {
  	  retval = stack.remove(stack.size() - 1);
  	  Arrays.fill(retval, 0);
  	}
  	//System.out.println("fetch buf " + retval);
    return (retval);
  }

  /***************************************************************************
  ** 
  ** Get an integer array
  */
  
  public synchronized byte[] fetchByteBuf(int size) {
  	
  	int useSize = size;
  	int rem = size % binSize_;
  	if (rem != 0) {
  		int quo = size / binSize_;
  		useSize = (quo + 1) * binSize_;
  	}
  	Integer key = Integer.valueOf(useSize);
  	
  	List<byte[]> stack = byteStack_.get(key);
  	if (stack == null) {
  		stack = new ArrayList<byte[]>();
  		byteStack_.put(key, stack);		
  	}
 
    byte[] retval;
  	if (stack.isEmpty()) {
  //		System.out.println("Creating new byte buf " + useSize);
  	  retval = new byte[useSize];
  	} else {
  	  retval = stack.remove(stack.size() - 1);
  	  Arrays.fill(retval, (byte)0);
  	}
    return (retval);
  }
  
  /***************************************************************************
  ** 
  ** Return a buffered image
  */
  
  public synchronized void returnImage(BufferedImage bi) {
  	if (bi == null) {
  		System.err.println("Returning null BufferedImage");
  		return;
  	}
  	int imgWidth = bi.getWidth();
  	int imgHeight = bi.getHeight();
  	int type = bi.getType();
  	List<BufferedImage> stack = null;
  	Iterator<StackKey> skit = biStack_.keySet().iterator();
  	while (skit.hasNext()) {
  		StackKey sk = skit.next();
  		if (sk.matches(imgWidth, imgHeight, type)) {
  			stack = biStack_.get(sk);
  			break;
  		}
  	}
  	
  	//
  	// Images coming off of a file may never have been seen before!
  	//
  	
  	if (stack == null) {
  		stack = new ArrayList<BufferedImage>();
  		biStack_.put(new StackKey(imgWidth, imgHeight, type), stack);		
  	}
    Iterator<BufferedImage> stit = stack.iterator();
  	while (stit.hasNext()) {
  		BufferedImage sk = stit.next();
  		if (sk.hashCode() == bi.hashCode()) {
  			throw new IllegalStateException();
  		}
  	}

  	stack.add(bi);
  	//System.out.println("Returning BI : size = " + stack.size() + " " +  + bi.hashCode());
    return;
  }
 
  /***************************************************************************
  ** 
  ** Return a buffer
  */
  
  public synchronized void returnBuf(int[] buf) {
  	if (buf == null) {
  		System.err.println("Returning null buf");
  		return;
  	}
  	List<int[]> stack = bufStack_.get(Integer.valueOf(buf.length));
  	if (stack == null) {
  		stack = new ArrayList<int[]>();
  		bufStack_.put(Integer.valueOf(buf.length), stack);		
  	} else if (stack.contains(buf)) {
  		throw new IllegalStateException();
  	}
  	
  	stack.add(buf);
  //	System.out.println("Returning buf: size = " + stack.size());
  	//System.out.println("returned buf " + buf);
    return;
  }
 
  /***************************************************************************
  ** 
  ** Return a buffer
  */
  
  public synchronized void returnByteBuf(byte[] buf) {
  	if (buf == null) {
  		System.err.println("Returning null buf");
  		return;
  	}
  	List<byte[]> stack = byteStack_.get(Integer.valueOf(buf.length));
  	if (stack == null) {
  		stack = new ArrayList<byte[]>();
  		byteStack_.put(Integer.valueOf(buf.length), stack);		
  	} else if (stack.contains(buf)) {
  		throw new IllegalStateException();
  	}
  	
  	stack.add(buf);
 // 	System.out.println("Returning byte buf: size = " + stack.size());
    return;
  } 
  
  ////////////////////////////////////////////////////////////////////////////
  //
  // INNER CLASSES
  //
  ////////////////////////////////////////////////////////////////////////////
  
  /***************************************************************************
  ** 
  ** Keys for stacks of different flavored images
  */ 
 
  private static class StackKey {
  
    int width;
    int height;
    int type;
  
    StackKey(int width, int height, int type) {
    	this.width = width;
      this.height = height;
      this.type = type;
	  }
 
   	boolean matches(int width, int height, int type) {
	    return ((this.width == width) && (this.height == height) && (this.type == type));
	  }
   
	  @Override
	  public int hashCode() {
	    return (width + height + type);
	  }
	  
	  @Override
	  public boolean equals(Object other) {
	    if (other == null) {
	      return (false);
	    }
	    if (other == this) {
	      return (true);
	    }
	    if (!(other instanceof StackKey)) {
	      return (false);
	    }
	    StackKey otherMM = (StackKey)other;
	    return ((this.width == otherMM.width) && (this.height == otherMM.height) && (this.type == otherMM.type));
	  }  
  }
}
```
