## Supplementary material for "BioFabric Visualization of Network Alignments": BioFabric Source Code: LinePath.java

```
/*
**    Copyright (C) 2003-2017 Institute for Systems Biology 
**                            Seattle, Washington, USA. 
**
**    This library is free software; you can redistribute it and/or
**    modify it under the terms of the GNU Lesser General Public
**    License as published by the Free Software Foundation; either
**    version 2.1 of the License, or (at your option) any later version.
**
**    This library is distributed in the hope that it will be useful,
**    but WITHOUT ANY WARRANTY; without even the implied warranty of
**    MERCHANTABILITY or FITNESS FOR A PARTICULAR PURPOSE.  See the GNU
**    Lesser General Public License for more details.
**
**    You should have received a copy of the GNU Lesser General Public
**    License along with this library; if not, write to the Free Software
**    Foundation, Inc., 59 Temple Place, Suite 330, Boston, MA  02111-1307  USA
*/

package org.systemsbiology.biofabric.ui.render;

import java.awt.Color;
import java.awt.Graphics2D;
import java.awt.Rectangle;
import java.awt.geom.Line2D;

import org.systemsbiology.biofabric.api.util.MinMax;

/****************************************************************************
**
** These are paths used for nodes and links
*/

public class LinePath {

	////////////////////////////////////////////////////////////////////////////
  //
  // PRIVATE INSTANCE MEMBERS
  //
  ////////////////////////////////////////////////////////////////////////////
 
	private Color col;
  private Line2D path;
  private int px;
  private int py;
  private MinMax range;
  private boolean reusing;
  
  ////////////////////////////////////////////////////////////////////////////
  //
  // PACKAGE CONSTRUCTORS
  //
  ////////////////////////////////////////////////////////////////////////////

  /***************************************************************************
  **
  ** For when we create one link path and reuse it:
  */ 

  LinePath() {
  	reusing = true;
  }

  /***************************************************************************
  **
  ** One constructor for both links and nodes.
  */ 

  LinePath(Color col, Line2D path, int x, int y, MinMax range) {
  	reset(col, path, x, y, range);
    reusing = false;
  }
  
  ////////////////////////////////////////////////////////////////////////////
  //
  // PACKAGE METHODS
  //
  ////////////////////////////////////////////////////////////////////////////
 
  /***************************************************************************
  **
  ** Reset the contents
  */ 
  
  public LinePath reset(Color col, Line2D path, int x, int y, MinMax range) {
    this.col = col;
    this.path = path;
    this.px = x;
    this.py = y;
    this.range = range;
    return (this);
  }  
  
  /***************************************************************************
  **
  ** Draw it
  */ 
  
  int paint(Graphics2D g2, Rectangle bounds) {
    int didPaint = 0;
    if (px == Integer.MIN_VALUE) {  // Horizontal (node) drawing
    	didPaint = paintAsNode(g2, bounds);   
    } else if (py == Integer.MIN_VALUE) {  // Vertical (link) drawing
    	didPaint = paintAsLink(g2, bounds);
    }
    return (didPaint);
  } 
  
  /***************************************************************************
  **
  ** Draw it
  */ 
  
  private int paintAsNode(Graphics2D g2, Rectangle bounds) {
    int didPaint = 0;
    g2.setPaint(col);
  	//
    // 4A: Horizontal (node) drawing:
    //   	      
    if ((bounds == null) || ((py >= bounds.y) && (py <= (bounds.y + bounds.height)))) {
      if ((bounds != null) && ((range.max < bounds.x) || (range.min > (bounds.x + bounds.width)))) {
        // do nothing
      } else {
      	// With very long lines (e.g. 100,000 units) we see fragmented draws on Mac. Make short, then restore:
      	double x1 = path.getX1();
      	double x2 = path.getX2();
      	double y1 = path.getY1();
      	double y2 = path.getY2();
      	boolean replace = false;
      	if (x2 < x1) {
      		throw new IllegalStateException();
      	} 
      	double useLowBound = x1;
      	double useHiBound = x2;
      	if ((x2 - x1) > 100000.0) {
      	  if (x1 < bounds.getMinX()) {
      		  useLowBound = bounds.getX() - 1000.0;
      		  replace = true;
      	  }
      	  if (x2 > bounds.getMaxX()) {
      		  useHiBound = bounds.getMaxX() + 1000.0;
      		  replace = true;
      	  }	
      	  path.setLine(useLowBound, y1, useHiBound, y2);
      	}
        g2.draw(path);
        if (replace) {
          path.setLine(x1, y1, x2, y2);
        }
        didPaint++;
      }
    }
    return (didPaint);
  } 
  
  /***************************************************************************
  **
  ** Draw it
  */ 
  
  private int paintAsLink(Graphics2D g2, Rectangle bounds) {
    int didPaint = 0;
    g2.setPaint(col);
    //
    // 4B: Vertical (link) drawing:
    //    
    if (py == Integer.MIN_VALUE) { // Vert line
      if ((bounds == null) || ((px >= bounds.x) && (px <= (bounds.x + bounds.width)))) { 
        if ((bounds != null) && ((range.max < bounds.y) || (range.min > (bounds.y + bounds.height)))) {
          // do nothing
        } else {
        	// With very long lines (e.g. 100,000 units) we see fragmented draws on Mac. Make short, then restore:
        	double x1 = path.getX1();
        	double x2 = path.getX2();
        	double y1 = path.getY1();
        	double y2 = path.getY2();
        	boolean replace = false;
        	if (y2 < y1) {
        		throw new IllegalStateException();
        	}
        	double useLowBound = y1;
        	double useHiBound = y2;
        	if ((y2 - y1) > 100000.0) {
        	  if (y1 < bounds.getMinY()) {
        		  useLowBound = bounds.getY() - 1000.0;
        		  replace = reusing;
        	  }
        	  if (y2 > bounds.getMaxY()) {
        		  useHiBound = bounds.getMaxY() + 1000.0;
        		  replace = reusing;
        	  }	
        	  path.setLine(x1, useLowBound, x2, useHiBound);
        	}
          g2.draw(path);
          if (replace) {
            path.setLine(x1, y1, x2, y2);
          }
          didPaint++;        
        }
      }
    }
    return (didPaint);
  }   
} 
```
