## Supplementary material for "BioFabric Visualization of Network Alignments": BioFabric Source Code: PaintCacheSmall.java

```
/*
**    Copyright (C) 2003-2018 Institute for Systems Biology 
**                            Seattle, Washington, USA. 
**
**    This library is free software; you can redistribute it and/or
**    modify it under the terms of the GNU Lesser General Public
**    License as published by the Free Software Foundation; either
**    version 2.1 of the License, or (at your option) any later version.
**
**    This library is distributed in the hope that it will be useful,
**    but WITHOUT ANY WARRANTY; without even the implied warranty of
**    MERCHANTABILITY or FITNESS FOR A PARTICULAR PURPOSE.  See the GNU
**    Lesser General Public License for more details.
**
**    You should have received a copy of the GNU Lesser General Public
**    License along with this library; if not, write to the Free Software
**    Foundation, Inc., 59 Temple Place, Suite 330, Boston, MA  02111-1307  USA
*/

package org.systemsbiology.biofabric.ui.render;

import java.awt.BasicStroke;
import java.awt.Color;
import java.awt.Font;
import java.awt.Graphics2D;
import java.awt.Rectangle;
import java.awt.font.FontRenderContext;
import java.awt.font.LineMetrics;
import java.awt.geom.AffineTransform;
import java.awt.geom.Line2D;
import java.awt.geom.Rectangle2D;
import java.util.ArrayList;
import java.util.Arrays;
import java.util.HashMap;
import java.util.HashSet;
import java.util.List;
import java.util.Map;
import java.util.Set;

import org.systemsbiology.biofabric.api.layout.AnnotColorSource;
import org.systemsbiology.biofabric.api.model.AnnotationSet;
import org.systemsbiology.biofabric.api.model.Annot;
import org.systemsbiology.biofabric.api.model.NetNode;
import org.systemsbiology.biofabric.api.util.MinMax;
import org.systemsbiology.biofabric.api.util.NID;
import org.systemsbiology.biofabric.api.worker.AsynchExitRequestException;
import org.systemsbiology.biofabric.api.worker.BTProgressMonitor;
import org.systemsbiology.biofabric.api.worker.LoopReporter;
import org.systemsbiology.biofabric.model.BioFabricNetwork;
import org.systemsbiology.biofabric.ui.FabricColorGenerator;
import org.systemsbiology.biofabric.ui.FabricDisplayOptions;
import org.systemsbiology.biofabric.ui.FabricDisplayOptionsManager;
import org.systemsbiology.biofabric.ui.display.BioFabricPanel;
import org.systemsbiology.biofabric.util.DoubMinMax;
import org.systemsbiology.biofabric.util.QuadTree;
import org.systemsbiology.biofabric.util.UiUtil;

/****************************************************************************
**
** This is the cache of simple paint objects.
*/

public class PaintCacheSmall {
  
  ////////////////////////////////////////////////////////////////////////////
  //
  // PRIVATE CONSTANTS
  //
  //////////////////////////////////////////////////////////////////////////// 
  
  private final static double BB_HALF_WIDTH_ = 5.0F;
  
  private final static double PERCENT_PAD_ = 0.10;
  private final static int MINIMUM_PAD_ = 10;
  
  private final static double DRAIN_ZONE_ROW_OFFSET_ = 0.5F; // Lifts drain zone text above the node line & boxes
  private final static double NODE_LABEL_X_SHIM_ = 5.0F;
  private final static double LABEL_FONT_HEIGHT_SCALE_ = 2.0 / 3.0;

  ////////////////////////////////////////////////////////////////////////////
  //
  // PUBLIC CONSTANTS
  //
  //////////////////////////////////////////////////////////////////////////// 
  
  public final static int STROKE_SIZE = 3;

  ////////////////////////////////////////////////////////////////////////////
  //
  // PRIVATE INSTANCE MEMBERS
  //
  ////////////////////////////////////////////////////////////////////////////
   
  private HashMap<TextPath.FontSizes, Font> fonts_; 
  private ArrayList<TextPath.FontSizes> bigToLittle_; 
 
  private List<BioFabricNetwork.NodeInfo> nodeRefs_;
  private List<BioFabricNetwork.LinkInfo> linkRefs_;
  private int[] linkIndex_;
  private int indexOffset_;
  private int[] nodeIndex_;
  private int nodeIndexOffset_;
  private boolean nodesForShadow_;
  private QuadTree names_;
  private HashMap<String, BoxPath> nameKeyToPaintZero_;
  private HashMap<String, BoxPath> nameKeyToPaintOneQuarter_;
  private HashMap<String, TextPath> nameKeyToPaintOneHalf_;
  private HashMap<String, BoxPath> nameKeyToPaintFirst_;
  private HashMap<String, TextPath> nameKeyToPaintSecond_;
  private HashMap<String, TextPath> nameKeyToPaintThird_;
  private int nameKeyCount_;
  private HashMap<String, NID> nameKeyToNodeID_;
  
  private FabricColorGenerator colGen_;
  private Color superLightPink_;
  private Color superLightBlue_;
  private AnnotColorSource.AnnotColor[] annotColors_;
  private Color[] annotGrays_;
  
  ////////////////////////////////////////////////////////////////////////////
  //
  // PUBLIC CONSTRUCTORS
  //
  ////////////////////////////////////////////////////////////////////////////

  /***************************************************************************
  **
  ** Constructor
  */

  public PaintCacheSmall(FabricColorGenerator colGen) {
  	fonts_ = new HashMap<TextPath.FontSizes, Font>();
  	bigToLittle_ = new ArrayList<TextPath.FontSizes>();
  	
  	bigToLittle_.add(TextPath.FontSizes.GINORMOUS);
    fonts_.put(TextPath.FontSizes.GINORMOUS, new Font("SansSerif", Font.PLAIN, 400));
  	bigToLittle_.add(TextPath.FontSizes.HUGE);
  	fonts_.put(TextPath.FontSizes.HUGE, new Font("SansSerif", Font.PLAIN, 200));
  	bigToLittle_.add(TextPath.FontSizes.MED);
  	fonts_.put(TextPath.FontSizes.MED, new Font("SansSerif", Font.PLAIN, 100));
  	bigToLittle_.add(TextPath.FontSizes.MED_SMALL);
  	fonts_.put(TextPath.FontSizes.MED_SMALL, new Font("SansSerif", Font.PLAIN, 70));
  	bigToLittle_.add(TextPath.FontSizes.SMALL);
  	fonts_.put(TextPath.FontSizes.SMALL, new Font("SansSerif", Font.PLAIN, 30));
  	bigToLittle_.add(TextPath.FontSizes.TINY);
    fonts_.put(TextPath.FontSizes.TINY, new Font("SansSerif", Font.PLAIN, 10));
  
    nameKeyToPaintZero_ = new HashMap<String, BoxPath>();
    nameKeyToPaintOneQuarter_ = new HashMap<String, BoxPath>();
    nameKeyToPaintOneHalf_ = new HashMap<String, TextPath>();
    nameKeyToPaintFirst_ = new HashMap<String, BoxPath>();
    nameKeyToPaintSecond_ = new HashMap<String, TextPath>();
    nameKeyToPaintThird_ = new HashMap<String, TextPath>();
    nameKeyToNodeID_ = new HashMap<String, NID>();

    colGen_ = colGen;
    superLightPink_ = new Color(255, 244, 244);
    superLightBlue_ = new Color(244, 244, 255);
     
    annotColors_ = AnnotColorSource.getColorCycle();
    annotGrays_ = AnnotColorSource.getGrayCycle();

  }
  
  ////////////////////////////////////////////////////////////////////////////
  //
  // PUBLIC METHODS
  //
  ////////////////////////////////////////////////////////////////////////////

  /***************************************************************************
  **
  **  Dump used memory
  */
  
  public void clear() {
    nameKeyToPaintZero_.clear();
    nameKeyToPaintOneQuarter_.clear();
    nameKeyToPaintOneHalf_.clear();
  	nameKeyToPaintFirst_.clear();
    nameKeyToPaintSecond_.clear();
    nameKeyToPaintThird_.clear();
    nameKeyToNodeID_.clear();
    nodeRefs_ = null;
    linkRefs_ = null;
    nodeIndex_ = null;
    nodeIndexOffset_ = 0;
    nodesForShadow_ = false;
    linkIndex_ = null;
    indexOffset_ = 0;
    if (names_ != null) {
    	names_.clear();
    }
  	return;
  }
 

  /***************************************************************************
  **
  **  paint it
  */
  
  public boolean paintIt(Graphics2D g2, Rectangle clip, Reduction reduce) {
    boolean retval = false;
    HashSet<String> pKeys = new HashSet<String>();
    if (names_ != null) {
      names_.getPayloadKeys(clip, pKeys);
    }
    
    //
    // Zero pass is node annotation rectangles, which are not drawn for selections:
    //
    
    if (reduce == null) {
      for (String pkey : pKeys) {
        BoxPath pp = nameKeyToPaintZero_.get(pkey);
        if (pp != null) {
          int result = pp.paint(g2, clip);
          retval = retval || (result > 0);
        }
      }
    }
    
    //
    // Next pass is link annotation rectangles, which are not drawn for selections:
    //
    
    if (reduce == null) {
      for (String pkey : pKeys) {
        BoxPath pp = nameKeyToPaintOneQuarter_.get(pkey);
        if (pp != null) {
          int result = pp.paint(g2, clip);
          retval = retval || (result > 0);
        }
      }
    }
 
    //
    // Next pass is annotation strings, which are not drawn for selections:
    //
    
    if (reduce == null) {
      for (String pkey : pKeys) {
        TextPath pp = nameKeyToPaintOneHalf_.get(pkey);
        if (pp != null) {
          int result = pp.paint(g2, clip, fonts_);
          retval = retval || (result > 0);
        }
      }
    }
       
    //
    // First pass is background rectangles, which are not drawn for selections:
    //
    
    if (reduce == null) {
	    for (String pkey : pKeys) {
	      BoxPath pp = nameKeyToPaintFirst_.get(pkey);
	    	if (pp != null) {
	        int result = pp.paint(g2, clip);
	        retval = retval || (result > 0);
	    	}
	    }
    }
    
    //
    // Draw strings for nodes, if in bounds and for selections, if selected:
    //
    
    for (String pkey : pKeys) {
    	NID forKey = nameKeyToNodeID_.get(pkey); 			
      if ((reduce == null) || reduce.paintNames.contains(forKey)) {
        TextPath pp = nameKeyToPaintSecond_.get(pkey);
    	  if (pp != null) {
          int result = pp.paint(g2, clip, fonts_);
          retval = retval || (result > 0);
    	  }
      }
    }
	    

    if (nodeRefs_ != null) {  	
    	double minY = clip.getMinY() / BioFabricPanel.GRID_SIZE;
    	double maxY = clip.getMaxY() / BioFabricPanel.GRID_SIZE;  
    	//
    	// Want a padding on either side based on percentage, though for tiny
    	// clips (e.g. for magnifier), we want that to not be smaller than a minimum:
    	//
    	int extraRows = (int)Math.round((maxY - minY) * PERCENT_PAD_ * 0.5);
    	if (extraRows < MINIMUM_PAD_) {
    		extraRows = MINIMUM_PAD_;
    	}
	    int startRow = (int)Math.floor(minY) - extraRows;
	    int endRow = (int)Math.floor(maxY) + extraRows;
	       
	    startRow = (startRow < nodeIndexOffset_) ? nodeIndexOffset_ : startRow;
	    endRow = (endRow >= (nodeIndex_.length + nodeIndexOffset_)) ? nodeIndex_.length - 1 + nodeIndexOffset_: endRow;
	    
	    Line2D line = new Line2D.Double();
      LinePath lp = new LinePath();
      MinMax mm = new MinMax();

	    for (int i = startRow; i <= endRow; i++) {
	    	// If we are not drawing contiguous nodes (subviews with gaps) we will hit non-node rows to skip:
	      if (nodeIndex_[i - nodeIndexOffset_] == -1) {
	      	continue;
	      }
	    	if ((reduce == null) || reduce.paintRows.contains(Integer.valueOf(i))) {
	    		BioFabricNetwork.NodeInfo ni = nodeRefs_.get(nodeIndex_[i - nodeIndexOffset_]);
	    		MinMax nmm = ni.getColRange(nodesForShadow_);
          int sCol = nmm.min;
          int eCol = nmm.max;
          Color paintCol = getColorForNode(ni, colGen_);
     
          int xStrt = sCol * BioFabricPanel.GRID_SIZE;
          int xEnd = eCol * BioFabricPanel.GRID_SIZE;
          int y = i * BioFabricPanel.GRID_SIZE;
  
          line.setLine(xStrt, y, xEnd, y);
          int result = lp.reset(paintCol, line, Integer.MIN_VALUE, y, mm.reset(xStrt, xEnd)).paint(g2, clip);
		      retval = retval || (result > 0);
	    	}
	    }
    }

    if (linkRefs_ != null) {  	
    	double minX = clip.getMinX() / BioFabricPanel.GRID_SIZE;
    	double maxX = clip.getMaxX() / BioFabricPanel.GRID_SIZE;   	
    	//
    	// Want a padding on either side based on percentage, though for tiny
    	// clips (e.g. for magnifier), we want that to not be smaller than a minimum:
    	//
    	int extraCols = (int)Math.round((maxX - minX) * PERCENT_PAD_ * 0.5);
    	if (extraCols < MINIMUM_PAD_) {
    		extraCols = MINIMUM_PAD_;
    	}
    	
	    int startCol = (int)Math.floor(minX) - extraCols;
	    int endCol = (int)Math.floor(maxX) + extraCols;
	    
	    startCol = (startCol < indexOffset_) ? indexOffset_ : startCol;
	    endCol = (endCol >= (linkIndex_.length + indexOffset_)) ? linkIndex_.length - 1 + indexOffset_: endCol;

	    Line2D line = new Line2D.Double();
      LinePath lp = new LinePath();
      MinMax mm = new MinMax();
      GlyphPath gp = new GlyphPath();

      int count = 0;
	    for (int i = startCol; i <= endCol; i++) {
	    	// If we are not drawing contiguous links (subviews with gaps) we will hit non-link columns to skip:
	      if (linkIndex_[i - indexOffset_] == -1) {
	      	continue;
	      }
	    	if ((reduce == null) || reduce.paintCols.contains(Integer.valueOf(i))) {

	    		BioFabricNetwork.LinkInfo li = linkRefs_.get(linkIndex_[i - indexOffset_]);
          int sRow = li.topRow();
          int eRow = li.bottomRow();
          Color paintCol = getColorForLink(li, colGen_);
     
          int yStrt = sRow * BioFabricPanel.GRID_SIZE;
          int yEnd = eRow * BioFabricPanel.GRID_SIZE;
          int x = i * BioFabricPanel.GRID_SIZE;
  
          line.setLine(x, yStrt, x, yEnd);
          int result = lp.reset(paintCol, line, x, Integer.MIN_VALUE, mm.reset(yStrt, yEnd)).paint(g2, clip);
		      retval = retval || (result > 0);
		      count += (result > 0) ? 1 : 0;
         
          if (!li.isDirected()) {
          	result = gp.reuse(paintCol, x, yStrt, yEnd, BB_HALF_WIDTH_, false).paint(g2, clip);
          	retval = retval || (result > 0);
          } else {
            int ySrc = li.getStartRow() * BioFabricPanel.GRID_SIZE;
            int yTrg = li.getEndRow() * BioFabricPanel.GRID_SIZE;
            result = gp.reuse(paintCol, x, ySrc, yTrg, BB_HALF_WIDTH_, true).paint(g2, clip);
            retval = retval || (result > 0);
          }
	    	}
	    }
    }

    //
    // Draw strings that appear on top of everything:
    //
    
    for (String pkey : pKeys) {
    	NID forKey = nameKeyToNodeID_.get(pkey); 			
      if ((reduce == null) || reduce.paintNames.contains(forKey)) {
        TextPath pp = nameKeyToPaintThird_.get(pkey);
    	  if (pp != null) {
          int result = pp.paint(g2, clip, fonts_);
          retval = retval || (result > 0);
    	  }
      }
    }

    return (retval);
  }
  
  /***************************************************************************
  **
  ** Drawing core
  */
  
  public void drawFloater(Graphics2D g2, FloaterSet floaters) {
    if ((floaters.floater == null) && (floaters.tourRect == null) && (floaters.currSelRect == null)) {
      return;
    }       
    BasicStroke selectedStroke = new BasicStroke(6, BasicStroke.CAP_SQUARE, BasicStroke.JOIN_MITER);    
    g2.setStroke(selectedStroke);    
    g2.setPaint(Color.BLACK);
    if (floaters.floater != null) {
      g2.drawRect(floaters.floater.x, floaters.floater.y, floaters.floater.width, floaters.floater.height); 
    }  
    if (floaters.tourRect != null) {
      g2.setPaint(new Color(0, 0, 255, 125));
      g2.drawArc(floaters.tourRect.x, floaters.tourRect.y, floaters.tourRect.width, floaters.tourRect.height, 0, 360); 
    }
    if (floaters.currSelRect != null) {
      g2.setPaint(new Color(Color.orange.getRed(), Color.orange.getGreen(), Color.orange.getBlue(), 125));
      g2.drawArc(floaters.currSelRect.x, floaters.currSelRect.y, floaters.currSelRect.width, floaters.currSelRect.height, 0, 360); 
    }
    return;
  }
 
  /***************************************************************************
  **
  ** Build objcache
  */
  
  public Rectangle2D buildObjCache(List<BioFabricNetwork.NodeInfo> targets, List<BioFabricNetwork.LinkInfo> links, 
				                           boolean shadeNodes, boolean showShadows, BioFabricNetwork.Extents ext, 
				                           Map<NetNode, Rectangle2D> nameMap, 
				                           Map<NetNode, List<Rectangle2D>> drainMap, Rectangle2D netBounds,
				                           AnnotationSet nodeAnnot, AnnotationSet linkAnnot, 
				                           BTProgressMonitor monitor) throws AsynchExitRequestException {
  	
  	
    nameKeyToPaintZero_.clear();
    nameKeyToPaintOneQuarter_.clear();
    nameKeyToPaintOneHalf_.clear();
    nameKeyToPaintFirst_.clear();
    nameKeyToPaintSecond_.clear();
    nameKeyToPaintThird_.clear();

    nameKeyCount_ = 0;
    
    //
    // Build the quad tree after we know the extents it has to cover:
    //
    
    ArrayList<QuadTree.Payload> qtpc = new ArrayList<QuadTree.Payload>();
    
    //
    // Need to find the maximum column in use for the links, depending on shadow display or not:
    //
     
   // nodes_ = new PaintedPath[targets.size()];

    FontRenderContext frc = new FontRenderContext(new AffineTransform(), true, true);
   
    int numLinks = links.size();
    Map<Integer, MinMax> linkExtents = ext.allLinkExtents.get(Boolean.valueOf(showShadows));
    MinMax linkCols = ext.allLinkFullRange.get(Boolean.valueOf(showShadows));
     
    linkIndex_ = new int[(numLinks == 0) ? 0 : linkCols.max + 1 - linkCols.min];
    indexOffset_ = linkCols.min;  // In subviews, links do NOT start at column 0!
    // And in subviews with non-contiguous links, we need to skip non-link columns, so init with -1:
    Arrays.fill(linkIndex_, -1);
    linkRefs_ = links;
    
    int numNodes = targets.size();
    
    LoopReporter lr = new LoopReporter(targets.size(), 20, monitor, 0.0, 1.0, "progress.buildNodeGraphics");
    HashMap<Integer, MinMax> nodeExtents = ext.allNodeExtents.get(Boolean.valueOf(showShadows));
    Integer singletonMinRow = ext.singletonNodeStart.get(Boolean.valueOf(showShadows));
    MinMax nodeRows = ext.allNodeFullRange.get(Boolean.valueOf(showShadows));
    for (int i = 0; i < numNodes; i++) {
      BioFabricNetwork.NodeInfo node = targets.get(i);
      int num = node.nodeRow;
      lr.report();
      buildNodeTextAndRect(node, frc, colGen_, linkExtents, shadeNodes, showShadows, nameMap, drainMap, qtpc);
    }
    
    nodeIndex_ = new int[(numNodes == 0) ? 0 : nodeRows.max + 1 - nodeRows.min];
    nodeIndexOffset_ = nodeRows.min;  // In subviews, links do NOT start at column 0!
    // And in subviews with non-contiguous links, we need to skip non-link columns, so init with -1:
    Arrays.fill(nodeIndex_, -1);
    nodeRefs_ = targets;
    nodesForShadow_ = showShadows;
      
    //
    // We do not build e.g. 10^6 link paths, but stock the drawing primitives while painting:
    //
    
    LoopReporter lr2 = new LoopReporter(numLinks, 20, monitor, 0.0, 1.0, "progress.buildLinkGraphics");
    for (int i = 0; i < numLinks; i++) {
      BioFabricNetwork.LinkInfo link = links.get(i);
      lr2.report();
      linkIndex_[link.getUseColumn(showShadows) - indexOffset_] = i;
    }
    
    //
    // Fill the node index
    //
    
    LoopReporter lr3 = new LoopReporter(numLinks, 20, monitor, 0.0, 1.0, "progress.buildNodeGraphicsToo");
    for (int i = 0; i < numNodes; i++) {
      BioFabricNetwork.NodeInfo node = targets.get(i);
      lr3.report();
      nodeIndex_[node.nodeRow - nodeIndexOffset_] = i;
    }
      
    int annotCount = 0;
    if (nodeAnnot != null) {
    	LoopReporter lr4 = new LoopReporter(nodeAnnot.size(), 20, monitor, 0.0, 1.0, "progress.buildNodeAnnots");
      for (Annot an : nodeAnnot) {
        AnnotColorSource.AnnotColor acol = an.getColor();
        AnnotColorSource.AnnotColor col = (acol == null) ? annotColors_[annotCount++ % annotColors_.length] : acol;
        lr4.report();
        buildAnAnnotationRect(an.getRange(), an.getName(), col.getColor(), true, nodeExtents, frc, linkCols, qtpc, null);
      }
    }
    
    annotCount = 0;
    if (linkAnnot != null) {
    	LoopReporter lr5 = new LoopReporter(linkAnnot.size(), 20, monitor, 0.0, 1.0, "progress.buildLinkAnnots");
      for (Annot an : linkAnnot) {
        Color col;
        if ((nodeAnnot != null) && (nodeAnnot.size() > 0)) {
          col = annotGrays_[annotCount++ % annotGrays_.length];
        } else {
          AnnotColorSource.AnnotColor acol = an.getColor();
          col = (acol == null) ? annotColors_[annotCount++ % annotColors_.length].getColor() : acol.getColor();
        }  
        lr5.report();
        buildAnAnnotationRect(an.getRange(), an.getName(), col, false, linkExtents, frc, nodeRows, qtpc, singletonMinRow);
      }
    }
    
    //
    // Now that we have built everything that is going into the quadTree, we can build it:
    //
    
    DoubMinMax dmmw = new DoubMinMax(netBounds.getMinX(), netBounds.getMaxX());
    DoubMinMax dmmh = new DoubMinMax(netBounds.getMinY(), netBounds.getMaxY());
  
    for (QuadTree.Payload qtp : qtpc) {
    	Rectangle2D qtpr = qtp.getRect();
    	dmmw.update(qtpr.getMinX());
    	dmmw.update(qtpr.getMaxX());
    	dmmh.update(qtpr.getMinY());
    	dmmh.update(qtpr.getMaxY());
    }
    
    Rectangle2D worldRect = new Rectangle2D.Double(dmmw.min, dmmh.min, dmmw.max - dmmw.min, dmmh.max - dmmh.min);
    names_ = new QuadTree(worldRect, 5);
    
    for (QuadTree.Payload qtp : qtpc) {   	
    	names_.insertPayload(qtp);
    }
    qtpc.clear();
    return (worldRect);
  }
 
  /***************************************************************************
  ** 
  ** Get a link color
  */

  public Color getColorForLink(BioFabricNetwork.LinkInfo link, FabricColorGenerator colGen) {
    return (colGen.getModifiedColor(link.getColorKey(), FabricColorGenerator.DARKER)); 
  }
  
  /***************************************************************************
  ** 
  ** Get a node color
  */

  public Color getColorForNode(BioFabricNetwork.NodeInfo node, FabricColorGenerator colGen) {
    return (colGen.getModifiedColor(node.colorKey, FabricColorGenerator.BRIGHTER)); 
  }

  /***************************************************************************
  **
  ** Build the text objects associated with a node
  */
  
  private void buildNodeTextAndRect(BioFabricNetwork.NodeInfo target,
                                    FontRenderContext frc,
                                    FabricColorGenerator colGen, Map<Integer, MinMax> linkExtents,
                                    boolean shadeNodes, boolean showShadows, Map<NetNode, Rectangle2D> nameMap,
                                    Map<NetNode, List<Rectangle2D>> drainMap, ArrayList<QuadTree.Payload> payloadCache) {
 
    //
    // Left end node label sizing and Y:
    //
    
    MinMax colmm = target.getColRange(showShadows);
    Rectangle2D labelBounds = fonts_.get(TextPath.FontSizes.TINY).getStringBounds(target.getNodeName(), frc);
    // Easiest font height hack is to scale it by ~.67: 
    double scaleHeight = labelBounds.getHeight() * LABEL_FONT_HEIGHT_SCALE_;
    double namey = (target.nodeRow * BioFabricPanel.GRID_SIZE) + (scaleHeight / 2.0);
    double namex = (colmm.min * BioFabricPanel.GRID_SIZE) - labelBounds.getWidth() - BB_HALF_WIDTH_ - NODE_LABEL_X_SHIM_;
    labelBounds.setRect(namex, namey - scaleHeight, labelBounds.getWidth(), scaleHeight);
    
    //
    // Get the name bounds into the map and the extents into the QuadTree:
    //
    
    nameMap.put(target.getNodeIDWithName(), (Rectangle2D) labelBounds.clone());
    TextPath npp = new TextPath(Color.BLACK, target.getNodeName(), namex, namey, labelBounds, false, TextPath.FontSizes.TINY);
    String nkk = Integer.toString(nameKeyCount_++);
    QuadTree.Payload pay = new QuadTree.Payload(labelBounds, nkk);
    payloadCache.add(pay);
    nameKeyToPaintSecond_.put(nkk, npp);
    nameKeyToNodeID_.put(nkk, target.getNodeID());
    
    //
    // Now process drain zones:
    //

    //
    // All the drain zone text rectangles associated with this node:
    //
    
    List<Rectangle2D> rectList = new ArrayList<Rectangle2D>();
    
    // Drain zone Y: Lifted slightly above node line and link boxes:
    
    double tnamey = (target.nodeRow - DRAIN_ZONE_ROW_OFFSET_) * BioFabricPanel.GRID_SIZE;
    
    //
    // Drain zone info for each zone:
    //
    
    List<BioFabricNetwork.DrainZone> zones = target.getDrainZones(showShadows);
 
    //
    // Process each zone:
    //
    
    for (int i = 0; i < zones.size(); i++) {
      DrainZoneInfo curr = new DrainZoneInfo(zones.get(i));
      if (curr.dzmm == null) {
        continue;
      }
      
      //
      // Check if drain zone can show text
      //
  
      FabricDisplayOptions options = FabricDisplayOptionsManager.getMgr().getDisplayOptions();
      curr.setGreaterThanMinDZ(curr.diff + 1 >= options.getMinDrainZone());
      // + 1 because #links in (a,b) is b-a+1
      
      // Drain zone sizing / rotation:
      
      curr.setTextSize(target, fonts_, bigToLittle_, frc);
      
	    //
	    // Create drain zone text if there is some:
	    //
	
	    float tnamex = 0;
	    if (curr.dumpRect != null) {
	      // Easiest font height hack is to scale it by ~.67:
	      double dumpScaleHeight = curr.dumpRect.getHeight() * LABEL_FONT_HEIGHT_SCALE_;
	      if (curr.doRotateName) {
          tnamex = (curr.dzmm.min * BioFabricPanel.GRID_SIZE) +
                   ((curr.diff * BioFabricPanel.GRID_SIZE) / 2.0F) +
                   (float)(dumpScaleHeight / 2.0);
          curr.dumpRect.setRect(tnamex - dumpScaleHeight, tnamey - curr.dumpRect.getWidth(),
                                dumpScaleHeight, curr.dumpRect.getWidth());
	      } else {
          tnamex = (curr.dzmm.min * BioFabricPanel.GRID_SIZE) +
                  ((curr.diff * BioFabricPanel.GRID_SIZE) / 2.0F) - ((int) curr.dumpRect.getWidth() / 2);
          curr.dumpRect.setRect(tnamex, tnamey - dumpScaleHeight, curr.dumpRect.getWidth(), dumpScaleHeight);
	      }
	      
	      //
	      // Build the node zone shade rectangle if one is requested:
	      //
	      
	      if (shadeNodes) {
	      	Color col = ((target.nodeRow % 2) == 0) ? superLightBlue_ : superLightPink_;
	        buildANodeShadeRect(curr.dzmm, linkExtents, curr.dumpRect, col, payloadCache);
	      }
	      
	    }
	    
	    if (curr.greaterThanMinDZ) { // only draw text if greater than min drain zone
       
	      //
          // Output the node label and (optional) drain zone label.  If we are using a tiny font for the drain
          // zone, it goes out last to get drawn above the links.
          //
          TextPath drain = new TextPath(Color.BLACK, target.getNodeName(), tnamex, tnamey,
                  curr.dumpRect, curr.doRotateName, curr.font);
          nkk = Integer.toString(nameKeyCount_++);
          pay = new QuadTree.Payload(curr.dumpRect, nkk);
          payloadCache.add(pay);
          if (curr.font == TextPath.FontSizes.TINY) {
            nameKeyToPaintThird_.put(nkk, drain);
            nameKeyToNodeID_.put(nkk, target.getNodeID());
          } else {
            nameKeyToPaintSecond_.put(nkk, drain);
            nameKeyToNodeID_.put(nkk, target.getNodeID());
          }
        }
      
      if (curr.dumpRect != null) {
        rectList.add((Rectangle2D) curr.dumpRect.clone());
      }
    }
    drainMap.put(target.getNodeIDWithName(), rectList);
    
    return;
  } 
  
  /***************************************************************************
   ** Contains the properties required to draw a drain zone
   */
  
  private static class DrainZoneInfo {
    
    private int diff;
    private TextPath.FontSizes font;
    private boolean doRotateName;
    private Rectangle2D dumpRect;
    private MinMax dzmm;
    private boolean greaterThanMinDZ;
    

    DrainZoneInfo(BioFabricNetwork.DrainZone dz) {
      this.dzmm = dz.getMinMax().clone();
      this.diff = this.dzmm.max - this.dzmm.min;
    }
  
    /***************************************************************************
     **
     ** Set if drain zone can show text or not
     */
    
    private void setGreaterThanMinDZ(boolean greaterThanMinDZ) {
      this.greaterThanMinDZ = greaterThanMinDZ;
    }
    
    /***************************************************************************
	  **
	  ** Set the drain zone text size
	  */
	  
	  private void setTextSize(BioFabricNetwork.NodeInfo target, Map<TextPath.FontSizes, Font> fonts, 
	  		                     List<TextPath.FontSizes> bigToSmall, FontRenderContext frc) { 
	  	this.doRotateName = false;      
	    this.diff = this.dzmm.max - this.dzmm.min;
	    
	    Rectangle2D bounds = null;
	    for (TextPath.FontSizes size : bigToSmall) {  
	      bounds = fonts.get(size).getStringBounds(target.getNodeName(), frc);
	      if (bounds.getWidth() <= (BioFabricPanel.GRID_SIZE * this.diff)) {
	        this.font = size;
	        this.dumpRect = bounds;
          bounds = null;
	        break;
	      }
	    }
	    
	    if (bounds != null) {
	      this.font = TextPath.FontSizes.TINY;
	      this.dumpRect = bounds;
	      this.doRotateName = true;
	    }
	    return;
	  }
  }
  
  /***************************************************************************
  **
  ** Build a backRect
  */
  
  private void buildANodeShadeRect(MinMax dzmm, Map<Integer, MinMax> linkExtents, 
  		                             Rectangle2D dumpRect, Color col, ArrayList<QuadTree.Payload> payloadCache) {  
    int minRow = Integer.MAX_VALUE;
    int maxRow = Integer.MIN_VALUE;       
    for (int i = dzmm.min; i <= dzmm.max; i++) {
      MinMax range = linkExtents.get(Integer.valueOf(i));
      if (range != null) {
        if (minRow > range.min) {
          minRow = range.min;
        }
        if (maxRow < range.max) {
          maxRow = range.max;
        }
      }
    }
    int rectLeft = (int)Math.floor((dzmm.min * BioFabricPanel.GRID_SIZE) - BB_HALF_WIDTH_ - (STROKE_SIZE / 2.0));
    int topRow = (int)Math.floor((minRow * BioFabricPanel.GRID_SIZE) - BB_HALF_WIDTH_ - (STROKE_SIZE / 2.0));
    int rectTop = (dumpRect == null) ? topRow : Math.min((int)dumpRect.getMinY(), topRow);
    int rectRight = (int)Math.ceil((dzmm.max * BioFabricPanel.GRID_SIZE) + BB_HALF_WIDTH_ + (STROKE_SIZE / 2.0));
    int rectWidth = rectRight - rectLeft;
    int rectBot = (int)Math.floor((maxRow * BioFabricPanel.GRID_SIZE) + BB_HALF_WIDTH_ + (STROKE_SIZE / 2.0));
    int rectHeight = rectBot - rectTop;
    Rectangle rect = new Rectangle(rectLeft, rectTop, rectWidth, rectHeight);
    BoxPath npp = new BoxPath(col, rect);
    String nkk = Integer.toString(nameKeyCount_++);
    QuadTree.Payload pay = new QuadTree.Payload(rect, nkk);
    payloadCache.add(pay);
    nameKeyToPaintFirst_.put(nkk, npp);
    return;
  }
 
  /***************************************************************************
  **
  ** Let others know how big we make annotation pads
  */

  public static int calcAnnotationPad(MinMax fullExtents) {  
    int minExtent = fullExtents.min; //Integer.MAX_VALUE;
    int maxExtent = fullExtents.max; //Integer.MIN_VALUE;
    int diff = maxExtent - minExtent;
    int pad = (int)(diff * BioFabricPanel.GRID_SIZE * 0.05);
    pad = UiUtil.forceToGridValueInt(Math.max(pad, 200), BioFabricPanel.GRID_SIZE);
    return (pad);
  }

  /***************************************************************************
  **
  ** Build an annotation backRect. For Nodes, isHoriz is true.
  */
  
  private void buildAnAnnotationRect(MinMax dzmm, String name, Color col, boolean isHoriz, 
                                     Map<Integer, MinMax> extents, FontRenderContext frc, 
                                     MinMax fullExtents, ArrayList<QuadTree.Payload> payloadCache,
                                     Integer extentCap) {  
    
    int minExtent = fullExtents.min; //Integer.MAX_VALUE;
    int maxExtent = (extentCap == null) ? fullExtents.max : extentCap - 1; //Integer.MIN_VALUE;
    
    int pad = calcAnnotationPad(fullExtents);
      
    int rectLeft;
    int rectRight;
    int rectTop;
    int rectBot;
    
    if (isHoriz) {
      rectLeft = (minExtent * BioFabricPanel.GRID_SIZE) - pad;
      rectRight = (maxExtent * BioFabricPanel.GRID_SIZE) + (BioFabricPanel.GRID_SIZE / 2);
      rectTop = (dzmm.min * BioFabricPanel.GRID_SIZE) - (BioFabricPanel.GRID_SIZE / 2);
      rectBot = (dzmm.max * BioFabricPanel.GRID_SIZE) + (BioFabricPanel.GRID_SIZE / 2);
    } else {
      rectLeft = (dzmm.min * BioFabricPanel.GRID_SIZE) - (BioFabricPanel.GRID_SIZE / 2);
      rectRight = (dzmm.max * BioFabricPanel.GRID_SIZE) + (BioFabricPanel.GRID_SIZE / 2);
      rectTop = (minExtent * BioFabricPanel.GRID_SIZE) - pad;
      rectBot = (maxExtent * BioFabricPanel.GRID_SIZE) + (BioFabricPanel.GRID_SIZE / 2);
    }
    int rectWidth = rectRight - rectLeft;
    int rectHeight = rectBot - rectTop;
    Rectangle rect = new Rectangle(rectLeft, rectTop, rectWidth, rectHeight);
    
    BoxPath npp = new BoxPath(col, rect);
    String nkk = Integer.toString(nameKeyCount_++);
    QuadTree.Payload pay = new QuadTree.Payload(rect, nkk);
    payloadCache.add(pay);
    if (isHoriz) {
      nameKeyToPaintZero_.put(nkk, npp);
    } else {
      nameKeyToPaintOneQuarter_.put(nkk, npp);
    }
    
    TextPath.FontSizes useFont = null;
    Rectangle2D useBounds = null;
    Rectangle2D bounds = null;
    boolean rotate = false;

    for (TextPath.FontSizes size : bigToLittle_) {  
      bounds = fonts_.get(size).getStringBounds(name, frc);
      double h = bounds.getHeight();
      double w = bounds.getWidth();
      
      
      if (isHoriz) {
      	if ((h < rect.getHeight()) && (w < pad)) {
          useFont = size;
          useBounds = bounds;
          break;
      	}
      } else {
        if ((w < rect.getWidth()) && (h < pad)) {
          useFont = size;
          useBounds = bounds;
          break;
      	}
      }
    }
       
    if (useBounds == null) {
      useFont = TextPath.FontSizes.TINY;
      useBounds = bounds;
      rotate = true;
    }
    
    double namey;
	  double namex;
	  double scaleHeight = useBounds.getHeight();
	  double scaleWidth = useBounds.getWidth();
	  
	  // Precise vertical placement in short annotation blocks:
	  LineMetrics metrics = fonts_.get(useFont).getLineMetrics(name, frc);
	  
    if (isHoriz) {
	    namey = rect.getCenterY() - (0.5 * scaleHeight) + metrics.getAscent();
	    namex = rect.getX();      
    } else {
    	if (rotate) {
    		namey = rect.getY() + (0.5 * pad) + (0.5 * scaleWidth);
	      namex = rect.getCenterX() - (0.5 * scaleHeight) + metrics.getAscent();
	      double hold = scaleHeight;
	      scaleHeight = scaleWidth;
	      scaleWidth = hold;
    	} else {
	      namey = rect.getY() + (0.5 * pad) - (0.5 * scaleHeight) + metrics.getAscent();
	      namex = rect.getCenterX() - (0.5 * scaleWidth);
    	}
    }
    useBounds.setRect(namex, namey, scaleWidth, scaleHeight);
    
    //
    // Get the name bounds into the map and the extents into the QuadTree:
    //
    
    TextPath tpp = new TextPath(Color.BLACK, name, namex, namey, useBounds, rotate, useFont);
    String tkk = Integer.toString(nameKeyCount_++);
    QuadTree.Payload textPay = new QuadTree.Payload(useBounds, tkk);
    payloadCache.add(textPay);
    nameKeyToPaintOneHalf_.put(nkk, tpp);
    
    return;
  }
  
    ////////////////////////////////////////////////////////////////////////////
  //
  // PUBLIC INNER CLASSES
  //
  ////////////////////////////////////////////////////////////////////////////

  /***************************************************************************
  **
  ** Floaters
  */  
  
  public static class FloaterSet {
    public Rectangle floater;
    public Rectangle tourRect;
    public Rectangle currSelRect;
    
    public FloaterSet(Rectangle floater, Rectangle tourRect, Rectangle currSelRect) {
      this.floater = floater;
      this.tourRect = tourRect;
      this.currSelRect = currSelRect;    
    }
    
    public void clear() {
      floater = null;
      tourRect = null;
      currSelRect = null;
      return;
    }
    
    public boolean isEmpty() {
      return ((floater == null) &&
              (tourRect == null) &&
              (currSelRect == null));
    }   
  } 
  
  /***************************************************************************
  **
  ** Used to reduce painting to a limited selection
  */  
  
  public static class Reduction {
    Set<Integer> paintRows;
    Set<Integer> paintCols;
    Set<NID> paintNames;
    
    public Reduction(Set<Integer> rows, Set<Integer> cols, Set<NID> names) {
      this.paintRows = rows;
      this.paintCols = cols;
      this.paintNames = names;    
    }
    
    public boolean somethingToPaint() {
    	return (!paintRows.isEmpty() || !paintCols.isEmpty() || !paintNames.isEmpty());	
    }
    
  } 
}
```
