## Supplementary material for "BioFabric Visualization of Network Alignments": BioFabric Source Code: RasterCache.java

```
/*
**    Copyright (C) 2003-2017 Institute for Systems Biology 
**                            Seattle, Washington, USA. 
**
**    This library is free software; you can redistribute it and/or
**    modify it under the terms of the GNU Lesser General Public
**    License as published by the Free Software Foundation; either
**    version 2.1 of the License, or (at your option) any later version.
**
**    This library is distributed in the hope that it will be useful,
**    but WITHOUT ANY WARRANTY; without even the implied warranty of
**    MERCHANTABILITY or FITNESS FOR A PARTICULAR PURPOSE.  See the GNU
**    Lesser General Public License for more details.
**
**    You should have received a copy of the GNU Lesser General Public
**    License along with this library; if not, write to the Free Software
**    Foundation, Inc., 59 Temple Place, Suite 330, Boston, MA  02111-1307  USA
*/

package org.systemsbiology.biofabric.ui.render;

import java.awt.image.BufferedImage;
import java.awt.image.DataBufferByte;
import java.awt.image.DataBufferInt;
import java.awt.image.DirectColorModel;
import java.awt.image.Raster;
import java.awt.image.WritableRaster;
import java.io.BufferedInputStream;
import java.io.BufferedOutputStream;
import java.io.File;
import java.io.FileInputStream;
import java.io.FileOutputStream;
import java.io.IOException;
import java.util.ArrayList;
import java.util.HashMap;
import java.util.zip.DataFormatException;
import java.util.zip.Deflater;
import java.util.zip.Inflater;

import org.systemsbiology.biofabric.util.UiUtil;


/****************************************************************************
**
** This is a cache to hold tiling images using their compressed underlying
** data buffers.
*/

public class RasterCache {
   
  ////////////////////////////////////////////////////////////////////////////
  //
  // PRIVATE INSTANCE MEMBERS
  //
  ////////////////////////////////////////////////////////////////////////////
   
  private String cachePref_;  
  private HashMap<String, BytesWithMeta> bufferCache_;
  private HashMap<String, String> handleToFile_;
  private HashMap<String, InfoForImage> infoForImage_;
  private ArrayList<String> queue_;
  private int nextHandle_;
  private int maxMeg_;
  private int currSize_;
  private ShiftData shiftData_;

  ////////////////////////////////////////////////////////////////////////////
  //
  // PUBLIC CONSTRUCTORS
  //
  ////////////////////////////////////////////////////////////////////////////

  /***************************************************************************
  **
  ** Constructor
  */

  public RasterCache(String cachePref, int maxMeg, DirectColorModel cMod) {
    cachePref_ = cachePref;
    bufferCache_ = new HashMap<String, BytesWithMeta>();
    handleToFile_ = new HashMap<String, String>();
    infoForImage_ = new HashMap<String, InfoForImage>();
    queue_ = new ArrayList<String>();
    nextHandle_ = 0;
    if (maxMeg == 0) {
      throw new IllegalArgumentException();
    }
    maxMeg_ = maxMeg * 1000000;
    currSize_ = 0;
    shiftData_ = new ShiftData(cMod);

  }

  ////////////////////////////////////////////////////////////////////////////
  //
  // PUBLIC METHODS
  //
  ////////////////////////////////////////////////////////////////////////////
  
  /***************************************************************************
  **
  ** Get an image from the cache; returns null if no image. Cache management
  ** now occurs at the raster level. This just uses the retrieved raster to stock the
  ** BufferedImage.
  */
  
  public BufferedImage getAnImage(String handle, ImgAndBufPool bis) throws IOException {
    BytesWithMeta bwm = getADataBuffer(handle, bis);
    if (bwm == null) {
    	return (null);
    }
    InfoForImage ifi = this.infoForImage_.get(handle); 
    BufferedImage bi = bufToImage(bwm, bis, ifi);
    return (bi);
  }
 
  /***************************************************************************
  **
  ** Cache an image, return a handle, *recycle the image*
  */
  
  public String cacheAnImage(BufferedImage bi, ImgAndBufPool bis) throws IOException {

  	InfoForImage ifi = new InfoForImage(bi);	
  	BytesWithMeta bwm = imageToBuf(bi, bis, ifi);
  	bis.returnImage(bi);
    maintainSize(bwm.buf.length, bis);
 
    String handle = Integer.toString(nextHandle_++);
    bufferCache_.put(handle, bwm);
    infoForImage_.put(handle, ifi);
    queue_.add(0, handle);
    currSize_ += bwm.buf.length;
    
    return (handle);
  }

  /***************************************************************************
  **
  ** Release resources
  */
  
  public void releaseResources()  {
  	UiUtil.fixMePrintout("DO THIS IN A THREAD_SAFE FASHION!!");
  	return;
  }
 
  /***************************************************************************
  **
  ** Write bytes into ints.
  */
  
  public static void threeBytesToOneInt(byte[] input, int[] output, ShiftData sd) {
    int byteIndex = 0;
    for (int intIndex = 0; intIndex < output.length; intIndex++) {
      output[intIndex] = 0;
      output[intIndex] |= (input[byteIndex++] & 0xFF) << sd.redShift_;
      output[intIndex] |= (input[byteIndex++] & 0xFF) << sd.greenShift_;
      output[intIndex] |= (input[byteIndex++] & 0xFF) << sd.blueShift_;
    }
    return;
  }
  
  /***************************************************************************
  **
  ** Determine shift needed to get contiguous mask to lowest bits
  */
  
  public static int shiftForMask(int mask) {
  	for (int i = 0; i < 32; i++) {
  		if (((mask >> i) & 0x01) == 0x01) {
  			return (i);
  		}
  	}
    throw new IllegalArgumentException();
  }
  
  /***************************************************************************
  **
  ** Get the ints for a TYPE_INT_RGB into bytes
  */
  
  public static void oneIntToThreeBytes(int[] input, byte[] output, ShiftData sd) {
    for (int intIndex = 0; intIndex < input.length; intIndex++) {
    	int byteBase = 3 * intIndex;
    	output[byteBase] = (byte)((input[intIndex] & sd.redMask_) >> sd.redShift_);
    	output[byteBase + 1] = (byte)((input[intIndex] & sd.greenMask_) >> sd.greenShift_);
    	output[byteBase + 2] = (byte)((input[intIndex] & sd.blueMask_) >> sd.blueShift_);
    }
    return;
  }

  /***************************************************************************
  **
  ** Drop the data associated with the handle. 
  */
  
  public void dropAnImage(String handle, ImgAndBufPool bis) throws IOException {
  	//
  	// Since we don't store images, we just need to drop the data buffer!
  	//
  	BytesWithMeta bye = getADataBuffer(handle, bis);
    if (bye == null) {
      return;
    }
    currSize_ -= bye.buf.length;
    if (currSize_ < 0) {
      currSize_ = 0;
    }
    
    queue_.remove(handle);
    bufferCache_.remove(handle);
    infoForImage_.remove(handle);
    bis.returnByteBuf(bye.buf);

    String fileName = handleToFile_.get(handle);
    if (fileName != null) {
      File dropFile = new File(fileName);
      if (dropFile.exists()) {
        dropFile.delete();
      }
      handleToFile_.remove(handle);
    }
    return;
  }
  
  /***************************************************************************
  **
  ** Replace the image with the given handle. *recycles the image*
  */
  
  public String replaceAnImage(String handle, BufferedImage bi, ImgAndBufPool bis) throws IOException {
  	
    BytesWithMeta bye = getADataBuffer(handle, bis);
    currSize_ -= bye.buf.length;
    if (currSize_ < 0) {
      currSize_ = 0;
    }
    bis.returnByteBuf(bye.buf);

    // Note above getADataBuffer put us at the front of the queue
    
    InfoForImage ifi = new InfoForImage(bi);
  	BytesWithMeta bwm = imageToBuf(bi, bis, ifi);
  	bis.returnImage(bi);
    maintainSize(bwm.buf.length, bis);
 
    bufferCache_.put(handle, bwm);
    infoForImage_.put(handle, ifi);
    queue_.add(0, handle);
    currSize_ += bwm.buf.length;
    
    String fileName = handleToFile_.get(handle);
    if (fileName != null) {
      File holdFile = new File(fileName);
      writeBufferToFile(bwm, holdFile, bis);
    }

    return (handle);
  }
   
  /***************************************************************************
  **
  ** Manage in-memory cache, toss least recently used. Tossed byte array recycled.
  */
  
  private void maintainSize(int sizeEst, ImgAndBufPool bis) throws IOException {
    while (((sizeEst + currSize_) > maxMeg_) && (queue_.size() > 0)) {
      String goodBye = queue_.remove(queue_.size() - 1);
      BytesWithMeta bwm = bufferCache_.remove(goodBye);
      String fileName = handleToFile_.get(goodBye);
      if (fileName == null) {
      	//System.out.println("Flushing to file cache to maintain size");
        File holdFile = getAFile();
        currSize_ -= bwm.buf.length;
        if (currSize_ < 0) {
          currSize_ = 0;
        }
        writeBufferToFile(bwm, holdFile, bis);
        bis.returnByteBuf(bwm.buf);
        handleToFile_.put(goodBye, holdFile.getAbsolutePath());
        fileCacheReport();
      }
    }
    // System.out.println("Curr mem cache: " + currSize_);
    return;
  }
  
  /***************************************************************************
  **
  ** Get temp file:
  ** On Mac 10.5.8, JDK 1.6, this seems to go into:
  ** /private/var/folders/[2 characters]/[Random string of characters]/-Tmp-
  ** Note: To cd into -Tmp-, use "cd -- -Tmp-"
  */
  
  private File getAFile() throws IOException {
    File file = null;
    File dir = null;
    if (cachePref_ != null) {
      dir = new File(cachePref_);
      if (!dir.exists() || !dir.isDirectory()) {
        throw new IOException();
      }
      file = File.createTempFile("BioFabric", ".tmp", dir);
    } else {
      file = File.createTempFile("BioFabric", ".tmp");
    }
    file.deleteOnExit();
    return (file);
  }

  /***************************************************************************
  ** 
  ** Stats on image file cache:
  */
  
  public long fileCacheReport() {
     
  	long length = 0;
  	int numFile = 0;
   
    for (String holdFilePath : handleToFile_.values()) {    
      File holdFile = new File(holdFilePath);
      if (!holdFile.exists()) {
        continue;
      }
      if (numFile == 0) {
        System.out.println("Directory: " + holdFile.getParentFile().getAbsolutePath());
      }
  	  length += holdFile.length();
  	  numFile++;
    }
    System.out.println("Total bytes used: " + length);
    System.out.println("Total file count: " + numFile);
    return (length);
  }

  /***************************************************************************
  ** 
  ** Write out a DataBuffer for file. Buffer is NOT recycled.
  */

  private void writeBufferToFile(BytesWithMeta bwm, File file, ImgAndBufPool bis) throws IOException {
  	long prewrite = Runtime.getRuntime().freeMemory();
  	BufferedOutputStream out = null;
  	try {
  	  out = new BufferedOutputStream(new FileOutputStream(file));
  	  out.write(bwm.buf, 0, bwm.used);
  	} finally {
  		if (out != null) {
  		  out.close();
  		}
  	}
    return;
  }
  
  /***************************************************************************
  ** 
  ** Read in a DataBuffer
  */

  private BytesWithMeta readBufferFromFile(File file, InfoForImage ifi, ImgAndBufPool bis) throws IOException {
  	long preread = Runtime.getRuntime().freeMemory();
  	byte[] buf = bis.fetchByteBuf(ifi.compressedNumBytes);
  	BufferedInputStream in = null;
  	int off = 0;
  	try {
	  	in = new BufferedInputStream(new FileInputStream(file));
	  	while (true) {
	  	  int num = in.read(buf, off, buf.length - off);
	  	  if (num == -1) {
	  	  	break;
	  	  }
	  	  off += num;
	  	}
  	} finally {
  		if (in != null) {
  		  in.close();
  		}
  	}
    if (ifi.compressedNumBytes != off) {
    	throw new IOException();
    }
    return (new BytesWithMeta(buf, off));
  }

  /***************************************************************************
  **
  ** Get a BytesWithMeta from the cache; returns null if no BytesWithMeta
  */
  
  private BytesWithMeta getADataBuffer(String handle, ImgAndBufPool bis) throws IOException {
  	
    //
  	// We have a hit in the memory cache. Retrieve it, and move it to the front of the
  	// queue since it now is the most recently used:
  	//
  	
    BytesWithMeta retval = bufferCache_.get(handle);
    if (retval != null) {
      queue_.remove(handle);
      queue_.add(0, handle);
      return (retval);
    }
    
    //
    // Not in memory cache, look for file:
    //
   
    String fileName = handleToFile_.get(handle);
   
    if (fileName == null) {
      return (null);
    }
    
    //
    // File is gone (should not happen...handleToFile_ should be authoritative):
    //
    
    File holdFile = new File(fileName);
    if (!holdFile.exists()) {
      return (null);
    }
    
    //
    // Get the image in from file:
    //
    
    InfoForImage ifi = infoForImage_.get(handle); 
    retval = readBufferFromFile(holdFile, ifi, bis);

    //
    // Manage in-memory cache size, toss least recently used:
    //

    int size = retval.buf.length;
    maintainSize(size, bis);
 
    //
    // Install it in the memory cache, again putting it as most recently used
    //

    bufferCache_.put(handle, retval);
    queue_.add(0, handle);
    currSize_ += size;
    return (retval);
  }
 
  /***************************************************************************
  ** 
  ** Convert an image into a buffer. Image is not recycled; caller is responsible.
  */

  private BytesWithMeta imageToBuf(BufferedImage bi, ImgAndBufPool bis, InfoForImage ifi)  { 
  	Raster rast = bi.getRaster();
  	DataBufferInt dbb = (DataBufferInt)rast.getDataBuffer();
  	int[] data = dbb.getData();
  	byte[] byteData = bis.fetchByteBuf(data.length * 3);
  	oneIntToThreeBytes(data, byteData, shiftData_);
  	
  	// We do not know how big the result will be. So we use a buffer the same size as the
  	// data. IS THAT A PROBLEM???? What about METADATA?
  	byte[] output = bis.fetchByteBuf(byteData.length);
    Deflater deflate = new Deflater();
    deflate.setInput(byteData);
    deflate.finish();
    int compressedDataLength = deflate.deflate(output);
    byte[] result = bis.fetchByteBuf(compressedDataLength);
    System.arraycopy(output, 0, result, 0, compressedDataLength);
    bis.returnByteBuf(output);
    bis.returnByteBuf(byteData);
    ifi.setCompressedSize(compressedDataLength);
    return (new BytesWithMeta(result, compressedDataLength));
  }
    
  /***************************************************************************
  ** 
  ** Convert a buffer into an image. Buffer is not recycled, caller is responsible.
  */

  private BufferedImage bufToImage(BytesWithMeta bwm, ImgAndBufPool bis, InfoForImage ifi) throws IOException {   	
  	Inflater inflate = new Inflater();
    byte[] decomp = bis.fetchByteBuf(ifi.uncompressedNumBytes);
    inflate.setInput(bwm.buf, 0, bwm.used);
    int resultLength;
    try {
      resultLength = inflate.inflate(decomp);
    } catch (DataFormatException dfex) {
    	throw new IOException();
    }
    inflate.end();
    
    int[] intData = bis.fetchBuf(decomp.length / 3);
    threeBytesToOneInt(decomp, intData, shiftData_);
 
    BufferedImage bi = bis.fetchImage(ifi.width, ifi.height, ifi.type);
  	WritableRaster biRast = bi.getRaster();
  	biRast.setDataElements(0, 0, ifi.width, ifi.height, intData);
    bis.returnByteBuf(decomp);
    bis.returnBuf(intData);
    return (bi);
  }
  
  ////////////////////////////////////////////////////////////////////////////
  //
  // INNER CLASSES
  //
  ////////////////////////////////////////////////////////////////////////////
  
  /***************************************************************************
  ** 
  ** Since we are working with compressed data of varying lengths, but
  ** canonical buffer lengths, we need to keep track of how much of the
  ** buffer holds useful data.
  */ 
 
  private static class BytesWithMeta {
  
    byte[] buf;
    int used;
  
    BytesWithMeta(byte[] buf, int used) {
    	this.buf = buf;
      this.used = used;
    }
  }
  
  /***************************************************************************
  ** 
  ** When we reconstitute an image from file, we need to know what image type
  ** to expect.
  */ 
 
  private static class InfoForImage {
  
  	int uncompressedNumBytes;
  	int compressedNumBytes;
    int width;
    int height;
    int type;
    
    InfoForImage(BufferedImage bi) {
      Raster rast = bi.getRaster();
      DataBufferInt dbb = (DataBufferInt)rast.getDataBuffer();
      int[] dest = dbb.getData();
      this.uncompressedNumBytes = dest.length * 3;	
    	this.compressedNumBytes = 0; // Don't know yet
    	this.width = bi.getWidth();
      this.height = bi.getHeight();
      this.type = bi.getType();
	  }
    
    void setCompressedSize(int size) {
    	this.compressedNumBytes = size;
	  } 
  }
  
   /***************************************************************************
  ** 
  ** Precalc shift information
  */ 
 
  public static class ShiftData {
  
    int redMask_;
    int greenMask_;
    int blueMask_;
    int redShift_;
    int greenShift_;
    int blueShift_;	
    
    public ShiftData(DirectColorModel cMod) {
      redMask_ = cMod.getRedMask();
      greenMask_ = cMod.getGreenMask();
      blueMask_ = cMod.getBlueMask();
      redShift_ = shiftForMask(redMask_);
      greenShift_ = shiftForMask(greenMask_);
      blueShift_ = shiftForMask(blueMask_);	
	  }
  }
}
```
