## Supplementary material for "BioFabric Visualization of Network Alignments": BioFabric Source Code: TextPath.java

```
/*
**    Copyright (C) 2003-2017 Institute for Systems Biology 
**                            Seattle, Washington, USA. 
**
**    This library is free software; you can redistribute it and/or
**    modify it under the terms of the GNU Lesser General Public
**    License as published by the Free Software Foundation; either
**    version 2.1 of the License, or (at your option) any later version.
**
**    This library is distributed in the hope that it will be useful,
**    but WITHOUT ANY WARRANTY; without even the implied warranty of
**    MERCHANTABILITY or FITNESS FOR A PARTICULAR PURPOSE.  See the GNU
**    Lesser General Public License for more details.
**
**    You should have received a copy of the GNU Lesser General Public
**    License along with this library; if not, write to the Free Software
**    Foundation, Inc., 59 Temple Place, Suite 330, Boston, MA  02111-1307  USA
*/

package org.systemsbiology.biofabric.ui.render;

import java.awt.Color;
import java.awt.Font;
import java.awt.Graphics2D;
import java.awt.Rectangle;
import java.awt.geom.AffineTransform;
import java.awt.geom.Rectangle2D;
import java.util.Map;

/****************************************************************************
**
** These are used to store strings to draw
*/

public class TextPath {
 
  ////////////////////////////////////////////////////////////////////////////
  //
  // PUBLIC CONSTANTS
  //
  //////////////////////////////////////////////////////////////////////////// 
  
	public enum FontSizes { TINY, HUGE, MED, MED_SMALL, SMALL, GINORMOUS }

  ////////////////////////////////////////////////////////////////////////////
  //
  // PRIVATE INSTANCE MEMBERS
  //
  //////////////////////////////////////////////////////////////////////////// 	
	
  private Color col;
  private String name;
  private double nameX;
  private double nameY;
  private boolean doRotateName;
  private Rectangle2D nameRect;
  private FontSizes font;

  ////////////////////////////////////////////////////////////////////////////
  //
  // PUBLIC CONSTRUCTORS
  //
  ////////////////////////////////////////////////////////////////////////////
  
  /***************************************************************************
  **
  ** Used for node labels for nodes without drain zones
  */
 
  public TextPath(Color col, String name, double x, double y, Rectangle2D nameRect, boolean doRotateName, FontSizes font) {
    this.col = col;
    this.name = name;
    nameX = x;
    nameY = y;
    this.nameRect = nameRect;
    this.doRotateName = doRotateName;
    this.font = font;
  }

  ////////////////////////////////////////////////////////////////////////////
  //
  // PUBLIC METHODS
  //
  ////////////////////////////////////////////////////////////////////////////

  /***************************************************************************
  **
  ** Draw it:
  */
 
  int paint(Graphics2D g2, Rectangle bounds, Map<TextPath.FontSizes, Font> fonts) {
    int didPaint = 0;
    g2.setPaint(col); 
    if (nameRect != null) {
      if ((bounds == null) ||
          ((nameRect.getMaxX() > bounds.getMinX()) && 
           (nameRect.getMinX() < bounds.getMaxX()) && 
           (nameRect.getMaxY() > bounds.getMinY()) && 
           (nameRect.getMinY() < bounds.getMaxY()))) {          
        if (doRotateName) {
          g2.setFont(fonts.get(FontSizes.TINY));
          drawRotString(g2, name, nameX, nameY, 0.0, 0.0);
          didPaint++;         
        } else {
          Font useit = fonts.get(font);
          g2.setFont(useit);
          g2.drawString(name, (int)nameX, (int)nameY); 
          didPaint++;
        }
      }
    }
    return (didPaint);
  } 
  
  ////////////////////////////////////////////////////////////////////////////
  //
  // PRIVATE METHODS
  //
  ////////////////////////////////////////////////////////////////////////////

  /***************************************************************************
  **
  ** Draw rotated strings:
  */ 
        
  private void drawRotString(Graphics2D g2, String name, double x, double y, double ptx, double pty) {
    AffineTransform sav = g2.getTransform();
    AffineTransform forRot = (AffineTransform)sav.clone();
    if ((ptx != 0.0) || (pty != 0.0)) {
      forRot.translate(ptx, pty);
    }
    forRot.rotate(-Math.PI / 2.0, x, y);
    g2.setTransform(forRot);
    g2.drawString(name, (float)x, (float)y);
    g2.setTransform(sav);
    return;
  }  

} 
```
