## Supplementary material for "BioFabric Visualization of Network Alignments": BioFabric Source Code: ZoomPresentation.java

```
/*
**    Copyright (C) 2003-2017 Institute for Systems Biology 
**                            Seattle, Washington, USA. 
**
**    This library is free software; you can redistribute it and/or
**    modify it under the terms of the GNU Lesser General Public
**    License as published by the Free Software Foundation; either
**    version 2.1 of the License, or (at your option) any later version.
**
**    This library is distributed in the hope that it will be useful,
**    but WITHOUT ANY WARRANTY; without even the implied warranty of
**    MERCHANTABILITY or FITNESS FOR A PARTICULAR PURPOSE.  See the GNU
**    Lesser General Public License for more details.
**
**    You should have received a copy of the GNU Lesser General Public
**    License along with this library; if not, write to the Free Software
**    Foundation, Inc., 59 Temple Place, Suite 330, Boston, MA  02111-1307  USA
*/

package org.systemsbiology.biofabric.ui;

import java.awt.Rectangle;

/****************************************************************************
**
** Interface for zoom presentation
*/

public interface ZoomPresentation {
 
  ////////////////////////////////////////////////////////////////////////////
  //
  // PUBLIC METHODS
  //
  ////////////////////////////////////////////////////////////////////////////

  /***************************************************************************
  **
  ** Return the required size of the layout
  */
  
  public Rectangle getRequiredSize();
 
  /***************************************************************************
  **
  ** Return the required size of the selected items.  Null if nothing is
  ** selected.
  */
  
  public Rectangle getSelectionSize(); 

  /***************************************************************************
  **
  ** Return the required size of the current zoom selection item.  Null if nothing is
  ** selected.
  */
  
  public Rectangle getCurrentSelectionSize();   
 
  /***************************************************************************
  **
  ** Answer if we have a current selection
  */
  
  public boolean haveCurrentSelection();
  
  /***************************************************************************
  **
  ** Answers if we have multiple selections (i.e. can cycle through selections) 
  */
  
  public boolean haveMultipleSelections();
 
  /***************************************************************************
  **
  ** Bump to the next selection for zoom
  */
  
  public void bumpNextSelection();
  
  /***************************************************************************
  **
  ** Bump to the previous selection for zoom
  */
  
  public void bumpPreviousSelection();
  
  /***************************************************************************
  **
  ** Set the zoom
  */
  
  public void setPresentationZoomFactor(double zoom);
    
}
```
