## Supplementary material for "BioFabric Visualization of Network Alignments": BioFabric Source Code: ChoiceContent.java

```
/*
**    Copyright (C) 2003-2018 Institute for Systems Biology 
**                            Seattle, Washington, USA. 
**
**    This library is free software; you can redistribute it and/or
**    modify it under the terms of the GNU Lesser General Public
**    License as published by the Free Software Foundation; either
**    version 2.1 of the License, or (at your option) any later version.
**
**    This library is distributed in the hope that it will be useful,
**    but WITHOUT ANY WARRANTY; without even the implied warranty of
**    MERCHANTABILITY or FITNESS FOR A PARTICULAR PURPOSE.  See the GNU
**    Lesser General Public License for more details.
**
**    You should have received a copy of the GNU Lesser General Public
**    License along with this library; if not, write to the Free Software
**    Foundation, Inc., 59 Temple Place, Suite 330, Boston, MA  02111-1307  USA
*/

package org.systemsbiology.biofabric.util;

/****************************************************************************
**
** Utility for choice menus
*/

public class ChoiceContent {
  
  public String name;
  public int val;
  
  public ChoiceContent(String name, int val) {
    this.name = name;
    this.val = val;
  }
  
  public String toString() {
    return (name);
  }
  
  public boolean equals(Object other) {
    if (this == other) {
      return (true);
    }
    if (!(other instanceof ChoiceContent)) {
      return (false);
    }
    ChoiceContent occ = (ChoiceContent)other;
    if (this.val != occ.val) {
      return (false);
    }
    
    if (name == null) {
      return (occ.name == null);
    }

    return (name.equals(occ.name));
  }
  
  public int hashCode() {
    return (val);
  }
}
```
