## Supplementary material for "BioFabric Visualization of Network Alignments": BioFabric Source Code: ColorLabel.java

```
/*
**    Copyright (C) 2003-2018 Institute for Systems Biology 
**                            Seattle, Washington, USA. 
**
**    This library is free software; you can redistribute it and/or
**    modify it under the terms of the GNU Lesser General Public
**    License as published by the Free Software Foundation; either
**    version 2.1 of the License, or (at your option) any later version.
**
**    This library is distributed in the hope that it will be useful,
**    but WITHOUT ANY WARRANTY; without even the implied warranty of
**    MERCHANTABILITY or FITNESS FOR A PARTICULAR PURPOSE.  See the GNU
**    Lesser General Public License for more details.
**
**    You should have received a copy of the GNU Lesser General Public
**    License along with this library; if not, write to the Free Software
**    Foundation, Inc., 59 Temple Place, Suite 330, Boston, MA  02111-1307  USA
*/

package org.systemsbiology.biofabric.util;

import java.awt.Color;
import java.awt.GridBagLayout;
import java.awt.GridBagConstraints;

import javax.swing.JPanel;
import javax.swing.JLabel;

/****************************************************************************
**
** Utility for bounds creation
*/

public class ColorLabel extends JPanel {

  ////////////////////////////////////////////////////////////////////////////
  //
  // PUBLIC CONSTRUCTORS
  //
  ////////////////////////////////////////////////////////////////////////////  

  private JPanel colorSwatch_;
  private JLabel label_;
  
  ////////////////////////////////////////////////////////////////////////////
  //
  // PUBLIC CONSTRUCTORS
  //
  ////////////////////////////////////////////////////////////////////////////

  /***************************************************************************
  **
  ** Constructor
  */

  public ColorLabel(Color col, String name) {
    
    setLayout(new GridBagLayout());
    GridBagConstraints gbc = new GridBagConstraints();     
    
    colorSwatch_ = new JPanel();
    colorSwatch_.setBackground(col);
    label_ = new JLabel(name);
    UiUtil.gbcSet(gbc, 0, 0, 2, 1, UiUtil.BO, 0, 0, 5, 5, 5, 5, UiUtil.W, 0.0, 1.0);       
    add(colorSwatch_, gbc);
    
    UiUtil.gbcSet(gbc, 2, 0, 5, 1, UiUtil.BO, 0, 0, 5, 5, 5, 5, UiUtil.E, 1.0, 1.0);
    add(label_, gbc);
  }   

  ////////////////////////////////////////////////////////////////////////////
  //
  // PUBLIC METHODS
  //
  ////////////////////////////////////////////////////////////////////////////
  
  /***************************************************************************
  **
  ** Set the color values
  */
      
  public void setColorValues(Color col, String name) {
    colorSwatch_.setBackground(col);
    label_.setText(name);
    return;
  }
}
```
