## Supplementary material for "BioFabric Visualization of Network Alignments": BioFabric Source Code: ColorListRenderer.java

```
/*
**    Copyright (C) 2003-2006 Institute for Systems Biology 
**                            Seattle, Washington, USA. 
**
**    This library is free software; you can redistribute it and/or
**    modify it under the terms of the GNU Lesser General Public
**    License as published by the Free Software Foundation; either
**    version 2.1 of the License, or (at your option) any later version.
**
**    This library is distributed in the hope that it will be useful,
**    but WITHOUT ANY WARRANTY; without even the implied warranty of
**    MERCHANTABILITY or FITNESS FOR A PARTICULAR PURPOSE.  See the GNU
**    Lesser General Public License for more details.
**
**    You should have received a copy of the GNU Lesser General Public
**    License along with this library; if not, write to the Free Software
**    Foundation, Inc., 59 Temple Place, Suite 330, Boston, MA  02111-1307  USA
*/

package org.systemsbiology.biofabric.util;

import java.awt.Color;
import java.awt.Component;
import java.util.List;
import javax.swing.JList;
import javax.swing.ListCellRenderer;

import org.systemsbiology.biofabric.api.util.ExceptionHandler;

/***************************************************************************
**
** Used for rendering a list with a color block
*/

public class ColorListRenderer extends ColorLabel implements ListCellRenderer {
    
  private List values_;

  public ColorListRenderer(List values) {
    super(Color.white, "");
    values_ = values;
  }

  public void setValues(List values) {
    values_ = values;
  }

  public Component getListCellRendererComponent(JList list, 
                                                Object value,
                                                int index,
                                                boolean isSelected, 
                                                boolean hasFocus) {
    try {                                              
      if (value == null) {
        return (this);
      }
      //
      // Java Swing book says this may be needed for combo boxes (not lists):
      //
      if (index == -1) {
        index = list.getSelectedIndex();
        if (index == -1) {
          return (this);
        }
      }
      ColorSource currCol = (ColorSource)values_.get(index);
      setColorValues(currCol.getColor(), currCol.getDescription());
      setBackground((isSelected) ? list.getSelectionBackground() : list.getBackground());
    } catch (Exception ex) {
      ExceptionHandler.getHandler().displayException(ex);
    }      
    return (this);             
  }
   
  public interface ColorSource {
    public Color getColor();
    public String getDescription();
  }
}
```
