## Supplementary material for "BioFabric Visualization of Network Alignments": BioFabric Source Code: ComboBoxEditor.java

```
/*
**    Copyright (C) 2003-2007 Institute for Systems Biology 
**                            Seattle, Washington, USA. 
**
**    This library is free software; you can redistribute it and/or
**    modify it under the terms of the GNU Lesser General Public
**    License as published by the Free Software Foundation; either
**    version 2.1 of the License, or (at your option) any later version.
**
**    This library is distributed in the hope that it will be useful,
**    but WITHOUT ANY WARRANTY; without even the implied warranty of
**    MERCHANTABILITY or FITNESS FOR A PARTICULAR PURPOSE.  See the GNU
**    Lesser General Public License for more details.
**
**    You should have received a copy of the GNU Lesser General Public
**    License along with this library; if not, write to the Free Software
**    Foundation, Inc., 59 Temple Place, Suite 330, Boston, MA  02111-1307  USA
*/


package org.systemsbiology.biofabric.util;

import java.awt.Component;
import javax.swing.JTable;
import javax.swing.JComboBox;
import javax.swing.table.TableCellEditor;

import org.systemsbiology.biofabric.api.util.ExceptionHandler;

import javax.swing.event.CellEditorListener;
import java.util.ArrayList;
import java.util.EventObject;
import java.util.Iterator;
import javax.swing.event.ChangeEvent;

/***************************************************************************
**
** Class for editing combo box table cells
*/
  
public abstract class ComboBoxEditor extends JComboBox implements TableCellEditor {
   
  private ArrayList listeners_;
  protected Object origValue_;
  private int currRow_;
  private int currColumn_;
   
  public ComboBoxEditor(Object valueObject) {
    super();
    valueFill(valueObject);
    listeners_ = new ArrayList();
    currRow_ = -1;
    currColumn_ = -1;      
  }
  
  public abstract void valueFill(Object valueObject);  
    
  public Component getTableCellEditorComponent(JTable table, Object value, 
                                               boolean isSelected,
                                               int row, int column) {
    try {
      if (value == null) {
        return (this);
      }
      currRow_ = row;
      currColumn_ = column;    
      setSelectedIndex(valueToIndex(value));
      table.setRowSelectionInterval(row, row);      
      table.setColumnSelectionInterval(column, column);
      origValue_ = value;
    } catch (Exception ex) {
      ExceptionHandler.getHandler().displayException(ex);
    }

    return (this);             
  }

  public abstract int valueToIndex(Object value);
    
  public int getCurrRow() {
    return (currRow_);
  }
    
  public int getCurrColumn() {
    return (currColumn_);
  }
  
  public void resetValue() {
    if (origValue_ != null) {
      setSelectedIndex(valueToIndex(origValue_));
    }
    return;
  }   

  public boolean entryIsValid() {  
    return ((currRow_ != -1) && (currColumn_ != -1));
  }
        
  public void addCellEditorListener(CellEditorListener l) {
    try {
      this.listeners_.add(l);
    } catch (Exception ex) {
      ExceptionHandler.getHandler().displayException(ex);
    }
    return;
  }
    
  public void cancelCellEditing() {
    try {
      editingCanceled();
    } catch (Exception ex) {
      ExceptionHandler.getHandler().displayException(ex);
    }
    return;
  }

  public abstract Object indexToValue(int index);
    
  public Object getCellEditorValue() {
    Object retval = null;
    try {
      retval = indexToValue(getSelectedIndex());
    } catch (Exception ex) {
      ExceptionHandler.getHandler().displayException(ex);
    }
    return (retval);
  }

  public boolean isCellEditable(EventObject anEvent) {
    return (true);
  }
    
  public void removeCellEditorListener(CellEditorListener l) {
    try {
      listeners_.remove(l);
    } catch (Exception ex) {
      ExceptionHandler.getHandler().displayException(ex);
    }
    return;
  }
    
  public boolean shouldSelectCell(EventObject anEvent) {
    return (true);
  }

  public boolean stopCellEditing() {
    try {
      editingStopped();
    } catch (Exception ex) {
      ExceptionHandler.getHandler().displayException(ex);
    }
    return (true);
  }

  private void editingCanceled() {
    currRow_ = -1;
    currColumn_ = -1;
    if (origValue_ != null) {
      setSelectedIndex(valueToIndex(origValue_));
    }
    ChangeEvent cev = new ChangeEvent(this);
    ArrayList lCopy = new ArrayList(listeners_);
    Iterator lit = lCopy.iterator();
    while (lit.hasNext()) {
      CellEditorListener cel = (CellEditorListener)lit.next();
      cel.editingCanceled(cev);
    }
    return;
  }
    
  private void editingStopped() {
    currRow_ = -1;
    currColumn_ = -1;
    ChangeEvent cev = new ChangeEvent(this);
    ArrayList lCopy = new ArrayList(listeners_);      
    Iterator lit = lCopy.iterator();
    while (lit.hasNext()) {
      CellEditorListener cel = (CellEditorListener)lit.next();
      cel.editingStopped(cev);
    }
    return;
  }    
}
  
```
