## Supplementary material for "BioFabric Visualization of Network Alignments": BioFabric Source Code: ComboBoxEditorTracker.java

```
/*
**    Copyright (C) 2003-2007 Institute for Systems Biology 
**                            Seattle, Washington, USA. 
**
**    This library is free software; you can redistribute it and/or
**    modify it under the terms of the GNU Lesser General Public
**    License as published by the Free Software Foundation; either
**    version 2.1 of the License, or (at your option) any later version.
**
**    This library is distributed in the hope that it will be useful,
**    but WITHOUT ANY WARRANTY; without even the implied warranty of
**    MERCHANTABILITY or FITNESS FOR A PARTICULAR PURPOSE.  See the GNU
**    Lesser General Public License for more details.
**
**    You should have received a copy of the GNU Lesser General Public
**    License along with this library; if not, write to the Free Software
**    Foundation, Inc., 59 Temple Place, Suite 330, Boston, MA  02111-1307  USA
*/


package org.systemsbiology.biofabric.util;

import java.awt.event.ActionEvent;
import java.awt.event.ActionListener;
import java.awt.event.ItemEvent;
import java.awt.event.ItemListener;
import javax.swing.table.AbstractTableModel;

import org.systemsbiology.biofabric.api.util.ExceptionHandler;

/****************************************************************************
**
** Helper to track combo box editor
*/

public class ComboBoxEditorTracker implements ItemListener, ActionListener {
  
  ////////////////////////////////////////////////////////////////////////////
  //
  // PRIVATE INSTANCE MEMBERS
  //
  ////////////////////////////////////////////////////////////////////////////  

  private ComboBoxEditor comboEdit_;
  private AbstractTableModel table_;
  private ComboFinishedTracker finished_;
  private boolean ignore_;
  
  ////////////////////////////////////////////////////////////////////////////
  //
  // PUBLIC CONSTRUCTORS
  //
  ////////////////////////////////////////////////////////////////////////////    

  /***************************************************************************
  **
  ** Constructor 
  */ 
    
  public ComboBoxEditorTracker(ComboBoxEditor comboEdit, AbstractTableModel table) {
    this(comboEdit, table, null);
  }
    
  public ComboBoxEditorTracker(ComboBoxEditor comboEdit, AbstractTableModel table, ComboFinishedTracker finished) { 
    comboEdit_ = comboEdit;
    table_ = table;
    finished_ = finished;
    ignore_ = false;    
    if (finished_ != null) {
      comboEdit_.addActionListener(this);
    } else {
      comboEdit_.addItemListener(this);
    }
  }
  
  //
  // Legacy tracking: But we may get two events per change, which
  // is bad when we are issuing dialogs:
  //
  
  public void itemStateChanged(ItemEvent e) {
    stateChangedGuts();
    return;
  }

  public void actionPerformed(ActionEvent ev) {
    stateChangedGuts();
    return;
  }
  
  private void stateChangedGuts() {
    try {
      if (ignore_) {
        return;
      }
      ignore_ = true;
      if (comboEdit_.entryIsValid()) {
        int cc = comboEdit_.getCurrColumn();
        int cr = comboEdit_.getCurrRow();
        int si = comboEdit_.getSelectedIndex();
        if (finished_ != null) {
          if (!finished_.comboEditingDone(cc, cr, comboEdit_.indexToValue(si))) {
            comboEdit_.resetValue();
          } else {
            table_.setValueAt(comboEdit_.indexToValue(si), cr, cc);
          }
        } else {
          table_.setValueAt(comboEdit_.indexToValue(si), cr, cc);
        }
      }
    } catch (Exception ex) {
      ExceptionHandler.getHandler().displayException(ex);
    } finally {
      ignore_ = false;
    }
    return;
  }  
}
```
