## Supplementary material for "BioFabric Visualization of Network Alignments": BioFabric Source Code: DataUtil.java

```
/*
**    Copyright (C) 2003-2018 Institute for Systems Biology 
**                            Seattle, Washington, USA. 
**
**    This library is free software; you can redistribute it and/or
**    modify it under the terms of the GNU Lesser General Public
**    License as published by the Free Software Foundation; either
**    version 2.1 of the License, or (at your option) any later version.
**
**    This library is distributed in the hope that it will be useful,
**    but WITHOUT ANY WARRANTY; without even the implied warranty of
**    MERCHANTABILITY or FITNESS FOR A PARTICULAR PURPOSE.  See the GNU
**    Lesser General Public License for more details.
**
**    You should have received a copy of the GNU Lesser General Public
**    License along with this library; if not, write to the Free Software
**    Foundation, Inc., 59 Temple Place, Suite 330, Boston, MA  02111-1307  USA
*/

package org.systemsbiology.biofabric.util;

import java.util.Collection;
import java.util.Collections;
import java.util.SortedSet;
import java.util.List;
import java.util.ArrayList;
import java.util.Iterator;
import java.net.URL;
import java.net.URLConnection;
import java.io.BufferedReader;
import java.io.InputStreamReader;
import java.net.URLEncoder;
import java.io.UnsupportedEncodingException;
import java.security.MessageDigest;
import java.security.NoSuchAlgorithmException;
import java.io.IOException;
import java.util.HashMap;
import java.util.HashSet;
import java.util.Map;
import java.util.Set;
import java.util.SortedMap;
import java.util.TreeMap;
import java.util.TreeSet;
import java.util.regex.Matcher;
import java.util.regex.Pattern;

import org.systemsbiology.biofabric.api.util.MinMax;

/****************************************************************************
**
** Utility for some data functions
*/

public class DataUtil {
  
  ////////////////////////////////////////////////////////////////////////////
  //
  // PUBLIC CONSTANTS
  //
  //////////////////////////////////////////////////////////////////////////// 

  ////////////////////////////////////////////////////////////////////////////
  //
  // PUBLIC CLASS METHODS
  //
  ////////////////////////////////////////////////////////////////////////////
 
  /***************************************************************************
  **
  ** Build a list when all we have is an iterator
  */  
  
  public static List listFromIterator(Iterator it) {
    ArrayList retval = new ArrayList();
    while (it.hasNext()) {
      retval.add(it.next());
    }
    return (retval);
  }
  
  /***************************************************************************
  **
  ** Build a set when all we have is an iterator
  */  
  
  public static Set setFromIterator(Iterator it) {
    HashSet retval = new HashSet();
    while (it.hasNext()) {
      retval.add(it.next());
    }
    return (retval);
  }
  
  /***************************************************************************
  **
  ** Handle incrementing a count map
  */  
  
  public static void bumpCountMap(Map map, String key) { 
    Integer countObj = (Integer)map.get(key);
    if (countObj == null) {
      countObj = new Integer(1);
    } else {
      countObj = new Integer(countObj.intValue() + 1);
    }
    map.put(key, countObj);
    return;
  }
  
  /***************************************************************************
  **
  ** Handle passwording
  */  
  
  public static String buildPasswordString(URL saltUrl, String user, String passAsString) {
    try {     
      URLConnection cc = saltUrl.openConnection();
      BufferedReader in = new BufferedReader(new InputStreamReader(cc.getInputStream()));
      String inputLine;

      ArrayList lines = new ArrayList();
      while ((inputLine = in.readLine()) != null) {
        lines.add(inputLine.trim());
      }
      in.close();

      if (lines.size() != 3) {
        return (null);
      }
      String time = (String)lines.get(0);
      String ip = (String)lines.get(1);
      String salt = (String)lines.get(2);
      user = URLEncoder.encode(user, "UTF-8");
      
      StringBuffer retval = new StringBuffer();
      retval.append("time=");
      retval.append(time);
      retval.append("&ip=");
      retval.append(ip);
      retval.append("&salt=");
      retval.append(salt);      
      retval.append("&uid=");
      retval.append(user);    
      retval.append("&hash=");
      
      String concat = time + ":" + ip + ":" + user + ":" + passAsString + ":" + salt;

      byte[] concatAsBytes = concat.getBytes("UTF-8");
      MessageDigest md = MessageDigest.getInstance("MD5");
      md.reset();
      md.update(concatAsBytes);
      byte[] digest = md.digest();
      String hexString = DataUtil.toHexString(digest);
      retval.append(hexString);
      return (retval.toString());
    } catch (UnsupportedEncodingException uee) {
      return (null);
    } catch (IOException ioe) {
      return (null);
    } catch (NoSuchAlgorithmException nsae) {
      return (null);    
    }
  }   
   
  /***************************************************************************
  **
  ** This conversion is key for matching PHP md5s.  Code snippet by "verdyp" at 
  ** http://forum.java.sun.com/thread.jspa?threadID=429739&messageID=1921162
  */

  public static String toHexString(byte[] v) {
    String digits = "0123456789abcdef";
    StringBuffer sb = new StringBuffer();
    for (int i = 0; i < v.length; i++) {
      int b = v[i] & 0xFF;
      sb.append(digits.charAt(b >>> 4)).append(digits.charAt(b & 0xF));
    }
    return sb.toString();
  }
  
  /***************************************************************************
  **
  ** Normalize the given key
  */
  
  public static String normKey(String key) {
    return (key.toUpperCase().replaceAll(" ", ""));
  }
  
  /***************************************************************************
  **
  ** Answers if normalized keys are equal
  */
  
  public static boolean keysEqual(String key1, String key2) {
    return (key1.replaceAll(" ", "").equalsIgnoreCase(key2.replaceAll(" ", "")));
  }  

  /***************************************************************************
  **
  ** Answers if normalized key is contained in the set of keys
  */
  
  public static boolean containsKey(Collection keys, String key) {
    Iterator ksit = keys.iterator();
    while (ksit.hasNext()) {
      String key1 = (String)ksit.next();
      if (key1.replaceAll(" ", "").equalsIgnoreCase(key.replaceAll(" ", ""))) {
        return (true);
      }
    }
    return (false);
  }
  
  /***************************************************************************
  **
  ** Return a list of normalized keys
  */
  
  public static List normalizeList(List keys) {
    ArrayList retval = new ArrayList();
    Iterator ksit = keys.iterator();
    while (ksit.hasNext()) {
      String key1 = (String)ksit.next();
      retval.add(key1.replaceAll(" ", "").toUpperCase());
    }
    return (retval);
  } 
  
  /***************************************************************************
  **
  ** Return a normalized key, turning greek characters into lower
  ** case (makes mismatches easier to spot)
  */
  
  public static String normKeyWithGreek(String key) {
    String retval = normKey(key);
    Pattern p = Pattern.compile(".*\\p{InGreek}.*");
    Matcher m = p.matcher(retval);
    if (m.matches()) {
      Pattern p1 = Pattern.compile("\\p{InGreek}");
      Matcher m1 = p1.matcher("");
      StringBuffer buf = new StringBuffer();
      int len = retval.length();     
      for (int i = 0; i < len; i++) {
        String oneChar = retval.substring(i, i + 1);
        m1.reset(oneChar);
        if (m1.matches()) {
          buf.append(oneChar.toLowerCase());
        } else {
          buf.append(oneChar);
        }
      }
      retval = buf.toString();
    }
    return (retval);
  }
  
  /***************************************************************************
  **
  ** Turn greek characters into lower case (makes mismatches easier to spot)
  */
  
  public static String greekToLowerCase(String val) {
    String retval = val;
    Pattern p = Pattern.compile(".*\\p{InGreek}.*");  // FIXME: make p, maybe m static class members
    Matcher m = p.matcher(retval);
    if (m.matches()) {
      Pattern p1 = Pattern.compile("\\p{InGreek}");
      Matcher m1 = p1.matcher("");
      StringBuffer buf = new StringBuffer();
      int len = retval.length();     
      for (int i = 0; i < len; i++) {
        String oneChar = retval.substring(i, i + 1);
        m1.reset(oneChar);
        if (m1.matches()) {
          buf.append(oneChar.toLowerCase());
        } else {
          buf.append(oneChar);
        }
      }
      retval = buf.toString();
    }
    return (retval);
  }
  
 
  /***************************************************************************
  **
  ** Return a map of normalized keys to original values
  */
  
  public static SortedMap normalizedMapToOrig(Collection keys) {
    TreeMap retval = new TreeMap();
    Iterator ksit = keys.iterator();
    while (ksit.hasNext()) {
      String key1 = (String)ksit.next();
      String normKey = normKey(key1);
      TreeSet setForKey = (TreeSet)retval.get(normKey);
      if (setForKey == null) {
        setForKey = new TreeSet(String.CASE_INSENSITIVE_ORDER);
        retval.put(normKey, setForKey);
      }
      setForKey.add(key1);
    }
    return (retval);
  } 
  
  /***************************************************************************
  **
  ** Get the maximum absolute value
  */  
 
  public static double maxAbsVal(List vals) {
    int num = vals.size(); 
    double maxVal = Double.NEGATIVE_INFINITY;
    for (int i = 0; i < num; i++) {
      Double val0bj = (Double)vals.get(i);
      if (val0bj.isNaN()) {
        return (val0bj.doubleValue());
      }
      double val = Math.abs(val0bj.doubleValue());
      if (val > maxVal) {
        maxVal = val;
      }
    }
    return ((maxVal == Double.NEGATIVE_INFINITY) ? Double.NaN : maxVal);
  }
  
  /***************************************************************************
  **
  ** Get the minimum absolute value
  */  
 
  public static double minAbsVal(List vals) {
    int num = vals.size(); 
    double minVal = Double.POSITIVE_INFINITY;
    for (int i = 0; i < num; i++) {
      Double val0bj = (Double)vals.get(i);
      if (val0bj.isNaN()) {
        return (val0bj.doubleValue());
      }
      double val = Math.abs(val0bj.doubleValue());
      if (val < minVal) {
        minVal = val;
      }
    }
    return ((minVal == Double.POSITIVE_INFINITY) ? Double.NaN : minVal);
  }
  
  /***************************************************************************
  **
  ** Average absolute value
  */  
 
  public static double avgAbsVal(Collection vals) {
    Iterator vit = vals.iterator(); 
    double sum = 0.0;
    int num = vals.size();
    if (num == 0) {
      return (Double.NaN);
    }
    while (vit.hasNext()) {
      Double val0bj = (Double)vit.next();
      if (val0bj.isNaN()) {
        return (val0bj.doubleValue());
      }
      double val = Math.abs(val0bj.doubleValue());
      sum += val;
    }
    return (sum / (double)num); 
  }
  
  /***************************************************************************
  **
  ** Average value
  */  
 
  public static double avgVal(List vals) {
    int num = vals.size();   
    double sum = 0.0;
    if (num == 0) {
      return sum;
    }
    for (int i = 0; i < num; i++) {
      Double val0bj = (Double)vals.get(i);
      sum += val0bj.doubleValue();
    }
    return (sum / (double)num); 
  }
 
  /***************************************************************************
  **
  ** Median absolute value
  */  
 
  public static double medianAbsVal(List vals) {
    if (vals.isEmpty()) {
      return (Double.NaN);
    }
    int num = vals.size();   
    ArrayList sorted = new ArrayList();
 
    for (int i = 0; i < num; i++) {
      Double val0bj = (Double)vals.get(i);
      if (val0bj.isNaN()) {
        return (Double.NaN);
      }
      sorted.add(new Double(Math.abs(val0bj.doubleValue())));
    }
    Collections.sort(sorted);
    int lindex = num / 2;    
    if ((num % 2) == 0) {
      Double lower = (Double)sorted.get(lindex - 1);
      Double upper = (Double)sorted.get(lindex);
      return ((lower.doubleValue() + upper.doubleValue()) / 2.0);
    } else {
      Double median = (Double)sorted.get(lindex);
      return (median.doubleValue());
    }
  }   
  
  /***************************************************************************
  **
  ** Return if average of numbers above threshold
  */  
 
  public static boolean avgValIsAboveThresh(List vals, double negThresh, double posThresh) {
    int num = vals.size();
    int count = 0;
    double sum = 0.0;
    for (int i = 0; i < num; i++) {
      Double val0bj = (Double)vals.get(i);
      if (val0bj.isNaN()) {
        continue;
      }
      double valv = val0bj.doubleValue();
      sum += valv;
      count++;
    }
    double avg = sum / (double)count;
    return ((avg <= negThresh) || (avg >= posThresh));
  }
  
  /***************************************************************************
  **
  ** Return median of numbers above threshold
  */  
 
  public static double medianAbsValAboveThresh(List vals, double thresh) {
    int num = vals.size();
    ArrayList above = new ArrayList();
    for (int i = 0; i < num; i++) {
      Double valObj = (Double)vals.get(i);
      if (valObj.isNaN()) {
        return (Double.NaN);
      }
      double val = Math.abs(valObj.doubleValue());
      if (val >= thresh) {
        above.add(valObj);
      }
    }
    return (medianAbsVal(above)); 
  }  
 
  /***************************************************************************
  **
  ** Return minimum value above (or at) threshold
  */  
 
  public static double minAbsValAboveThresh(List vals, double thresh) {
    int num = vals.size(); 
    double minVal = Double.POSITIVE_INFINITY;
    for (int i = 0; i < num; i++) {
      Double val0bj = (Double)vals.get(i);
      if (val0bj.isNaN()) {
        return (Double.NaN);
      }
      double val = Math.abs(val0bj.doubleValue());
      if ((val >= thresh) && (val < minVal)) {
        minVal = val;
      }
    }
    return ((minVal == Double.POSITIVE_INFINITY) ? Double.NaN : minVal);
  }

  /***************************************************************************
  **
  ** Look for mixed signs
  */  
 
  public static boolean signsMixed(List vals) {
    int num = vals.size();
    int runningSign = 0;    
    for (int i = 0; i < num; i++) {
      Double val = (Double)vals.get(i);
      double dVal = val.doubleValue();
      int currSign = 0;
      if (dVal < 0.0) {
        currSign = -1;
      } else if (dVal > 0.0) {
        currSign = 1;
      }
      if ((i == 0) || (runningSign == 0)) {
        runningSign = currSign;
      } else if (currSign == 0) {
        continue;
      } else if (currSign != runningSign) {
        return (true);
      }
    }
    return (false);
  }
  
  /***************************************************************************
  **
  ** Answer if list has a member above the threshold
  */

  public static boolean haveOne(List vals, double negThresh, double posThresh) {
    int numVals = vals.size();
    for (int i = 0; i < numVals; i++) {
      Double val = (Double)vals.get(i);
      double valv = val.doubleValue();
      if ((valv <= negThresh) || (valv >= posThresh)) {
        return (true);     
      }
    }
    return (false);
  }  
  
  /***************************************************************************
  **
  ** Answer if list has one true
  */

  public static boolean haveOneTrue(List vals) {
    int numVals = vals.size();
    for (int i = 0; i < numVals; i++) {
      Boolean val = (Boolean)vals.get(i);
      if (val.booleanValue()) {
        return (true);     
      }
    }
    return (false);
  }
 
  /***************************************************************************
  **
  ** Answer if list has one false
  */

  public static boolean haveOneFalse(List vals) {
    int numVals = vals.size();
    for (int i = 0; i < numVals; i++) {
      Boolean val = (Boolean)vals.get(i);
      if (!val.booleanValue()) {
        return (true);     
      }
    }
    return (false);
  }
  
  /***************************************************************************
  **
  ** Answer if list has a majority above the threshold
  */

  public static boolean haveMajority(List vals, double negThresh, double posThresh, boolean tieWins) {
    int numVals = vals.size();
    int goodCount = 0;
    int badCount = 0;
    for (int i = 0; i < numVals; i++) {
      Double val = (Double)vals.get(i);
      double valv = val.doubleValue();
      if ((valv <= negThresh) || (valv >= posThresh)) {
        goodCount++;
      } else {
        badCount++; 
      }
    }
    // Success for majority count:
    return ((((numVals % 2) == 0) && tieWins) ? (goodCount >= badCount) : (goodCount > badCount));
  }

  /***************************************************************************
  **
  ** Answer if list has a majority of trues
  */

  public static boolean haveMajorityTrue(List vals, boolean tieWins) {
    int numVals = vals.size();
    int goodCount = 0;
    int badCount = 0;
    for (int i = 0; i < numVals; i++) {
      Boolean val = (Boolean)vals.get(i);
      if (val.booleanValue()) {
        goodCount++;
      } else {
        badCount++; 
      }
    }
    // Success for majority count:
    return ((((numVals % 2) == 0) && tieWins) ? (goodCount >= badCount) : (goodCount > badCount));
  }
 
  /***************************************************************************
  **
  ** Answer if list has a NaN
  */

  public static boolean hasANaN(List vals) {
    int numVals = vals.size();
    for (int i = 0; i < numVals; i++) {
      Double val = (Double)vals.get(i);
      if (val.isNaN()) {
        return (true);
      }
    }
    return (false);
  }  
  
  /***************************************************************************
  **
  ** Return the count of elements at or above the threshold (abs val)
  */

  public static int numberAboveThresh(List vals, double thresh) {
    int numVals = vals.size();
    int count = 0;
    thresh = Math.abs(thresh);
    for (int i = 0; i < numVals; i++) {
      Double val = (Double)vals.get(i);
      if (val.isNaN()) {
        continue;
      }
      if (Math.abs(val.doubleValue()) >= thresh) {
        count++;
      }
    }
    return (count);
  } 
  
  /***************************************************************************
  **
  ** Return the geometric mean of the absolute values of elements
  */

  public static double geometricMean(List vals, double replacement) {
    int numVals = vals.size();
    double product = 1.0;
    for (int i = 0; i < numVals; i++) {
      Double val = (Double)vals.get(i);
      double dval = (val.isNaN()) ? replacement : val.doubleValue();
      if (dval == 0.0) {
        dval = replacement;
      }
      product *= Math.abs(dval);
    }
    return (Math.pow(product, 1.0 / (double)numVals));
  }
  
  /***************************************************************************
  **
  ** Fill the set of hours to be hourly
  */  
 
  public static SortedSet<Integer> fillOutHourly(SortedSet<Integer> hours) {
    TreeSet<Integer> retval = new TreeSet<Integer>(hours);
    if (hours.isEmpty() || (hours.size() == 1)) {
      return (retval);
    }
    
    int minTime = hours.first().intValue() + 1;
    int maxTime = hours.last().intValue() - 1;
    for (int i = minTime; i <= maxTime; i++) {
      retval.add(new Integer(i));
    }
    return (retval);
  }
  
  
  /***************************************************************************
  **
  ** Given a set of integers, find the closest to the requested value.  For ties,
  ** use the boolean directive;
  */  
 
  public static Integer closestInt(Set vals, int want, boolean tieBreakBelow) {
    
    if (vals.isEmpty()) {
      return (null);
    }
    Integer candBelow = null;
    Integer candAbove = null;
    int absDiffBelow = Integer.MAX_VALUE;
    int absDiffAbove = Integer.MAX_VALUE;
 
    Iterator vit = vals.iterator();
    while (vit.hasNext()) {
      Integer next = (Integer)vit.next();
      int nVal = next.intValue();
      if (nVal == want) {
        return (next);
      }
      int diff = want - nVal;
      int absDiff = Math.abs(diff);
      if (diff > 0) {
        if (absDiff < absDiffBelow) {
          candBelow = next;
          absDiffBelow = absDiff;
        }
      } else {
        if (diff < absDiffAbove) {
          candAbove = next;
          absDiffAbove = absDiff;
        }
      }  
    }
    if (absDiffBelow == absDiffAbove) {
      return ((tieBreakBelow) ? candBelow : candAbove);
    } else {
      return ((absDiffBelow < absDiffAbove) ? candBelow : candAbove);
    }
  }

  /***************************************************************************
  **
  ** Given a sorted set of integers, find the two values bounding the requested value.
  ** Above, below, or equal returns min == max
  */  
 
  public static MinMax boundingInts(SortedSet vals, int want) {
    
    MinMax retval = new MinMax();
    retval.init();
    
    if (vals.isEmpty()) {
      return (null);
    }
    
    Integer candBelow = null;
    Integer candAbove = null;
    int absDiffBelow = Integer.MAX_VALUE;
    int absDiffAbove = Integer.MAX_VALUE;
    boolean isFirst = true;
    
    Iterator vit = vals.iterator();
    while (vit.hasNext()) {
      Integer next = (Integer)vit.next();
      int nVal = next.intValue();
      if (nVal == want) {
        retval.update(nVal);
        return (retval);
      }
      if (isFirst) {
        isFirst = false;
        if (want < nVal) {
          retval.update(nVal);
          return (retval);         
        }
      }
      if (!vit.hasNext()) {
        if (want > nVal) {
          retval.update(nVal);
          return (retval);         
        }
      }

      int diff = want - nVal;
      int absDiff = Math.abs(diff);
      if (diff > 0) {
        if (absDiff < absDiffBelow) {
          candBelow = next;
          absDiffBelow = absDiff;
        }
      } else {
        if (diff < absDiffAbove) {
          candAbove = next;
          absDiffAbove = absDiff;
        }
      }  
    }
    retval.update(candAbove.intValue());
    retval.update(candBelow.intValue());
    return (retval);
  }
 
  /***************************************************************************
  **
  ** Trim the set on the top and bottom
  */  
 
  public static SortedSet trimOut(SortedSet argSet, int min, int max) {
    TreeSet retval = new TreeSet();
    if (argSet.isEmpty()) {
      return (retval);
    }
    
    Iterator asit = argSet.iterator();
    while (asit.hasNext()) {
      Integer nint = (Integer)asit.next();
      int intval = nint.intValue();
      if ((intval >= min) && (intval <= max)) {
        retval.add(nint);
      } 
    }
    return (retval);
  }
  
  
  /***************************************************************************
  **
  ** Do a doubling that creates a set of times where start and end times
  ** become distinct even/odd entities in the set.
  */  
 
  public static SortedSet doubleForDescreteStartAndEnds(SortedSet hours) {
    TreeSet retval = new TreeSet();
    if (hours.isEmpty()) {
      return (retval);
    }
    Iterator vit = hours.iterator();
    while (vit.hasNext()) {
      Integer time = (Integer)vit.next();
      int start = time.intValue() * 2;
      retval.add(new Integer(start));
      retval.add(new Integer(start + 1));
    }
    return (retval);
  }
  
  /***************************************************************************
  **
  ** Set intersection helper
  */  
 
  public static Set intersection(Set one, Set two, Set result) {
    result.clear();
    result.addAll(one);
    result.retainAll(two);
    return (result);
  }  
 
  /***************************************************************************
  **
  ** Set union helper
  */  
 
  public static Set union(Set one, Set two, Set result) {
    result.clear();
    result.addAll(one);
    result.addAll(two);
    return (result);
  }  
  
  /***************************************************************************
  **
  ** Merge the new set in, coalese so the set of sets remains disjoint
  */
  
  public static Set coaleseSets(Set setOfSets, Set newSet) {
    
    //
    // Merge together.  In general, the sets in the set are
    // no longer disjoint.
    //
    HashSet result = new HashSet(setOfSets);
    result.add(newSet);
    
    //
    // Get a fixed ordering of all set members.
    //
    
    HashSet allMembers = new HashSet();
    Iterator rit = result.iterator();
    while (rit.hasNext()) {
      Set nextSet = (Set)rit.next();
      allMembers.addAll(nextSet);
    }
    ArrayList fixedOrder = new ArrayList(allMembers);
    
    //
    // For each member, if it appears in two sets,
    // we extract those sets from the result, and
    // return the union.  Continue this until the
    // scan is clean for all members!
    //
    
    
    Iterator smit = fixedOrder.iterator();
    while (smit.hasNext()) {
      Object member = (Object)smit.next();     
      boolean done = false;
      while (!done) {
        Set firstSet = null;
        boolean collision = false;
        Iterator rsit = result.iterator();
        while (rsit.hasNext()) {
          Set nextSet = (Set)rsit.next();
          if (nextSet.contains(member)) {
            if (firstSet == null) {
              firstSet = nextSet;
            } else {
              result.remove(nextSet);
              result.remove(firstSet);
              HashSet union = new HashSet();
              DataUtil.union(nextSet, firstSet, union);
              result.add(union);
              collision = true;
              break;
            }
          }
        }
        if (!collision) {
          done = true;
        }
      }
    }
    return (result);
  }
  
  /***************************************************************************
  **
  ** Find index of first difference.  -1 is no diff:
  */  
 
  public static int strDiff(String str1, String str2) {
    int len1 = str1.length();
    int len2 = str2.length();
    
    // Only go to the end of the shorter:
    int uselen = (len1 > len2) ? len2 : len1;
    
    // find the diff:
    for (int i = 0; i < uselen; i++) {
      char ch1 = str1.charAt(i);
      char ch2 = str2.charAt(i);
      if (ch1 != ch2) {
        return (i);
      }
    }
    // no diff: Now depends on length
    return ((len1 == len2) ? -1 : uselen); 
  }  
   
  /***************************************************************************
  **
  ** Get a comma and & delineated display string
  **
  */
  
  public static String getMultiDisplayString(List asNames) {
    StringBuffer buf = new StringBuffer();
    int numSrc = asNames.size();
    for (int i = 0; i < numSrc; i++) {
      String invest = (String)asNames.get(i);
      buf.append(invest);
      if (i < (numSrc - 2)) {
        buf.append(", ");
      } else if (i < (numSrc - 1)) {
        buf.append(" & ");
      }
    }
    return (buf.toString());
  }
  
  /***************************************************************************
  **
  ** Derived from pseudocode at: http://en.wikipedia.org/wiki/Levenshtein_distance
  ** Used to determine edit distances between two strings
  */
  
  public static int levenshteinDistance(String s, String t) {
    
    int m = s.length();
    int n = t.length();
   
    int dtable[][] = new int[m + 1][n + 1]; 

    for (int i = 0; i <= m; i++) {
      dtable[i][0] = i; // deletion
    }
   
    for (int j = 0; j <= n; j++) {
      dtable[0][j] = j; // insertion
    }
   
    for (int j = 1; j <= n; j++) {
      for (int i = 1; i <= m; i++) {
        if (s.charAt(i - 1) == t.charAt(j - 1)) {
          dtable[i][j] = dtable[i - 1][j - 1];
        } else {
          int delDist = dtable[i - 1][j] + 1;  // deletion
          int insDist = dtable[i][j - 1] + 1;  // insertion
          int subDist = dtable[i - 1][j - 1] + 1; // substitution
          dtable[i][j] = Math.min(Math.min(delDist, insDist), subDist);
        }
      }
    }
  
    return (dtable[m][n]);
  }
  
  /***************************************************************************
  **
  ** Get the closest existing target name:
  */
  
  public static String getClosestStringToName(String targName, int distThresh, Set sources) {
    String normName = DataUtil.normKey(targName);
    int minDist = distThresh + 1;
    String minStr = null;
    Iterator ikit = sources.iterator();
    while (ikit.hasNext()) { 
      String trg = (String)ikit.next();
      int editDist = DataUtil.levenshteinDistance(normName, DataUtil.normKey(trg));
      if (editDist < minDist) {
        minDist = editDist;
        minStr = trg;
      }
    }
    return (minStr);
  }  
  
  /***************************************************************************
  **
  ** Set distance
  */
  
  public static int setDistance(SortedSet s1, SortedSet s2) {
    TreeSet union = new TreeSet(s1);
    union.addAll(s2);
    int sum = 0;
    Iterator uit = union.iterator();
    while (uit.hasNext()) {
      String unext = (String)uit.next();
      int increment = (s1.contains(unext) && s2.contains(unext)) ? 0 : 1;
      sum += increment;
    }
    return (sum);
  }
  
  /***************************************************************************
  **
  ** All (abs vals) above threshold?
  */
  
  public static boolean allAboveThreshold(List values, double threshold) {
    if ((values == null) || values.isEmpty()) {
      throw new IllegalArgumentException();
    }
    int numVal = values.size();
    for (int i = 0; i < numVal; i++) {
      Double val = (Double)values.get(i);
      if (Math.abs(val.doubleValue()) < threshold) {
        return (false);
      }
    }
    return (true); 
  }   

  /***************************************************************************
  **
  ** Get the average value for the list.  List size must be > 0.
  */
  
  public static double listAverage(List values) {
    if ((values == null) || values.isEmpty()) {
      throw new IllegalArgumentException();
    }
    double sum = 0.0;
    int count = 0;
    int numVal = values.size();
    for (int i = 0; i < numVal; i++) {
      Double val = (Double)values.get(i);
      sum += val.doubleValue();
      count++;
    }
    return (sum / (double)count); 
  } 
  
  /***************************************************************************
  **
  ** Get the value with the maximum absolute value. (Returned value may be negative)
  */
  
  public static double listMaxAbs(List values) {
    if ((values == null) || values.isEmpty()) {
      throw new IllegalArgumentException();
    }
    double maxAbs = 0.0;
    double max = 0.0;
    int numVal = values.size();
    for (int i = 0; i < numVal; i++) {
      Double valObj = (Double)values.get(i);
      double val = valObj.doubleValue();      
      double abs = Math.abs(val);
      if (abs > maxAbs) {
        maxAbs = abs;
        max = val;
      }
    }
    return (max); 
  } 
  
  /***************************************************************************
  **
  ** negate values of a map to Integers
  */
  
  public static Map negateMapToIntegers(Map values) {
    HashMap retval = new HashMap();
    Iterator vit = values.keySet().iterator();
    while (vit.hasNext()) {
      Object key = vit.next();
      Integer val = (Integer)values.get(key);
      retval.put(key, new Integer(-val.intValue()));
    }
    return (retval); 
  } 
  
  ////////////////////////////////////////////////////////////////////////////
  //
  // PRIVATE CONSTRUCTORS
  //
  ////////////////////////////////////////////////////////////////////////////

  /***************************************************************************
  **
  ** Null constructor.  Do not use
  */

  private DataUtil() {
  }
}
```
