## Supplementary material for "BioFabric Visualization of Network Alignments": BioFabric Source Code: DoubMinMax.java

```
/*
**    Copyright (C) 2003-2018 Institute for Systems Biology 
**                            Seattle, Washington, USA. 
**
**    This library is free software; you can redistribute it and/or
**    modify it under the terms of the GNU Lesser General Public
**    License as published by the Free Software Foundation; either
**    version 2.1 of the License, or (at your option) any later version.
**
**    This library is distributed in the hope that it will be useful,
**    but WITHOUT ANY WARRANTY; without even the implied warranty of
**    MERCHANTABILITY or FITNESS FOR A PARTICULAR PURPOSE.  See the GNU
**    Lesser General Public License for more details.
**
**    You should have received a copy of the GNU Lesser General Public
**    License along with this library; if not, write to the Free Software
**    Foundation, Inc., 59 Temple Place, Suite 330, Boston, MA  02111-1307  USA
*/

package org.systemsbiology.biofabric.util;

public class DoubMinMax implements Cloneable {
  
  public static final int EQUALS             = 0;   
  public static final int IS_PROPER_SUPERSET = 1;
  public static final int IS_PROPER_SUBSET   = 2;
  public static final int IS_DISJOINT        = 3;
  public static final int INTERSECTS         = 4;
  
  public double min;
  public double max;
  
  public DoubMinMax() {
  }

  public DoubMinMax(double val) {
    this.min = val;
    this.max = val;
  }  
  
  public DoubMinMax(double min, double max) {
    this.min = min;
    this.max = max;
  }
  
  public DoubMinMax(DoubMinMax other) {
    this.min = other.min;
    this.max = other.max;
  }
  
  public Object clone() {
    try {
      DoubMinMax newVal = (DoubMinMax)super.clone();
      return (newVal);
    } catch (CloneNotSupportedException ex) {
      throw new IllegalStateException();     
    }    
  }
  
  public int hashCode() {
    return ((int)Math.round(min + max));
  }
  
  public boolean equals(Object other) {
    if (other == null) {
      return (false);
    }
    if (other == this) {
      return (true);
    }
    if (!(other instanceof DoubMinMax)) {
      return (false);
    }
    DoubMinMax otherMM = (DoubMinMax)other;
    return ((this.min == otherMM.min) && (this.max == otherMM.max));
  }  
 
  public boolean contained(double val) {
    return ((val >= min) && (val <= max));
  } 
  
  public boolean outsideOrOnBoundary(double val) {
    return ((val <= min) || (val >= max));
  } 
   
  public DoubMinMax union(DoubMinMax other) {
    DoubMinMax retval = new DoubMinMax();
    retval.min = (this.min < other.min) ? this.min : other.min;
    retval.max = (this.max > other.max) ? this.max : other.max;    
    return (retval);
  } 

  public DoubMinMax init() {
    min = Double.NEGATIVE_INFINITY;
    max = Double.POSITIVE_INFINITY;
    return (this);
  }     
  
  public DoubMinMax inverseInit() {
    max = Double.NEGATIVE_INFINITY;
    min = Double.POSITIVE_INFINITY;
    return (this);
  }
  
  public void update(int newVal) {
    if (newVal < min) {
      min = newVal;
    }
    if (newVal > max) {
      max = newVal;
    }
    return;
  }
  
  public void update(double newVal) {
    if (newVal < min) {
      min = newVal;
    }
    if (newVal > max) {
      max = newVal;
    }
    return;
  }
 
  /**
  *** Answer how the given MinMax relates to this one.
  **/
  
  public int evaluate(DoubMinMax other) {
    if ((this.min == other.min) && (this.max == other.max)) {
      return (EQUALS);    
    } else if ((this.min >= other.min) && (this.max <= other.max)) {
      return (IS_PROPER_SUPERSET);
    } else if ((this.min <= other.min) && (this.max >= other.max)) {
      return (IS_PROPER_SUBSET);  
    } else if ((this.max < other.min) || (this.min > other.max)) {      
      return (IS_DISJOINT);
    } else {
      return (INTERSECTS);
    }
  }
 
  public String toString() {
    return ("min: " + min + " max: " + max);
  }
}
```
