## Supplementary figures and images for "BioFabric Visualization of Network Alignments"

### About24.gif

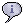

### Back24.gif

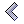

### BioFab16White.gif

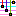

### C24.gif

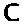

### ClearFabricSelected24.gif

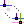

### CrossCursor32.gif

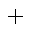

### D24.gif

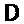

### ErrorCursor32.gif

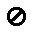

### Find24.gif

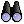

### Forward24.gif

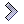

### H24.gif

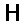

### L24.gif

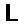

### MacBlank.gif

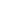

### MacC.gif

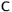

### MacD.gif

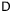

### MacGT.gif

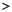

### MacL.gif

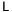

### MacLT.gif

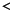

### MacN.gif

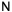

### MacR.gif

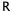

### MacS.gif

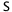

### MacU.gif

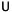

### MacZ.gif

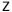

### N24.gif

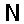

### P24.gif

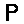

### P24Selected.gif

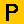

### PlusOneDeg24.gif

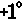

### Print24.gif

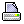

### PropagateSelected24.gif

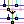

### S24.gif

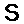

### Save24.gif

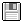

### SaveAs24.gif

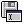

### Stop24.gif

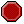

### U24.gif

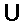

### U24Selected.gif

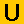

### ZoomIn24.gif

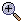

### ZoomOut24.gif

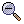

### ZoomToAllFabric24.gif

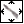

### ZoomToAllFabricSelected24.gif

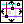

### ZoomToFabricRect24.gif

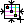

### ZoomToFabricSelected24.gif

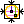
